# Abundant capped RNAs are derived from mRNA cleavage at 3’UTR G-Quadruplexes

**DOI:** 10.1101/2023.04.27.538568

**Authors:** Nejc Haberman, Holly Digby, Rupert Faraway, Rebecca Cheung, Andrew M Jobbins, Callum Parr, Kayoko Yasuzawa, Takeya Kasukawa, Chi Wai Yip, Masaki Kato, Hazuki Takahashi, Piero Carninci, Santiago Vernia, Jernej Ule, Christopher R Sibley, Aida Martinez-Sanchez, Boris Lenhard

## Abstract

The 3’ untranslated region (3’UTR) plays a crucial role in determining mRNA stability, localisation, translation and degradation. Cap analysis gene expression (CAGE), a method for the detection of capped 5’ ends of mRNAs, additionally reveals a large number of apparently 5’ capped RNAs derived from 3’UTRs. Here we provide the first direct evidence that these 3’UTR-derived RNAs are indeed capped and often more abundant than the corresponding full-length mRNAs. By using a combination of AGO2 enhanced individual nucleotide resolution UV crosslinking and immunoprecipitation (eiCLIP) and CAGE following siRNA knockdowns, we find that these 3’UTR-derived RNAs likely originate from AGO2-mediated cleavage, and most often occur at locations with potential to form RNA-G-quadruplexes and are enriched by RNA-binding protein UPF1. High-resolution imaging and long-read sequencing analysis validates several 3’UTR-derived RNAs, demonstrates their abundance and shows that they tend not to co-localise with the parental mRNAs. We also find that production of 3’UTR-derived RNA could explain the previously reported role of a 3’UTR G-quadruplex in regulating the production of APP protein. Taken together, we provide new insights into the origin and abundance of 3’UTR-derived RNAs, show the utility of CAGE-seq for their quantitative detection, and provide a rich dataset for exploring new biology of a poorly understood new class of RNAs.

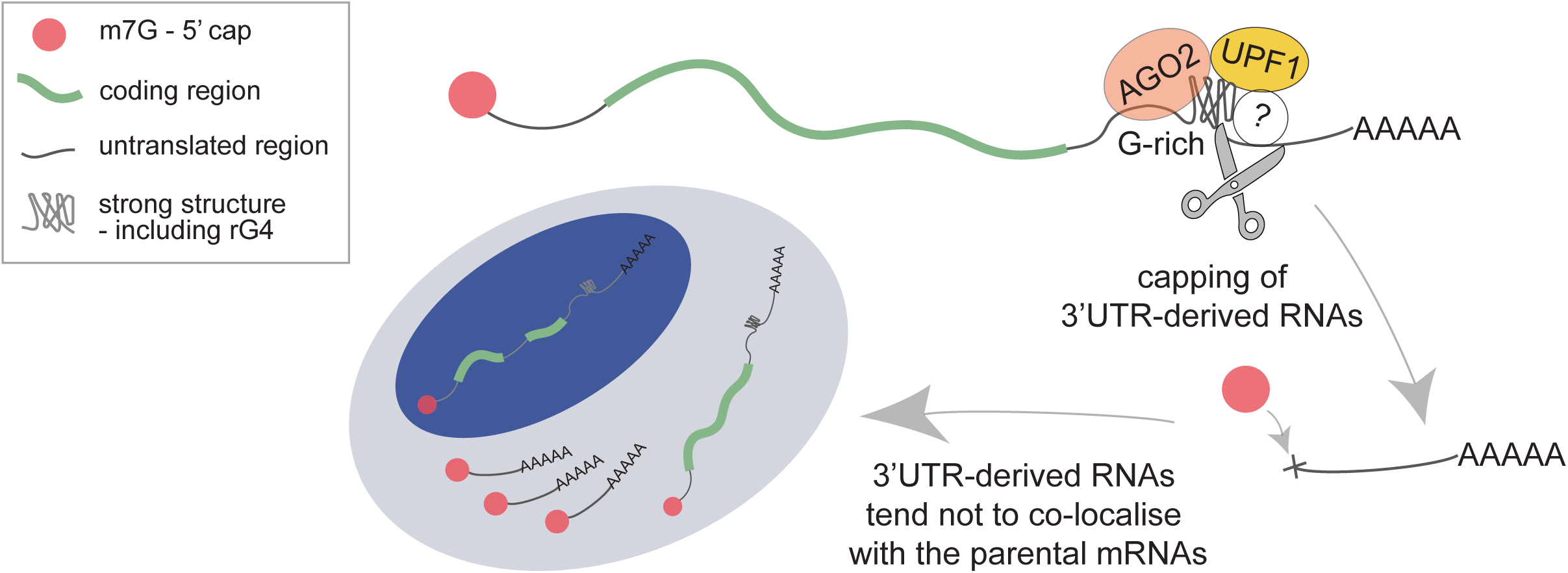

## Introduction

In all eukaryotes, mRNA molecules contain an evolutionarily conserved m7G cap (N7-methylated guanosine), which gets added at the 5’ end of nascent transcripts. Co-transcriptional capping is the first modification made to nascent RNA in the nucleus, which protects it from exonuclease cleavage while promoting cap-related biological functions such as pre-mRNA splicing, polyadenylation and nuclear export ^1^. In addition to the co-transcriptional capping of nascent mRNAs, there is evidence for a post-transcriptional capping mechanism, which adds cap to newly exposed 5’ ends of RNA fragments created upon endonucleolytic cleavage ^2–4^. However, little is known about the extent and biological roles of such post-transcriptional capping, and of its relation to other post-transcriptional RNA processing mechanisms.

Cap analysis of gene expression and deep-sequencing (CAGE-seq) was originally designed to precisely determine transcription start site (TSS) positions by capturing and sequencing 5’ ends of capped mRNA transcripts, and it can also be used to measure gene expression ^5^. However, several studies detected an unexpected, reproducible, and so-far unexplained enrichment (∼10-15%) of CAGE signal, or significant enrichment of RNA-seq reads, in the untranslated region within the 3’UTR, far away from the usual TSS ^6–14^. Previous studies have shown an absence of active promoter marks (i.e. no enrichment of modified histones or RNAPII) around these 3’UTR signals ^6, 8, 15^, arguing against the possibility that they are unannotated transcription start sites. Moreover, their expression profiles have been reported to be separated from the associated protein-coding sequence in a subcellular specific manner, with expression changes detected in several 3’UTRs during differentiation stages in mouse embryos ^15^. In addition, specific isolated 3’UTRs have been implicated in a growing number of physiological and pathological processes ^6, 16, 17^. Some processed and capped 3’UTRs have been reported to play important roles in regulating protein expression in trans, similar to long non-coding RNAs ^6, 8, 9^. It has been suggested that some 3’UTR CAGE signal arises as a consequence of post-transcriptional cleavage followed by capping mechanism ^2–4, 14^ rather than through conventional transcription initiation, and that this can lead to 3’UTR-derived RNAs that have been referred to as 3’UTR-associated RNAs (uaRNAs). To avoid potential misunderstandings we refer to these as 3’UTR-derived RNAs, as these newly generated RNAs are not known to be physically associated with 3’UTRs.

Here, we thoroughly examined the presence of these 3’UTR-derived RNAs across the transcriptome, and the molecular basis of their generation and regulation. We perform a genome-wide identification of 3’ UTR-derived RNAs based on their capped 5’ ends, and proceed to investigate the mechanisms involved in their formation. To this end, we combine CAGE, RNA-seq and cross-linking immunoprecipitation (CLIP)-based techniques from ENCODE and FANTOM consortia to detect 3’UTR-derived RNAs genome-wide. We show that those RNAs have biochemical properties expected of 5’ capped RNAs, that may originate by endonucleolytic cleavage of the host mRNA, and that they are often as abundant, or more so, as the protein-coding part of the host transcript. We support this by showing that the apparent cleavage sites near the 5’ ends of these 3’UTR-derived RNAs are bound by UPF1 and AGO2, and also have a tendency to form at RNA-G-quadruplexes. Moreover, some of those abundant 3’ UTR-derived RNAs show markedly different subcellular localisation than their protein-coding counterparts. Finally, we show that equivalent 3’ UTR capped RNAs can result from siRNA-mediated cleavage of RNA.

## Results

### CAGE-seq identifies non-promoter associated capped 3’UTR-derived RNAs

Ourselves and others ^6–11^ have previously reported the presence of CAGE-seq signals outside of annotated promoter regions in thousands of protein-coding genes, albeit their origin or biological relevance had not been interrogated. Here we first confirmed the abundance of these signals in human cell lines using CAGE data provided by the ENCODE consortium. As expected, we could detect a similar proportion of CAGE signals per genomic region in two human cell lines, as well as show that the CAGE signal is highly reproducible across replicates. This included in proportion, library size, position and distribution of the CAGE tags (Figure 1A, S1A,B,C). A similar ratio was also detected by other groups before, using the same protocol ^18^.

**Figure 1.**
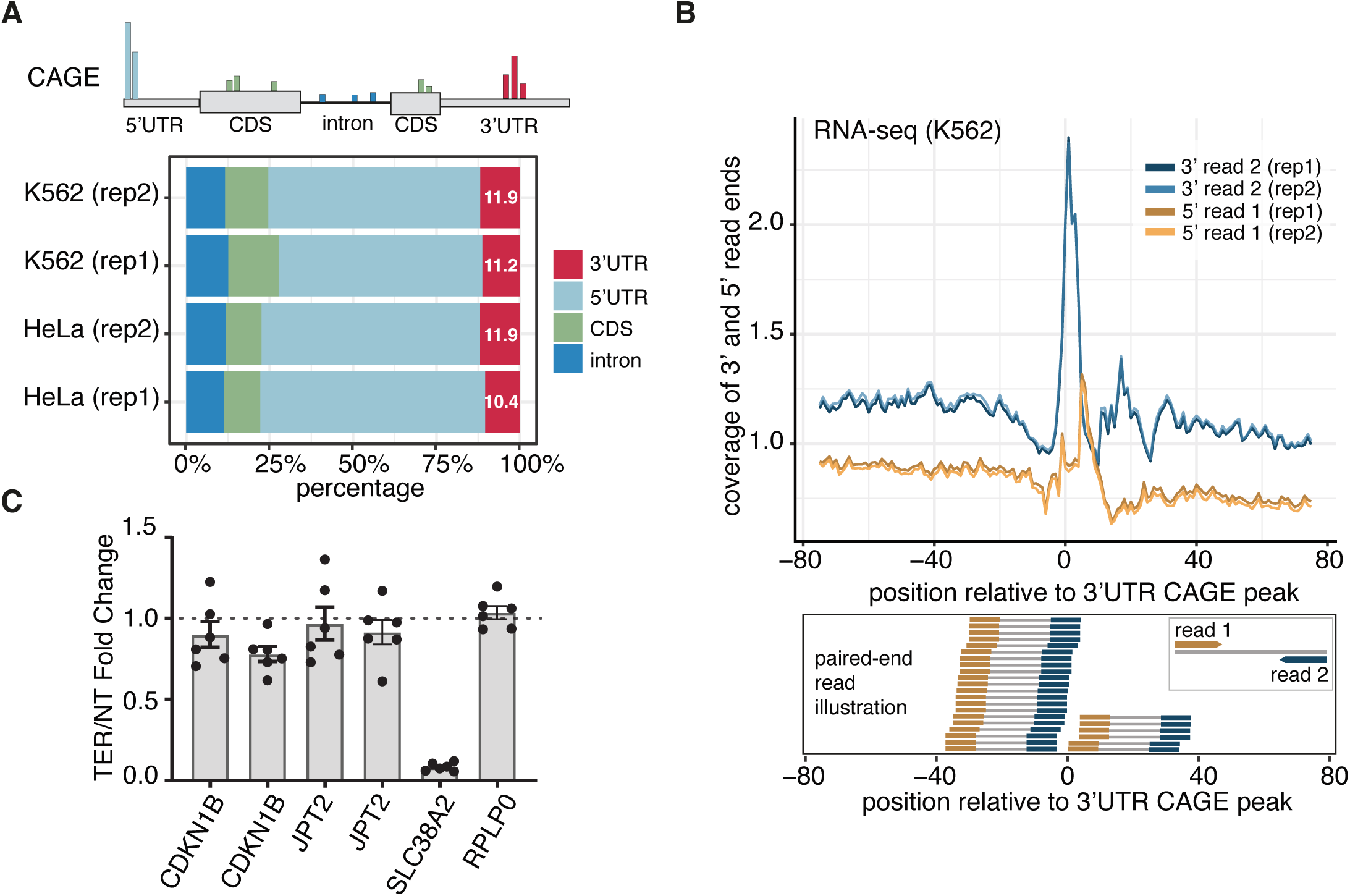
CAGE-seq identifies non-promoter associated capped 3’UTR-derived RNAs. (A) Schematic representation of CAGE signal across parts of transcripts, followed by proportions of tag clusters identified in CAGE-seq libraries of K562 and HeLa samples with two biological replicates each, provided by ENCODE. (B) The 3’ of downstream paired read (blue lines) and 5’ of upstream paired read (yellow lines) of RNA-seq (K562) normalised coverage around the 3’UTR CAGE (K562) peaks, followed by schematic representation of paired-end read positioning. The distribution represents an independent confirmation of cleavage sites by RNA-seq. (C) RT-qPCR data of gene expression ratios using primers amplifying regions immediately upstream (5’C) and downstream (3’C) of the 3‘UTR CAGE peak, except for SLC38A2 whose 3’ cleavage site results in uncapped downstream fragment. Data is presented as a fold-change of samples treated with TerminatorTM 5’-Phosphate-Dependent Exonuclease (TEX), which degrades uncapped RNAs, versus non-treated (NT).

The relative intensities of CAGE signal detected at different elements of the structural gene depend on the priming method for reverse transcription (oligo-dT, random hexamers, or mixtures thereof in different ratios) ^11^. Oligo-dT priming quantitatively favours shorter transcripts, while the reverse is true for random priming. We subsequently verified that 3’UTR CAGE signal is optimally detected when a combination of Oligo-dT and random primers is used, with the optimal inclusion ratio of 1 to 4 ratio of Oligo-dT to random primers ^18, 19^ (Figure S1D). Notably, the same ratio was used in ENCODE CAGE samples analysed in this study.

The CAGE signal is the strongest at 5’UTRs of known protein-coding genes ^18^ (Figure 1A, ∼65% of promoter signal). While low-level non-promoter CAGE signal (sometimes referred to as “exon painting”), can be detected along the entire length of transcripts, the signal at 3’ UTRs is consistently present and occurs in localised clusters, like at promoters (see Figure S1I for examples). We focused on the 3’UTR region, since a substantial amount (∼11%) of total CAGE reads map there (Figure 1A), and the significance of this is unknown. To identify robust CAGE signals with sufficient sensitivity, we used a 20nt window requiring at least two 5’ reads overlapping from two different replicates for each cell line separately. This revealed 32,065 unique 3’UTR CAGE clusters across all samples (Table 1). Moreover, these 3’UTR CAGE clusters showed high reproducibility, suggesting biological relevance; there was ∼0.9 correlation between replicates, and 0.98 between HeLa and K562 samples (Figure S1E). The latter correlation is higher than that for the 5’ UTR CAGE signal between the two cell types (0.79 correlation and Figure S1F). Together these analyses show that the transcripts whose 5’ end map to 3’UTR ends of protein-coding genes are abundant and reproducible across cell types, and that CAGE is a robust method for their quantitative detection.

### 3’UTR-derived RNAs are confirmed by RNA-seq, qPCR and long-read CAGE

We next wanted to investigate whether the 3’UTR CAGE signals originate from post-transcriptionally capped RNA fragments. First, we asked if there is support for the ends of the corresponding cleavage fragments sites in transcriptomic data produced by independent methods. For this we compared the CAGE signal with the RNA-seq signal of two different cell lines. To categorise CAGE peaks we first used the paraclu ^20^ peak caller to identify clusters of 5’ ends of capped RNAs, and within each cluster we selected the highest signal as dominant CAGE peak position. For comparison, we processed paired-end RNA-seq data from the same K562 and HeLa cell lines, then plotted read-starts and read-ends relative to the dominant 3’UTR CAGE peak per transcript (Figure 1B - in blue, and S1G). Both RNA-seq samples showed highly reproducible enrichments of read ends coinciding with dominant 3’UTR CAGE peaks. This reveals that the 3’UTR CAGE peaks are confirmed by the read-ends from RNA-seq data, which suggests that the signal could be originating from post-transcriptional cleavage sites. Notably, there is also a small enrichment of RNA-seq read-starts downstream from the 3’UTR CAGE peaks, which could represent the same RNA fragments detectable by the CAGE samples (Figure 1B in yellow). More importantly, these findings demonstrate that 3’UTR capped fragments identified by CAGE can also be detected by other, methodologically independent, high-throughput sequencing methods such as RNA-seq.

We next aimed to confirm the presence of transcripts initiating at the non-promoter 3’UTR CAGE peaks by an alternative experimental approach, not dependent on RNA library creation or high-throughput sequencing. We focussed on two genes, *CDKN1B* and *JPT2,* which showed a single strong 3’UTR CAGE peak and highly reproducible read coverage for CAGE and RNA-seq in both K562 and HeLa cells (Figure S1J). Two separate sets of primers were designed upstream and downstream the 3’UTR CAGE peak (see Methods) to quantify transcripts containing these regions. In agreement with CAGE and RNA-seq data (Figure S1J), RT-qPCR detected higher levels of these transcripts with the 3’ downstream primers (Figure S1H), suggesting an accumulation of abundant 3’UTR fragments.

Treatment of the samples with Terminator_TM_ 5’-Phosphate-Dependent Exonuclease (TEX), an enzyme capable of degrading uncapped RNAs, had none or little effect in the amount of *JPT2* and *CDKN1B* transcript detected either side of the 3’UTR CAGE peak within these cells. This was in sharp contrast with the known uncapped 3’ fragment of *SLC38A2* mRNA, previously described by Malka et al. ^7^, which was, as expected, sharply reduced upon TEX treatment (Figure 1C), thus further demonstrating that our studied 3’UTR fragments are capped.

We further confirmed that 3’UTR-derived RNAs can be detected by long-read Nanopore-sequencing CAGE. We were provided with such data from 10 genes in iPSC, neuron stem cell (NSC) and Cortical Neuron samples by the FANTOM6 consortium that contain HeLa and K562 3’UTR CAGE peaks (Figure S1K). In all of the 10 examples, the full length read sequencing CAGE identified highly abundant reads spanning from the start of our identified CAGE 3’UTR peaks till the end of the annotated transcripts (Figure S1K). This further confirms that the 3’UTR-derived RNAs originate from the full length mRNA. Altogether, these analyses confirm the presence of abundant, capped 3’UTR-derived RNAs that originate from cleavage of the full-length mRNAs.

### Capped 3’UTR-derived RNAs are predominantly cytoplasmic

Next, we asked if there is evidence of nuclear Cap Binding Complex (CBC) binding to the capped 5’ ends of 3’UTR fragments, as it is known to bind to 5’ ends of nascent protein-coding mRNA transcripts in the nucleus. Individual-nucleotide resolution UV crosslinking and immunoprecipitation (iCLIP) is a method that identifies protein-RNA crosslinking interactions with nucleotide resolution in a transcriptome-wide manner. We examined CBC-iCLIP data from HeLa cells, where the authors targeted nuclear cap-binding subunit CBP20 protein ^21^. CBP20 is a nuclear component of cap-binding complex (CBC), which binds co-transcriptionally to the 5’ cap of pre-mRNAs and interacts directly with the m7-G cap ^22, 23^. The CBP20 RNA binding data was analysed using standard iCLIP processing pipeline, where the nucleotide preceding cDNA-start position after PCR duplicate removal is reported as the crosslinking position (see Methods). The CBP20 crosslinking positions were then summarised across all dominant 5’UTR and 3’UTR CAGE peaks per transcript. As expected, CBP20 crosslinks were enriched around the dominant 5’UTR CAGE peaks where the TSS of full-length transcripts is positioned. However, the enrichment was very weak at the non-promoter 3’UTR CAGE peaks (Figure S2A). This strongly suggests that the 3’UTR capped fragments identified by CAGE are not part of nuclear CBC, but are likely a product of an independent post-transcriptional processing pathway.

Additionally, we analysed capCLIP data from HeLa cells. capCLIP is a version of CLIP that targets translation elongation factor eIF4E, a cytoplasmic protein which binds the 7-methyl-GTP moiety of the 5′-cap structure of RNAs for the efficient translation of almost all mRNAs ^24, 25^. The capCLIP data was analysed in the same way as CBP20-iCLIP. The enrichment of capCLIP signal at the non-promoter 3’UTR CAGE peaks was much stronger than in the CBC-iCLIP (Figure S2A, S2B), which suggests that the cap of the 3’UTR-derived RNAs is primarily bound by the cytoplasmic eIF4E, rather than the nuclear cap binding protein CBP20, suggesting that these RNAs are predominantly cytoplasmic.

### 5’ ends of 3’UTR-derived RNAs are enriched for G-rich motifs and strong secondary structures

Next, we wished to understand the sequence features that distinguish the CAGE peaks corresponding to co-transcriptional capping of transcription start sites from those originating from post-transcriptional capping of 3’UTR-derived RNAs. We first explored the possibility that 3’UTR fragments might be a side-product of nuclear polyadenylation and associated endonucleolytic cleavage. In this case, the identified 3’UTR CAGE peaks should be preceded by enrichment of the canonical polyA A[A/U]UAAA hexamers, which recruit the nuclear polyadenylation machinery. However, we only found such enrichment at the annotated 3’UTR ends, and not upstream of the 3’UTR CAGE peaks (Figure S2C). We observed a notable enrichment downstream from the 3’UTR CAGE peaks (Figure S2C - red line), most likely because some of the 3’UTR-derived RNAs are relatively short and their 5’ ends are close to the annotated 3’UTR ends.

Next, we examined whether there was any other distinguishing difference between the two types of CAGE peaks. Consistent with previous studies ^8, 12^, we detected a strong G-enrichment around the 5’ end of the CAGE reads present in non-promoter regions (Figure 2A, S2D), distinct from YR dinucleotide which is a feature of initiator signal at 5’ ends of genes. More surprisingly, CAGE peaks within the 3’UTR region showed a strong increase in internal pairing probability (see Methods: Secondary structure) in comparison to other regional groups (Figure 2B, S2E), suggesting structural preference is important for 3’UTR-derived RNAs. Notably, the surrounding region of CAGE peaks in 5’UTRs is more structured (light blue line in Figure 2B, S2E), which could be explained by the higher GC content that is present around all 5’UTRs in vertebrates ^26^, with a distinctive drop at -25 bps coinciding with the canonical TATA box position.

**Figure 2.**
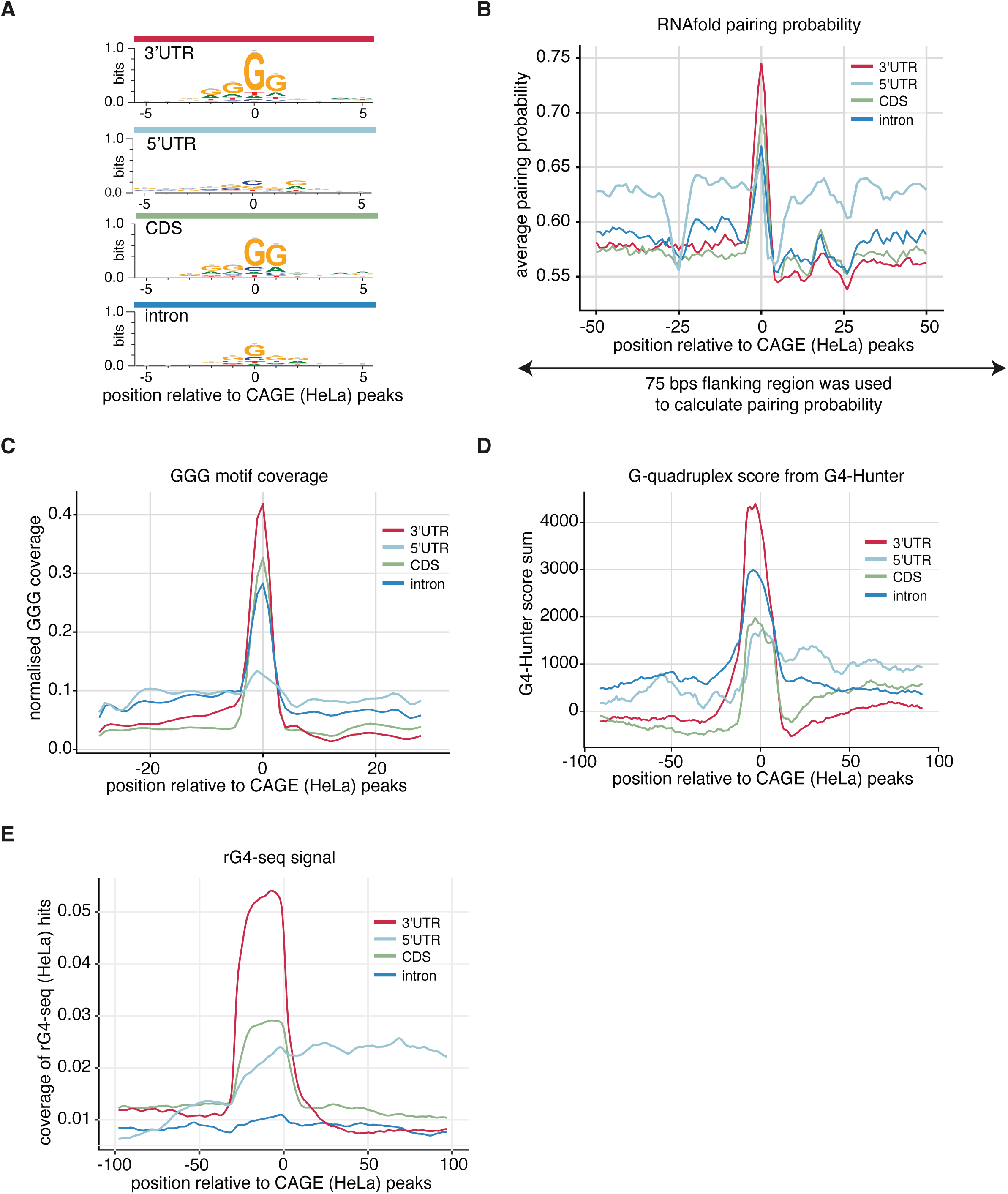
5’ ends of 3’UTR-derived RNAs are enriched for G-rich motifs, strong secondary structures and RNA-G-quadruplexes. (A) The sequence logos around CAGE (HeLa) peaks across the transcriptome regions. (B) The 75 nt region centred on CAGE (HeLa) peak to calculate pairing probability with the RNAfold program, and the average pairing probability of each nucleotide is shown for the 50 nt region around CAGE peaks. (C) GGG-motif enrichment relative to CAGE (HeLa) peaks. (D) Summarised score from G4-Hunter prediction tool in the region of 50 nts upstream and downstream relative to CAGE (HeLa) peaks. (E) Enrichment of RNA-G-quadruplex sequencing (rG4-seq) hits from HeLa cells relative to CAGE (HeLa) peaks.

### 3’UTR CAGE peaks coincide with RNA-G-quadruplexes and heavily structured regions

Motifs with G-rich repeats in the transcriptome can form non-canonical four-stranded structures (G4s) implicated in transcriptional regulation, mRNA processing, the regulation of translation and RNA translocation ^27^. Similar to web-logo motif analyses of CAGE peaks from different mRNA regions (Figure 2A, S2D), the nucleotide enrichment plot of GGG sequences showed the highest enrichment around 3’UTR CAGE peaks (Figure 2C, S2F). This suggests that the sequence around the 3’UTR CAGE peaks may show increased propensity to form RNA-G4 structures via the canonical G4 motif (GGG-{N-1:7}(3)-GGG) ^28^. To further explore the RNA G-quadruplexes formation profile, we integrated RNA-G-quadruplex sequencing (rG4-seq) data from HeLa cells ^29^ and ran G4-Hunter predictions ^30^ around CAGE peaks.

Indeed, both the rG4-seq data (HeLa) and G4-Hunter predictions (K562) showed the highest G4s enrichment in the 3’UTR region relative to CAGE peak (Figure 2D, S2G, 2E) with the highest number of G4s present in 3’UTRs (Figure S2H). Moreover, in 8 out of 10 examples with dominant 3’UTR CAGE peaks across multiple samples we identified rG4-seq clusters coinciding with 3’UTR CAGE peaks (Figure S1K).

Interestingly, beside the G4 preference, the top 3’UTR CAGE overlapping peak in both HeLa and K562 cell lines overlaps MALAT1-associated small cytoplasmic RNA (mascRNA, Figure S1E), which is extremely abundant, widely conserved among mammals, and is known to be upregulated in cancer cell lines ^31^. Notably it forms a triple helix structure at its 3′ end that makes it more stable from the rest of the ncRNAs ^32^. Overall, these results suggest that strong structures around 3’UTR CAGE peaks, including RNA-G4s, could play an important role in stabilising these RNA fragments. One possibility would be by making these fragments exoribonuclease-resistant ^33^ or by causing XRN1 to stall during 5’-3’ degradation ^34^.

### 3’UTR cleavage sites are flanked by enriched UPF1 binding

Based on the evidence outlined, we hypothesised that the capped 3’UTR-derived RNAs are formed post-transcriptionally. On that assumption, we aimed to determine whether specific RNA-binding proteins (RBPs) were involved in the process. To that end, we analysed publicly available enhanced CLIP (eCLIP) data for 80 different RBPs in the K562 cell line, produced by the ENCODE consortium ^35^. For each RBP, we calculated normalised cross-linking enrichment compared to other RBPs around maximum CAGE peak per annotated gene region (5’UTR, CDS, intron, 3’UTR). This identified a specific set of RBPs around CAGE peaks, with UPF1 (Regulator of nonsense transcripts 1) protein as the top candidate in 3’UTRs, and DDX3X (DEAD-Box Helicase 3 X-Linked) in 5’UTRs (Figure 3A, S3A). No specific RBP enrichment was detected in CDS and intronic regions. The DDX3X enrichment around 5’UTR CAGE peaks was no surprise since it is known to be involved in transcriptional process by interacting with transcription factors, in pre-mRNA-splicing by interacting with Spliceosomal B Complexes, and in RNA export by interacting with Cap-Binding-Complex (CBC) ^36^. UPF1 is a known factor of the Nonsense-Mediated Decay (NMD) pathway, where stalled UPF1 at CUG and GC-rich motifs activates its mRNA decay^37,38^.

**Figure 3.**
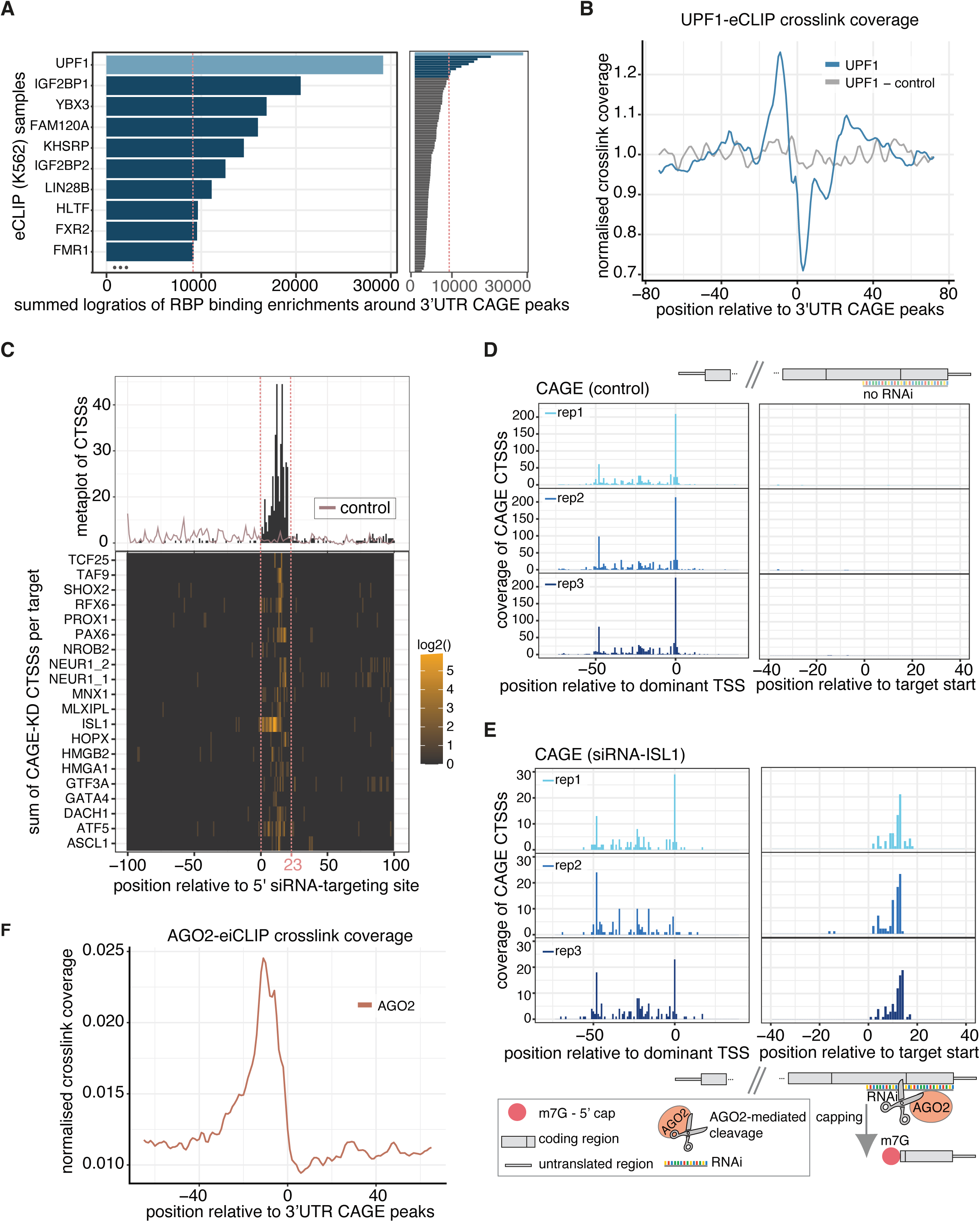
Enrichment of RNA binding proteins (RPB) at 3’UTR cleavage sites and capping at small interfering RNAs target sites. (A) Enrichment of eCLIP cross-linking clusters surrounding 3’UTR CAGE peaks from 80 different RBP samples (right panel) in K562 cells from ENCODE database using sum of log ratios. The red line represents the threshold of top 10 RBP targets (left panel). (B) RNA-map 66 showing normalised density of UPF1 crosslink sites relative to 3’UTR CAGE peaks and random positions of the same 3’UTRs as control. (C) Enrichment of CAGE transcription start sites (CTSS) relative to 5’ sites of small interfering RNAs for 20 different CAGE-KDs and merged control samples. (D) Enrichment of CAGE transcription start sites (CTSS) relative to dominant transcription start site (TSS) (left panel) and relative to 5’ of small interfering RNA of ISL1 target (right panel) for CAGE control samples with 3 replicates. (E) Enrichment of CAGE transcription start sites (CTSS) relative to dominant transcription start site (TSS) (left panel) and relative to 5’ of small interfering RNA of ISL1 target (right panel) for CAGE-ISL1-KD samples with 3 replicates. (F) RNA-map showing normalised density of eiCLIP-AGO2 crosslink sites relative to 3’UTR CAGE peaks.

Interestingly, the crosslinking enrichment of UPF1 around 3’UTR CAGE peaks is positioned just upstream from the peaks, followed by depletion downstream (Figure 3B). Moreover, there was a high correlation between the 3’UTR CAGE signal and UPF1 binding, which was not correlated with gene expression or 3’UTR length (Figure S3B), indicating that the 3’UTR CAGE signal could result from the post-transcriptional capping of NMD-mediated RNA degradation by-products ^8, 13^. This correlation could also be related to G-enrichment since 3’UTRs with UPF1 bindings are prone to having higher than average G content ^39^. More specifically, the strength of UPF1 binding coincides with the strength of the 3’UTR CAGE peaks and proximity to the peaks (Figure S3C), which suggest that the precise binding position of UPF1 relative to the cleavage/capping position could be important for the formation of these fragments. All in all, we find that 3’UTR-derived RNAs are not a simple by-product of high mRNA decay, but also find that the NMD factor UPF1 might regulate their generation.

### mRNA cleavage by small interfering RNAs generates newly capped RNA fragments

An alternative way in which mRNAs can be cleaved post-transcriptionally is through RNA interference (RNAi). Indeed, a common way to artificially accomplish gene silencing is to utilise small interfering RNAs (siRNAs) to induce endonucleolytic degradation of the target transcripts ^40, 41^. SiRNAs are usually 21-23 nt long and their sequence corresponds to an antisense mRNA target sequence. Silencing by siRNAs is mediated thanks to the RNAse III catalytic activity of Argonaute 2 (AGO2), a subunit of the RNA-induced gene-silencing complex (RISC) in the cytoplasm.

We hypothesised that siRNA silencing following AGO2 cleavage could lead to cytoplasmic capping of the cleaved RNA fragments instead of degradation. To follow up this hypothesis, we first investigated if CAGE could detect cleaved RNA fragments guided by siRNA. We analysed CAGE siRNA KDs from FANTOM5 dataset ^42^, for which we collected 28 samples with siRNA targeting sequences (20 siRNAs designed by ThermoFisher and 8 by the study authors) and 5 control samples. Surprisingly, in 20 of the 28 samples we detected CAGE 5’ end signal corresponding to the exact siRNA complementary genomic sequence in at least two replicates (Figure 3C).

The strongest enrichment of CAGE signal relative to the siRNA target start site was detected in the *ISL1*-KD sample, supported by all 3 biological replicates, and with no signal detected in control samples (Figure 3D,E, S3D). More interestingly, the dominant CAGE 5’ end signal was present in the middle of the siRNA target sequence (Figure 3E, S3D), where the AGO2 cleavage is known to take place ^43, 44^. Also, the TSS CAGE signal in 5’UTR of the corresponding protein-coding gene dropped by ∼75% compared to the control samples in all 3 replicates (Figure 3D,E), confirming that the KD of ISL1 transcript was efficient. Together these results indicate that siRNA mediated recruitment of AGO2 can lead to the generation of post-transcriptionally capped RNA fragments following mRNA cleavage.

### 3’UTR CAGE peaks coincide with AGO2 binding and RNA-G-quadruplexes

Since the endonuclease activity of AGO2 facilitates mRNA cleavage guided by siRNAs, we investigated if AGO2 binds also at the endogenous 3’UTR CAGE peaks. There was no publicly available AGO2 binding data for either HeLa or K562 cells, so it could not be detected in the analysis in Figure 3A. For that reason, we produced ‘enhanced individual nucleotide resolution’-CLIP (eiCLIP) ^45^ data for AGO2 (AGO2-eiCLIP) for HeLa cells, using the same pipeline with small adjustments (see Methods) as for UPF1-eCLIP. Analysis revealed 32.8% of crosslinking positions mapped to the 3’UTR region (Figure S3E), with a higher binding enrichment in known miRNA-regulated transcripts, and a clear miRNA-seed sequence enrichment downstream from the crosslinking site (Figure S3F,G). Similarly to UPF1, AGO2 crosslinks were enriched just upstream from the 3’UTR CAGE peaks; unlike UPF1, they were not depleted downstream of them (Figure 3F, S3H, Figure 3B).

In animals, endogenous RNAi is mainly mediated by microRNAs (miRNAs). MiRNAs are also ∼21-23 nucleotide (nt) long RNAs, but, in contrast to siRNAs, miRNAs recruit the miRNA induced silencing complex (miRISC) containing AGO1-4 to mRNAs with partial complementarity which results in translational repression and/or exonucleolytic cleavage ^40, 41^. To see if there is evidence compatible with AGO2 miRNA-guided cleavage of mRNA targets, which would hence be similar to the siRNA method of action, we first mapped reverse complements of miRNA sequences to the human genome, allowing 2 mismatches, to identify putative miRNA matches in 3’UTRs which could be mediated by miRNA-dependent endonucleolytic cleavage ^46, 47^. We identified 29 such targets but there was no CAGE signal present around them (data not shown). Accordingly, we instead explored the binding specificity of AGO2-eiCLIP data, and performed a motif analysis using HOMER motif finder. When analysing the 15 bp flanking region around AGO2-crosslinking peaks (see Methods), one of the most prominent motifs was highly enriched in Gs (Figure S3I - 2nd and 3rd). Notably, this also agrees with one of the first AGO2-CLIP studies performed on mouse embryonic stem cells, where the authors showed that, without the miRNA present, AGO2 binds preferentially to G-rich motifs ^48^. This suggests that miRNA-directed recruitment may not be necessary for AGO2 binding at the site of cleavage that generates 3’UTR-derived RNAs.

Our results demonstrate the AGO2 binding near 3’UTR CAGE peaks. Accordingly, given our previous observation that RNA-G-quadruplexes were enriched around 3’UTR-derived RNAs, we investigated whether AGO2 could be attracted by RNA-G-Quadruplexes in general. We first aligned AGO2-eiCLIP-HeLa cross-linking positions relative to 3’ end of rG4-seq-HeLa sites in different regions of primary transcripts. Similar to the 3’UTR CAGE peaks enrichment by RNA-G-Quadruplexes, AGO2 crosslink-binding sites are much more highly enriched at rG4-seq sites in the 3’UTRs relative to 5’UTRs, introns and coding sequence (Figure S3J,K). The mechanistic implications of the overlap between RNA-G4 structures and AGO2 binding close to the capped 3’UTR-derived RNAs remain to be experimentally interrogated.

### 3’UTR-derived RNAs could explain the previously reported regulation of APP protein

Additionally, we looked if there are any known RNA-G4s with regulatory features in 3’UTRs. Notably, there has been a proposed involvement of RNA-G4 in the regulation of amyloid precursor protein (APP) in Alzheimer’s disease ^49, 50^, where G4 motif in the 3’UTR of APP mRNA was found to suppress overproduction of APP protein, but the underlying mechanism remained unclear ^51^. Analysis of rG4-seq and CAGE data from HeLa cells in APP mRNA showed that the CAGE peak coinciding precisely with the 3’ end of the same G4 motif that was previously found to affect APP protein production (Figure S1K - APP, Figure 2E). Moreover, long-read sequencing CAGE from iPSC, neuron stem cell (NSC) and Cortical Neuron samples, confirmed that abundant 3’UTR-derived RNAs are present in all of these samples that start at the position of our identified CAGE peak and span till the end of the annotated APP gene (Figure S1K - APP). This indicates that the RNA-G4 very likely affects the production of the 3’UTR-derived RNAs, and that this mechanism accounts for its effect on the APP protein production.

### Capped 3’UTR fragments of CDKN1B and JPT2 transcripts do not co-localise with the parental mRNAs

Finally, we were interested to explore potential implications of 3’UTR derived RNAs. If a transcript is cleaved, it is possible for the two resultant RNA fragments to localise either together or independently from each other. To test this, we designed smFISH probes to simultaneously image the RNA upstream and downstream of the proposed post-transcriptional cleavage and capping site in CDKN1B and JPT2 using hybridisation chain reaction RNA-fluorescence in situ hybridization (HCR-FISH 3.0) ^52^. To account for the technical biases in detection, we also designed probes against the coding sequence (hereafter upstream) and 3’UTR (hereafter downstream) of a control mRNA, PGAM1, which does not contain CAGE peaks in the 3’UTR and contained a similar 3’UTR length to our targets.

We performed HCR-FISH in HeLa cells to determine whether putative 3’UTR-derived RNAs can be found independently of the RNA upstream of the cleavage site (Figure 4A, B). In the control transcript, PGAM1, we observed that 17.3% of upstream signals did not have a colocalising downstream signal and 21.3% of downstream signals did not have a colocalising upstream signal (Figure 4C, S4A). Interestingly though, the mRNAs that contain a 3’UTR CAGE signature were significantly more likely to show independent signals from the RNA downstream of the proposed cleavage site (CDKN1B: 53.3%, p adj. < 0.05; JPT2: 52.3%, p adj. < 0.05; Figure 4C). In the case of JPT2, we also observed significantly more independent signals from the upstream probes (29.3%, p adj. < 0.05; Figure 4C). These observations are consistent with the existence of cleaved 3’UTR fragments in the cell, and they reveal that these products may localise differently from their host transcripts.

**Figure 4.**
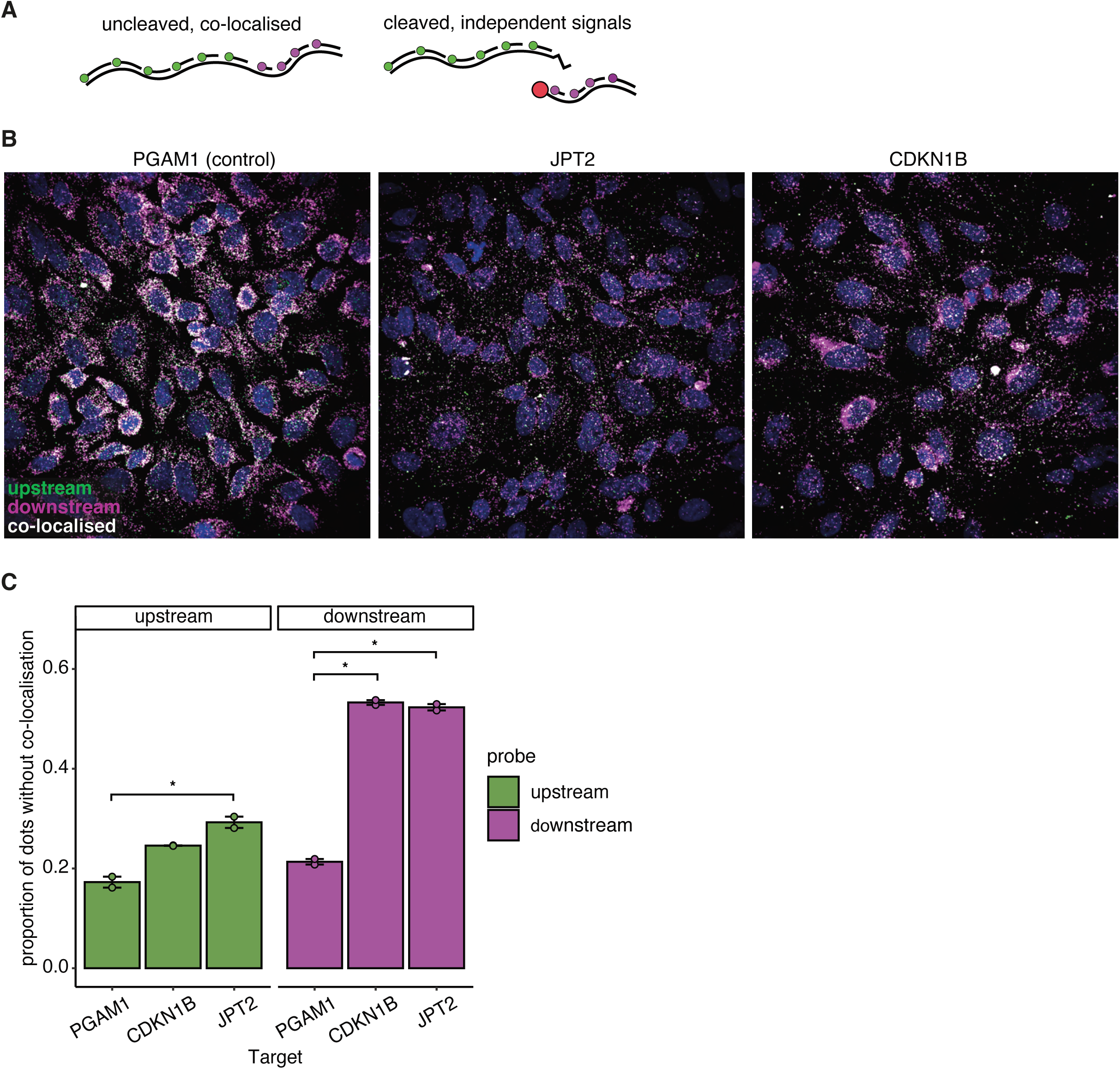
Capped 3’UTR fragments of CDKN1B and JPT2 transcripts do not co-localise with the parental mRNAs. (A) Schematic representation of probe design for HCR-FISH microscopy to separate CDS (green) and 3’UTR (purple) regions as cleaved, independent signals and uncleaved, co-localised signals. (B) Representative example of HCR-FISH Microscopy imaging for PGAM1-control, JPT2 and CDKN1B. Signal from CDS probes is shown in green and signal from 3’UTR probes is shown in purple, with colocalising signals appearing white. (C) Proportion of signal for each CDS or 3’UTR probe without a detected colocalising signal from the opposing probset (3’UTR or CDS, respectively). Error bars represent standard error. * p (adjusted) < 0.05.

## Discussion

We employed a combination of computational analyses of high throughput sequencing datasets from human cell lines that reveal capped 5’ ends of RNAs genome-wide (CAGE) and binding sites for dozens of RNA binding proteins. As the main resource we used large publicly available datasets from consortiums such as ENCODE (Encyclopedia of DNA Elements) and FANTOM together with new computational approaches and experimental validations. We identified several factors that show strong binding enrichment at the sites where 3’UTRs are cleaved to generate the capped 3’UTR-derived RNAs. Specifically, we compared the crosslinking enrichment of several RBPs including UPF1 and AGO2, which are both highly enriched around 3’UTR-derived RNAs beside RNA-G-Quadruplexes and heavily structured regions.

Other studies suggested that these capped RNAs originate as a consequence of incomplete degradation of the mRNA during the standard processes of mRNA decay ^6, 15, 17^, which would agree with the enrichment of RNA-seq read-starts at 3’UTR CAGE peaks (Figure 1B). However, it is challenging to use traditional fragmented-based sequencing methods such as RNA-seq and CAGE-seq for discovery and validation of 3’UTR-derived RNAs, because the reads of derived RNAs can not easily be distinguished from the parental mRNAs, and the only information available is the enrichment of starts of RNA-seq reads with the positions of capping detected by CAGE-seq. This could explain why most 3’UTR-derived RNAs have so far remained undetected. We now used multiple lines of evidence to complement CAGE and RNA-seq data, including long-read Nanopore sequencing, analysis of RNA structural features and RBP interactions relative to the cleavage of 3’UTR-derived RNAs, and HCR-FISH imaging. We show that the position and strength of capping is closely linked to RNA structure and RBP binding, and that the 3’UTR-derived RNAs are often abundant and do not co-localise with the parental mRNAs.

### Cytoplasmic capping of the siRNA-targeting cleaved fragments

In siRNA-KD CAGE samples we noticed that certain cleaved fragments which are involved in post-transcriptional cleavage processing, such as RNAi targeting, can form capped RNA fragments (Figure 3C,E). However, with the available data we can not quantify the efficiency of such capping, or identify all the factors that might be involved in the process. Understanding the cytoplasmic capping of cleaved fragments and their abundance will also give important insights for understanding viral RNA capping. Since the majority of RNA capping happens in the nucleus, viruses evolved to produce efficient capped RNAs in the cytoplasm by encoding their own capping machinery, or by or taking a capped 5’ fragment from the host’s mRNA, also known as cap snatching ^1^. Moreover, many new drugs which are based on RNAi and miRNA targeting are already in use or under active clinical trials for treating neurological or viral diseases, and in cancer treatments. Side products of the targeted mRNAs from these therapeutic drugs could still be subjected to a cytoplasmic capping mechanism and result in unwanted toxic side effects.

### The role of UPF1, AGO2 and RNA-G-Quadruplexes in capping 3’UTR-derived RNAs

On average 3’UTRs are shorter in cancer cells to evade miRNA-mediated repression ^53^ but we could not see any correlation between 3’ length and intensity of the CAGE signal (Figure S3B). It is known that UPF1 binds to GC-rich motifs in 3’UTRs ^37^, but it is still not known what the main trigger of UPF1-mediated mRNA decay is. Another study found evidence that a G content enrichment in 3’UTRs plays a more important role on mRNA destabilisation by inserting UPF1 binding motifs into non-UPF1 targets ^37^. This suggests that G enrichment plays a vital role in triggering UPF1-mediated mRNA decay. Meanwhile, significant overlaps in binding between UPF1 and AGO2 have been reported but with an unknown functional relationship ^39^. In the same study they also discovered preferential UPF1 binding in structured G-rich regions ^39^. We do not know if G enrichment is needed for the post-transcriptional capping process, but it has been shown in a previous study ^48^ that G enrichment could be important for AGO2 binding in the absence of miRNA guidance; it is also important for the RNA structure to form the hairpin-loop structure, which is necessary for AGO2 cleavage ^54^. Another recent study demonstrated 3’UTR cleavage site in rat cervical ganglion neurons, which are also cleaved post-transcriptionally but only expressed in axons and not in cell bodies, with AGO2 and UPF1 as the top two RBP targets ^55^. However, we now find that AGO2 binds to RNA-G4s in 3’UTRs (Figure S3J,K), but their functional relation remains unknown. Those AGO2 binding sites are less likely to be guided by miRNA, since it has been shown that RNA-G4s can also prevent miRNA binding from its target sites ^56^.

Moreover, RNA-G4s are known to form stable structures in vitro, but recent studies have suggested that they may be less stable in vivo due to active unwinding by RNA helicases ^57, 58^. However, Kharel et al. (2022) ^59^ have demonstrated that 3’UTR-G4s are dynamically regulated under cellular stress conditions and may play a positive role in mRNA stability for several transcripts, including the 3’UTR-G4 in APP (Figure S1K - APP). While the stability of RNA-G4s around 3’UTR cleavage sites is still unknown, our analysis shows that these sequences have a strong pairing probability and are capable of forming G4 structures (Figure 2B, S2E, 2D, S2G, 2E), suggesting a direct contribution to the formation of 3’UTR-derived RNAs.

### Capped 3’UTR fragment no longer co-localise in the cell with the rest of the mRNA in CDKN1B and JPT2

First, we show that 3’UTR-derived RNAs in CDKN1B and JPT2 are capped and highly expressed in the cell using qPCR primers (Figure 1C,S1H). Next, we produced additional HCR-FISH-probe experiments for JPT2 and CDKN1B targets, which demonstrate that 3’UTR-derived RNAs can be abundant in the cytoplasm without co-localising with parental mRNAs. In agreement with the CAGE data, the highest ratio of ∼2-fold of downstream vs. upstream probes is present in CDKN1B, where the 3’UTR CAGE peak is higher than the peak at the TSS (Figure 4A,B,C, S1I - CDKN1B).

Interestingly, some cells showed a strong perinuclear accumulation of 3’UTR-probes in CDKN1B (Figure 4B), whilst the signal was spread throughout the cytosol for most cells. An interesting possibility is a cell cycle-dependence, as it has already been observed for other aspects of regulation of p27 gene expression, including mRNA translation ^60, 61^. The role of these capped 3’UTR clusters in *CDKN1B* could also be related to cell cycle specific regulation of CDKN1B/p27kip1 (p27) protein expression. Two studies have demonstrated that rescue of splicing deficiency in CDKN1B improves protein production and leads to cell cycle arrest ^62, 63^. Additionally, another study suggested that high levels of 3’UTRs of NURR1 in proliferating cells could also be linked to cell cycle dynamics ^16^. It would be important to investigate further the dynamic nature of these isolated 3’UTRs and their impact on cellular functions in a cell cycle dependent manner.

### Methodological implications

Different abundance and localisation of 3’UTR derived RNAs relative to their parent transcript and the 5’ cleavage fragment containing protein coding sequence suggests that 3’UTR fragmented-based sequencing methods might be measuring the wrong RNA species in a significant proportion of cases. Even in the case of RNA-seq, quantitating the signal across the entire length of the uncleaved mRNA might measure a combination of protein-coding and non-coding RNA species. To increase the accuracy of quantitation of protein coding transcript levels, as well as those of 3’UTR-derived RNAs themselves, it may be necessary to develop new computational quantitation methods informed by the results of this paper, which will try to estimate the levels of protein-coding and 3’UTR fragments separately.

### Limitations

At the moment we do not have the ability to identify the full-length size of 3’UTR-derived RNAs since the main technique that we used (CAGE-seq) is based on 5’ end sequencing. Also, CAGE-seq method has a limitation on fragment size similar to other HT-sequencing methods with a minimum fragment size of 200 bps, which the long-read CAGE can overcome (Figure S1K). Using only experimental datasets has limitations in coverage, organisms, cell lines and can increase the number of false positives, which can result from experimental limitations and background noise. Developing computational methods to model these capped 3’UTR-derived RNAs genome-wide would be important for future studies, especially when applying it to other cells and organisms for which we do not have such large available datasets. Additionally, designing ideal HCR-FISH-probes that would span just over the 3’UTR capped region to detect uncleaved 3’UTRs would be very challenging because of the sequence space limitation and highly structured RNA which prevents probes from hybridising ^64^.

## Conclusions

3’UTR-derived RNAs are emerging as novel regulatory molecules, with potential implications in broad cellular processes such as cell cycle or neuronal homeostasis ^16, 17, 65^. However, the molecular mechanisms involved in the generation of these RNA species had been largely unknown. Our results shed light into these mechanisms showing that 3’UTR-derived RNAs are stabilised through strong structures including RNA-G-Quadruplexes, and both UPF1 and AGO2 play a key role in this regulation. Overall, our findings provide the framework for further investigations where their functions will surely emerge.

## Acknowledgments

This work was funded in part by The Wellcome Trust grants (106954/Z/15/Z) awarded to B.L., by the Medical Research Council (MRC) (MR/P023223/1) to A.M-S. and by (215593/Z/19/Z) to J.U., Medical Research Council (MRC) Core Funding (MC-A652-5QA10), by the Imperial College Research Fellowship awarded to N.H., and by the Francis Crick Institute which receives its core funding from Cancer Research UK (CC0102), the UK Medical Research Council (CC0102), and the Wellcome Trust (CC0102), and by an institutional budget from RIKEN, MEXT (Ministry of Education, Culture, Sports, Science and Technology) and from institutional budget from the Human Technopole. We thank the Crick Advanced Light Microscopy facility, especially Donald Bell, for their support. We thank Sarvesh Nikumbh for help with the CAGEr tool and other members of Lenhard’s group for helpful discussions and comments on the manuscript. For the purpose of Open Access, the author has applied a CC BY public copyright licence to any Author Accepted Manuscript version arising from this submission.

## Author contributions

N.H. and B.L. conceived the study. N.H. designed the experiments, analysed the data, and led the project. R.C. and A.M-S. performed and designed qPCR experiments. AGO2-eiCLIP was designed and performed by C.R.S and A.M-S. R.F. and H.D. designed and performed smFISH probe imaging supervised by J.U. Long-read CAGE examples were produced, processed and provided by C.P., K.Y., T.K., C.W.Y., M.K., H.T., P.C. N.H. wrote the manuscript, with contributions from R.F., H.D., A.M.J, S.V., C.R.S., J.U., A.M-S. and B.L.

## Figure legends

Visual Abstract Figure: Schematic representation of how 3’UTR-derived RNAs can stabilise through strong secondary structures including RNA-G-Quadruplexes and structure-specific RBP interactions such as AGO2.

**Figure S1: Related to Figure 1**

A. Pearson’s correlation of raw CAGE tag counts per TSS or consensus cluster, demonstrating high reproducibility across biological replicates and high correlation across different cell types..
B. Reverse cumulative distribution of CAGE tags after normalisation ^67^.
C. Total number of CAGE tags in each sample.
D. Percentage of CAGE tags per transcriptome region using Random primers, Oligod-T primers, and combination of both primers (1:4 oligod(T):Random Primers).
E. Pearson’s correlation between the replicates and different cell line samples in 3’UTRs.
F. Pearson’s correlation between the replicates and different cell line samples in 5’UTRs.
G. The 3’ of downstream paired read (blue line) and 5’ of upstream paired read (yellow line) of RNA-seq (HeLa) normalised coverage around the 3’UTR CAGE (HeLa) peaks followed by schematic representation of paired-end read positioning.
H. RT-qPCR data of gene expression using primers designed to amplify sequences located downstream (3’C), upstream (5’C) and overlapping (AC). Data represents fold detection using downstream *versus* upstream/overlapping primers relative to the 3‘UTR CAGE peaks. Primer target sequences relative to the 3’UTR CAGE peak are visualised using IGV genome browser.
I. Top gene examples with dominant 3’UTR CAGE peaks present in K562 and HeLa cell lines using IGV-genome browser.
J. Visualisation of RNA-seq reads relative to dominant 3’UTR CAGE peaks in CDKN1B and JPT2 gene using IGV-genome browser.
K. Visualisation of 10 gene examples with 3’UTR CAGE peaks (HeLa, K562), rG4-seq clusters (HeLa) and long-read CAGE (iPSC, neuron stem cell and Cortical Neuron) reads using IGV-genome browser.

**Figure S2: Related to Figure 2**

A. RNA-map showing normalised density of CBP20-iCLIP (HeLa) crosslink sites relative to dominant 3’UTR and 5’UTR CAGE peaks.
B. RNA-map showing normalised density of cap-CLIP (HeLa) crosslink sites relative to dominant 3’UTR and 5’UTR CAGE peaks.
C. Normalised motif enrichment of canonical PolyA motifs relative to 3’UTR ends and to the dominant 3’UTR CAGE peaks.
D. The composition of genomic nucleotides around CAGE (K562) peaks across the transcriptome regions.
E. The 75 nt region centred on CAGE (K562) peak to calculate pairing probability with the RNAfold program, and the average pairing probability of each nucleotide is shown for the 50 nt region around CAGE peaks.
F. GGG-motif enrichment relative to CAGE (K562) peaks.
G. Summarised score from G4-Hunter prediction tool in the region of 50 nts upstream and downstream relative to CAGE (K562) peaks.
H. Percentage of G-seq sites per transcriptome region.

**Figure S3: Related to Figure 3**

A. Enrichment of eCLIP cross-linking clusters surrounding 5’UTR CAGE peaks from 80 different RBP samples (right panel) in K562 cells from ENCODE database using sum of log ratios. The red line represents the threshold of top 10 RBP targets (left panel).
B. Pearson’s correlation between the 3’UTR CAGE (K562) tags and RNA-seq (K562) read coverage per gene (top-left). Pearson’s correlation between the 3’UTR crosslink coverage of UPF1-eCLIP (K562) and 3’UTR CAGE (K562) tags (top-right). Pearson’s correlation between the 3’UTR length and 3’UTR crosslink coverage of UPF1-eCLIP (K562) (bottom-left). Pearson’s correlation between the 3’UTR length and 3’UTR CAGE (K562) tags (bottom-right).
C. UPF1-eCLIP (K562) crosslink enrichment relative to the distance from the 3’UTR CAGE (K562) peaks.
D. Visualisation of CAGE transcription start sites (CTSS) relative to dominant transcription start site (TSS) and relative to 5’ of small interfering RNA of ISL1 target (in red) for CAGE-ISL1-KD and CAGE-control samples with 3 biological replicates using IGV-genome browser.
E. Percentage of AGO2-eiCLIP (HeLa) sites per transcriptome region.
F. Binding enrichment of AGO2-eiCLIP (HeLa) relative to miRNA-regulated transcripts and non-miRNA-regulated transcript in HeLa (data from ^68^).
G. Heatmap of miRNA-seed sequence enrichment in 30 nt flanking region showing the top 500 AGO2 binding sites relative to AGO-eiCLIP (HeLa) crosslink sites. Meta plot visualises the miRNA-seed sequence composition relative to the AGO2 crosslink site.
H. Heatmap of AGO2-eiCLIP (HeLa) crosslink site enrichment showing the top 500 3’UTR AGO2 targets in 100 nts flanking region relative to 3’UTR CAGE (HeLa) peaks.
I. Sequence logos and statistics of top 12 significantly enriched motifs of AGO2-eiCLIP (HeLa) binding sites using Homer for de novo motif discovery.
J. Enrichment of AGO2-eiCLIP (HeLa) cross-linking sites relative to the middle of the rG4-seq site (HeLa).
K. Heatmap for AGO2-eiCLIP (HeLa) crosslink site enrichment to show the top 500 3’UTR AGO2 targets in 100 nts flanking region relative to middle of rG4-seq (HeLa) site.

**Figure S4: Related to Figure 4**

A. Density plots showing the shortest distance per detected signal in pixels to a signal of the opposite colour. The dashed line shows the cutoff used to distinguish colocalising and non-colocalising signals.

## Tables

**Table 1:** This table contains genomic locations of 32,065 unique 3’UTR CAGE clusters across all CAGE samples from HeLa and K562 cell lines. Each sample contains a normalised value of 5’ read positions for each cluster.

## STAR Methods

### HCR-FISH Microscopy

HeLa cells (obtained from Cell Services at the Francis Crick Institute) were grown in DMEM supplemented with 10% FBS and plated into 8-well ibidi chambers. Cells were fixed for 10 minutes at room temperature using 4% paraformaldehyde/0.4% glyoxyl diluted in PBS before permeabilization overnight at –20°C in 70% EtOH. *In situ* HCR v3.0 with split-initiator probes was performed as described in ^52^, except amplification which used 30nM of each fluorescently labelled hairpin; cells were then stained with DAPI 1 µg/mL in PBS before mounting with Fluoromount-G^tm^ (Thermo Fisher). Cells were imaged on a spinning disk confocal microscope (Nikon CSU-W1 Spinning Disk) using 60x oil-immersion objective. 6 non-overlapping field z-stacks of 17 slices with 0.39um z-steps were taken per well. 8 HCR probe pairs per target were designed using the HCR 3.0 Probe Maker ^69^. Probes were designed for CDS and 3’UTR to be amplified by the B1 HCR-amplifier with Alexa594 or the B2 HCR-amplifier with Alexa674 (Molecular Technologies), respectively.

### HCR-FISH Analysis

We z-projected the images and segmented the nuclei and cytoplasms with Cellpose (v2.0.5, ^70^) using the DAPI signal and thresholded AlexaFluor594 signal. We then detected smFISH signal positions using the Fiji plugin RS-FISH (v2.3.0, ^71^). We excluded signals that fell outside of a cell mask. For each detected signal, the minimum distance to the nearest signal in the other channel was measured. Co-localisation was defined as a minimum distance of 3 or fewer pixels. We then filtered for high confidence signals with a signal intensity in the top 50% of all signals for that channel in that image. The proportion of independent signals was calculated for each replicate, and pairwise t-tests were calculated in R using the compare_means function from the ggpubr package (https://github.com/kassambara/ggpubr/) with Benjamini-Hochberg correction.

### Cell culture

K562 and HeLa cells were maintained in RPMI 1640 or DMEM medium, respectively, supplemented with 10% foetal bovine serum (FBS) and 100 U/mL penicillin/streptomycin at 37°C in 5% CO2 in a humidified incubator.

### Reverse transcription (RT) and quantitative PCR (qPCR)

K562 cells were lysed in Trizol^R^ (Thermo Fisher Scientific) and total RNA extracted as per manufacturer’s instructions. RNA was treated with RQ1 DNase (Promega) in the presence of RNasin ribonuclease inhibitors (Promega). When indicated, treatment of DNAse-treated DNA with TerminatorTM 5’-Phosphate-Dependent Exonuclease (Cambridge Biosciences) was performed in the presence of RNasin ribonuclease inhibitors (Promega) for 60 minutes at 30°C as per manufacturer’s instructions. 500 ng RNA were reversed transcribed with SuperScript III (Thermo Fisher Scientific) reverse transcriptase, following manufacturer’s instructions in the presence of 1:4 oligod(T):Random Primers (Thermo Fisher Scientific). Real-time PCR was performed using Fast SYBR™ Green Master Mix (Thermo Fisher Scientific) and specific primers designed within the proximal regions upstream or downstream of the 3’ CAGE signal identified for *CDKN1B*, *JPT2*. The sequences of the primers are:

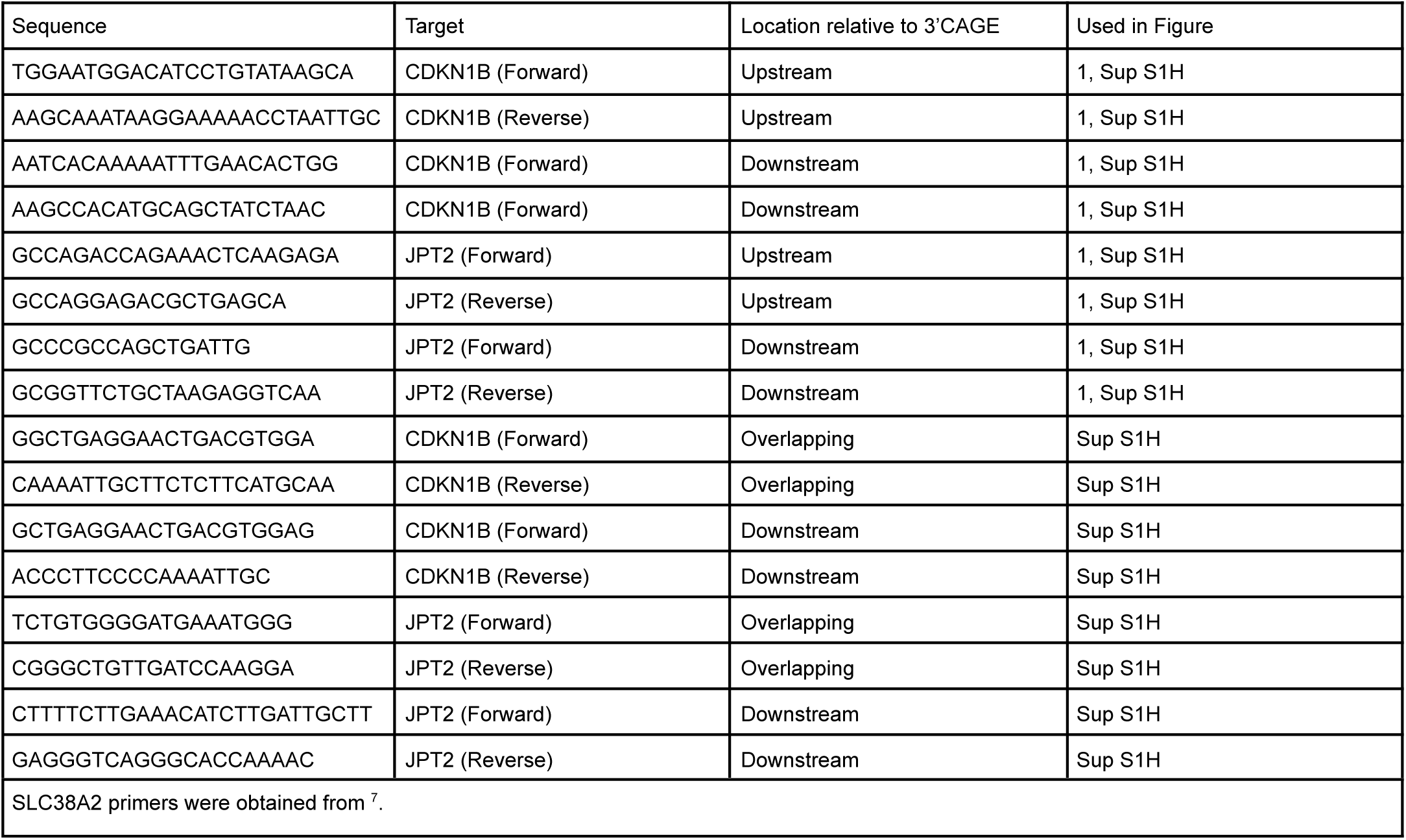

### AGO2-eiCLIP

AGO2-eiCLIP was performed as previously described ^45^. In brief, this involved following a previously described non-isotopic iCLIP workflow ^72^ which had additional modifications to enhance speed and efficiency. This included ligation of a new Cy5.5 labelled adapter (/5Phos/A[XXXXXX]NNNAGATCGGAAGAGCACACG/3Cy55Sp/) to bound RNA with high concentration T4 RNA ligase (New England Biolabs), use of RecJf exonuclease (New England Biolabs) to remove un-ligated adapter prior to SDS-PAGE analysis, reverse transcription with a biotinylated primer (/5BiotinTEG/CGTGTGCTCTTCCGA), exonuclease III (New England Biolabs) mediated removal of unextended RT-primer, cDNA capture with MyOne C1 streptavidin beads (ThermoFisher Scientific), 3’ adapter (/5Phos/ANNNNNNNAGATCGGAAGAGCGTCGTG/3ddC/) ligation instead of intramolecular ligation, and cDNA elution with nuclease and cation free water at high temperature. For a pellet of cells obtained from a 80% confluent 150mm dish, we used 100 µl Dynabeads Protein G (Thermo Fisher Scientific) conjugated to 1.5 µg anti-AGO2 antibody (MAB253, Sigma-Aldrich/ Merck). Samples of two biological replicates were sequenced with paired-end reads using NextSeq500.

### Mapping and processing of AGO2-eiCLIP

Pre-processing, mapping to hg38 gene annotation and removal of PCR duplicates of AGO2-eiCLIP data and peak calling was performed by using iMAPS (https://imaps.goodwright.com/) with default settings. Processed data was downloaded from the iMAPS in BEDgraph format where each count represents crosslinking position and was used for further analysis.

### miRNA analyses

For the genomic separations of crosslink positions we used GENCODE (v27 primary assembly) annotation and for the separation of transcripts with high and low miRNA targeting in HeLa cells we used ^68^ annotation. miRNA seed sequences were downloaded from ‘TargetScan’ (www.targetscan.org) database. Only miRNAs expressed in HeLa were selected from miRNA expression profile study ^73^ with the threshold of more than 10 reads in at least 2 replicates. The miRNA seed sequence heatmap was plotted by counting the expressed seed sequence motifs relative to the AGO2-eiCLIP dominant crosslink sites using the ‘ggplot2’ Bioconductor R package.

### CAGE data pre-processing

Paired-end sequenced CAGE data was downloaded from K562 (ENCSR000CJN) and HeLa (ENCSR000CJJ) cells was downloaded from ENCODE consortium with two biological replicates per sample. FASTQ files were mapped to the hg38 (GENCODE GRCh38.p10) gene annotation using STAR alignment tools (version 2.5.3a) by disabling 5’ read trimming function with the following command:

STAR --runMode alignReads --runThreadN $thread --genomeDir $genome_dir --readFilesIn ${path}${data1} ${path}${data2} --outSAMunmapped Within --outFilterMultimapNmax 1 --outFilterMultimapScoreRange 1 –outFileNamePrefix $path$data-STAR-Extend5pOfRead1/ --outSAMtype BAM SortedByCoordinate --outFilterType BySJout --outReadsUnmapped Fastx --outFilterScoreMin 10 --outSAMattrRGline ID:foo --alignEndsType Extend5pOfRead1 --clip5pNbases 9

### CAGE quality control

For the quality control we used Bioconductor CAGEr package (v2.0.2) by importing BAM files of mapped reads into R. The preprocessing was done by standardised pipeline provided by ENCODE, where trimming and adapter removal from raw reads was done by cutadapt (v4.2) tool followed by bowtie2 (v2.5.0) alignment tools. This type of mapping was needed to avoid junction reads, which are known to cause issues in certain R packages. Quality controls were then plotted with the following CAGEr functions `plotCorrelation2` and `plotReverseCumulatives`.

### CAGE data processing

The BAM files of mapped reads were converted into BED format using the bamtobed function from bedtools package (version v2.30.0). Each 5’ read position was then used for further analyses. The CAGE peaks were processed by using the *Paraclu* clustering tool (https://gitlab.com/mcfrith/paraclu). Default settings of minimum 5 reads filter for merged replicates was used followed by paraclu.cut.sh which removes:

1. Remove single-position clusters.
2. Remove clusters longer than 200. (Length = column_4 - column_3.)
3. Remove clusters with (maximum density / baseline density) < 2.
4. Remove any cluster that is contained in a larger cluster.
5. Single nucleotide clusters were added additionally

For each cluster the highest peak of 5’ CAGE reads was used as the max peak position.

### CAGE reproducibility of 3’UTR peaks

Mapped BAM samples from HeLa and K562 cell lines were converted to BED file format by using *bedtools* bamtobed conversion (v2.30.0) where 5’ read positions were used for further analyses. From each sample both replicates of 5’ read positions were used to define clusters within the 20 bps window by using bedtools (command: `bedtools merge -s -d 20`). For each cluster a maximum number of 5’ read-ends was defined as peak, with a threshold of minimum 2 reads per replicate. Read counts were then normalised by the library size factor function using Bioconductor DESeq2 R package. Correlation plots were then made with R (version 4.1.2) using Bioconductor ggplot2 package for scatter plots.

### eCLIP enrichment relative to 3’UTR CAGE peaks

ENCODE eCLIP data was processed by following a standardised guideline to study RBP-RNA interactions with CLIP Technologies ^66^. We mapped paired-end eCLIP samples to the human hg38 genome using annotation version GRCh38.p7 using the STAR (version 2.5.3a) alignment tool. For adapter removal the cutadapt tool (version 3.5) was used following the ENCODE guide line with two rounds of adapter removal in case there were double ligated adapters present. After mapping we removed PCR duplicates using the python script ‘barcode_collapse_pe.py’ provided by ENCODE. For the data format conversions between SAM, BAM and BED file types we used samtools (version 1.13) and bedtools (version v2.30.0).

For the eiCLIP-AGO2 samples we used a similar pipeline without double ligation removal, and additional custom script to swap random barcodes from the first 7 bps of the read sequence line to the header of the FASTQ read sequence. Uniquely mapped reads with the same genomic positions and non-unique barcode were treated as PCR duplicates by being discarded from the further analyses.

To identify RBP binding enrichments we first analysed input controls by using eCLIP mock samples from all 80 RBPs from K562 experiments provided by ENCODE consortium. For each sample we used False Discovery Rate peak finding algorithm from iCount (https://github.com/tomazc/iCount), by assessing the enrichment of crosslink sites at specific binding sites compared to shuffled data. The peak caller was set to 3nt peak window size to define binding regions genome-wide. Next, we merged all binding regions into one track and longer regions from 50 nts were evenly split into smaller clusters. For each binding site, RBP ratio was calculated relative to the maximum RBP enrichment. These ratios were then used to calculate the RBP enrichment profile around the 3’UTR CAGE peaks.

### RNA-seq

Raw reads of two biological replicates of stranded paired-end RNA-seq samples were downloaded from K562 (ENCODE: ENCFF044SJL, ENCFF728JKQ) and HeLa (GSE99169) cell lines. FASTQ files were then aligned to the human genome by STAR (version 2.5.3a) alignment tool using GENCODE annotation version GRCh38.p7. Soft-clipping was disabled to contain full length reads by using the following parameters:

*STAR --runMode alignReads --runThreadN $thread --genomeDir $genome_dir --readFilesIn ${FASTQ.read1} ${FASTQ.read2} --outSAMunmapped Within --outFilterMultimapNmax 1 --outFilterMultimapScoreRange 1 --outFileNamePrefix $path$data1-STAR/ --outSAMtype BAM SortedByCoordinate --outFilterType BySJout --outReadsUnmapped Fastx –outFilterScoreMin 10 --outSAMattrRGline ID:foo --alignEndsType EndToEnd*

Mapped paired-end reads were then converted from BAM to BED by using ‘bedtools bamtobed’ (version v2.30.0) function to extract both sides of each read. Read starts and read ends were then plotted as a metaplot relative to the 3’UTR CAGE peaks.

### CBP20-iCLIP

The CBP20-iCLIP data was downloaded from GEO (GSE94427) and analysed using standard iCLIP processing pipeline where each read was treated as truncated read to identify crosslinking positions of protein-RNA interactions ^74^. For the adapter removal we used the cutadapt tool (version 3.5) with removal of shorter reads than 18 bps.

`cutadapt --match-read-wildcards --times 1 -e 0.1 -O 1 --quality-cutoff 6 -m 18 -a AGATCGGAAG $data > ${data}.adapterTrim.fastq 2> $path$data.adapterTrim.metrics`

Random barcode from each read was then removed into a read header by using a custom python script. For mapping the read to human hg38 (GENCODE GRCh38.p7 annotation) genome we used STAR alignment tool (version 2.5.3a) with the following command:

*STAR --runMode alignReads --runThreadN $thread --genomeDir $genome_dir --readFilesIn ${data}.adapterTrim.barcodes.fastq --outSAMunmapped Within --outFilterMultimapNmax 1 --outFilterMultimapScoreRange 1 --outFileNamePrefix $data-STAR/ --outSAMattributes All --outStd BAM_SortedByCoordinate --outFilterType BySJout --outReadsUnmapped Fastx –outFilterScoreMin 10 --outSAMattrRGline ID:foo --alignEndsType EndToEnd*

BAM file of mapped reads was then converted into BED file using bedtools (version v2.30.0) bamtobed function followed by removal of PCR duplicates by collapsing identical reads with the same random barcode. For each read the read start position was used as the crosslinking position and was used for further analysis.

### rG4-seq

The processed RNA-G-quadruplex sequencing (rG4-seq) data from HeLa cells was downloaded from the genomics data repository (GSE77282). The rG4-seq hits were then lifted from the hg19 to hg38 genome using UCSC liftOver webtool. For Figure 2E we used middle positions of each rG4-seq target normalised by the number of CAGE (HeLa) peaks from each transcriptome region.

### RNA-maps of iCLIP, eCLIP, eiCLIP and RNA-seq reads start/ends

For the visualisation of all the CLIP based and RNA-seq methods we used previously developed RNA-map approach ^66, 74^ with small addition for RNA-seq read end positions by summarising the read start positions relative to the CAGE peaks, TSSs and G-Quadruplexes.

### Secondary structure

For each dominant CAGE peak we extracted a flanking region of 75 bps of the genomic sequence as an input to the RNAfold vienna package (version 2.4.17) with default settings. Each double stranded position was then plotted as a sum of all pairings in the region.

### Predictions of G-quadruplexes

To predict G-quadruplexes in the K562 and HeLa cell line we first selected CAGE peaks with a threshold of minimum 10 reads per peak in the region of 50 bps upstream and downstream from the peak. For the predictions we used sequence based prediction tool G4Hunter (https://github.com/AnimaTardeb/G4Hunter) with the following settings: `G4Hunter.py -i INPUT.fasta_sequence -o G4Hunter -w 25 -s 1.2`

### Motif discovery

For AGO2 binding motif discovery we used HOMER software for motif discovery and next-gen sequencing analysis (version 4.9), with default parameters for human genome hg38 and using a 15 bps window around crosslink positions of processed AGO2-eiCLIP-HeLa samples.

### Motif enrichment

For canonical polyA A[A/U]UAAA hexamers enrichment we first selected 3’UTR ending positions from GENCODE (v27) annotation. For each 3’UTR ending position we looked at the 100 nt flanking position and counted hexamer coverage per nucleotide. The same was done for 3’UTR CAGE (K562) peaks with removal of CAGE peaks that were in 200 nt into the 3’UTR region CAGE peaks were removed.

### siRNA-KD CAGE samples

For the capping of siRNA-targeting sites analyses we used 28 siRNA-KD samples and 5 Control samples with 3 replicates per sample from FANTOM5 ^42^. For individual KDs we plotted CAGE transcription start sites (CTSS) of Control and KDs around the siRNA targeting regions. For the Heatmap we first selected 20 out of 28 samples that had at least two overlapping replicates in the corresponding siRNA targeting region and then merged the replicates. Each siRNA targeting position was manually identified by using BLAST. For the control, we merged together all 5 samples (with 3 replicates per sample) into a metaplot (Figure S4A) normalised by the number of samples.

### Long-read CAGE

Long-read CAGE was based on the Cap-Trapper method with the full length cDNA sequencing using ONT MinION sequencer. After RNA extraction, 10 µg total RNAs from Human i^3^N-iPSC that harbours a doxycycline-inducible mouse Ngn2 transgene at an adeno-associated virus integration site 1 (AAVS1) safe-harbour locus of WTC11 iPSC line (https://www.ncbi.nlm.nih.gov/pmc/articles/PMC5639430/), differentiated neural stem cells and differentiated cortical neuron cells were polyadenylated with E-coli poly(A) Polymerase (PAP) (NEB M0276) at 37°C for 15 min and purified with AMPure RNA Clean XP beads. The PAP treated RNA (5 µg) was reverse transcribed with oligodT_16VN_UMI25_primer (GAGATGTCTCGTGGGCTCGGNNNNNNNNNNNNNNNNNNNNNNNNNCTACGTTTTTTTTTTTTTTTTVN) and Prime Script II Reverse Transcriptase (Takara Bio) at 42°C for 60 min. After purification with RNAClean XP beads, Cap-trapping from the RNA/cDNA hybrid was performed as previously described ^18^. RNA from the hybrid was depleted by RNase H (Takara Bio) digestion at 37°C for 30 min and the product was purified with AMPureXP beads. Then, 5’ linker (constituted of N6 up GTGGTATCAACGCAGAGTACNNNNNN-Phos, GN5 up GTGGTATCAACGCAGAGTACGNNNNN-Phos, down Phos-GTACTCTGCGTTGATACCAC-Phos) was ligated to the cDNA with Mighty Mix (Takara Bio) with overnight incubation and the ligated cDNA was purified with AMPure XP beads. Shrimp Alkaline Phosphatase (Takara Bio) was used to remove phosphates from the ligated linker and the product was purified with AMPureXP beads. The 5’ linker ligated cDNA was then second strand synthesised with KAPA HiFi mix (Roche) and the 2nd synthesis primer_UMI15 at 95°C for 5 min, 55°C for 5 min and 72°C for 30 min. Exonuclease I (Takara Bio) was added and incubated at 37°C for 30 min to remove excessive primer. Then, the cDNA/DNA hybrid was purified with AMPureXP and amplified with PrimerSTAR GXL DNA polymerase (Takara Bio) using PCR primers (fwd_CTACACTCGTCGGCAGCGTC, rev_GAGATGTCTCGTGGGCTCGG) for 7 cycles. The library was then subjected to the SQK-LSK110 (Oxford Nanopore Technologies) protocol according to manufacturer’s instructions and sequenced with R9.4 flowcell (FLO-MIN106) in MinION sequencer. Basecalling was processed by Guppy v5.0.14 basecaller software provided by Oxford Nanopore Technologies in high-accuracy mode to generate FASTQ files from FAST5 files. To prepare clean reads from FASTQ files, adapter sequences (VNP_GAGATGTCTCGTGGGCTCGGNNNNNNNNNNNNNNNCTACG and SSP_ CTACACTCGTCGGCAGCGTCNNNNNNNNNNNNNNNNNNNNNNNNNGTGGTATCAACGCAGAGTAC) and poly-A tails were trimmed by primer-chop (https://gitlab.com/mcfrith/primer-chop) and then oriented to original RNA strand. The clean FASTQ reads were mapped on our target genes.

### Data, pipelines and scripts

Raw sequencing files of eiCLIP-AGO2 experiment have been deposited in the ArrayExpress archive accessible at E-MTAB-12945. All the pipelines and scripts used in this study are deposited and available on github (https://github.com/nebo56/3pUTR-derived-RNAs).

