## Supplementary figure S1 for "Abundant capped RNAs are derived from mRNA cleavage at 3’UTR G-Quadruplexes"

**A**

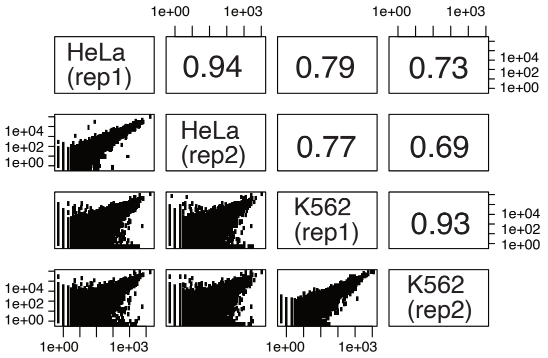


**B**

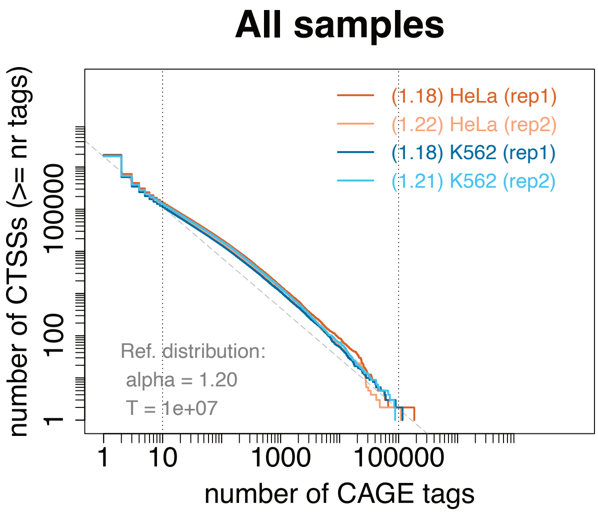


**C**

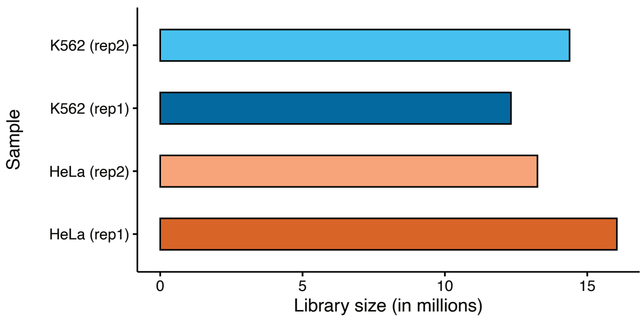


**D**

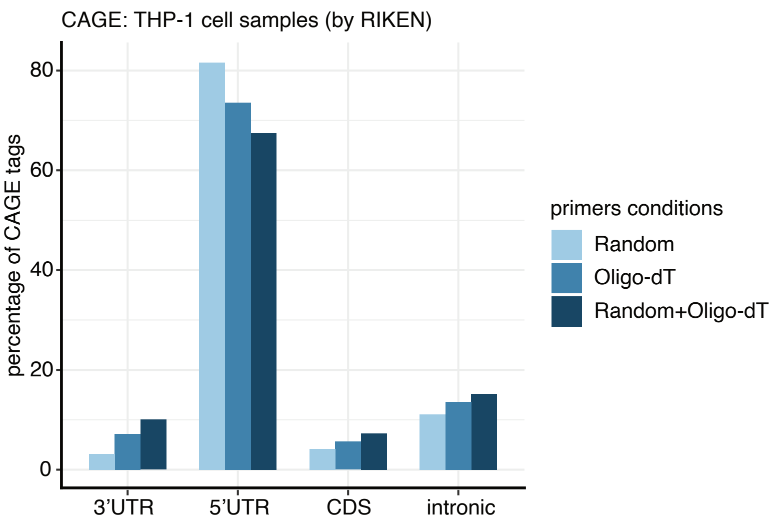

**E**

**3’UTR**

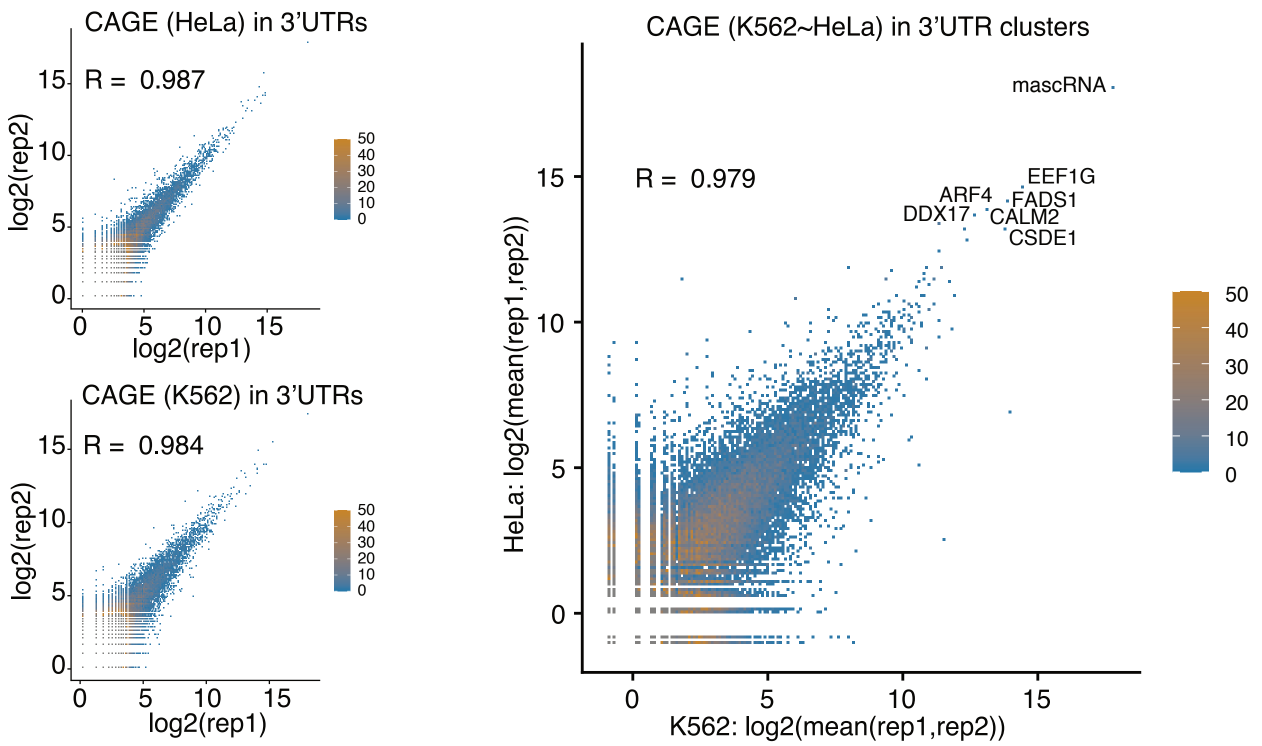
CAGE (HeLa) in 3’UTRs
R = 0.987
log2(rep2)
log2(rep1)
CAGE (K562) in 3’UTRs
R = 0.984
log2(rep2)
log2(rep1)
CAGE (K562~HeLa) in 3’UTR clusters
mascRNA
R = 0.979
EEF1G
ARF4
FADS1
DDX17
CALM2
CSDE1
HeLa: log2(mean(rep1,rep2))
K562: log2(mean(rep1,rep2))
50
40
30
20
10
0

**F**

**5’UTR**

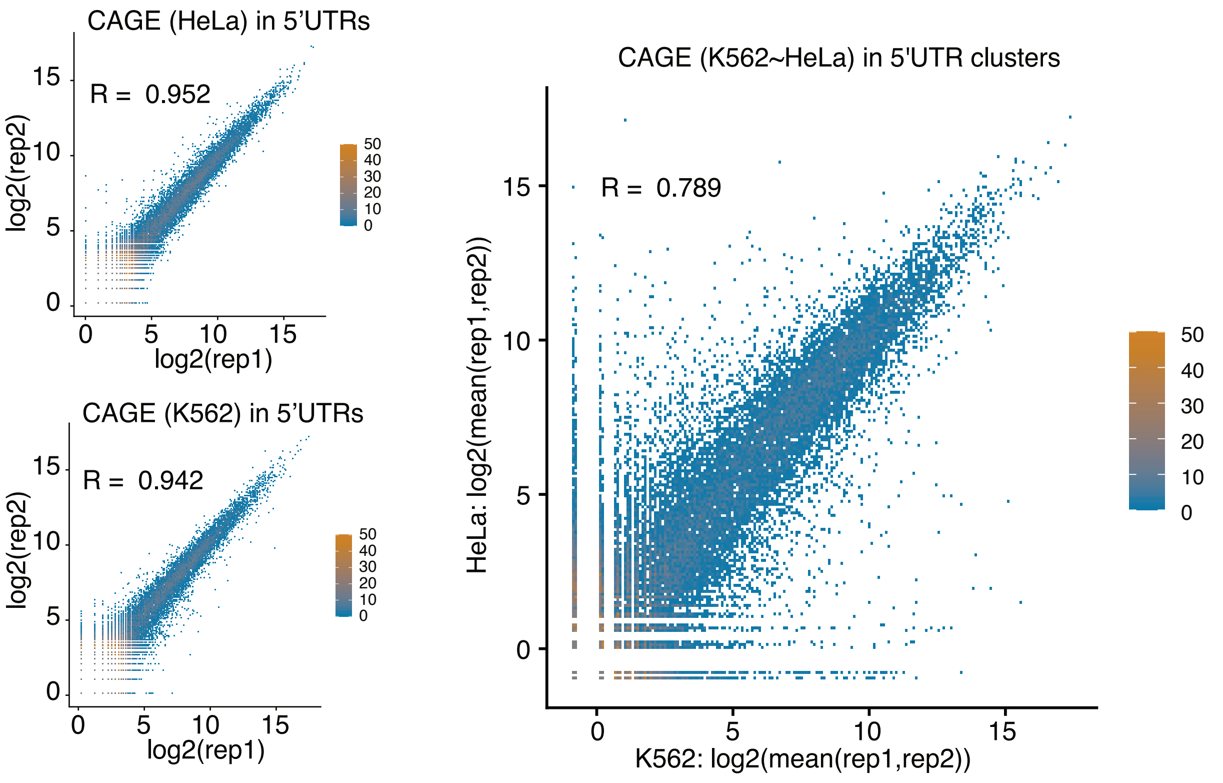
CAGE (HeLa) in 5’UTRs
R = 0.952
log2(rep2)
log2(rep1)
CAGE (K562) in 5’UTRs
R = 0.942
log2(rep2)
log2(rep1)
CAGE (K562~HeLa) in 5'UTR clusters
R = 0.789
HeLa: log2(mean(rep1,rep2))
K562: log2(mean(rep1,rep2))
50
40
30
20
10
0

G

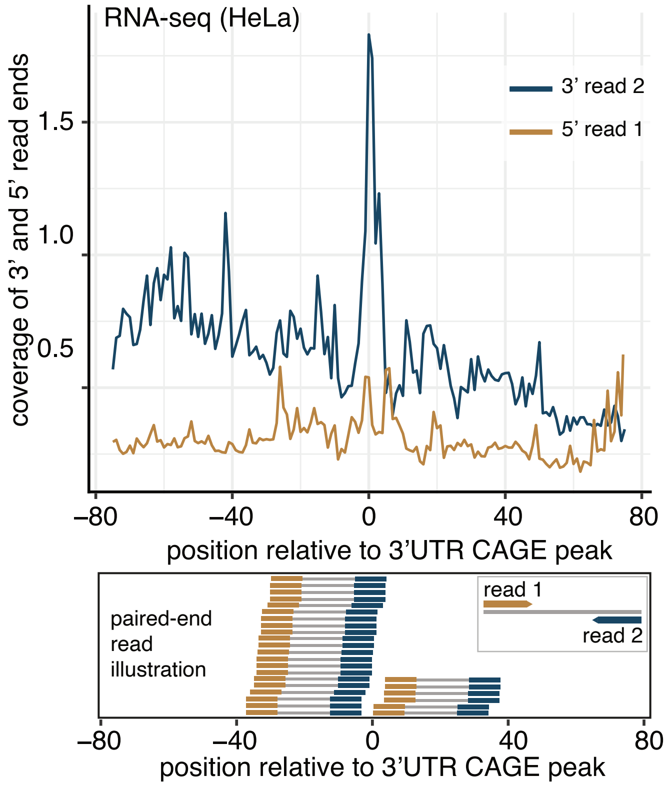

RNA-seq (HeLa)
3’ read 2
5’ read 1
coverage of 3’ and 5’ read ends
1.5
1.0
0.5
–80
–40
0
40
80
position relative to 3’UTR CAGE peak
paired-end
read
illustration
read 1
read 2
–80
–40
0
40
80
position relative to 3’UTR CAGE peak

**H**

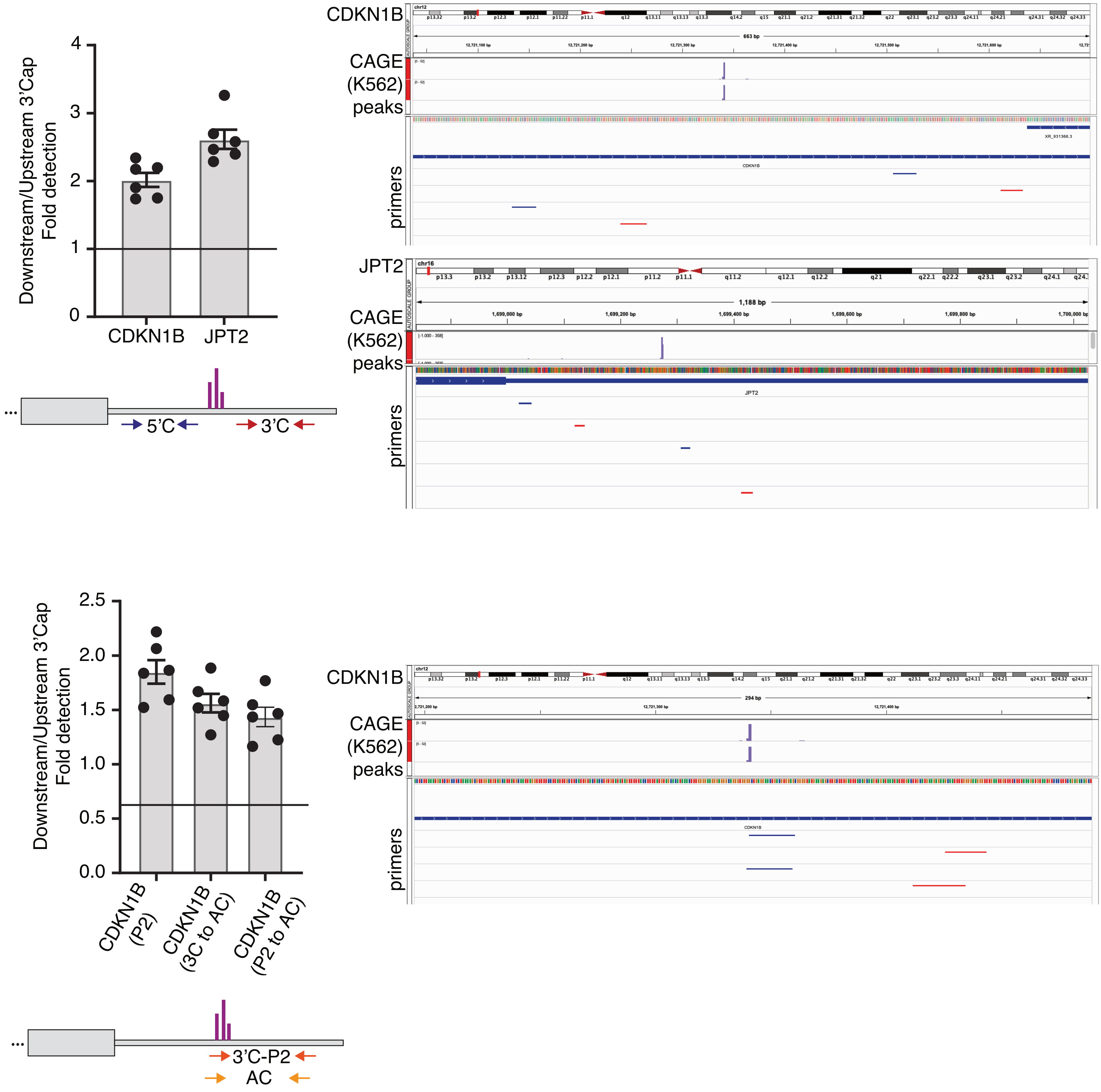
Downstream/Upstream 3'Cap Fold detection
0
1
2
3
4
CDKN1B
JPT2
5'C
3'C
CDKN1B
CAGE (K562) peaks
primers
chr12
663 bp
CDKN1B
XR_931366.3
JPT2
chr16
1,188 bp
Downstream/Upstream 3'Cap Fold detection
0.0
0.5
1.0
1.5
2.0
2.5
CDKN1B (P2)
CDKN1B (3C to AC)
CDKN1B (P2 to AC)
3'C-P2
AC
294 bp

I

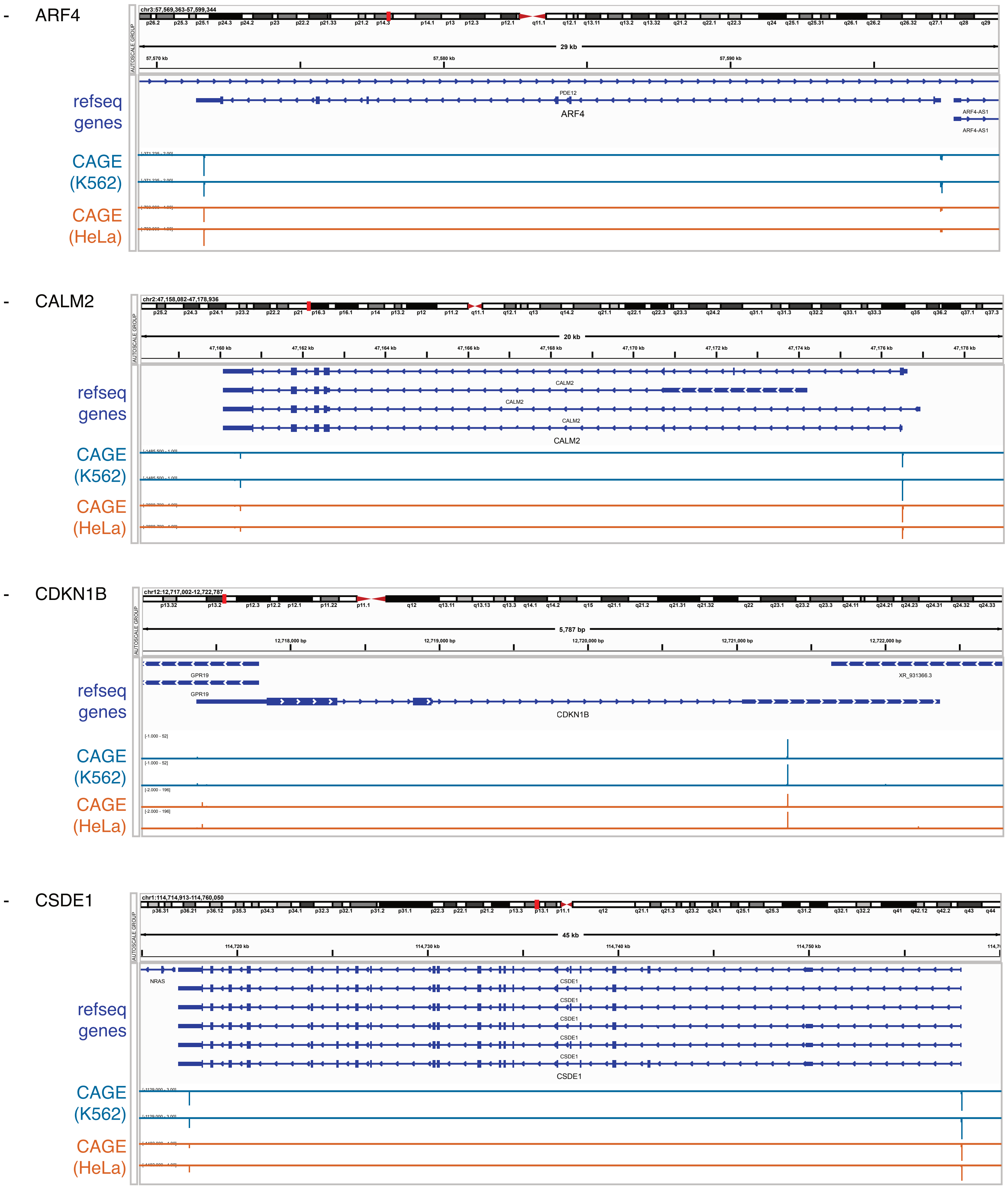

- ARF4
refseq genes
CAGE (K562)
CAGE (HeLa)
PDE12
ARF4
ARF4-AS1
29 kb
- CALM2
refseq genes
CAGE (K562)
CAGE (HeLa)
CALM2
20 kb
- CDKN1B
refseq genes
CAGE (K562)
CAGE (HeLa)
GPR19
XR_931366.3
CDKN1B
5,787 bp
- CSDE1
refseq genes
CAGE (K562)
CAGE (HeLa)
NRAS
CSDE1
45 kb

- DDX17

refseq genes

CAGE (K562)

CAGE (HeLa)

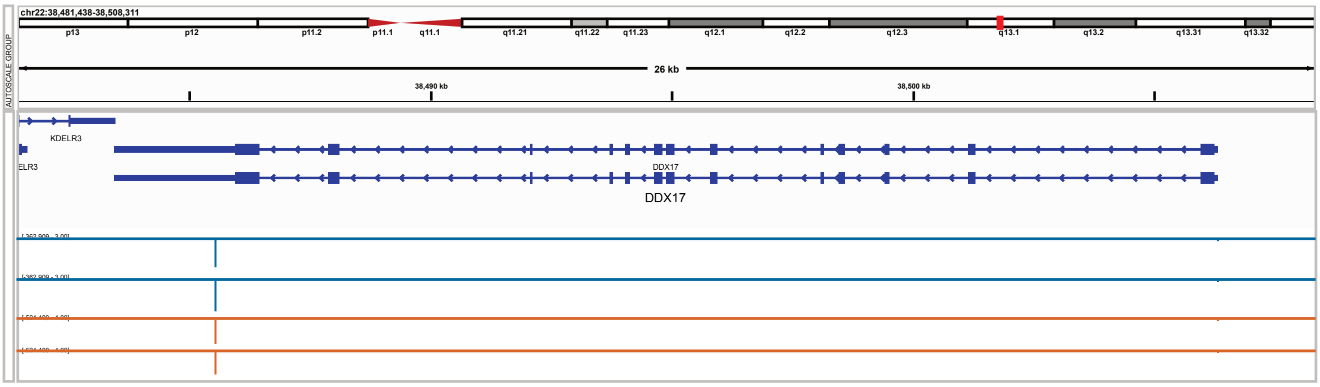


- FADS1

refseq genes

CAGE (K562)

CAGE (HeLa)

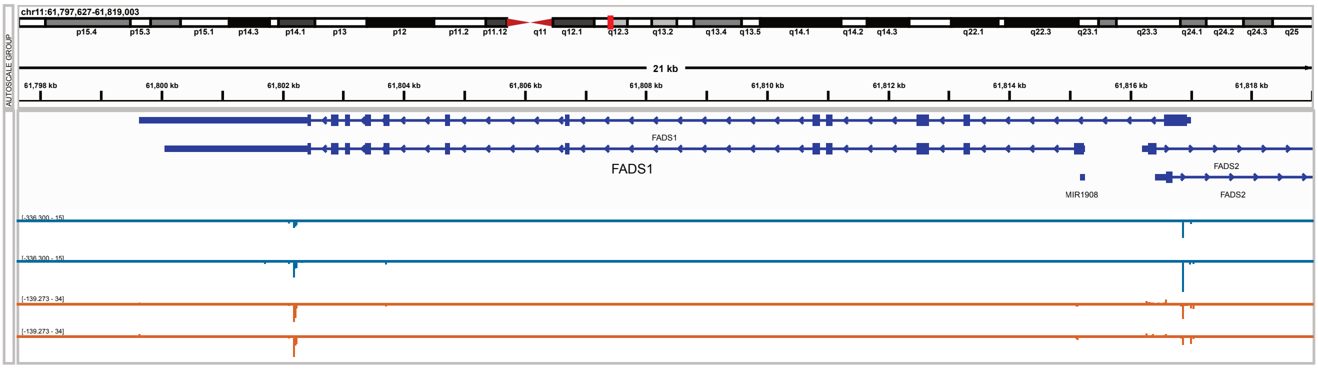


- JPT2

refseq genes

CAGE (K562)

CAGE (HeLa)

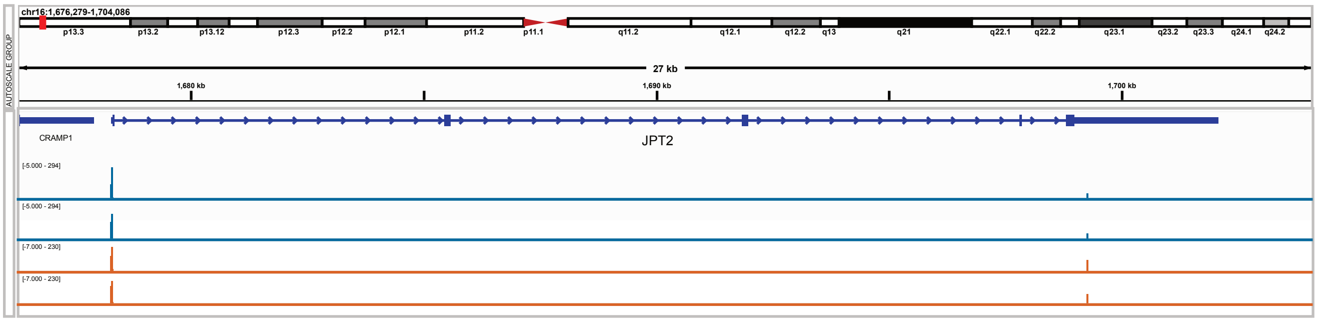


**J**

- 3'UTR of CDKN1B

refseq genes

CAGE (K562)

CAGE (HeLa)

RNA-seq (K562)

RNA-seq (HeLa)

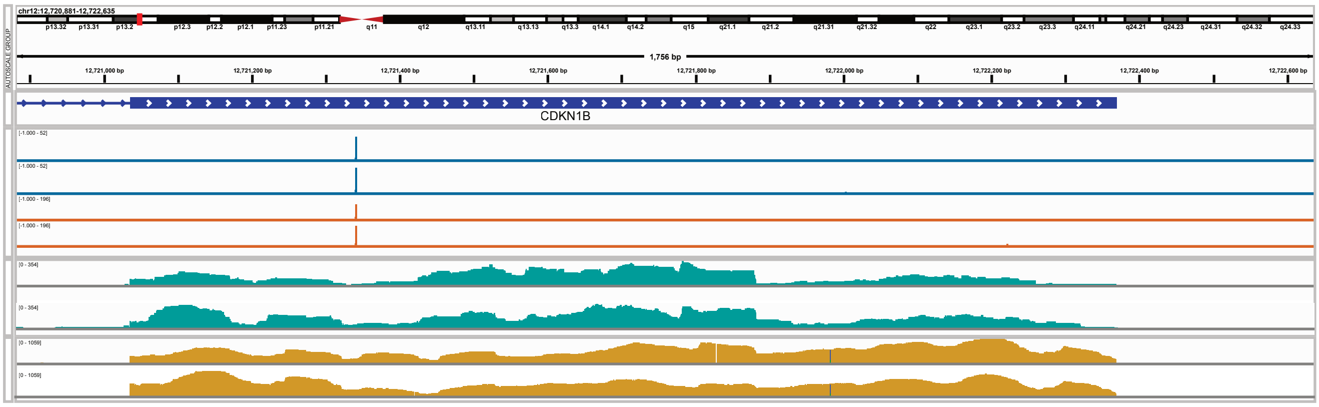


- 3'UTR of JPT2

refseq genes

CAGE (K562)

CAGE (HeLa)

RNA-seq (K562)

RNA-seq (HeLa)

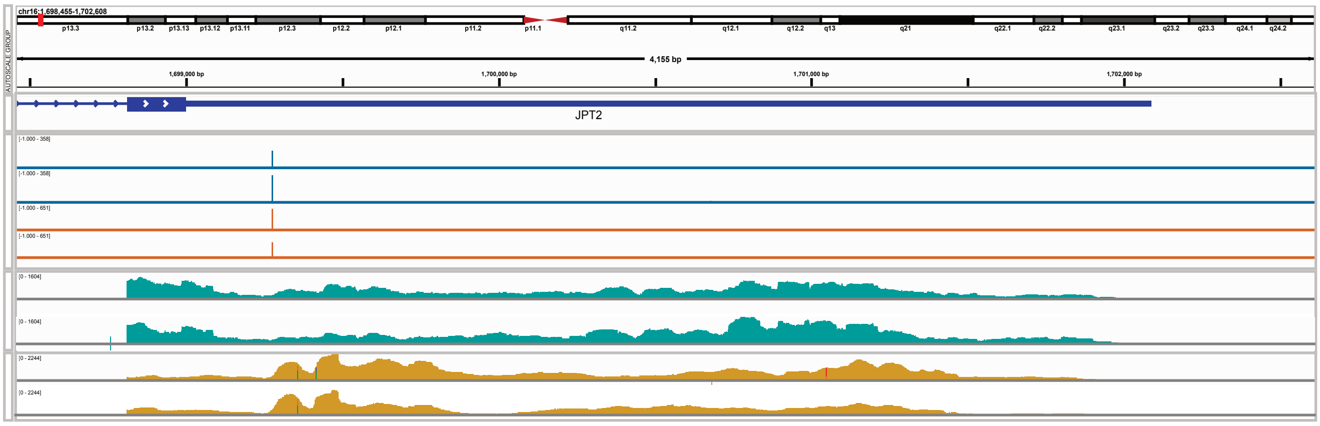

**K**

CDKN1B

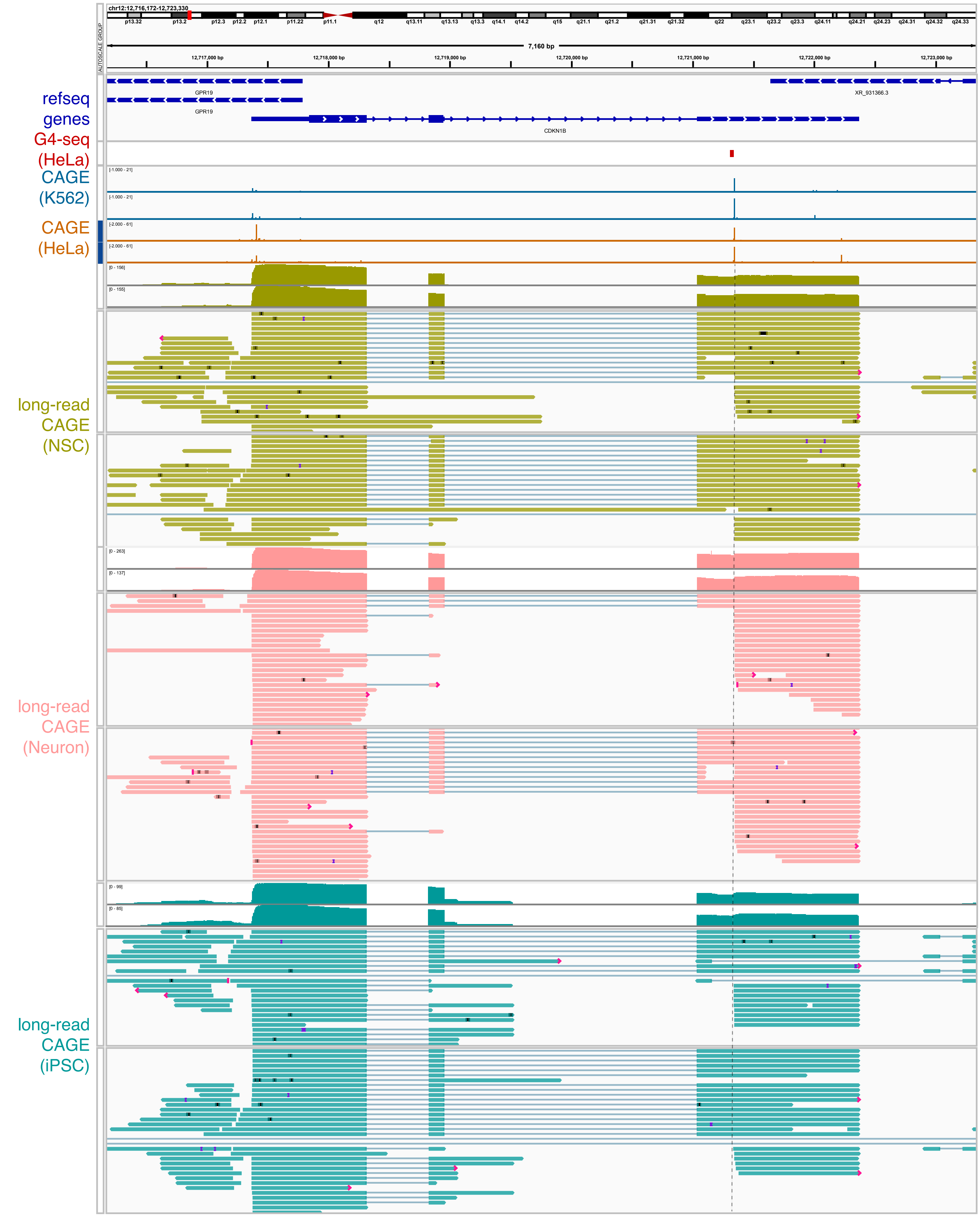
chr12:12,716,172-12,723,330
7,160 bp
12,717,000 bp
12,718,000 bp
12,719,000 bp
12,720,000 bp
12,721,000 bp
12,722,000 bp
12,723,000 bp
refseq genes
GPR19
GPR19
XR_931366.3
CDKN1B
G4-seq (HeLa)
CAGE (K562)
CAGE (HeLa)
long-read CAGE (NSC)
long-read CAGE (Neuron)
long-read CAGE (iPSC)

JPT2

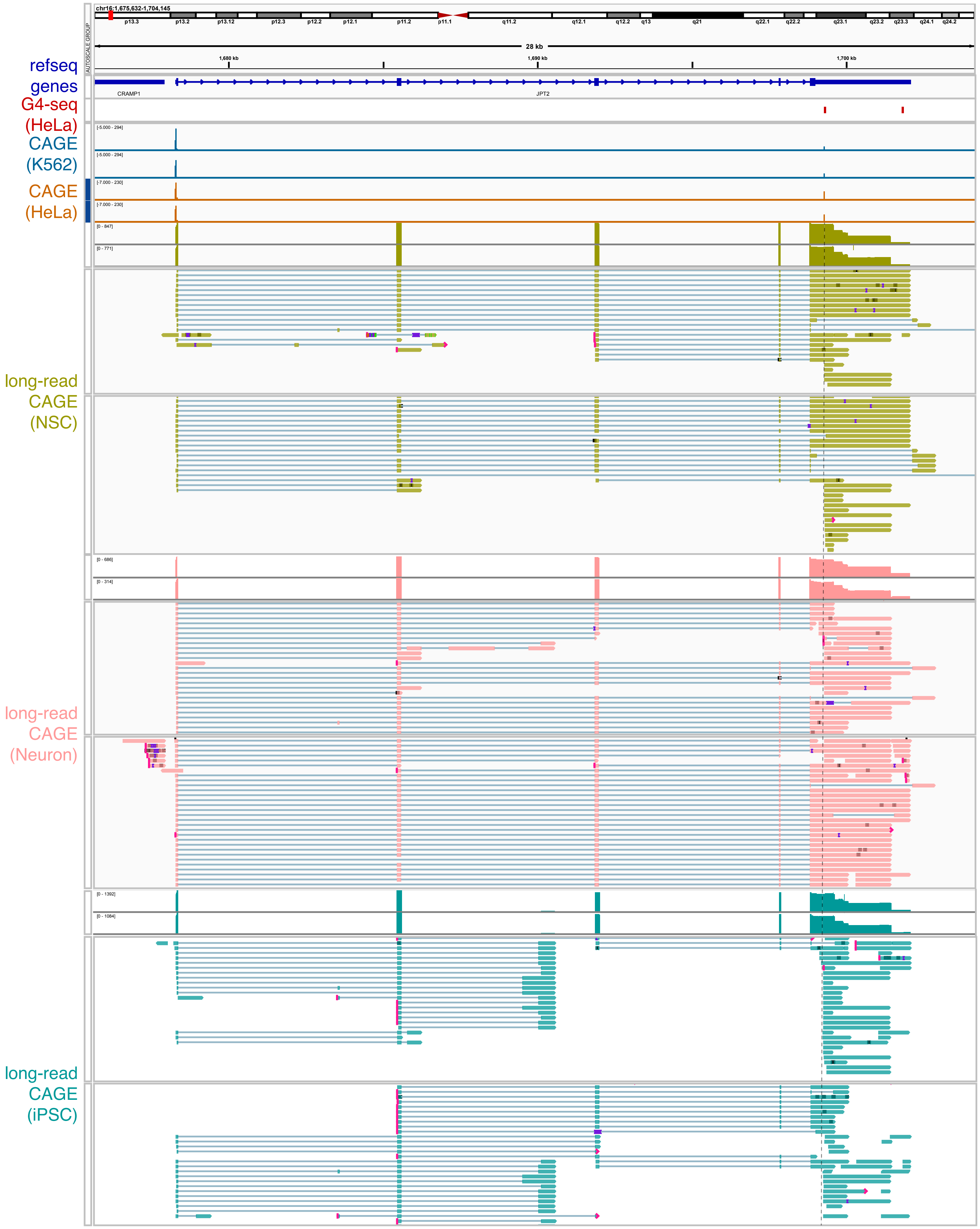

chr16:1,675,632-1,704,145
p13.3
p13.2
p13.12
p12.3
p12.2
p12.1
p11.2
p11.1
q11.2
q12.1
q12.2
q13
q21
q22.1
q22.2
q23.1
q23.2
q23.3
q24.1
q24.2
28 kb
1,680 kb
1,690 kb
1,700 kb
AUTOSCALE GROUP
refseq genes
CRAMP1
JPT2
G4-seq (HeLa)
CAGE (K562)
CAGE (HeLa)
long-read CAGE (NSC)
long-read CAGE (Neuron)
long-read CAGE (iPSC)

DDX17

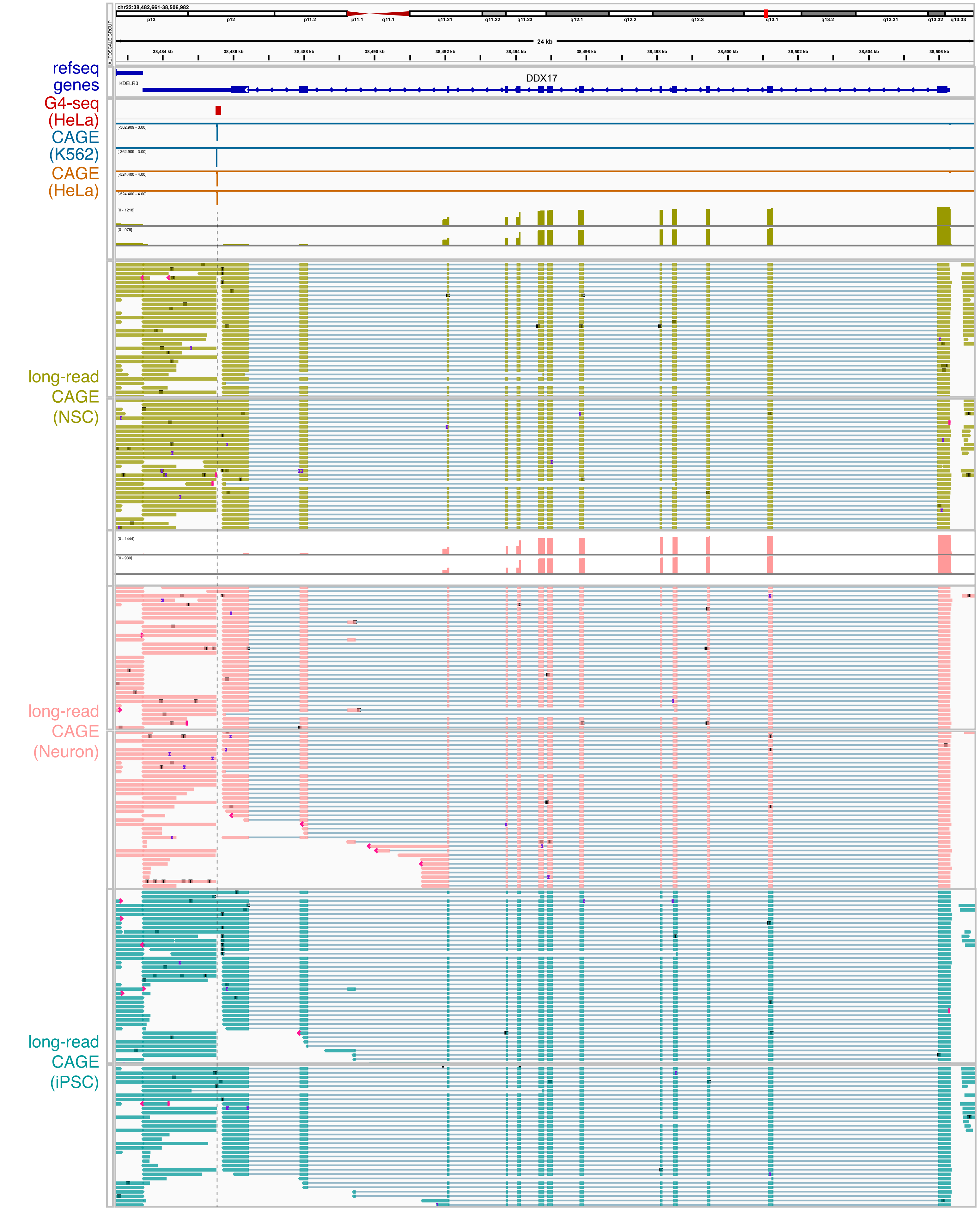
refseq genes
G4-seq (HeLa)
CAGE (K562)
CAGE (HeLa)
long-read CAGE (NSC)
long-read CAGE (Neuron)
long-read CAGE (iPSC)
chr22:38,482,661-38,506,982
24 kb
DDX17
KDELR3
AUTOSCALE GROUP

RAN

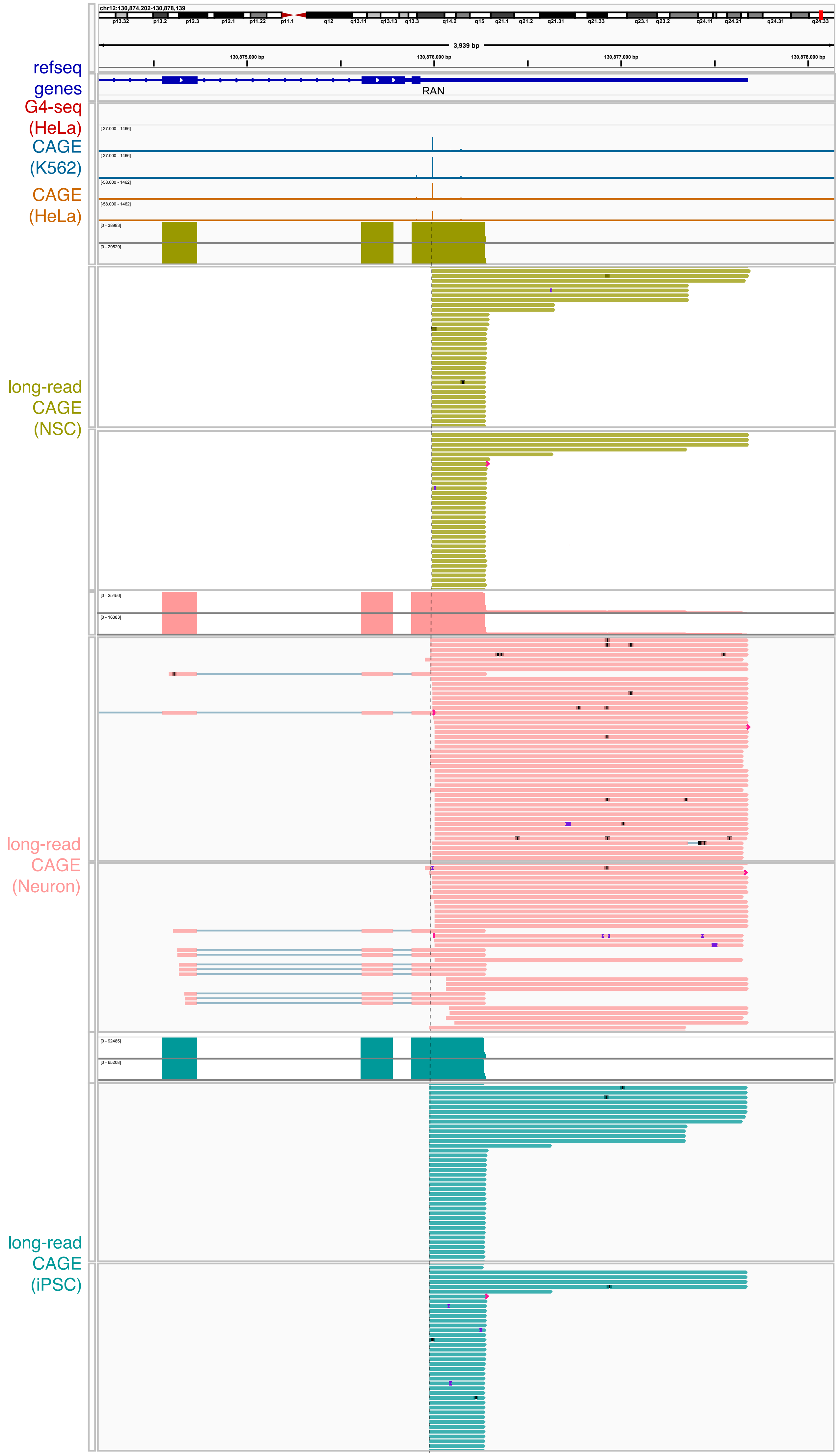

refseq genes
G4-seq (HeLa)
CAGE (K562)
CAGE (HeLa)
long-read CAGE (NSC)
long-read CAGE (Neuron)
long-read CAGE (iPSC)
chr12:130,874,202-130,878,139
3,939 bp
130,875,000 bp
130,876,000 bp
130,877,000 bp
130,878,000 bp
RAN

ARPC5L

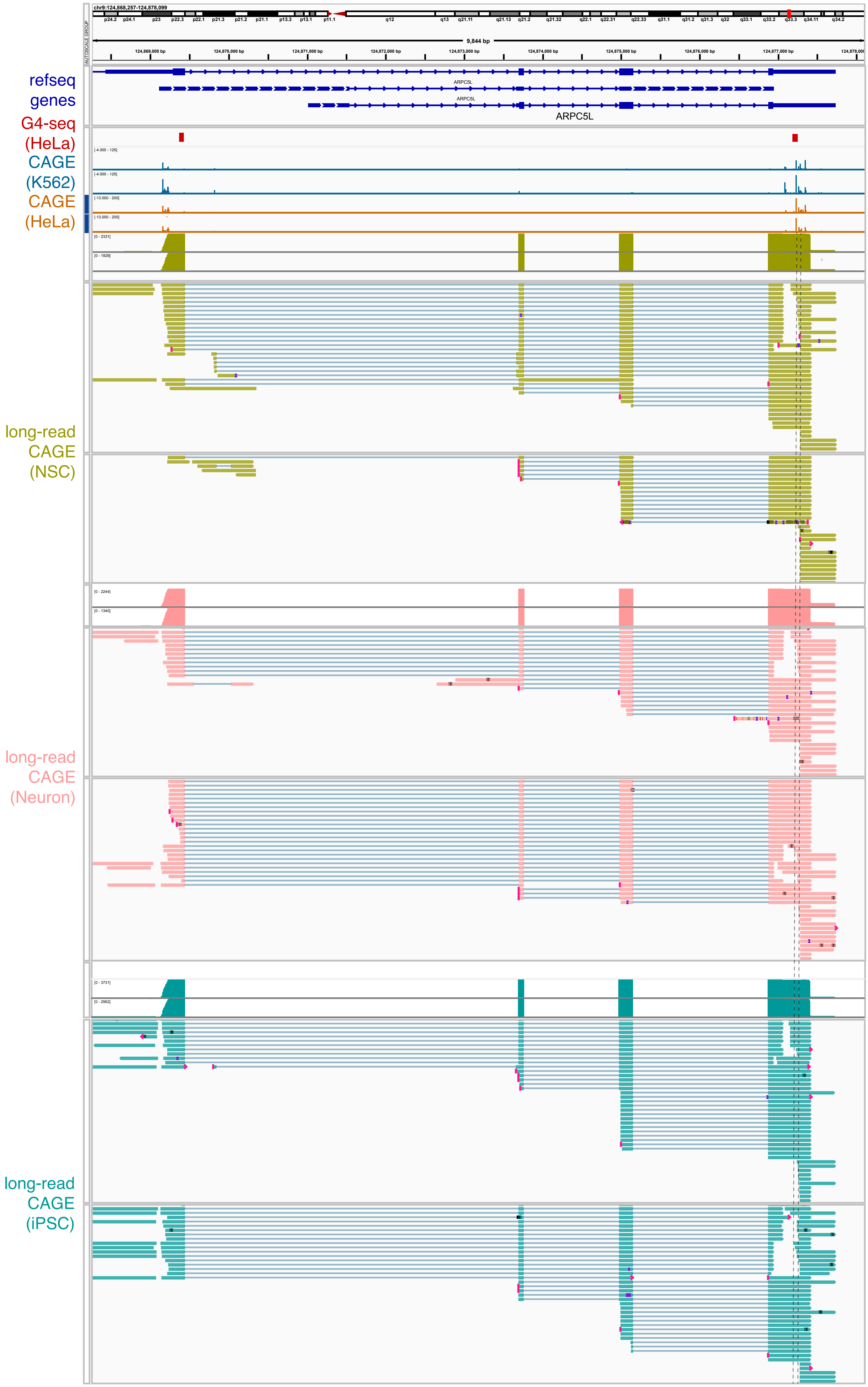

chr9:124,868,257-124,878,099
9,844 bp
124,869,000 bp
124,870,000 bp
124,871,000 bp
124,872,000 bp
124,873,000 bp
124,874,000 bp
124,875,000 bp
124,876,000 bp
124,877,000 bp
refseq genes
ARPC5L
G4-seq (HeLa)
CAGE (K562)
CAGE (HeLa)
long-read CAGE (NSC)
long-read CAGE (Neuron)
long-read CAGE (iPSC)

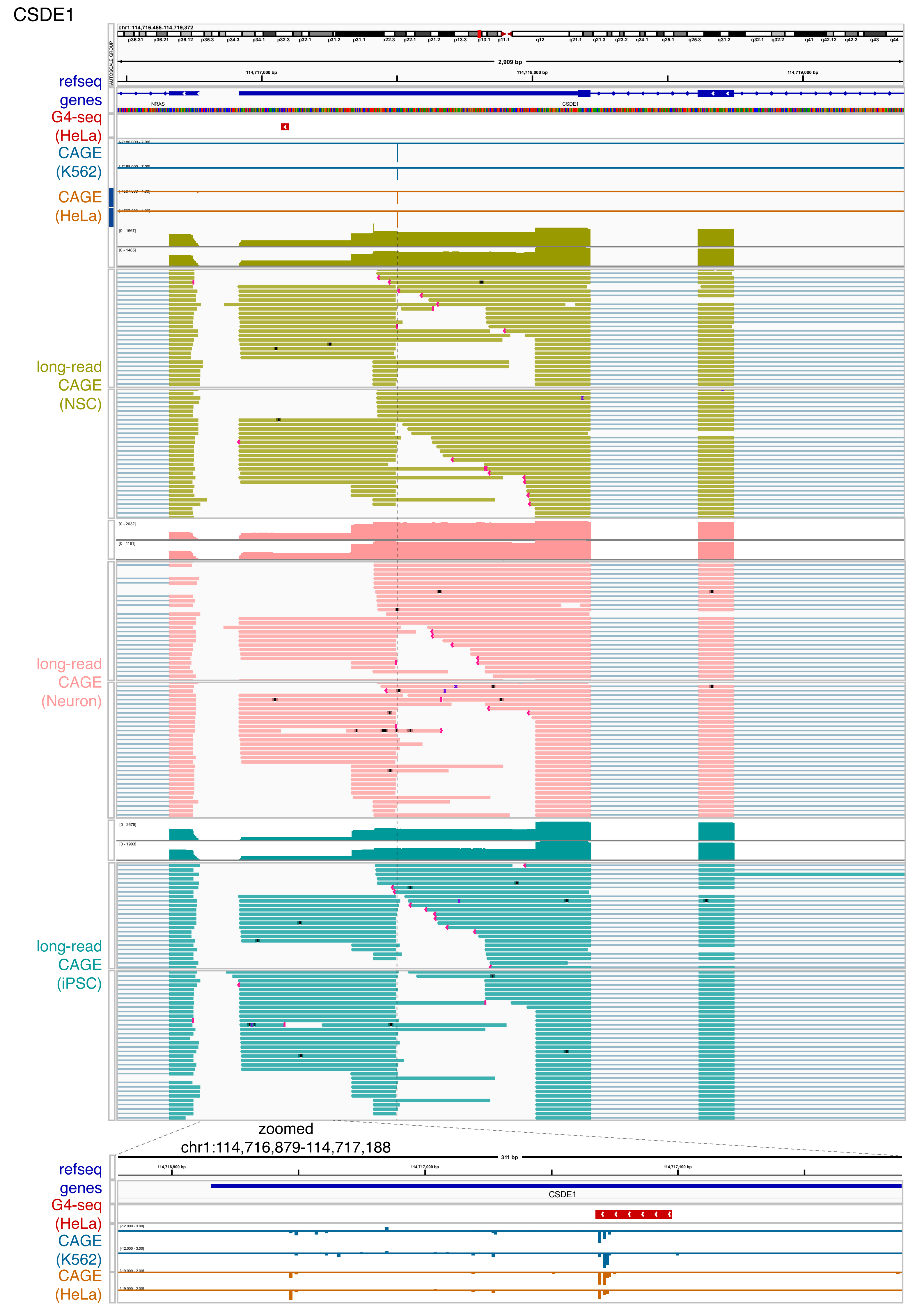
CSDE1
refseq genes
G4-seq (HeLa)
CAGE (K562)
CAGE (HeLa)
long-read CAGE (NSC)
long-read CAGE (Neuron)
long-read CAGE (iPSC)
chr1:114,716,465-114,719,372
2,909 bp
114,717,000 bp
114,718,000 bp
114,719,000 bp
NRAS
CSDE1
zoomed
chr1:114,716,879-114,717,188
311 bp
114,716,900 bp
114,717,000 bp
114,717,100 bp

### GHITM

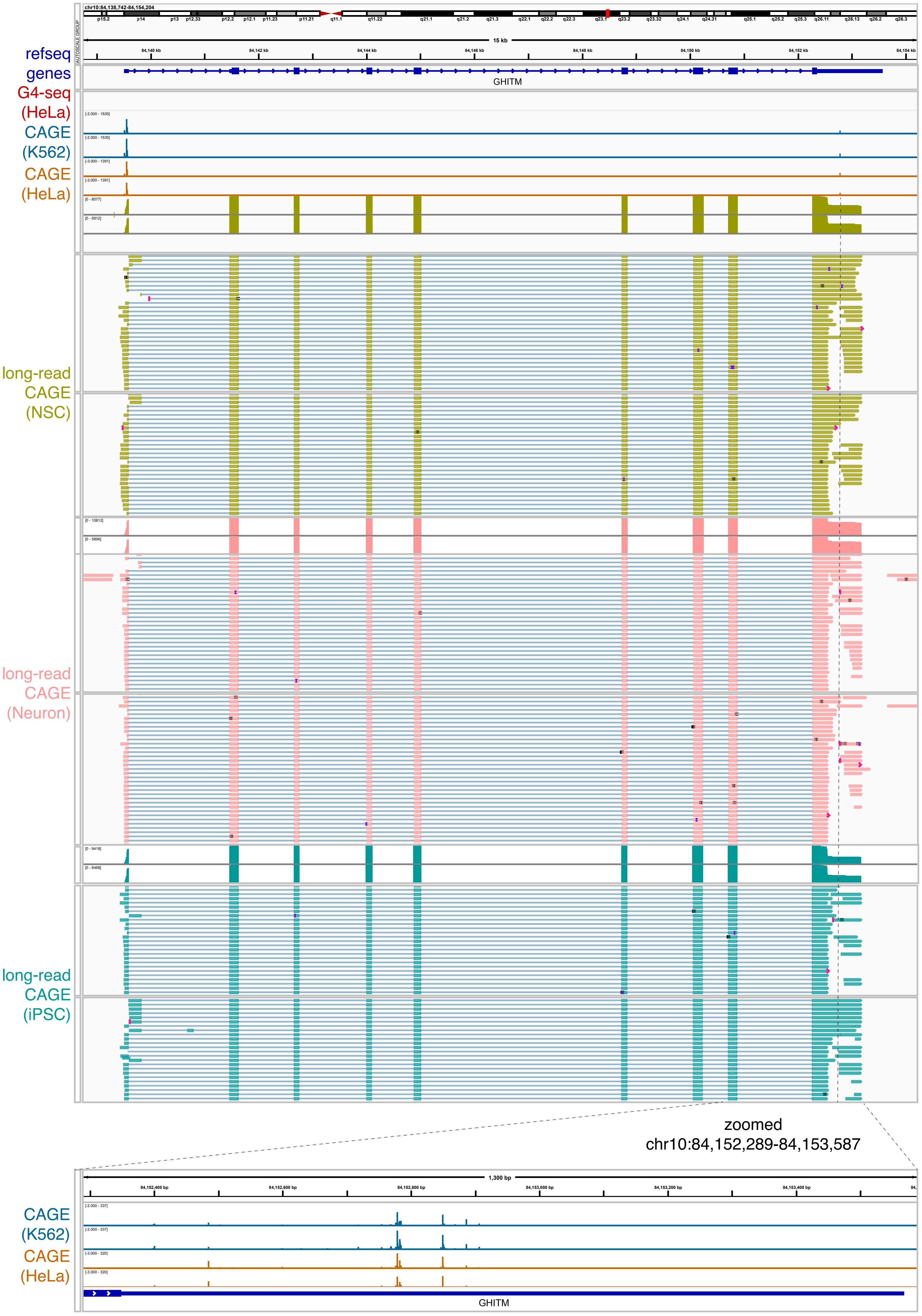
refseq genes
G4-seq (HeLa)
CAGE (K562)
CAGE (HeLa)
long-read CAGE (NSC)
long-read CAGE (Neuron)
long-read CAGE (iPSC)
GHITM
zoomed chr10:84,152,289-84,153,587
CAGE (K562)
CAGE (HeLa)
GHITM

### TNPO3

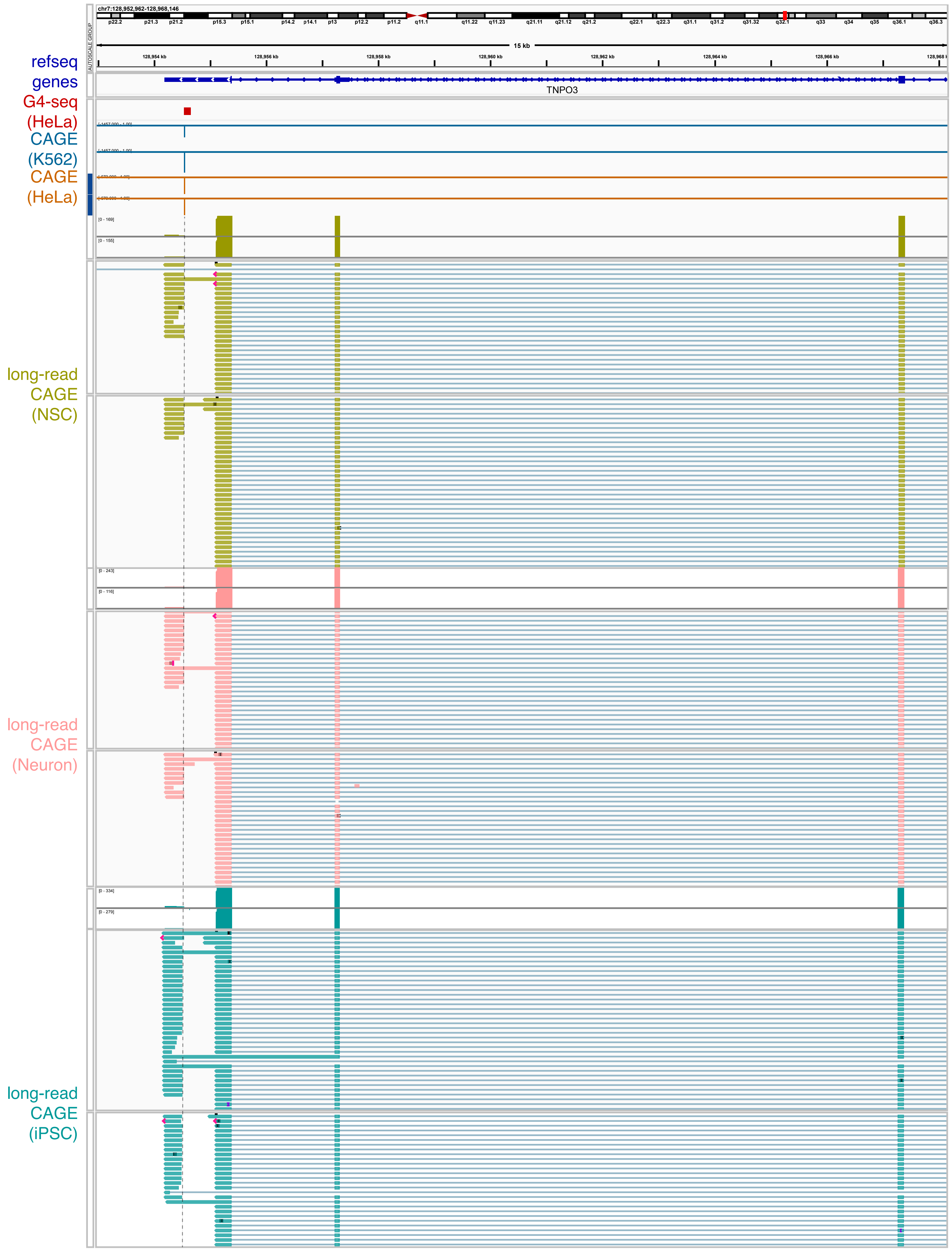

chr7:128,952,962-128,968,146
15 kb
128,954 kb
128,956 kb
128,958 kb
128,960 kb
128,962 kb
128,964 kb
128,966 kb
refseq genes
TNPO3
G4-seq (HeLa)
CAGE (K562)
CAGE (HeLa)
long-read CAGE (NSC)
long-read CAGE (Neuron)
long-read CAGE (iPSC)

ZAFND3

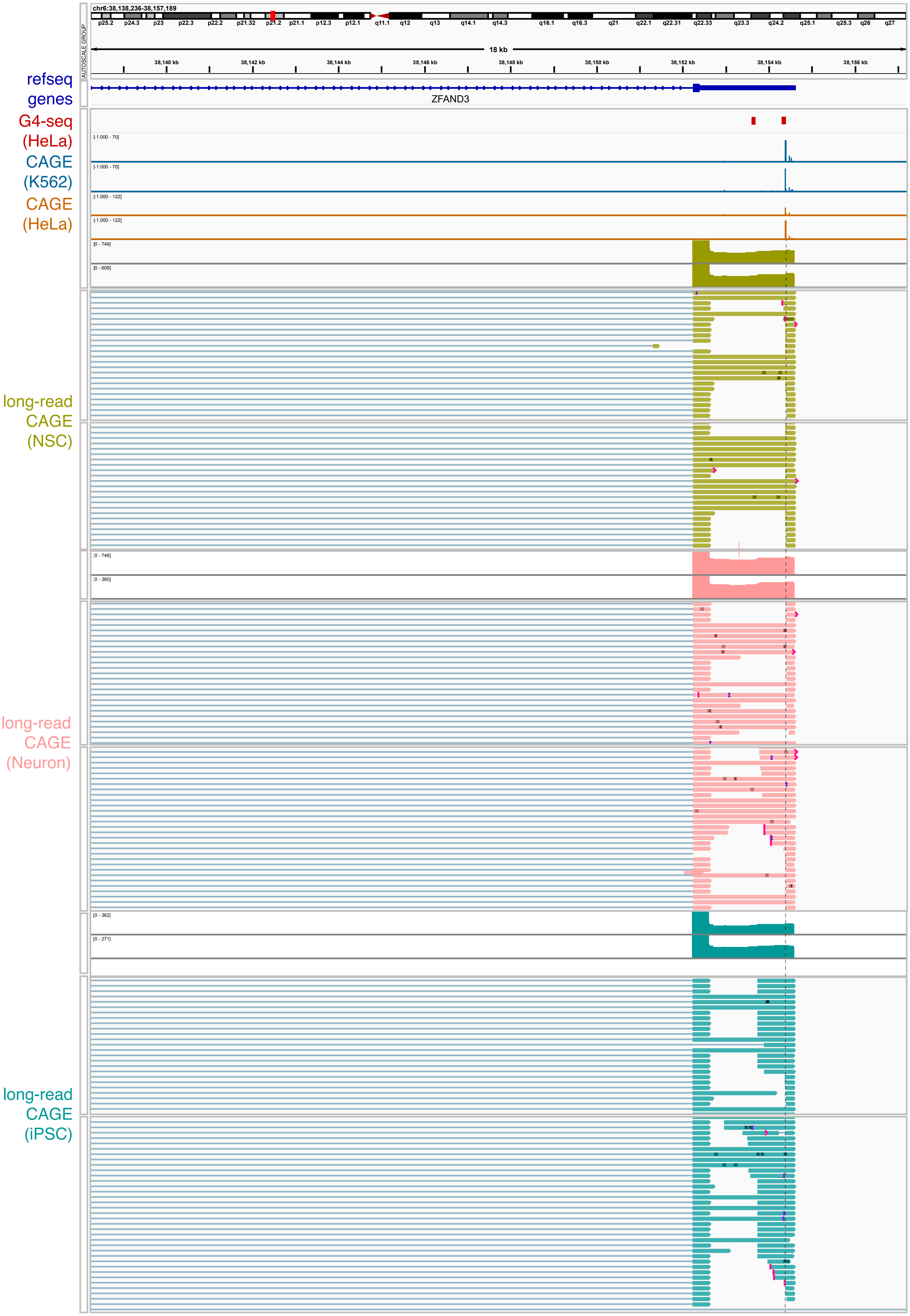

chr6:38,138,236-38,157,189
18 kb
38,140 kb
38,142 kb
38,144 kb
38,146 kb
38,148 kb
38,150 kb
38,152 kb
38,154 kb
38,156 kb
AUTOSCALE GROUP
ZFAND3
refseq genes
G4-seq (HeLa)
CAGE (K562)
CAGE (HeLa)
long-read CAGE (NSC)
long-read CAGE (Neuron)
long-read CAGE (iPSC)
