## Supplementary figures and images for "Abundant capped RNAs are derived from mRNA cleavage at 3’UTR G-Quadruplexes"

### Supplementary figure S2

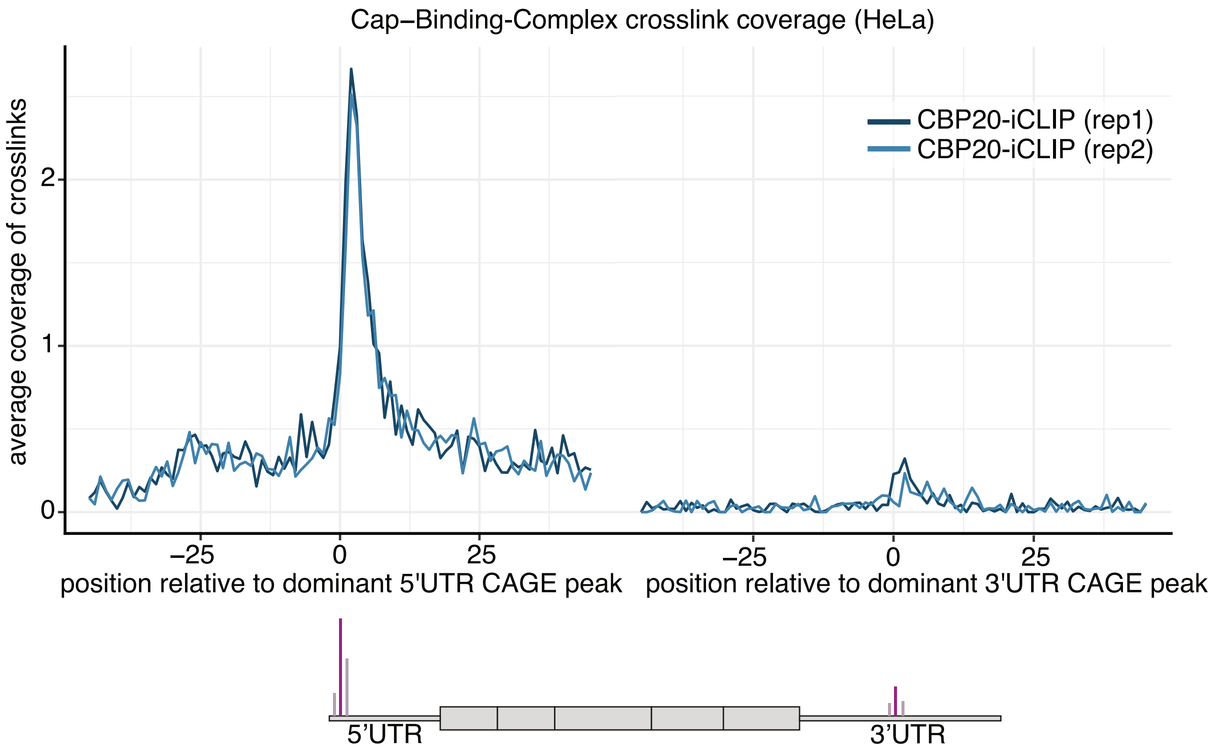

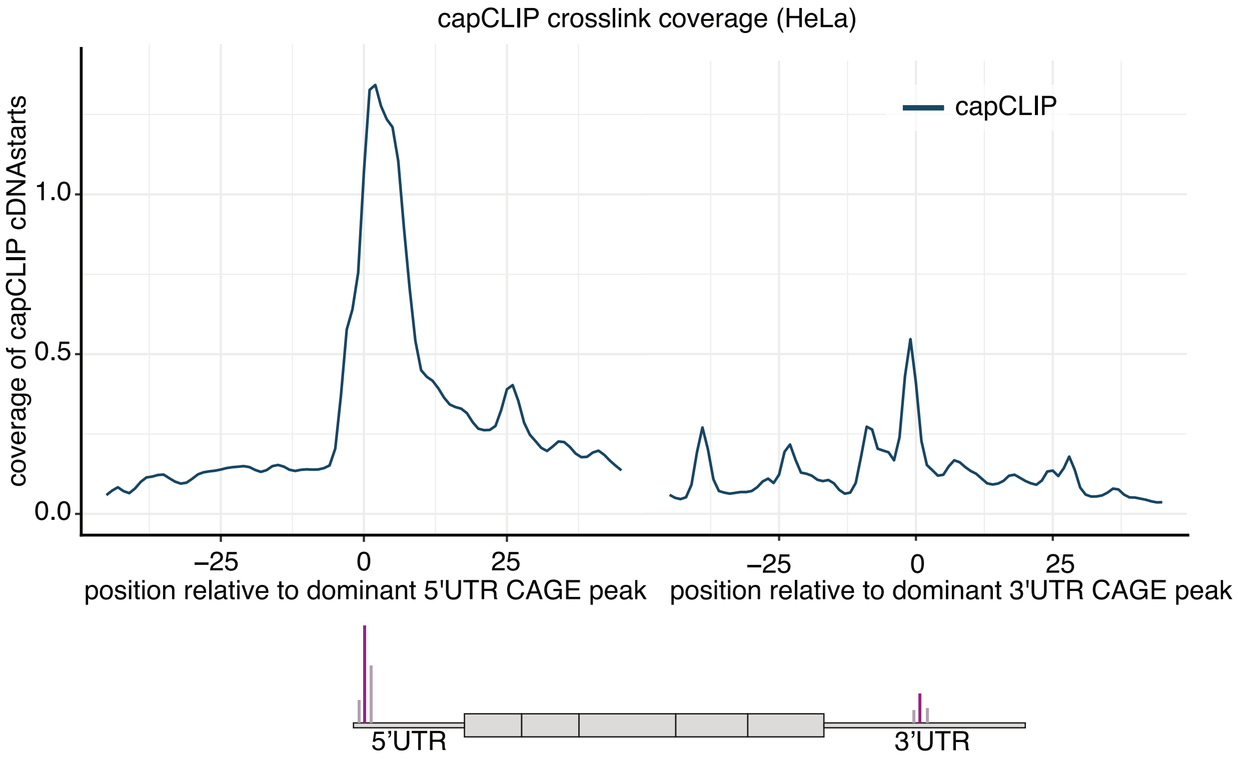

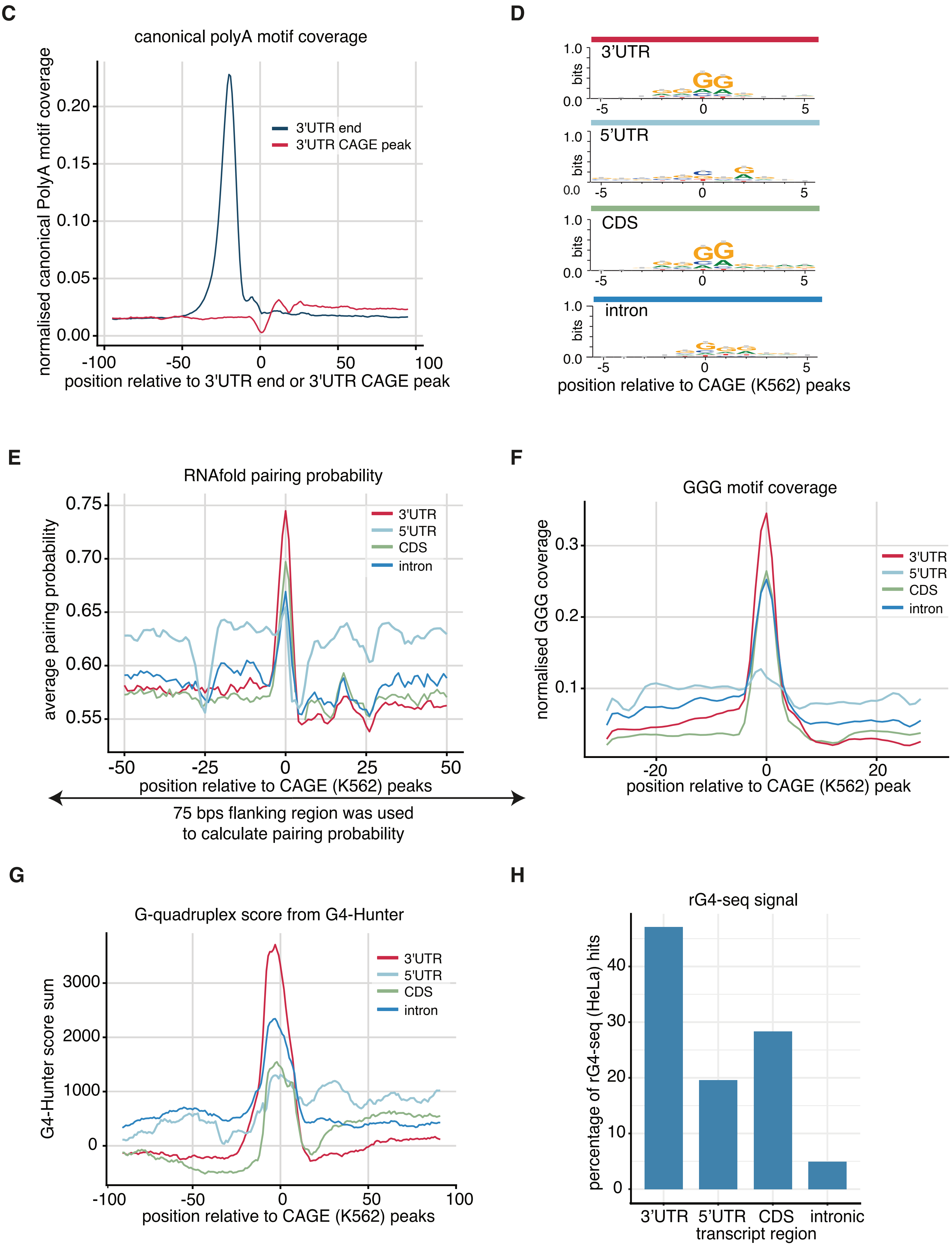

### Supplementary figure S4

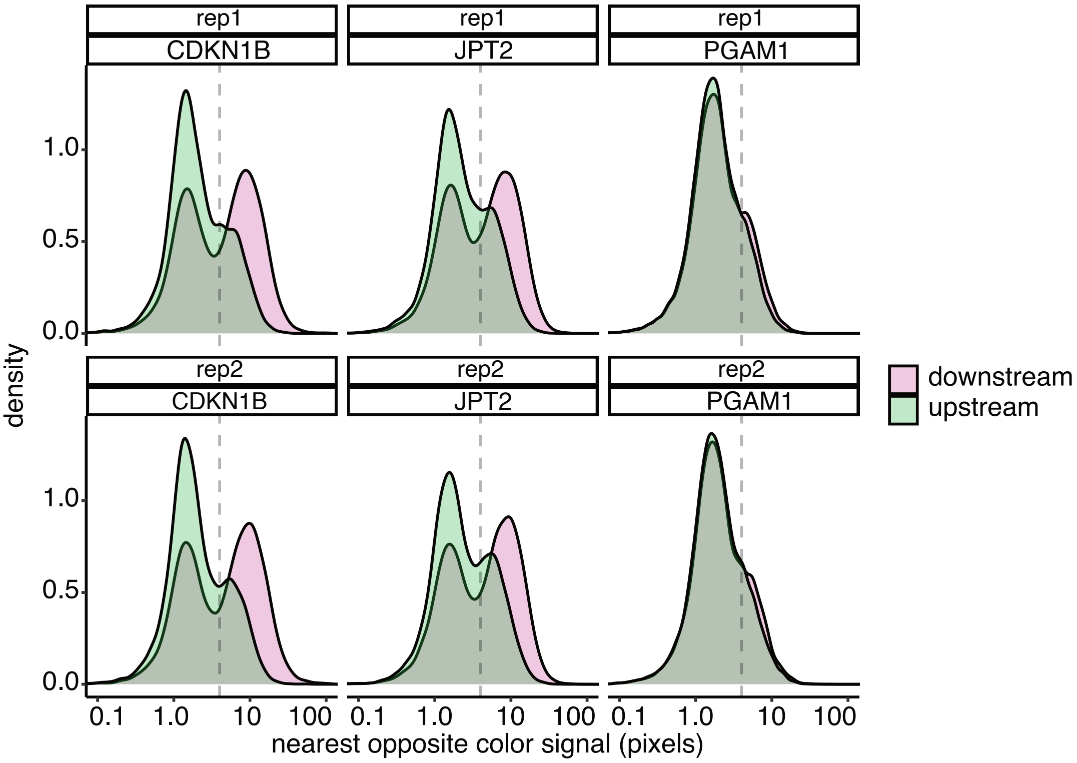
