## Supplementary figure S3 for "Abundant capped RNAs are derived from mRNA cleavage at 3’UTR G-Quadruplexes"

**A**

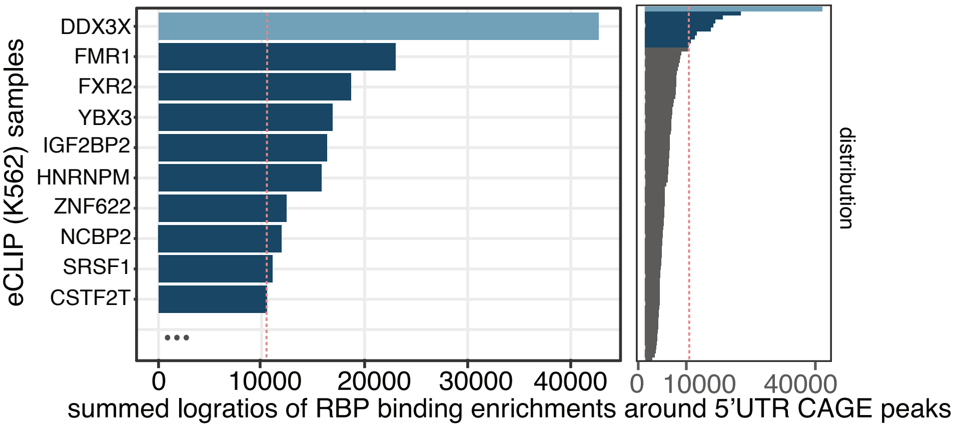


**B** K562: CAGE, RNA-seq and UPF1-eCLIP

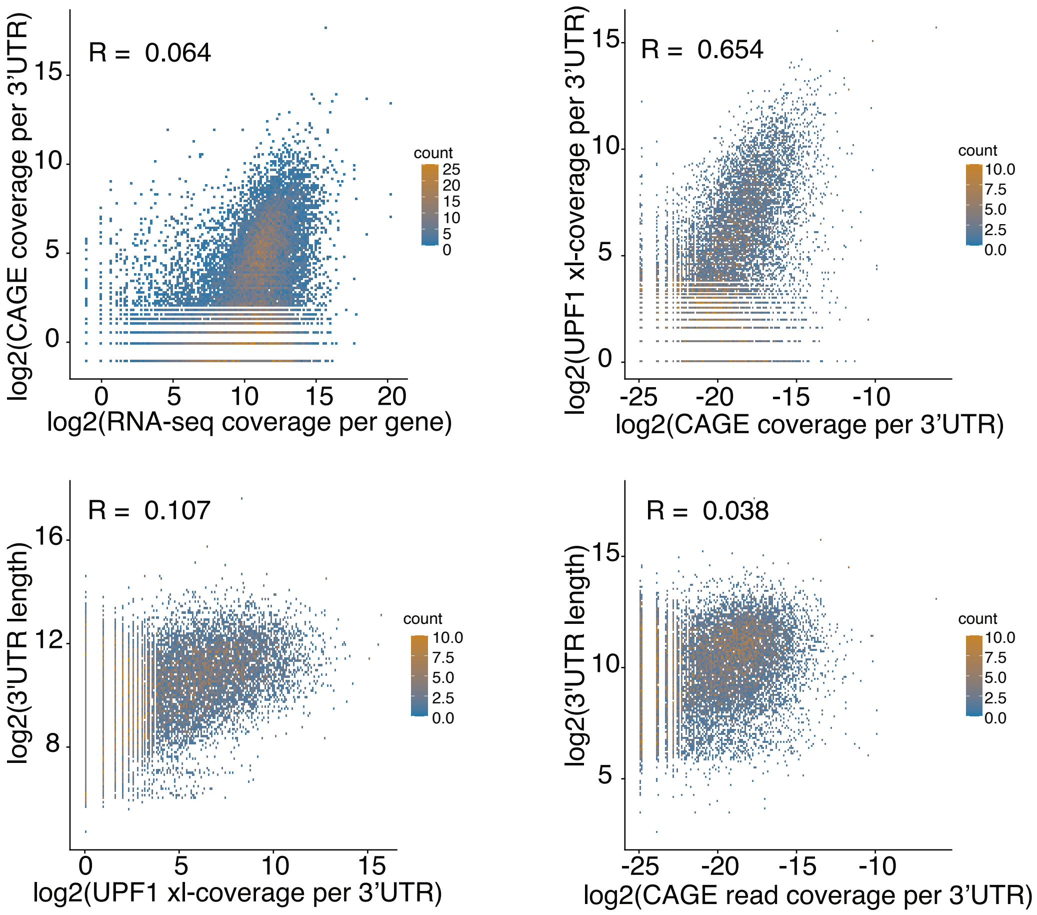


**C**

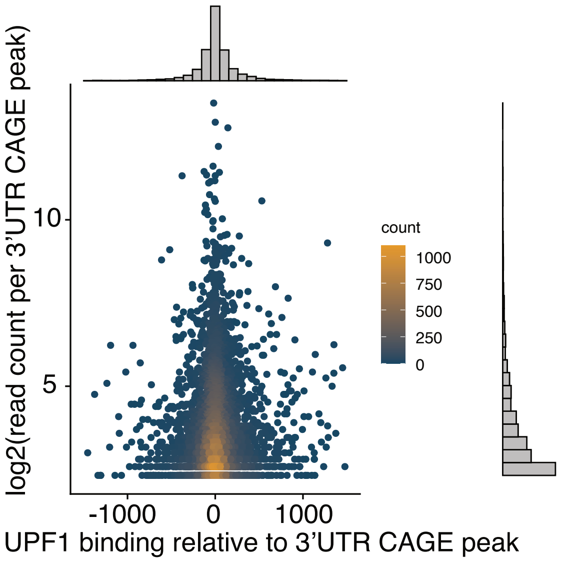

D

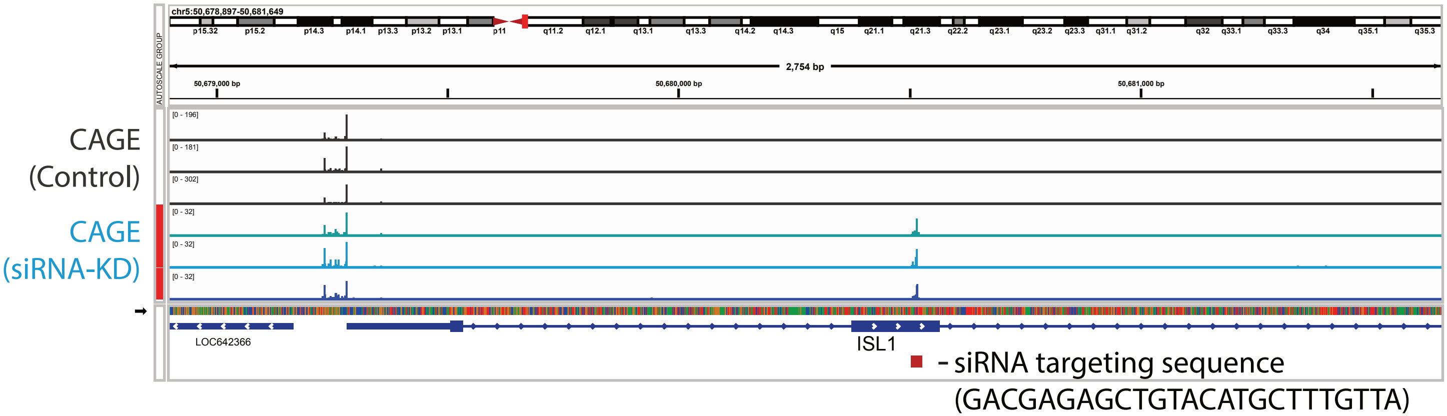
chr5:50,678,897-50,681,649
AUTOSCALE GROUP
p15.32 p15.2 p14.3 p14.1 p13.3 p13.2 p13.1 p11 q11.2 q12.1 q13.1 q13.3 q14.2 q14.3 q15 q21.1 q21.3 q22.2 q23.1 q23.2 q23.3 q31.1 q31.2 q32 q33.1 q33.3 q34 q35.1 q35.3
2,754 bp
50,679,000 bp
50,680,000 bp
50,681,000 bp
CAGE (Control)
CAGE (siRNA-KD)
[0 - 196]
[0 - 181]
[0 - 302]
[0 - 32]
[0 - 32]
[0 - 32]
LOC642366
ISL1
- siRNA targeting sequence (GACGAGAGCTGTACATGCTTTGTTA)

E

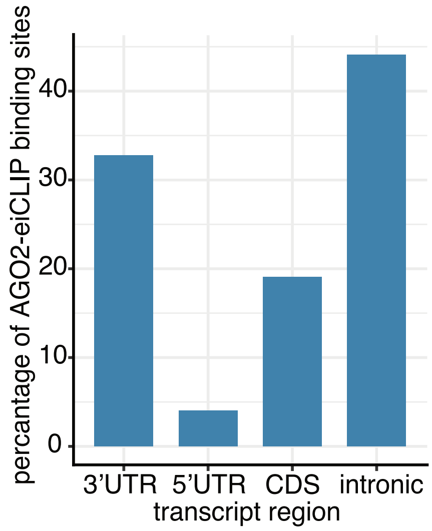


F

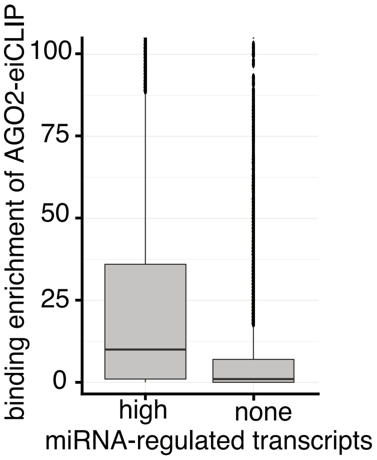


G

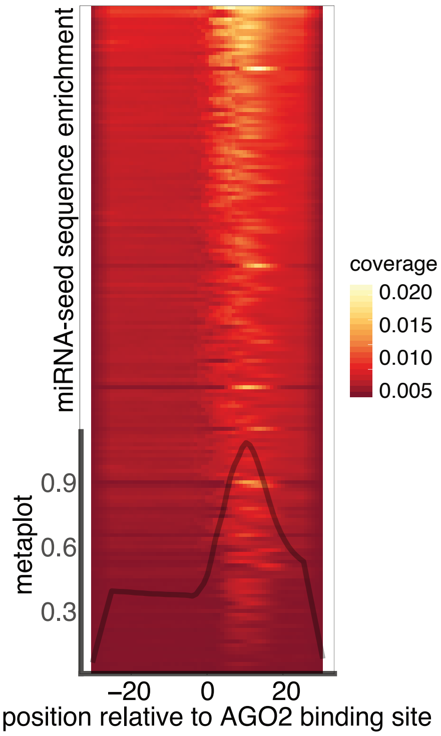


H

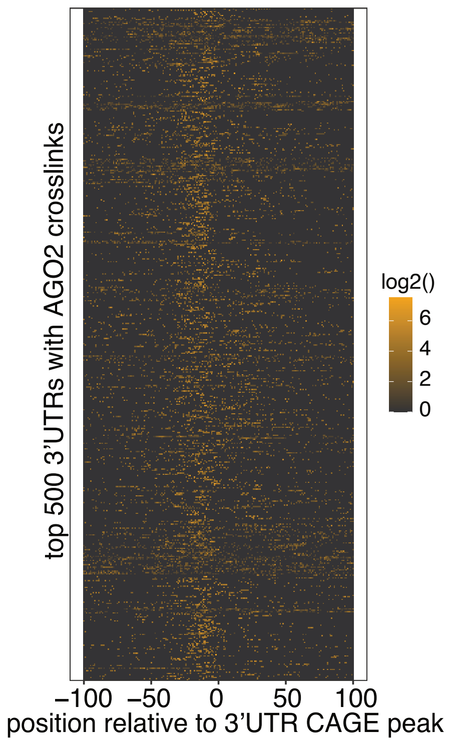


I

Homer de novo Motif Results
Total target sequences = 172773
Total background sequences = 177500
* - possible false positive

| Rank | Motif | P-value | log P-pvalue | % of Targets | % of Background | STD(Bg STD) | Best Match/Details |
|---|---|---|---|---|---|---|---|
| 1 | GUCGAAAU | 1e-5254 | -1.210e+04 | 1.06% | 0.00% | 1.0bp (0.0bp) | hsa-miR-1305 MIMAT0005893 Homo sapiens miR-1305 Targets (miRBase)(0.637) More Information \| Similar Motifs Found |
| 2 | GGUGAGGU | 1e-2229 | -5.133e+03 | 1.19% | 0.04% | 1.7bp (2.9bp) | hsa-miR-4777-3p MIMAT0019935 Homo sapiens miR-4777-3p Targets (miRBase)(0.743) More Information \| Similar Motifs Found |
| 3 | CGUGGGGG | 1e-817 | -1.882e+03 | 18.44% | 13.19% | 2.5bp (3.1bp) | hsa-miR-361-3p MIMAT0004682 Homo sapiens miR-361-3p Targets (miRBase)(0.701) More Information \| Similar Motifs Found |
| 4 | UUAAAGCA | 1e-759 | -1.749e+03 | 0.45% | 0.02% | 0.8bp (2.1bp) | hsa-miR-1273e MIMAT0018079 Homo sapiens miR-1273e Targets (miRBase)(0.699) More Information \| Similar Motifs Found |
| 5 | CCAGAAGA | 1e-689 | -1.589e+03 | 12.21% | 8.24% | 2.4bp (2.8bp) | hsa-miR-4659b-3p MIMAT0019734 Homo sapiens miR-4659b-3p Targets (miRBase)(0.684) More Information \| Similar Motifs Found |
| 6 | UAAUAUUG | 1e-331 | -7.632e+02 | 5.13% | 3.32% | 2.7bp (2.5bp) | hsa-miR-16-2* MIMAT0004518 Homo sapiens miR-16-2* Targets (miRBase)(0.724) More Information \| Similar Motifs Found |
| 7 | CCACUAAU | 1e-244 | -5.623e+02 | 17.25% | 14.35% | 2.4bp (2.2bp) | hsa-miR-4727-3p MIMAT0019848 Homo sapiens miR-4727-3p Targets (miRBase)(0.676) More Information \| Similar Motifs Found |
| 8 | UUGUUUGU | 1e-135 | -3.119e+02 | 0.97% | 0.50% | 2.7bp (2.7bp) | hsa-miR-495 MIMAT0002817 Homo sapiens miR-495 Targets (miRBase)(0.665) More Information \| Similar Motifs Found |
| 9 | CCCCCCCC | 1e-107 | -2.479e+02 | 4.08% | 3.12% | 2.4bp (2.0bp) | hsa-miR-4488 MIMAT0019022 Homo sapiens miR-4488 Targets (miRBase)(0.743) More Information \| Similar Motifs Found |
| 10 | AAACCCGU | 1e-79 | -1.824e+02 | 0.04% | 0.00% | 1.0bp (1.6bp) | hsa-miR-125b-1* MIMAT0004592 Homo sapiens miR-125b-1* Targets (miRBase)(0.699) More Information \| Similar Motifs Found |
| 11 | CUAAACGC | 1e-75 | -1.743e+02 | 0.03% | 0.00% | 1.3bp (1.5bp) | hsa-miR-629 MIMAT0004810 Homo sapiens miR-629 Targets (miRBase)(0.642) More Information \| Similar Motifs Found |
| 12 | GCAGGGAG | 1e-70 | -1.618e+02 | 0.25% | 0.09% | 2.3bp (2.9bp) | hsa-miR-3620 MIMAT0018001 Homo sapiens miR-3620 Targets (miRBase)(0.716) More Information \| Similar Motifs Found |

J

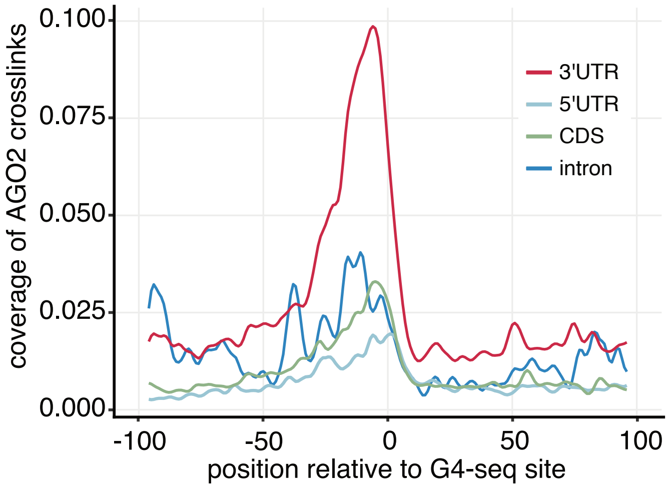
0.100
0.075
0.050
0.025
0.000
coverage of AGO2 crosslinks
3'UTR
5'UTR
CDS
intron
-100
-50
0
50
100
position relative to G4-seq site

K

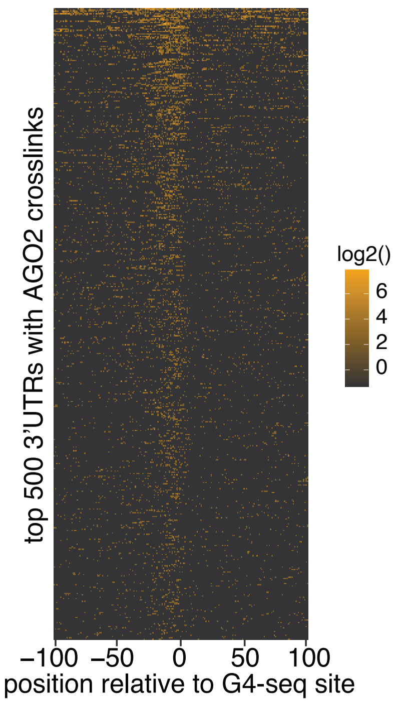
top 500 3'UTRs with AGO2 crosslinks
log2()
6
4
2
0
−100 −50 0 50 100
position relative to G4-seq site
