## Supplementary Table 1 for "Abundant capped RNAs are derived from mRNA cleavage at 3’UTR G-Quadruplexes"

| chromosome | CAGE.3UTR.cluster.start | CAGE.3UTR.cluster.end | CAGE.3UTR.strand | CAGE.HeLa.norm.read.counts.rep1 | CAGE.HeLa.norm.read.counts.rep2 | CAGE.K562.norm.read.counts.rep1 | CAGE.K562.norm.read.counts.rep2 |
| --- | --- | --- | --- | --- | --- | --- | --- |
| chr1 | 827709 | 827728 | + | 29.1148567293276 | 52.3018041349616 | 14.3187385244159 | 10.6710864697714 |
| chr1 | 827751 | 827762 | + | 5.01980288436682 | 4.54798296825753 | 3.57968463110397 | 6.40265188186281 |
| chr1 | 944223 | 944229 | - | 1.00396057687336 | 1.13699574206438 | 1.19322821036799 | 4.26843458790854 |
| chr1 | 944317 | 944394 | + | 29.1148567293276 | 30.6988850357383 | 2.38645642073598 | 1.06710864697714 |
| chr1 | 944335 | 944357 | - | 3.01188173062009 | 3.41098722619315 | 2.38645642073598 | 4.26843458790854 |
| chr1 | 944533 | 944596 | - | 73.2891221117556 | 97.7816338175368 | 297.113824381629 | 336.139223797798 |
| chr1 | 944627 | 944661 | - | 18.0712903837206 | 6.82197445238629 | 42.9562155732476 | 28.8119334683827 |
| chr1 | 998982 | 999032 | - | 43.1703048055547 | 96.6446380754725 | 27.4442488384638 | 28.8119334683827 |
| chr1 | 1014449 | 1014521 | + | 55.217831728035 | 51.1648083928972 | 14.3187385244159 | 42.6843458790854 |
| chr1 | 1054979 | 1055078 | + | 35.1386201905678 | 11.3699574206438 | 40.5697591525116 | 11.7381951167485 |
| chr1 | 1055116 | 1055153 | + | 2.00792115374673 | 1.13699574206438 | 9.54582568294392 | 5.33554323488568 |
| chr1 | 1055175 | 1055218 | + | 2.00792115374673 | 4.54798296825753 | 8.35259747257593 | 2.13421729395427 |
| chr1 | 1055335 | 1055415 | + | 9.03564519186028 | 6.82197445238629 | 10.7390538933119 | 17.0737383516342 |
| chr1 | 1056010 | 1056087 | + | 88.3485307648561 | 52.3018041349616 | 48.9223566250876 | 74.6976052883995 |
| chr1 | 1081850 | 1081860 | - | 6.02376346124019 | 9.09596593651506 | 5.96614105183995 | 14.9395210576799 |
| chr1 | 1216933 | 1217108 | - | 73.2891221117556 | 71.6307317500561 | 35.7968463110397 | 68.2949534065367 |
| chr1 | 1217139 | 1217155 | - | 8.03168461498692 | 6.82197445238629 | 3.57968463110397 | 3.20132594093141 |
| chr1 | 1233530 | 1233574 | + | 57.2257528817818 | 32.9728765198671 | 26.2510206280958 | 52.2883237018796 |
| chr1 | 1233637 | 1233693 | + | 5.01980288436682 | 9.09596593651506 | 5.96614105183995 | 3.20132594093141 |
| chr1 | 1254031 | 1254056 | - | 4.01584230749346 | 2.27399148412876 | 1.19322821036799 | 6.40265188186281 |
| chr1 | 1254931 | 1255002 | - | 47.1861471130481 | 52.3018041349616 | 36.9900745214077 | 57.6238669367653 |
| chr1 | 1311542 | 1311594 | + | 37.1465413443145 | 40.9318467143178 | 40.5697591525116 | 51.2212150549025 |
| chr1 | 1311608 | 1311671 | - | 24.0950538449607 | 40.9318467143178 | 69.2072362013434 | 53.3554323488568 |
| chr1 | 1311734 | 1311744 | - | 3.01188173062009 | 3.41098722619315 | 5.96614105183995 | 11.7381951167485 |
| chr1 | 1311853 | 1311855 | - | 1.00396057687336 | 1.13699574206438 | 1.19322821036799 | 2.13421729395427 |
| chr1 | 1318780 | 1318810 | - | 4.01584230749346 | 6.82197445238629 | 5.96614105183995 | 2.13421729395427 |
| chr1 | 1335316 | 1335335 | - | 1.00396057687336 | 1.13699574206438 | 1.19322821036799 | 1.06710864697714 |
| chr1 | 1335356 | 1335548 | - | 53.2099105742883 | 23.876910583352 | 9.54582568294392 | 24.5434988804741 |
| chr1 | 1335749 | 1335815 | - | 25.0990144218341 | 14.780944646837 | 25.0577924177278 | 38.4159112911769 |
| chr1 | 1335844 | 1335897 | - | 14.0554480762271 | 17.0549361309657 | 21.4781077866238 | 13.8724124107028 |
| chr1 | 1335945 | 1335955 | - | 9.03564519186028 | 4.54798296825753 | 2.38645642073598 | 3.20132594093141 |
| chr1 | 1336021 | 1336037 | - | 27.1069355755808 | 15.9179403889013 | 45.3426719939836 | 20.2750642925656 |
| chr1 | 1336085 | 1336107 | - | 1.00396057687336 | 1.13699574206438 | 8.35259747257593 | 1.06710864697714 |
| chr1 | 1373758 | 1373905 | - | 701.768443234482 | 401.359496948727 | 283.988314067581 | 846.217157052869 |
| chr1 | 1386522 | 1386582 | - | 10.0396057687336 | 11.3699574206438 | 10.7390538933119 | 7.46976052883995 |
| chr1 | 1386604 | 1386659 | - | 4.01584230749346 | 3.41098722619315 | 9.54582568294392 | 1.06710864697714 |
| chr1 | 1386708 | 1386789 | - | 2.00792115374673 | 13.6439489047726 | 7.15936926220794 | 10.6710864697714 |
| chr1 | 1386809 | 1386852 | - | 1.00396057687336 | 1.13699574206438 | 2.38645642073598 | 3.20132594093141 |
| chr1 | 1386887 | 1386919 | - | 3.01188173062009 | 2.27399148412876 | 2.38645642073598 | 2.13421729395427 |
| chr1 | 1386944 | 1387138 | - | 9.03564519186028 | 9.09596593651506 | 17.8984231555198 | 20.2750642925656 |
| chr1 | 1387187 | 1387268 | - | 7.02772403811355 | 5.68497871032191 | 15.5119667347839 | 19.2079556455884 |
| chr1 | 1392572 | 1392647 | - | 21.0831721143407 | 13.6439489047726 | 10.7390538933119 | 7.46976052883995 |
| chr1 | 1392674 | 1392697 | - | 6.02376346124019 | 2.27399148412876 | 1.19322821036799 | 5.33554323488568 |
| chr1 | 1392716 | 1392755 | - | 6.02376346124019 | 4.54798296825753 | 4.77291284147196 | 2.13421729395427 |
| chr1 | 1399513 | 1399621 | + | 353.394123059424 | 368.38662042886 | 184.950372607038 | 308.394398976392 |
| chr1 | 1400231 | 1400243 | + | 5.01980288436682 | 6.82197445238629 | 2.38645642073598 | 4.26843458790854 |
| chr1 | 1401136 | 1401204 | + | 3.01188173062009 | 14.780944646837 | 36.9900745214077 | 30.9461507623369 |
| chr1 | 1401232 | 1401264 | + | 1.00396057687336 | 1.13699574206438 | 2.38645642073598 | 9.60397782279422 |
| chr1 | 1401328 | 1401335 | + | 1.00396057687336 | 3.41098722619315 | 4.77291284147196 | 1.06710864697714 |
| chr1 | 1401476 | 1401538 | + | 14.0554480762271 | 31.8358807778027 | 25.0577924177278 | 32.0132594093141 |
| chr1 | 1401806 | 1401817 | + | 4.01584230749346 | 3.41098722619315 | 1.19322821036799 | 4.26843458790854 |
| chr1 | 1401839 | 1401871 | + | 2.00792115374673 | 1.13699574206438 | 1.19322821036799 | 4.26843458790854 |
| chr1 | 1401929 | 1401987 | - | 12.0475269224804 | 28.4248935516096 | 10.7390538933119 | 23.476390233497 |
| chr1 | 1402049 | 1402101 | - | 16.0633692299738 | 10.2329616785794 | 16.7051949451519 | 17.0737383516342 |
| chr1 | 1409645 | 1409647 | + | 2.00792115374673 | 1.13699574206438 | 2.38645642073598 | 9.60397782279422 |
| chr1 | 1418979 | 1418981 | - | 25.0990144218341 | 26.1509020674808 | 42.9562155732476 | 40.5501285851312 |
| chr1 | 1496141 | 1496172 | + | 16.0633692299738 | 12.5069531627082 | 9.54582568294392 | 4.26843458790854 |
| chr1 | 1534575 | 1534597 | + | 14.0554480762271 | 11.3699574206438 | 4.77291284147196 | 3.20132594093141 |
| chr1 | 1534639 | 1534656 | + | 30.1188173062009 | 39.7948509722534 | 15.5119667347839 | 21.3421729395427 |
| chr1 | 1541700 | 1541750 | - | 2150.48355566275 | 1646.36983450923 | 1285.10678256632 | 1977.35232284863 |
| chr1 | 1541770 | 1541835 | - | 45.1782259593014 | 42.0688424563821 | 21.4781077866238 | 39.483019938154 |
| chr1 | 1541846 | 1541872 | - | 36.1425807674411 | 44.3428339405109 | 26.2510206280958 | 28.8119334683827 |
| chr1 | 1541899 | 1541978 | - | 77.3049644192491 | 76.1787147183136 | 95.4582568294392 | 126.985928990279 |
| chr1 | 1542025 | 1542159 | - | 58.2297134586551 | 37.5208594881246 | 36.9900745214077 | 76.8318225823538 |
| chr1 | 1562393 | 1562398 | - | 1.00396057687336 | 1.13699574206438 | 1.19322821036799 | 4.26843458790854 |
| chr1 | 1661570 | 1661576 | - | 4.01584230749346 | 2.27399148412876 | 1.19322821036799 | 1.06710864697714 |
| chr1 | 1662189 | 1662199 | - | 2.00792115374673 | 1.13699574206438 | 3.57968463110397 | 2.13421729395427 |
| chr1 | 1752208 | 1752225 | - | 3.01188173062009 | 1.13699574206438 | 1.19322821036799 | 1.06710864697714 |
| chr1 | 1752240 | 1752243 | - | 1.00396057687336 | 1.13699574206438 | 2.38645642073598 | 2.13421729395427 |
| chr1 | 1752290 | 1752315 | - | 3.01188173062009 | 1.13699574206438 | 7.15936926220794 | 9.60397782279422 |
| chr1 | 1752751 | 1752852 | - | 28.1108961524542 | 18.1919318730301 | 40.5697591525116 | 28.8119334683827 |
| chr1 | 1752900 | 1752953 | - | 10.0396057687336 | 10.2329616785794 | 26.2510206280958 | 21.3421729395427 |
| chr1 | 1785316 | 1785364 | - | 6.02376346124019 | 12.5069531627082 | 17.8984231555198 | 9.60397782279422 |
| chr1 | 1785466 | 1785512 | - | 7.02772403811355 | 4.54798296825753 | 16.7051949451519 | 9.60397782279422 |
| chr1 | 1785554 | 1785555 | - | 2.00792115374673 | 2.27399148412876 | 1.19322821036799 | 1.06710864697714 |
| chr1 | 1785739 | 1785768 | - | 42.1663442286813 | 61.3977700714766 | 101.424397881279 | 93.9055609339879 |
| chr1 | 1785825 | 1785871 | - | 152.602007684751 | 90.9596593651506 | 165.858721241151 | 87.5029090521251 |
| chr1 | 1785907 | 1785940 | - | 2.00792115374673 | 2.27399148412876 | 7.15936926220794 | 6.40265188186281 |
| chr1 | 1785972 | 1786063 | - | 64.2534769198953 | 62.534765813541 | 106.197310722751 | 77.8989312293309 |
| chr1 | 1786129 | 1786196 | - | 21.0831721143407 | 15.9179403889013 | 54.8884976769275 | 68.2949534065367 |
| chr1 | 1786233 | 1786248 | - | 32.1267384599477 | 21.6029190992233 | 40.5697591525116 | 51.2212150549025 |
| chr1 | 1786279 | 1786303 | - | 1.00396057687336 | 3.41098722619315 | 9.54582568294392 | 6.40265188186281 |
| chr1 | 1786356 | 1786388 | - | 3.01188173062009 | 3.41098722619315 | 9.54582568294392 | 12.8053037637256 |
| chr1 | 1786470 | 1786528 | - | 6.02376346124019 | 2.27399148412876 | 8.35259747257593 | 8.53686917581709 |
| chr1 | 1786601 | 1786728 | - | 24.0950538449607 | 11.3699574206438 | 26.2510206280958 | 13.8724124107028 |
| chr1 | 1786782 | 1786798 | - | 5.01980288436682 | 17.0549361309657 | 11.9322821036799 | 2.13421729395427 |
| chr1 | 1786821 | 1786882 | - | 52.205949997415 | 57.9867828452835 | 71.5936926220794 | 125.918820343302 |
| chr1 | 1786911 | 1787053 | - | 28.1108961524542 | 12.5069531627082 | 51.3088130458235 | 46.952780466994 |
| chr1 | 2184543 | 2184561 | - | 3.01188173062009 | 1.13699574206438 | 2.38645642073598 | 2.13421729395427 |
| chr1 | 2189567 | 2189584 | - | 59.2336740355285 | 43.2058381984465 | 71.5936926220794 | 136.589906813073 |
| chr1 | 2306812 | 2306841 | + | 3.01188173062009 | 1.13699574206438 | 3.57968463110397 | 2.13421729395427 |
| chr1 | 2307181 | 2307210 | + | 12.0475269224804 | 2.27399148412876 | 9.54582568294392 | 4.26843458790854 |
| chr1 | 2307232 | 2307254 | + | 3.01188173062009 | 1.13699574206438 | 5.96614105183995 | 7.46976052883995 |
| chr1 | 2307558 | 2307560 | + | 9.03564519186028 | 2.27399148412876 | 2.38645642073598 | 3.20132594093141 |
| chr1 | 2307600 | 2307679 | + | 15.0594086531005 | 10.2329616785794 | 23.8645642073598 | 35.2145853502455 |
| chr1 | 2307904 | 2307907 | + | 5.01980288436682 | 6.82197445238629 | 7.15936926220794 | 6.40265188186281 |
| chr1 | 2308409 | 2308428 | + | 5.01980288436682 | 2.27399148412876 | 3.57968463110397 | 1.06710864697714 |
| chr1 | 2308516 | 2308521 | + | 2.00792115374673 | 1.13699574206438 | 2.38645642073598 | 9.60397782279422 |
| chr1 | 2309600 | 2309602 | + | 1.00396057687336 | 3.41098722619315 | 9.54582568294392 | 1.06710864697714 |
| chr1 | 2309858 | 2309927 | + | 37.1465413443145 | 39.7948509722534 | 125.288962088639 | 165.401840281456 |
| chr1 | 2309952 | 2309967 | + | 17.0673298068472 | 9.09596593651506 | 7.15936926220794 | 9.60397782279422 |
| chr1 | 2309986 | 2309997 | + | 8.03168461498692 | 3.41098722619315 | 1.19322821036799 | 6.40265188186281 |
| chr1 | 2310021 | 2310056 | + | 2.00792115374673 | 4.54798296825753 | 2.38645642073598 | 1.06710864697714 |
| chr1 | 2403072 | 2403091 | + | 3.01188173062009 | 1.13699574206438 | 2.38645642073598 | 2.13421729395427 |
| chr1 | 2403537 | 2403624 | + | 52.205949997415 | 62.534765813541 | 22.6713359969918 | 41.6172372321083 |
| chr1 | 2403646 | 2403664 | + | 11.043566345607 | 6.82197445238629 | 10.7390538933119 | 7.46976052883995 |
| chr1 | 2403687 | 2403728 | + | 82.3247673036159 | 98.9186295596012 | 36.9900745214077 | 78.966039876308 |
| chr1 | 2404139 | 2404154 | - | 21.0831721143407 | 13.6439489047726 | 9.54582568294392 | 17.0737383516342 |
| chr1 | 2404206 | 2404208 | + | 5.01980288436682 | 1.13699574206438 | 3.57968463110397 | 1.06710864697714 |
| chr1 | 2405012 | 2405035 | - | 4.01584230749346 | 3.41098722619315 | 3.57968463110397 | 1.06710864697714 |
| chr1 | 2405035 | 2405091 | + | 12.0475269224804 | 2.27399148412876 | 8.35259747257593 | 8.53686917581709 |
| chr1 | 2405217 | 2405229 | + | 4.01584230749346 | 2.27399148412876 | 4.77291284147196 | 1.06710864697714 |
| chr1 | 2405386 | 2405395 | + | 3.01188173062009 | 5.68497871032191 | 1.19322821036799 | 2.13421729395427 |
| chr1 | 3628673 | 3628692 | + | 12.0475269224804 | 5.68497871032191 | 2.38645642073598 | 1.06710864697714 |
| chr1 | 3628728 | 3628778 | + | 9.03564519186028 | 6.82197445238629 | 5.96614105183995 | 1.06710864697714 |
| chr1 | 3629972 | 3630083 | + | 12.0475269224804 | 7.95897019445067 | 1.19322821036799 | 3.20132594093141 |
| chr1 | 3630864 | 3630924 | - | 18.0712903837206 | 35.2468680039958 | 11.9322821036799 | 17.0737383516342 |
| chr1 | 3630944 | 3630947 | - | 3.01188173062009 | 2.27399148412876 | 1.19322821036799 | 1.06710864697714 |
| chr1 | 3780251 | 3780273 | - | 6.02376346124019 | 4.54798296825753 | 4.77291284147196 | 9.60397782279422 |
| chr1 | 3780297 | 3780422 | - | 238.942617295861 | 200.111250603331 | 336.490355323773 | 282.783791448941 |
| chr1 | 3780464 | 3780483 | - | 7.02772403811355 | 5.68497871032191 | 17.8984231555198 | 16.006629704657 |
| chr1 | 3780516 | 3780564 | - | 7.02772403811355 | 2.27399148412876 | 13.1255103140479 | 12.8053037637256 |
| chr1 | 3780588 | 3780678 | - | 38.1505019211879 | 21.6029190992233 | 66.8207797806074 | 62.959410171651 |
| chr1 | 3780725 | 3780731 | - | 8.03168461498692 | 1.13699574206438 | 1.19322821036799 | 1.06710864697714 |
| chr1 | 3780848 | 3780860 | - | 2.00792115374673 | 5.68497871032191 | 1.19322821036799 | 5.33554323488568 |
| chr1 | 3884408 | 3884410 | + | 1.00396057687336 | 1.13699574206438 | 5.96614105183995 | 2.13421729395427 |
| chr1 | 3884518 | 3884521 | + | 20.0792115374673 | 11.3699574206438 | 28.6374770488317 | 23.476390233497 |
| chr1 | 3885241 | 3885264 | + | 7.02772403811355 | 9.09596593651506 | 1.19322821036799 | 21.3421729395427 |
| chr1 | 3889199 | 3889203 | - | 7.02772403811355 | 7.95897019445067 | 7.15936926220794 | 9.60397782279422 |
| chr1 | 3889339 | 3889362 | - | 20.0792115374673 | 14.780944646837 | 10.7390538933119 | 16.006629704657 |
| chr1 | 3889412 | 3889418 | - | 3.01188173062009 | 1.13699574206438 | 2.38645642073598 | 5.33554323488568 |
| chr1 | 3889786 | 3889804 | - | 3.01188173062009 | 1.13699574206438 | 14.3187385244159 | 8.53686917581709 |
| chr1 | 3889957 | 3889993 | - | 4.01584230749346 | 2.27399148412876 | 7.15936926220794 | 8.53686917581709 |
| chr1 | 3900393 | 3900442 | + | 6.02376346124019 | 10.2329616785794 | 3.57968463110397 | 6.40265188186281 |
| chr1 | 5863009 | 5863026 | - | 4.01584230749346 | 2.27399148412876 | 1.19322821036799 | 3.20132594093141 |
| chr1 | 6100210 | 6100233 | + | 10.0396057687336 | 2.27399148412876 | 1.19322821036799 | 1.06710864697714 |
| chr1 | 6185040 | 6185077 | - | 16.0633692299738 | 7.95897019445067 | 1.19322821036799 | 1.06710864697714 |
| chr1 | 6185106 | 6185269 | - | 15.0594086531005 | 28.4248935516096 | 25.0577924177278 | 9.60397782279422 |
| chr1 | 6185472 | 6185496 | - | 2.00792115374673 | 4.54798296825753 | 1.19322821036799 | 2.13421729395427 |
| chr1 | 6186306 | 6186326 | - | 9.03564519186028 | 15.9179403889013 | 5.96614105183995 | 6.40265188186281 |
| chr1 | 6220690 | 6220710 | + | 5.01980288436682 | 7.95897019445067 | 8.35259747257593 | 4.26843458790854 |
| chr1 | 6221314 | 6221333 | - | 23.0910932680874 | 29.5618892936739 | 11.9322821036799 | 16.006629704657 |
| chr1 | 6221382 | 6221401 | - | 5.01980288436682 | 3.41098722619315 | 1.19322821036799 | 3.20132594093141 |
| chr1 | 6221433 | 6221515 | - | 43.1703048055547 | 21.6029190992233 | 10.7390538933119 | 7.46976052883995 |
| chr1 | 6221559 | 6221578 | - | 5.01980288436682 | 3.41098722619315 | 2.38645642073598 | 1.06710864697714 |
| chr1 | 6221651 | 6221670 | - | 39.1544624980612 | 40.9318467143178 | 15.5119667347839 | 9.60397782279422 |
| chr1 | 6221789 | 6221800 | - | 4.01584230749346 | 3.41098722619315 | 3.57968463110397 | 6.40265188186281 |
| chr1 | 6221982 | 6222041 | - | 15.0594086531005 | 12.5069531627082 | 10.7390538933119 | 9.60397782279422 |
| chr1 | 6222089 | 6222123 | - | 25.0990144218341 | 17.0549361309657 | 14.3187385244159 | 9.60397782279422 |
| chr1 | 6222146 | 6222159 | - | 70.2772403811355 | 48.8908169087684 | 54.8884976769275 | 48.0198891139711 |
| chr1 | 6223605 | 6223627 | - | 6.02376346124019 | 17.0549361309657 | 3.57968463110397 | 3.20132594093141 |
| chr1 | 6223669 | 6223681 | - | 16.0633692299738 | 17.0549361309657 | 5.96614105183995 | 4.26843458790854 |
| chr1 | 6223882 | 6223889 | - | 3.01188173062009 | 2.27399148412876 | 1.19322821036799 | 1.06710864697714 |
| chr1 | 6223893 | 6223913 | - | 3.01188173062009 | 1.13699574206438 | 5.96614105183995 | 3.20132594093141 |
| chr1 | 6223940 | 6223980 | - | 18.0712903837206 | 10.2329616785794 | 8.35259747257593 | 10.6710864697714 |
| chr1 | 6224193 | 6224222 | - | 5.01980288436682 | 4.54798296825753 | 7.15936926220794 | 1.06710864697714 |
| chr1 | 6224285 | 6224301 | - | 3.01188173062009 | 3.41098722619315 | 3.57968463110397 | 6.40265188186281 |
| chr1 | 6224324 | 6224327 | - | 9.03564519186028 | 5.68497871032191 | 5.96614105183995 | 2.13421729395427 |
| chr1 | 6224472 | 6224479 | - | 3.01188173062009 | 2.27399148412876 | 3.57968463110397 | 4.26843458790854 |
| chr1 | 6224662 | 6224690 | - | 7.02772403811355 | 6.82197445238629 | 7.15936926220794 | 8.53686917581709 |
| chr1 | 6224723 | 6224771 | - | 11.043566345607 | 18.1919318730301 | 17.8984231555198 | 2.13421729395427 |
| chr1 | 6224791 | 6224805 | - | 6.02376346124019 | 13.6439489047726 | 4.77291284147196 | 2.13421729395427 |
| chr1 | 6224837 | 6224905 | - | 17.0673298068472 | 27.2878978095452 | 20.2848795762558 | 18.1408469986113 |
| chr1 | 6224928 | 6224937 | - | 6.02376346124019 | 6.82197445238629 | 2.38645642073598 | 2.13421729395427 |
| chr1 | 6264309 | 6264387 | - | 111.439624032943 | 50.0278126508328 | 90.6853439879672 | 131.254363578188 |
| chr1 | 6264450 | 6264483 | - | 10.0396057687336 | 5.68497871032191 | 8.35259747257593 | 4.26843458790854 |
| chr1 | 6393530 | 6393561 | + | 1.00396057687336 | 9.09596593651506 | 5.96614105183995 | 3.20132594093141 |
| chr1 | 6460184 | 6460340 | + | 2.00792115374673 | 1.13699574206438 | 2.38645642073598 | 3.20132594093141 |
| chr1 | 6521453 | 6521523 | - | 21.0831721143407 | 6.82197445238629 | 11.9322821036799 | 20.2750642925656 |
| chr1 | 6521579 | 6521653 | - | 55.217831728035 | 56.8497871032191 | 72.7869208324474 | 89.6371263460794 |
| chr1 | 6521670 | 6521675 | - | 1.00396057687336 | 2.27399148412876 | 2.38645642073598 | 2.13421729395427 |
| chr1 | 6521697 | 6521752 | - | 22.087132691214 | 19.3289276150945 | 20.2848795762558 | 30.9461507623369 |
| chr1 | 6521829 | 6521830 | - | 1.00396057687336 | 1.13699574206438 | 1.19322821036799 | 1.06710864697714 |
| chr1 | 6521872 | 6521912 | - | 9.03564519186028 | 2.27399148412876 | 7.15936926220794 | 8.53686917581709 |
| chr1 | 6521932 | 6521949 | - | 4.01584230749346 | 3.41098722619315 | 5.96614105183995 | 5.33554323488568 |
| chr1 | 6522061 | 6522081 | - | 35.1386201905678 | 128.480518853275 | 78.7530618842873 | 57.6238669367653 |
| chr1 | 6522207 | 6522221 | - | 2.00792115374673 | 2.27399148412876 | 9.54582568294392 | 4.26843458790854 |
| chr1 | 6522270 | 6522272 | - | 2.00792115374673 | 4.54798296825753 | 2.38645642073598 | 2.13421729395427 |
| chr1 | 6522607 | 6522617 | - | 6.02376346124019 | 4.54798296825753 | 23.8645642073598 | 20.2750642925656 |
| chr1 | 6522915 | 6522927 | - | 2.00792115374673 | 3.41098722619315 | 13.1255103140479 | 13.8724124107028 |
| chr1 | 6523196 | 6523225 | - | 2.00792115374673 | 2.27399148412876 | 1.19322821036799 | 7.46976052883995 |
| chr1 | 6590827 | 6590842 | - | 196.776273067179 | 94.3706465913437 | 39.3765309421437 | 119.516168461439 |
| chr1 | 6591020 | 6591064 | - | 2.00792115374673 | 1.13699574206438 | 1.19322821036799 | 2.13421729395427 |
| chr1 | 6592277 | 6592286 | - | 4.01584230749346 | 1.13699574206438 | 7.15936926220794 | 3.20132594093141 |
| chr1 | 6592394 | 6592395 | - | 3.01188173062009 | 3.41098722619315 | 2.38645642073598 | 2.13421729395427 |
| chr1 | 6592446 | 6592459 | - | 9.03564519186028 | 1.13699574206438 | 11.9322821036799 | 5.33554323488568 |
| chr1 | 6621913 | 6621922 | + | 1.00396057687336 | 1.13699574206438 | 4.77291284147196 | 1.06710864697714 |
| chr1 | 6622582 | 6622613 | + | 2.00792115374673 | 4.54798296825753 | 5.96614105183995 | 10.6710864697714 |
| chr1 | 6622701 | 6622741 | + | 8.03168461498692 | 4.54798296825753 | 14.3187385244159 | 8.53686917581709 |
| chr1 | 6622778 | 6622942 | + | 73.2891221117556 | 105.740604011988 | 289.954455119421 | 152.59653651773 |
| chr1 | 6623822 | 6623831 | + | 1.00396057687336 | 2.27399148412876 | 3.57968463110397 | 7.46976052883995 |
| chr1 | 6633452 | 6633456 | + | 2.00792115374673 | 1.13699574206438 | 1.19322821036799 | 2.13421729395427 |
| chr1 | 6633504 | 6633539 | + | 1.00396057687336 | 12.5069531627082 | 4.77291284147196 | 7.46976052883995 |
| chr1 | 6634199 | 6634211 | - | 7.02772403811355 | 6.82197445238629 | 13.1255103140479 | 6.40265188186281 |
| chr1 | 6635276 | 6635299 | - | 4.01584230749346 | 5.68497871032191 | 4.77291284147196 | 5.33554323488568 |
| chr1 | 6635552 | 6635568 | - | 6.02376346124019 | 1.13699574206438 | 4.77291284147196 | 1.06710864697714 |
| chr1 | 6635589 | 6635602 | - | 2.00792115374673 | 1.13699574206438 | 7.15936926220794 | 8.53686917581709 |
| chr1 | 6635626 | 6635649 | - | 7.02772403811355 | 1.13699574206438 | 5.96614105183995 | 3.20132594093141 |
| chr1 | 6871757 | 6871773 | + | 5.01980288436682 | 4.54798296825753 | 3.57968463110397 | 11.7381951167485 |
| chr1 | 6871800 | 6871845 | + | 6.02376346124019 | 9.09596593651506 | 11.9322821036799 | 8.53686917581709 |
| chr1 | 6871923 | 6871974 | + | 17.0673298068472 | 14.780944646837 | 9.54582568294392 | 6.40265188186281 |
| chr1 | 7781260 | 7781273 | + | 2.00792115374673 | 5.68497871032191 | 2.38645642073598 | 3.20132594093141 |
| chr1 | 7781311 | 7781336 | + | 1.00396057687336 | 1.13699574206438 | 2.38645642073598 | 5.33554323488568 |
| chr1 | 7781383 | 7781409 | + | 13.0514874993537 | 7.95897019445067 | 8.35259747257593 | 4.26843458790854 |
| chr1 | 7844813 | 7844815 | + | 1.00396057687336 | 1.13699574206438 | 4.77291284147196 | 20.2750642925656 |
| chr1 | 7920331 | 7920333 | - | 6.02376346124019 | 4.54798296825753 | 2.38645642073598 | 4.26843458790854 |
| chr1 | 7985056 | 7985092 | + | 4.01584230749346 | 2.27399148412876 | 5.96614105183995 | 5.33554323488568 |
| chr1 | 7985131 | 7985230 | + | 19.0752509605939 | 10.2329616785794 | 16.7051949451519 | 11.7381951167485 |
| chr1 | 7985251 | 7985268 | + | 2.00792115374673 | 3.41098722619315 | 2.38645642073598 | 2.13421729395427 |
| chr1 | 8011945 | 8012924 | - | 173.685179799092 | 387.715548043954 | 19.0916513658878 | 23.476390233497 |
| chr1 | 8012945 | 8013874 | - | 79.3128855729958 | 185.330305956494 | 9.54582568294392 | 17.0737383516342 |
| chr1 | 8353096 | 8353102 | - | 1.00396057687336 | 1.13699574206438 | 1.19322821036799 | 8.53686917581709 |
| chr1 | 8353388 | 8353481 | - | 3.01188173062009 | 2.27399148412876 | 5.96614105183995 | 6.40265188186281 |
| chr1 | 8354652 | 8354681 | - | 2.00792115374673 | 2.27399148412876 | 1.19322821036799 | 11.7381951167485 |
| chr1 | 8861018 | 8861187 | - | 915.612046108508 | 1136.99574206438 | 643.150005388346 | 548.493844546248 |
| chr1 | 8861212 | 8861396 | - | 61.2415951892752 | 38.657855230189 | 59.6614105183995 | 103.509538756782 |
| chr1 | 8878814 | 8878904 | + | 15.0594086531005 | 34.1098722619315 | 19.0916513658878 | 20.2750642925656 |
| chr1 | 8879691 | 8879694 | + | 2.00792115374673 | 5.68497871032191 | 1.19322821036799 | 2.13421729395427 |
| chr1 | 8879775 | 8879779 | + | 10.0396057687336 | 37.5208594881246 | 21.4781077866238 | 20.2750642925656 |
| chr1 | 8879806 | 8879807 | + | 3.01188173062009 | 1.13699574206438 | 2.38645642073598 | 3.20132594093141 |
| chr1 | 9582632 | 9582667 | + | 2.00792115374673 | 1.13699574206438 | 2.38645642073598 | 4.26843458790854 |
| chr1 | 9582719 | 9582753 | + | 3.01188173062009 | 5.68497871032191 | 1.19322821036799 | 3.20132594093141 |
| chr1 | 9614763 | 9614783 | + | 3.01188173062009 | 1.13699574206438 | 2.38645642073598 | 2.13421729395427 |
| chr1 | 9729045 | 9729166 | - | 27.1069355755808 | 17.0549361309657 | 1.19322821036799 | 3.20132594093141 |
| chr1 | 9729403 | 9729410 | - | 46.1821865361748 | 17.0549361309657 | 11.9322821036799 | 14.9395210576799 |
| chr1 | 9729471 | 9729505 | - | 5.01980288436682 | 9.09596593651506 | 2.38645642073598 | 3.20132594093141 |
| chr1 | 9729774 | 9729831 | - | 19.0752509605939 | 17.0549361309657 | 26.2510206280958 | 5.33554323488568 |
| chr1 | 9730031 | 9730077 | - | 10.0396057687336 | 15.9179403889013 | 3.57968463110397 | 2.13421729395427 |
| chr1 | 9730123 | 9730166 | - | 37.1465413443145 | 25.0139063254164 | 17.8984231555198 | 9.60397782279422 |
| chr1 | 9850141 | 9850245 | - | 18.0712903837206 | 31.8358807778027 | 143.187385244159 | 49.0869977609482 |
| chr1 | 9850276 | 9850407 | - | 18.0712903837206 | 32.9728765198671 | 124.095733878271 | 69.3620620535138 |
| chr1 | 9850486 | 9850568 | - | 27.1069355755808 | 34.1098722619315 | 147.960298085631 | 184.609795927044 |
| chr1 | 9929517 | 9929522 | - | 7.02772403811355 | 5.68497871032191 | 2.38645642073598 | 1.06710864697714 |
| chr1 | 9930281 | 9930316 | - | 2.00792115374673 | 6.82197445238629 | 1.19322821036799 | 5.33554323488568 |
| chr1 | 10180084 | 10180142 | + | 9.03564519186028 | 13.6439489047726 | 23.8645642073598 | 38.4159112911769 |
| chr1 | 10180167 | 10180206 | + | 19.0752509605939 | 29.5618892936739 | 40.5697591525116 | 51.2212150549025 |
| chr1 | 10180228 | 10180292 | + | 148.586165377258 | 108.014595496116 | 221.940447128446 | 353.212962149432 |
| chr1 | 10180317 | 10180377 | + | 35.1386201905678 | 19.3289276150945 | 45.3426719939836 | 22.4092815865198 |
| chr1 | 10180406 | 10180408 | + | 3.01188173062009 | 1.13699574206438 | 1.19322821036799 | 1.06710864697714 |
| chr1 | 10180500 | 10180517 | + | 3.01188173062009 | 2.27399148412876 | 7.15936926220794 | 4.26843458790854 |
| chr1 | 10180553 | 10180587 | + | 6.02376346124019 | 6.82197445238629 | 10.7390538933119 | 4.26843458790854 |
| chr1 | 10180746 | 10180795 | + | 4.01584230749346 | 1.13699574206438 | 2.38645642073598 | 2.13421729395427 |
| chr1 | 10180878 | 10180911 | + | 2.00792115374673 | 3.41098722619315 | 1.19322821036799 | 2.13421729395427 |
| chr1 | 10181074 | 10181129 | + | 1.00396057687336 | 1.13699574206438 | 1.19322821036799 | 1.06710864697714 |
| chr1 | 10306537 | 10306547 | + | 5.01980288436682 | 3.41098722619315 | 2.38645642073598 | 4.26843458790854 |
| chr1 | 10307599 | 10307606 | + | 12.0475269224804 | 6.82197445238629 | 2.38645642073598 | 3.20132594093141 |
| chr1 | 10376666 | 10376693 | + | 5.01980288436682 | 20.4659233571589 | 5.96614105183995 | 1.06710864697714 |
| chr1 | 10376893 | 10376929 | + | 6.02376346124019 | 3.41098722619315 | 1.19322821036799 | 1.06710864697714 |
| chr1 | 10378855 | 10378872 | + | 5.01980288436682 | 2.27399148412876 | 3.57968463110397 | 2.13421729395427 |
| chr1 | 10379946 | 10379967 | + | 4.01584230749346 | 3.41098722619315 | 1.19322821036799 | 3.20132594093141 |
| chr1 | 10381533 | 10381570 | + | 2.00792115374673 | 5.68497871032191 | 2.38645642073598 | 1.06710864697714 |
| chr1 | 10419788 | 10419893 | + | 2.00792115374673 | 1.13699574206438 | 21.4781077866238 | 16.006629704657 |
| chr1 | 10419980 | 10420066 | + | 29.1148567293276 | 21.6029190992233 | 50.1155848354556 | 41.6172372321083 |
| chr1 | 10440382 | 10440414 | + | 10.0396057687336 | 5.68497871032191 | 9.54582568294392 | 3.20132594093141 |
| chr1 | 10442526 | 10442534 | + | 9.03564519186028 | 13.6439489047726 | 1.19322821036799 | 10.6710864697714 |
| chr1 | 10442641 | 10442658 | + | 9.03564519186028 | 3.41098722619315 | 1.19322821036799 | 5.33554323488568 |
| chr1 | 10442695 | 10442729 | + | 22.087132691214 | 12.5069531627082 | 3.57968463110397 | 8.53686917581709 |
| chr1 | 10442762 | 10442788 | + | 11.043566345607 | 20.4659233571589 | 9.54582568294392 | 10.6710864697714 |
| chr1 | 10451570 | 10451571 | + | 1.00396057687336 | 1.13699574206438 | 1.19322821036799 | 2.13421729395427 |
| chr1 | 10456718 | 10456731 | - | 5.01980288436682 | 5.68497871032191 | 2.38645642073598 | 4.26843458790854 |
| chr1 | 10456855 | 10456862 | - | 4.01584230749346 | 3.41098722619315 | 8.35259747257593 | 3.20132594093141 |
| chr1 | 10457533 | 10457539 | - | 3.01188173062009 | 5.68497871032191 | 9.54582568294392 | 9.60397782279422 |
| chr1 | 10458427 | 10458446 | - | 27.1069355755808 | 26.1509020674808 | 14.3187385244159 | 12.8053037637256 |
| chr1 | 10458610 | 10458628 | - | 15.0594086531005 | 11.3699574206438 | 21.4781077866238 | 6.40265188186281 |
| chr1 | 10459753 | 10459775 | - | 24.0950538449607 | 20.4659233571589 | 9.54582568294392 | 35.2145853502455 |
| chr1 | 10459796 | 10459808 | - | 1.00396057687336 | 1.13699574206438 | 2.38645642073598 | 12.8053037637256 |
| chr1 | 10459826 | 10459836 | - | 4.01584230749346 | 1.13699574206438 | 1.19322821036799 | 1.06710864697714 |
| chr1 | 10459980 | 10459985 | - | 15.0594086531005 | 9.09596593651506 | 2.38645642073598 | 6.40265188186281 |
| chr1 | 10461372 | 10461374 | - | 1.00396057687336 | 2.27399148412876 | 1.19322821036799 | 3.20132594093141 |
| chr1 | 10630610 | 10630627 | + | 2.00792115374673 | 2.27399148412876 | 4.77291284147196 | 5.33554323488568 |
| chr1 | 11022662 | 11022677 | + | 92.3643730723495 | 125.069531627082 | 215.974306076606 | 65.0936274656053 |
| chr1 | 11022814 | 11022818 | + | 2.00792115374673 | 3.41098722619315 | 2.38645642073598 | 2.13421729395427 |
| chr1 | 11023070 | 11023087 | + | 1.00396057687336 | 2.27399148412876 | 2.38645642073598 | 2.13421729395427 |
| chr1 | 11023214 | 11023217 | + | 2.00792115374673 | 2.27399148412876 | 3.57968463110397 | 2.13421729395427 |
| chr1 | 11023258 | 11023263 | + | 1.00396057687336 | 2.27399148412876 | 5.96614105183995 | 8.53686917581709 |
| chr1 | 11023325 | 11023371 | + | 101.40001826421 | 73.9047232341848 | 94.2650286190712 | 116.314842520508 |
| chr1 | 11023397 | 11023594 | + | 600.368424970272 | 561.675896579805 | 557.237574241851 | 643.466514127213 |
| chr1 | 11023645 | 11023700 | + | 94.3722942260963 | 78.4527062024424 | 69.2072362013434 | 75.7647139353766 |
| chr1 | 11023852 | 11023870 | + | 4.01584230749346 | 5.68497871032191 | 5.96614105183995 | 6.40265188186281 |
| chr1 | 11025200 | 11025227 | + | 6.02376346124019 | 7.95897019445067 | 2.38645642073598 | 4.26843458790854 |
| chr1 | 11054621 | 11054659 | - | 32.1267384599477 | 18.1919318730301 | 32.2171616799357 | 45.8856718200168 |
| chr1 | 11054680 | 11054887 | - | 181.716864414079 | 79.5897019445067 | 116.936364616063 | 201.683534278679 |
| chr1 | 11066665 | 11066700 | - | 20.0792115374673 | 38.657855230189 | 28.6374770488317 | 30.9461507623369 |
| chr1 | 11106673 | 11106695 | - | 7.02772403811355 | 4.54798296825753 | 15.5119667347839 | 4.26843458790854 |
| chr1 | 11106719 | 11106721 | - | 3.01188173062009 | 2.27399148412876 | 2.38645642073598 | 1.06710864697714 |
| chr1 | 11106743 | 11106771 | - | 74.293082688629 | 43.2058381984465 | 25.0577924177278 | 30.9461507623369 |
| chr1 | 11106794 | 11106817 | - | 76.3010038423757 | 46.6168254246397 | 13.1255103140479 | 62.959410171651 |
| chr1 | 11106837 | 11106865 | - | 3.01188173062009 | 4.54798296825753 | 2.38645642073598 | 8.53686917581709 |
| chr1 | 11107216 | 11107217 | - | 2.00792115374673 | 2.27399148412876 | 2.38645642073598 | 1.06710864697714 |
| chr1 | 11107402 | 11107406 | - | 3.01188173062009 | 2.27399148412876 | 4.77291284147196 | 1.06710864697714 |
| chr1 | 11107448 | 11107502 | - | 21.0831721143407 | 19.3289276150945 | 79.9462900946553 | 42.6843458790854 |
| chr1 | 11286221 | 11286250 | + | 40.1584230749346 | 43.2058381984465 | 41.7629873628796 | 75.7647139353766 |
| chr1 | 11648412 | 11648561 | - | 16.0633692299738 | 6.82197445238629 | 8.35259747257593 | 5.33554323488568 |
| chr1 | 11648643 | 11648772 | - | 5.01980288436682 | 7.95897019445067 | 7.15936926220794 | 1.06710864697714 |
| chr1 | 11662171 | 11662187 | + | 4.01584230749346 | 5.68497871032191 | 1.19322821036799 | 2.13421729395427 |
| chr1 | 11674290 | 11674302 | + | 2.00792115374673 | 3.41098722619315 | 23.8645642073598 | 16.006629704657 |
| chr1 | 11674499 | 11674561 | - | 10.0396057687336 | 12.5069531627082 | 10.7390538933119 | 11.7381951167485 |
| chr1 | 11674583 | 11674619 | - | 1.00396057687336 | 3.41098722619315 | 2.38645642073598 | 2.13421729395427 |
| chr1 | 11748572 | 11748607 | + | 8.03168461498692 | 5.68497871032191 | 19.0916513658878 | 7.46976052883995 |
| chr1 | 11750181 | 11750225 | + | 4.01584230749346 | 6.82197445238629 | 15.5119667347839 | 6.40265188186281 |
| chr1 | 11750244 | 11750254 | + | 1.00396057687336 | 2.27399148412876 | 2.38645642073598 | 3.20132594093141 |
| chr1 | 11750294 | 11750310 | + | 3.01188173062009 | 2.27399148412876 | 2.38645642073598 | 2.13421729395427 |
| chr1 | 11750333 | 11750349 | + | 3.01188173062009 | 2.27399148412876 | 1.19322821036799 | 2.13421729395427 |
| chr1 | 11750384 | 11750436 | + | 17.0673298068472 | 10.2329616785794 | 23.8645642073598 | 17.0737383516342 |
| chr1 | 11750457 | 11750485 | + | 2.00792115374673 | 4.54798296825753 | 2.38645642073598 | 18.1408469986113 |
| chr1 | 11750689 | 11750740 | + | 4.01584230749346 | 5.68497871032191 | 10.7390538933119 | 2.13421729395427 |
| chr1 | 11786304 | 11786352 | - | 2.00792115374673 | 9.09596593651506 | 15.5119667347839 | 56.5567582897882 |
| chr1 | 11786913 | 11787043 | - | 1.00396057687336 | 5.68497871032191 | 103.810854302015 | 73.6304966414224 |
| chr1 | 11787064 | 11787118 | - | 1.00396057687336 | 1.13699574206438 | 7.15936926220794 | 20.2750642925656 |
| chr1 | 11919679 | 11919695 | - | 12.0475269224804 | 14.780944646837 | 21.4781077866238 | 13.8724124107028 |
| chr1 | 11919862 | 11919972 | - | 10.0396057687336 | 4.54798296825753 | 13.1255103140479 | 29.8790421153598 |
| chr1 | 11920180 | 11920216 | - | 2.00792115374673 | 5.68497871032191 | 7.15936926220794 | 3.20132594093141 |
| chr1 | 11975407 | 11975526 | + | 37.1465413443145 | 42.0688424563821 | 53.6952694665595 | 110.979299285622 |
| chr1 | 12011595 | 12011601 | + | 4.01584230749346 | 6.82197445238629 | 8.35259747257593 | 1.06710864697714 |
| chr1 | 12011622 | 12011647 | + | 3.01188173062009 | 2.27399148412876 | 11.9322821036799 | 5.33554323488568 |
| chr1 | 12011789 | 12011832 | + | 10.0396057687336 | 6.82197445238629 | 27.4442488384638 | 25.6106075274513 |
| chr1 | 12011981 | 12012067 | + | 15.0594086531005 | 11.3699574206438 | 35.7968463110397 | 34.1474767032683 |
| chr1 | 12012097 | 12012126 | + | 17.0673298068472 | 10.2329616785794 | 16.7051949451519 | 13.8724124107028 |
| chr1 | 12012176 | 12012230 | + | 18.0712903837206 | 9.09596593651506 | 19.0916513658878 | 16.006629704657 |
| chr1 | 12012257 | 12012284 | + | 9.03564519186028 | 3.41098722619315 | 8.35259747257593 | 13.8724124107028 |
| chr1 | 12012308 | 12012324 | + | 6.02376346124019 | 2.27399148412876 | 3.57968463110397 | 1.06710864697714 |
| chr1 | 12012370 | 12012487 | + | 12.0475269224804 | 9.09596593651506 | 39.3765309421437 | 25.6106075274513 |
| chr1 | 12012511 | 12012578 | + | 11.043566345607 | 5.68497871032191 | 21.4781077866238 | 12.8053037637256 |
| chr1 | 12012640 | 12012671 | + | 9.03564519186028 | 4.54798296825753 | 29.8307052591997 | 14.9395210576799 |
| chr1 | 12012709 | 12012855 | + | 41.1623836518079 | 40.9318467143178 | 116.936364616063 | 93.9055609339879 |
| chr1 | 12012905 | 12012912 | + | 4.01584230749346 | 2.27399148412876 | 8.35259747257593 | 3.20132594093141 |
| chr1 | 12012939 | 12013150 | + | 94.3722942260963 | 90.9596593651506 | 118.129592826431 | 166.468948928433 |
| chr1 | 12013171 | 12013183 | + | 47.1861471130481 | 127.343523111211 | 26.2510206280958 | 68.2949534065367 |
| chr1 | 12013211 | 12013221 | + | 2.00792115374673 | 2.27399148412876 | 8.35259747257593 | 4.26843458790854 |
| chr1 | 12013243 | 12013266 | + | 60.2376346124019 | 60.2607743294122 | 76.3666054635513 | 86.435800405148 |
| chr1 | 12013302 | 12013333 | + | 222.879248065887 | 208.070220797782 | 286.374770488317 | 490.869977609482 |
| chr1 | 12013386 | 12013406 | + | 11.043566345607 | 11.3699574206438 | 23.8645642073598 | 32.0132594093141 |
| chr1 | 12013431 | 12013483 | + | 8.03168461498692 | 5.68497871032191 | 9.54582568294392 | 6.40265188186281 |
| chr1 | 12031808 | 12031840 | + | 7.02772403811355 | 9.09596593651506 | 22.6713359969918 | 20.2750642925656 |
| chr1 | 12031869 | 12031897 | + | 9.03564519186028 | 1.13699574206438 | 13.1255103140479 | 13.8724124107028 |
| chr1 | 12031913 | 12031933 | + | 3.01188173062009 | 2.27399148412876 | 4.77291284147196 | 7.46976052883995 |
| chr1 | 12031974 | 12032025 | + | 26.1029749987075 | 34.1098722619315 | 77.5598336739193 | 86.435800405148 |
| chr1 | 12509130 | 12509169 | + | 3.01188173062009 | 1.13699574206438 | 10.7390538933119 | 8.53686917581709 |
| chr1 | 12509205 | 12509227 | + | 3.01188173062009 | 2.27399148412876 | 8.35259747257593 | 3.20132594093141 |
| chr1 | 12509647 | 12509666 | + | 2.00792115374673 | 2.27399148412876 | 5.96614105183995 | 7.46976052883995 |
| chr1 | 12509784 | 12509786 | + | 1.00396057687336 | 4.54798296825753 | 3.57968463110397 | 6.40265188186281 |
| chr1 | 12510044 | 12510147 | + | 28.1108961524542 | 34.1098722619315 | 82.3327465153913 | 54.4225409958339 |
| chr1 | 12567937 | 12568022 | - | 3.01188173062009 | 2.27399148412876 | 3.57968463110397 | 10.6710864697714 |
| chr1 | 13787446 | 13787448 | + | 15.0594086531005 | 9.09596593651506 | 5.96614105183995 | 21.3421729395427 |
| chr1 | 15117202 | 15117203 | + | 2.00792115374673 | 2.27399148412876 | 1.19322821036799 | 1.06710864697714 |
| chr1 | 15428770 | 15428824 | + | 27.1069355755808 | 44.3428339405109 | 8.35259747257593 | 17.0737383516342 |
| chr1 | 15429818 | 15429843 | + | 7.02772403811355 | 7.95897019445067 | 2.38645642073598 | 4.26843458790854 |
| chr1 | 15429996 | 15430018 | + | 2.00792115374673 | 1.13699574206438 | 3.57968463110397 | 1.06710864697714 |
| chr1 | 15430128 | 15430183 | + | 30.1188173062009 | 47.753821166704 | 10.7390538933119 | 23.476390233497 |
| chr1 | 15492367 | 15492389 | - | 15.0594086531005 | 39.7948509722534 | 4.77291284147196 | 11.7381951167485 |
| chr1 | 15568229 | 15568269 | + | 1.00396057687336 | 2.27399148412876 | 7.15936926220794 | 3.20132594093141 |
| chr1 | 15568920 | 15568926 | + | 4.01584230749346 | 1.13699574206438 | 5.96614105183995 | 3.20132594093141 |
| chr1 | 15569495 | 15569512 | + | 6.02376346124019 | 1.13699574206438 | 5.96614105183995 | 20.2750642925656 |
| chr1 | 15569592 | 15569612 | + | 2.00792115374673 | 2.27399148412876 | 7.15936926220794 | 12.8053037637256 |
| chr1 | 15569929 | 15569946 | + | 1.00396057687336 | 10.2329616785794 | 3.57968463110397 | 1.06710864697714 |
| chr1 | 15570473 | 15570493 | + | 8.03168461498692 | 7.95897019445067 | 2.38645642073598 | 2.13421729395427 |
| chr1 | 15570608 | 15570619 | + | 3.01188173062009 | 2.27399148412876 | 5.96614105183995 | 1.06710864697714 |
| chr1 | 15570882 | 15570886 | + | 2.00792115374673 | 2.27399148412876 | 1.19322821036799 | 1.06710864697714 |
| chr1 | 15570960 | 15570971 | + | 8.03168461498692 | 12.5069531627082 | 15.5119667347839 | 4.26843458790854 |
| chr1 | 15571097 | 15571135 | + | 6.02376346124019 | 10.2329616785794 | 9.54582568294392 | 11.7381951167485 |
| chr1 | 15571165 | 15571214 | + | 8.03168461498692 | 9.09596593651506 | 17.8984231555198 | 14.9395210576799 |
| chr1 | 15571299 | 15571365 | + | 5.01980288436682 | 23.876910583352 | 16.7051949451519 | 16.006629704657 |
| chr1 | 15571417 | 15571443 | + | 9.03564519186028 | 10.2329616785794 | 8.35259747257593 | 6.40265188186281 |
| chr1 | 15571509 | 15571515 | + | 4.01584230749346 | 3.41098722619315 | 1.19322821036799 | 6.40265188186281 |
| chr1 | 15571544 | 15571550 | + | 4.01584230749346 | 1.13699574206438 | 2.38645642073598 | 3.20132594093141 |
| chr1 | 15660021 | 15660031 | + | 2.00792115374673 | 2.27399148412876 | 1.19322821036799 | 3.20132594093141 |
| chr1 | 15661726 | 15661738 | + | 2.00792115374673 | 4.54798296825753 | 4.77291284147196 | 3.20132594093141 |
| chr1 | 15661835 | 15661851 | + | 1.00396057687336 | 2.27399148412876 | 3.57968463110397 | 12.8053037637256 |
| chr1 | 15663804 | 15663806 | + | 13.0514874993537 | 7.95897019445067 | 5.96614105183995 | 1.06710864697714 |
| chr1 | 15664729 | 15664734 | + | 1.00396057687336 | 4.54798296825753 | 2.38645642073598 | 2.13421729395427 |
| chr1 | 15667936 | 15667951 | + | 2.00792115374673 | 1.13699574206438 | 2.38645642073598 | 5.33554323488568 |
| chr1 | 15668845 | 15668880 | + | 6.02376346124019 | 2.27399148412876 | 8.35259747257593 | 19.2079556455884 |
| chr1 | 15734707 | 15734723 | + | 57.2257528817818 | 70.4937360079917 | 35.7968463110397 | 48.0198891139711 |
| chr1 | 15939570 | 15939597 | + | 5.01980288436682 | 2.27399148412876 | 7.15936926220794 | 6.40265188186281 |
| chr1 | 15939658 | 15939682 | + | 23.0910932680874 | 17.0549361309657 | 29.8307052591997 | 49.0869977609482 |
| chr1 | 15939711 | 15939727 | + | 7.02772403811355 | 13.6439489047726 | 26.2510206280958 | 42.6843458790854 |
| chr1 | 15939910 | 15939919 | + | 220.87132691214 | 216.029190992233 | 226.713359969918 | 223.025707218221 |
| chr1 | 15940249 | 15940267 | + | 2.00792115374673 | 1.13699574206438 | 2.38645642073598 | 5.33554323488568 |
| chr1 | 15940296 | 15940320 | + | 6.02376346124019 | 7.95897019445067 | 8.35259747257593 | 3.20132594093141 |
| chr1 | 15941906 | 15941918 | - | 5.01980288436682 | 13.6439489047726 | 11.9322821036799 | 20.2750642925656 |
| chr1 | 16198146 | 16198153 | - | 6.02376346124019 | 9.09596593651506 | 11.9322821036799 | 18.1408469986113 |
| chr1 | 16198188 | 16198191 | - | 4.01584230749346 | 5.68497871032191 | 3.57968463110397 | 1.06710864697714 |
| chr1 | 16250442 | 16250444 | - | 2.00792115374673 | 1.13699574206438 | 1.19322821036799 | 3.20132594093141 |
| chr1 | 16395463 | 16395466 | + | 11.043566345607 | 5.68497871032191 | 3.57968463110397 | 2.13421729395427 |
| chr1 | 16395542 | 16395592 | + | 20.0792115374673 | 11.3699574206438 | 21.4781077866238 | 28.8119334683827 |
| chr1 | 16395947 | 16395952 | + | 5.01980288436682 | 4.54798296825753 | 1.19322821036799 | 2.13421729395427 |
| chr1 | 16396204 | 16396213 | + | 4.01584230749346 | 5.68497871032191 | 7.15936926220794 | 2.13421729395427 |
| chr1 | 16396251 | 16396254 | + | 7.02772403811355 | 7.95897019445067 | 2.38645642073598 | 4.26843458790854 |
| chr1 | 16396316 | 16396366 | + | 7.02772403811355 | 9.09596593651506 | 8.35259747257593 | 11.7381951167485 |
| chr1 | 16396915 | 16396967 | + | 17.0673298068472 | 22.7399148412876 | 9.54582568294392 | 9.60397782279422 |
| chr1 | 16396987 | 16397027 | + | 2.00792115374673 | 2.27399148412876 | 7.15936926220794 | 4.26843458790854 |
| chr1 | 16397032 | 16397140 | + | 22.087132691214 | 19.3289276150945 | 21.4781077866238 | 25.6106075274513 |
| chr1 | 16397161 | 16397223 | + | 42.1663442286813 | 28.4248935516096 | 40.5697591525116 | 23.476390233497 |
| chr1 | 16397244 | 16397345 | + | 101.40001826421 | 64.8087572976698 | 82.3327465153913 | 66.1607361125824 |
| chr1 | 16397354 | 16397461 | + | 1259.97052397607 | 579.867828452835 | 937.87737334924 | 923.048979635222 |
| chr1 | 16397503 | 16397524 | + | 25.0990144218341 | 6.82197445238629 | 5.96614105183995 | 18.1408469986113 |
| chr1 | 16397676 | 16397704 | + | 22.087132691214 | 9.09596593651506 | 32.2171616799357 | 12.8053037637256 |
| chr1 | 16397754 | 16397790 | + | 11.043566345607 | 1.13699574206438 | 13.1255103140479 | 12.8053037637256 |
| chr1 | 16397865 | 16397883 | + | 10.0396057687336 | 1.13699574206438 | 13.1255103140479 | 9.60397782279422 |
| chr1 | 16397945 | 16398009 | + | 24.0950538449607 | 12.5069531627082 | 19.0916513658878 | 48.0198891139711 |
| chr1 | 16459350 | 16459354 | + | 1.00396057687336 | 2.27399148412876 | 9.54582568294392 | 6.40265188186281 |
| chr1 | 16459386 | 16459435 | + | 3.01188173062009 | 5.68497871032191 | 21.4781077866238 | 9.60397782279422 |
| chr1 | 16459534 | 16459588 | + | 4.01584230749346 | 3.41098722619315 | 8.35259747257593 | 14.9395210576799 |
| chr1 | 16459655 | 16459716 | + | 31.1227778830743 | 31.8358807778027 | 38.1833027317757 | 67.2278447595595 |
| chr1 | 16459871 | 16459904 | + | 5.01980288436682 | 3.41098722619315 | 14.3187385244159 | 9.60397782279422 |
| chr1 | 16460026 | 16460037 | + | 2.00792115374673 | 6.82197445238629 | 7.15936926220794 | 6.40265188186281 |
| chr1 | 16986023 | 16986034 | - | 10.0396057687336 | 1.13699574206438 | 7.15936926220794 | 13.8724124107028 |
| chr1 | 17018745 | 17018855 | - | 16.0633692299738 | 32.9728765198671 | 50.1155848354556 | 35.2145853502455 |
| chr1 | 18871448 | 18871484 | - | 4.01584230749346 | 2.27399148412876 | 1.19322821036799 | 2.13421729395427 |
| chr1 | 18877513 | 18877564 | - | 1.00396057687336 | 1.13699574206438 | 1.19322821036799 | 1.06710864697714 |
| chr1 | 19074528 | 19074609 | - | 16.0633692299738 | 17.0549361309657 | 17.8984231555198 | 20.2750642925656 |
| chr1 | 19074634 | 19074694 | - | 3.01188173062009 | 5.68497871032191 | 2.38645642073598 | 12.8053037637256 |
| chr1 | 19112553 | 19112579 | - | 4.01584230749346 | 1.13699574206438 | 1.19322821036799 | 2.13421729395427 |
| chr1 | 19210341 | 19210554 | + | 112.443584609817 | 150.083437952498 | 223.133675338814 | 148.328101929822 |
| chr1 | 19218003 | 19218009 | - | 1.00396057687336 | 2.27399148412876 | 3.57968463110397 | 8.53686917581709 |
| chr1 | 19218121 | 19218175 | - | 8.03168461498692 | 7.95897019445067 | 4.77291284147196 | 1.06710864697714 |
| chr1 | 19218199 | 19218212 | - | 5.01980288436682 | 1.13699574206438 | 1.19322821036799 | 4.26843458790854 |
| chr1 | 19218265 | 19218327 | - | 89.3524913417294 | 69.3567402659273 | 54.8884976769275 | 76.8318225823538 |
| chr1 | 19218392 | 19218399 | - | 11.043566345607 | 14.780944646837 | 2.38645642073598 | 3.20132594093141 |
| chr1 | 19218475 | 19218482 | - | 5.01980288436682 | 6.82197445238629 | 3.57968463110397 | 4.26843458790854 |
| chr1 | 19218509 | 19218534 | - | 8.03168461498692 | 22.7399148412876 | 8.35259747257593 | 17.0737383516342 |
| chr1 | 19218559 | 19218577 | - | 2.00792115374673 | 4.54798296825753 | 1.19322821036799 | 4.26843458790854 |
| chr1 | 19218710 | 19218733 | - | 1.00396057687336 | 4.54798296825753 | 1.19322821036799 | 1.06710864697714 |
| chr1 | 19218955 | 19218975 | - | 6.02376346124019 | 4.54798296825753 | 3.57968463110397 | 9.60397782279422 |
| chr1 | 19219015 | 19219067 | - | 6.02376346124019 | 10.2329616785794 | 9.54582568294392 | 6.40265188186281 |
| chr1 | 19258728 | 19258850 | + | 43.1703048055547 | 22.7399148412876 | 68.0140079909754 | 105.643756050736 |
| chr1 | 19258909 | 19258910 | + | 1.00396057687336 | 1.13699574206438 | 3.57968463110397 | 2.13421729395427 |
| chr1 | 19259388 | 19259391 | + | 2.00792115374673 | 4.54798296825753 | 10.7390538933119 | 9.60397782279422 |
| chr1 | 19260045 | 19260055 | + | 2.00792115374673 | 1.13699574206438 | 2.38645642073598 | 8.53686917581709 |
| chr1 | 19260102 | 19260106 | + | 6.02376346124019 | 4.54798296825753 | 2.38645642073598 | 2.13421729395427 |
| chr1 | 19303991 | 19304055 | - | 22.087132691214 | 13.6439489047726 | 19.0916513658878 | 11.7381951167485 |
| chr1 | 19338885 | 19338975 | - | 53.2099105742883 | 44.3428339405109 | 76.3666054635513 | 84.3015831111937 |
| chr1 | 19339123 | 19339152 | - | 3.01188173062009 | 4.54798296825753 | 8.35259747257593 | 8.53686917581709 |
| chr1 | 19339381 | 19339448 | - | 3.01188173062009 | 1.13699574206438 | 4.77291284147196 | 4.26843458790854 |
| chr1 | 19626466 | 19626475 | + | 11.043566345607 | 17.0549361309657 | 8.35259747257593 | 20.2750642925656 |
| chr1 | 19626532 | 19626538 | + | 22.087132691214 | 10.2329616785794 | 2.38645642073598 | 7.46976052883995 |
| chr1 | 19626582 | 19626592 | + | 4.01584230749346 | 4.54798296825753 | 8.35259747257593 | 11.7381951167485 |
| chr1 | 19657271 | 19657381 | + | 349.378280751931 | 211.481208023975 | 11.9322821036799 | 13.8724124107028 |
| chr1 | 19657402 | 19657476 | + | 122.48319037855 | 106.877599754052 | 1.19322821036799 | 1.06710864697714 |
| chr1 | 19657678 | 19657700 | + | 16.0633692299738 | 14.780944646837 | 1.19322821036799 | 1.06710864697714 |
| chr1 | 19657809 | 19657819 | + | 4.01584230749346 | 1.13699574206438 | 1.19322821036799 | 1.06710864697714 |
| chr1 | 19911370 | 19911376 | + | 1.00396057687336 | 1.13699574206438 | 4.77291284147196 | 4.26843458790854 |
| chr1 | 19911665 | 19911670 | + | 3.01188173062009 | 2.27399148412876 | 10.7390538933119 | 1.06710864697714 |
| chr1 | 20499469 | 20499488 | - | 1.00396057687336 | 6.82197445238629 | 1.19322821036799 | 4.26843458790854 |
| chr1 | 20499508 | 20499531 | - | 12.0475269224804 | 19.3289276150945 | 21.4781077866238 | 22.4092815865198 |
| chr1 | 20499605 | 20499607 | - | 3.01188173062009 | 4.54798296825753 | 1.19322821036799 | 2.13421729395427 |
| chr1 | 20651790 | 20651795 | - | 2.00792115374673 | 4.54798296825753 | 3.57968463110397 | 4.26843458790854 |
| chr1 | 20651861 | 20651883 | - | 13.0514874993537 | 20.4659233571589 | 38.1833027317757 | 38.4159112911769 |
| chr1 | 20652190 | 20652412 | - | 112.443584609817 | 95.5076423334081 | 187.336829027774 | 212.35462074845 |
| chr1 | 20742745 | 20742881 | - | 85.336649034236 | 76.1787147183136 | 138.414472402687 | 54.4225409958339 |
| chr1 | 20742911 | 20742959 | - | 30.1188173062009 | 26.1509020674808 | 56.0817258872955 | 35.2145853502455 |
| chr1 | 20742990 | 20743014 | - | 14.0554480762271 | 15.9179403889013 | 8.35259747257593 | 9.60397782279422 |
| chr1 | 20743089 | 20743117 | - | 2.00792115374673 | 6.82197445238629 | 8.35259747257593 | 2.13421729395427 |
| chr1 | 20743153 | 20743176 | - | 2.00792115374673 | 4.54798296825753 | 4.77291284147196 | 4.26843458790854 |
| chr1 | 20743236 | 20743256 | - | 4.01584230749346 | 11.3699574206438 | 7.15936926220794 | 6.40265188186281 |
| chr1 | 20743284 | 20743296 | - | 2.00792115374673 | 4.54798296825753 | 2.38645642073598 | 5.33554323488568 |
| chr1 | 20743326 | 20743336 | - | 2.00792115374673 | 1.13699574206438 | 3.57968463110397 | 2.13421729395427 |
| chr1 | 20743381 | 20743413 | - | 9.03564519186028 | 1.13699574206438 | 4.77291284147196 | 7.46976052883995 |
| chr1 | 20743753 | 20743764 | - | 8.03168461498692 | 4.54798296825753 | 11.9322821036799 | 4.26843458790854 |
| chr1 | 20743788 | 20743804 | - | 22.087132691214 | 9.09596593651506 | 20.2848795762558 | 19.2079556455884 |
| chr1 | 20744496 | 20744521 | - | 12.0475269224804 | 15.9179403889013 | 8.35259747257593 | 44.8185631730397 |
| chr1 | 20744595 | 20744628 | - | 3.01188173062009 | 7.95897019445067 | 8.35259747257593 | 5.33554323488568 |
| chr1 | 20744651 | 20744675 | - | 44.174265382428 | 60.2607743294122 | 52.5020412561915 | 37.3488026441998 |
| chr1 | 20744735 | 20744781 | - | 4.01584230749346 | 10.2329616785794 | 9.54582568294392 | 9.60397782279422 |
| chr1 | 20775727 | 20775742 | - | 1.00396057687336 | 1.13699574206438 | 3.57968463110397 | 3.20132594093141 |
| chr1 | 20806816 | 20806833 | - | 55.217831728035 | 48.8908169087684 | 23.8645642073598 | 54.4225409958339 |
| chr1 | 20807113 | 20807120 | - | 2.00792115374673 | 1.13699574206438 | 2.38645642073598 | 2.13421729395427 |
| chr1 | 20807158 | 20807189 | - | 5.01980288436682 | 6.82197445238629 | 17.8984231555198 | 6.40265188186281 |
| chr1 | 21217271 | 21217289 | - | 3.01188173062009 | 7.95897019445067 | 1.19322821036799 | 1.06710864697714 |
| chr1 | 21217314 | 21217395 | - | 25.0990144218341 | 52.3018041349616 | 1.19322821036799 | 10.6710864697714 |
| chr1 | 21219568 | 21219571 | - | 20.0792115374673 | 19.3289276150945 | 7.15936926220794 | 10.6710864697714 |
| chr1 | 21219613 | 21219615 | - | 4.01584230749346 | 14.780944646837 | 2.38645642073598 | 2.13421729395427 |
| chr1 | 21596254 | 21596289 | - | 2.00792115374673 | 2.27399148412876 | 3.57968463110397 | 7.46976052883995 |
| chr1 | 21596511 | 21596534 | - | 5.01980288436682 | 2.27399148412876 | 52.5020412561915 | 7.46976052883995 |
| chr1 | 21597012 | 21597058 | - | 6.02376346124019 | 3.41098722619315 | 48.9223566250876 | 10.6710864697714 |
| chr1 | 21598008 | 21598036 | - | 1.00396057687336 | 2.27399148412876 | 5.96614105183995 | 4.26843458790854 |
| chr1 | 21678951 | 21679051 | - | 15.0594086531005 | 13.6439489047726 | 32.2171616799357 | 29.8790421153598 |
| chr1 | 21679083 | 21679102 | - | 52.205949997415 | 27.2878978095452 | 31.0239334695677 | 54.4225409958339 |
| chr1 | 21679142 | 21679176 | - | 218.863405758393 | 136.439489047726 | 140.800928823423 | 328.669463268958 |
| chr1 | 21679246 | 21679299 | - | 5.01980288436682 | 7.95897019445067 | 8.35259747257593 | 11.7381951167485 |
| chr1 | 21728227 | 21728246 | - | 3.01188173062009 | 7.95897019445067 | 2.38645642073598 | 7.46976052883995 |
| chr1 | 21728356 | 21728380 | - | 6.02376346124019 | 1.13699574206438 | 7.15936926220794 | 9.60397782279422 |
| chr1 | 21728542 | 21728711 | - | 74.293082688629 | 94.3706465913437 | 161.085808399679 | 217.690163983336 |
| chr1 | 21822285 | 21822334 | - | 11.043566345607 | 10.2329616785794 | 9.54582568294392 | 30.9461507623369 |
| chr1 | 21822412 | 21822415 | - | 2.00792115374673 | 1.13699574206438 | 1.19322821036799 | 1.06710864697714 |
| chr1 | 21822790 | 21822810 | - | 9.03564519186028 | 2.27399148412876 | 5.96614105183995 | 7.46976052883995 |
| chr1 | 21822844 | 21822898 | - | 3.01188173062009 | 1.13699574206438 | 3.57968463110397 | 3.20132594093141 |
| chr1 | 21822949 | 21822988 | - | 5.01980288436682 | 3.41098722619315 | 2.38645642073598 | 10.6710864697714 |
| chr1 | 21823236 | 21823287 | - | 3.01188173062009 | 2.27399148412876 | 5.96614105183995 | 3.20132594093141 |
| chr1 | 22090461 | 22090463 | + | 2.00792115374673 | 2.27399148412876 | 1.19322821036799 | 1.06710864697714 |
| chr1 | 22090475 | 22090492 | + | 21.0831721143407 | 14.780944646837 | 7.15936926220794 | 17.0737383516342 |
| chr1 | 22090519 | 22090521 | + | 16.0633692299738 | 6.82197445238629 | 1.19322821036799 | 6.40265188186281 |
| chr1 | 22090583 | 22090584 | + | 1.00396057687336 | 1.13699574206438 | 1.19322821036799 | 2.13421729395427 |
| chr1 | 22091725 | 22091760 | + | 13.0514874993537 | 10.2329616785794 | 17.8984231555198 | 85.3686917581709 |
| chr1 | 22092203 | 22092223 | + | 151.598047107878 | 104.603608269923 | 63.2410951495034 | 113.113516579576 |
| chr1 | 22092478 | 22092495 | + | 1.00396057687336 | 2.27399148412876 | 1.19322821036799 | 1.06710864697714 |
| chr1 | 22092597 | 22092649 | + | 228.903011527127 | 227.399148412876 | 108.583767143487 | 265.710053097307 |
| chr1 | 22092674 | 22092686 | + | 5.01980288436682 | 7.95897019445067 | 2.38645642073598 | 13.8724124107028 |
| chr1 | 22092721 | 22092762 | + | 4.01584230749346 | 3.41098722619315 | 10.7390538933119 | 12.8053037637256 |
| chr1 | 22092810 | 22092818 | + | 22.087132691214 | 20.4659233571589 | 26.2510206280958 | 7.46976052883995 |
| chr1 | 22092852 | 22092888 | + | 2.00792115374673 | 1.13699574206438 | 2.38645642073598 | 2.13421729395427 |
| chr1 | 22528084 | 22528085 | + | 1.00396057687336 | 2.27399148412876 | 1.19322821036799 | 1.06710864697714 |
| chr1 | 22528684 | 22528688 | + | 2.00792115374673 | 2.27399148412876 | 1.19322821036799 | 3.20132594093141 |
| chr1 | 22529189 | 22529195 | + | 7.02772403811355 | 2.27399148412876 | 8.35259747257593 | 1.06710864697714 |
| chr1 | 22530606 | 22530607 | + | 1.00396057687336 | 1.13699574206438 | 1.19322821036799 | 1.06710864697714 |
| chr1 | 23083362 | 23083417 | + | 2.00792115374673 | 1.13699574206438 | 1.19322821036799 | 3.20132594093141 |
| chr1 | 23085432 | 23085454 | - | 4.01584230749346 | 5.68497871032191 | 4.77291284147196 | 5.33554323488568 |
| chr1 | 23085652 | 23085675 | - | 5.01980288436682 | 2.27399148412876 | 1.19322821036799 | 9.60397782279422 |
| chr1 | 23085732 | 23085740 | - | 4.01584230749346 | 1.13699574206438 | 8.35259747257593 | 3.20132594093141 |
| chr1 | 23696351 | 23696400 | + | 227.899050950254 | 229.673139897005 | 421.2095582599 | 388.427547499677 |
| chr1 | 23759605 | 23759638 | + | 12.0475269224804 | 7.95897019445067 | 1.19322821036799 | 8.53686917581709 |
| chr1 | 23759803 | 23759862 | + | 14.0554480762271 | 7.95897019445067 | 8.35259747257593 | 5.33554323488568 |
| chr1 | 23759885 | 23759905 | + | 1.00396057687336 | 1.13699574206438 | 2.38645642073598 | 1.06710864697714 |
| chr1 | 23760143 | 23760161 | + | 2.00792115374673 | 2.27399148412876 | 4.77291284147196 | 4.26843458790854 |
| chr1 | 23760192 | 23760209 | + | 1.00396057687336 | 1.13699574206438 | 4.77291284147196 | 3.20132594093141 |
| chr1 | 23760244 | 23760293 | + | 28.1108961524542 | 12.5069531627082 | 9.54582568294392 | 17.0737383516342 |
| chr1 | 23787381 | 23787463 | + | 14.0554480762271 | 15.9179403889013 | 28.6374770488317 | 34.1474767032683 |
| chr1 | 23787502 | 23787635 | + | 26.1029749987075 | 30.6988850357383 | 88.2988875672312 | 88.5700176991023 |
| chr1 | 23787657 | 23787712 | + | 32.1267384599477 | 21.6029190992233 | 75.1733772531833 | 67.2278447595595 |
| chr1 | 23787749 | 23787791 | + | 6.02376346124019 | 4.54798296825753 | 21.4781077866238 | 10.6710864697714 |
| chr1 | 23787804 | 23787820 | + | 13.0514874993537 | 10.2329616785794 | 27.4442488384638 | 26.6777161744284 |
| chr1 | 23787870 | 23787919 | + | 7.02772403811355 | 3.41098722619315 | 20.2848795762558 | 21.3421729395427 |
| chr1 | 23788020 | 23788033 | + | 1.00396057687336 | 5.68497871032191 | 9.54582568294392 | 12.8053037637256 |
| chr1 | 23788189 | 23788204 | + | 1.00396057687336 | 3.41098722619315 | 2.38645642073598 | 3.20132594093141 |
| chr1 | 23794874 | 23794890 | + | 2.00792115374673 | 1.13699574206438 | 7.15936926220794 | 4.26843458790854 |
| chr1 | 23794929 | 23794941 | + | 8.03168461498692 | 3.41098722619315 | 9.54582568294392 | 17.0737383516342 |
| chr1 | 23795430 | 23795511 | + | 121.479229801677 | 103.466612527859 | 76.3666054635513 | 173.938709457273 |
| chr1 | 23795643 | 23795706 | - | 9.03564519186028 | 11.3699574206438 | 10.7390538933119 | 23.476390233497 |
| chr1 | 23801895 | 23802014 | - | 1.00396057687336 | 11.3699574206438 | 2.38645642073598 | 4.26843458790854 |
| chr1 | 23845231 | 23845278 | - | 6.02376346124019 | 10.2329616785794 | 8.35259747257593 | 18.1408469986113 |
| chr1 | 23961869 | 23961928 | + | 9.03564519186028 | 7.95897019445067 | 26.2510206280958 | 22.4092815865198 |
| chr1 | 23962130 | 23962154 | + | 20.0792115374673 | 40.9318467143178 | 59.6614105183995 | 49.0869977609482 |
| chr1 | 23962196 | 23962215 | + | 7.02772403811355 | 13.6439489047726 | 8.35259747257593 | 8.53686917581709 |
| chr1 | 23963208 | 23963224 | + | 5.01980288436682 | 6.82197445238629 | 9.54582568294392 | 11.7381951167485 |
| chr1 | 23963330 | 23963331 | + | 1.00396057687336 | 1.13699574206438 | 2.38645642073598 | 2.13421729395427 |
| chr1 | 23966473 | 23966494 | - | 31.1227778830743 | 52.3018041349616 | 20.2848795762558 | 19.2079556455884 |
| chr1 | 23966583 | 23966596 | - | 6.02376346124019 | 6.82197445238629 | 9.54582568294392 | 4.26843458790854 |
| chr1 | 23967296 | 23967305 | - | 1.00396057687336 | 4.54798296825753 | 2.38645642073598 | 3.20132594093141 |
| chr1 | 23967853 | 23967867 | - | 4.01584230749346 | 4.54798296825753 | 1.19322821036799 | 3.20132594093141 |
| chr1 | 23969386 | 23969399 | - | 2.00792115374673 | 1.13699574206438 | 4.77291284147196 | 2.13421729395427 |
| chr1 | 23970091 | 23970133 | - | 10.0396057687336 | 5.68497871032191 | 16.7051949451519 | 5.33554323488568 |
| chr1 | 23970149 | 23970172 | - | 10.0396057687336 | 19.3289276150945 | 7.15936926220794 | 8.53686917581709 |
| chr1 | 23970244 | 23970297 | - | 18.0712903837206 | 10.2329616785794 | 42.9562155732476 | 30.9461507623369 |
| chr1 | 23970326 | 23970364 | - | 10.0396057687336 | 3.41098722619315 | 20.2848795762558 | 13.8724124107028 |
| chr1 | 23970448 | 23970451 | - | 5.01980288436682 | 6.82197445238629 | 15.5119667347839 | 2.13421729395427 |
| chr1 | 23970858 | 23970874 | - | 5.01980288436682 | 4.54798296825753 | 2.38645642073598 | 3.20132594093141 |
| chr1 | 23971009 | 23971015 | - | 8.03168461498692 | 9.09596593651506 | 2.38645642073598 | 11.7381951167485 |
| chr1 | 23971114 | 23971171 | - | 2.00792115374673 | 2.27399148412876 | 3.57968463110397 | 2.13421729395427 |
| chr1 | 24471251 | 24471269 | + | 6.02376346124019 | 4.54798296825753 | 1.19322821036799 | 2.13421729395427 |
| chr1 | 24471484 | 24471491 | + | 5.01980288436682 | 4.54798296825753 | 1.19322821036799 | 5.33554323488568 |
| chr1 | 24471605 | 24471627 | + | 5.01980288436682 | 2.27399148412876 | 2.38645642073598 | 5.33554323488568 |
| chr1 | 24471765 | 24471769 | + | 16.0633692299738 | 9.09596593651506 | 7.15936926220794 | 2.13421729395427 |
| chr1 | 24672219 | 24672241 | + | 14.0554480762271 | 6.82197445238629 | 15.5119667347839 | 10.6710864697714 |
| chr1 | 24672504 | 24672518 | + | 2.00792115374673 | 5.68497871032191 | 9.54582568294392 | 7.46976052883995 |
| chr1 | 24672543 | 24672594 | + | 64.2534769198953 | 70.4937360079917 | 103.810854302015 | 70.429170700491 |
| chr1 | 24672932 | 24672952 | + | 4.01584230749346 | 6.82197445238629 | 3.57968463110397 | 6.40265188186281 |
| chr1 | 24673137 | 24673140 | + | 1.00396057687336 | 1.13699574206438 | 4.77291284147196 | 9.60397782279422 |
| chr1 | 24841338 | 24841354 | + | 9.03564519186028 | 3.41098722619315 | 1.19322821036799 | 6.40265188186281 |
| chr1 | 24841620 | 24841634 | + | 11.043566345607 | 3.41098722619315 | 3.57968463110397 | 6.40265188186281 |
| chr1 | 24841672 | 24841692 | + | 7.02772403811355 | 7.95897019445067 | 2.38645642073598 | 6.40265188186281 |
| chr1 | 24842054 | 24842193 | + | 42.1663442286813 | 47.753821166704 | 13.1255103140479 | 10.6710864697714 |
| chr1 | 24842243 | 24842283 | + | 5.01980288436682 | 7.95897019445067 | 3.57968463110397 | 6.40265188186281 |
| chr1 | 24842736 | 24842801 | + | 7.02772403811355 | 11.3699574206438 | 2.38645642073598 | 1.06710864697714 |
| chr1 | 24842926 | 24843002 | + | 19.0752509605939 | 14.780944646837 | 10.7390538933119 | 4.26843458790854 |
| chr1 | 24843973 | 24843997 | + | 35.1386201905678 | 62.534765813541 | 5.96614105183995 | 6.40265188186281 |
| chr1 | 24844045 | 24844059 | + | 12.0475269224804 | 14.780944646837 | 2.38645642073598 | 3.20132594093141 |
| chr1 | 25222735 | 25222784 | - | 20.0792115374673 | 55.7127913611547 | 15.5119667347839 | 25.6106075274513 |
| chr1 | 25232581 | 25232666 | + | 62.2455557661486 | 105.740604011988 | 45.3426719939836 | 69.3620620535138 |
| chr1 | 25234012 | 25234016 | + | 5.01980288436682 | 10.2329616785794 | 13.1255103140479 | 4.26843458790854 |
| chr1 | 25242282 | 25242305 | - | 2.00792115374673 | 11.3699574206438 | 20.2848795762558 | 3.20132594093141 |
| chr1 | 25242366 | 25242426 | - | 19.0752509605939 | 25.0139063254164 | 58.4681823080315 | 40.5501285851312 |
| chr1 | 25242494 | 25242497 | - | 6.02376346124019 | 12.5069531627082 | 4.77291284147196 | 8.53686917581709 |
| chr1 | 25242572 | 25242573 | - | 5.01980288436682 | 2.27399148412876 | 4.77291284147196 | 7.46976052883995 |
| chr1 | 25244076 | 25244105 | - | 6.02376346124019 | 4.54798296825753 | 3.57968463110397 | 3.20132594093141 |
| chr1 | 25244176 | 25244201 | - | 8.03168461498692 | 10.2329616785794 | 23.8645642073598 | 3.20132594093141 |
| chr1 | 25244427 | 25244455 | - | 2.00792115374673 | 1.13699574206438 | 4.77291284147196 | 5.33554323488568 |
| chr1 | 25244528 | 25244534 | - | 1.00396057687336 | 2.27399148412876 | 7.15936926220794 | 3.20132594093141 |
| chr1 | 25244790 | 25244804 | - | 2.00792115374673 | 1.13699574206438 | 1.19322821036799 | 4.26843458790854 |
| chr1 | 25360657 | 25360765 | + | 40.1584230749346 | 19.3289276150945 | 53.6952694665595 | 67.2278447595595 |
| chr1 | 25360809 | 25360823 | + | 1.00396057687336 | 2.27399148412876 | 5.96614105183995 | 2.13421729395427 |
| chr1 | 25360876 | 25360888 | + | 4.01584230749346 | 5.68497871032191 | 4.77291284147196 | 7.46976052883995 |
| chr1 | 25360910 | 25360930 | + | 26.1029749987075 | 25.0139063254164 | 46.5359002043516 | 29.8790421153598 |
| chr1 | 25361229 | 25361241 | + | 2.00792115374673 | 2.27399148412876 | 3.57968463110397 | 2.13421729395427 |
| chr1 | 25361457 | 25361499 | + | 3.01188173062009 | 5.68497871032191 | 9.54582568294392 | 12.8053037637256 |
| chr1 | 25361642 | 25361666 | + | 1.00396057687336 | 1.13699574206438 | 5.96614105183995 | 4.26843458790854 |
| chr1 | 25361893 | 25361956 | + | 30.1188173062009 | 22.7399148412876 | 16.7051949451519 | 33.0803680562912 |
| chr1 | 25362005 | 25362054 | + | 20.0792115374673 | 22.7399148412876 | 21.4781077866238 | 43.7514545260626 |
| chr1 | 25498567 | 25498604 | + | 18.0712903837206 | 17.0549361309657 | 29.8307052591997 | 32.0132594093141 |
| chr1 | 25499691 | 25499712 | + | 3.01188173062009 | 4.54798296825753 | 1.19322821036799 | 4.26843458790854 |
| chr1 | 25567136 | 25567151 | + | 6.02376346124019 | 5.68497871032191 | 17.8984231555198 | 5.33554323488568 |
| chr1 | 25567527 | 25567531 | + | 3.01188173062009 | 5.68497871032191 | 1.19322821036799 | 1.06710864697714 |
| chr1 | 25568776 | 25568779 | + | 22.087132691214 | 10.2329616785794 | 4.77291284147196 | 9.60397782279422 |
| chr1 | 25785912 | 25785915 | + | 3.01188173062009 | 1.13699574206438 | 1.19322821036799 | 3.20132594093141 |
| chr1 | 25817479 | 25817507 | + | 13.0514874993537 | 19.3289276150945 | 5.96614105183995 | 2.13421729395427 |
| chr1 | 25817722 | 25817725 | + | 2.00792115374673 | 1.13699574206438 | 2.38645642073598 | 2.13421729395427 |
| chr1 | 25818139 | 25818153 | + | 8.03168461498692 | 3.41098722619315 | 4.77291284147196 | 5.33554323488568 |
| chr1 | 25832164 | 25832172 | + | 1.00396057687336 | 1.13699574206438 | 11.9322821036799 | 6.40265188186281 |
| chr1 | 25832424 | 25832452 | + | 3.01188173062009 | 4.54798296825753 | 25.0577924177278 | 16.006629704657 |
| chr1 | 25832894 | 25832924 | + | 6.02376346124019 | 4.54798296825753 | 10.7390538933119 | 1.06710864697714 |
| chr1 | 25834052 | 25834055 | - | 7.02772403811355 | 2.27399148412876 | 2.38645642073598 | 1.06710864697714 |
| chr1 | 25834899 | 25834918 | - | 1.00396057687336 | 4.54798296825753 | 1.19322821036799 | 2.13421729395427 |
| chr1 | 25900150 | 25900308 | - | 1061.18632975515 | 880.034704357832 | 1374.59889834392 | 1358.42930760189 |
| chr1 | 25900341 | 25900374 | - | 19.0752509605939 | 17.0549361309657 | 31.0239334695677 | 30.9461507623369 |
| chr1 | 25900405 | 25900421 | - | 18.0712903837206 | 17.0549361309657 | 15.5119667347839 | 13.8724124107028 |
| chr1 | 25900459 | 25900510 | - | 22.087132691214 | 21.6029190992233 | 10.7390538933119 | 18.1408469986113 |
| chr1 | 25900559 | 25900608 | - | 32.1267384599477 | 30.6988850357383 | 38.1833027317757 | 19.2079556455884 |
| chr1 | 25900650 | 25900691 | - | 72.2851615348822 | 40.9318467143178 | 69.2072362013434 | 118.449059814462 |
| chr1 | 25900713 | 25900975 | - | 415.639678825573 | 367.249624686795 | 503.542304775292 | 672.278447595595 |
| chr1 | 25900998 | 25901040 | - | 11.043566345607 | 12.5069531627082 | 23.8645642073598 | 22.4092815865198 |
| chr1 | 26122680 | 26122686 | + | 5.01980288436682 | 3.41098722619315 | 2.38645642073598 | 5.33554323488568 |
| chr1 | 26123054 | 26123077 | + | 1.00396057687336 | 2.27399148412876 | 14.3187385244159 | 3.20132594093141 |
| chr1 | 26123159 | 26123160 | + | 2.00792115374673 | 2.27399148412876 | 1.19322821036799 | 3.20132594093141 |
| chr1 | 26124433 | 26124456 | + | 3.01188173062009 | 1.13699574206438 | 1.19322821036799 | 6.40265188186281 |
| chr1 | 26124677 | 26124700 | + | 2.00792115374673 | 3.41098722619315 | 25.0577924177278 | 9.60397782279422 |
| chr1 | 26124882 | 26124904 | + | 1.00396057687336 | 5.68497871032191 | 8.35259747257593 | 5.33554323488568 |
| chr1 | 26125097 | 26125246 | + | 83.3287278804893 | 162.590391115207 | 207.62170860403 | 197.41509969077 |
| chr1 | 26125474 | 26125476 | + | 3.01188173062009 | 2.27399148412876 | 7.15936926220794 | 10.6710864697714 |
| chr1 | 26162497 | 26162538 | + | 1.00396057687336 | 3.41098722619315 | 32.2171616799357 | 26.6777161744284 |
| chr1 | 26170815 | 26170839 | + | 2.00792115374673 | 9.09596593651506 | 2.38645642073598 | 16.006629704657 |
| chr1 | 26277503 | 26277506 | + | 2.00792115374673 | 1.13699574206438 | 11.9322821036799 | 4.26843458790854 |
| chr1 | 26278188 | 26278198 | + | 4.01584230749346 | 2.27399148412876 | 11.9322821036799 | 8.53686917581709 |
| chr1 | 26278434 | 26278445 | + | 2.00792115374673 | 1.13699574206438 | 15.5119667347839 | 12.8053037637256 |
| chr1 | 26278475 | 26278536 | + | 7.02772403811355 | 4.54798296825753 | 42.9562155732476 | 25.6106075274513 |
| chr1 | 26278679 | 26278703 | + | 1.00396057687336 | 1.13699574206438 | 10.7390538933119 | 9.60397782279422 |
| chr1 | 26281055 | 26281114 | + | 66.2613980736421 | 42.0688424563821 | 183.75714439667 | 110.979299285622 |
| chr1 | 26281315 | 26281366 | - | 20.0792115374673 | 17.0549361309657 | 27.4442488384638 | 44.8185631730397 |
| chr1 | 26281417 | 26281503 | + | 20.0792115374673 | 22.7399148412876 | 70.4004644117114 | 100.308212815851 |
| chr1 | 26281463 | 26281473 | - | 7.02772403811355 | 1.13699574206438 | 1.19322821036799 | 3.20132594093141 |
| chr1 | 26470130 | 26470133 | + | 6.02376346124019 | 1.13699574206438 | 7.15936926220794 | 3.20132594093141 |
| chr1 | 26471263 | 26471270 | + | 3.01188173062009 | 3.41098722619315 | 3.57968463110397 | 2.13421729395427 |
| chr1 | 26780760 | 26780765 | + | 7.02772403811355 | 2.27399148412876 | 4.77291284147196 | 7.46976052883995 |
| chr1 | 26780789 | 26780825 | + | 1.00396057687336 | 5.68497871032191 | 19.0916513658878 | 7.46976052883995 |
| chr1 | 26780854 | 26780896 | + | 12.0475269224804 | 18.1919318730301 | 26.2510206280958 | 28.8119334683827 |
| chr1 | 26780922 | 26781032 | + | 38.1505019211879 | 31.8358807778027 | 155.119667347839 | 94.9726695809651 |
| chr1 | 26781069 | 26781082 | + | 30.1188173062009 | 29.5618892936739 | 42.9562155732476 | 28.8119334683827 |
| chr1 | 26781113 | 26781142 | + | 10.0396057687336 | 10.2329616785794 | 28.6374770488317 | 28.8119334683827 |
| chr1 | 26781592 | 26781621 | + | 2.00792115374673 | 3.41098722619315 | 1.19322821036799 | 9.60397782279422 |
| chr1 | 26781684 | 26781784 | + | 10.0396057687336 | 12.5069531627082 | 33.4103898903037 | 54.4225409958339 |
| chr1 | 26781878 | 26781902 | + | 4.01584230749346 | 3.41098722619315 | 11.9322821036799 | 10.6710864697714 |
| chr1 | 26797858 | 26797890 | + | 19.0752509605939 | 27.2878978095452 | 13.1255103140479 | 7.46976052883995 |
| chr1 | 26854436 | 26854443 | + | 56.2217923049084 | 29.5618892936739 | 157.506123768575 | 57.6238669367653 |
| chr1 | 26854614 | 26854654 | + | 4.01584230749346 | 7.95897019445067 | 17.8984231555198 | 4.26843458790854 |
| chr1 | 26854766 | 26854786 | + | 3.01188173062009 | 3.41098722619315 | 4.77291284147196 | 10.6710864697714 |
| chr1 | 26855617 | 26855648 | + | 18.0712903837206 | 12.5069531627082 | 4.77291284147196 | 8.53686917581709 |
| chr1 | 26864010 | 26864217 | + | 362.429768251285 | 156.905412404885 | 5.96614105183995 | 2.13421729395427 |
| chr1 | 26864262 | 26864322 | + | 616.431794200246 | 260.372024932743 | 7.15936926220794 | 6.40265188186281 |
| chr1 | 26864354 | 26864430 | + | 132.522796147284 | 92.0966551072149 | 1.19322821036799 | 1.06710864697714 |
| chr1 | 26876158 | 26876190 | - | 4.01584230749346 | 4.54798296825753 | 2.38645642073598 | 2.13421729395427 |
| chr1 | 26877771 | 26877772 | - | 1.00396057687336 | 3.41098722619315 | 1.19322821036799 | 1.06710864697714 |
| chr1 | 26878217 | 26878224 | - | 1.00396057687336 | 1.13699574206438 | 2.38645642073598 | 4.26843458790854 |
| chr1 | 26879408 | 26879467 | - | 6.02376346124019 | 2.27399148412876 | 2.38645642073598 | 6.40265188186281 |
| chr1 | 26890519 | 26890538 | - | 5.01980288436682 | 6.82197445238629 | 1.19322821036799 | 5.33554323488568 |
| chr1 | 26946292 | 26946369 | + | 47.1861471130481 | 40.9318467143178 | 21.4781077866238 | 56.5567582897882 |
| chr1 | 27098913 | 27098942 | - | 38.1505019211879 | 31.8358807778027 | 16.7051949451519 | 14.9395210576799 |
| chr1 | 27098980 | 27099007 | - | 11.043566345607 | 9.09596593651506 | 7.15936926220794 | 5.33554323488568 |
| chr1 | 27099160 | 27099169 | - | 5.01980288436682 | 3.41098722619315 | 10.7390538933119 | 1.06710864697714 |
| chr1 | 27306512 | 27306528 | + | 4.01584230749346 | 6.82197445238629 | 7.15936926220794 | 3.20132594093141 |
| chr1 | 27306603 | 27306615 | + | 2.00792115374673 | 4.54798296825753 | 5.96614105183995 | 1.06710864697714 |
| chr1 | 27307080 | 27307087 | + | 13.0514874993537 | 10.2329616785794 | 32.2171616799357 | 2.13421729395427 |
| chr1 | 27307263 | 27307279 | + | 1.00396057687336 | 1.13699574206438 | 8.35259747257593 | 3.20132594093141 |
| chr1 | 27307535 | 27307537 | + | 1.00396057687336 | 2.27399148412876 | 4.77291284147196 | 16.006629704657 |
| chr1 | 27307726 | 27307755 | + | 6.02376346124019 | 4.54798296825753 | 26.2510206280958 | 5.33554323488568 |
| chr1 | 27308202 | 27308216 | + | 11.043566345607 | 17.0549361309657 | 20.2848795762558 | 13.8724124107028 |
| chr1 | 27335587 | 27335640 | + | 9.03564519186028 | 7.95897019445067 | 10.7390538933119 | 14.9395210576799 |
| chr1 | 27336357 | 27336378 | + | 9.03564519186028 | 14.780944646837 | 3.57968463110397 | 9.60397782279422 |
| chr1 | 27404272 | 27404274 | - | 1.00396057687336 | 3.41098722619315 | 2.38645642073598 | 2.13421729395427 |
| chr1 | 27404349 | 27404370 | - | 4.01584230749346 | 3.41098722619315 | 7.15936926220794 | 7.46976052883995 |
| chr1 | 27404920 | 27405005 | - | 23.0910932680874 | 18.1919318730301 | 57.2749540976635 | 75.7647139353766 |
| chr1 | 27405061 | 27405102 | - | 7.02772403811355 | 7.95897019445067 | 14.3187385244159 | 13.8724124107028 |
| chr1 | 27405144 | 27405162 | - | 5.01980288436682 | 5.68497871032191 | 7.15936926220794 | 2.13421729395427 |
| chr1 | 27405272 | 27405292 | - | 3.01188173062009 | 2.27399148412876 | 10.7390538933119 | 6.40265188186281 |
| chr1 | 27405313 | 27405321 | - | 3.01188173062009 | 2.27399148412876 | 3.57968463110397 | 2.13421729395427 |
| chr1 | 27405344 | 27405362 | - | 9.03564519186028 | 4.54798296825753 | 11.9322821036799 | 39.483019938154 |
| chr1 | 27405411 | 27405443 | - | 4.01584230749346 | 3.41098722619315 | 5.96614105183995 | 2.13421729395427 |
| chr1 | 27405468 | 27405481 | - | 2.00792115374673 | 3.41098722619315 | 5.96614105183995 | 5.33554323488568 |
| chr1 | 27405515 | 27405550 | - | 3.01188173062009 | 2.27399148412876 | 15.5119667347839 | 8.53686917581709 |
| chr1 | 27405667 | 27405690 | - | 91.3604124954762 | 56.8497871032191 | 107.390538933119 | 112.046407932599 |
| chr1 | 27405775 | 27405791 | - | 7.02772403811355 | 10.2329616785794 | 9.54582568294392 | 16.006629704657 |
| chr1 | 27405837 | 27405850 | - | 3.01188173062009 | 7.95897019445067 | 4.77291284147196 | 4.26843458790854 |
| chr1 | 27406636 | 27406670 | - | 1.00396057687336 | 1.13699574206438 | 21.4781077866238 | 23.476390233497 |
| chr1 | 27406734 | 27406740 | - | 1.00396057687336 | 2.27399148412876 | 3.57968463110397 | 3.20132594093141 |
| chr1 | 27406960 | 27406961 | - | 2.00792115374673 | 3.41098722619315 | 4.77291284147196 | 2.13421729395427 |
| chr1 | 27407245 | 27407265 | - | 2.00792115374673 | 1.13699574206438 | 3.57968463110397 | 3.20132594093141 |
| chr1 | 27407327 | 27407348 | - | 4.01584230749346 | 7.95897019445067 | 9.54582568294392 | 8.53686917581709 |
| chr1 | 27407525 | 27407547 | - | 3.01188173062009 | 1.13699574206438 | 4.77291284147196 | 2.13421729395427 |
| chr1 | 27407700 | 27407717 | - | 4.01584230749346 | 1.13699574206438 | 2.38645642073598 | 10.6710864697714 |
| chr1 | 27407768 | 27407787 | - | 6.02376346124019 | 4.54798296825753 | 4.77291284147196 | 7.46976052883995 |
| chr1 | 27407852 | 27407885 | - | 10.0396057687336 | 12.5069531627082 | 19.0916513658878 | 6.40265188186281 |
| chr1 | 27408096 | 27408125 | - | 4.01584230749346 | 4.54798296825753 | 2.38645642073598 | 2.13421729395427 |
| chr1 | 27408203 | 27408231 | - | 7.02772403811355 | 1.13699574206438 | 15.5119667347839 | 6.40265188186281 |
| chr1 | 27408299 | 27408335 | - | 11.043566345607 | 9.09596593651506 | 26.2510206280958 | 11.7381951167485 |
| chr1 | 27534409 | 27534435 | - | 18.0712903837206 | 14.780944646837 | 22.6713359969918 | 35.2145853502455 |
| chr1 | 27534551 | 27534575 | - | 2.00792115374673 | 1.13699574206438 | 4.77291284147196 | 7.46976052883995 |
| chr1 | 27534738 | 27534773 | - | 1.00396057687336 | 3.41098722619315 | 13.1255103140479 | 12.8053037637256 |
| chr1 | 27762733 | 27762747 | + | 6.02376346124019 | 4.54798296825753 | 8.35259747257593 | 13.8724124107028 |
| chr1 | 27823701 | 27823703 | + | 1.00396057687336 | 2.27399148412876 | 5.96614105183995 | 2.13421729395427 |
| chr1 | 27850471 | 27850520 | + | 125.495072109171 | 150.083437952498 | 65.6275515702394 | 70.429170700491 |
| chr1 | 27850597 | 27850605 | + | 11.043566345607 | 12.5069531627082 | 20.2848795762558 | 16.006629704657 |
| chr1 | 27850767 | 27850790 | + | 1.00396057687336 | 3.41098722619315 | 8.35259747257593 | 7.46976052883995 |
| chr1 | 27850823 | 27850833 | + | 7.02772403811355 | 9.09596593651506 | 16.7051949451519 | 16.006629704657 |
| chr1 | 27850860 | 27850886 | + | 8.03168461498692 | 11.3699574206438 | 16.7051949451519 | 9.60397782279422 |
| chr1 | 27850906 | 27850936 | + | 4.01584230749346 | 4.54798296825753 | 11.9322821036799 | 12.8053037637256 |
| chr1 | 27850961 | 27851022 | + | 30.1188173062009 | 51.1648083928972 | 78.7530618842873 | 72.5633879944452 |
| chr1 | 27851047 | 27851092 | + | 15.0594086531005 | 12.5069531627082 | 23.8645642073598 | 14.9395210576799 |
| chr1 | 27851225 | 27851260 | + | 3.01188173062009 | 3.41098722619315 | 1.19322821036799 | 2.13421729395427 |
| chr1 | 27851320 | 27851323 | + | 4.01584230749346 | 3.41098722619315 | 1.19322821036799 | 4.26843458790854 |
| chr1 | 27891599 | 27891633 | - | 22.087132691214 | 36.3838637460602 | 39.3765309421437 | 42.6843458790854 |
| chr1 | 27891794 | 27891804 | - | 12.0475269224804 | 7.95897019445067 | 4.77291284147196 | 19.2079556455884 |
| chr1 | 27891843 | 27891978 | - | 168.665376914725 | 187.604297440623 | 388.992396579965 | 428.977676084809 |
| chr1 | 27892012 | 27892035 | - | 3.01188173062009 | 4.54798296825753 | 8.35259747257593 | 6.40265188186281 |
| chr1 | 27892057 | 27892083 | - | 148.586165377258 | 213.755199508104 | 223.133675338814 | 328.669463268958 |
| chr1 | 27892107 | 27892169 | - | 28.1108961524542 | 29.5618892936739 | 51.3088130458235 | 35.2145853502455 |
| chr1 | 27970388 | 27970391 | - | 1.00396057687336 | 6.82197445238629 | 10.7390538933119 | 6.40265188186281 |
| chr1 | 27970426 | 27970427 | - | 1.00396057687336 | 3.41098722619315 | 1.19322821036799 | 1.06710864697714 |
| chr1 | 28200911 | 28200939 | - | 27.1069355755808 | 10.2329616785794 | 14.3187385244159 | 20.2750642925656 |
| chr1 | 28201050 | 28201072 | - | 12.0475269224804 | 3.41098722619315 | 8.35259747257593 | 11.7381951167485 |
| chr1 | 28237837 | 28237945 | + | 41.1623836518079 | 52.3018041349616 | 45.3426719939836 | 59.7580842307196 |
| chr1 | 28237950 | 28238080 | + | 672.653586505154 | 917.555563845956 | 346.036181006717 | 272.11270497917 |
| chr1 | 28281179 | 28281196 | + | 6.02376346124019 | 27.2878978095452 | 2.38645642073598 | 6.40265188186281 |
| chr1 | 28281430 | 28281445 | + | 2.00792115374673 | 12.5069531627082 | 19.0916513658878 | 7.46976052883995 |
| chr1 | 28281512 | 28281523 | + | 2.00792115374673 | 10.2329616785794 | 2.38645642073598 | 3.20132594093141 |
| chr1 | 28281688 | 28281707 | + | 1.00396057687336 | 2.27399148412876 | 4.77291284147196 | 6.40265188186281 |
| chr1 | 28281947 | 28281951 | + | 1.00396057687336 | 2.27399148412876 | 2.38645642073598 | 14.9395210576799 |
| chr1 | 28282315 | 28282366 | + | 4.01584230749346 | 30.6988850357383 | 7.15936926220794 | 13.8724124107028 |
| chr1 | 28335843 | 28335847 | + | 3.01188173062009 | 4.54798296825753 | 2.38645642073598 | 4.26843458790854 |
| chr1 | 28335906 | 28335929 | + | 5.01980288436682 | 7.95897019445067 | 5.96614105183995 | 1.06710864697714 |
| chr1 | 28498218 | 28498227 | + | 5.01980288436682 | 3.41098722619315 | 8.35259747257593 | 1.06710864697714 |
| chr1 | 28500115 | 28500134 | + | 73.2891221117556 | 75.0417189762492 | 20.2848795762558 | 49.0869977609482 |
| chr1 | 28506466 | 28506493 | + | 7.02772403811355 | 4.54798296825753 | 5.96614105183995 | 10.6710864697714 |
| chr1 | 28506534 | 28506579 | + | 63.249516343022 | 68.2197445238629 | 44.1494437836156 | 36.2816939972226 |
| chr1 | 28506599 | 28506654 | + | 18.0712903837206 | 4.54798296825753 | 44.1494437836156 | 21.3421729395427 |
| chr1 | 28506704 | 28506733 | + | 5.01980288436682 | 1.13699574206438 | 17.8984231555198 | 7.46976052883995 |
| chr1 | 28506937 | 28506981 | + | 3.01188173062009 | 2.27399148412876 | 5.96614105183995 | 11.7381951167485 |
| chr1 | 28507262 | 28507268 | + | 1.00396057687336 | 1.13699574206438 | 1.19322821036799 | 3.20132594093141 |
| chr1 | 28507546 | 28507581 | + | 4.01584230749346 | 1.13699574206438 | 9.54582568294392 | 13.8724124107028 |
| chr1 | 28508095 | 28508161 | + | 27.1069355755808 | 40.9318467143178 | 108.583767143487 | 24.5434988804741 |
| chr1 | 28508798 | 28508977 | + | 158.625771145992 | 145.535454984241 | 773.211880318457 | 478.064673845757 |
| chr1 | 28509028 | 28509044 | + | 8.03168461498692 | 6.82197445238629 | 25.0577924177278 | 22.4092815865198 |
| chr1 | 28509064 | 28509121 | + | 11.043566345607 | 9.09596593651506 | 50.1155848354556 | 33.0803680562912 |
| chr1 | 28509216 | 28509288 | + | 11.043566345607 | 3.41098722619315 | 19.0916513658878 | 5.33554323488568 |
| chr1 | 28509450 | 28509455 | + | 13.0514874993537 | 6.82197445238629 | 15.5119667347839 | 30.9461507623369 |
| chr1 | 28509503 | 28509560 | + | 81.3208067267425 | 67.0827487817985 | 108.583767143487 | 136.589906813073 |
| chr1 | 28509583 | 28509614 | + | 6.02376346124019 | 7.95897019445067 | 9.54582568294392 | 7.46976052883995 |
| chr1 | 28538051 | 28538107 | + | 11.043566345607 | 2.27399148412876 | 19.0916513658878 | 12.8053037637256 |
| chr1 | 28538211 | 28538252 | + | 2.00792115374673 | 1.13699574206438 | 1.19322821036799 | 7.46976052883995 |
| chr1 | 28538485 | 28538526 | + | 11.043566345607 | 3.41098722619315 | 4.77291284147196 | 8.53686917581709 |
| chr1 | 28538698 | 28538729 | + | 2.00792115374673 | 2.27399148412876 | 1.19322821036799 | 1.06710864697714 |
| chr1 | 28538732 | 28538901 | + | 151.598047107878 | 135.302493305661 | 128.868646719743 | 182.47557863309 |
| chr1 | 28538935 | 28538959 | + | 43.1703048055547 | 63.6717615556054 | 45.3426719939836 | 57.6238669367653 |
| chr1 | 28577640 | 28577683 | + | 39.1544624980612 | 38.657855230189 | 47.7291284147196 | 68.2949534065367 |
| chr1 | 28603300 | 28603320 | - | 4.01584230749346 | 9.09596593651506 | 2.38645642073598 | 7.46976052883995 |
| chr1 | 28603499 | 28603536 | - | 1.00396057687336 | 1.13699574206438 | 2.38645642073598 | 8.53686917581709 |
| chr1 | 28643227 | 28643244 | + | 5.01980288436682 | 1.13699574206438 | 9.54582568294392 | 6.40265188186281 |
| chr1 | 28648631 | 28648656 | + | 5.01980288436682 | 7.95897019445067 | 1.19322821036799 | 3.20132594093141 |
| chr1 | 28714749 | 28714778 | + | 2.00792115374673 | 7.95897019445067 | 2.38645642073598 | 6.40265188186281 |
| chr1 | 28716745 | 28716751 | + | 12.0475269224804 | 15.9179403889013 | 2.38645642073598 | 7.46976052883995 |
| chr1 | 28716850 | 28716853 | + | 8.03168461498692 | 3.41098722619315 | 1.19322821036799 | 4.26843458790854 |
| chr1 | 28716989 | 28716990 | + | 3.01188173062009 | 1.13699574206438 | 1.19322821036799 | 1.06710864697714 |
| chr1 | 28769461 | 28769493 | + | 11.043566345607 | 13.6439489047726 | 7.15936926220794 | 10.6710864697714 |
| chr1 | 28769606 | 28769614 | + | 1.00396057687336 | 1.13699574206438 | 3.57968463110397 | 3.20132594093141 |
| chr1 | 28769647 | 28769671 | + | 1.00396057687336 | 1.13699574206438 | 2.38645642073598 | 4.26843458790854 |
| chr1 | 29117692 | 29117708 | + | 3.01188173062009 | 1.13699574206438 | 7.15936926220794 | 6.40265188186281 |
| chr1 | 29117752 | 29117789 | + | 12.0475269224804 | 2.27399148412876 | 15.5119667347839 | 36.2816939972226 |
| chr1 | 29118020 | 29118032 | + | 4.01584230749346 | 3.41098722619315 | 11.9322821036799 | 13.8724124107028 |
| chr1 | 29118736 | 29118779 | + | 2.00792115374673 | 1.13699574206438 | 4.77291284147196 | 14.9395210576799 |
| chr1 | 29119061 | 29119069 | + | 1.00396057687336 | 1.13699574206438 | 3.57968463110397 | 7.46976052883995 |
| chr1 | 29119193 | 29119213 | + | 3.01188173062009 | 5.68497871032191 | 27.4442488384638 | 34.1474767032683 |
| chr1 | 29119234 | 29119263 | + | 16.0633692299738 | 9.09596593651506 | 87.1056593568632 | 50.1541064079254 |
| chr1 | 29119334 | 29119375 | + | 6.02376346124019 | 4.54798296825753 | 58.4681823080315 | 29.8790421153598 |
| chr1 | 29119611 | 29119811 | + | 261.029749987075 | 385.441556559825 | 1822.05947723192 | 1281.59748501954 |
| chr1 | 29119834 | 29119851 | + | 3.01188173062009 | 2.27399148412876 | 25.0577924177278 | 13.8724124107028 |
| chr1 | 29119875 | 29119909 | + | 34.1346596136944 | 34.1098722619315 | 291.147683329789 | 448.185631730397 |
| chr1 | 29119942 | 29119994 | + | 1.00396057687336 | 6.82197445238629 | 15.5119667347839 | 9.60397782279422 |
| chr1 | 29119979 | 29120022 | - | 4.01584230749346 | 4.54798296825753 | 15.5119667347839 | 18.1408469986113 |
| chr1 | 29147777 | 29147800 | - | 1.00396057687336 | 7.95897019445067 | 5.96614105183995 | 5.33554323488568 |
| chr1 | 29147876 | 29147956 | - | 20.0792115374673 | 37.5208594881246 | 27.4442488384638 | 28.8119334683827 |
| chr1 | 29148002 | 29148017 | - | 3.01188173062009 | 1.13699574206438 | 1.19322821036799 | 4.26843458790854 |
| chr1 | 29148066 | 29148130 | - | 21.0831721143407 | 31.8358807778027 | 28.6374770488317 | 41.6172372321083 |
| chr1 | 29148336 | 29148483 | - | 21.0831721143407 | 23.876910583352 | 65.6275515702394 | 75.7647139353766 |
| chr1 | 30718739 | 30718807 | + | 9.03564519186028 | 17.0549361309657 | 31.0239334695677 | 21.3421729395427 |
| chr1 | 30869592 | 30869625 | - | 4.01584230749346 | 4.54798296825753 | 3.57968463110397 | 11.7381951167485 |
| chr1 | 30871026 | 30871033 | - | 6.02376346124019 | 2.27399148412876 | 14.3187385244159 | 3.20132594093141 |
| chr1 | 30871367 | 30871385 | - | 5.01980288436682 | 5.68497871032191 | 11.9322821036799 | 5.33554323488568 |
| chr1 | 30871433 | 30871474 | - | 20.0792115374673 | 19.3289276150945 | 10.7390538933119 | 6.40265188186281 |
| chr1 | 30872506 | 30872518 | - | 11.043566345607 | 10.2329616785794 | 5.96614105183995 | 3.20132594093141 |
| chr1 | 30872541 | 30872562 | - | 10.0396057687336 | 5.68497871032191 | 17.8984231555198 | 4.26843458790854 |
| chr1 | 30872702 | 30872703 | - | 5.01980288436682 | 4.54798296825753 | 1.19322821036799 | 4.26843458790854 |
| chr1 | 30931566 | 30931628 | - | 4.01584230749346 | 6.82197445238629 | 7.15936926220794 | 5.33554323488568 |
| chr1 | 30931676 | 30931696 | - | 9.03564519186028 | 6.82197445238629 | 11.9322821036799 | 4.26843458790854 |
| chr1 | 30931835 | 30931878 | - | 4.01584230749346 | 3.41098722619315 | 1.19322821036799 | 4.26843458790854 |
| chr1 | 30932481 | 30932488 | - | 10.0396057687336 | 4.54798296825753 | 2.38645642073598 | 8.53686917581709 |
| chr1 | 30932869 | 30932890 | - | 7.02772403811355 | 4.54798296825753 | 3.57968463110397 | 1.06710864697714 |
| chr1 | 30933052 | 30933290 | - | 300.184212485136 | 297.892884420868 | 330.524214271933 | 346.810310267569 |
| chr1 | 31364030 | 31364058 | + | 1.00396057687336 | 7.95897019445067 | 21.4781077866238 | 21.3421729395427 |
| chr1 | 31364080 | 31364096 | + | 2.00792115374673 | 2.27399148412876 | 7.15936926220794 | 5.33554323488568 |
| chr1 | 31364151 | 31364217 | + | 11.043566345607 | 13.6439489047726 | 29.8307052591997 | 38.4159112911769 |
| chr1 | 31364310 | 31364380 | + | 11.043566345607 | 11.3699574206438 | 23.8645642073598 | 26.6777161744284 |
| chr1 | 31364528 | 31364566 | + | 14.0554480762271 | 9.09596593651506 | 31.0239334695677 | 20.2750642925656 |
| chr1 | 31364665 | 31364692 | + | 12.0475269224804 | 31.8358807778027 | 22.6713359969918 | 19.2079556455884 |
| chr1 | 31364748 | 31364761 | + | 1.00396057687336 | 1.13699574206438 | 5.96614105183995 | 6.40265188186281 |
| chr1 | 31364886 | 31364894 | + | 7.02772403811355 | 7.95897019445067 | 3.57968463110397 | 4.26843458790854 |
| chr1 | 31629914 | 31629918 | - | 3.01188173062009 | 5.68497871032191 | 7.15936926220794 | 6.40265188186281 |
| chr1 | 31906443 | 31906463 | - | 1.00396057687336 | 5.68497871032191 | 5.96614105183995 | 4.26843458790854 |
| chr1 | 31906574 | 31906609 | - | 3.01188173062009 | 5.68497871032191 | 2.38645642073598 | 1.06710864697714 |
| chr1 | 31906705 | 31906750 | - | 19.0752509605939 | 23.876910583352 | 7.15936926220794 | 23.476390233497 |
| chr1 | 31906858 | 31906892 | - | 6.02376346124019 | 10.2329616785794 | 5.96614105183995 | 6.40265188186281 |
| chr1 | 31907244 | 31907275 | - | 4.01584230749346 | 4.54798296825753 | 2.38645642073598 | 9.60397782279422 |
| chr1 | 31907311 | 31907520 | - | 136.538638454778 | 113.699574206438 | 205.235252183294 | 169.670274869365 |
| chr1 | 31907583 | 31907634 | - | 10.0396057687336 | 15.9179403889013 | 11.9322821036799 | 9.60397782279422 |
| chr1 | 31907674 | 31907679 | - | 3.01188173062009 | 2.27399148412876 | 11.9322821036799 | 1.06710864697714 |
| chr1 | 31907703 | 31907742 | - | 7.02772403811355 | 12.5069531627082 | 25.0577924177278 | 12.8053037637256 |
| chr1 | 31907816 | 31907880 | - | 59.2336740355285 | 76.1787147183136 | 232.679501021758 | 108.845081991668 |
| chr1 | 31907905 | 31908001 | - | 78.3089249961224 | 92.0966551072149 | 70.4004644117114 | 80.0331485232852 |
| chr1 | 31908025 | 31908051 | - | 27.1069355755808 | 25.0139063254164 | 25.0577924177278 | 55.4896496428111 |
| chr1 | 31908076 | 31908088 | - | 22.087132691214 | 29.5618892936739 | 17.8984231555198 | 25.6106075274513 |
| chr1 | 31908262 | 31908326 | - | 38.1505019211879 | 44.3428339405109 | 42.9562155732476 | 26.6777161744284 |
| chr1 | 31908393 | 31908483 | - | 483.908998052962 | 563.949888063933 | 477.291284147196 | 439.64876255458 |
| chr1 | 31908572 | 31908583 | - | 8.03168461498692 | 2.27399148412876 | 3.57968463110397 | 1.06710864697714 |
| chr1 | 31908740 | 31908756 | - | 14.0554480762271 | 30.6988850357383 | 15.5119667347839 | 11.7381951167485 |
| chr1 | 31908784 | 31908872 | - | 36.1425807674411 | 61.3977700714766 | 42.9562155732476 | 71.4962793474681 |
| chr1 | 31908944 | 31908957 | - | 22.087132691214 | 22.7399148412876 | 34.6036181006717 | 36.2816939972226 |
| chr1 | 32042634 | 32042692 | + | 37.1465413443145 | 42.0688424563821 | 42.9562155732476 | 54.4225409958339 |
| chr1 | 32042715 | 32042747 | + | 6.02376346124019 | 7.95897019445067 | 9.54582568294392 | 10.6710864697714 |
| chr1 | 32042785 | 32042866 | + | 15.0594086531005 | 18.1919318730301 | 11.9322821036799 | 23.476390233497 |
| chr1 | 32042898 | 32042904 | + | 2.00792115374673 | 1.13699574206438 | 8.35259747257593 | 4.26843458790854 |
| chr1 | 32042976 | 32043006 | + | 8.03168461498692 | 9.09596593651506 | 16.7051949451519 | 13.8724124107028 |
| chr1 | 32043092 | 32043096 | + | 5.01980288436682 | 3.41098722619315 | 13.1255103140479 | 5.33554323488568 |
| chr1 | 32043128 | 32043164 | + | 3.01188173062009 | 3.41098722619315 | 8.35259747257593 | 16.006629704657 |
| chr1 | 32043191 | 32043268 | + | 22.087132691214 | 21.6029190992233 | 27.4442488384638 | 32.0132594093141 |
| chr1 | 32043303 | 32043450 | + | 156.617849992245 | 140.987472015983 | 190.916513658878 | 279.58246550801 |
| chr1 | 32043505 | 32043702 | + | 99.3920971104631 | 144.398459242177 | 300.693509012733 | 310.528616270347 |
| chr1 | 32043727 | 32043811 | + | 29.1148567293276 | 59.1237785873479 | 96.6514850398071 | 29.8790421153598 |
| chr1 | 32043833 | 32043858 | + | 11.043566345607 | 25.0139063254164 | 27.4442488384638 | 25.6106075274513 |
| chr1 | 32102757 | 32102761 | + | 1.00396057687336 | 6.82197445238629 | 7.15936926220794 | 7.46976052883995 |
| chr1 | 32170731 | 32170778 | + | 9.03564519186028 | 6.82197445238629 | 15.5119667347839 | 4.26843458790854 |
| chr1 | 32170912 | 32170924 | + | 7.02772403811355 | 4.54798296825753 | 4.77291284147196 | 1.06710864697714 |
| chr1 | 32170964 | 32170966 | + | 17.0673298068472 | 3.41098722619315 | 5.96614105183995 | 4.26843458790854 |
| chr1 | 32171054 | 32171075 | + | 2.00792115374673 | 5.68497871032191 | 1.19322821036799 | 3.20132594093141 |
| chr1 | 32171105 | 32171125 | + | 13.0514874993537 | 10.2329616785794 | 8.35259747257593 | 12.8053037637256 |
| chr1 | 32171178 | 32171217 | + | 16.0633692299738 | 19.3289276150945 | 21.4781077866238 | 14.9395210576799 |
| chr1 | 32171256 | 32171260 | + | 4.01584230749346 | 4.54798296825753 | 5.96614105183995 | 8.53686917581709 |
| chr1 | 32171366 | 32171399 | + | 63.249516343022 | 50.0278126508328 | 51.3088130458235 | 21.3421729395427 |
| chr1 | 32171428 | 32171439 | + | 13.0514874993537 | 6.82197445238629 | 13.1255103140479 | 3.20132594093141 |
| chr1 | 32171613 | 32171631 | + | 12.0475269224804 | 5.68497871032191 | 3.57968463110397 | 2.13421729395427 |
| chr1 | 32171802 | 32171825 | + | 9.03564519186028 | 4.54798296825753 | 5.96614105183995 | 1.06710864697714 |
| chr1 | 32171851 | 32171856 | + | 5.01980288436682 | 2.27399148412876 | 5.96614105183995 | 1.06710864697714 |
| chr1 | 32171876 | 32171880 | + | 5.01980288436682 | 3.41098722619315 | 1.19322821036799 | 1.06710864697714 |
| chr1 | 32171913 | 32171957 | + | 24.0950538449607 | 10.2329616785794 | 32.2171616799357 | 14.9395210576799 |
| chr1 | 32172001 | 32172007 | + | 5.01980288436682 | 5.68497871032191 | 2.38645642073598 | 2.13421729395427 |
| chr1 | 32172073 | 32172132 | + | 10.0396057687336 | 6.82197445238629 | 31.0239334695677 | 8.53686917581709 |
| chr1 | 32172161 | 32172205 | + | 6.02376346124019 | 9.09596593651506 | 1.19322821036799 | 2.13421729395427 |
| chr1 | 32172306 | 32172313 | + | 4.01584230749346 | 3.41098722619315 | 3.57968463110397 | 1.06710864697714 |
| chr1 | 32172471 | 32172486 | + | 2.00792115374673 | 4.54798296825753 | 11.9322821036799 | 11.7381951167485 |
| chr1 | 32172980 | 32172995 | + | 11.043566345607 | 15.9179403889013 | 4.77291284147196 | 2.13421729395427 |
| chr1 | 32173042 | 32173082 | + | 9.03564519186028 | 5.68497871032191 | 1.19322821036799 | 1.06710864697714 |
| chr1 | 32173105 | 32173118 | + | 156.617849992245 | 120.521548658824 | 77.5598336739193 | 28.8119334683827 |
| chr1 | 32173494 | 32173513 | + | 3.01188173062009 | 2.27399148412876 | 3.57968463110397 | 1.06710864697714 |
| chr1 | 32173886 | 32173917 | + | 9.03564519186028 | 7.95897019445067 | 7.15936926220794 | 4.26843458790854 |
| chr1 | 32174197 | 32174212 | + | 7.02772403811355 | 6.82197445238629 | 1.19322821036799 | 5.33554323488568 |
| chr1 | 32174323 | 32174343 | + | 2.00792115374673 | 1.13699574206438 | 5.96614105183995 | 1.06710864697714 |
| chr1 | 32174668 | 32174674 | + | 1.00396057687336 | 1.13699574206438 | 5.96614105183995 | 5.33554323488568 |
| chr1 | 32174782 | 32174801 | + | 7.02772403811355 | 4.54798296825753 | 19.0916513658878 | 10.6710864697714 |
| chr1 | 32174917 | 32174936 | + | 1.00396057687336 | 1.13699574206438 | 3.57968463110397 | 3.20132594093141 |
| chr1 | 32175416 | 32175419 | + | 4.01584230749346 | 7.95897019445067 | 1.19322821036799 | 2.13421729395427 |
| chr1 | 32175459 | 32175462 | + | 4.01584230749346 | 3.41098722619315 | 1.19322821036799 | 2.13421729395427 |
| chr1 | 32176234 | 32176287 | + | 8.03168461498692 | 3.41098722619315 | 3.57968463110397 | 5.33554323488568 |
| chr1 | 32176420 | 32176446 | + | 7.02772403811355 | 1.13699574206438 | 1.19322821036799 | 2.13421729395427 |
| chr1 | 32195628 | 32195635 | + | 4.01584230749346 | 2.27399148412876 | 1.19322821036799 | 1.06710864697714 |
| chr1 | 32195736 | 32195763 | + | 3.01188173062009 | 5.68497871032191 | 9.54582568294392 | 29.8790421153598 |
| chr1 | 32195784 | 32195794 | + | 13.0514874993537 | 7.95897019445067 | 1.19322821036799 | 1.06710864697714 |
| chr1 | 32195839 | 32195863 | + | 4.01584230749346 | 6.82197445238629 | 4.77291284147196 | 4.26843458790854 |
| chr1 | 32195910 | 32195955 | + | 22.087132691214 | 25.0139063254164 | 20.2848795762558 | 49.0869977609482 |
| chr1 | 32196013 | 32196022 | + | 10.0396057687336 | 9.09596593651506 | 5.96614105183995 | 6.40265188186281 |
| chr1 | 32196044 | 32196090 | + | 13.0514874993537 | 12.5069531627082 | 14.3187385244159 | 5.33554323488568 |
| chr1 | 32196112 | 32196190 | + | 19.0752509605939 | 26.1509020674808 | 19.0916513658878 | 6.40265188186281 |
| chr1 | 32196221 | 32196242 | + | 7.02772403811355 | 4.54798296825753 | 4.77291284147196 | 1.06710864697714 |
| chr1 | 32196308 | 32196330 | + | 6.02376346124019 | 1.13699574206438 | 1.19322821036799 | 4.26843458790854 |
| chr1 | 32196354 | 32196413 | + | 12.0475269224804 | 5.68497871032191 | 3.57968463110397 | 1.06710864697714 |
| chr1 | 32196434 | 32196444 | + | 5.01980288436682 | 2.27399148412876 | 3.57968463110397 | 1.06710864697714 |
| chr1 | 32196616 | 32196628 | + | 2.00792115374673 | 1.13699574206438 | 3.57968463110397 | 2.13421729395427 |
| chr1 | 32196680 | 32196700 | + | 16.0633692299738 | 11.3699574206438 | 11.9322821036799 | 8.53686917581709 |
| chr1 | 32196769 | 32196782 | + | 2.00792115374673 | 3.41098722619315 | 1.19322821036799 | 1.06710864697714 |
| chr1 | 32197340 | 32197359 | + | 3.01188173062009 | 1.13699574206438 | 5.96614105183995 | 1.06710864697714 |
| chr1 | 32197453 | 32197474 | + | 7.02772403811355 | 4.54798296825753 | 8.35259747257593 | 11.7381951167485 |
| chr1 | 32197555 | 32197672 | + | 40.1584230749346 | 43.2058381984465 | 62.0478669391355 | 51.2212150549025 |
| chr1 | 32197692 | 32197812 | + | 42.1663442286813 | 38.657855230189 | 60.8546387287675 | 93.9055609339879 |
| chr1 | 32197835 | 32197887 | + | 6.02376346124019 | 10.2329616785794 | 20.2848795762558 | 70.429170700491 |
| chr1 | 32198111 | 32198122 | + | 4.01584230749346 | 5.68497871032191 | 4.77291284147196 | 4.26843458790854 |
| chr1 | 32198186 | 32198234 | + | 1.00396057687336 | 3.41098722619315 | 8.35259747257593 | 3.20132594093141 |
| chr1 | 32205346 | 32205357 | + | 1.00396057687336 | 2.27399148412876 | 27.4442488384638 | 44.8185631730397 |
| chr1 | 32208652 | 32208662 | + | 1.00396057687336 | 6.82197445238629 | 2.38645642073598 | 4.26843458790854 |
| chr1 | 32216229 | 32216236 | - | 6.02376346124019 | 4.54798296825753 | 2.38645642073598 | 3.20132594093141 |
| chr1 | 32231504 | 32231584 | + | 77.3049644192491 | 127.343523111211 | 192.109741869246 | 290.253551977781 |
| chr1 | 32333014 | 32333048 | + | 8.03168461498692 | 7.95897019445067 | 26.2510206280958 | 19.2079556455884 |
| chr1 | 32333249 | 32333295 | + | 2.00792115374673 | 2.27399148412876 | 7.15936926220794 | 10.6710864697714 |
| chr1 | 32333384 | 32333406 | + | 39.1544624980612 | 21.6029190992233 | 39.3765309421437 | 89.6371263460794 |
| chr1 | 32333536 | 32333597 | + | 38.1505019211879 | 29.5618892936739 | 79.9462900946553 | 72.5633879944452 |
| chr1 | 32333856 | 32333888 | - | 8.03168461498692 | 6.82197445238629 | 38.1833027317757 | 27.7448248214055 |
| chr1 | 32333932 | 32333947 | - | 1.00396057687336 | 1.13699574206438 | 14.3187385244159 | 42.6843458790854 |
| chr1 | 32333971 | 32334074 | - | 25.0990144218341 | 22.7399148412876 | 162.279036610047 | 321.199702740118 |
| chr1 | 32334110 | 32334160 | - | 6.02376346124019 | 3.41098722619315 | 20.2848795762558 | 71.4962793474681 |
| chr1 | 32365353 | 32365399 | - | 79.3128855729958 | 84.1376849127643 | 140.800928823423 | 108.845081991668 |
| chr1 | 32365426 | 32365451 | - | 4.01584230749346 | 1.13699574206438 | 5.96614105183995 | 4.26843458790854 |
| chr1 | 32365860 | 32365867 | - | 1.00396057687336 | 1.13699574206438 | 3.57968463110397 | 1.06710864697714 |
| chr1 | 32366379 | 32366449 | - | 4.01584230749346 | 10.2329616785794 | 31.0239334695677 | 11.7381951167485 |
| chr1 | 32366607 | 32366615 | - | 11.043566345607 | 2.27399148412876 | 4.77291284147196 | 5.33554323488568 |
| chr1 | 32367444 | 32367446 | - | 2.00792115374673 | 3.41098722619315 | 3.57968463110397 | 1.06710864697714 |
| chr1 | 32368311 | 32368325 | - | 1.00396057687336 | 1.13699574206438 | 2.38645642073598 | 5.33554323488568 |
| chr1 | 32621849 | 32621911 | - | 23.0910932680874 | 28.4248935516096 | 15.5119667347839 | 18.1408469986113 |
| chr1 | 32679654 | 32679720 | + | 3.01188173062009 | 3.41098722619315 | 8.35259747257593 | 2.13421729395427 |
| chr1 | 32679847 | 32679877 | + | 24.0950538449607 | 15.9179403889013 | 36.9900745214077 | 22.4092815865198 |
| chr1 | 32680324 | 32680333 | + | 3.01188173062009 | 6.82197445238629 | 14.3187385244159 | 8.53686917581709 |
| chr1 | 32680500 | 32680508 | + | 4.01584230749346 | 5.68497871032191 | 4.77291284147196 | 1.06710864697714 |
| chr1 | 32680573 | 32680593 | + | 12.0475269224804 | 14.780944646837 | 26.2510206280958 | 22.4092815865198 |
| chr1 | 32680629 | 32680633 | + | 21.0831721143407 | 62.534765813541 | 13.1255103140479 | 4.26843458790854 |
| chr1 | 32680654 | 32680658 | - | 38.1505019211879 | 21.6029190992233 | 1.19322821036799 | 1.06710864697714 |
| chr1 | 32680723 | 32680746 | - | 11.043566345607 | 3.41098722619315 | 1.19322821036799 | 2.13421729395427 |
| chr1 | 32683910 | 32683915 | + | 6.02376346124019 | 5.68497871032191 | 8.35259747257593 | 2.13421729395427 |
| chr1 | 32684046 | 32684051 | + | 1.00396057687336 | 2.27399148412876 | 4.77291284147196 | 2.13421729395427 |
| chr1 | 32684216 | 32684232 | + | 1.00396057687336 | 2.27399148412876 | 5.96614105183995 | 4.26843458790854 |
| chr1 | 32684637 | 32684638 | + | 2.00792115374673 | 1.13699574206438 | 1.19322821036799 | 2.13421729395427 |
| chr1 | 32684684 | 32684704 | + | 2.00792115374673 | 3.41098722619315 | 2.38645642073598 | 3.20132594093141 |
| chr1 | 32684973 | 32684978 | + | 4.01584230749346 | 3.41098722619315 | 1.19322821036799 | 1.06710864697714 |
| chr1 | 32685397 | 32685444 | + | 3.01188173062009 | 7.95897019445067 | 15.5119667347839 | 13.8724124107028 |
| chr1 | 32772984 | 32773035 | + | 1.00396057687336 | 4.54798296825753 | 2.38645642073598 | 1.06710864697714 |
| chr1 | 32773089 | 32773121 | + | 2.00792115374673 | 2.27399148412876 | 4.77291284147196 | 4.26843458790854 |
| chr1 | 32773202 | 32773226 | + | 12.0475269224804 | 22.7399148412876 | 21.4781077866238 | 4.26843458790854 |
| chr1 | 32773497 | 32773500 | + | 2.00792115374673 | 1.13699574206438 | 3.57968463110397 | 1.06710864697714 |
| chr1 | 32773766 | 32773804 | + | 12.0475269224804 | 11.3699574206438 | 21.4781077866238 | 4.26843458790854 |
| chr1 | 32773876 | 32773896 | + | 5.01980288436682 | 1.13699574206438 | 8.35259747257593 | 4.26843458790854 |
| chr1 | 32773947 | 32773966 | + | 3.01188173062009 | 6.82197445238629 | 8.35259747257593 | 9.60397782279422 |
| chr1 | 32773993 | 32773999 | + | 2.00792115374673 | 2.27399148412876 | 3.57968463110397 | 6.40265188186281 |
| chr1 | 32774026 | 32774072 | + | 15.0594086531005 | 11.3699574206438 | 27.4442488384638 | 10.6710864697714 |
| chr1 | 32774259 | 32774274 | + | 4.01584230749346 | 1.13699574206438 | 5.96614105183995 | 4.26843458790854 |
| chr1 | 32774307 | 32774351 | + | 10.0396057687336 | 2.27399148412876 | 14.3187385244159 | 24.5434988804741 |
| chr1 | 32774373 | 32774385 | + | 2.00792115374673 | 1.13699574206438 | 2.38645642073598 | 1.06710864697714 |
| chr1 | 32774435 | 32774442 | + | 9.03564519186028 | 3.41098722619315 | 8.35259747257593 | 4.26843458790854 |
| chr1 | 32774467 | 32774486 | + | 2.00792115374673 | 3.41098722619315 | 4.77291284147196 | 6.40265188186281 |
| chr1 | 32774522 | 32774551 | + | 4.01584230749346 | 1.13699574206438 | 3.57968463110397 | 8.53686917581709 |
| chr1 | 32774594 | 32774604 | + | 13.0514874993537 | 3.41098722619315 | 15.5119667347839 | 11.7381951167485 |
| chr1 | 32774713 | 32774754 | + | 34.1346596136944 | 30.6988850357383 | 13.1255103140479 | 19.2079556455884 |
| chr1 | 32775269 | 32775345 | - | 61.2415951892752 | 138.713480531855 | 59.6614105183995 | 88.5700176991023 |
| chr1 | 32775372 | 32775399 | - | 6.02376346124019 | 9.09596593651506 | 1.19322821036799 | 4.26843458790854 |
| chr1 | 32775704 | 32775762 | - | 12.0475269224804 | 13.6439489047726 | 34.6036181006717 | 25.6106075274513 |
| chr1 | 32775961 | 32776094 | - | 32.1267384599477 | 44.3428339405109 | 85.9124311464952 | 85.3686917581709 |
| chr1 | 32856258 | 32856272 | + | 2.00792115374673 | 4.54798296825753 | 2.38645642073598 | 14.9395210576799 |
| chr1 | 33008906 | 33008923 | - | 4.01584230749346 | 4.54798296825753 | 7.15936926220794 | 20.2750642925656 |
| chr1 | 33008951 | 33008959 | - | 13.0514874993537 | 10.2329616785794 | 15.5119667347839 | 11.7381951167485 |
| chr1 | 33010263 | 33010308 | - | 3.01188173062009 | 1.13699574206438 | 3.57968463110397 | 16.006629704657 |
| chr1 | 33010382 | 33010396 | - | 2.00792115374673 | 6.82197445238629 | 7.15936926220794 | 6.40265188186281 |
| chr1 | 33010575 | 33010583 | - | 10.0396057687336 | 12.5069531627082 | 10.7390538933119 | 12.8053037637256 |
| chr1 | 33010659 | 33010728 | - | 49.1940682667949 | 51.1648083928972 | 90.6853439879672 | 128.053037637256 |
| chr1 | 33010749 | 33010773 | - | 21.0831721143407 | 19.3289276150945 | 47.7291284147196 | 62.959410171651 |
| chr1 | 33010786 | 33010829 | - | 7.02772403811355 | 3.41098722619315 | 21.4781077866238 | 12.8053037637256 |
| chr1 | 33012072 | 33012076 | - | 5.01980288436682 | 12.5069531627082 | 17.8984231555198 | 8.53686917581709 |
| chr1 | 33013073 | 33013142 | - | 13.0514874993537 | 26.1509020674808 | 33.4103898903037 | 49.0869977609482 |
| chr1 | 33013164 | 33013207 | - | 8.03168461498692 | 14.780944646837 | 8.35259747257593 | 3.20132594093141 |
| chr1 | 33013270 | 33013299 | - | 16.0633692299738 | 13.6439489047726 | 39.3765309421437 | 36.2816939972226 |
| chr1 | 33013367 | 33013397 | - | 10.0396057687336 | 7.95897019445067 | 9.54582568294392 | 10.6710864697714 |
| chr1 | 33014521 | 33014523 | - | 5.01980288436682 | 2.27399148412876 | 4.77291284147196 | 2.13421729395427 |
| chr1 | 33323662 | 33323691 | - | 4.01584230749346 | 6.82197445238629 | 7.15936926220794 | 5.33554323488568 |
| chr1 | 33323725 | 33323748 | - | 3.01188173062009 | 7.95897019445067 | 4.77291284147196 | 9.60397782279422 |
| chr1 | 33324437 | 33324589 | - | 21.0831721143407 | 14.780944646837 | 54.8884976769275 | 60.8251928776967 |
| chr1 | 33324627 | 33324777 | - | 18.0712903837206 | 27.2878978095452 | 58.4681823080315 | 48.0198891139711 |
| chr1 | 34855079 | 34855080 | - | 3.01188173062009 | 5.68497871032191 | 1.19322821036799 | 1.06710864697714 |
| chr1 | 34855139 | 34855152 | - | 4.01584230749346 | 9.09596593651506 | 4.77291284147196 | 10.6710864697714 |
| chr1 | 34855481 | 34855484 | - | 1.00396057687336 | 1.13699574206438 | 3.57968463110397 | 3.20132594093141 |
| chr1 | 34855507 | 34855519 | - | 2.00792115374673 | 4.54798296825753 | 2.38645642073598 | 3.20132594093141 |
| chr1 | 34855662 | 34855687 | - | 1.00396057687336 | 3.41098722619315 | 14.3187385244159 | 5.33554323488568 |
| chr1 | 34981655 | 34981675 | - | 14.0554480762271 | 18.1919318730301 | 8.35259747257593 | 7.46976052883995 |
| chr1 | 34981815 | 34981853 | - | 3.01188173062009 | 5.68497871032191 | 2.38645642073598 | 5.33554323488568 |
| chr1 | 35115303 | 35115305 | + | 5.01980288436682 | 4.54798296825753 | 1.19322821036799 | 2.13421729395427 |
| chr1 | 35115433 | 35115479 | + | 39.1544624980612 | 54.5757956190903 | 54.8884976769275 | 37.3488026441998 |
| chr1 | 35115520 | 35115564 | + | 6.02376346124019 | 6.82197445238629 | 5.96614105183995 | 2.13421729395427 |
| chr1 | 35183636 | 35183656 | - | 4.01584230749346 | 10.2329616785794 | 9.54582568294392 | 1.06710864697714 |
| chr1 | 35183698 | 35183719 | - | 11.043566345607 | 6.82197445238629 | 15.5119667347839 | 12.8053037637256 |
| chr1 | 35183755 | 35183859 | - | 65.2574374967687 | 56.8497871032191 | 40.5697591525116 | 69.3620620535138 |
| chr1 | 35183891 | 35183895 | - | 22.087132691214 | 31.8358807778027 | 19.0916513658878 | 32.0132594093141 |
| chr1 | 35183935 | 35183951 | - | 2.00792115374673 | 2.27399148412876 | 4.77291284147196 | 3.20132594093141 |
| chr1 | 35183995 | 35184016 | - | 45.1782259593014 | 53.438799877026 | 36.9900745214077 | 29.8790421153598 |
| chr1 | 35184172 | 35184223 | - | 10.0396057687336 | 15.9179403889013 | 20.2848795762558 | 20.2750642925656 |
| chr1 | 35184275 | 35184394 | - | 90.3564519186028 | 75.0417189762492 | 89.4921157775992 | 158.999188399593 |
| chr1 | 35184436 | 35184588 | - | 96.380215379843 | 113.699574206438 | 100.231169670911 | 157.932079752616 |
| chr1 | 35419923 | 35419929 | + | 1.00396057687336 | 3.41098722619315 | 4.77291284147196 | 6.40265188186281 |
| chr1 | 35420086 | 35420094 | + | 2.00792115374673 | 3.41098722619315 | 4.77291284147196 | 5.33554323488568 |
| chr1 | 35420137 | 35420157 | + | 4.01584230749346 | 3.41098722619315 | 4.77291284147196 | 4.26843458790854 |
| chr1 | 35421809 | 35421811 | + | 3.01188173062009 | 5.68497871032191 | 4.77291284147196 | 8.53686917581709 |
| chr1 | 35433518 | 35433644 | - | 15.0594086531005 | 26.1509020674808 | 17.8984231555198 | 27.7448248214055 |
| chr1 | 35434470 | 35434472 | - | 1.00396057687336 | 2.27399148412876 | 1.19322821036799 | 2.13421729395427 |
| chr1 | 35434705 | 35434735 | - | 2.00792115374673 | 6.82197445238629 | 7.15936926220794 | 11.7381951167485 |
| chr1 | 35566240 | 35566269 | + | 10.0396057687336 | 6.82197445238629 | 5.96614105183995 | 10.6710864697714 |
| chr1 | 35601845 | 35601877 | - | 46.1821865361748 | 36.3838637460602 | 48.9223566250876 | 35.2145853502455 |
| chr1 | 35601903 | 35601921 | - | 10.0396057687336 | 10.2329616785794 | 22.6713359969918 | 69.3620620535138 |
| chr1 | 35601954 | 35601968 | - | 2.00792115374673 | 2.27399148412876 | 2.38645642073598 | 3.20132594093141 |
| chr1 | 35601996 | 35602000 | - | 1.00396057687336 | 2.27399148412876 | 3.57968463110397 | 3.20132594093141 |
| chr1 | 35602051 | 35602073 | - | 41.1623836518079 | 20.4659233571589 | 26.2510206280958 | 53.3554323488568 |
| chr1 | 35602187 | 35602207 | - | 1.00396057687336 | 2.27399148412876 | 4.77291284147196 | 14.9395210576799 |
| chr1 | 35602433 | 35602482 | - | 8.03168461498692 | 6.82197445238629 | 13.1255103140479 | 27.7448248214055 |
| chr1 | 35602505 | 35602507 | - | 9.03564519186028 | 10.2329616785794 | 9.54582568294392 | 3.20132594093141 |
| chr1 | 35603062 | 35603086 | - | 11.043566345607 | 2.27399148412876 | 15.5119667347839 | 4.26843458790854 |
| chr1 | 35603155 | 35603194 | - | 216.855484604647 | 304.714858873254 | 400.924678683644 | 409.76972043922 |
| chr1 | 35603219 | 35603377 | - | 99.3920971104631 | 128.480518853275 | 206.428480393662 | 284.918008742895 |
| chr1 | 35855614 | 35855634 | + | 6.02376346124019 | 4.54798296825753 | 4.77291284147196 | 6.40265188186281 |
| chr1 | 35857256 | 35857264 | + | 1.00396057687336 | 1.13699574206438 | 1.19322821036799 | 1.06710864697714 |
| chr1 | 35857540 | 35857546 | + | 2.00792115374673 | 1.13699574206438 | 3.57968463110397 | 1.06710864697714 |
| chr1 | 35920025 | 35920062 | + | 19.0752509605939 | 19.3289276150945 | 22.6713359969918 | 19.2079556455884 |
| chr1 | 35920370 | 35920372 | + | 1.00396057687336 | 1.13699574206438 | 5.96614105183995 | 1.06710864697714 |
| chr1 | 35920444 | 35920472 | + | 23.0910932680874 | 20.4659233571589 | 38.1833027317757 | 11.7381951167485 |
| chr1 | 35921001 | 35921004 | + | 1.00396057687336 | 2.27399148412876 | 5.96614105183995 | 1.06710864697714 |
| chr1 | 35921254 | 35921272 | + | 4.01584230749346 | 3.41098722619315 | 11.9322821036799 | 21.3421729395427 |
| chr1 | 35923095 | 35923106 | + | 3.01188173062009 | 1.13699574206438 | 1.19322821036799 | 1.06710864697714 |
| chr1 | 35923135 | 35923150 | + | 1.00396057687336 | 1.13699574206438 | 1.19322821036799 | 2.13421729395427 |
| chr1 | 35923576 | 35923595 | + | 2.00792115374673 | 2.27399148412876 | 5.96614105183995 | 12.8053037637256 |
| chr1 | 35923696 | 35923707 | + | 1.00396057687336 | 1.13699574206438 | 4.77291284147196 | 3.20132594093141 |
| chr1 | 35924003 | 35924014 | + | 9.03564519186028 | 6.82197445238629 | 5.96614105183995 | 9.60397782279422 |
| chr1 | 35924074 | 35924117 | + | 13.0514874993537 | 15.9179403889013 | 26.2510206280958 | 28.8119334683827 |
| chr1 | 35924280 | 35924304 | + | 5.01980288436682 | 2.27399148412876 | 10.7390538933119 | 11.7381951167485 |
| chr1 | 35924472 | 35924502 | + | 3.01188173062009 | 4.54798296825753 | 21.4781077866238 | 9.60397782279422 |
| chr1 | 35924523 | 35924546 | + | 4.01584230749346 | 1.13699574206438 | 9.54582568294392 | 9.60397782279422 |
| chr1 | 35924695 | 35924703 | + | 3.01188173062009 | 2.27399148412876 | 13.1255103140479 | 8.53686917581709 |
| chr1 | 35924767 | 35924815 | + | 5.01980288436682 | 6.82197445238629 | 29.8307052591997 | 24.5434988804741 |
| chr1 | 35925035 | 35925037 | + | 1.00396057687336 | 3.41098722619315 | 14.3187385244159 | 9.60397782279422 |
| chr1 | 35925848 | 35925851 | + | 5.01980288436682 | 1.13699574206438 | 4.77291284147196 | 4.26843458790854 |
| chr1 | 35925996 | 35925998 | + | 3.01188173062009 | 1.13699574206438 | 2.38645642073598 | 4.26843458790854 |
| chr1 | 35926037 | 35926047 | + | 1.00396057687336 | 1.13699574206438 | 9.54582568294392 | 3.20132594093141 |
| chr1 | 35928088 | 35928089 | + | 8.03168461498692 | 7.95897019445067 | 17.8984231555198 | 26.6777161744284 |
| chr1 | 35929359 | 35929362 | + | 2.00792115374673 | 1.13699574206438 | 2.38645642073598 | 3.20132594093141 |
| chr1 | 35929475 | 35929525 | + | 35.1386201905678 | 30.6988850357383 | 77.5598336739193 | 149.395210576799 |
| chr1 | 36056306 | 36056311 | + | 2.00792115374673 | 1.13699574206438 | 2.38645642073598 | 4.26843458790854 |
| chr1 | 36057018 | 36057024 | + | 3.01188173062009 | 6.82197445238629 | 14.3187385244159 | 3.20132594093141 |
| chr1 | 36058394 | 36058400 | + | 1.00396057687336 | 1.13699574206438 | 1.19322821036799 | 5.33554323488568 |
| chr1 | 36061617 | 36061620 | + | 4.01584230749346 | 2.27399148412876 | 2.38645642073598 | 1.06710864697714 |
| chr1 | 36061705 | 36061711 | + | 6.02376346124019 | 7.95897019445067 | 2.38645642073598 | 9.60397782279422 |
| chr1 | 36093820 | 36093862 | + | 2.00792115374673 | 3.41098722619315 | 1.19322821036799 | 1.06710864697714 |
| chr1 | 36093893 | 36093906 | + | 3.01188173062009 | 10.2329616785794 | 13.1255103140479 | 18.1408469986113 |
| chr1 | 36136629 | 36136633 | - | 49.1940682667949 | 43.2058381984465 | 33.4103898903037 | 39.483019938154 |
| chr1 | 36136654 | 36136658 | - | 4.01584230749346 | 4.54798296825753 | 1.19322821036799 | 6.40265188186281 |
| chr1 | 36136689 | 36136731 | - | 39.1544624980612 | 23.876910583352 | 46.5359002043516 | 64.0265188186281 |
| chr1 | 36136751 | 36136874 | - | 157.621810569118 | 127.343523111211 | 134.834787771583 | 145.12677598889 |
| chr1 | 36136936 | 36136952 | - | 4.01584230749346 | 1.13699574206438 | 4.77291284147196 | 1.06710864697714 |
| chr1 | 36136975 | 36136981 | - | 19.0752509605939 | 12.5069531627082 | 31.0239334695677 | 34.1474767032683 |
| chr1 | 36137046 | 36137057 | - | 3.01188173062009 | 2.27399148412876 | 5.96614105183995 | 3.20132594093141 |
| chr1 | 36137079 | 36137129 | - | 6.02376346124019 | 5.68497871032191 | 4.77291284147196 | 4.26843458790854 |
| chr1 | 36137918 | 36137979 | - | 10.0396057687336 | 6.82197445238629 | 7.15936926220794 | 7.46976052883995 |
| chr1 | 36180763 | 36180766 | + | 6.02376346124019 | 5.68497871032191 | 3.57968463110397 | 8.53686917581709 |
| chr1 | 36303926 | 36304110 | + | 62.2455557661486 | 51.1648083928972 | 66.8207797806074 | 40.5501285851312 |
| chr1 | 36304138 | 36304234 | + | 19.0752509605939 | 23.876910583352 | 38.1833027317757 | 24.5434988804741 |
| chr1 | 36304282 | 36304320 | + | 852.362529765486 | 656.046543171148 | 683.719764540858 | 700.023272417001 |
| chr1 | 36304342 | 36304441 | + | 32.1267384599477 | 39.7948509722534 | 69.2072362013434 | 56.5567582897882 |
| chr1 | 36304471 | 36304476 | + | 2.00792115374673 | 3.41098722619315 | 9.54582568294392 | 2.13421729395427 |
| chr1 | 36304502 | 36304523 | + | 7.02772403811355 | 3.41098722619315 | 9.54582568294392 | 5.33554323488568 |
| chr1 | 36304558 | 36304605 | + | 11.043566345607 | 7.95897019445067 | 13.1255103140479 | 3.20132594093141 |
| chr1 | 36304619 | 36304692 | + | 22.087132691214 | 19.3289276150945 | 27.4442488384638 | 33.0803680562912 |
| chr1 | 36304735 | 36304749 | + | 14.0554480762271 | 11.3699574206438 | 13.1255103140479 | 17.0737383516342 |
| chr1 | 36304970 | 36304988 | + | 2.00792115374673 | 1.13699574206438 | 3.57968463110397 | 22.4092815865198 |
| chr1 | 36305283 | 36305294 | + | 16.0633692299738 | 26.1509020674808 | 9.54582568294392 | 23.476390233497 |
| chr1 | 36322053 | 36322115 | - | 4.01584230749346 | 4.54798296825753 | 8.35259747257593 | 10.6710864697714 |
| chr1 | 36339650 | 36339672 | - | 2.00792115374673 | 1.13699574206438 | 11.9322821036799 | 17.0737383516342 |
| chr1 | 36339720 | 36339783 | - | 6.02376346124019 | 5.68497871032191 | 8.35259747257593 | 17.0737383516342 |
| chr1 | 36340008 | 36340058 | - | 4.01584230749346 | 2.27399148412876 | 11.9322821036799 | 9.60397782279422 |
| chr1 | 36340082 | 36340130 | - | 2.00792115374673 | 1.13699574206438 | 32.2171616799357 | 16.006629704657 |
| chr1 | 36340338 | 36340526 | - | 7.02772403811355 | 15.9179403889013 | 242.225326704702 | 125.918820343302 |
| chr1 | 36340578 | 36340618 | - | 3.01188173062009 | 3.41098722619315 | 10.7390538933119 | 6.40265188186281 |
| chr1 | 36340672 | 36340706 | - | 2.00792115374673 | 3.41098722619315 | 22.6713359969918 | 25.6106075274513 |
| chr1 | 36340771 | 36340796 | - | 1.00396057687336 | 1.13699574206438 | 21.4781077866238 | 12.8053037637256 |
| chr1 | 36340824 | 36340837 | - | 3.01188173062009 | 1.13699574206438 | 4.77291284147196 | 5.33554323488568 |
| chr1 | 36341212 | 36341242 | - | 6.02376346124019 | 1.13699574206438 | 13.1255103140479 | 4.26843458790854 |
| chr1 | 36341331 | 36341342 | - | 5.01980288436682 | 2.27399148412876 | 10.7390538933119 | 10.6710864697714 |
| chr1 | 36341387 | 36341422 | - | 1.00396057687336 | 2.27399148412876 | 8.35259747257593 | 10.6710864697714 |
| chr1 | 36393440 | 36393489 | - | 25.0990144218341 | 40.9318467143178 | 20.2848795762558 | 24.5434988804741 |
| chr1 | 36393541 | 36393574 | - | 32.1267384599477 | 34.1098722619315 | 29.8307052591997 | 41.6172372321083 |
| chr1 | 37484263 | 37484291 | + | 8.03168461498692 | 10.2329616785794 | 1.19322821036799 | 5.33554323488568 |
| chr1 | 37493388 | 37493426 | - | 9.03564519186028 | 3.41098722619315 | 7.15936926220794 | 5.33554323488568 |
| chr1 | 37493458 | 37493472 | - | 8.03168461498692 | 10.2329616785794 | 13.1255103140479 | 7.46976052883995 |
| chr1 | 37493497 | 37493507 | - | 6.02376346124019 | 5.68497871032191 | 1.19322821036799 | 5.33554323488568 |
| chr1 | 37493603 | 37493620 | - | 10.0396057687336 | 4.54798296825753 | 4.77291284147196 | 2.13421729395427 |
| chr1 | 37493640 | 37493731 | - | 228.903011527127 | 287.659922742289 | 393.765309421437 | 433.246110672717 |
| chr1 | 37493763 | 37493880 | - | 42.1663442286813 | 67.0827487817985 | 82.3327465153913 | 53.3554323488568 |
| chr1 | 37493911 | 37493950 | - | 15.0594086531005 | 20.4659233571589 | 47.7291284147196 | 16.006629704657 |
| chr1 | 37494065 | 37494069 | - | 2.00792115374673 | 1.13699574206438 | 3.57968463110397 | 3.20132594093141 |
| chr1 | 37537189 | 37537196 | - | 3.01188173062009 | 3.41098722619315 | 2.38645642073598 | 2.13421729395427 |
| chr1 | 37537650 | 37537673 | - | 10.0396057687336 | 9.09596593651506 | 57.2749540976635 | 6.40265188186281 |
| chr1 | 37682003 | 37682007 | - | 1.00396057687336 | 5.68497871032191 | 1.19322821036799 | 4.26843458790854 |
| chr1 | 37682470 | 37682474 | - | 1.00396057687336 | 2.27399148412876 | 2.38645642073598 | 6.40265188186281 |
| chr1 | 37682550 | 37682561 | - | 4.01584230749346 | 2.27399148412876 | 2.38645642073598 | 3.20132594093141 |
| chr1 | 37682803 | 37682817 | - | 3.01188173062009 | 2.27399148412876 | 3.57968463110397 | 3.20132594093141 |
| chr1 | 37708351 | 37708398 | + | 6.02376346124019 | 4.54798296825753 | 25.0577924177278 | 21.3421729395427 |
| chr1 | 37708434 | 37708497 | + | 9.03564519186028 | 6.82197445238629 | 31.0239334695677 | 12.8053037637256 |
| chr1 | 37708693 | 37708720 | + | 5.01980288436682 | 4.54798296825753 | 20.2848795762558 | 13.8724124107028 |
| chr1 | 37708769 | 37708813 | + | 4.01584230749346 | 9.09596593651506 | 28.6374770488317 | 12.8053037637256 |
| chr1 | 37708836 | 37708917 | + | 12.0475269224804 | 23.876910583352 | 73.9801490428153 | 56.5567582897882 |
| chr1 | 37708990 | 37709006 | + | 3.01188173062009 | 2.27399148412876 | 4.77291284147196 | 11.7381951167485 |
| chr1 | 37709292 | 37709319 | + | 4.01584230749346 | 2.27399148412876 | 11.9322821036799 | 12.8053037637256 |
| chr1 | 37709489 | 37709519 | + | 6.02376346124019 | 6.82197445238629 | 9.54582568294392 | 9.60397782279422 |
| chr1 | 37709669 | 37709698 | + | 5.01980288436682 | 10.2329616785794 | 13.1255103140479 | 10.6710864697714 |
| chr1 | 37803194 | 37803279 | - | 4.01584230749346 | 9.09596593651506 | 3.57968463110397 | 5.33554323488568 |
| chr1 | 37803737 | 37803739 | - | 7.02772403811355 | 6.82197445238629 | 1.19322821036799 | 3.20132594093141 |
| chr1 | 37803800 | 37803806 | - | 4.01584230749346 | 5.68497871032191 | 1.19322821036799 | 5.33554323488568 |
| chr1 | 37803823 | 37803864 | - | 2.00792115374673 | 11.3699574206438 | 16.7051949451519 | 13.8724124107028 |
| chr1 | 37803887 | 37803914 | - | 6.02376346124019 | 7.95897019445067 | 13.1255103140479 | 6.40265188186281 |
| chr1 | 37809081 | 37809082 | + | 1.00396057687336 | 1.13699574206438 | 2.38645642073598 | 2.13421729395427 |
| chr1 | 37809153 | 37809178 | + | 4.01584230749346 | 3.41098722619315 | 1.19322821036799 | 2.13421729395427 |
| chr1 | 37809334 | 37809353 | + | 4.01584230749346 | 1.13699574206438 | 1.19322821036799 | 2.13421729395427 |
| chr1 | 37809391 | 37809427 | + | 28.1108961524542 | 18.1919318730301 | 17.8984231555198 | 5.33554323488568 |
| chr1 | 37861352 | 37861367 | - | 1.00396057687336 | 1.13699574206438 | 4.77291284147196 | 3.20132594093141 |
| chr1 | 37957040 | 37957058 | - | 4.01584230749346 | 1.13699574206438 | 11.9322821036799 | 24.5434988804741 |
| chr1 | 37957120 | 37957171 | - | 30.1188173062009 | 18.1919318730301 | 42.9562155732476 | 55.4896496428111 |
| chr1 | 37957226 | 37957253 | - | 8.03168461498692 | 7.95897019445067 | 14.3187385244159 | 26.6777161744284 |
| chr1 | 37957279 | 37957389 | - | 37.1465413443145 | 46.6168254246397 | 115.743136405695 | 149.395210576799 |
| chr1 | 37957425 | 37957429 | - | 13.0514874993537 | 1.13699574206438 | 9.54582568294392 | 11.7381951167485 |
| chr1 | 37957463 | 37957480 | - | 3.01188173062009 | 3.41098722619315 | 22.6713359969918 | 17.0737383516342 |
| chr1 | 37957553 | 37957581 | - | 23.0910932680874 | 11.3699574206438 | 65.6275515702394 | 36.2816939972226 |
| chr1 | 37957668 | 37957671 | - | 2.00792115374673 | 1.13699574206438 | 1.19322821036799 | 2.13421729395427 |
| chr1 | 37957700 | 37957715 | - | 2.00792115374673 | 5.68497871032191 | 4.77291284147196 | 10.6710864697714 |
| chr1 | 37957837 | 37957843 | - | 4.01584230749346 | 2.27399148412876 | 1.19322821036799 | 5.33554323488568 |
| chr1 | 37957865 | 37957881 | - | 2.00792115374673 | 1.13699574206438 | 4.77291284147196 | 8.53686917581709 |
| chr1 | 37957925 | 37957986 | - | 5.01980288436682 | 10.2329616785794 | 14.3187385244159 | 25.6106075274513 |
| chr1 | 37958026 | 37958046 | - | 55.217831728035 | 80.7266976865711 | 57.2749540976635 | 87.5029090521251 |
| chr1 | 37958074 | 37958086 | - | 5.01980288436682 | 5.68497871032191 | 9.54582568294392 | 17.0737383516342 |
| chr1 | 37996812 | 37996823 | - | 1.00396057687336 | 5.68497871032191 | 7.15936926220794 | 12.8053037637256 |
| chr1 | 38023772 | 38023780 | + | 3.01188173062009 | 3.41098722619315 | 8.35259747257593 | 9.60397782279422 |
| chr1 | 38024698 | 38024707 | + | 15.0594086531005 | 3.41098722619315 | 7.15936926220794 | 10.6710864697714 |
| chr1 | 38024788 | 38024796 | + | 1.00396057687336 | 6.82197445238629 | 2.38645642073598 | 3.20132594093141 |
| chr1 | 38839427 | 38839429 | - | 2.00792115374673 | 2.27399148412876 | 4.77291284147196 | 1.06710864697714 |
| chr1 | 38839533 | 38839540 | - | 35.1386201905678 | 21.6029190992233 | 15.5119667347839 | 20.2750642925656 |
| chr1 | 38863517 | 38863534 | - | 1.00396057687336 | 5.68497871032191 | 3.57968463110397 | 2.13421729395427 |
| chr1 | 38863706 | 38863719 | - | 2.00792115374673 | 2.27399148412876 | 4.77291284147196 | 8.53686917581709 |
| chr1 | 38863791 | 38863804 | - | 4.01584230749346 | 2.27399148412876 | 2.38645642073598 | 3.20132594093141 |
| chr1 | 38863853 | 38863870 | - | 3.01188173062009 | 1.13699574206438 | 10.7390538933119 | 6.40265188186281 |
| chr1 | 38864678 | 38864689 | - | 14.0554480762271 | 12.5069531627082 | 33.4103898903037 | 32.0132594093141 |
| chr1 | 39004046 | 39004131 | + | 10.0396057687336 | 13.6439489047726 | 17.8984231555198 | 7.46976052883995 |
| chr1 | 39004159 | 39004221 | + | 4.01584230749346 | 11.3699574206438 | 4.77291284147196 | 4.26843458790854 |
| chr1 | 39004307 | 39004339 | + | 3.01188173062009 | 4.54798296825753 | 9.54582568294392 | 17.0737383516342 |
| chr1 | 39004365 | 39004437 | + | 6.02376346124019 | 5.68497871032191 | 21.4781077866238 | 25.6106075274513 |
| chr1 | 39005632 | 39005637 | + | 3.01188173062009 | 3.41098722619315 | 1.19322821036799 | 3.20132594093141 |
| chr1 | 39005758 | 39005781 | + | 2.00792115374673 | 2.27399148412876 | 3.57968463110397 | 2.13421729395427 |
| chr1 | 39034550 | 39034605 | + | 46.1821865361748 | 51.1648083928972 | 15.5119667347839 | 44.8185631730397 |
| chr1 | 39485828 | 39485921 | + | 26.1029749987075 | 27.2878978095452 | 54.8884976769275 | 28.8119334683827 |
| chr1 | 39486011 | 39486012 | + | 1.00396057687336 | 4.54798296825753 | 2.38645642073598 | 4.26843458790854 |
| chr1 | 39486043 | 39486050 | + | 1.00396057687336 | 1.13699574206438 | 4.77291284147196 | 3.20132594093141 |
| chr1 | 39486105 | 39486117 | + | 1.00396057687336 | 2.27399148412876 | 3.57968463110397 | 6.40265188186281 |
| chr1 | 39486279 | 39486285 | + | 7.02772403811355 | 5.68497871032191 | 16.7051949451519 | 13.8724124107028 |
| chr1 | 39486329 | 39486332 | + | 5.01980288436682 | 1.13699574206438 | 3.57968463110397 | 1.06710864697714 |
| chr1 | 39486433 | 39486488 | + | 5.01980288436682 | 10.2329616785794 | 16.7051949451519 | 17.0737383516342 |
| chr1 | 39486564 | 39486597 | + | 20.0792115374673 | 14.780944646837 | 17.8984231555198 | 16.006629704657 |
| chr1 | 39486640 | 39486682 | + | 26.1029749987075 | 13.6439489047726 | 14.3187385244159 | 20.2750642925656 |
| chr1 | 39486714 | 39486759 | + | 7.02772403811355 | 7.95897019445067 | 17.8984231555198 | 18.1408469986113 |
| chr1 | 39486804 | 39486851 | + | 26.1029749987075 | 14.780944646837 | 31.0239334695677 | 32.0132594093141 |
| chr1 | 39560901 | 39560934 | - | 3.01188173062009 | 2.27399148412876 | 1.19322821036799 | 1.06710864697714 |
| chr1 | 39560964 | 39561015 | - | 388.532743249992 | 238.76910583352 | 335.297127113405 | 241.166554216833 |
| chr1 | 39561038 | 39561139 | - | 80.3168461498692 | 64.8087572976698 | 88.2988875672312 | 74.6976052883995 |
| chr1 | 39753426 | 39753468 | + | 1.00396057687336 | 2.27399148412876 | 9.54582568294392 | 12.8053037637256 |
| chr1 | 39753549 | 39753596 | + | 51.2019894205416 | 38.657855230189 | 151.539982716735 | 350.011636208501 |
| chr1 | 39753633 | 39753677 | + | 1.00396057687336 | 5.68497871032191 | 9.54582568294392 | 10.6710864697714 |
| chr1 | 39799405 | 39799441 | + | 10.0396057687336 | 20.4659233571589 | 22.6713359969918 | 4.26843458790854 |
| chr1 | 39841132 | 39841439 | - | 868.42589899546 | 850.472815064158 | 3996.1212765224 | 3239.74185222258 |
| chr1 | 39841479 | 39841482 | - | 6.02376346124019 | 11.3699574206438 | 7.15936926220794 | 4.26843458790854 |
| chr1 | 39841509 | 39841564 | - | 62.2455557661486 | 73.9047232341848 | 114.549908195327 | 129.120146284233 |
| chr1 | 39841601 | 39841795 | - | 36.1425807674411 | 44.3428339405109 | 192.109741869246 | 119.516168461439 |
| chr1 | 40071785 | 40071798 | + | 6.02376346124019 | 5.68497871032191 | 17.8984231555198 | 17.0737383516342 |
| chr1 | 40071834 | 40071877 | + | 44.174265382428 | 37.5208594881246 | 65.6275515702394 | 25.6106075274513 |
| chr1 | 40071989 | 40072004 | + | 1.00396057687336 | 2.27399148412876 | 9.54582568294392 | 13.8724124107028 |
| chr1 | 40072033 | 40072057 | + | 7.02772403811355 | 5.68497871032191 | 87.1056593568632 | 32.0132594093141 |
| chr1 | 40072106 | 40072132 | + | 6.02376346124019 | 3.41098722619315 | 15.5119667347839 | 12.8053037637256 |
| chr1 | 40072494 | 40072497 | - | 1.00396057687336 | 2.27399148412876 | 8.35259747257593 | 1.06710864697714 |
| chr1 | 40072555 | 40072587 | + | 6.02376346124019 | 9.09596593651506 | 11.9322821036799 | 14.9395210576799 |
| chr1 | 40072735 | 40072789 | - | 99.3920971104631 | 144.398459242177 | 82.3327465153913 | 74.6976052883995 |
| chr1 | 40072833 | 40072978 | - | 40.1584230749346 | 40.9318467143178 | 16.7051949451519 | 32.0132594093141 |
| chr1 | 40072999 | 40073067 | - | 8.03168461498692 | 9.09596593651506 | 4.77291284147196 | 5.33554323488568 |
| chr1 | 40073096 | 40073139 | - | 10.0396057687336 | 13.6439489047726 | 4.77291284147196 | 3.20132594093141 |
| chr1 | 40073191 | 40073279 | - | 19.0752509605939 | 12.5069531627082 | 3.57968463110397 | 13.8724124107028 |
| chr1 | 40073423 | 40073431 | - | 6.02376346124019 | 3.41098722619315 | 5.96614105183995 | 5.33554323488568 |
| chr1 | 40073775 | 40073819 | - | 2.00792115374673 | 1.13699574206438 | 1.19322821036799 | 1.06710864697714 |
| chr1 | 40240783 | 40240797 | + | 4.01584230749346 | 2.27399148412876 | 8.35259747257593 | 6.40265188186281 |
| chr1 | 40293272 | 40293316 | + | 4.01584230749346 | 4.54798296825753 | 10.7390538933119 | 3.20132594093141 |
| chr1 | 40293628 | 40293635 | + | 16.0633692299738 | 15.9179403889013 | 3.57968463110397 | 11.7381951167485 |
| chr1 | 40293901 | 40293923 | + | 10.0396057687336 | 12.5069531627082 | 1.19322821036799 | 2.13421729395427 |
| chr1 | 40293959 | 40293975 | + | 11.043566345607 | 13.6439489047726 | 4.77291284147196 | 3.20132594093141 |
| chr1 | 40294032 | 40294045 | + | 4.01584230749346 | 10.2329616785794 | 8.35259747257593 | 12.8053037637256 |
| chr1 | 40422709 | 40422763 | + | 2.00792115374673 | 3.41098722619315 | 17.8984231555198 | 36.2816939972226 |
| chr1 | 40422812 | 40422953 | + | 19.0752509605939 | 11.3699574206438 | 310.239334695677 | 186.744013220999 |
| chr1 | 40422977 | 40423066 | + | 2.00792115374673 | 2.27399148412876 | 78.7530618842873 | 104.576647403759 |
| chr1 | 40423161 | 40423177 | + | 2.00792115374673 | 1.13699574206438 | 2.38645642073598 | 11.7381951167485 |
| chr1 | 40770892 | 40771003 | + | 20.0792115374673 | 13.6439489047726 | 31.0239334695677 | 18.1408469986113 |
| chr1 | 40771432 | 40771441 | + | 2.00792115374673 | 3.41098722619315 | 1.19322821036799 | 4.26843458790854 |
| chr1 | 40771562 | 40771582 | + | 2.00792115374673 | 2.27399148412876 | 5.96614105183995 | 1.06710864697714 |
| chr1 | 41010237 | 41010255 | + | 1.00396057687336 | 1.13699574206438 | 2.38645642073598 | 6.40265188186281 |
| chr1 | 41011657 | 41011690 | + | 6.02376346124019 | 13.6439489047726 | 7.15936926220794 | 11.7381951167485 |
| chr1 | 41011771 | 41011786 | + | 2.00792115374673 | 6.82197445238629 | 4.77291284147196 | 3.20132594093141 |
| chr1 | 41012352 | 41012355 | + | 4.01584230749346 | 13.6439489047726 | 2.38645642073598 | 5.33554323488568 |
| chr1 | 41012399 | 41012420 | + | 24.0950538449607 | 31.8358807778027 | 7.15936926220794 | 6.40265188186281 |
| chr1 | 41012509 | 41012523 | + | 5.01980288436682 | 17.0549361309657 | 2.38645642073598 | 2.13421729395427 |
| chr1 | 41027322 | 41027356 | - | 2.00792115374673 | 3.41098722619315 | 1.19322821036799 | 4.26843458790854 |
| chr1 | 41027568 | 41027604 | - | 4.01584230749346 | 1.13699574206438 | 5.96614105183995 | 2.13421729395427 |
| chr1 | 42176779 | 42177101 | - | 687.712995158255 | 764.061138667265 | 813.781639470969 | 737.372075061201 |
| chr1 | 42177127 | 42177150 | - | 2.00792115374673 | 3.41098722619315 | 8.35259747257593 | 19.2079556455884 |
| chr1 | 42177328 | 42177350 | - | 5.01980288436682 | 4.54798296825753 | 1.19322821036799 | 2.13421729395427 |
| chr1 | 42177501 | 42177507 | - | 4.01584230749346 | 2.27399148412876 | 11.9322821036799 | 3.20132594093141 |
| chr1 | 42177679 | 42177726 | - | 10.0396057687336 | 5.68497871032191 | 3.57968463110397 | 2.13421729395427 |
| chr1 | 42177962 | 42177965 | - | 1.00396057687336 | 2.27399148412876 | 1.19322821036799 | 1.06710864697714 |
| chr1 | 42178110 | 42178116 | - | 3.01188173062009 | 2.27399148412876 | 7.15936926220794 | 2.13421729395427 |
| chr1 | 42178260 | 42178264 | - | 5.01980288436682 | 10.2329616785794 | 10.7390538933119 | 12.8053037637256 |
| chr1 | 42178288 | 42178315 | - | 11.043566345607 | 9.09596593651506 | 16.7051949451519 | 16.006629704657 |
| chr1 | 42178499 | 42178521 | - | 9.03564519186028 | 15.9179403889013 | 13.1255103140479 | 8.53686917581709 |
| chr1 | 42178592 | 42178611 | - | 4.01584230749346 | 1.13699574206438 | 2.38645642073598 | 1.06710864697714 |
| chr1 | 42179037 | 42179046 | - | 3.01188173062009 | 3.41098722619315 | 2.38645642073598 | 5.33554323488568 |
| chr1 | 42460261 | 42460373 | + | 23.0910932680874 | 23.876910583352 | 2.38645642073598 | 3.20132594093141 |
| chr1 | 42667360 | 42667441 | + | 502.984249013556 | 392.263531012212 | 1493.92171938072 | 1630.54201258106 |
| chr1 | 42676573 | 42676586 | + | 25.0990144218341 | 9.09596593651506 | 65.6275515702394 | 46.952780466994 |
| chr1 | 42676613 | 42676633 | + | 21.0831721143407 | 7.95897019445067 | 9.54582568294392 | 30.9461507623369 |
| chr1 | 42676688 | 42676698 | + | 3.01188173062009 | 5.68497871032191 | 15.5119667347839 | 12.8053037637256 |
| chr1 | 42676724 | 42676733 | + | 2.00792115374673 | 4.54798296825753 | 3.57968463110397 | 1.06710864697714 |
| chr1 | 42702027 | 42702044 | + | 102.403978841083 | 104.603608269923 | 83.5259747257593 | 182.47557863309 |
| chr1 | 42702334 | 42702342 | + | 33.130699036821 | 73.9047232341848 | 42.9562155732476 | 6.40265188186281 |
| chr1 | 42775477 | 42775509 | + | 3.01188173062009 | 3.41098722619315 | 7.15936926220794 | 19.2079556455884 |
| chr1 | 42807083 | 42807153 | - | 7.02772403811355 | 3.41098722619315 | 35.7968463110397 | 27.7448248214055 |
| chr1 | 42807257 | 42807280 | - | 1.00396057687336 | 2.27399148412876 | 3.57968463110397 | 1.06710864697714 |
| chr1 | 42807369 | 42807412 | - | 3.01188173062009 | 1.13699574206438 | 3.57968463110397 | 2.13421729395427 |
| chr1 | 42852447 | 42852450 | + | 3.01188173062009 | 2.27399148412876 | 4.77291284147196 | 1.06710864697714 |
| chr1 | 42926294 | 42926492 | - | 14.0554480762271 | 11.3699574206438 | 39.3765309421437 | 51.2212150549025 |
| chr1 | 42926596 | 42926758 | - | 11.043566345607 | 6.82197445238629 | 66.8207797806074 | 59.7580842307196 |
| chr1 | 42926781 | 42927020 | - | 17.0673298068472 | 11.3699574206438 | 112.163451774591 | 133.388580872142 |
| chr1 | 42959028 | 42959103 | + | 3.01188173062009 | 1.13699574206438 | 33.4103898903037 | 32.0132594093141 |
| chr1 | 43164311 | 43164327 | - | 9.03564519186028 | 13.6439489047726 | 7.15936926220794 | 8.53686917581709 |
| chr1 | 43164352 | 43164416 | - | 118.467348071057 | 112.562578464374 | 87.1056593568632 | 97.1068868749194 |
| chr1 | 43363083 | 43363140 | + | 82.3247673036159 | 43.2058381984465 | 42.9562155732476 | 64.0265188186281 |
| chr1 | 43363172 | 43363179 | + | 6.02376346124019 | 5.68497871032191 | 5.96614105183995 | 1.06710864697714 |
| chr1 | 43363429 | 43363475 | - | 47.1861471130481 | 37.5208594881246 | 20.2848795762558 | 37.3488026441998 |
| chr1 | 43383936 | 43383979 | - | 53.2099105742883 | 104.603608269923 | 32.2171616799357 | 50.1541064079254 |
| chr1 | 43384077 | 43384084 | - | 8.03168461498692 | 6.82197445238629 | 10.7390538933119 | 8.53686917581709 |
| chr1 | 43384751 | 43384755 | - | 5.01980288436682 | 18.1919318730301 | 8.35259747257593 | 7.46976052883995 |
| chr1 | 43384777 | 43384817 | - | 47.1861471130481 | 81.8636934286355 | 50.1155848354556 | 29.8790421153598 |
| chr1 | 43384838 | 43384840 | - | 6.02376346124019 | 9.09596593651506 | 2.38645642073598 | 4.26843458790854 |
| chr1 | 43384876 | 43384895 | - | 6.02376346124019 | 7.95897019445067 | 2.38645642073598 | 8.53686917581709 |
| chr1 | 43384939 | 43384961 | - | 5.01980288436682 | 9.09596593651506 | 4.77291284147196 | 4.26843458790854 |
| chr1 | 43384993 | 43385017 | - | 6.02376346124019 | 6.82197445238629 | 9.54582568294392 | 8.53686917581709 |
| chr1 | 43385037 | 43385047 | - | 1.00396057687336 | 6.82197445238629 | 5.96614105183995 | 6.40265188186281 |
| chr1 | 43450793 | 43450795 | + | 3.01188173062009 | 2.27399148412876 | 1.19322821036799 | 2.13421729395427 |
| chr1 | 43451198 | 43451240 | - | 172.681219222219 | 175.097344277915 | 4.77291284147196 | 10.6710864697714 |
| chr1 | 43452492 | 43452495 | + | 8.03168461498692 | 4.54798296825753 | 5.96614105183995 | 2.13421729395427 |
| chr1 | 43622035 | 43622051 | + | 3.01188173062009 | 1.13699574206438 | 3.57968463110397 | 1.06710864697714 |
| chr1 | 43622244 | 43622364 | + | 36.1425807674411 | 38.657855230189 | 10.7390538933119 | 6.40265188186281 |
| chr1 | 43622392 | 43622436 | + | 61.2415951892752 | 29.5618892936739 | 10.7390538933119 | 1.06710864697714 |
| chr1 | 43622606 | 43622641 | + | 28.1108961524542 | 12.5069531627082 | 3.57968463110397 | 8.53686917581709 |
| chr1 | 43622699 | 43622724 | + | 11.043566345607 | 21.6029190992233 | 3.57968463110397 | 5.33554323488568 |
| chr1 | 43705120 | 43705127 | + | 1.00396057687336 | 3.41098722619315 | 2.38645642073598 | 3.20132594093141 |
| chr1 | 43705286 | 43705321 | + | 17.0673298068472 | 20.4659233571589 | 4.77291284147196 | 6.40265188186281 |
| chr1 | 43705373 | 43705376 | + | 12.0475269224804 | 21.6029190992233 | 7.15936926220794 | 9.60397782279422 |
| chr1 | 43930565 | 43930598 | + | 11.043566345607 | 31.8358807778027 | 64.4343233598714 | 11.7381951167485 |
| chr1 | 43967684 | 43967688 | + | 8.03168461498692 | 3.41098722619315 | 8.35259747257593 | 13.8724124107028 |
| chr1 | 43973079 | 43973082 | + | 4.01584230749346 | 6.82197445238629 | 3.57968463110397 | 7.46976052883995 |
| chr1 | 43973244 | 43973345 | + | 37.1465413443145 | 40.9318467143178 | 22.6713359969918 | 38.4159112911769 |
| chr1 | 43977948 | 43977957 | + | 2.00792115374673 | 1.13699574206438 | 9.54582568294392 | 3.20132594093141 |
| chr1 | 43977978 | 43978164 | + | 71.2812009580089 | 51.1648083928972 | 84.7192029361272 | 116.314842520508 |
| chr1 | 43978184 | 43978263 | + | 8.03168461498692 | 12.5069531627082 | 4.77291284147196 | 14.9395210576799 |
| chr1 | 43990429 | 43990461 | + | 3.01188173062009 | 2.27399148412876 | 5.96614105183995 | 4.26843458790854 |
| chr1 | 43990524 | 43990564 | + | 8.03168461498692 | 3.41098722619315 | 15.5119667347839 | 17.0737383516342 |
| chr1 | 43990712 | 43990722 | + | 6.02376346124019 | 4.54798296825753 | 11.9322821036799 | 77.8989312293309 |
| chr1 | 43990746 | 43990781 | + | 8.03168461498692 | 3.41098722619315 | 21.4781077866238 | 5.33554323488568 |
| chr1 | 43990808 | 43990861 | + | 49.1940682667949 | 31.8358807778027 | 120.516049247167 | 46.952780466994 |
| chr1 | 43990891 | 43990922 | + | 22.087132691214 | 18.1919318730301 | 100.231169670911 | 171.804492163319 |
| chr1 | 43991083 | 43991116 | + | 5.01980288436682 | 4.54798296825753 | 2.38645642073598 | 14.9395210576799 |
| chr1 | 43991139 | 43991145 | + | 6.02376346124019 | 14.780944646837 | 11.9322821036799 | 13.8724124107028 |
| chr1 | 44048268 | 44048469 | + | 465.837707669241 | 881.171700099896 | 15.5119667347839 | 11.7381951167485 |
| chr1 | 44220565 | 44220645 | + | 261.029749987075 | 260.372024932743 | 46.5359002043516 | 87.5029090521251 |
| chr1 | 44650744 | 44650871 | + | 23.0910932680874 | 20.4659233571589 | 34.6036181006717 | 16.006629704657 |
| chr1 | 44650993 | 44651003 | + | 22.087132691214 | 11.3699574206438 | 38.1833027317757 | 18.1408469986113 |
| chr1 | 44651035 | 44651073 | + | 10.0396057687336 | 9.09596593651506 | 14.3187385244159 | 22.4092815865198 |
| chr1 | 44651132 | 44651169 | + | 2.00792115374673 | 3.41098722619315 | 14.3187385244159 | 11.7381951167485 |
| chr1 | 44651204 | 44651224 | + | 6.02376346124019 | 1.13699574206438 | 5.96614105183995 | 2.13421729395427 |
| chr1 | 44651306 | 44651528 | + | 62.2455557661486 | 71.6307317500561 | 68.0140079909754 | 116.314842520508 |
| chr1 | 44651621 | 44651638 | + | 5.01980288436682 | 4.54798296825753 | 4.77291284147196 | 4.26843458790854 |
| chr1 | 44651663 | 44651665 | + | 1.00396057687336 | 4.54798296825753 | 1.19322821036799 | 2.13421729395427 |
| chr1 | 44767237 | 44767358 | + | 52.205949997415 | 52.3018041349616 | 188.530057238142 | 178.207144045182 |
| chr1 | 44767378 | 44767465 | + | 64.2534769198953 | 46.6168254246397 | 115.743136405695 | 228.361250453107 |
| chr1 | 44767488 | 44767549 | + | 313.23569998449 | 247.865071770035 | 261.31697807059 | 372.42091779502 |
| chr1 | 44767596 | 44767603 | + | 1.00396057687336 | 3.41098722619315 | 11.9322821036799 | 7.46976052883995 |
| chr1 | 44767639 | 44767641 | + | 1.00396057687336 | 1.13699574206438 | 2.38645642073598 | 1.06710864697714 |
| chr1 | 44767677 | 44767682 | + | 9.03564519186028 | 3.41098722619315 | 3.57968463110397 | 12.8053037637256 |
| chr1 | 44775860 | 44775866 | + | 27.1069355755808 | 23.876910583352 | 22.6713359969918 | 9.60397782279422 |
| chr1 | 44778712 | 44778752 | + | 305.204015369503 | 327.454773714542 | 350.809093848189 | 221.958598571244 |
| chr1 | 45002605 | 45002615 | - | 2.00792115374673 | 4.54798296825753 | 2.38645642073598 | 1.06710864697714 |
| chr1 | 45015336 | 45015373 | + | 2.00792115374673 | 1.13699574206438 | 11.9322821036799 | 6.40265188186281 |
| chr1 | 45015405 | 45015557 | + | 9.03564519186028 | 13.6439489047726 | 48.9223566250876 | 76.8318225823538 |
| chr1 | 45303878 | 45303917 | + | 6.02376346124019 | 5.68497871032191 | 13.1255103140479 | 13.8724124107028 |
| chr1 | 45343796 | 45343798 | + | 6.02376346124019 | 3.41098722619315 | 2.38645642073598 | 3.20132594093141 |
| chr1 | 45343873 | 45343874 | + | 1.00396057687336 | 3.41098722619315 | 1.19322821036799 | 1.06710864697714 |
| chr1 | 45509972 | 45509975 | + | 4.01584230749346 | 1.13699574206438 | 7.15936926220794 | 14.9395210576799 |
| chr1 | 45511066 | 45511089 | + | 21.0831721143407 | 12.5069531627082 | 2.38645642073598 | 11.7381951167485 |
| chr1 | 45511086 | 45511187 | - | 621.451597084613 | 389.989539528083 | 226.713359969918 | 454.58828361226 |
| chr1 | 45511215 | 45511227 | - | 7.02772403811355 | 2.27399148412876 | 7.15936926220794 | 3.20132594093141 |
| chr1 | 45511250 | 45511336 | - | 136.538638454778 | 134.165497563597 | 64.4343233598714 | 108.845081991668 |
| chr1 | 45567046 | 45567073 | + | 1.00396057687336 | 2.27399148412876 | 1.19322821036799 | 3.20132594093141 |
| chr1 | 45567440 | 45567444 | + | 1.00396057687336 | 3.41098722619315 | 5.96614105183995 | 2.13421729395427 |
| chr1 | 45569990 | 45570027 | + | 30.1188173062009 | 52.3018041349616 | 41.7629873628796 | 37.3488026441998 |
| chr1 | 45618243 | 45618255 | + | 3.01188173062009 | 2.27399148412876 | 3.57968463110397 | 5.33554323488568 |
| chr1 | 45627400 | 45627429 | - | 14.0554480762271 | 12.5069531627082 | 9.54582568294392 | 9.60397782279422 |
| chr1 | 45627513 | 45627548 | - | 12.0475269224804 | 34.1098722619315 | 38.1833027317757 | 42.6843458790854 |
| chr1 | 45627763 | 45627791 | - | 4.01584230749346 | 5.68497871032191 | 21.4781077866238 | 37.3488026441998 |
| chr1 | 45627824 | 45627870 | - | 2.00792115374673 | 2.27399148412876 | 5.96614105183995 | 4.26843458790854 |
| chr1 | 45628043 | 45628045 | - | 1.00396057687336 | 1.13699574206438 | 2.38645642073598 | 4.26843458790854 |
| chr1 | 45693965 | 45693975 | + | 2.00792115374673 | 1.13699574206438 | 1.19322821036799 | 2.13421729395427 |
| chr1 | 46035991 | 46036026 | + | 7.02772403811355 | 5.68497871032191 | 1.19322821036799 | 2.13421729395427 |
| chr1 | 46036077 | 46036116 | + | 5.01980288436682 | 2.27399148412876 | 1.19322821036799 | 1.06710864697714 |
| chr1 | 46041847 | 46041850 | - | 11.043566345607 | 11.3699574206438 | 7.15936926220794 | 2.13421729395427 |
| chr1 | 46188713 | 46188727 | - | 1.00396057687336 | 1.13699574206438 | 3.57968463110397 | 1.06710864697714 |
| chr1 | 46188911 | 46188915 | - | 5.01980288436682 | 1.13699574206438 | 3.57968463110397 | 1.06710864697714 |
| chr1 | 46189150 | 46189198 | - | 8.03168461498692 | 1.13699574206438 | 9.54582568294392 | 11.7381951167485 |
| chr1 | 46192177 | 46192179 | - | 2.00792115374673 | 2.27399148412876 | 2.38645642073598 | 1.06710864697714 |
| chr1 | 46192372 | 46192397 | - | 3.01188173062009 | 3.41098722619315 | 2.38645642073598 | 1.06710864697714 |
| chr1 | 46278515 | 46278536 | - | 1.00396057687336 | 5.68497871032191 | 7.15936926220794 | 2.13421729395427 |
| chr1 | 46278613 | 46278643 | - | 3.01188173062009 | 1.13699574206438 | 10.7390538933119 | 11.7381951167485 |
| chr1 | 46278720 | 46278817 | - | 8.03168461498692 | 3.41098722619315 | 36.9900745214077 | 25.6106075274513 |
| chr1 | 46278839 | 46278863 | - | 1.00396057687336 | 1.13699574206438 | 2.38645642073598 | 5.33554323488568 |
| chr1 | 46316582 | 46316647 | + | 53.2099105742883 | 76.1787147183136 | 140.800928823423 | 112.046407932599 |
| chr1 | 46316678 | 46316755 | + | 209.827760566533 | 225.125156928748 | 465.359002043516 | 624.258558481624 |
| chr1 | 46353008 | 46353029 | + | 5.01980288436682 | 3.41098722619315 | 15.5119667347839 | 4.26843458790854 |
| chr1 | 46360702 | 46360704 | + | 1.00396057687336 | 1.13699574206438 | 3.57968463110397 | 1.06710864697714 |
| chr1 | 46361587 | 46361603 | + | 1.00396057687336 | 3.41098722619315 | 8.35259747257593 | 2.13421729395427 |
| chr1 | 46361651 | 46361667 | + | 2.00792115374673 | 1.13699574206438 | 5.96614105183995 | 5.33554323488568 |
| chr1 | 46361700 | 46361736 | + | 1.00396057687336 | 5.68497871032191 | 8.35259747257593 | 10.6710864697714 |
| chr1 | 46361768 | 46361814 | + | 8.03168461498692 | 11.3699574206438 | 35.7968463110397 | 19.2079556455884 |
| chr1 | 46361830 | 46361837 | + | 2.00792115374673 | 2.27399148412876 | 4.77291284147196 | 1.06710864697714 |
| chr1 | 46361874 | 46361880 | + | 11.043566345607 | 11.3699574206438 | 8.35259747257593 | 4.26843458790854 |
| chr1 | 46361908 | 46361980 | + | 6.02376346124019 | 13.6439489047726 | 28.6374770488317 | 16.006629704657 |
| chr1 | 46362103 | 46362126 | + | 6.02376346124019 | 5.68497871032191 | 8.35259747257593 | 6.40265188186281 |
| chr1 | 46557603 | 46557651 | - | 238.942617295861 | 189.878288924752 | 193.302970079614 | 325.468137328026 |
| chr1 | 46557718 | 46557721 | - | 2.00792115374673 | 1.13699574206438 | 3.57968463110397 | 1.06710864697714 |
| chr1 | 46557800 | 46557807 | - | 3.01188173062009 | 4.54798296825753 | 11.9322821036799 | 5.33554323488568 |
| chr1 | 46607798 | 46607823 | - | 5.01980288436682 | 2.27399148412876 | 2.38645642073598 | 5.33554323488568 |
| chr1 | 46634951 | 46634954 | - | 4.01584230749346 | 6.82197445238629 | 9.54582568294392 | 6.40265188186281 |
| chr1 | 46635074 | 46635112 | - | 3.01188173062009 | 4.54798296825753 | 3.57968463110397 | 1.06710864697714 |
| chr1 | 46635456 | 46635486 | - | 8.03168461498692 | 5.68497871032191 | 3.57968463110397 | 3.20132594093141 |
| chr1 | 46635513 | 46635569 | - | 14.0554480762271 | 7.95897019445067 | 16.7051949451519 | 7.46976052883995 |
| chr1 | 46635625 | 46635697 | - | 3.01188173062009 | 6.82197445238629 | 11.9322821036799 | 6.40265188186281 |
| chr1 | 46676244 | 46676246 | - | 2.00792115374673 | 1.13699574206438 | 3.57968463110397 | 1.06710864697714 |
| chr1 | 46676470 | 46676473 | - | 8.03168461498692 | 1.13699574206438 | 4.77291284147196 | 1.06710864697714 |
| chr1 | 46676959 | 46676966 | - | 2.00792115374673 | 12.5069531627082 | 4.77291284147196 | 4.26843458790854 |
| chr1 | 46677003 | 46677007 | - | 5.01980288436682 | 4.54798296825753 | 4.77291284147196 | 6.40265188186281 |
| chr1 | 46677696 | 46677716 | - | 11.043566345607 | 1.13699574206438 | 1.19322821036799 | 2.13421729395427 |
| chr1 | 46678165 | 46678171 | - | 17.0673298068472 | 18.1919318730301 | 20.2848795762558 | 5.33554323488568 |
| chr1 | 46678201 | 46678218 | - | 6.02376346124019 | 3.41098722619315 | 9.54582568294392 | 8.53686917581709 |
| chr1 | 46678320 | 46678327 | - | 4.01584230749346 | 2.27399148412876 | 5.96614105183995 | 2.13421729395427 |
| chr1 | 46678431 | 46678452 | - | 12.0475269224804 | 10.2329616785794 | 7.15936926220794 | 8.53686917581709 |
| chr1 | 47179273 | 47179291 | + | 2.00792115374673 | 4.54798296825753 | 1.19322821036799 | 1.06710864697714 |
| chr1 | 47179322 | 47179421 | + | 55.217831728035 | 61.3977700714766 | 52.5020412561915 | 42.6843458790854 |
| chr1 | 47250290 | 47250296 | - | 3.01188173062009 | 2.27399148412876 | 9.54582568294392 | 10.6710864697714 |
| chr1 | 47377362 | 47377392 | + | 6.02376346124019 | 7.95897019445067 | 5.96614105183995 | 19.2079556455884 |
| chr1 | 47377428 | 47377453 | + | 1.00396057687336 | 4.54798296825753 | 13.1255103140479 | 20.2750642925656 |
| chr1 | 47377495 | 47377546 | + | 20.0792115374673 | 17.0549361309657 | 26.2510206280958 | 66.1607361125824 |
| chr1 | 47377567 | 47377590 | + | 1.00396057687336 | 2.27399148412876 | 5.96614105183995 | 4.26843458790854 |
| chr1 | 47377745 | 47377751 | + | 3.01188173062009 | 3.41098722619315 | 2.38645642073598 | 5.33554323488568 |
| chr1 | 47377921 | 47377928 | + | 3.01188173062009 | 3.41098722619315 | 1.19322821036799 | 1.06710864697714 |
| chr1 | 47378044 | 47378069 | + | 4.01584230749346 | 2.27399148412876 | 13.1255103140479 | 13.8724124107028 |
| chr1 | 47378114 | 47378122 | + | 1.00396057687336 | 1.13699574206438 | 2.38645642073598 | 1.06710864697714 |
| chr1 | 47378320 | 47378340 | + | 3.01188173062009 | 12.5069531627082 | 4.77291284147196 | 7.46976052883995 |
| chr1 | 47378391 | 47378449 | + | 3.01188173062009 | 5.68497871032191 | 5.96614105183995 | 5.33554323488568 |
| chr1 | 47378479 | 47378506 | + | 6.02376346124019 | 6.82197445238629 | 10.7390538933119 | 8.53686917581709 |
| chr1 | 47378534 | 47378586 | + | 5.01980288436682 | 7.95897019445067 | 14.3187385244159 | 16.006629704657 |
| chr1 | 47378617 | 47378663 | + | 2.00792115374673 | 7.95897019445067 | 14.3187385244159 | 6.40265188186281 |
| chr1 | 47378766 | 47378782 | + | 9.03564519186028 | 11.3699574206438 | 16.7051949451519 | 6.40265188186281 |
| chr1 | 50423651 | 50423652 | + | 7.02772403811355 | 6.82197445238629 | 1.19322821036799 | 3.20132594093141 |
| chr1 | 50441296 | 50441305 | - | 18.0712903837206 | 22.7399148412876 | 13.1255103140479 | 13.8724124107028 |
| chr1 | 50441370 | 50441385 | - | 14.0554480762271 | 10.2329616785794 | 11.9322821036799 | 18.1408469986113 |
| chr1 | 50974277 | 50974293 | + | 37.1465413443145 | 18.1919318730301 | 3.57968463110397 | 6.40265188186281 |
| chr1 | 50974564 | 50974602 | + | 8.03168461498692 | 21.6029190992233 | 2.38645642073598 | 1.06710864697714 |
| chr1 | 51271358 | 51271386 | + | 2.00792115374673 | 4.54798296825753 | 2.38645642073598 | 1.06710864697714 |
| chr1 | 51271724 | 51271749 | + | 1.00396057687336 | 2.27399148412876 | 2.38645642073598 | 3.20132594093141 |
| chr1 | 51272742 | 51272773 | + | 3.01188173062009 | 2.27399148412876 | 2.38645642073598 | 2.13421729395427 |
| chr1 | 51354299 | 51354312 | - | 1.00396057687336 | 3.41098722619315 | 1.19322821036799 | 4.26843458790854 |
| chr1 | 51354351 | 51354353 | - | 4.01584230749346 | 2.27399148412876 | 1.19322821036799 | 6.40265188186281 |
| chr1 | 51355077 | 51355079 | - | 2.00792115374673 | 3.41098722619315 | 1.19322821036799 | 7.46976052883995 |
| chr1 | 51355282 | 51355325 | - | 2.00792115374673 | 3.41098722619315 | 10.7390538933119 | 5.33554323488568 |
| chr1 | 51355749 | 51355763 | - | 4.01584230749346 | 2.27399148412876 | 9.54582568294392 | 9.60397782279422 |
| chr1 | 51787712 | 51787832 | + | 17.0673298068472 | 6.82197445238629 | 56.0817258872955 | 44.8185631730397 |
| chr1 | 51787861 | 51787870 | + | 12.0475269224804 | 5.68497871032191 | 33.4103898903037 | 29.8790421153598 |
| chr1 | 51787901 | 51787980 | + | 46.1821865361748 | 72.7677274921204 | 201.65556755219 | 158.999188399593 |
| chr1 | 51788001 | 51788035 | + | 15.0594086531005 | 15.9179403889013 | 53.6952694665595 | 33.0803680562912 |
| chr1 | 51788058 | 51788083 | + | 18.0712903837206 | 14.780944646837 | 26.2510206280958 | 28.8119334683827 |
| chr1 | 51788257 | 51788337 | + | 2.00792115374673 | 4.54798296825753 | 15.5119667347839 | 4.26843458790854 |
| chr1 | 51789243 | 51789322 | - | 28.1108961524542 | 25.0139063254164 | 40.5697591525116 | 37.3488026441998 |
| chr1 | 51789347 | 51789436 | - | 169.669337491599 | 143.261463500112 | 375.866886265917 | 331.870789209889 |
| chr1 | 51789573 | 51789655 | - | 212.839642297153 | 221.714169702554 | 460.586089202044 | 534.621432135545 |
| chr1 | 51790532 | 51790644 | - | 200.792115374673 | 216.029190992233 | 461.779317412412 | 316.931268152209 |
| chr1 | 51790907 | 51790921 | - | 2.00792115374673 | 1.13699574206438 | 20.2848795762558 | 5.33554323488568 |
| chr1 | 51790943 | 51790987 | - | 12.0475269224804 | 11.3699574206438 | 62.0478669391355 | 9.60397782279422 |
| chr1 | 51791577 | 51791664 | - | 20.0792115374673 | 13.6439489047726 | 91.8785721983352 | 56.5567582897882 |
| chr1 | 51792054 | 51792062 | - | 1.00396057687336 | 3.41098722619315 | 8.35259747257593 | 7.46976052883995 |
| chr1 | 52020169 | 52020293 | - | 40.1584230749346 | 51.1648083928972 | 36.9900745214077 | 35.2145853502455 |
| chr1 | 52020314 | 52020354 | - | 2.00792115374673 | 2.27399148412876 | 2.38645642073598 | 2.13421729395427 |
| chr1 | 52020732 | 52020810 | - | 3.01188173062009 | 2.27399148412876 | 1.19322821036799 | 2.13421729395427 |
| chr1 | 52020863 | 52020931 | - | 2.00792115374673 | 1.13699574206438 | 1.19322821036799 | 3.20132594093141 |
| chr1 | 52032178 | 52032181 | - | 1.00396057687336 | 3.41098722619315 | 1.19322821036799 | 2.13421729395427 |
| chr1 | 52032440 | 52032448 | - | 2.00792115374673 | 2.27399148412876 | 1.19322821036799 | 1.06710864697714 |
| chr1 | 52033782 | 52033827 | - | 76.3010038423757 | 53.438799877026 | 19.0916513658878 | 52.2883237018796 |
| chr1 | 52086978 | 52087002 | + | 8.03168461498692 | 10.2329616785794 | 9.54582568294392 | 3.20132594093141 |
| chr1 | 52088226 | 52088236 | + | 1.00396057687336 | 1.13699574206438 | 3.57968463110397 | 2.13421729395427 |
| chr1 | 52088387 | 52088396 | + | 6.02376346124019 | 10.2329616785794 | 7.15936926220794 | 7.46976052883995 |
| chr1 | 52346348 | 52346391 | + | 15.0594086531005 | 9.09596593651506 | 2.38645642073598 | 3.20132594093141 |
| chr1 | 52346603 | 52346614 | + | 17.0673298068472 | 14.780944646837 | 11.9322821036799 | 3.20132594093141 |
| chr1 | 52352580 | 52352619 | - | 4.01584230749346 | 17.0549361309657 | 5.96614105183995 | 4.26843458790854 |
| chr1 | 52352655 | 52352658 | - | 4.01584230749346 | 2.27399148412876 | 1.19322821036799 | 4.26843458790854 |
| chr1 | 52372860 | 52372885 | - | 2.00792115374673 | 5.68497871032191 | 11.9322821036799 | 7.46976052883995 |
| chr1 | 52372911 | 52372915 | - | 4.01584230749346 | 2.27399148412876 | 3.57968463110397 | 2.13421729395427 |
| chr1 | 52372965 | 52372972 | - | 3.01188173062009 | 3.41098722619315 | 1.19322821036799 | 6.40265188186281 |
| chr1 | 52416717 | 52416778 | + | 30.1188173062009 | 36.3838637460602 | 25.0577924177278 | 38.4159112911769 |
| chr1 | 52416832 | 52416843 | + | 9.03564519186028 | 28.4248935516096 | 23.8645642073598 | 32.0132594093141 |
| chr1 | 52416903 | 52416980 | + | 11.043566345607 | 30.6988850357383 | 34.6036181006717 | 39.483019938154 |
| chr1 | 52417267 | 52417268 | + | 9.03564519186028 | 10.2329616785794 | 13.1255103140479 | 7.46976052883995 |
| chr1 | 52417398 | 52417411 | + | 8.03168461498692 | 4.54798296825753 | 17.8984231555198 | 12.8053037637256 |
| chr1 | 52417434 | 52417440 | + | 4.01584230749346 | 4.54798296825753 | 13.1255103140479 | 5.33554323488568 |
| chr1 | 52417520 | 52417557 | + | 15.0594086531005 | 6.82197445238629 | 23.8645642073598 | 14.9395210576799 |
| chr1 | 52418062 | 52418105 | + | 6.02376346124019 | 13.6439489047726 | 11.9322821036799 | 10.6710864697714 |
| chr1 | 52423345 | 52423346 | - | 4.01584230749346 | 2.27399148412876 | 1.19322821036799 | 1.06710864697714 |
| chr1 | 52423867 | 52423870 | - | 6.02376346124019 | 6.82197445238629 | 3.57968463110397 | 6.40265188186281 |
| chr1 | 52423903 | 52423997 | - | 46.1821865361748 | 48.8908169087684 | 35.7968463110397 | 53.3554323488568 |
| chr1 | 52684521 | 52684544 | - | 14.0554480762271 | 27.2878978095452 | 3.57968463110397 | 7.46976052883995 |
| chr1 | 52684779 | 52684787 | - | 36.1425807674411 | 37.5208594881246 | 15.5119667347839 | 30.9461507623369 |
| chr1 | 52687515 | 52687517 | - | 2.00792115374673 | 2.27399148412876 | 1.19322821036799 | 1.06710864697714 |
| chr1 | 52821825 | 52821829 | + | 1.00396057687336 | 3.41098722619315 | 5.96614105183995 | 2.13421729395427 |
| chr1 | 52822291 | 52822297 | + | 3.01188173062009 | 5.68497871032191 | 4.77291284147196 | 2.13421729395427 |
| chr1 | 52823271 | 52823282 | + | 4.01584230749346 | 3.41098722619315 | 2.38645642073598 | 5.33554323488568 |
| chr1 | 52823418 | 52823504 | + | 8.03168461498692 | 9.09596593651506 | 20.2848795762558 | 131.254363578188 |
| chr1 | 52825293 | 52825314 | + | 7.02772403811355 | 9.09596593651506 | 5.96614105183995 | 1.06710864697714 |
| chr1 | 52825468 | 52825475 | + | 2.00792115374673 | 3.41098722619315 | 4.77291284147196 | 2.13421729395427 |
| chr1 | 52827185 | 52827187 | + | 1.00396057687336 | 1.13699574206438 | 1.19322821036799 | 2.13421729395427 |
| chr1 | 53047865 | 53047924 | + | 4.01584230749346 | 1.13699574206438 | 8.35259747257593 | 4.26843458790854 |
| chr1 | 53050607 | 53050657 | + | 1.00396057687336 | 1.13699574206438 | 21.4781077866238 | 17.0737383516342 |
| chr1 | 53050686 | 53050902 | + | 61.2415951892752 | 42.0688424563821 | 116.936364616063 | 68.2949534065367 |
| chr1 | 53050928 | 53050999 | + | 56.2217923049084 | 51.1648083928972 | 57.2749540976635 | 33.0803680562912 |
| chr1 | 53051314 | 53051346 | + | 5.01980288436682 | 1.13699574206438 | 1.19322821036799 | 4.26843458790854 |
| chr1 | 53051540 | 53051576 | + | 5.01980288436682 | 2.27399148412876 | 7.15936926220794 | 4.26843458790854 |
| chr1 | 53214126 | 53214180 | + | 5.01980288436682 | 11.3699574206438 | 11.9322821036799 | 7.46976052883995 |
| chr1 | 53214225 | 53214305 | - | 8.03168461498692 | 5.68497871032191 | 14.3187385244159 | 8.53686917581709 |
| chr1 | 53214442 | 53214471 | - | 1.00396057687336 | 1.13699574206438 | 4.77291284147196 | 8.53686917581709 |
| chr1 | 53226953 | 53227026 | - | 127.502993262917 | 170.549361309657 | 194.496198289982 | 237.965228275901 |
| chr1 | 53246303 | 53246308 | - | 3.01188173062009 | 1.13699574206438 | 3.57968463110397 | 5.33554323488568 |
| chr1 | 53246470 | 53246500 | - | 6.02376346124019 | 1.13699574206438 | 4.77291284147196 | 8.53686917581709 |
| chr1 | 53246641 | 53246667 | - | 69.2732798042621 | 31.8358807778027 | 58.4681823080315 | 40.5501285851312 |
| chr1 | 53246699 | 53246714 | - | 43.1703048055547 | 13.6439489047726 | 25.0577924177278 | 38.4159112911769 |
| chr1 | 53246847 | 53246873 | - | 4.01584230749346 | 6.82197445238629 | 11.9322821036799 | 9.60397782279422 |
| chr1 | 53246902 | 53246962 | - | 45.1782259593014 | 25.0139063254164 | 59.6614105183995 | 26.6777161744284 |
| chr1 | 53246991 | 53247033 | - | 15.0594086531005 | 5.68497871032191 | 10.7390538933119 | 12.8053037637256 |
| chr1 | 53765494 | 53765524 | - | 8.03168461498692 | 17.0549361309657 | 8.35259747257593 | 6.40265188186281 |
| chr1 | 53765582 | 53765642 | - | 2.00792115374673 | 7.95897019445067 | 7.15936926220794 | 1.06710864697714 |
| chr1 | 53765675 | 53765678 | - | 2.00792115374673 | 1.13699574206438 | 2.38645642073598 | 3.20132594093141 |
| chr1 | 53765865 | 53765914 | - | 9.03564519186028 | 14.780944646837 | 5.96614105183995 | 26.6777161744284 |
| chr1 | 53765984 | 53765998 | - | 18.0712903837206 | 18.1919318730301 | 7.15936926220794 | 11.7381951167485 |
| chr1 | 53767747 | 53767780 | - | 4.01584230749346 | 6.82197445238629 | 3.57968463110397 | 2.13421729395427 |
| chr1 | 53767813 | 53767824 | - | 2.00792115374673 | 2.27399148412876 | 5.96614105183995 | 6.40265188186281 |
| chr1 | 53851754 | 53851824 | - | 13.0514874993537 | 23.876910583352 | 17.8984231555198 | 17.0737383516342 |
| chr1 | 53851881 | 53851884 | - | 1.00396057687336 | 9.09596593651506 | 9.54582568294392 | 4.26843458790854 |
| chr1 | 53921588 | 53921605 | - | 16.0633692299738 | 23.876910583352 | 14.3187385244159 | 28.8119334683827 |
| chr1 | 53923698 | 53923711 | - | 3.01188173062009 | 2.27399148412876 | 20.2848795762558 | 40.5501285851312 |
| chr1 | 53967980 | 53968005 | + | 6.02376346124019 | 15.9179403889013 | 19.0916513658878 | 13.8724124107028 |
| chr1 | 53968071 | 53968104 | + | 26.1029749987075 | 47.753821166704 | 54.8884976769275 | 38.4159112911769 |
| chr1 | 54031699 | 54031740 | - | 32.1267384599477 | 31.8358807778027 | 15.5119667347839 | 10.6710864697714 |
| chr1 | 54031923 | 54032100 | - | 35.1386201905678 | 40.9318467143178 | 25.0577924177278 | 19.2079556455884 |
| chr1 | 54032125 | 54032234 | - | 33.130699036821 | 20.4659233571589 | 21.4781077866238 | 13.8724124107028 |
| chr1 | 54097367 | 54097377 | + | 2.00792115374673 | 2.27399148412876 | 4.77291284147196 | 1.06710864697714 |
| chr1 | 54218170 | 54218272 | + | 82.3247673036159 | 97.7816338175368 | 96.6514850398071 | 148.328101929822 |
| chr1 | 54218296 | 54218363 | + | 189.748549029066 | 87.5486721389574 | 51.3088130458235 | 193.146665102862 |
| chr1 | 54225429 | 54225444 | + | 1.00396057687336 | 5.68497871032191 | 1.19322821036799 | 5.33554323488568 |
| chr1 | 54225576 | 54225587 | - | 2.00792115374673 | 2.27399148412876 | 2.38645642073598 | 6.40265188186281 |
| chr1 | 54226659 | 54226667 | - | 3.01188173062009 | 3.41098722619315 | 3.57968463110397 | 1.06710864697714 |
| chr1 | 54226690 | 54226769 | - | 795.136776883705 | 1156.32466967948 | 1822.05947723192 | 1385.10702377632 |
| chr1 | 54226787 | 54226924 | - | 46.1821865361748 | 56.8497871032191 | 93.0718004087032 | 43.7514545260626 |
| chr1 | 54226944 | 54226946 | - | 1.00396057687336 | 1.13699574206438 | 7.15936926220794 | 3.20132594093141 |
| chr1 | 54227027 | 54227033 | - | 6.02376346124019 | 4.54798296825753 | 19.0916513658878 | 11.7381951167485 |
| chr1 | 54238378 | 54238379 | + | 40.1584230749346 | 18.1919318730301 | 2.38645642073598 | 9.60397782279422 |
| chr1 | 54722712 | 54722733 | + | 3.01188173062009 | 1.13699574206438 | 13.1255103140479 | 5.33554323488568 |
| chr1 | 54733704 | 54733711 | + | 9.03564519186028 | 3.41098722619315 | 9.54582568294392 | 13.8724124107028 |
| chr1 | 54741411 | 54741412 | + | 3.01188173062009 | 3.41098722619315 | 3.57968463110397 | 5.33554323488568 |
| chr1 | 54741594 | 54741626 | + | 5.01980288436682 | 3.41098722619315 | 7.15936926220794 | 6.40265188186281 |
| chr1 | 54741650 | 54741660 | + | 6.02376346124019 | 5.68497871032191 | 15.5119667347839 | 21.3421729395427 |
| chr1 | 54741690 | 54741712 | + | 3.01188173062009 | 2.27399148412876 | 3.57968463110397 | 1.06710864697714 |
| chr1 | 54742031 | 54742038 | + | 4.01584230749346 | 5.68497871032191 | 3.57968463110397 | 5.33554323488568 |
| chr1 | 54742593 | 54742595 | + | 6.02376346124019 | 10.2329616785794 | 2.38645642073598 | 9.60397782279422 |
| chr1 | 54849654 | 54849767 | - | 29.1148567293276 | 25.0139063254164 | 8.35259747257593 | 16.006629704657 |
| chr1 | 54849791 | 54850008 | - | 28.1108961524542 | 19.3289276150945 | 17.8984231555198 | 29.8790421153598 |
| chr1 | 54850033 | 54850084 | - | 6.02376346124019 | 4.54798296825753 | 4.77291284147196 | 14.9395210576799 |
| chr1 | 54850109 | 54850153 | - | 3.01188173062009 | 2.27399148412876 | 3.57968463110397 | 1.06710864697714 |
| chr1 | 54850456 | 54850565 | - | 7.02772403811355 | 11.3699574206438 | 14.3187385244159 | 16.006629704657 |
| chr1 | 54850688 | 54850809 | - | 2.00792115374673 | 2.27399148412876 | 1.19322821036799 | 2.13421729395427 |
| chr1 | 54850901 | 54850980 | - | 6.02376346124019 | 4.54798296825753 | 4.77291284147196 | 2.13421729395427 |
| chr1 | 54851216 | 54851298 | - | 3.01188173062009 | 5.68497871032191 | 3.57968463110397 | 3.20132594093141 |
| chr1 | 54851385 | 54851418 | - | 1.00396057687336 | 1.13699574206438 | 1.19322821036799 | 1.06710864697714 |
| chr1 | 54851913 | 54852007 | - | 3.01188173062009 | 2.27399148412876 | 1.19322821036799 | 2.13421729395427 |
| chr1 | 54852034 | 54852078 | - | 1.00396057687336 | 2.27399148412876 | 1.19322821036799 | 2.13421729395427 |
| chr1 | 55063599 | 55063633 | + | 8.03168461498692 | 12.5069531627082 | 15.5119667347839 | 5.33554323488568 |
| chr1 | 55063736 | 55063827 | + | 6.02376346124019 | 2.27399148412876 | 1.19322821036799 | 1.06710864697714 |
| chr1 | 55063994 | 55064011 | + | 11.043566345607 | 10.2329616785794 | 5.96614105183995 | 1.06710864697714 |
| chr1 | 55064636 | 55064647 | + | 14.0554480762271 | 9.09596593651506 | 8.35259747257593 | 3.20132594093141 |
| chr1 | 55064683 | 55064752 | + | 8.03168461498692 | 10.2329616785794 | 4.77291284147196 | 5.33554323488568 |
| chr1 | 55066407 | 55066409 | - | 3.01188173062009 | 1.13699574206438 | 1.19322821036799 | 3.20132594093141 |
| chr1 | 55066549 | 55066555 | - | 25.0990144218341 | 10.2329616785794 | 20.2848795762558 | 20.2750642925656 |
| chr1 | 55067169 | 55067186 | - | 2.00792115374673 | 1.13699574206438 | 5.96614105183995 | 7.46976052883995 |
| chr1 | 55067832 | 55067854 | - | 7.02772403811355 | 12.5069531627082 | 20.2848795762558 | 30.9461507623369 |
| chr1 | 55068470 | 55068472 | - | 2.00792115374673 | 1.13699574206438 | 2.38645642073598 | 2.13421729395427 |
| chr1 | 55068630 | 55068633 | - | 9.03564519186028 | 10.2329616785794 | 5.96614105183995 | 8.53686917581709 |
| chr1 | 55068803 | 55068819 | - | 1.00396057687336 | 1.13699574206438 | 2.38645642073598 | 2.13421729395427 |
| chr1 | 55068850 | 55068878 | - | 1.00396057687336 | 3.41098722619315 | 2.38645642073598 | 5.33554323488568 |
| chr1 | 55069021 | 55069026 | - | 2.00792115374673 | 1.13699574206438 | 2.38645642073598 | 2.13421729395427 |
| chr1 | 58480856 | 58480858 | - | 2.00792115374673 | 4.54798296825753 | 2.38645642073598 | 1.06710864697714 |
| chr1 | 58655325 | 58655328 | - | 2.00792115374673 | 3.41098722619315 | 1.19322821036799 | 1.06710864697714 |
| chr1 | 58655441 | 58655457 | - | 1.00396057687336 | 1.13699574206438 | 1.19322821036799 | 2.13421729395427 |
| chr1 | 58655915 | 58655960 | - | 3.01188173062009 | 2.27399148412876 | 16.7051949451519 | 25.6106075274513 |
| chr1 | 58659693 | 58659726 | - | 2.00792115374673 | 1.13699574206438 | 14.3187385244159 | 2.13421729395427 |
| chr1 | 58781416 | 58781471 | - | 1.00396057687336 | 1.13699574206438 | 4.77291284147196 | 1.06710864697714 |
| chr1 | 58781557 | 58781654 | - | 2.00792115374673 | 19.3289276150945 | 13.1255103140479 | 21.3421729395427 |
| chr1 | 58781717 | 58781725 | - | 7.02772403811355 | 14.780944646837 | 19.0916513658878 | 9.60397782279422 |
| chr1 | 61455634 | 61455641 | + | 2.00792115374673 | 4.54798296825753 | 1.19322821036799 | 3.20132594093141 |
| chr1 | 61455753 | 61455764 | + | 8.03168461498692 | 6.82197445238629 | 4.77291284147196 | 1.06710864697714 |
| chr1 | 61455809 | 61455823 | + | 20.0792115374673 | 11.3699574206438 | 7.15936926220794 | 4.26843458790854 |
| chr1 | 61455965 | 61455985 | + | 7.02772403811355 | 12.5069531627082 | 2.38645642073598 | 7.46976052883995 |
| chr1 | 61456097 | 61456115 | + | 5.01980288436682 | 13.6439489047726 | 4.77291284147196 | 5.33554323488568 |
| chr1 | 61456277 | 61456311 | + | 130.514874993537 | 95.5076423334081 | 38.1833027317757 | 43.7514545260626 |
| chr1 | 61456716 | 61456740 | + | 4.01584230749346 | 5.68497871032191 | 2.38645642073598 | 2.13421729395427 |
| chr1 | 61457458 | 61457502 | + | 10.0396057687336 | 9.09596593651506 | 2.38645642073598 | 3.20132594093141 |
| chr1 | 61459898 | 61459908 | + | 19.0752509605939 | 10.2329616785794 | 1.19322821036799 | 8.53686917581709 |
| chr1 | 61681139 | 61681182 | - | 6.02376346124019 | 5.68497871032191 | 7.15936926220794 | 2.13421729395427 |
| chr1 | 62451237 | 62451243 | + | 4.01584230749346 | 2.27399148412876 | 1.19322821036799 | 1.06710864697714 |
| chr1 | 62451432 | 62451479 | + | 18.0712903837206 | 13.6439489047726 | 7.15936926220794 | 3.20132594093141 |
| chr1 | 62451528 | 62451534 | + | 4.01584230749346 | 7.95897019445067 | 1.19322821036799 | 2.13421729395427 |
| chr1 | 62451602 | 62451608 | + | 3.01188173062009 | 17.0549361309657 | 4.77291284147196 | 2.13421729395427 |
| chr1 | 62451667 | 62451679 | + | 40.1584230749346 | 51.1648083928972 | 14.3187385244159 | 23.476390233497 |
| chr1 | 62451706 | 62451711 | + | 37.1465413443145 | 108.014595496116 | 25.0577924177278 | 10.6710864697714 |
| chr1 | 62864893 | 62864895 | + | 2.00792115374673 | 7.95897019445067 | 2.38645642073598 | 7.46976052883995 |
| chr1 | 63437146 | 63437155 | + | 1.00396057687336 | 1.13699574206438 | 2.38645642073598 | 7.46976052883995 |
| chr1 | 63440800 | 63440804 | - | 20.0792115374673 | 53.438799877026 | 29.8307052591997 | 49.0869977609482 |
| chr1 | 63440860 | 63440888 | - | 8.03168461498692 | 5.68497871032191 | 15.5119667347839 | 12.8053037637256 |
| chr1 | 64833261 | 64833288 | - | 1.00396057687336 | 2.27399148412876 | 2.38645642073598 | 2.13421729395427 |
| chr1 | 64833442 | 64833461 | - | 2.00792115374673 | 2.27399148412876 | 1.19322821036799 | 1.06710864697714 |
| chr1 | 64834214 | 64834238 | - | 4.01584230749346 | 1.13699574206438 | 7.15936926220794 | 4.26843458790854 |
| chr1 | 65226789 | 65226806 | + | 3.01188173062009 | 4.54798296825753 | 14.3187385244159 | 24.5434988804741 |
| chr1 | 65228696 | 65228706 | + | 6.02376346124019 | 4.54798296825753 | 29.8307052591997 | 10.6710864697714 |
| chr1 | 65432293 | 65432301 | + | 1.00396057687336 | 1.13699574206438 | 4.77291284147196 | 2.13421729395427 |
| chr1 | 65433091 | 65433109 | + | 3.01188173062009 | 1.13699574206438 | 15.5119667347839 | 3.20132594093141 |
| chr1 | 65433194 | 65433290 | + | 4.01584230749346 | 3.41098722619315 | 10.7390538933119 | 4.26843458790854 |
| chr1 | 65433296 | 65433374 | + | 3.01188173062009 | 2.27399148412876 | 7.15936926220794 | 3.20132594093141 |
| chr1 | 65433431 | 65433453 | + | 11.043566345607 | 13.6439489047726 | 11.9322821036799 | 19.2079556455884 |
| chr1 | 65433538 | 65433579 | + | 1.00396057687336 | 9.09596593651506 | 5.96614105183995 | 12.8053037637256 |
| chr1 | 65433603 | 65433633 | + | 1.00396057687336 | 7.95897019445067 | 4.77291284147196 | 8.53686917581709 |
| chr1 | 65433958 | 65433967 | + | 3.01188173062009 | 3.41098722619315 | 5.96614105183995 | 2.13421729395427 |
| chr1 | 65435960 | 65435968 | + | 2.00792115374673 | 1.13699574206438 | 3.57968463110397 | 5.33554323488568 |
| chr1 | 66985470 | 66985472 | + | 1.00396057687336 | 1.13699574206438 | 1.19322821036799 | 2.13421729395427 |
| chr1 | 66985654 | 66985695 | + | 5.01980288436682 | 10.2329616785794 | 4.77291284147196 | 1.06710864697714 |
| chr1 | 66985781 | 66985805 | + | 5.01980288436682 | 12.5069531627082 | 1.19322821036799 | 32.0132594093141 |
| chr1 | 66986051 | 66986063 | + | 27.1069355755808 | 23.876910583352 | 15.5119667347839 | 8.53686917581709 |
| chr1 | 66986724 | 66986731 | + | 7.02772403811355 | 5.68497871032191 | 5.96614105183995 | 5.33554323488568 |
| chr1 | 66986947 | 66986949 | + | 2.00792115374673 | 4.54798296825753 | 1.19322821036799 | 2.13421729395427 |
| chr1 | 66987351 | 66987352 | + | 1.00396057687336 | 4.54798296825753 | 3.57968463110397 | 2.13421729395427 |
| chr1 | 66987421 | 66987475 | + | 16.0633692299738 | 9.09596593651506 | 10.7390538933119 | 6.40265188186281 |
| chr1 | 66987619 | 66987640 | + | 2.00792115374673 | 6.82197445238629 | 3.57968463110397 | 3.20132594093141 |
| chr1 | 66987807 | 66987855 | + | 26.1029749987075 | 56.8497871032191 | 42.9562155732476 | 27.7448248214055 |
| chr1 | 67000517 | 67000571 | - | 34.1346596136944 | 30.6988850357383 | 4.77291284147196 | 7.46976052883995 |
| chr1 | 67000621 | 67000629 | - | 2.00792115374673 | 4.54798296825753 | 2.38645642073598 | 1.06710864697714 |
| chr1 | 67004162 | 67004181 | - | 5.01980288436682 | 10.2329616785794 | 2.38645642073598 | 2.13421729395427 |
| chr1 | 67004242 | 67004261 | - | 9.03564519186028 | 6.82197445238629 | 14.3187385244159 | 2.13421729395427 |
| chr1 | 67004333 | 67004357 | - | 11.043566345607 | 7.95897019445067 | 2.38645642073598 | 3.20132594093141 |
| chr1 | 67408353 | 67408363 | - | 7.02772403811355 | 6.82197445238629 | 5.96614105183995 | 3.20132594093141 |
| chr1 | 67409458 | 67409460 | - | 5.01980288436682 | 2.27399148412876 | 5.96614105183995 | 2.13421729395427 |
| chr1 | 67410516 | 67410530 | - | 1.00396057687336 | 1.13699574206438 | 1.19322821036799 | 5.33554323488568 |
| chr1 | 67411021 | 67411070 | - | 3.01188173062009 | 9.09596593651506 | 5.96614105183995 | 2.13421729395427 |
| chr1 | 67411096 | 67411132 | - | 9.03564519186028 | 14.780944646837 | 20.2848795762558 | 10.6710864697714 |
| chr1 | 67411180 | 67411201 | - | 9.03564519186028 | 6.82197445238629 | 7.15936926220794 | 2.13421729395427 |
| chr1 | 67411342 | 67411370 | - | 11.043566345607 | 6.82197445238629 | 10.7390538933119 | 1.06710864697714 |
| chr1 | 67411409 | 67411467 | - | 12.0475269224804 | 10.2329616785794 | 7.15936926220794 | 10.6710864697714 |
| chr1 | 67411632 | 67411688 | - | 19.0752509605939 | 15.9179403889013 | 8.35259747257593 | 7.46976052883995 |
| chr1 | 67411779 | 67411836 | - | 33.130699036821 | 30.6988850357383 | 13.1255103140479 | 36.2816939972226 |
| chr1 | 67411874 | 67411889 | - | 14.0554480762271 | 7.95897019445067 | 2.38645642073598 | 12.8053037637256 |
| chr1 | 67411910 | 67411916 | - | 20.0792115374673 | 15.9179403889013 | 16.7051949451519 | 24.5434988804741 |
| chr1 | 67411966 | 67411978 | - | 10.0396057687336 | 13.6439489047726 | 9.54582568294392 | 12.8053037637256 |
| chr1 | 67412001 | 67412021 | - | 9.03564519186028 | 4.54798296825753 | 3.57968463110397 | 6.40265188186281 |
| chr1 | 67412073 | 67412094 | - | 10.0396057687336 | 5.68497871032191 | 14.3187385244159 | 8.53686917581709 |
| chr1 | 67412156 | 67412163 | - | 26.1029749987075 | 28.4248935516096 | 27.4442488384638 | 26.6777161744284 |
| chr1 | 67412205 | 67412257 | - | 13.0514874993537 | 9.09596593651506 | 17.8984231555198 | 16.006629704657 |
| chr1 | 67412350 | 67412398 | - | 19.0752509605939 | 25.0139063254164 | 22.6713359969918 | 25.6106075274513 |
| chr1 | 67412523 | 67412529 | - | 5.01980288436682 | 1.13699574206438 | 3.57968463110397 | 6.40265188186281 |
| chr1 | 67413126 | 67413149 | - | 11.043566345607 | 6.82197445238629 | 9.54582568294392 | 12.8053037637256 |
| chr1 | 67413184 | 67413194 | - | 3.01188173062009 | 3.41098722619315 | 1.19322821036799 | 7.46976052883995 |
| chr1 | 67687876 | 67687895 | + | 6.02376346124019 | 6.82197445238629 | 106.197310722751 | 74.6976052883995 |
| chr1 | 67687921 | 67688047 | + | 8.03168461498692 | 11.3699574206438 | 100.231169670911 | 128.053037637256 |
| chr1 | 67688156 | 67688168 | + | 1.00396057687336 | 10.2329616785794 | 14.3187385244159 | 10.6710864697714 |
| chr1 | 67688261 | 67688316 | + | 1.00396057687336 | 6.82197445238629 | 11.9322821036799 | 9.60397782279422 |
| chr1 | 67701967 | 67701997 | - | 19.0752509605939 | 27.2878978095452 | 5.96614105183995 | 1.06710864697714 |
| chr1 | 67702042 | 67702060 | - | 7.02772403811355 | 11.3699574206438 | 1.19322821036799 | 2.13421729395427 |
| chr1 | 67702630 | 67702671 | - | 33.130699036821 | 42.0688424563821 | 7.15936926220794 | 11.7381951167485 |
| chr1 | 67704084 | 67704101 | - | 3.01188173062009 | 5.68497871032191 | 1.19322821036799 | 1.06710864697714 |
| chr1 | 68474602 | 68474614 | - | 6.02376346124019 | 6.82197445238629 | 1.19322821036799 | 2.13421729395427 |
| chr1 | 68474668 | 68474690 | - | 11.043566345607 | 6.82197445238629 | 1.19322821036799 | 3.20132594093141 |
| chr1 | 68475608 | 68475630 | - | 3.01188173062009 | 9.09596593651506 | 2.38645642073598 | 1.06710864697714 |
| chr1 | 68496669 | 68496700 | + | 54.2138711511617 | 62.534765813541 | 41.7629873628796 | 32.0132594093141 |
| chr1 | 68538458 | 68538460 | + | 8.03168461498692 | 14.780944646837 | 2.38645642073598 | 1.06710864697714 |
| chr1 | 70144910 | 70144911 | - | 2.00792115374673 | 1.13699574206438 | 1.19322821036799 | 1.06710864697714 |
| chr1 | 70250795 | 70250862 | + | 25.0990144218341 | 31.8358807778027 | 17.8984231555198 | 17.0737383516342 |
| chr1 | 70250980 | 70251019 | + | 3.01188173062009 | 3.41098722619315 | 8.35259747257593 | 4.26843458790854 |
| chr1 | 70251356 | 70251430 | + | 11.043566345607 | 17.0549361309657 | 8.35259747257593 | 10.6710864697714 |
| chr1 | 70251472 | 70251524 | + | 6.02376346124019 | 14.780944646837 | 3.57968463110397 | 4.26843458790854 |
| chr1 | 70251572 | 70251621 | + | 17.0673298068472 | 19.3289276150945 | 11.9322821036799 | 7.46976052883995 |
| chr1 | 70251679 | 70251721 | + | 20.0792115374673 | 31.8358807778027 | 20.2848795762558 | 45.8856718200168 |
| chr1 | 70251812 | 70251833 | + | 3.01188173062009 | 5.68497871032191 | 3.57968463110397 | 2.13421729395427 |
| chr1 | 70251929 | 70251993 | + | 7.02772403811355 | 12.5069531627082 | 13.1255103140479 | 5.33554323488568 |
| chr1 | 70439224 | 70439226 | + | 2.00792115374673 | 25.0139063254164 | 4.77291284147196 | 7.46976052883995 |
| chr1 | 70439374 | 70439400 | + | 18.0712903837206 | 72.7677274921204 | 19.0916513658878 | 18.1408469986113 |
| chr1 | 71063368 | 71063389 | - | 1.00396057687336 | 4.54798296825753 | 9.54582568294392 | 5.33554323488568 |
| chr1 | 71063435 | 71063478 | - | 1.00396057687336 | 3.41098722619315 | 4.77291284147196 | 6.40265188186281 |
| chr1 | 71063528 | 71063596 | - | 4.01584230749346 | 5.68497871032191 | 4.77291284147196 | 9.60397782279422 |
| chr1 | 71063678 | 71063682 | - | 5.01980288436682 | 1.13699574206438 | 2.38645642073598 | 5.33554323488568 |
| chr1 | 71063873 | 71064013 | - | 401.584230749346 | 452.524305341624 | 726.675980114106 | 618.923015246739 |
| chr1 | 71064038 | 71064057 | - | 4.01584230749346 | 10.2329616785794 | 20.2848795762558 | 5.33554323488568 |
| chr1 | 71064087 | 71064119 | - | 9.03564519186028 | 7.95897019445067 | 23.8645642073598 | 16.006629704657 |
| chr1 | 71064153 | 71064216 | - | 9.03564519186028 | 13.6439489047726 | 27.4442488384638 | 19.2079556455884 |
| chr1 | 71064244 | 71064287 | - | 7.02772403811355 | 13.6439489047726 | 35.7968463110397 | 28.8119334683827 |
| chr1 | 71064373 | 71064428 | - | 18.0712903837206 | 19.3289276150945 | 26.2510206280958 | 19.2079556455884 |
| chr1 | 71064450 | 71064476 | - | 25.0990144218341 | 38.657855230189 | 62.0478669391355 | 51.2212150549025 |
| chr1 | 71064529 | 71064555 | - | 16.0633692299738 | 14.780944646837 | 56.0817258872955 | 22.4092815865198 |
| chr1 | 71064580 | 71064603 | - | 3.01188173062009 | 5.68497871032191 | 19.0916513658878 | 4.26843458790854 |
| chr1 | 71064627 | 71064717 | - | 11.043566345607 | 14.780944646837 | 63.2410951495034 | 43.7514545260626 |
| chr1 | 71064727 | 71064752 | - | 5.01980288436682 | 3.41098722619315 | 4.77291284147196 | 1.06710864697714 |
| chr1 | 71064968 | 71064975 | - | 5.01980288436682 | 22.7399148412876 | 21.4781077866238 | 17.0737383516342 |
| chr1 | 71065008 | 71065043 | - | 9.03564519186028 | 4.54798296825753 | 27.4442488384638 | 10.6710864697714 |
| chr1 | 71065080 | 71065136 | - | 5.01980288436682 | 11.3699574206438 | 19.0916513658878 | 2.13421729395427 |
| chr1 | 71065706 | 71065713 | - | 6.02376346124019 | 4.54798296825753 | 5.96614105183995 | 2.13421729395427 |
| chr1 | 71081347 | 71081372 | + | 14.0554480762271 | 21.6029190992233 | 46.5359002043516 | 27.7448248214055 |
| chr1 | 74705589 | 74705598 | - | 25.0990144218341 | 25.0139063254164 | 56.0817258872955 | 33.0803680562912 |
| chr1 | 74705619 | 74705667 | - | 1.00396057687336 | 1.13699574206438 | 3.57968463110397 | 3.20132594093141 |
| chr1 | 74705695 | 74705742 | - | 4.01584230749346 | 1.13699574206438 | 8.35259747257593 | 2.13421729395427 |
| chr1 | 74705786 | 74705830 | - | 6.02376346124019 | 5.68497871032191 | 7.15936926220794 | 7.46976052883995 |
| chr1 | 74705884 | 74705950 | - | 2.00792115374673 | 1.13699574206438 | 1.19322821036799 | 3.20132594093141 |
| chr1 | 74706120 | 74706151 | - | 1.00396057687336 | 2.27399148412876 | 1.19322821036799 | 3.20132594093141 |
| chr1 | 74764129 | 74764174 | + | 6.02376346124019 | 4.54798296825753 | 3.57968463110397 | 3.20132594093141 |
| chr1 | 74765418 | 74765421 | + | 1.00396057687336 | 1.13699574206438 | 5.96614105183995 | 1.06710864697714 |
| chr1 | 74766397 | 74766401 | + | 7.02772403811355 | 9.09596593651506 | 3.57968463110397 | 4.26843458790854 |
| chr1 | 75762826 | 75762833 | + | 9.03564519186028 | 6.82197445238629 | 1.19322821036799 | 3.20132594093141 |
| chr1 | 75763088 | 75763091 | + | 4.01584230749346 | 3.41098722619315 | 1.19322821036799 | 2.13421729395427 |
| chr1 | 75763206 | 75763220 | + | 1.00396057687336 | 1.13699574206438 | 1.19322821036799 | 1.06710864697714 |
| chr1 | 75763396 | 75763421 | + | 9.03564519186028 | 4.54798296825753 | 4.77291284147196 | 2.13421729395427 |
| chr1 | 75787069 | 75787072 | + | 4.01584230749346 | 6.82197445238629 | 10.7390538933119 | 23.476390233497 |
| chr1 | 75787874 | 75787896 | + | 10.0396057687336 | 2.27399148412876 | 29.8307052591997 | 20.2750642925656 |
| chr1 | 75789542 | 75789554 | + | 8.03168461498692 | 6.82197445238629 | 7.15936926220794 | 7.46976052883995 |
| chr1 | 75790110 | 75790124 | + | 22.087132691214 | 18.1919318730301 | 50.1155848354556 | 23.476390233497 |
| chr1 | 75794602 | 75794771 | + | 29.1148567293276 | 38.657855230189 | 47.7291284147196 | 35.2145853502455 |
| chr1 | 77089047 | 77089050 | - | 2.00792115374673 | 4.54798296825753 | 2.38645642073598 | 1.06710864697714 |
| chr1 | 77089156 | 77089158 | - | 3.01188173062009 | 2.27399148412876 | 1.19322821036799 | 1.06710864697714 |
| chr1 | 77091143 | 77091161 | - | 10.0396057687336 | 12.5069531627082 | 3.57968463110397 | 14.9395210576799 |
| chr1 | 77091799 | 77091828 | - | 32.1267384599477 | 36.3838637460602 | 21.4781077866238 | 25.6106075274513 |
| chr1 | 77091865 | 77091874 | - | 6.02376346124019 | 13.6439489047726 | 10.7390538933119 | 2.13421729395427 |
| chr1 | 77091944 | 77092003 | - | 23.0910932680874 | 32.9728765198671 | 25.0577924177278 | 16.006629704657 |
| chr1 | 77092023 | 77092070 | - | 6.02376346124019 | 14.780944646837 | 2.38645642073598 | 4.26843458790854 |
| chr1 | 77092123 | 77092230 | - | 31.1227778830743 | 21.6029190992233 | 29.8307052591997 | 21.3421729395427 |
| chr1 | 77092332 | 77092480 | - | 88.3485307648561 | 80.7266976865711 | 54.8884976769275 | 53.3554323488568 |
| chr1 | 77564664 | 77564699 | - | 16.0633692299738 | 27.2878978095452 | 27.4442488384638 | 25.6106075274513 |
| chr1 | 77564799 | 77564844 | - | 10.0396057687336 | 19.3289276150945 | 14.3187385244159 | 9.60397782279422 |
| chr1 | 77564886 | 77564956 | - | 30.1188173062009 | 37.5208594881246 | 26.2510206280958 | 9.60397782279422 |
| chr1 | 77564990 | 77565037 | - | 84.3326884573626 | 139.850476273919 | 51.3088130458235 | 65.0936274656053 |
| chr1 | 77565112 | 77565164 | - | 12.0475269224804 | 9.09596593651506 | 3.57968463110397 | 8.53686917581709 |
| chr1 | 77696026 | 77696072 | - | 1.00396057687336 | 5.68497871032191 | 7.15936926220794 | 4.26843458790854 |
| chr1 | 77696233 | 77696240 | - | 1.00396057687336 | 1.13699574206438 | 1.19322821036799 | 3.20132594093141 |
| chr1 | 77696492 | 77696516 | - | 3.01188173062009 | 7.95897019445067 | 2.38645642073598 | 2.13421729395427 |
| chr1 | 77877627 | 77877650 | + | 65.2574374967687 | 67.0827487817985 | 7.15936926220794 | 20.2750642925656 |
| chr1 | 77877792 | 77877809 | + | 10.0396057687336 | 21.6029190992233 | 4.77291284147196 | 6.40265188186281 |
| chr1 | 77948489 | 77948514 | - | 19.0752509605939 | 9.09596593651506 | 10.7390538933119 | 4.26843458790854 |
| chr1 | 77948533 | 77948542 | - | 10.0396057687336 | 6.82197445238629 | 7.15936926220794 | 5.33554323488568 |
| chr1 | 77948572 | 77948601 | - | 54.2138711511617 | 72.7677274921204 | 31.0239334695677 | 21.3421729395427 |
| chr1 | 77948654 | 77948779 | - | 63.249516343022 | 44.3428339405109 | 50.1155848354556 | 38.4159112911769 |
| chr1 | 78016517 | 78016547 | + | 7.02772403811355 | 6.82197445238629 | 9.54582568294392 | 8.53686917581709 |
| chr1 | 78016662 | 78016682 | + | 4.01584230749346 | 7.95897019445067 | 2.38645642073598 | 1.06710864697714 |
| chr1 | 78016722 | 78016735 | + | 38.1505019211879 | 18.1919318730301 | 13.1255103140479 | 23.476390233497 |
| chr1 | 78016788 | 78016899 | + | 40.1584230749346 | 37.5208594881246 | 19.0916513658878 | 29.8790421153598 |
| chr1 | 78016943 | 78016951 | + | 6.02376346124019 | 6.82197445238629 | 7.15936926220794 | 2.13421729395427 |
| chr1 | 78016981 | 78016998 | + | 11.043566345607 | 9.09596593651506 | 7.15936926220794 | 4.26843458790854 |
| chr1 | 78017040 | 78017048 | + | 20.0792115374673 | 21.6029190992233 | 16.7051949451519 | 10.6710864697714 |
| chr1 | 78017136 | 78017179 | + | 11.043566345607 | 9.09596593651506 | 5.96614105183995 | 4.26843458790854 |
| chr1 | 78017345 | 78017364 | + | 3.01188173062009 | 3.41098722619315 | 2.38645642073598 | 3.20132594093141 |
| chr1 | 78017731 | 78017754 | + | 5.01980288436682 | 2.27399148412876 | 5.96614105183995 | 5.33554323488568 |
| chr1 | 78017861 | 78017874 | + | 7.02772403811355 | 9.09596593651506 | 4.77291284147196 | 3.20132594093141 |
| chr1 | 78017901 | 78017920 | + | 28.1108961524542 | 42.0688424563821 | 47.7291284147196 | 13.8724124107028 |
| chr1 | 84497500 | 84497509 | + | 1.00396057687336 | 3.41098722619315 | 2.38645642073598 | 6.40265188186281 |
| chr1 | 84497589 | 84497628 | + | 13.0514874993537 | 12.5069531627082 | 13.1255103140479 | 13.8724124107028 |
| chr1 | 84498343 | 84498361 | - | 38.1505019211879 | 45.4798296825753 | 199.269111131454 | 642.399405480236 |
| chr1 | 84498425 | 84498482 | - | 6.02376346124019 | 17.0549361309657 | 47.7291284147196 | 11.7381951167485 |
| chr1 | 84644644 | 84644680 | - | 6.02376346124019 | 10.2329616785794 | 29.8307052591997 | 22.4092815865198 |
| chr1 | 84644704 | 84644742 | - | 5.01980288436682 | 15.9179403889013 | 28.6374770488317 | 23.476390233497 |
| chr1 | 84644834 | 84644868 | - | 3.01188173062009 | 1.13699574206438 | 13.1255103140479 | 6.40265188186281 |
| chr1 | 84645169 | 84645172 | - | 1.00396057687336 | 3.41098722619315 | 2.38645642073598 | 1.06710864697714 |
| chr1 | 84646616 | 84646621 | - | 1.00396057687336 | 4.54798296825753 | 3.57968463110397 | 13.8724124107028 |
| chr1 | 84646998 | 84646999 | - | 1.00396057687336 | 1.13699574206438 | 1.19322821036799 | 2.13421729395427 |
| chr1 | 84647072 | 84647079 | - | 1.00396057687336 | 4.54798296825753 | 1.19322821036799 | 3.20132594093141 |
| chr1 | 85252245 | 85252271 | - | 4.01584230749346 | 6.82197445238629 | 4.77291284147196 | 7.46976052883995 |
| chr1 | 85252552 | 85252573 | - | 3.01188173062009 | 2.27399148412876 | 1.19322821036799 | 2.13421729395427 |
| chr1 | 85267729 | 85267734 | - | 6.02376346124019 | 10.2329616785794 | 3.57968463110397 | 4.26843458790854 |
| chr1 | 85583594 | 85583619 | + | 59.2336740355285 | 442.291343663045 | 4.77291284147196 | 1.06710864697714 |
| chr1 | 85652830 | 85652831 | - | 1.00396057687336 | 2.27399148412876 | 1.19322821036799 | 1.06710864697714 |
| chr1 | 85653058 | 85653070 | - | 3.01188173062009 | 11.3699574206438 | 3.57968463110397 | 5.33554323488568 |
| chr1 | 85653963 | 85653965 | - | 1.00396057687336 | 2.27399148412876 | 2.38645642073598 | 1.06710864697714 |
| chr1 | 86743239 | 86743255 | + | 5.01980288436682 | 7.95897019445067 | 4.77291284147196 | 8.53686917581709 |
| chr1 | 86743494 | 86743514 | + | 8.03168461498692 | 5.68497871032191 | 7.15936926220794 | 4.26843458790854 |
| chr1 | 86746929 | 86746952 | + | 3.01188173062009 | 1.13699574206438 | 1.19322821036799 | 1.06710864697714 |
| chr1 | 86862571 | 86862585 | - | 3.01188173062009 | 4.54798296825753 | 11.9322821036799 | 4.26843458790854 |
| chr1 | 86862616 | 86862618 | - | 3.01188173062009 | 1.13699574206438 | 2.38645642073598 | 7.46976052883995 |
| chr1 | 86862748 | 86862787 | - | 30.1188173062009 | 53.438799877026 | 58.4681823080315 | 44.8185631730397 |
| chr1 | 86862813 | 86862842 | - | 114.451505763564 | 180.782322988237 | 140.800928823423 | 139.791232754005 |
| chr1 | 86862864 | 86862874 | - | 2.00792115374673 | 2.27399148412876 | 2.38645642073598 | 1.06710864697714 |
| chr1 | 86862894 | 86862914 | - | 1.00396057687336 | 2.27399148412876 | 1.19322821036799 | 5.33554323488568 |
| chr1 | 86863023 | 86863234 | - | 25.0990144218341 | 61.3977700714766 | 46.5359002043516 | 53.3554323488568 |
| chr1 | 86863257 | 86863380 | - | 17.0673298068472 | 50.0278126508328 | 16.7051949451519 | 30.9461507623369 |
| chr1 | 86863456 | 86863570 | - | 10.0396057687336 | 11.3699574206438 | 21.4781077866238 | 11.7381951167485 |
| chr1 | 87105002 | 87105018 | + | 1.00396057687336 | 1.13699574206438 | 4.77291284147196 | 1.06710864697714 |
| chr1 | 87105057 | 87105193 | + | 38.1505019211879 | 23.876910583352 | 39.3765309421437 | 32.0132594093141 |
| chr1 | 87105375 | 87105417 | + | 4.01584230749346 | 2.27399148412876 | 5.96614105183995 | 3.20132594093141 |
| chr1 | 87105500 | 87105554 | + | 14.0554480762271 | 12.5069531627082 | 26.2510206280958 | 13.8724124107028 |
| chr1 | 87105850 | 87105917 | + | 16.0633692299738 | 14.780944646837 | 25.0577924177278 | 32.0132594093141 |
| chr1 | 87105944 | 87105954 | + | 12.0475269224804 | 7.95897019445067 | 19.0916513658878 | 20.2750642925656 |
| chr1 | 87106138 | 87106212 | + | 1.00396057687336 | 6.82197445238629 | 8.35259747257593 | 9.60397782279422 |
| chr1 | 87106337 | 87106365 | + | 4.01584230749346 | 9.09596593651506 | 7.15936926220794 | 7.46976052883995 |
| chr1 | 87106493 | 87106536 | + | 6.02376346124019 | 14.780944646837 | 22.6713359969918 | 23.476390233497 |
| chr1 | 87107346 | 87107373 | + | 3.01188173062009 | 7.95897019445067 | 8.35259747257593 | 2.13421729395427 |
| chr1 | 87107398 | 87107441 | + | 11.043566345607 | 12.5069531627082 | 15.5119667347839 | 20.2750642925656 |
| chr1 | 87107849 | 87107868 | + | 4.01584230749346 | 2.27399148412876 | 1.19322821036799 | 2.13421729395427 |
| chr1 | 87107910 | 87107932 | + | 1.00396057687336 | 2.27399148412876 | 3.57968463110397 | 3.20132594093141 |
| chr1 | 87109070 | 87109079 | + | 1.00396057687336 | 2.27399148412876 | 5.96614105183995 | 5.33554323488568 |
| chr1 | 87109168 | 87109191 | + | 20.0792115374673 | 12.5069531627082 | 32.2171616799357 | 26.6777161744284 |
| chr1 | 87109861 | 87109899 | + | 1.00396057687336 | 3.41098722619315 | 3.57968463110397 | 3.20132594093141 |
| chr1 | 87345049 | 87345091 | + | 30.1188173062009 | 131.891506079468 | 44.1494437836156 | 51.2212150549025 |
| chr1 | 87345380 | 87345402 | + | 2.00792115374673 | 14.780944646837 | 5.96614105183995 | 6.40265188186281 |
| chr1 | 88805910 | 88805911 | + | 4.01584230749346 | 7.95897019445067 | 3.57968463110397 | 1.06710864697714 |
| chr1 | 88833471 | 88833487 | + | 4.01584230749346 | 2.27399148412876 | 3.57968463110397 | 1.06710864697714 |
| chr1 | 88833520 | 88833535 | + | 5.01980288436682 | 2.27399148412876 | 8.35259747257593 | 1.06710864697714 |
| chr1 | 88833783 | 88833807 | + | 1.00396057687336 | 2.27399148412876 | 3.57968463110397 | 1.06710864697714 |
| chr1 | 88833950 | 88833977 | + | 46.1821865361748 | 80.7266976865711 | 21.4781077866238 | 17.0737383516342 |
| chr1 | 88834000 | 88834055 | + | 13.0514874993537 | 25.0139063254164 | 19.0916513658878 | 2.13421729395427 |
| chr1 | 88834080 | 88834105 | + | 14.0554480762271 | 18.1919318730301 | 9.54582568294392 | 9.60397782279422 |
| chr1 | 88834137 | 88834187 | + | 19.0752509605939 | 15.9179403889013 | 11.9322821036799 | 8.53686917581709 |
| chr1 | 88834221 | 88834254 | + | 8.03168461498692 | 10.2329616785794 | 3.57968463110397 | 1.06710864697714 |
| chr1 | 88834735 | 88834825 | + | 15.0594086531005 | 10.2329616785794 | 17.8984231555198 | 8.53686917581709 |
| chr1 | 88835351 | 88835353 | + | 1.00396057687336 | 2.27399148412876 | 2.38645642073598 | 1.06710864697714 |
| chr1 | 88852983 | 88852999 | - | 4.01584230749346 | 7.95897019445067 | 13.1255103140479 | 9.60397782279422 |
| chr1 | 88853090 | 88853113 | - | 9.03564519186028 | 14.780944646837 | 11.9322821036799 | 38.4159112911769 |
| chr1 | 88853140 | 88853143 | - | 2.00792115374673 | 6.82197445238629 | 7.15936926220794 | 4.26843458790854 |
| chr1 | 88936170 | 88936225 | - | 10.0396057687336 | 15.9179403889013 | 25.0577924177278 | 45.8856718200168 |
| chr1 | 88980377 | 88980382 | - | 6.02376346124019 | 9.09596593651506 | 11.9322821036799 | 8.53686917581709 |
| chr1 | 88981007 | 88981012 | - | 3.01188173062009 | 5.68497871032191 | 5.96614105183995 | 5.33554323488568 |
| chr1 | 89596416 | 89596419 | + | 1.00396057687336 | 2.27399148412876 | 3.57968463110397 | 3.20132594093141 |
| chr1 | 89719160 | 89719163 | + | 4.01584230749346 | 2.27399148412876 | 2.38645642073598 | 4.26843458790854 |
| chr1 | 89843629 | 89843684 | + | 87.3445701879827 | 43.2058381984465 | 5.96614105183995 | 3.20132594093141 |
| chr1 | 89933139 | 89933162 | + | 3.01188173062009 | 3.41098722619315 | 7.15936926220794 | 3.20132594093141 |
| chr1 | 89933223 | 89933224 | + | 3.01188173062009 | 4.54798296825753 | 5.96614105183995 | 1.06710864697714 |
| chr1 | 89935670 | 89935677 | + | 2.00792115374673 | 1.13699574206438 | 4.77291284147196 | 3.20132594093141 |
| chr1 | 89935763 | 89935784 | + | 5.01980288436682 | 3.41098722619315 | 3.57968463110397 | 10.6710864697714 |
| chr1 | 89935864 | 89935876 | + | 5.01980288436682 | 18.1919318730301 | 9.54582568294392 | 11.7381951167485 |
| chr1 | 90005979 | 90005982 | + | 3.01188173062009 | 2.27399148412876 | 2.38645642073598 | 1.06710864697714 |
| chr1 | 90027739 | 90027758 | + | 12.0475269224804 | 9.09596593651506 | 9.54582568294392 | 11.7381951167485 |
| chr1 | 90028037 | 90028039 | + | 2.00792115374673 | 3.41098722619315 | 2.38645642073598 | 4.26843458790854 |
| chr1 | 90028081 | 90028111 | + | 1.00396057687336 | 5.68497871032191 | 1.19322821036799 | 4.26843458790854 |
| chr1 | 90031363 | 90031389 | + | 1.00396057687336 | 1.13699574206438 | 5.96614105183995 | 3.20132594093141 |
| chr1 | 90031538 | 90031542 | + | 4.01584230749346 | 5.68497871032191 | 1.19322821036799 | 3.20132594093141 |
| chr1 | 90032190 | 90032191 | + | 2.00792115374673 | 1.13699574206438 | 1.19322821036799 | 1.06710864697714 |
| chr1 | 90032297 | 90032304 | + | 8.03168461498692 | 9.09596593651506 | 7.15936926220794 | 5.33554323488568 |
| chr1 | 90033385 | 90033400 | + | 6.02376346124019 | 1.13699574206438 | 3.57968463110397 | 6.40265188186281 |
| chr1 | 90915621 | 90915648 | - | 11.043566345607 | 13.6439489047726 | 10.7390538933119 | 8.53686917581709 |
| chr1 | 90916685 | 90916688 | - | 4.01584230749346 | 11.3699574206438 | 3.57968463110397 | 9.60397782279422 |
| chr1 | 90916762 | 90916813 | - | 3.01188173062009 | 7.95897019445067 | 5.96614105183995 | 7.46976052883995 |
| chr1 | 91524466 | 91524470 | + | 5.01980288436682 | 5.68497871032191 | 2.38645642073598 | 1.06710864697714 |
| chr1 | 91524601 | 91524609 | + | 5.01980288436682 | 2.27399148412876 | 3.57968463110397 | 3.20132594093141 |
| chr1 | 91524733 | 91524742 | + | 1.00396057687336 | 1.13699574206438 | 2.38645642073598 | 3.20132594093141 |
| chr1 | 91524766 | 91524774 | + | 4.01584230749346 | 1.13699574206438 | 10.7390538933119 | 9.60397782279422 |
| chr1 | 91524844 | 91524865 | + | 3.01188173062009 | 3.41098722619315 | 4.77291284147196 | 6.40265188186281 |
| chr1 | 91524950 | 91524969 | + | 7.02772403811355 | 18.1919318730301 | 19.0916513658878 | 17.0737383516342 |
| chr1 | 91525026 | 91525035 | + | 5.01980288436682 | 6.82197445238629 | 8.35259747257593 | 7.46976052883995 |
| chr1 | 91525105 | 91525139 | + | 4.01584230749346 | 7.95897019445067 | 11.9322821036799 | 18.1408469986113 |
| chr1 | 91525194 | 91525261 | + | 33.130699036821 | 57.9867828452835 | 36.9900745214077 | 55.4896496428111 |
| chr1 | 91525515 | 91525534 | + | 8.03168461498692 | 15.9179403889013 | 3.57968463110397 | 13.8724124107028 |
| chr1 | 91525579 | 91525587 | + | 3.01188173062009 | 2.27399148412876 | 1.19322821036799 | 1.06710864697714 |
| chr1 | 91680372 | 91680716 | - | 15.0594086531005 | 25.0139063254164 | 9.54582568294392 | 14.9395210576799 |
| chr1 | 91680737 | 91681156 | - | 44.174265382428 | 31.8358807778027 | 21.4781077866238 | 32.0132594093141 |
| chr1 | 91681212 | 91681318 | - | 8.03168461498692 | 3.41098722619315 | 4.77291284147196 | 4.26843458790854 |
| chr1 | 91681478 | 91681598 | - | 4.01584230749346 | 3.41098722619315 | 9.54582568294392 | 2.13421729395427 |
| chr1 | 91681624 | 91681721 | - | 2.00792115374673 | 1.13699574206438 | 2.38645642073598 | 6.40265188186281 |
| chr1 | 91681797 | 91681897 | - | 7.02772403811355 | 13.6439489047726 | 5.96614105183995 | 6.40265188186281 |
| chr1 | 91681919 | 91682262 | - | 31.1227778830743 | 29.5618892936739 | 19.0916513658878 | 56.5567582897882 |
| chr1 | 91682391 | 91682406 | - | 19.0752509605939 | 12.5069531627082 | 8.35259747257593 | 9.60397782279422 |
| chr1 | 91682439 | 91682525 | - | 12.0475269224804 | 4.54798296825753 | 14.3187385244159 | 4.26843458790854 |
| chr1 | 91682557 | 91682998 | - | 140.554480762271 | 103.466612527859 | 177.79100334483 | 81.1002571702623 |
| chr1 | 91683030 | 91683158 | - | 6.02376346124019 | 1.13699574206438 | 2.38645642073598 | 3.20132594093141 |
| chr1 | 92509243 | 92509279 | - | 10.0396057687336 | 13.6439489047726 | 8.35259747257593 | 9.60397782279422 |
| chr1 | 92509299 | 92509313 | - | 5.01980288436682 | 4.54798296825753 | 2.38645642073598 | 1.06710864697714 |
| chr1 | 92509334 | 92509351 | - | 2.00792115374673 | 4.54798296825753 | 1.19322821036799 | 6.40265188186281 |
| chr1 | 92837288 | 92837292 | + | 11.043566345607 | 3.41098722619315 | 25.0577924177278 | 29.8790421153598 |
| chr1 | 92841843 | 92841901 | + | 259.021828833328 | 285.38593125816 | 288.761226909053 | 326.535245975004 |
| chr1 | 93137091 | 93137101 | + | 3.01188173062009 | 4.54798296825753 | 9.54582568294392 | 5.33554323488568 |
| chr1 | 93137648 | 93137699 | + | 7.02772403811355 | 15.9179403889013 | 35.7968463110397 | 24.5434988804741 |
| chr1 | 93137740 | 93137863 | + | 22.087132691214 | 55.7127913611547 | 97.8447132501751 | 110.979299285622 |
| chr1 | 93137967 | 93137991 | + | 3.01188173062009 | 6.82197445238629 | 8.35259747257593 | 3.20132594093141 |
| chr1 | 93138006 | 93138079 | + | 8.03168461498692 | 7.95897019445067 | 62.0478669391355 | 37.3488026441998 |
| chr1 | 93138447 | 93138449 | + | 2.00792115374673 | 4.54798296825753 | 2.38645642073598 | 4.26843458790854 |
| chr1 | 93151878 | 93151942 | - | 15.0594086531005 | 11.3699574206438 | 9.54582568294392 | 5.33554323488568 |
| chr1 | 93152109 | 93152245 | - | 14.0554480762271 | 14.780944646837 | 11.9322821036799 | 17.0737383516342 |
| chr1 | 93152479 | 93152611 | - | 5.01980288436682 | 1.13699574206438 | 4.77291284147196 | 3.20132594093141 |
| chr1 | 93152633 | 93152705 | - | 5.01980288436682 | 3.41098722619315 | 1.19322821036799 | 2.13421729395427 |
| chr1 | 93152742 | 93152747 | - | 17.0673298068472 | 7.95897019445067 | 3.57968463110397 | 8.53686917581709 |
| chr1 | 93154537 | 93154593 | - | 10.0396057687336 | 12.5069531627082 | 8.35259747257593 | 6.40265188186281 |
| chr1 | 93154631 | 93154658 | - | 5.01980288436682 | 6.82197445238629 | 11.9322821036799 | 4.26843458790854 |
| chr1 | 93154679 | 93154821 | - | 116.45942691731 | 138.713480531855 | 139.607700613055 | 129.120146284233 |
| chr1 | 93361043 | 93361098 | + | 37.1465413443145 | 25.0139063254164 | 26.2510206280958 | 35.2145853502455 |
| chr1 | 93361213 | 93361249 | + | 7.02772403811355 | 6.82197445238629 | 5.96614105183995 | 6.40265188186281 |
| chr1 | 93362191 | 93362202 | + | 11.043566345607 | 7.95897019445067 | 3.57968463110397 | 12.8053037637256 |
| chr1 | 93362311 | 93362331 | + | 8.03168461498692 | 13.6439489047726 | 3.57968463110397 | 9.60397782279422 |
| chr1 | 93362509 | 93362512 | + | 1.00396057687336 | 3.41098722619315 | 5.96614105183995 | 1.06710864697714 |
| chr1 | 93364032 | 93364038 | + | 6.02376346124019 | 5.68497871032191 | 1.19322821036799 | 1.06710864697714 |
| chr1 | 93365152 | 93365153 | + | 2.00792115374673 | 2.27399148412876 | 4.77291284147196 | 2.13421729395427 |
| chr1 | 93367907 | 93367922 | + | 5.01980288436682 | 10.2329616785794 | 5.96614105183995 | 5.33554323488568 |
| chr1 | 93551972 | 93552000 | + | 8.03168461498692 | 15.9179403889013 | 8.35259747257593 | 1.06710864697714 |
| chr1 | 93552207 | 93552245 | + | 2.00792115374673 | 4.54798296825753 | 2.38645642073598 | 1.06710864697714 |
| chr1 | 93552476 | 93552531 | + | 18.0712903837206 | 18.1919318730301 | 11.9322821036799 | 4.26843458790854 |
| chr1 | 93847889 | 93847892 | + | 13.0514874993537 | 5.68497871032191 | 2.38645642073598 | 3.20132594093141 |
| chr1 | 93885467 | 93885469 | - | 2.00792115374673 | 9.09596593651506 | 2.38645642073598 | 5.33554323488568 |
| chr1 | 93888978 | 93889000 | - | 13.0514874993537 | 3.41098722619315 | 2.38645642073598 | 5.33554323488568 |
| chr1 | 94518109 | 94518129 | + | 8.03168461498692 | 21.6029190992233 | 1.19322821036799 | 4.26843458790854 |
| chr1 | 94518320 | 94518330 | + | 7.02772403811355 | 17.0549361309657 | 4.77291284147196 | 6.40265188186281 |
| chr1 | 94518463 | 94518486 | + | 7.02772403811355 | 6.82197445238629 | 3.57968463110397 | 3.20132594093141 |
| chr1 | 94983012 | 94983017 | - | 1.00396057687336 | 3.41098722619315 | 2.38645642073598 | 6.40265188186281 |
| chr1 | 95195301 | 95195308 | + | 3.01188173062009 | 1.13699574206438 | 2.38645642073598 | 7.46976052883995 |
| chr1 | 95246934 | 95246954 | + | 10.0396057687336 | 12.5069531627082 | 11.9322821036799 | 14.9395210576799 |
| chr1 | 95247075 | 95247099 | + | 6.02376346124019 | 6.82197445238629 | 5.96614105183995 | 5.33554323488568 |
| chr1 | 95247127 | 95247131 | + | 4.01584230749346 | 4.54798296825753 | 5.96614105183995 | 2.13421729395427 |
| chr1 | 95247185 | 95247190 | + | 2.00792115374673 | 1.13699574206438 | 4.77291284147196 | 6.40265188186281 |
| chr1 | 96814763 | 96814765 | + | 1.00396057687336 | 4.54798296825753 | 2.38645642073598 | 1.06710864697714 |
| chr1 | 99694288 | 99694292 | + | 2.00792115374673 | 6.82197445238629 | 1.19322821036799 | 2.13421729395427 |
| chr1 | 100022510 | 100022512 | + | 2.00792115374673 | 5.68497871032191 | 1.19322821036799 | 1.06710864697714 |
| chr1 | 100022691 | 100022708 | + | 1.00396057687336 | 1.13699574206438 | 4.77291284147196 | 2.13421729395427 |
| chr1 | 100023049 | 100023065 | + | 3.01188173062009 | 5.68497871032191 | 2.38645642073598 | 1.06710864697714 |
| chr1 | 100024456 | 100024459 | + | 1.00396057687336 | 2.27399148412876 | 2.38645642073598 | 6.40265188186281 |
| chr1 | 100083115 | 100083172 | + | 16.0633692299738 | 23.876910583352 | 19.0916513658878 | 18.1408469986113 |
| chr1 | 100083836 | 100083856 | - | 1.00396057687336 | 2.27399148412876 | 2.38645642073598 | 4.26843458790854 |
| chr1 | 100084168 | 100084181 | - | 3.01188173062009 | 2.27399148412876 | 3.57968463110397 | 5.33554323488568 |
| chr1 | 100150088 | 100150089 | + | 3.01188173062009 | 6.82197445238629 | 2.38645642073598 | 4.26843458790854 |
| chr1 | 100291639 | 100291647 | + | 3.01188173062009 | 5.68497871032191 | 3.57968463110397 | 3.20132594093141 |
| chr1 | 100291679 | 100291737 | + | 14.0554480762271 | 14.780944646837 | 22.6713359969918 | 20.2750642925656 |
| chr1 | 100519583 | 100519589 | + | 2.00792115374673 | 1.13699574206438 | 4.77291284147196 | 7.46976052883995 |
| chr1 | 100519922 | 100519927 | + | 2.00792115374673 | 2.27399148412876 | 8.35259747257593 | 30.9461507623369 |
| chr1 | 100974967 | 100974989 | + | 43.1703048055547 | 38.657855230189 | 11.9322821036799 | 11.7381951167485 |
| chr1 | 100975502 | 100975512 | + | 1.00396057687336 | 3.41098722619315 | 2.38645642073598 | 5.33554323488568 |
| chr1 | 100975920 | 100975953 | + | 6.02376346124019 | 12.5069531627082 | 5.96614105183995 | 2.13421729395427 |
| chr1 | 100981119 | 100981124 | + | 8.03168461498692 | 7.95897019445067 | 1.19322821036799 | 1.06710864697714 |
| chr1 | 100981587 | 100981589 | + | 2.00792115374673 | 3.41098722619315 | 1.19322821036799 | 1.06710864697714 |
| chr1 | 100990087 | 100990109 | - | 19.0752509605939 | 28.4248935516096 | 36.9900745214077 | 39.483019938154 |
| chr1 | 100990149 | 100990156 | - | 9.03564519186028 | 7.95897019445067 | 5.96614105183995 | 14.9395210576799 |
| chr1 | 101025837 | 101025906 | + | 104.41189999483 | 118.247557174696 | 176.597775134462 | 252.904749333581 |
| chr1 | 107058706 | 107058726 | + | 2.00792115374673 | 10.2329616785794 | 1.19322821036799 | 1.06710864697714 |
| chr1 | 107058814 | 107058845 | + | 17.0673298068472 | 9.09596593651506 | 2.38645642073598 | 4.26843458790854 |
| chr1 | 108635160 | 108635198 | + | 7.02772403811355 | 10.2329616785794 | 2.38645642073598 | 1.06710864697714 |
| chr1 | 108635347 | 108635365 | + | 5.01980288436682 | 1.13699574206438 | 1.19322821036799 | 1.06710864697714 |
| chr1 | 108635428 | 108635488 | + | 18.0712903837206 | 12.5069531627082 | 2.38645642073598 | 6.40265188186281 |
| chr1 | 108635509 | 108635552 | + | 38.1505019211879 | 53.438799877026 | 11.9322821036799 | 3.20132594093141 |
| chr1 | 108635647 | 108635725 | + | 28.1108961524542 | 40.9318467143178 | 4.77291284147196 | 6.40265188186281 |
| chr1 | 108635757 | 108635803 | + | 25.0990144218341 | 17.0549361309657 | 4.77291284147196 | 2.13421729395427 |
| chr1 | 108635953 | 108636014 | + | 34.1346596136944 | 28.4248935516096 | 8.35259747257593 | 1.06710864697714 |
| chr1 | 108636044 | 108636141 | + | 58.2297134586551 | 77.315710460378 | 13.1255103140479 | 5.33554323488568 |
| chr1 | 108636299 | 108636312 | + | 14.0554480762271 | 9.09596593651506 | 2.38645642073598 | 1.06710864697714 |
| chr1 | 108636426 | 108636477 | + | 37.1465413443145 | 46.6168254246397 | 3.57968463110397 | 7.46976052883995 |
| chr1 | 108636566 | 108636570 | + | 3.01188173062009 | 3.41098722619315 | 2.38645642073598 | 9.60397782279422 |
| chr1 | 108636658 | 108636691 | + | 51.2019894205416 | 51.1648083928972 | 9.54582568294392 | 6.40265188186281 |
| chr1 | 108636741 | 108636774 | + | 23.0910932680874 | 21.6029190992233 | 2.38645642073598 | 1.06710864697714 |
| chr1 | 108636810 | 108636829 | + | 11.043566345607 | 23.876910583352 | 1.19322821036799 | 2.13421729395427 |
| chr1 | 108636852 | 108636879 | + | 19.0752509605939 | 18.1919318730301 | 9.54582568294392 | 5.33554323488568 |
| chr1 | 108636937 | 108636973 | + | 20.0792115374673 | 7.95897019445067 | 2.38645642073598 | 3.20132594093141 |
| chr1 | 108637039 | 108637096 | + | 58.2297134586551 | 73.9047232341848 | 11.9322821036799 | 10.6710864697714 |
| chr1 | 108637278 | 108637355 | + | 20.0792115374673 | 29.5618892936739 | 3.57968463110397 | 1.06710864697714 |
| chr1 | 108637377 | 108637481 | + | 31.1227778830743 | 55.7127913611547 | 10.7390538933119 | 4.26843458790854 |
| chr1 | 108637594 | 108637670 | + | 22.087132691214 | 32.9728765198671 | 11.9322821036799 | 12.8053037637256 |
| chr1 | 108637693 | 108637778 | + | 31.1227778830743 | 34.1098722619315 | 7.15936926220794 | 6.40265188186281 |
| chr1 | 108637849 | 108637870 | + | 9.03564519186028 | 5.68497871032191 | 1.19322821036799 | 4.26843458790854 |
| chr1 | 108638229 | 108638236 | + | 3.01188173062009 | 1.13699574206438 | 2.38645642073598 | 1.06710864697714 |
| chr1 | 108638871 | 108638882 | + | 28.1108961524542 | 15.9179403889013 | 1.19322821036799 | 2.13421729395427 |
| chr1 | 108638992 | 108639040 | + | 10.0396057687336 | 20.4659233571589 | 1.19322821036799 | 4.26843458790854 |
| chr1 | 108639074 | 108639091 | + | 17.0673298068472 | 32.9728765198671 | 1.19322821036799 | 2.13421729395427 |
| chr1 | 108639158 | 108639167 | + | 6.02376346124019 | 15.9179403889013 | 3.57968463110397 | 2.13421729395427 |
| chr1 | 108639207 | 108639220 | + | 19.0752509605939 | 19.3289276150945 | 2.38645642073598 | 3.20132594093141 |
| chr1 | 108639283 | 108639299 | + | 13.0514874993537 | 38.657855230189 | 1.19322821036799 | 3.20132594093141 |
| chr1 | 108700153 | 108700165 | + | 1.00396057687336 | 2.27399148412876 | 4.77291284147196 | 6.40265188186281 |
| chr1 | 108700552 | 108700558 | + | 2.00792115374673 | 1.13699574206438 | 2.38645642073598 | 1.06710864697714 |
| chr1 | 108700868 | 108700892 | + | 17.0673298068472 | 34.1098722619315 | 8.35259747257593 | 16.006629704657 |
| chr1 | 108701154 | 108701192 | + | 5.01980288436682 | 5.68497871032191 | 2.38645642073598 | 1.06710864697714 |
| chr1 | 108808871 | 108808899 | + | 80.3168461498692 | 78.4527062024424 | 34.6036181006717 | 11.7381951167485 |
| chr1 | 108808929 | 108808946 | + | 6.02376346124019 | 7.95897019445067 | 14.3187385244159 | 2.13421729395427 |
| chr1 | 108808971 | 108808988 | + | 3.01188173062009 | 10.2329616785794 | 2.38645642073598 | 1.06710864697714 |
| chr1 | 108809005 | 108809032 | + | 8.03168461498692 | 9.09596593651506 | 9.54582568294392 | 2.13421729395427 |
| chr1 | 108809109 | 108809152 | + | 5.01980288436682 | 3.41098722619315 | 3.57968463110397 | 11.7381951167485 |
| chr1 | 108809219 | 108809239 | + | 1.00396057687336 | 1.13699574206438 | 1.19322821036799 | 1.06710864697714 |
| chr1 | 108809475 | 108809499 | + | 4.01584230749346 | 4.54798296825753 | 1.19322821036799 | 2.13421729395427 |
| chr1 | 108930169 | 108930170 | + | 15.0594086531005 | 14.780944646837 | 1.19322821036799 | 4.26843458790854 |
| chr1 | 108930318 | 108930327 | + | 1.00396057687336 | 1.13699574206438 | 2.38645642073598 | 3.20132594093141 |
| chr1 | 108931270 | 108931311 | - | 10.0396057687336 | 38.657855230189 | 22.6713359969918 | 14.9395210576799 |
| chr1 | 108931372 | 108931436 | - | 10.0396057687336 | 15.9179403889013 | 22.6713359969918 | 12.8053037637256 |
| chr1 | 108931656 | 108931664 | - | 3.01188173062009 | 3.41098722619315 | 9.54582568294392 | 25.6106075274513 |
| chr1 | 108932354 | 108932405 | - | 7.02772403811355 | 9.09596593651506 | 10.7390538933119 | 11.7381951167485 |
| chr1 | 108932488 | 108932489 | - | 4.01584230749346 | 10.2329616785794 | 4.77291284147196 | 4.26843458790854 |
| chr1 | 108970800 | 108970804 | - | 1.00396057687336 | 1.13699574206438 | 3.57968463110397 | 2.13421729395427 |
| chr1 | 109094536 | 109094579 | + | 3.01188173062009 | 2.27399148412876 | 2.38645642073598 | 1.06710864697714 |
| chr1 | 109094751 | 109094773 | + | 4.01584230749346 | 6.82197445238629 | 3.57968463110397 | 3.20132594093141 |
| chr1 | 109094943 | 109094977 | + | 3.01188173062009 | 10.2329616785794 | 2.38645642073598 | 7.46976052883995 |
| chr1 | 109096753 | 109096782 | + | 4.01584230749346 | 7.95897019445067 | 5.96614105183995 | 16.006629704657 |
| chr1 | 109096831 | 109096849 | + | 5.01980288436682 | 7.95897019445067 | 11.9322821036799 | 19.2079556455884 |
| chr1 | 109100188 | 109100239 | + | 29.1148567293276 | 31.8358807778027 | 29.8307052591997 | 28.8119334683827 |
| chr1 | 109237729 | 109237895 | + | 22.087132691214 | 51.1648083928972 | 140.800928823423 | 119.516168461439 |
| chr1 | 109238059 | 109238128 | + | 20.0792115374673 | 51.1648083928972 | 27.4442488384638 | 38.4159112911769 |
| chr1 | 109274281 | 109274318 | + | 5.01980288436682 | 3.41098722619315 | 4.77291284147196 | 5.33554323488568 |
| chr1 | 109274361 | 109274364 | + | 1.00396057687336 | 2.27399148412876 | 3.57968463110397 | 10.6710864697714 |
| chr1 | 109274732 | 109274736 | + | 2.00792115374673 | 3.41098722619315 | 2.38645642073598 | 10.6710864697714 |
| chr1 | 109274757 | 109274759 | + | 4.01584230749346 | 1.13699574206438 | 3.57968463110397 | 5.33554323488568 |
| chr1 | 109279601 | 109279633 | - | 22.087132691214 | 25.0139063254164 | 14.3187385244159 | 6.40265188186281 |
| chr1 | 109279675 | 109279695 | - | 7.02772403811355 | 4.54798296825753 | 7.15936926220794 | 3.20132594093141 |
| chr1 | 109279741 | 109279745 | - | 4.01584230749346 | 9.09596593651506 | 5.96614105183995 | 4.26843458790854 |
| chr1 | 109279922 | 109279933 | - | 3.01188173062009 | 3.41098722619315 | 4.77291284147196 | 3.20132594093141 |
| chr1 | 109279981 | 109280013 | - | 2.00792115374673 | 1.13699574206438 | 5.96614105183995 | 6.40265188186281 |
| chr1 | 109312536 | 109312594 | - | 6.02376346124019 | 2.27399148412876 | 8.35259747257593 | 28.8119334683827 |
| chr1 | 109312725 | 109312761 | - | 4.01584230749346 | 1.13699574206438 | 4.77291284147196 | 2.13421729395427 |
| chr1 | 109313105 | 109313123 | - | 7.02772403811355 | 1.13699574206438 | 10.7390538933119 | 17.0737383516342 |
| chr1 | 109313132 | 109313345 | - | 14.0554480762271 | 13.6439489047726 | 32.2171616799357 | 40.5501285851312 |
| chr1 | 109313444 | 109313463 | - | 2.00792115374673 | 1.13699574206438 | 8.35259747257593 | 16.006629704657 |
| chr1 | 109313483 | 109313610 | - | 91.3604124954762 | 111.425582722309 | 184.950372607038 | 260.374509862421 |
| chr1 | 109313723 | 109313927 | - | 58.2297134586551 | 48.8908169087684 | 146.767069875263 | 215.555946689381 |
| chr1 | 109313979 | 109314077 | - | 2.00792115374673 | 1.13699574206438 | 5.96614105183995 | 4.26843458790854 |
| chr1 | 109401872 | 109402092 | - | 390.540664403739 | 388.852543786019 | 355.582006689661 | 404.434177204334 |
| chr1 | 109592212 | 109592249 | + | 5.01980288436682 | 12.5069531627082 | 32.2171616799357 | 12.8053037637256 |
| chr1 | 109592351 | 109592562 | + | 55.217831728035 | 38.657855230189 | 77.5598336739193 | 91.7713436400337 |
| chr1 | 109592584 | 109592637 | + | 26.1029749987075 | 23.876910583352 | 42.9562155732476 | 29.8790421153598 |
| chr1 | 109592654 | 109592714 | + | 18.0712903837206 | 15.9179403889013 | 33.4103898903037 | 35.2145853502455 |
| chr1 | 109592777 | 109592899 | + | 47.1861471130481 | 46.6168254246397 | 85.9124311464952 | 32.0132594093141 |
| chr1 | 109592957 | 109592997 | + | 9.03564519186028 | 9.09596593651506 | 23.8645642073598 | 12.8053037637256 |
| chr1 | 109593027 | 109593038 | + | 16.0633692299738 | 9.09596593651506 | 3.57968463110397 | 6.40265188186281 |
| chr1 | 109593117 | 109593120 | + | 13.0514874993537 | 2.27399148412876 | 2.38645642073598 | 8.53686917581709 |
| chr1 | 109593212 | 109593249 | + | 5.01980288436682 | 5.68497871032191 | 3.57968463110397 | 11.7381951167485 |
| chr1 | 109593475 | 109593487 | + | 10.0396057687336 | 17.0549361309657 | 14.3187385244159 | 18.1408469986113 |
| chr1 | 109594104 | 109594106 | + | 5.01980288436682 | 3.41098722619315 | 3.57968463110397 | 4.26843458790854 |
| chr1 | 109594172 | 109594177 | + | 2.00792115374673 | 2.27399148412876 | 1.19322821036799 | 5.33554323488568 |
| chr1 | 109594252 | 109594256 | + | 1.00396057687336 | 3.41098722619315 | 8.35259747257593 | 1.06710864697714 |
| chr1 | 109597431 | 109597451 | + | 8.03168461498692 | 10.2329616785794 | 13.1255103140479 | 14.9395210576799 |
| chr1 | 109597894 | 109597907 | + | 2.00792115374673 | 2.27399148412876 | 8.35259747257593 | 11.7381951167485 |
| chr1 | 109631220 | 109631255 | + | 1.00396057687336 | 3.41098722619315 | 5.96614105183995 | 2.13421729395427 |
| chr1 | 109631874 | 109632028 | + | 7.02772403811355 | 10.2329616785794 | 9.54582568294392 | 20.2750642925656 |
| chr1 | 109737008 | 109737026 | - | 56.2217923049084 | 45.4798296825753 | 5.96614105183995 | 1.06710864697714 |
| chr1 | 110021671 | 110021784 | + | 126.499032686044 | 102.329616785794 | 63.2410951495034 | 100.308212815851 |
| chr1 | 110021829 | 110021885 | + | 95.3762548029696 | 70.4937360079917 | 34.6036181006717 | 50.1541064079254 |
| chr1 | 110021905 | 110021964 | + | 169.669337491599 | 220.57717396049 | 121.709277457535 | 118.449059814462 |
| chr1 | 110021998 | 110022028 | + | 26.1029749987075 | 14.780944646837 | 9.54582568294392 | 8.53686917581709 |
| chr1 | 110022064 | 110022065 | + | 1.00396057687336 | 1.13699574206438 | 1.19322821036799 | 1.06710864697714 |
| chr1 | 110022100 | 110022293 | + | 207.819839412786 | 198.974254861267 | 446.267350677628 | 167.53605757541 |
| chr1 | 110022389 | 110022452 | + | 27.1069355755808 | 14.780944646837 | 22.6713359969918 | 17.0737383516342 |
| chr1 | 110022477 | 110022505 | + | 4.01584230749346 | 9.09596593651506 | 4.77291284147196 | 3.20132594093141 |
| chr1 | 110022561 | 110022625 | + | 15.0594086531005 | 6.82197445238629 | 13.1255103140479 | 11.7381951167485 |
| chr1 | 110022651 | 110022658 | + | 2.00792115374673 | 2.27399148412876 | 11.9322821036799 | 4.26843458790854 |
| chr1 | 110022698 | 110022707 | + | 3.01188173062009 | 4.54798296825753 | 4.77291284147196 | 5.33554323488568 |
| chr1 | 110022734 | 110022783 | + | 45.1782259593014 | 29.5618892936739 | 41.7629873628796 | 49.0869977609482 |
| chr1 | 110022806 | 110022833 | + | 3.01188173062009 | 3.41098722619315 | 5.96614105183995 | 6.40265188186281 |
| chr1 | 110022877 | 110022923 | + | 5.01980288436682 | 9.09596593651506 | 17.8984231555198 | 13.8724124107028 |
| chr1 | 110022966 | 110022995 | + | 13.0514874993537 | 17.0549361309657 | 21.4781077866238 | 52.2883237018796 |
| chr1 | 110023025 | 110023083 | + | 7.02772403811355 | 14.780944646837 | 17.8984231555198 | 12.8053037637256 |
| chr1 | 110023130 | 110023150 | + | 9.03564519186028 | 13.6439489047726 | 11.9322821036799 | 12.8053037637256 |
| chr1 | 110023174 | 110023189 | + | 3.01188173062009 | 4.54798296825753 | 11.9322821036799 | 10.6710864697714 |
| chr1 | 110023215 | 110023217 | + | 4.01584230749346 | 5.68497871032191 | 7.15936926220794 | 4.26843458790854 |
| chr1 | 110023240 | 110023263 | + | 19.0752509605939 | 26.1509020674808 | 26.2510206280958 | 20.2750642925656 |
| chr1 | 110023323 | 110023437 | + | 21.0831721143407 | 23.876910583352 | 26.2510206280958 | 41.6172372321083 |
| chr1 | 110023465 | 110023473 | + | 4.01584230749346 | 3.41098722619315 | 1.19322821036799 | 2.13421729395427 |
| chr1 | 110023526 | 110023550 | + | 5.01980288436682 | 3.41098722619315 | 2.38645642073598 | 6.40265188186281 |
| chr1 | 110023616 | 110023643 | + | 39.1544624980612 | 80.7266976865711 | 27.4442488384638 | 91.7713436400337 |
| chr1 | 110023680 | 110023722 | + | 12.0475269224804 | 15.9179403889013 | 8.35259747257593 | 11.7381951167485 |
| chr1 | 110054375 | 110054383 | + | 1.00396057687336 | 9.09596593651506 | 1.19322821036799 | 3.20132594093141 |
| chr1 | 110054536 | 110054543 | + | 2.00792115374673 | 4.54798296825753 | 1.19322821036799 | 3.20132594093141 |
| chr1 | 110059257 | 110059258 | + | 1.00396057687336 | 4.54798296825753 | 9.54582568294392 | 10.6710864697714 |
| chr1 | 110060520 | 110060521 | + | 33.130699036821 | 13.6439489047726 | 23.8645642073598 | 53.3554323488568 |
| chr1 | 110342333 | 110342378 | + | 8.03168461498692 | 21.6029190992233 | 17.8984231555198 | 14.9395210576799 |
| chr1 | 110342481 | 110342544 | + | 14.0554480762271 | 13.6439489047726 | 10.7390538933119 | 16.006629704657 |
| chr1 | 110345538 | 110345650 | + | 14.0554480762271 | 13.6439489047726 | 26.2510206280958 | 29.8790421153598 |
| chr1 | 110370769 | 110370772 | + | 10.0396057687336 | 5.68497871032191 | 9.54582568294392 | 2.13421729395427 |
| chr1 | 110371286 | 110371310 | + | 3.01188173062009 | 6.82197445238629 | 9.54582568294392 | 3.20132594093141 |
| chr1 | 110371872 | 110371890 | + | 9.03564519186028 | 12.5069531627082 | 41.7629873628796 | 21.3421729395427 |
| chr1 | 110372102 | 110372105 | + | 1.00396057687336 | 2.27399148412876 | 1.19322821036799 | 3.20132594093141 |
| chr1 | 110372850 | 110372858 | + | 4.01584230749346 | 1.13699574206438 | 1.19322821036799 | 2.13421729395427 |
| chr1 | 110372949 | 110372965 | + | 5.01980288436682 | 6.82197445238629 | 4.77291284147196 | 6.40265188186281 |
| chr1 | 110401279 | 110401363 | - | 63.249516343022 | 103.466612527859 | 54.8884976769275 | 74.6976052883995 |
| chr1 | 110401383 | 110401438 | - | 7.02772403811355 | 14.780944646837 | 31.0239334695677 | 16.006629704657 |
| chr1 | 110401459 | 110401551 | - | 24.0950538449607 | 31.8358807778027 | 40.5697591525116 | 29.8790421153598 |
| chr1 | 110407782 | 110407847 | + | 36.1425807674411 | 47.753821166704 | 33.4103898903037 | 24.5434988804741 |
| chr1 | 110947220 | 110947257 | - | 4.01584230749346 | 9.09596593651506 | 5.96614105183995 | 5.33554323488568 |
| chr1 | 110947343 | 110947364 | - | 2.00792115374673 | 1.13699574206438 | 2.38645642073598 | 10.6710864697714 |
| chr1 | 110947572 | 110947575 | - | 2.00792115374673 | 1.13699574206438 | 1.19322821036799 | 3.20132594093141 |
| chr1 | 110947756 | 110947888 | - | 12.0475269224804 | 2.27399148412876 | 16.7051949451519 | 27.7448248214055 |
| chr1 | 110963301 | 110963359 | + | 14.0554480762271 | 11.3699574206438 | 2.38645642073598 | 2.13421729395427 |
| chr1 | 111117438 | 111117457 | - | 3.01188173062009 | 11.3699574206438 | 9.54582568294392 | 5.33554323488568 |
| chr1 | 111117479 | 111117481 | - | 5.01980288436682 | 11.3699574206438 | 8.35259747257593 | 4.26843458790854 |
| chr1 | 111117930 | 111117985 | - | 9.03564519186028 | 12.5069531627082 | 11.9322821036799 | 7.46976052883995 |
| chr1 | 111118010 | 111118022 | - | 1.00396057687336 | 1.13699574206438 | 3.57968463110397 | 1.06710864697714 |
| chr1 | 111118096 | 111118118 | - | 7.02772403811355 | 18.1919318730301 | 14.3187385244159 | 21.3421729395427 |
| chr1 | 111183461 | 111183555 | + | 3.01188173062009 | 2.27399148412876 | 8.35259747257593 | 3.20132594093141 |
| chr1 | 111184377 | 111184380 | + | 1.00396057687336 | 3.41098722619315 | 1.19322821036799 | 3.20132594093141 |
| chr1 | 111184777 | 111184781 | + | 2.00792115374673 | 5.68497871032191 | 1.19322821036799 | 5.33554323488568 |
| chr1 | 111184896 | 111184938 | + | 5.01980288436682 | 10.2329616785794 | 5.96614105183995 | 7.46976052883995 |
| chr1 | 111185012 | 111185013 | + | 3.01188173062009 | 3.41098722619315 | 1.19322821036799 | 1.06710864697714 |
| chr1 | 111440287 | 111440324 | - | 22.087132691214 | 14.780944646837 | 10.7390538933119 | 11.7381951167485 |
| chr1 | 111440399 | 111440420 | - | 16.0633692299738 | 4.54798296825753 | 2.38645642073598 | 16.006629704657 |
| chr1 | 111440461 | 111440465 | - | 4.01584230749346 | 1.13699574206438 | 2.38645642073598 | 1.06710864697714 |
| chr1 | 111440507 | 111440528 | - | 18.0712903837206 | 7.95897019445067 | 3.57968463110397 | 12.8053037637256 |
| chr1 | 111440971 | 111440979 | - | 11.043566345607 | 7.95897019445067 | 7.15936926220794 | 10.6710864697714 |
| chr1 | 111441002 | 111441080 | - | 16.0633692299738 | 13.6439489047726 | 19.0916513658878 | 24.5434988804741 |
| chr1 | 111461042 | 111461059 | + | 6.02376346124019 | 5.68497871032191 | 2.38645642073598 | 7.46976052883995 |
| chr1 | 111461195 | 111461200 | + | 6.02376346124019 | 4.54798296825753 | 2.38645642073598 | 1.06710864697714 |
| chr1 | 111461381 | 111461385 | + | 10.0396057687336 | 18.1919318730301 | 8.35259747257593 | 5.33554323488568 |
| chr1 | 111709259 | 111709265 | + | 8.03168461498692 | 2.27399148412876 | 8.35259747257593 | 4.26843458790854 |
| chr1 | 111712500 | 111712507 | + | 5.01980288436682 | 12.5069531627082 | 3.57968463110397 | 8.53686917581709 |
| chr1 | 111712571 | 111712574 | + | 8.03168461498692 | 7.95897019445067 | 1.19322821036799 | 3.20132594093141 |
| chr1 | 111712826 | 111712856 | + | 7.02772403811355 | 2.27399148412876 | 2.38645642073598 | 1.06710864697714 |
| chr1 | 111712960 | 111713010 | + | 20.0792115374673 | 12.5069531627082 | 28.6374770488317 | 6.40265188186281 |
| chr1 | 111713062 | 111713094 | + | 10.0396057687336 | 18.1919318730301 | 28.6374770488317 | 9.60397782279422 |
| chr1 | 111767163 | 111767166 | + | 3.01188173062009 | 2.27399148412876 | 2.38645642073598 | 2.13421729395427 |
| chr1 | 112670153 | 112670180 | + | 4.01584230749346 | 6.82197445238629 | 27.4442488384638 | 54.4225409958339 |
| chr1 | 112670227 | 112670348 | + | 226.89509037338 | 155.76841666282 | 289.954455119421 | 376.689352382929 |
| chr1 | 112670388 | 112670416 | + | 16.0633692299738 | 14.780944646837 | 46.5359002043516 | 27.7448248214055 |
| chr1 | 112670484 | 112670545 | + | 17.0673298068472 | 13.6439489047726 | 42.9562155732476 | 18.1408469986113 |
| chr1 | 112670566 | 112670606 | + | 13.0514874993537 | 5.68497871032191 | 34.6036181006717 | 14.9395210576799 |
| chr1 | 112670635 | 112670668 | + | 4.01584230749346 | 4.54798296825753 | 23.8645642073598 | 3.20132594093141 |
| chr1 | 112670755 | 112670790 | + | 11.043566345607 | 10.2329616785794 | 39.3765309421437 | 26.6777161744284 |
| chr1 | 112670823 | 112670843 | + | 16.0633692299738 | 4.54798296825753 | 62.0478669391355 | 12.8053037637256 |
| chr1 | 112670906 | 112670922 | + | 3.01188173062009 | 2.27399148412876 | 4.77291284147196 | 3.20132594093141 |
| chr1 | 112670950 | 112670960 | + | 2.00792115374673 | 1.13699574206438 | 2.38645642073598 | 3.20132594093141 |
| chr1 | 112671294 | 112671300 | + | 2.00792115374673 | 1.13699574206438 | 3.57968463110397 | 6.40265188186281 |
| chr1 | 112671346 | 112671352 | + | 19.0752509605939 | 20.4659233571589 | 41.7629873628796 | 57.6238669367653 |
| chr1 | 112671398 | 112671492 | + | 19.0752509605939 | 21.6029190992233 | 53.6952694665595 | 36.2816939972226 |
| chr1 | 112700591 | 112700618 | + | 24.0950538449607 | 23.876910583352 | 34.6036181006717 | 35.2145853502455 |
| chr1 | 112700654 | 112700719 | + | 53.2099105742883 | 60.2607743294122 | 76.3666054635513 | 93.9055609339879 |
| chr1 | 112701152 | 112701160 | - | 76.3010038423757 | 100.055625301666 | 103.810854302015 | 141.925450047959 |
| chr1 | 112701187 | 112701206 | - | 11.043566345607 | 9.09596593651506 | 5.96614105183995 | 25.6106075274513 |
| chr1 | 112701458 | 112701517 | - | 4.01584230749346 | 2.27399148412876 | 9.54582568294392 | 6.40265188186281 |
| chr1 | 113761893 | 113761908 | - | 1.00396057687336 | 2.27399148412876 | 4.77291284147196 | 2.13421729395427 |
| chr1 | 113762077 | 113762078 | - | 1.00396057687336 | 1.13699574206438 | 1.19322821036799 | 1.06710864697714 |
| chr1 | 113763968 | 113763981 | - | 1.00396057687336 | 1.13699574206438 | 3.57968463110397 | 4.26843458790854 |
| chr1 | 113764074 | 113764108 | - | 1.00396057687336 | 2.27399148412876 | 5.96614105183995 | 8.53686917581709 |
| chr1 | 113765524 | 113765536 | - | 1.00396057687336 | 6.82197445238629 | 4.77291284147196 | 5.33554323488568 |
| chr1 | 113973584 | 113973606 | + | 2.00792115374673 | 5.68497871032191 | 1.19322821036799 | 2.13421729395427 |
| chr1 | 113973633 | 113973699 | + | 11.043566345607 | 28.4248935516096 | 31.0239334695677 | 48.0198891139711 |
| chr1 | 113975007 | 113975043 | + | 13.0514874993537 | 6.82197445238629 | 2.38645642073598 | 5.33554323488568 |
| chr1 | 113975066 | 113975092 | + | 2.00792115374673 | 4.54798296825753 | 13.1255103140479 | 5.33554323488568 |
| chr1 | 113975231 | 113975256 | + | 3.01188173062009 | 2.27399148412876 | 5.96614105183995 | 4.26843458790854 |
| chr1 | 113975447 | 113975457 | + | 1.00396057687336 | 3.41098722619315 | 4.77291284147196 | 4.26843458790854 |
| chr1 | 113975491 | 113975498 | + | 2.00792115374673 | 3.41098722619315 | 1.19322821036799 | 1.06710864697714 |
| chr1 | 113975702 | 113975723 | + | 2.00792115374673 | 6.82197445238629 | 5.96614105183995 | 7.46976052883995 |
| chr1 | 113975752 | 113975771 | + | 6.02376346124019 | 6.82197445238629 | 7.15936926220794 | 2.13421729395427 |
| chr1 | 113975861 | 113975899 | + | 6.02376346124019 | 6.82197445238629 | 5.96614105183995 | 12.8053037637256 |
| chr1 | 113976012 | 113976050 | + | 8.03168461498692 | 7.95897019445067 | 8.35259747257593 | 14.9395210576799 |
| chr1 | 113976079 | 113976100 | + | 7.02772403811355 | 4.54798296825753 | 8.35259747257593 | 6.40265188186281 |
| chr1 | 113976125 | 113976146 | + | 7.02772403811355 | 4.54798296825753 | 1.19322821036799 | 1.06710864697714 |
| chr1 | 113976169 | 113976179 | + | 3.01188173062009 | 4.54798296825753 | 9.54582568294392 | 14.9395210576799 |
| chr1 | 113976412 | 113976414 | + | 1.00396057687336 | 1.13699574206438 | 4.77291284147196 | 2.13421729395427 |
| chr1 | 113976525 | 113976551 | + | 5.01980288436682 | 3.41098722619315 | 5.96614105183995 | 4.26843458790854 |
| chr1 | 113977206 | 113977221 | + | 5.01980288436682 | 6.82197445238629 | 3.57968463110397 | 11.7381951167485 |
| chr1 | 114392832 | 114392895 | - | 3.01188173062009 | 2.27399148412876 | 2.38645642073598 | 4.26843458790854 |
| chr1 | 114393194 | 114393202 | - | 43.1703048055547 | 30.6988850357383 | 1.19322821036799 | 3.20132594093141 |
| chr1 | 114393330 | 114393348 | - | 6.02376346124019 | 2.27399148412876 | 1.19322821036799 | 4.26843458790854 |
| chr1 | 114394305 | 114394345 | - | 10.0396057687336 | 11.3699574206438 | 11.9322821036799 | 9.60397782279422 |
| chr1 | 114394568 | 114394598 | - | 15.0594086531005 | 9.09596593651506 | 11.9322821036799 | 6.40265188186281 |
| chr1 | 114394652 | 114394686 | - | 7.02772403811355 | 2.27399148412876 | 5.96614105183995 | 4.26843458790854 |
| chr1 | 114394749 | 114394800 | - | 7.02772403811355 | 20.4659233571589 | 4.77291284147196 | 10.6710864697714 |
| chr1 | 114395377 | 114395393 | - | 2.00792115374673 | 4.54798296825753 | 4.77291284147196 | 2.13421729395427 |
| chr1 | 114395426 | 114395430 | - | 15.0594086531005 | 7.95897019445067 | 11.9322821036799 | 11.7381951167485 |
| chr1 | 114395553 | 114395592 | - | 7.02772403811355 | 4.54798296825753 | 5.96614105183995 | 5.33554323488568 |
| chr1 | 114395666 | 114395671 | - | 15.0594086531005 | 15.9179403889013 | 22.6713359969918 | 3.20132594093141 |
| chr1 | 114395828 | 114395884 | - | 51.2019894205416 | 62.534765813541 | 31.0239334695677 | 32.0132594093141 |
| chr1 | 114396248 | 114396274 | - | 6.02376346124019 | 10.2329616785794 | 7.15936926220794 | 3.20132594093141 |
| chr1 | 114396381 | 114396403 | - | 12.0475269224804 | 7.95897019445067 | 1.19322821036799 | 3.20132594093141 |
| chr1 | 114567593 | 114567614 | - | 2.00792115374673 | 1.13699574206438 | 1.19322821036799 | 2.13421729395427 |
| chr1 | 114567693 | 114567709 | - | 1.00396057687336 | 2.27399148412876 | 1.19322821036799 | 1.06710864697714 |
| chr1 | 114567785 | 114567799 | - | 5.01980288436682 | 2.27399148412876 | 2.38645642073598 | 4.26843458790854 |
| chr1 | 114568056 | 114568144 | - | 63.249516343022 | 38.657855230189 | 42.9562155732476 | 57.6238669367653 |
| chr1 | 114704490 | 114704492 | - | 3.01188173062009 | 2.27399148412876 | 1.19322821036799 | 3.20132594093141 |
| chr1 | 114704595 | 114704622 | - | 7.02772403811355 | 6.82197445238629 | 11.9322821036799 | 16.006629704657 |
| chr1 | 114704669 | 114704773 | - | 1106.36455571445 | 1032.39213379446 | 1010.66429418169 | 1355.22798166096 |
| chr1 | 114704796 | 114704817 | - | 30.1188173062009 | 36.3838637460602 | 25.0577924177278 | 19.2079556455884 |
| chr1 | 114704876 | 114704911 | - | 10.0396057687336 | 11.3699574206438 | 5.96614105183995 | 9.60397782279422 |
| chr1 | 114705031 | 114705038 | - | 10.0396057687336 | 4.54798296825753 | 7.15936926220794 | 9.60397782279422 |
| chr1 | 114705111 | 114705167 | - | 11.043566345607 | 11.3699574206438 | 22.6713359969918 | 23.476390233497 |
| chr1 | 114705326 | 114705345 | - | 3.01188173062009 | 1.13699574206438 | 7.15936926220794 | 5.33554323488568 |
| chr1 | 114705367 | 114705381 | - | 2.00792115374673 | 4.54798296825753 | 5.96614105183995 | 9.60397782279422 |
| chr1 | 114705527 | 114705585 | - | 2.00792115374673 | 4.54798296825753 | 11.9322821036799 | 12.8053037637256 |
| chr1 | 114705616 | 114705625 | - | 2.00792115374673 | 5.68497871032191 | 7.15936926220794 | 3.20132594093141 |
| chr1 | 114705939 | 114705941 | - | 3.01188173062009 | 3.41098722619315 | 1.19322821036799 | 1.06710864697714 |
| chr1 | 114705985 | 114705989 | - | 1.00396057687336 | 1.13699574206438 | 5.96614105183995 | 2.13421729395427 |
| chr1 | 114706030 | 114706061 | - | 2.00792115374673 | 4.54798296825753 | 27.4442488384638 | 8.53686917581709 |
| chr1 | 114706391 | 114706404 | - | 6.02376346124019 | 4.54798296825753 | 7.15936926220794 | 4.26843458790854 |
| chr1 | 114706435 | 114706514 | - | 7.02772403811355 | 3.41098722619315 | 4.77291284147196 | 22.4092815865198 |
| chr1 | 114706640 | 114706656 | - | 6.02376346124019 | 2.27399148412876 | 16.7051949451519 | 19.2079556455884 |
| chr1 | 114706685 | 114706719 | - | 9.03564519186028 | 9.09596593651506 | 19.0916513658878 | 36.2816939972226 |
| chr1 | 114706852 | 114706875 | - | 3.01188173062009 | 2.27399148412876 | 7.15936926220794 | 3.20132594093141 |
| chr1 | 114707000 | 114707031 | - | 3.01188173062009 | 5.68497871032191 | 3.57968463110397 | 8.53686917581709 |
| chr1 | 114707435 | 114707463 | - | 3.01188173062009 | 1.13699574206438 | 4.77291284147196 | 8.53686917581709 |
| chr1 | 114707726 | 114707749 | - | 17.0673298068472 | 7.95897019445067 | 14.3187385244159 | 13.8724124107028 |
| chr1 | 114707837 | 114707841 | - | 6.02376346124019 | 7.95897019445067 | 3.57968463110397 | 3.20132594093141 |
| chr1 | 114707863 | 114707894 | - | 6.02376346124019 | 5.68497871032191 | 8.35259747257593 | 2.13421729395427 |
| chr1 | 114707921 | 114707927 | - | 2.00792115374673 | 1.13699574206438 | 5.96614105183995 | 3.20132594093141 |
| chr1 | 114707973 | 114708035 | - | 28.1108961524542 | 26.1509020674808 | 76.3666054635513 | 23.476390233497 |
| chr1 | 114716946 | 114716966 | - | 13.0514874993537 | 22.7399148412876 | 14.3187385244159 | 8.53686917581709 |
| chr1 | 114716997 | 114717042 | - | 4.01584230749346 | 12.5069531627082 | 9.54582568294392 | 7.46976052883995 |
| chr1 | 114717050 | 114717126 | - | 59.2336740355285 | 37.5208594881246 | 29.8307052591997 | 38.4159112911769 |
| chr1 | 114717182 | 114717199 | - | 2.00792115374673 | 1.13699574206438 | 3.57968463110397 | 22.4092815865198 |
| chr1 | 114717290 | 114717308 | - | 1.00396057687336 | 1.13699574206438 | 5.96614105183995 | 3.20132594093141 |
| chr1 | 114717349 | 114717369 | - | 29.1148567293276 | 21.6029190992233 | 31.0239334695677 | 21.3421729395427 |
| chr1 | 114717410 | 114717599 | - | 8003.57371883446 | 11082.2974979015 | 17067.9363211037 | 10617.7310374225 |
| chr1 | 114717621 | 114717658 | - | 29.1148567293276 | 50.0278126508328 | 76.3666054635513 | 33.0803680562912 |
| chr1 | 114717675 | 114717715 | - | 531.09514516601 | 600.333751809994 | 571.556312766267 | 444.984305789466 |
| chr1 | 114717742 | 114717791 | - | 6.02376346124019 | 9.09596593651506 | 21.4781077866238 | 20.2750642925656 |
| chr1 | 114717815 | 114717992 | - | 98.3881365335897 | 87.5486721389574 | 88.2988875672312 | 89.6371263460794 |
| chr1 | 114718020 | 114718078 | - | 11.043566345607 | 25.0139063254164 | 20.2848795762558 | 10.6710864697714 |
| chr1 | 114718107 | 114718147 | - | 115.455466340437 | 106.877599754052 | 83.5259747257593 | 68.2949534065367 |
| chr1 | 114769516 | 114769542 | - | 5.01980288436682 | 6.82197445238629 | 3.57968463110397 | 1.06710864697714 |
| chr1 | 114769815 | 114769819 | - | 2.00792115374673 | 1.13699574206438 | 2.38645642073598 | 2.13421729395427 |
| chr1 | 114773313 | 114773333 | - | 7.02772403811355 | 1.13699574206438 | 9.54582568294392 | 41.6172372321083 |
| chr1 | 114773348 | 114773380 | - | 4.01584230749346 | 4.54798296825753 | 15.5119667347839 | 12.8053037637256 |
| chr1 | 114773451 | 114773476 | - | 4.01584230749346 | 4.54798296825753 | 2.38645642073598 | 7.46976052883995 |
| chr1 | 114773588 | 114773607 | - | 4.01584230749346 | 4.54798296825753 | 9.54582568294392 | 4.26843458790854 |
| chr1 | 114773667 | 114773741 | - | 9.03564519186028 | 15.9179403889013 | 16.7051949451519 | 8.53686917581709 |
| chr1 | 114774003 | 114774043 | - | 5.01980288436682 | 4.54798296825753 | 11.9322821036799 | 4.26843458790854 |
| chr1 | 114774120 | 114774155 | - | 2.00792115374673 | 4.54798296825753 | 9.54582568294392 | 9.60397782279422 |
| chr1 | 114774229 | 114774272 | - | 9.03564519186028 | 11.3699574206438 | 10.7390538933119 | 14.9395210576799 |
| chr1 | 115691445 | 115691452 | + | 4.01584230749346 | 4.54798296825753 | 1.19322821036799 | 5.33554323488568 |
| chr1 | 115691857 | 115691869 | + | 1.00396057687336 | 1.13699574206438 | 4.77291284147196 | 1.06710864697714 |
| chr1 | 116404459 | 116404757 | + | 241.954499026481 | 269.467990869259 | 215.974306076606 | 539.956975370431 |
| chr1 | 116514575 | 116514583 | - | 4.01584230749346 | 4.54798296825753 | 2.38645642073598 | 3.20132594093141 |
| chr1 | 116574759 | 116574813 | - | 9.03564519186028 | 3.41098722619315 | 9.54582568294392 | 4.26843458790854 |
| chr1 | 116576181 | 116576184 | - | 4.01584230749346 | 2.27399148412876 | 16.7051949451519 | 7.46976052883995 |
| chr1 | 116986975 | 116987015 | + | 3.01188173062009 | 3.41098722619315 | 9.54582568294392 | 4.26843458790854 |
| chr1 | 116987086 | 116987089 | + | 9.03564519186028 | 5.68497871032191 | 2.38645642073598 | 1.06710864697714 |
| chr1 | 116987130 | 116987166 | + | 8.03168461498692 | 1.13699574206438 | 9.54582568294392 | 17.0737383516342 |
| chr1 | 116987298 | 116987349 | + | 5.01980288436682 | 14.780944646837 | 7.15936926220794 | 18.1408469986113 |
| chr1 | 116987556 | 116987562 | + | 12.0475269224804 | 6.82197445238629 | 9.54582568294392 | 10.6710864697714 |
| chr1 | 116988009 | 116988013 | + | 7.02772403811355 | 1.13699574206438 | 3.57968463110397 | 1.06710864697714 |
| chr1 | 116988178 | 116988197 | + | 6.02376346124019 | 11.3699574206438 | 3.57968463110397 | 6.40265188186281 |
| chr1 | 116990073 | 116990095 | + | 6.02376346124019 | 4.54798296825753 | 2.38645642073598 | 4.26843458790854 |
| chr1 | 117102363 | 117102397 | + | 6.02376346124019 | 4.54798296825753 | 4.77291284147196 | 4.26843458790854 |
| chr1 | 117104787 | 117104799 | + | 13.0514874993537 | 7.95897019445067 | 13.1255103140479 | 3.20132594093141 |
| chr1 | 117105751 | 117105752 | + | 1.00396057687336 | 2.27399148412876 | 7.15936926220794 | 3.20132594093141 |
| chr1 | 117106295 | 117106328 | + | 10.0396057687336 | 11.3699574206438 | 4.77291284147196 | 2.13421729395427 |
| chr1 | 117107103 | 117107105 | + | 3.01188173062009 | 2.27399148412876 | 1.19322821036799 | 5.33554323488568 |
| chr1 | 117111661 | 117111664 | - | 3.01188173062009 | 4.54798296825753 | 2.38645642073598 | 3.20132594093141 |
| chr1 | 117524043 | 117524051 | + | 8.03168461498692 | 2.27399148412876 | 3.57968463110397 | 9.60397782279422 |
| chr1 | 117528731 | 117528733 | + | 3.01188173062009 | 4.54798296825753 | 5.96614105183995 | 2.13421729395427 |
| chr1 | 117934526 | 117934532 | + | 3.01188173062009 | 4.54798296825753 | 4.77291284147196 | 3.20132594093141 |
| chr1 | 117934554 | 117934562 | + | 4.01584230749346 | 2.27399148412876 | 10.7390538933119 | 2.13421729395427 |
| chr1 | 117934590 | 117934593 | + | 2.00792115374673 | 2.27399148412876 | 2.38645642073598 | 2.13421729395427 |
| chr1 | 117936777 | 117936792 | + | 1.00396057687336 | 1.13699574206438 | 2.38645642073598 | 4.26843458790854 |
| chr1 | 117936826 | 117936832 | + | 2.00792115374673 | 1.13699574206438 | 1.19322821036799 | 4.26843458790854 |
| chr1 | 117938492 | 117938508 | + | 1.00396057687336 | 3.41098722619315 | 4.77291284147196 | 9.60397782279422 |
| chr1 | 117939475 | 117939504 | + | 9.03564519186028 | 6.82197445238629 | 11.9322821036799 | 4.26843458790854 |
| chr1 | 117939541 | 117939551 | + | 3.01188173062009 | 5.68497871032191 | 4.77291284147196 | 1.06710864697714 |
| chr1 | 117940898 | 117940938 | + | 14.0554480762271 | 10.2329616785794 | 17.8984231555198 | 14.9395210576799 |
| chr1 | 117941159 | 117941167 | + | 1.00396057687336 | 3.41098722619315 | 4.77291284147196 | 6.40265188186281 |
| chr1 | 117959733 | 117959735 | + | 2.00792115374673 | 7.95897019445067 | 1.19322821036799 | 2.13421729395427 |
| chr1 | 117959955 | 117959958 | + | 1.00396057687336 | 4.54798296825753 | 2.38645642073598 | 2.13421729395427 |
| chr1 | 117960305 | 117960306 | + | 1.00396057687336 | 1.13699574206438 | 1.19322821036799 | 2.13421729395427 |
| chr1 | 119140373 | 119140450 | + | 312.231739407616 | 779.979079056166 | 146.767069875263 | 173.938709457273 |
| chr1 | 119140746 | 119140771 | + | 4.01584230749346 | 10.2329616785794 | 16.7051949451519 | 20.2750642925656 |
| chr1 | 119150652 | 119150653 | + | 69.2732798042621 | 62.534765813541 | 14.3187385244159 | 16.006629704657 |
| chr1 | 119742830 | 119743042 | + | 84.3326884573626 | 80.7266976865711 | 4.77291284147196 | 1.06710864697714 |
| chr1 | 119743934 | 119744113 | + | 130.514874993537 | 170.549361309657 | 5.96614105183995 | 2.13421729395427 |
| chr1 | 120151403 | 120151409 | - | 2.00792115374673 | 17.0549361309657 | 7.15936926220794 | 6.40265188186281 |
| chr1 | 120155781 | 120155812 | - | 8.03168461498692 | 17.0549361309657 | 7.15936926220794 | 12.8053037637256 |
| chr1 | 120156789 | 120156801 | - | 6.02376346124019 | 12.5069531627082 | 7.15936926220794 | 10.6710864697714 |
| chr1 | 144412572 | 144412578 | + | 179.708943260332 | 193.289276150945 | 33.4103898903037 | 19.2079556455884 |
| chr1 | 145281114 | 145281121 | + | 349.378280751931 | 405.907479916984 | 41.7629873628796 | 20.2750642925656 |
| chr1 | 145670346 | 145670349 | + | 3.01188173062009 | 9.09596593651506 | 1.19322821036799 | 8.53686917581709 |
| chr1 | 145744553 | 145744627 | - | 185.732706721572 | 280.837948289902 | 236.259185652862 | 250.770532039627 |
| chr1 | 145744665 | 145744688 | - | 8.03168461498692 | 6.82197445238629 | 7.15936926220794 | 6.40265188186281 |
| chr1 | 145744761 | 145744766 | - | 23.0910932680874 | 20.4659233571589 | 28.6374770488317 | 14.9395210576799 |
| chr1 | 145744805 | 145744806 | - | 3.01188173062009 | 2.27399148412876 | 3.57968463110397 | 2.13421729395427 |
| chr1 | 145744851 | 145744881 | - | 4.01584230749346 | 1.13699574206438 | 9.54582568294392 | 7.46976052883995 |
| chr1 | 145745077 | 145745081 | - | 2.00792115374673 | 2.27399148412876 | 4.77291284147196 | 1.06710864697714 |
| chr1 | 145746538 | 145746564 | - | 2.00792115374673 | 13.6439489047726 | 5.96614105183995 | 6.40265188186281 |
| chr1 | 145746604 | 145746654 | - | 20.0792115374673 | 14.780944646837 | 9.54582568294392 | 22.4092815865198 |
| chr1 | 145746672 | 145746698 | - | 14.0554480762271 | 7.95897019445067 | 10.7390538933119 | 10.6710864697714 |
| chr1 | 145746741 | 145746747 | - | 1.00396057687336 | 1.13699574206438 | 4.77291284147196 | 1.06710864697714 |
| chr1 | 145746833 | 145746842 | - | 22.087132691214 | 21.6029190992233 | 19.0916513658878 | 21.3421729395427 |
| chr1 | 145848854 | 145848863 | + | 2.00792115374673 | 4.54798296825753 | 1.19322821036799 | 2.13421729395427 |
| chr1 | 145911372 | 145911419 | - | 7.02772403811355 | 10.2329616785794 | 10.7390538933119 | 7.46976052883995 |
| chr1 | 145911668 | 145911678 | - | 2.00792115374673 | 2.27399148412876 | 1.19322821036799 | 1.06710864697714 |
| chr1 | 145911763 | 145911789 | - | 3.01188173062009 | 3.41098722619315 | 2.38645642073598 | 4.26843458790854 |
| chr1 | 145911951 | 145912039 | - | 5.01980288436682 | 14.780944646837 | 9.54582568294392 | 4.26843458790854 |
| chr1 | 145923347 | 145923385 | - | 23.0910932680874 | 11.3699574206438 | 4.77291284147196 | 3.20132594093141 |
| chr1 | 145925422 | 145925432 | - | 10.0396057687336 | 18.1919318730301 | 4.77291284147196 | 4.26843458790854 |
| chr1 | 145925773 | 145925800 | - | 16.0633692299738 | 36.3838637460602 | 16.7051949451519 | 10.6710864697714 |
| chr1 | 145927581 | 145927666 | + | 214.8475634509 | 264.920007901001 | 73.9801490428153 | 51.2212150549025 |
| chr1 | 145933617 | 145933648 | - | 12.0475269224804 | 6.82197445238629 | 1.19322821036799 | 7.46976052883995 |
| chr1 | 145934294 | 145934312 | - | 28.1108961524542 | 23.876910583352 | 9.54582568294392 | 4.26843458790854 |
| chr1 | 145935594 | 145935598 | - | 11.043566345607 | 15.9179403889013 | 4.77291284147196 | 4.26843458790854 |
| chr1 | 145960238 | 145960243 | - | 17.0673298068472 | 6.82197445238629 | 2.38645642073598 | 1.06710864697714 |
| chr1 | 145978784 | 145978813 | + | 12.0475269224804 | 4.54798296825753 | 17.8984231555198 | 46.952780466994 |
| chr1 | 145992465 | 145992492 | - | 5.01980288436682 | 69.3567402659273 | 7.15936926220794 | 16.006629704657 |
| chr1 | 145992568 | 145992609 | - | 9.03564519186028 | 73.9047232341848 | 27.4442488384638 | 53.3554323488568 |
| chr1 | 145992630 | 145992722 | - | 4.01584230749346 | 22.7399148412876 | 8.35259747257593 | 5.33554323488568 |
| chr1 | 145992923 | 145992947 | - | 3.01188173062009 | 1.13699574206438 | 10.7390538933119 | 4.26843458790854 |
| chr1 | 145992974 | 145993031 | - | 1.00396057687336 | 47.753821166704 | 51.3088130458235 | 20.2750642925656 |
| chr1 | 145993056 | 145993180 | - | 11.043566345607 | 96.6446380754725 | 156.312895558207 | 72.5633879944452 |
| chr1 | 145993287 | 145993311 | - | 4.01584230749346 | 25.0139063254164 | 36.9900745214077 | 19.2079556455884 |
| chr1 | 145993335 | 145993720 | - | 65.2574374967687 | 477.53821166704 | 347.229409217085 | 279.58246550801 |
| chr1 | 145993746 | 145993767 | - | 4.01584230749346 | 36.3838637460602 | 26.2510206280958 | 37.3488026441998 |
| chr1 | 145993806 | 145993909 | - | 3.01188173062009 | 29.5618892936739 | 59.6614105183995 | 39.483019938154 |
| chr1 | 146052602 | 146052625 | + | 1.00396057687336 | 4.54798296825753 | 3.57968463110397 | 3.20132594093141 |
| chr1 | 147156275 | 147156395 | - | 1.00396057687336 | 2.27399148412876 | 3.57968463110397 | 1.06710864697714 |
| chr1 | 147156433 | 147156453 | - | 1.00396057687336 | 3.41098722619315 | 1.19322821036799 | 2.13421729395427 |
| chr1 | 147156574 | 147156605 | - | 2.00792115374673 | 2.27399148412876 | 2.38645642073598 | 2.13421729395427 |
| chr1 | 147156895 | 147157004 | - | 2.00792115374673 | 2.27399148412876 | 2.38645642073598 | 5.33554323488568 |
| chr1 | 147172742 | 147172817 | + | 18.0712903837206 | 19.3289276150945 | 2.38645642073598 | 4.26843458790854 |
| chr1 | 147252649 | 147252728 | + | 7.02772403811355 | 4.54798296825753 | 7.15936926220794 | 11.7381951167485 |
| chr1 | 147265970 | 147265988 | + | 4.01584230749346 | 1.13699574206438 | 15.5119667347839 | 6.40265188186281 |
| chr1 | 147266049 | 147266077 | + | 2.00792115374673 | 3.41098722619315 | 10.7390538933119 | 5.33554323488568 |
| chr1 | 147267424 | 147267519 | + | 8.03168461498692 | 5.68497871032191 | 23.8645642073598 | 4.26843458790854 |
| chr1 | 147268745 | 147268805 | + | 11.043566345607 | 3.41098722619315 | 16.7051949451519 | 11.7381951167485 |
| chr1 | 147268845 | 147268855 | + | 1.00396057687336 | 3.41098722619315 | 3.57968463110397 | 5.33554323488568 |
| chr1 | 147268871 | 147268879 | + | 19.0752509605939 | 15.9179403889013 | 29.8307052591997 | 27.7448248214055 |
| chr1 | 147270931 | 147270944 | + | 3.01188173062009 | 2.27399148412876 | 13.1255103140479 | 6.40265188186281 |
| chr1 | 147270970 | 147271006 | + | 2.00792115374673 | 6.82197445238629 | 7.15936926220794 | 10.6710864697714 |
| chr1 | 147295597 | 147295734 | + | 8.03168461498692 | 15.9179403889013 | 13.1255103140479 | 22.4092815865198 |
| chr1 | 147625096 | 147625099 | + | 3.01188173062009 | 2.27399148412876 | 2.38645642073598 | 1.06710864697714 |
| chr1 | 147625594 | 147625598 | + | 4.01584230749346 | 5.68497871032191 | 2.38645642073598 | 6.40265188186281 |
| chr1 | 148263473 | 148263477 | + | 17.0673298068472 | 26.1509020674808 | 15.5119667347839 | 17.0737383516342 |
| chr1 | 149846036 | 149846038 | + | 3.01188173062009 | 2.27399148412876 | 1.19322821036799 | 4.26843458790854 |
| chr1 | 149886100 | 149886103 | - | 1.00396057687336 | 1.13699574206438 | 1.19322821036799 | 1.06710864697714 |
| chr1 | 149923451 | 149923460 | - | 13.0514874993537 | 46.6168254246397 | 15.5119667347839 | 42.6843458790854 |
| chr1 | 149940551 | 149940556 | - | 13.0514874993537 | 7.95897019445067 | 5.96614105183995 | 20.2750642925656 |
| chr1 | 149940583 | 149940592 | - | 5.01980288436682 | 1.13699574206438 | 1.19322821036799 | 3.20132594093141 |
| chr1 | 149940864 | 149940890 | - | 10.0396057687336 | 12.5069531627082 | 16.7051949451519 | 4.26843458790854 |
| chr1 | 149941059 | 149941091 | - | 6.02376346124019 | 17.0549361309657 | 2.38645642073598 | 5.33554323488568 |
| chr1 | 149941285 | 149941308 | - | 21.0831721143407 | 29.5618892936739 | 7.15936926220794 | 6.40265188186281 |
| chr1 | 149941893 | 149941899 | - | 6.02376346124019 | 12.5069531627082 | 3.57968463110397 | 3.20132594093141 |
| chr1 | 149942222 | 149942224 | - | 3.01188173062009 | 4.54798296825753 | 1.19322821036799 | 5.33554323488568 |
| chr1 | 149942941 | 149942991 | - | 42.1663442286813 | 50.0278126508328 | 26.2510206280958 | 28.8119334683827 |
| chr1 | 149943031 | 149943042 | - | 3.01188173062009 | 7.95897019445067 | 4.77291284147196 | 2.13421729395427 |
| chr1 | 149943223 | 149943239 | - | 10.0396057687336 | 14.780944646837 | 9.54582568294392 | 3.20132594093141 |
| chr1 | 149943271 | 149943296 | - | 45.1782259593014 | 56.8497871032191 | 35.7968463110397 | 26.6777161744284 |
| chr1 | 149943395 | 149943415 | - | 10.0396057687336 | 26.1509020674808 | 2.38645642073598 | 1.06710864697714 |
| chr1 | 149943481 | 149943541 | - | 98.3881365335897 | 137.57648478979 | 31.0239334695677 | 48.0198891139711 |
| chr1 | 150145277 | 150145285 | + | 3.01188173062009 | 6.82197445238629 | 1.19322821036799 | 3.20132594093141 |
| chr1 | 150218530 | 150218758 | - | 630.487242276473 | 618.525683683024 | 1100.15640995929 | 769.385334470515 |
| chr1 | 150218791 | 150218865 | - | 22.087132691214 | 18.1919318730301 | 32.2171616799357 | 20.2750642925656 |
| chr1 | 150218893 | 150218939 | - | 15.0594086531005 | 5.68497871032191 | 8.35259747257593 | 5.33554323488568 |
| chr1 | 150218968 | 150219030 | - | 186.736667298446 | 195.563267635074 | 59.6614105183995 | 92.8384522870108 |
| chr1 | 150219050 | 150219059 | - | 2.00792115374673 | 4.54798296825753 | 3.57968463110397 | 1.06710864697714 |
| chr1 | 150219149 | 150219194 | - | 7.02772403811355 | 12.5069531627082 | 13.1255103140479 | 6.40265188186281 |
| chr1 | 150219239 | 150219280 | - | 29.1148567293276 | 17.0549361309657 | 83.5259747257593 | 32.0132594093141 |
| chr1 | 150219331 | 150219395 | - | 28.1108961524542 | 30.6988850357383 | 58.4681823080315 | 67.2278447595595 |
| chr1 | 150219441 | 150219484 | - | 6.02376346124019 | 5.68497871032191 | 20.2848795762558 | 19.2079556455884 |
| chr1 | 150219591 | 150219605 | - | 7.02772403811355 | 4.54798296825753 | 3.57968463110397 | 1.06710864697714 |
| chr1 | 150219627 | 150219678 | - | 66.2613980736421 | 42.0688424563821 | 87.1056593568632 | 84.3015831111937 |
| chr1 | 150219770 | 150219808 | - | 13.0514874993537 | 9.09596593651506 | 20.2848795762558 | 11.7381951167485 |
| chr1 | 150219831 | 150219874 | - | 6.02376346124019 | 12.5069531627082 | 16.7051949451519 | 12.8053037637256 |
| chr1 | 150219936 | 150220011 | - | 24.0950538449607 | 28.4248935516096 | 64.4343233598714 | 42.6843458790854 |
| chr1 | 150220043 | 150220045 | - | 4.01584230749346 | 5.68497871032191 | 2.38645642073598 | 1.06710864697714 |
| chr1 | 150220070 | 150220120 | - | 18.0712903837206 | 18.1919318730301 | 23.8645642073598 | 16.006629704657 |
| chr1 | 150220141 | 150220148 | - | 4.01584230749346 | 5.68497871032191 | 2.38645642073598 | 1.06710864697714 |
| chr1 | 150220235 | 150220324 | - | 28.1108961524542 | 31.8358807778027 | 89.4921157775992 | 75.7647139353766 |
| chr1 | 150220353 | 150220431 | - | 27.1069355755808 | 31.8358807778027 | 46.5359002043516 | 57.6238669367653 |
| chr1 | 150220461 | 150220614 | - | 35.1386201905678 | 54.5757956190903 | 44.1494437836156 | 42.6843458790854 |
| chr1 | 150220622 | 150220667 | - | 16.0633692299738 | 29.5618892936739 | 8.35259747257593 | 18.1408469986113 |
| chr1 | 150220696 | 150220763 | - | 95.3762548029696 | 120.521548658824 | 71.5936926220794 | 109.912190638645 |
| chr1 | 150226616 | 150226798 | - | 472.865431707355 | 491.182160571813 | 841.225888309433 | 670.144230301641 |
| chr1 | 150265431 | 150265471 | - | 92.3643730723495 | 159.179403889013 | 34.6036181006717 | 78.966039876308 |
| chr1 | 150265499 | 150265642 | - | 5590.05249203089 | 5185.83757955565 | 2341.113748742 | 3041.25964388484 |
| chr1 | 150265685 | 150265739 | - | 73.2891221117556 | 64.8087572976698 | 33.4103898903037 | 26.6777161744284 |
| chr1 | 150265760 | 150265857 | - | 144.570323069764 | 148.946442210434 | 93.0718004087032 | 27.7448248214055 |
| chr1 | 150265890 | 150265919 | - | 25.0990144218341 | 22.7399148412876 | 13.1255103140479 | 7.46976052883995 |
| chr1 | 150265980 | 150266003 | - | 9.03564519186028 | 4.54798296825753 | 3.57968463110397 | 11.7381951167485 |
| chr1 | 150266080 | 150266132 | - | 12.0475269224804 | 5.68497871032191 | 3.57968463110397 | 4.26843458790854 |
| chr1 | 150308118 | 150308330 | + | 576.273371125311 | 407.044475659049 | 315.012247537149 | 473.796239257848 |
| chr1 | 150308353 | 150308357 | + | 8.03168461498692 | 11.3699574206438 | 1.19322821036799 | 3.20132594093141 |
| chr1 | 150353109 | 150353173 | + | 23.0910932680874 | 27.2878978095452 | 2.38645642073598 | 12.8053037637256 |
| chr1 | 150473267 | 150473397 | + | 238.942617295861 | 293.344901452611 | 275.635716595006 | 169.670274869365 |
| chr1 | 150473542 | 150473546 | + | 27.1069355755808 | 54.5757956190903 | 45.3426719939836 | 5.33554323488568 |
| chr1 | 150473617 | 150473635 | + | 19.0752509605939 | 23.876910583352 | 7.15936926220794 | 6.40265188186281 |
| chr1 | 150473864 | 150473900 | + | 17.0673298068472 | 40.9318467143178 | 8.35259747257593 | 16.006629704657 |
| chr1 | 150474035 | 150474044 | + | 8.03168461498692 | 17.0549361309657 | 9.54582568294392 | 2.13421729395427 |
| chr1 | 150474053 | 150474097 | + | 5.01980288436682 | 2.27399148412876 | 5.96614105183995 | 4.26843458790854 |
| chr1 | 150475260 | 150475264 | + | 30.1188173062009 | 50.0278126508328 | 19.0916513658878 | 24.5434988804741 |
| chr1 | 150574541 | 150574555 | - | 2.00792115374673 | 3.41098722619315 | 2.38645642073598 | 3.20132594093141 |
| chr1 | 150574583 | 150574662 | - | 30.1188173062009 | 42.0688424563821 | 25.0577924177278 | 24.5434988804741 |
| chr1 | 150574709 | 150574724 | - | 27.1069355755808 | 27.2878978095452 | 19.0916513658878 | 34.1474767032683 |
| chr1 | 150574787 | 150574838 | - | 2.00792115374673 | 6.82197445238629 | 5.96614105183995 | 12.8053037637256 |
| chr1 | 150574885 | 150574964 | - | 35.1386201905678 | 23.876910583352 | 20.2848795762558 | 58.6909755837425 |
| chr1 | 150574995 | 150575019 | - | 4.01584230749346 | 1.13699574206438 | 1.19322821036799 | 13.8724124107028 |
| chr1 | 150575050 | 150575079 | - | 2.00792115374673 | 3.41098722619315 | 1.19322821036799 | 6.40265188186281 |
| chr1 | 150575277 | 150575310 | - | 5.01980288436682 | 4.54798296825753 | 5.96614105183995 | 6.40265188186281 |
| chr1 | 150575552 | 150575555 | - | 2.00792115374673 | 3.41098722619315 | 4.77291284147196 | 6.40265188186281 |
| chr1 | 150575590 | 150575645 | - | 1.00396057687336 | 1.13699574206438 | 4.77291284147196 | 1.06710864697714 |
| chr1 | 150575773 | 150575806 | - | 5.01980288436682 | 3.41098722619315 | 7.15936926220794 | 18.1408469986113 |
| chr1 | 150575843 | 150576184 | - | 110.43566345607 | 122.795540142953 | 309.046106485309 | 203.817751572633 |
| chr1 | 150576207 | 150576349 | - | 40.1584230749346 | 42.0688424563821 | 62.0478669391355 | 49.0869977609482 |
| chr1 | 150576372 | 150576402 | - | 12.0475269224804 | 21.6029190992233 | 14.3187385244159 | 22.4092815865198 |
| chr1 | 150576444 | 150576534 | - | 18.0712903837206 | 30.6988850357383 | 29.8307052591997 | 30.9461507623369 |
| chr1 | 150576555 | 150576651 | - | 12.0475269224804 | 10.2329616785794 | 14.3187385244159 | 17.0737383516342 |
| chr1 | 150576706 | 150576719 | - | 2.00792115374673 | 1.13699574206438 | 9.54582568294392 | 12.8053037637256 |
| chr1 | 150576732 | 150576766 | - | 9.03564519186028 | 10.2329616785794 | 13.1255103140479 | 8.53686917581709 |
| chr1 | 150576801 | 150576816 | - | 3.01188173062009 | 3.41098722619315 | 5.96614105183995 | 3.20132594093141 |
| chr1 | 150576832 | 150577013 | - | 25.0990144218341 | 17.0549361309657 | 47.7291284147196 | 48.0198891139711 |
| chr1 | 150577035 | 150577061 | - | 10.0396057687336 | 2.27399148412876 | 5.96614105183995 | 5.33554323488568 |
| chr1 | 150577083 | 150577090 | - | 21.0831721143407 | 43.2058381984465 | 45.3426719939836 | 28.8119334683827 |
| chr1 | 150577114 | 150577132 | - | 13.0514874993537 | 15.9179403889013 | 23.8645642073598 | 21.3421729395427 |
| chr1 | 150577175 | 150577215 | - | 5.01980288436682 | 5.68497871032191 | 13.1255103140479 | 16.006629704657 |
| chr1 | 150577306 | 150577315 | - | 3.01188173062009 | 2.27399148412876 | 3.57968463110397 | 2.13421729395427 |
| chr1 | 150577356 | 150577421 | - | 14.0554480762271 | 18.1919318730301 | 17.8984231555198 | 11.7381951167485 |
| chr1 | 150622081 | 150622085 | - | 4.01584230749346 | 7.95897019445067 | 9.54582568294392 | 3.20132594093141 |
| chr1 | 150622144 | 150622250 | - | 79.3128855729958 | 87.5486721389574 | 51.3088130458235 | 91.7713436400337 |
| chr1 | 150622312 | 150622315 | - | 2.00792115374673 | 2.27399148412876 | 1.19322821036799 | 2.13421729395427 |
| chr1 | 150622387 | 150622394 | - | 4.01584230749346 | 1.13699574206438 | 2.38645642073598 | 1.06710864697714 |
| chr1 | 150622429 | 150622487 | - | 36.1425807674411 | 45.4798296825753 | 27.4442488384638 | 29.8790421153598 |
| chr1 | 150622519 | 150622585 | - | 406.604033633713 | 403.633488432856 | 139.607700613055 | 228.361250453107 |
| chr1 | 150622613 | 150622695 | - | 10.0396057687336 | 4.54798296825753 | 10.7390538933119 | 38.4159112911769 |
| chr1 | 150622712 | 150622724 | - | 1.00396057687336 | 6.82197445238629 | 2.38645642073598 | 4.26843458790854 |
| chr1 | 150623830 | 150623845 | - | 2.00792115374673 | 4.54798296825753 | 1.19322821036799 | 1.06710864697714 |
| chr1 | 150623933 | 150623951 | - | 11.043566345607 | 21.6029190992233 | 3.57968463110397 | 7.46976052883995 |
| chr1 | 150624897 | 150624900 | - | 2.00792115374673 | 1.13699574206438 | 1.19322821036799 | 1.06710864697714 |
| chr1 | 150625464 | 150625488 | - | 1.00396057687336 | 1.13699574206438 | 1.19322821036799 | 1.06710864697714 |
| chr1 | 150625641 | 150625647 | - | 8.03168461498692 | 1.13699574206438 | 1.19322821036799 | 1.06710864697714 |
| chr1 | 150646332 | 150646336 | - | 1.00396057687336 | 2.27399148412876 | 5.96614105183995 | 5.33554323488568 |
| chr1 | 150810133 | 150810171 | - | 1.00396057687336 | 2.27399148412876 | 4.77291284147196 | 7.46976052883995 |
| chr1 | 150810251 | 150810295 | - | 5.01980288436682 | 1.13699574206438 | 9.54582568294392 | 3.20132594093141 |
| chr1 | 150810661 | 150810686 | - | 2.00792115374673 | 1.13699574206438 | 5.96614105183995 | 9.60397782279422 |
| chr1 | 150811417 | 150811424 | - | 7.02772403811355 | 4.54798296825753 | 5.96614105183995 | 4.26843458790854 |
| chr1 | 150811604 | 150811632 | - | 15.0594086531005 | 12.5069531627082 | 7.15936926220794 | 9.60397782279422 |
| chr1 | 150811782 | 150811803 | - | 6.02376346124019 | 4.54798296825753 | 3.57968463110397 | 6.40265188186281 |
| chr1 | 150811832 | 150811841 | - | 1.00396057687336 | 4.54798296825753 | 1.19322821036799 | 2.13421729395427 |
| chr1 | 150811989 | 150812000 | - | 10.0396057687336 | 17.0549361309657 | 21.4781077866238 | 13.8724124107028 |
| chr1 | 150964569 | 150964580 | + | 4.01584230749346 | 7.95897019445067 | 5.96614105183995 | 4.26843458790854 |
| chr1 | 150965195 | 150965222 | - | 81.3208067267425 | 63.6717615556054 | 29.8307052591997 | 28.8119334683827 |
| chr1 | 150965312 | 150965364 | - | 24.0950538449607 | 18.1919318730301 | 9.54582568294392 | 24.5434988804741 |
| chr1 | 150965388 | 150965450 | - | 17.0673298068472 | 18.1919318730301 | 11.9322821036799 | 11.7381951167485 |
| chr1 | 150965479 | 150965536 | - | 15.0594086531005 | 9.09596593651506 | 7.15936926220794 | 9.60397782279422 |
| chr1 | 150965558 | 150965588 | - | 15.0594086531005 | 6.82197445238629 | 11.9322821036799 | 10.6710864697714 |
| chr1 | 151035561 | 151035588 | + | 26.1029749987075 | 32.9728765198671 | 23.8645642073598 | 39.483019938154 |
| chr1 | 151035611 | 151035628 | + | 4.01584230749346 | 11.3699574206438 | 1.19322821036799 | 3.20132594093141 |
| chr1 | 151051023 | 151051070 | - | 7.02772403811355 | 13.6439489047726 | 27.4442488384638 | 16.006629704657 |
| chr1 | 151051217 | 151051227 | - | 1.00396057687336 | 1.13699574206438 | 2.38645642073598 | 7.46976052883995 |
| chr1 | 151052558 | 151052591 | - | 1.00396057687336 | 1.13699574206438 | 13.1255103140479 | 3.20132594093141 |
| chr1 | 151052742 | 151052772 | - | 1.00396057687336 | 1.13699574206438 | 16.7051949451519 | 5.33554323488568 |
| chr1 | 151052840 | 151052856 | - | 17.0673298068472 | 37.5208594881246 | 168.245177661887 | 49.0869977609482 |
| chr1 | 151052922 | 151052971 | - | 4.01584230749346 | 5.68497871032191 | 16.7051949451519 | 11.7381951167485 |
| chr1 | 151053008 | 151053018 | - | 6.02376346124019 | 5.68497871032191 | 9.54582568294392 | 6.40265188186281 |
| chr1 | 151053100 | 151053105 | - | 1.00396057687336 | 3.41098722619315 | 3.57968463110397 | 4.26843458790854 |
| chr1 | 151053189 | 151053214 | - | 2.00792115374673 | 1.13699574206438 | 16.7051949451519 | 21.3421729395427 |
| chr1 | 151053241 | 151053289 | - | 1.00396057687336 | 2.27399148412876 | 9.54582568294392 | 5.33554323488568 |
| chr1 | 151118437 | 151118471 | + | 9.03564519186028 | 12.5069531627082 | 1.19322821036799 | 5.33554323488568 |
| chr1 | 151122296 | 151122319 | + | 1.00396057687336 | 6.82197445238629 | 1.19322821036799 | 4.26843458790854 |
| chr1 | 151168990 | 151169089 | + | 139.550520185398 | 136.439489047726 | 32.2171616799357 | 77.8989312293309 |
| chr1 | 151248338 | 151248380 | + | 48.1901076899215 | 57.9867828452835 | 1.19322821036799 | 4.26843458790854 |
| chr1 | 151249383 | 151249433 | + | 51.2019894205416 | 134.165497563597 | 4.77291284147196 | 9.60397782279422 |
| chr1 | 151267346 | 151267365 | + | 51.2019894205416 | 60.2607743294122 | 57.2749540976635 | 69.3620620535138 |
| chr1 | 151267435 | 151267452 | + | 9.03564519186028 | 13.6439489047726 | 9.54582568294392 | 2.13421729395427 |
| chr1 | 151291741 | 151291751 | + | 11.043566345607 | 17.0549361309657 | 3.57968463110397 | 6.40265188186281 |
| chr1 | 151291794 | 151291797 | + | 2.00792115374673 | 5.68497871032191 | 2.38645642073598 | 3.20132594093141 |
| chr1 | 151291854 | 151291866 | + | 3.01188173062009 | 2.27399148412876 | 5.96614105183995 | 6.40265188186281 |
| chr1 | 151292182 | 151292194 | - | 11.043566345607 | 22.7399148412876 | 22.6713359969918 | 28.8119334683827 |
| chr1 | 151292226 | 151292247 | - | 5.01980288436682 | 13.6439489047726 | 3.57968463110397 | 5.33554323488568 |
| chr1 | 151292446 | 151292472 | - | 5.01980288436682 | 2.27399148412876 | 10.7390538933119 | 10.6710864697714 |
| chr1 | 151292513 | 151292530 | - | 4.01584230749346 | 4.54798296825753 | 3.57968463110397 | 9.60397782279422 |
| chr1 | 151292649 | 151292685 | - | 8.03168461498692 | 9.09596593651506 | 2.38645642073598 | 6.40265188186281 |
| chr1 | 151340670 | 151340699 | - | 9.03564519186028 | 3.41098722619315 | 2.38645642073598 | 1.06710864697714 |
| chr1 | 151340802 | 151340806 | - | 4.01584230749346 | 5.68497871032191 | 1.19322821036799 | 2.13421729395427 |
| chr1 | 151340846 | 151340862 | - | 7.02772403811355 | 12.5069531627082 | 3.57968463110397 | 6.40265188186281 |
| chr1 | 151340946 | 151341007 | - | 17.0673298068472 | 23.876910583352 | 2.38645642073598 | 1.06710864697714 |
| chr1 | 151342165 | 151342175 | - | 10.0396057687336 | 7.95897019445067 | 1.19322821036799 | 3.20132594093141 |
| chr1 | 151346957 | 151347057 | + | 114.451505763564 | 155.76841666282 | 112.163451774591 | 87.5029090521251 |
| chr1 | 151364335 | 151364339 | - | 1.00396057687336 | 1.13699574206438 | 1.19322821036799 | 5.33554323488568 |
| chr1 | 151364491 | 151364493 | - | 4.01584230749346 | 1.13699574206438 | 1.19322821036799 | 2.13421729395427 |
| chr1 | 151401814 | 151401917 | + | 28.1108961524542 | 68.2197445238629 | 31.0239334695677 | 27.7448248214055 |
| chr1 | 151402791 | 151402858 | - | 9.03564519186028 | 20.4659233571589 | 7.15936926220794 | 11.7381951167485 |
| chr1 | 151403262 | 151403281 | - | 1.00396057687336 | 1.13699574206438 | 1.19322821036799 | 1.06710864697714 |
| chr1 | 151404465 | 151404470 | - | 2.00792115374673 | 5.68497871032191 | 2.38645642073598 | 1.06710864697714 |
| chr1 | 151404713 | 151404772 | - | 27.1069355755808 | 55.7127913611547 | 9.54582568294392 | 14.9395210576799 |
| chr1 | 151583478 | 151583492 | + | 10.0396057687336 | 14.780944646837 | 1.19322821036799 | 2.13421729395427 |
| chr1 | 151693664 | 151693671 | + | 4.01584230749346 | 6.82197445238629 | 1.19322821036799 | 1.06710864697714 |
| chr1 | 151694326 | 151694332 | + | 6.02376346124019 | 7.95897019445067 | 2.38645642073598 | 6.40265188186281 |
| chr1 | 151695740 | 151695743 | + | 2.00792115374673 | 2.27399148412876 | 2.38645642073598 | 2.13421729395427 |
| chr1 | 151696186 | 151696223 | + | 22.087132691214 | 25.0139063254164 | 5.96614105183995 | 4.26843458790854 |
| chr1 | 151696336 | 151696374 | + | 12.0475269224804 | 6.82197445238629 | 9.54582568294392 | 1.06710864697714 |
| chr1 | 151696394 | 151696416 | + | 25.0990144218341 | 21.6029190992233 | 23.8645642073598 | 24.5434988804741 |
| chr1 | 151759665 | 151759694 | - | 3.01188173062009 | 1.13699574206438 | 7.15936926220794 | 10.6710864697714 |
| chr1 | 151759721 | 151759753 | - | 6.02376346124019 | 9.09596593651506 | 19.0916513658878 | 22.4092815865198 |
| chr1 | 151759786 | 151759806 | - | 4.01584230749346 | 4.54798296825753 | 1.19322821036799 | 3.20132594093141 |
| chr1 | 151759833 | 151759866 | - | 7.02772403811355 | 11.3699574206438 | 7.15936926220794 | 10.6710864697714 |
| chr1 | 151790812 | 151790873 | + | 39.1544624980612 | 85.2746806548286 | 60.8546387287675 | 38.4159112911769 |
| chr1 | 151874414 | 151874416 | - | 2.00792115374673 | 1.13699574206438 | 2.38645642073598 | 8.53686917581709 |
| chr1 | 152032531 | 152032619 | - | 283.116882678289 | 238.76910583352 | 26.2510206280958 | 65.0936274656053 |
| chr1 | 152032651 | 152032675 | - | 68.2693192273888 | 37.5208594881246 | 3.57968463110397 | 14.9395210576799 |
| chr1 | 153534623 | 153534832 | - | 2617.32522390886 | 2397.92402001378 | 2.38645642073598 | 23.476390233497 |
| chr1 | 153618817 | 153618878 | - | 227.899050950254 | 266.057003643065 | 38.1833027317757 | 122.717494402371 |
| chr1 | 153643243 | 153643365 | + | 39.1544624980612 | 42.0688424563821 | 22.6713359969918 | 36.2816939972226 |
| chr1 | 153645063 | 153645132 | + | 23.0910932680874 | 29.5618892936739 | 13.1255103140479 | 12.8053037637256 |
| chr1 | 153645221 | 153645249 | + | 7.02772403811355 | 5.68497871032191 | 2.38645642073598 | 5.33554323488568 |
| chr1 | 153645285 | 153645290 | + | 3.01188173062009 | 3.41098722619315 | 1.19322821036799 | 3.20132594093141 |
| chr1 | 153645341 | 153645408 | + | 16.0633692299738 | 28.4248935516096 | 11.9322821036799 | 14.9395210576799 |
| chr1 | 153645442 | 153645463 | + | 11.043566345607 | 27.2878978095452 | 10.7390538933119 | 2.13421729395427 |
| chr1 | 153645505 | 153645511 | + | 28.1108961524542 | 21.6029190992233 | 9.54582568294392 | 5.33554323488568 |
| chr1 | 153645676 | 153645678 | + | 8.03168461498692 | 9.09596593651506 | 4.77291284147196 | 2.13421729395427 |
| chr1 | 153645705 | 153645721 | + | 7.02772403811355 | 5.68497871032191 | 1.19322821036799 | 1.06710864697714 |
| chr1 | 153645745 | 153645773 | + | 18.0712903837206 | 26.1509020674808 | 8.35259747257593 | 9.60397782279422 |
| chr1 | 153645832 | 153645864 | + | 4.01584230749346 | 5.68497871032191 | 2.38645642073598 | 7.46976052883995 |
| chr1 | 153645924 | 153645941 | + | 129.510914416664 | 171.686357051722 | 25.0577924177278 | 34.1474767032683 |
| chr1 | 153645983 | 153646078 | + | 395.560467288106 | 585.552807163157 | 223.133675338814 | 268.911379038238 |
| chr1 | 153661706 | 153661795 | + | 35.1386201905678 | 56.8497871032191 | 7.15936926220794 | 23.476390233497 |
| chr1 | 153662033 | 153662194 | - | 92.3643730723495 | 155.76841666282 | 71.5936926220794 | 90.7042349930565 |
| chr1 | 153662233 | 153662305 | - | 15.0594086531005 | 13.6439489047726 | 11.9322821036799 | 12.8053037637256 |
| chr1 | 153662325 | 153662368 | - | 10.0396057687336 | 2.27399148412876 | 2.38645642073598 | 4.26843458790854 |
| chr1 | 153929523 | 153929604 | - | 9.03564519186028 | 13.6439489047726 | 5.96614105183995 | 11.7381951167485 |
| chr1 | 153947734 | 153947748 | - | 6.02376346124019 | 9.09596593651506 | 3.57968463110397 | 4.26843458790854 |
| chr1 | 153959138 | 153959153 | - | 22.087132691214 | 18.1919318730301 | 8.35259747257593 | 6.40265188186281 |
| chr1 | 153959185 | 153959208 | - | 57.2257528817818 | 94.3706465913437 | 19.0916513658878 | 30.9461507623369 |
| chr1 | 153959236 | 153959244 | - | 30.1188173062009 | 25.0139063254164 | 1.19322821036799 | 18.1408469986113 |
| chr1 | 153959373 | 153959411 | - | 33.130699036821 | 25.0139063254164 | 28.6374770488317 | 16.006629704657 |
| chr1 | 153959438 | 153959455 | - | 43.1703048055547 | 9.09596593651506 | 14.3187385244159 | 11.7381951167485 |
| chr1 | 153959544 | 153959550 | - | 2.00792115374673 | 3.41098722619315 | 1.19322821036799 | 3.20132594093141 |
| chr1 | 153959571 | 153959610 | - | 19.0752509605939 | 4.54798296825753 | 5.96614105183995 | 7.46976052883995 |
| chr1 | 153959648 | 153959675 | - | 8.03168461498692 | 6.82197445238629 | 5.96614105183995 | 3.20132594093141 |
| chr1 | 153959744 | 153959756 | - | 13.0514874993537 | 10.2329616785794 | 8.35259747257593 | 3.20132594093141 |
| chr1 | 153959790 | 153959814 | - | 6.02376346124019 | 9.09596593651506 | 2.38645642073598 | 1.06710864697714 |
| chr1 | 153959831 | 153959894 | - | 29.1148567293276 | 23.876910583352 | 32.2171616799357 | 29.8790421153598 |
| chr1 | 153959922 | 153960159 | - | 178.704982683459 | 164.864382599335 | 132.448331350847 | 75.7647139353766 |
| chr1 | 153960206 | 153960248 | - | 5.01980288436682 | 7.95897019445067 | 3.57968463110397 | 4.26843458790854 |
| chr1 | 153960283 | 153960294 | - | 5.01980288436682 | 2.27399148412876 | 1.19322821036799 | 2.13421729395427 |
| chr1 | 153960348 | 153960384 | - | 10.0396057687336 | 2.27399148412876 | 9.54582568294392 | 3.20132594093141 |
| chr1 | 153960521 | 153960540 | - | 6.02376346124019 | 4.54798296825753 | 3.57968463110397 | 2.13421729395427 |
| chr1 | 153974311 | 153974338 | + | 12.0475269224804 | 18.1919318730301 | 3.57968463110397 | 2.13421729395427 |
| chr1 | 153974532 | 153974655 | - | 168.665376914725 | 147.80944646837 | 41.7629873628796 | 99.2411041688736 |
| chr1 | 153977732 | 153977750 | + | 7.02772403811355 | 5.68497871032191 | 14.3187385244159 | 4.26843458790854 |
| chr1 | 153981891 | 153981913 | - | 17.0673298068472 | 13.6439489047726 | 11.9322821036799 | 23.476390233497 |
| chr1 | 153981942 | 153981964 | - | 20.0792115374673 | 25.0139063254164 | 23.8645642073598 | 37.3488026441998 |
| chr1 | 153982002 | 153982074 | - | 527.079302858516 | 467.305249988461 | 591.841192342523 | 709.627250239795 |
| chr1 | 154156379 | 154156380 | - | 2.00792115374673 | 3.41098722619315 | 4.77291284147196 | 1.06710864697714 |
| chr1 | 154156589 | 154156597 | - | 13.0514874993537 | 4.54798296825753 | 9.54582568294392 | 28.8119334683827 |
| chr1 | 154156833 | 154156856 | - | 25.0990144218341 | 12.5069531627082 | 51.3088130458235 | 57.6238669367653 |
| chr1 | 154157138 | 154157217 | - | 208.82379998966 | 126.206527369146 | 244.611783125438 | 256.106075274513 |
| chr1 | 154157511 | 154157532 | - | 4.01584230749346 | 1.13699574206438 | 4.77291284147196 | 9.60397782279422 |
| chr1 | 154157571 | 154157590 | - | 20.0792115374673 | 12.5069531627082 | 22.6713359969918 | 19.2079556455884 |
| chr1 | 154157663 | 154157684 | - | 15.0594086531005 | 15.9179403889013 | 40.5697591525116 | 54.4225409958339 |
| chr1 | 154167811 | 154167815 | - | 12.0475269224804 | 6.82197445238629 | 11.9322821036799 | 26.6777161744284 |
| chr1 | 154206738 | 154206852 | - | 240.950538449607 | 353.605675782023 | 195.68942650035 | 136.589906813073 |
| chr1 | 154206888 | 154206924 | - | 12.0475269224804 | 23.876910583352 | 8.35259747257593 | 6.40265188186281 |
| chr1 | 154206955 | 154207098 | - | 147.582204800385 | 206.933225055718 | 60.8546387287675 | 89.6371263460794 |
| chr1 | 154207431 | 154207472 | - | 4.01584230749346 | 3.41098722619315 | 7.15936926220794 | 4.26843458790854 |
| chr1 | 154207609 | 154207651 | - | 3.01188173062009 | 7.95897019445067 | 5.96614105183995 | 4.26843458790854 |
| chr1 | 154212291 | 154212368 | - | 7.02772403811355 | 11.3699574206438 | 8.35259747257593 | 13.8724124107028 |
| chr1 | 154259725 | 154259727 | + | 29.1148567293276 | 26.1509020674808 | 63.2410951495034 | 25.6106075274513 |
| chr1 | 154263125 | 154263204 | + | 90.3564519186028 | 59.1237785873479 | 39.3765309421437 | 59.7580842307196 |
| chr1 | 154263242 | 154263355 | + | 119.47130864793 | 48.8908169087684 | 44.1494437836156 | 39.483019938154 |
| chr1 | 154263375 | 154263381 | + | 314.239660561363 | 139.850476273919 | 88.2988875672312 | 133.388580872142 |
| chr1 | 154263434 | 154263439 | + | 3.01188173062009 | 2.27399148412876 | 4.77291284147196 | 1.06710864697714 |
| chr1 | 154270618 | 154270624 | + | 3.01188173062009 | 3.41098722619315 | 10.7390538933119 | 3.20132594093141 |
| chr1 | 154275778 | 154275820 | + | 59.2336740355285 | 87.5486721389574 | 46.5359002043516 | 46.952780466994 |
| chr1 | 154548600 | 154548604 | - | 2.00792115374673 | 4.54798296825753 | 1.19322821036799 | 1.06710864697714 |
| chr1 | 154550010 | 154550050 | - | 24.0950538449607 | 62.534765813541 | 19.0916513658878 | 22.4092815865198 |
| chr1 | 154550105 | 154550139 | - | 11.043566345607 | 6.82197445238629 | 4.77291284147196 | 4.26843458790854 |
| chr1 | 154550274 | 154550304 | - | 5.01980288436682 | 7.95897019445067 | 1.19322821036799 | 6.40265188186281 |
| chr1 | 154550348 | 154550407 | - | 5.01980288436682 | 4.54798296825753 | 1.19322821036799 | 3.20132594093141 |
| chr1 | 154582096 | 154582183 | - | 29.1148567293276 | 25.0139063254164 | 8.35259747257593 | 11.7381951167485 |
| chr1 | 154582204 | 154582215 | - | 13.0514874993537 | 7.95897019445067 | 5.96614105183995 | 9.60397782279422 |
| chr1 | 154582445 | 154582463 | - | 13.0514874993537 | 12.5069531627082 | 3.57968463110397 | 12.8053037637256 |
| chr1 | 154582582 | 154582622 | - | 16.0633692299738 | 9.09596593651506 | 13.1255103140479 | 23.476390233497 |
| chr1 | 154582644 | 154582736 | - | 28.1108961524542 | 25.0139063254164 | 19.0916513658878 | 8.53686917581709 |
| chr1 | 154582798 | 154582827 | - | 14.0554480762271 | 7.95897019445067 | 9.54582568294392 | 7.46976052883995 |
| chr1 | 154582844 | 154582848 | - | 6.02376346124019 | 1.13699574206438 | 4.77291284147196 | 1.06710864697714 |
| chr1 | 154582969 | 154582982 | - | 1.00396057687336 | 1.13699574206438 | 2.38645642073598 | 1.06710864697714 |
| chr1 | 154583042 | 154583053 | - | 9.03564519186028 | 3.41098722619315 | 4.77291284147196 | 4.26843458790854 |
| chr1 | 154583105 | 154583115 | - | 5.01980288436682 | 2.27399148412876 | 1.19322821036799 | 2.13421729395427 |
| chr1 | 154583181 | 154583209 | - | 7.02772403811355 | 7.95897019445067 | 8.35259747257593 | 9.60397782279422 |
| chr1 | 154583223 | 154583268 | - | 14.0554480762271 | 7.95897019445067 | 3.57968463110397 | 4.26843458790854 |
| chr1 | 154583288 | 154583295 | - | 5.01980288436682 | 1.13699574206438 | 1.19322821036799 | 6.40265188186281 |
| chr1 | 154583317 | 154583378 | - | 23.0910932680874 | 19.3289276150945 | 10.7390538933119 | 14.9395210576799 |
| chr1 | 154583521 | 154583587 | - | 17.0673298068472 | 11.3699574206438 | 19.0916513658878 | 11.7381951167485 |
| chr1 | 154583628 | 154583653 | - | 3.01188173062009 | 3.41098722619315 | 2.38645642073598 | 10.6710864697714 |
| chr1 | 154583739 | 154583790 | - | 12.0475269224804 | 18.1919318730301 | 7.15936926220794 | 5.33554323488568 |
| chr1 | 154583813 | 154583896 | - | 18.0712903837206 | 11.3699574206438 | 9.54582568294392 | 8.53686917581709 |
| chr1 | 154584040 | 154584093 | - | 11.043566345607 | 7.95897019445067 | 3.57968463110397 | 6.40265188186281 |
| chr1 | 154584250 | 154584284 | - | 12.0475269224804 | 6.82197445238629 | 5.96614105183995 | 3.20132594093141 |
| chr1 | 154584495 | 154584518 | - | 11.043566345607 | 5.68497871032191 | 5.96614105183995 | 1.06710864697714 |
| chr1 | 154584719 | 154584795 | - | 60.2376346124019 | 50.0278126508328 | 46.5359002043516 | 121.650385755393 |
| chr1 | 154924788 | 154924793 | - | 7.02772403811355 | 2.27399148412876 | 1.19322821036799 | 3.20132594093141 |
| chr1 | 154944106 | 154944127 | - | 11.043566345607 | 13.6439489047726 | 3.57968463110397 | 7.46976052883995 |
| chr1 | 154957067 | 154957076 | - | 21.0831721143407 | 47.753821166704 | 10.7390538933119 | 13.8724124107028 |
| chr1 | 154958565 | 154958590 | - | 6.02376346124019 | 4.54798296825753 | 4.77291284147196 | 3.20132594093141 |
| chr1 | 154958700 | 154958748 | - | 7.02772403811355 | 11.3699574206438 | 5.96614105183995 | 5.33554323488568 |
| chr1 | 154962381 | 154962411 | - | 14.0554480762271 | 14.780944646837 | 5.96614105183995 | 5.33554323488568 |
| chr1 | 154963447 | 154963449 | - | 1.00396057687336 | 3.41098722619315 | 1.19322821036799 | 1.06710864697714 |
| chr1 | 154979124 | 154979143 | + | 98.3881365335897 | 168.275369825529 | 79.9462900946553 | 80.0331485232852 |
| chr1 | 154990403 | 154990448 | + | 5.01980288436682 | 7.95897019445067 | 4.77291284147196 | 5.33554323488568 |
| chr1 | 154993053 | 154993078 | + | 7.02772403811355 | 17.0549361309657 | 8.35259747257593 | 12.8053037637256 |
| chr1 | 155017096 | 155017103 | + | 7.02772403811355 | 5.68497871032191 | 5.96614105183995 | 1.06710864697714 |
| chr1 | 155017197 | 155017213 | + | 10.0396057687336 | 7.95897019445067 | 4.77291284147196 | 3.20132594093141 |
| chr1 | 155017482 | 155017492 | + | 14.0554480762271 | 11.3699574206438 | 3.57968463110397 | 4.26843458790854 |
| chr1 | 155017588 | 155017611 | + | 8.03168461498692 | 2.27399148412876 | 2.38645642073598 | 4.26843458790854 |
| chr1 | 155017860 | 155017899 | + | 12.0475269224804 | 11.3699574206438 | 8.35259747257593 | 8.53686917581709 |
| chr1 | 155017930 | 155017935 | + | 3.01188173062009 | 2.27399148412876 | 9.54582568294392 | 20.2750642925656 |
| chr1 | 155018212 | 155018228 | + | 6.02376346124019 | 2.27399148412876 | 4.77291284147196 | 1.06710864697714 |
| chr1 | 155018483 | 155018500 | + | 11.043566345607 | 22.7399148412876 | 3.57968463110397 | 13.8724124107028 |
| chr1 | 155062664 | 155062725 | + | 76.3010038423757 | 70.4937360079917 | 7.15936926220794 | 26.6777161744284 |
| chr1 | 155069403 | 155069428 | + | 2.00792115374673 | 4.54798296825753 | 4.77291284147196 | 5.33554323488568 |
| chr1 | 155087439 | 155087464 | + | 1.00396057687336 | 3.41098722619315 | 1.19322821036799 | 1.06710864697714 |
| chr1 | 155134626 | 155134859 | + | 17.0673298068472 | 128.480518853275 | 1.19322821036799 | 2.13421729395427 |
| chr1 | 155138369 | 155138388 | + | 4.01584230749346 | 2.27399148412876 | 2.38645642073598 | 9.60397782279422 |
| chr1 | 155138452 | 155138461 | + | 4.01584230749346 | 5.68497871032191 | 1.19322821036799 | 7.46976052883995 |
| chr1 | 155138764 | 155138784 | + | 2.00792115374673 | 11.3699574206438 | 5.96614105183995 | 7.46976052883995 |
| chr1 | 155139915 | 155139942 | - | 22.087132691214 | 30.6988850357383 | 9.54582568294392 | 12.8053037637256 |
| chr1 | 155169421 | 155169500 | - | 503.988209590429 | 440.017352178916 | 343.649724585981 | 525.017454312751 |
| chr1 | 155169539 | 155169546 | - | 14.0554480762271 | 3.41098722619315 | 2.38645642073598 | 5.33554323488568 |
| chr1 | 155172564 | 155172629 | - | 12.0475269224804 | 9.09596593651506 | 9.54582568294392 | 9.60397782279422 |
| chr1 | 155172658 | 155172690 | - | 6.02376346124019 | 4.54798296825753 | 9.54582568294392 | 11.7381951167485 |
| chr1 | 155172740 | 155172832 | - | 81.3208067267425 | 54.5757956190903 | 38.1833027317757 | 42.6843458790854 |
| chr1 | 155172867 | 155172938 | - | 58.2297134586551 | 38.657855230189 | 31.0239334695677 | 55.4896496428111 |
| chr1 | 155173133 | 155173360 | - | 13232.2004031909 | 9120.97984284047 | 7100.90107989991 | 11566.3906245852 |
| chr1 | 155185870 | 155185872 | - | 11.043566345607 | 13.6439489047726 | 1.19322821036799 | 1.06710864697714 |
| chr1 | 155185899 | 155185900 | - | 6.02376346124019 | 2.27399148412876 | 1.19322821036799 | 2.13421729395427 |
| chr1 | 155188039 | 155188045 | - | 4.01584230749346 | 5.68497871032191 | 2.38645642073598 | 10.6710864697714 |
| chr1 | 155320572 | 155320641 | + | 579.285252855931 | 818.636934286355 | 585.875051290683 | 590.111081778356 |
| chr1 | 155330572 | 155330594 | + | 2.00792115374673 | 5.68497871032191 | 1.19322821036799 | 1.06710864697714 |
| chr1 | 155331050 | 155331071 | + | 8.03168461498692 | 21.6029190992233 | 8.35259747257593 | 10.6710864697714 |
| chr1 | 155335869 | 155335902 | - | 45.1782259593014 | 40.9318467143178 | 10.7390538933119 | 20.2750642925656 |
| chr1 | 155336575 | 155336592 | - | 23.0910932680874 | 20.4659233571589 | 4.77291284147196 | 5.33554323488568 |
| chr1 | 155336992 | 155337021 | - | 27.1069355755808 | 15.9179403889013 | 14.3187385244159 | 4.26843458790854 |
| chr1 | 155337543 | 155337589 | - | 33.130699036821 | 44.3428339405109 | 9.54582568294392 | 9.60397782279422 |
| chr1 | 155562939 | 155563031 | + | 32.1267384599477 | 25.0139063254164 | 21.4781077866238 | 37.3488026441998 |
| chr1 | 155563202 | 155563265 | + | 95.3762548029696 | 38.657855230189 | 133.641559561215 | 113.113516579576 |
| chr1 | 155614884 | 155614887 | + | 4.01584230749346 | 6.82197445238629 | 1.19322821036799 | 10.6710864697714 |
| chr1 | 155659495 | 155659538 | - | 9.03564519186028 | 12.5069531627082 | 3.57968463110397 | 7.46976052883995 |
| chr1 | 155659563 | 155659599 | - | 6.02376346124019 | 6.82197445238629 | 1.19322821036799 | 4.26843458790854 |
| chr1 | 155738332 | 155738450 | + | 12.0475269224804 | 23.876910583352 | 9.54582568294392 | 23.476390233497 |
| chr1 | 155738493 | 155738499 | + | 4.01584230749346 | 4.54798296825753 | 4.77291284147196 | 2.13421729395427 |
| chr1 | 155899684 | 155899709 | - | 1.00396057687336 | 1.13699574206438 | 2.38645642073598 | 5.33554323488568 |
| chr1 | 155899735 | 155899744 | - | 2.00792115374673 | 5.68497871032191 | 1.19322821036799 | 10.6710864697714 |
| chr1 | 155900279 | 155900282 | - | 3.01188173062009 | 5.68497871032191 | 3.57968463110397 | 5.33554323488568 |
| chr1 | 155913104 | 155913136 | - | 4.01584230749346 | 11.3699574206438 | 11.9322821036799 | 4.26843458790854 |
| chr1 | 155913164 | 155913220 | - | 19.0752509605939 | 12.5069531627082 | 13.1255103140479 | 9.60397782279422 |
| chr1 | 155913256 | 155913447 | - | 129.510914416664 | 133.028501821533 | 147.960298085631 | 102.442430109805 |
| chr1 | 155913471 | 155913519 | - | 28.1108961524542 | 21.6029190992233 | 28.6374770488317 | 18.1408469986113 |
| chr1 | 155913553 | 155913580 | - | 16.0633692299738 | 13.6439489047726 | 19.0916513658878 | 10.6710864697714 |
| chr1 | 155913686 | 155913721 | - | 25.0990144218341 | 29.5618892936739 | 51.3088130458235 | 21.3421729395427 |
| chr1 | 155914035 | 155914044 | - | 2.00792115374673 | 1.13699574206438 | 2.38645642073598 | 5.33554323488568 |
| chr1 | 155914185 | 155914201 | - | 8.03168461498692 | 3.41098722619315 | 5.96614105183995 | 5.33554323488568 |
| chr1 | 155914237 | 155914247 | - | 3.01188173062009 | 2.27399148412876 | 11.9322821036799 | 2.13421729395427 |
| chr1 | 155914342 | 155914356 | - | 11.043566345607 | 9.09596593651506 | 25.0577924177278 | 8.53686917581709 |
| chr1 | 155914590 | 155914627 | - | 21.0831721143407 | 20.4659233571589 | 62.0478669391355 | 14.9395210576799 |
| chr1 | 155914679 | 155914686 | - | 3.01188173062009 | 3.41098722619315 | 4.77291284147196 | 5.33554323488568 |
| chr1 | 155914719 | 155914733 | - | 4.01584230749346 | 4.54798296825753 | 8.35259747257593 | 2.13421729395427 |
| chr1 | 155914832 | 155914835 | - | 5.01980288436682 | 7.95897019445067 | 2.38645642073598 | 7.46976052883995 |
| chr1 | 155914868 | 155914884 | - | 4.01584230749346 | 2.27399148412876 | 9.54582568294392 | 1.06710864697714 |
| chr1 | 155915045 | 155915054 | - | 1.00396057687336 | 5.68497871032191 | 1.19322821036799 | 2.13421729395427 |
| chr1 | 155915249 | 155915271 | - | 5.01980288436682 | 4.54798296825753 | 3.57968463110397 | 1.06710864697714 |
| chr1 | 155946925 | 155946979 | - | 5.01980288436682 | 15.9179403889013 | 9.54582568294392 | 16.006629704657 |
| chr1 | 155947142 | 155947150 | - | 1.00396057687336 | 1.13699574206438 | 2.38645642073598 | 11.7381951167485 |
| chr1 | 155947278 | 155947305 | - | 9.03564519186028 | 17.0549361309657 | 9.54582568294392 | 39.483019938154 |
| chr1 | 155947443 | 155947506 | - | 8.03168461498692 | 9.09596593651506 | 13.1255103140479 | 67.2278447595595 |
| chr1 | 155947648 | 155947656 | - | 4.01584230749346 | 9.09596593651506 | 13.1255103140479 | 2.13421729395427 |
| chr1 | 155947669 | 155947732 | - | 2.00792115374673 | 5.68497871032191 | 17.8984231555198 | 16.006629704657 |
| chr1 | 155947775 | 155947815 | - | 7.02772403811355 | 22.7399148412876 | 13.1255103140479 | 5.33554323488568 |
| chr1 | 155947879 | 155947910 | - | 20.0792115374673 | 38.657855230189 | 22.6713359969918 | 11.7381951167485 |
| chr1 | 155950341 | 155950345 | - | 3.01188173062009 | 4.54798296825753 | 2.38645642073598 | 1.06710864697714 |
| chr1 | 156009076 | 156009164 | - | 58.2297134586551 | 64.8087572976698 | 38.1833027317757 | 36.2816939972226 |
| chr1 | 156009260 | 156009403 | - | 6.02376346124019 | 5.68497871032191 | 5.96614105183995 | 6.40265188186281 |
| chr1 | 156009457 | 156009480 | - | 7.02772403811355 | 4.54798296825753 | 2.38645642073598 | 2.13421729395427 |
| chr1 | 156009534 | 156009649 | - | 11.043566345607 | 4.54798296825753 | 3.57968463110397 | 4.26843458790854 |
| chr1 | 156058419 | 156058429 | + | 4.01584230749346 | 10.2329616785794 | 1.19322821036799 | 17.0737383516342 |
| chr1 | 156058456 | 156058485 | + | 18.0712903837206 | 44.3428339405109 | 16.7051949451519 | 14.9395210576799 |
| chr1 | 156072070 | 156072078 | - | 2.00792115374673 | 2.27399148412876 | 2.38645642073598 | 5.33554323488568 |
| chr1 | 156073766 | 156073796 | - | 2.00792115374673 | 2.27399148412876 | 45.3426719939836 | 17.0737383516342 |
| chr1 | 156073817 | 156073836 | - | 28.1108961524542 | 11.3699574206438 | 155.119667347839 | 103.509538756782 |
| chr1 | 156074440 | 156074487 | - | 6.02376346124019 | 2.27399148412876 | 13.1255103140479 | 6.40265188186281 |
| chr1 | 156074569 | 156074597 | - | 3.01188173062009 | 3.41098722619315 | 31.0239334695677 | 11.7381951167485 |
| chr1 | 156074653 | 156074664 | - | 1.00396057687336 | 1.13699574206438 | 7.15936926220794 | 5.33554323488568 |
| chr1 | 156137800 | 156137828 | + | 26.1029749987075 | 12.5069531627082 | 2.38645642073598 | 9.60397782279422 |
| chr1 | 156139125 | 156139140 | + | 185.732706721572 | 122.795540142953 | 23.8645642073598 | 28.8119334683827 |
| chr1 | 156139178 | 156139226 | + | 820.235791305539 | 550.305939159161 | 99.0379414605431 | 83.2344744642166 |
| chr1 | 156139252 | 156139318 | + | 37.1465413443145 | 20.4659233571589 | 9.54582568294392 | 10.6710864697714 |
| chr1 | 156139375 | 156139449 | + | 45.1782259593014 | 29.5618892936739 | 31.0239334695677 | 21.3421729395427 |
| chr1 | 156139490 | 156139539 | + | 33.130699036821 | 12.5069531627082 | 5.96614105183995 | 10.6710864697714 |
| chr1 | 156139583 | 156139599 | + | 21.0831721143407 | 6.82197445238629 | 1.19322821036799 | 1.06710864697714 |
| chr1 | 156139637 | 156139654 | + | 1.00396057687336 | 1.13699574206438 | 2.38645642073598 | 3.20132594093141 |
| chr1 | 156139695 | 156139916 | + | 745.94270861691 | 441.15434792098 | 273.24926017427 | 239.032336922878 |
| chr1 | 156139956 | 156140005 | + | 37.1465413443145 | 30.6988850357383 | 5.96614105183995 | 13.8724124107028 |
| chr1 | 156140052 | 156140058 | + | 43.1703048055547 | 40.9318467143178 | 5.96614105183995 | 10.6710864697714 |
| chr1 | 156210931 | 156210933 | + | 2.00792115374673 | 2.27399148412876 | 1.19322821036799 | 1.06710864697714 |
| chr1 | 156211018 | 156211036 | + | 5.01980288436682 | 9.09596593651506 | 1.19322821036799 | 1.06710864697714 |
| chr1 | 156212558 | 156212572 | + | 12.0475269224804 | 9.09596593651506 | 1.19322821036799 | 2.13421729395427 |
| chr1 | 156212662 | 156212681 | + | 2.00792115374673 | 6.82197445238629 | 2.38645642073598 | 1.06710864697714 |
| chr1 | 156239546 | 156239612 | + | 26.1029749987075 | 9.09596593651506 | 20.2848795762558 | 23.476390233497 |
| chr1 | 156239665 | 156239703 | + | 34.1346596136944 | 34.1098722619315 | 56.0817258872955 | 27.7448248214055 |
| chr1 | 156239723 | 156239876 | + | 106.419821148577 | 51.1648083928972 | 78.7530618842873 | 73.6304966414224 |
| chr1 | 156239985 | 156240015 | + | 20.0792115374673 | 28.4248935516096 | 14.3187385244159 | 9.60397782279422 |
| chr1 | 156243502 | 156243523 | - | 2.00792115374673 | 2.27399148412876 | 1.19322821036799 | 1.06710864697714 |
| chr1 | 156249512 | 156249611 | - | 31.1227778830743 | 47.753821166704 | 57.2749540976635 | 90.7042349930565 |
| chr1 | 156249640 | 156249643 | - | 1.00396057687336 | 1.13699574206438 | 2.38645642073598 | 67.2278447595595 |
| chr1 | 156249677 | 156249715 | - | 3.01188173062009 | 7.95897019445067 | 8.35259747257593 | 18.1408469986113 |
| chr1 | 156249739 | 156249780 | - | 9.03564519186028 | 14.780944646837 | 22.6713359969918 | 23.476390233497 |
| chr1 | 156249955 | 156249959 | - | 4.01584230749346 | 3.41098722619315 | 2.38645642073598 | 5.33554323488568 |
| chr1 | 156249984 | 156250064 | - | 7.02772403811355 | 15.9179403889013 | 19.0916513658878 | 36.2816939972226 |
| chr1 | 156250131 | 156250201 | - | 21.0831721143407 | 32.9728765198671 | 35.7968463110397 | 13.8724124107028 |
| chr1 | 156250271 | 156250300 | - | 3.01188173062009 | 3.41098722619315 | 13.1255103140479 | 7.46976052883995 |
| chr1 | 156250351 | 156250417 | - | 23.0910932680874 | 31.8358807778027 | 39.3765309421437 | 17.0737383516342 |
| chr1 | 156250465 | 156250474 | - | 4.01584230749346 | 4.54798296825753 | 4.77291284147196 | 9.60397782279422 |
| chr1 | 156292339 | 156292362 | + | 4.01584230749346 | 15.9179403889013 | 2.38645642073598 | 6.40265188186281 |
| chr1 | 156292715 | 156292777 | - | 10.0396057687336 | 7.95897019445067 | 4.77291284147196 | 5.33554323488568 |
| chr1 | 156308986 | 156309121 | - | 334.31887209883 | 327.454773714542 | 219.55399070771 | 200.616425631702 |
| chr1 | 156309158 | 156309303 | - | 36.1425807674411 | 13.6439489047726 | 17.8984231555198 | 26.6777161744284 |
| chr1 | 156463903 | 156463929 | - | 10.0396057687336 | 12.5069531627082 | 2.38645642073598 | 2.13421729395427 |
| chr1 | 156526341 | 156526346 | - | 3.01188173062009 | 3.41098722619315 | 2.38645642073598 | 2.13421729395427 |
| chr1 | 156594096 | 156594100 | + | 14.0554480762271 | 10.2329616785794 | 5.96614105183995 | 2.13421729395427 |
| chr1 | 156594233 | 156594256 | + | 6.02376346124019 | 10.2329616785794 | 3.57968463110397 | 2.13421729395427 |
| chr1 | 156594670 | 156594724 | - | 6.02376346124019 | 12.5069531627082 | 1.19322821036799 | 3.20132594093141 |
| chr1 | 156594870 | 156595836 | - | 3930.50565845922 | 3316.6165796018 | 3121.48499832266 | 2082.99607889937 |
| chr1 | 156723231 | 156723237 | - | 3.01188173062009 | 4.54798296825753 | 4.77291284147196 | 3.20132594093141 |
| chr1 | 156736803 | 156736812 | + | 1.00396057687336 | 3.41098722619315 | 2.38645642073598 | 8.53686917581709 |
| chr1 | 156736857 | 156736865 | + | 3.01188173062009 | 5.68497871032191 | 4.77291284147196 | 2.13421729395427 |
| chr1 | 156737343 | 156737360 | - | 23.0910932680874 | 31.8358807778027 | 21.4781077866238 | 32.0132594093141 |
| chr1 | 156737375 | 156737393 | - | 23.0910932680874 | 28.4248935516096 | 11.9322821036799 | 33.0803680562912 |
| chr1 | 156742132 | 156742235 | - | 88.3485307648561 | 108.014595496116 | 33.4103898903037 | 42.6843458790854 |
| chr1 | 156742257 | 156742304 | - | 65.2574374967687 | 57.9867828452835 | 25.0577924177278 | 43.7514545260626 |
| chr1 | 156742326 | 156742350 | - | 8.03168461498692 | 7.95897019445067 | 4.77291284147196 | 7.46976052883995 |
| chr1 | 156742371 | 156742457 | - | 139.550520185398 | 113.699574206438 | 20.2848795762558 | 49.0869977609482 |
| chr1 | 156742487 | 156742512 | - | 2.00792115374673 | 1.13699574206438 | 2.38645642073598 | 2.13421729395427 |
| chr1 | 156742561 | 156742584 | - | 13.0514874993537 | 10.2329616785794 | 5.96614105183995 | 8.53686917581709 |
| chr1 | 156742608 | 156742782 | - | 55.217831728035 | 36.3838637460602 | 10.7390538933119 | 56.5567582897882 |
| chr1 | 156742963 | 156743088 | - | 79.3128855729958 | 34.1098722619315 | 19.0916513658878 | 38.4159112911769 |
| chr1 | 156743118 | 156743151 | - | 7.02772403811355 | 11.3699574206438 | 4.77291284147196 | 4.26843458790854 |
| chr1 | 156743184 | 156743349 | - | 862.40213553422 | 731.088262147398 | 1162.20427689842 | 296.656203859644 |
| chr1 | 156743371 | 156743409 | - | 35.1386201905678 | 29.5618892936739 | 20.2848795762558 | 33.0803680562912 |
| chr1 | 156800687 | 156800742 | + | 16.0633692299738 | 5.68497871032191 | 1.19322821036799 | 6.40265188186281 |
| chr1 | 156800780 | 156800791 | + | 4.01584230749346 | 9.09596593651506 | 1.19322821036799 | 2.13421729395427 |
| chr1 | 156806377 | 156806480 | - | 10.0396057687336 | 20.4659233571589 | 52.5020412561915 | 61.8923015246739 |
| chr1 | 156934997 | 156935072 | - | 43.1703048055547 | 60.2607743294122 | 70.4004644117114 | 23.476390233497 |
| chr1 | 156935103 | 156935125 | - | 29.1148567293276 | 12.5069531627082 | 11.9322821036799 | 13.8724124107028 |
| chr1 | 156935220 | 156935226 | - | 2.00792115374673 | 2.27399148412876 | 1.19322821036799 | 3.20132594093141 |
| chr1 | 157121278 | 157121282 | - | 1.00396057687336 | 3.41098722619315 | 3.57968463110397 | 2.13421729395427 |
| chr1 | 157121576 | 157121578 | - | 14.0554480762271 | 17.0549361309657 | 1.19322821036799 | 2.13421729395427 |
| chr1 | 157121731 | 157121748 | - | 6.02376346124019 | 4.54798296825753 | 3.57968463110397 | 9.60397782279422 |
| chr1 | 157124280 | 157124320 | - | 59.2336740355285 | 50.0278126508328 | 28.6374770488317 | 24.5434988804741 |
| chr1 | 157124543 | 157124561 | - | 3.01188173062009 | 1.13699574206438 | 3.57968463110397 | 4.26843458790854 |
| chr1 | 157124603 | 157124637 | - | 8.03168461498692 | 4.54798296825753 | 5.96614105183995 | 13.8724124107028 |
| chr1 | 157124727 | 157124758 | - | 20.0792115374673 | 21.6029190992233 | 25.0577924177278 | 7.46976052883995 |
| chr1 | 159918142 | 159918168 | - | 237.938656718987 | 252.413054738293 | 299.500280802365 | 295.589095212667 |
| chr1 | 159918196 | 159918217 | - | 36.1425807674411 | 31.8358807778027 | 19.0916513658878 | 66.1607361125824 |
| chr1 | 159918432 | 159918530 | - | 95.3762548029696 | 52.3018041349616 | 193.302970079614 | 195.280882396816 |
| chr1 | 159918556 | 159918647 | - | 48.1901076899215 | 45.4798296825753 | 245.805011335806 | 226.227033159153 |
| chr1 | 159918667 | 159918727 | - | 166.657455760979 | 70.4937360079917 | 138.414472402687 | 275.314030920101 |
| chr1 | 159918765 | 159918900 | - | 68.2693192273888 | 35.2468680039958 | 136.028015981951 | 123.784603049348 |
| chr1 | 160213701 | 160213750 | + | 20.0792115374673 | 32.9728765198671 | 16.7051949451519 | 20.2750642925656 |
| chr1 | 160213875 | 160213893 | + | 3.01188173062009 | 4.54798296825753 | 7.15936926220794 | 4.26843458790854 |
| chr1 | 160214142 | 160214159 | + | 3.01188173062009 | 6.82197445238629 | 1.19322821036799 | 7.46976052883995 |
| chr1 | 160214193 | 160214225 | + | 46.1821865361748 | 36.3838637460602 | 48.9223566250876 | 37.3488026441998 |
| chr1 | 160214260 | 160214277 | + | 6.02376346124019 | 4.54798296825753 | 4.77291284147196 | 2.13421729395427 |
| chr1 | 160214691 | 160214747 | + | 47.1861471130481 | 57.9867828452835 | 28.6374770488317 | 33.0803680562912 |
| chr1 | 160214791 | 160214803 | + | 4.01584230749346 | 3.41098722619315 | 1.19322821036799 | 9.60397782279422 |
| chr1 | 160214825 | 160214845 | + | 12.0475269224804 | 9.09596593651506 | 3.57968463110397 | 11.7381951167485 |
| chr1 | 160215071 | 160215161 | + | 12.0475269224804 | 51.1648083928972 | 13.1255103140479 | 29.8790421153598 |
| chr1 | 160215319 | 160215353 | + | 10.0396057687336 | 17.0549361309657 | 9.54582568294392 | 10.6710864697714 |
| chr1 | 160215739 | 160215796 | - | 17.0673298068472 | 15.9179403889013 | 9.54582568294392 | 9.60397782279422 |
| chr1 | 160215930 | 160215983 | - | 36.1425807674411 | 29.5618892936739 | 39.3765309421437 | 58.6909755837425 |
| chr1 | 160216048 | 160216133 | - | 19.0752509605939 | 7.95897019445067 | 21.4781077866238 | 26.6777161744284 |
| chr1 | 160216209 | 160216215 | - | 1.00396057687336 | 3.41098722619315 | 16.7051949451519 | 8.53686917581709 |
| chr1 | 160216408 | 160216446 | - | 5.01980288436682 | 4.54798296825753 | 5.96614105183995 | 5.33554323488568 |
| chr1 | 160216499 | 160216517 | - | 1.00396057687336 | 1.13699574206438 | 5.96614105183995 | 6.40265188186281 |
| chr1 | 160216715 | 160216740 | - | 3.01188173062009 | 2.27399148412876 | 2.38645642073598 | 2.13421729395427 |
| chr1 | 160216815 | 160216818 | - | 1.00396057687336 | 2.27399148412876 | 1.19322821036799 | 2.13421729395427 |
| chr1 | 160216860 | 160216919 | - | 11.043566345607 | 13.6439489047726 | 25.0577924177278 | 20.2750642925656 |
| chr1 | 160217185 | 160217234 | - | 18.0712903837206 | 15.9179403889013 | 14.3187385244159 | 19.2079556455884 |
| chr1 | 160261717 | 160261760 | + | 10.0396057687336 | 19.3289276150945 | 10.7390538933119 | 8.53686917581709 |
| chr1 | 160277022 | 160277077 | - | 15.0594086531005 | 17.0549361309657 | 29.8307052591997 | 18.1408469986113 |
| chr1 | 160277098 | 160277159 | - | 55.217831728035 | 39.7948509722534 | 56.0817258872955 | 66.1607361125824 |
| chr1 | 160277184 | 160277222 | - | 9.03564519186028 | 6.82197445238629 | 7.15936926220794 | 5.33554323488568 |
| chr1 | 160277267 | 160277295 | - | 2.00792115374673 | 3.41098722619315 | 2.38645642073598 | 2.13421729395427 |
| chr1 | 160277315 | 160277333 | - | 7.02772403811355 | 6.82197445238629 | 1.19322821036799 | 3.20132594093141 |
| chr1 | 160277356 | 160277372 | - | 4.01584230749346 | 1.13699574206438 | 1.19322821036799 | 3.20132594093141 |
| chr1 | 160277480 | 160277498 | - | 3.01188173062009 | 3.41098722619315 | 3.57968463110397 | 1.06710864697714 |
| chr1 | 160289283 | 160289311 | - | 1.00396057687336 | 1.13699574206438 | 1.19322821036799 | 1.06710864697714 |
| chr1 | 160289334 | 160289358 | - | 7.02772403811355 | 21.6029190992233 | 17.8984231555198 | 20.2750642925656 |
| chr1 | 160289383 | 160289413 | - | 3.01188173062009 | 1.13699574206438 | 4.77291284147196 | 4.26843458790854 |
| chr1 | 160289458 | 160289473 | - | 3.01188173062009 | 5.68497871032191 | 4.77291284147196 | 5.33554323488568 |
| chr1 | 160289495 | 160289502 | - | 2.00792115374673 | 4.54798296825753 | 2.38645642073598 | 6.40265188186281 |
| chr1 | 160289671 | 160289685 | - | 16.0633692299738 | 21.6029190992233 | 17.8984231555198 | 12.8053037637256 |
| chr1 | 160290132 | 160290157 | - | 2.00792115374673 | 2.27399148412876 | 1.19322821036799 | 4.26843458790854 |
| chr1 | 160358796 | 160358863 | + | 65.2574374967687 | 45.4798296825753 | 20.2848795762558 | 23.476390233497 |
| chr1 | 160996260 | 160996268 | - | 1.00396057687336 | 1.13699574206438 | 7.15936926220794 | 3.20132594093141 |
| chr1 | 160996360 | 160996448 | - | 4.01584230749346 | 6.82197445238629 | 15.5119667347839 | 36.2816939972226 |
| chr1 | 160997295 | 160997349 | - | 2.00792115374673 | 2.27399148412876 | 4.77291284147196 | 6.40265188186281 |
| chr1 | 160997406 | 160997468 | - | 1.00396057687336 | 1.13699574206438 | 4.77291284147196 | 2.13421729395427 |
| chr1 | 160997748 | 160997795 | - | 1.00396057687336 | 3.41098722619315 | 4.77291284147196 | 1.06710864697714 |
| chr1 | 160998365 | 160998452 | - | 2.00792115374673 | 4.54798296825753 | 2.38645642073598 | 2.13421729395427 |
| chr1 | 160998546 | 160998573 | - | 1.00396057687336 | 4.54798296825753 | 26.2510206280958 | 14.9395210576799 |
| chr1 | 160998717 | 160998758 | - | 4.01584230749346 | 2.27399148412876 | 11.9322821036799 | 9.60397782279422 |
| chr1 | 160998785 | 160998829 | - | 3.01188173062009 | 3.41098722619315 | 11.9322821036799 | 16.006629704657 |
| chr1 | 160998854 | 160998909 | - | 3.01188173062009 | 2.27399148412876 | 7.15936926220794 | 7.46976052883995 |
| chr1 | 160999691 | 160999742 | - | 4.01584230749346 | 1.13699574206438 | 9.54582568294392 | 5.33554323488568 |
| chr1 | 160999918 | 160999949 | - | 1.00396057687336 | 5.68497871032191 | 14.3187385244159 | 4.26843458790854 |
| chr1 | 161000149 | 161000175 | - | 3.01188173062009 | 1.13699574206438 | 1.19322821036799 | 3.20132594093141 |
| chr1 | 161000224 | 161000233 | - | 2.00792115374673 | 1.13699574206438 | 3.57968463110397 | 1.06710864697714 |
| chr1 | 161000637 | 161000674 | - | 4.01584230749346 | 1.13699574206438 | 5.96614105183995 | 3.20132594093141 |
| chr1 | 161021093 | 161021207 | - | 730.883299963809 | 700.389377111659 | 956.969024715128 | 1189.82614137951 |
| chr1 | 161100600 | 161100621 | - | 3.01188173062009 | 14.780944646837 | 8.35259747257593 | 8.53686917581709 |
| chr1 | 161100632 | 161100727 | - | 145.574283646638 | 150.083437952498 | 106.197310722751 | 144.059667341913 |
| chr1 | 161120822 | 161120986 | + | 233.922814411494 | 378.619582107439 | 214.781077866238 | 264.64294445033 |
| chr1 | 161121022 | 161121042 | + | 14.0554480762271 | 22.7399148412876 | 5.96614105183995 | 18.1408469986113 |
| chr1 | 161121081 | 161121084 | - | 3.01188173062009 | 4.54798296825753 | 1.19322821036799 | 2.13421729395427 |
| chr1 | 161121166 | 161121168 | - | 1.00396057687336 | 1.13699574206438 | 1.19322821036799 | 2.13421729395427 |
| chr1 | 161121379 | 161121393 | - | 2.00792115374673 | 3.41098722619315 | 7.15936926220794 | 10.6710864697714 |
| chr1 | 161121712 | 161121762 | - | 7.02772403811355 | 7.95897019445067 | 1.19322821036799 | 2.13421729395427 |
| chr1 | 161158512 | 161158540 | + | 12.0475269224804 | 11.3699574206438 | 1.19322821036799 | 16.006629704657 |
| chr1 | 161158586 | 161158640 | + | 8.03168461498692 | 11.3699574206438 | 9.54582568294392 | 13.8724124107028 |
| chr1 | 161158695 | 161158800 | + | 226.89509037338 | 262.646016416872 | 163.472264820415 | 211.287512101473 |
| chr1 | 161158820 | 161158836 | + | 14.0554480762271 | 20.4659233571589 | 4.77291284147196 | 4.26843458790854 |
| chr1 | 161165554 | 161165568 | + | 19.0752509605939 | 14.780944646837 | 8.35259747257593 | 9.60397782279422 |
| chr1 | 161171347 | 161171356 | - | 8.03168461498692 | 11.3699574206438 | 10.7390538933119 | 16.006629704657 |
| chr1 | 161171715 | 161171735 | - | 4.01584230749346 | 1.13699574206438 | 4.77291284147196 | 18.1408469986113 |
| chr1 | 161214154 | 161214259 | + | 46.1821865361748 | 45.4798296825753 | 15.5119667347839 | 50.1541064079254 |
| chr1 | 161214284 | 161214373 | + | 31.1227778830743 | 46.6168254246397 | 19.0916513658878 | 37.3488026441998 |
| chr1 | 161230221 | 161230222 | + | 1.00396057687336 | 2.27399148412876 | 1.19322821036799 | 1.06710864697714 |
| chr1 | 161230311 | 161230328 | + | 2.00792115374673 | 3.41098722619315 | 4.77291284147196 | 4.26843458790854 |
| chr1 | 161305229 | 161305260 | - | 1.00396057687336 | 2.27399148412876 | 3.57968463110397 | 14.9395210576799 |
| chr1 | 161362781 | 161362827 | + | 10.0396057687336 | 3.41098722619315 | 3.57968463110397 | 7.46976052883995 |
| chr1 | 161363030 | 161363056 | + | 7.02772403811355 | 10.2329616785794 | 11.9322821036799 | 3.20132594093141 |
| chr1 | 161363316 | 161363319 | + | 1.00396057687336 | 1.13699574206438 | 2.38645642073598 | 3.20132594093141 |
| chr1 | 161363567 | 161363573 | + | 2.00792115374673 | 1.13699574206438 | 2.38645642073598 | 2.13421729395427 |
| chr1 | 161363575 | 161363598 | + | 6.02376346124019 | 1.13699574206438 | 1.19322821036799 | 8.53686917581709 |
| chr1 | 161364068 | 161364070 | + | 1.00396057687336 | 1.13699574206438 | 1.19322821036799 | 1.06710864697714 |
| chr1 | 161364232 | 161364237 | + | 39.1544624980612 | 34.1098722619315 | 14.3187385244159 | 27.7448248214055 |
| chr1 | 161365343 | 161365345 | + | 3.01188173062009 | 2.27399148412876 | 3.57968463110397 | 1.06710864697714 |
| chr1 | 161367923 | 161367942 | + | 3.01188173062009 | 1.13699574206438 | 5.96614105183995 | 4.26843458790854 |
| chr1 | 161369353 | 161369374 | + | 2.00792115374673 | 2.27399148412876 | 4.77291284147196 | 2.13421729395427 |
| chr1 | 161399357 | 161399427 | + | 8.03168461498692 | 10.2329616785794 | 3.57968463110397 | 3.20132594093141 |
| chr1 | 161399776 | 161399869 | + | 17.0673298068472 | 27.2878978095452 | 65.6275515702394 | 64.0265188186281 |
| chr1 | 161422164 | 161422197 | + | 11.043566345607 | 10.2329616785794 | 5.96614105183995 | 6.40265188186281 |
| chr1 | 161422206 | 161422253 | + | 9.03564519186028 | 5.68497871032191 | 11.9322821036799 | 2.13421729395427 |
| chr1 | 161757054 | 161757056 | + | 2.00792115374673 | 10.2329616785794 | 8.35259747257593 | 12.8053037637256 |
| chr1 | 161757082 | 161757085 | + | 7.02772403811355 | 13.6439489047726 | 1.19322821036799 | 5.33554323488568 |
| chr1 | 161958743 | 161958755 | + | 4.01584230749346 | 1.13699574206438 | 2.38645642073598 | 6.40265188186281 |
| chr1 | 161958858 | 161958889 | + | 7.02772403811355 | 3.41098722619315 | 2.38645642073598 | 9.60397782279422 |
| chr1 | 161958904 | 161958914 | + | 6.02376346124019 | 5.68497871032191 | 2.38645642073598 | 5.33554323488568 |
| chr1 | 161958963 | 161958980 | + | 4.01584230749346 | 2.27399148412876 | 2.38645642073598 | 1.06710864697714 |
| chr1 | 161959291 | 161959293 | + | 4.01584230749346 | 4.54798296825753 | 1.19322821036799 | 3.20132594093141 |
| chr1 | 161959333 | 161959352 | + | 1.00396057687336 | 6.82197445238629 | 3.57968463110397 | 2.13421729395427 |
| chr1 | 161960792 | 161960814 | + | 3.01188173062009 | 4.54798296825753 | 2.38645642073598 | 4.26843458790854 |
| chr1 | 161961363 | 161961417 | + | 11.043566345607 | 4.54798296825753 | 4.77291284147196 | 8.53686917581709 |
| chr1 | 161961519 | 161961566 | + | 7.02772403811355 | 4.54798296825753 | 11.9322821036799 | 13.8724124107028 |
| chr1 | 161961588 | 161961628 | + | 6.02376346124019 | 3.41098722619315 | 2.38645642073598 | 3.20132594093141 |
| chr1 | 161961818 | 161961852 | + | 1.00396057687336 | 3.41098722619315 | 2.38645642073598 | 6.40265188186281 |
| chr1 | 161963820 | 161963841 | + | 6.02376346124019 | 5.68497871032191 | 7.15936926220794 | 3.20132594093141 |
| chr1 | 162385148 | 162385149 | + | 2.00792115374673 | 4.54798296825753 | 1.19322821036799 | 1.06710864697714 |
| chr1 | 162522723 | 162522808 | + | 21.0831721143407 | 14.780944646837 | 31.0239334695677 | 5.33554323488568 |
| chr1 | 162522849 | 162522868 | + | 2.00792115374673 | 1.13699574206438 | 13.1255103140479 | 3.20132594093141 |
| chr1 | 162522909 | 162522932 | + | 3.01188173062009 | 4.54798296825753 | 8.35259747257593 | 1.06710864697714 |
| chr1 | 162523091 | 162523116 | + | 2.00792115374673 | 2.27399148412876 | 13.1255103140479 | 20.2750642925656 |
| chr1 | 162523161 | 162523168 | + | 1.00396057687336 | 1.13699574206438 | 2.38645642073598 | 1.06710864697714 |
| chr1 | 162523195 | 162523223 | + | 12.0475269224804 | 3.41098722619315 | 5.96614105183995 | 2.13421729395427 |
| chr1 | 162523244 | 162523267 | + | 6.02376346124019 | 3.41098722619315 | 1.19322821036799 | 1.06710864697714 |
| chr1 | 162523398 | 162523428 | + | 3.01188173062009 | 1.13699574206438 | 11.9322821036799 | 1.06710864697714 |
| chr1 | 162523564 | 162523572 | + | 4.01584230749346 | 4.54798296825753 | 4.77291284147196 | 4.26843458790854 |
| chr1 | 162523631 | 162523672 | + | 18.0712903837206 | 23.876910583352 | 20.2848795762558 | 7.46976052883995 |
| chr1 | 162523772 | 162523852 | + | 30.1188173062009 | 30.6988850357383 | 50.1155848354556 | 26.6777161744284 |
| chr1 | 162523906 | 162524000 | + | 44.174265382428 | 50.0278126508328 | 124.095733878271 | 87.5029090521251 |
| chr1 | 162524021 | 162524082 | + | 18.0712903837206 | 19.3289276150945 | 59.6614105183995 | 20.2750642925656 |
| chr1 | 162524321 | 162524355 | + | 14.0554480762271 | 12.5069531627082 | 13.1255103140479 | 17.0737383516342 |
| chr1 | 162524387 | 162524395 | + | 3.01188173062009 | 1.13699574206438 | 13.1255103140479 | 1.06710864697714 |
| chr1 | 162524423 | 162524428 | + | 1.00396057687336 | 3.41098722619315 | 4.77291284147196 | 19.2079556455884 |
| chr1 | 162524835 | 162524856 | + | 9.03564519186028 | 3.41098722619315 | 3.57968463110397 | 3.20132594093141 |
| chr1 | 162525531 | 162525536 | + | 14.0554480762271 | 17.0549361309657 | 4.77291284147196 | 7.46976052883995 |
| chr1 | 162526025 | 162526032 | + | 1.00396057687336 | 1.13699574206438 | 3.57968463110397 | 3.20132594093141 |
| chr1 | 162526110 | 162526118 | + | 7.02772403811355 | 1.13699574206438 | 2.38645642073598 | 2.13421729395427 |
| chr1 | 162526408 | 162526426 | + | 4.01584230749346 | 3.41098722619315 | 4.77291284147196 | 3.20132594093141 |
| chr1 | 162526542 | 162526561 | + | 3.01188173062009 | 2.27399148412876 | 3.57968463110397 | 4.26843458790854 |
| chr1 | 162527106 | 162527119 | + | 1.00396057687336 | 1.13699574206438 | 4.77291284147196 | 23.476390233497 |
| chr1 | 162527276 | 162527349 | + | 6.02376346124019 | 3.41098722619315 | 15.5119667347839 | 20.2750642925656 |
| chr1 | 162527592 | 162527615 | + | 1.00396057687336 | 2.27399148412876 | 4.77291284147196 | 5.33554323488568 |
| chr1 | 162527648 | 162527665 | + | 4.01584230749346 | 5.68497871032191 | 8.35259747257593 | 9.60397782279422 |
| chr1 | 162527698 | 162527732 | + | 5.01980288436682 | 3.41098722619315 | 7.15936926220794 | 9.60397782279422 |
| chr1 | 162527751 | 162527798 | + | 3.01188173062009 | 9.09596593651506 | 10.7390538933119 | 3.20132594093141 |
| chr1 | 162527897 | 162527918 | + | 12.0475269224804 | 10.2329616785794 | 9.54582568294392 | 5.33554323488568 |
| chr1 | 162527940 | 162527949 | + | 107.42378172545 | 63.6717615556054 | 44.1494437836156 | 43.7514545260626 |
| chr1 | 162528034 | 162528041 | + | 21.0831721143407 | 18.1919318730301 | 28.6374770488317 | 3.20132594093141 |
| chr1 | 162528085 | 162528123 | + | 12.0475269224804 | 10.2329616785794 | 21.4781077866238 | 19.2079556455884 |
| chr1 | 162528192 | 162528215 | + | 6.02376346124019 | 6.82197445238629 | 13.1255103140479 | 5.33554323488568 |
| chr1 | 162528240 | 162528293 | + | 17.0673298068472 | 15.9179403889013 | 35.7968463110397 | 11.7381951167485 |
| chr1 | 162528461 | 162528468 | + | 3.01188173062009 | 4.54798296825753 | 5.96614105183995 | 2.13421729395427 |
| chr1 | 162528505 | 162528507 | + | 1.00396057687336 | 3.41098722619315 | 2.38645642073598 | 1.06710864697714 |
| chr1 | 162528541 | 162528559 | + | 6.02376346124019 | 3.41098722619315 | 9.54582568294392 | 8.53686917581709 |
| chr1 | 162528585 | 162528641 | + | 9.03564519186028 | 20.4659233571589 | 70.4004644117114 | 40.5501285851312 |
| chr1 | 162528674 | 162528683 | + | 1.00396057687336 | 2.27399148412876 | 8.35259747257593 | 5.33554323488568 |
| chr1 | 162528711 | 162528722 | + | 2.00792115374673 | 2.27399148412876 | 5.96614105183995 | 3.20132594093141 |
| chr1 | 162528814 | 162528873 | + | 21.0831721143407 | 20.4659233571589 | 21.4781077866238 | 13.8724124107028 |
| chr1 | 162528907 | 162528962 | + | 58.2297134586551 | 47.753821166704 | 66.8207797806074 | 58.6909755837425 |
| chr1 | 162529020 | 162529121 | + | 265.045592294568 | 269.467990869259 | 161.085808399679 | 151.529427870753 |
| chr1 | 162529145 | 162529247 | + | 128.506953839791 | 146.672450726305 | 195.68942650035 | 102.442430109805 |
| chr1 | 162529280 | 162529319 | + | 198.784194220926 | 179.645327246172 | 402.117906894012 | 114.180625226554 |
| chr1 | 162529437 | 162529456 | + | 5.01980288436682 | 1.13699574206438 | 7.15936926220794 | 1.06710864697714 |
| chr1 | 162529541 | 162529548 | + | 1.00396057687336 | 1.13699574206438 | 4.77291284147196 | 1.06710864697714 |
| chr1 | 162529580 | 162529584 | + | 2.00792115374673 | 3.41098722619315 | 4.77291284147196 | 3.20132594093141 |
| chr1 | 162599566 | 162599574 | + | 2.00792115374673 | 3.41098722619315 | 7.15936926220794 | 5.33554323488568 |
| chr1 | 162599635 | 162599652 | + | 1.00396057687336 | 1.13699574206438 | 7.15936926220794 | 7.46976052883995 |
| chr1 | 162599733 | 162599744 | + | 3.01188173062009 | 2.27399148412876 | 11.9322821036799 | 3.20132594093141 |
| chr1 | 162599802 | 162599814 | + | 8.03168461498692 | 5.68497871032191 | 26.2510206280958 | 41.6172372321083 |
| chr1 | 162797877 | 162797881 | + | 1.00396057687336 | 3.41098722619315 | 2.38645642073598 | 4.26843458790854 |
| chr1 | 163355530 | 163355544 | + | 4.01584230749346 | 1.13699574206438 | 7.15936926220794 | 3.20132594093141 |
| chr1 | 163355667 | 163355734 | + | 18.0712903837206 | 22.7399148412876 | 19.0916513658878 | 7.46976052883995 |
| chr1 | 165655513 | 165655518 | + | 11.043566345607 | 25.0139063254164 | 3.57968463110397 | 1.06710864697714 |
| chr1 | 165655598 | 165655602 | + | 8.03168461498692 | 7.95897019445067 | 4.77291284147196 | 4.26843458790854 |
| chr1 | 165662246 | 165662263 | - | 6.02376346124019 | 12.5069531627082 | 3.57968463110397 | 1.06710864697714 |
| chr1 | 165662293 | 165662299 | - | 21.0831721143407 | 15.9179403889013 | 3.57968463110397 | 5.33554323488568 |
| chr1 | 165662329 | 165662360 | - | 121.479229801677 | 135.302493305661 | 36.9900745214077 | 34.1474767032683 |
| chr1 | 165662393 | 165662467 | - | 807.184303806185 | 819.773930028419 | 167.051949451519 | 142.992558694936 |
| chr1 | 165662492 | 165662543 | - | 32.1267384599477 | 20.4659233571589 | 10.7390538933119 | 13.8724124107028 |
| chr1 | 165662591 | 165662606 | - | 5.01980288436682 | 4.54798296825753 | 8.35259747257593 | 4.26843458790854 |
| chr1 | 165662760 | 165662774 | - | 6.02376346124019 | 9.09596593651506 | 1.19322821036799 | 1.06710864697714 |
| chr1 | 165662826 | 165662844 | - | 3.01188173062009 | 3.41098722619315 | 2.38645642073598 | 6.40265188186281 |
| chr1 | 165662871 | 165662885 | - | 4.01584230749346 | 1.13699574206438 | 1.19322821036799 | 1.06710864697714 |
| chr1 | 165662923 | 165662944 | - | 25.0990144218341 | 30.6988850357383 | 5.96614105183995 | 1.06710864697714 |
| chr1 | 165662981 | 165663023 | - | 22.087132691214 | 13.6439489047726 | 21.4781077866238 | 3.20132594093141 |
| chr1 | 165727040 | 165727045 | - | 4.01584230749346 | 2.27399148412876 | 2.38645642073598 | 1.06710864697714 |
| chr1 | 165727097 | 165727106 | - | 3.01188173062009 | 4.54798296825753 | 3.57968463110397 | 9.60397782279422 |
| chr1 | 165727309 | 165727333 | - | 12.0475269224804 | 15.9179403889013 | 11.9322821036799 | 6.40265188186281 |
| chr1 | 165727365 | 165727372 | - | 9.03564519186028 | 4.54798296825753 | 4.77291284147196 | 7.46976052883995 |
| chr1 | 165727472 | 165727484 | - | 6.02376346124019 | 12.5069531627082 | 4.77291284147196 | 14.9395210576799 |
| chr1 | 165727509 | 165727569 | - | 26.1029749987075 | 21.6029190992233 | 33.4103898903037 | 50.1541064079254 |
| chr1 | 165727624 | 165727725 | - | 32.1267384599477 | 27.2878978095452 | 34.6036181006717 | 45.8856718200168 |
| chr1 | 165727789 | 165727851 | - | 16.0633692299738 | 6.82197445238629 | 13.1255103140479 | 9.60397782279422 |
| chr1 | 165727921 | 165727970 | - | 9.03564519186028 | 3.41098722619315 | 5.96614105183995 | 10.6710864697714 |
| chr1 | 165728015 | 165728067 | - | 3.01188173062009 | 4.54798296825753 | 7.15936926220794 | 6.40265188186281 |
| chr1 | 165768901 | 165768961 | + | 18.0712903837206 | 30.6988850357383 | 27.4442488384638 | 30.9461507623369 |
| chr1 | 165769010 | 165769029 | + | 28.1108961524542 | 22.7399148412876 | 8.35259747257593 | 5.33554323488568 |
| chr1 | 165907862 | 165907880 | + | 150.594086531005 | 102.329616785794 | 59.6614105183995 | 91.7713436400337 |
| chr1 | 165907913 | 165907951 | + | 306.207975946376 | 237.632110091456 | 167.051949451519 | 306.260181682438 |
| chr1 | 165907981 | 165908087 | + | 56.2217923049084 | 81.8636934286355 | 35.7968463110397 | 38.4159112911769 |
| chr1 | 166851655 | 166851660 | + | 7.02772403811355 | 9.09596593651506 | 5.96614105183995 | 3.20132594093141 |
| chr1 | 166851774 | 166851790 | + | 2.00792115374673 | 2.27399148412876 | 2.38645642073598 | 3.20132594093141 |
| chr1 | 166852898 | 166852925 | + | 1.00396057687336 | 6.82197445238629 | 1.19322821036799 | 1.06710864697714 |
| chr1 | 166852973 | 166852990 | + | 1.00396057687336 | 4.54798296825753 | 1.19322821036799 | 1.06710864697714 |
| chr1 | 166853279 | 166853299 | + | 6.02376346124019 | 2.27399148412876 | 3.57968463110397 | 3.20132594093141 |
| chr1 | 166853377 | 166853393 | + | 4.01584230749346 | 1.13699574206438 | 3.57968463110397 | 2.13421729395427 |
| chr1 | 166853494 | 166853507 | + | 8.03168461498692 | 5.68497871032191 | 7.15936926220794 | 6.40265188186281 |
| chr1 | 166853557 | 166853563 | + | 4.01584230749346 | 2.27399148412876 | 1.19322821036799 | 1.06710864697714 |
| chr1 | 166853724 | 166853726 | + | 1.00396057687336 | 1.13699574206438 | 1.19322821036799 | 2.13421729395427 |
| chr1 | 166853750 | 166853759 | + | 1.00396057687336 | 1.13699574206438 | 1.19322821036799 | 5.33554323488568 |
| chr1 | 166853857 | 166853900 | + | 55.217831728035 | 70.4937360079917 | 36.9900745214077 | 50.1541064079254 |
| chr1 | 166853973 | 166854006 | + | 8.03168461498692 | 17.0549361309657 | 9.54582568294392 | 3.20132594093141 |
| chr1 | 166854041 | 166854059 | + | 9.03564519186028 | 4.54798296825753 | 9.54582568294392 | 2.13421729395427 |
| chr1 | 166854085 | 166854175 | + | 41.1623836518079 | 51.1648083928972 | 16.7051949451519 | 40.5501285851312 |
| chr1 | 166854206 | 166854260 | + | 69.2732798042621 | 78.4527062024424 | 22.6713359969918 | 14.9395210576799 |
| chr1 | 166854285 | 166854304 | + | 10.0396057687336 | 2.27399148412876 | 2.38645642073598 | 2.13421729395427 |
| chr1 | 166854428 | 166854441 | + | 5.01980288436682 | 3.41098722619315 | 3.57968463110397 | 5.33554323488568 |
| chr1 | 167415813 | 167415821 | + | 6.02376346124019 | 15.9179403889013 | 3.57968463110397 | 3.20132594093141 |
| chr1 | 167415921 | 167415932 | + | 2.00792115374673 | 4.54798296825753 | 10.7390538933119 | 3.20132594093141 |
| chr1 | 167415997 | 167416008 | + | 4.01584230749346 | 3.41098722619315 | 11.9322821036799 | 1.06710864697714 |
| chr1 | 167416057 | 167416068 | + | 3.01188173062009 | 2.27399148412876 | 2.38645642073598 | 1.06710864697714 |
| chr1 | 167417746 | 167417750 | + | 2.00792115374673 | 6.82197445238629 | 5.96614105183995 | 1.06710864697714 |
| chr1 | 167417839 | 167417850 | + | 7.02772403811355 | 4.54798296825753 | 4.77291284147196 | 4.26843458790854 |
| chr1 | 167418039 | 167418044 | + | 3.01188173062009 | 1.13699574206438 | 3.57968463110397 | 7.46976052883995 |
| chr1 | 167421915 | 167421922 | + | 96.380215379843 | 134.165497563597 | 91.8785721983352 | 27.7448248214055 |
| chr1 | 167423018 | 167423034 | + | 4.01584230749346 | 12.5069531627082 | 9.54582568294392 | 3.20132594093141 |
| chr1 | 167423303 | 167423305 | + | 3.01188173062009 | 2.27399148412876 | 1.19322821036799 | 1.06710864697714 |
| chr1 | 167424876 | 167424889 | + | 17.0673298068472 | 4.54798296825753 | 9.54582568294392 | 7.46976052883995 |
| chr1 | 167425208 | 167425215 | + | 10.0396057687336 | 10.2329616785794 | 15.5119667347839 | 9.60397782279422 |
| chr1 | 167426030 | 167426041 | + | 7.02772403811355 | 1.13699574206438 | 2.38645642073598 | 2.13421729395427 |
| chr1 | 167426636 | 167426639 | + | 1.00396057687336 | 1.13699574206438 | 3.57968463110397 | 7.46976052883995 |
| chr1 | 167426668 | 167426690 | + | 10.0396057687336 | 10.2329616785794 | 9.54582568294392 | 7.46976052883995 |
| chr1 | 167427087 | 167427112 | + | 67.2653586505154 | 80.7266976865711 | 33.4103898903037 | 52.2883237018796 |
| chr1 | 167427154 | 167427218 | + | 281.108961524542 | 211.481208023975 | 250.577924177278 | 287.05222603685 |
| chr1 | 167541044 | 167541063 | - | 14.0554480762271 | 20.4659233571589 | 4.77291284147196 | 5.33554323488568 |
| chr1 | 167541157 | 167541209 | - | 26.1029749987075 | 23.876910583352 | 50.1155848354556 | 21.3421729395427 |
| chr1 | 167541269 | 167541276 | - | 3.01188173062009 | 5.68497871032191 | 5.96614105183995 | 2.13421729395427 |
| chr1 | 167541308 | 167541317 | - | 10.0396057687336 | 15.9179403889013 | 7.15936926220794 | 22.4092815865198 |
| chr1 | 167541369 | 167541400 | - | 2.00792115374673 | 4.54798296825753 | 1.19322821036799 | 5.33554323488568 |
| chr1 | 167541422 | 167541474 | - | 127.502993262917 | 81.8636934286355 | 20.2848795762558 | 92.8384522870108 |
| chr1 | 167541529 | 167541560 | - | 7.02772403811355 | 5.68497871032191 | 2.38645642073598 | 17.0737383516342 |
| chr1 | 167541684 | 167541718 | - | 18.0712903837206 | 17.0549361309657 | 2.38645642073598 | 16.006629704657 |
| chr1 | 167541774 | 167541809 | - | 7.02772403811355 | 2.27399148412876 | 5.96614105183995 | 3.20132594093141 |
| chr1 | 167541984 | 167542003 | - | 6.02376346124019 | 9.09596593651506 | 2.38645642073598 | 19.2079556455884 |
| chr1 | 167542098 | 167542112 | - | 11.043566345607 | 4.54798296825753 | 4.77291284147196 | 18.1408469986113 |
| chr1 | 167542153 | 167542193 | - | 3.01188173062009 | 5.68497871032191 | 3.57968463110397 | 2.13421729395427 |
| chr1 | 167542224 | 167542251 | - | 4.01584230749346 | 1.13699574206438 | 1.19322821036799 | 4.26843458790854 |
| chr1 | 167788049 | 167788072 | + | 3.01188173062009 | 4.54798296825753 | 7.15936926220794 | 7.46976052883995 |
| chr1 | 167788107 | 167788137 | + | 8.03168461498692 | 2.27399148412876 | 4.77291284147196 | 6.40265188186281 |
| chr1 | 167788386 | 167788407 | + | 1.00396057687336 | 3.41098722619315 | 2.38645642073598 | 8.53686917581709 |
| chr1 | 167788460 | 167788472 | + | 2.00792115374673 | 1.13699574206438 | 1.19322821036799 | 4.26843458790854 |
| chr1 | 167788508 | 167788527 | + | 11.043566345607 | 6.82197445238629 | 3.57968463110397 | 3.20132594093141 |
| chr1 | 167788622 | 167788643 | + | 3.01188173062009 | 3.41098722619315 | 3.57968463110397 | 7.46976052883995 |
| chr1 | 167788697 | 167788703 | + | 2.00792115374673 | 1.13699574206438 | 1.19322821036799 | 5.33554323488568 |
| chr1 | 167788761 | 167788784 | + | 11.043566345607 | 11.3699574206438 | 4.77291284147196 | 6.40265188186281 |
| chr1 | 167788859 | 167788870 | + | 40.1584230749346 | 25.0139063254164 | 38.1833027317757 | 21.3421729395427 |
| chr1 | 167788968 | 167788975 | + | 3.01188173062009 | 43.2058381984465 | 7.15936926220794 | 5.33554323488568 |
| chr1 | 167789006 | 167789015 | + | 8.03168461498692 | 4.54798296825753 | 5.96614105183995 | 2.13421729395427 |
| chr1 | 167789088 | 167789118 | + | 11.043566345607 | 6.82197445238629 | 7.15936926220794 | 5.33554323488568 |
| chr1 | 167789218 | 167789266 | + | 44.174265382428 | 21.6029190992233 | 26.2510206280958 | 19.2079556455884 |
| chr1 | 167789319 | 167789395 | + | 133.526756724157 | 93.2336508492793 | 101.424397881279 | 108.845081991668 |
| chr1 | 167789469 | 167789526 | + | 14.0554480762271 | 15.9179403889013 | 20.2848795762558 | 10.6710864697714 |
| chr1 | 167789536 | 167789589 | + | 12.0475269224804 | 19.3289276150945 | 20.2848795762558 | 14.9395210576799 |
| chr1 | 167789597 | 167789620 | + | 2.00792115374673 | 6.82197445238629 | 5.96614105183995 | 3.20132594093141 |
| chr1 | 167789679 | 167789683 | + | 3.01188173062009 | 2.27399148412876 | 2.38645642073598 | 3.20132594093141 |
| chr1 | 167790095 | 167790098 | + | 7.02772403811355 | 2.27399148412876 | 2.38645642073598 | 6.40265188186281 |
| chr1 | 167790587 | 167790591 | + | 8.03168461498692 | 5.68497871032191 | 2.38645642073598 | 4.26843458790854 |
| chr1 | 167790628 | 167790663 | + | 10.0396057687336 | 6.82197445238629 | 5.96614105183995 | 3.20132594093141 |
| chr1 | 167790737 | 167790791 | + | 32.1267384599477 | 19.3289276150945 | 4.77291284147196 | 8.53686917581709 |
| chr1 | 167820358 | 167820419 | + | 9.03564519186028 | 9.09596593651506 | 11.9322821036799 | 32.0132594093141 |
| chr1 | 167918092 | 167918240 | - | 102.403978841083 | 158.042408146949 | 155.119667347839 | 166.468948928433 |
| chr1 | 167918255 | 167918335 | - | 29.1148567293276 | 51.1648083928972 | 59.6614105183995 | 48.0198891139711 |
| chr1 | 168075422 | 168075451 | + | 9.03564519186028 | 2.27399148412876 | 10.7390538933119 | 23.476390233497 |
| chr1 | 168075575 | 168075576 | + | 2.00792115374673 | 1.13699574206438 | 1.19322821036799 | 1.06710864697714 |
| chr1 | 168075712 | 168075715 | + | 2.00792115374673 | 4.54798296825753 | 7.15936926220794 | 3.20132594093141 |
| chr1 | 168200166 | 168200183 | + | 4.01584230749346 | 10.2329616785794 | 5.96614105183995 | 9.60397782279422 |
| chr1 | 168201944 | 168201953 | + | 15.0594086531005 | 15.9179403889013 | 5.96614105183995 | 8.53686917581709 |
| chr1 | 168202082 | 168202084 | + | 4.01584230749346 | 6.82197445238629 | 1.19322821036799 | 3.20132594093141 |
| chr1 | 168243253 | 168243270 | + | 2.00792115374673 | 5.68497871032191 | 1.19322821036799 | 1.06710864697714 |
| chr1 | 168243298 | 168243299 | + | 7.02772403811355 | 2.27399148412876 | 2.38645642073598 | 1.06710864697714 |
| chr1 | 168244155 | 168244160 | + | 6.02376346124019 | 2.27399148412876 | 1.19322821036799 | 1.06710864697714 |
| chr1 | 168244729 | 168244735 | + | 5.01980288436682 | 1.13699574206438 | 1.19322821036799 | 1.06710864697714 |
| chr1 | 168244811 | 168244850 | + | 26.1029749987075 | 13.6439489047726 | 13.1255103140479 | 10.6710864697714 |
| chr1 | 168244992 | 168244996 | + | 6.02376346124019 | 4.54798296825753 | 2.38645642073598 | 2.13421729395427 |
| chr1 | 168245097 | 168245126 | + | 47.1861471130481 | 38.657855230189 | 36.9900745214077 | 9.60397782279422 |
| chr1 | 168246986 | 168247005 | + | 5.01980288436682 | 4.54798296825753 | 7.15936926220794 | 5.33554323488568 |
| chr1 | 168251572 | 168251578 | + | 1.00396057687336 | 1.13699574206438 | 2.38645642073598 | 16.006629704657 |
| chr1 | 168251661 | 168251674 | + | 5.01980288436682 | 4.54798296825753 | 2.38645642073598 | 1.06710864697714 |
| chr1 | 168252479 | 168252502 | + | 7.02772403811355 | 4.54798296825753 | 3.57968463110397 | 4.26843458790854 |
| chr1 | 168252661 | 168252697 | + | 11.043566345607 | 7.95897019445067 | 3.57968463110397 | 9.60397782279422 |
| chr1 | 168252934 | 168252945 | + | 5.01980288436682 | 2.27399148412876 | 1.19322821036799 | 2.13421729395427 |
| chr1 | 169132440 | 169132464 | + | 12.0475269224804 | 14.780944646837 | 1.19322821036799 | 1.06710864697714 |
| chr1 | 169132581 | 169132583 | - | 4.01584230749346 | 4.54798296825753 | 3.57968463110397 | 1.06710864697714 |
| chr1 | 169388205 | 169388207 | + | 5.01980288436682 | 13.6439489047726 | 3.57968463110397 | 2.13421729395427 |
| chr1 | 169464026 | 169464074 | - | 5.01980288436682 | 10.2329616785794 | 4.77291284147196 | 7.46976052883995 |
| chr1 | 169464224 | 169464266 | - | 10.0396057687336 | 14.780944646837 | 2.38645642073598 | 3.20132594093141 |
| chr1 | 169464308 | 169464386 | - | 22.087132691214 | 62.534765813541 | 11.9322821036799 | 20.2750642925656 |
| chr1 | 169464419 | 169464478 | - | 4.01584230749346 | 7.95897019445067 | 1.19322821036799 | 1.06710864697714 |
| chr1 | 169486108 | 169486194 | + | 40.1584230749346 | 67.0827487817985 | 63.2410951495034 | 94.9726695809651 |
| chr1 | 169849957 | 169849976 | - | 1.00396057687336 | 7.95897019445067 | 2.38645642073598 | 1.06710864697714 |
| chr1 | 171532341 | 171532904 | + | 2848.23615658973 | 3539.46774504642 | 6320.52983031924 | 4561.88946582725 |
| chr1 | 171591844 | 171591895 | + | 65.2574374967687 | 77.315710460378 | 109.776995353855 | 112.046407932599 |
| chr1 | 171591927 | 171591944 | + | 12.0475269224804 | 21.6029190992233 | 26.2510206280958 | 26.6777161744284 |
| chr1 | 171591966 | 171591987 | + | 10.0396057687336 | 3.41098722619315 | 4.77291284147196 | 13.8724124107028 |
| chr1 | 171592019 | 171592076 | + | 10.0396057687336 | 12.5069531627082 | 17.8984231555198 | 11.7381951167485 |
| chr1 | 171592184 | 171592255 | + | 16.0633692299738 | 10.2329616785794 | 44.1494437836156 | 93.9055609339879 |
| chr1 | 171592283 | 171592391 | + | 48.1901076899215 | 47.753821166704 | 103.810854302015 | 76.8318225823538 |
| chr1 | 171592423 | 171592477 | + | 22.087132691214 | 14.780944646837 | 29.8307052591997 | 19.2079556455884 |
| chr1 | 171592533 | 171592568 | + | 7.02772403811355 | 12.5069531627082 | 22.6713359969918 | 23.476390233497 |
| chr1 | 171592593 | 171592699 | + | 48.1901076899215 | 35.2468680039958 | 76.3666054635513 | 82.1673658172394 |
| chr1 | 171592730 | 171592769 | + | 12.0475269224804 | 5.68497871032191 | 38.1833027317757 | 20.2750642925656 |
| chr1 | 171592962 | 171593007 | + | 2.00792115374673 | 3.41098722619315 | 5.96614105183995 | 14.9395210576799 |
| chr1 | 171593153 | 171593159 | + | 11.043566345607 | 10.2329616785794 | 1.19322821036799 | 20.2750642925656 |
| chr1 | 171593378 | 171593398 | + | 6.02376346124019 | 6.82197445238629 | 7.15936926220794 | 2.13421729395427 |
| chr1 | 171593427 | 171593460 | + | 6.02376346124019 | 26.1509020674808 | 14.3187385244159 | 30.9461507623369 |
| chr1 | 171700591 | 171700613 | - | 1.00396057687336 | 5.68497871032191 | 1.19322821036799 | 1.06710864697714 |
| chr1 | 171796812 | 171796845 | + | 1.00396057687336 | 3.41098722619315 | 2.38645642073598 | 3.20132594093141 |
| chr1 | 171796897 | 171796934 | + | 5.01980288436682 | 6.82197445238629 | 4.77291284147196 | 11.7381951167485 |
| chr1 | 172441483 | 172441525 | - | 2.00792115374673 | 13.6439489047726 | 5.96614105183995 | 8.53686917581709 |
| chr1 | 172441566 | 172441576 | - | 5.01980288436682 | 4.54798296825753 | 4.77291284147196 | 10.6710864697714 |
| chr1 | 172611285 | 172611309 | + | 3.01188173062009 | 2.27399148412876 | 2.38645642073598 | 5.33554323488568 |
| chr1 | 172611650 | 172611663 | + | 10.0396057687336 | 9.09596593651506 | 4.77291284147196 | 9.60397782279422 |
| chr1 | 173487859 | 173488025 | + | 210.831721143407 | 206.933225055718 | 207.62170860403 | 179.274252692159 |
| chr1 | 173488299 | 173488361 | + | 12.0475269224804 | 9.09596593651506 | 9.54582568294392 | 18.1408469986113 |
| chr1 | 173488431 | 173488439 | + | 6.02376346124019 | 5.68497871032191 | 5.96614105183995 | 5.33554323488568 |
| chr1 | 173488548 | 173488704 | + | 83.3287278804893 | 48.8908169087684 | 38.1833027317757 | 69.3620620535138 |
| chr1 | 173488728 | 173488752 | + | 59.2336740355285 | 81.8636934286355 | 36.9900745214077 | 44.8185631730397 |
| chr1 | 173488774 | 173488778 | + | 25.0990144218341 | 38.657855230189 | 15.5119667347839 | 17.0737383516342 |
| chr1 | 173785285 | 173785306 | + | 24.0950538449607 | 25.0139063254164 | 10.7390538933119 | 10.6710864697714 |
| chr1 | 173799685 | 173799689 | - | 3.01188173062009 | 9.09596593651506 | 3.57968463110397 | 5.33554323488568 |
| chr1 | 173857813 | 173857832 | + | 2.00792115374673 | 2.27399148412876 | 3.57968463110397 | 10.6710864697714 |
| chr1 | 173858105 | 173858109 | + | 4.01584230749346 | 9.09596593651506 | 2.38645642073598 | 1.06710864697714 |
| chr1 | 173858191 | 173858194 | + | 5.01980288436682 | 1.13699574206438 | 3.57968463110397 | 7.46976052883995 |
| chr1 | 173858348 | 173858351 | + | 4.01584230749346 | 2.27399148412876 | 2.38645642073598 | 6.40265188186281 |
| chr1 | 173863918 | 173863922 | + | 2.00792115374673 | 11.3699574206438 | 4.77291284147196 | 1.06710864697714 |
| chr1 | 173903272 | 173903294 | + | 1.00396057687336 | 2.27399148412876 | 7.15936926220794 | 4.26843458790854 |
| chr1 | 173934003 | 173934024 | - | 4.01584230749346 | 5.68497871032191 | 4.77291284147196 | 1.06710864697714 |
| chr1 | 173935178 | 173935180 | - | 2.00792115374673 | 3.41098722619315 | 1.19322821036799 | 2.13421729395427 |
| chr1 | 173935372 | 173935380 | - | 2.00792115374673 | 1.13699574206438 | 4.77291284147196 | 2.13421729395427 |
| chr1 | 173935645 | 173935689 | - | 8.03168461498692 | 2.27399148412876 | 4.77291284147196 | 18.1408469986113 |
| chr1 | 173935748 | 173935791 | - | 1.00396057687336 | 4.54798296825753 | 3.57968463110397 | 20.2750642925656 |
| chr1 | 173936839 | 173936854 | - | 1.00396057687336 | 2.27399148412876 | 2.38645642073598 | 6.40265188186281 |
| chr1 | 173937440 | 173937445 | - | 4.01584230749346 | 7.95897019445067 | 2.38645642073598 | 3.20132594093141 |
| chr1 | 173937677 | 173937728 | - | 3.01188173062009 | 12.5069531627082 | 4.77291284147196 | 1.06710864697714 |
| chr1 | 173938664 | 173938666 | - | 1.00396057687336 | 2.27399148412876 | 4.77291284147196 | 5.33554323488568 |
| chr1 | 174991753 | 174991760 | + | 4.01584230749346 | 4.54798296825753 | 15.5119667347839 | 18.1408469986113 |
| chr1 | 174992525 | 174992528 | + | 4.01584230749346 | 5.68497871032191 | 10.7390538933119 | 9.60397782279422 |
| chr1 | 174993704 | 174993705 | + | 1.00396057687336 | 1.13699574206438 | 1.19322821036799 | 1.06710864697714 |
| chr1 | 175010204 | 175010206 | + | 1.00396057687336 | 3.41098722619315 | 8.35259747257593 | 5.33554323488568 |
| chr1 | 175010313 | 175010352 | + | 7.02772403811355 | 7.95897019445067 | 4.77291284147196 | 4.26843458790854 |
| chr1 | 175010395 | 175010407 | + | 3.01188173062009 | 1.13699574206438 | 1.19322821036799 | 1.06710864697714 |
| chr1 | 175010519 | 175010520 | + | 4.01584230749346 | 1.13699574206438 | 1.19322821036799 | 1.06710864697714 |
| chr1 | 175010705 | 175010706 | + | 4.01584230749346 | 4.54798296825753 | 4.77291284147196 | 2.13421729395427 |
| chr1 | 175010738 | 175010771 | + | 8.03168461498692 | 5.68497871032191 | 4.77291284147196 | 4.26843458790854 |
| chr1 | 175013555 | 175013559 | - | 6.02376346124019 | 4.54798296825753 | 4.77291284147196 | 2.13421729395427 |
| chr1 | 175013588 | 175013615 | - | 1.00396057687336 | 2.27399148412876 | 4.77291284147196 | 7.46976052883995 |
| chr1 | 175013843 | 175013858 | - | 36.1425807674411 | 36.3838637460602 | 31.0239334695677 | 32.0132594093141 |
| chr1 | 175014427 | 175014442 | - | 6.02376346124019 | 10.2329616785794 | 4.77291284147196 | 9.60397782279422 |
| chr1 | 175014541 | 175014572 | - | 2.00792115374673 | 4.54798296825753 | 10.7390538933119 | 12.8053037637256 |
| chr1 | 175945039 | 175945044 | - | 1.00396057687336 | 2.27399148412876 | 2.38645642073598 | 1.06710864697714 |
| chr1 | 176207661 | 176207712 | + | 8.03168461498692 | 13.6439489047726 | 34.6036181006717 | 20.2750642925656 |
| chr1 | 178916653 | 178916678 | + | 3.01188173062009 | 9.09596593651506 | 1.19322821036799 | 1.06710864697714 |
| chr1 | 178916738 | 178916744 | + | 13.0514874993537 | 28.4248935516096 | 5.96614105183995 | 1.06710864697714 |
| chr1 | 178916956 | 178916979 | + | 10.0396057687336 | 9.09596593651506 | 8.35259747257593 | 2.13421729395427 |
| chr1 | 178917001 | 178917004 | + | 19.0752509605939 | 7.95897019445067 | 2.38645642073598 | 3.20132594093141 |
| chr1 | 178918027 | 178918038 | + | 1.00396057687336 | 1.13699574206438 | 5.96614105183995 | 2.13421729395427 |
| chr1 | 178918140 | 178918174 | + | 10.0396057687336 | 11.3699574206438 | 15.5119667347839 | 6.40265188186281 |
| chr1 | 178919556 | 178919602 | + | 12.0475269224804 | 3.41098722619315 | 3.57968463110397 | 3.20132594093141 |
| chr1 | 178919828 | 178919856 | + | 44.174265382428 | 43.2058381984465 | 36.9900745214077 | 19.2079556455884 |
| chr1 | 178919903 | 178919933 | + | 7.02772403811355 | 18.1919318730301 | 10.7390538933119 | 1.06710864697714 |
| chr1 | 178919993 | 178920005 | + | 1.00396057687336 | 2.27399148412876 | 1.19322821036799 | 2.13421729395427 |
| chr1 | 179072256 | 179072267 | + | 1.00396057687336 | 2.27399148412876 | 7.15936926220794 | 6.40265188186281 |
| chr1 | 179072343 | 179072547 | + | 212.839642297153 | 218.303182476361 | 249.38469596691 | 231.562576394038 |
| chr1 | 179072607 | 179072616 | + | 3.01188173062009 | 2.27399148412876 | 3.57968463110397 | 5.33554323488568 |
| chr1 | 179072662 | 179072716 | + | 20.0792115374673 | 15.9179403889013 | 26.2510206280958 | 28.8119334683827 |
| chr1 | 179073650 | 179073670 | + | 2.00792115374673 | 2.27399148412876 | 7.15936926220794 | 6.40265188186281 |
| chr1 | 179073935 | 179073942 | + | 7.02772403811355 | 1.13699574206438 | 3.57968463110397 | 5.33554323488568 |
| chr1 | 179073975 | 179073996 | + | 1.00396057687336 | 1.13699574206438 | 11.9322821036799 | 3.20132594093141 |
| chr1 | 179075202 | 179075208 | + | 1.00396057687336 | 2.27399148412876 | 1.19322821036799 | 1.06710864697714 |
| chr1 | 179075511 | 179075513 | + | 4.01584230749346 | 3.41098722619315 | 3.57968463110397 | 4.26843458790854 |
| chr1 | 179076268 | 179076291 | + | 6.02376346124019 | 3.41098722619315 | 1.19322821036799 | 22.4092815865198 |
| chr1 | 179076334 | 179076395 | + | 5.01980288436682 | 5.68497871032191 | 17.8984231555198 | 22.4092815865198 |
| chr1 | 179095820 | 179095913 | + | 30.1188173062009 | 57.9867828452835 | 7.15936926220794 | 35.2145853502455 |
| chr1 | 179095934 | 179095966 | + | 46.1821865361748 | 106.877599754052 | 26.2510206280958 | 24.5434988804741 |
| chr1 | 179099648 | 179099660 | - | 1.00396057687336 | 1.13699574206438 | 1.19322821036799 | 3.20132594093141 |
| chr1 | 179101830 | 179101848 | - | 5.01980288436682 | 6.82197445238629 | 2.38645642073598 | 1.06710864697714 |
| chr1 | 179102656 | 179102694 | - | 1.00396057687336 | 3.41098722619315 | 1.19322821036799 | 5.33554323488568 |
| chr1 | 179102939 | 179102943 | - | 2.00792115374673 | 3.41098722619315 | 1.19322821036799 | 1.06710864697714 |
| chr1 | 179353865 | 179353933 | + | 69.2732798042621 | 57.9867828452835 | 3.57968463110397 | 8.53686917581709 |
| chr1 | 179354071 | 179354120 | + | 15.0594086531005 | 10.2329616785794 | 1.19322821036799 | 1.06710864697714 |
| chr1 | 179354931 | 179354967 | + | 61.2415951892752 | 43.2058381984465 | 1.19322821036799 | 4.26843458790854 |
| chr1 | 179355134 | 179355173 | + | 15.0594086531005 | 21.6029190992233 | 4.77291284147196 | 1.06710864697714 |
| chr1 | 179357166 | 179357168 | + | 9.03564519186028 | 3.41098722619315 | 1.19322821036799 | 4.26843458790854 |
| chr1 | 179357970 | 179357974 | + | 21.0831721143407 | 19.3289276150945 | 2.38645642073598 | 1.06710864697714 |
| chr1 | 179358036 | 179358113 | + | 33.130699036821 | 31.8358807778027 | 1.19322821036799 | 6.40265188186281 |
| chr1 | 179358137 | 179358197 | + | 54.2138711511617 | 56.8497871032191 | 2.38645642073598 | 1.06710864697714 |
| chr1 | 179358633 | 179358651 | + | 31.1227778830743 | 37.5208594881246 | 1.19322821036799 | 3.20132594093141 |
| chr1 | 179841150 | 179841190 | - | 5.01980288436682 | 3.41098722619315 | 3.57968463110397 | 4.26843458790854 |
| chr1 | 179842448 | 179842449 | - | 3.01188173062009 | 6.82197445238629 | 4.77291284147196 | 4.26843458790854 |
| chr1 | 179842749 | 179842781 | - | 14.0554480762271 | 35.2468680039958 | 9.54582568294392 | 7.46976052883995 |
| chr1 | 179844166 | 179844190 | - | 8.03168461498692 | 5.68497871032191 | 21.4781077866238 | 8.53686917581709 |
| chr1 | 179844302 | 179844328 | - | 15.0594086531005 | 9.09596593651506 | 17.8984231555198 | 19.2079556455884 |
| chr1 | 179844373 | 179844376 | - | 1.00396057687336 | 1.13699574206438 | 2.38645642073598 | 6.40265188186281 |
| chr1 | 179844406 | 179844437 | - | 7.02772403811355 | 5.68497871032191 | 4.77291284147196 | 1.06710864697714 |
| chr1 | 179844519 | 179844521 | - | 1.00396057687336 | 2.27399148412876 | 1.19322821036799 | 1.06710864697714 |
| chr1 | 179844582 | 179844603 | - | 11.043566345607 | 2.27399148412876 | 5.96614105183995 | 2.13421729395427 |
| chr1 | 179844634 | 179844730 | - | 28.1108961524542 | 21.6029190992233 | 14.3187385244159 | 19.2079556455884 |
| chr1 | 179844897 | 179844960 | - | 12.0475269224804 | 19.3289276150945 | 25.0577924177278 | 14.9395210576799 |
| chr1 | 179845275 | 179845278 | - | 2.00792115374673 | 1.13699574206438 | 1.19322821036799 | 3.20132594093141 |
| chr1 | 179845737 | 179845781 | - | 18.0712903837206 | 27.2878978095452 | 20.2848795762558 | 108.845081991668 |
| chr1 | 179845822 | 179845838 | - | 2.00792115374673 | 2.27399148412876 | 1.19322821036799 | 6.40265188186281 |
| chr1 | 179845878 | 179845883 | - | 4.01584230749346 | 3.41098722619315 | 4.77291284147196 | 21.3421729395427 |
| chr1 | 179845964 | 179845997 | - | 20.0792115374673 | 15.9179403889013 | 2.38645642073598 | 9.60397782279422 |
| chr1 | 179862484 | 179862535 | - | 7.02772403811355 | 10.2329616785794 | 8.35259747257593 | 7.46976052883995 |
| chr1 | 179863536 | 179863569 | - | 13.0514874993537 | 9.09596593651506 | 4.77291284147196 | 2.13421729395427 |
| chr1 | 179863739 | 179863749 | - | 1.00396057687336 | 3.41098722619315 | 9.54582568294392 | 28.8119334683827 |
| chr1 | 179863826 | 179863838 | - | 2.00792115374673 | 1.13699574206438 | 1.19322821036799 | 5.33554323488568 |
| chr1 | 179864005 | 179864041 | - | 8.03168461498692 | 13.6439489047726 | 8.35259747257593 | 4.26843458790854 |
| chr1 | 179864432 | 179864462 | - | 11.043566345607 | 30.6988850357383 | 4.77291284147196 | 10.6710864697714 |
| chr1 | 179864519 | 179864551 | - | 5.01980288436682 | 4.54798296825753 | 4.77291284147196 | 1.06710864697714 |
| chr1 | 179864671 | 179864690 | - | 5.01980288436682 | 3.41098722619315 | 1.19322821036799 | 1.06710864697714 |
| chr1 | 179864719 | 179864723 | - | 3.01188173062009 | 4.54798296825753 | 1.19322821036799 | 3.20132594093141 |
| chr1 | 179918508 | 179918547 | + | 11.043566345607 | 10.2329616785794 | 4.77291284147196 | 12.8053037637256 |
| chr1 | 179918642 | 179918647 | + | 2.00792115374673 | 5.68497871032191 | 1.19322821036799 | 19.2079556455884 |
| chr1 | 179918681 | 179918686 | + | 5.01980288436682 | 6.82197445238629 | 1.19322821036799 | 1.06710864697714 |
| chr1 | 179918707 | 179918743 | + | 11.043566345607 | 18.1919318730301 | 32.2171616799357 | 84.3015831111937 |
| chr1 | 180112910 | 180112911 | + | 1.00396057687336 | 1.13699574206438 | 3.57968463110397 | 1.06710864697714 |
| chr1 | 180112944 | 180112962 | + | 6.02376346124019 | 6.82197445238629 | 8.35259747257593 | 2.13421729395427 |
| chr1 | 180113004 | 180113011 | + | 8.03168461498692 | 3.41098722619315 | 4.77291284147196 | 1.06710864697714 |
| chr1 | 180114744 | 180114750 | + | 1.00396057687336 | 10.2329616785794 | 1.19322821036799 | 3.20132594093141 |
| chr1 | 180114780 | 180114792 | + | 4.01584230749346 | 1.13699574206438 | 1.19322821036799 | 2.13421729395427 |
| chr1 | 180197252 | 180197285 | + | 3.01188173062009 | 3.41098722619315 | 10.7390538933119 | 6.40265188186281 |
| chr1 | 180197294 | 180197348 | + | 1.00396057687336 | 1.13699574206438 | 8.35259747257593 | 2.13421729395427 |
| chr1 | 180197942 | 180198011 | + | 2.00792115374673 | 5.68497871032191 | 9.54582568294392 | 7.46976052883995 |
| chr1 | 180288259 | 180288277 | - | 3.01188173062009 | 5.68497871032191 | 2.38645642073598 | 2.13421729395427 |
| chr1 | 180884200 | 180884223 | + | 1.00396057687336 | 3.41098722619315 | 9.54582568294392 | 8.53686917581709 |
| chr1 | 180884501 | 180884505 | + | 1.00396057687336 | 1.13699574206438 | 4.77291284147196 | 2.13421729395427 |
| chr1 | 180884646 | 180884662 | + | 2.00792115374673 | 19.3289276150945 | 22.6713359969918 | 6.40265188186281 |
| chr1 | 180884739 | 180884795 | + | 17.0673298068472 | 12.5069531627082 | 63.2410951495034 | 54.4225409958339 |
| chr1 | 180884816 | 180884839 | + | 6.02376346124019 | 9.09596593651506 | 19.0916513658878 | 38.4159112911769 |
| chr1 | 180884961 | 180884981 | + | 1.00396057687336 | 5.68497871032191 | 1.19322821036799 | 4.26843458790854 |
| chr1 | 180885121 | 180885148 | + | 2.00792115374673 | 1.13699574206438 | 14.3187385244159 | 2.13421729395427 |
| chr1 | 180885226 | 180885228 | + | 5.01980288436682 | 10.2329616785794 | 9.54582568294392 | 8.53686917581709 |
| chr1 | 180885250 | 180885251 | + | 4.01584230749346 | 1.13699574206438 | 2.38645642073598 | 1.06710864697714 |
| chr1 | 180885733 | 180885740 | + | 5.01980288436682 | 6.82197445238629 | 4.77291284147196 | 9.60397782279422 |
| chr1 | 180885950 | 180885967 | + | 1.00396057687336 | 3.41098722619315 | 3.57968463110397 | 3.20132594093141 |
| chr1 | 180886067 | 180886074 | + | 5.01980288436682 | 11.3699574206438 | 15.5119667347839 | 14.9395210576799 |
| chr1 | 180886157 | 180886166 | + | 2.00792115374673 | 5.68497871032191 | 4.77291284147196 | 3.20132594093141 |
| chr1 | 180886230 | 180886233 | + | 7.02772403811355 | 2.27399148412876 | 11.9322821036799 | 4.26843458790854 |
| chr1 | 180975507 | 180975548 | - | 10.0396057687336 | 19.3289276150945 | 32.2171616799357 | 14.9395210576799 |
| chr1 | 180975579 | 180975638 | - | 56.2217923049084 | 47.753821166704 | 62.0478669391355 | 34.1474767032683 |
| chr1 | 180975731 | 180975833 | - | 11.043566345607 | 13.6439489047726 | 4.77291284147196 | 9.60397782279422 |
| chr1 | 180975908 | 180975919 | - | 4.01584230749346 | 11.3699574206438 | 7.15936926220794 | 1.06710864697714 |
| chr1 | 180976001 | 180976005 | - | 2.00792115374673 | 1.13699574206438 | 1.19322821036799 | 5.33554323488568 |
| chr1 | 180976523 | 180976526 | - | 2.00792115374673 | 2.27399148412876 | 1.19322821036799 | 1.06710864697714 |
| chr1 | 181090668 | 181090693 | + | 2.00792115374673 | 2.27399148412876 | 8.35259747257593 | 7.46976052883995 |
| chr1 | 181090727 | 181090737 | + | 11.043566345607 | 25.0139063254164 | 15.5119667347839 | 24.5434988804741 |
| chr1 | 182383155 | 182384336 | - | 132.522796147284 | 114.836569948503 | 15126.554022835 | 16121.8774385306 |
| chr1 | 182887726 | 182887728 | + | 2.00792115374673 | 2.27399148412876 | 1.19322821036799 | 3.20132594093141 |
| chr1 | 183552276 | 183552286 | + | 1.00396057687336 | 2.27399148412876 | 5.96614105183995 | 4.26843458790854 |
| chr1 | 183552352 | 183552369 | + | 3.01188173062009 | 5.68497871032191 | 4.77291284147196 | 1.06710864697714 |
| chr1 | 183552444 | 183552471 | + | 5.01980288436682 | 4.54798296825753 | 7.15936926220794 | 18.1408469986113 |
| chr1 | 183552495 | 183552537 | + | 3.01188173062009 | 5.68497871032191 | 14.3187385244159 | 7.46976052883995 |
| chr1 | 183552763 | 183552789 | + | 6.02376346124019 | 9.09596593651506 | 7.15936926220794 | 10.6710864697714 |
| chr1 | 183552816 | 183552862 | + | 13.0514874993537 | 9.09596593651506 | 15.5119667347839 | 14.9395210576799 |
| chr1 | 183552951 | 183553032 | + | 6.02376346124019 | 4.54798296825753 | 15.5119667347839 | 29.8790421153598 |
| chr1 | 183553125 | 183553143 | + | 3.01188173062009 | 6.82197445238629 | 1.19322821036799 | 10.6710864697714 |
| chr1 | 183553209 | 183553232 | + | 3.01188173062009 | 3.41098722619315 | 10.7390538933119 | 3.20132594093141 |
| chr1 | 183553289 | 183553319 | + | 2.00792115374673 | 5.68497871032191 | 5.96614105183995 | 9.60397782279422 |
| chr1 | 183553351 | 183553387 | + | 3.01188173062009 | 3.41098722619315 | 13.1255103140479 | 8.53686917581709 |
| chr1 | 183553433 | 183553471 | + | 4.01584230749346 | 2.27399148412876 | 7.15936926220794 | 11.7381951167485 |
| chr1 | 183553554 | 183553568 | + | 4.01584230749346 | 5.68497871032191 | 3.57968463110397 | 4.26843458790854 |
| chr1 | 183553616 | 183553656 | + | 8.03168461498692 | 9.09596593651506 | 15.5119667347839 | 29.8790421153598 |
| chr1 | 183553692 | 183553713 | + | 7.02772403811355 | 4.54798296825753 | 3.57968463110397 | 5.33554323488568 |
| chr1 | 183553873 | 183553885 | + | 1.00396057687336 | 4.54798296825753 | 1.19322821036799 | 8.53686917581709 |
| chr1 | 183553931 | 183553952 | + | 1.00396057687336 | 1.13699574206438 | 8.35259747257593 | 11.7381951167485 |
| chr1 | 183553962 | 183554106 | + | 807.184303806185 | 1105.15986128658 | 982.026817132855 | 978.538629278033 |
| chr1 | 183626346 | 183626381 | - | 21.0831721143407 | 21.6029190992233 | 26.2510206280958 | 11.7381951167485 |
| chr1 | 183626582 | 183626611 | - | 46.1821865361748 | 38.657855230189 | 4.77291284147196 | 14.9395210576799 |
| chr1 | 183626870 | 183626889 | - | 7.02772403811355 | 4.54798296825753 | 1.19322821036799 | 2.13421729395427 |
| chr1 | 183627496 | 183627538 | - | 19.0752509605939 | 11.3699574206438 | 4.77291284147196 | 10.6710864697714 |
| chr1 | 184072929 | 184072954 | + | 14.0554480762271 | 18.1919318730301 | 28.6374770488317 | 26.6777161744284 |
| chr1 | 184073005 | 184073007 | + | 1.00396057687336 | 1.13699574206438 | 7.15936926220794 | 3.20132594093141 |
| chr1 | 184073033 | 184073039 | + | 3.01188173062009 | 3.41098722619315 | 7.15936926220794 | 12.8053037637256 |
| chr1 | 184073105 | 184073137 | + | 7.02772403811355 | 10.2329616785794 | 22.6713359969918 | 11.7381951167485 |
| chr1 | 184073270 | 184073283 | + | 1.00396057687336 | 1.13699574206438 | 7.15936926220794 | 3.20132594093141 |
| chr1 | 184073423 | 184073426 | + | 1.00396057687336 | 2.27399148412876 | 7.15936926220794 | 5.33554323488568 |
| chr1 | 184073461 | 184073493 | + | 5.01980288436682 | 4.54798296825753 | 15.5119667347839 | 5.33554323488568 |
| chr1 | 184073529 | 184073557 | + | 7.02772403811355 | 3.41098722619315 | 28.6374770488317 | 25.6106075274513 |
| chr1 | 184073659 | 184073680 | + | 1.00396057687336 | 3.41098722619315 | 8.35259747257593 | 7.46976052883995 |
| chr1 | 184073703 | 184073711 | + | 16.0633692299738 | 31.8358807778027 | 40.5697591525116 | 42.6843458790854 |
| chr1 | 184073737 | 184073746 | + | 15.0594086531005 | 26.1509020674808 | 34.6036181006717 | 39.483019938154 |
| chr1 | 184073848 | 184073855 | + | 6.02376346124019 | 7.95897019445067 | 4.77291284147196 | 14.9395210576799 |
| chr1 | 184073887 | 184073912 | + | 4.01584230749346 | 1.13699574206438 | 5.96614105183995 | 8.53686917581709 |
| chr1 | 184074029 | 184074048 | + | 20.0792115374673 | 50.0278126508328 | 27.4442488384638 | 52.2883237018796 |
| chr1 | 184074100 | 184074158 | + | 3.01188173062009 | 29.5618892936739 | 28.6374770488317 | 28.8119334683827 |
| chr1 | 184690586 | 184690591 | - | 3.01188173062009 | 4.54798296825753 | 4.77291284147196 | 4.26843458790854 |
| chr1 | 184690829 | 184690848 | - | 14.0554480762271 | 7.95897019445067 | 7.15936926220794 | 6.40265188186281 |
| chr1 | 184690871 | 184690876 | - | 30.1188173062009 | 25.0139063254164 | 9.54582568294392 | 24.5434988804741 |
| chr1 | 184692229 | 184692237 | - | 2.00792115374673 | 4.54798296825753 | 4.77291284147196 | 6.40265188186281 |
| chr1 | 184692345 | 184692415 | - | 9.03564519186028 | 5.68497871032191 | 8.35259747257593 | 7.46976052883995 |
| chr1 | 184692684 | 184692722 | - | 2.00792115374673 | 1.13699574206438 | 7.15936926220794 | 9.60397782279422 |
| chr1 | 184692756 | 184692791 | - | 6.02376346124019 | 7.95897019445067 | 1.19322821036799 | 5.33554323488568 |
| chr1 | 185100477 | 185100485 | + | 48.1901076899215 | 65.9457530397342 | 110.970223564223 | 128.053037637256 |
| chr1 | 185100563 | 185100581 | + | 12.0475269224804 | 9.09596593651506 | 21.4781077866238 | 16.006629704657 |
| chr1 | 185100612 | 185100641 | + | 8.03168461498692 | 9.09596593651506 | 28.6374770488317 | 34.1474767032683 |
| chr1 | 185100719 | 185100725 | + | 4.01584230749346 | 13.6439489047726 | 39.3765309421437 | 19.2079556455884 |
| chr1 | 185100811 | 185100829 | + | 3.01188173062009 | 2.27399148412876 | 11.9322821036799 | 4.26843458790854 |
| chr1 | 185100878 | 185100897 | + | 8.03168461498692 | 10.2329616785794 | 21.4781077866238 | 19.2079556455884 |
| chr1 | 185100928 | 185100957 | + | 43.1703048055547 | 70.4937360079917 | 190.916513658878 | 150.462319223776 |
| chr1 | 185118855 | 185118891 | - | 28.1108961524542 | 36.3838637460602 | 59.6614105183995 | 30.9461507623369 |
| chr1 | 185119079 | 185119083 | - | 3.01188173062009 | 10.2329616785794 | 3.57968463110397 | 8.53686917581709 |
| chr1 | 185119141 | 185119187 | - | 2.00792115374673 | 1.13699574206438 | 10.7390538933119 | 6.40265188186281 |
| chr1 | 185119224 | 185119245 | - | 3.01188173062009 | 2.27399148412876 | 1.19322821036799 | 2.13421729395427 |
| chr1 | 185119302 | 185119305 | - | 1.00396057687336 | 1.13699574206438 | 2.38645642073598 | 5.33554323488568 |
| chr1 | 185296971 | 185296979 | - | 36.1425807674411 | 53.438799877026 | 17.8984231555198 | 35.2145853502455 |
| chr1 | 185297217 | 185297234 | - | 26.1029749987075 | 44.3428339405109 | 33.4103898903037 | 25.6106075274513 |
| chr1 | 185297401 | 185297406 | - | 2.00792115374673 | 5.68497871032191 | 1.19322821036799 | 5.33554323488568 |
| chr1 | 185297636 | 185297685 | - | 8.03168461498692 | 9.09596593651506 | 15.5119667347839 | 6.40265188186281 |
| chr1 | 185297827 | 185297892 | - | 8.03168461498692 | 11.3699574206438 | 13.1255103140479 | 11.7381951167485 |
| chr1 | 185297937 | 185297984 | - | 12.0475269224804 | 35.2468680039958 | 27.4442488384638 | 19.2079556455884 |
| chr1 | 185297987 | 185298008 | - | 5.01980288436682 | 5.68497871032191 | 1.19322821036799 | 2.13421729395427 |
| chr1 | 186313945 | 186314028 | - | 84.3326884573626 | 80.7266976865711 | 72.7869208324474 | 71.4962793474681 |
| chr1 | 186419045 | 186419140 | + | 125.495072109171 | 213.755199508104 | 331.717442482301 | 278.515356861032 |
| chr1 | 186420766 | 186420791 | + | 10.0396057687336 | 27.2878978095452 | 17.8984231555198 | 24.5434988804741 |
| chr1 | 186421069 | 186421073 | + | 5.01980288436682 | 9.09596593651506 | 5.96614105183995 | 2.13421729395427 |
| chr1 | 186421208 | 186421224 | + | 1.00396057687336 | 12.5069531627082 | 10.7390538933119 | 3.20132594093141 |
| chr1 | 187641242 | 187641286 | + | 16.0633692299738 | 7.95897019445067 | 1.19322821036799 | 2.13421729395427 |
| chr1 | 193014525 | 193014527 | - | 2.00792115374673 | 1.13699574206438 | 1.19322821036799 | 2.13421729395427 |
| chr1 | 193015505 | 193015526 | - | 11.043566345607 | 9.09596593651506 | 9.54582568294392 | 10.6710864697714 |
| chr1 | 193015702 | 193015742 | - | 2.00792115374673 | 3.41098722619315 | 7.15936926220794 | 5.33554323488568 |
| chr1 | 193015794 | 193015799 | - | 1.00396057687336 | 5.68497871032191 | 10.7390538933119 | 8.53686917581709 |
| chr1 | 193015832 | 193015962 | - | 148.586165377258 | 118.247557174696 | 159.892580189311 | 170.737383516342 |
| chr1 | 193015980 | 193016004 | - | 12.0475269224804 | 12.5069531627082 | 25.0577924177278 | 7.46976052883995 |
| chr1 | 193016053 | 193016113 | - | 17.0673298068472 | 14.780944646837 | 44.1494437836156 | 36.2816939972226 |
| chr1 | 193016139 | 193016187 | - | 19.0752509605939 | 17.0549361309657 | 62.0478669391355 | 44.8185631730397 |
| chr1 | 193016207 | 193016284 | - | 30.1188173062009 | 17.0549361309657 | 45.3426719939836 | 56.5567582897882 |
| chr1 | 193016301 | 193016323 | - | 4.01584230749346 | 3.41098722619315 | 4.77291284147196 | 2.13421729395427 |
| chr1 | 193084929 | 193084931 | + | 7.02772403811355 | 2.27399148412876 | 3.57968463110397 | 2.13421729395427 |
| chr1 | 193085208 | 193085232 | + | 1.00396057687336 | 6.82197445238629 | 2.38645642073598 | 7.46976052883995 |
| chr1 | 193085415 | 193085431 | + | 24.0950538449607 | 12.5069531627082 | 9.54582568294392 | 23.476390233497 |
| chr1 | 193085478 | 193085486 | + | 2.00792115374673 | 2.27399148412876 | 4.77291284147196 | 8.53686917581709 |
| chr1 | 193085856 | 193085877 | + | 1.00396057687336 | 2.27399148412876 | 7.15936926220794 | 7.46976052883995 |
| chr1 | 193086797 | 193086828 | + | 11.043566345607 | 2.27399148412876 | 8.35259747257593 | 28.8119334683827 |
| chr1 | 193086859 | 193086890 | + | 1.00396057687336 | 4.54798296825753 | 10.7390538933119 | 21.3421729395427 |
| chr1 | 193087714 | 193087738 | + | 11.043566345607 | 13.6439489047726 | 33.4103898903037 | 27.7448248214055 |
| chr1 | 193088665 | 193088687 | + | 6.02376346124019 | 4.54798296825753 | 5.96614105183995 | 20.2750642925656 |
| chr1 | 193088842 | 193088855 | + | 5.01980288436682 | 3.41098722619315 | 5.96614105183995 | 8.53686917581709 |
| chr1 | 193088970 | 193089017 | + | 2.00792115374673 | 1.13699574206438 | 4.77291284147196 | 12.8053037637256 |
| chr1 | 193089058 | 193089068 | + | 1.00396057687336 | 1.13699574206438 | 1.19322821036799 | 4.26843458790854 |
| chr1 | 193089140 | 193089193 | + | 2.00792115374673 | 5.68497871032191 | 7.15936926220794 | 5.33554323488568 |
| chr1 | 193089262 | 193089277 | + | 1.00396057687336 | 3.41098722619315 | 4.77291284147196 | 11.7381951167485 |
| chr1 | 193089343 | 193089427 | + | 19.0752509605939 | 23.876910583352 | 38.1833027317757 | 74.6976052883995 |
| chr1 | 193089597 | 193089618 | + | 5.01980288436682 | 4.54798296825753 | 7.15936926220794 | 8.53686917581709 |
| chr1 | 193090083 | 193090086 | + | 4.01584230749346 | 3.41098722619315 | 3.57968463110397 | 6.40265188186281 |
| chr1 | 193091692 | 193091729 | + | 1.00396057687336 | 2.27399148412876 | 14.3187385244159 | 12.8053037637256 |
| chr1 | 193251145 | 193251150 | + | 1.00396057687336 | 6.82197445238629 | 2.38645642073598 | 2.13421729395427 |
| chr1 | 193251470 | 193251475 | + | 2.00792115374673 | 21.6029190992233 | 5.96614105183995 | 8.53686917581709 |
| chr1 | 196701341 | 196701345 | + | 7.02772403811355 | 1.13699574206438 | 2.38645642073598 | 2.13421729395427 |
| chr1 | 196701367 | 196701534 | + | 708.796167272595 | 569.634866774255 | 157.506123768575 | 119.516168461439 |
| chr1 | 197153753 | 197153764 | - | 2.00792115374673 | 3.41098722619315 | 5.96614105183995 | 1.06710864697714 |
| chr1 | 197154670 | 197154678 | - | 4.01584230749346 | 2.27399148412876 | 4.77291284147196 | 5.33554323488568 |
| chr1 | 197155436 | 197155453 | - | 8.03168461498692 | 5.68497871032191 | 5.96614105183995 | 2.13421729395427 |
| chr1 | 197155898 | 197155903 | - | 3.01188173062009 | 6.82197445238629 | 2.38645642073598 | 2.13421729395427 |
| chr1 | 197156364 | 197156366 | - | 1.00396057687336 | 6.82197445238629 | 2.38645642073598 | 2.13421729395427 |
| chr1 | 197156579 | 197156606 | - | 7.02772403811355 | 5.68497871032191 | 14.3187385244159 | 4.26843458790854 |
| chr1 | 197156780 | 197156805 | - | 4.01584230749346 | 4.54798296825753 | 2.38645642073598 | 2.13421729395427 |
| chr1 | 197157157 | 197157160 | - | 3.01188173062009 | 6.82197445238629 | 3.57968463110397 | 1.06710864697714 |
| chr1 | 197157554 | 197157572 | - | 1.00396057687336 | 6.82197445238629 | 7.15936926220794 | 2.13421729395427 |
| chr1 | 197507759 | 197507761 | - | 1.00396057687336 | 2.27399148412876 | 4.77291284147196 | 1.06710864697714 |
| chr1 | 198319530 | 198319537 | + | 4.01584230749346 | 5.68497871032191 | 2.38645642073598 | 1.06710864697714 |
| chr1 | 198319569 | 198319570 | + | 2.00792115374673 | 1.13699574206438 | 1.19322821036799 | 1.06710864697714 |
| chr1 | 198319619 | 198319630 | + | 11.043566345607 | 13.6439489047726 | 3.57968463110397 | 3.20132594093141 |
| chr1 | 198319709 | 198319774 | + | 18.0712903837206 | 36.3838637460602 | 19.0916513658878 | 4.26843458790854 |
| chr1 | 198319830 | 198319852 | + | 21.0831721143407 | 38.657855230189 | 8.35259747257593 | 9.60397782279422 |
| chr1 | 198319883 | 198319964 | + | 16.0633692299738 | 23.876910583352 | 3.57968463110397 | 7.46976052883995 |
| chr1 | 198320010 | 198320050 | + | 3.01188173062009 | 11.3699574206438 | 1.19322821036799 | 1.06710864697714 |
| chr1 | 198320114 | 198320150 | + | 25.0990144218341 | 21.6029190992233 | 10.7390538933119 | 16.006629704657 |
| chr1 | 198320197 | 198320252 | + | 36.1425807674411 | 68.2197445238629 | 8.35259747257593 | 4.26843458790854 |
| chr1 | 198320635 | 198320705 | + | 20.0792115374673 | 28.4248935516096 | 8.35259747257593 | 6.40265188186281 |
| chr1 | 198320799 | 198320838 | + | 8.03168461498692 | 19.3289276150945 | 4.77291284147196 | 1.06710864697714 |
| chr1 | 198320842 | 198320927 | + | 26.1029749987075 | 30.6988850357383 | 16.7051949451519 | 7.46976052883995 |
| chr1 | 198320958 | 198320962 | + | 5.01980288436682 | 5.68497871032191 | 7.15936926220794 | 1.06710864697714 |
| chr1 | 198320983 | 198321057 | + | 19.0752509605939 | 28.4248935516096 | 10.7390538933119 | 2.13421729395427 |
| chr1 | 198321305 | 198321310 | + | 8.03168461498692 | 6.82197445238629 | 1.19322821036799 | 4.26843458790854 |
| chr1 | 198321971 | 198322001 | + | 22.087132691214 | 29.5618892936739 | 1.19322821036799 | 2.13421729395427 |
| chr1 | 198322136 | 198322141 | + | 9.03564519186028 | 5.68497871032191 | 1.19322821036799 | 2.13421729395427 |
| chr1 | 200405112 | 200405122 | - | 6.02376346124019 | 5.68497871032191 | 4.77291284147196 | 8.53686917581709 |
| chr1 | 200405162 | 200405163 | - | 6.02376346124019 | 3.41098722619315 | 3.57968463110397 | 3.20132594093141 |
| chr1 | 200405257 | 200405259 | - | 3.01188173062009 | 1.13699574206438 | 3.57968463110397 | 1.06710864697714 |
| chr1 | 200405668 | 200405690 | - | 1.00396057687336 | 3.41098722619315 | 1.19322821036799 | 4.26843458790854 |
| chr1 | 200406248 | 200406251 | - | 6.02376346124019 | 2.27399148412876 | 1.19322821036799 | 8.53686917581709 |
| chr1 | 200406746 | 200406755 | - | 7.02772403811355 | 2.27399148412876 | 4.77291284147196 | 3.20132594093141 |
| chr1 | 200406992 | 200407024 | - | 18.0712903837206 | 13.6439489047726 | 17.8984231555198 | 18.1408469986113 |
| chr1 | 200551585 | 200551589 | - | 3.01188173062009 | 5.68497871032191 | 4.77291284147196 | 5.33554323488568 |
| chr1 | 200551774 | 200551778 | - | 4.01584230749346 | 5.68497871032191 | 8.35259747257593 | 8.53686917581709 |
| chr1 | 200551806 | 200551807 | - | 4.01584230749346 | 1.13699574206438 | 5.96614105183995 | 7.46976052883995 |
| chr1 | 200552449 | 200552466 | - | 3.01188173062009 | 1.13699574206438 | 4.77291284147196 | 1.06710864697714 |
| chr1 | 200552528 | 200552531 | - | 3.01188173062009 | 4.54798296825753 | 8.35259747257593 | 9.60397782279422 |
| chr1 | 200552648 | 200552650 | - | 17.0673298068472 | 7.95897019445067 | 4.77291284147196 | 9.60397782279422 |
| chr1 | 200969626 | 200969630 | - | 3.01188173062009 | 1.13699574206438 | 9.54582568294392 | 19.2079556455884 |
| chr1 | 200969666 | 200969712 | - | 4.01584230749346 | 11.3699574206438 | 11.9322821036799 | 5.33554323488568 |
| chr1 | 200969821 | 200969824 | - | 2.00792115374673 | 1.13699574206438 | 5.96614105183995 | 1.06710864697714 |
| chr1 | 200972880 | 200972901 | - | 3.01188173062009 | 4.54798296825753 | 3.57968463110397 | 2.13421729395427 |
| chr1 | 201134798 | 201134953 | - | 35.1386201905678 | 60.2607743294122 | 59.6614105183995 | 33.0803680562912 |
| chr1 | 201134995 | 201135002 | - | 2.00792115374673 | 5.68497871032191 | 3.57968463110397 | 1.06710864697714 |
| chr1 | 201135074 | 201135140 | - | 12.0475269224804 | 14.780944646837 | 19.0916513658878 | 9.60397782279422 |
| chr1 | 201135222 | 201135336 | - | 20.0792115374673 | 21.6029190992233 | 40.5697591525116 | 22.4092815865198 |
| chr1 | 201135409 | 201135445 | - | 9.03564519186028 | 17.0549361309657 | 15.5119667347839 | 3.20132594093141 |
| chr1 | 201135467 | 201135518 | - | 26.1029749987075 | 32.9728765198671 | 25.0577924177278 | 17.0737383516342 |
| chr1 | 201135591 | 201135598 | - | 28.1108961524542 | 34.1098722619315 | 19.0916513658878 | 23.476390233497 |
| chr1 | 201507189 | 201507299 | + | 70.2772403811355 | 158.042408146949 | 16.7051949451519 | 21.3421729395427 |
| chr1 | 201876048 | 201876063 | + | 87.3445701879827 | 36.3838637460602 | 73.9801490428153 | 59.7580842307196 |
| chr1 | 201876093 | 201876130 | + | 3.01188173062009 | 3.41098722619315 | 8.35259747257593 | 11.7381951167485 |
| chr1 | 201876152 | 201876307 | + | 88.3485307648561 | 63.6717615556054 | 78.7530618842873 | 122.717494402371 |
| chr1 | 201876340 | 201876373 | + | 31.1227778830743 | 26.1509020674808 | 13.1255103140479 | 20.2750642925656 |
| chr1 | 201876407 | 201876419 | + | 135.534677877904 | 129.61751459534 | 53.6952694665595 | 133.388580872142 |
| chr1 | 201876470 | 201876500 | + | 10.0396057687336 | 5.68497871032191 | 5.96614105183995 | 8.53686917581709 |
| chr1 | 201876548 | 201876560 | + | 3.01188173062009 | 3.41098722619315 | 4.77291284147196 | 7.46976052883995 |
| chr1 | 201876632 | 201876641 | + | 6.02376346124019 | 6.82197445238629 | 4.77291284147196 | 4.26843458790854 |
| chr1 | 201877038 | 201877054 | + | 3.01188173062009 | 4.54798296825753 | 1.19322821036799 | 2.13421729395427 |
| chr1 | 201878624 | 201878651 | + | 13.0514874993537 | 7.95897019445067 | 5.96614105183995 | 2.13421729395427 |
| chr1 | 201878900 | 201878902 | + | 7.02772403811355 | 11.3699574206438 | 3.57968463110397 | 2.13421729395427 |
| chr1 | 201879016 | 201879039 | + | 6.02376346124019 | 6.82197445238629 | 2.38645642073598 | 3.20132594093141 |
| chr1 | 201879394 | 201879423 | + | 41.1623836518079 | 14.780944646837 | 2.38645642073598 | 12.8053037637256 |
| chr1 | 201880717 | 201880736 | + | 8.03168461498692 | 6.82197445238629 | 1.19322821036799 | 1.06710864697714 |
| chr1 | 201881854 | 201881864 | + | 2.00792115374673 | 5.68497871032191 | 1.19322821036799 | 2.13421729395427 |
| chr1 | 201883685 | 201883687 | + | 5.01980288436682 | 4.54798296825753 | 5.96614105183995 | 5.33554323488568 |
| chr1 | 201969647 | 201969666 | + | 12.0475269224804 | 11.3699574206438 | 7.15936926220794 | 17.0737383516342 |
| chr1 | 201969713 | 201969725 | + | 31.1227778830743 | 30.6988850357383 | 23.8645642073598 | 52.2883237018796 |
| chr1 | 201969828 | 201969838 | + | 1.00396057687336 | 3.41098722619315 | 1.19322821036799 | 3.20132594093141 |
| chr1 | 201969870 | 201969897 | + | 17.0673298068472 | 9.09596593651506 | 8.35259747257593 | 21.3421729395427 |
| chr1 | 201969924 | 201969930 | + | 10.0396057687336 | 13.6439489047726 | 11.9322821036799 | 20.2750642925656 |
| chr1 | 201970187 | 201970197 | + | 8.03168461498692 | 6.82197445238629 | 3.57968463110397 | 9.60397782279422 |
| chr1 | 201970263 | 201970331 | + | 28.1108961524542 | 42.0688424563821 | 57.2749540976635 | 36.2816939972226 |
| chr1 | 202005967 | 202005975 | + | 100.396057687336 | 56.8497871032191 | 15.5119667347839 | 32.0132594093141 |
| chr1 | 202006001 | 202006024 | + | 56.2217923049084 | 32.9728765198671 | 72.7869208324474 | 75.7647139353766 |
| chr1 | 202133580 | 202133601 | - | 54.2138711511617 | 68.2197445238629 | 52.5020412561915 | 28.8119334683827 |
| chr1 | 202134161 | 202134164 | - | 2.00792115374673 | 2.27399148412876 | 2.38645642073598 | 1.06710864697714 |
| chr1 | 202134350 | 202134356 | - | 1.00396057687336 | 2.27399148412876 | 4.77291284147196 | 1.06710864697714 |
| chr1 | 202331778 | 202331792 | - | 13.0514874993537 | 18.1919318730301 | 16.7051949451519 | 7.46976052883995 |
| chr1 | 202331820 | 202331844 | - | 66.2613980736421 | 83.0006891706999 | 35.7968463110397 | 51.2212150549025 |
| chr1 | 202588327 | 202588352 | + | 5.01980288436682 | 11.3699574206438 | 1.19322821036799 | 5.33554323488568 |
| chr1 | 202727671 | 202727695 | - | 1.00396057687336 | 1.13699574206438 | 3.57968463110397 | 3.20132594093141 |
| chr1 | 202880956 | 202880972 | - | 6.02376346124019 | 3.41098722619315 | 2.38645642073598 | 4.26843458790854 |
| chr1 | 202891552 | 202891559 | - | 61.2415951892752 | 68.2197445238629 | 76.3666054635513 | 154.730753811685 |
| chr1 | 202891604 | 202891626 | - | 16.0633692299738 | 35.2468680039958 | 32.2171616799357 | 30.9461507623369 |
| chr1 | 202891710 | 202891713 | - | 1.00396057687336 | 4.54798296825753 | 4.77291284147196 | 1.06710864697714 |
| chr1 | 202891819 | 202891840 | - | 1.00396057687336 | 3.41098722619315 | 13.1255103140479 | 3.20132594093141 |
| chr1 | 202891947 | 202891953 | - | 2.00792115374673 | 1.13699574206438 | 20.2848795762558 | 2.13421729395427 |
| chr1 | 202891974 | 202891988 | - | 3.01188173062009 | 6.82197445238629 | 23.8645642073598 | 8.53686917581709 |
| chr1 | 202892009 | 202892011 | - | 2.00792115374673 | 4.54798296825753 | 2.38645642073598 | 5.33554323488568 |
| chr1 | 202892110 | 202892184 | - | 17.0673298068472 | 37.5208594881246 | 41.7629873628796 | 53.3554323488568 |
| chr1 | 202940908 | 202940916 | - | 2.00792115374673 | 3.41098722619315 | 1.19322821036799 | 4.26843458790854 |
| chr1 | 202941004 | 202941114 | - | 61.2415951892752 | 69.3567402659273 | 34.6036181006717 | 96.0397782279422 |
| chr1 | 202941246 | 202941263 | - | 3.01188173062009 | 3.41098722619315 | 1.19322821036799 | 3.20132594093141 |
| chr1 | 202941290 | 202941321 | - | 1.00396057687336 | 3.41098722619315 | 4.77291284147196 | 2.13421729395427 |
| chr1 | 202941367 | 202941379 | - | 9.03564519186028 | 5.68497871032191 | 2.38645642073598 | 7.46976052883995 |
| chr1 | 202941546 | 202941551 | - | 19.0752509605939 | 10.2329616785794 | 10.7390538933119 | 8.53686917581709 |
| chr1 | 202942024 | 202942123 | - | 7.02772403811355 | 2.27399148412876 | 3.57968463110397 | 1.06710864697714 |
| chr1 | 202961966 | 202961987 | - | 4.01584230749346 | 5.68497871032191 | 4.77291284147196 | 4.26843458790854 |
| chr1 | 202961995 | 202962025 | - | 12.0475269224804 | 5.68497871032191 | 4.77291284147196 | 5.33554323488568 |
| chr1 | 202962048 | 202962113 | - | 9.03564519186028 | 20.4659233571589 | 20.2848795762558 | 16.006629704657 |
| chr1 | 202962162 | 202962177 | - | 5.01980288436682 | 4.54798296825753 | 3.57968463110397 | 4.26843458790854 |
| chr1 | 202962254 | 202962270 | - | 11.043566345607 | 9.09596593651506 | 10.7390538933119 | 9.60397782279422 |
| chr1 | 203024048 | 203024069 | + | 1.00396057687336 | 1.13699574206438 | 3.57968463110397 | 3.20132594093141 |
| chr1 | 203024818 | 203024821 | + | 7.02772403811355 | 6.82197445238629 | 9.54582568294392 | 8.53686917581709 |
| chr1 | 203307459 | 203307538 | + | 56.2217923049084 | 31.8358807778027 | 155.119667347839 | 180.341361339136 |
| chr1 | 203308120 | 203308148 | + | 3.01188173062009 | 3.41098722619315 | 31.0239334695677 | 16.006629704657 |
| chr1 | 203308226 | 203308229 | + | 2.00792115374673 | 1.13699574206438 | 2.38645642073598 | 4.26843458790854 |
| chr1 | 203308799 | 203308808 | + | 1.00396057687336 | 1.13699574206438 | 3.57968463110397 | 1.06710864697714 |
| chr1 | 203490542 | 203490544 | + | 55.217831728035 | 90.9596593651506 | 45.3426719939836 | 70.429170700491 |
| chr1 | 203739652 | 203739695 | + | 14.0554480762271 | 19.3289276150945 | 13.1255103140479 | 1.06710864697714 |
| chr1 | 203739813 | 203739839 | + | 7.02772403811355 | 5.68497871032191 | 5.96614105183995 | 8.53686917581709 |
| chr1 | 203740265 | 203740313 | + | 8.03168461498692 | 5.68497871032191 | 5.96614105183995 | 19.2079556455884 |
| chr1 | 203740396 | 203740425 | + | 3.01188173062009 | 7.95897019445067 | 8.35259747257593 | 3.20132594093141 |
| chr1 | 203740888 | 203740902 | + | 1.00396057687336 | 2.27399148412876 | 3.57968463110397 | 2.13421729395427 |
| chr1 | 203740960 | 203740962 | + | 9.03564519186028 | 6.82197445238629 | 2.38645642073598 | 4.26843458790854 |
| chr1 | 203741034 | 203741065 | + | 8.03168461498692 | 10.2329616785794 | 11.9322821036799 | 2.13421729395427 |
| chr1 | 203741367 | 203741370 | + | 3.01188173062009 | 3.41098722619315 | 4.77291284147196 | 1.06710864697714 |
| chr1 | 203741985 | 203742017 | + | 26.1029749987075 | 18.1919318730301 | 14.3187385244159 | 12.8053037637256 |
| chr1 | 203742181 | 203742198 | + | 4.01584230749346 | 5.68497871032191 | 2.38645642073598 | 2.13421729395427 |
| chr1 | 203742502 | 203742528 | + | 7.02772403811355 | 5.68497871032191 | 3.57968463110397 | 1.06710864697714 |
| chr1 | 203742772 | 203742796 | + | 11.043566345607 | 7.95897019445067 | 2.38645642073598 | 1.06710864697714 |
| chr1 | 203743184 | 203743208 | + | 11.043566345607 | 5.68497871032191 | 16.7051949451519 | 3.20132594093141 |
| chr1 | 203743340 | 203743364 | + | 19.0752509605939 | 19.3289276150945 | 16.7051949451519 | 14.9395210576799 |
| chr1 | 203743494 | 203743538 | + | 44.174265382428 | 60.2607743294122 | 29.8307052591997 | 29.8790421153598 |
| chr1 | 203743613 | 203743651 | + | 32.1267384599477 | 26.1509020674808 | 16.7051949451519 | 16.006629704657 |
| chr1 | 203743684 | 203743699 | + | 5.01980288436682 | 7.95897019445067 | 2.38645642073598 | 3.20132594093141 |
| chr1 | 203743723 | 203743832 | + | 39.1544624980612 | 45.4798296825753 | 44.1494437836156 | 36.2816939972226 |
| chr1 | 203743855 | 203743917 | + | 114.451505763564 | 120.521548658824 | 69.2072362013434 | 28.8119334683827 |
| chr1 | 203743944 | 203744019 | + | 169.669337491599 | 212.618203766039 | 190.916513658878 | 89.6371263460794 |
| chr1 | 203801628 | 203801640 | + | 3.01188173062009 | 4.54798296825753 | 1.19322821036799 | 4.26843458790854 |
| chr1 | 203801756 | 203801801 | + | 3.01188173062009 | 3.41098722619315 | 3.57968463110397 | 2.13421729395427 |
| chr1 | 203802079 | 203802127 | + | 3.01188173062009 | 1.13699574206438 | 4.77291284147196 | 4.26843458790854 |
| chr1 | 203802275 | 203802291 | + | 4.01584230749346 | 7.95897019445067 | 3.57968463110397 | 4.26843458790854 |
| chr1 | 203802372 | 203802390 | + | 10.0396057687336 | 5.68497871032191 | 16.7051949451519 | 7.46976052883995 |
| chr1 | 203802419 | 203802462 | + | 3.01188173062009 | 1.13699574206438 | 9.54582568294392 | 7.46976052883995 |
| chr1 | 203802525 | 203802544 | + | 23.0910932680874 | 19.3289276150945 | 13.1255103140479 | 17.0737383516342 |
| chr1 | 203802594 | 203802609 | + | 3.01188173062009 | 1.13699574206438 | 1.19322821036799 | 6.40265188186281 |
| chr1 | 203802650 | 203802700 | + | 2.00792115374673 | 5.68497871032191 | 4.77291284147196 | 12.8053037637256 |
| chr1 | 203802874 | 203802898 | + | 5.01980288436682 | 7.95897019445067 | 7.15936926220794 | 3.20132594093141 |
| chr1 | 203828414 | 203828423 | + | 3.01188173062009 | 4.54798296825753 | 1.19322821036799 | 1.06710864697714 |
| chr1 | 203833789 | 203833854 | + | 21.0831721143407 | 45.4798296825753 | 34.6036181006717 | 12.8053037637256 |
| chr1 | 203840361 | 203840375 | + | 11.043566345607 | 7.95897019445067 | 4.77291284147196 | 4.26843458790854 |
| chr1 | 203847243 | 203847264 | + | 6.02376346124019 | 10.2329616785794 | 14.3187385244159 | 6.40265188186281 |
| chr1 | 203847363 | 203847374 | + | 3.01188173062009 | 12.5069531627082 | 11.9322821036799 | 3.20132594093141 |
| chr1 | 203847477 | 203847560 | + | 11.043566345607 | 22.7399148412876 | 36.9900745214077 | 26.6777161744284 |
| chr1 | 203848328 | 203848399 | + | 17.0673298068472 | 26.1509020674808 | 68.0140079909754 | 42.6843458790854 |
| chr1 | 203849708 | 203849718 | + | 5.01980288436682 | 6.82197445238629 | 7.15936926220794 | 3.20132594093141 |
| chr1 | 203849750 | 203849777 | + | 5.01980288436682 | 10.2329616785794 | 13.1255103140479 | 6.40265188186281 |
| chr1 | 203849805 | 203849807 | + | 6.02376346124019 | 4.54798296825753 | 4.77291284147196 | 1.06710864697714 |
| chr1 | 203849943 | 203849987 | + | 10.0396057687336 | 13.6439489047726 | 27.4442488384638 | 7.46976052883995 |
| chr1 | 203850002 | 203850027 | + | 5.01980288436682 | 7.95897019445067 | 9.54582568294392 | 6.40265188186281 |
| chr1 | 203850525 | 203850534 | + | 5.01980288436682 | 1.13699574206438 | 3.57968463110397 | 2.13421729395427 |
| chr1 | 203852270 | 203852295 | + | 5.01980288436682 | 11.3699574206438 | 5.96614105183995 | 6.40265188186281 |
| chr1 | 203852344 | 203852373 | + | 8.03168461498692 | 12.5069531627082 | 5.96614105183995 | 8.53686917581709 |
| chr1 | 203852674 | 203852735 | + | 78.3089249961224 | 97.7816338175368 | 118.129592826431 | 60.8251928776967 |
| chr1 | 203853745 | 203853800 | + | 2.00792115374673 | 4.54798296825753 | 1.19322821036799 | 2.13421729395427 |
| chr1 | 203853876 | 203853919 | + | 2.00792115374673 | 11.3699574206438 | 5.96614105183995 | 10.6710864697714 |
| chr1 | 203869985 | 203870000 | + | 1.00396057687336 | 3.41098722619315 | 1.19322821036799 | 1.06710864697714 |
| chr1 | 203870030 | 203870068 | + | 23.0910932680874 | 38.657855230189 | 22.6713359969918 | 21.3421729395427 |
| chr1 | 203870268 | 203870284 | + | 3.01188173062009 | 12.5069531627082 | 5.96614105183995 | 4.26843458790854 |
| chr1 | 204403512 | 204403537 | - | 20.0792115374673 | 46.6168254246397 | 2.38645642073598 | 3.20132594093141 |
| chr1 | 204404337 | 204404346 | - | 1749.90328549027 | 2234.19663315651 | 1026.17626091647 | 730.969423179338 |
| chr1 | 204404522 | 204404562 | - | 11.043566345607 | 9.09596593651506 | 2.38645642073598 | 2.13421729395427 |
| chr1 | 204404664 | 204404667 | - | 4.01584230749346 | 4.54798296825753 | 1.19322821036799 | 1.06710864697714 |
| chr1 | 204404693 | 204404697 | - | 13.0514874993537 | 18.1919318730301 | 3.57968463110397 | 2.13421729395427 |
| chr1 | 204404769 | 204404814 | - | 25.0990144218341 | 35.2468680039958 | 2.38645642073598 | 7.46976052883995 |
| chr1 | 204404868 | 204404880 | - | 2.00792115374673 | 2.27399148412876 | 3.57968463110397 | 3.20132594093141 |
| chr1 | 204405686 | 204405713 | - | 3.01188173062009 | 4.54798296825753 | 2.38645642073598 | 3.20132594093141 |
| chr1 | 204405800 | 204405825 | - | 9.03564519186028 | 26.1509020674808 | 10.7390538933119 | 9.60397782279422 |
| chr1 | 204406013 | 204406048 | - | 18.0712903837206 | 31.8358807778027 | 9.54582568294392 | 10.6710864697714 |
| chr1 | 204406083 | 204406094 | - | 5.01980288436682 | 3.41098722619315 | 1.19322821036799 | 1.06710864697714 |
| chr1 | 204424508 | 204424522 | - | 61.2415951892752 | 67.0827487817985 | 14.3187385244159 | 1.06710864697714 |
| chr1 | 204542815 | 204542874 | + | 15.0594086531005 | 27.2878978095452 | 27.4442488384638 | 8.53686917581709 |
| chr1 | 204551976 | 204551977 | + | 13.0514874993537 | 15.9179403889013 | 8.35259747257593 | 12.8053037637256 |
| chr1 | 204552599 | 204552604 | + | 9.03564519186028 | 6.82197445238629 | 8.35259747257593 | 1.06710864697714 |
| chr1 | 204552836 | 204552852 | + | 3.01188173062009 | 7.95897019445067 | 13.1255103140479 | 3.20132594093141 |
| chr1 | 204553164 | 204553188 | + | 3.01188173062009 | 7.95897019445067 | 8.35259747257593 | 4.26843458790854 |
| chr1 | 204554080 | 204554091 | + | 3.01188173062009 | 3.41098722619315 | 3.57968463110397 | 1.06710864697714 |
| chr1 | 205088491 | 205088492 | - | 1.00396057687336 | 1.13699574206438 | 1.19322821036799 | 1.06710864697714 |
| chr1 | 205088657 | 205088666 | - | 3.01188173062009 | 2.27399148412876 | 1.19322821036799 | 3.20132594093141 |
| chr1 | 205142868 | 205142875 | - | 9.03564519186028 | 3.41098722619315 | 2.38645642073598 | 3.20132594093141 |
| chr1 | 205144768 | 205144772 | - | 3.01188173062009 | 2.27399148412876 | 1.19322821036799 | 1.06710864697714 |
| chr1 | 205531641 | 205531648 | + | 2.00792115374673 | 1.13699574206438 | 4.77291284147196 | 1.06710864697714 |
| chr1 | 205607978 | 205608005 | - | 2.00792115374673 | 3.41098722619315 | 4.77291284147196 | 3.20132594093141 |
| chr1 | 205608222 | 205608247 | - | 19.0752509605939 | 45.4798296825753 | 7.15936926220794 | 4.26843458790854 |
| chr1 | 205608303 | 205608319 | - | 21.0831721143407 | 79.5897019445067 | 50.1155848354556 | 18.1408469986113 |
| chr1 | 205610092 | 205610116 | - | 19.0752509605939 | 30.6988850357383 | 11.9322821036799 | 13.8724124107028 |
| chr1 | 205610160 | 205610183 | - | 11.043566345607 | 4.54798296825753 | 3.57968463110397 | 6.40265188186281 |
| chr1 | 205611346 | 205611371 | - | 7.02772403811355 | 11.3699574206438 | 4.77291284147196 | 4.26843458790854 |
| chr1 | 205611590 | 205611615 | - | 9.03564519186028 | 4.54798296825753 | 3.57968463110397 | 2.13421729395427 |
| chr1 | 205613100 | 205613121 | - | 24.0950538449607 | 22.7399148412876 | 9.54582568294392 | 29.8790421153598 |
| chr1 | 205613663 | 205613674 | - | 87.3445701879827 | 68.2197445238629 | 35.7968463110397 | 14.9395210576799 |
| chr1 | 205614497 | 205614499 | - | 1.00396057687336 | 5.68497871032191 | 3.57968463110397 | 1.06710864697714 |
| chr1 | 205615131 | 205615147 | - | 9.03564519186028 | 12.5069531627082 | 2.38645642073598 | 10.6710864697714 |
| chr1 | 205713321 | 205713356 | - | 15.0594086531005 | 27.2878978095452 | 19.0916513658878 | 22.4092815865198 |
| chr1 | 205713440 | 205713448 | - | 8.03168461498692 | 9.09596593651506 | 2.38645642073598 | 5.33554323488568 |
| chr1 | 205713589 | 205713615 | - | 30.1188173062009 | 60.2607743294122 | 9.54582568294392 | 13.8724124107028 |
| chr1 | 205713655 | 205713685 | - | 6.02376346124019 | 5.68497871032191 | 3.57968463110397 | 10.6710864697714 |
| chr1 | 205713792 | 205713802 | - | 8.03168461498692 | 4.54798296825753 | 4.77291284147196 | 19.2079556455884 |
| chr1 | 205713840 | 205713852 | - | 5.01980288436682 | 4.54798296825753 | 1.19322821036799 | 3.20132594093141 |
| chr1 | 205713921 | 205713965 | - | 5.01980288436682 | 1.13699574206438 | 2.38645642073598 | 2.13421729395427 |
| chr1 | 205714182 | 205714193 | - | 7.02772403811355 | 4.54798296825753 | 3.57968463110397 | 2.13421729395427 |
| chr1 | 205714242 | 205714329 | - | 39.1544624980612 | 38.657855230189 | 26.2510206280958 | 36.2816939972226 |
| chr1 | 205714525 | 205714533 | - | 2.00792115374673 | 2.27399148412876 | 2.38645642073598 | 2.13421729395427 |
| chr1 | 205714559 | 205714569 | - | 18.0712903837206 | 9.09596593651506 | 2.38645642073598 | 3.20132594093141 |
| chr1 | 205714643 | 205714675 | - | 3.01188173062009 | 2.27399148412876 | 3.57968463110397 | 2.13421729395427 |
| chr1 | 205714697 | 205714710 | - | 7.02772403811355 | 2.27399148412876 | 1.19322821036799 | 2.13421729395427 |
| chr1 | 205714781 | 205714792 | - | 2.00792115374673 | 1.13699574206438 | 5.96614105183995 | 2.13421729395427 |
| chr1 | 205714948 | 205714961 | - | 3.01188173062009 | 1.13699574206438 | 2.38645642073598 | 5.33554323488568 |
| chr1 | 205715033 | 205715139 | - | 19.0752509605939 | 17.0549361309657 | 45.3426719939836 | 33.0803680562912 |
| chr1 | 205715259 | 205715385 | - | 21.0831721143407 | 10.2329616785794 | 32.2171616799357 | 16.006629704657 |
| chr1 | 205715452 | 205715487 | - | 6.02376346124019 | 5.68497871032191 | 2.38645642073598 | 5.33554323488568 |
| chr1 | 205715599 | 205715647 | - | 10.0396057687336 | 10.2329616785794 | 13.1255103140479 | 10.6710864697714 |
| chr1 | 205715674 | 205715797 | - | 73.2891221117556 | 113.699574206438 | 218.360762497342 | 118.449059814462 |
| chr1 | 205715813 | 205715929 | - | 44.174265382428 | 54.5757956190903 | 145.573841664895 | 42.6843458790854 |
| chr1 | 205715954 | 205716070 | - | 21.0831721143407 | 25.0139063254164 | 60.8546387287675 | 24.5434988804741 |
| chr1 | 205716076 | 205716417 | - | 363.433728828158 | 425.236407532079 | 341.263268165245 | 250.770532039627 |
| chr1 | 205716442 | 205716567 | - | 33.130699036821 | 28.4248935516096 | 39.3765309421437 | 44.8185631730397 |
| chr1 | 205716631 | 205716648 | - | 1.00396057687336 | 2.27399148412876 | 5.96614105183995 | 5.33554323488568 |
| chr1 | 205716716 | 205716721 | - | 1.00396057687336 | 2.27399148412876 | 4.77291284147196 | 1.06710864697714 |
| chr1 | 205716745 | 205716776 | - | 17.0673298068472 | 22.7399148412876 | 4.77291284147196 | 18.1408469986113 |
| chr1 | 205717183 | 205717218 | - | 15.0594086531005 | 17.0549361309657 | 28.6374770488317 | 12.8053037637256 |
| chr1 | 205717225 | 205717240 | - | 3.01188173062009 | 4.54798296825753 | 2.38645642073598 | 1.06710864697714 |
| chr1 | 205717272 | 205717302 | - | 6.02376346124019 | 7.95897019445067 | 14.3187385244159 | 11.7381951167485 |
| chr1 | 205717331 | 205717381 | - | 10.0396057687336 | 13.6439489047726 | 15.5119667347839 | 10.6710864697714 |
| chr1 | 205717414 | 205717432 | - | 2.00792115374673 | 5.68497871032191 | 3.57968463110397 | 1.06710864697714 |
| chr1 | 205717475 | 205717562 | - | 47.1861471130481 | 40.9318467143178 | 78.7530618842873 | 50.1541064079254 |
| chr1 | 205717596 | 205717734 | - | 484.912958629835 | 527.566024317873 | 310.239334695677 | 521.816128371819 |
| chr1 | 205717776 | 205717847 | - | 14.0554480762271 | 5.68497871032191 | 7.15936926220794 | 3.20132594093141 |
| chr1 | 205717892 | 205717955 | - | 27.1069355755808 | 18.1919318730301 | 25.0577924177278 | 16.006629704657 |
| chr1 | 205717972 | 205718101 | - | 34.1346596136944 | 30.6988850357383 | 52.5020412561915 | 35.2145853502455 |
| chr1 | 205718125 | 205718238 | - | 69.2732798042621 | 75.0417189762492 | 124.095733878271 | 364.95115726618 |
| chr1 | 205718259 | 205718334 | - | 146.578244223511 | 133.028501821533 | 54.8884976769275 | 67.2278447595595 |
| chr1 | 205791163 | 205791221 | - | 6.02376346124019 | 7.95897019445067 | 3.57968463110397 | 28.8119334683827 |
| chr1 | 205978281 | 205978282 | - | 17.0673298068472 | 31.8358807778027 | 5.96614105183995 | 24.5434988804741 |
| chr1 | 206464381 | 206464401 | + | 4.01584230749346 | 6.82197445238629 | 2.38645642073598 | 2.13421729395427 |
| chr1 | 206543211 | 206543212 | + | 3.01188173062009 | 5.68497871032191 | 1.19322821036799 | 2.13421729395427 |
| chr1 | 206586941 | 206587003 | + | 1.00396057687336 | 3.41098722619315 | 21.4781077866238 | 5.33554323488568 |
| chr1 | 206588801 | 206588826 | + | 10.0396057687336 | 2.27399148412876 | 42.9562155732476 | 17.0737383516342 |
| chr1 | 206588899 | 206588960 | + | 26.1029749987075 | 14.780944646837 | 87.1056593568632 | 71.4962793474681 |
| chr1 | 206588981 | 206589116 | + | 34.1346596136944 | 32.9728765198671 | 176.597775134462 | 84.3015831111937 |
| chr1 | 206589141 | 206589150 | + | 2.00792115374673 | 2.27399148412876 | 7.15936926220794 | 8.53686917581709 |
| chr1 | 206591682 | 206591788 | - | 59.2336740355285 | 50.0278126508328 | 39.3765309421437 | 35.2145853502455 |
| chr1 | 206649127 | 206649128 | + | 1.00396057687336 | 1.13699574206438 | 2.38645642073598 | 3.20132594093141 |
| chr1 | 206732744 | 206732795 | + | 6.02376346124019 | 6.82197445238629 | 23.8645642073598 | 23.476390233497 |
| chr1 | 206732828 | 206732836 | + | 3.01188173062009 | 3.41098722619315 | 5.96614105183995 | 3.20132594093141 |
| chr1 | 206732948 | 206732972 | + | 1.00396057687336 | 4.54798296825753 | 7.15936926220794 | 4.26843458790854 |
| chr1 | 206733041 | 206733096 | + | 14.0554480762271 | 5.68497871032191 | 13.1255103140479 | 12.8053037637256 |
| chr1 | 206733155 | 206733208 | + | 21.0831721143407 | 12.5069531627082 | 14.3187385244159 | 13.8724124107028 |
| chr1 | 206733232 | 206733270 | + | 10.0396057687336 | 6.82197445238629 | 21.4781077866238 | 58.6909755837425 |
| chr1 | 206733291 | 206733320 | + | 4.01584230749346 | 1.13699574206438 | 8.35259747257593 | 8.53686917581709 |
| chr1 | 206734126 | 206734145 | + | 2.00792115374673 | 1.13699574206438 | 3.57968463110397 | 3.20132594093141 |
| chr1 | 206734166 | 206734220 | + | 12.0475269224804 | 21.6029190992233 | 11.9322821036799 | 26.6777161744284 |
| chr1 | 207044041 | 207044130 | - | 150.594086531005 | 247.865071770035 | 954.582568294392 | 794.995941997966 |
| chr1 | 207044158 | 207044211 | - | 49.1940682667949 | 48.8908169087684 | 107.390538933119 | 145.12677598889 |
| chr1 | 207044265 | 207044277 | - | 3.01188173062009 | 9.09596593651506 | 19.0916513658878 | 11.7381951167485 |
| chr1 | 207044319 | 207044341 | - | 8.03168461498692 | 10.2329616785794 | 26.2510206280958 | 25.6106075274513 |
| chr1 | 207044371 | 207044447 | - | 46.1821865361748 | 34.1098722619315 | 79.9462900946553 | 68.2949534065367 |
| chr1 | 207044464 | 207044521 | - | 7.02772403811355 | 3.41098722619315 | 29.8307052591997 | 16.006629704657 |
| chr1 | 207044542 | 207044559 | - | 3.01188173062009 | 2.27399148412876 | 33.4103898903037 | 32.0132594093141 |
| chr1 | 207044598 | 207044602 | - | 1.00396057687336 | 2.27399148412876 | 5.96614105183995 | 3.20132594093141 |
| chr1 | 207044811 | 207044832 | - | 2.00792115374673 | 1.13699574206438 | 11.9322821036799 | 16.006629704657 |
| chr1 | 207044881 | 207044910 | - | 2.00792115374673 | 1.13699574206438 | 13.1255103140479 | 3.20132594093141 |
| chr1 | 207045145 | 207045179 | - | 1.00396057687336 | 1.13699574206438 | 7.15936926220794 | 13.8724124107028 |
| chr1 | 207045463 | 207045478 | - | 4.01584230749346 | 1.13699574206438 | 4.77291284147196 | 4.26843458790854 |
| chr1 | 207045618 | 207045680 | - | 17.0673298068472 | 12.5069531627082 | 53.6952694665595 | 86.435800405148 |
| chr1 | 207045706 | 207045723 | - | 2.00792115374673 | 2.27399148412876 | 5.96614105183995 | 5.33554323488568 |
| chr1 | 207046086 | 207046113 | - | 2.00792115374673 | 7.95897019445067 | 17.8984231555198 | 6.40265188186281 |
| chr1 | 207046166 | 207046186 | - | 1.00396057687336 | 2.27399148412876 | 5.96614105183995 | 4.26843458790854 |
| chr1 | 207046208 | 207046227 | - | 2.00792115374673 | 1.13699574206438 | 8.35259747257593 | 6.40265188186281 |
| chr1 | 207046270 | 207046341 | - | 8.03168461498692 | 7.95897019445067 | 34.6036181006717 | 28.8119334683827 |
| chr1 | 207046408 | 207046439 | - | 12.0475269224804 | 11.3699574206438 | 22.6713359969918 | 27.7448248214055 |
| chr1 | 207046497 | 207046517 | - | 1.00396057687336 | 1.13699574206438 | 5.96614105183995 | 9.60397782279422 |
| chr1 | 207046610 | 207046630 | - | 1.00396057687336 | 5.68497871032191 | 9.54582568294392 | 4.26843458790854 |
| chr1 | 207046652 | 207046712 | - | 7.02772403811355 | 9.09596593651506 | 51.3088130458235 | 25.6106075274513 |
| chr1 | 207046755 | 207046776 | - | 2.00792115374673 | 2.27399148412876 | 8.35259747257593 | 7.46976052883995 |
| chr1 | 207047122 | 207047160 | - | 2.00792115374673 | 3.41098722619315 | 15.5119667347839 | 7.46976052883995 |
| chr1 | 207047383 | 207047407 | - | 2.00792115374673 | 7.95897019445067 | 4.77291284147196 | 7.46976052883995 |
| chr1 | 207047547 | 207047565 | - | 2.00792115374673 | 2.27399148412876 | 10.7390538933119 | 2.13421729395427 |
| chr1 | 207047603 | 207047611 | - | 2.00792115374673 | 2.27399148412876 | 4.77291284147196 | 9.60397782279422 |
| chr1 | 207048035 | 207048068 | - | 1.00396057687336 | 4.54798296825753 | 15.5119667347839 | 2.13421729395427 |
| chr1 | 207048999 | 207049005 | - | 4.01584230749346 | 6.82197445238629 | 5.96614105183995 | 3.20132594093141 |
| chr1 | 207073163 | 207073188 | + | 5.01980288436682 | 4.54798296825753 | 2.38645642073598 | 6.40265188186281 |
| chr1 | 207073449 | 207073463 | + | 1.00396057687336 | 9.09596593651506 | 2.38645642073598 | 1.06710864697714 |
| chr1 | 207073845 | 207073866 | + | 2.00792115374673 | 2.27399148412876 | 3.57968463110397 | 3.20132594093141 |
| chr1 | 207074463 | 207074469 | + | 3.01188173062009 | 3.41098722619315 | 1.19322821036799 | 2.13421729395427 |
| chr1 | 207074678 | 207074726 | + | 12.0475269224804 | 30.6988850357383 | 19.0916513658878 | 16.006629704657 |
| chr1 | 207074896 | 207074902 | + | 2.00792115374673 | 6.82197445238629 | 2.38645642073598 | 3.20132594093141 |
| chr1 | 207074935 | 207074958 | + | 5.01980288436682 | 14.780944646837 | 5.96614105183995 | 3.20132594093141 |
| chr1 | 207075046 | 207075083 | + | 2.00792115374673 | 10.2329616785794 | 10.7390538933119 | 8.53686917581709 |
| chr1 | 207075375 | 207075388 | + | 2.00792115374673 | 3.41098722619315 | 3.57968463110397 | 4.26843458790854 |
| chr1 | 207075414 | 207075421 | + | 2.00792115374673 | 3.41098722619315 | 2.38645642073598 | 1.06710864697714 |
| chr1 | 207077553 | 207077561 | + | 5.01980288436682 | 10.2329616785794 | 2.38645642073598 | 12.8053037637256 |
| chr1 | 207077584 | 207077610 | + | 2.00792115374673 | 5.68497871032191 | 1.19322821036799 | 4.26843458790854 |
| chr1 | 207359546 | 207359718 | + | 111.439624032943 | 160.316399631078 | 8.35259747257593 | 7.46976052883995 |
| chr1 | 207359832 | 207359922 | + | 42.1663442286813 | 53.438799877026 | 7.15936926220794 | 2.13421729395427 |
| chr1 | 207360039 | 207360199 | + | 779.073407653731 | 1232.50338439779 | 36.9900745214077 | 61.8923015246739 |
| chr1 | 207360222 | 207360306 | + | 36.1425807674411 | 81.8636934286355 | 3.57968463110397 | 4.26843458790854 |
| chr1 | 207360332 | 207360354 | + | 35.1386201905678 | 64.8087572976698 | 5.96614105183995 | 3.20132594093141 |
| chr1 | 207360399 | 207360483 | + | 71.2812009580089 | 142.124467758048 | 2.38645642073598 | 2.13421729395427 |
| chr1 | 207373025 | 207373026 | + | 1.00396057687336 | 1.13699574206438 | 1.19322821036799 | 1.06710864697714 |
| chr1 | 207793522 | 207793535 | + | 1.00396057687336 | 1.13699574206438 | 1.19322821036799 | 4.26843458790854 |
| chr1 | 207794132 | 207794373 | + | 32.1267384599477 | 26.1509020674808 | 10.7390538933119 | 6.40265188186281 |
| chr1 | 207794460 | 207794590 | + | 13.0514874993537 | 27.2878978095452 | 13.1255103140479 | 6.40265188186281 |
| chr1 | 207794614 | 207794747 | + | 28.1108961524542 | 30.6988850357383 | 38.1833027317757 | 21.3421729395427 |
| chr1 | 207794797 | 207794810 | + | 11.043566345607 | 2.27399148412876 | 2.38645642073598 | 2.13421729395427 |
| chr1 | 207794871 | 207795180 | + | 114.451505763564 | 115.973565690567 | 44.1494437836156 | 35.2145853502455 |
| chr1 | 207795308 | 207795366 | + | 28.1108961524542 | 22.7399148412876 | 19.0916513658878 | 9.60397782279422 |
| chr1 | 207795389 | 207795481 | + | 75.2970432655023 | 110.288586980245 | 35.7968463110397 | 16.006629704657 |
| chr1 | 208026997 | 208027006 | - | 10.0396057687336 | 5.68497871032191 | 1.19322821036799 | 1.06710864697714 |
| chr1 | 208730066 | 208730067 | + | 1.00396057687336 | 1.13699574206438 | 1.19322821036799 | 2.13421729395427 |
| chr1 | 209851664 | 209851670 | + | 24.0950538449607 | 15.9179403889013 | 4.77291284147196 | 7.46976052883995 |
| chr1 | 211373151 | 211373167 | + | 8.03168461498692 | 7.95897019445067 | 9.54582568294392 | 4.26843458790854 |
| chr1 | 211374422 | 211374442 | + | 3.01188173062009 | 5.68497871032191 | 1.19322821036799 | 2.13421729395427 |
| chr1 | 211382765 | 211382883 | + | 203.803997105293 | 465.031258504332 | 403.31113510438 | 424.7092414969 |
| chr1 | 211571810 | 211571820 | - | 19.0752509605939 | 19.3289276150945 | 2.38645642073598 | 2.13421729395427 |
| chr1 | 211572408 | 211572483 | - | 45.1782259593014 | 42.0688424563821 | 16.7051949451519 | 10.6710864697714 |
| chr1 | 211572528 | 211572556 | - | 17.0673298068472 | 13.6439489047726 | 11.9322821036799 | 3.20132594093141 |
| chr1 | 211572584 | 211572657 | - | 9.03564519186028 | 17.0549361309657 | 5.96614105183995 | 4.26843458790854 |
| chr1 | 211572829 | 211572850 | - | 8.03168461498692 | 6.82197445238629 | 4.77291284147196 | 1.06710864697714 |
| chr1 | 211572881 | 211572904 | - | 8.03168461498692 | 7.95897019445067 | 4.77291284147196 | 1.06710864697714 |
| chr1 | 211573191 | 211573205 | - | 14.0554480762271 | 10.2329616785794 | 4.77291284147196 | 5.33554323488568 |
| chr1 | 211573340 | 211573379 | - | 23.0910932680874 | 18.1919318730301 | 8.35259747257593 | 4.26843458790854 |
| chr1 | 211573457 | 211573463 | - | 2.00792115374673 | 7.95897019445067 | 3.57968463110397 | 2.13421729395427 |
| chr1 | 211573586 | 211573625 | - | 19.0752509605939 | 17.0549361309657 | 7.15936926220794 | 2.13421729395427 |
| chr1 | 211573649 | 211573684 | - | 12.0475269224804 | 11.3699574206438 | 3.57968463110397 | 1.06710864697714 |
| chr1 | 211573729 | 211573747 | - | 2.00792115374673 | 2.27399148412876 | 2.38645642073598 | 1.06710864697714 |
| chr1 | 211573836 | 211573858 | - | 23.0910932680874 | 46.6168254246397 | 5.96614105183995 | 4.26843458790854 |
| chr1 | 211573891 | 211573958 | - | 30.1188173062009 | 32.9728765198671 | 14.3187385244159 | 2.13421729395427 |
| chr1 | 211573981 | 211573996 | - | 6.02376346124019 | 4.54798296825753 | 1.19322821036799 | 1.06710864697714 |
| chr1 | 211574750 | 211574776 | - | 16.0633692299738 | 26.1509020674808 | 7.15936926220794 | 1.06710864697714 |
| chr1 | 211575041 | 211575064 | - | 27.1069355755808 | 62.534765813541 | 1.19322821036799 | 3.20132594093141 |
| chr1 | 211662824 | 211662835 | - | 17.0673298068472 | 12.5069531627082 | 4.77291284147196 | 7.46976052883995 |
| chr1 | 211663073 | 211663129 | - | 24.0950538449607 | 10.2329616785794 | 7.15936926220794 | 13.8724124107028 |
| chr1 | 211663292 | 211663294 | - | 1.00396057687336 | 3.41098722619315 | 1.19322821036799 | 1.06710864697714 |
| chr1 | 211743643 | 211743655 | - | 232.918853834621 | 178.508331504108 | 90.6853439879672 | 83.2344744642166 |
| chr1 | 211743719 | 211743764 | - | 10.0396057687336 | 12.5069531627082 | 7.15936926220794 | 3.20132594093141 |
| chr1 | 211743790 | 211743791 | - | 1.00396057687336 | 2.27399148412876 | 1.19322821036799 | 1.06710864697714 |
| chr1 | 211744036 | 211744043 | - | 2.00792115374673 | 1.13699574206438 | 1.19322821036799 | 1.06710864697714 |
| chr1 | 211745623 | 211745659 | - | 11.043566345607 | 2.27399148412876 | 4.77291284147196 | 10.6710864697714 |
| chr1 | 211746021 | 211746064 | - | 2.00792115374673 | 4.54798296825753 | 2.38645642073598 | 2.13421729395427 |
| chr1 | 211746091 | 211746107 | - | 2.00792115374673 | 1.13699574206438 | 4.77291284147196 | 13.8724124107028 |
| chr1 | 211746363 | 211746414 | - | 4.01584230749346 | 3.41098722619315 | 1.19322821036799 | 10.6710864697714 |
| chr1 | 211746826 | 211746827 | - | 2.00792115374673 | 3.41098722619315 | 4.77291284147196 | 1.06710864697714 |
| chr1 | 211747322 | 211747401 | - | 1.00396057687336 | 3.41098722619315 | 1.19322821036799 | 2.13421729395427 |
| chr1 | 211747882 | 211747918 | - | 4.01584230749346 | 2.27399148412876 | 5.96614105183995 | 6.40265188186281 |
| chr1 | 211748044 | 211748055 | - | 2.00792115374673 | 5.68497871032191 | 3.57968463110397 | 1.06710864697714 |
| chr1 | 211940431 | 211940461 | - | 2.00792115374673 | 3.41098722619315 | 1.19322821036799 | 1.06710864697714 |
| chr1 | 211941343 | 211941346 | - | 1.00396057687336 | 2.27399148412876 | 2.38645642073598 | 1.06710864697714 |
| chr1 | 212102950 | 212102965 | + | 5.01980288436682 | 12.5069531627082 | 3.57968463110397 | 1.06710864697714 |
| chr1 | 212102987 | 212102992 | + | 3.01188173062009 | 15.9179403889013 | 7.15936926220794 | 1.06710864697714 |
| chr1 | 212103023 | 212103028 | + | 7.02772403811355 | 7.95897019445067 | 5.96614105183995 | 6.40265188186281 |
| chr1 | 212103131 | 212103157 | + | 10.0396057687336 | 18.1919318730301 | 5.96614105183995 | 4.26843458790854 |
| chr1 | 212103248 | 212103269 | + | 23.0910932680874 | 20.4659233571589 | 15.5119667347839 | 11.7381951167485 |
| chr1 | 212103306 | 212103337 | + | 175.693100952839 | 296.755888678804 | 35.7968463110397 | 59.7580842307196 |
| chr1 | 212104192 | 212104195 | + | 6.02376346124019 | 2.27399148412876 | 1.19322821036799 | 4.26843458790854 |
| chr1 | 212104379 | 212104381 | + | 5.01980288436682 | 6.82197445238629 | 5.96614105183995 | 4.26843458790854 |
| chr1 | 212104578 | 212104581 | + | 3.01188173062009 | 13.6439489047726 | 2.38645642073598 | 3.20132594093141 |
| chr1 | 212104761 | 212104763 | + | 4.01584230749346 | 1.13699574206438 | 2.38645642073598 | 1.06710864697714 |
| chr1 | 212360757 | 212360820 | + | 4.01584230749346 | 2.27399148412876 | 15.5119667347839 | 12.8053037637256 |
| chr1 | 212360851 | 212360888 | + | 2.00792115374673 | 5.68497871032191 | 23.8645642073598 | 12.8053037637256 |
| chr1 | 212360918 | 212360957 | + | 3.01188173062009 | 4.54798296825753 | 29.8307052591997 | 32.0132594093141 |
| chr1 | 212360971 | 212360984 | + | 1.00396057687336 | 2.27399148412876 | 2.38645642073598 | 3.20132594093141 |
| chr1 | 212361005 | 212361115 | + | 17.0673298068472 | 12.5069531627082 | 66.8207797806074 | 35.2145853502455 |
| chr1 | 212361574 | 212361614 | + | 11.043566345607 | 14.780944646837 | 19.0916513658878 | 32.0132594093141 |
| chr1 | 212361682 | 212361718 | + | 8.03168461498692 | 2.27399148412876 | 20.2848795762558 | 9.60397782279422 |
| chr1 | 212361774 | 212361802 | + | 1.00396057687336 | 3.41098722619315 | 2.38645642073598 | 11.7381951167485 |
| chr1 | 212364535 | 212364538 | - | 2.00792115374673 | 3.41098722619315 | 2.38645642073598 | 4.26843458790854 |
| chr1 | 212364590 | 212364607 | - | 1.00396057687336 | 1.13699574206438 | 1.19322821036799 | 1.06710864697714 |
| chr1 | 212365048 | 212365049 | - | 2.00792115374673 | 2.27399148412876 | 3.57968463110397 | 1.06710864697714 |
| chr1 | 212446089 | 212446095 | + | 9.03564519186028 | 15.9179403889013 | 13.1255103140479 | 17.0737383516342 |
| chr1 | 212726290 | 212726312 | - | 4.01584230749346 | 3.41098722619315 | 4.77291284147196 | 3.20132594093141 |
| chr1 | 212737846 | 212737866 | - | 2.00792115374673 | 3.41098722619315 | 4.77291284147196 | 1.06710864697714 |
| chr1 | 212737889 | 212737908 | - | 1.00396057687336 | 5.68497871032191 | 2.38645642073598 | 4.26843458790854 |
| chr1 | 212738610 | 212738663 | - | 2.00792115374673 | 4.54798296825753 | 3.57968463110397 | 3.20132594093141 |
| chr1 | 212739542 | 212739594 | - | 1.00396057687336 | 1.13699574206438 | 4.77291284147196 | 7.46976052883995 |
| chr1 | 212804318 | 212804324 | + | 5.01980288436682 | 5.68497871032191 | 15.5119667347839 | 4.26843458790854 |
| chr1 | 212815192 | 212815197 | + | 2.00792115374673 | 1.13699574206438 | 4.77291284147196 | 6.40265188186281 |
| chr1 | 212815247 | 212815272 | + | 3.01188173062009 | 10.2329616785794 | 5.96614105183995 | 3.20132594093141 |
| chr1 | 212816577 | 212816586 | + | 1.00396057687336 | 4.54798296825753 | 1.19322821036799 | 2.13421729395427 |
| chr1 | 212896368 | 212896403 | + | 2.00792115374673 | 1.13699574206438 | 1.19322821036799 | 1.06710864697714 |
| chr1 | 212899182 | 212899192 | + | 4.01584230749346 | 2.27399148412876 | 3.57968463110397 | 6.40265188186281 |
| chr1 | 212992239 | 212992251 | - | 1.00396057687336 | 2.27399148412876 | 1.19322821036799 | 1.06710864697714 |
| chr1 | 212993806 | 212993824 | - | 7.02772403811355 | 10.2329616785794 | 3.57968463110397 | 3.20132594093141 |
| chr1 | 212993959 | 212993961 | - | 7.02772403811355 | 7.95897019445067 | 7.15936926220794 | 5.33554323488568 |
| chr1 | 212994018 | 212994021 | - | 6.02376346124019 | 9.09596593651506 | 2.38645642073598 | 11.7381951167485 |
| chr1 | 213273392 | 213273401 | + | 4.01584230749346 | 9.09596593651506 | 7.15936926220794 | 10.6710864697714 |
| chr1 | 214337006 | 214337054 | + | 9.03564519186028 | 1.13699574206438 | 3.57968463110397 | 7.46976052883995 |
| chr1 | 214337080 | 214337114 | + | 2.00792115374673 | 3.41098722619315 | 1.19322821036799 | 3.20132594093141 |
| chr1 | 214663770 | 214663806 | + | 8.03168461498692 | 1.13699574206438 | 2.38645642073598 | 1.06710864697714 |
| chr1 | 214663876 | 214663899 | + | 8.03168461498692 | 9.09596593651506 | 11.9322821036799 | 6.40265188186281 |
| chr1 | 215620829 | 215620887 | + | 8.03168461498692 | 9.09596593651506 | 9.54582568294392 | 14.9395210576799 |
| chr1 | 215621145 | 215621202 | + | 9.03564519186028 | 11.3699574206438 | 16.7051949451519 | 8.53686917581709 |
| chr1 | 215621226 | 215621234 | + | 9.03564519186028 | 11.3699574206438 | 8.35259747257593 | 6.40265188186281 |
| chr1 | 215621263 | 215621443 | + | 342.350556713817 | 417.277437337628 | 309.046106485309 | 417.23948096806 |
| chr1 | 215621463 | 215621492 | + | 21.0831721143407 | 15.9179403889013 | 20.2848795762558 | 34.1474767032683 |
| chr1 | 215621699 | 215621710 | + | 9.03564519186028 | 9.09596593651506 | 3.57968463110397 | 4.26843458790854 |
| chr1 | 218331161 | 218331173 | + | 4.01584230749346 | 4.54798296825753 | 2.38645642073598 | 5.33554323488568 |
| chr1 | 218331337 | 218331357 | + | 2.00792115374673 | 6.82197445238629 | 11.9322821036799 | 2.13421729395427 |
| chr1 | 218331531 | 218331550 | + | 15.0594086531005 | 25.0139063254164 | 23.8645642073598 | 22.4092815865198 |
| chr1 | 218331959 | 218331975 | + | 3.01188173062009 | 4.54798296825753 | 11.9322821036799 | 7.46976052883995 |
| chr1 | 219968619 | 219968664 | - | 6.02376346124019 | 10.2329616785794 | 8.35259747257593 | 3.20132594093141 |
| chr1 | 219968794 | 219968819 | - | 26.1029749987075 | 37.5208594881246 | 38.1833027317757 | 41.6172372321083 |
| chr1 | 220057859 | 220057906 | - | 3.01188173062009 | 6.82197445238629 | 7.15936926220794 | 12.8053037637256 |
| chr1 | 220058060 | 220058069 | - | 2.00792115374673 | 2.27399148412876 | 2.38645642073598 | 13.8724124107028 |
| chr1 | 220147651 | 220147799 | + | 16.0633692299738 | 17.0549361309657 | 26.2510206280958 | 28.8119334683827 |
| chr1 | 220147898 | 220147903 | + | 2.00792115374673 | 1.13699574206438 | 2.38645642073598 | 1.06710864697714 |
| chr1 | 220147947 | 220147957 | + | 1.00396057687336 | 2.27399148412876 | 5.96614105183995 | 9.60397782279422 |
| chr1 | 220148489 | 220148525 | - | 9.03564519186028 | 17.0549361309657 | 22.6713359969918 | 23.476390233497 |
| chr1 | 220148563 | 220148601 | - | 16.0633692299738 | 13.6439489047726 | 46.5359002043516 | 25.6106075274513 |
| chr1 | 220148692 | 220148709 | - | 6.02376346124019 | 7.95897019445067 | 13.1255103140479 | 5.33554323488568 |
| chr1 | 220148743 | 220148750 | - | 1.00396057687336 | 4.54798296825753 | 1.19322821036799 | 2.13421729395427 |
| chr1 | 220148793 | 220148835 | - | 42.1663442286813 | 51.1648083928972 | 33.4103898903037 | 41.6172372321083 |
| chr1 | 220149006 | 220149053 | - | 5.01980288436682 | 6.82197445238629 | 8.35259747257593 | 1.06710864697714 |
| chr1 | 220149407 | 220149417 | - | 6.02376346124019 | 11.3699574206438 | 14.3187385244159 | 25.6106075274513 |
| chr1 | 220150642 | 220150645 | - | 5.01980288436682 | 3.41098722619315 | 16.7051949451519 | 6.40265188186281 |
| chr1 | 220150670 | 220150705 | - | 19.0752509605939 | 21.6029190992233 | 27.4442488384638 | 35.2145853502455 |
| chr1 | 220150742 | 220150771 | - | 73.2891221117556 | 85.2746806548286 | 46.5359002043516 | 117.381951167485 |
| chr1 | 220150843 | 220150850 | - | 7.02772403811355 | 4.54798296825753 | 4.77291284147196 | 1.06710864697714 |
| chr1 | 220151088 | 220151100 | - | 2.00792115374673 | 3.41098722619315 | 2.38645642073598 | 2.13421729395427 |
| chr1 | 220151159 | 220151206 | - | 11.043566345607 | 7.95897019445067 | 28.6374770488317 | 28.8119334683827 |
| chr1 | 221702355 | 221702363 | - | 6.02376346124019 | 3.41098722619315 | 1.19322821036799 | 1.06710864697714 |
| chr1 | 222558484 | 222558508 | - | 5.01980288436682 | 12.5069531627082 | 8.35259747257593 | 5.33554323488568 |
| chr1 | 222558569 | 222558571 | - | 3.01188173062009 | 5.68497871032191 | 3.57968463110397 | 2.13421729395427 |
| chr1 | 222558608 | 222558621 | - | 3.01188173062009 | 4.54798296825753 | 5.96614105183995 | 6.40265188186281 |
| chr1 | 222589920 | 222590030 | + | 90.3564519186028 | 335.413743908993 | 217.167534286974 | 189.94533916193 |
| chr1 | 222590176 | 222590230 | + | 15.0594086531005 | 10.2329616785794 | 90.6853439879672 | 56.5567582897882 |
| chr1 | 222590400 | 222590408 | + | 5.01980288436682 | 4.54798296825753 | 3.57968463110397 | 1.06710864697714 |
| chr1 | 222665997 | 222666001 | + | 5.01980288436682 | 5.68497871032191 | 9.54582568294392 | 3.20132594093141 |
| chr1 | 222733634 | 222733637 | + | 1.00396057687336 | 1.13699574206438 | 1.19322821036799 | 1.06710864697714 |
| chr1 | 222734744 | 222734768 | + | 12.0475269224804 | 13.6439489047726 | 4.77291284147196 | 19.2079556455884 |
| chr1 | 222734818 | 222734843 | + | 5.01980288436682 | 10.2329616785794 | 3.57968463110397 | 2.13421729395427 |
| chr1 | 222814994 | 222815165 | + | 389.536703826865 | 712.896330274368 | 1.19322821036799 | 6.40265188186281 |
| chr1 | 223181236 | 223181237 | + | 7.02772403811355 | 19.3289276150945 | 3.57968463110397 | 4.26843458790854 |
| chr1 | 223775685 | 223775688 | + | 14.0554480762271 | 17.0549361309657 | 13.1255103140479 | 34.1474767032683 |
| chr1 | 223775807 | 223775826 | + | 9.03564519186028 | 12.5069531627082 | 2.38645642073598 | 5.33554323488568 |
| chr1 | 223780386 | 223780403 | - | 3.01188173062009 | 4.54798296825753 | 4.77291284147196 | 4.26843458790854 |
| chr1 | 224159008 | 224159011 | + | 2.00792115374673 | 3.41098722619315 | 1.19322821036799 | 1.06710864697714 |
| chr1 | 224161561 | 224161565 | + | 44.174265382428 | 47.753821166704 | 15.5119667347839 | 10.6710864697714 |
| chr1 | 224161593 | 224161617 | + | 17.0673298068472 | 11.3699574206438 | 3.57968463110397 | 3.20132594093141 |
| chr1 | 224193146 | 224193163 | + | 17.0673298068472 | 6.82197445238629 | 2.38645642073598 | 10.6710864697714 |
| chr1 | 224193215 | 224193243 | + | 414.6357182487 | 449.113318115431 | 227.906588180286 | 182.47557863309 |
| chr1 | 224193270 | 224193330 | + | 150.594086531005 | 119.38455291676 | 90.6853439879672 | 38.4159112911769 |
| chr1 | 224227442 | 224227501 | - | 58.2297134586551 | 73.9047232341848 | 62.0478669391355 | 59.7580842307196 |
| chr1 | 224227527 | 224227535 | - | 44.174265382428 | 56.8497871032191 | 32.2171616799357 | 33.0803680562912 |
| chr1 | 224227556 | 224227611 | - | 27.1069355755808 | 47.753821166704 | 16.7051949451519 | 22.4092815865198 |
| chr1 | 224375826 | 224375845 | + | 17.0673298068472 | 13.6439489047726 | 3.57968463110397 | 8.53686917581709 |
| chr1 | 224375867 | 224375882 | + | 24.0950538449607 | 23.876910583352 | 17.8984231555198 | 20.2750642925656 |
| chr1 | 224375905 | 224375939 | + | 7.02772403811355 | 9.09596593651506 | 3.57968463110397 | 6.40265188186281 |
| chr1 | 224376076 | 224376095 | + | 2.00792115374673 | 4.54798296825753 | 1.19322821036799 | 2.13421729395427 |
| chr1 | 224385560 | 224385601 | - | 1.00396057687336 | 1.13699574206438 | 7.15936926220794 | 6.40265188186281 |
| chr1 | 224386900 | 224386923 | - | 4.01584230749346 | 2.27399148412876 | 4.77291284147196 | 4.26843458790854 |
| chr1 | 224388126 | 224388146 | - | 9.03564519186028 | 11.3699574206438 | 9.54582568294392 | 20.2750642925656 |
| chr1 | 224388169 | 224388203 | - | 2.00792115374673 | 2.27399148412876 | 5.96614105183995 | 5.33554323488568 |
| chr1 | 224389175 | 224389199 | - | 2.00792115374673 | 9.09596593651506 | 1.19322821036799 | 4.26843458790854 |
| chr1 | 224967833 | 224967850 | + | 2.00792115374673 | 2.27399148412876 | 4.77291284147196 | 12.8053037637256 |
| chr1 | 225401556 | 225401559 | - | 3.01188173062009 | 5.68497871032191 | 9.54582568294392 | 25.6106075274513 |
| chr1 | 225401598 | 225401727 | - | 384.516900942499 | 220.57717396049 | 410.470504366588 | 774.720877705401 |
| chr1 | 225401763 | 225401926 | - | 25.0990144218341 | 19.3289276150945 | 40.5697591525116 | 71.4962793474681 |
| chr1 | 225401949 | 225402017 | - | 9.03564519186028 | 11.3699574206438 | 9.54582568294392 | 21.3421729395427 |
| chr1 | 225402140 | 225402179 | - | 4.01584230749346 | 4.54798296825753 | 3.57968463110397 | 8.53686917581709 |
| chr1 | 225402305 | 225402338 | - | 3.01188173062009 | 6.82197445238629 | 14.3187385244159 | 82.1673658172394 |
| chr1 | 225402619 | 225402681 | - | 2.00792115374673 | 1.13699574206438 | 7.15936926220794 | 9.60397782279422 |
| chr1 | 225402732 | 225402828 | - | 7.02772403811355 | 1.13699574206438 | 11.9322821036799 | 20.2750642925656 |
| chr1 | 225402879 | 225402898 | - | 3.01188173062009 | 3.41098722619315 | 11.9322821036799 | 42.6843458790854 |
| chr1 | 225402981 | 225403035 | - | 7.02772403811355 | 5.68497871032191 | 31.0239334695677 | 46.952780466994 |
| chr1 | 225403063 | 225403090 | - | 4.01584230749346 | 4.54798296825753 | 10.7390538933119 | 4.26843458790854 |
| chr1 | 225403135 | 225403265 | - | 44.174265382428 | 21.6029190992233 | 102.617626091647 | 201.683534278679 |
| chr1 | 225496288 | 225496289 | - | 96.380215379843 | 79.5897019445067 | 71.5936926220794 | 58.6909755837425 |
| chr1 | 225497735 | 225497757 | - | 3.01188173062009 | 1.13699574206438 | 1.19322821036799 | 1.06710864697714 |
| chr1 | 225789431 | 225789494 | + | 9.03564519186028 | 13.6439489047726 | 29.8307052591997 | 50.1541064079254 |
| chr1 | 225789803 | 225789816 | + | 1.00396057687336 | 3.41098722619315 | 3.57968463110397 | 1.06710864697714 |
| chr1 | 225790011 | 225790025 | + | 4.01584230749346 | 6.82197445238629 | 4.77291284147196 | 13.8724124107028 |
| chr1 | 225790211 | 225790226 | + | 106.419821148577 | 152.357429436627 | 161.085808399679 | 144.059667341913 |
| chr1 | 225790364 | 225790380 | + | 27.1069355755808 | 26.1509020674808 | 42.9562155732476 | 54.4225409958339 |
| chr1 | 225846112 | 225846140 | - | 1.00396057687336 | 4.54798296825753 | 11.9322821036799 | 6.40265188186281 |
| chr1 | 225919906 | 225919948 | - | 2.00792115374673 | 19.3289276150945 | 20.2848795762558 | 13.8724124107028 |
| chr1 | 225919972 | 225919994 | - | 2.00792115374673 | 2.27399148412876 | 9.54582568294392 | 11.7381951167485 |
| chr1 | 225920023 | 225920035 | - | 3.01188173062009 | 1.13699574206438 | 14.3187385244159 | 12.8053037637256 |
| chr1 | 225920038 | 225920140 | - | 65.2574374967687 | 93.2336508492793 | 147.960298085631 | 268.911379038238 |
| chr1 | 225982722 | 225982736 | - | 2.00792115374673 | 7.95897019445067 | 3.57968463110397 | 4.26843458790854 |
| chr1 | 225983588 | 225983624 | - | 3.01188173062009 | 10.2329616785794 | 9.54582568294392 | 9.60397782279422 |
| chr1 | 226144821 | 226144851 | - | 9.03564519186028 | 6.82197445238629 | 7.15936926220794 | 5.33554323488568 |
| chr1 | 226145012 | 226145035 | - | 7.02772403811355 | 9.09596593651506 | 14.3187385244159 | 9.60397782279422 |
| chr1 | 226145912 | 226145958 | - | 4.01584230749346 | 4.54798296825753 | 17.8984231555198 | 7.46976052883995 |
| chr1 | 226146029 | 226146074 | - | 3.01188173062009 | 3.41098722619315 | 7.15936926220794 | 24.5434988804741 |
| chr1 | 226232213 | 226232234 | - | 7.02772403811355 | 4.54798296825753 | 7.15936926220794 | 8.53686917581709 |
| chr1 | 226232372 | 226232401 | - | 4.01584230749346 | 6.82197445238629 | 15.5119667347839 | 5.33554323488568 |
| chr1 | 226360839 | 226360842 | - | 3.01188173062009 | 2.27399148412876 | 2.38645642073598 | 2.13421729395427 |
| chr1 | 226360909 | 226360932 | - | 5.01980288436682 | 7.95897019445067 | 5.96614105183995 | 8.53686917581709 |
| chr1 | 226361109 | 226361115 | - | 68.2693192273888 | 69.3567402659273 | 47.7291284147196 | 80.0331485232852 |
| chr1 | 226361190 | 226361196 | - | 1.00396057687336 | 5.68497871032191 | 4.77291284147196 | 8.53686917581709 |
| chr1 | 226392206 | 226392315 | - | 51.2019894205416 | 44.3428339405109 | 99.0379414605431 | 87.5029090521251 |
| chr1 | 226895813 | 226895823 | + | 1.00396057687336 | 1.13699574206438 | 1.19322821036799 | 4.26843458790854 |
| chr1 | 226895849 | 226895987 | + | 139.550520185398 | 133.028501821533 | 79.9462900946553 | 108.845081991668 |
| chr1 | 226986765 | 226986817 | + | 6.02376346124019 | 6.82197445238629 | 8.35259747257593 | 18.1408469986113 |
| chr1 | 226986847 | 226986852 | + | 2.00792115374673 | 5.68497871032191 | 3.57968463110397 | 3.20132594093141 |
| chr1 | 226986897 | 226986937 | + | 9.03564519186028 | 1.13699574206438 | 4.77291284147196 | 7.46976052883995 |
| chr1 | 226986965 | 226987002 | + | 8.03168461498692 | 5.68497871032191 | 10.7390538933119 | 16.006629704657 |
| chr1 | 226987107 | 226987182 | + | 13.0514874993537 | 10.2329616785794 | 21.4781077866238 | 25.6106075274513 |
| chr1 | 226987211 | 226987281 | + | 8.03168461498692 | 2.27399148412876 | 5.96614105183995 | 17.0737383516342 |
| chr1 | 226987334 | 226987357 | + | 3.01188173062009 | 2.27399148412876 | 2.38645642073598 | 6.40265188186281 |
| chr1 | 226987420 | 226987484 | + | 7.02772403811355 | 12.5069531627082 | 9.54582568294392 | 8.53686917581709 |
| chr1 | 226989967 | 226989970 | - | 2.00792115374673 | 3.41098722619315 | 1.19322821036799 | 3.20132594093141 |
| chr1 | 226990085 | 226990125 | - | 45.1782259593014 | 42.0688424563821 | 19.0916513658878 | 141.925450047959 |
| chr1 | 226990154 | 226990177 | - | 2.00792115374673 | 1.13699574206438 | 2.38645642073598 | 11.7381951167485 |
| chr1 | 226990274 | 226990294 | - | 16.0633692299738 | 13.6439489047726 | 7.15936926220794 | 32.0132594093141 |
| chr1 | 226991492 | 226991511 | - | 1.00396057687336 | 3.41098722619315 | 2.38645642073598 | 2.13421729395427 |
| chr1 | 227731929 | 227731930 | - | 1.00396057687336 | 1.13699574206438 | 2.38645642073598 | 1.06710864697714 |
| chr1 | 227731977 | 227731986 | - | 2.00792115374673 | 1.13699574206438 | 7.15936926220794 | 2.13421729395427 |
| chr1 | 228059704 | 228059706 | + | 56.2217923049084 | 61.3977700714766 | 16.7051949451519 | 43.7514545260626 |
| chr1 | 228097340 | 228097365 | + | 4.01584230749346 | 1.13699574206438 | 4.77291284147196 | 5.33554323488568 |
| chr1 | 228097950 | 228098018 | + | 4.01584230749346 | 2.27399148412876 | 7.15936926220794 | 5.33554323488568 |
| chr1 | 228098121 | 228098198 | + | 7.02772403811355 | 4.54798296825753 | 17.8984231555198 | 23.476390233497 |
| chr1 | 228098281 | 228098347 | + | 19.0752509605939 | 18.1919318730301 | 62.0478669391355 | 57.6238669367653 |
| chr1 | 228098370 | 228098414 | + | 4.01584230749346 | 7.95897019445067 | 20.2848795762558 | 10.6710864697714 |
| chr1 | 228098464 | 228098479 | + | 1.00396057687336 | 1.13699574206438 | 10.7390538933119 | 10.6710864697714 |
| chr1 | 228098676 | 228098705 | + | 8.03168461498692 | 1.13699574206438 | 15.5119667347839 | 20.2750642925656 |
| chr1 | 228098712 | 228098743 | + | 1.00396057687336 | 1.13699574206438 | 3.57968463110397 | 13.8724124107028 |
| chr1 | 228098886 | 228098916 | + | 11.043566345607 | 14.780944646837 | 8.35259747257593 | 27.7448248214055 |
| chr1 | 228098947 | 228099119 | + | 289.140646139529 | 322.906790746284 | 496.382935513084 | 558.097822369042 |
| chr1 | 228099149 | 228099193 | + | 14.0554480762271 | 21.6029190992233 | 31.0239334695677 | 12.8053037637256 |
| chr1 | 228100787 | 228100840 | - | 29.1148567293276 | 80.7266976865711 | 20.2848795762558 | 38.4159112911769 |
| chr1 | 228106715 | 228106763 | - | 24.0950538449607 | 37.5208594881246 | 57.2749540976635 | 53.3554323488568 |
| chr1 | 228148846 | 228148875 | + | 39.1544624980612 | 30.6988850357383 | 16.7051949451519 | 42.6843458790854 |
| chr1 | 228148900 | 228148934 | + | 384.516900942499 | 382.030569333632 | 262.510206280958 | 569.83601748579 |
| chr1 | 228179061 | 228179079 | + | 21.0831721143407 | 22.7399148412876 | 19.0916513658878 | 30.9461507623369 |
| chr1 | 228182008 | 228182009 | + | 2.00792115374673 | 2.27399148412876 | 1.19322821036799 | 5.33554323488568 |
| chr1 | 228393754 | 228393783 | - | 4.01584230749346 | 6.82197445238629 | 5.96614105183995 | 3.20132594093141 |
| chr1 | 228393805 | 228393821 | - | 1.00396057687336 | 2.27399148412876 | 1.19322821036799 | 5.33554323488568 |
| chr1 | 228393932 | 228393984 | - | 3.01188173062009 | 4.54798296825753 | 7.15936926220794 | 11.7381951167485 |
| chr1 | 228394036 | 228394236 | - | 21.0831721143407 | 22.7399148412876 | 84.7192029361272 | 135.522798166096 |
| chr1 | 228394277 | 228394311 | - | 3.01188173062009 | 2.27399148412876 | 9.54582568294392 | 10.6710864697714 |
| chr1 | 228457031 | 228457046 | - | 2.00792115374673 | 5.68497871032191 | 9.54582568294392 | 10.6710864697714 |
| chr1 | 228457080 | 228457102 | - | 2.00792115374673 | 6.82197445238629 | 8.35259747257593 | 6.40265188186281 |
| chr1 | 228494679 | 228494713 | + | 2.00792115374673 | 2.27399148412876 | 1.19322821036799 | 2.13421729395427 |
| chr1 | 228494773 | 228494946 | + | 23.0910932680874 | 67.0827487817985 | 31.0239334695677 | 45.8856718200168 |
| chr1 | 229303890 | 229303994 | + | 15.0594086531005 | 18.1919318730301 | 21.4781077866238 | 25.6106075274513 |
| chr1 | 229304078 | 229304144 | + | 1.00396057687336 | 2.27399148412876 | 3.57968463110397 | 8.53686917581709 |
| chr1 | 229304186 | 229304239 | + | 1.00396057687336 | 5.68497871032191 | 11.9322821036799 | 3.20132594093141 |
| chr1 | 229304310 | 229304424 | + | 7.02772403811355 | 2.27399148412876 | 19.0916513658878 | 10.6710864697714 |
| chr1 | 229304595 | 229304605 | + | 10.0396057687336 | 5.68497871032191 | 13.1255103140479 | 21.3421729395427 |
| chr1 | 229304626 | 229304640 | + | 2.00792115374673 | 3.41098722619315 | 8.35259747257593 | 3.20132594093141 |
| chr1 | 229304650 | 229304691 | + | 2.00792115374673 | 1.13699574206438 | 5.96614105183995 | 2.13421729395427 |
| chr1 | 229305342 | 229305350 | + | 1.00396057687336 | 1.13699574206438 | 3.57968463110397 | 2.13421729395427 |
| chr1 | 229323091 | 229323109 | - | 7.02772403811355 | 3.41098722619315 | 20.2848795762558 | 24.5434988804741 |
| chr1 | 229323151 | 229323162 | - | 3.01188173062009 | 5.68497871032191 | 2.38645642073598 | 1.06710864697714 |
| chr1 | 229323356 | 229323378 | - | 5.01980288436682 | 7.95897019445067 | 22.6713359969918 | 6.40265188186281 |
| chr1 | 229323580 | 229323581 | - | 1.00396057687336 | 1.13699574206438 | 1.19322821036799 | 1.06710864697714 |
| chr1 | 229323856 | 229323872 | - | 18.0712903837206 | 28.4248935516096 | 33.4103898903037 | 35.2145853502455 |
| chr1 | 229324461 | 229324469 | - | 4.01584230749346 | 2.27399148412876 | 7.15936926220794 | 6.40265188186281 |
| chr1 | 229324584 | 229324610 | - | 6.02376346124019 | 3.41098722619315 | 11.9322821036799 | 4.26843458790854 |
| chr1 | 229441346 | 229441348 | - | 2.00792115374673 | 4.54798296825753 | 3.57968463110397 | 6.40265188186281 |
| chr1 | 229441376 | 229441394 | - | 6.02376346124019 | 3.41098722619315 | 7.15936926220794 | 12.8053037637256 |
| chr1 | 229441456 | 229441466 | - | 2.00792115374673 | 5.68497871032191 | 2.38645642073598 | 4.26843458790854 |
| chr1 | 229517013 | 229517034 | - | 3.01188173062009 | 2.27399148412876 | 11.9322821036799 | 20.2750642925656 |
| chr1 | 229517487 | 229517511 | - | 3.01188173062009 | 2.27399148412876 | 3.57968463110397 | 8.53686917581709 |
| chr1 | 229517595 | 229517615 | - | 2.00792115374673 | 2.27399148412876 | 3.57968463110397 | 5.33554323488568 |
| chr1 | 229518110 | 229518136 | - | 2.00792115374673 | 1.13699574206438 | 14.3187385244159 | 7.46976052883995 |
| chr1 | 229593352 | 229593354 | - | 3.01188173062009 | 9.09596593651506 | 1.19322821036799 | 6.40265188186281 |
| chr1 | 229599474 | 229599476 | - | 1.00396057687336 | 1.13699574206438 | 1.19322821036799 | 2.13421729395427 |
| chr1 | 229659871 | 229659889 | + | 4.01584230749346 | 2.27399148412876 | 4.77291284147196 | 1.06710864697714 |
| chr1 | 230279823 | 230279868 | + | 56.2217923049084 | 45.4798296825753 | 26.2510206280958 | 8.53686917581709 |
| chr1 | 230279898 | 230279938 | + | 15.0594086531005 | 5.68497871032191 | 3.57968463110397 | 1.06710864697714 |
| chr1 | 230279987 | 230279994 | + | 11.043566345607 | 6.82197445238629 | 4.77291284147196 | 3.20132594093141 |
| chr1 | 230280025 | 230280117 | + | 69.2732798042621 | 65.9457530397342 | 14.3187385244159 | 11.7381951167485 |
| chr1 | 230280157 | 230280201 | + | 15.0594086531005 | 6.82197445238629 | 3.57968463110397 | 3.20132594093141 |
| chr1 | 230280250 | 230280415 | + | 119.47130864793 | 113.699574206438 | 36.9900745214077 | 23.476390233497 |
| chr1 | 230280470 | 230280564 | + | 16.0633692299738 | 20.4659233571589 | 14.3187385244159 | 5.33554323488568 |
| chr1 | 230280609 | 230280630 | + | 15.0594086531005 | 1.13699574206438 | 9.54582568294392 | 2.13421729395427 |
| chr1 | 230280673 | 230280777 | + | 66.2613980736421 | 32.9728765198671 | 21.4781077866238 | 11.7381951167485 |
| chr1 | 230280817 | 230280842 | + | 19.0752509605939 | 5.68497871032191 | 4.77291284147196 | 9.60397782279422 |
| chr1 | 230281098 | 230281155 | + | 25.0990144218341 | 7.95897019445067 | 8.35259747257593 | 5.33554323488568 |
| chr1 | 230281233 | 230281267 | + | 35.1386201905678 | 15.9179403889013 | 11.9322821036799 | 3.20132594093141 |
| chr1 | 230281289 | 230281336 | + | 20.0792115374673 | 7.95897019445067 | 5.96614105183995 | 16.006629704657 |
| chr1 | 230281369 | 230281450 | + | 105.415860571703 | 25.0139063254164 | 22.6713359969918 | 26.6777161744284 |
| chr1 | 230281490 | 230281523 | + | 28.1108961524542 | 15.9179403889013 | 15.5119667347839 | 21.3421729395427 |
| chr1 | 230281664 | 230281997 | + | 3111.27382773056 | 2244.42959483509 | 1866.20892101554 | 1228.24205267068 |
| chr1 | 230282068 | 230282088 | + | 7.02772403811355 | 6.82197445238629 | 4.77291284147196 | 4.26843458790854 |
| chr1 | 230693449 | 230693460 | + | 6.02376346124019 | 5.68497871032191 | 4.77291284147196 | 6.40265188186281 |
| chr1 | 230693585 | 230693606 | + | 2.00792115374673 | 2.27399148412876 | 2.38645642073598 | 4.26843458790854 |
| chr1 | 230693649 | 230693694 | + | 6.02376346124019 | 10.2329616785794 | 10.7390538933119 | 7.46976052883995 |
| chr1 | 230693789 | 230693790 | + | 2.00792115374673 | 1.13699574206438 | 1.19322821036799 | 1.06710864697714 |
| chr1 | 230693836 | 230693861 | + | 17.0673298068472 | 13.6439489047726 | 10.7390538933119 | 19.2079556455884 |
| chr1 | 230693934 | 230693936 | + | 1.00396057687336 | 1.13699574206438 | 1.19322821036799 | 4.26843458790854 |
| chr1 | 230839598 | 230839601 | - | 1.00396057687336 | 3.41098722619315 | 4.77291284147196 | 1.06710864697714 |
| chr1 | 231277553 | 231277629 | + | 2.00792115374673 | 1.13699574206438 | 1.19322821036799 | 4.26843458790854 |
| chr1 | 231277726 | 231277783 | + | 11.043566345607 | 7.95897019445067 | 8.35259747257593 | 13.8724124107028 |
| chr1 | 231333006 | 231333026 | - | 3.01188173062009 | 1.13699574206438 | 1.19322821036799 | 1.06710864697714 |
| chr1 | 231335046 | 231335057 | - | 2.00792115374673 | 2.27399148412876 | 4.77291284147196 | 3.20132594093141 |
| chr1 | 231354165 | 231354166 | + | 1.00396057687336 | 1.13699574206438 | 1.19322821036799 | 1.06710864697714 |
| chr1 | 231363890 | 231363902 | - | 1.00396057687336 | 6.82197445238629 | 11.9322821036799 | 9.60397782279422 |
| chr1 | 231364029 | 231364052 | - | 5.01980288436682 | 5.68497871032191 | 3.57968463110397 | 5.33554323488568 |
| chr1 | 231364163 | 231364185 | - | 11.043566345607 | 13.6439489047726 | 5.96614105183995 | 8.53686917581709 |
| chr1 | 231364814 | 231364818 | - | 1.00396057687336 | 2.27399148412876 | 1.19322821036799 | 3.20132594093141 |
| chr1 | 231366086 | 231366094 | - | 2.00792115374673 | 4.54798296825753 | 2.38645642073598 | 3.20132594093141 |
| chr1 | 231366170 | 231366201 | - | 16.0633692299738 | 15.9179403889013 | 7.15936926220794 | 13.8724124107028 |
| chr1 | 231366293 | 231366321 | - | 5.01980288436682 | 9.09596593651506 | 7.15936926220794 | 4.26843458790854 |
| chr1 | 231565848 | 231565864 | + | 8.03168461498692 | 6.82197445238629 | 8.35259747257593 | 14.9395210576799 |
| chr1 | 231566033 | 231566040 | + | 3.01188173062009 | 7.95897019445067 | 3.57968463110397 | 4.26843458790854 |
| chr1 | 231566451 | 231566455 | + | 5.01980288436682 | 6.82197445238629 | 10.7390538933119 | 21.3421729395427 |
| chr1 | 232978288 | 232978303 | + | 1.00396057687336 | 1.13699574206438 | 3.57968463110397 | 3.20132594093141 |
| chr1 | 232978336 | 232978363 | + | 4.01584230749346 | 6.82197445238629 | 8.35259747257593 | 14.9395210576799 |
| chr1 | 232978439 | 232978450 | + | 9.03564519186028 | 11.3699574206438 | 5.96614105183995 | 4.26843458790854 |
| chr1 | 232978579 | 232978596 | + | 10.0396057687336 | 25.0139063254164 | 16.7051949451519 | 26.6777161744284 |
| chr1 | 232978645 | 232978671 | + | 5.01980288436682 | 5.68497871032191 | 8.35259747257593 | 6.40265188186281 |
| chr1 | 232978730 | 232978760 | + | 7.02772403811355 | 9.09596593651506 | 9.54582568294392 | 4.26843458790854 |
| chr1 | 233671528 | 233671585 | + | 71.2812009580089 | 93.2336508492793 | 14.3187385244159 | 19.2079556455884 |
| chr1 | 233671693 | 233671744 | + | 21.0831721143407 | 53.438799877026 | 15.5119667347839 | 16.006629704657 |
| chr1 | 233671767 | 233671768 | + | 4.01584230749346 | 14.780944646837 | 3.57968463110397 | 4.26843458790854 |
| chr1 | 233671793 | 233671865 | + | 31.1227778830743 | 55.7127913611547 | 9.54582568294392 | 17.0737383516342 |
| chr1 | 233671891 | 233671976 | + | 39.1544624980612 | 48.8908169087684 | 17.8984231555198 | 26.6777161744284 |
| chr1 | 233672076 | 233672139 | + | 22.087132691214 | 35.2468680039958 | 14.3187385244159 | 9.60397782279422 |
| chr1 | 233672185 | 233672211 | + | 204.807957682166 | 387.715548043954 | 125.288962088639 | 81.1002571702623 |
| chr1 | 234383856 | 234383917 | + | 9.03564519186028 | 9.09596593651506 | 2.38645642073598 | 8.53686917581709 |
| chr1 | 234383946 | 234384026 | + | 16.0633692299738 | 18.1919318730301 | 20.2848795762558 | 12.8053037637256 |
| chr1 | 234391406 | 234391452 | - | 1.00396057687336 | 1.13699574206438 | 1.19322821036799 | 1.06710864697714 |
| chr1 | 234604640 | 234604673 | - | 33.130699036821 | 40.9318467143178 | 9.54582568294392 | 12.8053037637256 |
| chr1 | 234605838 | 234605955 | - | 3.01188173062009 | 5.68497871032191 | 1.19322821036799 | 1.06710864697714 |
| chr1 | 234605994 | 234606005 | - | 18.0712903837206 | 23.876910583352 | 8.35259747257593 | 14.9395210576799 |
| chr1 | 234606169 | 234606448 | - | 24.0950538449607 | 35.2468680039958 | 10.7390538933119 | 13.8724124107028 |
| chr1 | 234606481 | 234606516 | - | 13.0514874993537 | 10.2329616785794 | 11.9322821036799 | 8.53686917581709 |
| chr1 | 234606567 | 234606695 | - | 35.1386201905678 | 51.1648083928972 | 41.7629873628796 | 49.0869977609482 |
| chr1 | 234606742 | 234607053 | - | 51.2019894205416 | 92.0966551072149 | 85.9124311464952 | 100.308212815851 |
| chr1 | 234607095 | 234607287 | - | 107.42378172545 | 176.234340019979 | 99.0379414605431 | 65.0936274656053 |
| chr1 | 234725459 | 234725460 | + | 1.00396057687336 | 1.13699574206438 | 1.19322821036799 | 2.13421729395427 |
| chr1 | 235109466 | 235109505 | - | 5.01980288436682 | 3.41098722619315 | 8.35259747257593 | 6.40265188186281 |
| chr1 | 235109594 | 235109639 | - | 3.01188173062009 | 6.82197445238629 | 13.1255103140479 | 5.33554323488568 |
| chr1 | 235109739 | 235109800 | - | 5.01980288436682 | 2.27399148412876 | 8.35259747257593 | 13.8724124107028 |
| chr1 | 235109883 | 235109901 | - | 5.01980288436682 | 2.27399148412876 | 3.57968463110397 | 4.26843458790854 |
| chr1 | 235110120 | 235110185 | - | 5.01980288436682 | 15.9179403889013 | 31.0239334695677 | 17.0737383516342 |
| chr1 | 235110212 | 235110262 | - | 3.01188173062009 | 15.9179403889013 | 70.4004644117114 | 46.952780466994 |
| chr1 | 235110303 | 235110356 | - | 24.0950538449607 | 34.1098722619315 | 73.9801490428153 | 57.6238669367653 |
| chr1 | 235110468 | 235110513 | - | 24.0950538449607 | 35.2468680039958 | 152.733210927103 | 100.308212815851 |
| chr1 | 235110538 | 235110582 | - | 11.043566345607 | 1.13699574206438 | 35.7968463110397 | 17.0737383516342 |
| chr1 | 235110657 | 235110759 | - | 10.0396057687336 | 9.09596593651506 | 33.4103898903037 | 24.5434988804741 |
| chr1 | 235111012 | 235111122 | - | 3.01188173062009 | 11.3699574206438 | 10.7390538933119 | 13.8724124107028 |
| chr1 | 235111158 | 235111220 | - | 7.02772403811355 | 9.09596593651506 | 20.2848795762558 | 13.8724124107028 |
| chr1 | 235111265 | 235111278 | - | 5.01980288436682 | 2.27399148412876 | 10.7390538933119 | 4.26843458790854 |
| chr1 | 235111396 | 235111447 | - | 7.02772403811355 | 2.27399148412876 | 20.2848795762558 | 13.8724124107028 |
| chr1 | 235111479 | 235111532 | - | 7.02772403811355 | 13.6439489047726 | 46.5359002043516 | 56.5567582897882 |
| chr1 | 235111570 | 235111704 | - | 19.0752509605939 | 23.876910583352 | 122.902505667903 | 30.9461507623369 |
| chr1 | 235111797 | 235111882 | - | 6.02376346124019 | 4.54798296825753 | 28.6374770488317 | 6.40265188186281 |
| chr1 | 235131653 | 235131884 | - | 334.31887209883 | 487.77117334562 | 760.086370004409 | 514.346367842979 |
| chr1 | 235166916 | 235166932 | - | 1.00396057687336 | 3.41098722619315 | 1.19322821036799 | 4.26843458790854 |
| chr1 | 235167241 | 235167247 | - | 4.01584230749346 | 3.41098722619315 | 1.19322821036799 | 1.06710864697714 |
| chr1 | 235167504 | 235167513 | - | 1.00396057687336 | 3.41098722619315 | 4.77291284147196 | 5.33554323488568 |
| chr1 | 235342885 | 235342897 | + | 6.02376346124019 | 2.27399148412876 | 10.7390538933119 | 8.53686917581709 |
| chr1 | 235343014 | 235343060 | + | 2.00792115374673 | 1.13699574206438 | 2.38645642073598 | 6.40265188186281 |
| chr1 | 236483449 | 236483450 | + | 29.1148567293276 | 13.6439489047726 | 54.8884976769275 | 50.1541064079254 |
| chr1 | 236550691 | 236550695 | - | 2.00792115374673 | 3.41098722619315 | 5.96614105183995 | 3.20132594093141 |
| chr1 | 236550797 | 236550835 | - | 19.0752509605939 | 15.9179403889013 | 10.7390538933119 | 8.53686917581709 |
| chr1 | 236550865 | 236550930 | - | 17.0673298068472 | 7.95897019445067 | 13.1255103140479 | 22.4092815865198 |
| chr1 | 236898742 | 236898792 | + | 2.00792115374673 | 4.54798296825753 | 2.38645642073598 | 3.20132594093141 |
| chr1 | 236899417 | 236899428 | + | 4.01584230749346 | 7.95897019445067 | 2.38645642073598 | 1.06710864697714 |
| chr1 | 236900093 | 236900100 | + | 3.01188173062009 | 4.54798296825753 | 3.57968463110397 | 3.20132594093141 |
| chr1 | 236900313 | 236900327 | + | 20.0792115374673 | 15.9179403889013 | 10.7390538933119 | 16.006629704657 |
| chr1 | 236900466 | 236900547 | + | 24.0950538449607 | 39.7948509722534 | 46.5359002043516 | 27.7448248214055 |
| chr1 | 236900646 | 236900694 | + | 9.03564519186028 | 13.6439489047726 | 14.3187385244159 | 9.60397782279422 |
| chr1 | 236900770 | 236900779 | + | 4.01584230749346 | 3.41098722619315 | 3.57968463110397 | 4.26843458790854 |
| chr1 | 236900863 | 236900879 | + | 8.03168461498692 | 7.95897019445067 | 11.9322821036799 | 2.13421729395427 |
| chr1 | 236901099 | 236901119 | + | 1.00396057687336 | 2.27399148412876 | 1.19322821036799 | 2.13421729395427 |
| chr1 | 236901247 | 236901262 | + | 7.02772403811355 | 7.95897019445067 | 19.0916513658878 | 10.6710864697714 |
| chr1 | 236901298 | 236901312 | + | 5.01980288436682 | 6.82197445238629 | 11.9322821036799 | 7.46976052883995 |
| chr1 | 236901713 | 236901721 | + | 2.00792115374673 | 4.54798296825753 | 1.19322821036799 | 2.13421729395427 |
| chr1 | 236903189 | 236903228 | + | 16.0633692299738 | 22.7399148412876 | 25.0577924177278 | 23.476390233497 |
| chr1 | 236903328 | 236903333 | + | 17.0673298068472 | 38.657855230189 | 31.0239334695677 | 35.2145853502455 |
| chr1 | 236903353 | 236903398 | + | 7.02772403811355 | 6.82197445238629 | 9.54582568294392 | 7.46976052883995 |
| chr1 | 236903549 | 236903598 | + | 14.0554480762271 | 11.3699574206438 | 19.0916513658878 | 19.2079556455884 |
| chr1 | 241497669 | 241497681 | - | 23.0910932680874 | 36.3838637460602 | 14.3187385244159 | 12.8053037637256 |
| chr1 | 241497806 | 241497918 | - | 127.502993262917 | 183.056314472366 | 146.767069875263 | 139.791232754005 |
| chr1 | 241593242 | 241593245 | - | 1.00396057687336 | 3.41098722619315 | 1.19322821036799 | 1.06710864697714 |
| chr1 | 241593338 | 241593379 | - | 29.1148567293276 | 26.1509020674808 | 14.3187385244159 | 2.13421729395427 |
| chr1 | 241593497 | 241593517 | - | 7.02772403811355 | 7.95897019445067 | 7.15936926220794 | 1.06710864697714 |
| chr1 | 241593550 | 241593588 | - | 241.954499026481 | 187.604297440623 | 162.279036610047 | 102.442430109805 |
| chr1 | 241593634 | 241593638 | - | 39.1544624980612 | 28.4248935516096 | 4.77291284147196 | 7.46976052883995 |
| chr1 | 241593680 | 241593711 | - | 65.2574374967687 | 50.0278126508328 | 27.4442488384638 | 24.5434988804741 |
| chr1 | 241629160 | 241629164 | - | 16.0633692299738 | 18.1919318730301 | 9.54582568294392 | 14.9395210576799 |
| chr1 | 241629758 | 241629791 | - | 5.01980288436682 | 5.68497871032191 | 9.54582568294392 | 14.9395210576799 |
| chr1 | 241629846 | 241629857 | - | 4.01584230749346 | 5.68497871032191 | 16.7051949451519 | 6.40265188186281 |
| chr1 | 241629886 | 241630005 | - | 137.542599031651 | 154.631420920756 | 405.697591525116 | 287.05222603685 |
| chr1 | 241630093 | 241630142 | - | 33.130699036821 | 60.2607743294122 | 73.9801490428153 | 27.7448248214055 |
| chr1 | 241630175 | 241630192 | - | 10.0396057687336 | 12.5069531627082 | 17.8984231555198 | 14.9395210576799 |
| chr1 | 241630238 | 241630287 | - | 55.217831728035 | 69.3567402659273 | 84.7192029361272 | 104.576647403759 |
| chr1 | 241630411 | 241630418 | - | 8.03168461498692 | 5.68497871032191 | 7.15936926220794 | 1.06710864697714 |
| chr1 | 241630574 | 241630589 | - | 1.00396057687336 | 2.27399148412876 | 17.8984231555198 | 8.53686917581709 |
| chr1 | 241630653 | 241630693 | - | 3.01188173062009 | 11.3699574206438 | 4.77291284147196 | 11.7381951167485 |
| chr1 | 241630871 | 241630884 | - | 5.01980288436682 | 4.54798296825753 | 4.77291284147196 | 4.26843458790854 |
| chr1 | 241631037 | 241631042 | - | 3.01188173062009 | 5.68497871032191 | 5.96614105183995 | 7.46976052883995 |
| chr1 | 241631110 | 241631116 | - | 3.01188173062009 | 4.54798296825753 | 5.96614105183995 | 3.20132594093141 |
| chr1 | 241631145 | 241631149 | - | 8.03168461498692 | 5.68497871032191 | 5.96614105183995 | 3.20132594093141 |
| chr1 | 241631189 | 241631193 | - | 4.01584230749346 | 4.54798296825753 | 4.77291284147196 | 5.33554323488568 |
| chr1 | 241631224 | 241631252 | - | 2.00792115374673 | 3.41098722619315 | 7.15936926220794 | 3.20132594093141 |
| chr1 | 241631360 | 241631402 | - | 28.1108961524542 | 37.5208594881246 | 20.2848795762558 | 25.6106075274513 |
| chr1 | 241631590 | 241631598 | - | 3.01188173062009 | 5.68497871032191 | 8.35259747257593 | 1.06710864697714 |
| chr1 | 241631637 | 241631639 | - | 1.00396057687336 | 1.13699574206438 | 2.38645642073598 | 1.06710864697714 |
| chr1 | 241631779 | 241631821 | - | 102.403978841083 | 121.658544400889 | 60.8546387287675 | 29.8790421153598 |
| chr1 | 241631939 | 241631954 | - | 3.01188173062009 | 2.27399148412876 | 2.38645642073598 | 6.40265188186281 |
| chr1 | 241632121 | 241632127 | - | 6.02376346124019 | 5.68497871032191 | 8.35259747257593 | 4.26843458790854 |
| chr1 | 241632184 | 241632187 | - | 4.01584230749346 | 1.13699574206438 | 8.35259747257593 | 1.06710864697714 |
| chr1 | 241632216 | 241632225 | - | 1.00396057687336 | 4.54798296825753 | 7.15936926220794 | 3.20132594093141 |
| chr1 | 241632924 | 241632944 | - | 5.01980288436682 | 4.54798296825753 | 1.19322821036799 | 2.13421729395427 |
| chr1 | 241633303 | 241633308 | - | 1.00396057687336 | 10.2329616785794 | 3.57968463110397 | 2.13421729395427 |
| chr1 | 241635625 | 241635646 | - | 6.02376346124019 | 9.09596593651506 | 2.38645642073598 | 8.53686917581709 |
| chr1 | 241635668 | 241635679 | - | 9.03564519186028 | 11.3699574206438 | 13.1255103140479 | 7.46976052883995 |
| chr1 | 241635719 | 241635724 | - | 3.01188173062009 | 3.41098722619315 | 2.38645642073598 | 1.06710864697714 |
| chr1 | 241635760 | 241635762 | - | 18.0712903837206 | 7.95897019445067 | 8.35259747257593 | 12.8053037637256 |
| chr1 | 241635905 | 241635908 | - | 2.00792115374673 | 4.54798296825753 | 3.57968463110397 | 1.06710864697714 |
| chr1 | 241635974 | 241635976 | - | 1.00396057687336 | 1.13699574206438 | 1.19322821036799 | 1.06710864697714 |
| chr1 | 241889471 | 241889586 | + | 169.669337491599 | 341.098722619315 | 190.916513658878 | 184.609795927044 |
| chr1 | 241889662 | 241889683 | + | 5.01980288436682 | 7.95897019445067 | 1.19322821036799 | 3.20132594093141 |
| chr1 | 244408904 | 244408916 | - | 7.02772403811355 | 6.82197445238629 | 2.38645642073598 | 3.20132594093141 |
| chr1 | 244408990 | 244409012 | - | 26.1029749987075 | 18.1919318730301 | 9.54582568294392 | 6.40265188186281 |
| chr1 | 244409215 | 244409216 | - | 1.00396057687336 | 4.54798296825753 | 5.96614105183995 | 3.20132594093141 |
| chr1 | 244409303 | 244409358 | - | 7.02772403811355 | 5.68497871032191 | 20.2848795762558 | 12.8053037637256 |
| chr1 | 244409387 | 244409405 | - | 1.00396057687336 | 2.27399148412876 | 5.96614105183995 | 4.26843458790854 |
| chr1 | 244409439 | 244409488 | - | 10.0396057687336 | 5.68497871032191 | 26.2510206280958 | 11.7381951167485 |
| chr1 | 244409544 | 244409551 | - | 13.0514874993537 | 10.2329616785794 | 14.3187385244159 | 13.8724124107028 |
| chr1 | 244705958 | 244705985 | + | 6.02376346124019 | 4.54798296825753 | 9.54582568294392 | 7.46976052883995 |
| chr1 | 244706461 | 244706467 | + | 1.00396057687336 | 5.68497871032191 | 4.77291284147196 | 2.13421729395427 |
| chr1 | 244707430 | 244707487 | + | 13.0514874993537 | 6.82197445238629 | 8.35259747257593 | 16.006629704657 |
| chr1 | 244707648 | 244707663 | + | 5.01980288436682 | 1.13699574206438 | 9.54582568294392 | 2.13421729395427 |
| chr1 | 244707746 | 244707796 | + | 2.00792115374673 | 6.82197445238629 | 19.0916513658878 | 10.6710864697714 |
| chr1 | 244708196 | 244708236 | + | 3.01188173062009 | 7.95897019445067 | 5.96614105183995 | 4.26843458790854 |
| chr1 | 244708480 | 244708496 | + | 3.01188173062009 | 5.68497871032191 | 1.19322821036799 | 1.06710864697714 |
| chr1 | 244843147 | 244843189 | + | 2.00792115374673 | 1.13699574206438 | 2.38645642073598 | 8.53686917581709 |
| chr1 | 244843403 | 244843426 | + | 13.0514874993537 | 5.68497871032191 | 9.54582568294392 | 22.4092815865198 |
| chr1 | 244843682 | 244843701 | + | 6.02376346124019 | 7.95897019445067 | 2.38645642073598 | 2.13421729395427 |
| chr1 | 244843945 | 244843968 | + | 7.02772403811355 | 6.82197445238629 | 4.77291284147196 | 4.26843458790854 |
| chr1 | 244844074 | 244844076 | + | 4.01584230749346 | 2.27399148412876 | 7.15936926220794 | 2.13421729395427 |
| chr1 | 244844143 | 244844162 | + | 2.00792115374673 | 3.41098722619315 | 1.19322821036799 | 3.20132594093141 |
| chr1 | 244844281 | 244844301 | + | 6.02376346124019 | 7.95897019445067 | 5.96614105183995 | 10.6710864697714 |
| chr1 | 244844352 | 244844360 | + | 50.1980288436682 | 50.0278126508328 | 31.0239334695677 | 44.8185631730397 |
| chr1 | 244850389 | 244850390 | - | 1.00396057687336 | 2.27399148412876 | 2.38645642073598 | 1.06710864697714 |
| chr1 | 244850470 | 244850478 | - | 11.043566345607 | 9.09596593651506 | 10.7390538933119 | 4.26843458790854 |
| chr1 | 244851311 | 244851316 | - | 3.01188173062009 | 7.95897019445067 | 1.19322821036799 | 2.13421729395427 |
| chr1 | 244851426 | 244851450 | - | 7.02772403811355 | 6.82197445238629 | 9.54582568294392 | 4.26843458790854 |
| chr1 | 244851466 | 244851491 | - | 1.00396057687336 | 6.82197445238629 | 4.77291284147196 | 1.06710864697714 |
| chr1 | 244852472 | 244852506 | - | 3.01188173062009 | 2.27399148412876 | 3.57968463110397 | 2.13421729395427 |
| chr1 | 244852687 | 244852718 | - | 5.01980288436682 | 9.09596593651506 | 7.15936926220794 | 2.13421729395427 |
| chr1 | 244852795 | 244852816 | - | 4.01584230749346 | 6.82197445238629 | 1.19322821036799 | 1.06710864697714 |
| chr1 | 244853041 | 244853046 | - | 4.01584230749346 | 1.13699574206438 | 1.19322821036799 | 5.33554323488568 |
| chr1 | 244853138 | 244853160 | - | 5.01980288436682 | 1.13699574206438 | 1.19322821036799 | 3.20132594093141 |
| chr1 | 244853416 | 244853441 | - | 37.1465413443145 | 106.877599754052 | 27.4442488384638 | 70.429170700491 |
| chr1 | 244853483 | 244853486 | - | 1.00396057687336 | 5.68497871032191 | 4.77291284147196 | 5.33554323488568 |
| chr1 | 244853523 | 244853541 | - | 7.02772403811355 | 21.6029190992233 | 10.7390538933119 | 6.40265188186281 |
| chr1 | 244853567 | 244853703 | - | 159.629731722865 | 293.344901452611 | 215.974306076606 | 141.925450047959 |
| chr1 | 244853747 | 244853772 | - | 115.455466340437 | 193.289276150945 | 47.7291284147196 | 51.2212150549025 |
| chr1 | 244853816 | 244853844 | - | 27.1069355755808 | 47.753821166704 | 21.4781077866238 | 25.6106075274513 |
| chr1 | 244853869 | 244853963 | - | 75.2970432655023 | 104.603608269923 | 97.8447132501751 | 112.046407932599 |
| chr1 | 244853989 | 244854018 | - | 5.01980288436682 | 5.68497871032191 | 8.35259747257593 | 4.26843458790854 |
| chr1 | 244854077 | 244854093 | - | 5.01980288436682 | 6.82197445238629 | 9.54582568294392 | 24.5434988804741 |
| chr1 | 244854181 | 244854208 | - | 30.1188173062009 | 47.753821166704 | 22.6713359969918 | 27.7448248214055 |
| chr1 | 244854231 | 244854245 | - | 9.03564519186028 | 10.2329616785794 | 7.15936926220794 | 22.4092815865198 |
| chr1 | 244854312 | 244854326 | - | 85.336649034236 | 96.6446380754725 | 44.1494437836156 | 70.429170700491 |
| chr1 | 244854347 | 244854404 | - | 41.1623836518079 | 44.3428339405109 | 59.6614105183995 | 148.328101929822 |
| chr1 | 244854422 | 244854498 | - | 71.2812009580089 | 96.6446380754725 | 85.9124311464952 | 115.247733873531 |
| chr1 | 245087716 | 245087719 | + | 7.02772403811355 | 10.2329616785794 | 3.57968463110397 | 8.53686917581709 |
| chr1 | 245087801 | 245087809 | + | 2.00792115374673 | 1.13699574206438 | 5.96614105183995 | 3.20132594093141 |
| chr1 | 246540634 | 246540670 | - | 10.0396057687336 | 15.9179403889013 | 11.9322821036799 | 5.33554323488568 |
| chr1 | 246541003 | 246541009 | - | 35.1386201905678 | 42.0688424563821 | 9.54582568294392 | 17.0737383516342 |
| chr1 | 246668045 | 246668057 | + | 3.01188173062009 | 12.5069531627082 | 16.7051949451519 | 24.5434988804741 |
| chr1 | 246668372 | 246668380 | + | 4.01584230749346 | 1.13699574206438 | 8.35259747257593 | 9.60397782279422 |
| chr1 | 246668441 | 246668482 | + | 4.01584230749346 | 6.82197445238629 | 17.8984231555198 | 6.40265188186281 |
| chr1 | 247298369 | 247298390 | - | 3.01188173062009 | 7.95897019445067 | 10.7390538933119 | 5.33554323488568 |
| chr1 | 247298427 | 247298430 | - | 7.02772403811355 | 13.6439489047726 | 7.15936926220794 | 12.8053037637256 |
| chr1 | 247298465 | 247298484 | - | 1.00396057687336 | 4.54798296825753 | 7.15936926220794 | 6.40265188186281 |
| chr1 | 247298560 | 247298576 | - | 3.01188173062009 | 4.54798296825753 | 9.54582568294392 | 8.53686917581709 |
| chr1 | 247298658 | 247298673 | - | 4.01584230749346 | 2.27399148412876 | 8.35259747257593 | 5.33554323488568 |
| chr1 | 247298849 | 247298871 | - | 1.00396057687336 | 4.54798296825753 | 13.1255103140479 | 5.33554323488568 |
| chr1 | 247299167 | 247299172 | - | 2.00792115374673 | 5.68497871032191 | 3.57968463110397 | 3.20132594093141 |
| chr1 | 247299456 | 247299466 | - | 2.00792115374673 | 1.13699574206438 | 3.57968463110397 | 2.13421729395427 |
| chr1 | 247299802 | 247299817 | - | 4.01584230749346 | 3.41098722619315 | 8.35259747257593 | 2.13421729395427 |
| chr1 | 248522644 | 248522645 | + | 2.00792115374673 | 2.27399148412876 | 4.77291284147196 | 6.40265188186281 |
| chr1 | 248810490 | 248810500 | - | 6.02376346124019 | 10.2329616785794 | 5.96614105183995 | 7.46976052883995 |
| chr1 | 248810530 | 248810542 | - | 7.02772403811355 | 5.68497871032191 | 4.77291284147196 | 1.06710864697714 |
| chr1 | 248811476 | 248811481 | - | 4.01584230749346 | 2.27399148412876 | 3.57968463110397 | 1.06710864697714 |
| chr1 | 248849422 | 248849487 | + | 12.0475269224804 | 18.1919318730301 | 8.35259747257593 | 21.3421729395427 |
| chr1 | 248850032 | 248850099 | - | 34.1346596136944 | 60.2607743294122 | 57.2749540976635 | 77.8989312293309 |
| chr1 | 248918853 | 248918862 | + | 7.02772403811355 | 6.82197445238629 | 2.38645642073598 | 4.26843458790854 |
| chr10 | 252972 | 253018 | + | 4.01584230749346 | 4.54798296825753 | 1.19322821036799 | 2.13421729395427 |
| chr10 | 809692 | 809721 | - | 19.0752509605939 | 21.6029190992233 | 16.7051949451519 | 33.0803680562912 |
| chr10 | 809837 | 809855 | - | 1.00396057687336 | 1.13699574206438 | 1.19322821036799 | 4.26843458790854 |
| chr10 | 810200 | 810212 | - | 3.01188173062009 | 1.13699574206438 | 9.54582568294392 | 5.33554323488568 |
| chr10 | 810242 | 810243 | - | 3.01188173062009 | 4.54798296825753 | 4.77291284147196 | 7.46976052883995 |
| chr10 | 810835 | 810887 | - | 3.01188173062009 | 3.41098722619315 | 15.5119667347839 | 49.0869977609482 |
| chr10 | 811169 | 811176 | - | 6.02376346124019 | 3.41098722619315 | 2.38645642073598 | 7.46976052883995 |
| chr10 | 811222 | 811258 | - | 16.0633692299738 | 14.780944646837 | 14.3187385244159 | 14.9395210576799 |
| chr10 | 811323 | 811324 | - | 1.00396057687336 | 1.13699574206438 | 1.19322821036799 | 1.06710864697714 |
| chr10 | 811355 | 811367 | - | 5.01980288436682 | 3.41098722619315 | 5.96614105183995 | 4.26843458790854 |
| chr10 | 811545 | 811561 | - | 11.043566345607 | 6.82197445238629 | 13.1255103140479 | 14.9395210576799 |
| chr10 | 811932 | 811936 | - | 4.01584230749346 | 4.54798296825753 | 3.57968463110397 | 1.06710864697714 |
| chr10 | 811965 | 811996 | - | 5.01980288436682 | 2.27399148412876 | 4.77291284147196 | 8.53686917581709 |
| chr10 | 812567 | 812617 | - | 5.01980288436682 | 1.13699574206438 | 13.1255103140479 | 60.8251928776967 |
| chr10 | 812689 | 812705 | - | 7.02772403811355 | 3.41098722619315 | 8.35259747257593 | 8.53686917581709 |
| chr10 | 1017222 | 1017309 | + | 6.02376346124019 | 7.95897019445067 | 3.57968463110397 | 10.6710864697714 |
| chr10 | 1017480 | 1017487 | + | 3.01188173062009 | 4.54798296825753 | 1.19322821036799 | 3.20132594093141 |
| chr10 | 1017517 | 1017524 | + | 10.0396057687336 | 12.5069531627082 | 7.15936926220794 | 5.33554323488568 |
| chr10 | 1017549 | 1017636 | + | 3.01188173062009 | 12.5069531627082 | 13.1255103140479 | 14.9395210576799 |
| chr10 | 1040234 | 1040297 | - | 28.1108961524542 | 46.6168254246397 | 84.7192029361272 | 119.516168461439 |
| chr10 | 1040356 | 1040407 | - | 1.00396057687336 | 2.27399148412876 | 9.54582568294392 | 5.33554323488568 |
| chr10 | 1040796 | 1040817 | - | 3.01188173062009 | 2.27399148412876 | 3.57968463110397 | 5.33554323488568 |
| chr10 | 1040848 | 1040893 | - | 1.00396057687336 | 3.41098722619315 | 4.77291284147196 | 4.26843458790854 |
| chr10 | 1041024 | 1041025 | - | 4.01584230749346 | 1.13699574206438 | 1.19322821036799 | 1.06710864697714 |
| chr10 | 3136721 | 3136740 | + | 20.0792115374673 | 13.6439489047726 | 9.54582568294392 | 9.60397782279422 |
| chr10 | 3136766 | 3136773 | + | 13.0514874993537 | 23.876910583352 | 9.54582568294392 | 8.53686917581709 |
| chr10 | 3776779 | 3776885 | - | 5.01980288436682 | 3.41098722619315 | 3.57968463110397 | 14.9395210576799 |
| chr10 | 3777017 | 3777082 | - | 3.01188173062009 | 2.27399148412876 | 9.54582568294392 | 1.06710864697714 |
| chr10 | 3777281 | 3777319 | - | 4.01584230749346 | 3.41098722619315 | 5.96614105183995 | 13.8724124107028 |
| chr10 | 3777338 | 3777474 | - | 5.01980288436682 | 13.6439489047726 | 28.6374770488317 | 20.2750642925656 |
| chr10 | 3777503 | 3777564 | - | 2.00792115374673 | 3.41098722619315 | 5.96614105183995 | 2.13421729395427 |
| chr10 | 3777859 | 3777912 | - | 1.00396057687336 | 3.41098722619315 | 1.19322821036799 | 1.06710864697714 |
| chr10 | 3778238 | 3778269 | - | 12.0475269224804 | 7.95897019445067 | 33.4103898903037 | 7.46976052883995 |
| chr10 | 3778411 | 3778420 | - | 1.00396057687336 | 5.68497871032191 | 3.57968463110397 | 11.7381951167485 |
| chr10 | 3778464 | 3778493 | - | 8.03168461498692 | 3.41098722619315 | 4.77291284147196 | 6.40265188186281 |
| chr10 | 3778553 | 3778595 | - | 1.00396057687336 | 1.13699574206438 | 1.19322821036799 | 5.33554323488568 |
| chr10 | 3778647 | 3778664 | - | 2.00792115374673 | 1.13699574206438 | 2.38645642073598 | 3.20132594093141 |
| chr10 | 3778884 | 3778998 | - | 4.01584230749346 | 3.41098722619315 | 2.38645642073598 | 3.20132594093141 |
| chr10 | 3779077 | 3779090 | - | 79.3128855729958 | 110.288586980245 | 238.645642073598 | 105.643756050736 |
| chr10 | 3779206 | 3779304 | - | 22.087132691214 | 15.9179403889013 | 36.9900745214077 | 24.5434988804741 |
| chr10 | 3779330 | 3779407 | - | 23.0910932680874 | 25.0139063254164 | 20.2848795762558 | 19.2079556455884 |
| chr10 | 3779449 | 3779538 | - | 6.02376346124019 | 13.6439489047726 | 7.15936926220794 | 6.40265188186281 |
| chr10 | 5456987 | 5457045 | + | 3.01188173062009 | 5.68497871032191 | 23.8645642073598 | 10.6710864697714 |
| chr10 | 5457059 | 5457096 | + | 9.03564519186028 | 6.82197445238629 | 56.0817258872955 | 112.046407932599 |
| chr10 | 5457432 | 5457471 | + | 2.00792115374673 | 1.13699574206438 | 5.96614105183995 | 14.9395210576799 |
| chr10 | 5457495 | 5457523 | + | 2.00792115374673 | 3.41098722619315 | 26.2510206280958 | 32.0132594093141 |
| chr10 | 5457649 | 5457667 | + | 2.00792115374673 | 1.13699574206438 | 7.15936926220794 | 1.06710864697714 |
| chr10 | 5457833 | 5457855 | + | 4.01584230749346 | 1.13699574206438 | 17.8984231555198 | 9.60397782279422 |
| chr10 | 5457893 | 5457953 | + | 4.01584230749346 | 6.82197445238629 | 26.2510206280958 | 34.1474767032683 |
| chr10 | 5458033 | 5458099 | + | 8.03168461498692 | 3.41098722619315 | 27.4442488384638 | 53.3554323488568 |
| chr10 | 5458198 | 5458213 | + | 3.01188173062009 | 1.13699574206438 | 4.77291284147196 | 3.20132594093141 |
| chr10 | 5458281 | 5458334 | + | 36.1425807674411 | 83.0006891706999 | 158.699351978943 | 225.159924512176 |
| chr10 | 5458360 | 5458387 | + | 2.00792115374673 | 6.82197445238629 | 23.8645642073598 | 29.8790421153598 |
| chr10 | 5763671 | 5763699 | + | 3.01188173062009 | 5.68497871032191 | 2.38645642073598 | 1.06710864697714 |
| chr10 | 5765342 | 5765428 | - | 27.1069355755808 | 32.9728765198671 | 29.8307052591997 | 58.6909755837425 |
| chr10 | 5765469 | 5765495 | - | 3.01188173062009 | 3.41098722619315 | 2.38645642073598 | 3.20132594093141 |
| chr10 | 5765519 | 5765535 | - | 4.01584230749346 | 6.82197445238629 | 5.96614105183995 | 10.6710864697714 |
| chr10 | 5765566 | 5765601 | - | 5.01980288436682 | 3.41098722619315 | 3.57968463110397 | 7.46976052883995 |
| chr10 | 5765703 | 5766185 | - | 194.768351913433 | 210.344212281911 | 819.747780522809 | 924.116088282199 |
| chr10 | 5878155 | 5878174 | - | 1.00396057687336 | 2.27399148412876 | 2.38645642073598 | 2.13421729395427 |
| chr10 | 5937455 | 5937509 | + | 8.03168461498692 | 13.6439489047726 | 8.35259747257593 | 24.5434988804741 |
| chr10 | 6115554 | 6115600 | + | 1.00396057687336 | 1.13699574206438 | 1.19322821036799 | 6.40265188186281 |
| chr10 | 6115628 | 6115695 | + | 9.03564519186028 | 27.2878978095452 | 23.8645642073598 | 30.9461507623369 |
| chr10 | 6234149 | 6234166 | + | 1.00396057687336 | 3.41098722619315 | 3.57968463110397 | 5.33554323488568 |
| chr10 | 6234367 | 6234400 | + | 2.00792115374673 | 1.13699574206438 | 2.38645642073598 | 2.13421729395427 |
| chr10 | 6234478 | 6234486 | + | 1.00396057687336 | 1.13699574206438 | 1.19322821036799 | 3.20132594093141 |
| chr10 | 6235026 | 6235032 | + | 2.00792115374673 | 1.13699574206438 | 4.77291284147196 | 1.06710864697714 |
| chr10 | 6235343 | 6235403 | + | 6.02376346124019 | 1.13699574206438 | 1.19322821036799 | 3.20132594093141 |
| chr10 | 6427385 | 6427387 | - | 1.00396057687336 | 2.27399148412876 | 13.1255103140479 | 8.53686917581709 |
| chr10 | 6580375 | 6580577 | + | 50.1980288436682 | 45.4798296825753 | 239.838870283966 | 379.89067832386 |
| chr10 | 6584261 | 6584270 | + | 1.00396057687336 | 2.27399148412876 | 2.38645642073598 | 19.2079556455884 |
| chr10 | 7158833 | 7158844 | - | 7.02772403811355 | 10.2329616785794 | 3.57968463110397 | 2.13421729395427 |
| chr10 | 7284277 | 7284285 | - | 3.01188173062009 | 4.54798296825753 | 5.96614105183995 | 3.20132594093141 |
| chr10 | 7806984 | 7807011 | + | 21.0831721143407 | 20.4659233571589 | 52.5020412561915 | 77.8989312293309 |
| chr10 | 7807656 | 7807682 | + | 1311.17251339661 | 1300.72312892165 | 2720.56031963902 | 1735.11865998482 |
| chr10 | 7807705 | 7807770 | + | 30.1188173062009 | 47.753821166704 | 63.2410951495034 | 78.966039876308 |
| chr10 | 8014987 | 8014990 | + | 6.02376346124019 | 7.95897019445067 | 7.15936926220794 | 8.53686917581709 |
| chr10 | 8015020 | 8015044 | + | 9.03564519186028 | 9.09596593651506 | 7.15936926220794 | 7.46976052883995 |
| chr10 | 8015104 | 8015120 | + | 2.00792115374673 | 4.54798296825753 | 2.38645642073598 | 1.06710864697714 |
| chr10 | 8015212 | 8015234 | + | 6.02376346124019 | 15.9179403889013 | 4.77291284147196 | 9.60397782279422 |
| chr10 | 8015264 | 8015265 | + | 1.00396057687336 | 1.13699574206438 | 4.77291284147196 | 1.06710864697714 |
| chr10 | 8015296 | 8015318 | + | 4.01584230749346 | 2.27399148412876 | 4.77291284147196 | 2.13421729395427 |
| chr10 | 8015345 | 8015359 | + | 1.00396057687336 | 6.82197445238629 | 11.9322821036799 | 3.20132594093141 |
| chr10 | 11920213 | 11920251 | - | 9.03564519186028 | 6.82197445238629 | 9.54582568294392 | 10.6710864697714 |
| chr10 | 11920316 | 11920323 | - | 3.01188173062009 | 11.3699574206438 | 7.15936926220794 | 5.33554323488568 |
| chr10 | 11920386 | 11920401 | - | 7.02772403811355 | 3.41098722619315 | 4.77291284147196 | 11.7381951167485 |
| chr10 | 11920452 | 11920468 | - | 5.01980288436682 | 9.09596593651506 | 14.3187385244159 | 4.26843458790854 |
| chr10 | 11920683 | 11920691 | - | 3.01188173062009 | 2.27399148412876 | 4.77291284147196 | 1.06710864697714 |
| chr10 | 12165454 | 12165459 | - | 20.0792115374673 | 23.876910583352 | 8.35259747257593 | 5.33554323488568 |
| chr10 | 12165709 | 12165714 | - | 1.00396057687336 | 2.27399148412876 | 1.19322821036799 | 3.20132594093141 |
| chr10 | 12167602 | 12167627 | - | 4.01584230749346 | 4.54798296825753 | 5.96614105183995 | 2.13421729395427 |
| chr10 | 12167647 | 12167670 | - | 9.03564519186028 | 17.0549361309657 | 10.7390538933119 | 12.8053037637256 |
| chr10 | 12170716 | 12170718 | - | 3.01188173062009 | 3.41098722619315 | 2.38645642073598 | 1.06710864697714 |
| chr10 | 12170761 | 12170788 | - | 12.0475269224804 | 7.95897019445067 | 11.9322821036799 | 16.006629704657 |
| chr10 | 12250364 | 12250388 | + | 12.0475269224804 | 7.95897019445067 | 2.38645642073598 | 8.53686917581709 |
| chr10 | 12250425 | 12250440 | + | 9.03564519186028 | 5.68497871032191 | 8.35259747257593 | 21.3421729395427 |
| chr10 | 12250482 | 12250512 | + | 32.1267384599477 | 53.438799877026 | 68.0140079909754 | 68.2949534065367 |
| chr10 | 12250533 | 12250569 | + | 44.174265382428 | 57.9867828452835 | 147.960298085631 | 122.717494402371 |
| chr10 | 13211038 | 13211054 | + | 13.0514874993537 | 23.876910583352 | 10.7390538933119 | 23.476390233497 |
| chr10 | 13317453 | 13317472 | - | 1.00396057687336 | 1.13699574206438 | 1.19322821036799 | 1.06710864697714 |
| chr10 | 13317595 | 13317644 | - | 12.0475269224804 | 5.68497871032191 | 1.19322821036799 | 5.33554323488568 |
| chr10 | 13630610 | 13630616 | + | 7.02772403811355 | 5.68497871032191 | 5.96614105183995 | 2.13421729395427 |
| chr10 | 14843610 | 14843652 | + | 1.00396057687336 | 5.68497871032191 | 14.3187385244159 | 4.26843458790854 |
| chr10 | 14844243 | 14844245 | + | 1.00396057687336 | 9.09596593651506 | 2.38645642073598 | 1.06710864697714 |
| chr10 | 14844340 | 14844351 | + | 2.00792115374673 | 1.13699574206438 | 7.15936926220794 | 3.20132594093141 |
| chr10 | 14844388 | 14844439 | + | 7.02772403811355 | 4.54798296825753 | 16.7051949451519 | 10.6710864697714 |
| chr10 | 14871669 | 14871672 | + | 11.043566345607 | 15.9179403889013 | 15.5119667347839 | 10.6710864697714 |
| chr10 | 14898822 | 14898829 | - | 10.0396057687336 | 11.3699574206438 | 16.7051949451519 | 25.6106075274513 |
| chr10 | 14902527 | 14902537 | + | 6.02376346124019 | 5.68497871032191 | 21.4781077866238 | 11.7381951167485 |
| chr10 | 14902581 | 14902605 | + | 5.01980288436682 | 4.54798296825753 | 5.96614105183995 | 4.26843458790854 |
| chr10 | 14902660 | 14902710 | + | 88.3485307648561 | 110.288586980245 | 66.8207797806074 | 102.442430109805 |
| chr10 | 14902730 | 14902773 | + | 12.0475269224804 | 11.3699574206438 | 32.2171616799357 | 43.7514545260626 |
| chr10 | 14903296 | 14903297 | + | 2.00792115374673 | 1.13699574206438 | 1.19322821036799 | 1.06710864697714 |
| chr10 | 15104203 | 15104221 | + | 1.00396057687336 | 1.13699574206438 | 3.57968463110397 | 1.06710864697714 |
| chr10 | 15778197 | 15778207 | - | 1.00396057687336 | 3.41098722619315 | 7.15936926220794 | 3.20132594093141 |
| chr10 | 15778404 | 15778406 | - | 2.00792115374673 | 1.13699574206438 | 1.19322821036799 | 4.26843458790854 |
| chr10 | 16511390 | 16511392 | + | 5.01980288436682 | 5.68497871032191 | 2.38645642073598 | 4.26843458790854 |
| chr10 | 16511434 | 16511446 | + | 11.043566345607 | 9.09596593651506 | 3.57968463110397 | 2.13421729395427 |
| chr10 | 16511533 | 16511556 | + | 24.0950538449607 | 23.876910583352 | 50.1155848354556 | 36.2816939972226 |
| chr10 | 16511722 | 16511728 | + | 3.01188173062009 | 6.82197445238629 | 2.38645642073598 | 6.40265188186281 |
| chr10 | 16512165 | 16512198 | + | 11.043566345607 | 6.82197445238629 | 9.54582568294392 | 17.0737383516342 |
| chr10 | 16593123 | 16593125 | - | 1.00396057687336 | 4.54798296825753 | 2.38645642073598 | 4.26843458790854 |
| chr10 | 16593295 | 16593304 | - | 8.03168461498692 | 1.13699574206438 | 1.19322821036799 | 1.06710864697714 |
| chr10 | 17148255 | 17148257 | - | 1.00396057687336 | 2.27399148412876 | 2.38645642073598 | 2.13421729395427 |
| chr10 | 17237286 | 17237307 | + | 6.02376346124019 | 7.95897019445067 | 8.35259747257593 | 22.4092815865198 |
| chr10 | 17237335 | 17237344 | + | 3.01188173062009 | 4.54798296825753 | 2.38645642073598 | 2.13421729395427 |
| chr10 | 17237373 | 17237442 | + | 28.1108961524542 | 37.5208594881246 | 42.9562155732476 | 55.4896496428111 |
| chr10 | 17237472 | 17237519 | + | 27.1069355755808 | 29.5618892936739 | 22.6713359969918 | 17.0737383516342 |
| chr10 | 17237544 | 17237570 | + | 55.217831728035 | 57.9867828452835 | 65.6275515702394 | 16.006629704657 |
| chr10 | 18545757 | 18545770 | - | 4.01584230749346 | 9.09596593651506 | 13.1255103140479 | 30.9461507623369 |
| chr10 | 18675166 | 18675259 | + | 2.00792115374673 | 12.5069531627082 | 9.54582568294392 | 13.8724124107028 |
| chr10 | 18675325 | 18675367 | + | 7.02772403811355 | 9.09596593651506 | 11.9322821036799 | 21.3421729395427 |
| chr10 | 18676046 | 18676061 | + | 1.00396057687336 | 1.13699574206438 | 7.15936926220794 | 1.06710864697714 |
| chr10 | 18676097 | 18676232 | + | 8.03168461498692 | 15.9179403889013 | 54.8884976769275 | 13.8724124107028 |
| chr10 | 18676991 | 18677050 | + | 1.00396057687336 | 1.13699574206438 | 2.38645642073598 | 2.13421729395427 |
| chr10 | 18677440 | 18677515 | + | 2.00792115374673 | 6.82197445238629 | 3.57968463110397 | 1.06710864697714 |
| chr10 | 18677860 | 18677897 | + | 3.01188173062009 | 3.41098722619315 | 2.38645642073598 | 11.7381951167485 |
| chr10 | 18677952 | 18678025 | + | 1.00396057687336 | 3.41098722619315 | 5.96614105183995 | 6.40265188186281 |
| chr10 | 18678063 | 18678107 | + | 2.00792115374673 | 2.27399148412876 | 2.38645642073598 | 5.33554323488568 |
| chr10 | 18680681 | 18680715 | + | 1.00396057687336 | 1.13699574206438 | 1.19322821036799 | 3.20132594093141 |
| chr10 | 18681082 | 18681146 | + | 2.00792115374673 | 5.68497871032191 | 1.19322821036799 | 1.06710864697714 |
| chr10 | 18681329 | 18681346 | + | 3.01188173062009 | 4.54798296825753 | 8.35259747257593 | 14.9395210576799 |
| chr10 | 18681393 | 18681431 | + | 1.00396057687336 | 2.27399148412876 | 2.38645642073598 | 5.33554323488568 |
| chr10 | 18681501 | 18681579 | + | 2.00792115374673 | 4.54798296825753 | 3.57968463110397 | 3.20132594093141 |
| chr10 | 20288247 | 20288248 | + | 3.01188173062009 | 2.27399148412876 | 7.15936926220794 | 2.13421729395427 |
| chr10 | 21514886 | 21514899 | - | 8.03168461498692 | 11.3699574206438 | 7.15936926220794 | 6.40265188186281 |
| chr10 | 21556773 | 21556824 | + | 5.01980288436682 | 2.27399148412876 | 1.19322821036799 | 4.26843458790854 |
| chr10 | 21556932 | 21556936 | + | 1.00396057687336 | 2.27399148412876 | 1.19322821036799 | 6.40265188186281 |
| chr10 | 21742118 | 21742135 | + | 1.00396057687336 | 1.13699574206438 | 2.38645642073598 | 1.06710864697714 |
| chr10 | 21742547 | 21742580 | + | 3.01188173062009 | 2.27399148412876 | 9.54582568294392 | 6.40265188186281 |
| chr10 | 21756579 | 21756608 | - | 5.01980288436682 | 12.5069531627082 | 15.5119667347839 | 13.8724124107028 |
| chr10 | 21756674 | 21756679 | - | 8.03168461498692 | 21.6029190992233 | 17.8984231555198 | 18.1408469986113 |
| chr10 | 22320003 | 22320005 | + | 4.01584230749346 | 1.13699574206438 | 2.38645642073598 | 3.20132594093141 |
| chr10 | 22320076 | 22320079 | + | 1.00396057687336 | 4.54798296825753 | 10.7390538933119 | 5.33554323488568 |
| chr10 | 22320163 | 22320178 | + | 12.0475269224804 | 20.4659233571589 | 28.6374770488317 | 52.2883237018796 |
| chr10 | 22320256 | 22320290 | + | 7.02772403811355 | 14.780944646837 | 25.0577924177278 | 34.1474767032683 |
| chr10 | 22330394 | 22330415 | + | 1.00396057687336 | 4.54798296825753 | 4.77291284147196 | 2.13421729395427 |
| chr10 | 22330519 | 22330550 | + | 1.00396057687336 | 2.27399148412876 | 14.3187385244159 | 6.40265188186281 |
| chr10 | 22536946 | 22536964 | - | 4.01584230749346 | 4.54798296825753 | 26.2510206280958 | 20.2750642925656 |
| chr10 | 23120875 | 23120879 | + | 4.01584230749346 | 1.13699574206438 | 1.19322821036799 | 1.06710864697714 |
| chr10 | 24586017 | 24586030 | - | 4.01584230749346 | 6.82197445238629 | 11.9322821036799 | 19.2079556455884 |
| chr10 | 24586070 | 24586086 | - | 7.02772403811355 | 15.9179403889013 | 26.2510206280958 | 23.476390233497 |
| chr10 | 24908456 | 24908458 | - | 2.00792115374673 | 6.82197445238629 | 5.96614105183995 | 7.46976052883995 |
| chr10 | 26746600 | 26746603 | + | 1.00396057687336 | 3.41098722619315 | 5.96614105183995 | 7.46976052883995 |
| chr10 | 26746972 | 26746976 | - | 1.00396057687336 | 3.41098722619315 | 1.19322821036799 | 1.06710864697714 |
| chr10 | 26747290 | 26747300 | - | 6.02376346124019 | 7.95897019445067 | 9.54582568294392 | 14.9395210576799 |
| chr10 | 26748286 | 26748297 | - | 1.00396057687336 | 6.82197445238629 | 3.57968463110397 | 8.53686917581709 |
| chr10 | 27110488 | 27110491 | - | 9.03564519186028 | 20.4659233571589 | 10.7390538933119 | 10.6710864697714 |
| chr10 | 27110641 | 27110664 | - | 2.00792115374673 | 1.13699574206438 | 2.38645642073598 | 4.26843458790854 |
| chr10 | 27111862 | 27111877 | - | 4.01584230749346 | 2.27399148412876 | 3.57968463110397 | 12.8053037637256 |
| chr10 | 27145496 | 27145498 | - | 1.00396057687336 | 3.41098722619315 | 4.77291284147196 | 1.06710864697714 |
| chr10 | 27145571 | 27145577 | - | 3.01188173062009 | 1.13699574206438 | 4.77291284147196 | 5.33554323488568 |
| chr10 | 27186622 | 27186625 | + | 13.0514874993537 | 25.0139063254164 | 10.7390538933119 | 13.8724124107028 |
| chr10 | 27538456 | 27538473 | + | 1.00396057687336 | 3.41098722619315 | 3.57968463110397 | 3.20132594093141 |
| chr10 | 27539323 | 27539330 | + | 1.00396057687336 | 2.27399148412876 | 1.19322821036799 | 1.06710864697714 |
| chr10 | 28590784 | 28590833 | + | 35.1386201905678 | 31.8358807778027 | 35.7968463110397 | 11.7381951167485 |
| chr10 | 28595880 | 28595894 | + | 5.01980288436682 | 5.68497871032191 | 1.19322821036799 | 1.06710864697714 |
| chr10 | 28595969 | 28595987 | + | 3.01188173062009 | 5.68497871032191 | 4.77291284147196 | 5.33554323488568 |
| chr10 | 28596019 | 28596036 | + | 3.01188173062009 | 3.41098722619315 | 2.38645642073598 | 4.26843458790854 |
| chr10 | 28608223 | 28608269 | + | 4.01584230749346 | 9.09596593651506 | 14.3187385244159 | 7.46976052883995 |
| chr10 | 28608288 | 28608333 | + | 5.01980288436682 | 9.09596593651506 | 14.3187385244159 | 8.53686917581709 |
| chr10 | 28608340 | 28608349 | + | 1.00396057687336 | 4.54798296825753 | 5.96614105183995 | 4.26843458790854 |
| chr10 | 28616178 | 28616181 | + | 2.00792115374673 | 1.13699574206438 | 2.38645642073598 | 2.13421729395427 |
| chr10 | 28617654 | 28617702 | + | 14.0554480762271 | 21.6029190992233 | 17.8984231555198 | 19.2079556455884 |
| chr10 | 28617733 | 28617785 | + | 8.03168461498692 | 21.6029190992233 | 13.1255103140479 | 27.7448248214055 |
| chr10 | 28619516 | 28619549 | + | 3.01188173062009 | 4.54798296825753 | 9.54582568294392 | 2.13421729395427 |
| chr10 | 28619562 | 28619613 | + | 18.0712903837206 | 37.5208594881246 | 19.0916513658878 | 44.8185631730397 |
| chr10 | 28619632 | 28619645 | + | 8.03168461498692 | 2.27399148412876 | 14.3187385244159 | 9.60397782279422 |
| chr10 | 28619702 | 28619775 | + | 181.716864414079 | 203.522237829524 | 250.577924177278 | 331.870789209889 |
| chr10 | 28619810 | 28619875 | + | 9.03564519186028 | 11.3699574206438 | 23.8645642073598 | 20.2750642925656 |
| chr10 | 28619899 | 28619922 | + | 21.0831721143407 | 19.3289276150945 | 81.1395183050233 | 48.0198891139711 |
| chr10 | 28619978 | 28619988 | + | 37.1465413443145 | 65.9457530397342 | 34.6036181006717 | 17.0737383516342 |
| chr10 | 28620034 | 28620085 | + | 26.1029749987075 | 55.7127913611547 | 27.4442488384638 | 30.9461507623369 |
| chr10 | 28620125 | 28620197 | + | 27.1069355755808 | 50.0278126508328 | 45.3426719939836 | 35.2145853502455 |
| chr10 | 28620246 | 28620265 | + | 5.01980288436682 | 20.4659233571589 | 19.0916513658878 | 3.20132594093141 |
| chr10 | 28620290 | 28620340 | + | 15.0594086531005 | 21.6029190992233 | 5.96614105183995 | 14.9395210576799 |
| chr10 | 28620396 | 28620513 | + | 326.287187483843 | 472.990228698783 | 298.307052591997 | 530.352997547636 |
| chr10 | 28620564 | 28620614 | + | 134.530717301031 | 204.659233571589 | 85.9124311464952 | 115.247733873531 |
| chr10 | 28620636 | 28620700 | + | 34.1346596136944 | 88.6856678810218 | 83.5259747257593 | 80.0331485232852 |
| chr10 | 28620728 | 28620828 | + | 79.3128855729958 | 97.7816338175368 | 120.516049247167 | 55.4896496428111 |
| chr10 | 28620853 | 28620881 | + | 10.0396057687336 | 9.09596593651506 | 11.9322821036799 | 7.46976052883995 |
| chr10 | 28620906 | 28620935 | + | 4.01584230749346 | 3.41098722619315 | 1.19322821036799 | 1.06710864697714 |
| chr10 | 28682332 | 28682433 | + | 15.0594086531005 | 4.54798296825753 | 20.2848795762558 | 32.0132594093141 |
| chr10 | 28682645 | 28682673 | + | 3.01188173062009 | 5.68497871032191 | 14.3187385244159 | 20.2750642925656 |
| chr10 | 29409532 | 29409627 | + | 107.42378172545 | 115.973565690567 | 84.7192029361272 | 92.8384522870108 |
| chr10 | 30312984 | 30312991 | - | 4.01584230749346 | 5.68497871032191 | 7.15936926220794 | 1.06710864697714 |
| chr10 | 30313238 | 30313242 | - | 3.01188173062009 | 3.41098722619315 | 9.54582568294392 | 7.46976052883995 |
| chr10 | 32009030 | 32009062 | - | 4.01584230749346 | 2.27399148412876 | 2.38645642073598 | 4.26843458790854 |
| chr10 | 32009091 | 32009124 | - | 5.01980288436682 | 3.41098722619315 | 8.35259747257593 | 2.13421729395427 |
| chr10 | 32009151 | 32009233 | - | 6.02376346124019 | 6.82197445238629 | 7.15936926220794 | 8.53686917581709 |
| chr10 | 32009260 | 32009322 | - | 2.00792115374673 | 15.9179403889013 | 7.15936926220794 | 3.20132594093141 |
| chr10 | 32009420 | 32009634 | - | 45.1782259593014 | 50.0278126508328 | 16.7051949451519 | 20.2750642925656 |
| chr10 | 32009673 | 32009711 | - | 15.0594086531005 | 13.6439489047726 | 9.54582568294392 | 14.9395210576799 |
| chr10 | 32009737 | 32009762 | - | 6.02376346124019 | 5.68497871032191 | 4.77291284147196 | 13.8724124107028 |
| chr10 | 32009827 | 32009922 | - | 6.02376346124019 | 4.54798296825753 | 3.57968463110397 | 7.46976052883995 |
| chr10 | 32009982 | 32010074 | - | 16.0633692299738 | 5.68497871032191 | 15.5119667347839 | 5.33554323488568 |
| chr10 | 32010109 | 32010172 | - | 17.0673298068472 | 10.2329616785794 | 20.2848795762558 | 16.006629704657 |
| chr10 | 32010200 | 32010214 | - | 32.1267384599477 | 26.1509020674808 | 13.1255103140479 | 39.483019938154 |
| chr10 | 32010515 | 32010535 | - | 1.00396057687336 | 1.13699574206438 | 3.57968463110397 | 2.13421729395427 |
| chr10 | 32010552 | 32010639 | - | 6.02376346124019 | 13.6439489047726 | 5.96614105183995 | 3.20132594093141 |
| chr10 | 32010660 | 32010694 | - | 7.02772403811355 | 11.3699574206438 | 4.77291284147196 | 5.33554323488568 |
| chr10 | 32010766 | 32010798 | - | 20.0792115374673 | 18.1919318730301 | 8.35259747257593 | 8.53686917581709 |
| chr10 | 32010824 | 32010885 | - | 15.0594086531005 | 7.95897019445067 | 8.35259747257593 | 10.6710864697714 |
| chr10 | 32010959 | 32010979 | - | 11.043566345607 | 9.09596593651506 | 3.57968463110397 | 5.33554323488568 |
| chr10 | 32011060 | 32011117 | - | 1.00396057687336 | 10.2329616785794 | 3.57968463110397 | 5.33554323488568 |
| chr10 | 32011131 | 32011161 | - | 5.01980288436682 | 3.41098722619315 | 7.15936926220794 | 5.33554323488568 |
| chr10 | 32011177 | 32011334 | - | 30.1188173062009 | 23.876910583352 | 26.2510206280958 | 34.1474767032683 |
| chr10 | 32011380 | 32011394 | - | 3.01188173062009 | 2.27399148412876 | 1.19322821036799 | 2.13421729395427 |
| chr10 | 32011437 | 32011443 | - | 14.0554480762271 | 3.41098722619315 | 9.54582568294392 | 1.06710864697714 |
| chr10 | 32011476 | 32011496 | - | 10.0396057687336 | 10.2329616785794 | 17.8984231555198 | 11.7381951167485 |
| chr10 | 32015508 | 32015555 | - | 8.03168461498692 | 9.09596593651506 | 22.6713359969918 | 13.8724124107028 |
| chr10 | 32268701 | 32268759 | - | 125.495072109171 | 256.96103770655 | 298.307052591997 | 243.300771510787 |
| chr10 | 32900864 | 32900884 | - | 11.043566345607 | 10.2329616785794 | 5.96614105183995 | 5.33554323488568 |
| chr10 | 32901446 | 32901497 | - | 3.01188173062009 | 5.68497871032191 | 5.96614105183995 | 2.13421729395427 |
| chr10 | 35009923 | 35009979 | - | 26.1029749987075 | 36.3838637460602 | 54.8884976769275 | 16.006629704657 |
| chr10 | 35010011 | 35010090 | - | 234.926774988367 | 202.38524208746 | 422.402786470268 | 319.065485446164 |
| chr10 | 35010116 | 35010162 | - | 14.0554480762271 | 10.2329616785794 | 22.6713359969918 | 25.6106075274513 |
| chr10 | 35010286 | 35010292 | - | 2.00792115374673 | 1.13699574206438 | 3.57968463110397 | 5.33554323488568 |
| chr10 | 35211709 | 35211729 | + | 12.0475269224804 | 15.9179403889013 | 2.38645642073598 | 3.20132594093141 |
| chr10 | 35211754 | 35211756 | + | 2.00792115374673 | 9.09596593651506 | 1.19322821036799 | 3.20132594093141 |
| chr10 | 35211833 | 35211869 | + | 13.0514874993537 | 36.3838637460602 | 4.77291284147196 | 3.20132594093141 |
| chr10 | 35212373 | 35212396 | + | 6.02376346124019 | 10.2329616785794 | 2.38645642073598 | 1.06710864697714 |
| chr10 | 35569185 | 35569189 | + | 4.01584230749346 | 1.13699574206438 | 3.57968463110397 | 4.26843458790854 |
| chr10 | 35569210 | 35569246 | + | 19.0752509605939 | 17.0549361309657 | 68.0140079909754 | 30.9461507623369 |
| chr10 | 35569381 | 35569462 | + | 3.01188173062009 | 7.95897019445067 | 21.4781077866238 | 23.476390233497 |
| chr10 | 35569498 | 35569510 | + | 2.00792115374673 | 1.13699574206438 | 3.57968463110397 | 16.006629704657 |
| chr10 | 35569531 | 35569570 | + | 8.03168461498692 | 6.82197445238629 | 21.4781077866238 | 23.476390233497 |
| chr10 | 35569699 | 35569724 | + | 2.00792115374673 | 1.13699574206438 | 3.57968463110397 | 3.20132594093141 |
| chr10 | 35569783 | 35569785 | + | 2.00792115374673 | 1.13699574206438 | 4.77291284147196 | 4.26843458790854 |
| chr10 | 35570003 | 35570037 | + | 4.01584230749346 | 1.13699574206438 | 10.7390538933119 | 13.8724124107028 |
| chr10 | 35570054 | 35570106 | + | 4.01584230749346 | 3.41098722619315 | 11.9322821036799 | 16.006629704657 |
| chr10 | 35570243 | 35570301 | + | 15.0594086531005 | 10.2329616785794 | 51.3088130458235 | 35.2145853502455 |
| chr10 | 35570519 | 35570572 | + | 4.01584230749346 | 1.13699574206438 | 11.9322821036799 | 10.6710864697714 |
| chr10 | 35570870 | 35570891 | + | 3.01188173062009 | 3.41098722619315 | 15.5119667347839 | 22.4092815865198 |
| chr10 | 35570933 | 35570984 | + | 18.0712903837206 | 14.780944646837 | 45.3426719939836 | 42.6843458790854 |
| chr10 | 35571013 | 35571057 | + | 3.01188173062009 | 3.41098722619315 | 20.2848795762558 | 4.26843458790854 |
| chr10 | 35571088 | 35571119 | + | 40.1584230749346 | 40.9318467143178 | 141.994157033791 | 90.7042349930565 |
| chr10 | 35571151 | 35571159 | + | 5.01980288436682 | 6.82197445238629 | 25.0577924177278 | 10.6710864697714 |
| chr10 | 35571190 | 35571225 | + | 5.01980288436682 | 7.95897019445067 | 5.96614105183995 | 10.6710864697714 |
| chr10 | 37829161 | 37829208 | - | 20.0792115374673 | 19.3289276150945 | 35.7968463110397 | 29.8790421153598 |
| chr10 | 37829249 | 37829264 | - | 3.01188173062009 | 6.82197445238629 | 7.15936926220794 | 6.40265188186281 |
| chr10 | 37829460 | 37829466 | - | 1.00396057687336 | 3.41098722619315 | 8.35259747257593 | 4.26843458790854 |
| chr10 | 38057897 | 38057900 | + | 4.01584230749346 | 4.54798296825753 | 1.19322821036799 | 1.06710864697714 |
| chr10 | 38057932 | 38057938 | + | 4.01584230749346 | 3.41098722619315 | 3.57968463110397 | 2.13421729395427 |
| chr10 | 38058925 | 38058974 | + | 9.03564519186028 | 5.68497871032191 | 4.77291284147196 | 5.33554323488568 |
| chr10 | 38059197 | 38059218 | + | 6.02376346124019 | 9.09596593651506 | 4.77291284147196 | 4.26843458790854 |
| chr10 | 38059303 | 38059310 | + | 2.00792115374673 | 1.13699574206438 | 11.9322821036799 | 1.06710864697714 |
| chr10 | 38059342 | 38059359 | + | 2.00792115374673 | 1.13699574206438 | 4.77291284147196 | 2.13421729395427 |
| chr10 | 38059946 | 38059964 | + | 4.01584230749346 | 10.2329616785794 | 5.96614105183995 | 6.40265188186281 |
| chr10 | 38120718 | 38120724 | + | 17.0673298068472 | 17.0549361309657 | 9.54582568294392 | 18.1408469986113 |
| chr10 | 38120831 | 38120838 | + | 2.00792115374673 | 1.13699574206438 | 1.19322821036799 | 5.33554323488568 |
| chr10 | 38121450 | 38121475 | + | 4.01584230749346 | 3.41098722619315 | 5.96614105183995 | 5.33554323488568 |
| chr10 | 38121543 | 38121571 | + | 4.01584230749346 | 5.68497871032191 | 4.77291284147196 | 4.26843458790854 |
| chr10 | 38123137 | 38123145 | + | 1.00396057687336 | 1.13699574206438 | 1.19322821036799 | 1.06710864697714 |
| chr10 | 42589215 | 42589244 | - | 3.01188173062009 | 6.82197445238629 | 2.38645642073598 | 1.06710864697714 |
| chr10 | 42589763 | 42589770 | - | 2.00792115374673 | 3.41098722619315 | 7.15936926220794 | 5.33554323488568 |
| chr10 | 42589964 | 42589975 | - | 1.00396057687336 | 4.54798296825753 | 3.57968463110397 | 4.26843458790854 |
| chr10 | 42591210 | 42591212 | - | 2.00792115374673 | 1.13699574206438 | 1.19322821036799 | 1.06710864697714 |
| chr10 | 42831106 | 42831137 | + | 83.3287278804893 | 69.3567402659273 | 124.095733878271 | 257.17318392149 |
| chr10 | 42831164 | 42831205 | + | 71.2812009580089 | 37.5208594881246 | 96.6514850398071 | 177.140035398205 |
| chr10 | 42831229 | 42831236 | + | 16.0633692299738 | 6.82197445238629 | 7.15936926220794 | 9.60397782279422 |
| chr10 | 42831258 | 42831266 | + | 5.01980288436682 | 2.27399148412876 | 2.38645642073598 | 2.13421729395427 |
| chr10 | 42831310 | 42831316 | + | 2.00792115374673 | 1.13699574206438 | 2.38645642073598 | 2.13421729395427 |
| chr10 | 42831369 | 42831380 | + | 4.01584230749346 | 2.27399148412876 | 2.38645642073598 | 5.33554323488568 |
| chr10 | 43385872 | 43385890 | - | 20.0792115374673 | 23.876910583352 | 9.54582568294392 | 12.8053037637256 |
| chr10 | 43385961 | 43386002 | - | 10.0396057687336 | 30.6988850357383 | 15.5119667347839 | 12.8053037637256 |
| chr10 | 43386088 | 43386218 | - | 66.2613980736421 | 77.315710460378 | 114.549908195327 | 147.260993282845 |
| chr10 | 43386235 | 43386329 | - | 101.40001826421 | 109.151591238181 | 116.936364616063 | 187.811121867976 |
| chr10 | 43386350 | 43386355 | - | 2.00792115374673 | 4.54798296825753 | 2.38645642073598 | 4.26843458790854 |
| chr10 | 43386376 | 43386415 | - | 9.03564519186028 | 14.780944646837 | 11.9322821036799 | 8.53686917581709 |
| chr10 | 43386515 | 43386600 | - | 16.0633692299738 | 13.6439489047726 | 17.8984231555198 | 32.0132594093141 |
| chr10 | 43386625 | 43386718 | - | 9.03564519186028 | 4.54798296825753 | 21.4781077866238 | 32.0132594093141 |
| chr10 | 43643927 | 43643930 | - | 1.00396057687336 | 1.13699574206438 | 2.38645642073598 | 5.33554323488568 |
| chr10 | 46003325 | 46003363 | + | 4.01584230749346 | 2.27399148412876 | 3.57968463110397 | 5.33554323488568 |
| chr10 | 46003621 | 46003630 | + | 4.01584230749346 | 6.82197445238629 | 7.15936926220794 | 9.60397782279422 |
| chr10 | 46003675 | 46003719 | + | 26.1029749987075 | 27.2878978095452 | 21.4781077866238 | 22.4092815865198 |
| chr10 | 46005106 | 46005320 | - | 154.609928838498 | 155.76841666282 | 91.8785721983352 | 133.388580872142 |
| chr10 | 46005397 | 46005449 | - | 78.3089249961224 | 70.4937360079917 | 22.6713359969918 | 56.5567582897882 |
| chr10 | 46005479 | 46005607 | - | 72.2851615348822 | 68.2197445238629 | 22.6713359969918 | 66.1607361125824 |
| chr10 | 46005638 | 46005729 | - | 4.01584230749346 | 2.27399148412876 | 2.38645642073598 | 5.33554323488568 |
| chr10 | 46005753 | 46005794 | - | 1.00396057687336 | 6.82197445238629 | 2.38645642073598 | 7.46976052883995 |
| chr10 | 46006126 | 46006205 | - | 2.00792115374673 | 2.27399148412876 | 7.15936926220794 | 1.06710864697714 |
| chr10 | 46006235 | 46006358 | - | 2.00792115374673 | 6.82197445238629 | 1.19322821036799 | 5.33554323488568 |
| chr10 | 46006379 | 46006463 | - | 3.01188173062009 | 4.54798296825753 | 2.38645642073598 | 3.20132594093141 |
| chr10 | 46006588 | 46006600 | - | 4.01584230749346 | 4.54798296825753 | 3.57968463110397 | 12.8053037637256 |
| chr10 | 48435049 | 48435120 | + | 290.144606716402 | 530.977011544066 | 1034.52885838905 | 681.88242541839 |
| chr10 | 48435180 | 48435205 | + | 1.00396057687336 | 2.27399148412876 | 8.35259747257593 | 10.6710864697714 |
| chr10 | 48435705 | 48435713 | + | 3.01188173062009 | 3.41098722619315 | 5.96614105183995 | 7.46976052883995 |
| chr10 | 48436224 | 48436244 | + | 6.02376346124019 | 3.41098722619315 | 14.3187385244159 | 4.26843458790854 |
| chr10 | 48439199 | 48439217 | + | 1.00396057687336 | 1.13699574206438 | 1.19322821036799 | 1.06710864697714 |
| chr10 | 49456816 | 49456827 | - | 7.02772403811355 | 9.09596593651506 | 3.57968463110397 | 2.13421729395427 |
| chr10 | 49456936 | 49456946 | - | 9.03564519186028 | 3.41098722619315 | 3.57968463110397 | 9.60397782279422 |
| chr10 | 49457302 | 49457329 | - | 4.01584230749346 | 4.54798296825753 | 4.77291284147196 | 3.20132594093141 |
| chr10 | 49457467 | 49457487 | - | 7.02772403811355 | 3.41098722619315 | 8.35259747257593 | 11.7381951167485 |
| chr10 | 50305747 | 50305751 | - | 6.02376346124019 | 11.3699574206438 | 11.9322821036799 | 14.9395210576799 |
| chr10 | 50305814 | 50305822 | - | 1.00396057687336 | 1.13699574206438 | 7.15936926220794 | 16.006629704657 |
| chr10 | 50305852 | 50305856 | - | 6.02376346124019 | 5.68497871032191 | 8.35259747257593 | 6.40265188186281 |
| chr10 | 50305878 | 50305885 | - | 3.01188173062009 | 7.95897019445067 | 16.7051949451519 | 4.26843458790854 |
| chr10 | 50305910 | 50305914 | - | 12.0475269224804 | 28.4248935516096 | 38.1833027317757 | 37.3488026441998 |
| chr10 | 50306603 | 50306630 | - | 8.03168461498692 | 1.13699574206438 | 3.57968463110397 | 6.40265188186281 |
| chr10 | 50306679 | 50306702 | - | 4.01584230749346 | 4.54798296825753 | 2.38645642073598 | 6.40265188186281 |
| chr10 | 50307124 | 50307128 | - | 1.00396057687336 | 3.41098722619315 | 4.77291284147196 | 4.26843458790854 |
| chr10 | 50307283 | 50307284 | - | 1.00396057687336 | 1.13699574206438 | 2.38645642073598 | 3.20132594093141 |
| chr10 | 51697502 | 51697521 | - | 49.1940682667949 | 50.0278126508328 | 23.8645642073598 | 50.1541064079254 |
| chr10 | 56357483 | 56357504 | - | 7.02772403811355 | 10.2329616785794 | 9.54582568294392 | 4.26843458790854 |
| chr10 | 56357534 | 56357549 | - | 33.130699036821 | 44.3428339405109 | 26.2510206280958 | 23.476390233497 |
| chr10 | 56357624 | 56357631 | - | 2.00792115374673 | 10.2329616785794 | 7.15936926220794 | 7.46976052883995 |
| chr10 | 56357711 | 56357750 | - | 11.043566345607 | 12.5069531627082 | 9.54582568294392 | 3.20132594093141 |
| chr10 | 56357784 | 56357817 | - | 16.0633692299738 | 9.09596593651506 | 5.96614105183995 | 12.8053037637256 |
| chr10 | 56357847 | 56357908 | - | 139.550520185398 | 173.96034853585 | 81.1395183050233 | 173.938709457273 |
| chr10 | 56357929 | 56357958 | - | 16.0633692299738 | 11.3699574206438 | 8.35259747257593 | 14.9395210576799 |
| chr10 | 58287657 | 58287686 | + | 3.01188173062009 | 1.13699574206438 | 5.96614105183995 | 4.26843458790854 |
| chr10 | 58287961 | 58287963 | + | 4.01584230749346 | 10.2329616785794 | 4.77291284147196 | 2.13421729395427 |
| chr10 | 58304813 | 58304814 | + | 10.0396057687336 | 9.09596593651506 | 15.5119667347839 | 3.20132594093141 |
| chr10 | 58369162 | 58369167 | + | 2.00792115374673 | 1.13699574206438 | 1.19322821036799 | 1.06710864697714 |
| chr10 | 58394994 | 58395136 | + | 23.0910932680874 | 46.6168254246397 | 66.8207797806074 | 62.959410171651 |
| chr10 | 58395152 | 58395186 | + | 10.0396057687336 | 15.9179403889013 | 10.7390538933119 | 10.6710864697714 |
| chr10 | 58395204 | 58395220 | + | 1.00396057687336 | 1.13699574206438 | 7.15936926220794 | 5.33554323488568 |
| chr10 | 58395254 | 58395263 | + | 3.01188173062009 | 3.41098722619315 | 3.57968463110397 | 4.26843458790854 |
| chr10 | 58395293 | 58395352 | + | 48.1901076899215 | 53.438799877026 | 54.8884976769275 | 43.7514545260626 |
| chr10 | 58395372 | 58395427 | + | 37.1465413443145 | 53.438799877026 | 54.8884976769275 | 92.8384522870108 |
| chr10 | 58395459 | 58395646 | + | 230.910932680874 | 339.96172687725 | 313.819019326781 | 534.621432135545 |
| chr10 | 58395677 | 58395716 | + | 9.03564519186028 | 27.2878978095452 | 14.3187385244159 | 17.0737383516342 |
| chr10 | 58395763 | 58395806 | + | 21.0831721143407 | 52.3018041349616 | 69.2072362013434 | 58.6909755837425 |
| chr10 | 58395837 | 58395910 | + | 41.1623836518079 | 90.9596593651506 | 108.583767143487 | 90.7042349930565 |
| chr10 | 58395939 | 58395967 | + | 4.01584230749346 | 11.3699574206438 | 10.7390538933119 | 10.6710864697714 |
| chr10 | 58395989 | 58396007 | + | 2.00792115374673 | 2.27399148412876 | 1.19322821036799 | 7.46976052883995 |
| chr10 | 58396033 | 58396042 | + | 3.01188173062009 | 2.27399148412876 | 4.77291284147196 | 1.06710864697714 |
| chr10 | 58396985 | 58397015 | + | 4.01584230749346 | 1.13699574206438 | 3.57968463110397 | 5.33554323488568 |
| chr10 | 58398185 | 58398208 | + | 2.00792115374673 | 13.6439489047726 | 4.77291284147196 | 7.46976052883995 |
| chr10 | 58398407 | 58398426 | + | 13.0514874993537 | 31.8358807778027 | 20.2848795762558 | 19.2079556455884 |
| chr10 | 58398512 | 58398518 | + | 3.01188173062009 | 4.54798296825753 | 4.77291284147196 | 1.06710864697714 |
| chr10 | 59792239 | 59792252 | - | 1.00396057687336 | 9.09596593651506 | 4.77291284147196 | 2.13421729395427 |
| chr10 | 59792273 | 59792301 | - | 2.00792115374673 | 2.27399148412876 | 8.35259747257593 | 3.20132594093141 |
| chr10 | 59792395 | 59792406 | - | 2.00792115374673 | 6.82197445238629 | 7.15936926220794 | 6.40265188186281 |
| chr10 | 59792512 | 59792521 | - | 3.01188173062009 | 7.95897019445067 | 7.15936926220794 | 1.06710864697714 |
| chr10 | 59792565 | 59792690 | - | 56.2217923049084 | 48.8908169087684 | 51.3088130458235 | 38.4159112911769 |
| chr10 | 59792706 | 59792759 | - | 15.0594086531005 | 10.2329616785794 | 13.1255103140479 | 9.60397782279422 |
| chr10 | 59792785 | 59792839 | - | 11.043566345607 | 4.54798296825753 | 7.15936926220794 | 6.40265188186281 |
| chr10 | 59792868 | 59792888 | - | 7.02772403811355 | 4.54798296825753 | 7.15936926220794 | 4.26843458790854 |
| chr10 | 60028743 | 60028749 | - | 7.02772403811355 | 11.3699574206438 | 10.7390538933119 | 35.2145853502455 |
| chr10 | 60028796 | 60028799 | - | 3.01188173062009 | 4.54798296825753 | 1.19322821036799 | 3.20132594093141 |
| chr10 | 60029199 | 60029205 | - | 3.01188173062009 | 6.82197445238629 | 2.38645642073598 | 8.53686917581709 |
| chr10 | 60792169 | 60792198 | + | 2.00792115374673 | 3.41098722619315 | 4.77291284147196 | 8.53686917581709 |
| chr10 | 60792271 | 60792290 | + | 7.02772403811355 | 5.68497871032191 | 17.8984231555198 | 4.26843458790854 |
| chr10 | 60793881 | 60793916 | + | 5.01980288436682 | 4.54798296825753 | 1.19322821036799 | 7.46976052883995 |
| chr10 | 60793951 | 60793972 | + | 11.043566345607 | 6.82197445238629 | 9.54582568294392 | 4.26843458790854 |
| chr10 | 60794157 | 60794173 | + | 18.0712903837206 | 25.0139063254164 | 17.8984231555198 | 26.6777161744284 |
| chr10 | 60794214 | 60794233 | + | 2.00792115374673 | 5.68497871032191 | 3.57968463110397 | 7.46976052883995 |
| chr10 | 60794318 | 60794340 | + | 1.00396057687336 | 3.41098722619315 | 4.77291284147196 | 1.06710864697714 |
| chr10 | 60794399 | 60794441 | + | 6.02376346124019 | 11.3699574206438 | 8.35259747257593 | 9.60397782279422 |
| chr10 | 60794554 | 60794632 | + | 30.1188173062009 | 37.5208594881246 | 47.7291284147196 | 72.5633879944452 |
| chr10 | 60794693 | 60794737 | + | 50.1980288436682 | 63.6717615556054 | 65.6275515702394 | 92.8384522870108 |
| chr10 | 60794792 | 60794830 | + | 21.0831721143407 | 25.0139063254164 | 20.2848795762558 | 10.6710864697714 |
| chr10 | 60870035 | 60870081 | - | 9.03564519186028 | 9.09596593651506 | 2.38645642073598 | 12.8053037637256 |
| chr10 | 62806050 | 62806061 | + | 22.087132691214 | 32.9728765198671 | 9.54582568294392 | 12.8053037637256 |
| chr10 | 62807027 | 62807028 | + | 1.00396057687336 | 4.54798296825753 | 1.19322821036799 | 1.06710864697714 |
| chr10 | 62807234 | 62807243 | + | 1.00396057687336 | 2.27399148412876 | 3.57968463110397 | 2.13421729395427 |
| chr10 | 63154619 | 63154622 | + | 3.01188173062009 | 6.82197445238629 | 2.38645642073598 | 5.33554323488568 |
| chr10 | 63154871 | 63154875 | + | 2.00792115374673 | 4.54798296825753 | 1.19322821036799 | 6.40265188186281 |
| chr10 | 63154962 | 63154983 | + | 2.00792115374673 | 14.780944646837 | 8.35259747257593 | 9.60397782279422 |
| chr10 | 63167429 | 63167450 | - | 1.00396057687336 | 2.27399148412876 | 3.57968463110397 | 1.06710864697714 |
| chr10 | 63620938 | 63620946 | + | 3.01188173062009 | 7.95897019445067 | 3.57968463110397 | 1.06710864697714 |
| chr10 | 63623959 | 63623962 | + | 3.01188173062009 | 1.13699574206438 | 1.19322821036799 | 1.06710864697714 |
| chr10 | 65570520 | 65570523 | + | 5.01980288436682 | 7.95897019445067 | 32.2171616799357 | 2.13421729395427 |
| chr10 | 67796694 | 67796698 | - | 88.3485307648561 | 202.38524208746 | 10.7390538933119 | 8.53686917581709 |
| chr10 | 67811487 | 67811525 | - | 9.03564519186028 | 11.3699574206438 | 1.19322821036799 | 1.06710864697714 |
| chr10 | 67916591 | 67916644 | + | 13.0514874993537 | 25.0139063254164 | 3.57968463110397 | 4.26843458790854 |
| chr10 | 67916676 | 67916741 | + | 41.1623836518079 | 75.0417189762492 | 8.35259747257593 | 20.2750642925656 |
| chr10 | 67916866 | 67916892 | + | 1.00396057687336 | 5.68497871032191 | 2.38645642073598 | 1.06710864697714 |
| chr10 | 67917183 | 67917201 | + | 4.01584230749346 | 12.5069531627082 | 8.35259747257593 | 6.40265188186281 |
| chr10 | 67917445 | 67917458 | + | 2.00792115374673 | 14.780944646837 | 4.77291284147196 | 3.20132594093141 |
| chr10 | 67922118 | 67922119 | - | 9.03564519186028 | 2.27399148412876 | 1.19322821036799 | 2.13421729395427 |
| chr10 | 68342199 | 68342247 | + | 28.1108961524542 | 44.3428339405109 | 20.2848795762558 | 24.5434988804741 |
| chr10 | 68342790 | 68342792 | + | 10.0396057687336 | 5.68497871032191 | 2.38645642073598 | 2.13421729395427 |
| chr10 | 68342971 | 68342978 | + | 1.00396057687336 | 1.13699574206438 | 7.15936926220794 | 1.06710864697714 |
| chr10 | 68343032 | 68343042 | + | 5.01980288436682 | 3.41098722619315 | 4.77291284147196 | 5.33554323488568 |
| chr10 | 68483359 | 68483364 | - | 11.043566345607 | 6.82197445238629 | 5.96614105183995 | 6.40265188186281 |
| chr10 | 68757349 | 68757384 | + | 31.1227778830743 | 43.2058381984465 | 22.6713359969918 | 14.9395210576799 |
| chr10 | 68761005 | 68761057 | + | 38.1505019211879 | 43.2058381984465 | 48.9223566250876 | 50.1541064079254 |
| chr10 | 68761069 | 68761184 | + | 80.3168461498692 | 57.9867828452835 | 118.129592826431 | 110.979299285622 |
| chr10 | 68765887 | 68766077 | + | 124.491111532297 | 90.9596593651506 | 226.713359969918 | 120.583277108416 |
| chr10 | 68771186 | 68771442 | + | 548.162474972857 | 446.839326631302 | 1035.72208659941 | 708.560141592818 |
| chr10 | 68772986 | 68773101 | + | 34.1346596136944 | 14.780944646837 | 68.0140079909754 | 56.5567582897882 |
| chr10 | 68786140 | 68786147 | + | 13.0514874993537 | 19.3289276150945 | 25.0577924177278 | 27.7448248214055 |
| chr10 | 68786581 | 68786601 | + | 3.01188173062009 | 2.27399148412876 | 8.35259747257593 | 3.20132594093141 |
| chr10 | 68786639 | 68786648 | + | 3.01188173062009 | 4.54798296825753 | 1.19322821036799 | 2.13421729395427 |
| chr10 | 68786684 | 68786687 | + | 6.02376346124019 | 9.09596593651506 | 3.57968463110397 | 4.26843458790854 |
| chr10 | 68787957 | 68787976 | + | 4.01584230749346 | 4.54798296825753 | 5.96614105183995 | 2.13421729395427 |
| chr10 | 68788170 | 68788199 | + | 11.043566345607 | 10.2329616785794 | 11.9322821036799 | 13.8724124107028 |
| chr10 | 68788225 | 68788243 | + | 5.01980288436682 | 3.41098722619315 | 4.77291284147196 | 3.20132594093141 |
| chr10 | 68788273 | 68788304 | + | 2.00792115374673 | 3.41098722619315 | 5.96614105183995 | 8.53686917581709 |
| chr10 | 68789708 | 68789718 | + | 2.00792115374673 | 4.54798296825753 | 1.19322821036799 | 1.06710864697714 |
| chr10 | 68789888 | 68789890 | + | 2.00792115374673 | 3.41098722619315 | 1.19322821036799 | 3.20132594093141 |
| chr10 | 68791221 | 68791260 | + | 8.03168461498692 | 4.54798296825753 | 1.19322821036799 | 6.40265188186281 |
| chr10 | 68791286 | 68791294 | + | 4.01584230749346 | 1.13699574206438 | 3.57968463110397 | 1.06710864697714 |
| chr10 | 68791340 | 68791344 | + | 4.01584230749346 | 7.95897019445067 | 2.38645642073598 | 10.6710864697714 |
| chr10 | 68946652 | 68946657 | + | 6.02376346124019 | 2.27399148412876 | 10.7390538933119 | 14.9395210576799 |
| chr10 | 68946782 | 68946784 | + | 1.00396057687336 | 7.95897019445067 | 9.54582568294392 | 5.33554323488568 |
| chr10 | 68983169 | 68983307 | + | 66.2613980736421 | 56.8497871032191 | 26.2510206280958 | 74.6976052883995 |
| chr10 | 68983337 | 68983428 | + | 107.42378172545 | 128.480518853275 | 91.8785721983352 | 69.3620620535138 |
| chr10 | 68983449 | 68983458 | + | 6.02376346124019 | 6.82197445238629 | 7.15936926220794 | 17.0737383516342 |
| chr10 | 68983483 | 68983599 | + | 32.1267384599477 | 38.657855230189 | 62.0478669391355 | 53.3554323488568 |
| chr10 | 68983650 | 68983687 | + | 92.3643730723495 | 65.9457530397342 | 137.221244192319 | 161.133405693548 |
| chr10 | 68983772 | 68983802 | + | 9.03564519186028 | 3.41098722619315 | 2.38645642073598 | 5.33554323488568 |
| chr10 | 68983856 | 68983928 | + | 7.02772403811355 | 15.9179403889013 | 10.7390538933119 | 34.1474767032683 |
| chr10 | 68983967 | 68984034 | + | 7.02772403811355 | 6.82197445238629 | 23.8645642073598 | 9.60397782279422 |
| chr10 | 68984081 | 68984123 | + | 15.0594086531005 | 14.780944646837 | 21.4781077866238 | 22.4092815865198 |
| chr10 | 68984149 | 68984288 | + | 40.1584230749346 | 26.1509020674808 | 79.9462900946553 | 76.8318225823538 |
| chr10 | 68984309 | 68984431 | + | 26.1029749987075 | 37.5208594881246 | 57.2749540976635 | 33.0803680562912 |
| chr10 | 68984528 | 68984578 | + | 46.1821865361748 | 63.6717615556054 | 58.4681823080315 | 84.3015831111937 |
| chr10 | 68984604 | 68984615 | + | 8.03168461498692 | 12.5069531627082 | 20.2848795762558 | 20.2750642925656 |
| chr10 | 68984645 | 68984932 | + | 1587.26167203679 | 2391.1020455614 | 5246.62444098805 | 2197.17670412592 |
| chr10 | 68984952 | 68985045 | + | 312.231739407616 | 358.15365875028 | 628.83126686393 | 493.004194903437 |
| chr10 | 69016435 | 69016436 | + | 1.00396057687336 | 1.13699574206438 | 1.19322821036799 | 1.06710864697714 |
| chr10 | 69171278 | 69171294 | + | 1.00396057687336 | 2.27399148412876 | 4.77291284147196 | 1.06710864697714 |
| chr10 | 69171347 | 69171386 | + | 11.043566345607 | 5.68497871032191 | 9.54582568294392 | 9.60397782279422 |
| chr10 | 69171482 | 69171498 | + | 10.0396057687336 | 4.54798296825753 | 10.7390538933119 | 10.6710864697714 |
| chr10 | 69171528 | 69171529 | + | 1.00396057687336 | 1.13699574206438 | 1.19322821036799 | 1.06710864697714 |
| chr10 | 69171579 | 69171589 | + | 17.0673298068472 | 10.2329616785794 | 8.35259747257593 | 14.9395210576799 |
| chr10 | 69171617 | 69171622 | + | 2.00792115374673 | 1.13699574206438 | 1.19322821036799 | 1.06710864697714 |
| chr10 | 69171645 | 69171676 | + | 11.043566345607 | 21.6029190992233 | 13.1255103140479 | 11.7381951167485 |
| chr10 | 69171883 | 69171902 | + | 2.00792115374673 | 10.2329616785794 | 7.15936926220794 | 2.13421729395427 |
| chr10 | 69171919 | 69171944 | + | 12.0475269224804 | 12.5069531627082 | 10.7390538933119 | 8.53686917581709 |
| chr10 | 69171990 | 69172027 | + | 4.01584230749346 | 9.09596593651506 | 4.77291284147196 | 9.60397782279422 |
| chr10 | 69172163 | 69172229 | + | 32.1267384599477 | 48.8908169087684 | 45.3426719939836 | 56.5567582897882 |
| chr10 | 69172371 | 69172407 | + | 35.1386201905678 | 29.5618892936739 | 15.5119667347839 | 16.006629704657 |
| chr10 | 69172622 | 69172640 | + | 5.01980288436682 | 3.41098722619315 | 3.57968463110397 | 4.26843458790854 |
| chr10 | 69172645 | 69172648 | + | 2.00792115374673 | 2.27399148412876 | 1.19322821036799 | 6.40265188186281 |
| chr10 | 69172664 | 69172679 | + | 3.01188173062009 | 4.54798296825753 | 4.77291284147196 | 1.06710864697714 |
| chr10 | 69172795 | 69172833 | + | 3.01188173062009 | 7.95897019445067 | 3.57968463110397 | 2.13421729395427 |
| chr10 | 69209020 | 69209036 | + | 11.043566345607 | 9.09596593651506 | 8.35259747257593 | 3.20132594093141 |
| chr10 | 69401218 | 69401254 | + | 2.00792115374673 | 1.13699574206438 | 16.7051949451519 | 6.40265188186281 |
| chr10 | 69401739 | 69401762 | + | 16.0633692299738 | 5.68497871032191 | 144.380613454527 | 359.615614031295 |
| chr10 | 69401840 | 69401859 | + | 2.00792115374673 | 4.54798296825753 | 77.5598336739193 | 68.2949534065367 |
| chr10 | 70139231 | 70139260 | - | 23.0910932680874 | 40.9318467143178 | 2.38645642073598 | 1.06710864697714 |
| chr10 | 70150262 | 70150299 | - | 3.01188173062009 | 11.3699574206438 | 1.19322821036799 | 10.6710864697714 |
| chr10 | 70150332 | 70150381 | - | 15.0594086531005 | 6.82197445238629 | 3.57968463110397 | 4.26843458790854 |
| chr10 | 70150419 | 70150551 | - | 226.89509037338 | 183.056314472366 | 110.970223564223 | 98.1739955218965 |
| chr10 | 70150583 | 70150644 | - | 146.578244223511 | 102.329616785794 | 82.3327465153913 | 98.1739955218965 |
| chr10 | 70150672 | 70150688 | - | 15.0594086531005 | 20.4659233571589 | 11.9322821036799 | 10.6710864697714 |
| chr10 | 70150710 | 70150816 | - | 219.867366335267 | 171.686357051722 | 72.7869208324474 | 46.952780466994 |
| chr10 | 70150848 | 70150999 | - | 131.518835570411 | 86.411676396893 | 118.129592826431 | 59.7580842307196 |
| chr10 | 70151055 | 70151081 | - | 9.03564519186028 | 11.3699574206438 | 1.19322821036799 | 3.20132594093141 |
| chr10 | 70151098 | 70151125 | - | 17.0673298068472 | 7.95897019445067 | 11.9322821036799 | 7.46976052883995 |
| chr10 | 70151218 | 70151226 | - | 17.0673298068472 | 5.68497871032191 | 8.35259747257593 | 11.7381951167485 |
| chr10 | 70151302 | 70151328 | - | 9.03564519186028 | 4.54798296825753 | 1.19322821036799 | 2.13421729395427 |
| chr10 | 70151369 | 70151388 | - | 25.0990144218341 | 13.6439489047726 | 10.7390538933119 | 3.20132594093141 |
| chr10 | 70151456 | 70151472 | - | 4.01584230749346 | 5.68497871032191 | 2.38645642073598 | 1.06710864697714 |
| chr10 | 70151538 | 70151553 | - | 8.03168461498692 | 5.68497871032191 | 5.96614105183995 | 7.46976052883995 |
| chr10 | 70151696 | 70151865 | - | 110.43566345607 | 96.6446380754725 | 212.394621445502 | 99.2411041688736 |
| chr10 | 70151890 | 70151900 | - | 8.03168461498692 | 9.09596593651506 | 1.19322821036799 | 6.40265188186281 |
| chr10 | 70152063 | 70152315 | - | 208.82379998966 | 181.919318730301 | 169.438405872254 | 130.187254931211 |
| chr10 | 70152362 | 70152369 | - | 7.02772403811355 | 1.13699574206438 | 3.57968463110397 | 3.20132594093141 |
| chr10 | 70152451 | 70152543 | - | 69.2732798042621 | 79.5897019445067 | 42.9562155732476 | 51.2212150549025 |
| chr10 | 70202855 | 70202863 | - | 22.087132691214 | 40.9318467143178 | 11.9322821036799 | 6.40265188186281 |
| chr10 | 70202949 | 70202968 | - | 3.01188173062009 | 4.54798296825753 | 1.19322821036799 | 2.13421729395427 |
| chr10 | 70203026 | 70203032 | - | 3.01188173062009 | 1.13699574206438 | 2.38645642073598 | 1.06710864697714 |
| chr10 | 70203092 | 70203189 | - | 646.550611506447 | 600.333751809994 | 678.946851699386 | 386.293330205723 |
| chr10 | 70299195 | 70299255 | - | 7.02772403811355 | 4.54798296825753 | 2.38645642073598 | 1.06710864697714 |
| chr10 | 70301222 | 70301253 | - | 1.00396057687336 | 5.68497871032191 | 8.35259747257593 | 3.20132594093141 |
| chr10 | 70422007 | 70422022 | + | 1.00396057687336 | 1.13699574206438 | 7.15936926220794 | 4.26843458790854 |
| chr10 | 70422136 | 70422176 | + | 2.00792115374673 | 6.82197445238629 | 27.4442488384638 | 8.53686917581709 |
| chr10 | 70422240 | 70422248 | + | 2.00792115374673 | 1.13699574206438 | 2.38645642073598 | 1.06710864697714 |
| chr10 | 70422259 | 70422288 | + | 3.01188173062009 | 4.54798296825753 | 4.77291284147196 | 7.46976052883995 |
| chr10 | 70422332 | 70422381 | + | 3.01188173062009 | 3.41098722619315 | 14.3187385244159 | 8.53686917581709 |
| chr10 | 70422496 | 70422499 | + | 2.00792115374673 | 1.13699574206438 | 1.19322821036799 | 6.40265188186281 |
| chr10 | 70422596 | 70422781 | + | 91.3604124954762 | 61.3977700714766 | 175.404546924094 | 260.374509862421 |
| chr10 | 70422898 | 70422943 | + | 15.0594086531005 | 14.780944646837 | 16.7051949451519 | 6.40265188186281 |
| chr10 | 70423050 | 70423059 | + | 2.00792115374673 | 2.27399148412876 | 7.15936926220794 | 5.33554323488568 |
| chr10 | 70423440 | 70423486 | + | 4.01584230749346 | 1.13699574206438 | 10.7390538933119 | 16.006629704657 |
| chr10 | 70423593 | 70423599 | + | 1.00396057687336 | 3.41098722619315 | 4.77291284147196 | 6.40265188186281 |
| chr10 | 70423808 | 70423812 | + | 2.00792115374673 | 5.68497871032191 | 4.77291284147196 | 4.26843458790854 |
| chr10 | 70425438 | 70425464 | + | 7.02772403811355 | 2.27399148412876 | 4.77291284147196 | 7.46976052883995 |
| chr10 | 70426558 | 70426580 | + | 221.875287489014 | 154.631420920756 | 17.8984231555198 | 27.7448248214055 |
| chr10 | 70426683 | 70426713 | + | 3.01188173062009 | 1.13699574206438 | 5.96614105183995 | 8.53686917581709 |
| chr10 | 70426733 | 70426738 | + | 6.02376346124019 | 4.54798296825753 | 5.96614105183995 | 1.06710864697714 |
| chr10 | 70426774 | 70426818 | + | 8.03168461498692 | 5.68497871032191 | 15.5119667347839 | 8.53686917581709 |
| chr10 | 70428577 | 70428581 | + | 2.00792115374673 | 1.13699574206438 | 2.38645642073598 | 2.13421729395427 |
| chr10 | 70883528 | 70883662 | - | 24.0950538449607 | 17.0549361309657 | 13.1255103140479 | 10.6710864697714 |
| chr10 | 70883695 | 70883739 | - | 61.2415951892752 | 51.1648083928972 | 32.2171616799357 | 44.8185631730397 |
| chr10 | 70883794 | 70883831 | - | 8.03168461498692 | 1.13699574206438 | 3.57968463110397 | 7.46976052883995 |
| chr10 | 71363360 | 71363362 | + | 3.01188173062009 | 4.54798296825753 | 1.19322821036799 | 1.06710864697714 |
| chr10 | 71816318 | 71816500 | - | 107.42378172545 | 143.261463500112 | 102.617626091647 | 118.449059814462 |
| chr10 | 71816518 | 71817172 | - | 614.423873046499 | 462.757267020203 | 726.675980114106 | 662.674469772801 |
| chr10 | 71817198 | 71817477 | - | 268.057474025188 | 222.851165444619 | 378.253342686653 | 277.448248214055 |
| chr10 | 72096613 | 72096615 | - | 5.01980288436682 | 4.54798296825753 | 1.19322821036799 | 5.33554323488568 |
| chr10 | 72233128 | 72233169 | + | 10.0396057687336 | 23.876910583352 | 25.0577924177278 | 18.1408469986113 |
| chr10 | 72233413 | 72233443 | + | 1.00396057687336 | 4.54798296825753 | 1.19322821036799 | 2.13421729395427 |
| chr10 | 72233464 | 72233489 | + | 1.00396057687336 | 1.13699574206438 | 3.57968463110397 | 5.33554323488568 |
| chr10 | 72233548 | 72233590 | + | 3.01188173062009 | 4.54798296825753 | 19.0916513658878 | 11.7381951167485 |
| chr10 | 72233778 | 72233786 | + | 2.00792115374673 | 4.54798296825753 | 3.57968463110397 | 1.06710864697714 |
| chr10 | 72275176 | 72275287 | + | 15.0594086531005 | 29.5618892936739 | 52.5020412561915 | 26.6777161744284 |
| chr10 | 72275297 | 72275380 | + | 18.0712903837206 | 43.2058381984465 | 46.5359002043516 | 37.3488026441998 |
| chr10 | 72275405 | 72275459 | + | 2.00792115374673 | 18.1919318730301 | 22.6713359969918 | 23.476390233497 |
| chr10 | 72275492 | 72275497 | + | 3.01188173062009 | 1.13699574206438 | 2.38645642073598 | 16.006629704657 |
| chr10 | 72275544 | 72275560 | + | 7.02772403811355 | 15.9179403889013 | 14.3187385244159 | 18.1408469986113 |
| chr10 | 72275581 | 72275646 | + | 56.2217923049084 | 155.76841666282 | 194.496198289982 | 88.5700176991023 |
| chr10 | 72275708 | 72275766 | + | 4.01584230749346 | 7.95897019445067 | 17.8984231555198 | 11.7381951167485 |
| chr10 | 72275863 | 72275912 | + | 68.2693192273888 | 336.550739651057 | 215.974306076606 | 251.837640686604 |
| chr10 | 72275935 | 72276010 | + | 16.0633692299738 | 86.411676396893 | 58.4681823080315 | 18.1408469986113 |
| chr10 | 72334569 | 72334572 | - | 1.00396057687336 | 2.27399148412876 | 1.19322821036799 | 2.13421729395427 |
| chr10 | 72367406 | 72367449 | - | 7.02772403811355 | 2.27399148412876 | 3.57968463110397 | 18.1408469986113 |
| chr10 | 72367854 | 72367943 | - | 10.0396057687336 | 10.2329616785794 | 23.8645642073598 | 33.0803680562912 |
| chr10 | 72368061 | 72368105 | - | 4.01584230749346 | 2.27399148412876 | 16.7051949451519 | 6.40265188186281 |
| chr10 | 72368174 | 72368178 | - | 4.01584230749346 | 1.13699574206438 | 5.96614105183995 | 3.20132594093141 |
| chr10 | 72721085 | 72721088 | + | 1.00396057687336 | 1.13699574206438 | 5.96614105183995 | 4.26843458790854 |
| chr10 | 72885912 | 72885915 | + | 18.0712903837206 | 18.1919318730301 | 13.1255103140479 | 5.33554323488568 |
| chr10 | 72886118 | 72886121 | + | 4.01584230749346 | 7.95897019445067 | 1.19322821036799 | 5.33554323488568 |
| chr10 | 72886507 | 72886522 | + | 2.00792115374673 | 3.41098722619315 | 5.96614105183995 | 8.53686917581709 |
| chr10 | 72886877 | 72886934 | + | 32.1267384599477 | 22.7399148412876 | 21.4781077866238 | 21.3421729395427 |
| chr10 | 72887102 | 72887128 | + | 26.1029749987075 | 12.5069531627082 | 33.4103898903037 | 13.8724124107028 |
| chr10 | 72887182 | 72887189 | + | 7.02772403811355 | 4.54798296825753 | 9.54582568294392 | 1.06710864697714 |
| chr10 | 72887525 | 72887575 | + | 17.0673298068472 | 27.2878978095452 | 15.5119667347839 | 21.3421729395427 |
| chr10 | 73007237 | 73007258 | - | 2.00792115374673 | 2.27399148412876 | 10.7390538933119 | 1.06710864697714 |
| chr10 | 73007334 | 73007354 | - | 4.01584230749346 | 13.6439489047726 | 14.3187385244159 | 8.53686917581709 |
| chr10 | 73007416 | 73007424 | - | 1.00396057687336 | 6.82197445238629 | 10.7390538933119 | 24.5434988804741 |
| chr10 | 73007456 | 73007499 | - | 6.02376346124019 | 1.13699574206438 | 54.8884976769275 | 57.6238669367653 |
| chr10 | 73007602 | 73007670 | - | 20.0792115374673 | 43.2058381984465 | 94.2650286190712 | 80.0331485232852 |
| chr10 | 73007696 | 73007756 | - | 6.02376346124019 | 7.95897019445067 | 50.1155848354556 | 29.8790421153598 |
| chr10 | 73007787 | 73008003 | - | 8.03168461498692 | 22.7399148412876 | 78.7530618842873 | 59.7580842307196 |
| chr10 | 73242862 | 73242879 | - | 8.03168461498692 | 17.0549361309657 | 25.0577924177278 | 8.53686917581709 |
| chr10 | 73243084 | 73243090 | - | 3.01188173062009 | 4.54798296825753 | 1.19322821036799 | 3.20132594093141 |
| chr10 | 73248914 | 73248964 | - | 6.02376346124019 | 11.3699574206438 | 25.0577924177278 | 18.1408469986113 |
| chr10 | 73249739 | 73249747 | - | 17.0673298068472 | 13.6439489047726 | 3.57968463110397 | 4.26843458790854 |
| chr10 | 73249813 | 73249819 | - | 8.03168461498692 | 10.2329616785794 | 3.57968463110397 | 6.40265188186281 |
| chr10 | 73249939 | 73249947 | - | 17.0673298068472 | 12.5069531627082 | 4.77291284147196 | 11.7381951167485 |
| chr10 | 73250238 | 73250240 | - | 10.0396057687336 | 9.09596593651506 | 9.54582568294392 | 5.33554323488568 |
| chr10 | 73250267 | 73250268 | - | 2.00792115374673 | 1.13699574206438 | 4.77291284147196 | 4.26843458790854 |
| chr10 | 73250410 | 73250442 | - | 37.1465413443145 | 65.9457530397342 | 28.6374770488317 | 24.5434988804741 |
| chr10 | 73250661 | 73250703 | - | 13.0514874993537 | 22.7399148412876 | 21.4781077866238 | 17.0737383516342 |
| chr10 | 73250750 | 73250824 | - | 38.1505019211879 | 40.9318467143178 | 29.8307052591997 | 21.3421729395427 |
| chr10 | 73250849 | 73250882 | - | 7.02772403811355 | 10.2329616785794 | 3.57968463110397 | 17.0737383516342 |
| chr10 | 73252727 | 73252840 | + | 89.3524913417294 | 154.631420920756 | 108.583767143487 | 173.938709457273 |
| chr10 | 73253242 | 73253256 | + | 5.01980288436682 | 14.780944646837 | 8.35259747257593 | 11.7381951167485 |
| chr10 | 73254165 | 73254167 | + | 3.01188173062009 | 2.27399148412876 | 3.57968463110397 | 1.06710864697714 |
| chr10 | 73254283 | 73254321 | + | 4.01584230749346 | 32.9728765198671 | 5.96614105183995 | 7.46976052883995 |
| chr10 | 73375513 | 73375560 | - | 20.0792115374673 | 20.4659233571589 | 20.2848795762558 | 8.53686917581709 |
| chr10 | 73375695 | 73375713 | - | 5.01980288436682 | 2.27399148412876 | 4.77291284147196 | 2.13421729395427 |
| chr10 | 73375802 | 73375814 | - | 5.01980288436682 | 7.95897019445067 | 3.57968463110397 | 4.26843458790854 |
| chr10 | 73375933 | 73375998 | - | 11.043566345607 | 7.95897019445067 | 4.77291284147196 | 7.46976052883995 |
| chr10 | 73437331 | 73437364 | - | 2.00792115374673 | 1.13699574206438 | 1.19322821036799 | 3.20132594093141 |
| chr10 | 73437990 | 73438019 | - | 2.00792115374673 | 5.68497871032191 | 9.54582568294392 | 10.6710864697714 |
| chr10 | 73443880 | 73443883 | - | 3.01188173062009 | 4.54798296825753 | 3.57968463110397 | 3.20132594093141 |
| chr10 | 73495693 | 73495766 | + | 85.336649034236 | 93.2336508492793 | 16.7051949451519 | 44.8185631730397 |
| chr10 | 73626061 | 73626062 | + | 1.00396057687336 | 1.13699574206438 | 1.19322821036799 | 1.06710864697714 |
| chr10 | 73626070 | 73626120 | + | 14.0554480762271 | 19.3289276150945 | 11.9322821036799 | 22.4092815865198 |
| chr10 | 73771068 | 73771165 | + | 5.01980288436682 | 5.68497871032191 | 28.6374770488317 | 21.3421729395427 |
| chr10 | 73771196 | 73771245 | + | 24.0950538449607 | 13.6439489047726 | 50.1155848354556 | 39.483019938154 |
| chr10 | 73771329 | 73771416 | + | 17.0673298068472 | 4.54798296825753 | 19.0916513658878 | 23.476390233497 |
| chr10 | 73771446 | 73771494 | + | 8.03168461498692 | 5.68497871032191 | 36.9900745214077 | 43.7514545260626 |
| chr10 | 73771515 | 73771602 | + | 22.087132691214 | 18.1919318730301 | 54.8884976769275 | 51.2212150549025 |
| chr10 | 73771674 | 73771719 | + | 7.02772403811355 | 6.82197445238629 | 29.8307052591997 | 29.8790421153598 |
| chr10 | 73771837 | 73771900 | + | 10.0396057687336 | 2.27399148412876 | 23.8645642073598 | 25.6106075274513 |
| chr10 | 73772007 | 73772106 | + | 12.0475269224804 | 5.68497871032191 | 14.3187385244159 | 26.6777161744284 |
| chr10 | 73775586 | 73775602 | + | 5.01980288436682 | 4.54798296825753 | 7.15936926220794 | 1.06710864697714 |
| chr10 | 73775879 | 73775912 | + | 4.01584230749346 | 5.68497871032191 | 9.54582568294392 | 8.53686917581709 |
| chr10 | 73775945 | 73775959 | + | 3.01188173062009 | 1.13699574206438 | 3.57968463110397 | 3.20132594093141 |
| chr10 | 73776028 | 73776033 | + | 3.01188173062009 | 3.41098722619315 | 7.15936926220794 | 4.26843458790854 |
| chr10 | 73776134 | 73776161 | + | 3.01188173062009 | 1.13699574206438 | 3.57968463110397 | 4.26843458790854 |
| chr10 | 73783040 | 73783041 | + | 4.01584230749346 | 1.13699574206438 | 2.38645642073598 | 8.53686917581709 |
| chr10 | 73783078 | 73783122 | + | 4.01584230749346 | 11.3699574206438 | 3.57968463110397 | 8.53686917581709 |
| chr10 | 73783191 | 73783213 | + | 2.00792115374673 | 2.27399148412876 | 2.38645642073598 | 3.20132594093141 |
| chr10 | 73783280 | 73783351 | + | 115.455466340437 | 169.412365567593 | 125.288962088639 | 228.361250453107 |
| chr10 | 73801706 | 73801761 | + | 23.0910932680874 | 11.3699574206438 | 4.77291284147196 | 17.0737383516342 |
| chr10 | 73801940 | 73801972 | - | 1.00396057687336 | 6.82197445238629 | 7.15936926220794 | 4.26843458790854 |
| chr10 | 73813458 | 73813479 | - | 5.01980288436682 | 4.54798296825753 | 13.1255103140479 | 7.46976052883995 |
| chr10 | 73813684 | 73813705 | - | 2.00792115374673 | 2.27399148412876 | 11.9322821036799 | 8.53686917581709 |
| chr10 | 73814105 | 73814111 | - | 3.01188173062009 | 2.27399148412876 | 3.57968463110397 | 4.26843458790854 |
| chr10 | 73814143 | 73814144 | - | 4.01584230749346 | 2.27399148412876 | 3.57968463110397 | 2.13421729395427 |
| chr10 | 73814432 | 73814452 | - | 5.01980288436682 | 2.27399148412876 | 14.3187385244159 | 4.26843458790854 |
| chr10 | 73814480 | 73814508 | - | 15.0594086531005 | 20.4659233571589 | 29.8307052591997 | 20.2750642925656 |
| chr10 | 73911204 | 73911205 | - | 6.02376346124019 | 4.54798296825753 | 3.57968463110397 | 9.60397782279422 |
| chr10 | 73917388 | 73917393 | + | 2.00792115374673 | 1.13699574206438 | 3.57968463110397 | 1.06710864697714 |
| chr10 | 74118301 | 74118303 | + | 15.0594086531005 | 3.41098722619315 | 10.7390538933119 | 4.26843458790854 |
| chr10 | 74118685 | 74118754 | + | 11.043566345607 | 4.54798296825753 | 28.6374770488317 | 33.0803680562912 |
| chr10 | 74118859 | 74118933 | + | 2.00792115374673 | 3.41098722619315 | 25.0577924177278 | 25.6106075274513 |
| chr10 | 74119021 | 74119089 | + | 17.0673298068472 | 9.09596593651506 | 54.8884976769275 | 61.8923015246739 |
| chr10 | 74119114 | 74119136 | + | 4.01584230749346 | 1.13699574206438 | 16.7051949451519 | 11.7381951167485 |
| chr10 | 74119165 | 74119278 | + | 23.0910932680874 | 20.4659233571589 | 89.4921157775992 | 75.7647139353766 |
| chr10 | 74119303 | 74119351 | + | 9.03564519186028 | 3.41098722619315 | 25.0577924177278 | 49.0869977609482 |
| chr10 | 74119377 | 74119402 | + | 2.00792115374673 | 2.27399148412876 | 3.57968463110397 | 4.26843458790854 |
| chr10 | 74119468 | 74119484 | + | 4.01584230749346 | 6.82197445238629 | 5.96614105183995 | 26.6777161744284 |
| chr10 | 74119538 | 74119544 | + | 9.03564519186028 | 6.82197445238629 | 28.6374770488317 | 59.7580842307196 |
| chr10 | 74119633 | 74119651 | + | 3.01188173062009 | 3.41098722619315 | 4.77291284147196 | 3.20132594093141 |
| chr10 | 74119714 | 74119734 | + | 7.02772403811355 | 4.54798296825753 | 13.1255103140479 | 14.9395210576799 |
| chr10 | 74119758 | 74119801 | + | 15.0594086531005 | 15.9179403889013 | 32.2171616799357 | 42.6843458790854 |
| chr10 | 74119890 | 74119892 | + | 1.00396057687336 | 1.13699574206438 | 1.19322821036799 | 2.13421729395427 |
| chr10 | 74119931 | 74119933 | + | 2.00792115374673 | 1.13699574206438 | 3.57968463110397 | 2.13421729395427 |
| chr10 | 74121910 | 74121913 | - | 4.01584230749346 | 3.41098722619315 | 5.96614105183995 | 6.40265188186281 |
| chr10 | 74122085 | 74122165 | - | 50.1980288436682 | 48.8908169087684 | 45.3426719939836 | 34.1474767032683 |
| chr10 | 74122167 | 74122195 | - | 1.00396057687336 | 2.27399148412876 | 4.77291284147196 | 3.20132594093141 |
| chr10 | 74122217 | 74122271 | - | 29.1148567293276 | 25.0139063254164 | 23.8645642073598 | 34.1474767032683 |
| chr10 | 74122426 | 74122450 | - | 1.00396057687336 | 2.27399148412876 | 8.35259747257593 | 2.13421729395427 |
| chr10 | 74122488 | 74122491 | - | 1.00396057687336 | 1.13699574206438 | 4.77291284147196 | 1.06710864697714 |
| chr10 | 74122587 | 74122600 | - | 9.03564519186028 | 2.27399148412876 | 3.57968463110397 | 3.20132594093141 |
| chr10 | 74122616 | 74122638 | - | 2.00792115374673 | 2.27399148412876 | 4.77291284147196 | 8.53686917581709 |
| chr10 | 74122754 | 74122759 | - | 4.01584230749346 | 9.09596593651506 | 10.7390538933119 | 6.40265188186281 |
| chr10 | 74122826 | 74122851 | - | 2.00792115374673 | 1.13699574206438 | 9.54582568294392 | 13.8724124107028 |
| chr10 | 74123019 | 74123042 | - | 5.01980288436682 | 2.27399148412876 | 4.77291284147196 | 6.40265188186281 |
| chr10 | 74123066 | 74123069 | - | 6.02376346124019 | 5.68497871032191 | 3.57968463110397 | 4.26843458790854 |
| chr10 | 74123371 | 74123373 | - | 5.01980288436682 | 1.13699574206438 | 2.38645642073598 | 4.26843458790854 |
| chr10 | 74123520 | 74123590 | - | 8.03168461498692 | 9.09596593651506 | 32.2171616799357 | 9.60397782279422 |
| chr10 | 74123619 | 74123624 | - | 1.00396057687336 | 2.27399148412876 | 3.57968463110397 | 2.13421729395427 |
| chr10 | 74123646 | 74123663 | - | 5.01980288436682 | 1.13699574206438 | 19.0916513658878 | 4.26843458790854 |
| chr10 | 74123731 | 74123740 | - | 1.00396057687336 | 2.27399148412876 | 8.35259747257593 | 4.26843458790854 |
| chr10 | 74123770 | 74123896 | - | 12.0475269224804 | 18.1919318730301 | 26.2510206280958 | 22.4092815865198 |
| chr10 | 74708404 | 74708503 | + | 5.01980288436682 | 5.68497871032191 | 7.15936926220794 | 16.006629704657 |
| chr10 | 74708581 | 74708592 | + | 10.0396057687336 | 6.82197445238629 | 7.15936926220794 | 16.006629704657 |
| chr10 | 75176573 | 75176586 | + | 2.00792115374673 | 2.27399148412876 | 1.19322821036799 | 2.13421729395427 |
| chr10 | 75177409 | 75177411 | + | 3.01188173062009 | 5.68497871032191 | 1.19322821036799 | 1.06710864697714 |
| chr10 | 75177595 | 75177614 | + | 10.0396057687336 | 10.2329616785794 | 1.19322821036799 | 4.26843458790854 |
| chr10 | 75179870 | 75179953 | + | 7.02772403811355 | 13.6439489047726 | 11.9322821036799 | 6.40265188186281 |
| chr10 | 75179986 | 75180000 | + | 4.01584230749346 | 6.82197445238629 | 9.54582568294392 | 19.2079556455884 |
| chr10 | 75181181 | 75181194 | + | 2.00792115374673 | 4.54798296825753 | 1.19322821036799 | 2.13421729395427 |
| chr10 | 75181842 | 75181890 | + | 6.02376346124019 | 5.68497871032191 | 1.19322821036799 | 2.13421729395427 |
| chr10 | 75231139 | 75231155 | + | 6.02376346124019 | 3.41098722619315 | 5.96614105183995 | 7.46976052883995 |
| chr10 | 75231252 | 75231289 | + | 14.0554480762271 | 26.1509020674808 | 35.7968463110397 | 8.53686917581709 |
| chr10 | 75231324 | 75231418 | + | 34.1346596136944 | 46.6168254246397 | 41.7629873628796 | 44.8185631730397 |
| chr10 | 75234031 | 75234045 | - | 6.02376346124019 | 6.82197445238629 | 9.54582568294392 | 7.46976052883995 |
| chr10 | 78040204 | 78040218 | + | 17.0673298068472 | 9.09596593651506 | 23.8645642073598 | 20.2750642925656 |
| chr10 | 78040613 | 78040678 | + | 525.07138170477 | 674.238475044178 | 615.705756549883 | 640.265188186281 |
| chr10 | 79313594 | 79313597 | + | 4.01584230749346 | 2.27399148412876 | 1.19322821036799 | 2.13421729395427 |
| chr10 | 79315961 | 79315990 | + | 1.00396057687336 | 1.13699574206438 | 3.57968463110397 | 6.40265188186281 |
| chr10 | 79353779 | 79353873 | + | 17.0673298068472 | 10.2329616785794 | 22.6713359969918 | 27.7448248214055 |
| chr10 | 79353900 | 79353908 | + | 7.02772403811355 | 4.54798296825753 | 3.57968463110397 | 3.20132594093141 |
| chr10 | 79353958 | 79354002 | + | 16.0633692299738 | 12.5069531627082 | 14.3187385244159 | 16.006629704657 |
| chr10 | 79354038 | 79354066 | + | 4.01584230749346 | 5.68497871032191 | 7.15936926220794 | 12.8053037637256 |
| chr10 | 79354097 | 79354105 | + | 1.00396057687336 | 1.13699574206438 | 1.19322821036799 | 1.06710864697714 |
| chr10 | 79354128 | 79354279 | + | 53.2099105742883 | 57.9867828452835 | 133.641559561215 | 139.791232754005 |
| chr10 | 79354320 | 79354414 | + | 25.0990144218341 | 28.4248935516096 | 34.6036181006717 | 58.6909755837425 |
| chr10 | 79354456 | 79354611 | + | 29.1148567293276 | 40.9318467143178 | 19.0916513658878 | 57.6238669367653 |
| chr10 | 79354633 | 79354697 | + | 3.01188173062009 | 4.54798296825753 | 5.96614105183995 | 3.20132594093141 |
| chr10 | 79354728 | 79354757 | + | 1.00396057687336 | 1.13699574206438 | 3.57968463110397 | 1.06710864697714 |
| chr10 | 79383919 | 79383985 | - | 1.00396057687336 | 3.41098722619315 | 4.77291284147196 | 11.7381951167485 |
| chr10 | 79513735 | 79513738 | + | 11.043566345607 | 11.3699574206438 | 10.7390538933119 | 5.33554323488568 |
| chr10 | 79905028 | 79905029 | + | 2.00792115374673 | 1.13699574206438 | 1.19322821036799 | 1.06710864697714 |
| chr10 | 80152688 | 80152706 | - | 2.00792115374673 | 3.41098722619315 | 8.35259747257593 | 4.26843458790854 |
| chr10 | 80155617 | 80155633 | - | 10.0396057687336 | 10.2329616785794 | 7.15936926220794 | 1.06710864697714 |
| chr10 | 80155656 | 80155685 | - | 25.0990144218341 | 23.876910583352 | 19.0916513658878 | 21.3421729395427 |
| chr10 | 80518681 | 80518735 | + | 2.00792115374673 | 1.13699574206438 | 1.19322821036799 | 1.06710864697714 |
| chr10 | 80518757 | 80518785 | + | 7.02772403811355 | 2.27399148412876 | 7.15936926220794 | 2.13421729395427 |
| chr10 | 80519432 | 80519505 | + | 17.0673298068472 | 12.5069531627082 | 29.8307052591997 | 78.966039876308 |
| chr10 | 80519539 | 80519603 | + | 2.00792115374673 | 3.41098722619315 | 11.9322821036799 | 11.7381951167485 |
| chr10 | 80519704 | 80519728 | + | 3.01188173062009 | 1.13699574206438 | 4.77291284147196 | 7.46976052883995 |
| chr10 | 80520535 | 80520563 | + | 6.02376346124019 | 9.09596593651506 | 1.19322821036799 | 3.20132594093141 |
| chr10 | 84152262 | 84153022 | + | 2133.4162258559 | 1984.05756990235 | 2499.81310072094 | 2771.28115619962 |
| chr10 | 84153033 | 84153064 | + | 2.00792115374673 | 4.54798296825753 | 7.15936926220794 | 4.26843458790854 |
| chr10 | 84153087 | 84153089 | + | 5.01980288436682 | 4.54798296825753 | 5.96614105183995 | 2.13421729395427 |
| chr10 | 84153110 | 84153153 | + | 8.03168461498692 | 6.82197445238629 | 7.15936926220794 | 6.40265188186281 |
| chr10 | 84518074 | 84518076 | + | 2.00792115374673 | 3.41098722619315 | 1.19322821036799 | 1.06710864697714 |
| chr10 | 86435344 | 86435364 | - | 1.00396057687336 | 6.82197445238629 | 4.77291284147196 | 5.33554323488568 |
| chr10 | 86435587 | 86435605 | - | 13.0514874993537 | 13.6439489047726 | 2.38645642073598 | 6.40265188186281 |
| chr10 | 86436747 | 86436769 | - | 7.02772403811355 | 5.68497871032191 | 2.38645642073598 | 2.13421729395427 |
| chr10 | 86437075 | 86437154 | - | 11.043566345607 | 18.1919318730301 | 10.7390538933119 | 17.0737383516342 |
| chr10 | 86437219 | 86437240 | - | 1.00396057687336 | 6.82197445238629 | 3.57968463110397 | 6.40265188186281 |
| chr10 | 86437271 | 86437276 | - | 7.02772403811355 | 15.9179403889013 | 2.38645642073598 | 16.006629704657 |
| chr10 | 86437299 | 86437334 | - | 15.0594086531005 | 13.6439489047726 | 2.38645642073598 | 8.53686917581709 |
| chr10 | 86437420 | 86437448 | - | 4.01584230749346 | 9.09596593651506 | 1.19322821036799 | 1.06710864697714 |
| chr10 | 86521937 | 86521988 | + | 7.02772403811355 | 10.2329616785794 | 7.15936926220794 | 3.20132594093141 |
| chr10 | 86963153 | 86963209 | + | 10.0396057687336 | 2.27399148412876 | 21.4781077866238 | 144.059667341913 |
| chr10 | 87050503 | 87050523 | - | 8.03168461498692 | 11.3699574206438 | 4.77291284147196 | 4.26843458790854 |
| chr10 | 87050581 | 87050661 | - | 10.0396057687336 | 7.95897019445067 | 14.3187385244159 | 7.46976052883995 |
| chr10 | 87050695 | 87050712 | - | 1.00396057687336 | 2.27399148412876 | 1.19322821036799 | 2.13421729395427 |
| chr10 | 87050918 | 87051044 | - | 3.01188173062009 | 2.27399148412876 | 1.19322821036799 | 1.06710864697714 |
| chr10 | 87051224 | 87051281 | - | 6.02376346124019 | 6.82197445238629 | 10.7390538933119 | 5.33554323488568 |
| chr10 | 87051343 | 87051462 | - | 59.2336740355285 | 51.1648083928972 | 87.1056593568632 | 69.3620620535138 |
| chr10 | 87051569 | 87051622 | - | 12.0475269224804 | 10.2329616785794 | 20.2848795762558 | 13.8724124107028 |
| chr10 | 87190992 | 87191024 | + | 17.0673298068472 | 18.1919318730301 | 15.5119667347839 | 14.9395210576799 |
| chr10 | 87342918 | 87342920 | + | 1.00396057687336 | 4.54798296825753 | 1.19322821036799 | 2.13421729395427 |
| chr10 | 87552802 | 87552831 | + | 1.00396057687336 | 1.13699574206438 | 2.38645642073598 | 5.33554323488568 |
| chr10 | 87552932 | 87552966 | + | 1.00396057687336 | 1.13699574206438 | 2.38645642073598 | 5.33554323488568 |
| chr10 | 87752068 | 87752079 | - | 23.0910932680874 | 43.2058381984465 | 4.77291284147196 | 2.13421729395427 |
| chr10 | 87753217 | 87753238 | - | 5.01980288436682 | 20.4659233571589 | 3.57968463110397 | 2.13421729395427 |
| chr10 | 87753265 | 87753286 | - | 19.0752509605939 | 61.3977700714766 | 2.38645642073598 | 7.46976052883995 |
| chr10 | 87754492 | 87754494 | - | 2.00792115374673 | 4.54798296825753 | 1.19322821036799 | 2.13421729395427 |
| chr10 | 87754683 | 87754702 | - | 13.0514874993537 | 12.5069531627082 | 1.19322821036799 | 1.06710864697714 |
| chr10 | 89584317 | 89584342 | - | 4.01584230749346 | 6.82197445238629 | 1.19322821036799 | 1.06710864697714 |
| chr10 | 89774886 | 89774888 | + | 3.01188173062009 | 7.95897019445067 | 5.96614105183995 | 3.20132594093141 |
| chr10 | 89838328 | 89838329 | + | 14.0554480762271 | 21.6029190992233 | 31.0239334695677 | 29.8790421153598 |
| chr10 | 91279417 | 91279430 | + | 1.00396057687336 | 2.27399148412876 | 7.15936926220794 | 2.13421729395427 |
| chr10 | 91281633 | 91281661 | + | 6.02376346124019 | 12.5069531627082 | 4.77291284147196 | 1.06710864697714 |
| chr10 | 91283550 | 91283575 | + | 1.00396057687336 | 3.41098722619315 | 1.19322821036799 | 6.40265188186281 |
| chr10 | 91283596 | 91283615 | + | 1.00396057687336 | 1.13699574206438 | 3.57968463110397 | 2.13421729395427 |
| chr10 | 91283649 | 91283661 | + | 2.00792115374673 | 2.27399148412876 | 5.96614105183995 | 2.13421729395427 |
| chr10 | 91865071 | 91865123 | + | 121.479229801677 | 136.439489047726 | 42.9562155732476 | 43.7514545260626 |
| chr10 | 91908019 | 91908066 | - | 14.0554480762271 | 10.2329616785794 | 13.1255103140479 | 12.8053037637256 |
| chr10 | 91908096 | 91908119 | - | 3.01188173062009 | 9.09596593651506 | 4.77291284147196 | 7.46976052883995 |
| chr10 | 91908149 | 91908163 | - | 2.00792115374673 | 1.13699574206438 | 1.19322821036799 | 4.26843458790854 |
| chr10 | 92029223 | 92029231 | + | 6.02376346124019 | 4.54798296825753 | 9.54582568294392 | 3.20132594093141 |
| chr10 | 92029458 | 92029462 | + | 3.01188173062009 | 1.13699574206438 | 2.38645642073598 | 1.06710864697714 |
| chr10 | 92029934 | 92029980 | + | 6.02376346124019 | 10.2329616785794 | 2.38645642073598 | 1.06710864697714 |
| chr10 | 92030254 | 92030296 | + | 3.01188173062009 | 3.41098722619315 | 2.38645642073598 | 1.06710864697714 |
| chr10 | 92049624 | 92049629 | - | 4.01584230749346 | 26.1509020674808 | 5.96614105183995 | 4.26843458790854 |
| chr10 | 92351209 | 92351258 | + | 9.03564519186028 | 7.95897019445067 | 9.54582568294392 | 1.06710864697714 |
| chr10 | 92351653 | 92351674 | + | 10.0396057687336 | 12.5069531627082 | 2.38645642073598 | 2.13421729395427 |
| chr10 | 92353583 | 92353642 | + | 12.0475269224804 | 9.09596593651506 | 1.19322821036799 | 1.06710864697714 |
| chr10 | 92452280 | 92452294 | - | 7.02772403811355 | 4.54798296825753 | 9.54582568294392 | 9.60397782279422 |
| chr10 | 92452316 | 92452318 | - | 5.01980288436682 | 5.68497871032191 | 4.77291284147196 | 7.46976052883995 |
| chr10 | 92452640 | 92452644 | - | 5.01980288436682 | 4.54798296825753 | 4.77291284147196 | 3.20132594093141 |
| chr10 | 92452672 | 92452696 | - | 21.0831721143407 | 17.0549361309657 | 8.35259747257593 | 12.8053037637256 |
| chr10 | 92452741 | 92452744 | - | 3.01188173062009 | 1.13699574206438 | 3.57968463110397 | 1.06710864697714 |
| chr10 | 92452844 | 92452848 | - | 3.01188173062009 | 5.68497871032191 | 3.57968463110397 | 3.20132594093141 |
| chr10 | 92452946 | 92452967 | - | 6.02376346124019 | 2.27399148412876 | 2.38645642073598 | 3.20132594093141 |
| chr10 | 92453310 | 92453321 | - | 3.01188173062009 | 6.82197445238629 | 4.77291284147196 | 1.06710864697714 |
| chr10 | 92453901 | 92453920 | - | 4.01584230749346 | 2.27399148412876 | 1.19322821036799 | 2.13421729395427 |
| chr10 | 92694683 | 92694806 | + | 1.00396057687336 | 1.13699574206438 | 39.3765309421437 | 35.2145853502455 |
| chr10 | 92694836 | 92694996 | + | 4.01584230749346 | 1.13699574206438 | 76.3666054635513 | 41.6172372321083 |
| chr10 | 92695096 | 92695132 | + | 1.00396057687336 | 1.13699574206438 | 20.2848795762558 | 8.53686917581709 |
| chr10 | 92695167 | 92695235 | + | 5.01980288436682 | 2.27399148412876 | 58.4681823080315 | 39.483019938154 |
| chr10 | 92695394 | 92695412 | + | 7.02772403811355 | 2.27399148412876 | 5.96614105183995 | 13.8724124107028 |
| chr10 | 93059174 | 93059176 | + | 1.00396057687336 | 5.68497871032191 | 4.77291284147196 | 6.40265188186281 |
| chr10 | 93528605 | 93528615 | + | 44.174265382428 | 28.4248935516096 | 8.35259747257593 | 8.53686917581709 |
| chr10 | 93669811 | 93669841 | - | 2.00792115374673 | 1.13699574206438 | 5.96614105183995 | 1.06710864697714 |
| chr10 | 94333286 | 94333319 | - | 27.1069355755808 | 40.9318467143178 | 4.77291284147196 | 32.0132594093141 |
| chr10 | 94333389 | 94333435 | - | 12.0475269224804 | 12.5069531627082 | 7.15936926220794 | 7.46976052883995 |
| chr10 | 94333461 | 94333503 | - | 22.087132691214 | 23.876910583352 | 7.15936926220794 | 19.2079556455884 |
| chr10 | 94334116 | 94334137 | - | 1.00396057687336 | 2.27399148412876 | 1.19322821036799 | 2.13421729395427 |
| chr10 | 94533658 | 94533661 | + | 1.00396057687336 | 1.13699574206438 | 1.19322821036799 | 1.06710864697714 |
| chr10 | 94601998 | 94602010 | + | 26.1029749987075 | 18.1919318730301 | 1.19322821036799 | 1.06710864697714 |
| chr10 | 95237594 | 95238114 | - | 116.45942691731 | 101.19262104373 | 509.508445827131 | 543.158301311362 |
| chr10 | 95605964 | 95606006 | - | 8.03168461498692 | 7.95897019445067 | 1.19322821036799 | 1.06710864697714 |
| chr10 | 95606062 | 95606186 | - | 22.087132691214 | 29.5618892936739 | 29.8307052591997 | 41.6172372321083 |
| chr10 | 95606213 | 95606310 | - | 33.130699036821 | 32.9728765198671 | 36.9900745214077 | 32.0132594093141 |
| chr10 | 95663436 | 95663438 | - | 2.00792115374673 | 2.27399148412876 | 3.57968463110397 | 1.06710864697714 |
| chr10 | 95663901 | 95663952 | - | 8.03168461498692 | 5.68497871032191 | 4.77291284147196 | 3.20132594093141 |
| chr10 | 96520691 | 96520761 | - | 18.0712903837206 | 25.0139063254164 | 9.54582568294392 | 6.40265188186281 |
| chr10 | 96520929 | 96520931 | - | 2.00792115374673 | 1.13699574206438 | 5.96614105183995 | 3.20132594093141 |
| chr10 | 96521015 | 96521050 | - | 8.03168461498692 | 3.41098722619315 | 2.38645642073598 | 1.06710864697714 |
| chr10 | 96521131 | 96521183 | - | 38.1505019211879 | 39.7948509722534 | 25.0577924177278 | 39.483019938154 |
| chr10 | 96521213 | 96521313 | - | 13.0514874993537 | 14.780944646837 | 16.7051949451519 | 17.0737383516342 |
| chr10 | 96521345 | 96521356 | - | 9.03564519186028 | 14.780944646837 | 4.77291284147196 | 10.6710864697714 |
| chr10 | 96521473 | 96521532 | - | 5.01980288436682 | 9.09596593651506 | 1.19322821036799 | 3.20132594093141 |
| chr10 | 96521691 | 96521717 | - | 6.02376346124019 | 3.41098722619315 | 8.35259747257593 | 17.0737383516342 |
| chr10 | 96521749 | 96522054 | - | 281.108961524542 | 278.563956805774 | 187.336829027774 | 165.401840281456 |
| chr10 | 96522069 | 96522186 | - | 19.0752509605939 | 21.6029190992233 | 38.1833027317757 | 25.6106075274513 |
| chr10 | 96522212 | 96522264 | - | 15.0594086531005 | 11.3699574206438 | 25.0577924177278 | 33.0803680562912 |
| chr10 | 96956191 | 96956217 | + | 6.02376346124019 | 4.54798296825753 | 3.57968463110397 | 3.20132594093141 |
| chr10 | 96957147 | 96957161 | + | 17.0673298068472 | 10.2329616785794 | 7.15936926220794 | 2.13421729395427 |
| chr10 | 96958431 | 96958440 | + | 6.02376346124019 | 4.54798296825753 | 1.19322821036799 | 1.06710864697714 |
| chr10 | 96958501 | 96958529 | + | 1.00396057687336 | 4.54798296825753 | 1.19322821036799 | 1.06710864697714 |
| chr10 | 96959066 | 96959108 | + | 1.00396057687336 | 2.27399148412876 | 1.19322821036799 | 3.20132594093141 |
| chr10 | 96959603 | 96959610 | + | 1.00396057687336 | 1.13699574206438 | 1.19322821036799 | 1.06710864697714 |
| chr10 | 96961707 | 96961730 | + | 13.0514874993537 | 18.1919318730301 | 13.1255103140479 | 5.33554323488568 |
| chr10 | 96964340 | 96964404 | + | 1.00396057687336 | 4.54798296825753 | 2.38645642073598 | 3.20132594093141 |
| chr10 | 96990840 | 96990845 | + | 1.00396057687336 | 1.13699574206438 | 3.57968463110397 | 2.13421729395427 |
| chr10 | 96990880 | 96990897 | + | 7.02772403811355 | 6.82197445238629 | 16.7051949451519 | 1.06710864697714 |
| chr10 | 96993925 | 96993933 | + | 2.00792115374673 | 1.13699574206438 | 1.19322821036799 | 1.06710864697714 |
| chr10 | 96994684 | 96994713 | + | 11.043566345607 | 18.1919318730301 | 7.15936926220794 | 9.60397782279422 |
| chr10 | 97222357 | 97222401 | - | 4.01584230749346 | 10.2329616785794 | 13.1255103140479 | 16.006629704657 |
| chr10 | 97223709 | 97223716 | - | 8.03168461498692 | 1.13699574206438 | 17.8984231555198 | 16.006629704657 |
| chr10 | 97223770 | 97223793 | - | 1.00396057687336 | 2.27399148412876 | 2.38645642073598 | 8.53686917581709 |
| chr10 | 97223954 | 97224020 | - | 13.0514874993537 | 9.09596593651506 | 39.3765309421437 | 21.3421729395427 |
| chr10 | 97224053 | 97224055 | - | 2.00792115374673 | 1.13699574206438 | 1.19322821036799 | 1.06710864697714 |
| chr10 | 97321739 | 97321758 | + | 1.00396057687336 | 3.41098722619315 | 2.38645642073598 | 1.06710864697714 |
| chr10 | 97332546 | 97332725 | - | 718.835773041329 | 689.019419691016 | 278.022173015742 | 283.850900095918 |
| chr10 | 97332746 | 97332770 | - | 19.0752509605939 | 14.780944646837 | 9.54582568294392 | 2.13421729395427 |
| chr10 | 97332807 | 97332831 | - | 83.3287278804893 | 48.8908169087684 | 23.8645642073598 | 42.6843458790854 |
| chr10 | 97332857 | 97332894 | - | 14.0554480762271 | 3.41098722619315 | 1.19322821036799 | 5.33554323488568 |
| chr10 | 97333161 | 97333167 | - | 6.02376346124019 | 1.13699574206438 | 4.77291284147196 | 2.13421729395427 |
| chr10 | 97333248 | 97333286 | - | 19.0752509605939 | 7.95897019445067 | 11.9322821036799 | 7.46976052883995 |
| chr10 | 97333304 | 97333312 | - | 2.00792115374673 | 6.82197445238629 | 1.19322821036799 | 3.20132594093141 |
| chr10 | 97333353 | 97333389 | - | 39.1544624980612 | 28.4248935516096 | 14.3187385244159 | 21.3421729395427 |
| chr10 | 97333487 | 97333517 | - | 36.1425807674411 | 15.9179403889013 | 15.5119667347839 | 11.7381951167485 |
| chr10 | 97333596 | 97333609 | - | 2.00792115374673 | 4.54798296825753 | 1.19322821036799 | 1.06710864697714 |
| chr10 | 97333698 | 97333705 | - | 3.01188173062009 | 2.27399148412876 | 3.57968463110397 | 1.06710864697714 |
| chr10 | 97333739 | 97333752 | - | 12.0475269224804 | 18.1919318730301 | 13.1255103140479 | 10.6710864697714 |
| chr10 | 97333801 | 97333811 | - | 4.01584230749346 | 1.13699574206438 | 3.57968463110397 | 1.06710864697714 |
| chr10 | 97356747 | 97356755 | - | 7.02772403811355 | 11.3699574206438 | 7.15936926220794 | 5.33554323488568 |
| chr10 | 97436309 | 97436320 | - | 1.00396057687336 | 11.3699574206438 | 2.38645642073598 | 7.46976052883995 |
| chr10 | 97436355 | 97436362 | - | 2.00792115374673 | 4.54798296825753 | 2.38645642073598 | 4.26843458790854 |
| chr10 | 97436412 | 97436425 | - | 365.441649981905 | 369.523616170924 | 139.607700613055 | 173.938709457273 |
| chr10 | 97456944 | 97456976 | + | 6.02376346124019 | 7.95897019445067 | 5.96614105183995 | 6.40265188186281 |
| chr10 | 97456989 | 97457033 | + | 21.0831721143407 | 12.5069531627082 | 3.57968463110397 | 4.26843458790854 |
| chr10 | 97457049 | 97457086 | + | 11.043566345607 | 9.09596593651506 | 9.54582568294392 | 5.33554323488568 |
| chr10 | 97457113 | 97457145 | + | 36.1425807674411 | 30.6988850357383 | 13.1255103140479 | 25.6106075274513 |
| chr10 | 97457167 | 97457278 | + | 78.3089249961224 | 70.4937360079917 | 19.0916513658878 | 38.4159112911769 |
| chr10 | 97457299 | 97457306 | + | 4.01584230749346 | 4.54798296825753 | 1.19322821036799 | 2.13421729395427 |
| chr10 | 97457328 | 97457349 | + | 7.02772403811355 | 17.0549361309657 | 3.57968463110397 | 4.26843458790854 |
| chr10 | 97458373 | 97458582 | - | 329.299069214464 | 261.509020674808 | 248.191467756542 | 326.535245975004 |
| chr10 | 97458604 | 97458655 | - | 18.0712903837206 | 12.5069531627082 | 10.7390538933119 | 10.6710864697714 |
| chr10 | 97673945 | 97673948 | + | 1.00396057687336 | 3.41098722619315 | 1.19322821036799 | 6.40265188186281 |
| chr10 | 97674025 | 97674026 | + | 1.00396057687336 | 2.27399148412876 | 2.38645642073598 | 2.13421729395427 |
| chr10 | 97675945 | 97675965 | + | 2.00792115374673 | 3.41098722619315 | 4.77291284147196 | 2.13421729395427 |
| chr10 | 97676020 | 97676021 | + | 1.00396057687336 | 4.54798296825753 | 1.19322821036799 | 1.06710864697714 |
| chr10 | 97677445 | 97677477 | - | 67.2653586505154 | 210.344212281911 | 3.57968463110397 | 4.26843458790854 |
| chr10 | 97677577 | 97677614 | - | 114.451505763564 | 216.029190992233 | 7.15936926220794 | 6.40265188186281 |
| chr10 | 97677722 | 97677865 | - | 33.130699036821 | 68.2197445238629 | 1.19322821036799 | 1.06710864697714 |
| chr10 | 97714432 | 97714437 | + | 10.0396057687336 | 5.68497871032191 | 2.38645642073598 | 1.06710864697714 |
| chr10 | 97718062 | 97718085 | + | 43.1703048055547 | 28.4248935516096 | 8.35259747257593 | 12.8053037637256 |
| chr10 | 97760736 | 97760764 | + | 4.01584230749346 | 1.13699574206438 | 1.19322821036799 | 2.13421729395427 |
| chr10 | 97760798 | 97760874 | + | 11.043566345607 | 7.95897019445067 | 3.57968463110397 | 2.13421729395427 |
| chr10 | 98244779 | 98244783 | + | 6.02376346124019 | 18.1919318730301 | 1.19322821036799 | 6.40265188186281 |
| chr10 | 98244867 | 98244880 | + | 2.00792115374673 | 20.4659233571589 | 5.96614105183995 | 5.33554323488568 |
| chr10 | 98429177 | 98429180 | - | 4.01584230749346 | 5.68497871032191 | 4.77291284147196 | 3.20132594093141 |
| chr10 | 99396900 | 99396952 | - | 4.01584230749346 | 12.5069531627082 | 3.57968463110397 | 4.26843458790854 |
| chr10 | 99397002 | 99397009 | - | 3.01188173062009 | 5.68497871032191 | 1.19322821036799 | 4.26843458790854 |
| chr10 | 99397031 | 99397048 | - | 2.00792115374673 | 4.54798296825753 | 1.19322821036799 | 6.40265188186281 |
| chr10 | 99397076 | 99397114 | - | 1.00396057687336 | 5.68497871032191 | 3.57968463110397 | 5.33554323488568 |
| chr10 | 99397139 | 99397170 | - | 8.03168461498692 | 6.82197445238629 | 2.38645642073598 | 8.53686917581709 |
| chr10 | 99397218 | 99397255 | - | 4.01584230749346 | 10.2329616785794 | 3.57968463110397 | 9.60397782279422 |
| chr10 | 99397333 | 99397374 | - | 5.01980288436682 | 14.780944646837 | 4.77291284147196 | 1.06710864697714 |
| chr10 | 99610549 | 99610572 | - | 37.1465413443145 | 48.8908169087684 | 4.77291284147196 | 8.53686917581709 |
| chr10 | 99610738 | 99610747 | - | 2.00792115374673 | 2.27399148412876 | 1.19322821036799 | 1.06710864697714 |
| chr10 | 99705252 | 99705281 | + | 5.01980288436682 | 1.13699574206438 | 2.38645642073598 | 10.6710864697714 |
| chr10 | 99705446 | 99705485 | + | 6.02376346124019 | 2.27399148412876 | 11.9322821036799 | 7.46976052883995 |
| chr10 | 99705598 | 99705663 | + | 5.01980288436682 | 12.5069531627082 | 10.7390538933119 | 8.53686917581709 |
| chr10 | 99712115 | 99712120 | - | 11.043566345607 | 12.5069531627082 | 4.77291284147196 | 14.9395210576799 |
| chr10 | 99713550 | 99713585 | - | 21.0831721143407 | 14.780944646837 | 32.2171616799357 | 19.2079556455884 |
| chr10 | 99713605 | 99713670 | - | 66.2613980736421 | 71.6307317500561 | 72.7869208324474 | 50.1541064079254 |
| chr10 | 99713690 | 99713712 | - | 14.0554480762271 | 13.6439489047726 | 8.35259747257593 | 10.6710864697714 |
| chr10 | 99713999 | 99714082 | - | 27.1069355755808 | 15.9179403889013 | 27.4442488384638 | 18.1408469986113 |
| chr10 | 99714406 | 99714442 | - | 2.00792115374673 | 3.41098722619315 | 9.54582568294392 | 3.20132594093141 |
| chr10 | 99755993 | 99756012 | + | 5.01980288436682 | 3.41098722619315 | 1.19322821036799 | 2.13421729395427 |
| chr10 | 99875712 | 99875730 | - | 10.0396057687336 | 9.09596593651506 | 2.38645642073598 | 7.46976052883995 |
| chr10 | 100150117 | 100150249 | - | 15.0594086531005 | 43.2058381984465 | 8.35259747257593 | 17.0737383516342 |
| chr10 | 100150284 | 100150324 | - | 1.00396057687336 | 6.82197445238629 | 2.38645642073598 | 2.13421729395427 |
| chr10 | 100188651 | 100188652 | - | 2.00792115374673 | 2.27399148412876 | 1.19322821036799 | 1.06710864697714 |
| chr10 | 100188695 | 100188716 | - | 1.00396057687336 | 1.13699574206438 | 1.19322821036799 | 6.40265188186281 |
| chr10 | 100188939 | 100188948 | - | 11.043566345607 | 5.68497871032191 | 8.35259747257593 | 3.20132594093141 |
| chr10 | 100188987 | 100188996 | - | 15.0594086531005 | 31.8358807778027 | 10.7390538933119 | 2.13421729395427 |
| chr10 | 100189052 | 100189084 | - | 5.01980288436682 | 11.3699574206438 | 9.54582568294392 | 3.20132594093141 |
| chr10 | 100189496 | 100189497 | - | 1.00396057687336 | 1.13699574206438 | 1.19322821036799 | 1.06710864697714 |
| chr10 | 100233189 | 100233243 | - | 32.1267384599477 | 30.6988850357383 | 17.8984231555198 | 42.6843458790854 |
| chr10 | 100274735 | 100274769 | - | 12.0475269224804 | 25.0139063254164 | 4.77291284147196 | 12.8053037637256 |
| chr10 | 100274852 | 100274886 | - | 9.03564519186028 | 17.0549361309657 | 7.15936926220794 | 13.8724124107028 |
| chr10 | 100275127 | 100275155 | - | 17.0673298068472 | 15.9179403889013 | 4.77291284147196 | 8.53686917581709 |
| chr10 | 100360887 | 100360957 | + | 242.958459603354 | 201.248246345396 | 181.370687975934 | 50.1541064079254 |
| chr10 | 100361031 | 100361071 | + | 93.3683336492229 | 90.9596593651506 | 51.3088130458235 | 13.8724124107028 |
| chr10 | 100361090 | 100361121 | + | 57.2257528817818 | 57.9867828452835 | 85.9124311464952 | 23.476390233497 |
| chr10 | 100361153 | 100361179 | + | 6.02376346124019 | 3.41098722619315 | 8.35259747257593 | 1.06710864697714 |
| chr10 | 100361181 | 100361236 | + | 30.1188173062009 | 21.6029190992233 | 34.6036181006717 | 14.9395210576799 |
| chr10 | 100361272 | 100361292 | + | 32.1267384599477 | 28.4248935516096 | 32.2171616799357 | 11.7381951167485 |
| chr10 | 100361366 | 100361392 | + | 40.1584230749346 | 30.6988850357383 | 41.7629873628796 | 7.46976052883995 |
| chr10 | 100361448 | 100361474 | + | 33.130699036821 | 27.2878978095452 | 39.3765309421437 | 16.006629704657 |
| chr10 | 100361510 | 100361568 | + | 68.2693192273888 | 67.0827487817985 | 94.2650286190712 | 36.2816939972226 |
| chr10 | 100361592 | 100361700 | + | 233.922814411494 | 143.261463500112 | 184.950372607038 | 103.509538756782 |
| chr10 | 100361761 | 100361872 | + | 139.550520185398 | 142.124467758048 | 305.466421854205 | 360.682722678272 |
| chr10 | 100361893 | 100362000 | + | 166.657455760979 | 152.357429436627 | 292.340911540157 | 102.442430109805 |
| chr10 | 100362040 | 100362082 | + | 86.3406096111093 | 70.4937360079917 | 118.129592826431 | 45.8856718200168 |
| chr10 | 100362116 | 100362231 | + | 84.3326884573626 | 62.534765813541 | 91.8785721983352 | 68.2949534065367 |
| chr10 | 100362327 | 100362424 | + | 137.542599031651 | 117.110561432631 | 257.737293439486 | 39.483019938154 |
| chr10 | 100362465 | 100362517 | + | 24.0950538449607 | 19.3289276150945 | 36.9900745214077 | 13.8724124107028 |
| chr10 | 100362554 | 100362574 | + | 8.03168461498692 | 12.5069531627082 | 23.8645642073598 | 13.8724124107028 |
| chr10 | 100362599 | 100362724 | + | 45.1782259593014 | 25.0139063254164 | 40.5697591525116 | 24.5434988804741 |
| chr10 | 100362753 | 100362780 | + | 11.043566345607 | 6.82197445238629 | 26.2510206280958 | 20.2750642925656 |
| chr10 | 100362818 | 100362968 | + | 160.633692299738 | 106.877599754052 | 221.940447128446 | 217.690163983336 |
| chr10 | 100363039 | 100363106 | + | 59.2336740355285 | 50.0278126508328 | 132.448331350847 | 72.5633879944452 |
| chr10 | 100363137 | 100363293 | + | 125.495072109171 | 137.57648478979 | 287.567998698685 | 187.811121867976 |
| chr10 | 100363344 | 100363496 | + | 573.261489394691 | 590.100790131414 | 1029.75594554757 | 579.439995308585 |
| chr10 | 100363513 | 100363588 | + | 74.293082688629 | 64.8087572976698 | 108.583767143487 | 40.5501285851312 |
| chr10 | 100363659 | 100363672 | + | 14.0554480762271 | 4.54798296825753 | 15.5119667347839 | 22.4092815865198 |
| chr10 | 100363697 | 100363708 | + | 10.0396057687336 | 2.27399148412876 | 3.57968463110397 | 6.40265188186281 |
| chr10 | 100363731 | 100363742 | + | 1.00396057687336 | 1.13699574206438 | 2.38645642073598 | 3.20132594093141 |
| chr10 | 100363834 | 100363861 | + | 11.043566345607 | 1.13699574206438 | 11.9322821036799 | 13.8724124107028 |
| chr10 | 100363971 | 100364001 | + | 4.01584230749346 | 1.13699574206438 | 16.7051949451519 | 9.60397782279422 |
| chr10 | 100364034 | 100364108 | + | 57.2257528817818 | 52.3018041349616 | 76.3666054635513 | 90.7042349930565 |
| chr10 | 100364160 | 100364287 | + | 8.03168461498692 | 3.41098722619315 | 9.54582568294392 | 6.40265188186281 |
| chr10 | 100364358 | 100364381 | + | 22.087132691214 | 28.4248935516096 | 8.35259747257593 | 7.46976052883995 |
| chr10 | 100364411 | 100364518 | + | 15.0594086531005 | 11.3699574206438 | 8.35259747257593 | 21.3421729395427 |
| chr10 | 100364542 | 100364637 | + | 52.205949997415 | 77.315710460378 | 63.2410951495034 | 100.308212815851 |
| chr10 | 100364682 | 100364709 | + | 22.087132691214 | 14.780944646837 | 15.5119667347839 | 13.8724124107028 |
| chr10 | 100364777 | 100364807 | + | 4.01584230749346 | 14.780944646837 | 5.96614105183995 | 2.13421729395427 |
| chr10 | 100373491 | 100373723 | + | 385.520861519372 | 687.882423948951 | 533.373010034491 | 434.313219319694 |
| chr10 | 100373804 | 100373900 | + | 120.475269224804 | 119.38455291676 | 27.4442488384638 | 34.1474767032683 |
| chr10 | 100374005 | 100374013 | + | 7.02772403811355 | 6.82197445238629 | 8.35259747257593 | 11.7381951167485 |
| chr10 | 100383029 | 100383079 | + | 7.02772403811355 | 15.9179403889013 | 26.2510206280958 | 43.7514545260626 |
| chr10 | 100523800 | 100523860 | - | 53.2099105742883 | 60.2607743294122 | 16.7051949451519 | 39.483019938154 |
| chr10 | 100549719 | 100549737 | + | 51.2019894205416 | 39.7948509722534 | 3.57968463110397 | 3.20132594093141 |
| chr10 | 100551131 | 100551144 | + | 4.01584230749346 | 4.54798296825753 | 1.19322821036799 | 1.06710864697714 |
| chr10 | 100553799 | 100553850 | + | 4.01584230749346 | 10.2329616785794 | 1.19322821036799 | 1.06710864697714 |
| chr10 | 100962567 | 100962585 | + | 2.00792115374673 | 3.41098722619315 | 1.19322821036799 | 1.06710864697714 |
| chr10 | 100963043 | 100963065 | + | 4.01584230749346 | 7.95897019445067 | 7.15936926220794 | 4.26843458790854 |
| chr10 | 100963462 | 100963485 | + | 1.00396057687336 | 1.13699574206438 | 2.38645642073598 | 4.26843458790854 |
| chr10 | 100963579 | 100963602 | + | 15.0594086531005 | 12.5069531627082 | 7.15936926220794 | 3.20132594093141 |
| chr10 | 100963757 | 100963771 | + | 32.1267384599477 | 45.4798296825753 | 25.0577924177278 | 12.8053037637256 |
| chr10 | 100981396 | 100981401 | - | 7.02772403811355 | 13.6439489047726 | 1.19322821036799 | 1.06710864697714 |
| chr10 | 100986334 | 100986431 | - | 111.439624032943 | 97.7816338175368 | 60.8546387287675 | 88.5700176991023 |
| chr10 | 100986473 | 100986522 | - | 69.2732798042621 | 67.0827487817985 | 44.1494437836156 | 51.2212150549025 |
| chr10 | 100986527 | 100986561 | - | 40.1584230749346 | 51.1648083928972 | 36.9900745214077 | 33.0803680562912 |
| chr10 | 100986591 | 100986630 | - | 9.03564519186028 | 3.41098722619315 | 3.57968463110397 | 7.46976052883995 |
| chr10 | 100986664 | 100986706 | - | 9.03564519186028 | 5.68497871032191 | 4.77291284147196 | 4.26843458790854 |
| chr10 | 100993289 | 100993301 | + | 5.01980288436682 | 2.27399148412876 | 1.19322821036799 | 1.06710864697714 |
| chr10 | 100993575 | 100993576 | + | 2.00792115374673 | 1.13699574206438 | 2.38645642073598 | 1.06710864697714 |
| chr10 | 100993776 | 100993792 | + | 2.00792115374673 | 2.27399148412876 | 2.38645642073598 | 2.13421729395427 |
| chr10 | 100993887 | 100993921 | + | 5.01980288436682 | 4.54798296825753 | 2.38645642073598 | 1.06710864697714 |
| chr10 | 100994063 | 100994075 | + | 17.0673298068472 | 4.54798296825753 | 1.19322821036799 | 4.26843458790854 |
| chr10 | 100994231 | 100994319 | + | 81.3208067267425 | 56.8497871032191 | 23.8645642073598 | 49.0869977609482 |
| chr10 | 101007655 | 101007686 | + | 5.01980288436682 | 7.95897019445067 | 1.19322821036799 | 8.53686917581709 |
| chr10 | 101007795 | 101007801 | + | 7.02772403811355 | 9.09596593651506 | 3.57968463110397 | 6.40265188186281 |
| chr10 | 101554053 | 101554073 | + | 3.01188173062009 | 5.68497871032191 | 1.19322821036799 | 3.20132594093141 |
| chr10 | 101578903 | 101578918 | - | 5.01980288436682 | 14.780944646837 | 5.96614105183995 | 4.26843458790854 |
| chr10 | 101611223 | 101611262 | - | 4.01584230749346 | 10.2329616785794 | 4.77291284147196 | 1.06710864697714 |
| chr10 | 101781393 | 101781423 | - | 13.0514874993537 | 5.68497871032191 | 10.7390538933119 | 12.8053037637256 |
| chr10 | 101781493 | 101781526 | - | 58.2297134586551 | 38.657855230189 | 14.3187385244159 | 62.959410171651 |
| chr10 | 101784470 | 101784474 | - | 12.0475269224804 | 30.6988850357383 | 22.6713359969918 | 12.8053037637256 |
| chr10 | 101784495 | 101784516 | - | 1.00396057687336 | 1.13699574206438 | 3.57968463110397 | 1.06710864697714 |
| chr10 | 101784548 | 101784588 | - | 2.00792115374673 | 3.41098722619315 | 3.57968463110397 | 6.40265188186281 |
| chr10 | 101785068 | 101785072 | - | 1.00396057687336 | 1.13699574206438 | 2.38645642073598 | 1.06710864697714 |
| chr10 | 101785554 | 101785556 | - | 1.00396057687336 | 3.41098722619315 | 1.19322821036799 | 1.06710864697714 |
| chr10 | 102107653 | 102107656 | - | 10.0396057687336 | 6.82197445238629 | 3.57968463110397 | 12.8053037637256 |
| chr10 | 102107737 | 102107748 | - | 3.01188173062009 | 4.54798296825753 | 11.9322821036799 | 6.40265188186281 |
| chr10 | 102150067 | 102150071 | + | 13.0514874993537 | 7.95897019445067 | 1.19322821036799 | 9.60397782279422 |
| chr10 | 102150105 | 102150151 | + | 18.0712903837206 | 20.4659233571589 | 8.35259747257593 | 17.0737383516342 |
| chr10 | 102162335 | 102162391 | + | 19.0752509605939 | 12.5069531627082 | 9.54582568294392 | 16.006629704657 |
| chr10 | 102162404 | 102162441 | + | 3.01188173062009 | 4.54798296825753 | 5.96614105183995 | 11.7381951167485 |
| chr10 | 102162469 | 102162492 | + | 6.02376346124019 | 29.5618892936739 | 3.57968463110397 | 8.53686917581709 |
| chr10 | 102162649 | 102162677 | + | 4.01584230749346 | 2.27399148412876 | 1.19322821036799 | 3.20132594093141 |
| chr10 | 102162764 | 102162768 | + | 16.0633692299738 | 28.4248935516096 | 3.57968463110397 | 9.60397782279422 |
| chr10 | 102162824 | 102162947 | + | 45.1782259593014 | 48.8908169087684 | 41.7629873628796 | 54.4225409958339 |
| chr10 | 102163283 | 102163305 | + | 1.00396057687336 | 1.13699574206438 | 2.38645642073598 | 3.20132594093141 |
| chr10 | 102163597 | 102163606 | + | 38.1505019211879 | 36.3838637460602 | 8.35259747257593 | 40.5501285851312 |
| chr10 | 102163627 | 102163676 | + | 16.0633692299738 | 17.0549361309657 | 8.35259747257593 | 17.0737383516342 |
| chr10 | 102163799 | 102163825 | + | 23.0910932680874 | 38.657855230189 | 7.15936926220794 | 3.20132594093141 |
| chr10 | 102382348 | 102382358 | + | 2.00792115374673 | 2.27399148412876 | 1.19322821036799 | 8.53686917581709 |
| chr10 | 102382800 | 102382859 | + | 38.1505019211879 | 31.8358807778027 | 19.0916513658878 | 25.6106075274513 |
| chr10 | 102402500 | 102402506 | + | 21.0831721143407 | 44.3428339405109 | 4.77291284147196 | 6.40265188186281 |
| chr10 | 102423082 | 102423108 | + | 1.00396057687336 | 6.82197445238629 | 2.38645642073598 | 10.6710864697714 |
| chr10 | 102449854 | 102449876 | + | 44.174265382428 | 20.4659233571589 | 51.3088130458235 | 53.3554323488568 |
| chr10 | 102449953 | 102449959 | + | 7.02772403811355 | 21.6029190992233 | 2.38645642073598 | 5.33554323488568 |
| chr10 | 102455616 | 102455622 | + | 15.0594086531005 | 23.876910583352 | 41.7629873628796 | 36.2816939972226 |
| chr10 | 102455754 | 102455758 | + | 3.01188173062009 | 6.82197445238629 | 7.15936926220794 | 9.60397782279422 |
| chr10 | 102456202 | 102456208 | + | 3.01188173062009 | 1.13699574206438 | 2.38645642073598 | 4.26843458790854 |
| chr10 | 102479246 | 102479304 | - | 45.1782259593014 | 68.2197445238629 | 28.6374770488317 | 33.0803680562912 |
| chr10 | 102479348 | 102479361 | - | 12.0475269224804 | 7.95897019445067 | 2.38645642073598 | 2.13421729395427 |
| chr10 | 102479761 | 102479776 | - | 4.01584230749346 | 3.41098722619315 | 3.57968463110397 | 2.13421729395427 |
| chr10 | 102480233 | 102480247 | - | 4.01584230749346 | 1.13699574206438 | 4.77291284147196 | 2.13421729395427 |
| chr10 | 102480361 | 102480422 | - | 9.03564519186028 | 13.6439489047726 | 9.54582568294392 | 6.40265188186281 |
| chr10 | 102480553 | 102480583 | - | 10.0396057687336 | 6.82197445238629 | 8.35259747257593 | 10.6710864697714 |
| chr10 | 102480654 | 102480696 | - | 6.02376346124019 | 7.95897019445067 | 2.38645642073598 | 5.33554323488568 |
| chr10 | 102480766 | 102480803 | - | 2.00792115374673 | 4.54798296825753 | 13.1255103140479 | 8.53686917581709 |
| chr10 | 102480855 | 102480862 | - | 3.01188173062009 | 7.95897019445067 | 5.96614105183995 | 6.40265188186281 |
| chr10 | 102633479 | 102633480 | + | 2.00792115374673 | 2.27399148412876 | 2.38645642073598 | 5.33554323488568 |
| chr10 | 102657566 | 102657633 | + | 1.00396057687336 | 1.13699574206438 | 2.38645642073598 | 1.06710864697714 |
| chr10 | 102657746 | 102657847 | + | 23.0910932680874 | 15.9179403889013 | 11.9322821036799 | 11.7381951167485 |
| chr10 | 102657924 | 102658055 | + | 21.0831721143407 | 26.1509020674808 | 3.57968463110397 | 4.26843458790854 |
| chr10 | 102658200 | 102658247 | + | 4.01584230749346 | 4.54798296825753 | 1.19322821036799 | 1.06710864697714 |
| chr10 | 102676686 | 102676737 | - | 53.2099105742883 | 73.9047232341848 | 15.5119667347839 | 10.6710864697714 |
| chr10 | 102676761 | 102676772 | - | 13.0514874993537 | 26.1509020674808 | 10.7390538933119 | 7.46976052883995 |
| chr10 | 102676849 | 102676871 | - | 5.01980288436682 | 3.41098722619315 | 5.96614105183995 | 4.26843458790854 |
| chr10 | 102739058 | 102739076 | + | 7.02772403811355 | 7.95897019445067 | 3.57968463110397 | 17.0737383516342 |
| chr10 | 103088183 | 103088220 | - | 32.1267384599477 | 42.0688424563821 | 9.54582568294392 | 7.46976052883995 |
| chr10 | 103089305 | 103089311 | - | 3.01188173062009 | 1.13699574206438 | 1.19322821036799 | 1.06710864697714 |
| chr10 | 103388427 | 103388449 | + | 8.03168461498692 | 19.3289276150945 | 17.8984231555198 | 13.8724124107028 |
| chr10 | 103388471 | 103388475 | + | 3.01188173062009 | 5.68497871032191 | 2.38645642073598 | 3.20132594093141 |
| chr10 | 103389147 | 103389167 | - | 3.01188173062009 | 1.13699574206438 | 1.19322821036799 | 2.13421729395427 |
| chr10 | 103392190 | 103392231 | - | 15.0594086531005 | 10.2329616785794 | 4.77291284147196 | 6.40265188186281 |
| chr10 | 103445566 | 103445595 | + | 2.00792115374673 | 1.13699574206438 | 3.57968463110397 | 8.53686917581709 |
| chr10 | 103445672 | 103445699 | + | 9.03564519186028 | 9.09596593651506 | 9.54582568294392 | 14.9395210576799 |
| chr10 | 103445821 | 103445840 | + | 1.00396057687336 | 1.13699574206438 | 1.19322821036799 | 5.33554323488568 |
| chr10 | 103445880 | 103445896 | + | 4.01584230749346 | 2.27399148412876 | 9.54582568294392 | 5.33554323488568 |
| chr10 | 103445935 | 103445962 | + | 7.02772403811355 | 5.68497871032191 | 13.1255103140479 | 5.33554323488568 |
| chr10 | 103445982 | 103445992 | + | 2.00792115374673 | 1.13699574206438 | 3.57968463110397 | 1.06710864697714 |
| chr10 | 103446017 | 103446062 | + | 7.02772403811355 | 6.82197445238629 | 3.57968463110397 | 14.9395210576799 |
| chr10 | 103446240 | 103446272 | + | 1.00396057687336 | 4.54798296825753 | 1.19322821036799 | 3.20132594093141 |
| chr10 | 103592408 | 103592448 | + | 16.0633692299738 | 32.9728765198671 | 1.19322821036799 | 1.06710864697714 |
| chr10 | 103597897 | 103597899 | - | 2.00792115374673 | 3.41098722619315 | 1.19322821036799 | 1.06710864697714 |
| chr10 | 103599879 | 103599926 | - | 32.1267384599477 | 30.6988850357383 | 3.57968463110397 | 11.7381951167485 |
| chr10 | 103600850 | 103600924 | - | 104.41189999483 | 68.2197445238629 | 29.8307052591997 | 28.8119334683827 |
| chr10 | 103600953 | 103601032 | - | 23.0910932680874 | 19.3289276150945 | 8.35259747257593 | 12.8053037637256 |
| chr10 | 103601061 | 103601075 | - | 4.01584230749346 | 9.09596593651506 | 1.19322821036799 | 1.06710864697714 |
| chr10 | 103601708 | 103601717 | - | 192.760430759686 | 150.083437952498 | 72.7869208324474 | 34.1474767032683 |
| chr10 | 103882576 | 103882604 | - | 4.01584230749346 | 2.27399148412876 | 7.15936926220794 | 6.40265188186281 |
| chr10 | 104025720 | 104025752 | + | 44.174265382428 | 32.9728765198671 | 19.0916513658878 | 28.8119334683827 |
| chr10 | 104025801 | 104025838 | + | 21.0831721143407 | 9.09596593651506 | 15.5119667347839 | 32.0132594093141 |
| chr10 | 104025888 | 104025953 | + | 26.1029749987075 | 23.876910583352 | 16.7051949451519 | 19.2079556455884 |
| chr10 | 104026804 | 104026813 | + | 2.00792115374673 | 1.13699574206438 | 2.38645642073598 | 1.06710864697714 |
| chr10 | 104027124 | 104027128 | + | 4.01584230749346 | 2.27399148412876 | 1.19322821036799 | 1.06710864697714 |
| chr10 | 104027154 | 104027156 | + | 2.00792115374673 | 2.27399148412876 | 1.19322821036799 | 1.06710864697714 |
| chr10 | 104027317 | 104027347 | + | 1.00396057687336 | 2.27399148412876 | 2.38645642073598 | 2.13421729395427 |
| chr10 | 104027474 | 104027475 | + | 2.00792115374673 | 4.54798296825753 | 1.19322821036799 | 1.06710864697714 |
| chr10 | 104027529 | 104027533 | + | 3.01188173062009 | 4.54798296825753 | 1.19322821036799 | 3.20132594093141 |
| chr10 | 104027865 | 104027872 | + | 4.01584230749346 | 4.54798296825753 | 3.57968463110397 | 1.06710864697714 |
| chr10 | 104028856 | 104028858 | + | 4.01584230749346 | 1.13699574206438 | 2.38645642073598 | 1.06710864697714 |
| chr10 | 104028920 | 104028925 | + | 54.2138711511617 | 52.3018041349616 | 5.96614105183995 | 20.2750642925656 |
| chr10 | 104029068 | 104029070 | + | 2.00792115374673 | 6.82197445238629 | 4.77291284147196 | 3.20132594093141 |
| chr10 | 104267398 | 104267440 | + | 99.3920971104631 | 108.014595496116 | 180.177459765566 | 296.656203859644 |
| chr10 | 104313548 | 104313552 | - | 2.00792115374673 | 4.54798296825753 | 2.38645642073598 | 4.26843458790854 |
| chr10 | 104313913 | 104313935 | - | 1.00396057687336 | 1.13699574206438 | 5.96614105183995 | 6.40265188186281 |
| chr10 | 109864843 | 109864882 | - | 11.043566345607 | 9.09596593651506 | 5.96614105183995 | 7.46976052883995 |
| chr10 | 109865125 | 109865139 | - | 2.00792115374673 | 3.41098722619315 | 1.19322821036799 | 4.26843458790854 |
| chr10 | 110134329 | 110134333 | + | 6.02376346124019 | 7.95897019445067 | 17.8984231555198 | 4.26843458790854 |
| chr10 | 110134475 | 110134479 | + | 1.00396057687336 | 1.13699574206438 | 3.57968463110397 | 2.13421729395427 |
| chr10 | 110135160 | 110135171 | + | 5.01980288436682 | 3.41098722619315 | 4.77291284147196 | 7.46976052883995 |
| chr10 | 110135198 | 110135236 | + | 4.01584230749346 | 12.5069531627082 | 7.15936926220794 | 8.53686917581709 |
| chr10 | 110285011 | 110285039 | + | 5.01980288436682 | 6.82197445238629 | 9.54582568294392 | 6.40265188186281 |
| chr10 | 110285116 | 110285134 | + | 2.00792115374673 | 1.13699574206438 | 2.38645642073598 | 1.06710864697714 |
| chr10 | 110285163 | 110285187 | + | 3.01188173062009 | 1.13699574206438 | 11.9322821036799 | 3.20132594093141 |
| chr10 | 110285342 | 110285360 | + | 2.00792115374673 | 2.27399148412876 | 8.35259747257593 | 1.06710864697714 |
| chr10 | 110285476 | 110285515 | + | 8.03168461498692 | 9.09596593651506 | 26.2510206280958 | 12.8053037637256 |
| chr10 | 110285569 | 110285571 | + | 1.00396057687336 | 1.13699574206438 | 4.77291284147196 | 1.06710864697714 |
| chr10 | 110285986 | 110285988 | + | 5.01980288436682 | 3.41098722619315 | 3.57968463110397 | 2.13421729395427 |
| chr10 | 110286073 | 110286096 | + | 13.0514874993537 | 10.2329616785794 | 3.57968463110397 | 13.8724124107028 |
| chr10 | 110286418 | 110286451 | + | 4.01584230749346 | 10.2329616785794 | 4.77291284147196 | 9.60397782279422 |
| chr10 | 110286587 | 110286599 | + | 2.00792115374673 | 11.3699574206438 | 2.38645642073598 | 4.26843458790854 |
| chr10 | 110293100 | 110293150 | - | 8.03168461498692 | 18.1919318730301 | 7.15936926220794 | 6.40265188186281 |
| chr10 | 110293527 | 110293533 | - | 4.01584230749346 | 3.41098722619315 | 10.7390538933119 | 8.53686917581709 |
| chr10 | 110294116 | 110294163 | - | 3.01188173062009 | 1.13699574206438 | 4.77291284147196 | 2.13421729395427 |
| chr10 | 110604294 | 110604331 | + | 12.0475269224804 | 22.7399148412876 | 7.15936926220794 | 10.6710864697714 |
| chr10 | 110604361 | 110604363 | + | 1.00396057687336 | 3.41098722619315 | 2.38645642073598 | 2.13421729395427 |
| chr10 | 110604464 | 110604484 | + | 2.00792115374673 | 1.13699574206438 | 1.19322821036799 | 1.06710864697714 |
| chr10 | 110898025 | 110898090 | + | 31.1227778830743 | 54.5757956190903 | 21.4781077866238 | 36.2816939972226 |
| chr10 | 110898127 | 110898149 | + | 19.0752509605939 | 34.1098722619315 | 13.1255103140479 | 6.40265188186281 |
| chr10 | 110898188 | 110898249 | + | 152.602007684751 | 275.15296957958 | 139.607700613055 | 78.966039876308 |
| chr10 | 110898277 | 110898360 | + | 66.2613980736421 | 115.973565690567 | 46.5359002043516 | 26.6777161744284 |
| chr10 | 110898412 | 110898430 | + | 2.00792115374673 | 11.3699574206438 | 3.57968463110397 | 9.60397782279422 |
| chr10 | 110898462 | 110898475 | + | 379.497098058132 | 577.593836968706 | 79.9462900946553 | 66.1607361125824 |
| chr10 | 110898508 | 110898531 | + | 4.01584230749346 | 12.5069531627082 | 1.19322821036799 | 2.13421729395427 |
| chr10 | 110898949 | 110898952 | - | 5.01980288436682 | 4.54798296825753 | 1.19322821036799 | 1.06710864697714 |
| chr10 | 110900338 | 110900340 | - | 3.01188173062009 | 6.82197445238629 | 1.19322821036799 | 5.33554323488568 |
| chr10 | 111012411 | 111012413 | + | 1.00396057687336 | 1.13699574206438 | 1.19322821036799 | 1.06710864697714 |
| chr10 | 111012538 | 111012556 | + | 7.02772403811355 | 17.0549361309657 | 5.96614105183995 | 9.60397782279422 |
| chr10 | 111013087 | 111013128 | + | 8.03168461498692 | 14.780944646837 | 10.7390538933119 | 5.33554323488568 |
| chr10 | 111013184 | 111013237 | + | 10.0396057687336 | 19.3289276150945 | 4.77291284147196 | 7.46976052883995 |
| chr10 | 111013278 | 111013297 | + | 2.00792115374673 | 11.3699574206438 | 13.1255103140479 | 5.33554323488568 |
| chr10 | 111013324 | 111013336 | + | 89.3524913417294 | 186.467301698559 | 13.1255103140479 | 14.9395210576799 |
| chr10 | 111013379 | 111013414 | + | 12.0475269224804 | 35.2468680039958 | 8.35259747257593 | 6.40265188186281 |
| chr10 | 111013436 | 111013580 | + | 310.22381825387 | 623.073666651281 | 230.293044601022 | 193.146665102862 |
| chr10 | 112151757 | 112151810 | - | 15.0594086531005 | 15.9179403889013 | 4.77291284147196 | 11.7381951167485 |
| chr10 | 112153339 | 112153343 | - | 3.01188173062009 | 6.82197445238629 | 1.19322821036799 | 2.13421729395427 |
| chr10 | 112430457 | 112430474 | - | 5.01980288436682 | 14.780944646837 | 1.19322821036799 | 1.06710864697714 |
| chr10 | 112818398 | 112818401 | + | 14.0554480762271 | 11.3699574206438 | 1.19322821036799 | 3.20132594093141 |
| chr10 | 112827369 | 112827372 | + | 3.01188173062009 | 3.41098722619315 | 2.38645642073598 | 1.06710864697714 |
| chr10 | 112827400 | 112827402 | + | 3.01188173062009 | 2.27399148412876 | 1.19322821036799 | 1.06710864697714 |
| chr10 | 113834752 | 113834758 | - | 3.01188173062009 | 25.0139063254164 | 4.77291284147196 | 1.06710864697714 |
| chr10 | 113834908 | 113834910 | - | 6.02376346124019 | 3.41098722619315 | 2.38645642073598 | 1.06710864697714 |
| chr10 | 113910479 | 113910505 | + | 12.0475269224804 | 11.3699574206438 | 5.96614105183995 | 2.13421729395427 |
| chr10 | 113910562 | 113910563 | + | 1.00396057687336 | 3.41098722619315 | 2.38645642073598 | 1.06710864697714 |
| chr10 | 113911926 | 113911929 | + | 6.02376346124019 | 3.41098722619315 | 2.38645642073598 | 6.40265188186281 |
| chr10 | 113912185 | 113912188 | + | 2.00792115374673 | 2.27399148412876 | 2.38645642073598 | 1.06710864697714 |
| chr10 | 113912534 | 113912536 | + | 6.02376346124019 | 9.09596593651506 | 7.15936926220794 | 1.06710864697714 |
| chr10 | 113912704 | 113912729 | + | 3.01188173062009 | 4.54798296825753 | 5.96614105183995 | 3.20132594093141 |
| chr10 | 113912845 | 113912880 | + | 11.043566345607 | 4.54798296825753 | 8.35259747257593 | 11.7381951167485 |
| chr10 | 113912910 | 113912912 | + | 1.00396057687336 | 1.13699574206438 | 2.38645642073598 | 1.06710864697714 |
| chr10 | 113913139 | 113913141 | + | 17.0673298068472 | 18.1919318730301 | 10.7390538933119 | 6.40265188186281 |
| chr10 | 113913192 | 113913193 | + | 1.00396057687336 | 1.13699574206438 | 1.19322821036799 | 1.06710864697714 |
| chr10 | 113916385 | 113916420 | + | 5.01980288436682 | 9.09596593651506 | 14.3187385244159 | 4.26843458790854 |
| chr10 | 113916455 | 113916459 | + | 7.02772403811355 | 10.2329616785794 | 8.35259747257593 | 3.20132594093141 |
| chr10 | 113916498 | 113916509 | + | 3.01188173062009 | 7.95897019445067 | 4.77291284147196 | 2.13421729395427 |
| chr10 | 113916630 | 113916650 | + | 7.02772403811355 | 10.2329616785794 | 3.57968463110397 | 6.40265188186281 |
| chr10 | 114975561 | 114975584 | + | 1.00396057687336 | 11.3699574206438 | 2.38645642073598 | 3.20132594093141 |
| chr10 | 114975788 | 114975800 | + | 3.01188173062009 | 2.27399148412876 | 1.19322821036799 | 1.06710864697714 |
| chr10 | 114976917 | 114976983 | + | 9.03564519186028 | 6.82197445238629 | 4.77291284147196 | 8.53686917581709 |
| chr10 | 114977442 | 114977448 | + | 5.01980288436682 | 10.2329616785794 | 1.19322821036799 | 6.40265188186281 |
| chr10 | 116883166 | 116883191 | - | 12.0475269224804 | 3.41098722619315 | 9.54582568294392 | 7.46976052883995 |
| chr10 | 116885073 | 116885111 | - | 5.01980288436682 | 7.95897019445067 | 1.19322821036799 | 6.40265188186281 |
| chr10 | 116885142 | 116885234 | - | 7.02772403811355 | 9.09596593651506 | 3.57968463110397 | 3.20132594093141 |
| chr10 | 116885524 | 116885548 | - | 18.0712903837206 | 10.2329616785794 | 3.57968463110397 | 1.06710864697714 |
| chr10 | 116885662 | 116885673 | - | 11.043566345607 | 15.9179403889013 | 10.7390538933119 | 7.46976052883995 |
| chr10 | 116885695 | 116885709 | - | 21.0831721143407 | 10.2329616785794 | 4.77291284147196 | 4.26843458790854 |
| chr10 | 116885979 | 116885994 | - | 3.01188173062009 | 2.27399148412876 | 3.57968463110397 | 2.13421729395427 |
| chr10 | 116886266 | 116886289 | - | 6.02376346124019 | 2.27399148412876 | 2.38645642073598 | 1.06710864697714 |
| chr10 | 116886414 | 116886425 | - | 8.03168461498692 | 7.95897019445067 | 8.35259747257593 | 2.13421729395427 |
| chr10 | 117280635 | 117280638 | - | 8.03168461498692 | 3.41098722619315 | 13.1255103140479 | 27.7448248214055 |
| chr10 | 117282291 | 117282295 | - | 3.01188173062009 | 3.41098722619315 | 1.19322821036799 | 1.06710864697714 |
| chr10 | 117282732 | 117282743 | - | 45.1782259593014 | 62.534765813541 | 41.7629873628796 | 34.1474767032683 |
| chr10 | 117282812 | 117282824 | - | 8.03168461498692 | 11.3699574206438 | 8.35259747257593 | 22.4092815865198 |
| chr10 | 117283232 | 117283238 | - | 2.00792115374673 | 5.68497871032191 | 4.77291284147196 | 6.40265188186281 |
| chr10 | 118006841 | 118006843 | - | 4.01584230749346 | 3.41098722619315 | 1.19322821036799 | 2.13421729395427 |
| chr10 | 118046984 | 118047035 | + | 18.0712903837206 | 39.7948509722534 | 9.54582568294392 | 5.33554323488568 |
| chr10 | 118681548 | 118681563 | - | 10.0396057687336 | 28.4248935516096 | 7.15936926220794 | 1.06710864697714 |
| chr10 | 118681669 | 118681676 | - | 56.2217923049084 | 38.657855230189 | 14.3187385244159 | 13.8724124107028 |
| chr10 | 118681694 | 118681848 | - | 191.756470182813 | 169.412365567593 | 58.4681823080315 | 58.6909755837425 |
| chr10 | 118681961 | 118682001 | - | 28.1108961524542 | 27.2878978095452 | 9.54582568294392 | 6.40265188186281 |
| chr10 | 118682205 | 118682224 | - | 8.03168461498692 | 4.54798296825753 | 3.57968463110397 | 2.13421729395427 |
| chr10 | 118682422 | 118682436 | - | 3.01188173062009 | 5.68497871032191 | 3.57968463110397 | 2.13421729395427 |
| chr10 | 118682589 | 118682597 | - | 5.01980288436682 | 6.82197445238629 | 5.96614105183995 | 1.06710864697714 |
| chr10 | 118684623 | 118684691 | - | 29.1148567293276 | 27.2878978095452 | 7.15936926220794 | 7.46976052883995 |
| chr10 | 118685229 | 118685237 | - | 5.01980288436682 | 1.13699574206438 | 3.57968463110397 | 2.13421729395427 |
| chr10 | 118685909 | 118685928 | - | 13.0514874993537 | 11.3699574206438 | 13.1255103140479 | 5.33554323488568 |
| chr10 | 118686001 | 118686006 | - | 2.00792115374673 | 3.41098722619315 | 4.77291284147196 | 1.06710864697714 |
| chr10 | 118686033 | 118686055 | - | 8.03168461498692 | 10.2329616785794 | 32.2171616799357 | 4.26843458790854 |
| chr10 | 118686126 | 118686154 | - | 5.01980288436682 | 7.95897019445067 | 3.57968463110397 | 2.13421729395427 |
| chr10 | 119033783 | 119033785 | - | 1.00396057687336 | 1.13699574206438 | 2.38645642073598 | 2.13421729395427 |
| chr10 | 119034504 | 119034518 | - | 1.00396057687336 | 2.27399148412876 | 1.19322821036799 | 6.40265188186281 |
| chr10 | 119034635 | 119034651 | - | 5.01980288436682 | 3.41098722619315 | 1.19322821036799 | 3.20132594093141 |
| chr10 | 119035296 | 119035322 | - | 10.0396057687336 | 10.2329616785794 | 16.7051949451519 | 13.8724124107028 |
| chr10 | 119035582 | 119035597 | - | 152.602007684751 | 209.207216539846 | 62.0478669391355 | 81.1002571702623 |
| chr10 | 119035654 | 119035673 | - | 20.0792115374673 | 17.0549361309657 | 5.96614105183995 | 6.40265188186281 |
| chr10 | 119035699 | 119035724 | - | 31.1227778830743 | 36.3838637460602 | 5.96614105183995 | 1.06710864697714 |
| chr10 | 119035898 | 119035913 | - | 1.00396057687336 | 1.13699574206438 | 7.15936926220794 | 2.13421729395427 |
| chr10 | 119035968 | 119036075 | - | 29.1148567293276 | 38.657855230189 | 40.5697591525116 | 28.8119334683827 |
| chr10 | 119140974 | 119141044 | - | 45.1782259593014 | 64.8087572976698 | 36.9900745214077 | 69.3620620535138 |
| chr10 | 119167734 | 119167882 | - | 36.1425807674411 | 55.7127913611547 | 22.6713359969918 | 14.9395210576799 |
| chr10 | 119167901 | 119167934 | - | 9.03564519186028 | 13.6439489047726 | 5.96614105183995 | 4.26843458790854 |
| chr10 | 119167973 | 119168039 | - | 18.0712903837206 | 21.6029190992233 | 7.15936926220794 | 6.40265188186281 |
| chr10 | 119168115 | 119168162 | - | 10.0396057687336 | 3.41098722619315 | 1.19322821036799 | 2.13421729395427 |
| chr10 | 119168168 | 119168216 | - | 27.1069355755808 | 22.7399148412876 | 4.77291284147196 | 9.60397782279422 |
| chr10 | 119168238 | 119168476 | - | 72.2851615348822 | 64.8087572976698 | 22.6713359969918 | 38.4159112911769 |
| chr10 | 119499942 | 119499949 | - | 6.02376346124019 | 4.54798296825753 | 29.8307052591997 | 43.7514545260626 |
| chr10 | 119500019 | 119500025 | - | 3.01188173062009 | 5.68497871032191 | 4.77291284147196 | 16.006629704657 |
| chr10 | 119571922 | 119571923 | - | 4.01584230749346 | 1.13699574206438 | 1.19322821036799 | 1.06710864697714 |
| chr10 | 119571976 | 119571999 | - | 10.0396057687336 | 13.6439489047726 | 1.19322821036799 | 2.13421729395427 |
| chr10 | 119572038 | 119572044 | - | 1.00396057687336 | 4.54798296825753 | 2.38645642073598 | 1.06710864697714 |
| chr10 | 119572094 | 119572105 | - | 8.03168461498692 | 3.41098722619315 | 2.38645642073598 | 3.20132594093141 |
| chr10 | 119572148 | 119572160 | - | 3.01188173062009 | 3.41098722619315 | 4.77291284147196 | 3.20132594093141 |
| chr10 | 119573845 | 119573867 | - | 16.0633692299738 | 15.9179403889013 | 4.77291284147196 | 9.60397782279422 |
| chr10 | 119574630 | 119574633 | - | 4.01584230749346 | 1.13699574206438 | 3.57968463110397 | 1.06710864697714 |
| chr10 | 119574732 | 119574736 | - | 1.00396057687336 | 1.13699574206438 | 1.19322821036799 | 2.13421729395427 |
| chr10 | 119575500 | 119575521 | - | 13.0514874993537 | 9.09596593651506 | 11.9322821036799 | 19.2079556455884 |
| chr10 | 119575627 | 119575634 | - | 2.00792115374673 | 4.54798296825753 | 1.19322821036799 | 3.20132594093141 |
| chr10 | 119575660 | 119575698 | - | 3.01188173062009 | 1.13699574206438 | 7.15936926220794 | 5.33554323488568 |
| chr10 | 119677409 | 119677439 | + | 3.01188173062009 | 2.27399148412876 | 3.57968463110397 | 2.13421729395427 |
| chr10 | 119677721 | 119677775 | + | 14.0554480762271 | 4.54798296825753 | 10.7390538933119 | 3.20132594093141 |
| chr10 | 119830176 | 119830185 | - | 7.02772403811355 | 4.54798296825753 | 2.38645642073598 | 1.06710864697714 |
| chr10 | 119831446 | 119831449 | - | 2.00792115374673 | 4.54798296825753 | 1.19322821036799 | 2.13421729395427 |
| chr10 | 119940937 | 119940943 | + | 2.00792115374673 | 2.27399148412876 | 3.57968463110397 | 1.06710864697714 |
| chr10 | 119941563 | 119941572 | + | 4.01584230749346 | 4.54798296825753 | 1.19322821036799 | 4.26843458790854 |
| chr10 | 120909279 | 120909333 | + | 7.02772403811355 | 3.41098722619315 | 4.77291284147196 | 2.13421729395427 |
| chr10 | 120909429 | 120909465 | + | 2.00792115374673 | 5.68497871032191 | 1.19322821036799 | 2.13421729395427 |
| chr10 | 121957109 | 121957133 | - | 12.0475269224804 | 13.6439489047726 | 5.96614105183995 | 5.33554323488568 |
| chr10 | 122430016 | 122430035 | + | 22.087132691214 | 15.9179403889013 | 1.19322821036799 | 2.13421729395427 |
| chr10 | 122430161 | 122430230 | + | 6.02376346124019 | 7.95897019445067 | 2.38645642073598 | 1.06710864697714 |
| chr10 | 122432039 | 122432072 | + | 17.0673298068472 | 32.9728765198671 | 2.38645642073598 | 1.06710864697714 |
| chr10 | 122992988 | 122993010 | - | 3.01188173062009 | 3.41098722619315 | 3.57968463110397 | 2.13421729395427 |
| chr10 | 123054996 | 123055064 | + | 4.01584230749346 | 1.13699574206438 | 3.57968463110397 | 2.13421729395427 |
| chr10 | 123163820 | 123163865 | + | 2.00792115374673 | 7.95897019445067 | 16.7051949451519 | 11.7381951167485 |
| chr10 | 123163894 | 123163900 | + | 3.01188173062009 | 2.27399148412876 | 5.96614105183995 | 4.26843458790854 |
| chr10 | 123163923 | 123163926 | + | 5.01980288436682 | 2.27399148412876 | 2.38645642073598 | 1.06710864697714 |
| chr10 | 123163948 | 123163976 | + | 9.03564519186028 | 19.3289276150945 | 14.3187385244159 | 39.483019938154 |
| chr10 | 123164071 | 123164100 | + | 3.01188173062009 | 10.2329616785794 | 5.96614105183995 | 6.40265188186281 |
| chr10 | 123164199 | 123164213 | + | 7.02772403811355 | 1.13699574206438 | 3.57968463110397 | 3.20132594093141 |
| chr10 | 123164510 | 123164512 | + | 2.00792115374673 | 1.13699574206438 | 5.96614105183995 | 4.26843458790854 |
| chr10 | 123164536 | 123164553 | + | 4.01584230749346 | 1.13699574206438 | 4.77291284147196 | 4.26843458790854 |
| chr10 | 123164873 | 123164930 | + | 9.03564519186028 | 12.5069531627082 | 5.96614105183995 | 10.6710864697714 |
| chr10 | 123165113 | 123165150 | + | 49.1940682667949 | 54.5757956190903 | 36.9900745214077 | 33.0803680562912 |
| chr10 | 123165207 | 123165245 | + | 26.1029749987075 | 31.8358807778027 | 16.7051949451519 | 16.006629704657 |
| chr10 | 123165264 | 123165299 | + | 20.0792115374673 | 19.3289276150945 | 4.77291284147196 | 3.20132594093141 |
| chr10 | 123165324 | 123165350 | + | 10.0396057687336 | 5.68497871032191 | 2.38645642073598 | 2.13421729395427 |
| chr10 | 124397447 | 124397452 | - | 10.0396057687336 | 7.95897019445067 | 25.0577924177278 | 16.006629704657 |
| chr10 | 124397503 | 124397517 | - | 1.00396057687336 | 4.54798296825753 | 4.77291284147196 | 1.06710864697714 |
| chr10 | 124397540 | 124397624 | - | 4.01584230749346 | 1.13699574206438 | 11.9322821036799 | 2.13421729395427 |
| chr10 | 124397686 | 124397711 | - | 1.00396057687336 | 7.95897019445067 | 9.54582568294392 | 8.53686917581709 |
| chr10 | 124620063 | 124620066 | - | 2.00792115374673 | 2.27399148412876 | 1.19322821036799 | 1.06710864697714 |
| chr10 | 124620159 | 124620182 | - | 21.0831721143407 | 6.82197445238629 | 17.8984231555198 | 9.60397782279422 |
| chr10 | 124621031 | 124621033 | - | 2.00792115374673 | 1.13699574206438 | 1.19322821036799 | 1.06710864697714 |
| chr10 | 124760216 | 124760241 | - | 11.043566345607 | 5.68497871032191 | 3.57968463110397 | 3.20132594093141 |
| chr10 | 124836378 | 124836381 | + | 7.02772403811355 | 7.95897019445067 | 3.57968463110397 | 2.13421729395427 |
| chr10 | 125788612 | 125788641 | - | 19.0752509605939 | 17.0549361309657 | 3.57968463110397 | 3.20132594093141 |
| chr10 | 125836329 | 125836333 | + | 7.02772403811355 | 5.68497871032191 | 8.35259747257593 | 3.20132594093141 |
| chr10 | 125836388 | 125836410 | + | 10.0396057687336 | 5.68497871032191 | 1.19322821036799 | 1.06710864697714 |
| chr10 | 125836457 | 125836461 | + | 8.03168461498692 | 9.09596593651506 | 3.57968463110397 | 2.13421729395427 |
| chr10 | 125836482 | 125836525 | + | 68.2693192273888 | 114.836569948503 | 33.4103898903037 | 26.6777161744284 |
| chr10 | 125836578 | 125836627 | - | 7.02772403811355 | 17.0549361309657 | 2.38645642073598 | 3.20132594093141 |
| chr10 | 125836657 | 125836672 | - | 3.01188173062009 | 6.82197445238629 | 1.19322821036799 | 4.26843458790854 |
| chr10 | 125841428 | 125841430 | + | 7.02772403811355 | 1.13699574206438 | 1.19322821036799 | 6.40265188186281 |
| chr10 | 125842443 | 125842446 | + | 2.00792115374673 | 1.13699574206438 | 10.7390538933119 | 8.53686917581709 |
| chr10 | 125842525 | 125842526 | + | 3.01188173062009 | 1.13699574206438 | 1.19322821036799 | 2.13421729395427 |
| chr10 | 125842637 | 125842638 | + | 3.01188173062009 | 1.13699574206438 | 1.19322821036799 | 2.13421729395427 |
| chr10 | 125842708 | 125842709 | + | 1.00396057687336 | 4.54798296825753 | 2.38645642073598 | 5.33554323488568 |
| chr10 | 125853264 | 125853297 | + | 7.02772403811355 | 7.95897019445067 | 3.57968463110397 | 4.26843458790854 |
| chr10 | 125853385 | 125853394 | + | 5.01980288436682 | 6.82197445238629 | 17.8984231555198 | 4.26843458790854 |
| chr10 | 125853460 | 125853463 | + | 2.00792115374673 | 4.54798296825753 | 3.57968463110397 | 1.06710864697714 |
| chr10 | 128096690 | 128096699 | - | 5.01980288436682 | 9.09596593651506 | 3.57968463110397 | 3.20132594093141 |
| chr10 | 128097058 | 128097097 | - | 17.0673298068472 | 2.27399148412876 | 2.38645642073598 | 2.13421729395427 |
| chr10 | 128097276 | 128097280 | - | 11.043566345607 | 7.95897019445067 | 3.57968463110397 | 1.06710864697714 |
| chr10 | 128097393 | 128097411 | - | 11.043566345607 | 2.27399148412876 | 2.38645642073598 | 8.53686917581709 |
| chr10 | 128097708 | 128097715 | - | 12.0475269224804 | 7.95897019445067 | 2.38645642073598 | 3.20132594093141 |
| chr10 | 128097772 | 128097822 | - | 15.0594086531005 | 13.6439489047726 | 4.77291284147196 | 1.06710864697714 |
| chr10 | 128097932 | 128097938 | - | 2.00792115374673 | 4.54798296825753 | 2.38645642073598 | 7.46976052883995 |
| chr10 | 128097977 | 128097980 | - | 30.1188173062009 | 13.6439489047726 | 4.77291284147196 | 4.26843458790854 |
| chr10 | 128099052 | 128099113 | - | 19.0752509605939 | 7.95897019445067 | 3.57968463110397 | 9.60397782279422 |
| chr10 | 128099131 | 128099175 | - | 32.1267384599477 | 30.6988850357383 | 19.0916513658878 | 11.7381951167485 |
| chr10 | 130179473 | 130179487 | + | 7.02772403811355 | 7.95897019445067 | 4.77291284147196 | 4.26843458790854 |
| chr10 | 130179519 | 130179550 | + | 4.01584230749346 | 10.2329616785794 | 2.38645642073598 | 1.06710864697714 |
| chr10 | 131955964 | 131955979 | + | 3.01188173062009 | 4.54798296825753 | 4.77291284147196 | 3.20132594093141 |
| chr10 | 131956500 | 131956508 | + | 4.01584230749346 | 3.41098722619315 | 2.38645642073598 | 1.06710864697714 |
| chr10 | 131967791 | 131967891 | - | 5.01980288436682 | 13.6439489047726 | 57.2749540976635 | 49.0869977609482 |
| chr10 | 131967930 | 131967977 | - | 8.03168461498692 | 2.27399148412876 | 93.0718004087032 | 30.9461507623369 |
| chr10 | 131968013 | 131968039 | - | 4.01584230749346 | 2.27399148412876 | 27.4442488384638 | 8.53686917581709 |
| chr10 | 131968062 | 131968082 | - | 3.01188173062009 | 2.27399148412876 | 17.8984231555198 | 7.46976052883995 |
| chr10 | 131968175 | 131968238 | - | 8.03168461498692 | 1.13699574206438 | 59.6614105183995 | 24.5434988804741 |
| chr10 | 132330356 | 132330377 | - | 9.03564519186028 | 11.3699574206438 | 8.35259747257593 | 2.13421729395427 |
| chr10 | 132782498 | 132782518 | + | 14.0554480762271 | 5.68497871032191 | 5.96614105183995 | 1.06710864697714 |
| chr10 | 132782823 | 132782826 | + | 13.0514874993537 | 2.27399148412876 | 4.77291284147196 | 4.26843458790854 |
| chr10 | 132782850 | 132782895 | + | 15.0594086531005 | 11.3699574206438 | 10.7390538933119 | 8.53686917581709 |
| chr10 | 132783144 | 132783183 | + | 4.01584230749346 | 2.27399148412876 | 5.96614105183995 | 6.40265188186281 |
| chr10 | 132783229 | 132783275 | + | 30.1188173062009 | 21.6029190992233 | 9.54582568294392 | 18.1408469986113 |
| chr10 | 133279666 | 133279746 | - | 46.1821865361748 | 52.3018041349616 | 23.8645642073598 | 52.2883237018796 |
| chr10 | 133312825 | 133312859 | + | 3.01188173062009 | 1.13699574206438 | 9.54582568294392 | 12.8053037637256 |
| chr10 | 133313055 | 133313122 | + | 21.0831721143407 | 20.4659233571589 | 4.77291284147196 | 13.8724124107028 |
| chr10 | 133355185 | 133355257 | - | 6.02376346124019 | 5.68497871032191 | 7.15936926220794 | 23.476390233497 |
| chr10 | 133362503 | 133362648 | - | 1030.06355187207 | 593.511777357607 | 329.330986061565 | 481.265999786688 |
| chr10 | 133362672 | 133362689 | - | 46.1821865361748 | 32.9728765198671 | 7.15936926220794 | 21.3421729395427 |
| chr10 | 133362729 | 133362779 | - | 39.1544624980612 | 15.9179403889013 | 22.6713359969918 | 43.7514545260626 |
| chr10 | 133362800 | 133362836 | - | 45.1782259593014 | 23.876910583352 | 39.3765309421437 | 46.952780466994 |
| chr10 | 133362861 | 133362932 | - | 28.1108961524542 | 25.0139063254164 | 33.4103898903037 | 38.4159112911769 |
| chr10 | 133420334 | 133420455 | + | 3.01188173062009 | 2.27399148412876 | 1.19322821036799 | 4.26843458790854 |
| chr11 | 202974 | 203013 | - | 3.01188173062009 | 3.41098722619315 | 4.77291284147196 | 2.13421729395427 |
| chr11 | 203035 | 203056 | - | 2.00792115374673 | 2.27399148412876 | 4.77291284147196 | 10.6710864697714 |
| chr11 | 203276 | 203383 | - | 5.01980288436682 | 13.6439489047726 | 26.2510206280958 | 23.476390233497 |
| chr11 | 203557 | 203569 | - | 4.01584230749346 | 1.13699574206438 | 1.19322821036799 | 3.20132594093141 |
| chr11 | 204242 | 204295 | - | 2.00792115374673 | 6.82197445238629 | 7.15936926220794 | 7.46976052883995 |
| chr11 | 214408 | 214422 | + | 3.01188173062009 | 1.13699574206438 | 2.38645642073598 | 3.20132594093141 |
| chr11 | 214881 | 214889 | + | 3.01188173062009 | 4.54798296825753 | 1.19322821036799 | 9.60397782279422 |
| chr11 | 216166 | 216224 | - | 2.00792115374673 | 1.13699574206438 | 3.57968463110397 | 1.06710864697714 |
| chr11 | 252776 | 252780 | + | 1.00396057687336 | 1.13699574206438 | 1.19322821036799 | 3.20132594093141 |
| chr11 | 252812 | 252867 | + | 10.0396057687336 | 3.41098722619315 | 17.8984231555198 | 14.9395210576799 |
| chr11 | 252892 | 252899 | + | 8.03168461498692 | 3.41098722619315 | 10.7390538933119 | 8.53686917581709 |
| chr11 | 252923 | 252958 | + | 30.1188173062009 | 35.2468680039958 | 39.3765309421437 | 41.6172372321083 |
| chr11 | 295160 | 295185 | + | 2.00792115374673 | 1.13699574206438 | 11.9322821036799 | 2.13421729395427 |
| chr11 | 295314 | 295351 | + | 1.00396057687336 | 1.13699574206438 | 20.2848795762558 | 16.006629704657 |
| chr11 | 309183 | 309331 | + | 279.101040370795 | 206.933225055718 | 409.27727615622 | 89.6371263460794 |
| chr11 | 315155 | 315259 | + | 86.3406096111093 | 53.438799877026 | 335.297127113405 | 250.770532039627 |
| chr11 | 319740 | 319753 | - | 39.1544624980612 | 36.3838637460602 | 11.9322821036799 | 6.40265188186281 |
| chr11 | 327284 | 327303 | + | 1.00396057687336 | 3.41098722619315 | 13.1255103140479 | 6.40265188186281 |
| chr11 | 404773 | 404779 | + | 4.01584230749346 | 1.13699574206438 | 2.38645642073598 | 4.26843458790854 |
| chr11 | 404835 | 404846 | + | 4.01584230749346 | 9.09596593651506 | 7.15936926220794 | 5.33554323488568 |
| chr11 | 405740 | 405777 | - | 16.0633692299738 | 3.41098722619315 | 10.7390538933119 | 33.0803680562912 |
| chr11 | 726899 | 727093 | + | 36.1425807674411 | 27.2878978095452 | 51.3088130458235 | 24.5434988804741 |
| chr11 | 727562 | 727702 | + | 20.0792115374673 | 26.1509020674808 | 7.15936926220794 | 14.9395210576799 |
| chr11 | 764860 | 764905 | + | 11.043566345607 | 3.41098722619315 | 20.2848795762558 | 12.8053037637256 |
| chr11 | 767248 | 767303 | - | 42.1663442286813 | 56.8497871032191 | 48.9223566250876 | 61.8923015246739 |
| chr11 | 767324 | 767361 | - | 4.01584230749346 | 6.82197445238629 | 9.54582568294392 | 7.46976052883995 |
| chr11 | 768506 | 768511 | - | 2.00792115374673 | 1.13699574206438 | 3.57968463110397 | 4.26843458790854 |
| chr11 | 768681 | 768707 | - | 5.01980288436682 | 3.41098722619315 | 10.7390538933119 | 11.7381951167485 |
| chr11 | 768822 | 768841 | - | 2.00792115374673 | 2.27399148412876 | 5.96614105183995 | 6.40265188186281 |
| chr11 | 777548 | 777662 | + | 148.586165377258 | 114.836569948503 | 32.2171616799357 | 104.576647403759 |
| chr11 | 783594 | 783623 | + | 11.043566345607 | 7.95897019445067 | 3.57968463110397 | 9.60397782279422 |
| chr11 | 790604 | 790687 | - | 87.3445701879827 | 30.6988850357383 | 42.9562155732476 | 103.509538756782 |
| chr11 | 790950 | 790957 | - | 10.0396057687336 | 3.41098722619315 | 7.15936926220794 | 3.20132594093141 |
| chr11 | 791066 | 791083 | - | 4.01584230749346 | 3.41098722619315 | 3.57968463110397 | 20.2750642925656 |
| chr11 | 791285 | 791298 | - | 2.00792115374673 | 2.27399148412876 | 4.77291284147196 | 2.13421729395427 |
| chr11 | 791380 | 791388 | - | 3.01188173062009 | 3.41098722619315 | 5.96614105183995 | 3.20132594093141 |
| chr11 | 791723 | 791743 | - | 11.043566345607 | 4.54798296825753 | 3.57968463110397 | 1.06710864697714 |
| chr11 | 812848 | 812861 | + | 27.1069355755808 | 37.5208594881246 | 83.5259747257593 | 51.2212150549025 |
| chr11 | 824846 | 824897 | + | 14.0554480762271 | 9.09596593651506 | 9.54582568294392 | 7.46976052883995 |
| chr11 | 824994 | 825128 | + | 280.105000947669 | 150.083437952498 | 58.4681823080315 | 101.375321462828 |
| chr11 | 825151 | 825192 | + | 2.00792115374673 | 6.82197445238629 | 2.38645642073598 | 6.40265188186281 |
| chr11 | 838217 | 838254 | + | 2.00792115374673 | 1.13699574206438 | 4.77291284147196 | 5.33554323488568 |
| chr11 | 838302 | 838305 | + | 2.00792115374673 | 1.13699574206438 | 3.57968463110397 | 2.13421729395427 |
| chr11 | 838360 | 838458 | + | 67.2653586505154 | 76.1787147183136 | 116.936364616063 | 69.3620620535138 |
| chr11 | 838465 | 838540 | + | 35.1386201905678 | 22.7399148412876 | 66.8207797806074 | 37.3488026441998 |
| chr11 | 838580 | 838634 | + | 10.0396057687336 | 4.54798296825753 | 13.1255103140479 | 24.5434988804741 |
| chr11 | 838675 | 838811 | + | 66.2613980736421 | 29.5618892936739 | 34.6036181006717 | 41.6172372321083 |
| chr11 | 840222 | 840304 | - | 127.502993262917 | 106.877599754052 | 93.0718004087032 | 128.053037637256 |
| chr11 | 840314 | 840419 | - | 10.0396057687336 | 9.09596593651506 | 9.54582568294392 | 7.46976052883995 |
| chr11 | 867065 | 867086 | + | 5.01980288436682 | 21.6029190992233 | 1.19322821036799 | 5.33554323488568 |
| chr11 | 868721 | 868737 | - | 3.01188173062009 | 1.13699574206438 | 20.2848795762558 | 12.8053037637256 |
| chr11 | 869782 | 869811 | - | 12.0475269224804 | 10.2329616785794 | 10.7390538933119 | 18.1408469986113 |
| chr11 | 1275117 | 1275139 | - | 2.00792115374673 | 1.13699574206438 | 8.35259747257593 | 10.6710864697714 |
| chr11 | 1275611 | 1275622 | - | 1.00396057687336 | 1.13699574206438 | 1.19322821036799 | 1.06710864697714 |
| chr11 | 1276648 | 1276664 | - | 1.00396057687336 | 1.13699574206438 | 1.19322821036799 | 1.06710864697714 |
| chr11 | 1469956 | 1470008 | - | 4.01584230749346 | 9.09596593651506 | 20.2848795762558 | 14.9395210576799 |
| chr11 | 1470043 | 1470079 | - | 4.01584230749346 | 4.54798296825753 | 13.1255103140479 | 6.40265188186281 |
| chr11 | 1470154 | 1470232 | - | 4.01584230749346 | 10.2329616785794 | 27.4442488384638 | 23.476390233497 |
| chr11 | 1752775 | 1753386 | - | 65.2574374967687 | 47.753821166704 | 15.5119667347839 | 19.2079556455884 |
| chr11 | 1753456 | 1753669 | - | 20.0792115374673 | 6.82197445238629 | 2.38645642073598 | 2.13421729395427 |
| chr11 | 2396886 | 2396899 | + | 1.00396057687336 | 2.27399148412876 | 1.19322821036799 | 8.53686917581709 |
| chr11 | 2396927 | 2396933 | + | 2.00792115374673 | 1.13699574206438 | 3.57968463110397 | 8.53686917581709 |
| chr11 | 2397235 | 2397252 | + | 1.00396057687336 | 1.13699574206438 | 4.77291284147196 | 10.6710864697714 |
| chr11 | 2397278 | 2397292 | + | 8.03168461498692 | 7.95897019445067 | 13.1255103140479 | 21.3421729395427 |
| chr11 | 2403816 | 2403824 | + | 14.0554480762271 | 6.82197445238629 | 5.96614105183995 | 3.20132594093141 |
| chr11 | 2945069 | 2945088 | - | 133.526756724157 | 122.795540142953 | 116.936364616063 | 107.777973344691 |
| chr11 | 3675136 | 3675184 | - | 11.043566345607 | 12.5069531627082 | 5.96614105183995 | 14.9395210576799 |
| chr11 | 3675242 | 3675275 | - | 9.03564519186028 | 6.82197445238629 | 2.38645642073598 | 3.20132594093141 |
| chr11 | 3675350 | 3675398 | - | 14.0554480762271 | 10.2329616785794 | 11.9322821036799 | 11.7381951167485 |
| chr11 | 3675407 | 3675466 | - | 21.0831721143407 | 5.68497871032191 | 8.35259747257593 | 2.13421729395427 |
| chr11 | 3675557 | 3675595 | - | 60.2376346124019 | 31.8358807778027 | 33.4103898903037 | 45.8856718200168 |
| chr11 | 3675619 | 3675622 | - | 3.01188173062009 | 3.41098722619315 | 2.38645642073598 | 1.06710864697714 |
| chr11 | 3675695 | 3675708 | - | 4.01584230749346 | 2.27399148412876 | 13.1255103140479 | 2.13421729395427 |
| chr11 | 3675746 | 3675759 | - | 5.01980288436682 | 1.13699574206438 | 2.38645642073598 | 6.40265188186281 |
| chr11 | 3675861 | 3675863 | - | 2.00792115374673 | 1.13699574206438 | 1.19322821036799 | 1.06710864697714 |
| chr11 | 3675932 | 3675980 | - | 6.02376346124019 | 1.13699574206438 | 10.7390538933119 | 11.7381951167485 |
| chr11 | 3676101 | 3676115 | - | 3.01188173062009 | 3.41098722619315 | 3.57968463110397 | 7.46976052883995 |
| chr11 | 3676151 | 3676185 | - | 5.01980288436682 | 2.27399148412876 | 5.96614105183995 | 9.60397782279422 |
| chr11 | 3711918 | 3711932 | - | 4.01584230749346 | 5.68497871032191 | 3.57968463110397 | 6.40265188186281 |
| chr11 | 3711967 | 3711968 | - | 1.00396057687336 | 1.13699574206438 | 1.19322821036799 | 1.06710864697714 |
| chr11 | 3712005 | 3712019 | - | 13.0514874993537 | 23.876910583352 | 9.54582568294392 | 11.7381951167485 |
| chr11 | 3712100 | 3712102 | - | 4.01584230749346 | 9.09596593651506 | 1.19322821036799 | 2.13421729395427 |
| chr11 | 3712248 | 3712255 | - | 4.01584230749346 | 14.780944646837 | 5.96614105183995 | 2.13421729395427 |
| chr11 | 3712398 | 3712417 | - | 20.0792115374673 | 6.82197445238629 | 7.15936926220794 | 17.0737383516342 |
| chr11 | 3826217 | 3826272 | + | 93.3683336492229 | 37.5208594881246 | 27.4442488384638 | 55.4896496428111 |
| chr11 | 3827003 | 3827029 | - | 39.1544624980612 | 60.2607743294122 | 36.9900745214077 | 75.7647139353766 |
| chr11 | 3827102 | 3827124 | - | 2.00792115374673 | 6.82197445238629 | 1.19322821036799 | 14.9395210576799 |
| chr11 | 3855139 | 3855222 | + | 17.0673298068472 | 28.4248935516096 | 13.1255103140479 | 18.1408469986113 |
| chr11 | 3856098 | 3856137 | + | 9.03564519186028 | 14.780944646837 | 13.1255103140479 | 2.13421729395427 |
| chr11 | 4091698 | 4091724 | + | 1.00396057687336 | 2.27399148412876 | 9.54582568294392 | 2.13421729395427 |
| chr11 | 4092198 | 4092241 | + | 2.00792115374673 | 3.41098722619315 | 1.19322821036799 | 5.33554323488568 |
| chr11 | 4093066 | 4093147 | + | 6.02376346124019 | 12.5069531627082 | 7.15936926220794 | 10.6710864697714 |
| chr11 | 4138372 | 4138524 | + | 191.756470182813 | 218.303182476361 | 46.5359002043516 | 103.509538756782 |
| chr11 | 4138668 | 4138677 | + | 37.1465413443145 | 50.0278126508328 | 7.15936926220794 | 13.8724124107028 |
| chr11 | 4138765 | 4138858 | + | 140.554480762271 | 241.043097317649 | 68.0140079909754 | 43.7514545260626 |
| chr11 | 4384933 | 4384937 | - | 6.02376346124019 | 1.13699574206438 | 3.57968463110397 | 5.33554323488568 |
| chr11 | 4598894 | 4598929 | - | 6.02376346124019 | 6.82197445238629 | 3.57968463110397 | 5.33554323488568 |
| chr11 | 5268367 | 5268600 | - | 3.01188173062009 | 1.13699574206438 | 647.922918229818 | 4639.78839705659 |
| chr11 | 5663927 | 5663928 | - | 1.00396057687336 | 1.13699574206438 | 1.19322821036799 | 1.06710864697714 |
| chr11 | 6211423 | 6211432 | - | 2.00792115374673 | 4.54798296825753 | 1.19322821036799 | 2.13421729395427 |
| chr11 | 6211455 | 6211464 | - | 3.01188173062009 | 4.54798296825753 | 3.57968463110397 | 9.60397782279422 |
| chr11 | 6394778 | 6394811 | + | 12.0475269224804 | 11.3699574206438 | 2.38645642073598 | 5.33554323488568 |
| chr11 | 6394849 | 6394859 | + | 12.0475269224804 | 17.0549361309657 | 5.96614105183995 | 6.40265188186281 |
| chr11 | 6394906 | 6394917 | + | 3.01188173062009 | 2.27399148412876 | 1.19322821036799 | 2.13421729395427 |
| chr11 | 6476539 | 6476569 | - | 16.0633692299738 | 17.0549361309657 | 7.15936926220794 | 4.26843458790854 |
| chr11 | 6478129 | 6478152 | - | 8.03168461498692 | 4.54798296825753 | 21.4781077866238 | 5.33554323488568 |
| chr11 | 6478194 | 6478201 | - | 2.00792115374673 | 4.54798296825753 | 8.35259747257593 | 4.26843458790854 |
| chr11 | 6482240 | 6482246 | + | 4.01584230749346 | 1.13699574206438 | 1.19322821036799 | 1.06710864697714 |
| chr11 | 6482277 | 6482357 | + | 8.03168461498692 | 9.09596593651506 | 21.4781077866238 | 25.6106075274513 |
| chr11 | 6482481 | 6482549 | + | 3.01188173062009 | 6.82197445238629 | 8.35259747257593 | 7.46976052883995 |
| chr11 | 6482568 | 6482575 | + | 2.00792115374673 | 1.13699574206438 | 10.7390538933119 | 4.26843458790854 |
| chr11 | 6482612 | 6482697 | + | 6.02376346124019 | 7.95897019445067 | 28.6374770488317 | 28.8119334683827 |
| chr11 | 6482718 | 6482731 | + | 8.03168461498692 | 7.95897019445067 | 17.8984231555198 | 22.4092815865198 |
| chr11 | 6482899 | 6482925 | + | 2.00792115374673 | 3.41098722619315 | 4.77291284147196 | 2.13421729395427 |
| chr11 | 6482945 | 6482973 | + | 1.00396057687336 | 1.13699574206438 | 5.96614105183995 | 2.13421729395427 |
| chr11 | 6483034 | 6483038 | + | 2.00792115374673 | 1.13699574206438 | 2.38645642073598 | 6.40265188186281 |
| chr11 | 6483079 | 6483111 | + | 1.00396057687336 | 7.95897019445067 | 14.3187385244159 | 8.53686917581709 |
| chr11 | 6483136 | 6483238 | + | 11.043566345607 | 13.6439489047726 | 34.6036181006717 | 10.6710864697714 |
| chr11 | 6483603 | 6483615 | + | 1.00396057687336 | 3.41098722619315 | 3.57968463110397 | 8.53686917581709 |
| chr11 | 6484368 | 6484420 | + | 23.0910932680874 | 17.0549361309657 | 13.1255103140479 | 32.0132594093141 |
| chr11 | 6484437 | 6484534 | + | 58.2297134586551 | 47.753821166704 | 75.1733772531833 | 97.1068868749194 |
| chr11 | 6484553 | 6484563 | + | 3.01188173062009 | 6.82197445238629 | 10.7390538933119 | 4.26843458790854 |
| chr11 | 6484588 | 6484625 | + | 13.0514874993537 | 12.5069531627082 | 17.8984231555198 | 30.9461507623369 |
| chr11 | 6598106 | 6598144 | - | 6.02376346124019 | 10.2329616785794 | 10.7390538933119 | 9.60397782279422 |
| chr11 | 6598341 | 6598359 | - | 5.01980288436682 | 3.41098722619315 | 1.19322821036799 | 1.06710864697714 |
| chr11 | 6599952 | 6599956 | - | 19.0752509605939 | 40.9318467143178 | 8.35259747257593 | 26.6777161744284 |
| chr11 | 6600033 | 6600035 | - | 3.01188173062009 | 19.3289276150945 | 2.38645642073598 | 13.8724124107028 |
| chr11 | 6610775 | 6610791 | + | 6.02376346124019 | 2.27399148412876 | 2.38645642073598 | 3.20132594093141 |
| chr11 | 6610810 | 6610866 | + | 133.526756724157 | 119.38455291676 | 19.0916513658878 | 33.0803680562912 |
| chr11 | 6610872 | 6610944 | - | 11.043566345607 | 11.3699574206438 | 8.35259747257593 | 17.0737383516342 |
| chr11 | 6612791 | 6613251 | - | 909.588282647268 | 750.417189762492 | 218.360762497342 | 312.662833564301 |
| chr11 | 6613284 | 6613295 | - | 7.02772403811355 | 3.41098722619315 | 1.19322821036799 | 3.20132594093141 |
| chr11 | 6613843 | 6613860 | - | 3.01188173062009 | 2.27399148412876 | 3.57968463110397 | 2.13421729395427 |
| chr11 | 6613972 | 6613996 | - | 7.02772403811355 | 2.27399148412876 | 3.57968463110397 | 7.46976052883995 |
| chr11 | 6614034 | 6614101 | - | 14.0554480762271 | 12.5069531627082 | 1.19322821036799 | 4.26843458790854 |
| chr11 | 6614122 | 6614144 | - | 5.01980288436682 | 9.09596593651506 | 1.19322821036799 | 4.26843458790854 |
| chr11 | 6614224 | 6614263 | - | 5.01980288436682 | 11.3699574206438 | 2.38645642073598 | 3.20132594093141 |
| chr11 | 6614280 | 6614323 | - | 7.02772403811355 | 1.13699574206438 | 1.19322821036799 | 1.06710864697714 |
| chr11 | 6614397 | 6614442 | - | 9.03564519186028 | 3.41098722619315 | 1.19322821036799 | 3.20132594093141 |
| chr11 | 6614493 | 6614571 | - | 25.0990144218341 | 22.7399148412876 | 13.1255103140479 | 11.7381951167485 |
| chr11 | 6618831 | 6618916 | - | 13.0514874993537 | 9.09596593651506 | 8.35259747257593 | 1.06710864697714 |
| chr11 | 6681938 | 6682045 | - | 71.2812009580089 | 90.9596593651506 | 21.4781077866238 | 62.959410171651 |
| chr11 | 8614073 | 8614078 | - | 13.0514874993537 | 12.5069531627082 | 7.15936926220794 | 3.20132594093141 |
| chr11 | 8685438 | 8685439 | + | 2.00792115374673 | 2.27399148412876 | 7.15936926220794 | 1.06710864697714 |
| chr11 | 8685784 | 8685834 | + | 1645.49138549544 | 2048.86632720002 | 1118.05483311481 | 1054.30334321341 |
| chr11 | 8685903 | 8685943 | + | 7.02772403811355 | 9.09596593651506 | 3.57968463110397 | 1.06710864697714 |
| chr11 | 8685970 | 8685997 | + | 8.03168461498692 | 5.68497871032191 | 8.35259747257593 | 2.13421729395427 |
| chr11 | 8686714 | 8686716 | + | 4.01584230749346 | 4.54798296825753 | 1.19322821036799 | 2.13421729395427 |
| chr11 | 8687972 | 8687973 | + | 3.01188173062009 | 1.13699574206438 | 2.38645642073598 | 5.33554323488568 |
| chr11 | 8689098 | 8689162 | + | 9.03564519186028 | 21.6029190992233 | 20.2848795762558 | 5.33554323488568 |
| chr11 | 8919958 | 8919968 | + | 22.087132691214 | 36.3838637460602 | 16.7051949451519 | 26.6777161744284 |
| chr11 | 8947795 | 8947817 | - | 6.02376346124019 | 10.2329616785794 | 2.38645642073598 | 1.06710864697714 |
| chr11 | 8947905 | 8947938 | - | 68.2693192273888 | 100.055625301666 | 21.4781077866238 | 27.7448248214055 |
| chr11 | 8948007 | 8948029 | - | 1.00396057687336 | 2.27399148412876 | 2.38645642073598 | 5.33554323488568 |
| chr11 | 8948247 | 8948262 | - | 16.0633692299738 | 4.54798296825753 | 7.15936926220794 | 2.13421729395427 |
| chr11 | 9138851 | 9138873 | - | 5.01980288436682 | 6.82197445238629 | 5.96614105183995 | 8.53686917581709 |
| chr11 | 9138907 | 9138936 | - | 10.0396057687336 | 5.68497871032191 | 7.15936926220794 | 12.8053037637256 |
| chr11 | 9138969 | 9138976 | - | 2.00792115374673 | 2.27399148412876 | 1.19322821036799 | 2.13421729395427 |
| chr11 | 9139094 | 9139117 | - | 1.00396057687336 | 3.41098722619315 | 2.38645642073598 | 3.20132594093141 |
| chr11 | 9139154 | 9139161 | - | 2.00792115374673 | 2.27399148412876 | 1.19322821036799 | 3.20132594093141 |
| chr11 | 9280810 | 9280825 | - | 4.01584230749346 | 1.13699574206438 | 1.19322821036799 | 5.33554323488568 |
| chr11 | 9281436 | 9281447 | - | 1.00396057687336 | 3.41098722619315 | 1.19322821036799 | 4.26843458790854 |
| chr11 | 9281652 | 9281662 | - | 4.01584230749346 | 2.27399148412876 | 3.57968463110397 | 2.13421729395427 |
| chr11 | 9282609 | 9282626 | - | 4.01584230749346 | 3.41098722619315 | 4.77291284147196 | 2.13421729395427 |
| chr11 | 9282654 | 9282662 | - | 12.0475269224804 | 56.8497871032191 | 17.8984231555198 | 18.1408469986113 |
| chr11 | 9283204 | 9283208 | - | 1.00396057687336 | 2.27399148412876 | 3.57968463110397 | 1.06710864697714 |
| chr11 | 9283249 | 9283270 | - | 2.00792115374673 | 4.54798296825753 | 5.96614105183995 | 17.0737383516342 |
| chr11 | 9445386 | 9445417 | + | 1078.25365956199 | 1010.78921469524 | 496.382935513084 | 884.633068344045 |
| chr11 | 9445648 | 9445692 | + | 5.01980288436682 | 6.82197445238629 | 1.19322821036799 | 3.20132594093141 |
| chr11 | 9445876 | 9445887 | + | 10.0396057687336 | 13.6439489047726 | 5.96614105183995 | 19.2079556455884 |
| chr11 | 9446162 | 9446181 | + | 2.00792115374673 | 9.09596593651506 | 1.19322821036799 | 3.20132594093141 |
| chr11 | 9446546 | 9446566 | + | 2.00792115374673 | 4.54798296825753 | 2.38645642073598 | 3.20132594093141 |
| chr11 | 9446680 | 9446686 | + | 5.01980288436682 | 14.780944646837 | 4.77291284147196 | 5.33554323488568 |
| chr11 | 9446830 | 9446837 | + | 7.02772403811355 | 12.5069531627082 | 2.38645642073598 | 10.6710864697714 |
| chr11 | 9446993 | 9447004 | + | 4.01584230749346 | 7.95897019445067 | 1.19322821036799 | 1.06710864697714 |
| chr11 | 9447064 | 9447092 | + | 10.0396057687336 | 15.9179403889013 | 7.15936926220794 | 10.6710864697714 |
| chr11 | 9447587 | 9447590 | + | 5.01980288436682 | 7.95897019445067 | 3.57968463110397 | 9.60397782279422 |
| chr11 | 9527950 | 9527952 | + | 3.01188173062009 | 1.13699574206438 | 4.77291284147196 | 2.13421729395427 |
| chr11 | 9588862 | 9588868 | + | 1.00396057687336 | 3.41098722619315 | 2.38645642073598 | 3.20132594093141 |
| chr11 | 9588974 | 9588975 | + | 2.00792115374673 | 2.27399148412876 | 1.19322821036799 | 2.13421729395427 |
| chr11 | 9589110 | 9589126 | + | 6.02376346124019 | 1.13699574206438 | 1.19322821036799 | 1.06710864697714 |
| chr11 | 9589232 | 9589266 | + | 2.00792115374673 | 10.2329616785794 | 5.96614105183995 | 13.8724124107028 |
| chr11 | 9589385 | 9589406 | + | 4.01584230749346 | 6.82197445238629 | 1.19322821036799 | 8.53686917581709 |
| chr11 | 9751692 | 9751693 | + | 2.00792115374673 | 1.13699574206438 | 1.19322821036799 | 1.06710864697714 |
| chr11 | 9758263 | 9758367 | + | 36.1425807674411 | 92.0966551072149 | 14.3187385244159 | 14.9395210576799 |
| chr11 | 10511988 | 10512008 | - | 1.00396057687336 | 2.27399148412876 | 2.38645642073598 | 4.26843458790854 |
| chr11 | 10512128 | 10512129 | - | 2.00792115374673 | 2.27399148412876 | 1.19322821036799 | 1.06710864697714 |
| chr11 | 10514824 | 10514883 | - | 5.01980288436682 | 6.82197445238629 | 4.77291284147196 | 5.33554323488568 |
| chr11 | 10779405 | 10779410 | + | 2.00792115374673 | 1.13699574206438 | 3.57968463110397 | 5.33554323488568 |
| chr11 | 10779549 | 10779550 | + | 3.01188173062009 | 1.13699574206438 | 2.38645642073598 | 1.06710864697714 |
| chr11 | 10779651 | 10779673 | + | 1.00396057687336 | 1.13699574206438 | 7.15936926220794 | 2.13421729395427 |
| chr11 | 10797082 | 10797257 | - | 143.566362492891 | 146.672450726305 | 60.8546387287675 | 74.6976052883995 |
| chr11 | 10797278 | 10797472 | - | 35.1386201905678 | 39.7948509722534 | 31.0239334695677 | 26.6777161744284 |
| chr11 | 10797502 | 10797552 | - | 14.0554480762271 | 19.3289276150945 | 5.96614105183995 | 18.1408469986113 |
| chr11 | 10797577 | 10797597 | - | 4.01584230749346 | 2.27399148412876 | 1.19322821036799 | 1.06710864697714 |
| chr11 | 10797808 | 10797852 | - | 11.043566345607 | 5.68497871032191 | 5.96614105183995 | 5.33554323488568 |
| chr11 | 10852829 | 10852857 | - | 11.043566345607 | 13.6439489047726 | 1.19322821036799 | 1.06710864697714 |
| chr11 | 11956179 | 11956204 | + | 13.0514874993537 | 19.3289276150945 | 5.96614105183995 | 6.40265188186281 |
| chr11 | 11956258 | 11956271 | + | 5.01980288436682 | 5.68497871032191 | 3.57968463110397 | 2.13421729395427 |
| chr11 | 11956379 | 11956404 | + | 1.00396057687336 | 12.5069531627082 | 3.57968463110397 | 3.20132594093141 |
| chr11 | 11956452 | 11956468 | + | 7.02772403811355 | 12.5069531627082 | 4.77291284147196 | 5.33554323488568 |
| chr11 | 11956705 | 11956723 | + | 4.01584230749346 | 2.27399148412876 | 5.96614105183995 | 5.33554323488568 |
| chr11 | 11958157 | 11958159 | + | 2.00792115374673 | 2.27399148412876 | 1.19322821036799 | 2.13421729395427 |
| chr11 | 12937499 | 12937577 | + | 42.1663442286813 | 34.1098722619315 | 2.38645642073598 | 4.26843458790854 |
| chr11 | 12937731 | 12937764 | + | 66.2613980736421 | 88.6856678810218 | 2.38645642073598 | 8.53686917581709 |
| chr11 | 12941767 | 12941802 | + | 4.01584230749346 | 7.95897019445067 | 1.19322821036799 | 1.06710864697714 |
| chr11 | 12942870 | 12942886 | + | 41.1623836518079 | 39.7948509722534 | 5.96614105183995 | 3.20132594093141 |
| chr11 | 12943644 | 12943669 | + | 27.1069355755808 | 20.4659233571589 | 3.57968463110397 | 6.40265188186281 |
| chr11 | 12944066 | 12944128 | + | 41.1623836518079 | 29.5618892936739 | 1.19322821036799 | 3.20132594093141 |
| chr11 | 12944230 | 12944248 | + | 38.1505019211879 | 52.3018041349616 | 1.19322821036799 | 2.13421729395427 |
| chr11 | 13729031 | 13729032 | + | 1.00396057687336 | 1.13699574206438 | 4.77291284147196 | 1.06710864697714 |
| chr11 | 13730712 | 13730724 | + | 6.02376346124019 | 2.27399148412876 | 4.77291284147196 | 2.13421729395427 |
| chr11 | 13731445 | 13731448 | + | 4.01584230749346 | 9.09596593651506 | 2.38645642073598 | 3.20132594093141 |
| chr11 | 13732292 | 13732293 | + | 6.02376346124019 | 5.68497871032191 | 8.35259747257593 | 8.53686917581709 |
| chr11 | 14457560 | 14457592 | - | 5.01980288436682 | 2.27399148412876 | 2.38645642073598 | 2.13421729395427 |
| chr11 | 14457647 | 14457693 | - | 4.01584230749346 | 2.27399148412876 | 4.77291284147196 | 5.33554323488568 |
| chr11 | 14457767 | 14457811 | - | 20.0792115374673 | 13.6439489047726 | 22.6713359969918 | 18.1408469986113 |
| chr11 | 14493780 | 14493812 | - | 3.01188173062009 | 2.27399148412876 | 8.35259747257593 | 5.33554323488568 |
| chr11 | 14494214 | 14494262 | - | 10.0396057687336 | 13.6439489047726 | 10.7390538933119 | 16.006629704657 |
| chr11 | 14494284 | 14494322 | - | 5.01980288436682 | 1.13699574206438 | 3.57968463110397 | 5.33554323488568 |
| chr11 | 14494354 | 14494382 | - | 2.00792115374673 | 4.54798296825753 | 3.57968463110397 | 4.26843458790854 |
| chr11 | 14494411 | 14494435 | - | 5.01980288436682 | 2.27399148412876 | 5.96614105183995 | 2.13421729395427 |
| chr11 | 14498914 | 14498924 | - | 7.02772403811355 | 5.68497871032191 | 13.1255103140479 | 8.53686917581709 |
| chr11 | 14504963 | 14504971 | - | 5.01980288436682 | 7.95897019445067 | 4.77291284147196 | 3.20132594093141 |
| chr11 | 14504993 | 14505033 | - | 9.03564519186028 | 10.2329616785794 | 9.54582568294392 | 11.7381951167485 |
| chr11 | 14505084 | 14505098 | - | 12.0475269224804 | 15.9179403889013 | 16.7051949451519 | 10.6710864697714 |
| chr11 | 14507667 | 14507758 | - | 68.2693192273888 | 86.411676396893 | 89.4921157775992 | 60.8251928776967 |
| chr11 | 14510871 | 14510954 | - | 50.1980288436682 | 37.5208594881246 | 119.322821036799 | 59.7580842307196 |
| chr11 | 14869981 | 14869987 | + | 5.01980288436682 | 6.82197445238629 | 1.19322821036799 | 1.06710864697714 |
| chr11 | 14870032 | 14870044 | + | 13.0514874993537 | 5.68497871032191 | 2.38645642073598 | 2.13421729395427 |
| chr11 | 14870224 | 14870262 | + | 6.02376346124019 | 3.41098722619315 | 3.57968463110397 | 5.33554323488568 |
| chr11 | 14870998 | 14871040 | + | 2.00792115374673 | 2.27399148412876 | 1.19322821036799 | 2.13421729395427 |
| chr11 | 15966580 | 15966601 | - | 2.00792115374673 | 3.41098722619315 | 44.1494437836156 | 21.3421729395427 |
| chr11 | 15967888 | 15967895 | - | 5.01980288436682 | 7.95897019445067 | 44.1494437836156 | 43.7514545260626 |
| chr11 | 15969106 | 15969109 | - | 5.01980288436682 | 2.27399148412876 | 7.15936926220794 | 3.20132594093141 |
| chr11 | 16754868 | 16754877 | + | 17.0673298068472 | 19.3289276150945 | 16.7051949451519 | 16.006629704657 |
| chr11 | 16754924 | 16754974 | + | 30.1188173062009 | 25.0139063254164 | 15.5119667347839 | 10.6710864697714 |
| chr11 | 16755137 | 16755165 | + | 26.1029749987075 | 36.3838637460602 | 33.4103898903037 | 37.3488026441998 |
| chr11 | 16755290 | 16755331 | + | 11.043566345607 | 37.5208594881246 | 23.8645642073598 | 11.7381951167485 |
| chr11 | 16755452 | 16755470 | + | 22.087132691214 | 38.657855230189 | 13.1255103140479 | 5.33554323488568 |
| chr11 | 16755499 | 16755511 | + | 43.1703048055547 | 37.5208594881246 | 13.1255103140479 | 34.1474767032683 |
| chr11 | 16755594 | 16755595 | + | 4.01584230749346 | 4.54798296825753 | 1.19322821036799 | 2.13421729395427 |
| chr11 | 16755698 | 16755724 | + | 5.01980288436682 | 3.41098722619315 | 3.57968463110397 | 2.13421729395427 |
| chr11 | 16755754 | 16755774 | + | 1.00396057687336 | 3.41098722619315 | 3.57968463110397 | 1.06710864697714 |
| chr11 | 17087315 | 17087338 | - | 4.01584230749346 | 4.54798296825753 | 1.19322821036799 | 2.13421729395427 |
| chr11 | 17087894 | 17087942 | - | 1.00396057687336 | 22.7399148412876 | 4.77291284147196 | 7.46976052883995 |
| chr11 | 17331421 | 17331451 | + | 10.0396057687336 | 7.95897019445067 | 14.3187385244159 | 17.0737383516342 |
| chr11 | 17372582 | 17372592 | + | 1.00396057687336 | 1.13699574206438 | 1.19322821036799 | 1.06710864697714 |
| chr11 | 17376772 | 17376793 | + | 1.00396057687336 | 1.13699574206438 | 2.38645642073598 | 1.06710864697714 |
| chr11 | 17377132 | 17377134 | + | 5.01980288436682 | 1.13699574206438 | 3.57968463110397 | 4.26843458790854 |
| chr11 | 17381287 | 17381332 | + | 1.00396057687336 | 1.13699574206438 | 3.57968463110397 | 3.20132594093141 |
| chr11 | 17788080 | 17788097 | - | 1.00396057687336 | 9.09596593651506 | 2.38645642073598 | 5.33554323488568 |
| chr11 | 18365870 | 18365874 | + | 1.00396057687336 | 2.27399148412876 | 1.19322821036799 | 2.13421729395427 |
| chr11 | 18366047 | 18366069 | + | 3.01188173062009 | 3.41098722619315 | 5.96614105183995 | 4.26843458790854 |
| chr11 | 18366580 | 18366601 | + | 2.00792115374673 | 3.41098722619315 | 1.19322821036799 | 3.20132594093141 |
| chr11 | 18366753 | 18366756 | + | 2.00792115374673 | 9.09596593651506 | 2.38645642073598 | 1.06710864697714 |
| chr11 | 18407185 | 18407333 | + | 52.205949997415 | 29.5618892936739 | 89.4921157775992 | 128.053037637256 |
| chr11 | 18407395 | 18407433 | + | 74.293082688629 | 71.6307317500561 | 73.9801490428153 | 90.7042349930565 |
| chr11 | 18407469 | 18407557 | + | 68.2693192273888 | 40.9318467143178 | 66.8207797806074 | 102.442430109805 |
| chr11 | 18407627 | 18407689 | + | 103.407939417957 | 112.562578464374 | 159.892580189311 | 146.193884635868 |
| chr11 | 18407756 | 18407782 | + | 15.0594086531005 | 28.4248935516096 | 22.6713359969918 | 8.53686917581709 |
| chr11 | 18407824 | 18407846 | + | 1.00396057687336 | 1.13699574206438 | 2.38645642073598 | 1.06710864697714 |
| chr11 | 18480332 | 18480350 | - | 19.0752509605939 | 32.9728765198671 | 9.54582568294392 | 21.3421729395427 |
| chr11 | 18480373 | 18480402 | - | 3.01188173062009 | 7.95897019445067 | 3.57968463110397 | 5.33554323488568 |
| chr11 | 18480428 | 18480570 | - | 81.3208067267425 | 128.480518853275 | 71.5936926220794 | 84.3015831111937 |
| chr11 | 18532278 | 18532299 | - | 1.00396057687336 | 2.27399148412876 | 2.38645642073598 | 1.06710864697714 |
| chr11 | 18608099 | 18608102 | - | 1.00396057687336 | 1.13699574206438 | 2.38645642073598 | 4.26843458790854 |
| chr11 | 18609681 | 18609686 | - | 5.01980288436682 | 1.13699574206438 | 1.19322821036799 | 6.40265188186281 |
| chr11 | 19224216 | 19224218 | - | 1.00396057687336 | 11.3699574206438 | 2.38645642073598 | 1.06710864697714 |
| chr11 | 20121048 | 20121127 | + | 67.2653586505154 | 160.316399631078 | 3.57968463110397 | 1.06710864697714 |
| chr11 | 20383454 | 20383456 | + | 2.00792115374673 | 5.68497871032191 | 3.57968463110397 | 1.06710864697714 |
| chr11 | 20383515 | 20383538 | + | 64.2534769198953 | 64.8087572976698 | 10.7390538933119 | 18.1408469986113 |
| chr11 | 20383709 | 20383766 | + | 99.3920971104631 | 168.275369825529 | 36.9900745214077 | 37.3488026441998 |
| chr11 | 20508438 | 20508479 | + | 2.00792115374673 | 5.68497871032191 | 3.57968463110397 | 6.40265188186281 |
| chr11 | 20508718 | 20508726 | + | 2.00792115374673 | 3.41098722619315 | 1.19322821036799 | 10.6710864697714 |
| chr11 | 20508843 | 20508850 | + | 13.0514874993537 | 23.876910583352 | 5.96614105183995 | 23.476390233497 |
| chr11 | 20508880 | 20508881 | + | 1.00396057687336 | 1.13699574206438 | 1.19322821036799 | 2.13421729395427 |
| chr11 | 22820751 | 22820800 | - | 25.0990144218341 | 20.4659233571589 | 36.9900745214077 | 33.0803680562912 |
| chr11 | 22820885 | 22820887 | - | 1.00396057687336 | 3.41098722619315 | 3.57968463110397 | 2.13421729395427 |
| chr11 | 22820973 | 22820978 | - | 1.00396057687336 | 7.95897019445067 | 8.35259747257593 | 8.53686917581709 |
| chr11 | 22821179 | 22821201 | - | 23.0910932680874 | 23.876910583352 | 13.1255103140479 | 21.3421729395427 |
| chr11 | 22821233 | 22821290 | - | 51.2019894205416 | 75.0417189762492 | 66.8207797806074 | 89.6371263460794 |
| chr11 | 22821459 | 22821461 | - | 2.00792115374673 | 1.13699574206438 | 1.19322821036799 | 2.13421729395427 |
| chr11 | 22821693 | 22821714 | - | 8.03168461498692 | 1.13699574206438 | 4.77291284147196 | 9.60397782279422 |
| chr11 | 22822615 | 22822630 | - | 2.00792115374673 | 4.54798296825753 | 9.54582568294392 | 9.60397782279422 |
| chr11 | 22822889 | 22822894 | - | 2.00792115374673 | 3.41098722619315 | 1.19322821036799 | 3.20132594093141 |
| chr11 | 22822940 | 22822953 | - | 2.00792115374673 | 5.68497871032191 | 2.38645642073598 | 11.7381951167485 |
| chr11 | 22823058 | 22823061 | - | 2.00792115374673 | 3.41098722619315 | 2.38645642073598 | 5.33554323488568 |
| chr11 | 22829410 | 22829424 | + | 17.0673298068472 | 23.876910583352 | 9.54582568294392 | 5.33554323488568 |
| chr11 | 27338594 | 27338605 | - | 10.0396057687336 | 7.95897019445067 | 4.77291284147196 | 6.40265188186281 |
| chr11 | 27338670 | 27338693 | - | 13.0514874993537 | 3.41098722619315 | 7.15936926220794 | 3.20132594093141 |
| chr11 | 27366210 | 27366269 | - | 15.0594086531005 | 12.5069531627082 | 1.19322821036799 | 3.20132594093141 |
| chr11 | 27366426 | 27366439 | - | 16.0633692299738 | 6.82197445238629 | 1.19322821036799 | 1.06710864697714 |
| chr11 | 27494546 | 27494582 | - | 4.01584230749346 | 7.95897019445067 | 7.15936926220794 | 1.06710864697714 |
| chr11 | 27494661 | 27494681 | - | 7.02772403811355 | 13.6439489047726 | 1.19322821036799 | 3.20132594093141 |
| chr11 | 27494706 | 27494754 | - | 31.1227778830743 | 35.2468680039958 | 25.0577924177278 | 12.8053037637256 |
| chr11 | 27494782 | 27494801 | - | 10.0396057687336 | 13.6439489047726 | 5.96614105183995 | 5.33554323488568 |
| chr11 | 27494856 | 27494880 | - | 6.02376346124019 | 2.27399148412876 | 4.77291284147196 | 1.06710864697714 |
| chr11 | 27494962 | 27495018 | - | 25.0990144218341 | 30.6988850357383 | 17.8984231555198 | 14.9395210576799 |
| chr11 | 27495044 | 27495074 | - | 9.03564519186028 | 13.6439489047726 | 2.38645642073598 | 1.06710864697714 |
| chr11 | 27495769 | 27495816 | - | 64.2534769198953 | 73.9047232341848 | 36.9900745214077 | 30.9461507623369 |
| chr11 | 27495846 | 27495849 | - | 16.0633692299738 | 22.7399148412876 | 1.19322821036799 | 5.33554323488568 |
| chr11 | 27496042 | 27496056 | - | 5.01980288436682 | 1.13699574206438 | 2.38645642073598 | 5.33554323488568 |
| chr11 | 27496121 | 27496141 | - | 4.01584230749346 | 5.68497871032191 | 1.19322821036799 | 1.06710864697714 |
| chr11 | 27496182 | 27496206 | - | 8.03168461498692 | 25.0139063254164 | 3.57968463110397 | 7.46976052883995 |
| chr11 | 27497047 | 27497065 | - | 3.01188173062009 | 5.68497871032191 | 1.19322821036799 | 1.06710864697714 |
| chr11 | 27497094 | 27497108 | - | 8.03168461498692 | 7.95897019445067 | 3.57968463110397 | 14.9395210576799 |
| chr11 | 27497185 | 27497263 | - | 23.0910932680874 | 26.1509020674808 | 14.3187385244159 | 9.60397782279422 |
| chr11 | 27497311 | 27497321 | - | 4.01584230749346 | 5.68497871032191 | 1.19322821036799 | 1.06710864697714 |
| chr11 | 27497386 | 27497393 | - | 11.043566345607 | 21.6029190992233 | 2.38645642073598 | 5.33554323488568 |
| chr11 | 27497716 | 27497750 | - | 8.03168461498692 | 13.6439489047726 | 3.57968463110397 | 6.40265188186281 |
| chr11 | 27497857 | 27497868 | - | 4.01584230749346 | 4.54798296825753 | 1.19322821036799 | 2.13421729395427 |
| chr11 | 27498266 | 27498285 | - | 7.02772403811355 | 5.68497871032191 | 1.19322821036799 | 2.13421729395427 |
| chr11 | 27498363 | 27498448 | - | 252.998065372088 | 352.468680039958 | 190.916513658878 | 97.1068868749194 |
| chr11 | 27498478 | 27498494 | - | 14.0554480762271 | 5.68497871032191 | 13.1255103140479 | 11.7381951167485 |
| chr11 | 27498566 | 27498626 | - | 10.0396057687336 | 14.780944646837 | 15.5119667347839 | 6.40265188186281 |
| chr11 | 27506836 | 27506874 | + | 32.1267384599477 | 38.657855230189 | 13.1255103140479 | 7.46976052883995 |
| chr11 | 30337003 | 30337030 | + | 7.02772403811355 | 4.54798296825753 | 4.77291284147196 | 5.33554323488568 |
| chr11 | 30337112 | 30337118 | + | 1.00396057687336 | 2.27399148412876 | 2.38645642073598 | 6.40265188186281 |
| chr11 | 30338063 | 30338099 | + | 6.02376346124019 | 12.5069531627082 | 5.96614105183995 | 10.6710864697714 |
| chr11 | 31431633 | 31431639 | + | 1.00396057687336 | 2.27399148412876 | 5.96614105183995 | 4.26843458790854 |
| chr11 | 31432431 | 31432433 | - | 21.0831721143407 | 52.3018041349616 | 8.35259747257593 | 16.006629704657 |
| chr11 | 31783468 | 31783506 | + | 4.01584230749346 | 6.82197445238629 | 14.3187385244159 | 6.40265188186281 |
| chr11 | 31783524 | 31783548 | + | 6.02376346124019 | 7.95897019445067 | 1.19322821036799 | 6.40265188186281 |
| chr11 | 31783651 | 31783652 | + | 3.01188173062009 | 1.13699574206438 | 4.77291284147196 | 6.40265188186281 |
| chr11 | 31783684 | 31783697 | + | 5.01980288436682 | 6.82197445238629 | 7.15936926220794 | 14.9395210576799 |
| chr11 | 31783730 | 31783733 | + | 1.00396057687336 | 2.27399148412876 | 2.38645642073598 | 4.26843458790854 |
| chr11 | 32104678 | 32104696 | + | 18.0712903837206 | 19.3289276150945 | 2.38645642073598 | 1.06710864697714 |
| chr11 | 32104722 | 32104729 | + | 14.0554480762271 | 11.3699574206438 | 1.19322821036799 | 1.06710864697714 |
| chr11 | 32602354 | 32602393 | - | 33.130699036821 | 55.7127913611547 | 33.4103898903037 | 75.7647139353766 |
| chr11 | 32602426 | 32602437 | + | 12.0475269224804 | 17.0549361309657 | 13.1255103140479 | 18.1408469986113 |
| chr11 | 32976527 | 32976532 | + | 5.01980288436682 | 5.68497871032191 | 1.19322821036799 | 3.20132594093141 |
| chr11 | 32976622 | 32976637 | + | 2.00792115374673 | 2.27399148412876 | 2.38645642073598 | 4.26843458790854 |
| chr11 | 32976655 | 32976663 | + | 3.01188173062009 | 1.13699574206438 | 3.57968463110397 | 1.06710864697714 |
| chr11 | 32976742 | 32976775 | + | 2.00792115374673 | 2.27399148412876 | 3.57968463110397 | 26.6777161744284 |
| chr11 | 32977676 | 32977687 | + | 1.00396057687336 | 1.13699574206438 | 4.77291284147196 | 4.26843458790854 |
| chr11 | 32978316 | 32978337 | + | 1.00396057687336 | 4.54798296825753 | 4.77291284147196 | 2.13421729395427 |
| chr11 | 32978704 | 32978711 | + | 4.01584230749346 | 1.13699574206438 | 11.9322821036799 | 2.13421729395427 |
| chr11 | 32979139 | 32979143 | + | 3.01188173062009 | 1.13699574206438 | 1.19322821036799 | 4.26843458790854 |
| chr11 | 32979217 | 32979220 | + | 3.01188173062009 | 1.13699574206438 | 5.96614105183995 | 1.06710864697714 |
| chr11 | 32979299 | 32979301 | + | 1.00396057687336 | 1.13699574206438 | 1.19322821036799 | 1.06710864697714 |
| chr11 | 32979375 | 32979381 | + | 3.01188173062009 | 1.13699574206438 | 1.19322821036799 | 1.06710864697714 |
| chr11 | 33033532 | 33033558 | + | 18.0712903837206 | 23.876910583352 | 4.77291284147196 | 4.26843458790854 |
| chr11 | 33141425 | 33141432 | - | 5.01980288436682 | 7.95897019445067 | 1.19322821036799 | 1.06710864697714 |
| chr11 | 33141461 | 33141463 | - | 6.02376346124019 | 2.27399148412876 | 1.19322821036799 | 1.06710864697714 |
| chr11 | 33141526 | 33141534 | - | 4.01584230749346 | 11.3699574206438 | 9.54582568294392 | 3.20132594093141 |
| chr11 | 33141666 | 33141700 | - | 19.0752509605939 | 15.9179403889013 | 27.4442488384638 | 25.6106075274513 |
| chr11 | 33141725 | 33141743 | - | 10.0396057687336 | 4.54798296825753 | 4.77291284147196 | 11.7381951167485 |
| chr11 | 33161622 | 33161690 | + | 6.02376346124019 | 19.3289276150945 | 36.9900745214077 | 40.5501285851312 |
| chr11 | 33353959 | 33354007 | + | 7.02772403811355 | 13.6439489047726 | 2.38645642073598 | 1.06710864697714 |
| chr11 | 33354138 | 33354220 | + | 13.0514874993537 | 13.6439489047726 | 8.35259747257593 | 4.26843458790854 |
| chr11 | 33354239 | 33354259 | + | 8.03168461498692 | 7.95897019445067 | 7.15936926220794 | 1.06710864697714 |
| chr11 | 33354779 | 33354785 | + | 8.03168461498692 | 6.82197445238629 | 4.77291284147196 | 19.2079556455884 |
| chr11 | 33356806 | 33356821 | + | 1.00396057687336 | 3.41098722619315 | 1.19322821036799 | 4.26843458790854 |
| chr11 | 33356907 | 33356951 | + | 3.01188173062009 | 5.68497871032191 | 1.19322821036799 | 1.06710864697714 |
| chr11 | 33670748 | 33670749 | + | 8.03168461498692 | 3.41098722619315 | 4.77291284147196 | 3.20132594093141 |
| chr11 | 33708684 | 33708825 | - | 66.2613980736421 | 88.6856678810218 | 16.7051949451519 | 27.7448248214055 |
| chr11 | 33708890 | 33708969 | - | 92.3643730723495 | 161.453395373142 | 27.4442488384638 | 32.0132594093141 |
| chr11 | 33709044 | 33709048 | - | 80.3168461498692 | 81.8636934286355 | 17.8984231555198 | 35.2145853502455 |
| chr11 | 33709198 | 33709205 | - | 40.1584230749346 | 37.5208594881246 | 3.57968463110397 | 6.40265188186281 |
| chr11 | 33709264 | 33709270 | - | 3.01188173062009 | 4.54798296825753 | 2.38645642073598 | 1.06710864697714 |
| chr11 | 33709444 | 33709513 | - | 75.2970432655023 | 112.562578464374 | 8.35259747257593 | 17.0737383516342 |
| chr11 | 33709532 | 33709647 | - | 76.3010038423757 | 65.9457530397342 | 15.5119667347839 | 29.8790421153598 |
| chr11 | 33709730 | 33709753 | - | 18.0712903837206 | 9.09596593651506 | 3.57968463110397 | 8.53686917581709 |
| chr11 | 33710015 | 33710056 | - | 96.380215379843 | 330.865760940735 | 38.1833027317757 | 34.1474767032683 |
| chr11 | 33710077 | 33710116 | - | 49.1940682667949 | 52.3018041349616 | 5.96614105183995 | 20.2750642925656 |
| chr11 | 33710147 | 33710169 | - | 4.01584230749346 | 1.13699574206438 | 1.19322821036799 | 1.06710864697714 |
| chr11 | 33710197 | 33710251 | - | 117.463387494184 | 110.288586980245 | 28.6374770488317 | 41.6172372321083 |
| chr11 | 33710272 | 33710346 | - | 96.380215379843 | 65.9457530397342 | 16.7051949451519 | 27.7448248214055 |
| chr11 | 33741703 | 33741720 | - | 4.01584230749346 | 1.13699574206438 | 3.57968463110397 | 4.26843458790854 |
| chr11 | 33741959 | 33741964 | - | 5.01980288436682 | 9.09596593651506 | 2.38645642073598 | 6.40265188186281 |
| chr11 | 33742001 | 33742064 | - | 8.03168461498692 | 4.54798296825753 | 3.57968463110397 | 6.40265188186281 |
| chr11 | 33744369 | 33744416 | - | 16.0633692299738 | 7.95897019445067 | 8.35259747257593 | 14.9395210576799 |
| chr11 | 33744593 | 33744603 | - | 7.02772403811355 | 3.41098722619315 | 3.57968463110397 | 8.53686917581709 |
| chr11 | 33746997 | 33747000 | - | 15.0594086531005 | 14.780944646837 | 13.1255103140479 | 12.8053037637256 |
| chr11 | 34097695 | 34097797 | + | 30.1188173062009 | 14.780944646837 | 25.0577924177278 | 14.9395210576799 |
| chr11 | 34097817 | 34097821 | + | 4.01584230749346 | 1.13699574206438 | 7.15936926220794 | 2.13421729395427 |
| chr11 | 34098548 | 34098559 | + | 2.00792115374673 | 6.82197445238629 | 3.57968463110397 | 5.33554323488568 |
| chr11 | 34098632 | 34098646 | + | 4.01584230749346 | 2.27399148412876 | 4.77291284147196 | 3.20132594093141 |
| chr11 | 34098677 | 34098699 | + | 43.1703048055547 | 42.0688424563821 | 38.1833027317757 | 13.8724124107028 |
| chr11 | 34098826 | 34098875 | + | 5.01980288436682 | 6.82197445238629 | 20.2848795762558 | 11.7381951167485 |
| chr11 | 34099432 | 34099477 | + | 7.02772403811355 | 6.82197445238629 | 15.5119667347839 | 10.6710864697714 |
| chr11 | 34099652 | 34099662 | + | 6.02376346124019 | 12.5069531627082 | 21.4781077866238 | 13.8724124107028 |
| chr11 | 34099696 | 34099706 | + | 6.02376346124019 | 3.41098722619315 | 5.96614105183995 | 1.06710864697714 |
| chr11 | 34099785 | 34099805 | + | 24.0950538449607 | 10.2329616785794 | 19.0916513658878 | 9.60397782279422 |
| chr11 | 34099846 | 34099861 | + | 8.03168461498692 | 3.41098722619315 | 5.96614105183995 | 8.53686917581709 |
| chr11 | 34099897 | 34100328 | + | 350.382241328804 | 227.399148412876 | 387.799168369597 | 221.958598571244 |
| chr11 | 34100439 | 34100482 | + | 6.02376346124019 | 7.95897019445067 | 2.38645642073598 | 7.46976052883995 |
| chr11 | 34100704 | 34100777 | + | 45.1782259593014 | 27.2878978095452 | 14.3187385244159 | 36.2816939972226 |
| chr11 | 34100881 | 34100888 | + | 2.00792115374673 | 2.27399148412876 | 2.38645642073598 | 7.46976052883995 |
| chr11 | 34146082 | 34146148 | + | 15.0594086531005 | 6.82197445238629 | 36.9900745214077 | 22.4092815865198 |
| chr11 | 34146176 | 34146192 | + | 2.00792115374673 | 1.13699574206438 | 10.7390538933119 | 2.13421729395427 |
| chr11 | 34146324 | 34146408 | + | 16.0633692299738 | 11.3699574206438 | 26.2510206280958 | 28.8119334683827 |
| chr11 | 34146448 | 34146475 | + | 2.00792115374673 | 3.41098722619315 | 17.8984231555198 | 12.8053037637256 |
| chr11 | 34146638 | 34146693 | + | 17.0673298068472 | 12.5069531627082 | 4.77291284147196 | 6.40265188186281 |
| chr11 | 34146821 | 34146876 | + | 45.1782259593014 | 76.1787147183136 | 45.3426719939836 | 65.0936274656053 |
| chr11 | 34471408 | 34471503 | + | 2.00792115374673 | 1.13699574206438 | 1.19322821036799 | 6.40265188186281 |
| chr11 | 34471605 | 34471624 | + | 37.1465413443145 | 54.5757956190903 | 9.54582568294392 | 25.6106075274513 |
| chr11 | 34471659 | 34471664 | + | 5.01980288436682 | 4.54798296825753 | 2.38645642073598 | 3.20132594093141 |
| chr11 | 34471693 | 34471711 | + | 2.00792115374673 | 7.95897019445067 | 9.54582568294392 | 5.33554323488568 |
| chr11 | 34471767 | 34471823 | + | 9.03564519186028 | 10.2329616785794 | 4.77291284147196 | 9.60397782279422 |
| chr11 | 34471896 | 34471946 | + | 16.0633692299738 | 11.3699574206438 | 3.57968463110397 | 6.40265188186281 |
| chr11 | 34471990 | 34472007 | + | 4.01584230749346 | 13.6439489047726 | 4.77291284147196 | 4.26843458790854 |
| chr11 | 34882392 | 34882401 | - | 6.02376346124019 | 9.09596593651506 | 11.9322821036799 | 14.9395210576799 |
| chr11 | 34882524 | 34882526 | - | 1.00396057687336 | 3.41098722619315 | 5.96614105183995 | 5.33554323488568 |
| chr11 | 34995315 | 34995333 | + | 1.00396057687336 | 1.13699574206438 | 2.38645642073598 | 1.06710864697714 |
| chr11 | 34995568 | 34995609 | + | 2.00792115374673 | 3.41098722619315 | 5.96614105183995 | 7.46976052883995 |
| chr11 | 34995759 | 34995765 | + | 5.01980288436682 | 13.6439489047726 | 9.54582568294392 | 6.40265188186281 |
| chr11 | 34995914 | 34995933 | + | 3.01188173062009 | 3.41098722619315 | 5.96614105183995 | 7.46976052883995 |
| chr11 | 35229388 | 35229449 | + | 22.087132691214 | 12.5069531627082 | 1.19322821036799 | 1.06710864697714 |
| chr11 | 35806359 | 35806405 | + | 3.01188173062009 | 4.54798296825753 | 2.38645642073598 | 4.26843458790854 |
| chr11 | 35806624 | 35806625 | + | 4.01584230749346 | 4.54798296825753 | 1.19322821036799 | 1.06710864697714 |
| chr11 | 35806828 | 35806859 | + | 13.0514874993537 | 5.68497871032191 | 20.2848795762558 | 11.7381951167485 |
| chr11 | 35806939 | 35806960 | + | 5.01980288436682 | 6.82197445238629 | 4.77291284147196 | 1.06710864697714 |
| chr11 | 35807043 | 35807079 | + | 14.0554480762271 | 13.6439489047726 | 5.96614105183995 | 4.26843458790854 |
| chr11 | 35807906 | 35807928 | + | 104.41189999483 | 70.4937360079917 | 52.5020412561915 | 67.2278447595595 |
| chr11 | 35807958 | 35808199 | + | 52.205949997415 | 52.3018041349616 | 164.665493030783 | 25.6106075274513 |
| chr11 | 35808349 | 35808372 | + | 1.00396057687336 | 1.13699574206438 | 4.77291284147196 | 5.33554323488568 |
| chr11 | 35810888 | 35810925 | + | 8.03168461498692 | 11.3699574206438 | 3.57968463110397 | 6.40265188186281 |
| chr11 | 35810990 | 35811026 | + | 9.03564519186028 | 7.95897019445067 | 4.77291284147196 | 8.53686917581709 |
| chr11 | 36273980 | 36274005 | - | 26.1029749987075 | 29.5618892936739 | 11.9322821036799 | 12.8053037637256 |
| chr11 | 36274069 | 36274070 | - | 4.01584230749346 | 3.41098722619315 | 2.38645642073598 | 3.20132594093141 |
| chr11 | 36274349 | 36274357 | - | 10.0396057687336 | 6.82197445238629 | 2.38645642073598 | 8.53686917581709 |
| chr11 | 36274413 | 36274416 | - | 1.00396057687336 | 1.13699574206438 | 2.38645642073598 | 2.13421729395427 |
| chr11 | 36510351 | 36510371 | + | 1.00396057687336 | 1.13699574206438 | 5.96614105183995 | 5.33554323488568 |
| chr11 | 36659195 | 36659214 | + | 5.01980288436682 | 7.95897019445067 | 3.57968463110397 | 6.40265188186281 |
| chr11 | 43342493 | 43342495 | + | 9.03564519186028 | 2.27399148412876 | 10.7390538933119 | 3.20132594093141 |
| chr11 | 43342708 | 43342714 | + | 4.01584230749346 | 2.27399148412876 | 2.38645642073598 | 3.20132594093141 |
| chr11 | 43342868 | 43342873 | + | 2.00792115374673 | 1.13699574206438 | 8.35259747257593 | 3.20132594093141 |
| chr11 | 43343311 | 43343312 | + | 1.00396057687336 | 2.27399148412876 | 1.19322821036799 | 2.13421729395427 |
| chr11 | 43343501 | 43343513 | + | 3.01188173062009 | 1.13699574206438 | 4.77291284147196 | 5.33554323488568 |
| chr11 | 43344329 | 43344337 | + | 7.02772403811355 | 3.41098722619315 | 8.35259747257593 | 3.20132594093141 |
| chr11 | 43344482 | 43344496 | + | 6.02376346124019 | 6.82197445238629 | 9.54582568294392 | 3.20132594093141 |
| chr11 | 43494023 | 43494046 | + | 4.01584230749346 | 4.54798296825753 | 5.96614105183995 | 5.33554323488568 |
| chr11 | 43494151 | 43494188 | + | 3.01188173062009 | 2.27399148412876 | 1.19322821036799 | 2.13421729395427 |
| chr11 | 43494255 | 43494265 | + | 2.00792115374673 | 1.13699574206438 | 3.57968463110397 | 2.13421729395427 |
| chr11 | 43494795 | 43494819 | + | 6.02376346124019 | 5.68497871032191 | 1.19322821036799 | 5.33554323488568 |
| chr11 | 43494888 | 43494902 | + | 3.01188173062009 | 3.41098722619315 | 1.19322821036799 | 1.06710864697714 |
| chr11 | 43840028 | 43840053 | + | 2.00792115374673 | 4.54798296825753 | 1.19322821036799 | 1.06710864697714 |
| chr11 | 43855496 | 43855502 | + | 1.00396057687336 | 1.13699574206438 | 1.19322821036799 | 1.06710864697714 |
| chr11 | 43856010 | 43856033 | + | 3.01188173062009 | 1.13699574206438 | 1.19322821036799 | 3.20132594093141 |
| chr11 | 43856532 | 43856590 | + | 1.00396057687336 | 2.27399148412876 | 8.35259747257593 | 3.20132594093141 |
| chr11 | 44244324 | 44244336 | + | 4.01584230749346 | 1.13699574206438 | 7.15936926220794 | 10.6710864697714 |
| chr11 | 44244363 | 44244387 | + | 7.02772403811355 | 11.3699574206438 | 17.8984231555198 | 9.60397782279422 |
| chr11 | 44244470 | 44244514 | + | 3.01188173062009 | 1.13699574206438 | 2.38645642073598 | 2.13421729395427 |
| chr11 | 44244812 | 44244896 | + | 2.00792115374673 | 3.41098722619315 | 3.57968463110397 | 5.33554323488568 |
| chr11 | 44245260 | 44245285 | + | 4.01584230749346 | 3.41098722619315 | 1.19322821036799 | 1.06710864697714 |
| chr11 | 45812287 | 45812309 | + | 4.01584230749346 | 2.27399148412876 | 5.96614105183995 | 3.20132594093141 |
| chr11 | 45812704 | 45812707 | + | 1.00396057687336 | 1.13699574206438 | 2.38645642073598 | 1.06710864697714 |
| chr11 | 45812825 | 45812837 | + | 5.01980288436682 | 3.41098722619315 | 3.57968463110397 | 5.33554323488568 |
| chr11 | 45812898 | 45812907 | + | 68.2693192273888 | 15.9179403889013 | 11.9322821036799 | 33.0803680562912 |
| chr11 | 45812969 | 45812987 | + | 2.00792115374673 | 4.54798296825753 | 1.19322821036799 | 1.06710864697714 |
| chr11 | 45883173 | 45883182 | + | 3.01188173062009 | 5.68497871032191 | 4.77291284147196 | 10.6710864697714 |
| chr11 | 45932730 | 45932753 | - | 1.00396057687336 | 11.3699574206438 | 3.57968463110397 | 1.06710864697714 |
| chr11 | 45933007 | 45933029 | - | 1.00396057687336 | 5.68497871032191 | 1.19322821036799 | 3.20132594093141 |
| chr11 | 46320842 | 46320894 | + | 4.01584230749346 | 2.27399148412876 | 63.2410951495034 | 26.6777161744284 |
| chr11 | 46320915 | 46320923 | + | 1.00396057687336 | 2.27399148412876 | 11.9322821036799 | 12.8053037637256 |
| chr11 | 46379971 | 46379989 | + | 4.01584230749346 | 3.41098722619315 | 3.57968463110397 | 3.20132594093141 |
| chr11 | 46380472 | 46380531 | + | 41.1623836518079 | 52.3018041349616 | 34.6036181006717 | 49.0869977609482 |
| chr11 | 46383661 | 46383799 | + | 1.00396057687336 | 2.27399148412876 | 124.095733878271 | 234.76390233497 |
| chr11 | 46396455 | 46396488 | - | 11.043566345607 | 19.3289276150945 | 7.15936926220794 | 6.40265188186281 |
| chr11 | 46396664 | 46396671 | - | 2.00792115374673 | 1.13699574206438 | 1.19322821036799 | 1.06710864697714 |
| chr11 | 46672432 | 46672464 | + | 5.01980288436682 | 6.82197445238629 | 10.7390538933119 | 2.13421729395427 |
| chr11 | 46672959 | 46673026 | + | 12.0475269224804 | 11.3699574206438 | 10.7390538933119 | 4.26843458790854 |
| chr11 | 46673138 | 46673174 | + | 3.01188173062009 | 3.41098722619315 | 9.54582568294392 | 14.9395210576799 |
| chr11 | 46677128 | 46677216 | - | 14.0554480762271 | 17.0549361309657 | 5.96614105183995 | 18.1408469986113 |
| chr11 | 46677246 | 46677258 | - | 7.02772403811355 | 4.54798296825753 | 1.19322821036799 | 3.20132594093141 |
| chr11 | 46677615 | 46677632 | - | 7.02772403811355 | 4.54798296825753 | 4.77291284147196 | 8.53686917581709 |
| chr11 | 46677661 | 46677678 | - | 9.03564519186028 | 9.09596593651506 | 13.1255103140479 | 11.7381951167485 |
| chr11 | 46677966 | 46678019 | - | 7.02772403811355 | 3.41098722619315 | 4.77291284147196 | 11.7381951167485 |
| chr11 | 46743570 | 46743596 | - | 4.01584230749346 | 2.27399148412876 | 1.19322821036799 | 6.40265188186281 |
| chr11 | 46743619 | 46743622 | - | 15.0594086531005 | 10.2329616785794 | 9.54582568294392 | 17.0737383516342 |
| chr11 | 46743760 | 46743820 | - | 12.0475269224804 | 9.09596593651506 | 8.35259747257593 | 9.60397782279422 |
| chr11 | 46743865 | 46744025 | - | 309.219857676996 | 241.043097317649 | 230.293044601022 | 312.662833564301 |
| chr11 | 47161543 | 47161559 | + | 31.1227778830743 | 21.6029190992233 | 9.54582568294392 | 2.13421729395427 |
| chr11 | 47162078 | 47162148 | + | 16.0633692299738 | 11.3699574206438 | 1.19322821036799 | 5.33554323488568 |
| chr11 | 47164321 | 47164353 | - | 5.01980288436682 | 13.6439489047726 | 15.5119667347839 | 12.8053037637256 |
| chr11 | 47164376 | 47164396 | - | 3.01188173062009 | 9.09596593651506 | 4.77291284147196 | 6.40265188186281 |
| chr11 | 47164756 | 47164772 | - | 2.00792115374673 | 1.13699574206438 | 5.96614105183995 | 11.7381951167485 |
| chr11 | 47164798 | 47164855 | - | 6.02376346124019 | 4.54798296825753 | 25.0577924177278 | 37.3488026441998 |
| chr11 | 47164876 | 47164948 | - | 8.03168461498692 | 18.1919318730301 | 34.6036181006717 | 21.3421729395427 |
| chr11 | 47165035 | 47165071 | - | 3.01188173062009 | 12.5069531627082 | 13.1255103140479 | 7.46976052883995 |
| chr11 | 47165117 | 47165212 | - | 23.0910932680874 | 21.6029190992233 | 51.3088130458235 | 50.1541064079254 |
| chr11 | 47165250 | 47165253 | - | 2.00792115374673 | 2.27399148412876 | 2.38645642073598 | 5.33554323488568 |
| chr11 | 47165321 | 47165344 | - | 9.03564519186028 | 15.9179403889013 | 15.5119667347839 | 6.40265188186281 |
| chr11 | 47165390 | 47165418 | - | 8.03168461498692 | 12.5069531627082 | 19.0916513658878 | 7.46976052883995 |
| chr11 | 47165441 | 47165465 | - | 2.00792115374673 | 7.95897019445067 | 5.96614105183995 | 10.6710864697714 |
| chr11 | 47177551 | 47177691 | - | 37.1465413443145 | 47.753821166704 | 3.57968463110397 | 3.20132594093141 |
| chr11 | 47177746 | 47177784 | - | 8.03168461498692 | 3.41098722619315 | 1.19322821036799 | 1.06710864697714 |
| chr11 | 47239140 | 47239197 | + | 12.0475269224804 | 31.8358807778027 | 1.19322821036799 | 1.06710864697714 |
| chr11 | 47239323 | 47239357 | - | 15.0594086531005 | 13.6439489047726 | 3.57968463110397 | 6.40265188186281 |
| chr11 | 47268796 | 47268809 | + | 3.01188173062009 | 4.54798296825753 | 2.38645642073598 | 3.20132594093141 |
| chr11 | 47329278 | 47329295 | + | 2.00792115374673 | 5.68497871032191 | 1.19322821036799 | 1.06710864697714 |
| chr11 | 47329300 | 47329315 | + | 2.00792115374673 | 5.68497871032191 | 3.57968463110397 | 1.06710864697714 |
| chr11 | 47329376 | 47329436 | + | 15.0594086531005 | 17.0549361309657 | 16.7051949451519 | 10.6710864697714 |
| chr11 | 47329471 | 47329473 | + | 1.00396057687336 | 1.13699574206438 | 2.38645642073598 | 1.06710864697714 |
| chr11 | 47329566 | 47329643 | + | 56.2217923049084 | 45.4798296825753 | 9.54582568294392 | 23.476390233497 |
| chr11 | 47329912 | 47329956 | + | 14.0554480762271 | 18.1919318730301 | 13.1255103140479 | 11.7381951167485 |
| chr11 | 47416224 | 47416256 | + | 20.0792115374673 | 11.3699574206438 | 5.96614105183995 | 1.06710864697714 |
| chr11 | 47416384 | 47416435 | + | 10.0396057687336 | 7.95897019445067 | 4.77291284147196 | 3.20132594093141 |
| chr11 | 47418799 | 47418804 | - | 6.02376346124019 | 2.27399148412876 | 2.38645642073598 | 3.20132594093141 |
| chr11 | 47466090 | 47466124 | - | 11.043566345607 | 15.9179403889013 | 4.77291284147196 | 5.33554323488568 |
| chr11 | 47466195 | 47466203 | - | 2.00792115374673 | 1.13699574206438 | 3.57968463110397 | 3.20132594093141 |
| chr11 | 47466324 | 47466396 | - | 7.02772403811355 | 6.82197445238629 | 3.57968463110397 | 10.6710864697714 |
| chr11 | 47466491 | 47466511 | - | 13.0514874993537 | 15.9179403889013 | 4.77291284147196 | 2.13421729395427 |
| chr11 | 47466704 | 47466709 | - | 1.00396057687336 | 1.13699574206438 | 1.19322821036799 | 2.13421729395427 |
| chr11 | 47466819 | 47466823 | - | 3.01188173062009 | 3.41098722619315 | 3.57968463110397 | 2.13421729395427 |
| chr11 | 47466900 | 47466932 | - | 24.0950538449607 | 28.4248935516096 | 9.54582568294392 | 7.46976052883995 |
| chr11 | 47467046 | 47467166 | - | 193.764391336559 | 263.783012158937 | 116.936364616063 | 116.314842520508 |
| chr11 | 47467194 | 47467443 | - | 206.815878835913 | 264.920007901001 | 182.563916186302 | 153.663645164708 |
| chr11 | 47467478 | 47467497 | - | 7.02772403811355 | 6.82197445238629 | 3.57968463110397 | 3.20132594093141 |
| chr11 | 47467534 | 47467541 | - | 1.00396057687336 | 1.13699574206438 | 3.57968463110397 | 1.06710864697714 |
| chr11 | 47467587 | 47467637 | - | 14.0554480762271 | 6.82197445238629 | 8.35259747257593 | 2.13421729395427 |
| chr11 | 47467685 | 47467784 | - | 94.3722942260963 | 105.740604011988 | 140.800928823423 | 99.2411041688736 |
| chr11 | 47467867 | 47467918 | - | 15.0594086531005 | 13.6439489047726 | 25.0577924177278 | 7.46976052883995 |
| chr11 | 47468096 | 47468213 | - | 124.491111532297 | 96.6446380754725 | 57.2749540976635 | 153.663645164708 |
| chr11 | 47468236 | 47468256 | - | 4.01584230749346 | 3.41098722619315 | 4.77291284147196 | 9.60397782279422 |
| chr11 | 47468294 | 47468300 | - | 19.0752509605939 | 14.780944646837 | 11.9322821036799 | 10.6710864697714 |
| chr11 | 47468341 | 47468367 | - | 39.1544624980612 | 35.2468680039958 | 25.0577924177278 | 14.9395210576799 |
| chr11 | 47468472 | 47468486 | - | 9.03564519186028 | 11.3699574206438 | 1.19322821036799 | 3.20132594093141 |
| chr11 | 47468512 | 47468537 | - | 8.03168461498692 | 10.2329616785794 | 4.77291284147196 | 3.20132594093141 |
| chr11 | 47468627 | 47468652 | - | 45.1782259593014 | 75.0417189762492 | 23.8645642073598 | 13.8724124107028 |
| chr11 | 47468734 | 47468999 | - | 377.489176904385 | 411.592458627306 | 218.360762497342 | 260.374509862421 |
| chr11 | 47469022 | 47469088 | - | 18.0712903837206 | 23.876910583352 | 2.38645642073598 | 3.20132594093141 |
| chr11 | 47469157 | 47469209 | - | 40.1584230749346 | 46.6168254246397 | 19.0916513658878 | 17.0737383516342 |
| chr11 | 47469524 | 47469551 | - | 10.0396057687336 | 5.68497871032191 | 4.77291284147196 | 1.06710864697714 |
| chr11 | 47469573 | 47469595 | - | 8.03168461498692 | 4.54798296825753 | 2.38645642073598 | 1.06710864697714 |
| chr11 | 47469682 | 47469750 | - | 31.1227778830743 | 28.4248935516096 | 19.0916513658878 | 3.20132594093141 |
| chr11 | 47469803 | 47469809 | - | 3.01188173062009 | 3.41098722619315 | 3.57968463110397 | 2.13421729395427 |
| chr11 | 47469970 | 47469986 | - | 14.0554480762271 | 11.3699574206438 | 11.9322821036799 | 1.06710864697714 |
| chr11 | 47470035 | 47470040 | - | 1.00396057687336 | 2.27399148412876 | 3.57968463110397 | 4.26843458790854 |
| chr11 | 47470086 | 47470103 | - | 5.01980288436682 | 4.54798296825753 | 1.19322821036799 | 1.06710864697714 |
| chr11 | 47470185 | 47470192 | - | 12.0475269224804 | 20.4659233571589 | 11.9322821036799 | 2.13421729395427 |
| chr11 | 47470261 | 47470278 | - | 15.0594086531005 | 21.6029190992233 | 5.96614105183995 | 3.20132594093141 |
| chr11 | 47470350 | 47470442 | - | 139.550520185398 | 156.905412404885 | 126.482190299007 | 21.3421729395427 |
| chr11 | 47470465 | 47470558 | - | 234.926774988367 | 251.276058996228 | 93.0718004087032 | 53.3554323488568 |
| chr11 | 47470580 | 47470669 | - | 256.009947102708 | 300.166875904997 | 81.1395183050233 | 64.0265188186281 |
| chr11 | 47470717 | 47470823 | - | 20.0792115374673 | 19.3289276150945 | 11.9322821036799 | 6.40265188186281 |
| chr11 | 47471641 | 47471643 | - | 1.00396057687336 | 4.54798296825753 | 1.19322821036799 | 1.06710864697714 |
| chr11 | 47471652 | 47471673 | - | 2.00792115374673 | 3.41098722619315 | 4.77291284147196 | 4.26843458790854 |
| chr11 | 47471695 | 47471776 | - | 26.1029749987075 | 20.4659233571589 | 27.4442488384638 | 23.476390233497 |
| chr11 | 47471990 | 47472006 | - | 14.0554480762271 | 12.5069531627082 | 9.54582568294392 | 33.0803680562912 |
| chr11 | 47472037 | 47472064 | - | 5.01980288436682 | 5.68497871032191 | 5.96614105183995 | 10.6710864697714 |
| chr11 | 47472177 | 47472245 | - | 88.3485307648561 | 97.7816338175368 | 25.0577924177278 | 23.476390233497 |
| chr11 | 47571584 | 47571641 | + | 17.0673298068472 | 18.1919318730301 | 8.35259747257593 | 2.13421729395427 |
| chr11 | 47571663 | 47571747 | + | 16.0633692299738 | 18.1919318730301 | 7.15936926220794 | 12.8053037637256 |
| chr11 | 47571854 | 47571863 | + | 6.02376346124019 | 13.6439489047726 | 2.38645642073598 | 6.40265188186281 |
| chr11 | 47572789 | 47572793 | - | 4.01584230749346 | 1.13699574206438 | 2.38645642073598 | 2.13421729395427 |
| chr11 | 47572910 | 47572916 | - | 20.0792115374673 | 13.6439489047726 | 5.96614105183995 | 8.53686917581709 |
| chr11 | 47580832 | 47580985 | + | 77.3049644192491 | 44.3428339405109 | 126.482190299007 | 92.8384522870108 |
| chr11 | 47584385 | 47584538 | + | 49.1940682667949 | 61.3977700714766 | 22.6713359969918 | 34.1474767032683 |
| chr11 | 47618738 | 47618921 | - | 75.2970432655023 | 76.1787147183136 | 73.9801490428153 | 117.381951167485 |
| chr11 | 47716590 | 47716638 | - | 16.0633692299738 | 6.82197445238629 | 2.38645642073598 | 5.33554323488568 |
| chr11 | 47778760 | 47778771 | - | 7.02772403811355 | 9.09596593651506 | 2.38645642073598 | 1.06710864697714 |
| chr11 | 47778956 | 47778960 | - | 3.01188173062009 | 1.13699574206438 | 2.38645642073598 | 4.26843458790854 |
| chr11 | 47839881 | 47839884 | - | 10.0396057687336 | 21.6029190992233 | 4.77291284147196 | 5.33554323488568 |
| chr11 | 57326003 | 57326102 | - | 92.3643730723495 | 140.987472015983 | 57.2749540976635 | 87.5029090521251 |
| chr11 | 57326171 | 57326357 | - | 137.542599031651 | 118.247557174696 | 175.404546924094 | 228.361250453107 |
| chr11 | 57326379 | 57326497 | - | 67.2653586505154 | 38.657855230189 | 85.9124311464952 | 80.0331485232852 |
| chr11 | 57484556 | 57484632 | - | 1.00396057687336 | 3.41098722619315 | 11.9322821036799 | 26.6777161744284 |
| chr11 | 57528538 | 57528590 | - | 30.1188173062009 | 57.9867828452835 | 14.3187385244159 | 16.006629704657 |
| chr11 | 57528646 | 57528661 | - | 2.00792115374673 | 2.27399148412876 | 1.19322821036799 | 8.53686917581709 |
| chr11 | 57528689 | 57528693 | - | 5.01980288436682 | 11.3699574206438 | 8.35259747257593 | 21.3421729395427 |
| chr11 | 57551697 | 57551716 | - | 1.00396057687336 | 1.13699574206438 | 4.77291284147196 | 14.9395210576799 |
| chr11 | 57638570 | 57638587 | + | 12.0475269224804 | 12.5069531627082 | 4.77291284147196 | 14.9395210576799 |
| chr11 | 57661798 | 57661802 | + | 22.087132691214 | 30.6988850357383 | 14.3187385244159 | 26.6777161744284 |
| chr11 | 57700152 | 57700165 | + | 7.02772403811355 | 4.54798296825753 | 7.15936926220794 | 3.20132594093141 |
| chr11 | 57700185 | 57700242 | + | 19.0752509605939 | 23.876910583352 | 42.9562155732476 | 12.8053037637256 |
| chr11 | 57700264 | 57700365 | + | 30.1188173062009 | 28.4248935516096 | 21.4781077866238 | 20.2750642925656 |
| chr11 | 57700394 | 57700404 | + | 7.02772403811355 | 11.3699574206438 | 4.77291284147196 | 9.60397782279422 |
| chr11 | 57700476 | 57700503 | + | 16.0633692299738 | 27.2878978095452 | 26.2510206280958 | 14.9395210576799 |
| chr11 | 57700537 | 57700548 | + | 1.00396057687336 | 1.13699574206438 | 13.1255103140479 | 3.20132594093141 |
| chr11 | 57701054 | 57701115 | + | 8.03168461498692 | 18.1919318730301 | 5.96614105183995 | 13.8724124107028 |
| chr11 | 57703806 | 57703831 | - | 1.00396057687336 | 4.54798296825753 | 2.38645642073598 | 4.26843458790854 |
| chr11 | 57703870 | 57703872 | - | 4.01584230749346 | 10.2329616785794 | 1.19322821036799 | 1.06710864697714 |
| chr11 | 57740757 | 57740760 | + | 3.01188173062009 | 5.68497871032191 | 4.77291284147196 | 5.33554323488568 |
| chr11 | 57740892 | 57740940 | + | 6.02376346124019 | 7.95897019445067 | 7.15936926220794 | 1.06710864697714 |
| chr11 | 57742213 | 57742245 | + | 20.0792115374673 | 45.4798296825753 | 9.54582568294392 | 36.2816939972226 |
| chr11 | 57742874 | 57742936 | + | 16.0633692299738 | 30.6988850357383 | 11.9322821036799 | 10.6710864697714 |
| chr11 | 57742959 | 57743014 | + | 224.887169219634 | 388.852543786019 | 192.109741869246 | 129.120146284233 |
| chr11 | 57816295 | 57816372 | + | 28.1108961524542 | 21.6029190992233 | 20.2848795762558 | 22.4092815865198 |
| chr11 | 57816578 | 57816586 | + | 4.01584230749346 | 4.54798296825753 | 9.54582568294392 | 3.20132594093141 |
| chr11 | 57816608 | 57816693 | + | 14.0554480762271 | 27.2878978095452 | 33.4103898903037 | 19.2079556455884 |
| chr11 | 57816754 | 57816766 | + | 12.0475269224804 | 13.6439489047726 | 13.1255103140479 | 13.8724124107028 |
| chr11 | 57818121 | 57818250 | + | 16.0633692299738 | 12.5069531627082 | 13.1255103140479 | 14.9395210576799 |
| chr11 | 57818344 | 57818358 | + | 37.1465413443145 | 20.4659233571589 | 16.7051949451519 | 30.9461507623369 |
| chr11 | 57819007 | 57819082 | + | 5.01980288436682 | 3.41098722619315 | 5.96614105183995 | 6.40265188186281 |
| chr11 | 57819103 | 57819149 | + | 1.00396057687336 | 4.54798296825753 | 4.77291284147196 | 4.26843458790854 |
| chr11 | 58617655 | 58617732 | + | 59.2336740355285 | 102.329616785794 | 65.6275515702394 | 14.9395210576799 |
| chr11 | 58617808 | 58617816 | + | 21.0831721143407 | 22.7399148412876 | 15.5119667347839 | 3.20132594093141 |
| chr11 | 58617875 | 58617912 | + | 6.02376346124019 | 15.9179403889013 | 7.15936926220794 | 1.06710864697714 |
| chr11 | 58617952 | 58617990 | + | 20.0792115374673 | 22.7399148412876 | 14.3187385244159 | 10.6710864697714 |
| chr11 | 58618412 | 58618422 | + | 20.0792115374673 | 13.6439489047726 | 17.8984231555198 | 13.8724124107028 |
| chr11 | 58618450 | 58618572 | + | 41.1623836518079 | 65.9457530397342 | 41.7629873628796 | 81.1002571702623 |
| chr11 | 58618610 | 58618703 | + | 35.1386201905678 | 50.0278126508328 | 13.1255103140479 | 27.7448248214055 |
| chr11 | 58618785 | 58618809 | + | 6.02376346124019 | 5.68497871032191 | 5.96614105183995 | 2.13421729395427 |
| chr11 | 58618826 | 58618841 | + | 6.02376346124019 | 1.13699574206438 | 4.77291284147196 | 1.06710864697714 |
| chr11 | 58618880 | 58618938 | + | 69.2732798042621 | 121.658544400889 | 65.6275515702394 | 62.959410171651 |
| chr11 | 58618972 | 58619008 | + | 6.02376346124019 | 2.27399148412876 | 8.35259747257593 | 7.46976052883995 |
| chr11 | 58620669 | 58620672 | + | 20.0792115374673 | 60.2607743294122 | 3.57968463110397 | 20.2750642925656 |
| chr11 | 58620722 | 58620724 | + | 6.02376346124019 | 2.27399148412876 | 1.19322821036799 | 1.06710864697714 |
| chr11 | 58620854 | 58620910 | + | 5.01980288436682 | 15.9179403889013 | 4.77291284147196 | 9.60397782279422 |
| chr11 | 59574510 | 59574512 | - | 1.00396057687336 | 2.27399148412876 | 1.19322821036799 | 1.06710864697714 |
| chr11 | 59574535 | 59574536 | - | 2.00792115374673 | 2.27399148412876 | 1.19322821036799 | 2.13421729395427 |
| chr11 | 59574831 | 59574895 | - | 14.0554480762271 | 21.6029190992233 | 29.8307052591997 | 34.1474767032683 |
| chr11 | 59574950 | 59574985 | - | 5.01980288436682 | 3.41098722619315 | 11.9322821036799 | 14.9395210576799 |
| chr11 | 59575039 | 59575066 | - | 3.01188173062009 | 3.41098722619315 | 10.7390538933119 | 8.53686917581709 |
| chr11 | 59575089 | 59575122 | - | 12.0475269224804 | 13.6439489047726 | 25.0577924177278 | 30.9461507623369 |
| chr11 | 59575148 | 59575157 | - | 1.00396057687336 | 2.27399148412876 | 10.7390538933119 | 5.33554323488568 |
| chr11 | 59575415 | 59575449 | - | 1.00396057687336 | 5.68497871032191 | 10.7390538933119 | 6.40265188186281 |
| chr11 | 59575812 | 59575818 | - | 2.00792115374673 | 3.41098722619315 | 1.19322821036799 | 2.13421729395427 |
| chr11 | 59575836 | 59575860 | - | 6.02376346124019 | 4.54798296825753 | 11.9322821036799 | 7.46976052883995 |
| chr11 | 59575950 | 59576012 | - | 1.00396057687336 | 4.54798296825753 | 11.9322821036799 | 11.7381951167485 |
| chr11 | 59576093 | 59576105 | - | 1.00396057687336 | 2.27399148412876 | 4.77291284147196 | 1.06710864697714 |
| chr11 | 59636903 | 59636971 | - | 10.0396057687336 | 10.2329616785794 | 4.77291284147196 | 4.26843458790854 |
| chr11 | 59637084 | 59637097 | - | 14.0554480762271 | 17.0549361309657 | 8.35259747257593 | 8.53686917581709 |
| chr11 | 59637264 | 59637289 | - | 4.01584230749346 | 9.09596593651506 | 7.15936926220794 | 9.60397782279422 |
| chr11 | 59637679 | 59637699 | - | 4.01584230749346 | 3.41098722619315 | 2.38645642073598 | 2.13421729395427 |
| chr11 | 59637764 | 59637769 | - | 5.01980288436682 | 5.68497871032191 | 2.38645642073598 | 1.06710864697714 |
| chr11 | 59637884 | 59637906 | - | 6.02376346124019 | 4.54798296825753 | 1.19322821036799 | 9.60397782279422 |
| chr11 | 59638034 | 59638059 | - | 3.01188173062009 | 6.82197445238629 | 5.96614105183995 | 8.53686917581709 |
| chr11 | 59638148 | 59638167 | - | 1.00396057687336 | 2.27399148412876 | 7.15936926220794 | 2.13421729395427 |
| chr11 | 59638247 | 59638270 | - | 31.1227778830743 | 31.8358807778027 | 19.0916513658878 | 25.6106075274513 |
| chr11 | 59638298 | 59638327 | - | 15.0594086531005 | 6.82197445238629 | 14.3187385244159 | 8.53686917581709 |
| chr11 | 59801438 | 59801445 | + | 6.02376346124019 | 6.82197445238629 | 23.8645642073598 | 11.7381951167485 |
| chr11 | 59802128 | 59802149 | + | 1.00396057687336 | 3.41098722619315 | 17.8984231555198 | 5.33554323488568 |
| chr11 | 59802179 | 59802199 | + | 2.00792115374673 | 4.54798296825753 | 13.1255103140479 | 1.06710864697714 |
| chr11 | 59802251 | 59802271 | + | 1.00396057687336 | 1.13699574206438 | 2.38645642073598 | 1.06710864697714 |
| chr11 | 59802580 | 59802581 | + | 1.00396057687336 | 1.13699574206438 | 3.57968463110397 | 3.20132594093141 |
| chr11 | 59806184 | 59806185 | - | 2.00792115374673 | 3.41098722619315 | 4.77291284147196 | 2.13421729395427 |
| chr11 | 59806224 | 59806268 | - | 19.0752509605939 | 23.876910583352 | 28.6374770488317 | 16.006629704657 |
| chr11 | 59806306 | 59806311 | - | 1.00396057687336 | 6.82197445238629 | 1.19322821036799 | 3.20132594093141 |
| chr11 | 59806330 | 59806344 | - | 12.0475269224804 | 4.54798296825753 | 8.35259747257593 | 11.7381951167485 |
| chr11 | 60850541 | 60850548 | + | 1.00396057687336 | 2.27399148412876 | 4.77291284147196 | 4.26843458790854 |
| chr11 | 60850842 | 60850862 | + | 19.0752509605939 | 18.1919318730301 | 8.35259747257593 | 20.2750642925656 |
| chr11 | 60850976 | 60850997 | + | 9.03564519186028 | 2.27399148412876 | 4.77291284147196 | 7.46976052883995 |
| chr11 | 60851011 | 60851056 | + | 43.1703048055547 | 31.8358807778027 | 22.6713359969918 | 25.6106075274513 |
| chr11 | 60890758 | 60890848 | - | 91.3604124954762 | 110.288586980245 | 42.9562155732476 | 86.435800405148 |
| chr11 | 60890867 | 60890879 | - | 56.2217923049084 | 55.7127913611547 | 20.2848795762558 | 74.6976052883995 |
| chr11 | 60891049 | 60891101 | - | 274.081237486429 | 142.124467758048 | 65.6275515702394 | 139.791232754005 |
| chr11 | 60922082 | 60922191 | + | 51.2019894205416 | 44.3428339405109 | 22.6713359969918 | 14.9395210576799 |
| chr11 | 60922217 | 60922220 | + | 6.02376346124019 | 1.13699574206438 | 1.19322821036799 | 1.06710864697714 |
| chr11 | 60922277 | 60922309 | + | 1.00396057687336 | 1.13699574206438 | 3.57968463110397 | 4.26843458790854 |
| chr11 | 60922311 | 60922322 | + | 2.00792115374673 | 2.27399148412876 | 1.19322821036799 | 1.06710864697714 |
| chr11 | 60922365 | 60922380 | + | 13.0514874993537 | 5.68497871032191 | 2.38645642073598 | 6.40265188186281 |
| chr11 | 60922423 | 60922448 | + | 15.0594086531005 | 7.95897019445067 | 10.7390538933119 | 5.33554323488568 |
| chr11 | 60922527 | 60922556 | + | 11.043566345607 | 7.95897019445067 | 7.15936926220794 | 2.13421729395427 |
| chr11 | 60922687 | 60922702 | + | 16.0633692299738 | 14.780944646837 | 13.1255103140479 | 2.13421729395427 |
| chr11 | 60922751 | 60922781 | + | 15.0594086531005 | 15.9179403889013 | 21.4781077866238 | 9.60397782279422 |
| chr11 | 60922839 | 60922878 | + | 21.0831721143407 | 14.780944646837 | 4.77291284147196 | 5.33554323488568 |
| chr11 | 60923247 | 60923280 | + | 51.2019894205416 | 68.2197445238629 | 13.1255103140479 | 25.6106075274513 |
| chr11 | 60923334 | 60923352 | + | 2.00792115374673 | 4.54798296825753 | 2.38645642073598 | 1.06710864697714 |
| chr11 | 61130315 | 61130347 | - | 1.00396057687336 | 2.27399148412876 | 2.38645642073598 | 1.06710864697714 |
| chr11 | 61130556 | 61130606 | - | 5.01980288436682 | 1.13699574206438 | 14.3187385244159 | 73.6304966414224 |
| chr11 | 61130613 | 61130686 | - | 30.1188173062009 | 18.1919318730301 | 27.4442488384638 | 38.4159112911769 |
| chr11 | 61130722 | 61130738 | - | 2.00792115374673 | 4.54798296825753 | 14.3187385244159 | 6.40265188186281 |
| chr11 | 61130764 | 61130767 | - | 3.01188173062009 | 2.27399148412876 | 2.38645642073598 | 4.26843458790854 |
| chr11 | 61131291 | 61131317 | - | 3.01188173062009 | 7.95897019445067 | 3.57968463110397 | 2.13421729395427 |
| chr11 | 61299481 | 61299640 | - | 149.590125954131 | 151.220433694563 | 122.902505667903 | 232.629685041016 |
| chr11 | 61299670 | 61299790 | - | 31.1227778830743 | 15.9179403889013 | 16.7051949451519 | 17.0737383516342 |
| chr11 | 61299820 | 61299908 | - | 19.0752509605939 | 9.09596593651506 | 4.77291284147196 | 22.4092815865198 |
| chr11 | 61299946 | 61299988 | - | 15.0594086531005 | 2.27399148412876 | 8.35259747257593 | 17.0737383516342 |
| chr11 | 61300127 | 61300222 | - | 45.1782259593014 | 21.6029190992233 | 41.7629873628796 | 87.5029090521251 |
| chr11 | 61346774 | 61346892 | + | 5.01980288436682 | 6.82197445238629 | 4.77291284147196 | 2.13421729395427 |
| chr11 | 61348833 | 61348838 | - | 6.02376346124019 | 6.82197445238629 | 1.19322821036799 | 3.20132594093141 |
| chr11 | 61348861 | 61348864 | - | 13.0514874993537 | 20.4659233571589 | 8.35259747257593 | 10.6710864697714 |
| chr11 | 61349169 | 61349202 | - | 16.0633692299738 | 11.3699574206438 | 16.7051949451519 | 71.4962793474681 |
| chr11 | 61349307 | 61349313 | - | 6.02376346124019 | 5.68497871032191 | 7.15936926220794 | 13.8724124107028 |
| chr11 | 61349489 | 61349519 | - | 7.02772403811355 | 4.54798296825753 | 7.15936926220794 | 11.7381951167485 |
| chr11 | 61349754 | 61349793 | - | 6.02376346124019 | 4.54798296825753 | 16.7051949451519 | 23.476390233497 |
| chr11 | 61349899 | 61349911 | - | 6.02376346124019 | 1.13699574206438 | 5.96614105183995 | 4.26843458790854 |
| chr11 | 61350387 | 61350409 | - | 1.00396057687336 | 1.13699574206438 | 7.15936926220794 | 3.20132594093141 |
| chr11 | 61369038 | 61369061 | + | 6.02376346124019 | 7.95897019445067 | 3.57968463110397 | 8.53686917581709 |
| chr11 | 61369128 | 61369157 | + | 4.01584230749346 | 7.95897019445067 | 3.57968463110397 | 5.33554323488568 |
| chr11 | 61398469 | 61398513 | + | 2.00792115374673 | 5.68497871032191 | 1.19322821036799 | 2.13421729395427 |
| chr11 | 61402667 | 61402668 | - | 1.00396057687336 | 6.82197445238629 | 1.19322821036799 | 2.13421729395427 |
| chr11 | 61402787 | 61402817 | - | 16.0633692299738 | 13.6439489047726 | 5.96614105183995 | 9.60397782279422 |
| chr11 | 61402918 | 61402936 | - | 22.087132691214 | 20.4659233571589 | 5.96614105183995 | 7.46976052883995 |
| chr11 | 61402957 | 61402997 | - | 2.00792115374673 | 9.09596593651506 | 2.38645642073598 | 2.13421729395427 |
| chr11 | 61403030 | 61403037 | - | 2.00792115374673 | 3.41098722619315 | 1.19322821036799 | 4.26843458790854 |
| chr11 | 61403441 | 61403533 | - | 19.0752509605939 | 19.3289276150945 | 26.2510206280958 | 49.0869977609482 |
| chr11 | 61403580 | 61403620 | - | 9.03564519186028 | 4.54798296825753 | 2.38645642073598 | 2.13421729395427 |
| chr11 | 61403641 | 61403682 | - | 5.01980288436682 | 15.9179403889013 | 7.15936926220794 | 5.33554323488568 |
| chr11 | 61403698 | 61403766 | - | 19.0752509605939 | 29.5618892936739 | 14.3187385244159 | 26.6777161744284 |
| chr11 | 61403800 | 61403834 | - | 15.0594086531005 | 28.4248935516096 | 7.15936926220794 | 7.46976052883995 |
| chr11 | 61404090 | 61404120 | - | 2.00792115374673 | 6.82197445238629 | 3.57968463110397 | 1.06710864697714 |
| chr11 | 61404161 | 61404182 | - | 2.00792115374673 | 2.27399148412876 | 4.77291284147196 | 7.46976052883995 |
| chr11 | 61404260 | 61404316 | - | 6.02376346124019 | 7.95897019445067 | 8.35259747257593 | 10.6710864697714 |
| chr11 | 61404483 | 61404506 | - | 17.0673298068472 | 9.09596593651506 | 7.15936926220794 | 13.8724124107028 |
| chr11 | 61404555 | 61404576 | - | 2.00792115374673 | 5.68497871032191 | 3.57968463110397 | 3.20132594093141 |
| chr11 | 61404690 | 61404704 | - | 4.01584230749346 | 14.780944646837 | 2.38645642073598 | 3.20132594093141 |
| chr11 | 61420015 | 61420050 | - | 5.01980288436682 | 5.68497871032191 | 1.19322821036799 | 3.20132594093141 |
| chr11 | 61420069 | 61420085 | - | 5.01980288436682 | 6.82197445238629 | 5.96614105183995 | 1.06710864697714 |
| chr11 | 61446495 | 61446511 | + | 3.01188173062009 | 1.13699574206438 | 2.38645642073598 | 2.13421729395427 |
| chr11 | 61446550 | 61446575 | + | 3.01188173062009 | 1.13699574206438 | 1.19322821036799 | 2.13421729395427 |
| chr11 | 61446610 | 61446704 | + | 33.130699036821 | 59.1237785873479 | 29.8307052591997 | 22.4092815865198 |
| chr11 | 61789154 | 61789178 | - | 24.0950538449607 | 14.780944646837 | 25.0577924177278 | 19.2079556455884 |
| chr11 | 61789200 | 61789268 | - | 1552.12305184622 | 1357.57291602487 | 1755.23869745131 | 1513.16006141358 |
| chr11 | 61789785 | 61789917 | - | 381.505019211879 | 363.838637460602 | 788.723847053241 | 517.547693783911 |
| chr11 | 61796545 | 61796610 | + | 191.756470182813 | 181.919318730301 | 116.936364616063 | 149.395210576799 |
| chr11 | 61796721 | 61796749 | + | 7.02772403811355 | 6.82197445238629 | 9.54582568294392 | 12.8053037637256 |
| chr11 | 61796817 | 61796842 | + | 20.0792115374673 | 30.6988850357383 | 34.6036181006717 | 52.2883237018796 |
| chr11 | 61796875 | 61796940 | + | 666.629823043914 | 541.209973222646 | 235.065957442494 | 443.917197142488 |
| chr11 | 61796976 | 61797023 | + | 45.1782259593014 | 21.6029190992233 | 10.7390538933119 | 41.6172372321083 |
| chr11 | 61797051 | 61797072 | + | 54.2138711511617 | 43.2058381984465 | 27.4442488384638 | 77.8989312293309 |
| chr11 | 61797113 | 61797155 | + | 28.1108961524542 | 45.4798296825753 | 27.4442488384638 | 38.4159112911769 |
| chr11 | 61797184 | 61797185 | + | 2.00792115374673 | 5.68497871032191 | 2.38645642073598 | 4.26843458790854 |
| chr11 | 61797206 | 61797217 | + | 13.0514874993537 | 14.780944646837 | 3.57968463110397 | 5.33554323488568 |
| chr11 | 61799648 | 61799687 | - | 11.043566345607 | 32.9728765198671 | 39.3765309421437 | 29.8790421153598 |
| chr11 | 61799708 | 61799742 | - | 13.0514874993537 | 15.9179403889013 | 26.2510206280958 | 62.959410171651 |
| chr11 | 61799785 | 61799855 | - | 23.0910932680874 | 20.4659233571589 | 53.6952694665595 | 82.1673658172394 |
| chr11 | 61799914 | 61799933 | - | 3.01188173062009 | 7.95897019445067 | 5.96614105183995 | 20.2750642925656 |
| chr11 | 61800284 | 61800292 | - | 3.01188173062009 | 1.13699574206438 | 9.54582568294392 | 7.46976052883995 |
| chr11 | 61800367 | 61800717 | - | 174.689140375965 | 186.467301698559 | 192.109741869246 | 411.903937733174 |
| chr11 | 61800768 | 61800871 | - | 28.1108961524542 | 19.3289276150945 | 75.1733772531833 | 78.966039876308 |
| chr11 | 61801029 | 61801081 | - | 12.0475269224804 | 6.82197445238629 | 11.9322821036799 | 22.4092815865198 |
| chr11 | 61801153 | 61801263 | - | 20.0792115374673 | 15.9179403889013 | 42.9562155732476 | 61.8923015246739 |
| chr11 | 61801312 | 61801328 | - | 13.0514874993537 | 20.4659233571589 | 13.1255103140479 | 9.60397782279422 |
| chr11 | 61801351 | 61801416 | - | 25.0990144218341 | 18.1919318730301 | 27.4442488384638 | 42.6843458790854 |
| chr11 | 61801484 | 61801533 | - | 9.03564519186028 | 10.2329616785794 | 19.0916513658878 | 23.476390233497 |
| chr11 | 61801581 | 61801626 | - | 3.01188173062009 | 4.54798296825753 | 5.96614105183995 | 9.60397782279422 |
| chr11 | 61801689 | 61801839 | - | 45.1782259593014 | 54.5757956190903 | 152.733210927103 | 218.757272630313 |
| chr11 | 61801864 | 61801882 | - | 1.00396057687336 | 3.41098722619315 | 22.6713359969918 | 3.20132594093141 |
| chr11 | 61802046 | 61802247 | - | 2331.19645949995 | 2944.81897194675 | 2588.11198828817 | 4053.94574986614 |
| chr11 | 61802293 | 61802315 | - | 57.2257528817818 | 60.2607743294122 | 126.482190299007 | 177.140035398205 |
| chr11 | 61802337 | 61802389 | - | 22.087132691214 | 25.0139063254164 | 69.2072362013434 | 55.4896496428111 |
| chr11 | 61865712 | 61865750 | + | 9.03564519186028 | 9.09596593651506 | 1.19322821036799 | 1.06710864697714 |
| chr11 | 61866326 | 61866334 | + | 14.0554480762271 | 7.95897019445067 | 13.1255103140479 | 1.06710864697714 |
| chr11 | 61866479 | 61866501 | + | 17.0673298068472 | 10.2329616785794 | 10.7390538933119 | 19.2079556455884 |
| chr11 | 61866529 | 61866557 | + | 5.01980288436682 | 2.27399148412876 | 7.15936926220794 | 19.2079556455884 |
| chr11 | 61866634 | 61866672 | + | 8.03168461498692 | 6.82197445238629 | 7.15936926220794 | 8.53686917581709 |
| chr11 | 61866761 | 61866824 | + | 18.0712903837206 | 9.09596593651506 | 9.54582568294392 | 2.13421729395427 |
| chr11 | 61866872 | 61867039 | + | 157.621810569118 | 71.6307317500561 | 71.5936926220794 | 78.966039876308 |
| chr11 | 61867148 | 61867166 | + | 4.01584230749346 | 1.13699574206438 | 1.19322821036799 | 1.06710864697714 |
| chr11 | 61867254 | 61867303 | + | 17.0673298068472 | 12.5069531627082 | 3.57968463110397 | 8.53686917581709 |
| chr11 | 61873558 | 61873630 | - | 70.2772403811355 | 98.9186295596012 | 186.143600817406 | 235.831010981947 |
| chr11 | 61897334 | 61897335 | - | 2.00792115374673 | 11.3699574206438 | 2.38645642073598 | 4.26843458790854 |
| chr11 | 61964721 | 61964745 | - | 7.02772403811355 | 7.95897019445067 | 10.7390538933119 | 14.9395210576799 |
| chr11 | 61967769 | 61967815 | + | 5.01980288436682 | 3.41098722619315 | 7.15936926220794 | 10.6710864697714 |
| chr11 | 62152010 | 62152038 | + | 1.00396057687336 | 2.27399148412876 | 5.96614105183995 | 6.40265188186281 |
| chr11 | 62152183 | 62152191 | + | 103.407939417957 | 60.2607743294122 | 39.3765309421437 | 19.2079556455884 |
| chr11 | 62152219 | 62152229 | + | 6.02376346124019 | 1.13699574206438 | 3.57968463110397 | 8.53686917581709 |
| chr11 | 62152295 | 62152300 | + | 14.0554480762271 | 9.09596593651506 | 9.54582568294392 | 16.006629704657 |
| chr11 | 62152545 | 62152577 | + | 12.0475269224804 | 3.41098722619315 | 1.19322821036799 | 3.20132594093141 |
| chr11 | 62152731 | 62152754 | + | 9.03564519186028 | 1.13699574206438 | 7.15936926220794 | 5.33554323488568 |
| chr11 | 62357003 | 62357062 | + | 2.00792115374673 | 3.41098722619315 | 1.19322821036799 | 1.06710864697714 |
| chr11 | 62392180 | 62392211 | + | 11.043566345607 | 9.09596593651506 | 1.19322821036799 | 5.33554323488568 |
| chr11 | 62392497 | 62392510 | + | 1.00396057687336 | 2.27399148412876 | 3.57968463110397 | 1.06710864697714 |
| chr11 | 62392681 | 62392750 | + | 8.03168461498692 | 10.2329616785794 | 10.7390538933119 | 4.26843458790854 |
| chr11 | 62393080 | 62393165 | + | 7.02772403811355 | 4.54798296825753 | 3.57968463110397 | 4.26843458790854 |
| chr11 | 62393197 | 62393261 | + | 28.1108961524542 | 29.5618892936739 | 26.2510206280958 | 35.2145853502455 |
| chr11 | 62433741 | 62433742 | - | 1.00396057687336 | 1.13699574206438 | 1.19322821036799 | 1.06710864697714 |
| chr11 | 62515921 | 62516099 | - | 54.2138711511617 | 63.6717615556054 | 20.2848795762558 | 16.006629704657 |
| chr11 | 62516173 | 62516254 | - | 32.1267384599477 | 29.5618892936739 | 8.35259747257593 | 10.6710864697714 |
| chr11 | 62516283 | 62516416 | - | 164.649534607232 | 79.5897019445067 | 16.7051949451519 | 9.60397782279422 |
| chr11 | 62516439 | 62516464 | - | 1.00396057687336 | 6.82197445238629 | 2.38645642073598 | 2.13421729395427 |
| chr11 | 62516502 | 62516618 | - | 34.1346596136944 | 19.3289276150945 | 22.6713359969918 | 14.9395210576799 |
| chr11 | 62516650 | 62516705 | - | 12.0475269224804 | 11.3699574206438 | 10.7390538933119 | 4.26843458790854 |
| chr11 | 62516728 | 62516810 | - | 26.1029749987075 | 17.0549361309657 | 13.1255103140479 | 5.33554323488568 |
| chr11 | 62559616 | 62559626 | - | 3.01188173062009 | 4.54798296825753 | 3.57968463110397 | 6.40265188186281 |
| chr11 | 62559697 | 62559716 | - | 58.2297134586551 | 79.5897019445067 | 65.6275515702394 | 125.918820343302 |
| chr11 | 62559777 | 62559801 | - | 21.0831721143407 | 17.0549361309657 | 7.15936926220794 | 19.2079556455884 |
| chr11 | 62560068 | 62560069 | - | 5.01980288436682 | 4.54798296825753 | 2.38645642073598 | 12.8053037637256 |
| chr11 | 62566805 | 62566806 | - | 5.01980288436682 | 2.27399148412876 | 1.19322821036799 | 1.06710864697714 |
| chr11 | 62566990 | 62567013 | - | 47.1861471130481 | 42.0688424563821 | 103.810854302015 | 103.509538756782 |
| chr11 | 62567398 | 62567403 | - | 19.0752509605939 | 20.4659233571589 | 31.0239334695677 | 16.006629704657 |
| chr11 | 62567436 | 62567463 | - | 60.2376346124019 | 42.0688424563821 | 118.129592826431 | 70.429170700491 |
| chr11 | 62567510 | 62567531 | - | 32.1267384599477 | 42.0688424563821 | 79.9462900946553 | 40.5501285851312 |
| chr11 | 62572605 | 62572643 | - | 57.2257528817818 | 63.6717615556054 | 113.356679984959 | 59.7580842307196 |
| chr11 | 62572670 | 62572693 | - | 125.495072109171 | 155.76841666282 | 365.127832372605 | 151.529427870753 |
| chr11 | 62573740 | 62573766 | - | 9.03564519186028 | 11.3699574206438 | 1.19322821036799 | 6.40265188186281 |
| chr11 | 62573796 | 62574097 | - | 21471.7048575906 | 29458.4226811461 | 27561.1852030798 | 15704.6379575625 |
| chr11 | 62593261 | 62593266 | - | 1.00396057687336 | 1.13699574206438 | 3.57968463110397 | 2.13421729395427 |
| chr11 | 62593295 | 62593326 | - | 8.03168461498692 | 6.82197445238629 | 4.77291284147196 | 3.20132594093141 |
| chr11 | 62593351 | 62593357 | - | 1.00396057687336 | 1.13699574206438 | 1.19322821036799 | 2.13421729395427 |
| chr11 | 62593477 | 62593492 | - | 10.0396057687336 | 7.95897019445067 | 9.54582568294392 | 26.6777161744284 |
| chr11 | 62593512 | 62593549 | - | 5.01980288436682 | 10.2329616785794 | 2.38645642073598 | 5.33554323488568 |
| chr11 | 62593583 | 62593592 | - | 33.130699036821 | 31.8358807778027 | 20.2848795762558 | 26.6777161744284 |
| chr11 | 62593623 | 62593656 | - | 32.1267384599477 | 29.5618892936739 | 10.7390538933119 | 32.0132594093141 |
| chr11 | 62593785 | 62593789 | - | 4.01584230749346 | 3.41098722619315 | 1.19322821036799 | 2.13421729395427 |
| chr11 | 62602240 | 62602244 | - | 7.02772403811355 | 22.7399148412876 | 17.8984231555198 | 19.2079556455884 |
| chr11 | 62602283 | 62602395 | - | 277.093119217049 | 152.357429436627 | 39.3765309421437 | 107.777973344691 |
| chr11 | 62615359 | 62615380 | - | 1.00396057687336 | 1.13699574206438 | 3.57968463110397 | 4.26843458790854 |
| chr11 | 62615628 | 62615641 | - | 3.01188173062009 | 2.27399148412876 | 5.96614105183995 | 14.9395210576799 |
| chr11 | 62616100 | 62616101 | - | 62.2455557661486 | 64.8087572976698 | 36.9900745214077 | 23.476390233497 |
| chr11 | 62624854 | 62624889 | - | 30.1188173062009 | 17.0549361309657 | 10.7390538933119 | 6.40265188186281 |
| chr11 | 62624923 | 62624955 | - | 201.796075951546 | 234.221122865263 | 77.5598336739193 | 123.784603049348 |
| chr11 | 62625017 | 62625083 | - | 9.03564519186028 | 12.5069531627082 | 4.77291284147196 | 24.5434988804741 |
| chr11 | 62625235 | 62625320 | - | 56.2217923049084 | 53.438799877026 | 68.0140079909754 | 73.6304966414224 |
| chr11 | 62625342 | 62625496 | - | 240.950538449607 | 202.38524208746 | 332.910670692669 | 269.978487685215 |
| chr11 | 62625545 | 62625926 | - | 627.475360545853 | 537.798985996453 | 381.833027317757 | 194.213773749839 |
| chr11 | 62646911 | 62646914 | - | 4.01584230749346 | 6.82197445238629 | 2.38645642073598 | 4.26843458790854 |
| chr11 | 62646943 | 62646970 | - | 9.03564519186028 | 4.54798296825753 | 1.19322821036799 | 10.6710864697714 |
| chr11 | 62662855 | 62662864 | - | 1.00396057687336 | 2.27399148412876 | 8.35259747257593 | 6.40265188186281 |
| chr11 | 62662892 | 62662913 | - | 40.1584230749346 | 30.6988850357383 | 22.6713359969918 | 41.6172372321083 |
| chr11 | 62663005 | 62663022 | - | 1.00396057687336 | 6.82197445238629 | 3.57968463110397 | 9.60397782279422 |
| chr11 | 62672234 | 62672258 | + | 10.0396057687336 | 7.95897019445067 | 3.57968463110397 | 6.40265188186281 |
| chr11 | 62672307 | 62672345 | + | 79.3128855729958 | 102.329616785794 | 50.1155848354556 | 99.2411041688736 |
| chr11 | 62676529 | 62676612 | - | 44.174265382428 | 59.1237785873479 | 25.0577924177278 | 29.8790421153598 |
| chr11 | 62690338 | 62690360 | - | 9.03564519186028 | 6.82197445238629 | 5.96614105183995 | 6.40265188186281 |
| chr11 | 62690794 | 62690854 | - | 10.0396057687336 | 6.82197445238629 | 26.2510206280958 | 23.476390233497 |
| chr11 | 62691074 | 62691084 | - | 9.03564519186028 | 1.13699574206438 | 4.77291284147196 | 14.9395210576799 |
| chr11 | 62691304 | 62691337 | - | 22.087132691214 | 12.5069531627082 | 26.2510206280958 | 16.006629704657 |
| chr11 | 62694600 | 62694603 | - | 1.00396057687336 | 3.41098722619315 | 1.19322821036799 | 1.06710864697714 |
| chr11 | 62714138 | 62714146 | - | 84.3326884573626 | 63.6717615556054 | 21.4781077866238 | 30.9461507623369 |
| chr11 | 62714347 | 62714366 | - | 4.01584230749346 | 4.54798296825753 | 2.38645642073598 | 2.13421729395427 |
| chr11 | 62714412 | 62714421 | - | 12.0475269224804 | 13.6439489047726 | 13.1255103140479 | 1.06710864697714 |
| chr11 | 62714554 | 62714574 | - | 17.0673298068472 | 11.3699574206438 | 5.96614105183995 | 1.06710864697714 |
| chr11 | 62714773 | 62714922 | - | 90.3564519186028 | 110.288586980245 | 65.6275515702394 | 99.2411041688736 |
| chr11 | 62714961 | 62714973 | - | 2.00792115374673 | 2.27399148412876 | 2.38645642073598 | 12.8053037637256 |
| chr11 | 62715128 | 62715178 | - | 730.883299963809 | 690.15641543308 | 635.990636126138 | 688.285077300253 |
| chr11 | 62738440 | 62738464 | + | 7.02772403811355 | 10.2329616785794 | 2.38645642073598 | 2.13421729395427 |
| chr11 | 62738488 | 62738493 | + | 9.03564519186028 | 9.09596593651506 | 1.19322821036799 | 2.13421729395427 |
| chr11 | 62738598 | 62738616 | + | 12.0475269224804 | 5.68497871032191 | 5.96614105183995 | 3.20132594093141 |
| chr11 | 62766491 | 62766578 | + | 187.740627875319 | 109.151591238181 | 134.834787771583 | 188.878230514953 |
| chr11 | 62766600 | 62766682 | + | 34.1346596136944 | 43.2058381984465 | 29.8307052591997 | 23.476390233497 |
| chr11 | 62787338 | 62787439 | + | 207.819839412786 | 216.029190992233 | 198.075882921086 | 358.548505384318 |
| chr11 | 62790156 | 62790221 | + | 27.1069355755808 | 29.5618892936739 | 11.9322821036799 | 21.3421729395427 |
| chr11 | 62790326 | 62790356 | + | 2.00792115374673 | 7.95897019445067 | 4.77291284147196 | 11.7381951167485 |
| chr11 | 62790344 | 62790363 | - | 11.043566345607 | 17.0549361309657 | 7.15936926220794 | 7.46976052883995 |
| chr11 | 62790387 | 62790404 | - | 18.0712903837206 | 52.3018041349616 | 4.77291284147196 | 14.9395210576799 |
| chr11 | 62790465 | 62790518 | - | 15.0594086531005 | 31.8358807778027 | 7.15936926220794 | 14.9395210576799 |
| chr11 | 62792203 | 62792257 | - | 16.0633692299738 | 19.3289276150945 | 8.35259747257593 | 3.20132594093141 |
| chr11 | 62794934 | 62794938 | - | 19.0752509605939 | 29.5618892936739 | 39.3765309421437 | 24.5434988804741 |
| chr11 | 62797225 | 62797241 | - | 6.02376346124019 | 2.27399148412876 | 2.38645642073598 | 4.26843458790854 |
| chr11 | 62798684 | 62798698 | - | 2.00792115374673 | 1.13699574206438 | 3.57968463110397 | 5.33554323488568 |
| chr11 | 62798727 | 62798746 | - | 3.01188173062009 | 3.41098722619315 | 8.35259747257593 | 3.20132594093141 |
| chr11 | 62798858 | 62798867 | - | 1.00396057687336 | 4.54798296825753 | 3.57968463110397 | 2.13421729395427 |
| chr11 | 62806991 | 62807079 | - | 86.3406096111093 | 75.0417189762492 | 46.5359002043516 | 43.7514545260626 |
| chr11 | 62807109 | 62807137 | - | 21.0831721143407 | 23.876910583352 | 10.7390538933119 | 8.53686917581709 |
| chr11 | 62807169 | 62807196 | - | 12.0475269224804 | 13.6439489047726 | 8.35259747257593 | 7.46976052883995 |
| chr11 | 62807218 | 62807234 | - | 2.00792115374673 | 2.27399148412876 | 2.38645642073598 | 7.46976052883995 |
| chr11 | 62807353 | 62807369 | - | 2.00792115374673 | 3.41098722619315 | 1.19322821036799 | 3.20132594093141 |
| chr11 | 62832936 | 62833011 | - | 458.809983631128 | 247.865071770035 | 136.028015981951 | 265.710053097307 |
| chr11 | 62888573 | 62888748 | + | 117.463387494184 | 153.494425178692 | 45.3426719939836 | 62.959410171651 |
| chr11 | 62888790 | 62888831 | + | 22.087132691214 | 55.7127913611547 | 9.54582568294392 | 6.40265188186281 |
| chr11 | 62888854 | 62888861 | + | 8.03168461498692 | 15.9179403889013 | 1.19322821036799 | 2.13421729395427 |
| chr11 | 63624173 | 63624177 | - | 7.02772403811355 | 13.6439489047726 | 2.38645642073598 | 4.26843458790854 |
| chr11 | 63624374 | 63624386 | - | 19.0752509605939 | 20.4659233571589 | 3.57968463110397 | 6.40265188186281 |
| chr11 | 63627597 | 63627606 | - | 14.0554480762271 | 5.68497871032191 | 3.57968463110397 | 2.13421729395427 |
| chr11 | 63628042 | 63628048 | - | 17.0673298068472 | 17.0549361309657 | 3.57968463110397 | 3.20132594093141 |
| chr11 | 63628341 | 63628385 | - | 50.1980288436682 | 59.1237785873479 | 13.1255103140479 | 17.0737383516342 |
| chr11 | 63628543 | 63628568 | - | 42.1663442286813 | 81.8636934286355 | 15.5119667347839 | 8.53686917581709 |
| chr11 | 63629175 | 63629255 | - | 24.0950538449607 | 11.3699574206438 | 4.77291284147196 | 18.1408469986113 |
| chr11 | 63758192 | 63758209 | + | 3.01188173062009 | 4.54798296825753 | 4.77291284147196 | 7.46976052883995 |
| chr11 | 63758256 | 63758278 | + | 15.0594086531005 | 25.0139063254164 | 7.15936926220794 | 12.8053037637256 |
| chr11 | 63758424 | 63758440 | + | 2.00792115374673 | 5.68497871032191 | 2.38645642073598 | 6.40265188186281 |
| chr11 | 63758484 | 63758534 | + | 55.217831728035 | 79.5897019445067 | 51.3088130458235 | 32.0132594093141 |
| chr11 | 63758576 | 63758652 | + | 112.443584609817 | 97.7816338175368 | 124.095733878271 | 53.3554323488568 |
| chr11 | 63758688 | 63758731 | + | 15.0594086531005 | 13.6439489047726 | 27.4442488384638 | 11.7381951167485 |
| chr11 | 63758740 | 63758808 | + | 14.0554480762271 | 15.9179403889013 | 22.6713359969918 | 14.9395210576799 |
| chr11 | 63758830 | 63758867 | + | 183.724785567826 | 228.536144154941 | 124.095733878271 | 117.381951167485 |
| chr11 | 63758892 | 63758956 | + | 197.780233644053 | 192.152280408881 | 212.394621445502 | 128.053037637256 |
| chr11 | 63758980 | 63759042 | + | 498.968406706062 | 511.648083928972 | 205.235252183294 | 100.308212815851 |
| chr11 | 63759075 | 63759115 | + | 22.087132691214 | 19.3289276150945 | 10.7390538933119 | 19.2079556455884 |
| chr11 | 63759268 | 63759270 | + | 2.00792115374673 | 2.27399148412876 | 1.19322821036799 | 1.06710864697714 |
| chr11 | 63759417 | 63759491 | + | 60.2376346124019 | 46.6168254246397 | 73.9801490428153 | 58.6909755837425 |
| chr11 | 63759624 | 63759654 | + | 219.867366335267 | 162.590391115207 | 87.1056593568632 | 160.06629704657 |
| chr11 | 63759785 | 63759790 | + | 4.01584230749346 | 4.54798296825753 | 7.15936926220794 | 6.40265188186281 |
| chr11 | 63759828 | 63759860 | + | 17.0673298068472 | 27.2878978095452 | 2.38645642073598 | 7.46976052883995 |
| chr11 | 63762215 | 63762217 | - | 2.00792115374673 | 2.27399148412876 | 1.19322821036799 | 1.06710864697714 |
| chr11 | 63762273 | 63762285 | - | 10.0396057687336 | 11.3699574206438 | 3.57968463110397 | 1.06710864697714 |
| chr11 | 63827617 | 63827647 | + | 22.087132691214 | 17.0549361309657 | 9.54582568294392 | 19.2079556455884 |
| chr11 | 63909502 | 63909556 | + | 330.303029791337 | 230.81013563907 | 41.7629873628796 | 50.1541064079254 |
| chr11 | 63909629 | 63909677 | + | 16.0633692299738 | 7.95897019445067 | 5.96614105183995 | 4.26843458790854 |
| chr11 | 63909843 | 63909852 | + | 25.0990144218341 | 14.780944646837 | 13.1255103140479 | 3.20132594093141 |
| chr11 | 63910002 | 63910005 | + | 1.00396057687336 | 1.13699574206438 | 1.19322821036799 | 2.13421729395427 |
| chr11 | 63910087 | 63910193 | + | 41.1623836518079 | 36.3838637460602 | 28.6374770488317 | 7.46976052883995 |
| chr11 | 63910242 | 63910278 | + | 3.01188173062009 | 6.82197445238629 | 5.96614105183995 | 2.13421729395427 |
| chr11 | 63910462 | 63910510 | + | 39.1544624980612 | 17.0549361309657 | 10.7390538933119 | 11.7381951167485 |
| chr11 | 63910564 | 63910625 | + | 16.0633692299738 | 12.5069531627082 | 2.38645642073598 | 32.0132594093141 |
| chr11 | 63955372 | 63955394 | + | 1.00396057687336 | 4.54798296825753 | 2.38645642073598 | 8.53686917581709 |
| chr11 | 63955712 | 63955713 | + | 1.00396057687336 | 1.13699574206438 | 1.19322821036799 | 1.06710864697714 |
| chr11 | 63957114 | 63957138 | + | 5.01980288436682 | 10.2329616785794 | 1.19322821036799 | 6.40265188186281 |
| chr11 | 63957254 | 63957294 | + | 13.0514874993537 | 3.41098722619315 | 5.96614105183995 | 1.06710864697714 |
| chr11 | 63976307 | 63976411 | + | 35.1386201905678 | 31.8358807778027 | 23.8645642073598 | 54.4225409958339 |
| chr11 | 63976433 | 63976524 | + | 408.611954787459 | 367.249624686795 | 335.297127113405 | 372.42091779502 |
| chr11 | 63997588 | 63997631 | + | 18.0712903837206 | 14.780944646837 | 17.8984231555198 | 14.9395210576799 |
| chr11 | 63998101 | 63998108 | + | 2.00792115374673 | 1.13699574206438 | 4.77291284147196 | 2.13421729395427 |
| chr11 | 63998152 | 63998172 | + | 5.01980288436682 | 5.68497871032191 | 9.54582568294392 | 3.20132594093141 |
| chr11 | 63998202 | 63998211 | + | 7.02772403811355 | 6.82197445238629 | 5.96614105183995 | 2.13421729395427 |
| chr11 | 63998237 | 63998249 | + | 2.00792115374673 | 1.13699574206438 | 2.38645642073598 | 1.06710864697714 |
| chr11 | 63998272 | 63998296 | + | 17.0673298068472 | 6.82197445238629 | 7.15936926220794 | 13.8724124107028 |
| chr11 | 63998344 | 63998382 | + | 39.1544624980612 | 15.9179403889013 | 8.35259747257593 | 32.0132594093141 |
| chr11 | 63998581 | 63998599 | - | 15.0594086531005 | 13.6439489047726 | 8.35259747257593 | 6.40265188186281 |
| chr11 | 64204193 | 64204228 | + | 3.01188173062009 | 1.13699574206438 | 1.19322821036799 | 10.6710864697714 |
| chr11 | 64204372 | 64204403 | + | 17.0673298068472 | 2.27399148412876 | 3.57968463110397 | 10.6710864697714 |
| chr11 | 64204430 | 64204488 | + | 64.2534769198953 | 26.1509020674808 | 26.2510206280958 | 28.8119334683827 |
| chr11 | 64234172 | 64234214 | + | 2.00792115374673 | 2.27399148412876 | 3.57968463110397 | 2.13421729395427 |
| chr11 | 64234240 | 64234262 | + | 7.02772403811355 | 7.95897019445067 | 7.15936926220794 | 6.40265188186281 |
| chr11 | 64238728 | 64238767 | + | 4.01584230749346 | 5.68497871032191 | 3.57968463110397 | 4.26843458790854 |
| chr11 | 64244041 | 64244073 | + | 97.3841759567164 | 88.6856678810218 | 62.0478669391355 | 114.180625226554 |
| chr11 | 64245903 | 64245993 | + | 924.647691300369 | 744.73221105217 | 47.7291284147196 | 73.6304966414224 |
| chr11 | 64247023 | 64247113 | + | 48.1901076899215 | 46.6168254246397 | 54.8884976769275 | 105.643756050736 |
| chr11 | 64267606 | 64267614 | + | 3.01188173062009 | 1.13699574206438 | 8.35259747257593 | 5.33554323488568 |
| chr11 | 64267821 | 64267825 | + | 4.01584230749346 | 1.13699574206438 | 1.19322821036799 | 1.06710864697714 |
| chr11 | 64267849 | 64267860 | + | 5.01980288436682 | 3.41098722619315 | 1.19322821036799 | 4.26843458790854 |
| chr11 | 64316101 | 64316103 | + | 2.00792115374673 | 1.13699574206438 | 1.19322821036799 | 1.06710864697714 |
| chr11 | 64316134 | 64316158 | + | 21.0831721143407 | 4.54798296825753 | 13.1255103140479 | 26.6777161744284 |
| chr11 | 64316218 | 64316300 | + | 33.130699036821 | 21.6029190992233 | 29.8307052591997 | 20.2750642925656 |
| chr11 | 64316711 | 64316735 | + | 35.1386201905678 | 29.5618892936739 | 10.7390538933119 | 17.0737383516342 |
| chr11 | 64316731 | 64316772 | - | 42.1663442286813 | 48.8908169087684 | 20.2848795762558 | 10.6710864697714 |
| chr11 | 64316804 | 64316852 | - | 18.0712903837206 | 13.6439489047726 | 8.35259747257593 | 9.60397782279422 |
| chr11 | 64317094 | 64317153 | - | 262.033710563948 | 192.152280408881 | 108.583767143487 | 106.710864697714 |
| chr11 | 64317269 | 64317360 | - | 246.974301910848 | 158.042408146949 | 169.438405872254 | 153.663645164708 |
| chr11 | 64321742 | 64321785 | + | 38.1505019211879 | 30.6988850357383 | 5.96614105183995 | 11.7381951167485 |
| chr11 | 64357499 | 64357503 | + | 4.01584230749346 | 9.09596593651506 | 5.96614105183995 | 6.40265188186281 |
| chr11 | 64371513 | 64371579 | + | 38.1505019211879 | 23.876910583352 | 4.77291284147196 | 2.13421729395427 |
| chr11 | 64371618 | 64371643 | + | 13.0514874993537 | 5.68497871032191 | 2.38645642073598 | 16.006629704657 |
| chr11 | 64727065 | 64727066 | - | 3.01188173062009 | 1.13699574206438 | 3.57968463110397 | 1.06710864697714 |
| chr11 | 64764625 | 64764674 | - | 15.0594086531005 | 20.4659233571589 | 19.0916513658878 | 21.3421729395427 |
| chr11 | 64764696 | 64764742 | - | 3.01188173062009 | 2.27399148412876 | 4.77291284147196 | 4.26843458790854 |
| chr11 | 64764791 | 64764820 | - | 16.0633692299738 | 5.68497871032191 | 1.19322821036799 | 14.9395210576799 |
| chr11 | 64764866 | 64764886 | - | 2.00792115374673 | 3.41098722619315 | 2.38645642073598 | 3.20132594093141 |
| chr11 | 64764922 | 64764974 | - | 4.01584230749346 | 3.41098722619315 | 2.38645642073598 | 1.06710864697714 |
| chr11 | 64765190 | 64765191 | - | 4.01584230749346 | 1.13699574206438 | 2.38645642073598 | 1.06710864697714 |
| chr11 | 64765345 | 64765506 | - | 136.538638454778 | 88.6856678810218 | 176.597775134462 | 210.220403454496 |
| chr11 | 64765567 | 64765597 | - | 11.043566345607 | 12.5069531627082 | 31.0239334695677 | 25.6106075274513 |
| chr11 | 64765633 | 64765646 | - | 6.02376346124019 | 2.27399148412876 | 3.57968463110397 | 12.8053037637256 |
| chr11 | 64765698 | 64765733 | - | 5.01980288436682 | 2.27399148412876 | 7.15936926220794 | 22.4092815865198 |
| chr11 | 64765759 | 64765762 | - | 1.00396057687336 | 1.13699574206438 | 1.19322821036799 | 16.006629704657 |
| chr11 | 64765784 | 64765896 | - | 44.174265382428 | 13.6439489047726 | 46.5359002043516 | 45.8856718200168 |
| chr11 | 64789362 | 64789377 | - | 12.0475269224804 | 5.68497871032191 | 2.38645642073598 | 2.13421729395427 |
| chr11 | 64803537 | 64803546 | - | 1.00396057687336 | 3.41098722619315 | 3.57968463110397 | 4.26843458790854 |
| chr11 | 64803575 | 64803637 | - | 27.1069355755808 | 18.1919318730301 | 16.7051949451519 | 16.006629704657 |
| chr11 | 64823912 | 64823924 | - | 3.01188173062009 | 2.27399148412876 | 5.96614105183995 | 5.33554323488568 |
| chr11 | 64823955 | 64823988 | - | 6.02376346124019 | 6.82197445238629 | 13.1255103140479 | 10.6710864697714 |
| chr11 | 64824100 | 64824137 | - | 6.02376346124019 | 5.68497871032191 | 15.5119667347839 | 12.8053037637256 |
| chr11 | 64824202 | 64824204 | - | 4.01584230749346 | 2.27399148412876 | 5.96614105183995 | 1.06710864697714 |
| chr11 | 64852757 | 64852803 | - | 8.03168461498692 | 4.54798296825753 | 5.96614105183995 | 8.53686917581709 |
| chr11 | 64852825 | 64852854 | - | 3.01188173062009 | 1.13699574206438 | 1.19322821036799 | 6.40265188186281 |
| chr11 | 64853678 | 64853680 | - | 1.00396057687336 | 1.13699574206438 | 2.38645642073598 | 2.13421729395427 |
| chr11 | 64854272 | 64854292 | - | 5.01980288436682 | 1.13699574206438 | 1.19322821036799 | 1.06710864697714 |
| chr11 | 64894618 | 64894649 | - | 5.01980288436682 | 9.09596593651506 | 4.77291284147196 | 4.26843458790854 |
| chr11 | 64894796 | 64894816 | - | 9.03564519186028 | 2.27399148412876 | 3.57968463110397 | 3.20132594093141 |
| chr11 | 64894947 | 64894967 | - | 9.03564519186028 | 7.95897019445067 | 3.57968463110397 | 3.20132594093141 |
| chr11 | 64934446 | 64934449 | + | 5.01980288436682 | 19.3289276150945 | 22.6713359969918 | 13.8724124107028 |
| chr11 | 65018568 | 65018647 | + | 30.1188173062009 | 12.5069531627082 | 23.8645642073598 | 21.3421729395427 |
| chr11 | 65021885 | 65021941 | + | 29.1148567293276 | 10.2329616785794 | 35.7968463110397 | 37.3488026441998 |
| chr11 | 65022020 | 65022085 | + | 95.3762548029696 | 50.0278126508328 | 53.6952694665595 | 81.1002571702623 |
| chr11 | 65022108 | 65022166 | + | 28.1108961524542 | 31.8358807778027 | 25.0577924177278 | 23.476390233497 |
| chr11 | 65027404 | 65027552 | + | 189.748549029066 | 145.535454984241 | 68.0140079909754 | 114.180625226554 |
| chr11 | 65034879 | 65034888 | + | 3.01188173062009 | 2.27399148412876 | 1.19322821036799 | 1.06710864697714 |
| chr11 | 65040429 | 65040442 | + | 6.02376346124019 | 9.09596593651506 | 2.38645642073598 | 6.40265188186281 |
| chr11 | 65040488 | 65040537 | + | 15.0594086531005 | 22.7399148412876 | 9.54582568294392 | 8.53686917581709 |
| chr11 | 65040541 | 65040544 | + | 1.00396057687336 | 3.41098722619315 | 1.19322821036799 | 1.06710864697714 |
| chr11 | 65044729 | 65044811 | + | 18.0712903837206 | 31.8358807778027 | 5.96614105183995 | 13.8724124107028 |
| chr11 | 65077478 | 65077531 | - | 13.0514874993537 | 18.1919318730301 | 8.35259747257593 | 6.40265188186281 |
| chr11 | 65077553 | 65077569 | - | 6.02376346124019 | 21.6029190992233 | 7.15936926220794 | 3.20132594093141 |
| chr11 | 65077612 | 65077636 | - | 8.03168461498692 | 13.6439489047726 | 1.19322821036799 | 5.33554323488568 |
| chr11 | 65077676 | 65077690 | - | 6.02376346124019 | 14.780944646837 | 7.15936926220794 | 7.46976052883995 |
| chr11 | 65077722 | 65077929 | - | 259.021828833328 | 272.878978095452 | 115.743136405695 | 197.41509969077 |
| chr11 | 65077954 | 65077986 | - | 11.043566345607 | 9.09596593651506 | 20.2848795762558 | 26.6777161744284 |
| chr11 | 65078015 | 65078040 | - | 12.0475269224804 | 6.82197445238629 | 4.77291284147196 | 10.6710864697714 |
| chr11 | 65078090 | 65078098 | - | 15.0594086531005 | 15.9179403889013 | 25.0577924177278 | 16.006629704657 |
| chr11 | 65078124 | 65078143 | - | 1.00396057687336 | 5.68497871032191 | 2.38645642073598 | 4.26843458790854 |
| chr11 | 65078241 | 65078270 | - | 16.0633692299738 | 17.0549361309657 | 13.1255103140479 | 11.7381951167485 |
| chr11 | 65078330 | 65078405 | - | 31.1227778830743 | 25.0139063254164 | 10.7390538933119 | 25.6106075274513 |
| chr11 | 65078421 | 65078430 | - | 3.01188173062009 | 7.95897019445067 | 5.96614105183995 | 4.26843458790854 |
| chr11 | 65078452 | 65078586 | - | 67.2653586505154 | 60.2607743294122 | 64.4343233598714 | 49.0869977609482 |
| chr11 | 65078603 | 65078641 | - | 12.0475269224804 | 17.0549361309657 | 8.35259747257593 | 20.2750642925656 |
| chr11 | 65078662 | 65078681 | - | 13.0514874993537 | 6.82197445238629 | 5.96614105183995 | 7.46976052883995 |
| chr11 | 65078709 | 65078831 | - | 37.1465413443145 | 43.2058381984465 | 33.4103898903037 | 51.2212150549025 |
| chr11 | 65078989 | 65079029 | - | 12.0475269224804 | 6.82197445238629 | 7.15936926220794 | 3.20132594093141 |
| chr11 | 65079080 | 65079103 | - | 13.0514874993537 | 20.4659233571589 | 9.54582568294392 | 22.4092815865198 |
| chr11 | 65088360 | 65088373 | + | 29.1148567293276 | 65.9457530397342 | 22.6713359969918 | 17.0737383516342 |
| chr11 | 65111581 | 65111609 | + | 3.01188173062009 | 1.13699574206438 | 1.19322821036799 | 2.13421729395427 |
| chr11 | 65111666 | 65111683 | + | 2.00792115374673 | 3.41098722619315 | 9.54582568294392 | 4.26843458790854 |
| chr11 | 65111831 | 65111892 | + | 17.0673298068472 | 9.09596593651506 | 119.322821036799 | 96.0397782279422 |
| chr11 | 65116167 | 65116205 | + | 1.00396057687336 | 6.82197445238629 | 10.7390538933119 | 5.33554323488568 |
| chr11 | 65120657 | 65120801 | - | 138.546559608524 | 128.480518853275 | 153.926439137471 | 208.086186160541 |
| chr11 | 65125594 | 65125603 | + | 6.02376346124019 | 4.54798296825753 | 5.96614105183995 | 4.26843458790854 |
| chr11 | 65125914 | 65125968 | + | 3.01188173062009 | 15.9179403889013 | 4.77291284147196 | 12.8053037637256 |
| chr11 | 65127022 | 65127024 | + | 2.00792115374673 | 4.54798296825753 | 3.57968463110397 | 7.46976052883995 |
| chr11 | 65127076 | 65127093 | + | 9.03564519186028 | 4.54798296825753 | 2.38645642073598 | 4.26843458790854 |
| chr11 | 65127105 | 65127117 | + | 19.0752509605939 | 22.7399148412876 | 25.0577924177278 | 18.1408469986113 |
| chr11 | 65127146 | 65127196 | + | 2.00792115374673 | 7.95897019445067 | 8.35259747257593 | 4.26843458790854 |
| chr11 | 65127231 | 65127254 | + | 3.01188173062009 | 9.09596593651506 | 7.15936926220794 | 2.13421729395427 |
| chr11 | 65127366 | 65127392 | - | 7.02772403811355 | 10.2329616785794 | 5.96614105183995 | 8.53686917581709 |
| chr11 | 65127424 | 65127443 | - | 18.0712903837206 | 21.6029190992233 | 8.35259747257593 | 23.476390233497 |
| chr11 | 65128226 | 65128256 | - | 4.01584230749346 | 1.13699574206438 | 1.19322821036799 | 1.06710864697714 |
| chr11 | 65128283 | 65128307 | - | 4.01584230749346 | 2.27399148412876 | 5.96614105183995 | 2.13421729395427 |
| chr11 | 65211395 | 65211401 | + | 3.01188173062009 | 1.13699574206438 | 3.57968463110397 | 9.60397782279422 |
| chr11 | 65211456 | 65211459 | + | 4.01584230749346 | 1.13699574206438 | 5.96614105183995 | 2.13421729395427 |
| chr11 | 65211490 | 65211505 | + | 6.02376346124019 | 1.13699574206438 | 3.57968463110397 | 3.20132594093141 |
| chr11 | 65211564 | 65211597 | + | 7.02772403811355 | 5.68497871032191 | 20.2848795762558 | 12.8053037637256 |
| chr11 | 65211814 | 65211839 | + | 5.01980288436682 | 2.27399148412876 | 2.38645642073598 | 3.20132594093141 |
| chr11 | 65211884 | 65211989 | + | 21.0831721143407 | 12.5069531627082 | 13.1255103140479 | 40.5501285851312 |
| chr11 | 65297544 | 65297551 | + | 6.02376346124019 | 17.0549361309657 | 5.96614105183995 | 6.40265188186281 |
| chr11 | 65297941 | 65297946 | + | 5.01980288436682 | 2.27399148412876 | 3.57968463110397 | 3.20132594093141 |
| chr11 | 65321984 | 65322020 | + | 44.174265382428 | 57.9867828452835 | 14.3187385244159 | 8.53686917581709 |
| chr11 | 65322048 | 65322065 | + | 13.0514874993537 | 13.6439489047726 | 5.96614105183995 | 1.06710864697714 |
| chr11 | 65322375 | 65322388 | + | 4.01584230749346 | 10.2329616785794 | 1.19322821036799 | 1.06710864697714 |
| chr11 | 65352826 | 65352827 | + | 2.00792115374673 | 2.27399148412876 | 2.38645642073598 | 1.06710864697714 |
| chr11 | 65352874 | 65352911 | + | 22.087132691214 | 27.2878978095452 | 13.1255103140479 | 19.2079556455884 |
| chr11 | 65411376 | 65411463 | + | 16.0633692299738 | 3.41098722619315 | 20.2848795762558 | 13.8724124107028 |
| chr11 | 65412267 | 65412302 | + | 3.01188173062009 | 1.13699574206438 | 17.8984231555198 | 11.7381951167485 |
| chr11 | 65412762 | 65412809 | + | 5.01980288436682 | 2.27399148412876 | 8.35259747257593 | 17.0737383516342 |
| chr11 | 65413315 | 65413371 | + | 11.043566345607 | 12.5069531627082 | 3.57968463110397 | 14.9395210576799 |
| chr11 | 65422792 | 65423198 | + | 7662.22712269752 | 13995.2805890705 | 1711.0892536677 | 3626.03518242831 |
| chr11 | 65423223 | 65423252 | + | 7.02772403811355 | 29.5618892936739 | 2.38645642073598 | 2.13421729395427 |
| chr11 | 65423590 | 65423665 | + | 17.0673298068472 | 22.7399148412876 | 2.38645642073598 | 16.006629704657 |
| chr11 | 65423739 | 65423905 | + | 114.451505763564 | 188.741293182687 | 27.4442488384638 | 26.6777161744284 |
| chr11 | 65423954 | 65424073 | + | 37.1465413443145 | 61.3977700714766 | 8.35259747257593 | 10.6710864697714 |
| chr11 | 65424091 | 65424111 | + | 9.03564519186028 | 26.1509020674808 | 7.15936926220794 | 4.26843458790854 |
| chr11 | 65424142 | 65424204 | + | 17.0673298068472 | 38.657855230189 | 8.35259747257593 | 3.20132594093141 |
| chr11 | 65424290 | 65424317 | + | 71.2812009580089 | 88.6856678810218 | 8.35259747257593 | 8.53686917581709 |
| chr11 | 65424351 | 65425085 | + | 2204.69742681391 | 3433.72714103443 | 477.291284147196 | 880.364633756137 |
| chr11 | 65425140 | 65425179 | + | 18.0712903837206 | 25.0139063254164 | 3.57968463110397 | 3.20132594093141 |
| chr11 | 65425219 | 65425254 | + | 13.0514874993537 | 31.8358807778027 | 7.15936926220794 | 13.8724124107028 |
| chr11 | 65425284 | 65425347 | + | 52.205949997415 | 128.480518853275 | 23.8645642073598 | 32.0132594093141 |
| chr11 | 65425369 | 65425418 | + | 60.2376346124019 | 133.028501821533 | 7.15936926220794 | 35.2145853502455 |
| chr11 | 65425441 | 65425552 | + | 87.3445701879827 | 123.932535885018 | 35.7968463110397 | 86.435800405148 |
| chr11 | 65425574 | 65425980 | + | 172.681219222219 | 363.838637460602 | 159.892580189311 | 151.529427870753 |
| chr11 | 65426012 | 65426157 | + | 209.827760566533 | 402.496492690791 | 42.9562155732476 | 98.1739955218965 |
| chr11 | 65426174 | 65426246 | + | 50.1980288436682 | 92.0966551072149 | 11.9322821036799 | 20.2750642925656 |
| chr11 | 65426265 | 65426468 | + | 329.299069214464 | 666.279504849728 | 70.4004644117114 | 147.260993282845 |
| chr11 | 65426489 | 65426509 | + | 5.01980288436682 | 13.6439489047726 | 1.19322821036799 | 1.06710864697714 |
| chr11 | 65426946 | 65426972 | + | 1.00396057687336 | 1.13699574206438 | 4.77291284147196 | 1.06710864697714 |
| chr11 | 65437245 | 65437320 | + | 1.00396057687336 | 2.27399148412876 | 4.77291284147196 | 4.26843458790854 |
| chr11 | 65439757 | 65439801 | + | 1.00396057687336 | 1.13699574206438 | 11.9322821036799 | 9.60397782279422 |
| chr11 | 65440931 | 65440986 | + | 4.01584230749346 | 2.27399148412876 | 3.57968463110397 | 1.06710864697714 |
| chr11 | 65441263 | 65441285 | + | 1.00396057687336 | 3.41098722619315 | 4.77291284147196 | 2.13421729395427 |
| chr11 | 65497737 | 65497845 | + | 22.087132691214 | 23.876910583352 | 94.2650286190712 | 107.777973344691 |
| chr11 | 65498208 | 65498209 | + | 28.1108961524542 | 23.876910583352 | 2.38645642073598 | 5.33554323488568 |
| chr11 | 65498620 | 65498624 | + | 4.01584230749346 | 3.41098722619315 | 4.77291284147196 | 3.20132594093141 |
| chr11 | 65498657 | 65498715 | + | 21.0831721143407 | 18.1919318730301 | 14.3187385244159 | 11.7381951167485 |
| chr11 | 65498838 | 65498852 | + | 3.01188173062009 | 2.27399148412876 | 2.38645642073598 | 1.06710864697714 |
| chr11 | 65498937 | 65501254 | + | 311773.933384562 | 238408.678183286 | 275503.268263655 | 172287.892380399 |
| chr11 | 65501276 | 65502809 | + | 18582.3063173491 | 17433.5557130732 | 17122.8248187806 | 13103.0270762322 |
| chr11 | 65502833 | 65502949 | + | 99.3920971104631 | 118.247557174696 | 106.197310722751 | 65.0936274656053 |
| chr11 | 65502975 | 65503411 | + | 890.513031686674 | 808.403972607776 | 690.879133803066 | 363.884048619203 |
| chr11 | 65503432 | 65503870 | + | 1725.80823164531 | 1561.0951538544 | 1351.92756234693 | 693.620620535138 |
| chr11 | 65503903 | 65503987 | + | 40.1584230749346 | 35.2468680039958 | 53.6952694665595 | 33.0803680562912 |
| chr11 | 65504020 | 65504059 | + | 6.02376346124019 | 5.68497871032191 | 14.3187385244159 | 4.26843458790854 |
| chr11 | 65504084 | 65504869 | + | 9919.13049950884 | 8804.89502654657 | 5658.28817356501 | 4030.46935963264 |
| chr11 | 65504907 | 65505039 | + | 292.152527870149 | 221.714169702554 | 190.916513658878 | 132.321472225165 |
| chr11 | 65505051 | 65505068 | + | 8.03168461498692 | 5.68497871032191 | 4.77291284147196 | 12.8053037637256 |
| chr11 | 65505098 | 65505118 | + | 4.01584230749346 | 9.09596593651506 | 8.35259747257593 | 13.8724124107028 |
| chr11 | 65505134 | 65505163 | + | 580.289213432805 | 382.030569333632 | 251.771152387646 | 289.186443330804 |
| chr11 | 65505251 | 65505298 | + | 11.043566345607 | 5.68497871032191 | 2.38645642073598 | 1.06710864697714 |
| chr11 | 65505347 | 65505364 | + | 5.01980288436682 | 9.09596593651506 | 1.19322821036799 | 2.13421729395427 |
| chr11 | 65505467 | 65505485 | + | 28.1108961524542 | 26.1509020674808 | 7.15936926220794 | 10.6710864697714 |
| chr11 | 65505618 | 65505709 | + | 50.1980288436682 | 62.534765813541 | 46.5359002043516 | 35.2145853502455 |
| chr11 | 65505738 | 65505742 | + | 2.00792115374673 | 1.13699574206438 | 5.96614105183995 | 16.006629704657 |
| chr11 | 65505836 | 65505872 | + | 15.0594086531005 | 9.09596593651506 | 14.3187385244159 | 8.53686917581709 |
| chr11 | 65506115 | 65506117 | + | 66.2613980736421 | 25.0139063254164 | 21.4781077866238 | 56.5567582897882 |
| chr11 | 65506150 | 65506202 | + | 75.2970432655023 | 76.1787147183136 | 72.7869208324474 | 17.0737383516342 |
| chr11 | 65506229 | 65506423 | + | 1314.18439512723 | 1108.57084851277 | 1209.93340531314 | 915.579219106382 |
| chr11 | 65538550 | 65538597 | + | 4.01584230749346 | 2.27399148412876 | 1.19322821036799 | 5.33554323488568 |
| chr11 | 65538640 | 65538686 | + | 22.087132691214 | 39.7948509722534 | 22.6713359969918 | 27.7448248214055 |
| chr11 | 65538856 | 65538914 | - | 7.02772403811355 | 9.09596593651506 | 7.15936926220794 | 10.6710864697714 |
| chr11 | 65571695 | 65571750 | + | 62.2455557661486 | 34.1098722619315 | 51.3088130458235 | 90.7042349930565 |
| chr11 | 65572511 | 65572573 | + | 105.415860571703 | 72.7677274921204 | 44.1494437836156 | 76.8318225823538 |
| chr11 | 65573761 | 65573780 | + | 3.01188173062009 | 3.41098722619315 | 1.19322821036799 | 8.53686917581709 |
| chr11 | 65574146 | 65574168 | + | 11.043566345607 | 14.780944646837 | 7.15936926220794 | 13.8724124107028 |
| chr11 | 65592558 | 65592626 | + | 9.03564519186028 | 11.3699574206438 | 14.3187385244159 | 13.8724124107028 |
| chr11 | 65597783 | 65597824 | - | 7.02772403811355 | 6.82197445238629 | 2.38645642073598 | 10.6710864697714 |
| chr11 | 65597861 | 65597874 | - | 12.0475269224804 | 4.54798296825753 | 5.96614105183995 | 14.9395210576799 |
| chr11 | 65597953 | 65598006 | - | 17.0673298068472 | 6.82197445238629 | 11.9322821036799 | 12.8053037637256 |
| chr11 | 65598036 | 65598066 | - | 5.01980288436682 | 3.41098722619315 | 4.77291284147196 | 1.06710864697714 |
| chr11 | 65598121 | 65598169 | - | 47.1861471130481 | 37.5208594881246 | 56.0817258872955 | 37.3488026441998 |
| chr11 | 65598252 | 65598271 | - | 1.00396057687336 | 1.13699574206438 | 2.38645642073598 | 3.20132594093141 |
| chr11 | 65636360 | 65636361 | + | 3.01188173062009 | 4.54798296825753 | 4.77291284147196 | 7.46976052883995 |
| chr11 | 65636971 | 65636989 | + | 5.01980288436682 | 1.13699574206438 | 3.57968463110397 | 2.13421729395427 |
| chr11 | 65637314 | 65637350 | + | 28.1108961524542 | 14.780944646837 | 8.35259747257593 | 6.40265188186281 |
| chr11 | 65653643 | 65653710 | - | 21.0831721143407 | 19.3289276150945 | 11.9322821036799 | 18.1408469986113 |
| chr11 | 65654248 | 65654253 | - | 1.00396057687336 | 1.13699574206438 | 1.19322821036799 | 1.06710864697714 |
| chr11 | 65654422 | 65654459 | - | 5.01980288436682 | 1.13699574206438 | 1.19322821036799 | 2.13421729395427 |
| chr11 | 65654953 | 65654970 | - | 5.01980288436682 | 4.54798296825753 | 1.19322821036799 | 1.06710864697714 |
| chr11 | 65719464 | 65719482 | + | 1.00396057687336 | 7.95897019445067 | 2.38645642073598 | 8.53686917581709 |
| chr11 | 65719506 | 65719557 | + | 38.1505019211879 | 48.8908169087684 | 20.2848795762558 | 34.1474767032683 |
| chr11 | 65719719 | 65719739 | - | 13.0514874993537 | 19.3289276150945 | 4.77291284147196 | 17.0737383516342 |
| chr11 | 65719757 | 65719799 | - | 4.01584230749346 | 14.780944646837 | 3.57968463110397 | 11.7381951167485 |
| chr11 | 65777655 | 65777658 | - | 2.00792115374673 | 13.6439489047726 | 7.15936926220794 | 12.8053037637256 |
| chr11 | 65854824 | 65854932 | - | 723.855575925696 | 676.512466528307 | 317.398703957885 | 374.555135088975 |
| chr11 | 65854965 | 65854989 | - | 24.0950538449607 | 20.4659233571589 | 10.7390538933119 | 18.1408469986113 |
| chr11 | 65855017 | 65855070 | - | 97.3841759567164 | 60.2607743294122 | 34.6036181006717 | 81.1002571702623 |
| chr11 | 65855095 | 65855102 | - | 3.01188173062009 | 2.27399148412876 | 1.19322821036799 | 8.53686917581709 |
| chr11 | 65855129 | 65855141 | - | 4.01584230749346 | 1.13699574206438 | 3.57968463110397 | 6.40265188186281 |
| chr11 | 65855199 | 65855227 | - | 7.02772403811355 | 1.13699574206438 | 3.57968463110397 | 2.13421729395427 |
| chr11 | 65855255 | 65855340 | - | 25.0990144218341 | 10.2329616785794 | 9.54582568294392 | 33.0803680562912 |
| chr11 | 65866282 | 65866310 | + | 18.0712903837206 | 22.7399148412876 | 11.9322821036799 | 12.8053037637256 |
| chr11 | 65866365 | 65866377 | + | 8.03168461498692 | 18.1919318730301 | 14.3187385244159 | 7.46976052883995 |
| chr11 | 65891449 | 65891464 | + | 7.02772403811355 | 5.68497871032191 | 1.19322821036799 | 3.20132594093141 |
| chr11 | 65891488 | 65891505 | + | 2.00792115374673 | 1.13699574206438 | 2.38645642073598 | 7.46976052883995 |
| chr11 | 65891564 | 65891613 | + | 45.1782259593014 | 56.8497871032191 | 16.7051949451519 | 30.9461507623369 |
| chr11 | 65916834 | 65916867 | - | 2.00792115374673 | 4.54798296825753 | 2.38645642073598 | 4.26843458790854 |
| chr11 | 65916988 | 65917000 | - | 1.00396057687336 | 4.54798296825753 | 1.19322821036799 | 1.06710864697714 |
| chr11 | 65921454 | 65921513 | + | 260.025789410201 | 283.111939774031 | 564.396943504059 | 682.949534065367 |
| chr11 | 65921543 | 65921547 | + | 4.01584230749346 | 1.13699574206438 | 1.19322821036799 | 2.13421729395427 |
| chr11 | 65979011 | 65979062 | + | 6.02376346124019 | 2.27399148412876 | 3.57968463110397 | 2.13421729395427 |
| chr11 | 65996723 | 65996726 | - | 2.00792115374673 | 2.27399148412876 | 1.19322821036799 | 1.06710864697714 |
| chr11 | 65997386 | 65997398 | - | 3.01188173062009 | 2.27399148412876 | 4.77291284147196 | 6.40265188186281 |
| chr11 | 65997966 | 65997974 | - | 3.01188173062009 | 4.54798296825753 | 4.77291284147196 | 1.06710864697714 |
| chr11 | 65998486 | 65998487 | - | 1.00396057687336 | 2.27399148412876 | 1.19322821036799 | 1.06710864697714 |
| chr11 | 66068707 | 66068861 | + | 47.1861471130481 | 88.6856678810218 | 39.3765309421437 | 41.6172372321083 |
| chr11 | 66243436 | 66243455 | + | 4.01584230749346 | 1.13699574206438 | 2.38645642073598 | 1.06710864697714 |
| chr11 | 66244173 | 66244186 | + | 16.0633692299738 | 7.95897019445067 | 2.38645642073598 | 2.13421729395427 |
| chr11 | 66244489 | 66244530 | + | 12.0475269224804 | 10.2329616785794 | 7.15936926220794 | 4.26843458790854 |
| chr11 | 66244557 | 66244575 | + | 2.00792115374673 | 4.54798296825753 | 4.77291284147196 | 2.13421729395427 |
| chr11 | 66244676 | 66244709 | + | 13.0514874993537 | 15.9179403889013 | 4.77291284147196 | 21.3421729395427 |
| chr11 | 66276220 | 66276243 | + | 9.03564519186028 | 4.54798296825753 | 8.35259747257593 | 2.13421729395427 |
| chr11 | 66276255 | 66276314 | + | 22.087132691214 | 15.9179403889013 | 23.8645642073598 | 11.7381951167485 |
| chr11 | 66276349 | 66276362 | + | 10.0396057687336 | 2.27399148412876 | 7.15936926220794 | 6.40265188186281 |
| chr11 | 66276471 | 66276533 | + | 26.1029749987075 | 19.3289276150945 | 20.2848795762558 | 24.5434988804741 |
| chr11 | 66277416 | 66277426 | + | 2.00792115374673 | 3.41098722619315 | 2.38645642073598 | 2.13421729395427 |
| chr11 | 66284600 | 66284697 | - | 48.1901076899215 | 39.7948509722534 | 28.6374770488317 | 72.5633879944452 |
| chr11 | 66337728 | 66337754 | - | 4.01584230749346 | 4.54798296825753 | 1.19322821036799 | 1.06710864697714 |
| chr11 | 66345702 | 66345816 | - | 6.02376346124019 | 1.13699574206438 | 4.77291284147196 | 1.06710864697714 |
| chr11 | 66345907 | 66345997 | - | 1.00396057687336 | 1.13699574206438 | 1.19322821036799 | 1.06710864697714 |
| chr11 | 66362592 | 66362602 | - | 8.03168461498692 | 13.6439489047726 | 8.35259747257593 | 22.4092815865198 |
| chr11 | 66435856 | 66435879 | - | 57.2257528817818 | 73.9047232341848 | 48.9223566250876 | 81.1002571702623 |
| chr11 | 66435971 | 66435991 | - | 2.00792115374673 | 1.13699574206438 | 2.38645642073598 | 1.06710864697714 |
| chr11 | 66436024 | 66436049 | - | 4.01584230749346 | 3.41098722619315 | 1.19322821036799 | 3.20132594093141 |
| chr11 | 66509377 | 66509384 | + | 1.00396057687336 | 1.13699574206438 | 2.38645642073598 | 6.40265188186281 |
| chr11 | 66536136 | 66536144 | - | 32.1267384599477 | 37.5208594881246 | 44.1494437836156 | 49.0869977609482 |
| chr11 | 66536292 | 66536298 | - | 7.02772403811355 | 1.13699574206438 | 2.38645642073598 | 1.06710864697714 |
| chr11 | 66539286 | 66539307 | - | 10.0396057687336 | 12.5069531627082 | 1.19322821036799 | 3.20132594093141 |
| chr11 | 66563928 | 66563929 | - | 2.00792115374673 | 1.13699574206438 | 1.19322821036799 | 1.06710864697714 |
| chr11 | 66605911 | 66605953 | + | 31.1227778830743 | 27.2878978095452 | 11.9322821036799 | 14.9395210576799 |
| chr11 | 66605973 | 66605997 | + | 2.00792115374673 | 7.95897019445067 | 4.77291284147196 | 1.06710864697714 |
| chr11 | 66624268 | 66624310 | + | 6.02376346124019 | 11.3699574206438 | 5.96614105183995 | 1.06710864697714 |
| chr11 | 66624327 | 66624396 | + | 25.0990144218341 | 26.1509020674808 | 28.6374770488317 | 18.1408469986113 |
| chr11 | 66626623 | 66626656 | + | 2.00792115374673 | 2.27399148412876 | 7.15936926220794 | 7.46976052883995 |
| chr11 | 66626662 | 66626665 | + | 3.01188173062009 | 2.27399148412876 | 1.19322821036799 | 1.06710864697714 |
| chr11 | 66626677 | 66626719 | + | 10.0396057687336 | 12.5069531627082 | 27.4442488384638 | 12.8053037637256 |
| chr11 | 66626765 | 66626812 | + | 19.0752509605939 | 12.5069531627082 | 13.1255103140479 | 13.8724124107028 |
| chr11 | 66626880 | 66626931 | + | 15.0594086531005 | 17.0549361309657 | 16.7051949451519 | 29.8790421153598 |
| chr11 | 66626959 | 66627074 | + | 31.1227778830743 | 21.6029190992233 | 53.6952694665595 | 61.8923015246739 |
| chr11 | 66627118 | 66627173 | + | 8.03168461498692 | 10.2329616785794 | 7.15936926220794 | 6.40265188186281 |
| chr11 | 66627263 | 66627270 | + | 3.01188173062009 | 5.68497871032191 | 1.19322821036799 | 1.06710864697714 |
| chr11 | 66640756 | 66640759 | + | 5.01980288436682 | 3.41098722619315 | 3.57968463110397 | 1.06710864697714 |
| chr11 | 66646339 | 66646450 | + | 89.3524913417294 | 154.631420920756 | 69.2072362013434 | 55.4896496428111 |
| chr11 | 66665134 | 66665156 | - | 5.01980288436682 | 1.13699574206438 | 3.57968463110397 | 1.06710864697714 |
| chr11 | 66668204 | 66668206 | + | 1.00396057687336 | 2.27399148412876 | 2.38645642073598 | 3.20132594093141 |
| chr11 | 66669206 | 66669214 | - | 1.00396057687336 | 1.13699574206438 | 1.19322821036799 | 4.26843458790854 |
| chr11 | 66685274 | 66685278 | - | 5.01980288436682 | 22.7399148412876 | 2.38645642073598 | 4.26843458790854 |
| chr11 | 66685920 | 66685922 | - | 1.00396057687336 | 2.27399148412876 | 2.38645642073598 | 1.06710864697714 |
| chr11 | 66685950 | 66685958 | - | 8.03168461498692 | 1.13699574206438 | 3.57968463110397 | 1.06710864697714 |
| chr11 | 66846204 | 66846218 | + | 4.01584230749346 | 3.41098722619315 | 1.19322821036799 | 5.33554323488568 |
| chr11 | 66846269 | 66846291 | + | 17.0673298068472 | 11.3699574206438 | 2.38645642073598 | 9.60397782279422 |
| chr11 | 66848555 | 66848616 | - | 17.0673298068472 | 15.9179403889013 | 1.19322821036799 | 2.13421729395427 |
| chr11 | 66860412 | 66860435 | + | 18.0712903837206 | 20.4659233571589 | 3.57968463110397 | 12.8053037637256 |
| chr11 | 67256217 | 67256263 | + | 53.2099105742883 | 42.0688424563821 | 57.2749540976635 | 22.4092815865198 |
| chr11 | 67256369 | 67256413 | + | 8.03168461498692 | 5.68497871032191 | 4.77291284147196 | 1.06710864697714 |
| chr11 | 67256492 | 67256495 | + | 1.00396057687336 | 2.27399148412876 | 2.38645642073598 | 2.13421729395427 |
| chr11 | 67256642 | 67256646 | + | 2.00792115374673 | 3.41098722619315 | 5.96614105183995 | 4.26843458790854 |
| chr11 | 67256789 | 67256798 | + | 2.00792115374673 | 3.41098722619315 | 3.57968463110397 | 1.06710864697714 |
| chr11 | 67257123 | 67257160 | + | 1.00396057687336 | 5.68497871032191 | 2.38645642073598 | 8.53686917581709 |
| chr11 | 67257497 | 67257527 | + | 8.03168461498692 | 1.13699574206438 | 1.19322821036799 | 4.26843458790854 |
| chr11 | 67257555 | 67257580 | + | 37.1465413443145 | 17.0549361309657 | 11.9322821036799 | 17.0737383516342 |
| chr11 | 67285389 | 67285489 | + | 19.0752509605939 | 17.0549361309657 | 21.4781077866238 | 6.40265188186281 |
| chr11 | 67285821 | 67285839 | + | 4.01584230749346 | 6.82197445238629 | 29.8307052591997 | 58.6909755837425 |
| chr11 | 67285922 | 67285960 | + | 6.02376346124019 | 6.82197445238629 | 34.6036181006717 | 9.60397782279422 |
| chr11 | 67285976 | 67286009 | + | 2.00792115374673 | 5.68497871032191 | 7.15936926220794 | 11.7381951167485 |
| chr11 | 67286039 | 67286048 | + | 3.01188173062009 | 3.41098722619315 | 4.77291284147196 | 2.13421729395427 |
| chr11 | 67286408 | 67286533 | + | 34.1346596136944 | 30.6988850357383 | 29.8307052591997 | 39.483019938154 |
| chr11 | 67302344 | 67302371 | + | 10.0396057687336 | 2.27399148412876 | 5.96614105183995 | 4.26843458790854 |
| chr11 | 67302442 | 67302461 | + | 3.01188173062009 | 2.27399148412876 | 2.38645642073598 | 4.26843458790854 |
| chr11 | 67351578 | 67351616 | - | 16.0633692299738 | 26.1509020674808 | 11.9322821036799 | 19.2079556455884 |
| chr11 | 67351750 | 67351808 | - | 2.00792115374673 | 7.95897019445067 | 8.35259747257593 | 19.2079556455884 |
| chr11 | 67398198 | 67398313 | - | 56.2217923049084 | 61.3977700714766 | 32.2171616799357 | 89.6371263460794 |
| chr11 | 67398363 | 67398376 | + | 29.1148567293276 | 47.753821166704 | 14.3187385244159 | 26.6777161744284 |
| chr11 | 67398414 | 67398434 | - | 14.0554480762271 | 5.68497871032191 | 21.4781077866238 | 33.0803680562912 |
| chr11 | 67398462 | 67398469 | - | 5.01980288436682 | 1.13699574206438 | 2.38645642073598 | 2.13421729395427 |
| chr11 | 67435130 | 67435188 | + | 45.1782259593014 | 22.7399148412876 | 83.5259747257593 | 54.4225409958339 |
| chr11 | 67435226 | 67435278 | + | 39.1544624980612 | 18.1919318730301 | 23.8645642073598 | 29.8790421153598 |
| chr11 | 67435341 | 67435376 | + | 4.01584230749346 | 18.1919318730301 | 3.57968463110397 | 11.7381951167485 |
| chr11 | 67438088 | 67438101 | - | 48.1901076899215 | 39.7948509722534 | 38.1833027317757 | 44.8185631730397 |
| chr11 | 67438151 | 67438172 | - | 3.01188173062009 | 1.13699574206438 | 2.38645642073598 | 4.26843458790854 |
| chr11 | 67438210 | 67438218 | - | 1.00396057687336 | 1.13699574206438 | 1.19322821036799 | 1.06710864697714 |
| chr11 | 67464372 | 67464417 | - | 3.01188173062009 | 6.82197445238629 | 7.15936926220794 | 5.33554323488568 |
| chr11 | 67464477 | 67464500 | - | 3.01188173062009 | 1.13699574206438 | 1.19322821036799 | 4.26843458790854 |
| chr11 | 67490902 | 67491006 | + | 299.180251908263 | 272.878978095452 | 102.617626091647 | 105.643756050736 |
| chr11 | 67491052 | 67491071 | + | 7.02772403811355 | 19.3289276150945 | 2.38645642073598 | 5.33554323488568 |
| chr11 | 67491792 | 67491845 | - | 5.01980288436682 | 7.95897019445067 | 3.57968463110397 | 5.33554323488568 |
| chr11 | 67506537 | 67506619 | - | 20.0792115374673 | 35.2468680039958 | 10.7390538933119 | 21.3421729395427 |
| chr11 | 67612474 | 67612514 | + | 9.03564519186028 | 15.9179403889013 | 4.77291284147196 | 6.40265188186281 |
| chr11 | 67628300 | 67628303 | - | 9.03564519186028 | 6.82197445238629 | 2.38645642073598 | 3.20132594093141 |
| chr11 | 68036460 | 68036625 | + | 389.536703826865 | 319.495803520091 | 161.085808399679 | 377.756461029906 |
| chr11 | 68052939 | 68053159 | - | 142.562401916018 | 88.6856678810218 | 26.2510206280958 | 30.9461507623369 |
| chr11 | 68052945 | 68052965 | + | 2.00792115374673 | 3.41098722619315 | 2.38645642073598 | 1.06710864697714 |
| chr11 | 68053585 | 68053659 | - | 13.0514874993537 | 5.68497871032191 | 3.57968463110397 | 6.40265188186281 |
| chr11 | 68053681 | 68053704 | - | 5.01980288436682 | 1.13699574206438 | 7.15936926220794 | 1.06710864697714 |
| chr11 | 68053737 | 68053775 | - | 8.03168461498692 | 6.82197445238629 | 2.38645642073598 | 2.13421729395427 |
| chr11 | 68053814 | 68053950 | - | 47.1861471130481 | 15.9179403889013 | 11.9322821036799 | 16.006629704657 |
| chr11 | 68053977 | 68054010 | - | 32.1267384599477 | 20.4659233571589 | 15.5119667347839 | 13.8724124107028 |
| chr11 | 68155272 | 68155303 | - | 8.03168461498692 | 9.09596593651506 | 1.19322821036799 | 2.13421729395427 |
| chr11 | 68155333 | 68155350 | - | 2.00792115374673 | 5.68497871032191 | 2.38645642073598 | 2.13421729395427 |
| chr11 | 68157028 | 68157034 | - | 5.01980288436682 | 3.41098722619315 | 2.38645642073598 | 8.53686917581709 |
| chr11 | 68165819 | 68165830 | - | 4.01584230749346 | 3.41098722619315 | 2.38645642073598 | 2.13421729395427 |
| chr11 | 68165911 | 68165933 | - | 44.174265382428 | 38.657855230189 | 23.8645642073598 | 36.2816939972226 |
| chr11 | 68165965 | 68165969 | - | 11.043566345607 | 10.2329616785794 | 9.54582568294392 | 14.9395210576799 |
| chr11 | 68166027 | 68166045 | - | 3.01188173062009 | 3.41098722619315 | 3.57968463110397 | 5.33554323488568 |
| chr11 | 68166122 | 68166127 | - | 2.00792115374673 | 1.13699574206438 | 2.38645642073598 | 3.20132594093141 |
| chr11 | 68166254 | 68166298 | - | 1.00396057687336 | 3.41098722619315 | 3.57968463110397 | 11.7381951167485 |
| chr11 | 68166339 | 68166351 | - | 24.0950538449607 | 18.1919318730301 | 46.5359002043516 | 21.3421729395427 |
| chr11 | 68166677 | 68166695 | - | 2.00792115374673 | 2.27399148412876 | 1.19322821036799 | 3.20132594093141 |
| chr11 | 68166958 | 68166983 | - | 1.00396057687336 | 4.54798296825753 | 2.38645642073598 | 4.26843458790854 |
| chr11 | 68261369 | 68261410 | - | 7.02772403811355 | 3.41098722619315 | 1.19322821036799 | 2.13421729395427 |
| chr11 | 68261480 | 68261487 | - | 5.01980288436682 | 4.54798296825753 | 3.57968463110397 | 2.13421729395427 |
| chr11 | 68261543 | 68261585 | - | 6.02376346124019 | 7.95897019445067 | 2.38645642073598 | 4.26843458790854 |
| chr11 | 68261614 | 68261647 | - | 14.0554480762271 | 3.41098722619315 | 1.19322821036799 | 8.53686917581709 |
| chr11 | 68449009 | 68449131 | + | 3.01188173062009 | 3.41098722619315 | 4.77291284147196 | 5.33554323488568 |
| chr11 | 68449167 | 68449245 | + | 33.130699036821 | 10.2329616785794 | 17.8984231555198 | 7.46976052883995 |
| chr11 | 68613170 | 68613182 | + | 54.2138711511617 | 37.5208594881246 | 13.1255103140479 | 11.7381951167485 |
| chr11 | 68613661 | 68613662 | + | 2.00792115374673 | 1.13699574206438 | 1.19322821036799 | 1.06710864697714 |
| chr11 | 68613774 | 68613790 | + | 4.01584230749346 | 4.54798296825753 | 1.19322821036799 | 2.13421729395427 |
| chr11 | 68613862 | 68613888 | + | 13.0514874993537 | 15.9179403889013 | 17.8984231555198 | 8.53686917581709 |
| chr11 | 68614472 | 68614512 | + | 4.01584230749346 | 3.41098722619315 | 7.15936926220794 | 1.06710864697714 |
| chr11 | 68614543 | 68614604 | + | 11.043566345607 | 22.7399148412876 | 14.3187385244159 | 9.60397782279422 |
| chr11 | 68614627 | 68614647 | + | 1.00396057687336 | 1.13699574206438 | 4.77291284147196 | 6.40265188186281 |
| chr11 | 68614683 | 68614701 | + | 4.01584230749346 | 2.27399148412876 | 3.57968463110397 | 2.13421729395427 |
| chr11 | 68614792 | 68614809 | + | 3.01188173062009 | 1.13699574206438 | 1.19322821036799 | 13.8724124107028 |
| chr11 | 68614848 | 68614862 | + | 3.01188173062009 | 5.68497871032191 | 2.38645642073598 | 3.20132594093141 |
| chr11 | 68614897 | 68614928 | + | 14.0554480762271 | 6.82197445238629 | 7.15936926220794 | 5.33554323488568 |
| chr11 | 68614994 | 68615018 | + | 11.043566345607 | 4.54798296825753 | 4.77291284147196 | 7.46976052883995 |
| chr11 | 68615072 | 68615102 | + | 7.02772403811355 | 4.54798296825753 | 1.19322821036799 | 6.40265188186281 |
| chr11 | 68615130 | 68615206 | + | 14.0554480762271 | 25.0139063254164 | 9.54582568294392 | 25.6106075274513 |
| chr11 | 68690979 | 68691015 | + | 58.2297134586551 | 22.7399148412876 | 21.4781077866238 | 48.0198891139711 |
| chr11 | 68691056 | 68691111 | + | 56.2217923049084 | 54.5757956190903 | 34.6036181006717 | 55.4896496428111 |
| chr11 | 68691151 | 68691156 | + | 58.2297134586551 | 92.0966551072149 | 33.4103898903037 | 43.7514545260626 |
| chr11 | 68736321 | 68736325 | - | 1.00396057687336 | 1.13699574206438 | 5.96614105183995 | 9.60397782279422 |
| chr11 | 68754928 | 68755010 | - | 12.0475269224804 | 5.68497871032191 | 1.19322821036799 | 1.06710864697714 |
| chr11 | 68755062 | 68755072 | - | 36.1425807674411 | 38.657855230189 | 2.38645642073598 | 11.7381951167485 |
| chr11 | 68891305 | 68891389 | - | 34.1346596136944 | 36.3838637460602 | 8.35259747257593 | 14.9395210576799 |
| chr11 | 69416228 | 69416233 | + | 23.0910932680874 | 21.6029190992233 | 7.15936926220794 | 17.0737383516342 |
| chr11 | 69651297 | 69651413 | + | 3.01188173062009 | 2.27399148412876 | 5.96614105183995 | 6.40265188186281 |
| chr11 | 69652414 | 69652515 | + | 6.02376346124019 | 3.41098722619315 | 4.77291284147196 | 3.20132594093141 |
| chr11 | 69653054 | 69653225 | + | 6.02376346124019 | 1.13699574206438 | 2.38645642073598 | 5.33554323488568 |
| chr11 | 69653780 | 69653887 | + | 2.00792115374673 | 3.41098722619315 | 3.57968463110397 | 2.13421729395427 |
| chr11 | 69654334 | 69654448 | + | 1.00396057687336 | 1.13699574206438 | 3.57968463110397 | 9.60397782279422 |
| chr11 | 69665672 | 69665682 | - | 11.043566345607 | 18.1919318730301 | 2.38645642073598 | 4.26843458790854 |
| chr11 | 70206474 | 70206475 | + | 3.01188173062009 | 2.27399148412876 | 1.19322821036799 | 2.13421729395427 |
| chr11 | 70206577 | 70206584 | + | 3.01188173062009 | 2.27399148412876 | 2.38645642073598 | 2.13421729395427 |
| chr11 | 70206710 | 70206736 | + | 11.043566345607 | 2.27399148412876 | 1.19322821036799 | 1.06710864697714 |
| chr11 | 70206813 | 70206817 | + | 2.00792115374673 | 2.27399148412876 | 1.19322821036799 | 1.06710864697714 |
| chr11 | 70206841 | 70206871 | + | 14.0554480762271 | 7.95897019445067 | 3.57968463110397 | 18.1408469986113 |
| chr11 | 70207007 | 70207072 | + | 86.3406096111093 | 42.0688424563821 | 4.77291284147196 | 39.483019938154 |
| chr11 | 70207093 | 70207096 | + | 1.00396057687336 | 2.27399148412876 | 1.19322821036799 | 1.06710864697714 |
| chr11 | 70207122 | 70207130 | + | 22.087132691214 | 12.5069531627082 | 1.19322821036799 | 24.5434988804741 |
| chr11 | 70207298 | 70207345 | + | 10.0396057687336 | 20.4659233571589 | 9.54582568294392 | 9.60397782279422 |
| chr11 | 70383959 | 70383967 | + | 3.01188173062009 | 2.27399148412876 | 2.38645642073598 | 1.06710864697714 |
| chr11 | 70384192 | 70384219 | + | 1.00396057687336 | 5.68497871032191 | 4.77291284147196 | 3.20132594093141 |
| chr11 | 70435201 | 70435241 | + | 28.1108961524542 | 23.876910583352 | 65.6275515702394 | 217.690163983336 |
| chr11 | 70435278 | 70435420 | + | 307.21193652325 | 150.083437952498 | 221.940447128446 | 114.180625226554 |
| chr11 | 70435469 | 70435473 | + | 6.02376346124019 | 6.82197445238629 | 9.54582568294392 | 12.8053037637256 |
| chr11 | 70435569 | 70435597 | + | 19.0752509605939 | 6.82197445238629 | 5.96614105183995 | 4.26843458790854 |
| chr11 | 70435676 | 70435702 | + | 5.01980288436682 | 2.27399148412876 | 3.57968463110397 | 10.6710864697714 |
| chr11 | 70435736 | 70435760 | + | 15.0594086531005 | 2.27399148412876 | 7.15936926220794 | 26.6777161744284 |
| chr11 | 70435845 | 70435975 | + | 45.1782259593014 | 13.6439489047726 | 26.2510206280958 | 20.2750642925656 |
| chr11 | 70436021 | 70436034 | + | 8.03168461498692 | 1.13699574206438 | 4.77291284147196 | 3.20132594093141 |
| chr11 | 70436335 | 70436346 | + | 12.0475269224804 | 3.41098722619315 | 1.19322821036799 | 7.46976052883995 |
| chr11 | 70436368 | 70436431 | + | 19.0752509605939 | 14.780944646837 | 15.5119667347839 | 23.476390233497 |
| chr11 | 70436466 | 70436489 | + | 8.03168461498692 | 10.2329616785794 | 10.7390538933119 | 9.60397782279422 |
| chr11 | 71434434 | 71434593 | - | 51.2019894205416 | 77.315710460378 | 99.0379414605431 | 139.791232754005 |
| chr11 | 71434821 | 71434898 | - | 21.0831721143407 | 7.95897019445067 | 28.6374770488317 | 38.4159112911769 |
| chr11 | 71434929 | 71434971 | - | 6.02376346124019 | 7.95897019445067 | 26.2510206280958 | 26.6777161744284 |
| chr11 | 71434993 | 71435027 | - | 1.00396057687336 | 3.41098722619315 | 23.8645642073598 | 13.8724124107028 |
| chr11 | 71435055 | 71435169 | - | 9.03564519186028 | 11.3699574206438 | 54.8884976769275 | 38.4159112911769 |
| chr11 | 71435217 | 71435260 | - | 8.03168461498692 | 3.41098722619315 | 23.8645642073598 | 10.6710864697714 |
| chr11 | 71435345 | 71435394 | - | 4.01584230749346 | 3.41098722619315 | 17.8984231555198 | 12.8053037637256 |
| chr11 | 71448575 | 71448707 | + | 77.3049644192491 | 56.8497871032191 | 357.968463110397 | 229.428359100084 |
| chr11 | 71501476 | 71501483 | + | 5.01980288436682 | 11.3699574206438 | 1.19322821036799 | 2.13421729395427 |
| chr11 | 71996458 | 71996461 | + | 1.00396057687336 | 3.41098722619315 | 1.19322821036799 | 3.20132594093141 |
| chr11 | 71996499 | 71996508 | + | 1.00396057687336 | 11.3699574206438 | 1.19322821036799 | 7.46976052883995 |
| chr11 | 72002887 | 72002960 | - | 94.3722942260963 | 95.5076423334081 | 53.6952694665595 | 70.429170700491 |
| chr11 | 72002995 | 72003018 | - | 12.0475269224804 | 3.41098722619315 | 4.77291284147196 | 9.60397782279422 |
| chr11 | 72003205 | 72003241 | - | 6.02376346124019 | 3.41098722619315 | 10.7390538933119 | 7.46976052883995 |
| chr11 | 72003270 | 72003345 | - | 24.0950538449607 | 21.6029190992233 | 41.7629873628796 | 36.2816939972226 |
| chr11 | 72003397 | 72003450 | - | 4.01584230749346 | 5.68497871032191 | 10.7390538933119 | 30.9461507623369 |
| chr11 | 72003472 | 72003481 | - | 2.00792115374673 | 2.27399148412876 | 1.19322821036799 | 1.06710864697714 |
| chr11 | 72097314 | 72097454 | - | 65.2574374967687 | 71.6307317500561 | 38.1833027317757 | 36.2816939972226 |
| chr11 | 72097508 | 72097533 | - | 15.0594086531005 | 10.2329616785794 | 2.38645642073598 | 10.6710864697714 |
| chr11 | 72097807 | 72097903 | - | 16.0633692299738 | 7.95897019445067 | 2.38645642073598 | 11.7381951167485 |
| chr11 | 72109626 | 72109660 | - | 13.0514874993537 | 14.780944646837 | 2.38645642073598 | 3.20132594093141 |
| chr11 | 72238439 | 72238484 | + | 1.00396057687336 | 1.13699574206438 | 1.19322821036799 | 1.06710864697714 |
| chr11 | 72238779 | 72238794 | + | 2.00792115374673 | 3.41098722619315 | 1.19322821036799 | 3.20132594093141 |
| chr11 | 72238819 | 72238827 | + | 5.01980288436682 | 7.95897019445067 | 1.19322821036799 | 3.20132594093141 |
| chr11 | 72238858 | 72238914 | + | 25.0990144218341 | 26.1509020674808 | 9.54582568294392 | 13.8724124107028 |
| chr11 | 72239023 | 72239078 | + | 46.1821865361748 | 45.4798296825753 | 23.8645642073598 | 22.4092815865198 |
| chr11 | 72685103 | 72685150 | - | 8.03168461498692 | 4.54798296825753 | 14.3187385244159 | 8.53686917581709 |
| chr11 | 72829547 | 72829558 | + | 4.01584230749346 | 11.3699574206438 | 2.38645642073598 | 3.20132594093141 |
| chr11 | 73676300 | 73676321 | - | 24.0950538449607 | 36.3838637460602 | 11.9322821036799 | 13.8724124107028 |
| chr11 | 73676503 | 73676592 | - | 31.1227778830743 | 50.0278126508328 | 81.1395183050233 | 25.6106075274513 |
| chr11 | 73676853 | 73676891 | - | 11.043566345607 | 6.82197445238629 | 19.0916513658878 | 14.9395210576799 |
| chr11 | 73676925 | 73676940 | - | 2.00792115374673 | 1.13699574206438 | 8.35259747257593 | 4.26843458790854 |
| chr11 | 73677058 | 73677076 | - | 3.01188173062009 | 4.54798296825753 | 1.19322821036799 | 1.06710864697714 |
| chr11 | 73677253 | 73677368 | - | 8.03168461498692 | 10.2329616785794 | 7.15936926220794 | 8.53686917581709 |
| chr11 | 73677744 | 73677756 | - | 6.02376346124019 | 5.68497871032191 | 1.19322821036799 | 1.06710864697714 |
| chr11 | 73677889 | 73677934 | - | 3.01188173062009 | 2.27399148412876 | 5.96614105183995 | 3.20132594093141 |
| chr11 | 73864294 | 73864469 | + | 243.962420180228 | 230.81013563907 | 233.872729232126 | 162.200514340525 |
| chr11 | 73864491 | 73864567 | + | 1156.56258455812 | 1119.94080593342 | 788.723847053241 | 411.903937733174 |
| chr11 | 73872683 | 73872721 | - | 17.0673298068472 | 13.6439489047726 | 13.1255103140479 | 24.5434988804741 |
| chr11 | 73872805 | 73872835 | - | 50.1980288436682 | 47.753821166704 | 21.4781077866238 | 39.483019938154 |
| chr11 | 73872859 | 73872867 | - | 8.03168461498692 | 2.27399148412876 | 2.38645642073598 | 14.9395210576799 |
| chr11 | 73872974 | 73873053 | - | 23.0910932680874 | 13.6439489047726 | 4.77291284147196 | 18.1408469986113 |
| chr11 | 73974716 | 73974721 | - | 3.01188173062009 | 1.13699574206438 | 3.57968463110397 | 2.13421729395427 |
| chr11 | 73974955 | 73974961 | - | 9.03564519186028 | 12.5069531627082 | 16.7051949451519 | 22.4092815865198 |
| chr11 | 73975037 | 73975124 | - | 18.0712903837206 | 28.4248935516096 | 9.54582568294392 | 25.6106075274513 |
| chr11 | 74253662 | 74253684 | + | 2.00792115374673 | 1.13699574206438 | 2.38645642073598 | 1.06710864697714 |
| chr11 | 74253871 | 74253882 | + | 2.00792115374673 | 1.13699574206438 | 1.19322821036799 | 2.13421729395427 |
| chr11 | 74253936 | 74253948 | + | 5.01980288436682 | 7.95897019445067 | 19.0916513658878 | 16.006629704657 |
| chr11 | 74253983 | 74253986 | + | 1.00396057687336 | 1.13699574206438 | 4.77291284147196 | 5.33554323488568 |
| chr11 | 74254021 | 74254035 | + | 2.00792115374673 | 4.54798296825753 | 8.35259747257593 | 13.8724124107028 |
| chr11 | 74254072 | 74254094 | + | 4.01584230749346 | 1.13699574206438 | 2.38645642073598 | 8.53686917581709 |
| chr11 | 74254209 | 74254212 | + | 3.01188173062009 | 2.27399148412876 | 4.77291284147196 | 5.33554323488568 |
| chr11 | 74254337 | 74254395 | + | 7.02772403811355 | 12.5069531627082 | 20.2848795762558 | 14.9395210576799 |
| chr11 | 74254477 | 74254577 | + | 365.441649981905 | 264.920007901001 | 264.896662701694 | 361.749831325249 |
| chr11 | 74254626 | 74254642 | + | 9.03564519186028 | 3.41098722619315 | 17.8984231555198 | 10.6710864697714 |
| chr11 | 74493357 | 74493407 | + | 46.1821865361748 | 72.7677274921204 | 28.6374770488317 | 20.2750642925656 |
| chr11 | 74493768 | 74493798 | + | 2.00792115374673 | 3.41098722619315 | 4.77291284147196 | 1.06710864697714 |
| chr11 | 74641550 | 74641575 | + | 23.0910932680874 | 13.6439489047726 | 8.35259747257593 | 5.33554323488568 |
| chr11 | 74641822 | 74641855 | + | 10.0396057687336 | 15.9179403889013 | 4.77291284147196 | 3.20132594093141 |
| chr11 | 74641959 | 74641996 | + | 4.01584230749346 | 7.95897019445067 | 4.77291284147196 | 7.46976052883995 |
| chr11 | 74642387 | 74642388 | + | 4.01584230749346 | 12.5069531627082 | 1.19322821036799 | 2.13421729395427 |
| chr11 | 74642631 | 74642643 | + | 9.03564519186028 | 14.780944646837 | 3.57968463110397 | 1.06710864697714 |
| chr11 | 74836851 | 74836863 | + | 2.00792115374673 | 2.27399148412876 | 1.19322821036799 | 1.06710864697714 |
| chr11 | 74836983 | 74837004 | + | 4.01584230749346 | 10.2329616785794 | 7.15936926220794 | 1.06710864697714 |
| chr11 | 74837034 | 74837038 | + | 31.1227778830743 | 57.9867828452835 | 25.0577924177278 | 9.60397782279422 |
| chr11 | 74837079 | 74837082 | + | 5.01980288436682 | 6.82197445238629 | 3.57968463110397 | 2.13421729395427 |
| chr11 | 74837815 | 74837828 | + | 2.00792115374673 | 10.2329616785794 | 4.77291284147196 | 2.13421729395427 |
| chr11 | 74838788 | 74838796 | + | 15.0594086531005 | 18.1919318730301 | 8.35259747257593 | 16.006629704657 |
| chr11 | 74838819 | 74838823 | + | 4.01584230749346 | 2.27399148412876 | 8.35259747257593 | 1.06710864697714 |
| chr11 | 74838884 | 74838899 | + | 1.00396057687336 | 4.54798296825753 | 7.15936926220794 | 2.13421729395427 |
| chr11 | 74839252 | 74839276 | + | 26.1029749987075 | 13.6439489047726 | 16.7051949451519 | 4.26843458790854 |
| chr11 | 74839420 | 74839446 | + | 6.02376346124019 | 5.68497871032191 | 8.35259747257593 | 2.13421729395427 |
| chr11 | 74839488 | 74839524 | + | 9.03564519186028 | 17.0549361309657 | 11.9322821036799 | 8.53686917581709 |
| chr11 | 74839608 | 74839714 | + | 27.1069355755808 | 29.5618892936739 | 71.5936926220794 | 27.7448248214055 |
| chr11 | 74839733 | 74839739 | + | 1.00396057687336 | 1.13699574206438 | 2.38645642073598 | 4.26843458790854 |
| chr11 | 74839882 | 74839930 | + | 51.2019894205416 | 77.315710460378 | 52.5020412561915 | 37.3488026441998 |
| chr11 | 74840553 | 74840557 | + | 4.01584230749346 | 1.13699574206438 | 9.54582568294392 | 4.26843458790854 |
| chr11 | 74840719 | 74840742 | + | 2.00792115374673 | 4.54798296825753 | 5.96614105183995 | 11.7381951167485 |
| chr11 | 74840856 | 74840860 | + | 1.00396057687336 | 4.54798296825753 | 5.96614105183995 | 7.46976052883995 |
| chr11 | 74840914 | 74840940 | + | 10.0396057687336 | 10.2329616785794 | 9.54582568294392 | 13.8724124107028 |
| chr11 | 74841026 | 74841035 | + | 2.00792115374673 | 1.13699574206438 | 4.77291284147196 | 8.53686917581709 |
| chr11 | 74841066 | 74841089 | + | 3.01188173062009 | 4.54798296825753 | 2.38645642073598 | 2.13421729395427 |
| chr11 | 74841261 | 74841324 | + | 12.0475269224804 | 15.9179403889013 | 32.2171616799357 | 12.8053037637256 |
| chr11 | 74841976 | 74842000 | + | 4.01584230749346 | 7.95897019445067 | 7.15936926220794 | 3.20132594093141 |
| chr11 | 74842141 | 74842155 | + | 2.00792115374673 | 3.41098722619315 | 4.77291284147196 | 3.20132594093141 |
| chr11 | 74842371 | 74842391 | + | 2.00792115374673 | 4.54798296825753 | 1.19322821036799 | 3.20132594093141 |
| chr11 | 74977162 | 74977188 | + | 32.1267384599477 | 31.8358807778027 | 11.9322821036799 | 13.8724124107028 |
| chr11 | 75007438 | 75007440 | + | 5.01980288436682 | 6.82197445238629 | 8.35259747257593 | 2.13421729395427 |
| chr11 | 75008516 | 75008540 | + | 4.01584230749346 | 2.27399148412876 | 5.96614105183995 | 11.7381951167485 |
| chr11 | 75008922 | 75008926 | + | 2.00792115374673 | 3.41098722619315 | 1.19322821036799 | 1.06710864697714 |
| chr11 | 75008950 | 75008958 | + | 3.01188173062009 | 1.13699574206438 | 4.77291284147196 | 4.26843458790854 |
| chr11 | 75009048 | 75009101 | + | 14.0554480762271 | 18.1919318730301 | 23.8645642073598 | 16.006629704657 |
| chr11 | 75010289 | 75010290 | + | 2.00792115374673 | 1.13699574206438 | 1.19322821036799 | 1.06710864697714 |
| chr11 | 75010430 | 75010453 | + | 7.02772403811355 | 6.82197445238629 | 8.35259747257593 | 10.6710864697714 |
| chr11 | 75010774 | 75010785 | + | 2.00792115374673 | 1.13699574206438 | 3.57968463110397 | 3.20132594093141 |
| chr11 | 75265269 | 75265280 | - | 1.00396057687336 | 1.13699574206438 | 1.19322821036799 | 2.13421729395427 |
| chr11 | 75402437 | 75402468 | + | 18.0712903837206 | 17.0549361309657 | 41.7629873628796 | 38.4159112911769 |
| chr11 | 75403492 | 75403494 | + | 4.01584230749346 | 1.13699574206438 | 2.38645642073598 | 3.20132594093141 |
| chr11 | 75404813 | 75404869 | + | 100.396057687336 | 75.0417189762492 | 57.2749540976635 | 113.113516579576 |
| chr11 | 75405630 | 75405669 | + | 587.316937470918 | 959.624406302338 | 533.373010034491 | 375.622243735952 |
| chr11 | 75405725 | 75405728 | + | 4.01584230749346 | 1.13699574206438 | 1.19322821036799 | 2.13421729395427 |
| chr11 | 75405920 | 75405928 | + | 5.01980288436682 | 7.95897019445067 | 3.57968463110397 | 1.06710864697714 |
| chr11 | 75434664 | 75434759 | - | 7.02772403811355 | 2.27399148412876 | 1.19322821036799 | 5.33554323488568 |
| chr11 | 75436954 | 75436966 | - | 1.00396057687336 | 3.41098722619315 | 3.57968463110397 | 6.40265188186281 |
| chr11 | 75441766 | 75441767 | - | 1.00396057687336 | 1.13699574206438 | 3.57968463110397 | 1.06710864697714 |
| chr11 | 75572157 | 75572165 | + | 12.0475269224804 | 2.27399148412876 | 10.7390538933119 | 5.33554323488568 |
| chr11 | 75572188 | 75572248 | + | 28.1108961524542 | 20.4659233571589 | 8.35259747257593 | 8.53686917581709 |
| chr11 | 75572435 | 75572449 | + | 2.00792115374673 | 4.54798296825753 | 4.77291284147196 | 3.20132594093141 |
| chr11 | 75572577 | 75572649 | + | 51.2019894205416 | 28.4248935516096 | 32.2171616799357 | 23.476390233497 |
| chr11 | 75572696 | 75572741 | + | 153.605968261625 | 128.480518853275 | 105.004082512383 | 132.321472225165 |
| chr11 | 76350394 | 76350395 | - | 6.02376346124019 | 14.780944646837 | 9.54582568294392 | 13.8724124107028 |
| chr11 | 76350500 | 76350505 | - | 12.0475269224804 | 15.9179403889013 | 19.0916513658878 | 12.8053037637256 |
| chr11 | 76350626 | 76350641 | - | 4.01584230749346 | 4.54798296825753 | 8.35259747257593 | 3.20132594093141 |
| chr11 | 76350664 | 76350674 | - | 8.03168461498692 | 1.13699574206438 | 7.15936926220794 | 4.26843458790854 |
| chr11 | 76350807 | 76350817 | - | 2.00792115374673 | 10.2329616785794 | 8.35259747257593 | 3.20132594093141 |
| chr11 | 76551261 | 76551267 | + | 18.0712903837206 | 12.5069531627082 | 9.54582568294392 | 12.8053037637256 |
| chr11 | 76797080 | 76797097 | + | 4.01584230749346 | 3.41098722619315 | 3.57968463110397 | 1.06710864697714 |
| chr11 | 76797819 | 76797826 | + | 9.03564519186028 | 11.3699574206438 | 4.77291284147196 | 3.20132594093141 |
| chr11 | 76798064 | 76798071 | + | 6.02376346124019 | 3.41098722619315 | 1.19322821036799 | 2.13421729395427 |
| chr11 | 77020438 | 77020441 | + | 2.00792115374673 | 3.41098722619315 | 2.38645642073598 | 1.06710864697714 |
| chr11 | 77020496 | 77020499 | + | 3.01188173062009 | 3.41098722619315 | 2.38645642073598 | 3.20132594093141 |
| chr11 | 77021252 | 77021271 | + | 10.0396057687336 | 9.09596593651506 | 2.38645642073598 | 1.06710864697714 |
| chr11 | 77022368 | 77022408 | + | 27.1069355755808 | 37.5208594881246 | 16.7051949451519 | 22.4092815865198 |
| chr11 | 77022438 | 77022459 | + | 10.0396057687336 | 10.2329616785794 | 13.1255103140479 | 16.006629704657 |
| chr11 | 77022573 | 77022582 | + | 5.01980288436682 | 4.54798296825753 | 3.57968463110397 | 3.20132594093141 |
| chr11 | 77022636 | 77022653 | + | 3.01188173062009 | 4.54798296825753 | 8.35259747257593 | 2.13421729395427 |
| chr11 | 77023284 | 77023329 | + | 10.0396057687336 | 14.780944646837 | 15.5119667347839 | 9.60397782279422 |
| chr11 | 77616336 | 77616445 | - | 380.501058635005 | 326.317777972478 | 283.988314067581 | 368.152483207112 |
| chr11 | 77616490 | 77616503 | - | 127.502993262917 | 152.357429436627 | 97.8447132501751 | 71.4962793474681 |
| chr11 | 77616531 | 77616553 | - | 3.01188173062009 | 11.3699574206438 | 20.2848795762558 | 13.8724124107028 |
| chr11 | 77616582 | 77616608 | - | 6.02376346124019 | 6.82197445238629 | 28.6374770488317 | 18.1408469986113 |
| chr11 | 77616640 | 77616684 | - | 146.578244223511 | 218.303182476361 | 287.567998698685 | 163.267622987502 |
| chr11 | 77619603 | 77619639 | - | 7.02772403811355 | 6.82197445238629 | 17.8984231555198 | 13.8724124107028 |
| chr11 | 77660405 | 77660412 | - | 5.01980288436682 | 4.54798296825753 | 2.38645642073598 | 3.20132594093141 |
| chr11 | 77661829 | 77661831 | - | 2.00792115374673 | 5.68497871032191 | 2.38645642073598 | 1.06710864697714 |
| chr11 | 77662349 | 77662385 | - | 6.02376346124019 | 4.54798296825753 | 2.38645642073598 | 3.20132594093141 |
| chr11 | 77662490 | 77662492 | - | 4.01584230749346 | 4.54798296825753 | 1.19322821036799 | 1.06710864697714 |
| chr11 | 77662678 | 77662709 | - | 6.02376346124019 | 11.3699574206438 | 3.57968463110397 | 5.33554323488568 |
| chr11 | 77663275 | 77663277 | - | 4.01584230749346 | 4.54798296825753 | 8.35259747257593 | 3.20132594093141 |
| chr11 | 77665340 | 77665378 | - | 15.0594086531005 | 9.09596593651506 | 4.77291284147196 | 11.7381951167485 |
| chr11 | 77666529 | 77666534 | - | 5.01980288436682 | 7.95897019445067 | 9.54582568294392 | 3.20132594093141 |
| chr11 | 77666860 | 77666882 | - | 3.01188173062009 | 4.54798296825753 | 1.19322821036799 | 11.7381951167485 |
| chr11 | 77872329 | 77872332 | + | 1.00396057687336 | 3.41098722619315 | 2.38645642073598 | 1.06710864697714 |
| chr11 | 78069873 | 78070038 | - | 1411.56857108395 | 1008.51522321111 | 695.652046644538 | 444.984305789466 |
| chr11 | 78100965 | 78101050 | - | 19.0752509605939 | 15.9179403889013 | 7.15936926220794 | 10.6710864697714 |
| chr11 | 78101087 | 78101128 | - | 146.578244223511 | 121.658544400889 | 48.9223566250876 | 109.912190638645 |
| chr11 | 78101153 | 78101197 | - | 65.2574374967687 | 38.657855230189 | 21.4781077866238 | 50.1541064079254 |
| chr11 | 78139745 | 78139873 | + | 85.336649034236 | 106.877599754052 | 19.0916513658878 | 41.6172372321083 |
| chr11 | 78214393 | 78214417 | + | 5.01980288436682 | 1.13699574206438 | 8.35259747257593 | 2.13421729395427 |
| chr11 | 78216049 | 78216054 | - | 1.00396057687336 | 1.13699574206438 | 7.15936926220794 | 1.06710864697714 |
| chr11 | 78217705 | 78217710 | - | 1.00396057687336 | 2.27399148412876 | 1.19322821036799 | 3.20132594093141 |
| chr11 | 78218279 | 78218300 | - | 3.01188173062009 | 3.41098722619315 | 2.38645642073598 | 1.06710864697714 |
| chr11 | 78218692 | 78218694 | - | 1.00396057687336 | 3.41098722619315 | 5.96614105183995 | 3.20132594093141 |
| chr11 | 78436264 | 78436319 | - | 5.01980288436682 | 6.82197445238629 | 3.57968463110397 | 5.33554323488568 |
| chr11 | 78436445 | 78436498 | - | 7.02772403811355 | 2.27399148412876 | 9.54582568294392 | 19.2079556455884 |
| chr11 | 82824409 | 82824468 | - | 4.01584230749346 | 3.41098722619315 | 1.19322821036799 | 2.13421729395427 |
| chr11 | 82824544 | 82824563 | - | 9.03564519186028 | 10.2329616785794 | 2.38645642073598 | 1.06710864697714 |
| chr11 | 82824611 | 82824621 | - | 2.00792115374673 | 2.27399148412876 | 2.38645642073598 | 4.26843458790854 |
| chr11 | 82824736 | 82824799 | - | 5.01980288436682 | 2.27399148412876 | 1.19322821036799 | 1.06710864697714 |
| chr11 | 82933029 | 82933036 | + | 1.00396057687336 | 4.54798296825753 | 3.57968463110397 | 12.8053037637256 |
| chr11 | 82933982 | 82933994 | + | 2.00792115374673 | 4.54798296825753 | 2.38645642073598 | 2.13421729395427 |
| chr11 | 82934279 | 82934321 | + | 7.02772403811355 | 11.3699574206438 | 1.19322821036799 | 6.40265188186281 |
| chr11 | 82934358 | 82934402 | + | 124.491111532297 | 129.61751459534 | 47.7291284147196 | 87.5029090521251 |
| chr11 | 82934463 | 82934466 | + | 4.01584230749346 | 6.82197445238629 | 4.77291284147196 | 10.6710864697714 |
| chr11 | 83072053 | 83072149 | + | 444.7545355549 | 590.100790131414 | 350.809093848189 | 495.138412197391 |
| chr11 | 83072621 | 83072640 | + | 6.02376346124019 | 3.41098722619315 | 1.19322821036799 | 4.26843458790854 |
| chr11 | 83072716 | 83072718 | + | 6.02376346124019 | 4.54798296825753 | 2.38645642073598 | 5.33554323488568 |
| chr11 | 83073102 | 83073109 | + | 4.01584230749346 | 2.27399148412876 | 2.38645642073598 | 4.26843458790854 |
| chr11 | 83073318 | 83073323 | + | 1.00396057687336 | 2.27399148412876 | 2.38645642073598 | 1.06710864697714 |
| chr11 | 83073344 | 83073354 | + | 2.00792115374673 | 4.54798296825753 | 2.38645642073598 | 2.13421729395427 |
| chr11 | 83073446 | 83073447 | + | 1.00396057687336 | 5.68497871032191 | 1.19322821036799 | 1.06710864697714 |
| chr11 | 83073471 | 83073474 | + | 2.00792115374673 | 4.54798296825753 | 1.19322821036799 | 1.06710864697714 |
| chr11 | 83073571 | 83073577 | + | 8.03168461498692 | 18.1919318730301 | 2.38645642073598 | 8.53686917581709 |
| chr11 | 83180365 | 83180382 | + | 7.02772403811355 | 4.54798296825753 | 8.35259747257593 | 12.8053037637256 |
| chr11 | 83184950 | 83184952 | + | 2.00792115374673 | 1.13699574206438 | 2.38645642073598 | 3.20132594093141 |
| chr11 | 83185357 | 83185359 | + | 7.02772403811355 | 3.41098722619315 | 1.19322821036799 | 1.06710864697714 |
| chr11 | 83212793 | 83212829 | + | 7.02772403811355 | 7.95897019445067 | 2.38645642073598 | 1.06710864697714 |
| chr11 | 83212951 | 83212959 | + | 170.673298068472 | 112.562578464374 | 165.858721241151 | 106.710864697714 |
| chr11 | 83214377 | 83214379 | + | 4.01584230749346 | 6.82197445238629 | 2.38645642073598 | 1.06710864697714 |
| chr11 | 83261719 | 83261728 | - | 3.01188173062009 | 2.27399148412876 | 3.57968463110397 | 3.20132594093141 |
| chr11 | 83261857 | 83261862 | - | 10.0396057687336 | 18.1919318730301 | 10.7390538933119 | 17.0737383516342 |
| chr11 | 83261959 | 83261968 | - | 4.01584230749346 | 3.41098722619315 | 14.3187385244159 | 5.33554323488568 |
| chr11 | 83265886 | 83265903 | - | 1.00396057687336 | 1.13699574206438 | 3.57968463110397 | 1.06710864697714 |
| chr11 | 83286085 | 83286151 | + | 19.0752509605939 | 40.9318467143178 | 7.15936926220794 | 7.46976052883995 |
| chr11 | 85636074 | 85636100 | + | 6.02376346124019 | 6.82197445238629 | 2.38645642073598 | 5.33554323488568 |
| chr11 | 85636277 | 85636296 | + | 2.00792115374673 | 3.41098722619315 | 5.96614105183995 | 1.06710864697714 |
| chr11 | 85636423 | 85636460 | + | 18.0712903837206 | 20.4659233571589 | 13.1255103140479 | 19.2079556455884 |
| chr11 | 85660274 | 85660276 | - | 5.01980288436682 | 1.13699574206438 | 4.77291284147196 | 1.06710864697714 |
| chr11 | 85660316 | 85660325 | - | 32.1267384599477 | 31.8358807778027 | 27.4442488384638 | 26.6777161744284 |
| chr11 | 85660349 | 85660387 | - | 18.0712903837206 | 13.6439489047726 | 25.0577924177278 | 6.40265188186281 |
| chr11 | 85660414 | 85660428 | - | 4.01584230749346 | 3.41098722619315 | 9.54582568294392 | 3.20132594093141 |
| chr11 | 85660528 | 85660548 | - | 9.03564519186028 | 9.09596593651506 | 3.57968463110397 | 6.40265188186281 |
| chr11 | 85661084 | 85661137 | - | 6.02376346124019 | 3.41098722619315 | 7.15936926220794 | 3.20132594093141 |
| chr11 | 85661169 | 85661244 | - | 16.0633692299738 | 18.1919318730301 | 14.3187385244159 | 4.26843458790854 |
| chr11 | 85661349 | 85661442 | - | 26.1029749987075 | 38.657855230189 | 34.6036181006717 | 7.46976052883995 |
| chr11 | 85661492 | 85661495 | - | 2.00792115374673 | 2.27399148412876 | 5.96614105183995 | 4.26843458790854 |
| chr11 | 85661511 | 85661558 | - | 21.0831721143407 | 26.1509020674808 | 16.7051949451519 | 9.60397782279422 |
| chr11 | 85661643 | 85661652 | - | 1.00396057687336 | 2.27399148412876 | 3.57968463110397 | 1.06710864697714 |
| chr11 | 85661659 | 85661666 | - | 3.01188173062009 | 1.13699574206438 | 1.19322821036799 | 1.06710864697714 |
| chr11 | 85662015 | 85662049 | - | 12.0475269224804 | 7.95897019445067 | 9.54582568294392 | 10.6710864697714 |
| chr11 | 85662371 | 85662420 | - | 20.0792115374673 | 10.2329616785794 | 10.7390538933119 | 5.33554323488568 |
| chr11 | 85662484 | 85662485 | - | 9.03564519186028 | 7.95897019445067 | 13.1255103140479 | 10.6710864697714 |
| chr11 | 85662605 | 85662611 | - | 6.02376346124019 | 13.6439489047726 | 10.7390538933119 | 4.26843458790854 |
| chr11 | 85662779 | 85662784 | - | 3.01188173062009 | 2.27399148412876 | 1.19322821036799 | 1.06710864697714 |
| chr11 | 85662889 | 85662927 | - | 4.01584230749346 | 3.41098722619315 | 4.77291284147196 | 2.13421729395427 |
| chr11 | 85662948 | 85662953 | - | 6.02376346124019 | 1.13699574206438 | 1.19322821036799 | 1.06710864697714 |
| chr11 | 85663049 | 85663055 | - | 2.00792115374673 | 3.41098722619315 | 2.38645642073598 | 2.13421729395427 |
| chr11 | 85663072 | 85663121 | - | 12.0475269224804 | 18.1919318730301 | 7.15936926220794 | 9.60397782279422 |
| chr11 | 85663142 | 85663210 | - | 21.0831721143407 | 17.0549361309657 | 8.35259747257593 | 10.6710864697714 |
| chr11 | 85663246 | 85663260 | - | 2.00792115374673 | 1.13699574206438 | 1.19322821036799 | 3.20132594093141 |
| chr11 | 85663523 | 85663524 | - | 1.00396057687336 | 2.27399148412876 | 1.19322821036799 | 2.13421729395427 |
| chr11 | 85663557 | 85663563 | - | 7.02772403811355 | 3.41098722619315 | 2.38645642073598 | 1.06710864697714 |
| chr11 | 85663589 | 85663592 | - | 5.01980288436682 | 2.27399148412876 | 5.96614105183995 | 3.20132594093141 |
| chr11 | 85663618 | 85663626 | - | 1.00396057687336 | 3.41098722619315 | 1.19322821036799 | 1.06710864697714 |
| chr11 | 85663710 | 85663714 | - | 1.00396057687336 | 1.13699574206438 | 2.38645642073598 | 11.7381951167485 |
| chr11 | 85663736 | 85663739 | - | 4.01584230749346 | 4.54798296825753 | 2.38645642073598 | 3.20132594093141 |
| chr11 | 85663804 | 85663923 | - | 110.43566345607 | 97.7816338175368 | 79.9462900946553 | 48.0198891139711 |
| chr11 | 85957714 | 85957736 | - | 9.03564519186028 | 21.6029190992233 | 32.2171616799357 | 23.476390233497 |
| chr11 | 85957780 | 85957928 | - | 49.1940682667949 | 50.0278126508328 | 56.0817258872955 | 131.254363578188 |
| chr11 | 85957953 | 85957961 | - | 15.0594086531005 | 19.3289276150945 | 4.77291284147196 | 16.006629704657 |
| chr11 | 85958014 | 85958049 | - | 9.03564519186028 | 9.09596593651506 | 13.1255103140479 | 32.0132594093141 |
| chr11 | 85958441 | 85958442 | - | 1.00396057687336 | 1.13699574206438 | 1.19322821036799 | 1.06710864697714 |
| chr11 | 85958739 | 85958758 | - | 1.00396057687336 | 1.13699574206438 | 2.38645642073598 | 8.53686917581709 |
| chr11 | 85959035 | 85959083 | - | 8.03168461498692 | 6.82197445238629 | 27.4442488384638 | 17.0737383516342 |
| chr11 | 85974713 | 85974807 | - | 20.0792115374673 | 23.876910583352 | 118.129592826431 | 119.516168461439 |
| chr11 | 85976619 | 85976684 | - | 35.1386201905678 | 42.0688424563821 | 84.7192029361272 | 66.1607361125824 |
| chr11 | 85981128 | 85981231 | - | 43.1703048055547 | 34.1098722619315 | 125.288962088639 | 104.576647403759 |
| chr11 | 85981744 | 85981770 | - | 9.03564519186028 | 11.3699574206438 | 23.8645642073598 | 28.8119334683827 |
| chr11 | 85981874 | 85982006 | - | 54.2138711511617 | 25.0139063254164 | 68.0140079909754 | 80.0331485232852 |
| chr11 | 85983865 | 85983885 | - | 4.01584230749346 | 1.13699574206438 | 9.54582568294392 | 8.53686917581709 |
| chr11 | 85983902 | 85983937 | - | 11.043566345607 | 6.82197445238629 | 17.8984231555198 | 14.9395210576799 |
| chr11 | 85990249 | 85990250 | - | 5.01980288436682 | 2.27399148412876 | 3.57968463110397 | 5.33554323488568 |
| chr11 | 85990322 | 85990331 | - | 8.03168461498692 | 5.68497871032191 | 10.7390538933119 | 12.8053037637256 |
| chr11 | 85996816 | 85996845 | - | 45.1782259593014 | 46.6168254246397 | 124.095733878271 | 133.388580872142 |
| chr11 | 86000674 | 86000726 | - | 24.0950538449607 | 35.2468680039958 | 41.7629873628796 | 36.2816939972226 |
| chr11 | 86001053 | 86001161 | - | 29.1148567293276 | 23.876910583352 | 71.5936926220794 | 48.0198891139711 |
| chr11 | 86007541 | 86007586 | - | 5.01980288436682 | 11.3699574206438 | 15.5119667347839 | 20.2750642925656 |
| chr11 | 86011029 | 86011036 | - | 3.01188173062009 | 6.82197445238629 | 9.54582568294392 | 1.06710864697714 |
| chr11 | 86011063 | 86011115 | - | 3.01188173062009 | 4.54798296825753 | 11.9322821036799 | 13.8724124107028 |
| chr11 | 86012299 | 86012331 | - | 6.02376346124019 | 4.54798296825753 | 3.57968463110397 | 4.26843458790854 |
| chr11 | 86012356 | 86012386 | - | 10.0396057687336 | 7.95897019445067 | 9.54582568294392 | 6.40265188186281 |
| chr11 | 86014948 | 86014965 | - | 11.043566345607 | 6.82197445238629 | 13.1255103140479 | 21.3421729395427 |
| chr11 | 86022366 | 86022378 | - | 11.043566345607 | 11.3699574206438 | 15.5119667347839 | 5.33554323488568 |
| chr11 | 86278486 | 86278521 | + | 10.0396057687336 | 9.09596593651506 | 4.77291284147196 | 14.9395210576799 |
| chr11 | 86278599 | 86278603 | + | 1.00396057687336 | 2.27399148412876 | 4.77291284147196 | 4.26843458790854 |
| chr11 | 86278645 | 86278658 | + | 1.00396057687336 | 5.68497871032191 | 5.96614105183995 | 7.46976052883995 |
| chr11 | 86278697 | 86278707 | + | 6.02376346124019 | 7.95897019445067 | 10.7390538933119 | 9.60397782279422 |
| chr11 | 86345582 | 86345603 | + | 2.00792115374673 | 7.95897019445067 | 8.35259747257593 | 2.13421729395427 |
| chr11 | 86345714 | 86345716 | + | 3.01188173062009 | 9.09596593651506 | 1.19322821036799 | 5.33554323488568 |
| chr11 | 86345863 | 86345883 | + | 2.00792115374673 | 3.41098722619315 | 1.19322821036799 | 1.06710864697714 |
| chr11 | 86945945 | 86946105 | - | 10.0396057687336 | 10.2329616785794 | 4.77291284147196 | 2.13421729395427 |
| chr11 | 86946129 | 86946390 | - | 6.02376346124019 | 12.5069531627082 | 1.19322821036799 | 3.20132594093141 |
| chr11 | 86949440 | 86949477 | - | 3.01188173062009 | 3.41098722619315 | 3.57968463110397 | 1.06710864697714 |
| chr11 | 86950766 | 86950855 | - | 4.01584230749346 | 22.7399148412876 | 1.19322821036799 | 2.13421729395427 |
| chr11 | 87321402 | 87321441 | + | 3.01188173062009 | 2.27399148412876 | 15.5119667347839 | 6.40265188186281 |
| chr11 | 87321828 | 87321838 | + | 4.01584230749346 | 3.41098722619315 | 3.57968463110397 | 5.33554323488568 |
| chr11 | 87321852 | 87321911 | + | 2.00792115374673 | 3.41098722619315 | 5.96614105183995 | 5.33554323488568 |
| chr11 | 87323338 | 87323353 | + | 1.00396057687336 | 1.13699574206438 | 2.38645642073598 | 2.13421729395427 |
| chr11 | 88293629 | 88293670 | - | 8.03168461498692 | 7.95897019445067 | 7.15936926220794 | 5.33554323488568 |
| chr11 | 88293758 | 88293803 | - | 10.0396057687336 | 14.780944646837 | 14.3187385244159 | 24.5434988804741 |
| chr11 | 88293934 | 88293981 | - | 55.217831728035 | 62.534765813541 | 16.7051949451519 | 46.952780466994 |
| chr11 | 88294003 | 88294110 | - | 16.0633692299738 | 12.5069531627082 | 8.35259747257593 | 23.476390233497 |
| chr11 | 88326129 | 88326165 | - | 4.01584230749346 | 3.41098722619315 | 7.15936926220794 | 8.53686917581709 |
| chr11 | 88326242 | 88326264 | - | 1.00396057687336 | 1.13699574206438 | 3.57968463110397 | 4.26843458790854 |
| chr11 | 90201928 | 90201934 | - | 18.0712903837206 | 11.3699574206438 | 2.38645642073598 | 6.40265188186281 |
| chr11 | 90202144 | 90202153 | - | 6.02376346124019 | 7.95897019445067 | 4.77291284147196 | 4.26843458790854 |
| chr11 | 90202378 | 90202415 | - | 70.2772403811355 | 64.8087572976698 | 56.0817258872955 | 56.5567582897882 |
| chr11 | 90213368 | 90213372 | - | 1.00396057687336 | 2.27399148412876 | 2.38645642073598 | 25.6106075274513 |
| chr11 | 93478545 | 93478570 | - | 14.0554480762271 | 21.6029190992233 | 5.96614105183995 | 2.13421729395427 |
| chr11 | 93478638 | 93478669 | - | 5.01980288436682 | 7.95897019445067 | 4.77291284147196 | 5.33554323488568 |
| chr11 | 93721512 | 93721515 | + | 8.03168461498692 | 12.5069531627082 | 36.9900745214077 | 1.06710864697714 |
| chr11 | 93721796 | 93721797 | + | 2.00792115374673 | 6.82197445238629 | 4.77291284147196 | 4.26843458790854 |
| chr11 | 93735960 | 93736005 | - | 3.01188173062009 | 2.27399148412876 | 1.19322821036799 | 1.06710864697714 |
| chr11 | 93736047 | 93736091 | - | 19.0752509605939 | 27.2878978095452 | 10.7390538933119 | 11.7381951167485 |
| chr11 | 93736130 | 93736176 | - | 90.3564519186028 | 43.2058381984465 | 26.2510206280958 | 24.5434988804741 |
| chr11 | 93764399 | 93764411 | + | 23.0910932680874 | 22.7399148412876 | 8.35259747257593 | 7.46976052883995 |
| chr11 | 94181667 | 94181668 | + | 1.00396057687336 | 1.13699574206438 | 2.38645642073598 | 1.06710864697714 |
| chr11 | 94181897 | 94181930 | + | 9.03564519186028 | 11.3699574206438 | 4.77291284147196 | 7.46976052883995 |
| chr11 | 94419912 | 94419913 | - | 2.00792115374673 | 2.27399148412876 | 2.38645642073598 | 2.13421729395427 |
| chr11 | 94498464 | 94498466 | + | 4.01584230749346 | 1.13699574206438 | 3.57968463110397 | 4.26843458790854 |
| chr11 | 94499450 | 94499488 | + | 8.03168461498692 | 6.82197445238629 | 7.15936926220794 | 4.26843458790854 |
| chr11 | 94963144 | 94963164 | - | 3.01188173062009 | 9.09596593651506 | 5.96614105183995 | 2.13421729395427 |
| chr11 | 94963219 | 94963222 | - | 1.00396057687336 | 3.41098722619315 | 1.19322821036799 | 1.06710864697714 |
| chr11 | 95068942 | 95068968 | + | 1.00396057687336 | 5.68497871032191 | 5.96614105183995 | 4.26843458790854 |
| chr11 | 95069454 | 95069477 | + | 2.00792115374673 | 4.54798296825753 | 3.57968463110397 | 9.60397782279422 |
| chr11 | 95129689 | 95129693 | + | 1.00396057687336 | 1.13699574206438 | 7.15936926220794 | 5.33554323488568 |
| chr11 | 95771222 | 95771243 | - | 3.01188173062009 | 6.82197445238629 | 8.35259747257593 | 16.006629704657 |
| chr11 | 95822548 | 95822579 | + | 12.0475269224804 | 14.780944646837 | 13.1255103140479 | 10.6710864697714 |
| chr11 | 95831553 | 95831557 | + | 4.01584230749346 | 1.13699574206438 | 1.19322821036799 | 4.26843458790854 |
| chr11 | 95831670 | 95831701 | + | 8.03168461498692 | 6.82197445238629 | 7.15936926220794 | 11.7381951167485 |
| chr11 | 95832454 | 95832467 | + | 2.00792115374673 | 3.41098722619315 | 3.57968463110397 | 1.06710864697714 |
| chr11 | 95833334 | 95833341 | - | 9.03564519186028 | 5.68497871032191 | 9.54582568294392 | 5.33554323488568 |
| chr11 | 95833434 | 95833446 | - | 55.217831728035 | 92.0966551072149 | 36.9900745214077 | 41.6172372321083 |
| chr11 | 95834143 | 95834159 | - | 4.01584230749346 | 5.68497871032191 | 2.38645642073598 | 5.33554323488568 |
| chr11 | 95834376 | 95834395 | - | 28.1108961524542 | 23.876910583352 | 13.1255103140479 | 14.9395210576799 |
| chr11 | 95834595 | 95834650 | - | 98.3881365335897 | 90.9596593651506 | 16.7051949451519 | 53.3554323488568 |
| chr11 | 95834720 | 95834749 | - | 6.02376346124019 | 4.54798296825753 | 1.19322821036799 | 4.26843458790854 |
| chr11 | 95977062 | 95977107 | - | 1.00396057687336 | 2.27399148412876 | 2.38645642073598 | 4.26843458790854 |
| chr11 | 96352998 | 96353011 | - | 5.01980288436682 | 2.27399148412876 | 8.35259747257593 | 4.26843458790854 |
| chr11 | 96353032 | 96353066 | - | 17.0673298068472 | 22.7399148412876 | 20.2848795762558 | 19.2079556455884 |
| chr11 | 96353111 | 96353126 | - | 13.0514874993537 | 35.2468680039958 | 41.7629873628796 | 22.4092815865198 |
| chr11 | 96353188 | 96353197 | - | 9.03564519186028 | 10.2329616785794 | 5.96614105183995 | 2.13421729395427 |
| chr11 | 96353268 | 96353301 | - | 13.0514874993537 | 13.6439489047726 | 9.54582568294392 | 7.46976052883995 |
| chr11 | 96353349 | 96353384 | - | 2.00792115374673 | 6.82197445238629 | 4.77291284147196 | 5.33554323488568 |
| chr11 | 96353488 | 96353499 | - | 3.01188173062009 | 6.82197445238629 | 2.38645642073598 | 6.40265188186281 |
| chr11 | 96353603 | 96353606 | - | 3.01188173062009 | 1.13699574206438 | 1.19322821036799 | 2.13421729395427 |
| chr11 | 101129152 | 101129153 | + | 11.043566345607 | 13.6439489047726 | 7.15936926220794 | 9.60397782279422 |
| chr11 | 102378427 | 102378433 | + | 4.01584230749346 | 12.5069531627082 | 2.38645642073598 | 8.53686917581709 |
| chr11 | 102378544 | 102378581 | + | 2.00792115374673 | 4.54798296825753 | 3.57968463110397 | 1.06710864697714 |
| chr11 | 102378630 | 102378642 | + | 1.00396057687336 | 2.27399148412876 | 1.19322821036799 | 2.13421729395427 |
| chr11 | 102396348 | 102396431 | - | 2.00792115374673 | 7.95897019445067 | 4.77291284147196 | 1.06710864697714 |
| chr11 | 102396461 | 102396921 | - | 29.1148567293276 | 21.6029190992233 | 23.8645642073598 | 13.8724124107028 |
| chr11 | 102397033 | 102397408 | - | 15.0594086531005 | 22.7399148412876 | 20.2848795762558 | 27.7448248214055 |
| chr11 | 102397429 | 102397593 | - | 4.01584230749346 | 1.13699574206438 | 3.57968463110397 | 6.40265188186281 |
| chr11 | 102397950 | 102398092 | - | 4.01584230749346 | 4.54798296825753 | 5.96614105183995 | 5.33554323488568 |
| chr11 | 102398113 | 102398266 | - | 5.01980288436682 | 4.54798296825753 | 11.9322821036799 | 19.2079556455884 |
| chr11 | 102398323 | 102398893 | - | 11.043566345607 | 18.1919318730301 | 34.6036181006717 | 43.7514545260626 |
| chr11 | 102452918 | 102452942 | + | 1.00396057687336 | 1.13699574206438 | 2.38645642073598 | 3.20132594093141 |
| chr11 | 103062135 | 103062214 | - | 11.043566345607 | 10.2329616785794 | 29.8307052591997 | 42.6843458790854 |
| chr11 | 103062251 | 103062295 | - | 18.0712903837206 | 23.876910583352 | 27.4442488384638 | 72.5633879944452 |
| chr11 | 106008009 | 106008010 | - | 1.00396057687336 | 1.13699574206438 | 3.57968463110397 | 1.06710864697714 |
| chr11 | 106096835 | 106096942 | + | 2.00792115374673 | 9.09596593651506 | 13.1255103140479 | 14.9395210576799 |
| chr11 | 106097123 | 106097155 | + | 2.00792115374673 | 2.27399148412876 | 11.9322821036799 | 5.33554323488568 |
| chr11 | 106097373 | 106097452 | + | 3.01188173062009 | 1.13699574206438 | 22.6713359969918 | 14.9395210576799 |
| chr11 | 106097502 | 106097523 | + | 2.00792115374673 | 1.13699574206438 | 10.7390538933119 | 13.8724124107028 |
| chr11 | 106098351 | 106098406 | + | 4.01584230749346 | 5.68497871032191 | 5.96614105183995 | 10.6710864697714 |
| chr11 | 106098563 | 106098642 | + | 7.02772403811355 | 12.5069531627082 | 22.6713359969918 | 14.9395210576799 |
| chr11 | 107791036 | 107791039 | - | 10.0396057687336 | 12.5069531627082 | 7.15936926220794 | 20.2750642925656 |
| chr11 | 107792193 | 107792197 | - | 3.01188173062009 | 1.13699574206438 | 15.5119667347839 | 61.8923015246739 |
| chr11 | 107792650 | 107792664 | - | 6.02376346124019 | 5.68497871032191 | 9.54582568294392 | 9.60397782279422 |
| chr11 | 108104458 | 108104461 | + | 1.00396057687336 | 1.13699574206438 | 2.38645642073598 | 1.06710864697714 |
| chr11 | 108104690 | 108104705 | + | 39.1544624980612 | 42.0688424563821 | 63.2410951495034 | 27.7448248214055 |
| chr11 | 108104842 | 108104911 | + | 4.01584230749346 | 9.09596593651506 | 19.0916513658878 | 17.0737383516342 |
| chr11 | 108105012 | 108105060 | + | 7.02772403811355 | 12.5069531627082 | 16.7051949451519 | 20.2750642925656 |
| chr11 | 108105395 | 108105397 | + | 2.00792115374673 | 1.13699574206438 | 1.19322821036799 | 1.06710864697714 |
| chr11 | 108105430 | 108105434 | + | 3.01188173062009 | 4.54798296825753 | 13.1255103140479 | 11.7381951167485 |
| chr11 | 108105567 | 108105587 | + | 1.00396057687336 | 4.54798296825753 | 1.19322821036799 | 7.46976052883995 |
| chr11 | 108105628 | 108105711 | + | 9.03564519186028 | 20.4659233571589 | 42.9562155732476 | 21.3421729395427 |
| chr11 | 108105789 | 108105838 | + | 3.01188173062009 | 2.27399148412876 | 20.2848795762558 | 8.53686917581709 |
| chr11 | 108106254 | 108106296 | + | 13.0514874993537 | 9.09596593651506 | 2.38645642073598 | 3.20132594093141 |
| chr11 | 108106350 | 108106438 | + | 40.1584230749346 | 37.5208594881246 | 70.4004644117114 | 71.4962793474681 |
| chr11 | 108106481 | 108106527 | + | 16.0633692299738 | 11.3699574206438 | 48.9223566250876 | 38.4159112911769 |
| chr11 | 108107141 | 108107147 | + | 2.00792115374673 | 1.13699574206438 | 3.57968463110397 | 7.46976052883995 |
| chr11 | 108107360 | 108107364 | + | 2.00792115374673 | 3.41098722619315 | 2.38645642073598 | 7.46976052883995 |
| chr11 | 108107476 | 108107500 | + | 2.00792115374673 | 1.13699574206438 | 1.19322821036799 | 1.06710864697714 |
| chr11 | 108107524 | 108107538 | + | 4.01584230749346 | 2.27399148412876 | 2.38645642073598 | 3.20132594093141 |
| chr11 | 108107601 | 108107632 | + | 4.01584230749346 | 1.13699574206438 | 1.19322821036799 | 3.20132594093141 |
| chr11 | 108147315 | 108147417 | + | 247.978262487721 | 214.892195250168 | 206.428480393662 | 314.797050858255 |
| chr11 | 108147448 | 108147485 | + | 1.00396057687336 | 3.41098722619315 | 4.77291284147196 | 7.46976052883995 |
| chr11 | 108366310 | 108366317 | + | 3.01188173062009 | 6.82197445238629 | 11.9322821036799 | 7.46976052883995 |
| chr11 | 108367557 | 108367630 | + | 9.03564519186028 | 17.0549361309657 | 9.54582568294392 | 3.20132594093141 |
| chr11 | 108368076 | 108368092 | + | 2.00792115374673 | 1.13699574206438 | 9.54582568294392 | 1.06710864697714 |
| chr11 | 108368160 | 108368197 | + | 6.02376346124019 | 2.27399148412876 | 9.54582568294392 | 7.46976052883995 |
| chr11 | 108368360 | 108368407 | + | 7.02772403811355 | 4.54798296825753 | 10.7390538933119 | 7.46976052883995 |
| chr11 | 108472367 | 108472410 | - | 29.1148567293276 | 43.2058381984465 | 31.0239334695677 | 28.8119334683827 |
| chr11 | 108940622 | 108940627 | + | 3.01188173062009 | 2.27399148412876 | 3.57968463110397 | 2.13421729395427 |
| chr11 | 110165915 | 110165919 | + | 1.00396057687336 | 3.41098722619315 | 1.19322821036799 | 1.06710864697714 |
| chr11 | 110168984 | 110168987 | + | 4.01584230749346 | 2.27399148412876 | 1.19322821036799 | 1.06710864697714 |
| chr11 | 110229923 | 110229935 | - | 17.0673298068472 | 15.9179403889013 | 4.77291284147196 | 7.46976052883995 |
| chr11 | 110230561 | 110230598 | - | 5.01980288436682 | 10.2329616785794 | 8.35259747257593 | 6.40265188186281 |
| chr11 | 110230625 | 110230657 | - | 28.1108961524542 | 31.8358807778027 | 8.35259747257593 | 17.0737383516342 |
| chr11 | 110231635 | 110231657 | - | 16.0633692299738 | 3.41098722619315 | 7.15936926220794 | 10.6710864697714 |
| chr11 | 110231691 | 110231725 | - | 4.01584230749346 | 2.27399148412876 | 3.57968463110397 | 8.53686917581709 |
| chr11 | 110462550 | 110462559 | + | 2.00792115374673 | 1.13699574206438 | 4.77291284147196 | 4.26843458790854 |
| chr11 | 111724881 | 111724896 | + | 3.01188173062009 | 2.27399148412876 | 3.57968463110397 | 5.33554323488568 |
| chr11 | 111739328 | 111739330 | - | 1.00396057687336 | 1.13699574206438 | 1.19322821036799 | 8.53686917581709 |
| chr11 | 111741025 | 111741055 | - | 1.00396057687336 | 2.27399148412876 | 3.57968463110397 | 17.0737383516342 |
| chr11 | 111883988 | 111884009 | + | 2.00792115374673 | 4.54798296825753 | 23.8645642073598 | 23.476390233497 |
| chr11 | 111918860 | 111918966 | + | 4.01584230749346 | 5.68497871032191 | 127.675418509375 | 70.429170700491 |
| chr11 | 112062609 | 112062615 | + | 1.00396057687336 | 1.13699574206438 | 7.15936926220794 | 18.1408469986113 |
| chr11 | 112062971 | 112062980 | + | 4.01584230749346 | 5.68497871032191 | 3.57968463110397 | 5.33554323488568 |
| chr11 | 112063518 | 112063521 | + | 3.01188173062009 | 4.54798296825753 | 3.57968463110397 | 3.20132594093141 |
| chr11 | 112063800 | 112063821 | + | 1.00396057687336 | 2.27399148412876 | 5.96614105183995 | 1.06710864697714 |
| chr11 | 112083103 | 112083128 | + | 6.02376346124019 | 4.54798296825753 | 7.15936926220794 | 4.26843458790854 |
| chr11 | 112083158 | 112083182 | + | 4.01584230749346 | 5.68497871032191 | 17.8984231555198 | 14.9395210576799 |
| chr11 | 112083320 | 112083347 | + | 1.00396057687336 | 3.41098722619315 | 15.5119667347839 | 14.9395210576799 |
| chr11 | 112083470 | 112083505 | + | 7.02772403811355 | 7.95897019445067 | 16.7051949451519 | 14.9395210576799 |
| chr11 | 112084782 | 112084784 | + | 1.00396057687336 | 4.54798296825753 | 1.19322821036799 | 3.20132594093141 |
| chr11 | 112084930 | 112084932 | - | 3.01188173062009 | 5.68497871032191 | 5.96614105183995 | 2.13421729395427 |
| chr11 | 112085173 | 112085197 | - | 141.558441339144 | 168.275369825529 | 163.472264820415 | 179.274252692159 |
| chr11 | 112085223 | 112085240 | - | 248.982223064594 | 288.796918484353 | 223.133675338814 | 465.259370082031 |
| chr11 | 112085280 | 112085283 | - | 4.01584230749346 | 1.13699574206438 | 5.96614105183995 | 6.40265188186281 |
| chr11 | 112095526 | 112095540 | + | 1.00396057687336 | 1.13699574206438 | 7.15936926220794 | 6.40265188186281 |
| chr11 | 112095567 | 112095591 | + | 7.02772403811355 | 10.2329616785794 | 9.54582568294392 | 16.006629704657 |
| chr11 | 112233228 | 112233238 | + | 2.00792115374673 | 1.13699574206438 | 3.57968463110397 | 1.06710864697714 |
| chr11 | 112233561 | 112233573 | + | 53.2099105742883 | 75.0417189762492 | 9.54582568294392 | 21.3421729395427 |
| chr11 | 112233633 | 112233677 | + | 11.043566345607 | 20.4659233571589 | 16.7051949451519 | 11.7381951167485 |
| chr11 | 112233752 | 112233770 | + | 1.00396057687336 | 4.54798296825753 | 1.19322821036799 | 4.26843458790854 |
| chr11 | 112290273 | 112290280 | + | 2.00792115374673 | 2.27399148412876 | 1.19322821036799 | 1.06710864697714 |
| chr11 | 112795638 | 112795639 | + | 2.00792115374673 | 2.27399148412876 | 5.96614105183995 | 4.26843458790854 |
| chr11 | 113733230 | 113733233 | - | 1.00396057687336 | 2.27399148412876 | 4.77291284147196 | 1.06710864697714 |
| chr11 | 113733262 | 113733291 | - | 2.00792115374673 | 3.41098722619315 | 2.38645642073598 | 6.40265188186281 |
| chr11 | 113798100 | 113798107 | - | 3.01188173062009 | 7.95897019445067 | 3.57968463110397 | 3.20132594093141 |
| chr11 | 113798135 | 113798136 | - | 1.00396057687336 | 2.27399148412876 | 2.38645642073598 | 1.06710864697714 |
| chr11 | 113798992 | 113799030 | - | 5.01980288436682 | 3.41098722619315 | 7.15936926220794 | 3.20132594093141 |
| chr11 | 113799127 | 113799158 | - | 4.01584230749346 | 2.27399148412876 | 1.19322821036799 | 1.06710864697714 |
| chr11 | 114312452 | 114312497 | + | 60.2376346124019 | 64.8087572976698 | 2.38645642073598 | 8.53686917581709 |
| chr11 | 114407477 | 114407479 | + | 3.01188173062009 | 2.27399148412876 | 2.38645642073598 | 1.06710864697714 |
| chr11 | 114408679 | 114408687 | + | 2.00792115374673 | 2.27399148412876 | 1.19322821036799 | 2.13421729395427 |
| chr11 | 114408909 | 114408918 | + | 9.03564519186028 | 7.95897019445067 | 11.9322821036799 | 12.8053037637256 |
| chr11 | 114443885 | 114443934 | + | 8.03168461498692 | 6.82197445238629 | 15.5119667347839 | 18.1408469986113 |
| chr11 | 114449916 | 114449960 | + | 6.02376346124019 | 6.82197445238629 | 13.1255103140479 | 10.6710864697714 |
| chr11 | 114450011 | 114450014 | + | 7.02772403811355 | 1.13699574206438 | 8.35259747257593 | 2.13421729395427 |
| chr11 | 114450047 | 114450068 | + | 9.03564519186028 | 9.09596593651506 | 9.54582568294392 | 14.9395210576799 |
| chr11 | 114450100 | 114450112 | + | 3.01188173062009 | 4.54798296825753 | 3.57968463110397 | 9.60397782279422 |
| chr11 | 114450155 | 114450208 | + | 35.1386201905678 | 79.5897019445067 | 40.5697591525116 | 99.2411041688736 |
| chr11 | 114450248 | 114450258 | + | 5.01980288436682 | 5.68497871032191 | 4.77291284147196 | 6.40265188186281 |
| chr11 | 116748223 | 116748238 | - | 3.01188173062009 | 4.54798296825753 | 4.77291284147196 | 7.46976052883995 |
| chr11 | 116777893 | 116777904 | - | 4.01584230749346 | 3.41098722619315 | 13.1255103140479 | 9.60397782279422 |
| chr11 | 116777960 | 116777982 | - | 9.03564519186028 | 9.09596593651506 | 15.5119667347839 | 20.2750642925656 |
| chr11 | 116778022 | 116778042 | - | 7.02772403811355 | 1.13699574206438 | 11.9322821036799 | 10.6710864697714 |
| chr11 | 116778110 | 116778245 | - | 8.03168461498692 | 9.09596593651506 | 34.6036181006717 | 39.483019938154 |
| chr11 | 116778496 | 116778500 | - | 1.00396057687336 | 2.27399148412876 | 8.35259747257593 | 10.6710864697714 |
| chr11 | 116778577 | 116778606 | - | 7.02772403811355 | 3.41098722619315 | 15.5119667347839 | 14.9395210576799 |
| chr11 | 116778668 | 116778694 | - | 4.01584230749346 | 3.41098722619315 | 4.77291284147196 | 11.7381951167485 |
| chr11 | 116843654 | 116843655 | - | 1.00396057687336 | 2.27399148412876 | 1.19322821036799 | 1.06710864697714 |
| chr11 | 116845353 | 116845355 | - | 3.01188173062009 | 18.1919318730301 | 7.15936926220794 | 7.46976052883995 |
| chr11 | 117167833 | 117167869 | + | 12.0475269224804 | 12.5069531627082 | 15.5119667347839 | 23.476390233497 |
| chr11 | 117167932 | 117167940 | + | 13.0514874993537 | 11.3699574206438 | 23.8645642073598 | 26.6777161744284 |
| chr11 | 117168230 | 117168253 | + | 3.01188173062009 | 4.54798296825753 | 26.2510206280958 | 11.7381951167485 |
| chr11 | 117168286 | 117168315 | + | 1.00396057687336 | 1.13699574206438 | 8.35259747257593 | 24.5434988804741 |
| chr11 | 117168683 | 117168716 | + | 10.0396057687336 | 4.54798296825753 | 3.57968463110397 | 7.46976052883995 |
| chr11 | 117168744 | 117168780 | + | 1.00396057687336 | 1.13699574206438 | 4.77291284147196 | 7.46976052883995 |
| chr11 | 117169080 | 117169101 | + | 2.00792115374673 | 1.13699574206438 | 10.7390538933119 | 8.53686917581709 |
| chr11 | 117169494 | 117169568 | + | 24.0950538449607 | 12.5069531627082 | 10.7390538933119 | 16.006629704657 |
| chr11 | 117169714 | 117169724 | + | 5.01980288436682 | 5.68497871032191 | 4.77291284147196 | 9.60397782279422 |
| chr11 | 117169756 | 117169773 | + | 1.00396057687336 | 1.13699574206438 | 3.57968463110397 | 5.33554323488568 |
| chr11 | 117169841 | 117169851 | + | 1.00396057687336 | 1.13699574206438 | 5.96614105183995 | 9.60397782279422 |
| chr11 | 117169917 | 117169951 | + | 4.01584230749346 | 6.82197445238629 | 4.77291284147196 | 5.33554323488568 |
| chr11 | 117170002 | 117170036 | + | 1.00396057687336 | 1.13699574206438 | 2.38645642073598 | 8.53686917581709 |
| chr11 | 117170151 | 117170185 | + | 6.02376346124019 | 4.54798296825753 | 3.57968463110397 | 5.33554323488568 |
| chr11 | 117170373 | 117170550 | + | 61.2415951892752 | 40.9318467143178 | 144.380613454527 | 100.308212815851 |
| chr11 | 117170587 | 117170606 | + | 10.0396057687336 | 12.5069531627082 | 25.0577924177278 | 11.7381951167485 |
| chr11 | 117170645 | 117170652 | + | 5.01980288436682 | 3.41098722619315 | 13.1255103140479 | 3.20132594093141 |
| chr11 | 117170723 | 117170987 | + | 157.621810569118 | 153.494425178692 | 189.72328544851 | 314.797050858255 |
| chr11 | 117197335 | 117197392 | + | 1.00396057687336 | 1.13699574206438 | 2.38645642073598 | 8.53686917581709 |
| chr11 | 117205225 | 117205269 | - | 5.01980288436682 | 3.41098722619315 | 1.19322821036799 | 5.33554323488568 |
| chr11 | 117285377 | 117285398 | + | 1.00396057687336 | 9.09596593651506 | 5.96614105183995 | 5.33554323488568 |
| chr11 | 117413182 | 117413212 | + | 4.01584230749346 | 12.5069531627082 | 3.57968463110397 | 3.20132594093141 |
| chr11 | 118226766 | 118226783 | - | 1.00396057687336 | 2.27399148412876 | 2.38645642073598 | 1.06710864697714 |
| chr11 | 118396467 | 118396522 | + | 15.0594086531005 | 14.780944646837 | 14.3187385244159 | 13.8724124107028 |
| chr11 | 118398167 | 118398235 | + | 4.01584230749346 | 1.13699574206438 | 16.7051949451519 | 21.3421729395427 |
| chr11 | 118398969 | 118399000 | + | 3.01188173062009 | 2.27399148412876 | 5.96614105183995 | 6.40265188186281 |
| chr11 | 118399132 | 118399138 | + | 3.01188173062009 | 2.27399148412876 | 3.57968463110397 | 3.20132594093141 |
| chr11 | 118409013 | 118409104 | + | 48.1901076899215 | 39.7948509722534 | 70.4004644117114 | 78.966039876308 |
| chr11 | 118409136 | 118409142 | + | 3.01188173062009 | 2.27399148412876 | 2.38645642073598 | 1.06710864697714 |
| chr11 | 118409417 | 118409430 | + | 7.02772403811355 | 4.54798296825753 | 3.57968463110397 | 5.33554323488568 |
| chr11 | 118409630 | 118409634 | + | 2.00792115374673 | 3.41098722619315 | 4.77291284147196 | 3.20132594093141 |
| chr11 | 118523343 | 118523346 | + | 3.01188173062009 | 1.13699574206438 | 2.38645642073598 | 2.13421729395427 |
| chr11 | 118523366 | 118523384 | + | 1.00396057687336 | 2.27399148412876 | 4.77291284147196 | 1.06710864697714 |
| chr11 | 118523611 | 118523668 | + | 6.02376346124019 | 9.09596593651506 | 7.15936926220794 | 4.26843458790854 |
| chr11 | 118524007 | 118524008 | + | 1.00396057687336 | 3.41098722619315 | 14.3187385244159 | 2.13421729395427 |
| chr11 | 118525472 | 118525476 | + | 7.02772403811355 | 3.41098722619315 | 7.15936926220794 | 5.33554323488568 |
| chr11 | 118526297 | 118526319 | + | 4.01584230749346 | 5.68497871032191 | 2.38645642073598 | 5.33554323488568 |
| chr11 | 118544563 | 118544566 | - | 1.00396057687336 | 2.27399148412876 | 2.38645642073598 | 1.06710864697714 |
| chr11 | 118544631 | 118544646 | - | 5.01980288436682 | 12.5069531627082 | 3.57968463110397 | 4.26843458790854 |
| chr11 | 118600712 | 118600933 | + | 13.0514874993537 | 12.5069531627082 | 90.6853439879672 | 54.4225409958339 |
| chr11 | 118601002 | 118601030 | + | 5.01980288436682 | 1.13699574206438 | 16.7051949451519 | 8.53686917581709 |
| chr11 | 118601343 | 118601363 | + | 3.01188173062009 | 5.68497871032191 | 10.7390538933119 | 3.20132594093141 |
| chr11 | 118601442 | 118601478 | + | 1.00396057687336 | 1.13699574206438 | 10.7390538933119 | 1.06710864697714 |
| chr11 | 118601503 | 118601559 | + | 10.0396057687336 | 6.82197445238629 | 20.2848795762558 | 19.2079556455884 |
| chr11 | 118601594 | 118601644 | + | 18.0712903837206 | 12.5069531627082 | 35.7968463110397 | 27.7448248214055 |
| chr11 | 118601671 | 118601728 | + | 1.00396057687336 | 2.27399148412876 | 16.7051949451519 | 6.40265188186281 |
| chr11 | 118601754 | 118601838 | + | 7.02772403811355 | 3.41098722619315 | 27.4442488384638 | 23.476390233497 |
| chr11 | 118602665 | 118602714 | + | 9.03564519186028 | 21.6029190992233 | 15.5119667347839 | 29.8790421153598 |
| chr11 | 118602809 | 118602829 | + | 3.01188173062009 | 2.27399148412876 | 7.15936926220794 | 3.20132594093141 |
| chr11 | 118602855 | 118602892 | + | 5.01980288436682 | 10.2329616785794 | 19.0916513658878 | 16.006629704657 |
| chr11 | 118747835 | 118747913 | - | 141.558441339144 | 122.795540142953 | 233.872729232126 | 186.744013220999 |
| chr11 | 118747961 | 118747972 | - | 6.02376346124019 | 5.68497871032191 | 33.4103898903037 | 76.8318225823538 |
| chr11 | 118748076 | 118748084 | - | 2.00792115374673 | 5.68497871032191 | 11.9322821036799 | 34.1474767032683 |
| chr11 | 118748101 | 118748200 | - | 61.2415951892752 | 36.3838637460602 | 144.380613454527 | 129.120146284233 |
| chr11 | 118748222 | 118748245 | - | 1.00396057687336 | 2.27399148412876 | 7.15936926220794 | 21.3421729395427 |
| chr11 | 118748275 | 118748291 | - | 13.0514874993537 | 10.2329616785794 | 33.4103898903037 | 42.6843458790854 |
| chr11 | 118748315 | 118748328 | - | 3.01188173062009 | 1.13699574206438 | 8.35259747257593 | 2.13421729395427 |
| chr11 | 118748448 | 118748495 | - | 1.00396057687336 | 3.41098722619315 | 11.9322821036799 | 10.6710864697714 |
| chr11 | 118748504 | 118748545 | - | 4.01584230749346 | 3.41098722619315 | 8.35259747257593 | 4.26843458790854 |
| chr11 | 118748621 | 118748623 | - | 2.00792115374673 | 2.27399148412876 | 3.57968463110397 | 2.13421729395427 |
| chr11 | 118748695 | 118748736 | - | 330.303029791337 | 235.358118607327 | 93.0718004087032 | 56.5567582897882 |
| chr11 | 118748990 | 118749022 | - | 2.00792115374673 | 2.27399148412876 | 4.77291284147196 | 3.20132594093141 |
| chr11 | 118750277 | 118750300 | - | 1.00396057687336 | 1.13699574206438 | 9.54582568294392 | 21.3421729395427 |
| chr11 | 118750477 | 118750478 | - | 1.00396057687336 | 1.13699574206438 | 2.38645642073598 | 4.26843458790854 |
| chr11 | 118751133 | 118751158 | - | 2.00792115374673 | 3.41098722619315 | 4.77291284147196 | 10.6710864697714 |
| chr11 | 118751450 | 118751497 | - | 2.00792115374673 | 4.54798296825753 | 23.8645642073598 | 114.180625226554 |
| chr11 | 118751583 | 118751628 | - | 9.03564519186028 | 10.2329616785794 | 54.8884976769275 | 110.979299285622 |
| chr11 | 118751829 | 118751842 | - | 8.03168461498692 | 12.5069531627082 | 15.5119667347839 | 11.7381951167485 |
| chr11 | 118751926 | 118752004 | - | 11.043566345607 | 10.2329616785794 | 14.3187385244159 | 20.2750642925656 |
| chr11 | 118752031 | 118752067 | - | 11.043566345607 | 7.95897019445067 | 63.2410951495034 | 57.6238669367653 |
| chr11 | 118754637 | 118754638 | - | 1.00396057687336 | 3.41098722619315 | 1.19322821036799 | 1.06710864697714 |
| chr11 | 118791246 | 118791254 | + | 5.01980288436682 | 4.54798296825753 | 2.38645642073598 | 1.06710864697714 |
| chr11 | 118896198 | 118896211 | - | 21.0831721143407 | 18.1919318730301 | 1.19322821036799 | 1.06710864697714 |
| chr11 | 118898273 | 118898316 | - | 81.3208067267425 | 39.7948509722534 | 2.38645642073598 | 7.46976052883995 |
| chr11 | 119093468 | 119093513 | + | 2.00792115374673 | 9.09596593651506 | 13.1255103140479 | 23.476390233497 |
| chr11 | 119093905 | 119093957 | - | 19.0752509605939 | 30.6988850357383 | 17.8984231555198 | 26.6777161744284 |
| chr11 | 119096530 | 119096557 | - | 21.0831721143407 | 32.9728765198671 | 27.4442488384638 | 42.6843458790854 |
| chr11 | 119096632 | 119096650 | - | 5.01980288436682 | 2.27399148412876 | 1.19322821036799 | 2.13421729395427 |
| chr11 | 119306702 | 119306703 | + | 2.00792115374673 | 2.27399148412876 | 1.19322821036799 | 2.13421729395427 |
| chr11 | 119306820 | 119306843 | + | 9.03564519186028 | 15.9179403889013 | 41.7629873628796 | 23.476390233497 |
| chr11 | 119307684 | 119307688 | + | 8.03168461498692 | 17.0549361309657 | 4.77291284147196 | 34.1474767032683 |
| chr11 | 119307738 | 119307750 | + | 1.00396057687336 | 1.13699574206438 | 5.96614105183995 | 12.8053037637256 |
| chr11 | 119308622 | 119308647 | - | 20.0792115374673 | 31.8358807778027 | 4.77291284147196 | 3.20132594093141 |
| chr11 | 119308678 | 119308734 | - | 34.1346596136944 | 35.2468680039958 | 10.7390538933119 | 34.1474767032683 |
| chr11 | 119309217 | 119309218 | - | 3.01188173062009 | 1.13699574206438 | 3.57968463110397 | 3.20132594093141 |
| chr11 | 119309795 | 119309819 | - | 1.00396057687336 | 5.68497871032191 | 3.57968463110397 | 3.20132594093141 |
| chr11 | 119337238 | 119337275 | + | 23.0910932680874 | 10.2329616785794 | 9.54582568294392 | 21.3421729395427 |
| chr11 | 119381766 | 119381842 | + | 188.744588452193 | 201.248246345396 | 198.075882921086 | 221.958598571244 |
| chr11 | 119641489 | 119641490 | + | 3.01188173062009 | 10.2329616785794 | 16.7051949451519 | 7.46976052883995 |
| chr11 | 120229705 | 120229922 | + | 20.0792115374673 | 13.6439489047726 | 9.54582568294392 | 6.40265188186281 |
| chr11 | 120330623 | 120330624 | + | 1.00396057687336 | 1.13699574206438 | 1.19322821036799 | 1.06710864697714 |
| chr11 | 120485363 | 120485368 | + | 2.00792115374673 | 3.41098722619315 | 4.77291284147196 | 1.06710864697714 |
| chr11 | 120485618 | 120485620 | + | 1.00396057687336 | 2.27399148412876 | 1.19322821036799 | 1.06710864697714 |
| chr11 | 120485900 | 120485911 | + | 47.1861471130481 | 50.0278126508328 | 72.7869208324474 | 107.777973344691 |
| chr11 | 120486066 | 120486118 | + | 266.049552871442 | 312.673829067705 | 113.356679984959 | 128.053037637256 |
| chr11 | 120486369 | 120486372 | + | 4.01584230749346 | 4.54798296825753 | 3.57968463110397 | 7.46976052883995 |
| chr11 | 120486479 | 120486485 | + | 3.01188173062009 | 1.13699574206438 | 2.38645642073598 | 5.33554323488568 |
| chr11 | 120486732 | 120486740 | + | 3.01188173062009 | 3.41098722619315 | 2.38645642073598 | 2.13421729395427 |
| chr11 | 120486960 | 120486976 | + | 1.00396057687336 | 1.13699574206438 | 2.38645642073598 | 3.20132594093141 |
| chr11 | 120487097 | 120487115 | + | 1.00396057687336 | 1.13699574206438 | 1.19322821036799 | 7.46976052883995 |
| chr11 | 120487157 | 120487160 | + | 1.00396057687336 | 1.13699574206438 | 5.96614105183995 | 1.06710864697714 |
| chr11 | 120487541 | 120487580 | + | 1.00396057687336 | 1.13699574206438 | 2.38645642073598 | 7.46976052883995 |
| chr11 | 120487592 | 120487613 | + | 1.00396057687336 | 1.13699574206438 | 7.15936926220794 | 5.33554323488568 |
| chr11 | 120487649 | 120487661 | + | 6.02376346124019 | 5.68497871032191 | 1.19322821036799 | 10.6710864697714 |
| chr11 | 120488978 | 120489004 | + | 2.00792115374673 | 13.6439489047726 | 1.19322821036799 | 8.53686917581709 |
| chr11 | 121307532 | 121307549 | + | 5.01980288436682 | 5.68497871032191 | 4.77291284147196 | 7.46976052883995 |
| chr11 | 121308096 | 121308128 | + | 14.0554480762271 | 10.2329616785794 | 2.38645642073598 | 5.33554323488568 |
| chr11 | 121308435 | 121308449 | + | 2.00792115374673 | 2.27399148412876 | 1.19322821036799 | 1.06710864697714 |
| chr11 | 121308576 | 121308585 | + | 3.01188173062009 | 4.54798296825753 | 1.19322821036799 | 5.33554323488568 |
| chr11 | 123057722 | 123057749 | - | 23.0910932680874 | 10.2329616785794 | 25.0577924177278 | 27.7448248214055 |
| chr11 | 124124440 | 124124446 | + | 5.01980288436682 | 11.3699574206438 | 1.19322821036799 | 1.06710864697714 |
| chr11 | 124632246 | 124632248 | + | 1.00396057687336 | 3.41098722619315 | 1.19322821036799 | 1.06710864697714 |
| chr11 | 124632389 | 124632401 | + | 4.01584230749346 | 4.54798296825753 | 1.19322821036799 | 4.26843458790854 |
| chr11 | 124633453 | 124633458 | + | 4.01584230749346 | 5.68497871032191 | 1.19322821036799 | 1.06710864697714 |
| chr11 | 124636464 | 124636482 | - | 5.01980288436682 | 4.54798296825753 | 2.38645642073598 | 3.20132594093141 |
| chr11 | 125583664 | 125583798 | + | 171.677258645345 | 98.9186295596012 | 93.0718004087032 | 364.95115726618 |
| chr11 | 125583832 | 125583879 | + | 33.130699036821 | 17.0549361309657 | 47.7291284147196 | 80.0331485232852 |
| chr11 | 125583950 | 125583993 | + | 76.3010038423757 | 57.9867828452835 | 212.394621445502 | 148.328101929822 |
| chr11 | 125584015 | 125584079 | + | 35.1386201905678 | 22.7399148412876 | 84.7192029361272 | 37.3488026441998 |
| chr11 | 125584113 | 125584134 | + | 30.1188173062009 | 21.6029190992233 | 33.4103898903037 | 21.3421729395427 |
| chr11 | 125584206 | 125584218 | + | 47.1861471130481 | 60.2607743294122 | 50.1155848354556 | 35.2145853502455 |
| chr11 | 125584473 | 125584505 | + | 3.01188173062009 | 1.13699574206438 | 2.38645642073598 | 1.06710864697714 |
| chr11 | 125584532 | 125584558 | + | 9.03564519186028 | 1.13699574206438 | 1.19322821036799 | 7.46976052883995 |
| chr11 | 125584615 | 125584626 | + | 58.2297134586551 | 39.7948509722534 | 64.4343233598714 | 176.072926751227 |
| chr11 | 125620837 | 125620860 | + | 4.01584230749346 | 9.09596593651506 | 3.57968463110397 | 26.6777161744284 |
| chr11 | 125655526 | 125655528 | + | 1.00396057687336 | 1.13699574206438 | 1.19322821036799 | 3.20132594093141 |
| chr11 | 125655616 | 125655618 | + | 2.00792115374673 | 7.95897019445067 | 5.96614105183995 | 5.33554323488568 |
| chr11 | 125656850 | 125656871 | + | 4.01584230749346 | 9.09596593651506 | 1.19322821036799 | 2.13421729395427 |
| chr11 | 126202433 | 126202561 | - | 25.0990144218341 | 23.876910583352 | 27.4442488384638 | 39.483019938154 |
| chr11 | 126203063 | 126203156 | - | 4.01584230749346 | 1.13699574206438 | 4.77291284147196 | 6.40265188186281 |
| chr11 | 126203277 | 126203317 | - | 2.00792115374673 | 1.13699574206438 | 5.96614105183995 | 6.40265188186281 |
| chr11 | 126203408 | 126203449 | - | 2.00792115374673 | 2.27399148412876 | 1.19322821036799 | 7.46976052883995 |
| chr11 | 126262977 | 126263023 | - | 12.0475269224804 | 26.1509020674808 | 5.96614105183995 | 24.5434988804741 |
| chr11 | 126278040 | 126278085 | + | 10.0396057687336 | 17.0549361309657 | 14.3187385244159 | 21.3421729395427 |
| chr11 | 126345623 | 126345687 | + | 119.47130864793 | 195.563267635074 | 34.6036181006717 | 61.8923015246739 |
| chr11 | 129899763 | 129899766 | - | 1.00396057687336 | 12.5069531627082 | 1.19322821036799 | 1.06710864697714 |
| chr11 | 129900056 | 129900078 | - | 1.00396057687336 | 10.2329616785794 | 7.15936926220794 | 4.26843458790854 |
| chr11 | 129900109 | 129900156 | - | 7.02772403811355 | 10.2329616785794 | 4.77291284147196 | 3.20132594093141 |
| chr11 | 129901863 | 129901903 | - | 5.01980288436682 | 3.41098722619315 | 4.77291284147196 | 4.26843458790854 |
| chr11 | 129901933 | 129901967 | - | 3.01188173062009 | 5.68497871032191 | 7.15936926220794 | 1.06710864697714 |
| chr11 | 129902065 | 129902088 | - | 6.02376346124019 | 4.54798296825753 | 3.57968463110397 | 5.33554323488568 |
| chr11 | 130143346 | 130144238 | + | 104.41189999483 | 86.411676396893 | 140.800928823423 | 188.878230514953 |
| chr11 | 130144366 | 130144576 | + | 37.1465413443145 | 28.4248935516096 | 31.0239334695677 | 62.959410171651 |
| chr11 | 130144607 | 130144652 | + | 23.0910932680874 | 21.6029190992233 | 7.15936926220794 | 23.476390233497 |
| chr11 | 130144674 | 130144716 | + | 3.01188173062009 | 2.27399148412876 | 4.77291284147196 | 1.06710864697714 |
| chr11 | 130144737 | 130144788 | + | 2.00792115374673 | 10.2329616785794 | 5.96614105183995 | 12.8053037637256 |
| chr11 | 130226760 | 130226784 | - | 6.02376346124019 | 2.27399148412876 | 2.38645642073598 | 1.06710864697714 |
| chr11 | 130228789 | 130228805 | - | 4.01584230749346 | 6.82197445238629 | 3.57968463110397 | 4.26843458790854 |
| chr11 | 130230685 | 130230701 | - | 11.043566345607 | 7.95897019445067 | 10.7390538933119 | 9.60397782279422 |
| chr11 | 130230792 | 130230853 | - | 20.0792115374673 | 19.3289276150945 | 4.77291284147196 | 8.53686917581709 |
| chr11 | 130230895 | 130230901 | - | 14.0554480762271 | 7.95897019445067 | 4.77291284147196 | 6.40265188186281 |
| chr11 | 130231413 | 130231434 | - | 1.00396057687336 | 1.13699574206438 | 2.38645642073598 | 1.06710864697714 |
| chr11 | 130231492 | 130231496 | - | 4.01584230749346 | 2.27399148412876 | 1.19322821036799 | 1.06710864697714 |
| chr11 | 130231640 | 130231669 | - | 5.01980288436682 | 4.54798296825753 | 3.57968463110397 | 2.13421729395427 |
| chr11 | 130231689 | 130231712 | - | 5.01980288436682 | 5.68497871032191 | 4.77291284147196 | 6.40265188186281 |
| chr11 | 130231830 | 130231868 | - | 1.00396057687336 | 3.41098722619315 | 1.19322821036799 | 9.60397782279422 |
| chr11 | 130232127 | 130232129 | - | 5.01980288436682 | 1.13699574206438 | 2.38645642073598 | 1.06710864697714 |
| chr11 | 130232735 | 130232737 | - | 1.00396057687336 | 1.13699574206438 | 4.77291284147196 | 1.06710864697714 |
| chr11 | 130233034 | 130233043 | - | 16.0633692299738 | 7.95897019445067 | 3.57968463110397 | 5.33554323488568 |
| chr11 | 130233084 | 130233090 | - | 4.01584230749346 | 2.27399148412876 | 3.57968463110397 | 3.20132594093141 |
| chr11 | 130876645 | 130876647 | - | 2.00792115374673 | 5.68497871032191 | 10.7390538933119 | 10.6710864697714 |
| chr11 | 134032763 | 134032764 | + | 1.00396057687336 | 3.41098722619315 | 1.19322821036799 | 1.06710864697714 |
| chr11 | 134152485 | 134152506 | - | 2.00792115374673 | 3.41098722619315 | 1.19322821036799 | 3.20132594093141 |
| chr11 | 134245658 | 134245692 | + | 14.0554480762271 | 7.95897019445067 | 8.35259747257593 | 19.2079556455884 |
| chr11 | 134248311 | 134248363 | - | 12.0475269224804 | 13.6439489047726 | 13.1255103140479 | 32.0132594093141 |
| chr11 | 134248448 | 134248459 | - | 13.0514874993537 | 11.3699574206438 | 4.77291284147196 | 17.0737383516342 |
| chr11 | 134265586 | 134265587 | + | 1.00396057687336 | 3.41098722619315 | 2.38645642073598 | 1.06710864697714 |
| chr11 | 134265608 | 134265618 | + | 4.01584230749346 | 10.2329616785794 | 2.38645642073598 | 5.33554323488568 |
| chr12 | 282543 | 282548 | - | 1.00396057687336 | 1.13699574206438 | 2.38645642073598 | 3.20132594093141 |
| chr12 | 283701 | 283703 | - | 2.00792115374673 | 3.41098722619315 | 1.19322821036799 | 1.06710864697714 |
| chr12 | 284163 | 284166 | - | 5.01980288436682 | 5.68497871032191 | 2.38645642073598 | 1.06710864697714 |
| chr12 | 284363 | 284391 | - | 29.1148567293276 | 31.8358807778027 | 7.15936926220794 | 8.53686917581709 |
| chr12 | 284571 | 284588 | - | 4.01584230749346 | 4.54798296825753 | 3.57968463110397 | 4.26843458790854 |
| chr12 | 284678 | 284696 | - | 5.01980288436682 | 5.68497871032191 | 5.96614105183995 | 2.13421729395427 |
| chr12 | 284736 | 284759 | - | 11.043566345607 | 10.2329616785794 | 15.5119667347839 | 3.20132594093141 |
| chr12 | 284809 | 284830 | - | 6.02376346124019 | 4.54798296825753 | 5.96614105183995 | 2.13421729395427 |
| chr12 | 442463 | 442482 | + | 2.00792115374673 | 4.54798296825753 | 2.38645642073598 | 4.26843458790854 |
| chr12 | 442522 | 442531 | + | 14.0554480762271 | 9.09596593651506 | 3.57968463110397 | 6.40265188186281 |
| chr12 | 645152 | 645154 | + | 1.00396057687336 | 3.41098722619315 | 2.38645642073598 | 3.20132594093141 |
| chr12 | 645180 | 645182 | + | 2.00792115374673 | 2.27399148412876 | 2.38645642073598 | 2.13421729395427 |
| chr12 | 908850 | 908864 | + | 15.0594086531005 | 17.0549361309657 | 1.19322821036799 | 4.26843458790854 |
| chr12 | 908918 | 908921 | + | 1.00396057687336 | 2.27399148412876 | 1.19322821036799 | 6.40265188186281 |
| chr12 | 909193 | 909267 | + | 26.1029749987075 | 27.2878978095452 | 1.19322821036799 | 12.8053037637256 |
| chr12 | 909318 | 909379 | + | 35.1386201905678 | 54.5757956190903 | 7.15936926220794 | 13.8724124107028 |
| chr12 | 909431 | 909497 | + | 9.03564519186028 | 7.95897019445067 | 4.77291284147196 | 4.26843458790854 |
| chr12 | 909674 | 909703 | + | 1.00396057687336 | 7.95897019445067 | 3.57968463110397 | 2.13421729395427 |
| chr12 | 909752 | 909822 | + | 6.02376346124019 | 19.3289276150945 | 13.1255103140479 | 6.40265188186281 |
| chr12 | 909855 | 909910 | + | 8.03168461498692 | 2.27399148412876 | 1.19322821036799 | 10.6710864697714 |
| chr12 | 909943 | 909972 | + | 10.0396057687336 | 12.5069531627082 | 9.54582568294392 | 7.46976052883995 |
| chr12 | 910924 | 910963 | + | 3.01188173062009 | 4.54798296825753 | 1.19322821036799 | 10.6710864697714 |
| chr12 | 910995 | 911008 | + | 18.0712903837206 | 17.0549361309657 | 5.96614105183995 | 18.1408469986113 |
| chr12 | 911350 | 911366 | + | 3.01188173062009 | 3.41098722619315 | 1.19322821036799 | 1.06710864697714 |
| chr12 | 1491598 | 1491624 | + | 6.02376346124019 | 5.68497871032191 | 3.57968463110397 | 1.06710864697714 |
| chr12 | 1495725 | 1495750 | + | 40.1584230749346 | 20.4659233571589 | 4.77291284147196 | 10.6710864697714 |
| chr12 | 1786188 | 1786209 | + | 2.00792115374673 | 3.41098722619315 | 2.38645642073598 | 3.20132594093141 |
| chr12 | 1786608 | 1786620 | + | 33.130699036821 | 35.2468680039958 | 13.1255103140479 | 16.006629704657 |
| chr12 | 1786831 | 1786863 | + | 1.00396057687336 | 2.27399148412876 | 4.77291284147196 | 3.20132594093141 |
| chr12 | 1787825 | 1787844 | + | 1.00396057687336 | 2.27399148412876 | 2.38645642073598 | 1.06710864697714 |
| chr12 | 1788262 | 1788273 | + | 2.00792115374673 | 3.41098722619315 | 2.38645642073598 | 1.06710864697714 |
| chr12 | 1788303 | 1788304 | + | 1.00396057687336 | 2.27399148412876 | 1.19322821036799 | 1.06710864697714 |
| chr12 | 1788369 | 1788420 | + | 16.0633692299738 | 21.6029190992233 | 4.77291284147196 | 10.6710864697714 |
| chr12 | 1788452 | 1788462 | + | 13.0514874993537 | 20.4659233571589 | 8.35259747257593 | 7.46976052883995 |
| chr12 | 1788565 | 1788649 | + | 21.0831721143407 | 68.2197445238629 | 25.0577924177278 | 22.4092815865198 |
| chr12 | 2803284 | 2803301 | + | 3.01188173062009 | 1.13699574206438 | 2.38645642073598 | 2.13421729395427 |
| chr12 | 2803401 | 2803434 | + | 17.0673298068472 | 3.41098722619315 | 7.15936926220794 | 11.7381951167485 |
| chr12 | 2803678 | 2803711 | + | 15.0594086531005 | 4.54798296825753 | 5.96614105183995 | 19.2079556455884 |
| chr12 | 2803752 | 2803778 | + | 3.01188173062009 | 1.13699574206438 | 1.19322821036799 | 5.33554323488568 |
| chr12 | 2803818 | 2803843 | + | 11.043566345607 | 2.27399148412876 | 2.38645642073598 | 2.13421729395427 |
| chr12 | 2803885 | 2803896 | + | 25.0990144218341 | 13.6439489047726 | 9.54582568294392 | 12.8053037637256 |
| chr12 | 2857711 | 2857735 | - | 25.0990144218341 | 25.0139063254164 | 2.38645642073598 | 2.13421729395427 |
| chr12 | 2888575 | 2888593 | + | 6.02376346124019 | 4.54798296825753 | 4.77291284147196 | 1.06710864697714 |
| chr12 | 2888776 | 2888809 | + | 4.01584230749346 | 3.41098722619315 | 3.57968463110397 | 2.13421729395427 |
| chr12 | 2889013 | 2889018 | + | 3.01188173062009 | 3.41098722619315 | 2.38645642073598 | 1.06710864697714 |
| chr12 | 2889106 | 2889141 | + | 22.087132691214 | 28.4248935516096 | 7.15936926220794 | 17.0737383516342 |
| chr12 | 2939884 | 2939925 | + | 6.02376346124019 | 17.0549361309657 | 9.54582568294392 | 2.13421729395427 |
| chr12 | 2940982 | 2940989 | + | 7.02772403811355 | 7.95897019445067 | 10.7390538933119 | 7.46976052883995 |
| chr12 | 3040536 | 3040658 | + | 13.0514874993537 | 19.3289276150945 | 4.77291284147196 | 16.006629704657 |
| chr12 | 3043807 | 3043846 | + | 3.01188173062009 | 2.27399148412876 | 3.57968463110397 | 1.06710864697714 |
| chr12 | 4488289 | 4488292 | - | 6.02376346124019 | 2.27399148412876 | 1.19322821036799 | 1.06710864697714 |
| chr12 | 4488574 | 4488577 | - | 1.00396057687336 | 1.13699574206438 | 3.57968463110397 | 1.06710864697714 |
| chr12 | 4559002 | 4559022 | + | 2.00792115374673 | 1.13699574206438 | 3.57968463110397 | 5.33554323488568 |
| chr12 | 4559352 | 4559359 | + | 4.01584230749346 | 17.0549361309657 | 1.19322821036799 | 5.33554323488568 |
| chr12 | 4559608 | 4559612 | + | 17.0673298068472 | 27.2878978095452 | 7.15936926220794 | 7.46976052883995 |
| chr12 | 4559691 | 4559700 | + | 2.00792115374673 | 4.54798296825753 | 2.38645642073598 | 2.13421729395427 |
| chr12 | 4613763 | 4613790 | + | 8.03168461498692 | 12.5069531627082 | 5.96614105183995 | 3.20132594093141 |
| chr12 | 4654372 | 4654407 | + | 9.03564519186028 | 7.95897019445067 | 10.7390538933119 | 13.8724124107028 |
| chr12 | 4654420 | 4654442 | + | 3.01188173062009 | 6.82197445238629 | 10.7390538933119 | 17.0737383516342 |
| chr12 | 4654833 | 4654836 | + | 7.02772403811355 | 4.54798296825753 | 13.1255103140479 | 6.40265188186281 |
| chr12 | 4654894 | 4654905 | + | 12.0475269224804 | 2.27399148412876 | 4.77291284147196 | 4.26843458790854 |
| chr12 | 4657758 | 4657805 | + | 14.0554480762271 | 10.2329616785794 | 13.1255103140479 | 11.7381951167485 |
| chr12 | 4659096 | 4659106 | + | 5.01980288436682 | 11.3699574206438 | 8.35259747257593 | 5.33554323488568 |
| chr12 | 4662531 | 4662572 | + | 34.1346596136944 | 30.6988850357383 | 21.4781077866238 | 22.4092815865198 |
| chr12 | 4662586 | 4662611 | + | 13.0514874993537 | 20.4659233571589 | 3.57968463110397 | 11.7381951167485 |
| chr12 | 4687157 | 4687193 | + | 44.174265382428 | 57.9867828452835 | 22.6713359969918 | 26.6777161744284 |
| chr12 | 6238027 | 6238043 | + | 138.546559608524 | 102.329616785794 | 1.19322821036799 | 2.13421729395427 |
| chr12 | 6328790 | 6328850 | - | 40.1584230749346 | 55.7127913611547 | 8.35259747257593 | 13.8724124107028 |
| chr12 | 6328973 | 6329016 | - | 6.02376346124019 | 3.41098722619315 | 2.38645642073598 | 9.60397782279422 |
| chr12 | 6386434 | 6386445 | + | 7.02772403811355 | 3.41098722619315 | 2.38645642073598 | 3.20132594093141 |
| chr12 | 6390959 | 6391007 | + | 7.02772403811355 | 3.41098722619315 | 8.35259747257593 | 8.53686917581709 |
| chr12 | 6391015 | 6391056 | + | 10.0396057687336 | 13.6439489047726 | 11.9322821036799 | 14.9395210576799 |
| chr12 | 6391097 | 6391105 | + | 5.01980288436682 | 2.27399148412876 | 2.38645642073598 | 3.20132594093141 |
| chr12 | 6391440 | 6391521 | + | 88.3485307648561 | 97.7816338175368 | 29.8307052591997 | 78.966039876308 |
| chr12 | 6492189 | 6492291 | - | 95.3762548029696 | 108.014595496116 | 58.4681823080315 | 104.576647403759 |
| chr12 | 6531778 | 6531792 | + | 4.01584230749346 | 6.82197445238629 | 3.57968463110397 | 11.7381951167485 |
| chr12 | 6531822 | 6531846 | + | 23.0910932680874 | 14.780944646837 | 5.96614105183995 | 6.40265188186281 |
| chr12 | 6531874 | 6531912 | + | 12.0475269224804 | 7.95897019445067 | 7.15936926220794 | 5.33554323488568 |
| chr12 | 6538169 | 6538194 | + | 47.1861471130481 | 17.0549361309657 | 44.1494437836156 | 49.0869977609482 |
| chr12 | 6538262 | 6538346 | + | 954.76650860657 | 842.513844869707 | 696.845274854906 | 1115.12853609111 |
| chr12 | 6556899 | 6557003 | - | 23.0910932680874 | 21.6029190992233 | 13.1255103140479 | 8.53686917581709 |
| chr12 | 6557088 | 6557213 | - | 74.293082688629 | 34.1098722619315 | 11.9322821036799 | 42.6843458790854 |
| chr12 | 6557247 | 6557399 | - | 34.1346596136944 | 26.1509020674808 | 45.3426719939836 | 40.5501285851312 |
| chr12 | 6557544 | 6557584 | - | 5.01980288436682 | 5.68497871032191 | 8.35259747257593 | 7.46976052883995 |
| chr12 | 6566100 | 6566269 | - | 107.42378172545 | 57.9867828452835 | 116.936364616063 | 35.2145853502455 |
| chr12 | 6566299 | 6566326 | - | 7.02772403811355 | 7.95897019445067 | 10.7390538933119 | 2.13421729395427 |
| chr12 | 6570107 | 6570120 | - | 3.01188173062009 | 5.68497871032191 | 1.19322821036799 | 2.13421729395427 |
| chr12 | 6570143 | 6570260 | - | 53.2099105742883 | 55.7127913611547 | 82.3327465153913 | 104.576647403759 |
| chr12 | 6570356 | 6570388 | - | 11.043566345607 | 9.09596593651506 | 5.96614105183995 | 16.006629704657 |
| chr12 | 6570441 | 6570446 | - | 17.0673298068472 | 14.780944646837 | 10.7390538933119 | 36.2816939972226 |
| chr12 | 6570510 | 6570530 | - | 3.01188173062009 | 1.13699574206438 | 3.57968463110397 | 3.20132594093141 |
| chr12 | 6650612 | 6650620 | - | 2.00792115374673 | 11.3699574206438 | 3.57968463110397 | 3.20132594093141 |
| chr12 | 6650649 | 6650662 | - | 7.02772403811355 | 12.5069531627082 | 13.1255103140479 | 9.60397782279422 |
| chr12 | 6666884 | 6666923 | - | 2.00792115374673 | 3.41098722619315 | 2.38645642073598 | 1.06710864697714 |
| chr12 | 6667423 | 6667449 | - | 8.03168461498692 | 4.54798296825753 | 10.7390538933119 | 5.33554323488568 |
| chr12 | 6667642 | 6667644 | - | 1.00396057687336 | 3.41098722619315 | 3.57968463110397 | 4.26843458790854 |
| chr12 | 6667669 | 6667691 | - | 3.01188173062009 | 3.41098722619315 | 5.96614105183995 | 1.06710864697714 |
| chr12 | 6731063 | 6731078 | + | 10.0396057687336 | 1.13699574206438 | 2.38645642073598 | 1.06710864697714 |
| chr12 | 6731180 | 6731229 | + | 13.0514874993537 | 11.3699574206438 | 14.3187385244159 | 3.20132594093141 |
| chr12 | 6731361 | 6731377 | + | 7.02772403811355 | 2.27399148412876 | 3.57968463110397 | 1.06710864697714 |
| chr12 | 6731459 | 6731479 | + | 4.01584230749346 | 3.41098722619315 | 5.96614105183995 | 3.20132594093141 |
| chr12 | 6731516 | 6731574 | + | 32.1267384599477 | 17.0549361309657 | 7.15936926220794 | 13.8724124107028 |
| chr12 | 6731691 | 6731734 | + | 9.03564519186028 | 18.1919318730301 | 3.57968463110397 | 2.13421729395427 |
| chr12 | 6731759 | 6731767 | + | 10.0396057687336 | 7.95897019445067 | 3.57968463110397 | 1.06710864697714 |
| chr12 | 6748007 | 6748087 | - | 7.02772403811355 | 13.6439489047726 | 15.5119667347839 | 10.6710864697714 |
| chr12 | 6748127 | 6748166 | - | 18.0712903837206 | 14.780944646837 | 20.2848795762558 | 41.6172372321083 |
| chr12 | 6748414 | 6748439 | - | 6.02376346124019 | 2.27399148412876 | 16.7051949451519 | 7.46976052883995 |
| chr12 | 6748517 | 6748534 | - | 1.00396057687336 | 2.27399148412876 | 1.19322821036799 | 3.20132594093141 |
| chr12 | 6770493 | 6770514 | + | 50.1980288436682 | 21.6029190992233 | 13.1255103140479 | 12.8053037637256 |
| chr12 | 6770541 | 6770625 | + | 37.1465413443145 | 17.0549361309657 | 11.9322821036799 | 7.46976052883995 |
| chr12 | 6770867 | 6770908 | + | 17.0673298068472 | 3.41098722619315 | 3.57968463110397 | 10.6710864697714 |
| chr12 | 6770927 | 6770928 | + | 1.00396057687336 | 1.13699574206438 | 1.19322821036799 | 1.06710864697714 |
| chr12 | 6848859 | 6848860 | - | 13.0514874993537 | 6.82197445238629 | 3.57968463110397 | 5.33554323488568 |
| chr12 | 6848886 | 6848918 | - | 51.2019894205416 | 27.2878978095452 | 17.8984231555198 | 17.0737383516342 |
| chr12 | 6848954 | 6848970 | - | 14.0554480762271 | 3.41098722619315 | 8.35259747257593 | 5.33554323488568 |
| chr12 | 6866566 | 6866611 | + | 14.0554480762271 | 21.6029190992233 | 4.77291284147196 | 4.26843458790854 |
| chr12 | 6870430 | 6870464 | + | 3.01188173062009 | 3.41098722619315 | 2.38645642073598 | 4.26843458790854 |
| chr12 | 6870489 | 6870515 | + | 2.00792115374673 | 1.13699574206438 | 2.38645642073598 | 3.20132594093141 |
| chr12 | 6870624 | 6870643 | + | 7.02772403811355 | 7.95897019445067 | 2.38645642073598 | 14.9395210576799 |
| chr12 | 6870776 | 6870835 | + | 21.0831721143407 | 11.3699574206438 | 5.96614105183995 | 23.476390233497 |
| chr12 | 6870882 | 6870896 | + | 4.01584230749346 | 1.13699574206438 | 3.57968463110397 | 2.13421729395427 |
| chr12 | 6871015 | 6871026 | - | 5.01980288436682 | 3.41098722619315 | 1.19322821036799 | 1.06710864697714 |
| chr12 | 6914188 | 6914198 | + | 2.00792115374673 | 2.27399148412876 | 2.38645642073598 | 1.06710864697714 |
| chr12 | 6923523 | 6923568 | + | 1.00396057687336 | 3.41098722619315 | 2.38645642073598 | 3.20132594093141 |
| chr12 | 6942181 | 6942219 | + | 27.1069355755808 | 21.6029190992233 | 8.35259747257593 | 6.40265188186281 |
| chr12 | 6942246 | 6942298 | + | 18.0712903837206 | 28.4248935516096 | 5.96614105183995 | 4.26843458790854 |
| chr12 | 6945951 | 6945976 | + | 97.3841759567164 | 133.028501821533 | 65.6275515702394 | 59.7580842307196 |
| chr12 | 6965370 | 6965410 | - | 445.758496131774 | 545.757956190903 | 457.00640457094 | 709.627250239795 |
| chr12 | 6965450 | 6965477 | - | 38.1505019211879 | 26.1509020674808 | 22.6713359969918 | 28.8119334683827 |
| chr12 | 6965504 | 6965536 | - | 16.0633692299738 | 17.0549361309657 | 5.96614105183995 | 10.6710864697714 |
| chr12 | 6965654 | 6965659 | - | 3.01188173062009 | 1.13699574206438 | 2.38645642073598 | 3.20132594093141 |
| chr12 | 6966423 | 6966492 | - | 4.01584230749346 | 5.68497871032191 | 4.77291284147196 | 11.7381951167485 |
| chr12 | 6975749 | 6975849 | + | 1055.16256629391 | 948.254448881695 | 515.474586878971 | 600.782168248127 |
| chr12 | 6975881 | 6975944 | + | 10.0396057687336 | 32.9728765198671 | 7.15936926220794 | 36.2816939972226 |
| chr12 | 6976799 | 6976826 | - | 2.00792115374673 | 2.27399148412876 | 4.77291284147196 | 2.13421729395427 |
| chr12 | 6977760 | 6977767 | + | 1.00396057687336 | 2.27399148412876 | 4.77291284147196 | 2.13421729395427 |
| chr12 | 6979655 | 6979671 | + | 6.02376346124019 | 2.27399148412876 | 2.38645642073598 | 3.20132594093141 |
| chr12 | 6979906 | 6979909 | + | 6.02376346124019 | 10.2329616785794 | 5.96614105183995 | 1.06710864697714 |
| chr12 | 7096272 | 7096297 | - | 10.0396057687336 | 7.95897019445067 | 4.77291284147196 | 7.46976052883995 |
| chr12 | 7108542 | 7108700 | + | 387.528782673119 | 486.634177603555 | 99.0379414605431 | 90.7042349930565 |
| chr12 | 7158800 | 7158909 | + | 23.0910932680874 | 25.0139063254164 | 17.8984231555198 | 23.476390233497 |
| chr12 | 7210314 | 7210378 | + | 15.0594086531005 | 10.2329616785794 | 16.7051949451519 | 13.8724124107028 |
| chr12 | 7210446 | 7210456 | + | 1.00396057687336 | 2.27399148412876 | 4.77291284147196 | 2.13421729395427 |
| chr12 | 7210480 | 7210507 | + | 1.00396057687336 | 1.13699574206438 | 5.96614105183995 | 5.33554323488568 |
| chr12 | 7210525 | 7210529 | + | 2.00792115374673 | 2.27399148412876 | 2.38645642073598 | 3.20132594093141 |
| chr12 | 7211238 | 7211279 | + | 2.00792115374673 | 2.27399148412876 | 3.57968463110397 | 10.6710864697714 |
| chr12 | 7919328 | 7919351 | - | 1.00396057687336 | 3.41098722619315 | 17.8984231555198 | 20.2750642925656 |
| chr12 | 7920542 | 7920619 | - | 1.00396057687336 | 10.2329616785794 | 7.15936926220794 | 28.8119334683827 |
| chr12 | 7920697 | 7920762 | - | 1.00396057687336 | 6.82197445238629 | 4.77291284147196 | 22.4092815865198 |
| chr12 | 7920812 | 7920860 | - | 2.00792115374673 | 1.13699574206438 | 23.8645642073598 | 74.6976052883995 |
| chr12 | 7921260 | 7921280 | - | 5.01980288436682 | 5.68497871032191 | 5.96614105183995 | 10.6710864697714 |
| chr12 | 7921306 | 7921359 | - | 1.00396057687336 | 2.27399148412876 | 7.15936926220794 | 10.6710864697714 |
| chr12 | 8052905 | 8052906 | + | 1.00396057687336 | 2.27399148412876 | 1.19322821036799 | 1.06710864697714 |
| chr12 | 8055433 | 8055443 | + | 8.03168461498692 | 6.82197445238629 | 7.15936926220794 | 2.13421729395427 |
| chr12 | 8096384 | 8096388 | + | 12.0475269224804 | 13.6439489047726 | 10.7390538933119 | 6.40265188186281 |
| chr12 | 8097593 | 8097607 | + | 4.01584230749346 | 7.95897019445067 | 3.57968463110397 | 2.13421729395427 |
| chr12 | 8773919 | 8773935 | + | 11.043566345607 | 25.0139063254164 | 3.57968463110397 | 3.20132594093141 |
| chr12 | 8773995 | 8774013 | + | 18.0712903837206 | 20.4659233571589 | 2.38645642073598 | 1.06710864697714 |
| chr12 | 8774042 | 8774075 | + | 13.0514874993537 | 27.2878978095452 | 4.77291284147196 | 4.26843458790854 |
| chr12 | 8774410 | 8774483 | + | 6.02376346124019 | 30.6988850357383 | 3.57968463110397 | 1.06710864697714 |
| chr12 | 8774508 | 8774563 | + | 72.2851615348822 | 113.699574206438 | 7.15936926220794 | 7.46976052883995 |
| chr12 | 8774624 | 8774688 | + | 22.087132691214 | 21.6029190992233 | 4.77291284147196 | 8.53686917581709 |
| chr12 | 8775026 | 8775088 | + | 38.1505019211879 | 57.9867828452835 | 2.38645642073598 | 4.26843458790854 |
| chr12 | 8775627 | 8775647 | + | 19.0752509605939 | 34.1098722619315 | 1.19322821036799 | 2.13421729395427 |
| chr12 | 8775852 | 8775880 | + | 10.0396057687336 | 37.5208594881246 | 1.19322821036799 | 1.06710864697714 |
| chr12 | 8776128 | 8776153 | + | 113.44754518669 | 129.61751459534 | 14.3187385244159 | 5.33554323488568 |
| chr12 | 8776318 | 8776322 | + | 11.043566345607 | 11.3699574206438 | 4.77291284147196 | 2.13421729395427 |
| chr12 | 8776344 | 8776381 | + | 20.0792115374673 | 46.6168254246397 | 7.15936926220794 | 1.06710864697714 |
| chr12 | 8776403 | 8776415 | + | 6.02376346124019 | 9.09596593651506 | 2.38645642073598 | 1.06710864697714 |
| chr12 | 8776605 | 8776617 | + | 15.0594086531005 | 20.4659233571589 | 3.57968463110397 | 2.13421729395427 |
| chr12 | 8776640 | 8776645 | + | 5.01980288436682 | 9.09596593651506 | 2.38645642073598 | 1.06710864697714 |
| chr12 | 8776920 | 8777038 | + | 178.704982683459 | 523.018041349616 | 11.9322821036799 | 17.0737383516342 |
| chr12 | 8780688 | 8780709 | + | 10.0396057687336 | 18.1919318730301 | 2.38645642073598 | 1.06710864697714 |
| chr12 | 8940388 | 8940406 | + | 18.0712903837206 | 32.9728765198671 | 5.96614105183995 | 25.6106075274513 |
| chr12 | 8941405 | 8941417 | - | 5.01980288436682 | 2.27399148412876 | 1.19322821036799 | 3.20132594093141 |
| chr12 | 8941499 | 8941517 | - | 10.0396057687336 | 6.82197445238629 | 4.77291284147196 | 8.53686917581709 |
| chr12 | 8941579 | 8941624 | - | 14.0554480762271 | 13.6439489047726 | 16.7051949451519 | 5.33554323488568 |
| chr12 | 9694939 | 9694942 | + | 4.01584230749346 | 4.54798296825753 | 38.1833027317757 | 8.53686917581709 |
| chr12 | 9695282 | 9695286 | + | 213.843602874027 | 111.425582722309 | 76.3666054635513 | 199.549316984724 |
| chr12 | 9695652 | 9695654 | + | 3.01188173062009 | 4.54798296825753 | 11.9322821036799 | 10.6710864697714 |
| chr12 | 9853279 | 9853281 | - | 15.0594086531005 | 11.3699574206438 | 3.57968463110397 | 1.06710864697714 |
| chr12 | 10221855 | 10221960 | + | 42.1663442286813 | 57.9867828452835 | 2.38645642073598 | 2.13421729395427 |
| chr12 | 10222420 | 10222430 | + | 4.01584230749346 | 17.0549361309657 | 1.19322821036799 | 1.06710864697714 |
| chr12 | 10222691 | 10222744 | + | 50.1980288436682 | 142.124467758048 | 1.19322821036799 | 4.26843458790854 |
| chr12 | 10223015 | 10223034 | + | 35.1386201905678 | 97.7816338175368 | 5.96614105183995 | 3.20132594093141 |
| chr12 | 10379835 | 10379842 | + | 1.00396057687336 | 5.68497871032191 | 1.19322821036799 | 2.13421729395427 |
| chr12 | 10606091 | 10606101 | - | 10.0396057687336 | 9.09596593651506 | 3.57968463110397 | 12.8053037637256 |
| chr12 | 10699124 | 10699154 | - | 13.0514874993537 | 17.0549361309657 | 11.9322821036799 | 2.13421729395427 |
| chr12 | 10699231 | 10699261 | - | 51.2019894205416 | 56.8497871032191 | 13.1255103140479 | 16.006629704657 |
| chr12 | 10699294 | 10699315 | - | 5.01980288436682 | 4.54798296825753 | 2.38645642073598 | 1.06710864697714 |
| chr12 | 10699402 | 10699471 | - | 43.1703048055547 | 29.5618892936739 | 23.8645642073598 | 33.0803680562912 |
| chr12 | 10699636 | 10699657 | - | 10.0396057687336 | 14.780944646837 | 2.38645642073598 | 6.40265188186281 |
| chr12 | 10701257 | 10701275 | - | 3.01188173062009 | 3.41098722619315 | 1.19322821036799 | 2.13421729395427 |
| chr12 | 11553908 | 11553911 | + | 2.00792115374673 | 1.13699574206438 | 1.19322821036799 | 1.06710864697714 |
| chr12 | 11892462 | 11892471 | + | 8.03168461498692 | 11.3699574206438 | 2.38645642073598 | 1.06710864697714 |
| chr12 | 11893382 | 11893384 | + | 3.01188173062009 | 2.27399148412876 | 3.57968463110397 | 3.20132594093141 |
| chr12 | 11893612 | 11893614 | + | 4.01584230749346 | 3.41098722619315 | 1.19322821036799 | 2.13421729395427 |
| chr12 | 11895244 | 11895261 | + | 1.00396057687336 | 10.2329616785794 | 4.77291284147196 | 2.13421729395427 |
| chr12 | 12644498 | 12644502 | + | 18.0712903837206 | 36.3838637460602 | 7.15936926220794 | 10.6710864697714 |
| chr12 | 12645009 | 12645014 | + | 5.01980288436682 | 11.3699574206438 | 2.38645642073598 | 1.06710864697714 |
| chr12 | 12721329 | 12721369 | + | 208.82379998966 | 288.796918484353 | 68.0140079909754 | 74.6976052883995 |
| chr12 | 12721715 | 12721777 | + | 11.043566345607 | 11.3699574206438 | 1.19322821036799 | 4.26843458790854 |
| chr12 | 12721989 | 12722031 | + | 5.01980288436682 | 13.6439489047726 | 5.96614105183995 | 4.26843458790854 |
| chr12 | 12722185 | 12722189 | + | 1.00396057687336 | 2.27399148412876 | 2.38645642073598 | 1.06710864697714 |
| chr12 | 12722210 | 12722224 | + | 21.0831721143407 | 52.3018041349616 | 3.57968463110397 | 6.40265188186281 |
| chr12 | 12722270 | 12722275 | + | 1.00396057687336 | 12.5069531627082 | 1.19322821036799 | 1.06710864697714 |
| chr12 | 12722307 | 12722341 | + | 6.02376346124019 | 5.68497871032191 | 1.19322821036799 | 1.06710864697714 |
| chr12 | 12788979 | 12788984 | + | 1.00396057687336 | 4.54798296825753 | 3.57968463110397 | 1.06710864697714 |
| chr12 | 12829699 | 12829700 | + | 5.01980288436682 | 10.2329616785794 | 1.19322821036799 | 3.20132594093141 |
| chr12 | 12829846 | 12829893 | + | 1.00396057687336 | 5.68497871032191 | 1.19322821036799 | 1.06710864697714 |
| chr12 | 12829923 | 12829959 | + | 7.02772403811355 | 10.2329616785794 | 3.57968463110397 | 1.06710864697714 |
| chr12 | 12927700 | 12927771 | + | 15.0594086531005 | 28.4248935516096 | 161.085808399679 | 166.468948928433 |
| chr12 | 12974918 | 12974969 | - | 36.1425807674411 | 98.9186295596012 | 3.57968463110397 | 6.40265188186281 |
| chr12 | 13081017 | 13081018 | + | 1.00396057687336 | 1.13699574206438 | 1.19322821036799 | 1.06710864697714 |
| chr12 | 14786567 | 14786582 | - | 4.01584230749346 | 14.780944646837 | 13.1255103140479 | 8.53686917581709 |
| chr12 | 14786724 | 14786733 | - | 8.03168461498692 | 5.68497871032191 | 5.96614105183995 | 18.1408469986113 |
| chr12 | 15902972 | 15902996 | + | 1.00396057687336 | 1.13699574206438 | 1.19322821036799 | 2.13421729395427 |
| chr12 | 15903110 | 15903137 | + | 4.01584230749346 | 2.27399148412876 | 2.38645642073598 | 1.06710864697714 |
| chr12 | 15903157 | 15903183 | + | 86.3406096111093 | 72.7677274921204 | 31.0239334695677 | 74.6976052883995 |
| chr12 | 15903243 | 15903337 | + | 122.48319037855 | 152.357429436627 | 110.970223564223 | 132.321472225165 |
| chr12 | 15903368 | 15903452 | + | 22.087132691214 | 44.3428339405109 | 23.8645642073598 | 12.8053037637256 |
| chr12 | 16036763 | 16036769 | + | 5.01980288436682 | 12.5069531627082 | 1.19322821036799 | 1.06710864697714 |
| chr12 | 16036835 | 16036893 | + | 55.217831728035 | 94.3706465913437 | 4.77291284147196 | 8.53686917581709 |
| chr12 | 16036953 | 16036985 | + | 22.087132691214 | 47.753821166704 | 8.35259747257593 | 4.26843458790854 |
| chr12 | 16037046 | 16037140 | + | 163.645574030358 | 181.919318730301 | 17.8984231555198 | 14.9395210576799 |
| chr12 | 16037238 | 16037260 | + | 23.0910932680874 | 25.0139063254164 | 1.19322821036799 | 2.13421729395427 |
| chr12 | 16363792 | 16364386 | + | 3470.69171425122 | 2671.9399938513 | 1557.16281453023 | 983.874172512919 |
| chr12 | 19151126 | 19151128 | + | 1.00396057687336 | 1.13699574206438 | 1.19322821036799 | 1.06710864697714 |
| chr12 | 19375784 | 19375864 | + | 8.03168461498692 | 20.4659233571589 | 9.54582568294392 | 8.53686917581709 |
| chr12 | 19518798 | 19518832 | + | 2.00792115374673 | 9.09596593651506 | 3.57968463110397 | 10.6710864697714 |
| chr12 | 19519462 | 19519487 | + | 5.01980288436682 | 4.54798296825753 | 1.19322821036799 | 2.13421729395427 |
| chr12 | 19519766 | 19519813 | + | 156.617849992245 | 181.919318730301 | 8.35259747257593 | 23.476390233497 |
| chr12 | 19519907 | 19519919 | + | 1.00396057687336 | 7.95897019445067 | 1.19322821036799 | 1.06710864697714 |
| chr12 | 21515803 | 21515818 | + | 4.01584230749346 | 1.13699574206438 | 2.38645642073598 | 4.26843458790854 |
| chr12 | 21515915 | 21515972 | + | 15.0594086531005 | 20.4659233571589 | 8.35259747257593 | 7.46976052883995 |
| chr12 | 21515995 | 21516032 | + | 15.0594086531005 | 14.780944646837 | 19.0916513658878 | 8.53686917581709 |
| chr12 | 21516053 | 21516066 | + | 4.01584230749346 | 3.41098722619315 | 11.9322821036799 | 6.40265188186281 |
| chr12 | 21516107 | 21516163 | + | 8.03168461498692 | 19.3289276150945 | 10.7390538933119 | 8.53686917581709 |
| chr12 | 21516192 | 21516208 | + | 5.01980288436682 | 3.41098722619315 | 10.7390538933119 | 1.06710864697714 |
| chr12 | 21516237 | 21516265 | + | 7.02772403811355 | 11.3699574206438 | 5.96614105183995 | 3.20132594093141 |
| chr12 | 21516287 | 21516290 | + | 1.00396057687336 | 3.41098722619315 | 2.38645642073598 | 1.06710864697714 |
| chr12 | 21516392 | 21516420 | + | 7.02772403811355 | 3.41098722619315 | 5.96614105183995 | 6.40265188186281 |
| chr12 | 21516489 | 21516497 | + | 10.0396057687336 | 3.41098722619315 | 7.15936926220794 | 2.13421729395427 |
| chr12 | 21516553 | 21516582 | + | 6.02376346124019 | 5.68497871032191 | 13.1255103140479 | 13.8724124107028 |
| chr12 | 21516724 | 21516728 | + | 3.01188173062009 | 4.54798296825753 | 2.38645642073598 | 2.13421729395427 |
| chr12 | 21516943 | 21516955 | + | 1.00396057687336 | 1.13699574206438 | 3.57968463110397 | 6.40265188186281 |
| chr12 | 21517010 | 21517013 | + | 4.01584230749346 | 3.41098722619315 | 7.15936926220794 | 1.06710864697714 |
| chr12 | 21517227 | 21517266 | + | 5.01980288436682 | 4.54798296825753 | 10.7390538933119 | 3.20132594093141 |
| chr12 | 21517300 | 21517318 | + | 5.01980288436682 | 4.54798296825753 | 1.19322821036799 | 1.06710864697714 |
| chr12 | 21517369 | 21517380 | + | 3.01188173062009 | 3.41098722619315 | 2.38645642073598 | 2.13421729395427 |
| chr12 | 21517420 | 21517423 | + | 1.00396057687336 | 1.13699574206438 | 3.57968463110397 | 1.06710864697714 |
| chr12 | 21517546 | 21517566 | + | 10.0396057687336 | 13.6439489047726 | 13.1255103140479 | 12.8053037637256 |
| chr12 | 21517676 | 21517684 | + | 28.1108961524542 | 61.3977700714766 | 25.0577924177278 | 7.46976052883995 |
| chr12 | 21517705 | 21517789 | + | 90.3564519186028 | 114.836569948503 | 96.6514850398071 | 118.449059814462 |
| chr12 | 21517816 | 21517821 | + | 2.00792115374673 | 3.41098722619315 | 1.19322821036799 | 2.13421729395427 |
| chr12 | 21517854 | 21517873 | + | 6.02376346124019 | 7.95897019445067 | 11.9322821036799 | 7.46976052883995 |
| chr12 | 21517896 | 21517901 | + | 2.00792115374673 | 3.41098722619315 | 4.77291284147196 | 1.06710864697714 |
| chr12 | 21517925 | 21518015 | + | 259.021828833328 | 518.470058381358 | 397.34499405254 | 227.29414180613 |
| chr12 | 21635375 | 21635430 | - | 206.815878835913 | 237.632110091456 | 177.79100334483 | 142.992558694936 |
| chr12 | 21635451 | 21635501 | - | 47.1861471130481 | 45.4798296825753 | 50.1155848354556 | 32.0132594093141 |
| chr12 | 22065587 | 22065635 | + | 15.0594086531005 | 9.09596593651506 | 48.9223566250876 | 41.6172372321083 |
| chr12 | 22448871 | 22448913 | - | 4.01584230749346 | 5.68497871032191 | 1.19322821036799 | 3.20132594093141 |
| chr12 | 22685500 | 22685538 | + | 6.02376346124019 | 3.41098722619315 | 8.35259747257593 | 11.7381951167485 |
| chr12 | 22685572 | 22685574 | + | 2.00792115374673 | 3.41098722619315 | 2.38645642073598 | 3.20132594093141 |
| chr12 | 22686023 | 22686156 | + | 9.03564519186028 | 4.54798296825753 | 75.1733772531833 | 101.375321462828 |
| chr12 | 25204121 | 25204122 | + | 4.01584230749346 | 2.27399148412876 | 1.19322821036799 | 3.20132594093141 |
| chr12 | 25204390 | 25204393 | + | 1.00396057687336 | 3.41098722619315 | 4.77291284147196 | 8.53686917581709 |
| chr12 | 25204466 | 25204490 | + | 18.0712903837206 | 28.4248935516096 | 22.6713359969918 | 41.6172372321083 |
| chr12 | 25204642 | 25204672 | + | 4.01584230749346 | 21.6029190992233 | 22.6713359969918 | 16.006629704657 |
| chr12 | 25204686 | 25204692 | + | 1.00396057687336 | 1.13699574206438 | 3.57968463110397 | 1.06710864697714 |
| chr12 | 25204714 | 25204734 | + | 10.0396057687336 | 17.0549361309657 | 19.0916513658878 | 8.53686917581709 |
| chr12 | 25204791 | 25204793 | + | 5.01980288436682 | 5.68497871032191 | 8.35259747257593 | 5.33554323488568 |
| chr12 | 25204833 | 25204837 | + | 10.0396057687336 | 17.0549361309657 | 28.6374770488317 | 17.0737383516342 |
| chr12 | 25204867 | 25204876 | + | 7.02772403811355 | 18.1919318730301 | 42.9562155732476 | 25.6106075274513 |
| chr12 | 25204940 | 25204960 | + | 2.00792115374673 | 3.41098722619315 | 3.57968463110397 | 5.33554323488568 |
| chr12 | 25205329 | 25205343 | - | 2.00792115374673 | 3.41098722619315 | 1.19322821036799 | 3.20132594093141 |
| chr12 | 25206464 | 25206468 | - | 5.01980288436682 | 1.13699574206438 | 1.19322821036799 | 3.20132594093141 |
| chr12 | 25206580 | 25206636 | - | 6.02376346124019 | 3.41098722619315 | 8.35259747257593 | 48.0198891139711 |
| chr12 | 25208004 | 25208058 | - | 6.02376346124019 | 6.82197445238629 | 14.3187385244159 | 8.53686917581709 |
| chr12 | 25208195 | 25208201 | - | 5.01980288436682 | 6.82197445238629 | 3.57968463110397 | 7.46976052883995 |
| chr12 | 25208234 | 25208330 | - | 15.0594086531005 | 26.1509020674808 | 19.0916513658878 | 21.3421729395427 |
| chr12 | 25208532 | 25208540 | - | 17.0673298068472 | 12.5069531627082 | 8.35259747257593 | 9.60397782279422 |
| chr12 | 25208680 | 25208717 | - | 7.02772403811355 | 5.68497871032191 | 7.15936926220794 | 5.33554323488568 |
| chr12 | 25209909 | 25209911 | - | 1.00396057687336 | 1.13699574206438 | 1.19322821036799 | 1.06710864697714 |
| chr12 | 26337316 | 26337328 | - | 2.00792115374673 | 3.41098722619315 | 1.19322821036799 | 1.06710864697714 |
| chr12 | 26905214 | 26905215 | - | 8.03168461498692 | 13.6439489047726 | 1.19322821036799 | 2.13421729395427 |
| chr12 | 26905397 | 26905419 | - | 25.0990144218341 | 11.3699574206438 | 4.77291284147196 | 11.7381951167485 |
| chr12 | 26905446 | 26905450 | - | 3.01188173062009 | 6.82197445238629 | 4.77291284147196 | 5.33554323488568 |
| chr12 | 26905494 | 26905502 | - | 3.01188173062009 | 5.68497871032191 | 2.38645642073598 | 4.26843458790854 |
| chr12 | 26960656 | 26960709 | + | 28.1108961524542 | 26.1509020674808 | 8.35259747257593 | 17.0737383516342 |
| chr12 | 26964706 | 26964762 | + | 6.02376346124019 | 5.68497871032191 | 17.8984231555198 | 8.53686917581709 |
| chr12 | 26965766 | 26965799 | + | 6.02376346124019 | 7.95897019445067 | 14.3187385244159 | 13.8724124107028 |
| chr12 | 26966443 | 26966446 | + | 7.02772403811355 | 1.13699574206438 | 1.19322821036799 | 1.06710864697714 |
| chr12 | 26973749 | 26973852 | - | 30.1188173062009 | 35.2468680039958 | 7.15936926220794 | 9.60397782279422 |
| chr12 | 26973874 | 26973915 | - | 10.0396057687336 | 5.68497871032191 | 1.19322821036799 | 2.13421729395427 |
| chr12 | 26973935 | 26973957 | - | 12.0475269224804 | 4.54798296825753 | 1.19322821036799 | 4.26843458790854 |
| chr12 | 27028477 | 27028486 | + | 3.01188173062009 | 5.68497871032191 | 3.57968463110397 | 2.13421729395427 |
| chr12 | 27029472 | 27029494 | + | 8.03168461498692 | 15.9179403889013 | 2.38645642073598 | 3.20132594093141 |
| chr12 | 27038721 | 27038723 | + | 6.02376346124019 | 7.95897019445067 | 3.57968463110397 | 3.20132594093141 |
| chr12 | 27323729 | 27323751 | + | 4.01584230749346 | 1.13699574206438 | 1.19322821036799 | 1.06710864697714 |
| chr12 | 27695401 | 27695471 | + | 26.1029749987075 | 23.876910583352 | 2.38645642073598 | 4.26843458790854 |
| chr12 | 27737537 | 27737609 | + | 34.1346596136944 | 32.9728765198671 | 90.6853439879672 | 120.583277108416 |
| chr12 | 27755186 | 27755253 | + | 28.1108961524542 | 43.2058381984465 | 65.6275515702394 | 67.2278447595595 |
| chr12 | 27755303 | 27755467 | + | 31.1227778830743 | 28.4248935516096 | 79.9462900946553 | 36.2816939972226 |
| chr12 | 27755490 | 27755500 | + | 6.02376346124019 | 3.41098722619315 | 2.38645642073598 | 1.06710864697714 |
| chr12 | 27755588 | 27755601 | + | 8.03168461498692 | 9.09596593651506 | 25.0577924177278 | 7.46976052883995 |
| chr12 | 27755629 | 27755665 | + | 12.0475269224804 | 13.6439489047726 | 28.6374770488317 | 13.8724124107028 |
| chr12 | 27755695 | 27755714 | + | 5.01980288436682 | 3.41098722619315 | 9.54582568294392 | 2.13421729395427 |
| chr12 | 27755769 | 27755786 | + | 2.00792115374673 | 1.13699574206438 | 7.15936926220794 | 5.33554323488568 |
| chr12 | 27755814 | 27755825 | + | 7.02772403811355 | 1.13699574206438 | 7.15936926220794 | 6.40265188186281 |
| chr12 | 27756011 | 27756028 | + | 7.02772403811355 | 4.54798296825753 | 2.38645642073598 | 12.8053037637256 |
| chr12 | 27756153 | 27756197 | + | 8.03168461498692 | 6.82197445238629 | 8.35259747257593 | 8.53686917581709 |
| chr12 | 27756223 | 27756253 | + | 4.01584230749346 | 7.95897019445067 | 7.15936926220794 | 5.33554323488568 |
| chr12 | 27800284 | 27800292 | + | 82.3247673036159 | 85.2746806548286 | 13.1255103140479 | 6.40265188186281 |
| chr12 | 27802009 | 27802018 | + | 1.00396057687336 | 2.27399148412876 | 5.96614105183995 | 3.20132594093141 |
| chr12 | 28549800 | 28549812 | + | 9.03564519186028 | 10.2329616785794 | 5.96614105183995 | 8.53686917581709 |
| chr12 | 29338914 | 29338929 | - | 5.01980288436682 | 4.54798296825753 | 1.19322821036799 | 4.26843458790854 |
| chr12 | 29339353 | 29339356 | - | 1.00396057687336 | 3.41098722619315 | 1.19322821036799 | 2.13421729395427 |
| chr12 | 29339989 | 29340010 | - | 18.0712903837206 | 38.657855230189 | 31.0239334695677 | 11.7381951167485 |
| chr12 | 29340162 | 29340173 | - | 10.0396057687336 | 6.82197445238629 | 8.35259747257593 | 3.20132594093141 |
| chr12 | 29340233 | 29340249 | - | 3.01188173062009 | 5.68497871032191 | 3.57968463110397 | 4.26843458790854 |
| chr12 | 29340335 | 29340336 | - | 10.0396057687336 | 12.5069531627082 | 1.19322821036799 | 1.06710864697714 |
| chr12 | 29340418 | 29340502 | - | 31.1227778830743 | 20.4659233571589 | 16.7051949451519 | 4.26843458790854 |
| chr12 | 29340669 | 29340685 | - | 3.01188173062009 | 3.41098722619315 | 2.38645642073598 | 1.06710864697714 |
| chr12 | 29340714 | 29340744 | - | 4.01584230749346 | 9.09596593651506 | 7.15936926220794 | 3.20132594093141 |
| chr12 | 29340772 | 29340820 | - | 38.1505019211879 | 47.753821166704 | 19.0916513658878 | 28.8119334683827 |
| chr12 | 29340859 | 29340876 | - | 9.03564519186028 | 10.2329616785794 | 4.77291284147196 | 9.60397782279422 |
| chr12 | 29340900 | 29340931 | - | 7.02772403811355 | 10.2329616785794 | 2.38645642073598 | 2.13421729395427 |
| chr12 | 29341046 | 29341088 | - | 9.03564519186028 | 5.68497871032191 | 10.7390538933119 | 7.46976052883995 |
| chr12 | 29341116 | 29341149 | - | 30.1188173062009 | 31.8358807778027 | 17.8984231555198 | 14.9395210576799 |
| chr12 | 30629003 | 30629037 | - | 5.01980288436682 | 2.27399148412876 | 2.38645642073598 | 4.26843458790854 |
| chr12 | 30629101 | 30629105 | - | 2.00792115374673 | 2.27399148412876 | 3.57968463110397 | 1.06710864697714 |
| chr12 | 30629141 | 30629151 | - | 1.00396057687336 | 2.27399148412876 | 2.38645642073598 | 4.26843458790854 |
| chr12 | 30630570 | 30630575 | - | 7.02772403811355 | 1.13699574206438 | 1.19322821036799 | 2.13421729395427 |
| chr12 | 30630618 | 30630628 | - | 1.00396057687336 | 1.13699574206438 | 3.57968463110397 | 7.46976052883995 |
| chr12 | 30630775 | 30630778 | - | 1.00396057687336 | 1.13699574206438 | 2.38645642073598 | 2.13421729395427 |
| chr12 | 30709780 | 30709788 | - | 4.01584230749346 | 7.95897019445067 | 4.77291284147196 | 3.20132594093141 |
| chr12 | 31104481 | 31104490 | + | 2.00792115374673 | 3.41098722619315 | 2.38645642073598 | 1.06710864697714 |
| chr12 | 31104665 | 31104667 | + | 8.03168461498692 | 4.54798296825753 | 3.57968463110397 | 6.40265188186281 |
| chr12 | 31104714 | 31104765 | + | 154.609928838498 | 106.877599754052 | 127.675418509375 | 104.576647403759 |
| chr12 | 31729225 | 31729287 | + | 24.0950538449607 | 43.2058381984465 | 227.906588180286 | 326.535245975004 |
| chr12 | 31729306 | 31729338 | + | 3.01188173062009 | 7.95897019445067 | 23.8645642073598 | 34.1474767032683 |
| chr12 | 31992840 | 31992845 | + | 8.03168461498692 | 5.68497871032191 | 11.9322821036799 | 32.0132594093141 |
| chr12 | 32743389 | 32743432 | + | 12.0475269224804 | 18.1919318730301 | 20.2848795762558 | 6.40265188186281 |
| chr12 | 32743451 | 32743475 | + | 6.02376346124019 | 3.41098722619315 | 5.96614105183995 | 5.33554323488568 |
| chr12 | 32743524 | 32743545 | + | 1.00396057687336 | 4.54798296825753 | 3.57968463110397 | 3.20132594093141 |
| chr12 | 32743716 | 32743730 | + | 2.00792115374673 | 3.41098722619315 | 3.57968463110397 | 5.33554323488568 |
| chr12 | 32744790 | 32744796 | + | 1.00396057687336 | 1.13699574206438 | 2.38645642073598 | 1.06710864697714 |
| chr12 | 32745282 | 32745283 | + | 7.02772403811355 | 9.09596593651506 | 4.77291284147196 | 3.20132594093141 |
| chr12 | 32747046 | 32747091 | - | 58.2297134586551 | 54.5757956190903 | 20.2848795762558 | 10.6710864697714 |
| chr12 | 34027013 | 34027023 | + | 10.0396057687336 | 11.3699574206438 | 3.57968463110397 | 3.20132594093141 |
| chr12 | 39293255 | 39293263 | - | 11.043566345607 | 13.6439489047726 | 2.38645642073598 | 5.33554323488568 |
| chr12 | 39293452 | 39293525 | - | 20.0792115374673 | 46.6168254246397 | 10.7390538933119 | 26.6777161744284 |
| chr12 | 39293638 | 39293674 | - | 9.03564519186028 | 5.68497871032191 | 5.96614105183995 | 4.26843458790854 |
| chr12 | 39294393 | 39294418 | - | 4.01584230749346 | 6.82197445238629 | 4.77291284147196 | 8.53686917581709 |
| chr12 | 42083843 | 42083866 | - | 4.01584230749346 | 5.68497871032191 | 2.38645642073598 | 3.20132594093141 |
| chr12 | 42084362 | 42084396 | - | 6.02376346124019 | 10.2329616785794 | 8.35259747257593 | 11.7381951167485 |
| chr12 | 42084428 | 42084467 | - | 33.130699036821 | 27.2878978095452 | 14.3187385244159 | 10.6710864697714 |
| chr12 | 42084490 | 42084527 | - | 7.02772403811355 | 11.3699574206438 | 1.19322821036799 | 1.06710864697714 |
| chr12 | 42084877 | 42084917 | - | 5.01980288436682 | 6.82197445238629 | 2.38645642073598 | 3.20132594093141 |
| chr12 | 42085989 | 42085993 | - | 2.00792115374673 | 2.27399148412876 | 1.19322821036799 | 3.20132594093141 |
| chr12 | 42087149 | 42087190 | - | 3.01188173062009 | 10.2329616785794 | 3.57968463110397 | 2.13421729395427 |
| chr12 | 42160143 | 42160165 | - | 3.01188173062009 | 3.41098722619315 | 8.35259747257593 | 6.40265188186281 |
| chr12 | 42160218 | 42160231 | - | 5.01980288436682 | 4.54798296825753 | 7.15936926220794 | 3.20132594093141 |
| chr12 | 42160255 | 42160266 | - | 8.03168461498692 | 11.3699574206438 | 11.9322821036799 | 4.26843458790854 |
| chr12 | 42160288 | 42160360 | - | 29.1148567293276 | 56.8497871032191 | 33.4103898903037 | 18.1408469986113 |
| chr12 | 42235781 | 42235789 | - | 6.02376346124019 | 18.1919318730301 | 3.57968463110397 | 6.40265188186281 |
| chr12 | 42312436 | 42312438 | - | 2.00792115374673 | 2.27399148412876 | 2.38645642073598 | 2.13421729395427 |
| chr12 | 42312808 | 42312831 | - | 2.00792115374673 | 3.41098722619315 | 1.19322821036799 | 3.20132594093141 |
| chr12 | 42312863 | 42312872 | - | 3.01188173062009 | 6.82197445238629 | 4.77291284147196 | 2.13421729395427 |
| chr12 | 42313012 | 42313013 | - | 3.01188173062009 | 3.41098722619315 | 2.38645642073598 | 2.13421729395427 |
| chr12 | 42441842 | 42441893 | + | 5.01980288436682 | 3.41098722619315 | 8.35259747257593 | 5.33554323488568 |
| chr12 | 42446698 | 42446704 | + | 1.00396057687336 | 1.13699574206438 | 3.57968463110397 | 1.06710864697714 |
| chr12 | 42447158 | 42447160 | + | 3.01188173062009 | 3.41098722619315 | 1.19322821036799 | 1.06710864697714 |
| chr12 | 42447189 | 42447217 | + | 11.043566345607 | 5.68497871032191 | 7.15936926220794 | 3.20132594093141 |
| chr12 | 42448536 | 42448558 | + | 1.00396057687336 | 1.13699574206438 | 2.38645642073598 | 4.26843458790854 |
| chr12 | 43721059 | 43721061 | - | 5.01980288436682 | 1.13699574206438 | 1.19322821036799 | 3.20132594093141 |
| chr12 | 43724068 | 43724087 | - | 5.01980288436682 | 1.13699574206438 | 1.19322821036799 | 8.53686917581709 |
| chr12 | 43728683 | 43728695 | - | 8.03168461498692 | 11.3699574206438 | 3.57968463110397 | 4.26843458790854 |
| chr12 | 43728905 | 43728908 | - | 1.00396057687336 | 2.27399148412876 | 1.19322821036799 | 3.20132594093141 |
| chr12 | 43729441 | 43729456 | - | 6.02376346124019 | 6.82197445238629 | 3.57968463110397 | 2.13421729395427 |
| chr12 | 45429503 | 45429558 | + | 5.01980288436682 | 4.54798296825753 | 5.96614105183995 | 3.20132594093141 |
| chr12 | 45429639 | 45429680 | + | 2.00792115374673 | 3.41098722619315 | 3.57968463110397 | 2.13421729395427 |
| chr12 | 45429709 | 45429749 | + | 3.01188173062009 | 3.41098722619315 | 1.19322821036799 | 1.06710864697714 |
| chr12 | 45429925 | 45429982 | + | 1.00396057687336 | 4.54798296825753 | 11.9322821036799 | 1.06710864697714 |
| chr12 | 45430366 | 45430386 | + | 1.00396057687336 | 1.13699574206438 | 3.57968463110397 | 2.13421729395427 |
| chr12 | 45430796 | 45430888 | + | 3.01188173062009 | 1.13699574206438 | 11.9322821036799 | 4.26843458790854 |
| chr12 | 45431302 | 45431308 | + | 2.00792115374673 | 2.27399148412876 | 1.19322821036799 | 1.06710864697714 |
| chr12 | 45432082 | 45432086 | + | 2.00792115374673 | 6.82197445238629 | 1.19322821036799 | 3.20132594093141 |
| chr12 | 45432145 | 45432177 | + | 14.0554480762271 | 25.0139063254164 | 8.35259747257593 | 11.7381951167485 |
| chr12 | 45904988 | 45905005 | + | 4.01584230749346 | 1.13699574206438 | 3.57968463110397 | 1.06710864697714 |
| chr12 | 45905095 | 45905109 | + | 23.0910932680874 | 12.5069531627082 | 52.5020412561915 | 26.6777161744284 |
| chr12 | 45905200 | 45905214 | + | 1.00396057687336 | 1.13699574206438 | 5.96614105183995 | 3.20132594093141 |
| chr12 | 45905559 | 45905586 | + | 8.03168461498692 | 4.54798296825753 | 14.3187385244159 | 14.9395210576799 |
| chr12 | 45905875 | 45905887 | + | 1.00396057687336 | 1.13699574206438 | 7.15936926220794 | 19.2079556455884 |
| chr12 | 45905911 | 45905919 | + | 2.00792115374673 | 4.54798296825753 | 4.77291284147196 | 7.46976052883995 |
| chr12 | 45905995 | 45906010 | + | 12.0475269224804 | 13.6439489047726 | 5.96614105183995 | 8.53686917581709 |
| chr12 | 45906101 | 45906122 | + | 4.01584230749346 | 2.27399148412876 | 3.57968463110397 | 9.60397782279422 |
| chr12 | 45906202 | 45906209 | + | 10.0396057687336 | 10.2329616785794 | 5.96614105183995 | 3.20132594093141 |
| chr12 | 45906300 | 45906314 | + | 5.01980288436682 | 6.82197445238629 | 3.57968463110397 | 5.33554323488568 |
| chr12 | 45906342 | 45906353 | + | 7.02772403811355 | 2.27399148412876 | 4.77291284147196 | 4.26843458790854 |
| chr12 | 45906400 | 45906408 | + | 16.0633692299738 | 10.2329616785794 | 7.15936926220794 | 8.53686917581709 |
| chr12 | 45919164 | 45919173 | - | 7.02772403811355 | 9.09596593651506 | 2.38645642073598 | 2.13421729395427 |
| chr12 | 45919207 | 45919220 | - | 4.01584230749346 | 9.09596593651506 | 3.57968463110397 | 8.53686917581709 |
| chr12 | 45919240 | 45919266 | - | 17.0673298068472 | 9.09596593651506 | 8.35259747257593 | 10.6710864697714 |
| chr12 | 45919280 | 45919299 | - | 4.01584230749346 | 5.68497871032191 | 1.19322821036799 | 3.20132594093141 |
| chr12 | 45919401 | 45919534 | - | 43.1703048055547 | 67.0827487817985 | 69.2072362013434 | 37.3488026441998 |
| chr12 | 45919610 | 45919644 | - | 2.00792115374673 | 11.3699574206438 | 2.38645642073598 | 1.06710864697714 |
| chr12 | 45919691 | 45919748 | - | 60.2376346124019 | 36.3838637460602 | 34.6036181006717 | 39.483019938154 |
| chr12 | 45919790 | 45919822 | - | 7.02772403811355 | 11.3699574206438 | 4.77291284147196 | 11.7381951167485 |
| chr12 | 45919888 | 45919896 | - | 7.02772403811355 | 2.27399148412876 | 2.38645642073598 | 1.06710864697714 |
| chr12 | 45920088 | 45920094 | - | 2.00792115374673 | 4.54798296825753 | 2.38645642073598 | 1.06710864697714 |
| chr12 | 45920150 | 45920152 | - | 1.00396057687336 | 1.13699574206438 | 4.77291284147196 | 1.06710864697714 |
| chr12 | 45920232 | 45920253 | - | 3.01188173062009 | 2.27399148412876 | 2.38645642073598 | 4.26843458790854 |
| chr12 | 45920627 | 45920679 | - | 20.0792115374673 | 19.3289276150945 | 28.6374770488317 | 53.3554323488568 |
| chr12 | 45920733 | 45920759 | - | 3.01188173062009 | 3.41098722619315 | 5.96614105183995 | 6.40265188186281 |
| chr12 | 45921067 | 45921071 | - | 1.00396057687336 | 2.27399148412876 | 2.38645642073598 | 2.13421729395427 |
| chr12 | 45921733 | 45921742 | - | 1.00396057687336 | 2.27399148412876 | 4.77291284147196 | 1.06710864697714 |
| chr12 | 45921809 | 45921829 | - | 1.00396057687336 | 3.41098722619315 | 2.38645642073598 | 2.13421729395427 |
| chr12 | 45922027 | 45922053 | - | 4.01584230749346 | 5.68497871032191 | 2.38645642073598 | 4.26843458790854 |
| chr12 | 45954933 | 45954938 | - | 16.0633692299738 | 11.3699574206438 | 14.3187385244159 | 14.9395210576799 |
| chr12 | 45956027 | 45956029 | - | 2.00792115374673 | 1.13699574206438 | 8.35259747257593 | 1.06710864697714 |
| chr12 | 46183217 | 46183234 | - | 4.01584230749346 | 14.780944646837 | 1.19322821036799 | 1.06710864697714 |
| chr12 | 46183318 | 46183353 | - | 11.043566345607 | 18.1919318730301 | 3.57968463110397 | 4.26843458790854 |
| chr12 | 46183462 | 46183520 | - | 27.1069355755808 | 51.1648083928972 | 2.38645642073598 | 7.46976052883995 |
| chr12 | 46183580 | 46183621 | - | 27.1069355755808 | 57.9867828452835 | 5.96614105183995 | 10.6710864697714 |
| chr12 | 46183856 | 46183857 | - | 2.00792115374673 | 5.68497871032191 | 1.19322821036799 | 1.06710864697714 |
| chr12 | 46184148 | 46184161 | - | 1.00396057687336 | 1.13699574206438 | 3.57968463110397 | 4.26843458790854 |
| chr12 | 46184864 | 46184884 | - | 9.03564519186028 | 11.3699574206438 | 2.38645642073598 | 2.13421729395427 |
| chr12 | 46185666 | 46185709 | - | 35.1386201905678 | 32.9728765198671 | 2.38645642073598 | 3.20132594093141 |
| chr12 | 46185740 | 46185847 | - | 82.3247673036159 | 57.9867828452835 | 9.54582568294392 | 13.8724124107028 |
| chr12 | 46185878 | 46185905 | - | 33.130699036821 | 37.5208594881246 | 7.15936926220794 | 2.13421729395427 |
| chr12 | 46185958 | 46185986 | - | 21.0831721143407 | 26.1509020674808 | 5.96614105183995 | 30.9461507623369 |
| chr12 | 46186013 | 46186103 | - | 21.0831721143407 | 20.4659233571589 | 1.19322821036799 | 4.26843458790854 |
| chr12 | 46186211 | 46186243 | - | 30.1188173062009 | 28.4248935516096 | 2.38645642073598 | 4.26843458790854 |
| chr12 | 46186354 | 46186550 | - | 46.1821865361748 | 59.1237785873479 | 5.96614105183995 | 12.8053037637256 |
| chr12 | 46186901 | 46186911 | - | 22.087132691214 | 25.0139063254164 | 3.57968463110397 | 1.06710864697714 |
| chr12 | 46186934 | 46186964 | - | 23.0910932680874 | 22.7399148412876 | 1.19322821036799 | 2.13421729395427 |
| chr12 | 46188055 | 46188066 | - | 76.3010038423757 | 115.973565690567 | 2.38645642073598 | 4.26843458790854 |
| chr12 | 46188352 | 46188366 | - | 1.00396057687336 | 5.68497871032191 | 2.38645642073598 | 1.06710864697714 |
| chr12 | 46188459 | 46188513 | - | 18.0712903837206 | 22.7399148412876 | 2.38645642073598 | 2.13421729395427 |
| chr12 | 46188534 | 46188554 | - | 11.043566345607 | 6.82197445238629 | 1.19322821036799 | 3.20132594093141 |
| chr12 | 46188659 | 46188727 | - | 14.0554480762271 | 14.780944646837 | 3.57968463110397 | 3.20132594093141 |
| chr12 | 46188765 | 46188781 | - | 3.01188173062009 | 9.09596593651506 | 1.19322821036799 | 2.13421729395427 |
| chr12 | 46188959 | 46189019 | - | 15.0594086531005 | 30.6988850357383 | 3.57968463110397 | 1.06710864697714 |
| chr12 | 46358213 | 46358447 | - | 54.2138711511617 | 134.165497563597 | 21.4781077866238 | 21.3421729395427 |
| chr12 | 46358472 | 46359023 | - | 177.701022106586 | 374.071599139182 | 32.2171616799357 | 27.7448248214055 |
| chr12 | 46359044 | 46359814 | - | 106.419821148577 | 223.988161186683 | 59.6614105183995 | 182.47557863309 |
| chr12 | 46360130 | 46360402 | - | 29.1148567293276 | 44.3428339405109 | 5.96614105183995 | 9.60397782279422 |
| chr12 | 46360426 | 46360471 | - | 2.00792115374673 | 11.3699574206438 | 2.38645642073598 | 1.06710864697714 |
| chr12 | 46360497 | 46360612 | - | 7.02772403811355 | 12.5069531627082 | 3.57968463110397 | 3.20132594093141 |
| chr12 | 46360698 | 46360947 | - | 75.2970432655023 | 184.19331021443 | 26.2510206280958 | 17.0737383516342 |
| chr12 | 46360968 | 46361247 | - | 337.33075382945 | 565.086883805998 | 76.3666054635513 | 102.442430109805 |
| chr12 | 46383647 | 46383708 | + | 38.1505019211879 | 112.562578464374 | 213.58784965587 | 216.623055336359 |
| chr12 | 47663330 | 47663370 | - | 6.02376346124019 | 6.82197445238629 | 3.57968463110397 | 8.53686917581709 |
| chr12 | 47663392 | 47663409 | - | 5.01980288436682 | 1.13699574206438 | 2.38645642073598 | 4.26843458790854 |
| chr12 | 47706081 | 47706135 | + | 13.0514874993537 | 22.7399148412876 | 15.5119667347839 | 21.3421729395427 |
| chr12 | 47780287 | 47780312 | + | 6.02376346124019 | 4.54798296825753 | 2.38645642073598 | 3.20132594093141 |
| chr12 | 47781263 | 47781271 | + | 4.01584230749346 | 4.54798296825753 | 1.19322821036799 | 2.13421729395427 |
| chr12 | 47781526 | 47781543 | + | 6.02376346124019 | 15.9179403889013 | 4.77291284147196 | 17.0737383516342 |
| chr12 | 47781573 | 47781574 | + | 2.00792115374673 | 4.54798296825753 | 1.19322821036799 | 7.46976052883995 |
| chr12 | 47782703 | 47782704 | + | 4.01584230749346 | 2.27399148412876 | 3.57968463110397 | 3.20132594093141 |
| chr12 | 47782728 | 47782747 | + | 5.01980288436682 | 7.95897019445067 | 7.15936926220794 | 5.33554323488568 |
| chr12 | 47782814 | 47782868 | - | 13.0514874993537 | 10.2329616785794 | 9.54582568294392 | 18.1408469986113 |
| chr12 | 47782898 | 47782930 | - | 22.087132691214 | 25.0139063254164 | 14.3187385244159 | 36.2816939972226 |
| chr12 | 47782952 | 47782994 | - | 7.02772403811355 | 12.5069531627082 | 11.9322821036799 | 29.8790421153598 |
| chr12 | 47841744 | 47841756 | - | 8.03168461498692 | 10.2329616785794 | 2.38645642073598 | 4.26843458790854 |
| chr12 | 47841863 | 47841871 | - | 14.0554480762271 | 21.6029190992233 | 3.57968463110397 | 16.006629704657 |
| chr12 | 47841955 | 47841995 | - | 24.0950538449607 | 55.7127913611547 | 5.96614105183995 | 14.9395210576799 |
| chr12 | 47843010 | 47843012 | - | 9.03564519186028 | 7.95897019445067 | 1.19322821036799 | 3.20132594093141 |
| chr12 | 47968326 | 47968334 | + | 17.0673298068472 | 39.7948509722534 | 4.77291284147196 | 2.13421729395427 |
| chr12 | 47968791 | 47968860 | + | 220.87132691214 | 294.481897194675 | 21.4781077866238 | 18.1408469986113 |
| chr12 | 48043693 | 48043705 | - | 6.02376346124019 | 11.3699574206438 | 13.1255103140479 | 6.40265188186281 |
| chr12 | 48043975 | 48043980 | - | 3.01188173062009 | 3.41098722619315 | 2.38645642073598 | 4.26843458790854 |
| chr12 | 48044119 | 48044125 | - | 2.00792115374673 | 5.68497871032191 | 1.19322821036799 | 1.06710864697714 |
| chr12 | 48044156 | 48044158 | - | 2.00792115374673 | 4.54798296825753 | 1.19322821036799 | 3.20132594093141 |
| chr12 | 48045081 | 48045114 | - | 4.01584230749346 | 5.68497871032191 | 4.77291284147196 | 14.9395210576799 |
| chr12 | 48045285 | 48045292 | - | 7.02772403811355 | 4.54798296825753 | 7.15936926220794 | 6.40265188186281 |
| chr12 | 48145861 | 48145877 | + | 8.03168461498692 | 5.68497871032191 | 2.38645642073598 | 4.26843458790854 |
| chr12 | 48145939 | 48145964 | + | 9.03564519186028 | 11.3699574206438 | 1.19322821036799 | 4.26843458790854 |
| chr12 | 48146028 | 48146037 | + | 16.0633692299738 | 18.1919318730301 | 3.57968463110397 | 5.33554323488568 |
| chr12 | 48148077 | 48148100 | - | 3.01188173062009 | 5.68497871032191 | 1.19322821036799 | 4.26843458790854 |
| chr12 | 48341962 | 48341992 | - | 13.0514874993537 | 39.7948509722534 | 3.57968463110397 | 1.06710864697714 |
| chr12 | 48342319 | 48342343 | - | 15.0594086531005 | 14.780944646837 | 3.57968463110397 | 1.06710864697714 |
| chr12 | 48653533 | 48653557 | - | 21.0831721143407 | 25.0139063254164 | 11.9322821036799 | 10.6710864697714 |
| chr12 | 48653595 | 48653621 | - | 3.01188173062009 | 3.41098722619315 | 4.77291284147196 | 2.13421729395427 |
| chr12 | 48688622 | 48688685 | - | 84.3326884573626 | 95.5076423334081 | 70.4004644117114 | 74.6976052883995 |
| chr12 | 48688716 | 48688788 | - | 24.0950538449607 | 25.0139063254164 | 25.0577924177278 | 20.2750642925656 |
| chr12 | 48688810 | 48688827 | - | 9.03564519186028 | 10.2329616785794 | 4.77291284147196 | 9.60397782279422 |
| chr12 | 48688926 | 48688974 | - | 15.0594086531005 | 26.1509020674808 | 17.8984231555198 | 12.8053037637256 |
| chr12 | 48688996 | 48689144 | - | 39.1544624980612 | 57.9867828452835 | 59.6614105183995 | 32.0132594093141 |
| chr12 | 48689211 | 48689241 | - | 3.01188173062009 | 3.41098722619315 | 4.77291284147196 | 8.53686917581709 |
| chr12 | 48689282 | 48689366 | - | 27.1069355755808 | 35.2468680039958 | 21.4781077866238 | 16.006629704657 |
| chr12 | 48689540 | 48689544 | - | 2.00792115374673 | 2.27399148412876 | 1.19322821036799 | 3.20132594093141 |
| chr12 | 48689686 | 48689756 | - | 23.0910932680874 | 17.0549361309657 | 21.4781077866238 | 12.8053037637256 |
| chr12 | 48690012 | 48690035 | - | 8.03168461498692 | 9.09596593651506 | 13.1255103140479 | 28.8119334683827 |
| chr12 | 48690063 | 48690101 | - | 4.01584230749346 | 5.68497871032191 | 13.1255103140479 | 4.26843458790854 |
| chr12 | 48690134 | 48690209 | - | 6.02376346124019 | 11.3699574206438 | 5.96614105183995 | 6.40265188186281 |
| chr12 | 48690255 | 48690275 | - | 5.01980288436682 | 9.09596593651506 | 7.15936926220794 | 5.33554323488568 |
| chr12 | 48690361 | 48690370 | - | 3.01188173062009 | 5.68497871032191 | 3.57968463110397 | 6.40265188186281 |
| chr12 | 48690409 | 48690418 | - | 9.03564519186028 | 7.95897019445067 | 5.96614105183995 | 10.6710864697714 |
| chr12 | 48690664 | 48690684 | - | 1.00396057687336 | 1.13699574206438 | 5.96614105183995 | 1.06710864697714 |
| chr12 | 48690957 | 48691002 | - | 6.02376346124019 | 5.68497871032191 | 9.54582568294392 | 5.33554323488568 |
| chr12 | 48691187 | 48691210 | - | 6.02376346124019 | 5.68497871032191 | 2.38645642073598 | 4.26843458790854 |
| chr12 | 48691352 | 48691377 | - | 3.01188173062009 | 3.41098722619315 | 9.54582568294392 | 7.46976052883995 |
| chr12 | 48691569 | 48691609 | - | 6.02376346124019 | 6.82197445238629 | 11.9322821036799 | 6.40265188186281 |
| chr12 | 48691807 | 48691813 | - | 7.02772403811355 | 9.09596593651506 | 5.96614105183995 | 8.53686917581709 |
| chr12 | 48691958 | 48691967 | - | 2.00792115374673 | 2.27399148412876 | 2.38645642073598 | 1.06710864697714 |
| chr12 | 48692368 | 48692390 | - | 4.01584230749346 | 5.68497871032191 | 4.77291284147196 | 3.20132594093141 |
| chr12 | 48692457 | 48692497 | - | 95.3762548029696 | 104.603608269923 | 53.6952694665595 | 77.8989312293309 |
| chr12 | 48693466 | 48693470 | - | 12.0475269224804 | 5.68497871032191 | 4.77291284147196 | 1.06710864697714 |
| chr12 | 48693868 | 48693887 | - | 3.01188173062009 | 4.54798296825753 | 1.19322821036799 | 1.06710864697714 |
| chr12 | 48829784 | 48829808 | - | 20.0792115374673 | 20.4659233571589 | 8.35259747257593 | 7.46976052883995 |
| chr12 | 48829854 | 48829875 | - | 19.0752509605939 | 5.68497871032191 | 8.35259747257593 | 14.9395210576799 |
| chr12 | 48922051 | 48922129 | - | 33.130699036821 | 26.1509020674808 | 1.19322821036799 | 3.20132594093141 |
| chr12 | 48936231 | 48936271 | - | 37.1465413443145 | 30.6988850357383 | 20.2848795762558 | 19.2079556455884 |
| chr12 | 48936335 | 48936383 | - | 15.0594086531005 | 27.2878978095452 | 7.15936926220794 | 4.26843458790854 |
| chr12 | 48936519 | 48936527 | - | 20.0792115374673 | 11.3699574206438 | 3.57968463110397 | 10.6710864697714 |
| chr12 | 48936754 | 48936773 | - | 2.00792115374673 | 1.13699574206438 | 1.19322821036799 | 1.06710864697714 |
| chr12 | 48936914 | 48936958 | - | 4.01584230749346 | 7.95897019445067 | 14.3187385244159 | 8.53686917581709 |
| chr12 | 48936980 | 48937170 | - | 833.287278804893 | 480.949198893234 | 1288.68646719743 | 1210.10120567207 |
| chr12 | 48937194 | 48937314 | - | 33.130699036821 | 21.6029190992233 | 60.8546387287675 | 26.6777161744284 |
| chr12 | 48937356 | 48937382 | - | 5.01980288436682 | 7.95897019445067 | 4.77291284147196 | 4.26843458790854 |
| chr12 | 48937511 | 48937540 | - | 9.03564519186028 | 6.82197445238629 | 2.38645642073598 | 4.26843458790854 |
| chr12 | 48937561 | 48937575 | - | 7.02772403811355 | 11.3699574206438 | 3.57968463110397 | 2.13421729395427 |
| chr12 | 48937774 | 48937785 | - | 3.01188173062009 | 4.54798296825753 | 2.38645642073598 | 1.06710864697714 |
| chr12 | 48937997 | 48938019 | - | 10.0396057687336 | 3.41098722619315 | 10.7390538933119 | 1.06710864697714 |
| chr12 | 48938739 | 48938749 | - | 4.01584230749346 | 4.54798296825753 | 2.38645642073598 | 3.20132594093141 |
| chr12 | 49002306 | 49002371 | - | 62.2455557661486 | 69.3567402659273 | 23.8645642073598 | 37.3488026441998 |
| chr12 | 49002401 | 49002422 | - | 45.1782259593014 | 54.5757956190903 | 9.54582568294392 | 20.2750642925656 |
| chr12 | 49002464 | 49002469 | - | 8.03168461498692 | 19.3289276150945 | 1.19322821036799 | 10.6710864697714 |
| chr12 | 49002886 | 49002894 | - | 6.02376346124019 | 3.41098722619315 | 11.9322821036799 | 4.26843458790854 |
| chr12 | 49019044 | 49019055 | - | 6.02376346124019 | 11.3699574206438 | 7.15936926220794 | 8.53686917581709 |
| chr12 | 49019554 | 49019555 | - | 2.00792115374673 | 1.13699574206438 | 2.38645642073598 | 2.13421729395427 |
| chr12 | 49019814 | 49019837 | - | 15.0594086531005 | 17.0549361309657 | 41.7629873628796 | 32.0132594093141 |
| chr12 | 49019868 | 49019900 | - | 17.0673298068472 | 7.95897019445067 | 25.0577924177278 | 9.60397782279422 |
| chr12 | 49019979 | 49019998 | - | 6.02376346124019 | 2.27399148412876 | 5.96614105183995 | 4.26843458790854 |
| chr12 | 49020069 | 49020112 | - | 28.1108961524542 | 11.3699574206438 | 45.3426719939836 | 13.8724124107028 |
| chr12 | 49020160 | 49020182 | - | 5.01980288436682 | 7.95897019445067 | 9.54582568294392 | 13.8724124107028 |
| chr12 | 49020243 | 49020268 | - | 7.02772403811355 | 3.41098722619315 | 5.96614105183995 | 2.13421729395427 |
| chr12 | 49020472 | 49020480 | - | 4.01584230749346 | 12.5069531627082 | 9.54582568294392 | 6.40265188186281 |
| chr12 | 49020510 | 49020518 | - | 8.03168461498692 | 4.54798296825753 | 7.15936926220794 | 1.06710864697714 |
| chr12 | 49020544 | 49020584 | - | 58.2297134586551 | 85.2746806548286 | 32.2171616799357 | 22.4092815865198 |
| chr12 | 49020605 | 49020627 | - | 15.0594086531005 | 12.5069531627082 | 4.77291284147196 | 7.46976052883995 |
| chr12 | 49020713 | 49020740 | - | 38.1505019211879 | 31.8358807778027 | 3.57968463110397 | 3.20132594093141 |
| chr12 | 49064720 | 49064724 | - | 3.01188173062009 | 10.2329616785794 | 2.38645642073598 | 12.8053037637256 |
| chr12 | 49097159 | 49097208 | - | 6.02376346124019 | 4.54798296825753 | 10.7390538933119 | 10.6710864697714 |
| chr12 | 49097233 | 49097235 | - | 1.00396057687336 | 1.13699574206438 | 1.19322821036799 | 1.06710864697714 |
| chr12 | 49097440 | 49097466 | - | 3.01188173062009 | 1.13699574206438 | 9.54582568294392 | 10.6710864697714 |
| chr12 | 49097526 | 49097612 | - | 3.01188173062009 | 5.68497871032191 | 41.7629873628796 | 16.006629704657 |
| chr12 | 49097658 | 49097667 | - | 1.00396057687336 | 1.13699574206438 | 13.1255103140479 | 5.33554323488568 |
| chr12 | 49127801 | 49127982 | - | 674.661507658901 | 698.11538562753 | 1011.85752239206 | 852.619808934731 |
| chr12 | 49127814 | 49127952 | + | 138.546559608524 | 133.028501821533 | 215.974306076606 | 325.468137328026 |
| chr12 | 49184827 | 49184829 | - | 1.00396057687336 | 1.13699574206438 | 5.96614105183995 | 1.06710864697714 |
| chr12 | 49273227 | 49273303 | + | 209.827760566533 | 155.76841666282 | 368.707517003709 | 361.749831325249 |
| chr12 | 49324035 | 49324038 | + | 4.01584230749346 | 3.41098722619315 | 2.38645642073598 | 3.20132594093141 |
| chr12 | 49324298 | 49324328 | + | 78.3089249961224 | 108.014595496116 | 13.1255103140479 | 28.8119334683827 |
| chr12 | 49331686 | 49331711 | + | 24.0950538449607 | 11.3699574206438 | 9.54582568294392 | 13.8724124107028 |
| chr12 | 49352776 | 49352822 | + | 3.01188173062009 | 3.41098722619315 | 1.19322821036799 | 1.06710864697714 |
| chr12 | 49526430 | 49526432 | + | 2.00792115374673 | 1.13699574206438 | 2.38645642073598 | 2.13421729395427 |
| chr12 | 49558353 | 49558398 | - | 12.0475269224804 | 17.0549361309657 | 4.77291284147196 | 10.6710864697714 |
| chr12 | 49558435 | 49558470 | - | 4.01584230749346 | 5.68497871032191 | 1.19322821036799 | 5.33554323488568 |
| chr12 | 49762871 | 49763037 | + | 101.40001826421 | 111.425582722309 | 36.9900745214077 | 60.8251928776967 |
| chr12 | 49763056 | 49763228 | + | 41.1623836518079 | 29.5618892936739 | 21.4781077866238 | 11.7381951167485 |
| chr12 | 49763243 | 49763504 | + | 90.3564519186028 | 50.0278126508328 | 52.5020412561915 | 34.1474767032683 |
| chr12 | 49763534 | 49763883 | + | 334.31887209883 | 275.15296957958 | 316.205475747517 | 290.253551977781 |
| chr12 | 49763907 | 49764558 | + | 1211.78041628615 | 1096.06389535006 | 763.666054635513 | 706.425924298864 |
| chr12 | 49764584 | 49764615 | + | 28.1108961524542 | 17.0549361309657 | 15.5119667347839 | 17.0737383516342 |
| chr12 | 49764629 | 49764705 | + | 85.336649034236 | 76.1787147183136 | 29.8307052591997 | 29.8790421153598 |
| chr12 | 49764821 | 49764899 | + | 29.1148567293276 | 43.2058381984465 | 19.0916513658878 | 12.8053037637256 |
| chr12 | 49791236 | 49791245 | - | 3.01188173062009 | 3.41098722619315 | 1.19322821036799 | 1.06710864697714 |
| chr12 | 49837943 | 49837947 | - | 14.0554480762271 | 45.4798296825753 | 9.54582568294392 | 13.8724124107028 |
| chr12 | 49955761 | 49955770 | + | 6.02376346124019 | 7.95897019445067 | 8.35259747257593 | 2.13421729395427 |
| chr12 | 49989410 | 49989416 | - | 12.0475269224804 | 11.3699574206438 | 2.38645642073598 | 1.06710864697714 |
| chr12 | 49989558 | 49989570 | - | 14.0554480762271 | 20.4659233571589 | 2.38645642073598 | 2.13421729395427 |
| chr12 | 49990021 | 49990025 | - | 4.01584230749346 | 2.27399148412876 | 1.19322821036799 | 5.33554323488568 |
| chr12 | 50099130 | 50099140 | + | 1.00396057687336 | 1.13699574206438 | 4.77291284147196 | 1.06710864697714 |
| chr12 | 50099484 | 50099486 | + | 2.00792115374673 | 1.13699574206438 | 1.19322821036799 | 1.06710864697714 |
| chr12 | 50099619 | 50099641 | + | 1.00396057687336 | 6.82197445238629 | 7.15936926220794 | 2.13421729395427 |
| chr12 | 50099728 | 50099736 | + | 9.03564519186028 | 3.41098722619315 | 13.1255103140479 | 5.33554323488568 |
| chr12 | 50100490 | 50100528 | + | 6.02376346124019 | 2.27399148412876 | 2.38645642073598 | 1.06710864697714 |
| chr12 | 50100575 | 50100618 | + | 4.01584230749346 | 10.2329616785794 | 1.19322821036799 | 2.13421729395427 |
| chr12 | 50100646 | 50100686 | + | 9.03564519186028 | 6.82197445238629 | 2.38645642073598 | 7.46976052883995 |
| chr12 | 50112141 | 50112279 | + | 859.3902538036 | 568.497871032191 | 242.225326704702 | 442.850088495511 |
| chr12 | 50120248 | 50120268 | + | 6.02376346124019 | 11.3699574206438 | 2.38645642073598 | 4.26843458790854 |
| chr12 | 50120412 | 50120431 | + | 89.3524913417294 | 170.549361309657 | 44.1494437836156 | 76.8318225823538 |
| chr12 | 50129842 | 50129891 | - | 4.01584230749346 | 15.9179403889013 | 10.7390538933119 | 16.006629704657 |
| chr12 | 50175931 | 50175942 | - | 68.2693192273888 | 110.288586980245 | 20.2848795762558 | 86.435800405148 |
| chr12 | 50176001 | 50176029 | - | 8.03168461498692 | 19.3289276150945 | 1.19322821036799 | 4.26843458790854 |
| chr12 | 50176062 | 50176064 | - | 6.02376346124019 | 5.68497871032191 | 2.38645642073598 | 6.40265188186281 |
| chr12 | 50176574 | 50176609 | - | 6.02376346124019 | 5.68497871032191 | 5.96614105183995 | 4.26843458790854 |
| chr12 | 50176631 | 50176681 | - | 28.1108961524542 | 28.4248935516096 | 31.0239334695677 | 16.006629704657 |
| chr12 | 50176786 | 50176824 | - | 11.043566345607 | 6.82197445238629 | 1.19322821036799 | 7.46976052883995 |
| chr12 | 50454393 | 50454418 | + | 3.01188173062009 | 4.54798296825753 | 1.19322821036799 | 4.26843458790854 |
| chr12 | 50462579 | 50462582 | + | 4.01584230749346 | 5.68497871032191 | 4.77291284147196 | 4.26843458790854 |
| chr12 | 50462604 | 50462623 | + | 8.03168461498692 | 2.27399148412876 | 8.35259747257593 | 3.20132594093141 |
| chr12 | 50474105 | 50474108 | + | 6.02376346124019 | 6.82197445238629 | 7.15936926220794 | 7.46976052883995 |
| chr12 | 50475729 | 50475769 | + | 3.01188173062009 | 7.95897019445067 | 10.7390538933119 | 10.6710864697714 |
| chr12 | 50475938 | 50475979 | + | 57.2257528817818 | 75.0417189762492 | 47.7291284147196 | 112.046407932599 |
| chr12 | 50476025 | 50476055 | + | 1.00396057687336 | 5.68497871032191 | 4.77291284147196 | 3.20132594093141 |
| chr12 | 50476233 | 50476239 | + | 1.00396057687336 | 3.41098722619315 | 2.38645642073598 | 1.06710864697714 |
| chr12 | 50477295 | 50477298 | + | 2.00792115374673 | 2.27399148412876 | 2.38645642073598 | 1.06710864697714 |
| chr12 | 50477806 | 50477812 | + | 4.01584230749346 | 3.41098722619315 | 4.77291284147196 | 14.9395210576799 |
| chr12 | 50479949 | 50479962 | + | 2.00792115374673 | 9.09596593651506 | 9.54582568294392 | 4.26843458790854 |
| chr12 | 50746679 | 50746736 | + | 7.02772403811355 | 1.13699574206438 | 2.38645642073598 | 4.26843458790854 |
| chr12 | 50748308 | 50748344 | + | 2.00792115374673 | 2.27399148412876 | 1.19322821036799 | 5.33554323488568 |
| chr12 | 50748375 | 50748447 | + | 1.00396057687336 | 5.68497871032191 | 7.15936926220794 | 11.7381951167485 |
| chr12 | 50748511 | 50748517 | + | 1.00396057687336 | 2.27399148412876 | 1.19322821036799 | 1.06710864697714 |
| chr12 | 50748587 | 50748626 | + | 2.00792115374673 | 3.41098722619315 | 1.19322821036799 | 5.33554323488568 |
| chr12 | 50819824 | 50819857 | + | 3.01188173062009 | 3.41098722619315 | 5.96614105183995 | 2.13421729395427 |
| chr12 | 50819914 | 50819977 | + | 32.1267384599477 | 77.315710460378 | 35.7968463110397 | 25.6106075274513 |
| chr12 | 50820006 | 50820014 | + | 7.02772403811355 | 11.3699574206438 | 8.35259747257593 | 6.40265188186281 |
| chr12 | 50820071 | 50820080 | + | 9.03564519186028 | 17.0549361309657 | 8.35259747257593 | 6.40265188186281 |
| chr12 | 50820175 | 50820184 | + | 10.0396057687336 | 5.68497871032191 | 7.15936926220794 | 1.06710864697714 |
| chr12 | 50820218 | 50820224 | + | 1.00396057687336 | 3.41098722619315 | 3.57968463110397 | 6.40265188186281 |
| chr12 | 50820306 | 50820417 | + | 101.40001826421 | 176.234340019979 | 79.9462900946553 | 43.7514545260626 |
| chr12 | 50820437 | 50820479 | + | 17.0673298068472 | 37.5208594881246 | 10.7390538933119 | 9.60397782279422 |
| chr12 | 50820773 | 50820778 | + | 3.01188173062009 | 5.68497871032191 | 1.19322821036799 | 2.13421729395427 |
| chr12 | 50930108 | 50930178 | + | 11.043566345607 | 12.5069531627082 | 2.38645642073598 | 3.20132594093141 |
| chr12 | 50930990 | 50931077 | + | 2.00792115374673 | 2.27399148412876 | 7.15936926220794 | 4.26843458790854 |
| chr12 | 50931168 | 50931211 | + | 6.02376346124019 | 1.13699574206438 | 1.19322821036799 | 1.06710864697714 |
| chr12 | 50979963 | 50979964 | - | 2.00792115374673 | 2.27399148412876 | 1.19322821036799 | 2.13421729395427 |
| chr12 | 50980029 | 50980030 | - | 3.01188173062009 | 2.27399148412876 | 2.38645642073598 | 4.26843458790854 |
| chr12 | 50980148 | 50980149 | - | 2.00792115374673 | 1.13699574206438 | 1.19322821036799 | 1.06710864697714 |
| chr12 | 50986114 | 50986124 | - | 3.01188173062009 | 6.82197445238629 | 3.57968463110397 | 3.20132594093141 |
| chr12 | 50986520 | 50986532 | - | 7.02772403811355 | 11.3699574206438 | 2.38645642073598 | 2.13421729395427 |
| chr12 | 50987130 | 50987174 | - | 4.01584230749346 | 2.27399148412876 | 1.19322821036799 | 1.06710864697714 |
| chr12 | 50987414 | 50987471 | - | 8.03168461498692 | 6.82197445238629 | 5.96614105183995 | 4.26843458790854 |
| chr12 | 50987480 | 50987505 | - | 1.00396057687336 | 11.3699574206438 | 3.57968463110397 | 2.13421729395427 |
| chr12 | 50987703 | 50987726 | - | 9.03564519186028 | 3.41098722619315 | 5.96614105183995 | 2.13421729395427 |
| chr12 | 50988013 | 50988053 | - | 1.00396057687336 | 3.41098722619315 | 2.38645642073598 | 3.20132594093141 |
| chr12 | 51059475 | 51059486 | + | 1.00396057687336 | 1.13699574206438 | 7.15936926220794 | 1.06710864697714 |
| chr12 | 51059539 | 51059572 | + | 8.03168461498692 | 5.68497871032191 | 10.7390538933119 | 4.26843458790854 |
| chr12 | 51059638 | 51059657 | + | 1.00396057687336 | 2.27399148412876 | 13.1255103140479 | 3.20132594093141 |
| chr12 | 51059752 | 51059939 | + | 30.1188173062009 | 53.438799877026 | 79.9462900946553 | 38.4159112911769 |
| chr12 | 51059970 | 51060094 | + | 43.1703048055547 | 75.0417189762492 | 109.776995353855 | 70.429170700491 |
| chr12 | 51060133 | 51060137 | + | 6.02376346124019 | 2.27399148412876 | 4.77291284147196 | 8.53686917581709 |
| chr12 | 51060201 | 51060309 | + | 21.0831721143407 | 27.2878978095452 | 25.0577924177278 | 21.3421729395427 |
| chr12 | 51060332 | 51060402 | + | 6.02376346124019 | 11.3699574206438 | 9.54582568294392 | 3.20132594093141 |
| chr12 | 51094883 | 51094947 | - | 9.03564519186028 | 7.95897019445067 | 1.19322821036799 | 7.46976052883995 |
| chr12 | 51242572 | 51242585 | + | 7.02772403811355 | 4.54798296825753 | 1.19322821036799 | 1.06710864697714 |
| chr12 | 51242695 | 51242706 | + | 15.0594086531005 | 11.3699574206438 | 8.35259747257593 | 2.13421729395427 |
| chr12 | 51242787 | 51242798 | + | 12.0475269224804 | 5.68497871032191 | 11.9322821036799 | 3.20132594093141 |
| chr12 | 51243288 | 51243320 | + | 3.01188173062009 | 3.41098722619315 | 8.35259747257593 | 6.40265188186281 |
| chr12 | 51243362 | 51243367 | + | 2.00792115374673 | 1.13699574206438 | 2.38645642073598 | 3.20132594093141 |
| chr12 | 51243409 | 51243419 | + | 9.03564519186028 | 10.2329616785794 | 1.19322821036799 | 3.20132594093141 |
| chr12 | 51243446 | 51243491 | + | 3.01188173062009 | 7.95897019445067 | 5.96614105183995 | 6.40265188186281 |
| chr12 | 51243498 | 51243521 | + | 6.02376346124019 | 9.09596593651506 | 4.77291284147196 | 6.40265188186281 |
| chr12 | 51243611 | 51243637 | + | 6.02376346124019 | 3.41098722619315 | 3.57968463110397 | 4.26843458790854 |
| chr12 | 51243659 | 51243713 | + | 27.1069355755808 | 27.2878978095452 | 5.96614105183995 | 5.33554323488568 |
| chr12 | 51243754 | 51243800 | + | 31.1227778830743 | 11.3699574206438 | 17.8984231555198 | 10.6710864697714 |
| chr12 | 51245391 | 51245431 | - | 2.00792115374673 | 2.27399148412876 | 11.9322821036799 | 11.7381951167485 |
| chr12 | 51245451 | 51245489 | - | 6.02376346124019 | 5.68497871032191 | 9.54582568294392 | 19.2079556455884 |
| chr12 | 51271095 | 51271118 | + | 2.00792115374673 | 4.54798296825753 | 1.19322821036799 | 9.60397782279422 |
| chr12 | 51271140 | 51271141 | + | 4.01584230749346 | 3.41098722619315 | 2.38645642073598 | 1.06710864697714 |
| chr12 | 51995273 | 51995337 | + | 19.0752509605939 | 20.4659233571589 | 8.35259747257593 | 8.53686917581709 |
| chr12 | 51995415 | 51995430 | + | 1.00396057687336 | 4.54798296825753 | 1.19322821036799 | 2.13421729395427 |
| chr12 | 51995479 | 51995491 | + | 6.02376346124019 | 3.41098722619315 | 1.19322821036799 | 2.13421729395427 |
| chr12 | 51995524 | 51995553 | + | 6.02376346124019 | 6.82197445238629 | 2.38645642073598 | 1.06710864697714 |
| chr12 | 51995582 | 51995585 | + | 6.02376346124019 | 4.54798296825753 | 7.15936926220794 | 3.20132594093141 |
| chr12 | 51996690 | 51996716 | + | 6.02376346124019 | 4.54798296825753 | 4.77291284147196 | 6.40265188186281 |
| chr12 | 51996748 | 51996815 | + | 20.0792115374673 | 18.1919318730301 | 19.0916513658878 | 27.7448248214055 |
| chr12 | 51996873 | 51996902 | + | 89.3524913417294 | 95.5076423334081 | 128.868646719743 | 82.1673658172394 |
| chr12 | 51996928 | 51996954 | + | 1.00396057687336 | 13.6439489047726 | 8.35259747257593 | 8.53686917581709 |
| chr12 | 51997003 | 51997006 | + | 7.02772403811355 | 4.54798296825753 | 5.96614105183995 | 6.40265188186281 |
| chr12 | 52059263 | 52059302 | + | 36.1425807674411 | 32.9728765198671 | 1.19322821036799 | 2.13421729395427 |
| chr12 | 52077445 | 52077465 | + | 22.087132691214 | 61.3977700714766 | 5.96614105183995 | 11.7381951167485 |
| chr12 | 52897366 | 52897390 | - | 4.01584230749346 | 1.13699574206438 | 2.38645642073598 | 4.26843458790854 |
| chr12 | 52947899 | 52947906 | - | 2.00792115374673 | 3.41098722619315 | 4.77291284147196 | 1.06710864697714 |
| chr12 | 52952873 | 52952883 | + | 635.50704516084 | 404.77048417492 | 571.556312766267 | 773.653769058423 |
| chr12 | 53040328 | 53040333 | + | 13.0514874993537 | 14.780944646837 | 26.2510206280958 | 12.8053037637256 |
| chr12 | 53040530 | 53040554 | + | 6.02376346124019 | 2.27399148412876 | 11.9322821036799 | 8.53686917581709 |
| chr12 | 53040688 | 53040712 | + | 12.0475269224804 | 4.54798296825753 | 19.0916513658878 | 30.9461507623369 |
| chr12 | 53040757 | 53040783 | + | 186.736667298446 | 168.275369825529 | 48.9223566250876 | 168.603166222387 |
| chr12 | 53040997 | 53041009 | + | 15.0594086531005 | 20.4659233571589 | 13.1255103140479 | 19.2079556455884 |
| chr12 | 53041261 | 53041307 | + | 12.0475269224804 | 2.27399148412876 | 9.54582568294392 | 6.40265188186281 |
| chr12 | 53041332 | 53041357 | + | 2.00792115374673 | 1.13699574206438 | 9.54582568294392 | 4.26843458790854 |
| chr12 | 53041390 | 53041416 | + | 13.0514874993537 | 17.0549361309657 | 17.8984231555198 | 14.9395210576799 |
| chr12 | 53041477 | 53041500 | + | 13.0514874993537 | 6.82197445238629 | 47.7291284147196 | 303.058855741507 |
| chr12 | 53041852 | 53041873 | + | 9.03564519186028 | 3.41098722619315 | 11.9322821036799 | 17.0737383516342 |
| chr12 | 53064414 | 53064421 | - | 3.01188173062009 | 3.41098722619315 | 2.38645642073598 | 1.06710864697714 |
| chr12 | 53064727 | 53064786 | - | 22.087132691214 | 18.1919318730301 | 5.96614105183995 | 8.53686917581709 |
| chr12 | 53065127 | 53065134 | - | 2.00792115374673 | 3.41098722619315 | 4.77291284147196 | 1.06710864697714 |
| chr12 | 53187931 | 53187950 | + | 2.00792115374673 | 2.27399148412876 | 1.19322821036799 | 1.06710864697714 |
| chr12 | 53188671 | 53188683 | + | 6.02376346124019 | 12.5069531627082 | 5.96614105183995 | 3.20132594093141 |
| chr12 | 53188752 | 53188777 | + | 12.0475269224804 | 28.4248935516096 | 11.9322821036799 | 6.40265188186281 |
| chr12 | 53189085 | 53189095 | + | 9.03564519186028 | 3.41098722619315 | 2.38645642073598 | 7.46976052883995 |
| chr12 | 53189363 | 53189368 | + | 2.00792115374673 | 3.41098722619315 | 4.77291284147196 | 2.13421729395427 |
| chr12 | 53189459 | 53189508 | + | 675.665468235774 | 1312.0930863423 | 115.743136405695 | 80.0331485232852 |
| chr12 | 53189906 | 53189918 | + | 2.00792115374673 | 1.13699574206438 | 4.77291284147196 | 1.06710864697714 |
| chr12 | 53190029 | 53190062 | + | 60.2376346124019 | 118.247557174696 | 21.4781077866238 | 19.2079556455884 |
| chr12 | 53190100 | 53190143 | + | 15.0594086531005 | 5.68497871032191 | 7.15936926220794 | 4.26843458790854 |
| chr12 | 53190165 | 53190175 | + | 4.01584230749346 | 2.27399148412876 | 2.38645642073598 | 2.13421729395427 |
| chr12 | 53190227 | 53190239 | + | 3.01188173062009 | 1.13699574206438 | 1.19322821036799 | 6.40265188186281 |
| chr12 | 53190320 | 53190333 | + | 2.00792115374673 | 6.82197445238629 | 3.57968463110397 | 1.06710864697714 |
| chr12 | 53190362 | 53190397 | + | 6.02376346124019 | 15.9179403889013 | 7.15936926220794 | 9.60397782279422 |
| chr12 | 53190580 | 53190588 | + | 3.01188173062009 | 1.13699574206438 | 2.38645642073598 | 10.6710864697714 |
| chr12 | 53190712 | 53190732 | + | 20.0792115374673 | 48.8908169087684 | 4.77291284147196 | 11.7381951167485 |
| chr12 | 53254315 | 53254349 | + | 11.043566345607 | 5.68497871032191 | 3.57968463110397 | 10.6710864697714 |
| chr12 | 53299316 | 53299424 | + | 110.43566345607 | 119.38455291676 | 77.5598336739193 | 58.6909755837425 |
| chr12 | 53411699 | 53411742 | + | 2.00792115374673 | 18.1919318730301 | 9.54582568294392 | 6.40265188186281 |
| chr12 | 53411916 | 53411927 | + | 3.01188173062009 | 10.2329616785794 | 2.38645642073598 | 2.13421729395427 |
| chr12 | 53412202 | 53412217 | + | 3.01188173062009 | 3.41098722619315 | 4.77291284147196 | 2.13421729395427 |
| chr12 | 53412298 | 53412335 | + | 10.0396057687336 | 20.4659233571589 | 11.9322821036799 | 6.40265188186281 |
| chr12 | 53412879 | 53412918 | + | 12.0475269224804 | 9.09596593651506 | 2.38645642073598 | 5.33554323488568 |
| chr12 | 53412949 | 53412984 | + | 29.1148567293276 | 43.2058381984465 | 14.3187385244159 | 8.53686917581709 |
| chr12 | 53413052 | 53413084 | + | 34.1346596136944 | 34.1098722619315 | 9.54582568294392 | 16.006629704657 |
| chr12 | 53413166 | 53413169 | + | 2.00792115374673 | 4.54798296825753 | 2.38645642073598 | 2.13421729395427 |
| chr12 | 53413199 | 53413216 | + | 12.0475269224804 | 20.4659233571589 | 4.77291284147196 | 2.13421729395427 |
| chr12 | 53413253 | 53413268 | + | 24.0950538449607 | 39.7948509722534 | 20.2848795762558 | 11.7381951167485 |
| chr12 | 53413307 | 53413308 | + | 1.00396057687336 | 2.27399148412876 | 1.19322821036799 | 1.06710864697714 |
| chr12 | 53413333 | 53413338 | + | 3.01188173062009 | 3.41098722619315 | 7.15936926220794 | 2.13421729395427 |
| chr12 | 53413678 | 53413759 | + | 24.0950538449607 | 35.2468680039958 | 27.4442488384638 | 10.6710864697714 |
| chr12 | 53413811 | 53413838 | + | 5.01980288436682 | 7.95897019445067 | 5.96614105183995 | 1.06710864697714 |
| chr12 | 53414076 | 53414081 | + | 5.01980288436682 | 5.68497871032191 | 4.77291284147196 | 5.33554323488568 |
| chr12 | 53414124 | 53414152 | + | 7.02772403811355 | 4.54798296825753 | 1.19322821036799 | 27.7448248214055 |
| chr12 | 53414443 | 53414450 | + | 6.02376346124019 | 1.13699574206438 | 3.57968463110397 | 5.33554323488568 |
| chr12 | 53414491 | 53414554 | + | 86.3406096111093 | 121.658544400889 | 76.3666054635513 | 38.4159112911769 |
| chr12 | 53414664 | 53414666 | + | 2.00792115374673 | 1.13699574206438 | 3.57968463110397 | 2.13421729395427 |
| chr12 | 53414770 | 53414826 | + | 32.1267384599477 | 38.657855230189 | 19.0916513658878 | 18.1408469986113 |
| chr12 | 53414940 | 53415019 | + | 30.1188173062009 | 57.9867828452835 | 52.5020412561915 | 26.6777161744284 |
| chr12 | 53415048 | 53415068 | + | 6.02376346124019 | 2.27399148412876 | 8.35259747257593 | 16.006629704657 |
| chr12 | 53415102 | 53415112 | + | 4.01584230749346 | 1.13699574206438 | 3.57968463110397 | 4.26843458790854 |
| chr12 | 53415176 | 53415215 | + | 10.0396057687336 | 13.6439489047726 | 9.54582568294392 | 2.13421729395427 |
| chr12 | 53415258 | 53415283 | + | 8.03168461498692 | 7.95897019445067 | 2.38645642073598 | 9.60397782279422 |
| chr12 | 53415407 | 53415428 | + | 64.2534769198953 | 70.4937360079917 | 54.8884976769275 | 19.2079556455884 |
| chr12 | 53415478 | 53415500 | + | 3.01188173062009 | 2.27399148412876 | 16.7051949451519 | 6.40265188186281 |
| chr12 | 53415573 | 53415579 | + | 2.00792115374673 | 1.13699574206438 | 2.38645642073598 | 3.20132594093141 |
| chr12 | 53415663 | 53415681 | + | 6.02376346124019 | 11.3699574206438 | 10.7390538933119 | 3.20132594093141 |
| chr12 | 53441580 | 53441748 | + | 1382.45371435462 | 1199.53050787792 | 3613.09502099427 | 2900.40130248386 |
| chr12 | 53443660 | 53443686 | + | 16.0633692299738 | 12.5069531627082 | 97.8447132501751 | 68.2949534065367 |
| chr12 | 53443745 | 53443769 | + | 3.01188173062009 | 1.13699574206438 | 10.7390538933119 | 11.7381951167485 |
| chr12 | 53446204 | 53446215 | + | 6.02376346124019 | 10.2329616785794 | 15.5119667347839 | 13.8724124107028 |
| chr12 | 53446236 | 53446261 | + | 6.02376346124019 | 1.13699574206438 | 8.35259747257593 | 16.006629704657 |
| chr12 | 53446574 | 53446600 | + | 9.03564519186028 | 4.54798296825753 | 11.9322821036799 | 10.6710864697714 |
| chr12 | 53454785 | 53454846 | + | 52.205949997415 | 75.0417189762492 | 44.1494437836156 | 22.4092815865198 |
| chr12 | 53455477 | 53455494 | + | 16.0633692299738 | 17.0549361309657 | 16.7051949451519 | 2.13421729395427 |
| chr12 | 53455882 | 53455895 | + | 14.0554480762271 | 26.1509020674808 | 29.8307052591997 | 3.20132594093141 |
| chr12 | 53455916 | 53455969 | + | 17.0673298068472 | 15.9179403889013 | 7.15936926220794 | 3.20132594093141 |
| chr12 | 53455998 | 53455999 | + | 8.03168461498692 | 5.68497871032191 | 8.35259747257593 | 1.06710864697714 |
| chr12 | 53459268 | 53459319 | + | 20.0792115374673 | 13.6439489047726 | 2.38645642073598 | 8.53686917581709 |
| chr12 | 53459352 | 53459404 | + | 49.1940682667949 | 45.4798296825753 | 22.6713359969918 | 18.1408469986113 |
| chr12 | 53461012 | 53461144 | + | 107.42378172545 | 97.7816338175368 | 84.7192029361272 | 73.6304966414224 |
| chr12 | 53462482 | 53462568 | + | 107.42378172545 | 78.4527062024424 | 60.8546387287675 | 37.3488026441998 |
| chr12 | 53464769 | 53464850 | + | 73.2891221117556 | 110.288586980245 | 82.3327465153913 | 44.8185631730397 |
| chr12 | 53465929 | 53465974 | + | 35.1386201905678 | 35.2468680039958 | 54.8884976769275 | 53.3554323488568 |
| chr12 | 53467221 | 53467223 | + | 4.01584230749346 | 4.54798296825753 | 2.38645642073598 | 1.06710864697714 |
| chr12 | 53467257 | 53467307 | + | 23.0910932680874 | 23.876910583352 | 25.0577924177278 | 26.6777161744284 |
| chr12 | 53467319 | 53467325 | + | 2.00792115374673 | 2.27399148412876 | 1.19322821036799 | 1.06710864697714 |
| chr12 | 53467521 | 53467558 | + | 24.0950538449607 | 23.876910583352 | 14.3187385244159 | 11.7381951167485 |
| chr12 | 53479429 | 53479453 | + | 1234.87150955424 | 1021.02217637382 | 1100.15640995929 | 1645.48153363874 |
| chr12 | 53479501 | 53479526 | + | 7.02772403811355 | 10.2329616785794 | 7.15936926220794 | 16.006629704657 |
| chr12 | 53479568 | 53479661 | + | 28.1108961524542 | 23.876910583352 | 17.8984231555198 | 28.8119334683827 |
| chr12 | 53479682 | 53479848 | + | 64.2534769198953 | 78.4527062024424 | 40.5697591525116 | 50.1541064079254 |
| chr12 | 53479901 | 53479930 | + | 4.01584230749346 | 6.82197445238629 | 1.19322821036799 | 2.13421729395427 |
| chr12 | 53480079 | 53480082 | + | 12.0475269224804 | 4.54798296825753 | 2.38645642073598 | 4.26843458790854 |
| chr12 | 53480107 | 53480121 | + | 18.0712903837206 | 23.876910583352 | 4.77291284147196 | 7.46976052883995 |
| chr12 | 53480735 | 53480739 | + | 1.00396057687336 | 7.95897019445067 | 1.19322821036799 | 3.20132594093141 |
| chr12 | 53481069 | 53481076 | + | 2.00792115374673 | 2.27399148412876 | 1.19322821036799 | 3.20132594093141 |
| chr12 | 53506381 | 53506402 | + | 18.0712903837206 | 39.7948509722534 | 14.3187385244159 | 20.2750642925656 |
| chr12 | 53514179 | 53514201 | - | 4.01584230749346 | 1.13699574206438 | 4.77291284147196 | 1.06710864697714 |
| chr12 | 53515619 | 53515620 | - | 5.01980288436682 | 4.54798296825753 | 2.38645642073598 | 1.06710864697714 |
| chr12 | 53543088 | 53543090 | - | 12.0475269224804 | 9.09596593651506 | 1.19322821036799 | 3.20132594093141 |
| chr12 | 53543224 | 53543226 | - | 4.01584230749346 | 4.54798296825753 | 1.19322821036799 | 4.26843458790854 |
| chr12 | 53665195 | 53665225 | - | 744.938748040036 | 1202.94149510412 | 439.10798141542 | 335.072115150821 |
| chr12 | 53665267 | 53665312 | - | 57.2257528817818 | 51.1648083928972 | 35.7968463110397 | 66.1607361125824 |
| chr12 | 54002938 | 54002939 | + | 3.01188173062009 | 2.27399148412876 | 2.38645642073598 | 1.06710864697714 |
| chr12 | 54017165 | 54017182 | + | 10.0396057687336 | 38.657855230189 | 1.19322821036799 | 4.26843458790854 |
| chr12 | 54030525 | 54030528 | + | 4.01584230749346 | 10.2329616785794 | 1.19322821036799 | 5.33554323488568 |
| chr12 | 54030660 | 54030680 | + | 7.02772403811355 | 14.780944646837 | 2.38645642073598 | 3.20132594093141 |
| chr12 | 54181015 | 54181030 | - | 15.0594086531005 | 10.2329616785794 | 9.54582568294392 | 10.6710864697714 |
| chr12 | 54181515 | 54181558 | - | 2.00792115374673 | 1.13699574206438 | 3.57968463110397 | 6.40265188186281 |
| chr12 | 54231181 | 54231195 | - | 39.1544624980612 | 38.657855230189 | 28.6374770488317 | 64.0265188186281 |
| chr12 | 54233038 | 54233057 | - | 3.01188173062009 | 3.41098722619315 | 3.57968463110397 | 1.06710864697714 |
| chr12 | 54233304 | 54233312 | - | 16.0633692299738 | 2.27399148412876 | 3.57968463110397 | 6.40265188186281 |
| chr12 | 54233609 | 54233652 | - | 9.03564519186028 | 5.68497871032191 | 10.7390538933119 | 4.26843458790854 |
| chr12 | 54233734 | 54233737 | - | 10.0396057687336 | 7.95897019445067 | 8.35259747257593 | 1.06710864697714 |
| chr12 | 54233793 | 54233826 | - | 31.1227778830743 | 25.0139063254164 | 34.6036181006717 | 9.60397782279422 |
| chr12 | 54234393 | 54234435 | - | 4.01584230749346 | 4.54798296825753 | 14.3187385244159 | 77.8989312293309 |
| chr12 | 54234682 | 54234709 | - | 6.02376346124019 | 3.41098722619315 | 4.77291284147196 | 3.20132594093141 |
| chr12 | 54236170 | 54236188 | - | 3.01188173062009 | 1.13699574206438 | 5.96614105183995 | 2.13421729395427 |
| chr12 | 54236805 | 54236806 | - | 1.00396057687336 | 2.27399148412876 | 5.96614105183995 | 1.06710864697714 |
| chr12 | 54237136 | 54237138 | - | 2.00792115374673 | 2.27399148412876 | 2.38645642073598 | 3.20132594093141 |
| chr12 | 54238651 | 54238709 | - | 8.03168461498692 | 5.68497871032191 | 10.7390538933119 | 14.9395210576799 |
| chr12 | 54239381 | 54239400 | - | 2.00792115374673 | 6.82197445238629 | 10.7390538933119 | 17.0737383516342 |
| chr12 | 54239474 | 54239475 | - | 2.00792115374673 | 1.13699574206438 | 3.57968463110397 | 1.06710864697714 |
| chr12 | 54239678 | 54239690 | - | 9.03564519186028 | 9.09596593651506 | 7.15936926220794 | 8.53686917581709 |
| chr12 | 54239957 | 54239965 | - | 13.0514874993537 | 5.68497871032191 | 11.9322821036799 | 16.006629704657 |
| chr12 | 54240138 | 54240147 | - | 4.01584230749346 | 1.13699574206438 | 1.19322821036799 | 4.26843458790854 |
| chr12 | 54240179 | 54240183 | - | 10.0396057687336 | 3.41098722619315 | 9.54582568294392 | 2.13421729395427 |
| chr12 | 54240284 | 54240359 | - | 6.02376346124019 | 4.54798296825753 | 17.8984231555198 | 11.7381951167485 |
| chr12 | 54240431 | 54240441 | - | 10.0396057687336 | 12.5069531627082 | 26.2510206280958 | 10.6710864697714 |
| chr12 | 54240469 | 54240476 | - | 8.03168461498692 | 9.09596593651506 | 8.35259747257593 | 10.6710864697714 |
| chr12 | 54240683 | 54240710 | - | 6.02376346124019 | 6.82197445238629 | 7.15936926220794 | 8.53686917581709 |
| chr12 | 54241742 | 54241753 | - | 5.01980288436682 | 10.2329616785794 | 4.77291284147196 | 11.7381951167485 |
| chr12 | 54284292 | 54284344 | + | 52.205949997415 | 51.1648083928972 | 58.4681823080315 | 36.2816939972226 |
| chr12 | 54284796 | 54284797 | + | 1.00396057687336 | 1.13699574206438 | 1.19322821036799 | 1.06710864697714 |
| chr12 | 54284826 | 54284858 | + | 34.1346596136944 | 39.7948509722534 | 34.6036181006717 | 34.1474767032683 |
| chr12 | 54284882 | 54284905 | + | 14.0554480762271 | 11.3699574206438 | 4.77291284147196 | 5.33554323488568 |
| chr12 | 54285181 | 54285202 | + | 145.574283646638 | 150.083437952498 | 150.346754506367 | 112.046407932599 |
| chr12 | 54285876 | 54285916 | + | 124.491111532297 | 32.9728765198671 | 93.0718004087032 | 87.5029090521251 |
| chr12 | 54285952 | 54285976 | + | 1.00396057687336 | 2.27399148412876 | 1.19322821036799 | 1.06710864697714 |
| chr12 | 54286246 | 54286283 | + | 3.01188173062009 | 1.13699574206438 | 8.35259747257593 | 4.26843458790854 |
| chr12 | 54286568 | 54286628 | + | 29.1148567293276 | 18.1919318730301 | 9.54582568294392 | 9.60397782279422 |
| chr12 | 54286659 | 54286672 | + | 2.00792115374673 | 11.3699574206438 | 2.38645642073598 | 4.26843458790854 |
| chr12 | 54286930 | 54286979 | + | 1.00396057687336 | 1.13699574206438 | 3.57968463110397 | 1.06710864697714 |
| chr12 | 54287021 | 54287048 | + | 3.01188173062009 | 1.13699574206438 | 3.57968463110397 | 2.13421729395427 |
| chr12 | 54350473 | 54350541 | + | 20.0792115374673 | 17.0549361309657 | 23.8645642073598 | 20.2750642925656 |
| chr12 | 54350754 | 54350764 | + | 10.0396057687336 | 6.82197445238629 | 9.54582568294392 | 6.40265188186281 |
| chr12 | 54350831 | 54350847 | + | 2.00792115374673 | 1.13699574206438 | 1.19322821036799 | 1.06710864697714 |
| chr12 | 54351725 | 54351727 | + | 2.00792115374673 | 1.13699574206438 | 3.57968463110397 | 1.06710864697714 |
| chr12 | 54351781 | 54351793 | + | 17.0673298068472 | 18.1919318730301 | 21.4781077866238 | 13.8724124107028 |
| chr12 | 54351815 | 54351827 | + | 40.1584230749346 | 56.8497871032191 | 27.4442488384638 | 16.006629704657 |
| chr12 | 54369268 | 54369287 | - | 1.00396057687336 | 1.13699574206438 | 4.77291284147196 | 8.53686917581709 |
| chr12 | 54369777 | 54369786 | - | 2.00792115374673 | 1.13699574206438 | 2.38645642073598 | 3.20132594093141 |
| chr12 | 54395290 | 54395305 | - | 1.00396057687336 | 3.41098722619315 | 2.38645642073598 | 3.20132594093141 |
| chr12 | 54395437 | 54395438 | - | 2.00792115374673 | 3.41098722619315 | 2.38645642073598 | 3.20132594093141 |
| chr12 | 55719654 | 55719683 | + | 26.1029749987075 | 34.1098722619315 | 7.15936926220794 | 32.0132594093141 |
| chr12 | 55725467 | 55725595 | - | 301.188173062009 | 281.974944031967 | 204.042023972926 | 289.186443330804 |
| chr12 | 55729092 | 55729130 | + | 11.043566345607 | 10.2329616785794 | 3.57968463110397 | 5.33554323488568 |
| chr12 | 55729201 | 55729221 | + | 4.01584230749346 | 4.54798296825753 | 2.38645642073598 | 1.06710864697714 |
| chr12 | 55729788 | 55729789 | + | 1.00396057687336 | 1.13699574206438 | 1.19322821036799 | 2.13421729395427 |
| chr12 | 55730590 | 55730599 | + | 11.043566345607 | 4.54798296825753 | 2.38645642073598 | 1.06710864697714 |
| chr12 | 55750000 | 55750151 | + | 301.188173062009 | 270.604986611323 | 267.28311912243 | 209.153294807519 |
| chr12 | 55750313 | 55750316 | + | 6.02376346124019 | 3.41098722619315 | 3.57968463110397 | 2.13421729395427 |
| chr12 | 55751106 | 55751127 | + | 4.01584230749346 | 3.41098722619315 | 1.19322821036799 | 6.40265188186281 |
| chr12 | 55751169 | 55751178 | + | 1.00396057687336 | 2.27399148412876 | 5.96614105183995 | 1.06710864697714 |
| chr12 | 55752387 | 55752394 | + | 4.01584230749346 | 2.27399148412876 | 1.19322821036799 | 5.33554323488568 |
| chr12 | 55757296 | 55757330 | - | 44.174265382428 | 69.3567402659273 | 76.3666054635513 | 18.1408469986113 |
| chr12 | 55757362 | 55757404 | - | 15.0594086531005 | 29.5618892936739 | 23.8645642073598 | 23.476390233497 |
| chr12 | 55757454 | 55757464 | - | 2.00792115374673 | 3.41098722619315 | 5.96614105183995 | 4.26843458790854 |
| chr12 | 55757489 | 55757538 | - | 16.0633692299738 | 29.5618892936739 | 23.8645642073598 | 20.2750642925656 |
| chr12 | 55820427 | 55820451 | + | 21.0831721143407 | 23.876910583352 | 2.38645642073598 | 7.46976052883995 |
| chr12 | 55901437 | 55901446 | - | 2.00792115374673 | 4.54798296825753 | 1.19322821036799 | 1.06710864697714 |
| chr12 | 55901481 | 55901503 | - | 10.0396057687336 | 10.2329616785794 | 1.19322821036799 | 1.06710864697714 |
| chr12 | 55901540 | 55901549 | - | 3.01188173062009 | 3.41098722619315 | 2.38645642073598 | 2.13421729395427 |
| chr12 | 55901828 | 55902000 | - | 45.1782259593014 | 34.1098722619315 | 58.4681823080315 | 39.483019938154 |
| chr12 | 55902129 | 55902161 | - | 4.01584230749346 | 1.13699574206438 | 9.54582568294392 | 8.53686917581709 |
| chr12 | 55902252 | 55902255 | - | 2.00792115374673 | 2.27399148412876 | 3.57968463110397 | 3.20132594093141 |
| chr12 | 55953873 | 55953879 | + | 7.02772403811355 | 5.68497871032191 | 8.35259747257593 | 9.60397782279422 |
| chr12 | 55971668 | 55971724 | + | 57.2257528817818 | 48.8908169087684 | 34.6036181006717 | 52.2883237018796 |
| chr12 | 55971748 | 55971830 | + | 32.1267384599477 | 43.2058381984465 | 73.9801490428153 | 45.8856718200168 |
| chr12 | 55971868 | 55971889 | + | 41.1623836518079 | 46.6168254246397 | 62.0478669391355 | 11.7381951167485 |
| chr12 | 55971922 | 55971934 | + | 3.01188173062009 | 9.09596593651506 | 2.38645642073598 | 5.33554323488568 |
| chr12 | 55972472 | 55972509 | + | 7.02772403811355 | 15.9179403889013 | 5.96614105183995 | 18.1408469986113 |
| chr12 | 55972539 | 55972576 | + | 3.01188173062009 | 11.3699574206438 | 9.54582568294392 | 27.7448248214055 |
| chr12 | 55972612 | 55972618 | + | 15.0594086531005 | 12.5069531627082 | 3.57968463110397 | 5.33554323488568 |
| chr12 | 55972641 | 55972659 | + | 27.1069355755808 | 32.9728765198671 | 5.96614105183995 | 1.06710864697714 |
| chr12 | 55972707 | 55972718 | + | 14.0554480762271 | 19.3289276150945 | 8.35259747257593 | 5.33554323488568 |
| chr12 | 55992134 | 55992160 | + | 7.02772403811355 | 4.54798296825753 | 5.96614105183995 | 5.33554323488568 |
| chr12 | 55992425 | 55992450 | + | 43.1703048055547 | 19.3289276150945 | 9.54582568294392 | 5.33554323488568 |
| chr12 | 55992725 | 55992732 | + | 15.0594086531005 | 9.09596593651506 | 13.1255103140479 | 5.33554323488568 |
| chr12 | 55992902 | 55992945 | + | 11.043566345607 | 4.54798296825753 | 4.77291284147196 | 5.33554323488568 |
| chr12 | 55992994 | 55993009 | + | 11.043566345607 | 9.09596593651506 | 5.96614105183995 | 5.33554323488568 |
| chr12 | 55993071 | 55993086 | + | 8.03168461498692 | 5.68497871032191 | 4.77291284147196 | 6.40265188186281 |
| chr12 | 55993185 | 55993205 | + | 9.03564519186028 | 9.09596593651506 | 3.57968463110397 | 5.33554323488568 |
| chr12 | 55993503 | 55993508 | + | 7.02772403811355 | 5.68497871032191 | 5.96614105183995 | 1.06710864697714 |
| chr12 | 55993745 | 55993748 | + | 4.01584230749346 | 5.68497871032191 | 4.77291284147196 | 14.9395210576799 |
| chr12 | 55993808 | 55993857 | + | 8.03168461498692 | 3.41098722619315 | 17.8984231555198 | 5.33554323488568 |
| chr12 | 55994167 | 55994170 | + | 2.00792115374673 | 1.13699574206438 | 2.38645642073598 | 3.20132594093141 |
| chr12 | 55994248 | 55994264 | + | 4.01584230749346 | 4.54798296825753 | 5.96614105183995 | 7.46976052883995 |
| chr12 | 55994290 | 55994336 | + | 6.02376346124019 | 2.27399148412876 | 9.54582568294392 | 9.60397782279422 |
| chr12 | 55994357 | 55994359 | + | 3.01188173062009 | 3.41098722619315 | 3.57968463110397 | 2.13421729395427 |
| chr12 | 55994397 | 55994450 | + | 59.2336740355285 | 28.4248935516096 | 38.1833027317757 | 38.4159112911769 |
| chr12 | 55994470 | 55994500 | + | 177.701022106586 | 120.521548658824 | 168.245177661887 | 88.5700176991023 |
| chr12 | 55994524 | 55994568 | + | 1323.22004031909 | 983.50131688569 | 1274.36772867301 | 706.425924298864 |
| chr12 | 55994592 | 55994605 | + | 13.0514874993537 | 17.0549361309657 | 25.0577924177278 | 14.9395210576799 |
| chr12 | 55994650 | 55994672 | + | 8.03168461498692 | 10.2329616785794 | 3.57968463110397 | 1.06710864697714 |
| chr12 | 56005454 | 56005467 | + | 4.01584230749346 | 6.82197445238629 | 4.77291284147196 | 11.7381951167485 |
| chr12 | 56037065 | 56037097 | + | 1.00396057687336 | 2.27399148412876 | 10.7390538933119 | 5.33554323488568 |
| chr12 | 56037233 | 56037285 | + | 4.01584230749346 | 3.41098722619315 | 16.7051949451519 | 6.40265188186281 |
| chr12 | 56037705 | 56037709 | + | 2.00792115374673 | 2.27399148412876 | 10.7390538933119 | 4.26843458790854 |
| chr12 | 56102439 | 56102471 | + | 2.00792115374673 | 3.41098722619315 | 3.57968463110397 | 1.06710864697714 |
| chr12 | 56103159 | 56103316 | + | 1.00396057687336 | 6.82197445238629 | 3.57968463110397 | 1.06710864697714 |
| chr12 | 56112996 | 56113001 | + | 9.03564519186028 | 1.13699574206438 | 3.57968463110397 | 8.53686917581709 |
| chr12 | 56113075 | 56113097 | + | 167.661416337852 | 191.015284666816 | 145.573841664895 | 317.998376799186 |
| chr12 | 56113701 | 56113729 | + | 20.0792115374673 | 17.0549361309657 | 9.54582568294392 | 39.483019938154 |
| chr12 | 56113747 | 56113802 | + | 17.0673298068472 | 25.0139063254164 | 16.7051949451519 | 13.8724124107028 |
| chr12 | 56117805 | 56117823 | + | 7.02772403811355 | 17.0549361309657 | 14.3187385244159 | 7.46976052883995 |
| chr12 | 56144113 | 56144186 | + | 5.01980288436682 | 6.82197445238629 | 21.4781077866238 | 42.6843458790854 |
| chr12 | 56144207 | 56144246 | + | 4.01584230749346 | 2.27399148412876 | 21.4781077866238 | 28.8119334683827 |
| chr12 | 56144267 | 56144306 | + | 4.01584230749346 | 2.27399148412876 | 7.15936926220794 | 12.8053037637256 |
| chr12 | 56144414 | 56144455 | + | 3.01188173062009 | 5.68497871032191 | 22.6713359969918 | 13.8724124107028 |
| chr12 | 56144517 | 56144648 | + | 45.1782259593014 | 45.4798296825753 | 76.3666054635513 | 87.5029090521251 |
| chr12 | 56157785 | 56157830 | + | 26.1029749987075 | 18.1919318730301 | 3.57968463110397 | 2.13421729395427 |
| chr12 | 56157912 | 56158035 | + | 99.3920971104631 | 100.055625301666 | 42.9562155732476 | 44.8185631730397 |
| chr12 | 56160366 | 56160368 | + | 2.00792115374673 | 1.13699574206438 | 1.19322821036799 | 1.06710864697714 |
| chr12 | 56160502 | 56160523 | + | 12.0475269224804 | 11.3699574206438 | 9.54582568294392 | 2.13421729395427 |
| chr12 | 56161391 | 56161556 | + | 112.443584609817 | 143.261463500112 | 42.9562155732476 | 54.4225409958339 |
| chr12 | 56163382 | 56163387 | - | 5.01980288436682 | 2.27399148412876 | 2.38645642073598 | 1.06710864697714 |
| chr12 | 56163575 | 56163597 | - | 2.00792115374673 | 4.54798296825753 | 2.38645642073598 | 6.40265188186281 |
| chr12 | 56204535 | 56204537 | - | 1.00396057687336 | 1.13699574206438 | 2.38645642073598 | 1.06710864697714 |
| chr12 | 56205392 | 56205399 | - | 26.1029749987075 | 61.3977700714766 | 3.57968463110397 | 7.46976052883995 |
| chr12 | 56206270 | 56206276 | - | 1.00396057687336 | 2.27399148412876 | 3.57968463110397 | 1.06710864697714 |
| chr12 | 56229800 | 56229834 | + | 20.0792115374673 | 23.876910583352 | 8.35259747257593 | 8.53686917581709 |
| chr12 | 56237834 | 56237842 | - | 37.1465413443145 | 39.7948509722534 | 11.9322821036799 | 3.20132594093141 |
| chr12 | 56238557 | 56238565 | - | 26.1029749987075 | 23.876910583352 | 11.9322821036799 | 13.8724124107028 |
| chr12 | 56238734 | 56238747 | - | 9.03564519186028 | 6.82197445238629 | 2.38645642073598 | 1.06710864697714 |
| chr12 | 56241210 | 56241221 | - | 8.03168461498692 | 3.41098722619315 | 1.19322821036799 | 1.06710864697714 |
| chr12 | 56241339 | 56241341 | - | 7.02772403811355 | 4.54798296825753 | 1.19322821036799 | 2.13421729395427 |
| chr12 | 56270858 | 56270901 | + | 3.01188173062009 | 5.68497871032191 | 2.38645642073598 | 1.06710864697714 |
| chr12 | 56270923 | 56270925 | + | 1.00396057687336 | 1.13699574206438 | 1.19322821036799 | 4.26843458790854 |
| chr12 | 56271725 | 56271798 | - | 21.0831721143407 | 34.1098722619315 | 14.3187385244159 | 19.2079556455884 |
| chr12 | 56271962 | 56271976 | - | 15.0594086531005 | 10.2329616785794 | 31.0239334695677 | 20.2750642925656 |
| chr12 | 56272042 | 56272066 | - | 47.1861471130481 | 52.3018041349616 | 94.2650286190712 | 49.0869977609482 |
| chr12 | 56272146 | 56272235 | - | 25.0990144218341 | 20.4659233571589 | 38.1833027317757 | 35.2145853502455 |
| chr12 | 56272270 | 56272288 | - | 2.00792115374673 | 7.95897019445067 | 10.7390538933119 | 5.33554323488568 |
| chr12 | 56272351 | 56272393 | - | 12.0475269224804 | 9.09596593651506 | 11.9322821036799 | 8.53686917581709 |
| chr12 | 56272619 | 56272641 | - | 4.01584230749346 | 1.13699574206438 | 5.96614105183995 | 10.6710864697714 |
| chr12 | 56272717 | 56272762 | - | 18.0712903837206 | 10.2329616785794 | 11.9322821036799 | 7.46976052883995 |
| chr12 | 56272898 | 56272918 | - | 4.01584230749346 | 3.41098722619315 | 7.15936926220794 | 9.60397782279422 |
| chr12 | 56273070 | 56273092 | - | 2.00792115374673 | 9.09596593651506 | 14.3187385244159 | 5.33554323488568 |
| chr12 | 56300097 | 56300153 | + | 30.1188173062009 | 37.5208594881246 | 39.3765309421437 | 23.476390233497 |
| chr12 | 56310454 | 56310590 | - | 148.586165377258 | 178.508331504108 | 35.7968463110397 | 52.2883237018796 |
| chr12 | 56316240 | 56316254 | - | 4.01584230749346 | 1.13699574206438 | 4.77291284147196 | 6.40265188186281 |
| chr12 | 56316303 | 56316369 | - | 32.1267384599477 | 32.9728765198671 | 70.4004644117114 | 61.8923015246739 |
| chr12 | 56317110 | 56317188 | - | 57.2257528817818 | 70.4937360079917 | 41.7629873628796 | 39.483019938154 |
| chr12 | 56317300 | 56317325 | - | 5.01980288436682 | 6.82197445238629 | 4.77291284147196 | 2.13421729395427 |
| chr12 | 56317484 | 56317501 | - | 5.01980288436682 | 12.5069531627082 | 7.15936926220794 | 3.20132594093141 |
| chr12 | 56417151 | 56417202 | - | 28.1108961524542 | 35.2468680039958 | 39.3765309421437 | 27.7448248214055 |
| chr12 | 56417306 | 56417354 | - | 35.1386201905678 | 30.6988850357383 | 17.8984231555198 | 30.9461507623369 |
| chr12 | 56417586 | 56417612 | - | 5.01980288436682 | 7.95897019445067 | 11.9322821036799 | 21.3421729395427 |
| chr12 | 56417639 | 56417788 | - | 44.174265382428 | 34.1098722619315 | 114.549908195327 | 67.2278447595595 |
| chr12 | 56595958 | 56596009 | - | 2.00792115374673 | 3.41098722619315 | 10.7390538933119 | 5.33554323488568 |
| chr12 | 56596123 | 56596189 | - | 3.01188173062009 | 7.95897019445067 | 2.38645642073598 | 6.40265188186281 |
| chr12 | 56596222 | 56596231 | - | 3.01188173062009 | 3.41098722619315 | 2.38645642073598 | 7.46976052883995 |
| chr12 | 56597157 | 56597227 | - | 38.1505019211879 | 30.6988850357383 | 35.7968463110397 | 24.5434988804741 |
| chr12 | 56597316 | 56597424 | - | 1.00396057687336 | 4.54798296825753 | 27.4442488384638 | 13.8724124107028 |
| chr12 | 56597645 | 56597652 | - | 5.01980288436682 | 2.27399148412876 | 4.77291284147196 | 2.13421729395427 |
| chr12 | 56597685 | 56597712 | - | 4.01584230749346 | 9.09596593651506 | 9.54582568294392 | 18.1408469986113 |
| chr12 | 56597804 | 56597829 | - | 10.0396057687336 | 5.68497871032191 | 3.57968463110397 | 7.46976052883995 |
| chr12 | 56598000 | 56598007 | - | 6.02376346124019 | 4.54798296825753 | 5.96614105183995 | 2.13421729395427 |
| chr12 | 56598315 | 56598318 | - | 26.1029749987075 | 22.7399148412876 | 17.8984231555198 | 12.8053037637256 |
| chr12 | 56598392 | 56598419 | - | 2.00792115374673 | 4.54798296825753 | 1.19322821036799 | 3.20132594093141 |
| chr12 | 56598589 | 56598611 | - | 7.02772403811355 | 9.09596593651506 | 7.15936926220794 | 5.33554323488568 |
| chr12 | 56638201 | 56638426 | - | 1690.66961145475 | 1686.16468548148 | 1125.21420237701 | 1000.94791086455 |
| chr12 | 56663362 | 56663370 | - | 3.01188173062009 | 5.68497871032191 | 8.35259747257593 | 6.40265188186281 |
| chr12 | 56663392 | 56663452 | - | 77.3049644192491 | 109.151591238181 | 108.583767143487 | 109.912190638645 |
| chr12 | 56663471 | 56663480 | - | 7.02772403811355 | 6.82197445238629 | 5.96614105183995 | 1.06710864697714 |
| chr12 | 56663533 | 56663543 | - | 3.01188173062009 | 9.09596593651506 | 21.4781077866238 | 22.4092815865198 |
| chr12 | 56663645 | 56663657 | - | 37.1465413443145 | 52.3018041349616 | 44.1494437836156 | 66.1607361125824 |
| chr12 | 56663710 | 56663745 | - | 22.087132691214 | 25.0139063254164 | 32.2171616799357 | 11.7381951167485 |
| chr12 | 56663775 | 56663784 | - | 1.00396057687336 | 2.27399148412876 | 4.77291284147196 | 1.06710864697714 |
| chr12 | 56663826 | 56663836 | - | 2.00792115374673 | 11.3699574206438 | 14.3187385244159 | 21.3421729395427 |
| chr12 | 56663872 | 56663896 | - | 5.01980288436682 | 2.27399148412876 | 9.54582568294392 | 10.6710864697714 |
| chr12 | 56663930 | 56663968 | - | 17.0673298068472 | 11.3699574206438 | 25.0577924177278 | 46.952780466994 |
| chr12 | 56664028 | 56664044 | - | 3.01188173062009 | 2.27399148412876 | 3.57968463110397 | 5.33554323488568 |
| chr12 | 56664090 | 56664134 | - | 4.01584230749346 | 2.27399148412876 | 9.54582568294392 | 3.20132594093141 |
| chr12 | 56664198 | 56664228 | - | 8.03168461498692 | 3.41098722619315 | 14.3187385244159 | 9.60397782279422 |
| chr12 | 56664381 | 56664421 | - | 7.02772403811355 | 5.68497871032191 | 25.0577924177278 | 6.40265188186281 |
| chr12 | 56664477 | 56664485 | - | 8.03168461498692 | 15.9179403889013 | 10.7390538933119 | 4.26843458790854 |
| chr12 | 56712530 | 56712555 | - | 7.02772403811355 | 7.95897019445067 | 17.8984231555198 | 8.53686917581709 |
| chr12 | 56731616 | 56731633 | - | 1.00396057687336 | 3.41098722619315 | 2.38645642073598 | 3.20132594093141 |
| chr12 | 57000067 | 57000069 | - | 2.00792115374673 | 2.27399148412876 | 7.15936926220794 | 2.13421729395427 |
| chr12 | 57055750 | 57055771 | - | 7.02772403811355 | 12.5069531627082 | 9.54582568294392 | 4.26843458790854 |
| chr12 | 57056289 | 57056303 | - | 4.01584230749346 | 3.41098722619315 | 2.38645642073598 | 5.33554323488568 |
| chr12 | 57056350 | 57056361 | - | 6.02376346124019 | 11.3699574206438 | 26.2510206280958 | 26.6777161744284 |
| chr12 | 57056383 | 57056402 | - | 7.02772403811355 | 13.6439489047726 | 25.0577924177278 | 36.2816939972226 |
| chr12 | 57056458 | 57056484 | - | 2.00792115374673 | 5.68497871032191 | 7.15936926220794 | 11.7381951167485 |
| chr12 | 57056578 | 57056600 | - | 8.03168461498692 | 12.5069531627082 | 15.5119667347839 | 16.006629704657 |
| chr12 | 57056656 | 57056666 | - | 4.01584230749346 | 12.5069531627082 | 14.3187385244159 | 8.53686917581709 |
| chr12 | 57057218 | 57057226 | - | 3.01188173062009 | 2.27399148412876 | 10.7390538933119 | 8.53686917581709 |
| chr12 | 57057311 | 57057324 | - | 5.01980288436682 | 4.54798296825753 | 7.15936926220794 | 1.06710864697714 |
| chr12 | 57057848 | 57057857 | - | 8.03168461498692 | 3.41098722619315 | 10.7390538933119 | 4.26843458790854 |
| chr12 | 57058226 | 57058229 | - | 3.01188173062009 | 4.54798296825753 | 1.19322821036799 | 1.06710864697714 |
| chr12 | 57058496 | 57058497 | - | 1.00396057687336 | 1.13699574206438 | 1.19322821036799 | 1.06710864697714 |
| chr12 | 57095364 | 57095367 | + | 6.02376346124019 | 5.68497871032191 | 4.77291284147196 | 10.6710864697714 |
| chr12 | 57095483 | 57095487 | - | 8.03168461498692 | 20.4659233571589 | 7.15936926220794 | 5.33554323488568 |
| chr12 | 57095598 | 57095622 | - | 8.03168461498692 | 6.82197445238629 | 2.38645642073598 | 2.13421729395427 |
| chr12 | 57095919 | 57095927 | - | 8.03168461498692 | 2.27399148412876 | 9.54582568294392 | 5.33554323488568 |
| chr12 | 57095960 | 57095973 | - | 10.0396057687336 | 4.54798296825753 | 2.38645642073598 | 3.20132594093141 |
| chr12 | 57096206 | 57096251 | - | 14.0554480762271 | 10.2329616785794 | 9.54582568294392 | 2.13421729395427 |
| chr12 | 57226414 | 57226424 | + | 10.0396057687336 | 11.3699574206438 | 4.77291284147196 | 2.13421729395427 |
| chr12 | 57234349 | 57234380 | + | 3.01188173062009 | 2.27399148412876 | 4.77291284147196 | 2.13421729395427 |
| chr12 | 57234458 | 57234523 | + | 5.01980288436682 | 6.82197445238629 | 7.15936926220794 | 16.006629704657 |
| chr12 | 57234607 | 57234698 | + | 67.2653586505154 | 86.411676396893 | 35.7968463110397 | 80.0331485232852 |
| chr12 | 57234881 | 57234908 | + | 5.01980288436682 | 6.82197445238629 | 1.19322821036799 | 3.20132594093141 |
| chr12 | 57253806 | 57253825 | - | 4.01584230749346 | 11.3699574206438 | 13.1255103140479 | 13.8724124107028 |
| chr12 | 57256502 | 57256513 | - | 2.00792115374673 | 2.27399148412876 | 16.7051949451519 | 2.13421729395427 |
| chr12 | 57268971 | 57268974 | - | 2.00792115374673 | 1.13699574206438 | 3.57968463110397 | 4.26843458790854 |
| chr12 | 57490237 | 57490267 | + | 5.01980288436682 | 4.54798296825753 | 9.54582568294392 | 2.13421729395427 |
| chr12 | 57490285 | 57490304 | + | 2.00792115374673 | 4.54798296825753 | 2.38645642073598 | 1.06710864697714 |
| chr12 | 57490349 | 57490375 | + | 6.02376346124019 | 12.5069531627082 | 28.6374770488317 | 5.33554323488568 |
| chr12 | 57490537 | 57490568 | + | 17.0673298068472 | 29.5618892936739 | 34.6036181006717 | 27.7448248214055 |
| chr12 | 57490588 | 57490602 | + | 1.00396057687336 | 14.780944646837 | 13.1255103140479 | 6.40265188186281 |
| chr12 | 57490632 | 57490645 | + | 7.02772403811355 | 6.82197445238629 | 4.77291284147196 | 6.40265188186281 |
| chr12 | 57498166 | 57498214 | + | 5.01980288436682 | 6.82197445238629 | 15.5119667347839 | 7.46976052883995 |
| chr12 | 57498437 | 57498476 | + | 10.0396057687336 | 18.1919318730301 | 20.2848795762558 | 10.6710864697714 |
| chr12 | 57498500 | 57498553 | + | 5.01980288436682 | 18.1919318730301 | 47.7291284147196 | 18.1408469986113 |
| chr12 | 57500403 | 57500523 | + | 19.0752509605939 | 18.1919318730301 | 31.0239334695677 | 34.1474767032683 |
| chr12 | 57504257 | 57504297 | + | 7.02772403811355 | 4.54798296825753 | 9.54582568294392 | 8.53686917581709 |
| chr12 | 57511696 | 57511761 | + | 17.0673298068472 | 22.7399148412876 | 17.8984231555198 | 25.6106075274513 |
| chr12 | 57511828 | 57511866 | + | 13.0514874993537 | 17.0549361309657 | 38.1833027317757 | 21.3421729395427 |
| chr12 | 57512006 | 57512099 | + | 17.0673298068472 | 20.4659233571589 | 23.8645642073598 | 35.2145853502455 |
| chr12 | 57512233 | 57512407 | + | 24.0950538449607 | 34.1098722619315 | 76.3666054635513 | 43.7514545260626 |
| chr12 | 57512748 | 57512813 | + | 10.0396057687336 | 31.8358807778027 | 42.9562155732476 | 28.8119334683827 |
| chr12 | 57512859 | 57512885 | + | 2.00792115374673 | 6.82197445238629 | 22.6713359969918 | 3.20132594093141 |
| chr12 | 57512913 | 57512965 | + | 23.0910932680874 | 22.7399148412876 | 20.2848795762558 | 22.4092815865198 |
| chr12 | 57514717 | 57514765 | + | 17.0673298068472 | 31.8358807778027 | 47.7291284147196 | 45.8856718200168 |
| chr12 | 57514797 | 57514816 | + | 2.00792115374673 | 3.41098722619315 | 7.15936926220794 | 4.26843458790854 |
| chr12 | 57514952 | 57514971 | + | 8.03168461498692 | 9.09596593651506 | 17.8984231555198 | 11.7381951167485 |
| chr12 | 57514991 | 57515051 | + | 24.0950538449607 | 28.4248935516096 | 41.7629873628796 | 78.966039876308 |
| chr12 | 57515147 | 57515190 | + | 15.0594086531005 | 22.7399148412876 | 41.7629873628796 | 27.7448248214055 |
| chr12 | 57515980 | 57515992 | + | 8.03168461498692 | 7.95897019445067 | 16.7051949451519 | 12.8053037637256 |
| chr12 | 57516481 | 57516633 | + | 156.617849992245 | 369.523616170924 | 184.950372607038 | 203.817751572633 |
| chr12 | 57516618 | 57516696 | - | 6.02376346124019 | 328.591769456606 | 89.4921157775992 | 81.1002571702623 |
| chr12 | 57516778 | 57516801 | - | 1.00396057687336 | 12.5069531627082 | 4.77291284147196 | 3.20132594093141 |
| chr12 | 57529460 | 57529477 | + | 5.01980288436682 | 2.27399148412876 | 17.8984231555198 | 5.33554323488568 |
| chr12 | 57529501 | 57529533 | + | 25.0990144218341 | 23.876910583352 | 33.4103898903037 | 21.3421729395427 |
| chr12 | 57529984 | 57530011 | + | 2.00792115374673 | 4.54798296825753 | 1.19322821036799 | 7.46976052883995 |
| chr12 | 57530083 | 57530128 | + | 7.02772403811355 | 10.2329616785794 | 7.15936926220794 | 5.33554323488568 |
| chr12 | 57530270 | 57530286 | - | 3.01188173062009 | 2.27399148412876 | 3.57968463110397 | 3.20132594093141 |
| chr12 | 57530330 | 57530410 | - | 98.3881365335897 | 86.411676396893 | 89.4921157775992 | 115.247733873531 |
| chr12 | 57530438 | 57530521 | - | 21.0831721143407 | 15.9179403889013 | 14.3187385244159 | 23.476390233497 |
| chr12 | 57530547 | 57530573 | - | 6.02376346124019 | 1.13699574206438 | 4.77291284147196 | 8.53686917581709 |
| chr12 | 57530678 | 57530691 | - | 2.00792115374673 | 3.41098722619315 | 1.19322821036799 | 2.13421729395427 |
| chr12 | 57720824 | 57720943 | + | 47.1861471130481 | 50.0278126508328 | 106.197310722751 | 29.8790421153598 |
| chr12 | 57720996 | 57721029 | + | 14.0554480762271 | 7.95897019445067 | 3.57968463110397 | 8.53686917581709 |
| chr12 | 57721084 | 57721126 | + | 33.130699036821 | 13.6439489047726 | 17.8984231555198 | 14.9395210576799 |
| chr12 | 57721268 | 57721364 | + | 9.03564519186028 | 6.82197445238629 | 1.19322821036799 | 4.26843458790854 |
| chr12 | 57721416 | 57721457 | + | 6.02376346124019 | 14.780944646837 | 11.9322821036799 | 17.0737383516342 |
| chr12 | 57721488 | 57721533 | + | 10.0396057687336 | 14.780944646837 | 11.9322821036799 | 6.40265188186281 |
| chr12 | 57748253 | 57748297 | - | 24.0950538449607 | 46.6168254246397 | 21.4781077866238 | 21.3421729395427 |
| chr12 | 57748302 | 57748325 | - | 134.530717301031 | 188.741293182687 | 54.8884976769275 | 103.509538756782 |
| chr12 | 57748355 | 57748358 | - | 4.01584230749346 | 5.68497871032191 | 1.19322821036799 | 4.26843458790854 |
| chr12 | 57748402 | 57748439 | - | 22.087132691214 | 21.6029190992233 | 13.1255103140479 | 12.8053037637256 |
| chr12 | 57749212 | 57749320 | - | 33.130699036821 | 34.1098722619315 | 31.0239334695677 | 34.1474767032683 |
| chr12 | 57759386 | 57759389 | + | 2.00792115374673 | 11.3699574206438 | 3.57968463110397 | 7.46976052883995 |
| chr12 | 57780364 | 57780366 | + | 1.00396057687336 | 1.13699574206438 | 3.57968463110397 | 3.20132594093141 |
| chr12 | 57796208 | 57796218 | + | 5.01980288436682 | 2.27399148412876 | 2.38645642073598 | 2.13421729395427 |
| chr12 | 57796430 | 57796472 | + | 9.03564519186028 | 7.95897019445067 | 13.1255103140479 | 6.40265188186281 |
| chr12 | 57796474 | 57796573 | + | 60.2376346124019 | 54.5757956190903 | 17.8984231555198 | 36.2816939972226 |
| chr12 | 57796603 | 57796646 | + | 19.0752509605939 | 20.4659233571589 | 15.5119667347839 | 13.8724124107028 |
| chr12 | 57796671 | 57796702 | + | 51.2019894205416 | 88.6856678810218 | 13.1255103140479 | 8.53686917581709 |
| chr12 | 57820432 | 57820453 | - | 5.01980288436682 | 4.54798296825753 | 4.77291284147196 | 9.60397782279422 |
| chr12 | 57820548 | 57820597 | - | 8.03168461498692 | 4.54798296825753 | 4.77291284147196 | 12.8053037637256 |
| chr12 | 57820739 | 57820754 | - | 2.00792115374673 | 4.54798296825753 | 5.96614105183995 | 5.33554323488568 |
| chr12 | 57820800 | 57820815 | - | 8.03168461498692 | 19.3289276150945 | 21.4781077866238 | 27.7448248214055 |
| chr12 | 57821118 | 57821149 | - | 6.02376346124019 | 11.3699574206438 | 8.35259747257593 | 12.8053037637256 |
| chr12 | 57821686 | 57821737 | - | 3.01188173062009 | 4.54798296825753 | 10.7390538933119 | 5.33554323488568 |
| chr12 | 57822002 | 57822020 | - | 3.01188173062009 | 3.41098722619315 | 8.35259747257593 | 2.13421729395427 |
| chr12 | 57822084 | 57822104 | - | 13.0514874993537 | 20.4659233571589 | 57.2749540976635 | 9.60397782279422 |
| chr12 | 57822137 | 57822155 | - | 4.01584230749346 | 5.68497871032191 | 13.1255103140479 | 3.20132594093141 |
| chr12 | 57822530 | 57822549 | - | 5.01980288436682 | 3.41098722619315 | 3.57968463110397 | 2.13421729395427 |
| chr12 | 57822607 | 57822619 | - | 3.01188173062009 | 3.41098722619315 | 2.38645642073598 | 3.20132594093141 |
| chr12 | 57822658 | 57822698 | - | 9.03564519186028 | 10.2329616785794 | 5.96614105183995 | 1.06710864697714 |
| chr12 | 57822854 | 57822890 | - | 14.0554480762271 | 15.9179403889013 | 19.0916513658878 | 11.7381951167485 |
| chr12 | 57822964 | 57823017 | - | 5.01980288436682 | 1.13699574206438 | 2.38645642073598 | 2.13421729395427 |
| chr12 | 57823210 | 57823235 | - | 2.00792115374673 | 3.41098722619315 | 5.96614105183995 | 1.06710864697714 |
| chr12 | 57823291 | 57823311 | - | 6.02376346124019 | 11.3699574206438 | 9.54582568294392 | 3.20132594093141 |
| chr12 | 57823333 | 57823403 | - | 22.087132691214 | 12.5069531627082 | 13.1255103140479 | 5.33554323488568 |
| chr12 | 57823437 | 57823581 | - | 31.1227778830743 | 30.6988850357383 | 14.3187385244159 | 20.2750642925656 |
| chr12 | 62260328 | 62260435 | + | 658.598138428927 | 933.473504234858 | 1559.54927095096 | 1877.04411003278 |
| chr12 | 62404199 | 62404390 | + | 76.3010038423757 | 118.247557174696 | 223.133675338814 | 224.092815865199 |
| chr12 | 62404411 | 62404436 | + | 16.0633692299738 | 23.876910583352 | 40.5697591525116 | 35.2145853502455 |
| chr12 | 62404466 | 62404490 | + | 26.1029749987075 | 56.8497871032191 | 97.8447132501751 | 76.8318225823538 |
| chr12 | 62404521 | 62404615 | + | 10.0396057687336 | 22.7399148412876 | 38.1833027317757 | 25.6106075274513 |
| chr12 | 62409545 | 62409552 | + | 11.043566345607 | 5.68497871032191 | 13.1255103140479 | 8.53686917581709 |
| chr12 | 62602814 | 62602867 | + | 23.0910932680874 | 18.1919318730301 | 5.96614105183995 | 8.53686917581709 |
| chr12 | 62603766 | 62603777 | + | 8.03168461498692 | 10.2329616785794 | 15.5119667347839 | 7.46976052883995 |
| chr12 | 62604163 | 62604164 | + | 1.00396057687336 | 1.13699574206438 | 1.19322821036799 | 2.13421729395427 |
| chr12 | 62647317 | 62647320 | - | 2.00792115374673 | 3.41098722619315 | 4.77291284147196 | 1.06710864697714 |
| chr12 | 64193445 | 64193446 | - | 1.00396057687336 | 2.27399148412876 | 1.19322821036799 | 2.13421729395427 |
| chr12 | 64448117 | 64448149 | + | 2.00792115374673 | 3.41098722619315 | 2.38645642073598 | 4.26843458790854 |
| chr12 | 64448186 | 64448190 | + | 3.01188173062009 | 2.27399148412876 | 1.19322821036799 | 4.26843458790854 |
| chr12 | 64448243 | 64448262 | + | 2.00792115374673 | 7.95897019445067 | 3.57968463110397 | 2.13421729395427 |
| chr12 | 64448352 | 64448387 | + | 5.01980288436682 | 7.95897019445067 | 5.96614105183995 | 6.40265188186281 |
| chr12 | 64448418 | 64448472 | + | 3.01188173062009 | 9.09596593651506 | 4.77291284147196 | 11.7381951167485 |
| chr12 | 64448523 | 64448533 | + | 73.2891221117556 | 201.248246345396 | 71.5936926220794 | 108.845081991668 |
| chr12 | 64448621 | 64448634 | + | 10.0396057687336 | 21.6029190992233 | 8.35259747257593 | 6.40265188186281 |
| chr12 | 64501806 | 64501923 | + | 238.942617295861 | 185.330305956494 | 33.4103898903037 | 107.777973344691 |
| chr12 | 64695165 | 64695203 | + | 31.1227778830743 | 35.2468680039958 | 11.9322821036799 | 30.9461507623369 |
| chr12 | 64695229 | 64695238 | + | 4.01584230749346 | 4.54798296825753 | 7.15936926220794 | 11.7381951167485 |
| chr12 | 64695285 | 64695327 | + | 9.03564519186028 | 5.68497871032191 | 10.7390538933119 | 22.4092815865198 |
| chr12 | 64695395 | 64695457 | + | 19.0752509605939 | 20.4659233571589 | 9.54582568294392 | 11.7381951167485 |
| chr12 | 64695521 | 64695561 | + | 30.1188173062009 | 22.7399148412876 | 10.7390538933119 | 5.33554323488568 |
| chr12 | 64695591 | 64695618 | + | 4.01584230749346 | 4.54798296825753 | 5.96614105183995 | 8.53686917581709 |
| chr12 | 64695875 | 64695879 | + | 2.00792115374673 | 1.13699574206438 | 2.38645642073598 | 5.33554323488568 |
| chr12 | 64695929 | 64695957 | + | 10.0396057687336 | 2.27399148412876 | 11.9322821036799 | 10.6710864697714 |
| chr12 | 64697373 | 64697420 | + | 10.0396057687336 | 13.6439489047726 | 4.77291284147196 | 10.6710864697714 |
| chr12 | 64697471 | 64697534 | + | 5.01980288436682 | 4.54798296825753 | 2.38645642073598 | 5.33554323488568 |
| chr12 | 64713514 | 64713654 | - | 369.457492289398 | 263.783012158937 | 96.6514850398071 | 92.8384522870108 |
| chr12 | 64713675 | 64713823 | - | 154.609928838498 | 118.247557174696 | 40.5697591525116 | 40.5501285851312 |
| chr12 | 64713850 | 64713870 | - | 46.1821865361748 | 23.876910583352 | 2.38645642073598 | 14.9395210576799 |
| chr12 | 64714152 | 64714200 | - | 8.03168461498692 | 4.54798296825753 | 4.77291284147196 | 7.46976052883995 |
| chr12 | 64714580 | 64714608 | - | 5.01980288436682 | 11.3699574206438 | 4.77291284147196 | 2.13421729395427 |
| chr12 | 64714925 | 64714944 | - | 11.043566345607 | 7.95897019445067 | 4.77291284147196 | 4.26843458790854 |
| chr12 | 64715027 | 64715048 | - | 3.01188173062009 | 2.27399148412876 | 1.19322821036799 | 2.13421729395427 |
| chr12 | 64715092 | 64715187 | - | 37.1465413443145 | 26.1509020674808 | 21.4781077866238 | 23.476390233497 |
| chr12 | 64715234 | 64715260 | - | 11.043566345607 | 15.9179403889013 | 10.7390538933119 | 8.53686917581709 |
| chr12 | 65248217 | 65248220 | + | 3.01188173062009 | 3.41098722619315 | 4.77291284147196 | 3.20132594093141 |
| chr12 | 65963424 | 65963470 | + | 35.1386201905678 | 40.9318467143178 | 4.77291284147196 | 19.2079556455884 |
| chr12 | 66123105 | 66123111 | - | 3.01188173062009 | 2.27399148412876 | 1.19322821036799 | 10.6710864697714 |
| chr12 | 66123798 | 66123810 | - | 7.02772403811355 | 10.2329616785794 | 2.38645642073598 | 5.33554323488568 |
| chr12 | 66128895 | 66128896 | - | 5.01980288436682 | 2.27399148412876 | 2.38645642073598 | 5.33554323488568 |
| chr12 | 66130787 | 66130811 | + | 5.01980288436682 | 6.82197445238629 | 23.8645642073598 | 32.0132594093141 |
| chr12 | 66130847 | 66130922 | + | 39.1544624980612 | 52.3018041349616 | 51.3088130458235 | 48.0198891139711 |
| chr12 | 66137866 | 66137914 | - | 28.1108961524542 | 31.8358807778027 | 14.3187385244159 | 29.8790421153598 |
| chr12 | 66137976 | 66138000 | - | 5.01980288436682 | 11.3699574206438 | 10.7390538933119 | 9.60397782279422 |
| chr12 | 66138037 | 66138063 | - | 4.01584230749346 | 7.95897019445067 | 10.7390538933119 | 11.7381951167485 |
| chr12 | 66138118 | 66138143 | - | 14.0554480762271 | 10.2329616785794 | 5.96614105183995 | 14.9395210576799 |
| chr12 | 67312846 | 67312849 | + | 6.02376346124019 | 2.27399148412876 | 5.96614105183995 | 4.26843458790854 |
| chr12 | 67312881 | 67312925 | + | 4.01584230749346 | 9.09596593651506 | 5.96614105183995 | 7.46976052883995 |
| chr12 | 67312953 | 67312967 | + | 4.01584230749346 | 4.54798296825753 | 13.1255103140479 | 11.7381951167485 |
| chr12 | 67313031 | 67313035 | + | 4.01584230749346 | 4.54798296825753 | 5.96614105183995 | 1.06710864697714 |
| chr12 | 67313057 | 67313061 | + | 1.00396057687336 | 2.27399148412876 | 1.19322821036799 | 1.06710864697714 |
| chr12 | 67313087 | 67313146 | + | 43.1703048055547 | 50.0278126508328 | 40.5697591525116 | 49.0869977609482 |
| chr12 | 67313562 | 67313585 | + | 3.01188173062009 | 1.13699574206438 | 5.96614105183995 | 5.33554323488568 |
| chr12 | 67313862 | 67313871 | + | 2.00792115374673 | 3.41098722619315 | 7.15936926220794 | 2.13421729395427 |
| chr12 | 67313920 | 67314007 | + | 31.1227778830743 | 54.5757956190903 | 46.5359002043516 | 36.2816939972226 |
| chr12 | 67314049 | 67314094 | + | 7.02772403811355 | 11.3699574206438 | 7.15936926220794 | 8.53686917581709 |
| chr12 | 67314143 | 67314196 | + | 12.0475269224804 | 7.95897019445067 | 29.8307052591997 | 23.476390233497 |
| chr12 | 67314538 | 67314549 | + | 1.00396057687336 | 5.68497871032191 | 3.57968463110397 | 2.13421729395427 |
| chr12 | 67662001 | 67662017 | + | 3.01188173062009 | 5.68497871032191 | 1.19322821036799 | 3.20132594093141 |
| chr12 | 67662268 | 67662271 | + | 15.0594086531005 | 12.5069531627082 | 7.15936926220794 | 10.6710864697714 |
| chr12 | 67662354 | 67662391 | + | 12.0475269224804 | 9.09596593651506 | 11.9322821036799 | 16.006629704657 |
| chr12 | 67662445 | 67662453 | + | 6.02376346124019 | 3.41098722619315 | 3.57968463110397 | 4.26843458790854 |
| chr12 | 67664873 | 67664877 | + | 1.00396057687336 | 1.13699574206438 | 1.19322821036799 | 2.13421729395427 |
| chr12 | 68659813 | 68659822 | + | 1.00396057687336 | 1.13699574206438 | 10.7390538933119 | 9.60397782279422 |
| chr12 | 68660450 | 68660480 | + | 17.0673298068472 | 18.1919318730301 | 11.9322821036799 | 25.6106075274513 |
| chr12 | 68660508 | 68660510 | + | 2.00792115374673 | 2.27399148412876 | 1.19322821036799 | 1.06710864697714 |
| chr12 | 68742614 | 68742656 | + | 13.0514874993537 | 22.7399148412876 | 16.7051949451519 | 18.1408469986113 |
| chr12 | 68840433 | 68840449 | + | 30.1188173062009 | 85.2746806548286 | 7.15936926220794 | 9.60397782279422 |
| chr12 | 68842286 | 68842288 | + | 1.00396057687336 | 1.13699574206438 | 1.19322821036799 | 1.06710864697714 |
| chr12 | 68842365 | 68842378 | + | 3.01188173062009 | 3.41098722619315 | 1.19322821036799 | 9.60397782279422 |
| chr12 | 68842850 | 68842880 | + | 1.00396057687336 | 1.13699574206438 | 2.38645642073598 | 3.20132594093141 |
| chr12 | 68843031 | 68843041 | + | 1.00396057687336 | 1.13699574206438 | 2.38645642073598 | 10.6710864697714 |
| chr12 | 68843733 | 68843765 | + | 9.03564519186028 | 11.3699574206438 | 11.9322821036799 | 10.6710864697714 |
| chr12 | 68843832 | 68843854 | + | 3.01188173062009 | 4.54798296825753 | 5.96614105183995 | 1.06710864697714 |
| chr12 | 68843932 | 68843948 | + | 2.00792115374673 | 3.41098722619315 | 2.38645642073598 | 7.46976052883995 |
| chr12 | 68844052 | 68844060 | + | 3.01188173062009 | 1.13699574206438 | 7.15936926220794 | 5.33554323488568 |
| chr12 | 69269517 | 69269521 | + | 6.02376346124019 | 2.27399148412876 | 1.19322821036799 | 1.06710864697714 |
| chr12 | 69269544 | 69269558 | + | 28.1108961524542 | 22.7399148412876 | 19.0916513658878 | 29.8790421153598 |
| chr12 | 69269619 | 69269641 | + | 2.00792115374673 | 2.27399148412876 | 3.57968463110397 | 7.46976052883995 |
| chr12 | 69269779 | 69269824 | + | 3.01188173062009 | 4.54798296825753 | 3.57968463110397 | 2.13421729395427 |
| chr12 | 69269982 | 69269988 | + | 2.00792115374673 | 1.13699574206438 | 9.54582568294392 | 2.13421729395427 |
| chr12 | 69270535 | 69270538 | + | 1.00396057687336 | 1.13699574206438 | 4.77291284147196 | 2.13421729395427 |
| chr12 | 69270574 | 69270593 | + | 1.00396057687336 | 1.13699574206438 | 8.35259747257593 | 2.13421729395427 |
| chr12 | 69270622 | 69270641 | + | 1.00396057687336 | 1.13699574206438 | 4.77291284147196 | 1.06710864697714 |
| chr12 | 69270795 | 69270831 | + | 28.1108961524542 | 27.2878978095452 | 52.5020412561915 | 34.1474767032683 |
| chr12 | 69270917 | 69270920 | + | 1.00396057687336 | 3.41098722619315 | 3.57968463110397 | 2.13421729395427 |
| chr12 | 69271161 | 69271172 | + | 4.01584230749346 | 1.13699574206438 | 9.54582568294392 | 3.20132594093141 |
| chr12 | 69271799 | 69271824 | + | 4.01584230749346 | 3.41098722619315 | 3.57968463110397 | 1.06710864697714 |
| chr12 | 69272084 | 69272148 | + | 1.00396057687336 | 10.2329616785794 | 17.8984231555198 | 11.7381951167485 |
| chr12 | 69272181 | 69272199 | + | 1.00396057687336 | 4.54798296825753 | 4.77291284147196 | 7.46976052883995 |
| chr12 | 69272326 | 69272330 | + | 1.00396057687336 | 3.41098722619315 | 7.15936926220794 | 4.26843458790854 |
| chr12 | 69272421 | 69272439 | + | 6.02376346124019 | 3.41098722619315 | 10.7390538933119 | 5.33554323488568 |
| chr12 | 69272630 | 69272659 | + | 1.00396057687336 | 3.41098722619315 | 13.1255103140479 | 3.20132594093141 |
| chr12 | 69272811 | 69272847 | + | 4.01584230749346 | 5.68497871032191 | 3.57968463110397 | 8.53686917581709 |
| chr12 | 69272880 | 69272886 | + | 2.00792115374673 | 4.54798296825753 | 7.15936926220794 | 7.46976052883995 |
| chr12 | 69273024 | 69273025 | + | 1.00396057687336 | 1.13699574206438 | 1.19322821036799 | 1.06710864697714 |
| chr12 | 69273153 | 69273158 | + | 20.0792115374673 | 32.9728765198671 | 57.2749540976635 | 42.6843458790854 |
| chr12 | 69273231 | 69273267 | + | 3.01188173062009 | 2.27399148412876 | 11.9322821036799 | 2.13421729395427 |
| chr12 | 69577059 | 69577114 | + | 70.2772403811355 | 79.5897019445067 | 27.4442488384638 | 10.6710864697714 |
| chr12 | 69601334 | 69601372 | + | 20.0792115374673 | 21.6029190992233 | 20.2848795762558 | 30.9461507623369 |
| chr12 | 69601395 | 69601546 | + | 103.407939417957 | 97.7816338175368 | 95.4582568294392 | 88.5700176991023 |
| chr12 | 69815457 | 69815460 | + | 3.01188173062009 | 9.09596593651506 | 2.38645642073598 | 2.13421729395427 |
| chr12 | 69815653 | 69815665 | + | 2.00792115374673 | 7.95897019445067 | 1.19322821036799 | 3.20132594093141 |
| chr12 | 70353953 | 70353971 | + | 10.0396057687336 | 4.54798296825753 | 16.7051949451519 | 6.40265188186281 |
| chr12 | 70353990 | 70354011 | + | 3.01188173062009 | 4.54798296825753 | 8.35259747257593 | 7.46976052883995 |
| chr12 | 70354138 | 70354145 | + | 7.02772403811355 | 3.41098722619315 | 11.9322821036799 | 8.53686917581709 |
| chr12 | 70354385 | 70354395 | + | 1.00396057687336 | 2.27399148412876 | 3.57968463110397 | 2.13421729395427 |
| chr12 | 70354513 | 70354563 | + | 12.0475269224804 | 15.9179403889013 | 19.0916513658878 | 14.9395210576799 |
| chr12 | 70431072 | 70431089 | + | 2.00792115374673 | 1.13699574206438 | 1.19322821036799 | 1.06710864697714 |
| chr12 | 70542566 | 70542567 | + | 196.776273067179 | 660.594526139406 | 182.563916186302 | 460.990935494123 |
| chr12 | 71609759 | 71609780 | - | 19.0752509605939 | 18.1919318730301 | 14.3187385244159 | 17.0737383516342 |
| chr12 | 71609825 | 71609826 | - | 2.00792115374673 | 1.13699574206438 | 2.38645642073598 | 1.06710864697714 |
| chr12 | 71609862 | 71609866 | - | 5.01980288436682 | 1.13699574206438 | 1.19322821036799 | 1.06710864697714 |
| chr12 | 71701089 | 71701121 | + | 5.01980288436682 | 4.54798296825753 | 13.1255103140479 | 2.13421729395427 |
| chr12 | 71703845 | 71703864 | + | 3.01188173062009 | 1.13699574206438 | 1.19322821036799 | 1.06710864697714 |
| chr12 | 71785841 | 71785858 | + | 5.01980288436682 | 6.82197445238629 | 3.57968463110397 | 3.20132594093141 |
| chr12 | 71786189 | 71786197 | + | 20.0792115374673 | 87.5486721389574 | 5.96614105183995 | 9.60397782279422 |
| chr12 | 71786221 | 71786231 | + | 1.00396057687336 | 6.82197445238629 | 1.19322821036799 | 2.13421729395427 |
| chr12 | 71786263 | 71786279 | + | 2.00792115374673 | 3.41098722619315 | 1.19322821036799 | 2.13421729395427 |
| chr12 | 71786330 | 71786389 | + | 13.0514874993537 | 9.09596593651506 | 10.7390538933119 | 5.33554323488568 |
| chr12 | 71786463 | 71786481 | + | 3.01188173062009 | 1.13699574206438 | 1.19322821036799 | 5.33554323488568 |
| chr12 | 71786638 | 71786652 | + | 6.02376346124019 | 11.3699574206438 | 7.15936926220794 | 4.26843458790854 |
| chr12 | 71786699 | 71786719 | + | 8.03168461498692 | 15.9179403889013 | 9.54582568294392 | 18.1408469986113 |
| chr12 | 71786782 | 71786829 | + | 10.0396057687336 | 19.3289276150945 | 11.9322821036799 | 11.7381951167485 |
| chr12 | 71787158 | 71787199 | + | 6.02376346124019 | 5.68497871032191 | 4.77291284147196 | 4.26843458790854 |
| chr12 | 71787395 | 71787402 | + | 3.01188173062009 | 1.13699574206438 | 1.19322821036799 | 1.06710864697714 |
| chr12 | 71787726 | 71787757 | + | 7.02772403811355 | 3.41098722619315 | 5.96614105183995 | 10.6710864697714 |
| chr12 | 71787837 | 71787851 | + | 6.02376346124019 | 7.95897019445067 | 3.57968463110397 | 6.40265188186281 |
| chr12 | 71788193 | 71788196 | + | 2.00792115374673 | 1.13699574206438 | 4.77291284147196 | 1.06710864697714 |
| chr12 | 71788275 | 71788285 | + | 37.1465413443145 | 62.534765813541 | 70.4004644117114 | 29.8790421153598 |
| chr12 | 71788945 | 71788983 | + | 23.0910932680874 | 23.876910583352 | 17.8984231555198 | 18.1408469986113 |
| chr12 | 74538551 | 74538552 | + | 1.00396057687336 | 2.27399148412876 | 2.38645642073598 | 1.06710864697714 |
| chr12 | 74539236 | 74539255 | + | 2.00792115374673 | 5.68497871032191 | 4.77291284147196 | 2.13421729395427 |
| chr12 | 74539399 | 74539412 | + | 1.00396057687336 | 2.27399148412876 | 3.57968463110397 | 2.13421729395427 |
| chr12 | 74539465 | 74539497 | + | 1.00396057687336 | 3.41098722619315 | 5.96614105183995 | 1.06710864697714 |
| chr12 | 74539619 | 74539643 | + | 8.03168461498692 | 4.54798296825753 | 8.35259747257593 | 2.13421729395427 |
| chr12 | 74539816 | 74539833 | + | 3.01188173062009 | 3.41098722619315 | 4.77291284147196 | 3.20132594093141 |
| chr12 | 74539892 | 74539919 | + | 5.01980288436682 | 1.13699574206438 | 3.57968463110397 | 11.7381951167485 |
| chr12 | 74539955 | 74540068 | + | 13.0514874993537 | 15.9179403889013 | 17.8984231555198 | 34.1474767032683 |
| chr12 | 74540101 | 74540178 | + | 8.03168461498692 | 4.54798296825753 | 16.7051949451519 | 7.46976052883995 |
| chr12 | 74540239 | 74540275 | + | 5.01980288436682 | 6.82197445238629 | 5.96614105183995 | 5.33554323488568 |
| chr12 | 74540336 | 74540367 | + | 17.0673298068472 | 15.9179403889013 | 23.8645642073598 | 29.8790421153598 |
| chr12 | 74540416 | 74540442 | + | 7.02772403811355 | 9.09596593651506 | 9.54582568294392 | 6.40265188186281 |
| chr12 | 74540534 | 74540554 | + | 12.0475269224804 | 21.6029190992233 | 26.2510206280958 | 14.9395210576799 |
| chr12 | 74540591 | 74540606 | + | 3.01188173062009 | 3.41098722619315 | 3.57968463110397 | 2.13421729395427 |
| chr12 | 74540641 | 74540683 | + | 3.01188173062009 | 6.82197445238629 | 15.5119667347839 | 3.20132594093141 |
| chr12 | 74540712 | 74540715 | + | 1.00396057687336 | 1.13699574206438 | 2.38645642073598 | 4.26843458790854 |
| chr12 | 74540766 | 74540782 | + | 1.00396057687336 | 4.54798296825753 | 7.15936926220794 | 9.60397782279422 |
| chr12 | 74540819 | 74540867 | + | 10.0396057687336 | 5.68497871032191 | 5.96614105183995 | 7.46976052883995 |
| chr12 | 74540938 | 74540999 | + | 22.087132691214 | 28.4248935516096 | 51.3088130458235 | 62.959410171651 |
| chr12 | 74541019 | 74541024 | + | 3.01188173062009 | 7.95897019445067 | 8.35259747257593 | 17.0737383516342 |
| chr12 | 74541061 | 74541213 | + | 256.009947102708 | 309.262841841512 | 188.530057238142 | 212.35462074845 |
| chr12 | 74541234 | 74541365 | + | 619.443675930866 | 827.73290022287 | 1316.13071603589 | 1062.84021238923 |
| chr12 | 74541395 | 74541430 | + | 5.01980288436682 | 12.5069531627082 | 13.1255103140479 | 5.33554323488568 |
| chr12 | 75494487 | 75494490 | - | 6.02376346124019 | 3.41098722619315 | 3.57968463110397 | 3.20132594093141 |
| chr12 | 75497236 | 75497243 | - | 3.01188173062009 | 2.27399148412876 | 3.57968463110397 | 5.33554323488568 |
| chr12 | 75499214 | 75499217 | - | 1.00396057687336 | 1.13699574206438 | 1.19322821036799 | 2.13421729395427 |
| chr12 | 75499673 | 75499699 | - | 5.01980288436682 | 9.09596593651506 | 8.35259747257593 | 6.40265188186281 |
| chr12 | 75499727 | 75499771 | - | 6.02376346124019 | 5.68497871032191 | 4.77291284147196 | 4.26843458790854 |
| chr12 | 76028550 | 76028619 | - | 2.00792115374673 | 2.27399148412876 | 2.38645642073598 | 1.06710864697714 |
| chr12 | 76029573 | 76029679 | - | 18.0712903837206 | 34.1098722619315 | 7.15936926220794 | 9.60397782279422 |
| chr12 | 76029807 | 76029931 | - | 8.03168461498692 | 20.4659233571589 | 8.35259747257593 | 16.006629704657 |
| chr12 | 76029996 | 76030006 | - | 1.00396057687336 | 7.95897019445067 | 1.19322821036799 | 3.20132594093141 |
| chr12 | 76030580 | 76030652 | + | 13.0514874993537 | 11.3699574206438 | 7.15936926220794 | 5.33554323488568 |
| chr12 | 76046366 | 76046367 | - | 10.0396057687336 | 15.9179403889013 | 13.1255103140479 | 23.476390233497 |
| chr12 | 76046683 | 76046688 | - | 1.00396057687336 | 2.27399148412876 | 3.57968463110397 | 3.20132594093141 |
| chr12 | 76046762 | 76046790 | - | 8.03168461498692 | 4.54798296825753 | 7.15936926220794 | 21.3421729395427 |
| chr12 | 76046952 | 76046993 | - | 12.0475269224804 | 4.54798296825753 | 22.6713359969918 | 22.4092815865198 |
| chr12 | 76047042 | 76047047 | - | 5.01980288436682 | 2.27399148412876 | 2.38645642073598 | 4.26843458790854 |
| chr12 | 76047467 | 76047498 | - | 17.0673298068472 | 14.780944646837 | 48.9223566250876 | 41.6172372321083 |
| chr12 | 76047521 | 76047553 | - | 12.0475269224804 | 22.7399148412876 | 39.3765309421437 | 25.6106075274513 |
| chr12 | 76047620 | 76047645 | - | 8.03168461498692 | 12.5069531627082 | 9.54582568294392 | 6.40265188186281 |
| chr12 | 76047782 | 76047819 | - | 8.03168461498692 | 17.0549361309657 | 11.9322821036799 | 28.8119334683827 |
| chr12 | 76047883 | 76047900 | - | 4.01584230749346 | 2.27399148412876 | 5.96614105183995 | 4.26843458790854 |
| chr12 | 76048122 | 76048136 | - | 4.01584230749346 | 3.41098722619315 | 15.5119667347839 | 39.483019938154 |
| chr12 | 76048395 | 76048426 | - | 15.0594086531005 | 20.4659233571589 | 33.4103898903037 | 32.0132594093141 |
| chr12 | 76049199 | 76049211 | - | 3.01188173062009 | 1.13699574206438 | 2.38645642073598 | 8.53686917581709 |
| chr12 | 76049241 | 76049253 | - | 84.3326884573626 | 70.4937360079917 | 84.7192029361272 | 133.388580872142 |
| chr12 | 76049524 | 76049527 | - | 3.01188173062009 | 9.09596593651506 | 1.19322821036799 | 1.06710864697714 |
| chr12 | 76049585 | 76049663 | - | 25.0990144218341 | 28.4248935516096 | 82.3327465153913 | 22.4092815865198 |
| chr12 | 76049720 | 76049781 | - | 64.2534769198953 | 56.8497871032191 | 69.2072362013434 | 81.1002571702623 |
| chr12 | 76344844 | 76344854 | - | 8.03168461498692 | 5.68497871032191 | 9.54582568294392 | 11.7381951167485 |
| chr12 | 76352444 | 76352446 | - | 2.00792115374673 | 7.95897019445067 | 1.19322821036799 | 2.13421729395427 |
| chr12 | 76352515 | 76352517 | - | 4.01584230749346 | 2.27399148412876 | 3.57968463110397 | 17.0737383516342 |
| chr12 | 76352553 | 76352581 | - | 16.0633692299738 | 12.5069531627082 | 15.5119667347839 | 48.0198891139711 |
| chr12 | 76353018 | 76353021 | - | 1.00396057687336 | 2.27399148412876 | 1.19322821036799 | 1.06710864697714 |
| chr12 | 76353481 | 76353514 | - | 5.01980288436682 | 4.54798296825753 | 17.8984231555198 | 6.40265188186281 |
| chr12 | 76353942 | 76353982 | - | 7.02772403811355 | 3.41098722619315 | 2.38645642073598 | 6.40265188186281 |
| chr12 | 76354705 | 76354728 | - | 4.01584230749346 | 2.27399148412876 | 4.77291284147196 | 7.46976052883995 |
| chr12 | 76355110 | 76355146 | - | 2.00792115374673 | 2.27399148412876 | 4.77291284147196 | 3.20132594093141 |
| chr12 | 76355302 | 76355327 | - | 4.01584230749346 | 3.41098722619315 | 9.54582568294392 | 11.7381951167485 |
| chr12 | 76355825 | 76355845 | - | 2.00792115374673 | 2.27399148412876 | 2.38645642073598 | 3.20132594093141 |
| chr12 | 76852292 | 76852316 | + | 5.01980288436682 | 5.68497871032191 | 1.19322821036799 | 3.20132594093141 |
| chr12 | 76853487 | 76853507 | + | 3.01188173062009 | 7.95897019445067 | 7.15936926220794 | 10.6710864697714 |
| chr12 | 77021964 | 77021988 | - | 7.02772403811355 | 5.68497871032191 | 3.57968463110397 | 2.13421729395427 |
| chr12 | 77022132 | 77022138 | - | 1.00396057687336 | 2.27399148412876 | 4.77291284147196 | 4.26843458790854 |
| chr12 | 77022580 | 77022598 | - | 5.01980288436682 | 3.41098722619315 | 4.77291284147196 | 1.06710864697714 |
| chr12 | 77023553 | 77023574 | - | 1.00396057687336 | 3.41098722619315 | 1.19322821036799 | 3.20132594093141 |
| chr12 | 77023678 | 77023703 | - | 21.0831721143407 | 17.0549361309657 | 3.57968463110397 | 13.8724124107028 |
| chr12 | 79774048 | 79774116 | - | 3.01188173062009 | 3.41098722619315 | 2.38645642073598 | 2.13421729395427 |
| chr12 | 79774139 | 79774152 | - | 8.03168461498692 | 10.2329616785794 | 7.15936926220794 | 3.20132594093141 |
| chr12 | 79774176 | 79774195 | - | 1.00396057687336 | 1.13699574206438 | 1.19322821036799 | 1.06710864697714 |
| chr12 | 79774218 | 79774220 | - | 2.00792115374673 | 1.13699574206438 | 1.19322821036799 | 1.06710864697714 |
| chr12 | 79774336 | 79774356 | - | 1.00396057687336 | 4.54798296825753 | 3.57968463110397 | 5.33554323488568 |
| chr12 | 79775201 | 79775224 | - | 7.02772403811355 | 2.27399148412876 | 3.57968463110397 | 5.33554323488568 |
| chr12 | 79775746 | 79775770 | - | 9.03564519186028 | 15.9179403889013 | 16.7051949451519 | 18.1408469986113 |
| chr12 | 79775805 | 79775810 | - | 1.00396057687336 | 6.82197445238629 | 11.9322821036799 | 9.60397782279422 |
| chr12 | 79775836 | 79775883 | - | 7.02772403811355 | 11.3699574206438 | 23.8645642073598 | 22.4092815865198 |
| chr12 | 79935249 | 79935395 | + | 83.3287278804893 | 139.850476273919 | 298.307052591997 | 340.407658385706 |
| chr12 | 82353133 | 82353135 | - | 2.00792115374673 | 4.54798296825753 | 1.19322821036799 | 1.06710864697714 |
| chr12 | 88198183 | 88198193 | + | 5.01980288436682 | 10.2329616785794 | 3.57968463110397 | 1.06710864697714 |
| chr12 | 89419827 | 89419831 | - | 3.01188173062009 | 3.41098722619315 | 3.57968463110397 | 1.06710864697714 |
| chr12 | 89420029 | 89420031 | - | 2.00792115374673 | 3.41098722619315 | 1.19322821036799 | 3.20132594093141 |
| chr12 | 89420065 | 89420071 | - | 3.01188173062009 | 6.82197445238629 | 2.38645642073598 | 5.33554323488568 |
| chr12 | 89522856 | 89522886 | - | 3.01188173062009 | 1.13699574206438 | 2.38645642073598 | 1.06710864697714 |
| chr12 | 89589890 | 89589913 | - | 8.03168461498692 | 18.1919318730301 | 3.57968463110397 | 6.40265188186281 |
| chr12 | 89590488 | 89590530 | - | 12.0475269224804 | 20.4659233571589 | 13.1255103140479 | 6.40265188186281 |
| chr12 | 89590617 | 89590662 | - | 39.1544624980612 | 62.534765813541 | 22.6713359969918 | 38.4159112911769 |
| chr12 | 89590735 | 89590751 | - | 3.01188173062009 | 2.27399148412876 | 1.19322821036799 | 7.46976052883995 |
| chr12 | 89590780 | 89590805 | - | 69.2732798042621 | 64.8087572976698 | 36.9900745214077 | 44.8185631730397 |
| chr12 | 89590869 | 89590946 | - | 15.0594086531005 | 23.876910583352 | 10.7390538933119 | 7.46976052883995 |
| chr12 | 89590968 | 89591019 | - | 6.02376346124019 | 9.09596593651506 | 1.19322821036799 | 1.06710864697714 |
| chr12 | 89591136 | 89591164 | - | 6.02376346124019 | 11.3699574206438 | 3.57968463110397 | 2.13421729395427 |
| chr12 | 89591186 | 89591196 | - | 3.01188173062009 | 5.68497871032191 | 1.19322821036799 | 1.06710864697714 |
| chr12 | 92143174 | 92143181 | - | 18.0712903837206 | 42.0688424563821 | 9.54582568294392 | 8.53686917581709 |
| chr12 | 92143202 | 92143244 | - | 8.03168461498692 | 30.6988850357383 | 13.1255103140479 | 8.53686917581709 |
| chr12 | 92143345 | 92143359 | - | 5.01980288436682 | 15.9179403889013 | 4.77291284147196 | 7.46976052883995 |
| chr12 | 92143543 | 92143556 | - | 4.01584230749346 | 12.5069531627082 | 1.19322821036799 | 1.06710864697714 |
| chr12 | 92143813 | 92143838 | - | 2.00792115374673 | 2.27399148412876 | 1.19322821036799 | 2.13421729395427 |
| chr12 | 92146174 | 92146193 | + | 2.00792115374673 | 3.41098722619315 | 31.0239334695677 | 10.6710864697714 |
| chr12 | 92774081 | 92774101 | - | 1.00396057687336 | 5.68497871032191 | 5.96614105183995 | 2.13421729395427 |
| chr12 | 92775291 | 92775320 | - | 10.0396057687336 | 3.41098722619315 | 4.77291284147196 | 3.20132594093141 |
| chr12 | 93399789 | 93399795 | + | 2.00792115374673 | 2.27399148412876 | 2.38645642073598 | 6.40265188186281 |
| chr12 | 93401203 | 93401214 | + | 2.00792115374673 | 2.27399148412876 | 15.5119667347839 | 7.46976052883995 |
| chr12 | 93401622 | 93401637 | + | 1.00396057687336 | 1.13699574206438 | 22.6713359969918 | 10.6710864697714 |
| chr12 | 93402393 | 93402395 | + | 4.01584230749346 | 9.09596593651506 | 29.8307052591997 | 55.4896496428111 |
| chr12 | 93402420 | 93402460 | + | 1.00396057687336 | 2.27399148412876 | 8.35259747257593 | 6.40265188186281 |
| chr12 | 93402555 | 93402558 | + | 14.0554480762271 | 21.6029190992233 | 84.7192029361272 | 73.6304966414224 |
| chr12 | 93402581 | 93402621 | + | 6.02376346124019 | 14.780944646837 | 81.1395183050233 | 48.0198891139711 |
| chr12 | 93408401 | 93408408 | - | 4.01584230749346 | 1.13699574206438 | 2.38645642073598 | 2.13421729395427 |
| chr12 | 93408433 | 93408452 | - | 2.00792115374673 | 12.5069531627082 | 7.15936926220794 | 3.20132594093141 |
| chr12 | 93409389 | 93409406 | - | 2.00792115374673 | 4.54798296825753 | 5.96614105183995 | 1.06710864697714 |
| chr12 | 93409459 | 93409468 | - | 1.00396057687336 | 3.41098722619315 | 3.57968463110397 | 6.40265188186281 |
| chr12 | 93409642 | 93409667 | - | 35.1386201905678 | 32.9728765198671 | 25.0577924177278 | 64.0265188186281 |
| chr12 | 93409706 | 93409735 | - | 6.02376346124019 | 7.95897019445067 | 9.54582568294392 | 33.0803680562912 |
| chr12 | 93501215 | 93501320 | + | 45.1782259593014 | 51.1648083928972 | 47.7291284147196 | 66.1607361125824 |
| chr12 | 93501347 | 93501396 | + | 9.03564519186028 | 17.0549361309657 | 14.3187385244159 | 23.476390233497 |
| chr12 | 93501438 | 93501445 | + | 4.01584230749346 | 2.27399148412876 | 4.77291284147196 | 3.20132594093141 |
| chr12 | 93501542 | 93501549 | + | 2.00792115374673 | 3.41098722619315 | 3.57968463110397 | 3.20132594093141 |
| chr12 | 93501737 | 93501792 | + | 6.02376346124019 | 2.27399148412876 | 15.5119667347839 | 5.33554323488568 |
| chr12 | 93501968 | 93502160 | + | 160.633692299738 | 140.987472015983 | 241.032098494334 | 344.676092973615 |
| chr12 | 93502188 | 93502262 | + | 23.0910932680874 | 35.2468680039958 | 36.9900745214077 | 32.0132594093141 |
| chr12 | 93502343 | 93502369 | + | 52.205949997415 | 65.9457530397342 | 103.810854302015 | 81.1002571702623 |
| chr12 | 93503211 | 93503229 | + | 8.03168461498692 | 4.54798296825753 | 2.38645642073598 | 1.06710864697714 |
| chr12 | 93503341 | 93503372 | + | 2.00792115374673 | 1.13699574206438 | 8.35259747257593 | 12.8053037637256 |
| chr12 | 93503674 | 93503680 | + | 8.03168461498692 | 1.13699574206438 | 11.9322821036799 | 10.6710864697714 |
| chr12 | 93503842 | 93503843 | + | 5.01980288436682 | 1.13699574206438 | 1.19322821036799 | 2.13421729395427 |
| chr12 | 93505474 | 93505476 | + | 2.00792115374673 | 3.41098722619315 | 1.19322821036799 | 5.33554323488568 |
| chr12 | 93510099 | 93510133 | + | 4.01584230749346 | 4.54798296825753 | 5.96614105183995 | 2.13421729395427 |
| chr12 | 93575509 | 93575567 | + | 2.00792115374673 | 4.54798296825753 | 65.6275515702394 | 45.8856718200168 |
| chr12 | 93575767 | 93575816 | + | 1.00396057687336 | 5.68497871032191 | 72.7869208324474 | 99.2411041688736 |
| chr12 | 93575843 | 93575854 | + | 2.00792115374673 | 10.2329616785794 | 54.8884976769275 | 78.966039876308 |
| chr12 | 93575885 | 93575981 | + | 3.01188173062009 | 23.876910583352 | 115.743136405695 | 98.1739955218965 |
| chr12 | 93576046 | 93576084 | + | 5.01980288436682 | 7.95897019445067 | 114.549908195327 | 75.7647139353766 |
| chr12 | 93576114 | 93576122 | + | 54.2138711511617 | 68.2197445238629 | 1405.62283181349 | 962.531999573376 |
| chr12 | 93850293 | 93850346 | + | 1.00396057687336 | 2.27399148412876 | 14.3187385244159 | 23.476390233497 |
| chr12 | 94308151 | 94308155 | - | 4.01584230749346 | 3.41098722619315 | 5.96614105183995 | 8.53686917581709 |
| chr12 | 94308782 | 94308783 | - | 5.01980288436682 | 2.27399148412876 | 14.3187385244159 | 7.46976052883995 |
| chr12 | 94971363 | 94971433 | - | 35.1386201905678 | 67.0827487817985 | 46.5359002043516 | 67.2278447595595 |
| chr12 | 94971465 | 94971473 | - | 39.1544624980612 | 36.3838637460602 | 51.3088130458235 | 70.429170700491 |
| chr12 | 94971545 | 94971551 | - | 2.00792115374673 | 1.13699574206438 | 2.38645642073598 | 7.46976052883995 |
| chr12 | 95021936 | 95021988 | - | 1.00396057687336 | 3.41098722619315 | 7.15936926220794 | 4.26843458790854 |
| chr12 | 95078078 | 95078079 | - | 8.03168461498692 | 2.27399148412876 | 1.19322821036799 | 2.13421729395427 |
| chr12 | 95300166 | 95300196 | + | 2.00792115374673 | 7.95897019445067 | 2.38645642073598 | 8.53686917581709 |
| chr12 | 95300241 | 95300271 | + | 3.01188173062009 | 12.5069531627082 | 9.54582568294392 | 1.06710864697714 |
| chr12 | 95300295 | 95300327 | + | 7.02772403811355 | 5.68497871032191 | 1.19322821036799 | 2.13421729395427 |
| chr12 | 95300600 | 95300626 | + | 61.2415951892752 | 109.151591238181 | 34.6036181006717 | 58.6909755837425 |
| chr12 | 95300655 | 95300665 | + | 2.00792115374673 | 5.68497871032191 | 1.19322821036799 | 4.26843458790854 |
| chr12 | 95300687 | 95300698 | + | 2.00792115374673 | 11.3699574206438 | 2.38645642073598 | 9.60397782279422 |
| chr12 | 95300720 | 95300754 | + | 35.1386201905678 | 71.6307317500561 | 14.3187385244159 | 16.006629704657 |
| chr12 | 95300780 | 95300806 | + | 19.0752509605939 | 30.6988850357383 | 13.1255103140479 | 7.46976052883995 |
| chr12 | 95300870 | 95300876 | + | 8.03168461498692 | 12.5069531627082 | 7.15936926220794 | 3.20132594093141 |
| chr12 | 95301000 | 95301007 | + | 4.01584230749346 | 3.41098722619315 | 4.77291284147196 | 3.20132594093141 |
| chr12 | 95308499 | 95308501 | + | 5.01980288436682 | 1.13699574206438 | 2.38645642073598 | 2.13421729395427 |
| chr12 | 95309968 | 95309981 | + | 5.01980288436682 | 5.68497871032191 | 15.5119667347839 | 5.33554323488568 |
| chr12 | 95514081 | 95514286 | + | 24.0950538449607 | 40.9318467143178 | 45.3426719939836 | 19.2079556455884 |
| chr12 | 95515455 | 95515456 | + | 2.00792115374673 | 2.27399148412876 | 2.38645642073598 | 2.13421729395427 |
| chr12 | 95866029 | 95866088 | + | 49.1940682667949 | 101.19262104373 | 100.231169670911 | 130.187254931211 |
| chr12 | 96000963 | 96001052 | - | 57.2257528817818 | 60.2607743294122 | 34.6036181006717 | 23.476390233497 |
| chr12 | 96002959 | 96003057 | - | 2.00792115374673 | 7.95897019445067 | 2.38645642073598 | 4.26843458790854 |
| chr12 | 96278498 | 96278520 | - | 3.01188173062009 | 13.6439489047726 | 3.57968463110397 | 6.40265188186281 |
| chr12 | 96278730 | 96278768 | - | 32.1267384599477 | 64.8087572976698 | 70.4004644117114 | 32.0132594093141 |
| chr12 | 96278798 | 96278850 | - | 64.2534769198953 | 179.645327246172 | 91.8785721983352 | 62.959410171651 |
| chr12 | 96278876 | 96278888 | - | 3.01188173062009 | 6.82197445238629 | 5.96614105183995 | 5.33554323488568 |
| chr12 | 96278918 | 96278943 | - | 105.415860571703 | 213.755199508104 | 46.5359002043516 | 76.8318225823538 |
| chr12 | 96279017 | 96279039 | - | 11.043566345607 | 15.9179403889013 | 5.96614105183995 | 5.33554323488568 |
| chr12 | 96279067 | 96279085 | - | 1.00396057687336 | 2.27399148412876 | 8.35259747257593 | 3.20132594093141 |
| chr12 | 96279116 | 96279124 | - | 15.0594086531005 | 43.2058381984465 | 7.15936926220794 | 14.9395210576799 |
| chr12 | 96279198 | 96279200 | - | 10.0396057687336 | 9.09596593651506 | 1.19322821036799 | 4.26843458790854 |
| chr12 | 96279258 | 96279272 | - | 1.00396057687336 | 1.13699574206438 | 1.19322821036799 | 1.06710864697714 |
| chr12 | 96279321 | 96279362 | - | 8.03168461498692 | 14.780944646837 | 20.2848795762558 | 5.33554323488568 |
| chr12 | 96279408 | 96279429 | - | 3.01188173062009 | 4.54798296825753 | 3.57968463110397 | 1.06710864697714 |
| chr12 | 96279850 | 96279875 | - | 7.02772403811355 | 10.2329616785794 | 3.57968463110397 | 4.26843458790854 |
| chr12 | 96279995 | 96280013 | - | 2.00792115374673 | 4.54798296825753 | 2.38645642073598 | 2.13421729395427 |
| chr12 | 96280036 | 96280051 | - | 2.00792115374673 | 5.68497871032191 | 3.57968463110397 | 3.20132594093141 |
| chr12 | 96280155 | 96280180 | - | 2.00792115374673 | 3.41098722619315 | 8.35259747257593 | 3.20132594093141 |
| chr12 | 96952834 | 96952836 | + | 3.01188173062009 | 1.13699574206438 | 2.38645642073598 | 2.13421729395427 |
| chr12 | 96953095 | 96953100 | + | 10.0396057687336 | 14.780944646837 | 11.9322821036799 | 20.2750642925656 |
| chr12 | 96953589 | 96953590 | + | 1.00396057687336 | 4.54798296825753 | 1.19322821036799 | 1.06710864697714 |
| chr12 | 98535431 | 98535442 | + | 4.01584230749346 | 4.54798296825753 | 2.38645642073598 | 2.13421729395427 |
| chr12 | 98535463 | 98535504 | + | 8.03168461498692 | 5.68497871032191 | 8.35259747257593 | 5.33554323488568 |
| chr12 | 98535558 | 98535562 | + | 11.043566345607 | 9.09596593651506 | 4.77291284147196 | 9.60397782279422 |
| chr12 | 98535587 | 98535595 | + | 2.00792115374673 | 3.41098722619315 | 1.19322821036799 | 1.06710864697714 |
| chr12 | 98547832 | 98547887 | + | 29.1148567293276 | 14.780944646837 | 29.8307052591997 | 20.2750642925656 |
| chr12 | 98547920 | 98548016 | + | 17.0673298068472 | 27.2878978095452 | 40.5697591525116 | 28.8119334683827 |
| chr12 | 98548067 | 98548129 | + | 19.0752509605939 | 31.8358807778027 | 48.9223566250876 | 34.1474767032683 |
| chr12 | 98548156 | 98548206 | + | 18.0712903837206 | 17.0549361309657 | 19.0916513658878 | 12.8053037637256 |
| chr12 | 98548498 | 98548523 | + | 2.00792115374673 | 3.41098722619315 | 5.96614105183995 | 13.8724124107028 |
| chr12 | 98549437 | 98549442 | + | 8.03168461498692 | 3.41098722619315 | 3.57968463110397 | 12.8053037637256 |
| chr12 | 98549485 | 98549505 | + | 10.0396057687336 | 13.6439489047726 | 9.54582568294392 | 3.20132594093141 |
| chr12 | 98549942 | 98549961 | + | 168.665376914725 | 152.357429436627 | 53.6952694665595 | 150.462319223776 |
| chr12 | 98550062 | 98550066 | + | 3.01188173062009 | 3.41098722619315 | 8.35259747257593 | 5.33554323488568 |
| chr12 | 98550116 | 98550151 | + | 16.0633692299738 | 26.1509020674808 | 22.6713359969918 | 17.0737383516342 |
| chr12 | 98550171 | 98550191 | + | 5.01980288436682 | 3.41098722619315 | 3.57968463110397 | 4.26843458790854 |
| chr12 | 98601368 | 98601593 | + | 81.3208067267425 | 101.19262104373 | 65.6275515702394 | 75.7647139353766 |
| chr12 | 98601618 | 98601687 | + | 53.2099105742883 | 84.1376849127643 | 33.4103898903037 | 20.2750642925656 |
| chr12 | 98624377 | 98624396 | - | 4.01584230749346 | 18.1919318730301 | 7.15936926220794 | 11.7381951167485 |
| chr12 | 98624417 | 98624447 | - | 10.0396057687336 | 1.13699574206438 | 1.19322821036799 | 5.33554323488568 |
| chr12 | 98624775 | 98624777 | - | 1.00396057687336 | 3.41098722619315 | 1.19322821036799 | 2.13421729395427 |
| chr12 | 98625594 | 98625596 | - | 7.02772403811355 | 3.41098722619315 | 3.57968463110397 | 3.20132594093141 |
| chr12 | 98735163 | 98735184 | + | 5.01980288436682 | 5.68497871032191 | 5.96614105183995 | 8.53686917581709 |
| chr12 | 100224152 | 100224159 | + | 5.01980288436682 | 9.09596593651506 | 1.19322821036799 | 2.13421729395427 |
| chr12 | 100224387 | 100224394 | + | 8.03168461498692 | 17.0549361309657 | 27.4442488384638 | 36.2816939972226 |
| chr12 | 100339435 | 100339439 | + | 2.00792115374673 | 4.54798296825753 | 1.19322821036799 | 1.06710864697714 |
| chr12 | 100339487 | 100339529 | + | 54.2138711511617 | 63.6717615556054 | 44.1494437836156 | 65.0936274656053 |
| chr12 | 100339557 | 100339578 | + | 7.02772403811355 | 6.82197445238629 | 13.1255103140479 | 5.33554323488568 |
| chr12 | 100339677 | 100339680 | + | 3.01188173062009 | 1.13699574206438 | 2.38645642073598 | 2.13421729395427 |
| chr12 | 100339729 | 100339734 | + | 4.01584230749346 | 5.68497871032191 | 2.38645642073598 | 4.26843458790854 |
| chr12 | 100339836 | 100339844 | + | 1.00396057687336 | 4.54798296825753 | 1.19322821036799 | 2.13421729395427 |
| chr12 | 100339873 | 100339888 | + | 5.01980288436682 | 3.41098722619315 | 8.35259747257593 | 4.26843458790854 |
| chr12 | 100339910 | 100339971 | + | 59.2336740355285 | 77.315710460378 | 150.346754506367 | 84.3015831111937 |
| chr12 | 100340014 | 100340033 | + | 41.1623836518079 | 44.3428339405109 | 53.6952694665595 | 27.7448248214055 |
| chr12 | 100341680 | 100341682 | + | 4.01584230749346 | 10.2329616785794 | 4.77291284147196 | 2.13421729395427 |
| chr12 | 101386493 | 101386508 | + | 10.0396057687336 | 3.41098722619315 | 4.77291284147196 | 4.26843458790854 |
| chr12 | 101393229 | 101393238 | - | 3.01188173062009 | 2.27399148412876 | 2.38645642073598 | 2.13421729395427 |
| chr12 | 101394884 | 101394911 | - | 2.00792115374673 | 2.27399148412876 | 7.15936926220794 | 3.20132594093141 |
| chr12 | 101394928 | 101394957 | - | 1.00396057687336 | 2.27399148412876 | 1.19322821036799 | 2.13421729395427 |
| chr12 | 101395198 | 101395217 | - | 1.00396057687336 | 1.13699574206438 | 3.57968463110397 | 2.13421729395427 |
| chr12 | 101395470 | 101395497 | - | 21.0831721143407 | 18.1919318730301 | 16.7051949451519 | 12.8053037637256 |
| chr12 | 101726291 | 101726498 | + | 74.293082688629 | 90.9596593651506 | 242.225326704702 | 145.12677598889 |
| chr12 | 101728874 | 101728956 | + | 58.2297134586551 | 60.2607743294122 | 145.573841664895 | 72.5633879944452 |
| chr12 | 101728979 | 101729055 | + | 4.01584230749346 | 4.54798296825753 | 14.3187385244159 | 17.0737383516342 |
| chr12 | 102012988 | 102013010 | - | 5.01980288436682 | 9.09596593651506 | 2.38645642073598 | 4.26843458790854 |
| chr12 | 102074234 | 102074252 | - | 1.00396057687336 | 2.27399148412876 | 1.19322821036799 | 1.06710864697714 |
| chr12 | 102197353 | 102197356 | + | 1.00396057687336 | 4.54798296825753 | 2.38645642073598 | 2.13421729395427 |
| chr12 | 103841443 | 103841494 | + | 2.00792115374673 | 1.13699574206438 | 9.54582568294392 | 26.6777161744284 |
| chr12 | 103930312 | 103930531 | + | 23131.2516911623 | 12999.2723190221 | 18290.9952367309 | 30274.9394233883 |
| chr12 | 103930553 | 103930578 | + | 2.00792115374673 | 4.54798296825753 | 11.9322821036799 | 4.26843458790854 |
| chr12 | 103947307 | 103947416 | + | 41.1623836518079 | 35.2468680039958 | 140.800928823423 | 186.744013220999 |
| chr12 | 103947660 | 103947819 | + | 99.3920971104631 | 83.0006891706999 | 109.776995353855 | 112.046407932599 |
| chr12 | 103947873 | 103947903 | + | 39.1544624980612 | 52.3018041349616 | 33.4103898903037 | 6.40265188186281 |
| chr12 | 103951404 | 103951413 | - | 5.01980288436682 | 2.27399148412876 | 7.15936926220794 | 4.26843458790854 |
| chr12 | 103951534 | 103951538 | - | 2.00792115374673 | 3.41098722619315 | 1.19322821036799 | 2.13421729395427 |
| chr12 | 103987067 | 103987091 | + | 3.01188173062009 | 1.13699574206438 | 2.38645642073598 | 12.8053037637256 |
| chr12 | 103988668 | 103988669 | + | 2.00792115374673 | 2.27399148412876 | 1.19322821036799 | 1.06710864697714 |
| chr12 | 104118087 | 104118109 | - | 17.0673298068472 | 35.2468680039958 | 46.5359002043516 | 22.4092815865198 |
| chr12 | 104118152 | 104118164 | - | 12.0475269224804 | 21.6029190992233 | 21.4781077866238 | 6.40265188186281 |
| chr12 | 104118278 | 104118283 | - | 48.1901076899215 | 108.014595496116 | 68.0140079909754 | 96.0397782279422 |
| chr12 | 104119370 | 104119378 | - | 7.02772403811355 | 5.68497871032191 | 2.38645642073598 | 2.13421729395427 |
| chr12 | 104348682 | 104348701 | + | 22.087132691214 | 13.6439489047726 | 2.38645642073598 | 2.13421729395427 |
| chr12 | 104348728 | 104348781 | + | 22.087132691214 | 13.6439489047726 | 10.7390538933119 | 32.0132594093141 |
| chr12 | 104348801 | 104348811 | + | 4.01584230749346 | 3.41098722619315 | 1.19322821036799 | 1.06710864697714 |
| chr12 | 104348863 | 104348950 | + | 45.1782259593014 | 18.1919318730301 | 10.7390538933119 | 11.7381951167485 |
| chr12 | 104348967 | 104348981 | + | 7.02772403811355 | 2.27399148412876 | 2.38645642073598 | 4.26843458790854 |
| chr12 | 104349078 | 104349087 | + | 9.03564519186028 | 1.13699574206438 | 7.15936926220794 | 7.46976052883995 |
| chr12 | 104349112 | 104349117 | + | 4.01584230749346 | 2.27399148412876 | 1.19322821036799 | 1.06710864697714 |
| chr12 | 104349190 | 104349217 | + | 48.1901076899215 | 22.7399148412876 | 5.96614105183995 | 9.60397782279422 |
| chr12 | 104349313 | 104349320 | + | 1.00396057687336 | 6.82197445238629 | 1.19322821036799 | 2.13421729395427 |
| chr12 | 104349353 | 104349358 | + | 6.02376346124019 | 3.41098722619315 | 2.38645642073598 | 7.46976052883995 |
| chr12 | 104349385 | 104349390 | + | 3.01188173062009 | 2.27399148412876 | 3.57968463110397 | 1.06710864697714 |
| chr12 | 104349490 | 104349504 | + | 12.0475269224804 | 4.54798296825753 | 2.38645642073598 | 3.20132594093141 |
| chr12 | 104349699 | 104349729 | + | 14.0554480762271 | 6.82197445238629 | 1.19322821036799 | 3.20132594093141 |
| chr12 | 104349897 | 104349901 | + | 3.01188173062009 | 2.27399148412876 | 2.38645642073598 | 2.13421729395427 |
| chr12 | 104349925 | 104349986 | + | 130.514874993537 | 70.4937360079917 | 26.2510206280958 | 55.4896496428111 |
| chr12 | 104350059 | 104350139 | + | 340.342635560071 | 280.837948289902 | 60.8546387287675 | 99.2411041688736 |
| chr12 | 104759481 | 104759484 | + | 16.0633692299738 | 14.780944646837 | 466.552230253884 | 348.944527561523 |
| chr12 | 104994640 | 104994658 | + | 3.01188173062009 | 15.9179403889013 | 10.7390538933119 | 17.0737383516342 |
| chr12 | 104995983 | 104995985 | + | 1.00396057687336 | 4.54798296825753 | 1.19322821036799 | 1.06710864697714 |
| chr12 | 105167197 | 105167200 | + | 2.00792115374673 | 6.82197445238629 | 1.19322821036799 | 2.13421729395427 |
| chr12 | 105173591 | 105173592 | - | 1.00396057687336 | 5.68497871032191 | 1.19322821036799 | 2.13421729395427 |
| chr12 | 106971136 | 106971250 | + | 31.1227778830743 | 40.9318467143178 | 76.3666054635513 | 37.3488026441998 |
| chr12 | 106971503 | 106971521 | + | 3.01188173062009 | 3.41098722619315 | 2.38645642073598 | 2.13421729395427 |
| chr12 | 106971623 | 106971655 | + | 4.01584230749346 | 4.54798296825753 | 2.38645642073598 | 1.06710864697714 |
| chr12 | 106971766 | 106971821 | + | 9.03564519186028 | 9.09596593651506 | 15.5119667347839 | 9.60397782279422 |
| chr12 | 106971844 | 106971919 | + | 6.02376346124019 | 12.5069531627082 | 20.2848795762558 | 7.46976052883995 |
| chr12 | 106972024 | 106972063 | + | 2.00792115374673 | 1.13699574206438 | 7.15936926220794 | 8.53686917581709 |
| chr12 | 106972605 | 106972670 | + | 3.01188173062009 | 14.780944646837 | 19.0916513658878 | 7.46976052883995 |
| chr12 | 106972797 | 106972805 | + | 5.01980288436682 | 7.95897019445067 | 1.19322821036799 | 8.53686917581709 |
| chr12 | 106972879 | 106972993 | + | 40.1584230749346 | 36.3838637460602 | 82.3327465153913 | 61.8923015246739 |
| chr12 | 106973008 | 106973026 | + | 1.00396057687336 | 2.27399148412876 | 3.57968463110397 | 8.53686917581709 |
| chr12 | 106973124 | 106973141 | + | 2.00792115374673 | 2.27399148412876 | 3.57968463110397 | 2.13421729395427 |
| chr12 | 106973163 | 106973187 | + | 5.01980288436682 | 3.41098722619315 | 3.57968463110397 | 4.26843458790854 |
| chr12 | 106973227 | 106973235 | + | 2.00792115374673 | 1.13699574206438 | 1.19322821036799 | 3.20132594093141 |
| chr12 | 106973271 | 106973322 | + | 11.043566345607 | 6.82197445238629 | 10.7390538933119 | 5.33554323488568 |
| chr12 | 106973341 | 106973421 | + | 3.01188173062009 | 3.41098722619315 | 9.54582568294392 | 8.53686917581709 |
| chr12 | 106973552 | 106973569 | + | 1.00396057687336 | 2.27399148412876 | 1.19322821036799 | 2.13421729395427 |
| chr12 | 106973711 | 106973770 | + | 3.01188173062009 | 1.13699574206438 | 7.15936926220794 | 4.26843458790854 |
| chr12 | 106973799 | 106973849 | + | 21.0831721143407 | 18.1919318730301 | 14.3187385244159 | 12.8053037637256 |
| chr12 | 106973933 | 106973954 | + | 21.0831721143407 | 59.1237785873479 | 57.2749540976635 | 27.7448248214055 |
| chr12 | 106973985 | 106973990 | + | 8.03168461498692 | 10.2329616785794 | 9.54582568294392 | 7.46976052883995 |
| chr12 | 106991428 | 106991454 | - | 7.02772403811355 | 13.6439489047726 | 15.5119667347839 | 7.46976052883995 |
| chr12 | 106991958 | 106991963 | - | 7.02772403811355 | 7.95897019445067 | 5.96614105183995 | 2.13421729395427 |
| chr12 | 107712115 | 107712227 | + | 93.3683336492229 | 70.4937360079917 | 50.1155848354556 | 87.5029090521251 |
| chr12 | 107712356 | 107712379 | + | 11.043566345607 | 7.95897019445067 | 5.96614105183995 | 19.2079556455884 |
| chr12 | 107712399 | 107712425 | + | 17.0673298068472 | 35.2468680039958 | 19.0916513658878 | 20.2750642925656 |
| chr12 | 107733328 | 107733344 | - | 2.00792115374673 | 2.27399148412876 | 1.19322821036799 | 2.13421729395427 |
| chr12 | 108522627 | 108522630 | - | 6.02376346124019 | 4.54798296825753 | 7.15936926220794 | 4.26843458790854 |
| chr12 | 108522867 | 108522882 | - | 4.01584230749346 | 1.13699574206438 | 9.54582568294392 | 8.53686917581709 |
| chr12 | 108522903 | 108522913 | - | 5.01980288436682 | 3.41098722619315 | 11.9322821036799 | 6.40265188186281 |
| chr12 | 108522925 | 108523043 | - | 350.382241328804 | 320.632799262156 | 453.426719939836 | 509.010824608094 |
| chr12 | 108523106 | 108523118 | - | 4.01584230749346 | 2.27399148412876 | 1.19322821036799 | 2.13421729395427 |
| chr12 | 108523179 | 108523185 | - | 1.00396057687336 | 1.13699574206438 | 3.57968463110397 | 2.13421729395427 |
| chr12 | 108523214 | 108523238 | - | 7.02772403811355 | 4.54798296825753 | 14.3187385244159 | 8.53686917581709 |
| chr12 | 108523306 | 108523324 | - | 13.0514874993537 | 5.68497871032191 | 8.35259747257593 | 11.7381951167485 |
| chr12 | 108568831 | 108568943 | + | 34.1346596136944 | 38.657855230189 | 32.2171616799357 | 59.7580842307196 |
| chr12 | 108569184 | 108569218 | + | 1.00396057687336 | 1.13699574206438 | 4.77291284147196 | 3.20132594093141 |
| chr12 | 108569252 | 108569297 | + | 63.249516343022 | 81.8636934286355 | 51.3088130458235 | 43.7514545260626 |
| chr12 | 108569318 | 108569322 | + | 2.00792115374673 | 1.13699574206438 | 3.57968463110397 | 2.13421729395427 |
| chr12 | 108645131 | 108645211 | - | 14.0554480762271 | 6.82197445238629 | 25.0577924177278 | 34.1474767032683 |
| chr12 | 108645266 | 108645293 | - | 2.00792115374673 | 9.09596593651506 | 13.1255103140479 | 16.006629704657 |
| chr12 | 108645360 | 108645379 | - | 7.02772403811355 | 2.27399148412876 | 7.15936926220794 | 13.8724124107028 |
| chr12 | 108645523 | 108645986 | - | 228.903011527127 | 115.973565690567 | 791.110303473977 | 682.949534065367 |
| chr12 | 108646112 | 108646191 | - | 26.1029749987075 | 22.7399148412876 | 64.4343233598714 | 51.2212150549025 |
| chr12 | 108646219 | 108646232 | - | 1.00396057687336 | 1.13699574206438 | 10.7390538933119 | 5.33554323488568 |
| chr12 | 108646295 | 108646304 | - | 13.0514874993537 | 4.54798296825753 | 36.9900745214077 | 10.6710864697714 |
| chr12 | 108646345 | 108646420 | - | 29.1148567293276 | 5.68497871032191 | 65.6275515702394 | 117.381951167485 |
| chr12 | 108646473 | 108646504 | - | 5.01980288436682 | 3.41098722619315 | 21.4781077866238 | 11.7381951167485 |
| chr12 | 108646524 | 108646564 | - | 6.02376346124019 | 4.54798296825753 | 8.35259747257593 | 18.1408469986113 |
| chr12 | 108646589 | 108646645 | - | 23.0910932680874 | 13.6439489047726 | 53.6952694665595 | 37.3488026441998 |
| chr12 | 108646675 | 108646707 | - | 15.0594086531005 | 9.09596593651506 | 63.2410951495034 | 90.7042349930565 |
| chr12 | 108646730 | 108646834 | - | 10.0396057687336 | 4.54798296825753 | 44.1494437836156 | 50.1541064079254 |
| chr12 | 108646857 | 108646956 | - | 15.0594086531005 | 7.95897019445067 | 75.1733772531833 | 22.4092815865198 |
| chr12 | 108646989 | 108647080 | - | 39.1544624980612 | 22.7399148412876 | 73.9801490428153 | 94.9726695809651 |
| chr12 | 108647141 | 108647221 | - | 8.03168461498692 | 6.82197445238629 | 31.0239334695677 | 22.4092815865198 |
| chr12 | 108647243 | 108647268 | - | 2.00792115374673 | 2.27399148412876 | 11.9322821036799 | 6.40265188186281 |
| chr12 | 108647295 | 108647374 | - | 24.0950538449607 | 14.780944646837 | 59.6614105183995 | 44.8185631730397 |
| chr12 | 108647401 | 108647462 | - | 7.02772403811355 | 6.82197445238629 | 20.2848795762558 | 27.7448248214055 |
| chr12 | 109086251 | 109086265 | + | 1.00396057687336 | 1.13699574206438 | 3.57968463110397 | 1.06710864697714 |
| chr12 | 109087695 | 109087719 | + | 18.0712903837206 | 12.5069531627082 | 42.9562155732476 | 30.9461507623369 |
| chr12 | 109087757 | 109087797 | + | 8.03168461498692 | 9.09596593651506 | 8.35259747257593 | 12.8053037637256 |
| chr12 | 109087892 | 109087903 | + | 7.02772403811355 | 12.5069531627082 | 5.96614105183995 | 2.13421729395427 |
| chr12 | 109088231 | 109088264 | - | 60.2376346124019 | 131.891506079468 | 39.3765309421437 | 56.5567582897882 |
| chr12 | 109110075 | 109110106 | + | 2.00792115374673 | 2.27399148412876 | 3.57968463110397 | 1.06710864697714 |
| chr12 | 109110199 | 109110235 | + | 7.02772403811355 | 7.95897019445067 | 17.8984231555198 | 17.0737383516342 |
| chr12 | 109110258 | 109110274 | + | 2.00792115374673 | 6.82197445238629 | 1.19322821036799 | 4.26843458790854 |
| chr12 | 109110644 | 109110718 | + | 11.043566345607 | 27.2878978095452 | 11.9322821036799 | 12.8053037637256 |
| chr12 | 109110940 | 109110966 | + | 22.087132691214 | 31.8358807778027 | 13.1255103140479 | 13.8724124107028 |
| chr12 | 109450161 | 109450261 | - | 28.1108961524542 | 11.3699574206438 | 23.8645642073598 | 32.0132594093141 |
| chr12 | 109450419 | 109450437 | - | 2.00792115374673 | 1.13699574206438 | 3.57968463110397 | 1.06710864697714 |
| chr12 | 109450441 | 109450480 | - | 10.0396057687336 | 2.27399148412876 | 7.15936926220794 | 2.13421729395427 |
| chr12 | 109450620 | 109450626 | - | 3.01188173062009 | 1.13699574206438 | 3.57968463110397 | 14.9395210576799 |
| chr12 | 109535050 | 109535062 | + | 3.01188173062009 | 10.2329616785794 | 14.3187385244159 | 8.53686917581709 |
| chr12 | 109536477 | 109536500 | + | 14.0554480762271 | 26.1509020674808 | 11.9322821036799 | 21.3421729395427 |
| chr12 | 109556719 | 109556736 | - | 14.0554480762271 | 13.6439489047726 | 11.9322821036799 | 8.53686917581709 |
| chr12 | 109586019 | 109586066 | + | 7.02772403811355 | 2.27399148412876 | 9.54582568294392 | 14.9395210576799 |
| chr12 | 109596918 | 109597023 | + | 6.02376346124019 | 2.27399148412876 | 28.6374770488317 | 38.4159112911769 |
| chr12 | 109597120 | 109597192 | + | 6.02376346124019 | 14.780944646837 | 5.96614105183995 | 20.2750642925656 |
| chr12 | 109597213 | 109597240 | + | 2.00792115374673 | 21.6029190992233 | 16.7051949451519 | 9.60397782279422 |
| chr12 | 109852666 | 109852684 | - | 4.01584230749346 | 5.68497871032191 | 2.38645642073598 | 1.06710864697714 |
| chr12 | 109855714 | 109855724 | - | 9.03564519186028 | 1.13699574206438 | 1.19322821036799 | 2.13421729395427 |
| chr12 | 109918036 | 109918043 | + | 1.00396057687336 | 2.27399148412876 | 3.57968463110397 | 1.06710864697714 |
| chr12 | 110037775 | 110037787 | + | 1.00396057687336 | 2.27399148412876 | 1.19322821036799 | 3.20132594093141 |
| chr12 | 110039281 | 110039356 | + | 4.01584230749346 | 4.54798296825753 | 2.38645642073598 | 1.06710864697714 |
| chr12 | 110041460 | 110041465 | - | 3.01188173062009 | 3.41098722619315 | 4.77291284147196 | 3.20132594093141 |
| chr12 | 110296679 | 110296738 | + | 13.0514874993537 | 9.09596593651506 | 4.77291284147196 | 9.60397782279422 |
| chr12 | 110346220 | 110346498 | + | 180.712903837206 | 171.686357051722 | 245.805011335806 | 171.804492163319 |
| chr12 | 110346546 | 110346591 | + | 34.1346596136944 | 59.1237785873479 | 62.0478669391355 | 42.6843458790854 |
| chr12 | 110346623 | 110346634 | + | 68.2693192273888 | 94.3706465913437 | 106.197310722751 | 55.4896496428111 |
| chr12 | 110346718 | 110346737 | + | 2.00792115374673 | 10.2329616785794 | 4.77291284147196 | 4.26843458790854 |
| chr12 | 110346807 | 110346834 | + | 8.03168461498692 | 4.54798296825753 | 5.96614105183995 | 10.6710864697714 |
| chr12 | 110346859 | 110346891 | + | 4.01584230749346 | 2.27399148412876 | 5.96614105183995 | 6.40265188186281 |
| chr12 | 110347054 | 110347088 | + | 9.03564519186028 | 10.2329616785794 | 4.77291284147196 | 10.6710864697714 |
| chr12 | 110347139 | 110347202 | + | 7.02772403811355 | 11.3699574206438 | 8.35259747257593 | 2.13421729395427 |
| chr12 | 110347267 | 110347279 | + | 2.00792115374673 | 7.95897019445067 | 7.15936926220794 | 7.46976052883995 |
| chr12 | 110350451 | 110350476 | + | 3.01188173062009 | 2.27399148412876 | 1.19322821036799 | 1.06710864697714 |
| chr12 | 110350792 | 110350817 | + | 4.01584230749346 | 3.41098722619315 | 3.57968463110397 | 2.13421729395427 |
| chr12 | 110373454 | 110373476 | - | 5.01980288436682 | 11.3699574206438 | 5.96614105183995 | 4.26843458790854 |
| chr12 | 110373507 | 110373522 | - | 5.01980288436682 | 19.3289276150945 | 17.8984231555198 | 19.2079556455884 |
| chr12 | 110434939 | 110434999 | - | 34.1346596136944 | 40.9318467143178 | 45.3426719939836 | 28.8119334683827 |
| chr12 | 110452741 | 110452759 | - | 13.0514874993537 | 12.5069531627082 | 8.35259747257593 | 6.40265188186281 |
| chr12 | 110452949 | 110453007 | - | 18.0712903837206 | 30.6988850357383 | 48.9223566250876 | 58.6909755837425 |
| chr12 | 110490171 | 110490217 | + | 32.1267384599477 | 45.4798296825753 | 58.4681823080315 | 75.7647139353766 |
| chr12 | 110490264 | 110490298 | + | 8.03168461498692 | 7.95897019445067 | 1.19322821036799 | 4.26843458790854 |
| chr12 | 110490350 | 110490363 | + | 1.00396057687336 | 2.27399148412876 | 1.19322821036799 | 3.20132594093141 |
| chr12 | 110491678 | 110491702 | - | 6.02376346124019 | 7.95897019445067 | 4.77291284147196 | 4.26843458790854 |
| chr12 | 110491994 | 110492001 | - | 25.0990144218341 | 23.876910583352 | 17.8984231555198 | 40.5501285851312 |
| chr12 | 110533310 | 110533354 | - | 2.00792115374673 | 5.68497871032191 | 4.77291284147196 | 2.13421729395427 |
| chr12 | 110535185 | 110535203 | - | 26.1029749987075 | 9.09596593651506 | 5.96614105183995 | 6.40265188186281 |
| chr12 | 110535902 | 110535905 | - | 2.00792115374673 | 2.27399148412876 | 4.77291284147196 | 1.06710864697714 |
| chr12 | 110536293 | 110536304 | - | 5.01980288436682 | 6.82197445238629 | 4.77291284147196 | 5.33554323488568 |
| chr12 | 110719832 | 110719869 | - | 4.01584230749346 | 7.95897019445067 | 10.7390538933119 | 8.53686917581709 |
| chr12 | 110719910 | 110719972 | - | 14.0554480762271 | 23.876910583352 | 26.2510206280958 | 37.3488026441998 |
| chr12 | 110720061 | 110720173 | - | 15.0594086531005 | 6.82197445238629 | 20.2848795762558 | 24.5434988804741 |
| chr12 | 110720195 | 110720206 | - | 2.00792115374673 | 1.13699574206438 | 1.19322821036799 | 2.13421729395427 |
| chr12 | 110720320 | 110720458 | - | 39.1544624980612 | 18.1919318730301 | 14.3187385244159 | 24.5434988804741 |
| chr12 | 110720504 | 110720656 | - | 11.043566345607 | 9.09596593651506 | 13.1255103140479 | 14.9395210576799 |
| chr12 | 110720682 | 110720719 | - | 5.01980288436682 | 6.82197445238629 | 13.1255103140479 | 9.60397782279422 |
| chr12 | 110720751 | 110720850 | - | 13.0514874993537 | 5.68497871032191 | 16.7051949451519 | 17.0737383516342 |
| chr12 | 110720945 | 110721066 | - | 4.01584230749346 | 3.41098722619315 | 22.6713359969918 | 9.60397782279422 |
| chr12 | 110722031 | 110722032 | - | 1.00396057687336 | 4.54798296825753 | 1.19322821036799 | 2.13421729395427 |
| chr12 | 110722134 | 110722272 | - | 5.01980288436682 | 9.09596593651506 | 16.7051949451519 | 22.4092815865198 |
| chr12 | 111360701 | 111360738 | - | 4.01584230749346 | 2.27399148412876 | 8.35259747257593 | 3.20132594093141 |
| chr12 | 111448558 | 111448561 | + | 2.00792115374673 | 1.13699574206438 | 5.96614105183995 | 9.60397782279422 |
| chr12 | 111448604 | 111448616 | + | 1.00396057687336 | 1.13699574206438 | 8.35259747257593 | 9.60397782279422 |
| chr12 | 111448687 | 111448723 | + | 3.01188173062009 | 6.82197445238629 | 23.8645642073598 | 29.8790421153598 |
| chr12 | 111449897 | 111449920 | + | 2.00792115374673 | 3.41098722619315 | 3.57968463110397 | 30.9461507623369 |
| chr12 | 111452292 | 111452332 | - | 1.00396057687336 | 2.27399148412876 | 5.96614105183995 | 2.13421729395427 |
| chr12 | 111452548 | 111452581 | - | 8.03168461498692 | 3.41098722619315 | 5.96614105183995 | 5.33554323488568 |
| chr12 | 111452609 | 111452670 | - | 25.0990144218341 | 23.876910583352 | 15.5119667347839 | 17.0737383516342 |
| chr12 | 111642214 | 111642222 | - | 12.0475269224804 | 13.6439489047726 | 8.35259747257593 | 7.46976052883995 |
| chr12 | 111643026 | 111643030 | - | 5.01980288436682 | 9.09596593651506 | 7.15936926220794 | 7.46976052883995 |
| chr12 | 111643136 | 111643140 | - | 22.087132691214 | 7.95897019445067 | 25.0577924177278 | 13.8724124107028 |
| chr12 | 111644114 | 111644121 | - | 7.02772403811355 | 6.82197445238629 | 5.96614105183995 | 18.1408469986113 |
| chr12 | 111809654 | 111809808 | + | 6.02376346124019 | 3.41098722619315 | 11.9322821036799 | 24.5434988804741 |
| chr12 | 111892965 | 111893077 | + | 54.2138711511617 | 86.411676396893 | 94.2650286190712 | 66.1607361125824 |
| chr12 | 111893173 | 111893195 | + | 12.0475269224804 | 29.5618892936739 | 13.1255103140479 | 14.9395210576799 |
| chr12 | 111893369 | 111893375 | + | 3.01188173062009 | 1.13699574206438 | 3.57968463110397 | 3.20132594093141 |
| chr12 | 112022171 | 112022247 | + | 27.1069355755808 | 50.0278126508328 | 58.4681823080315 | 83.2344744642166 |
| chr12 | 112022293 | 112022562 | + | 713.815970156962 | 1119.94080593342 | 962.935165766968 | 528.218780253682 |
| chr12 | 112022583 | 112022605 | + | 21.0831721143407 | 36.3838637460602 | 44.1494437836156 | 19.2079556455884 |
| chr12 | 112022620 | 112022688 | + | 162.641613453485 | 158.042408146949 | 141.994157033791 | 151.529427870753 |
| chr12 | 112022720 | 112022757 | + | 25.0990144218341 | 39.7948509722534 | 17.8984231555198 | 21.3421729395427 |
| chr12 | 112022782 | 112022950 | + | 124.491111532297 | 208.070220797782 | 105.004082512383 | 120.583277108416 |
| chr12 | 112027252 | 112027273 | - | 4.01584230749346 | 2.27399148412876 | 2.38645642073598 | 3.20132594093141 |
| chr12 | 112027388 | 112027405 | - | 14.0554480762271 | 13.6439489047726 | 16.7051949451519 | 12.8053037637256 |
| chr12 | 112027499 | 112027510 | - | 14.0554480762271 | 13.6439489047726 | 2.38645642073598 | 2.13421729395427 |
| chr12 | 112027546 | 112027549 | - | 4.01584230749346 | 2.27399148412876 | 5.96614105183995 | 2.13421729395427 |
| chr12 | 112027602 | 112027623 | - | 2.00792115374673 | 2.27399148412876 | 10.7390538933119 | 11.7381951167485 |
| chr12 | 112027737 | 112027761 | - | 23.0910932680874 | 22.7399148412876 | 50.1155848354556 | 46.952780466994 |
| chr12 | 112027850 | 112027861 | - | 2.00792115374673 | 2.27399148412876 | 7.15936926220794 | 6.40265188186281 |
| chr12 | 112027883 | 112027887 | - | 1.00396057687336 | 2.27399148412876 | 2.38645642073598 | 2.13421729395427 |
| chr12 | 112028547 | 112028577 | - | 3.01188173062009 | 5.68497871032191 | 5.96614105183995 | 7.46976052883995 |
| chr12 | 112028711 | 112028746 | - | 1.00396057687336 | 2.27399148412876 | 8.35259747257593 | 3.20132594093141 |
| chr12 | 112028808 | 112028814 | - | 2.00792115374673 | 5.68497871032191 | 9.54582568294392 | 7.46976052883995 |
| chr12 | 112029081 | 112029098 | - | 9.03564519186028 | 9.09596593651506 | 8.35259747257593 | 8.53686917581709 |
| chr12 | 112029220 | 112029233 | - | 4.01584230749346 | 7.95897019445067 | 8.35259747257593 | 6.40265188186281 |
| chr12 | 112029337 | 112029338 | - | 1.00396057687336 | 2.27399148412876 | 2.38645642073598 | 1.06710864697714 |
| chr12 | 112152953 | 112152974 | + | 3.01188173062009 | 1.13699574206438 | 2.38645642073598 | 3.20132594093141 |
| chr12 | 112160592 | 112160629 | - | 21.0831721143407 | 29.5618892936739 | 5.96614105183995 | 12.8053037637256 |
| chr12 | 112160741 | 112160775 | - | 12.0475269224804 | 10.2329616785794 | 4.77291284147196 | 4.26843458790854 |
| chr12 | 112506065 | 112506160 | + | 39.1544624980612 | 42.0688424563821 | 32.2171616799357 | 24.5434988804741 |
| chr12 | 112506340 | 112506360 | + | 1.00396057687336 | 2.27399148412876 | 10.7390538933119 | 5.33554323488568 |
| chr12 | 112506397 | 112506432 | + | 20.0792115374673 | 20.4659233571589 | 32.2171616799357 | 153.663645164708 |
| chr12 | 112506727 | 112506749 | + | 10.0396057687336 | 1.13699574206438 | 11.9322821036799 | 1.06710864697714 |
| chr12 | 112506972 | 112506987 | + | 14.0554480762271 | 6.82197445238629 | 10.7390538933119 | 2.13421729395427 |
| chr12 | 112507334 | 112507335 | + | 2.00792115374673 | 1.13699574206438 | 1.19322821036799 | 1.06710864697714 |
| chr12 | 112507385 | 112507409 | + | 5.01980288436682 | 5.68497871032191 | 5.96614105183995 | 4.26843458790854 |
| chr12 | 112507478 | 112507489 | + | 8.03168461498692 | 3.41098722619315 | 2.38645642073598 | 10.6710864697714 |
| chr12 | 112507729 | 112507734 | + | 6.02376346124019 | 3.41098722619315 | 5.96614105183995 | 1.06710864697714 |
| chr12 | 112507765 | 112507775 | + | 7.02772403811355 | 1.13699574206438 | 2.38645642073598 | 2.13421729395427 |
| chr12 | 112508381 | 112508391 | + | 1.00396057687336 | 1.13699574206438 | 2.38645642073598 | 2.13421729395427 |
| chr12 | 112509104 | 112509107 | + | 7.02772403811355 | 4.54798296825753 | 1.19322821036799 | 3.20132594093141 |
| chr12 | 112972760 | 112972846 | + | 148.586165377258 | 67.0827487817985 | 1.19322821036799 | 3.20132594093141 |
| chr12 | 113157507 | 113157510 | - | 5.01980288436682 | 3.41098722619315 | 1.19322821036799 | 2.13421729395427 |
| chr12 | 113158526 | 113158586 | - | 23.0910932680874 | 20.4659233571589 | 27.4442488384638 | 29.8790421153598 |
| chr12 | 113158634 | 113158713 | - | 38.1505019211879 | 11.3699574206438 | 33.4103898903037 | 66.1607361125824 |
| chr12 | 113158738 | 113158797 | - | 34.1346596136944 | 18.1919318730301 | 70.4004644117114 | 32.0132594093141 |
| chr12 | 113158821 | 113158944 | - | 45.1782259593014 | 25.0139063254164 | 101.424397881279 | 94.9726695809651 |
| chr12 | 113192089 | 113192091 | + | 3.01188173062009 | 1.13699574206438 | 1.19322821036799 | 3.20132594093141 |
| chr12 | 113192265 | 113192287 | + | 4.01584230749346 | 5.68497871032191 | 1.19322821036799 | 3.20132594093141 |
| chr12 | 113297252 | 113297260 | + | 1.00396057687336 | 4.54798296825753 | 1.19322821036799 | 1.06710864697714 |
| chr12 | 113298456 | 113298554 | + | 7.02772403811355 | 7.95897019445067 | 1.19322821036799 | 2.13421729395427 |
| chr12 | 113298917 | 113298936 | - | 3.01188173062009 | 7.95897019445067 | 3.57968463110397 | 4.26843458790854 |
| chr12 | 113388723 | 113388746 | + | 8.03168461498692 | 1.13699574206438 | 7.15936926220794 | 4.26843458790854 |
| chr12 | 113389424 | 113389442 | + | 6.02376346124019 | 6.82197445238629 | 7.15936926220794 | 5.33554323488568 |
| chr12 | 113389987 | 113390004 | + | 1.00396057687336 | 1.13699574206438 | 2.38645642073598 | 2.13421729395427 |
| chr12 | 113391434 | 113391516 | + | 7.02772403811355 | 4.54798296825753 | 3.57968463110397 | 16.006629704657 |
| chr12 | 113438251 | 113438253 | + | 1.00396057687336 | 11.3699574206438 | 3.57968463110397 | 5.33554323488568 |
| chr12 | 113816832 | 113816845 | - | 14.0554480762271 | 17.0549361309657 | 5.96614105183995 | 5.33554323488568 |
| chr12 | 115958626 | 115958635 | - | 2.00792115374673 | 2.27399148412876 | 5.96614105183995 | 3.20132594093141 |
| chr12 | 115958716 | 115958718 | - | 2.00792115374673 | 1.13699574206438 | 1.19322821036799 | 1.06710864697714 |
| chr12 | 115958921 | 115958932 | - | 9.03564519186028 | 10.2329616785794 | 11.9322821036799 | 25.6106075274513 |
| chr12 | 115959027 | 115959046 | - | 1.00396057687336 | 4.54798296825753 | 4.77291284147196 | 16.006629704657 |
| chr12 | 115959670 | 115959674 | - | 3.01188173062009 | 9.09596593651506 | 8.35259747257593 | 17.0737383516342 |
| chr12 | 115959941 | 115959944 | - | 1.00396057687336 | 1.13699574206438 | 2.38645642073598 | 8.53686917581709 |
| chr12 | 115960680 | 115960734 | - | 1.00396057687336 | 5.68497871032191 | 8.35259747257593 | 8.53686917581709 |
| chr12 | 116713728 | 116713762 | - | 11.043566345607 | 13.6439489047726 | 8.35259747257593 | 8.53686917581709 |
| chr12 | 116713854 | 116713862 | - | 10.0396057687336 | 9.09596593651506 | 7.15936926220794 | 2.13421729395427 |
| chr12 | 116713908 | 116713911 | - | 3.01188173062009 | 2.27399148412876 | 3.57968463110397 | 5.33554323488568 |
| chr12 | 116713938 | 116714048 | - | 26.1029749987075 | 17.0549361309657 | 19.0916513658878 | 13.8724124107028 |
| chr12 | 116716380 | 116716401 | - | 8.03168461498692 | 3.41098722619315 | 2.38645642073598 | 3.20132594093141 |
| chr12 | 116717457 | 116717501 | - | 2.00792115374673 | 4.54798296825753 | 2.38645642073598 | 3.20132594093141 |
| chr12 | 116850134 | 116850137 | + | 9.03564519186028 | 5.68497871032191 | 4.77291284147196 | 1.06710864697714 |
| chr12 | 116852524 | 116852542 | + | 2.00792115374673 | 2.27399148412876 | 4.77291284147196 | 7.46976052883995 |
| chr12 | 116852663 | 116852701 | + | 5.01980288436682 | 5.68497871032191 | 10.7390538933119 | 3.20132594093141 |
| chr12 | 116852777 | 116852778 | + | 2.00792115374673 | 2.27399148412876 | 4.77291284147196 | 2.13421729395427 |
| chr12 | 117143980 | 117143985 | - | 3.01188173062009 | 2.27399148412876 | 1.19322821036799 | 1.06710864697714 |
| chr12 | 117144013 | 117144056 | - | 6.02376346124019 | 11.3699574206438 | 1.19322821036799 | 3.20132594093141 |
| chr12 | 117144256 | 117144275 | - | 16.0633692299738 | 9.09596593651506 | 1.19322821036799 | 6.40265188186281 |
| chr12 | 117144988 | 117145022 | - | 6.02376346124019 | 2.27399148412876 | 2.38645642073598 | 3.20132594093141 |
| chr12 | 117145528 | 117145636 | - | 78.3089249961224 | 68.2197445238629 | 14.3187385244159 | 21.3421729395427 |
| chr12 | 118031269 | 118031289 | + | 8.03168461498692 | 5.68497871032191 | 7.15936926220794 | 7.46976052883995 |
| chr12 | 118031319 | 118031324 | + | 4.01584230749346 | 1.13699574206438 | 5.96614105183995 | 2.13421729395427 |
| chr12 | 118031355 | 118031369 | + | 5.01980288436682 | 4.54798296825753 | 1.19322821036799 | 2.13421729395427 |
| chr12 | 118031511 | 118031519 | + | 5.01980288436682 | 17.0549361309657 | 3.57968463110397 | 8.53686917581709 |
| chr12 | 118031579 | 118031584 | + | 3.01188173062009 | 5.68497871032191 | 1.19322821036799 | 8.53686917581709 |
| chr12 | 118031772 | 118031778 | + | 1.00396057687336 | 1.13699574206438 | 4.77291284147196 | 2.13421729395427 |
| chr12 | 118031964 | 118031967 | + | 7.02772403811355 | 14.780944646837 | 3.57968463110397 | 6.40265188186281 |
| chr12 | 118032004 | 118032017 | + | 8.03168461498692 | 3.41098722619315 | 2.38645642073598 | 2.13421729395427 |
| chr12 | 118032108 | 118032123 | + | 1.00396057687336 | 5.68497871032191 | 4.77291284147196 | 3.20132594093141 |
| chr12 | 118032150 | 118032152 | + | 8.03168461498692 | 1.13699574206438 | 4.77291284147196 | 8.53686917581709 |
| chr12 | 118032933 | 118032948 | - | 4.01584230749346 | 5.68497871032191 | 7.15936926220794 | 6.40265188186281 |
| chr12 | 118033289 | 118033365 | - | 14.0554480762271 | 11.3699574206438 | 29.8307052591997 | 13.8724124107028 |
| chr12 | 118033393 | 118033407 | - | 5.01980288436682 | 5.68497871032191 | 1.19322821036799 | 10.6710864697714 |
| chr12 | 118033431 | 118033495 | - | 9.03564519186028 | 12.5069531627082 | 26.2510206280958 | 32.0132594093141 |
| chr12 | 118033521 | 118033549 | - | 4.01584230749346 | 3.41098722619315 | 10.7390538933119 | 16.006629704657 |
| chr12 | 118033574 | 118033615 | - | 11.043566345607 | 4.54798296825753 | 26.2510206280958 | 65.0936274656053 |
| chr12 | 118033644 | 118033679 | - | 3.01188173062009 | 2.27399148412876 | 15.5119667347839 | 12.8053037637256 |
| chr12 | 118033710 | 118033734 | - | 5.01980288436682 | 2.27399148412876 | 5.96614105183995 | 2.13421729395427 |
| chr12 | 118033763 | 118033771 | - | 5.01980288436682 | 1.13699574206438 | 4.77291284147196 | 12.8053037637256 |
| chr12 | 118033790 | 118033812 | - | 9.03564519186028 | 1.13699574206438 | 13.1255103140479 | 5.33554323488568 |
| chr12 | 118033894 | 118033979 | - | 13.0514874993537 | 3.41098722619315 | 25.0577924177278 | 24.5434988804741 |
| chr12 | 118034050 | 118034181 | - | 15.0594086531005 | 10.2329616785794 | 38.1833027317757 | 28.8119334683827 |
| chr12 | 118144660 | 118144828 | + | 151.598047107878 | 92.0966551072149 | 152.733210927103 | 164.334731634479 |
| chr12 | 118144867 | 118144933 | + | 226.89509037338 | 186.467301698559 | 81.1395183050233 | 198.482208337747 |
| chr12 | 118145003 | 118145061 | + | 565.229804779704 | 392.263531012212 | 140.800928823423 | 145.12677598889 |
| chr12 | 118145082 | 118145116 | + | 49.1940682667949 | 27.2878978095452 | 46.5359002043516 | 20.2750642925656 |
| chr12 | 118145216 | 118145240 | + | 4.01584230749346 | 5.68497871032191 | 1.19322821036799 | 2.13421729395427 |
| chr12 | 118145271 | 118145339 | + | 25.0990144218341 | 34.1098722619315 | 15.5119667347839 | 22.4092815865198 |
| chr12 | 118145361 | 118145407 | + | 84.3326884573626 | 75.0417189762492 | 36.9900745214077 | 46.952780466994 |
| chr12 | 118145435 | 118145484 | + | 128.506953839791 | 85.2746806548286 | 81.1395183050233 | 58.6909755837425 |
| chr12 | 118145507 | 118145563 | + | 326.287187483843 | 329.728765198671 | 137.221244192319 | 116.314842520508 |
| chr12 | 118150176 | 118150183 | - | 2.00792115374673 | 4.54798296825753 | 8.35259747257593 | 5.33554323488568 |
| chr12 | 118150331 | 118150340 | - | 1.00396057687336 | 6.82197445238629 | 14.3187385244159 | 13.8724124107028 |
| chr12 | 118150369 | 118150384 | - | 94.3722942260963 | 133.028501821533 | 628.83126686393 | 488.735760315528 |
| chr12 | 118150412 | 118150418 | - | 4.01584230749346 | 1.13699574206438 | 25.0577924177278 | 91.7713436400337 |
| chr12 | 118150536 | 118150543 | - | 4.01584230749346 | 7.95897019445067 | 20.2848795762558 | 9.60397782279422 |
| chr12 | 118150575 | 118150590 | - | 39.1544624980612 | 46.6168254246397 | 107.390538933119 | 113.113516579576 |
| chr12 | 118150823 | 118150828 | - | 5.01980288436682 | 4.54798296825753 | 10.7390538933119 | 9.60397782279422 |
| chr12 | 118150902 | 118150905 | - | 5.01980288436682 | 3.41098722619315 | 4.77291284147196 | 7.46976052883995 |
| chr12 | 118150983 | 118151015 | - | 9.03564519186028 | 9.09596593651506 | 40.5697591525116 | 16.006629704657 |
| chr12 | 118414468 | 118414471 | + | 4.01584230749346 | 1.13699574206438 | 2.38645642073598 | 1.06710864697714 |
| chr12 | 118414527 | 118414531 | + | 2.00792115374673 | 3.41098722619315 | 2.38645642073598 | 2.13421729395427 |
| chr12 | 118414759 | 118414765 | + | 7.02772403811355 | 6.82197445238629 | 5.96614105183995 | 4.26843458790854 |
| chr12 | 118414808 | 118414820 | + | 4.01584230749346 | 5.68497871032191 | 3.57968463110397 | 1.06710864697714 |
| chr12 | 118415141 | 118415144 | + | 2.00792115374673 | 1.13699574206438 | 3.57968463110397 | 2.13421729395427 |
| chr12 | 118415280 | 118415301 | + | 2.00792115374673 | 6.82197445238629 | 9.54582568294392 | 4.26843458790854 |
| chr12 | 118417341 | 118417355 | + | 7.02772403811355 | 4.54798296825753 | 2.38645642073598 | 2.13421729395427 |
| chr12 | 118417914 | 118417915 | + | 1.00396057687336 | 1.13699574206438 | 2.38645642073598 | 1.06710864697714 |
| chr12 | 119640849 | 119640850 | + | 5.01980288436682 | 2.27399148412876 | 2.38645642073598 | 2.13421729395427 |
| chr12 | 119680574 | 119680588 | + | 1.00396057687336 | 1.13699574206438 | 5.96614105183995 | 2.13421729395427 |
| chr12 | 119680610 | 119680629 | + | 2.00792115374673 | 4.54798296825753 | 9.54582568294392 | 8.53686917581709 |
| chr12 | 119680839 | 119680859 | + | 3.01188173062009 | 1.13699574206438 | 25.0577924177278 | 8.53686917581709 |
| chr12 | 119680927 | 119680997 | + | 3.01188173062009 | 2.27399148412876 | 17.8984231555198 | 19.2079556455884 |
| chr12 | 119681479 | 119681493 | + | 3.01188173062009 | 6.82197445238629 | 13.1255103140479 | 19.2079556455884 |
| chr12 | 119681536 | 119681539 | + | 8.03168461498692 | 31.8358807778027 | 35.7968463110397 | 94.9726695809651 |
| chr12 | 119681582 | 119681589 | + | 8.03168461498692 | 9.09596593651506 | 20.2848795762558 | 18.1408469986113 |
| chr12 | 119685883 | 119685887 | - | 3.01188173062009 | 2.27399148412876 | 1.19322821036799 | 4.26843458790854 |
| chr12 | 119686021 | 119686030 | - | 13.0514874993537 | 12.5069531627082 | 4.77291284147196 | 17.0737383516342 |
| chr12 | 119686511 | 119686513 | - | 1.00396057687336 | 1.13699574206438 | 1.19322821036799 | 1.06710864697714 |
| chr12 | 119687321 | 119687329 | - | 5.01980288436682 | 2.27399148412876 | 2.38645642073598 | 2.13421729395427 |
| chr12 | 119687476 | 119687478 | - | 1.00396057687336 | 2.27399148412876 | 1.19322821036799 | 1.06710864697714 |
| chr12 | 119687847 | 119687910 | - | 20.0792115374673 | 20.4659233571589 | 25.0577924177278 | 30.9461507623369 |
| chr12 | 119688056 | 119688076 | - | 4.01584230749346 | 5.68497871032191 | 3.57968463110397 | 7.46976052883995 |
| chr12 | 120095225 | 120095265 | - | 8.03168461498692 | 2.27399148412876 | 1.19322821036799 | 3.20132594093141 |
| chr12 | 120096425 | 120096492 | - | 46.1821865361748 | 29.5618892936739 | 48.9223566250876 | 30.9461507623369 |
| chr12 | 120097166 | 120097175 | - | 16.0633692299738 | 10.2329616785794 | 8.35259747257593 | 4.26843458790854 |
| chr12 | 120127285 | 120127353 | - | 52.205949997415 | 87.5486721389574 | 68.0140079909754 | 90.7042349930565 |
| chr12 | 120127400 | 120127402 | - | 2.00792115374673 | 3.41098722619315 | 1.19322821036799 | 1.06710864697714 |
| chr12 | 120127432 | 120127454 | - | 4.01584230749346 | 1.13699574206438 | 2.38645642073598 | 9.60397782279422 |
| chr12 | 120196717 | 120196771 | - | 26.1029749987075 | 28.4248935516096 | 17.8984231555198 | 22.4092815865198 |
| chr12 | 120201258 | 120201312 | + | 74.293082688629 | 77.315710460378 | 34.6036181006717 | 57.6238669367653 |
| chr12 | 120201342 | 120201371 | + | 51.2019894205416 | 45.4798296825753 | 57.2749540976635 | 82.1673658172394 |
| chr12 | 120210558 | 120210595 | - | 25.0990144218341 | 37.5208594881246 | 33.4103898903037 | 41.6172372321083 |
| chr12 | 120210691 | 120210714 | - | 4.01584230749346 | 2.27399148412876 | 1.19322821036799 | 4.26843458790854 |
| chr12 | 120211393 | 120211437 | - | 23.0910932680874 | 19.3289276150945 | 31.0239334695677 | 17.0737383516342 |
| chr12 | 120211463 | 120211483 | - | 9.03564519186028 | 3.41098722619315 | 4.77291284147196 | 2.13421729395427 |
| chr12 | 120211522 | 120211537 | - | 3.01188173062009 | 2.27399148412876 | 3.57968463110397 | 1.06710864697714 |
| chr12 | 120211568 | 120211593 | - | 10.0396057687336 | 2.27399148412876 | 11.9322821036799 | 1.06710864697714 |
| chr12 | 120211620 | 120211676 | - | 6.02376346124019 | 9.09596593651506 | 11.9322821036799 | 8.53686917581709 |
| chr12 | 120211790 | 120211800 | - | 2.00792115374673 | 2.27399148412876 | 2.38645642073598 | 2.13421729395427 |
| chr12 | 120211865 | 120211868 | - | 2.00792115374673 | 2.27399148412876 | 3.57968463110397 | 2.13421729395427 |
| chr12 | 120211924 | 120211957 | - | 20.0792115374673 | 14.780944646837 | 36.9900745214077 | 7.46976052883995 |
| chr12 | 120212052 | 120212070 | - | 8.03168461498692 | 1.13699574206438 | 9.54582568294392 | 8.53686917581709 |
| chr12 | 120212710 | 120212725 | + | 4.01584230749346 | 7.95897019445067 | 3.57968463110397 | 5.33554323488568 |
| chr12 | 120440565 | 120440591 | + | 8.03168461498692 | 9.09596593651506 | 11.9322821036799 | 8.53686917581709 |
| chr12 | 120440672 | 120440708 | + | 18.0712903837206 | 32.9728765198671 | 45.3426719939836 | 49.0869977609482 |
| chr12 | 120444050 | 120444075 | - | 34.1346596136944 | 31.8358807778027 | 22.6713359969918 | 40.5501285851312 |
| chr12 | 120444214 | 120444251 | - | 5.01980288436682 | 5.68497871032191 | 1.19322821036799 | 5.33554323488568 |
| chr12 | 120444273 | 120444290 | - | 2.00792115374673 | 1.13699574206438 | 3.57968463110397 | 2.13421729395427 |
| chr12 | 120444393 | 120444500 | - | 4.01584230749346 | 4.54798296825753 | 3.57968463110397 | 9.60397782279422 |
| chr12 | 120444550 | 120444639 | - | 11.043566345607 | 14.780944646837 | 10.7390538933119 | 7.46976052883995 |
| chr12 | 120444660 | 120444684 | - | 8.03168461498692 | 5.68497871032191 | 7.15936926220794 | 2.13421729395427 |
| chr12 | 120444715 | 120444740 | - | 1.00396057687336 | 1.13699574206438 | 3.57968463110397 | 4.26843458790854 |
| chr12 | 120444757 | 120444801 | - | 7.02772403811355 | 14.780944646837 | 16.7051949451519 | 16.006629704657 |
| chr12 | 120444827 | 120444834 | - | 3.01188173062009 | 6.82197445238629 | 3.57968463110397 | 4.26843458790854 |
| chr12 | 120459978 | 120459999 | + | 20.0792115374673 | 7.95897019445067 | 4.77291284147196 | 8.53686917581709 |
| chr12 | 120460443 | 120460445 | + | 2.00792115374673 | 3.41098722619315 | 3.57968463110397 | 1.06710864697714 |
| chr12 | 120461211 | 120461231 | + | 2.00792115374673 | 1.13699574206438 | 2.38645642073598 | 5.33554323488568 |
| chr12 | 120461576 | 120461578 | + | 3.01188173062009 | 2.27399148412876 | 3.57968463110397 | 2.13421729395427 |
| chr12 | 120461694 | 120461768 | - | 32.1267384599477 | 38.657855230189 | 22.6713359969918 | 21.3421729395427 |
| chr12 | 120461925 | 120461976 | - | 12.0475269224804 | 20.4659233571589 | 23.8645642073598 | 26.6777161744284 |
| chr12 | 120462186 | 120462196 | + | 13.0514874993537 | 3.41098722619315 | 9.54582568294392 | 4.26843458790854 |
| chr12 | 120463590 | 120463612 | + | 7.02772403811355 | 13.6439489047726 | 1.19322821036799 | 8.53686917581709 |
| chr12 | 120463641 | 120463647 | + | 4.01584230749346 | 4.54798296825753 | 7.15936926220794 | 1.06710864697714 |
| chr12 | 120498348 | 120498435 | + | 53.2099105742883 | 78.4527062024424 | 33.4103898903037 | 40.5501285851312 |
| chr12 | 120503308 | 120503358 | - | 5.01980288436682 | 2.27399148412876 | 5.96614105183995 | 6.40265188186281 |
| chr12 | 120503433 | 120503453 | - | 3.01188173062009 | 5.68497871032191 | 2.38645642073598 | 3.20132594093141 |
| chr12 | 120503474 | 120503482 | - | 4.01584230749346 | 2.27399148412876 | 1.19322821036799 | 2.13421729395427 |
| chr12 | 120503491 | 120503524 | - | 125.495072109171 | 120.521548658824 | 26.2510206280958 | 20.2750642925656 |
| chr12 | 120503559 | 120503652 | - | 13.0514874993537 | 17.0549361309657 | 13.1255103140479 | 28.8119334683827 |
| chr12 | 120503675 | 120503700 | - | 8.03168461498692 | 12.5069531627082 | 5.96614105183995 | 10.6710864697714 |
| chr12 | 120503724 | 120503736 | - | 3.01188173062009 | 12.5069531627082 | 4.77291284147196 | 2.13421729395427 |
| chr12 | 120503767 | 120503788 | - | 2.00792115374673 | 3.41098722619315 | 2.38645642073598 | 5.33554323488568 |
| chr12 | 120576590 | 120576694 | + | 52.205949997415 | 47.753821166704 | 100.231169670911 | 102.442430109805 |
| chr12 | 120576770 | 120576863 | + | 26.1029749987075 | 15.9179403889013 | 17.8984231555198 | 26.6777161744284 |
| chr12 | 120579075 | 120579151 | - | 55.217831728035 | 70.4937360079917 | 13.1255103140479 | 17.0737383516342 |
| chr12 | 120696553 | 120696598 | + | 18.0712903837206 | 20.4659233571589 | 13.1255103140479 | 4.26843458790854 |
| chr12 | 120696650 | 120696652 | + | 1.00396057687336 | 3.41098722619315 | 3.57968463110397 | 1.06710864697714 |
| chr12 | 120696942 | 120696953 | + | 4.01584230749346 | 6.82197445238629 | 1.19322821036799 | 10.6710864697714 |
| chr12 | 120697275 | 120697310 | + | 12.0475269224804 | 18.1919318730301 | 2.38645642073598 | 3.20132594093141 |
| chr12 | 120697547 | 120697558 | + | 25.0990144218341 | 13.6439489047726 | 9.54582568294392 | 3.20132594093141 |
| chr12 | 120697808 | 120697838 | + | 7.02772403811355 | 3.41098722619315 | 2.38645642073598 | 1.06710864697714 |
| chr12 | 120698677 | 120698701 | + | 21.0831721143407 | 11.3699574206438 | 4.77291284147196 | 1.06710864697714 |
| chr12 | 120698944 | 120699045 | + | 79.3128855729958 | 38.657855230189 | 40.5697591525116 | 135.522798166096 |
| chr12 | 120699077 | 120699172 | + | 62.2455557661486 | 52.3018041349616 | 32.2171616799357 | 20.2750642925656 |
| chr12 | 120699220 | 120699312 | + | 25.0990144218341 | 17.0549361309657 | 10.7390538933119 | 9.60397782279422 |
| chr12 | 120699380 | 120699394 | + | 15.0594086531005 | 9.09596593651506 | 9.54582568294392 | 7.46976052883995 |
| chr12 | 120700392 | 120700436 | + | 42.1663442286813 | 13.6439489047726 | 20.2848795762558 | 74.6976052883995 |
| chr12 | 120700900 | 120700937 | + | 7.02772403811355 | 4.54798296825753 | 2.38645642073598 | 4.26843458790854 |
| chr12 | 120701708 | 120701828 | + | 14.0554480762271 | 11.3699574206438 | 2.38645642073598 | 9.60397782279422 |
| chr12 | 120720156 | 120720159 | + | 2.00792115374673 | 2.27399148412876 | 1.19322821036799 | 1.06710864697714 |
| chr12 | 120720568 | 120720572 | + | 8.03168461498692 | 3.41098722619315 | 4.77291284147196 | 2.13421729395427 |
| chr12 | 120720815 | 120720842 | + | 6.02376346124019 | 3.41098722619315 | 9.54582568294392 | 6.40265188186281 |
| chr12 | 120720878 | 120720926 | + | 4.01584230749346 | 6.82197445238629 | 11.9322821036799 | 18.1408469986113 |
| chr12 | 120720957 | 120720973 | + | 9.03564519186028 | 3.41098722619315 | 2.38645642073598 | 4.26843458790854 |
| chr12 | 120721998 | 120722041 | + | 6.02376346124019 | 4.54798296825753 | 4.77291284147196 | 3.20132594093141 |
| chr12 | 120722125 | 120722147 | + | 1.00396057687336 | 2.27399148412876 | 10.7390538933119 | 7.46976052883995 |
| chr12 | 120722182 | 120722213 | + | 6.02376346124019 | 5.68497871032191 | 8.35259747257593 | 8.53686917581709 |
| chr12 | 120722817 | 120722830 | + | 5.01980288436682 | 4.54798296825753 | 4.77291284147196 | 3.20132594093141 |
| chr12 | 120722986 | 120723007 | + | 5.01980288436682 | 5.68497871032191 | 4.77291284147196 | 5.33554323488568 |
| chr12 | 120723088 | 120723105 | + | 3.01188173062009 | 3.41098722619315 | 2.38645642073598 | 2.13421729395427 |
| chr12 | 120723199 | 120723214 | + | 8.03168461498692 | 2.27399148412876 | 4.77291284147196 | 7.46976052883995 |
| chr12 | 120723373 | 120723381 | + | 18.0712903837206 | 22.7399148412876 | 8.35259747257593 | 22.4092815865198 |
| chr12 | 120739951 | 120739959 | + | 1.00396057687336 | 1.13699574206438 | 1.19322821036799 | 7.46976052883995 |
| chr12 | 120763312 | 120763314 | - | 6.02376346124019 | 2.27399148412876 | 1.19322821036799 | 4.26843458790854 |
| chr12 | 120763954 | 120763970 | - | 3.01188173062009 | 7.95897019445067 | 5.96614105183995 | 1.06710864697714 |
| chr12 | 120764002 | 120764007 | - | 5.01980288436682 | 5.68497871032191 | 5.96614105183995 | 3.20132594093141 |
| chr12 | 120764055 | 120764062 | - | 3.01188173062009 | 5.68497871032191 | 3.57968463110397 | 1.06710864697714 |
| chr12 | 120764222 | 120764224 | - | 1.00396057687336 | 5.68497871032191 | 3.57968463110397 | 2.13421729395427 |
| chr12 | 120764350 | 120764377 | - | 5.01980288436682 | 13.6439489047726 | 7.15936926220794 | 3.20132594093141 |
| chr12 | 120764525 | 120764531 | - | 1.00396057687336 | 1.13699574206438 | 2.38645642073598 | 1.06710864697714 |
| chr12 | 120764556 | 120764579 | - | 4.01584230749346 | 3.41098722619315 | 1.19322821036799 | 3.20132594093141 |
| chr12 | 121237918 | 121237935 | - | 8.03168461498692 | 5.68497871032191 | 2.38645642073598 | 5.33554323488568 |
| chr12 | 121238586 | 121238604 | - | 21.0831721143407 | 26.1509020674808 | 13.1255103140479 | 34.1474767032683 |
| chr12 | 121240137 | 121240201 | - | 9.03564519186028 | 17.0549361309657 | 29.8307052591997 | 32.0132594093141 |
| chr12 | 121240233 | 121240273 | - | 20.0792115374673 | 19.3289276150945 | 16.7051949451519 | 10.6710864697714 |
| chr12 | 121240434 | 121240446 | - | 3.01188173062009 | 2.27399148412876 | 1.19322821036799 | 1.06710864697714 |
| chr12 | 121240467 | 121240474 | - | 4.01584230749346 | 3.41098722619315 | 1.19322821036799 | 3.20132594093141 |
| chr12 | 121308366 | 121308477 | - | 22.087132691214 | 22.7399148412876 | 25.0577924177278 | 20.2750642925656 |
| chr12 | 121424145 | 121424150 | + | 6.02376346124019 | 7.95897019445067 | 1.19322821036799 | 1.06710864697714 |
| chr12 | 121424231 | 121424234 | + | 2.00792115374673 | 4.54798296825753 | 4.77291284147196 | 2.13421729395427 |
| chr12 | 121429250 | 121429260 | - | 2.00792115374673 | 4.54798296825753 | 3.57968463110397 | 3.20132594093141 |
| chr12 | 121429963 | 121430068 | - | 20.0792115374673 | 9.09596593651506 | 36.9900745214077 | 44.8185631730397 |
| chr12 | 121430112 | 121430157 | - | 12.0475269224804 | 9.09596593651506 | 26.2510206280958 | 24.5434988804741 |
| chr12 | 121430227 | 121430264 | - | 9.03564519186028 | 22.7399148412876 | 27.4442488384638 | 18.1408469986113 |
| chr12 | 121430553 | 121430563 | + | 11.043566345607 | 5.68497871032191 | 1.19322821036799 | 2.13421729395427 |
| chr12 | 121780339 | 121780340 | + | 2.00792115374673 | 1.13699574206438 | 2.38645642073598 | 2.13421729395427 |
| chr12 | 121801026 | 121801028 | + | 21.0831721143407 | 15.9179403889013 | 2.38645642073598 | 17.0737383516342 |
| chr12 | 121803089 | 121803090 | + | 5.01980288436682 | 1.13699574206438 | 3.57968463110397 | 7.46976052883995 |
| chr12 | 121831090 | 121831092 | + | 11.043566345607 | 18.1919318730301 | 5.96614105183995 | 5.33554323488568 |
| chr12 | 121832479 | 121832494 | + | 32.1267384599477 | 71.6307317500561 | 17.8984231555198 | 23.476390233497 |
| chr12 | 121832524 | 121832548 | + | 5.01980288436682 | 4.54798296825753 | 1.19322821036799 | 1.06710864697714 |
| chr12 | 121903004 | 121903045 | + | 16.0633692299738 | 13.6439489047726 | 32.2171616799357 | 10.6710864697714 |
| chr12 | 121903080 | 121903105 | + | 12.0475269224804 | 2.27399148412876 | 9.54582568294392 | 19.2079556455884 |
| chr12 | 121916321 | 121916365 | + | 5.01980288436682 | 3.41098722619315 | 4.77291284147196 | 6.40265188186281 |
| chr12 | 121916531 | 121916599 | + | 95.3762548029696 | 105.740604011988 | 53.6952694665595 | 99.2411041688736 |
| chr12 | 122146154 | 122146219 | + | 4.01584230749346 | 3.41098722619315 | 26.2510206280958 | 6.40265188186281 |
| chr12 | 122146245 | 122146264 | + | 3.01188173062009 | 1.13699574206438 | 11.9322821036799 | 13.8724124107028 |
| chr12 | 122146809 | 122146867 | + | 5.01980288436682 | 1.13699574206438 | 11.9322821036799 | 17.0737383516342 |
| chr12 | 122146895 | 122146913 | + | 1.00396057687336 | 1.13699574206438 | 4.77291284147196 | 10.6710864697714 |
| chr12 | 122146940 | 122146954 | + | 2.00792115374673 | 3.41098722619315 | 9.54582568294392 | 10.6710864697714 |
| chr12 | 122147030 | 122147083 | + | 6.02376346124019 | 4.54798296825753 | 25.0577924177278 | 13.8724124107028 |
| chr12 | 122147110 | 122147218 | + | 46.1821865361748 | 35.2468680039958 | 119.322821036799 | 117.381951167485 |
| chr12 | 122207795 | 122207829 | - | 19.0752509605939 | 29.5618892936739 | 25.0577924177278 | 42.6843458790854 |
| chr12 | 122207855 | 122207877 | - | 5.01980288436682 | 13.6439489047726 | 10.7390538933119 | 5.33554323488568 |
| chr12 | 122207951 | 122207965 | - | 2.00792115374673 | 5.68497871032191 | 4.77291284147196 | 10.6710864697714 |
| chr12 | 122208073 | 122208090 | - | 1.00396057687336 | 4.54798296825753 | 5.96614105183995 | 9.60397782279422 |
| chr12 | 122216487 | 122216489 | - | 7.02772403811355 | 3.41098722619315 | 9.54582568294392 | 14.9395210576799 |
| chr12 | 122231582 | 122231586 | - | 4.01584230749346 | 2.27399148412876 | 4.77291284147196 | 5.33554323488568 |
| chr12 | 122231650 | 122231653 | - | 3.01188173062009 | 1.13699574206438 | 1.19322821036799 | 4.26843458790854 |
| chr12 | 122232129 | 122232131 | - | 3.01188173062009 | 1.13699574206438 | 3.57968463110397 | 3.20132594093141 |
| chr12 | 122271957 | 122271963 | - | 9.03564519186028 | 9.09596593651506 | 1.19322821036799 | 1.06710864697714 |
| chr12 | 122272023 | 122272059 | - | 3.01188173062009 | 10.2329616785794 | 4.77291284147196 | 2.13421729395427 |
| chr12 | 122272496 | 122272502 | - | 1.00396057687336 | 3.41098722619315 | 4.77291284147196 | 2.13421729395427 |
| chr12 | 122272790 | 122272810 | - | 2.00792115374673 | 5.68497871032191 | 3.57968463110397 | 1.06710864697714 |
| chr12 | 122272852 | 122272862 | - | 8.03168461498692 | 4.54798296825753 | 1.19322821036799 | 1.06710864697714 |
| chr12 | 122471816 | 122471817 | - | 1.00396057687336 | 2.27399148412876 | 1.19322821036799 | 2.13421729395427 |
| chr12 | 122472921 | 122472945 | - | 2.00792115374673 | 7.95897019445067 | 3.57968463110397 | 1.06710864697714 |
| chr12 | 122473032 | 122473060 | - | 9.03564519186028 | 17.0549361309657 | 3.57968463110397 | 6.40265188186281 |
| chr12 | 122473211 | 122473218 | - | 4.01584230749346 | 4.54798296825753 | 4.77291284147196 | 1.06710864697714 |
| chr12 | 122473260 | 122473294 | - | 31.1227778830743 | 42.0688424563821 | 16.7051949451519 | 33.0803680562912 |
| chr12 | 122505082 | 122505091 | - | 92.3643730723495 | 318.358807778027 | 121.709277457535 | 90.7042349930565 |
| chr12 | 122505250 | 122505263 | - | 16.0633692299738 | 23.876910583352 | 4.77291284147196 | 5.33554323488568 |
| chr12 | 122626249 | 122626283 | + | 5.01980288436682 | 11.3699574206438 | 11.9322821036799 | 17.0737383516342 |
| chr12 | 122769782 | 122769806 | + | 9.03564519186028 | 9.09596593651506 | 14.3187385244159 | 22.4092815865198 |
| chr12 | 122769849 | 122769874 | + | 15.0594086531005 | 2.27399148412876 | 20.2848795762558 | 21.3421729395427 |
| chr12 | 122769982 | 122770010 | + | 2.00792115374673 | 9.09596593651506 | 10.7390538933119 | 7.46976052883995 |
| chr12 | 122770031 | 122770078 | + | 5.01980288436682 | 5.68497871032191 | 20.2848795762558 | 9.60397782279422 |
| chr12 | 122770272 | 122770316 | + | 22.087132691214 | 23.876910583352 | 22.6713359969918 | 22.4092815865198 |
| chr12 | 122770365 | 122770392 | + | 15.0594086531005 | 13.6439489047726 | 14.3187385244159 | 12.8053037637256 |
| chr12 | 122770437 | 122770449 | + | 10.0396057687336 | 7.95897019445067 | 13.1255103140479 | 23.476390233497 |
| chr12 | 122770476 | 122770504 | + | 14.0554480762271 | 14.780944646837 | 8.35259747257593 | 5.33554323488568 |
| chr12 | 122770531 | 122770559 | + | 6.02376346124019 | 6.82197445238629 | 2.38645642073598 | 6.40265188186281 |
| chr12 | 122770582 | 122770601 | + | 2.00792115374673 | 3.41098722619315 | 3.57968463110397 | 2.13421729395427 |
| chr12 | 122770628 | 122770681 | + | 21.0831721143407 | 28.4248935516096 | 25.0577924177278 | 43.7514545260626 |
| chr12 | 122770705 | 122770751 | + | 17.0673298068472 | 18.1919318730301 | 40.5697591525116 | 37.3488026441998 |
| chr12 | 122770773 | 122770872 | + | 45.1782259593014 | 59.1237785873479 | 76.3666054635513 | 74.6976052883995 |
| chr12 | 122975182 | 122975357 | + | 317.251542291983 | 536.661990254388 | 472.518371305724 | 420.440806908991 |
| chr12 | 122976667 | 122976689 | + | 1.00396057687336 | 3.41098722619315 | 9.54582568294392 | 5.33554323488568 |
| chr12 | 122978023 | 122978025 | + | 1.00396057687336 | 2.27399148412876 | 2.38645642073598 | 1.06710864697714 |
| chr12 | 122979367 | 122979409 | + | 2.00792115374673 | 2.27399148412876 | 7.15936926220794 | 8.53686917581709 |
| chr12 | 122980014 | 122980026 | + | 3.01188173062009 | 9.09596593651506 | 7.15936926220794 | 3.20132594093141 |
| chr12 | 122980213 | 122980398 | + | 4.01584230749346 | 4.54798296825753 | 8.35259747257593 | 34.1474767032683 |
| chr12 | 122980645 | 122980864 | + | 882.481347071687 | 867.527751195123 | 844.805572940537 | 1315.74496172281 |
| chr12 | 122980892 | 122980968 | + | 78.3089249961224 | 50.0278126508328 | 58.4681823080315 | 102.442430109805 |
| chr12 | 122981126 | 122981143 | + | 7.02772403811355 | 5.68497871032191 | 15.5119667347839 | 9.60397782279422 |
| chr12 | 122981168 | 122981241 | + | 41.1623836518079 | 42.0688424563821 | 75.1733772531833 | 61.8923015246739 |
| chr12 | 122981263 | 122981296 | + | 11.043566345607 | 6.82197445238629 | 31.0239334695677 | 27.7448248214055 |
| chr12 | 122981594 | 122981626 | + | 16.0633692299738 | 21.6029190992233 | 39.3765309421437 | 37.3488026441998 |
| chr12 | 122981702 | 122981812 | + | 53.2099105742883 | 59.1237785873479 | 66.8207797806074 | 59.7580842307196 |
| chr12 | 122981835 | 122981870 | + | 6.02376346124019 | 9.09596593651506 | 14.3187385244159 | 23.476390233497 |
| chr12 | 122982466 | 122982529 | + | 6.02376346124019 | 5.68497871032191 | 2.38645642073598 | 7.46976052883995 |
| chr12 | 122982704 | 122982762 | + | 3.01188173062009 | 1.13699574206438 | 2.38645642073598 | 3.20132594093141 |
| chr12 | 122982817 | 122982888 | + | 70.2772403811355 | 53.438799877026 | 34.6036181006717 | 66.1607361125824 |
| chr12 | 122983630 | 122983633 | - | 4.01584230749346 | 1.13699574206438 | 1.19322821036799 | 1.06710864697714 |
| chr12 | 122983924 | 122983927 | - | 3.01188173062009 | 2.27399148412876 | 4.77291284147196 | 1.06710864697714 |
| chr12 | 122984505 | 122984522 | - | 1.00396057687336 | 3.41098722619315 | 11.9322821036799 | 3.20132594093141 |
| chr12 | 122984650 | 122984660 | - | 5.01980288436682 | 2.27399148412876 | 2.38645642073598 | 2.13421729395427 |
| chr12 | 123084588 | 123084589 | + | 2.00792115374673 | 11.3699574206438 | 9.54582568294392 | 1.06710864697714 |
| chr12 | 123154471 | 123154517 | - | 2.00792115374673 | 4.54798296825753 | 15.5119667347839 | 7.46976052883995 |
| chr12 | 123154622 | 123154629 | - | 1.00396057687336 | 1.13699574206438 | 4.77291284147196 | 9.60397782279422 |
| chr12 | 123154876 | 123154877 | - | 1.00396057687336 | 2.27399148412876 | 1.19322821036799 | 4.26843458790854 |
| chr12 | 123155020 | 123155060 | - | 32.1267384599477 | 46.6168254246397 | 42.9562155732476 | 21.3421729395427 |
| chr12 | 123257770 | 123257775 | + | 1.00396057687336 | 1.13699574206438 | 2.38645642073598 | 1.06710864697714 |
| chr12 | 123257804 | 123257863 | + | 17.0673298068472 | 45.4798296825753 | 20.2848795762558 | 21.3421729395427 |
| chr12 | 123261120 | 123261157 | - | 7.02772403811355 | 6.82197445238629 | 1.19322821036799 | 2.13421729395427 |
| chr12 | 123261269 | 123261371 | - | 42.1663442286813 | 36.3838637460602 | 11.9322821036799 | 9.60397782279422 |
| chr12 | 123261481 | 123261485 | - | 8.03168461498692 | 5.68497871032191 | 7.15936926220794 | 7.46976052883995 |
| chr12 | 123261527 | 123261565 | - | 20.0792115374673 | 17.0549361309657 | 4.77291284147196 | 8.53686917581709 |
| chr12 | 123261605 | 123261694 | - | 13.0514874993537 | 6.82197445238629 | 1.19322821036799 | 1.06710864697714 |
| chr12 | 123291196 | 123291199 | - | 2.00792115374673 | 1.13699574206438 | 3.57968463110397 | 7.46976052883995 |
| chr12 | 123291530 | 123291535 | - | 4.01584230749346 | 2.27399148412876 | 8.35259747257593 | 11.7381951167485 |
| chr12 | 123292176 | 123292180 | - | 1.00396057687336 | 1.13699574206438 | 16.7051949451519 | 4.26843458790854 |
| chr12 | 123292242 | 123292258 | - | 1.00396057687336 | 2.27399148412876 | 13.1255103140479 | 3.20132594093141 |
| chr12 | 123292366 | 123292369 | - | 7.02772403811355 | 1.13699574206438 | 14.3187385244159 | 9.60397782279422 |
| chr12 | 123292885 | 123292888 | - | 2.00792115374673 | 1.13699574206438 | 1.19322821036799 | 5.33554323488568 |
| chr12 | 123292981 | 123292984 | - | 1.00396057687336 | 4.54798296825753 | 3.57968463110397 | 1.06710864697714 |
| chr12 | 123294846 | 123294868 | - | 1.00396057687336 | 2.27399148412876 | 8.35259747257593 | 1.06710864697714 |
| chr12 | 123294952 | 123295099 | - | 14.0554480762271 | 12.5069531627082 | 75.1733772531833 | 34.1474767032683 |
| chr12 | 123295170 | 123295186 | - | 21.0831721143407 | 6.82197445238629 | 22.6713359969918 | 32.0132594093141 |
| chr12 | 123295213 | 123295236 | - | 6.02376346124019 | 1.13699574206438 | 5.96614105183995 | 5.33554323488568 |
| chr12 | 123295293 | 123295311 | - | 1.00396057687336 | 3.41098722619315 | 3.57968463110397 | 6.40265188186281 |
| chr12 | 123295479 | 123295500 | - | 3.01188173062009 | 1.13699574206438 | 5.96614105183995 | 4.26843458790854 |
| chr12 | 123295637 | 123295655 | - | 7.02772403811355 | 2.27399148412876 | 3.57968463110397 | 12.8053037637256 |
| chr12 | 123295722 | 123295735 | - | 2.00792115374673 | 1.13699574206438 | 5.96614105183995 | 8.53686917581709 |
| chr12 | 123295837 | 123295842 | - | 2.00792115374673 | 1.13699574206438 | 10.7390538933119 | 7.46976052883995 |
| chr12 | 123363867 | 123363920 | + | 8.03168461498692 | 9.09596593651506 | 1.19322821036799 | 5.33554323488568 |
| chr12 | 123364946 | 123364968 | + | 4.01584230749346 | 5.68497871032191 | 1.19322821036799 | 1.06710864697714 |
| chr12 | 123408028 | 123408043 | + | 3.01188173062009 | 1.13699574206438 | 2.38645642073598 | 6.40265188186281 |
| chr12 | 123408146 | 123408153 | + | 18.0712903837206 | 17.0549361309657 | 25.0577924177278 | 41.6172372321083 |
| chr12 | 123408231 | 123408237 | + | 4.01584230749346 | 1.13699574206438 | 1.19322821036799 | 1.06710864697714 |
| chr12 | 123408295 | 123408296 | + | 3.01188173062009 | 5.68497871032191 | 11.9322821036799 | 12.8053037637256 |
| chr12 | 123408317 | 123408356 | + | 13.0514874993537 | 15.9179403889013 | 21.4781077866238 | 32.0132594093141 |
| chr12 | 123409130 | 123409142 | + | 1.00396057687336 | 4.54798296825753 | 2.38645642073598 | 1.06710864697714 |
| chr12 | 123415578 | 123415587 | - | 2.00792115374673 | 3.41098722619315 | 3.57968463110397 | 3.20132594093141 |
| chr12 | 123415629 | 123415636 | - | 7.02772403811355 | 2.27399148412876 | 1.19322821036799 | 3.20132594093141 |
| chr12 | 123466330 | 123466369 | + | 51.2019894205416 | 60.2607743294122 | 16.7051949451519 | 28.8119334683827 |
| chr12 | 123472334 | 123472349 | - | 4.01584230749346 | 2.27399148412876 | 2.38645642073598 | 3.20132594093141 |
| chr12 | 123596677 | 123596801 | + | 9.03564519186028 | 10.2329616785794 | 10.7390538933119 | 9.60397782279422 |
| chr12 | 123596822 | 123596932 | + | 5.01980288436682 | 19.3289276150945 | 17.8984231555198 | 5.33554323488568 |
| chr12 | 123596955 | 123597000 | + | 6.02376346124019 | 6.82197445238629 | 11.9322821036799 | 9.60397782279422 |
| chr12 | 123597325 | 123597338 | + | 2.00792115374673 | 3.41098722619315 | 1.19322821036799 | 3.20132594093141 |
| chr12 | 123597374 | 123597454 | + | 11.043566345607 | 1.13699574206438 | 3.57968463110397 | 4.26843458790854 |
| chr12 | 123597581 | 123597594 | + | 1.00396057687336 | 3.41098722619315 | 5.96614105183995 | 11.7381951167485 |
| chr12 | 123597625 | 123597666 | + | 14.0554480762271 | 14.780944646837 | 31.0239334695677 | 107.777973344691 |
| chr12 | 123597688 | 123597721 | + | 7.02772403811355 | 2.27399148412876 | 8.35259747257593 | 4.26843458790854 |
| chr12 | 123597739 | 123597765 | + | 3.01188173062009 | 3.41098722619315 | 4.77291284147196 | 1.06710864697714 |
| chr12 | 123597782 | 123597837 | + | 9.03564519186028 | 14.780944646837 | 19.0916513658878 | 18.1408469986113 |
| chr12 | 123597872 | 123597897 | + | 20.0792115374673 | 14.780944646837 | 7.15936926220794 | 22.4092815865198 |
| chr12 | 123597924 | 123597969 | + | 17.0673298068472 | 31.8358807778027 | 19.0916513658878 | 26.6777161744284 |
| chr12 | 123597992 | 123598120 | + | 367.449571135651 | 695.841394143402 | 311.432562906045 | 241.166554216833 |
| chr12 | 123598344 | 123598352 | + | 1.00396057687336 | 2.27399148412876 | 2.38645642073598 | 3.20132594093141 |
| chr12 | 123598388 | 123598390 | + | 6.02376346124019 | 1.13699574206438 | 3.57968463110397 | 1.06710864697714 |
| chr12 | 123598428 | 123598431 | + | 2.00792115374673 | 1.13699574206438 | 1.19322821036799 | 2.13421729395427 |
| chr12 | 123598490 | 123598501 | + | 7.02772403811355 | 5.68497871032191 | 16.7051949451519 | 8.53686917581709 |
| chr12 | 123620421 | 123620444 | + | 2.00792115374673 | 6.82197445238629 | 10.7390538933119 | 16.006629704657 |
| chr12 | 123620846 | 123620891 | + | 5.01980288436682 | 7.95897019445067 | 3.57968463110397 | 5.33554323488568 |
| chr12 | 123621096 | 123621127 | - | 7.02772403811355 | 11.3699574206438 | 1.19322821036799 | 6.40265188186281 |
| chr12 | 123621173 | 123621182 | - | 2.00792115374673 | 4.54798296825753 | 1.19322821036799 | 3.20132594093141 |
| chr12 | 123621331 | 123621350 | - | 4.01584230749346 | 4.54798296825753 | 5.96614105183995 | 5.33554323488568 |
| chr12 | 123621390 | 123621394 | - | 1.00396057687336 | 6.82197445238629 | 4.77291284147196 | 7.46976052883995 |
| chr12 | 123626424 | 123626426 | - | 5.01980288436682 | 1.13699574206438 | 4.77291284147196 | 4.26843458790854 |
| chr12 | 123660399 | 123660418 | + | 1.00396057687336 | 1.13699574206438 | 4.77291284147196 | 3.20132594093141 |
| chr12 | 123662192 | 123662202 | + | 2.00792115374673 | 2.27399148412876 | 2.38645642073598 | 6.40265188186281 |
| chr12 | 123662254 | 123662259 | + | 3.01188173062009 | 7.95897019445067 | 2.38645642073598 | 5.33554323488568 |
| chr12 | 123662468 | 123662478 | + | 1.00396057687336 | 3.41098722619315 | 1.19322821036799 | 2.13421729395427 |
| chr12 | 123758301 | 123758317 | + | 5.01980288436682 | 3.41098722619315 | 13.1255103140479 | 10.6710864697714 |
| chr12 | 123758436 | 123758449 | + | 1.00396057687336 | 3.41098722619315 | 5.96614105183995 | 6.40265188186281 |
| chr12 | 123761615 | 123761635 | + | 5.01980288436682 | 10.2329616785794 | 1.19322821036799 | 9.60397782279422 |
| chr12 | 124013483 | 124013499 | + | 1.00396057687336 | 1.13699574206438 | 1.19322821036799 | 1.06710864697714 |
| chr12 | 124014177 | 124014255 | + | 442.746614401154 | 442.291343663045 | 204.042023972926 | 202.750642925656 |
| chr12 | 124014295 | 124014318 | + | 20.0792115374673 | 11.3699574206438 | 3.57968463110397 | 14.9395210576799 |
| chr12 | 124014345 | 124014396 | + | 56.2217923049084 | 63.6717615556054 | 35.7968463110397 | 92.8384522870108 |
| chr12 | 124014492 | 124014508 | + | 3.01188173062009 | 1.13699574206438 | 5.96614105183995 | 3.20132594093141 |
| chr12 | 124014556 | 124014626 | + | 4.01584230749346 | 6.82197445238629 | 10.7390538933119 | 17.0737383516342 |
| chr12 | 124014647 | 124014712 | + | 5.01980288436682 | 10.2329616785794 | 14.3187385244159 | 12.8053037637256 |
| chr12 | 124014842 | 124014847 | + | 16.0633692299738 | 10.2329616785794 | 8.35259747257593 | 4.26843458790854 |
| chr12 | 124014940 | 124014943 | + | 2.00792115374673 | 3.41098722619315 | 1.19322821036799 | 7.46976052883995 |
| chr12 | 124015038 | 124015055 | + | 3.01188173062009 | 3.41098722619315 | 4.77291284147196 | 2.13421729395427 |
| chr12 | 124015287 | 124015298 | + | 11.043566345607 | 31.8358807778027 | 2.38645642073598 | 6.40265188186281 |
| chr12 | 124324446 | 124324490 | - | 12.0475269224804 | 14.780944646837 | 10.7390538933119 | 11.7381951167485 |
| chr12 | 124324517 | 124324520 | - | 2.00792115374673 | 1.13699574206438 | 1.19322821036799 | 4.26843458790854 |
| chr12 | 124324541 | 124324558 | - | 1.00396057687336 | 5.68497871032191 | 1.19322821036799 | 12.8053037637256 |
| chr12 | 124324600 | 124324622 | - | 1.00396057687336 | 1.13699574206438 | 4.77291284147196 | 5.33554323488568 |
| chr12 | 124324682 | 124324696 | - | 1.00396057687336 | 2.27399148412876 | 5.96614105183995 | 1.06710864697714 |
| chr12 | 124324717 | 124324777 | - | 18.0712903837206 | 15.9179403889013 | 44.1494437836156 | 71.4962793474681 |
| chr12 | 124324803 | 124324866 | - | 47.1861471130481 | 38.657855230189 | 69.2072362013434 | 53.3554323488568 |
| chr12 | 124325086 | 124325101 | - | 5.01980288436682 | 4.54798296825753 | 5.96614105183995 | 5.33554323488568 |
| chr12 | 124325133 | 124325151 | - | 6.02376346124019 | 3.41098722619315 | 7.15936926220794 | 4.26843458790854 |
| chr12 | 124325169 | 124325211 | - | 3.01188173062009 | 12.5069531627082 | 11.9322821036799 | 30.9461507623369 |
| chr12 | 124325370 | 124325384 | - | 43.1703048055547 | 20.4659233571589 | 40.5697591525116 | 49.0869977609482 |
| chr12 | 124777651 | 124777779 | - | 34.1346596136944 | 35.2468680039958 | 28.6374770488317 | 53.3554323488568 |
| chr12 | 124778528 | 124778578 | - | 2.00792115374673 | 1.13699574206438 | 2.38645642073598 | 3.20132594093141 |
| chr12 | 124785879 | 124785893 | - | 11.043566345607 | 4.54798296825753 | 8.35259747257593 | 2.13421729395427 |
| chr12 | 124785945 | 124785975 | - | 6.02376346124019 | 3.41098722619315 | 15.5119667347839 | 1.06710864697714 |
| chr12 | 124911664 | 124911770 | - | 887.501149956054 | 1003.96724024285 | 451.0402635191 | 579.439995308585 |
| chr12 | 124946845 | 124946861 | - | 2.00792115374673 | 3.41098722619315 | 2.38645642073598 | 2.13421729395427 |
| chr12 | 125025443 | 125025449 | + | 7.02772403811355 | 2.27399148412876 | 2.38645642073598 | 2.13421729395427 |
| chr12 | 125025491 | 125025493 | + | 19.0752509605939 | 10.2329616785794 | 1.19322821036799 | 3.20132594093141 |
| chr12 | 125025616 | 125025655 | + | 14.0554480762271 | 13.6439489047726 | 5.96614105183995 | 6.40265188186281 |
| chr12 | 125025822 | 125025826 | + | 15.0594086531005 | 6.82197445238629 | 10.7390538933119 | 3.20132594093141 |
| chr12 | 125026297 | 125026299 | + | 1.00396057687336 | 1.13699574206438 | 1.19322821036799 | 5.33554323488568 |
| chr12 | 125028921 | 125028922 | + | 16.0633692299738 | 12.5069531627082 | 3.57968463110397 | 9.60397782279422 |
| chr12 | 125029143 | 125029153 | + | 13.0514874993537 | 2.27399148412876 | 7.15936926220794 | 5.33554323488568 |
| chr12 | 125030750 | 125030759 | + | 7.02772403811355 | 2.27399148412876 | 2.38645642073598 | 3.20132594093141 |
| chr12 | 125143261 | 125143277 | + | 12.0475269224804 | 14.780944646837 | 2.38645642073598 | 7.46976052883995 |
| chr12 | 127148543 | 127148549 | + | 1.00396057687336 | 2.27399148412876 | 4.77291284147196 | 3.20132594093141 |
| chr12 | 128244545 | 128244546 | + | 5.01980288436682 | 5.68497871032191 | 4.77291284147196 | 2.13421729395427 |
| chr12 | 128793863 | 128793881 | - | 3.01188173062009 | 2.27399148412876 | 1.19322821036799 | 4.26843458790854 |
| chr12 | 128793980 | 128793987 | - | 3.01188173062009 | 1.13699574206438 | 1.19322821036799 | 1.06710864697714 |
| chr12 | 130668442 | 130668443 | + | 4.01584230749346 | 3.41098722619315 | 14.3187385244159 | 12.8053037637256 |
| chr12 | 130789646 | 130789648 | - | 4.01584230749346 | 11.3699574206438 | 7.15936926220794 | 10.6710864697714 |
| chr12 | 130875882 | 130875905 | + | 199.7881547978 | 136.439489047726 | 133.641559561215 | 329.736571915935 |
| chr12 | 130875963 | 130875990 | + | 1559.15077588433 | 992.597282822205 | 1319.710400667 | 1766.06481074716 |
| chr12 | 130876079 | 130876105 | + | 201.796075951546 | 193.289276150945 | 424.789242891004 | 422.575024202946 |
| chr12 | 130876133 | 130876178 | + | 641.53080862208 | 619.662679425088 | 762.472826425145 | 669.077121654664 |
| chr12 | 130876558 | 130876576 | + | 20.0792115374673 | 23.876910583352 | 7.15936926220794 | 14.9395210576799 |
| chr12 | 130876601 | 130876618 | + | 12.0475269224804 | 17.0549361309657 | 1.19322821036799 | 1.06710864697714 |
| chr12 | 130876698 | 130876759 | + | 40.1584230749346 | 47.753821166704 | 36.9900745214077 | 14.9395210576799 |
| chr12 | 130876782 | 130876872 | + | 8.03168461498692 | 29.5618892936739 | 22.6713359969918 | 10.6710864697714 |
| chr12 | 130877314 | 130877322 | + | 14.0554480762271 | 18.1919318730301 | 5.96614105183995 | 6.40265188186281 |
| chr12 | 130877563 | 130877613 | + | 24.0950538449607 | 79.5897019445067 | 10.7390538933119 | 9.60397782279422 |
| chr12 | 131799503 | 131799504 | + | 5.01980288436682 | 3.41098722619315 | 1.19322821036799 | 3.20132594093141 |
| chr12 | 131799525 | 131799555 | + | 1.00396057687336 | 3.41098722619315 | 4.77291284147196 | 4.26843458790854 |
| chr12 | 131799714 | 131799719 | + | 13.0514874993537 | 35.2468680039958 | 34.6036181006717 | 23.476390233497 |
| chr12 | 131921556 | 131921582 | + | 7.02772403811355 | 10.2329616785794 | 3.57968463110397 | 2.13421729395427 |
| chr12 | 131921627 | 131921631 | + | 9.03564519186028 | 10.2329616785794 | 1.19322821036799 | 2.13421729395427 |
| chr12 | 131921752 | 131921774 | + | 4.01584230749346 | 2.27399148412876 | 2.38645642073598 | 1.06710864697714 |
| chr12 | 131922347 | 131922349 | + | 1.00396057687336 | 2.27399148412876 | 2.38645642073598 | 1.06710864697714 |
| chr12 | 131922847 | 131922872 | + | 25.0990144218341 | 46.6168254246397 | 7.15936926220794 | 8.53686917581709 |
| chr12 | 131922894 | 131923056 | + | 185.732706721572 | 176.234340019979 | 23.8645642073598 | 33.0803680562912 |
| chr12 | 131943696 | 131943751 | + | 24.0950538449607 | 44.3428339405109 | 11.9322821036799 | 14.9395210576799 |
| chr12 | 131943818 | 131943839 | + | 3.01188173062009 | 9.09596593651506 | 1.19322821036799 | 1.06710864697714 |
| chr12 | 132078832 | 132078841 | + | 2.00792115374673 | 3.41098722619315 | 2.38645642073598 | 2.13421729395427 |
| chr12 | 132080237 | 132080261 | + | 3.01188173062009 | 2.27399148412876 | 1.19322821036799 | 3.20132594093141 |
| chr12 | 132136632 | 132136658 | - | 2.00792115374673 | 4.54798296825753 | 2.38645642073598 | 2.13421729395427 |
| chr12 | 132566393 | 132566399 | + | 1.00396057687336 | 2.27399148412876 | 5.96614105183995 | 2.13421729395427 |
| chr12 | 132583910 | 132583936 | + | 15.0594086531005 | 4.54798296825753 | 2.38645642073598 | 4.26843458790854 |
| chr12 | 132583987 | 132583988 | + | 2.00792115374673 | 1.13699574206438 | 1.19322821036799 | 2.13421729395427 |
| chr12 | 132584010 | 132584031 | + | 3.01188173062009 | 3.41098722619315 | 2.38645642073598 | 4.26843458790854 |
| chr12 | 132584236 | 132584268 | + | 9.03564519186028 | 1.13699574206438 | 3.57968463110397 | 3.20132594093141 |
| chr12 | 132623801 | 132623812 | - | 8.03168461498692 | 5.68497871032191 | 11.9322821036799 | 14.9395210576799 |
| chr12 | 132623835 | 132623896 | - | 11.043566345607 | 10.2329616785794 | 5.96614105183995 | 18.1408469986113 |
| chr12 | 132624086 | 132624089 | - | 1.00396057687336 | 1.13699574206438 | 8.35259747257593 | 5.33554323488568 |
| chr12 | 132704679 | 132704776 | + | 12.0475269224804 | 6.82197445238629 | 2.38645642073598 | 10.6710864697714 |
| chr12 | 132704803 | 132704827 | + | 3.01188173062009 | 1.13699574206438 | 2.38645642073598 | 1.06710864697714 |
| chr12 | 132704886 | 132704889 | + | 3.01188173062009 | 4.54798296825753 | 1.19322821036799 | 1.06710864697714 |
| chr12 | 132721884 | 132721901 | + | 11.043566345607 | 5.68497871032191 | 10.7390538933119 | 18.1408469986113 |
| chr12 | 132722626 | 132722629 | + | 2.00792115374673 | 1.13699574206438 | 2.38645642073598 | 2.13421729395427 |
| chr12 | 132722655 | 132722664 | + | 3.01188173062009 | 2.27399148412876 | 2.38645642073598 | 7.46976052883995 |
| chr12 | 132725881 | 132725882 | - | 1.00396057687336 | 1.13699574206438 | 1.19322821036799 | 3.20132594093141 |
| chr12 | 132726525 | 132726537 | - | 5.01980288436682 | 6.82197445238629 | 1.19322821036799 | 1.06710864697714 |
| chr12 | 132726566 | 132726611 | - | 15.0594086531005 | 13.6439489047726 | 9.54582568294392 | 4.26843458790854 |
| chr12 | 132726678 | 132726710 | - | 34.1346596136944 | 50.0278126508328 | 14.3187385244159 | 26.6777161744284 |
| chr12 | 132726731 | 132726753 | - | 3.01188173062009 | 2.27399148412876 | 2.38645642073598 | 4.26843458790854 |
| chr12 | 132726757 | 132726783 | - | 10.0396057687336 | 20.4659233571589 | 5.96614105183995 | 4.26843458790854 |
| chr12 | 132726854 | 132726883 | - | 3.01188173062009 | 7.95897019445067 | 4.77291284147196 | 4.26843458790854 |
| chr12 | 132726983 | 132726986 | - | 2.00792115374673 | 1.13699574206438 | 2.38645642073598 | 2.13421729395427 |
| chr12 | 132727193 | 132727199 | - | 2.00792115374673 | 6.82197445238629 | 1.19322821036799 | 3.20132594093141 |
| chr12 | 132727222 | 132727270 | - | 9.03564519186028 | 10.2329616785794 | 4.77291284147196 | 3.20132594093141 |
| chr12 | 132768937 | 132768965 | - | 3.01188173062009 | 4.54798296825753 | 10.7390538933119 | 5.33554323488568 |
| chr12 | 132769012 | 132769042 | - | 2.00792115374673 | 3.41098722619315 | 1.19322821036799 | 4.26843458790854 |
| chr12 | 132769099 | 132769145 | - | 6.02376346124019 | 7.95897019445067 | 11.9322821036799 | 16.006629704657 |
| chr12 | 132772265 | 132772266 | - | 1.00396057687336 | 3.41098722619315 | 2.38645642073598 | 2.13421729395427 |
| chr12 | 133059248 | 133059281 | + | 5.01980288436682 | 12.5069531627082 | 2.38645642073598 | 4.26843458790854 |
| chr12 | 133059554 | 133059574 | + | 1.00396057687336 | 4.54798296825753 | 1.19322821036799 | 1.06710864697714 |
| chr12 | 133118660 | 133118663 | - | 6.02376346124019 | 10.2329616785794 | 2.38645642073598 | 4.26843458790854 |
| chr12 | 133118784 | 133118785 | - | 2.00792115374673 | 3.41098722619315 | 1.19322821036799 | 1.06710864697714 |
| chr13 | 20086369 | 20086379 | + | 3.01188173062009 | 6.82197445238629 | 8.35259747257593 | 5.33554323488568 |
| chr13 | 20088212 | 20088218 | + | 2.00792115374673 | 10.2329616785794 | 3.57968463110397 | 10.6710864697714 |
| chr13 | 20403700 | 20403746 | - | 2.00792115374673 | 3.41098722619315 | 2.38645642073598 | 3.20132594093141 |
| chr13 | 20779238 | 20779249 | - | 4.01584230749346 | 1.13699574206438 | 1.19322821036799 | 4.26843458790854 |
| chr13 | 20779312 | 20779359 | - | 19.0752509605939 | 11.3699574206438 | 11.9322821036799 | 17.0737383516342 |
| chr13 | 20779501 | 20779511 | - | 6.02376346124019 | 6.82197445238629 | 3.57968463110397 | 11.7381951167485 |
| chr13 | 20780255 | 20780257 | - | 1.00396057687336 | 1.13699574206438 | 3.57968463110397 | 2.13421729395427 |
| chr13 | 20783056 | 20783084 | - | 14.0554480762271 | 14.780944646837 | 15.5119667347839 | 13.8724124107028 |
| chr13 | 20973972 | 20973979 | - | 12.0475269224804 | 22.7399148412876 | 3.57968463110397 | 1.06710864697714 |
| chr13 | 20974782 | 20974826 | - | 11.043566345607 | 21.6029190992233 | 3.57968463110397 | 4.26843458790854 |
| chr13 | 21147304 | 21147424 | + | 50.1980288436682 | 45.4798296825753 | 32.2171616799357 | 59.7580842307196 |
| chr13 | 21147471 | 21147512 | + | 17.0673298068472 | 19.3289276150945 | 11.9322821036799 | 9.60397782279422 |
| chr13 | 21147541 | 21147582 | + | 14.0554480762271 | 20.4659233571589 | 22.6713359969918 | 13.8724124107028 |
| chr13 | 21147601 | 21147703 | + | 22.087132691214 | 22.7399148412876 | 29.8307052591997 | 30.9461507623369 |
| chr13 | 21147923 | 21147945 | + | 2.00792115374673 | 1.13699574206438 | 9.54582568294392 | 1.06710864697714 |
| chr13 | 21148024 | 21148096 | + | 8.03168461498692 | 10.2329616785794 | 13.1255103140479 | 6.40265188186281 |
| chr13 | 21148130 | 21148136 | + | 6.02376346124019 | 10.2329616785794 | 4.77291284147196 | 3.20132594093141 |
| chr13 | 21148804 | 21148806 | + | 1.00396057687336 | 1.13699574206438 | 1.19322821036799 | 1.06710864697714 |
| chr13 | 21153963 | 21153964 | - | 3.01188173062009 | 4.54798296825753 | 2.38645642073598 | 1.06710864697714 |
| chr13 | 21154049 | 21154052 | - | 2.00792115374673 | 6.82197445238629 | 3.57968463110397 | 13.8724124107028 |
| chr13 | 21177229 | 21177333 | + | 50.1980288436682 | 64.8087572976698 | 33.4103898903037 | 48.0198891139711 |
| chr13 | 21374065 | 21374079 | - | 4.01584230749346 | 10.2329616785794 | 4.77291284147196 | 9.60397782279422 |
| chr13 | 21376185 | 21376208 | - | 6.02376346124019 | 4.54798296825753 | 1.19322821036799 | 2.13421729395427 |
| chr13 | 21492716 | 21492726 | - | 6.02376346124019 | 4.54798296825753 | 4.77291284147196 | 6.40265188186281 |
| chr13 | 21493037 | 21493040 | - | 1.00396057687336 | 6.82197445238629 | 1.19322821036799 | 3.20132594093141 |
| chr13 | 23328902 | 23328946 | - | 35.1386201905678 | 31.8358807778027 | 57.2749540976635 | 23.476390233497 |
| chr13 | 23328968 | 23329031 | - | 29.1148567293276 | 15.9179403889013 | 53.6952694665595 | 35.2145853502455 |
| chr13 | 23329053 | 23329066 | - | 12.0475269224804 | 2.27399148412876 | 4.77291284147196 | 9.60397782279422 |
| chr13 | 23329094 | 23329100 | - | 5.01980288436682 | 4.54798296825753 | 13.1255103140479 | 6.40265188186281 |
| chr13 | 23329123 | 23329128 | - | 3.01188173062009 | 2.27399148412876 | 7.15936926220794 | 3.20132594093141 |
| chr13 | 23329152 | 23329170 | - | 8.03168461498692 | 6.82197445238629 | 15.5119667347839 | 5.33554323488568 |
| chr13 | 23329266 | 23329328 | - | 31.1227778830743 | 18.1919318730301 | 22.6713359969918 | 30.9461507623369 |
| chr13 | 23329440 | 23329462 | - | 14.0554480762271 | 2.27399148412876 | 8.35259747257593 | 22.4092815865198 |
| chr13 | 23329726 | 23329734 | - | 4.01584230749346 | 3.41098722619315 | 9.54582568294392 | 3.20132594093141 |
| chr13 | 23329768 | 23329786 | - | 4.01584230749346 | 1.13699574206438 | 7.15936926220794 | 4.26843458790854 |
| chr13 | 23329808 | 23329854 | - | 6.02376346124019 | 6.82197445238629 | 13.1255103140479 | 10.6710864697714 |
| chr13 | 23329893 | 23329936 | - | 27.1069355755808 | 1.13699574206438 | 15.5119667347839 | 10.6710864697714 |
| chr13 | 23329976 | 23329978 | - | 3.01188173062009 | 2.27399148412876 | 4.77291284147196 | 4.26843458790854 |
| chr13 | 23730211 | 23730223 | - | 13.0514874993537 | 18.1919318730301 | 7.15936926220794 | 11.7381951167485 |
| chr13 | 24420955 | 24420986 | - | 3.01188173062009 | 3.41098722619315 | 2.38645642073598 | 3.20132594093141 |
| chr13 | 24883183 | 24883208 | - | 1.00396057687336 | 4.54798296825753 | 5.96614105183995 | 9.60397782279422 |
| chr13 | 25247394 | 25247402 | - | 4.01584230749346 | 2.27399148412876 | 1.19322821036799 | 2.13421729395427 |
| chr13 | 25247608 | 25247611 | - | 2.00792115374673 | 4.54798296825753 | 2.38645642073598 | 2.13421729395427 |
| chr13 | 25248622 | 25248623 | - | 1.00396057687336 | 1.13699574206438 | 1.19322821036799 | 1.06710864697714 |
| chr13 | 25333819 | 25333839 | + | 1.00396057687336 | 1.13699574206438 | 16.7051949451519 | 4.26843458790854 |
| chr13 | 25333959 | 25334007 | + | 2.00792115374673 | 5.68497871032191 | 17.8984231555198 | 4.26843458790854 |
| chr13 | 25336175 | 25336177 | + | 2.00792115374673 | 1.13699574206438 | 3.57968463110397 | 2.13421729395427 |
| chr13 | 25341997 | 25342001 | + | 4.01584230749346 | 1.13699574206438 | 2.38645642073598 | 9.60397782279422 |
| chr13 | 26212806 | 26212843 | - | 1.00396057687336 | 2.27399148412876 | 3.57968463110397 | 1.06710864697714 |
| chr13 | 26212885 | 26212887 | - | 7.02772403811355 | 13.6439489047726 | 1.19322821036799 | 3.20132594093141 |
| chr13 | 26404980 | 26404984 | + | 3.01188173062009 | 1.13699574206438 | 3.57968463110397 | 4.26843458790854 |
| chr13 | 26405067 | 26405075 | + | 3.01188173062009 | 5.68497871032191 | 4.77291284147196 | 6.40265188186281 |
| chr13 | 27067005 | 27067014 | - | 3.01188173062009 | 4.54798296825753 | 5.96614105183995 | 5.33554323488568 |
| chr13 | 27067753 | 27067759 | - | 2.00792115374673 | 1.13699574206438 | 3.57968463110397 | 2.13421729395427 |
| chr13 | 27067976 | 27067995 | - | 9.03564519186028 | 14.780944646837 | 8.35259747257593 | 4.26843458790854 |
| chr13 | 27068421 | 27068430 | - | 1.00396057687336 | 4.54798296825753 | 3.57968463110397 | 1.06710864697714 |
| chr13 | 27069052 | 27069056 | - | 4.01584230749346 | 3.41098722619315 | 7.15936926220794 | 2.13421729395427 |
| chr13 | 27255064 | 27255066 | + | 1.00396057687336 | 2.27399148412876 | 3.57968463110397 | 1.06710864697714 |
| chr13 | 27435619 | 27435659 | + | 74.293082688629 | 70.4937360079917 | 119.322821036799 | 152.59653651773 |
| chr13 | 27435664 | 27435685 | - | 1.00396057687336 | 3.41098722619315 | 3.57968463110397 | 1.06710864697714 |
| chr13 | 27623269 | 27623429 | + | 15.0594086531005 | 29.5618892936739 | 26.2510206280958 | 13.8724124107028 |
| chr13 | 27665983 | 27666051 | + | 10.0396057687336 | 12.5069531627082 | 38.1833027317757 | 33.0803680562912 |
| chr13 | 27666125 | 27666157 | + | 4.01584230749346 | 1.13699574206438 | 7.15936926220794 | 4.26843458790854 |
| chr13 | 27666529 | 27666543 | + | 1.00396057687336 | 2.27399148412876 | 4.77291284147196 | 1.06710864697714 |
| chr13 | 27667139 | 27667154 | + | 2.00792115374673 | 1.13699574206438 | 4.77291284147196 | 2.13421729395427 |
| chr13 | 27945552 | 27945564 | + | 87.3445701879827 | 135.302493305661 | 75.1733772531833 | 36.2816939972226 |
| chr13 | 28292659 | 28292724 | + | 6.02376346124019 | 4.54798296825753 | 4.77291284147196 | 7.46976052883995 |
| chr13 | 28293416 | 28293436 | + | 18.0712903837206 | 19.3289276150945 | 51.3088130458235 | 14.9395210576799 |
| chr13 | 28294911 | 28294915 | + | 3.01188173062009 | 7.95897019445067 | 4.77291284147196 | 10.6710864697714 |
| chr13 | 28295155 | 28295158 | + | 1.00396057687336 | 14.780944646837 | 5.96614105183995 | 7.46976052883995 |
| chr13 | 28295226 | 28295253 | + | 3.01188173062009 | 7.95897019445067 | 2.38645642073598 | 1.06710864697714 |
| chr13 | 28678046 | 28678198 | + | 334.31887209883 | 214.892195250168 | 534.566238244859 | 1011.61899733432 |
| chr13 | 28678243 | 28678247 | + | 20.0792115374673 | 17.0549361309657 | 13.1255103140479 | 36.2816939972226 |
| chr13 | 28678285 | 28678304 | + | 18.0712903837206 | 19.3289276150945 | 25.0577924177278 | 26.6777161744284 |
| chr13 | 29509620 | 29509684 | - | 13.0514874993537 | 23.876910583352 | 17.8984231555198 | 6.40265188186281 |
| chr13 | 29509705 | 29510104 | - | 858.386293226727 | 1674.79472806083 | 640.76354896761 | 656.271817890938 |
| chr13 | 29510110 | 29510235 | - | 14.0554480762271 | 29.5618892936739 | 27.4442488384638 | 13.8724124107028 |
| chr13 | 29510552 | 29510558 | - | 4.01584230749346 | 4.54798296825753 | 1.19322821036799 | 2.13421729395427 |
| chr13 | 29510591 | 29510608 | - | 11.043566345607 | 23.876910583352 | 9.54582568294392 | 9.60397782279422 |
| chr13 | 29511540 | 29511579 | - | 4.01584230749346 | 9.09596593651506 | 3.57968463110397 | 4.26843458790854 |
| chr13 | 29511850 | 29511877 | - | 4.01584230749346 | 5.68497871032191 | 1.19322821036799 | 1.06710864697714 |
| chr13 | 29511906 | 29511908 | - | 2.00792115374673 | 2.27399148412876 | 2.38645642073598 | 1.06710864697714 |
| chr13 | 29512035 | 29512183 | - | 72.2851615348822 | 126.206527369146 | 65.6275515702394 | 34.1474767032683 |
| chr13 | 29512228 | 29512234 | - | 4.01584230749346 | 5.68497871032191 | 3.57968463110397 | 1.06710864697714 |
| chr13 | 29512490 | 29512549 | - | 88.3485307648561 | 167.138374083464 | 45.3426719939836 | 60.8251928776967 |
| chr13 | 29512577 | 29512724 | - | 27.1069355755808 | 62.534765813541 | 15.5119667347839 | 20.2750642925656 |
| chr13 | 29512758 | 29512830 | - | 9.03564519186028 | 7.95897019445067 | 1.19322821036799 | 5.33554323488568 |
| chr13 | 30458768 | 30458771 | - | 1.00396057687336 | 1.13699574206438 | 2.38645642073598 | 1.06710864697714 |
| chr13 | 30459956 | 30459979 | - | 3.01188173062009 | 3.41098722619315 | 5.96614105183995 | 1.06710864697714 |
| chr13 | 30460087 | 30460111 | - | 4.01584230749346 | 5.68497871032191 | 4.77291284147196 | 8.53686917581709 |
| chr13 | 30460182 | 30460201 | - | 6.02376346124019 | 7.95897019445067 | 5.96614105183995 | 13.8724124107028 |
| chr13 | 30460916 | 30460926 | - | 9.03564519186028 | 10.2329616785794 | 13.1255103140479 | 4.26843458790854 |
| chr13 | 31136038 | 31136041 | - | 3.01188173062009 | 2.27399148412876 | 4.77291284147196 | 2.13421729395427 |
| chr13 | 31136915 | 31136928 | - | 40.1584230749346 | 19.3289276150945 | 11.9322821036799 | 34.1474767032683 |
| chr13 | 31136995 | 31137000 | - | 25.0990144218341 | 7.95897019445067 | 7.15936926220794 | 19.2079556455884 |
| chr13 | 32517646 | 32517754 | - | 8.03168461498692 | 22.7399148412876 | 4.77291284147196 | 4.26843458790854 |
| chr13 | 32775039 | 32775084 | + | 13.0514874993537 | 14.780944646837 | 8.35259747257593 | 8.53686917581709 |
| chr13 | 32775645 | 32775683 | + | 1.00396057687336 | 1.13699574206438 | 1.19322821036799 | 1.06710864697714 |
| chr13 | 32784551 | 32784552 | + | 13.0514874993537 | 10.2329616785794 | 2.38645642073598 | 1.06710864697714 |
| chr13 | 33836814 | 33836816 | + | 2.00792115374673 | 1.13699574206438 | 1.19322821036799 | 1.06710864697714 |
| chr13 | 33837129 | 33837130 | + | 3.01188173062009 | 9.09596593651506 | 2.38645642073598 | 3.20132594093141 |
| chr13 | 33837173 | 33837174 | + | 5.01980288436682 | 10.2329616785794 | 1.19322821036799 | 2.13421729395427 |
| chr13 | 33837213 | 33837253 | + | 9.03564519186028 | 9.09596593651506 | 1.19322821036799 | 4.26843458790854 |
| chr13 | 33837443 | 33837464 | + | 6.02376346124019 | 2.27399148412876 | 7.15936926220794 | 4.26843458790854 |
| chr13 | 33847405 | 33847430 | + | 13.0514874993537 | 11.3699574206438 | 3.57968463110397 | 9.60397782279422 |
| chr13 | 33966301 | 33966310 | + | 4.01584230749346 | 2.27399148412876 | 1.19322821036799 | 2.13421729395427 |
| chr13 | 36303759 | 36303771 | - | 3.01188173062009 | 4.54798296825753 | 3.57968463110397 | 4.26843458790854 |
| chr13 | 36303824 | 36303827 | - | 2.00792115374673 | 2.27399148412876 | 4.77291284147196 | 1.06710864697714 |
| chr13 | 36304085 | 36304134 | - | 33.130699036821 | 36.3838637460602 | 19.0916513658878 | 19.2079556455884 |
| chr13 | 36304158 | 36304226 | - | 88.3485307648561 | 63.6717615556054 | 62.0478669391355 | 73.6304966414224 |
| chr13 | 36304294 | 36304364 | - | 67.2653586505154 | 67.0827487817985 | 44.1494437836156 | 45.8856718200168 |
| chr13 | 36828198 | 36828201 | + | 2.00792115374673 | 2.27399148412876 | 2.38645642073598 | 1.06710864697714 |
| chr13 | 36828476 | 36828499 | + | 12.0475269224804 | 7.95897019445067 | 1.19322821036799 | 1.06710864697714 |
| chr13 | 36949902 | 36949972 | - | 12.0475269224804 | 12.5069531627082 | 19.0916513658878 | 6.40265188186281 |
| chr13 | 37009266 | 37009304 | + | 5.01980288436682 | 18.1919318730301 | 8.35259747257593 | 10.6710864697714 |
| chr13 | 37009351 | 37009363 | + | 3.01188173062009 | 7.95897019445067 | 2.38645642073598 | 9.60397782279422 |
| chr13 | 37009484 | 37009511 | - | 2.00792115374673 | 7.95897019445067 | 9.54582568294392 | 6.40265188186281 |
| chr13 | 38361020 | 38361068 | + | 9.03564519186028 | 15.9179403889013 | 26.2510206280958 | 29.8790421153598 |
| chr13 | 38361239 | 38361283 | + | 3.01188173062009 | 9.09596593651506 | 4.77291284147196 | 3.20132594093141 |
| chr13 | 38361442 | 38361459 | + | 5.01980288436682 | 6.82197445238629 | 3.57968463110397 | 9.60397782279422 |
| chr13 | 38361523 | 38361556 | + | 10.0396057687336 | 3.41098722619315 | 16.7051949451519 | 3.20132594093141 |
| chr13 | 38361577 | 38361653 | + | 14.0554480762271 | 13.6439489047726 | 20.2848795762558 | 8.53686917581709 |
| chr13 | 38361678 | 38361687 | + | 3.01188173062009 | 3.41098722619315 | 8.35259747257593 | 7.46976052883995 |
| chr13 | 38361725 | 38361860 | + | 50.1980288436682 | 54.5757956190903 | 70.4004644117114 | 82.1673658172394 |
| chr13 | 38361997 | 38362043 | + | 17.0673298068472 | 23.876910583352 | 27.4442488384638 | 21.3421729395427 |
| chr13 | 38362097 | 38362147 | + | 9.03564519186028 | 22.7399148412876 | 29.8307052591997 | 21.3421729395427 |
| chr13 | 38362176 | 38362201 | + | 5.01980288436682 | 6.82197445238629 | 19.0916513658878 | 13.8724124107028 |
| chr13 | 38362232 | 38362339 | + | 34.1346596136944 | 62.534765813541 | 100.231169670911 | 62.959410171651 |
| chr13 | 38362380 | 38362440 | + | 16.0633692299738 | 35.2468680039958 | 50.1155848354556 | 20.2750642925656 |
| chr13 | 38362571 | 38362592 | + | 20.0792115374673 | 17.0549361309657 | 41.7629873628796 | 37.3488026441998 |
| chr13 | 38362618 | 38362702 | + | 64.2534769198953 | 135.302493305661 | 232.679501021758 | 88.5700176991023 |
| chr13 | 39010195 | 39010196 | - | 1.00396057687336 | 2.27399148412876 | 1.19322821036799 | 1.06710864697714 |
| chr13 | 39010365 | 39010387 | - | 5.01980288436682 | 7.95897019445067 | 11.9322821036799 | 16.006629704657 |
| chr13 | 39010873 | 39010879 | - | 2.00792115374673 | 2.27399148412876 | 7.15936926220794 | 3.20132594093141 |
| chr13 | 39010926 | 39010934 | - | 1.00396057687336 | 1.13699574206438 | 1.19322821036799 | 2.13421729395427 |
| chr13 | 39010956 | 39010979 | - | 5.01980288436682 | 3.41098722619315 | 4.77291284147196 | 5.33554323488568 |
| chr13 | 39011157 | 39011160 | - | 1.00396057687336 | 1.13699574206438 | 1.19322821036799 | 1.06710864697714 |
| chr13 | 39011245 | 39011265 | - | 3.01188173062009 | 7.95897019445067 | 5.96614105183995 | 3.20132594093141 |
| chr13 | 39048810 | 39048830 | + | 1.00396057687336 | 2.27399148412876 | 1.19322821036799 | 3.20132594093141 |
| chr13 | 39049712 | 39049759 | + | 1.00396057687336 | 1.13699574206438 | 2.38645642073598 | 1.06710864697714 |
| chr13 | 39049829 | 39049844 | + | 3.01188173062009 | 3.41098722619315 | 3.57968463110397 | 1.06710864697714 |
| chr13 | 39050054 | 39050074 | + | 1.00396057687336 | 1.13699574206438 | 1.19322821036799 | 1.06710864697714 |
| chr13 | 40557823 | 40557825 | - | 3.01188173062009 | 3.41098722619315 | 2.38645642073598 | 1.06710864697714 |
| chr13 | 40729332 | 40729366 | - | 7.02772403811355 | 7.95897019445067 | 9.54582568294392 | 12.8053037637256 |
| chr13 | 40932325 | 40932331 | - | 5.01980288436682 | 6.82197445238629 | 3.57968463110397 | 6.40265188186281 |
| chr13 | 41083490 | 41083492 | + | 3.01188173062009 | 3.41098722619315 | 1.19322821036799 | 1.06710864697714 |
| chr13 | 41132903 | 41133016 | + | 28.1108961524542 | 35.2468680039958 | 17.8984231555198 | 13.8724124107028 |
| chr13 | 41191919 | 41191925 | - | 4.01584230749346 | 3.41098722619315 | 2.38645642073598 | 1.06710864697714 |
| chr13 | 41376768 | 41376773 | + | 1.00396057687336 | 4.54798296825753 | 1.19322821036799 | 4.26843458790854 |
| chr13 | 41376963 | 41376966 | + | 1.00396057687336 | 3.41098722619315 | 1.19322821036799 | 1.06710864697714 |
| chr13 | 42236062 | 42236064 | + | 9.03564519186028 | 12.5069531627082 | 14.3187385244159 | 5.33554323488568 |
| chr13 | 42236968 | 42236970 | + | 2.00792115374673 | 1.13699574206438 | 2.38645642073598 | 2.13421729395427 |
| chr13 | 42319953 | 42319955 | + | 2.00792115374673 | 2.27399148412876 | 1.19322821036799 | 4.26843458790854 |
| chr13 | 42320593 | 42320605 | + | 1.00396057687336 | 3.41098722619315 | 2.38645642073598 | 1.06710864697714 |
| chr13 | 42322773 | 42322815 | + | 3.01188173062009 | 10.2329616785794 | 4.77291284147196 | 5.33554323488568 |
| chr13 | 44433557 | 44433585 | - | 38.1505019211879 | 69.3567402659273 | 1.19322821036799 | 2.13421729395427 |
| chr13 | 44433633 | 44433675 | - | 69.2732798042621 | 169.412365567593 | 4.77291284147196 | 1.06710864697714 |
| chr13 | 44433868 | 44434090 | - | 395.560467288106 | 741.321223825977 | 39.3765309421437 | 53.3554323488568 |
| chr13 | 44434146 | 44434196 | - | 76.3010038423757 | 130.754510337404 | 10.7390538933119 | 11.7381951167485 |
| chr13 | 44434222 | 44434339 | - | 44.174265382428 | 73.9047232341848 | 5.96614105183995 | 12.8053037637256 |
| chr13 | 44434362 | 44434455 | - | 36.1425807674411 | 45.4798296825753 | 4.77291284147196 | 2.13421729395427 |
| chr13 | 44434482 | 44434532 | - | 22.087132691214 | 19.3289276150945 | 3.57968463110397 | 3.20132594093141 |
| chr13 | 45029693 | 45029730 | + | 5.01980288436682 | 3.41098722619315 | 2.38645642073598 | 2.13421729395427 |
| chr13 | 45283600 | 45283619 | + | 1.00396057687336 | 1.13699574206438 | 2.38645642073598 | 3.20132594093141 |
| chr13 | 45283653 | 45283658 | + | 6.02376346124019 | 7.95897019445067 | 1.19322821036799 | 4.26843458790854 |
| chr13 | 45283738 | 45283746 | + | 1.00396057687336 | 1.13699574206438 | 1.19322821036799 | 1.06710864697714 |
| chr13 | 45283804 | 45283823 | + | 8.03168461498692 | 11.3699574206438 | 3.57968463110397 | 10.6710864697714 |
| chr13 | 45283905 | 45283932 | + | 4.01584230749346 | 2.27399148412876 | 9.54582568294392 | 10.6710864697714 |
| chr13 | 45284027 | 45284061 | + | 2.00792115374673 | 1.13699574206438 | 1.19322821036799 | 1.06710864697714 |
| chr13 | 45334510 | 45334516 | - | 3.01188173062009 | 1.13699574206438 | 2.38645642073598 | 2.13421729395427 |
| chr13 | 45334579 | 45334586 | - | 1.00396057687336 | 3.41098722619315 | 2.38645642073598 | 3.20132594093141 |
| chr13 | 45336688 | 45336708 | - | 3.01188173062009 | 1.13699574206438 | 7.15936926220794 | 4.26843458790854 |
| chr13 | 45336874 | 45337066 | - | 42.1663442286813 | 98.9186295596012 | 34.6036181006717 | 64.0265188186281 |
| chr13 | 45337089 | 45337177 | - | 4.01584230749346 | 6.82197445238629 | 5.96614105183995 | 3.20132594093141 |
| chr13 | 45337259 | 45337281 | - | 15.0594086531005 | 11.3699574206438 | 19.0916513658878 | 14.9395210576799 |
| chr13 | 45337350 | 45337391 | - | 13.0514874993537 | 15.9179403889013 | 26.2510206280958 | 18.1408469986113 |
| chr13 | 45341547 | 45341617 | + | 42.1663442286813 | 44.3428339405109 | 58.4681823080315 | 33.0803680562912 |
| chr13 | 45391839 | 45391863 | + | 3.01188173062009 | 4.54798296825753 | 2.38645642073598 | 1.06710864697714 |
| chr13 | 45395236 | 45395316 | - | 2.00792115374673 | 5.68497871032191 | 8.35259747257593 | 1.06710864697714 |
| chr13 | 45954525 | 45954526 | - | 2.00792115374673 | 2.27399148412876 | 2.38645642073598 | 1.06710864697714 |
| chr13 | 45956656 | 45956680 | - | 17.0673298068472 | 22.7399148412876 | 13.1255103140479 | 29.8790421153598 |
| chr13 | 46052813 | 46052876 | + | 9.03564519186028 | 9.09596593651506 | 54.8884976769275 | 35.2145853502455 |
| chr13 | 46752427 | 46752454 | + | 13.0514874993537 | 13.6439489047726 | 10.7390538933119 | 4.26843458790854 |
| chr13 | 46771275 | 46771305 | - | 2.00792115374673 | 2.27399148412876 | 8.35259747257593 | 3.20132594093141 |
| chr13 | 47942685 | 47942705 | - | 3.01188173062009 | 2.27399148412876 | 2.38645642073598 | 1.06710864697714 |
| chr13 | 47942726 | 47942749 | - | 11.043566345607 | 6.82197445238629 | 1.19322821036799 | 2.13421729395427 |
| chr13 | 48046866 | 48046882 | + | 1.00396057687336 | 6.82197445238629 | 1.19322821036799 | 2.13421729395427 |
| chr13 | 48046925 | 48046969 | + | 11.043566345607 | 14.780944646837 | 11.9322821036799 | 6.40265188186281 |
| chr13 | 48047035 | 48047079 | + | 6.02376346124019 | 11.3699574206438 | 5.96614105183995 | 5.33554323488568 |
| chr13 | 48076085 | 48076088 | - | 1.00396057687336 | 9.09596593651506 | 9.54582568294392 | 5.33554323488568 |
| chr13 | 48076139 | 48076142 | - | 2.00792115374673 | 7.95897019445067 | 2.38645642073598 | 2.13421729395427 |
| chr13 | 48076299 | 48076302 | - | 8.03168461498692 | 17.0549361309657 | 23.8645642073598 | 18.1408469986113 |
| chr13 | 48076345 | 48076350 | - | 3.01188173062009 | 1.13699574206438 | 4.77291284147196 | 1.06710864697714 |
| chr13 | 48076448 | 48076460 | - | 4.01584230749346 | 11.3699574206438 | 3.57968463110397 | 1.06710864697714 |
| chr13 | 48076602 | 48076668 | - | 13.0514874993537 | 20.4659233571589 | 19.0916513658878 | 22.4092815865198 |
| chr13 | 48076713 | 48076742 | - | 1.00396057687336 | 1.13699574206438 | 1.19322821036799 | 2.13421729395427 |
| chr13 | 48076815 | 48076819 | - | 4.01584230749346 | 6.82197445238629 | 5.96614105183995 | 13.8724124107028 |
| chr13 | 48076924 | 48076929 | - | 13.0514874993537 | 25.0139063254164 | 10.7390538933119 | 7.46976052883995 |
| chr13 | 48077073 | 48077079 | - | 4.01584230749346 | 1.13699574206438 | 4.77291284147196 | 4.26843458790854 |
| chr13 | 48261137 | 48261511 | + | 336.326793252577 | 392.263531012212 | 180.177459765566 | 77.8989312293309 |
| chr13 | 48261537 | 48261655 | + | 57.2257528817818 | 72.7677274921204 | 57.2749540976635 | 18.1408469986113 |
| chr13 | 48261679 | 48261739 | + | 35.1386201905678 | 35.2468680039958 | 25.0577924177278 | 19.2079556455884 |
| chr13 | 48481655 | 48481679 | + | 17.0673298068472 | 35.2468680039958 | 4.77291284147196 | 14.9395210576799 |
| chr13 | 49207435 | 49207509 | + | 2.00792115374673 | 15.9179403889013 | 7.15936926220794 | 2.13421729395427 |
| chr13 | 49207521 | 49207555 | + | 7.02772403811355 | 10.2329616785794 | 2.38645642073598 | 2.13421729395427 |
| chr13 | 49207605 | 49207617 | + | 8.03168461498692 | 13.6439489047726 | 4.77291284147196 | 2.13421729395427 |
| chr13 | 49207873 | 49207948 | + | 33.130699036821 | 71.6307317500561 | 15.5119667347839 | 5.33554323488568 |
| chr13 | 49209356 | 49209366 | + | 5.01980288436682 | 6.82197445238629 | 3.57968463110397 | 5.33554323488568 |
| chr13 | 49491905 | 49491931 | + | 1.00396057687336 | 4.54798296825753 | 2.38645642073598 | 3.20132594093141 |
| chr13 | 49532220 | 49532237 | - | 3.01188173062009 | 10.2329616785794 | 4.77291284147196 | 2.13421729395427 |
| chr13 | 49533653 | 49533674 | - | 3.01188173062009 | 1.13699574206438 | 1.19322821036799 | 1.06710864697714 |
| chr13 | 49533713 | 49533744 | - | 1.00396057687336 | 3.41098722619315 | 5.96614105183995 | 3.20132594093141 |
| chr13 | 49533842 | 49533901 | - | 1.00396057687336 | 7.95897019445067 | 4.77291284147196 | 5.33554323488568 |
| chr13 | 49533961 | 49533979 | - | 1.00396057687336 | 1.13699574206438 | 4.77291284147196 | 2.13421729395427 |
| chr13 | 49534084 | 49534101 | - | 2.00792115374673 | 2.27399148412876 | 1.19322821036799 | 1.06710864697714 |
| chr13 | 49660757 | 49660828 | - | 27.1069355755808 | 40.9318467143178 | 8.35259747257593 | 24.5434988804741 |
| chr13 | 49660972 | 49661038 | - | 4.01584230749346 | 2.27399148412876 | 3.57968463110397 | 7.46976052883995 |
| chr13 | 49661089 | 49661111 | - | 2.00792115374673 | 1.13699574206438 | 1.19322821036799 | 4.26843458790854 |
| chr13 | 49701127 | 49701157 | - | 17.0673298068472 | 10.2329616785794 | 27.4442488384638 | 14.9395210576799 |
| chr13 | 49701277 | 49701285 | - | 5.01980288436682 | 6.82197445238629 | 31.0239334695677 | 3.20132594093141 |
| chr13 | 49701370 | 49701398 | - | 105.415860571703 | 77.315710460378 | 64.4343233598714 | 51.2212150549025 |
| chr13 | 49701433 | 49701444 | - | 20.0792115374673 | 10.2329616785794 | 1.19322821036799 | 6.40265188186281 |
| chr13 | 49914716 | 49914723 | - | 1.00396057687336 | 1.13699574206438 | 2.38645642073598 | 1.06710864697714 |
| chr13 | 50014063 | 50014069 | + | 3.01188173062009 | 5.68497871032191 | 3.57968463110397 | 2.13421729395427 |
| chr13 | 50015045 | 50015050 | + | 4.01584230749346 | 3.41098722619315 | 2.38645642073598 | 1.06710864697714 |
| chr13 | 50015529 | 50015530 | + | 1.00396057687336 | 2.27399148412876 | 1.19322821036799 | 1.06710864697714 |
| chr13 | 50082159 | 50082311 | + | 713.815970156962 | 668.553496333857 | 929.524775876664 | 1116.19564473808 |
| chr13 | 50082425 | 50082441 | + | 1.00396057687336 | 1.13699574206438 | 3.57968463110397 | 6.40265188186281 |
| chr13 | 50082518 | 50082523 | + | 8.03168461498692 | 6.82197445238629 | 8.35259747257593 | 5.33554323488568 |
| chr13 | 50104775 | 50104785 | + | 3.01188173062009 | 1.13699574206438 | 2.38645642073598 | 2.13421729395427 |
| chr13 | 50104886 | 50104942 | + | 4.01584230749346 | 3.41098722619315 | 11.9322821036799 | 12.8053037637256 |
| chr13 | 50104962 | 50105081 | + | 39.1544624980612 | 51.1648083928972 | 101.424397881279 | 73.6304966414224 |
| chr13 | 50105084 | 50105117 | + | 6.02376346124019 | 10.2329616785794 | 25.0577924177278 | 5.33554323488568 |
| chr13 | 50105181 | 50105205 | + | 229.906972104 | 259.235029190679 | 558.430802452219 | 437.514545260626 |
| chr13 | 50956393 | 50956481 | + | 58.2297134586551 | 101.19262104373 | 46.5359002043516 | 50.1541064079254 |
| chr13 | 51362904 | 51362921 | - | 1.00396057687336 | 5.68497871032191 | 2.38645642073598 | 1.06710864697714 |
| chr13 | 51447238 | 51447241 | - | 2.00792115374673 | 3.41098722619315 | 1.19322821036799 | 1.06710864697714 |
| chr13 | 51450530 | 51450548 | - | 6.02376346124019 | 5.68497871032191 | 1.19322821036799 | 1.06710864697714 |
| chr13 | 51453303 | 51453390 | + | 22.087132691214 | 31.8358807778027 | 38.1833027317757 | 58.6909755837425 |
| chr13 | 51760243 | 51760277 | + | 12.0475269224804 | 29.5618892936739 | 8.35259747257593 | 8.53686917581709 |
| chr13 | 51760302 | 51760308 | + | 1.00396057687336 | 4.54798296825753 | 5.96614105183995 | 1.06710864697714 |
| chr13 | 51760360 | 51760365 | + | 3.01188173062009 | 4.54798296825753 | 3.57968463110397 | 1.06710864697714 |
| chr13 | 51760933 | 51760941 | + | 4.01584230749346 | 4.54798296825753 | 1.19322821036799 | 1.06710864697714 |
| chr13 | 51766393 | 51766415 | + | 7.02772403811355 | 6.82197445238629 | 3.57968463110397 | 1.06710864697714 |
| chr13 | 51766508 | 51766534 | + | 8.03168461498692 | 12.5069531627082 | 8.35259747257593 | 3.20132594093141 |
| chr13 | 51766562 | 51766617 | + | 4.01584230749346 | 13.6439489047726 | 4.77291284147196 | 1.06710864697714 |
| chr13 | 51803812 | 51803854 | + | 19.0752509605939 | 19.3289276150945 | 3.57968463110397 | 17.0737383516342 |
| chr13 | 51804217 | 51804235 | + | 2.00792115374673 | 1.13699574206438 | 7.15936926220794 | 9.60397782279422 |
| chr13 | 51932763 | 51932770 | - | 2.00792115374673 | 1.13699574206438 | 2.38645642073598 | 3.20132594093141 |
| chr13 | 52032668 | 52032686 | + | 5.01980288436682 | 6.82197445238629 | 3.57968463110397 | 8.53686917581709 |
| chr13 | 52032749 | 52032774 | + | 12.0475269224804 | 14.780944646837 | 7.15936926220794 | 2.13421729395427 |
| chr13 | 52032893 | 52032933 | + | 7.02772403811355 | 13.6439489047726 | 7.15936926220794 | 6.40265188186281 |
| chr13 | 52032976 | 52033028 | + | 7.02772403811355 | 9.09596593651506 | 14.3187385244159 | 12.8053037637256 |
| chr13 | 52033110 | 52033141 | + | 3.01188173062009 | 5.68497871032191 | 14.3187385244159 | 14.9395210576799 |
| chr13 | 52033180 | 52033225 | + | 9.03564519186028 | 9.09596593651506 | 19.0916513658878 | 12.8053037637256 |
| chr13 | 52132741 | 52132774 | - | 1.00396057687336 | 3.41098722619315 | 1.19322821036799 | 4.26843458790854 |
| chr13 | 52412745 | 52412747 | - | 1.00396057687336 | 2.27399148412876 | 1.19322821036799 | 2.13421729395427 |
| chr13 | 52412799 | 52412824 | - | 2.00792115374673 | 6.82197445238629 | 7.15936926220794 | 11.7381951167485 |
| chr13 | 52415418 | 52415420 | - | 1.00396057687336 | 2.27399148412876 | 1.19322821036799 | 2.13421729395427 |
| chr13 | 52415779 | 52415781 | - | 1.00396057687336 | 4.54798296825753 | 2.38645642073598 | 1.06710864697714 |
| chr13 | 52475928 | 52475930 | + | 2.00792115374673 | 2.27399148412876 | 1.19322821036799 | 4.26843458790854 |
| chr13 | 52476291 | 52476310 | + | 5.01980288436682 | 2.27399148412876 | 9.54582568294392 | 6.40265188186281 |
| chr13 | 52643672 | 52643678 | + | 3.01188173062009 | 5.68497871032191 | 1.19322821036799 | 4.26843458790854 |
| chr13 | 52688109 | 52688113 | + | 1.00396057687336 | 1.13699574206438 | 5.96614105183995 | 7.46976052883995 |
| chr13 | 52688508 | 52688530 | + | 3.01188173062009 | 3.41098722619315 | 4.77291284147196 | 3.20132594093141 |
| chr13 | 52688560 | 52688588 | + | 8.03168461498692 | 3.41098722619315 | 14.3187385244159 | 8.53686917581709 |
| chr13 | 52697894 | 52697902 | + | 5.01980288436682 | 7.95897019445067 | 2.38645642073598 | 4.26843458790854 |
| chr13 | 59666413 | 59666435 | - | 1.00396057687336 | 2.27399148412876 | 2.38645642073598 | 5.33554323488568 |
| chr13 | 60573657 | 60573659 | + | 5.01980288436682 | 5.68497871032191 | 1.19322821036799 | 1.06710864697714 |
| chr13 | 70139338 | 70139339 | + | 11.043566345607 | 7.95897019445067 | 17.8984231555198 | 10.6710864697714 |
| chr13 | 72708512 | 72708546 | - | 12.0475269224804 | 18.1919318730301 | 10.7390538933119 | 16.006629704657 |
| chr13 | 72709562 | 72709573 | - | 2.00792115374673 | 2.27399148412876 | 2.38645642073598 | 1.06710864697714 |
| chr13 | 72709703 | 72709709 | - | 2.00792115374673 | 1.13699574206438 | 2.38645642073598 | 4.26843458790854 |
| chr13 | 72709984 | 72709987 | - | 9.03564519186028 | 12.5069531627082 | 2.38645642073598 | 3.20132594093141 |
| chr13 | 72710075 | 72710101 | - | 3.01188173062009 | 2.27399148412876 | 17.8984231555198 | 5.33554323488568 |
| chr13 | 72710128 | 72710183 | - | 11.043566345607 | 20.4659233571589 | 53.6952694665595 | 36.2816939972226 |
| chr13 | 72710221 | 72710316 | - | 30.1188173062009 | 59.1237785873479 | 116.936364616063 | 57.6238669367653 |
| chr13 | 72755401 | 72755404 | + | 1.00396057687336 | 1.13699574206438 | 2.38645642073598 | 1.06710864697714 |
| chr13 | 72758984 | 72758986 | - | 6.02376346124019 | 19.3289276150945 | 5.96614105183995 | 8.53686917581709 |
| chr13 | 72759009 | 72759013 | - | 4.01584230749346 | 4.54798296825753 | 3.57968463110397 | 2.13421729395427 |
| chr13 | 72759162 | 72759184 | - | 2.00792115374673 | 2.27399148412876 | 2.38645642073598 | 1.06710864697714 |
| chr13 | 72759233 | 72759246 | - | 5.01980288436682 | 4.54798296825753 | 1.19322821036799 | 2.13421729395427 |
| chr13 | 72759348 | 72759355 | - | 4.01584230749346 | 6.82197445238629 | 2.38645642073598 | 1.06710864697714 |
| chr13 | 75526484 | 75526544 | - | 10.0396057687336 | 22.7399148412876 | 3.57968463110397 | 5.33554323488568 |
| chr13 | 75526571 | 75526596 | - | 17.0673298068472 | 14.780944646837 | 5.96614105183995 | 6.40265188186281 |
| chr13 | 75549738 | 75549827 | + | 228.903011527127 | 219.440178218426 | 306.659650064573 | 329.736571915935 |
| chr13 | 75605760 | 75605826 | + | 11.043566345607 | 12.5069531627082 | 9.54582568294392 | 8.53686917581709 |
| chr13 | 77001949 | 77001951 | + | 1.00396057687336 | 2.27399148412876 | 1.19322821036799 | 2.13421729395427 |
| chr13 | 77002261 | 77002263 | + | 1.00396057687336 | 1.13699574206438 | 1.19322821036799 | 1.06710864697714 |
| chr13 | 77045287 | 77045297 | - | 3.01188173062009 | 2.27399148412876 | 1.19322821036799 | 1.06710864697714 |
| chr13 | 77045318 | 77045325 | - | 8.03168461498692 | 11.3699574206438 | 1.19322821036799 | 2.13421729395427 |
| chr13 | 78614485 | 78614506 | - | 17.0673298068472 | 34.1098722619315 | 9.54582568294392 | 12.8053037637256 |
| chr13 | 79319868 | 79319870 | - | 2.00792115374673 | 1.13699574206438 | 1.19322821036799 | 1.06710864697714 |
| chr13 | 79319983 | 79320053 | - | 23.0910932680874 | 27.2878978095452 | 22.6713359969918 | 12.8053037637256 |
| chr13 | 79320073 | 79320115 | - | 14.0554480762271 | 5.68497871032191 | 4.77291284147196 | 3.20132594093141 |
| chr13 | 79320481 | 79320482 | - | 1.00396057687336 | 3.41098722619315 | 3.57968463110397 | 2.13421729395427 |
| chr13 | 79406284 | 79406336 | + | 64.2534769198953 | 120.521548658824 | 97.8447132501751 | 109.912190638645 |
| chr13 | 79552632 | 79552659 | + | 6.02376346124019 | 1.13699574206438 | 1.19322821036799 | 1.06710864697714 |
| chr13 | 79553359 | 79553368 | + | 4.01584230749346 | 2.27399148412876 | 4.77291284147196 | 3.20132594093141 |
| chr13 | 79553404 | 79553408 | + | 6.02376346124019 | 3.41098722619315 | 5.96614105183995 | 4.26843458790854 |
| chr13 | 79553702 | 79553749 | + | 34.1346596136944 | 32.9728765198671 | 15.5119667347839 | 28.8119334683827 |
| chr13 | 79553960 | 79553981 | + | 3.01188173062009 | 9.09596593651506 | 4.77291284147196 | 3.20132594093141 |
| chr13 | 79555489 | 79555515 | + | 29.1148567293276 | 47.753821166704 | 57.2749540976635 | 42.6843458790854 |
| chr13 | 79555553 | 79555568 | + | 14.0554480762271 | 15.9179403889013 | 19.0916513658878 | 10.6710864697714 |
| chr13 | 79555983 | 79555984 | + | 1.00396057687336 | 2.27399148412876 | 2.38645642073598 | 1.06710864697714 |
| chr13 | 91350961 | 91350983 | + | 1.00396057687336 | 1.13699574206438 | 17.8984231555198 | 18.1408469986113 |
| chr13 | 91351381 | 91351406 | + | 32.1267384599477 | 27.2878978095452 | 320.978388588989 | 149.395210576799 |
| chr13 | 91351536 | 91351556 | + | 4.01584230749346 | 2.27399148412876 | 4.77291284147196 | 11.7381951167485 |
| chr13 | 94574460 | 94574486 | - | 2.00792115374673 | 3.41098722619315 | 1.19322821036799 | 6.40265188186281 |
| chr13 | 95792143 | 95792159 | + | 4.01584230749346 | 6.82197445238629 | 1.19322821036799 | 1.06710864697714 |
| chr13 | 95792344 | 95792448 | + | 10.0396057687336 | 11.3699574206438 | 17.8984231555198 | 6.40265188186281 |
| chr13 | 95794724 | 95794808 | + | 5.01980288436682 | 3.41098722619315 | 2.38645642073598 | 1.06710864697714 |
| chr13 | 95794926 | 95794929 | + | 1.00396057687336 | 5.68497871032191 | 3.57968463110397 | 1.06710864697714 |
| chr13 | 95801764 | 95801799 | - | 17.0673298068472 | 19.3289276150945 | 2.38645642073598 | 11.7381951167485 |
| chr13 | 97392282 | 97392312 | + | 4.01584230749346 | 3.41098722619315 | 1.19322821036799 | 2.13421729395427 |
| chr13 | 97393009 | 97393028 | + | 6.02376346124019 | 14.780944646837 | 5.96614105183995 | 5.33554323488568 |
| chr13 | 97393076 | 97393203 | + | 14.0554480762271 | 15.9179403889013 | 15.5119667347839 | 8.53686917581709 |
| chr13 | 97393226 | 97393251 | + | 6.02376346124019 | 4.54798296825753 | 4.77291284147196 | 3.20132594093141 |
| chr13 | 97393331 | 97393383 | + | 7.02772403811355 | 27.2878978095452 | 17.8984231555198 | 12.8053037637256 |
| chr13 | 97394026 | 97394032 | + | 4.01584230749346 | 7.95897019445067 | 2.38645642073598 | 1.06710864697714 |
| chr13 | 97467486 | 97467503 | + | 2.00792115374673 | 1.13699574206438 | 1.19322821036799 | 3.20132594093141 |
| chr13 | 98021833 | 98021883 | + | 3.01188173062009 | 3.41098722619315 | 2.38645642073598 | 3.20132594093141 |
| chr13 | 98021889 | 98021928 | + | 2.00792115374673 | 2.27399148412876 | 4.77291284147196 | 2.13421729395427 |
| chr13 | 98021970 | 98021997 | + | 1.00396057687336 | 2.27399148412876 | 1.19322821036799 | 1.06710864697714 |
| chr13 | 98022016 | 98022039 | + | 3.01188173062009 | 1.13699574206438 | 4.77291284147196 | 6.40265188186281 |
| chr13 | 98022086 | 98022100 | + | 19.0752509605939 | 22.7399148412876 | 11.9322821036799 | 34.1474767032683 |
| chr13 | 98022509 | 98022516 | + | 5.01980288436682 | 1.13699574206438 | 1.19322821036799 | 3.20132594093141 |
| chr13 | 98452196 | 98452246 | - | 1.00396057687336 | 5.68497871032191 | 3.57968463110397 | 1.06710864697714 |
| chr13 | 98452267 | 98452287 | - | 1.00396057687336 | 3.41098722619315 | 5.96614105183995 | 3.20132594093141 |
| chr13 | 98452351 | 98452784 | - | 311.227778830743 | 484.360186119427 | 260.123749860222 | 185.676904574022 |
| chr13 | 98452954 | 98452960 | - | 7.02772403811355 | 7.95897019445067 | 2.38645642073598 | 3.20132594093141 |
| chr13 | 98453153 | 98453161 | - | 1.00396057687336 | 5.68497871032191 | 2.38645642073598 | 3.20132594093141 |
| chr13 | 98793585 | 98793586 | - | 6.02376346124019 | 2.27399148412876 | 3.57968463110397 | 1.06710864697714 |
| chr13 | 98793638 | 98793700 | - | 27.1069355755808 | 22.7399148412876 | 46.5359002043516 | 41.6172372321083 |
| chr13 | 98793766 | 98793772 | - | 6.02376346124019 | 5.68497871032191 | 8.35259747257593 | 6.40265188186281 |
| chr13 | 98793865 | 98793868 | - | 4.01584230749346 | 5.68497871032191 | 1.19322821036799 | 4.26843458790854 |
| chr13 | 98794228 | 98794274 | - | 8.03168461498692 | 11.3699574206438 | 8.35259747257593 | 23.476390233497 |
| chr13 | 99385711 | 99385719 | + | 2.00792115374673 | 3.41098722619315 | 2.38645642073598 | 2.13421729395427 |
| chr13 | 99385826 | 99385832 | + | 1.00396057687336 | 2.27399148412876 | 4.77291284147196 | 4.26843458790854 |
| chr13 | 99385901 | 99385927 | + | 1.00396057687336 | 1.13699574206438 | 2.38645642073598 | 5.33554323488568 |
| chr13 | 99386230 | 99386312 | + | 32.1267384599477 | 37.5208594881246 | 20.2848795762558 | 48.0198891139711 |
| chr13 | 99386333 | 99386399 | + | 22.087132691214 | 19.3289276150945 | 5.96614105183995 | 11.7381951167485 |
| chr13 | 99498735 | 99498914 | + | 2.00792115374673 | 3.41098722619315 | 73.9801490428153 | 33.0803680562912 |
| chr13 | 99562698 | 99562786 | + | 15.0594086531005 | 17.0549361309657 | 4.77291284147196 | 11.7381951167485 |
| chr13 | 99562979 | 99563014 | + | 4.01584230749346 | 2.27399148412876 | 1.19322821036799 | 2.13421729395427 |
| chr13 | 99563059 | 99563108 | + | 4.01584230749346 | 4.54798296825753 | 3.57968463110397 | 3.20132594093141 |
| chr13 | 99563133 | 99563168 | + | 8.03168461498692 | 9.09596593651506 | 2.38645642073598 | 14.9395210576799 |
| chr13 | 99563198 | 99563368 | + | 61.2415951892752 | 95.5076423334081 | 60.8546387287675 | 42.6843458790854 |
| chr13 | 99577276 | 99577277 | + | 6.02376346124019 | 17.0549361309657 | 16.7051949451519 | 14.9395210576799 |
| chr13 | 99965108 | 99965119 | - | 21.0831721143407 | 20.4659233571589 | 9.54582568294392 | 5.33554323488568 |
| chr13 | 99965227 | 99965243 | - | 1.00396057687336 | 1.13699574206438 | 2.38645642073598 | 3.20132594093141 |
| chr13 | 100530096 | 100530168 | + | 87.3445701879827 | 80.7266976865711 | 82.3327465153913 | 66.1607361125824 |
| chr13 | 100530251 | 100530354 | + | 18.0712903837206 | 25.0139063254164 | 42.9562155732476 | 35.2145853502455 |
| chr13 | 100532013 | 100532019 | - | 2.00792115374673 | 2.27399148412876 | 2.38645642073598 | 3.20132594093141 |
| chr13 | 100604080 | 100604121 | - | 12.0475269224804 | 17.0549361309657 | 27.4442488384638 | 11.7381951167485 |
| chr13 | 100604148 | 100604223 | - | 19.0752509605939 | 19.3289276150945 | 71.5936926220794 | 60.8251928776967 |
| chr13 | 100604252 | 100604302 | - | 1.00396057687336 | 4.54798296825753 | 5.96614105183995 | 17.0737383516342 |
| chr13 | 100604387 | 100604393 | - | 1.00396057687336 | 3.41098722619315 | 1.19322821036799 | 3.20132594093141 |
| chr13 | 102394693 | 102394694 | + | 1.00396057687336 | 4.54798296825753 | 2.38645642073598 | 1.06710864697714 |
| chr13 | 102678543 | 102678551 | + | 3.01188173062009 | 5.68497871032191 | 2.38645642073598 | 3.20132594093141 |
| chr13 | 102766181 | 102766219 | - | 30.1188173062009 | 105.740604011988 | 53.6952694665595 | 56.5567582897882 |
| chr13 | 102766488 | 102766494 | - | 10.0396057687336 | 7.95897019445067 | 5.96614105183995 | 5.33554323488568 |
| chr13 | 102766517 | 102766519 | - | 2.00792115374673 | 4.54798296825753 | 2.38645642073598 | 3.20132594093141 |
| chr13 | 102840012 | 102840035 | + | 4.01584230749346 | 1.13699574206438 | 5.96614105183995 | 2.13421729395427 |
| chr13 | 102840988 | 102841035 | + | 6.02376346124019 | 13.6439489047726 | 11.9322821036799 | 14.9395210576799 |
| chr13 | 102841055 | 102841100 | + | 6.02376346124019 | 14.780944646837 | 8.35259747257593 | 12.8053037637256 |
| chr13 | 102841286 | 102841292 | + | 2.00792115374673 | 1.13699574206438 | 2.38645642073598 | 3.20132594093141 |
| chr13 | 102841379 | 102841401 | + | 8.03168461498692 | 17.0549361309657 | 3.57968463110397 | 1.06710864697714 |
| chr13 | 106543426 | 106543583 | - | 15.0594086531005 | 22.7399148412876 | 20.2848795762558 | 8.53686917581709 |
| chr13 | 106543690 | 106543706 | - | 5.01980288436682 | 4.54798296825753 | 1.19322821036799 | 1.06710864697714 |
| chr13 | 106543729 | 106543743 | - | 2.00792115374673 | 2.27399148412876 | 1.19322821036799 | 3.20132594093141 |
| chr13 | 106543777 | 106543794 | - | 9.03564519186028 | 11.3699574206438 | 3.57968463110397 | 3.20132594093141 |
| chr13 | 106543852 | 106543883 | - | 8.03168461498692 | 2.27399148412876 | 2.38645642073598 | 8.53686917581709 |
| chr13 | 106543936 | 106543950 | - | 3.01188173062009 | 1.13699574206438 | 3.57968463110397 | 9.60397782279422 |
| chr13 | 109754152 | 109754181 | - | 36.1425807674411 | 52.3018041349616 | 1.19322821036799 | 2.13421729395427 |
| chr13 | 109754681 | 109754717 | - | 7.02772403811355 | 13.6439489047726 | 1.19322821036799 | 2.13421729395427 |
| chr13 | 109754882 | 109754910 | - | 14.0554480762271 | 26.1509020674808 | 4.77291284147196 | 3.20132594093141 |
| chr13 | 109755250 | 109755260 | - | 18.0712903837206 | 19.3289276150945 | 1.19322821036799 | 1.06710864697714 |
| chr13 | 109755293 | 109755349 | - | 113.44754518669 | 164.864382599335 | 14.3187385244159 | 1.06710864697714 |
| chr13 | 109755510 | 109755551 | - | 790.116973999338 | 1106.29685702864 | 44.1494437836156 | 62.959410171651 |
| chr13 | 109755615 | 109755718 | - | 86.3406096111093 | 108.014595496116 | 1.19322821036799 | 1.06710864697714 |
| chr13 | 110512896 | 110512898 | + | 7.02772403811355 | 6.82197445238629 | 5.96614105183995 | 5.33554323488568 |
| chr13 | 110512951 | 110512955 | + | 10.0396057687336 | 11.3699574206438 | 4.77291284147196 | 5.33554323488568 |
| chr13 | 110523208 | 110523243 | - | 2.00792115374673 | 1.13699574206438 | 1.19322821036799 | 1.06710864697714 |
| chr13 | 110638551 | 110638583 | + | 6.02376346124019 | 3.41098722619315 | 4.77291284147196 | 3.20132594093141 |
| chr13 | 110641441 | 110641453 | - | 10.0396057687336 | 2.27399148412876 | 2.38645642073598 | 2.13421729395427 |
| chr13 | 110719954 | 110719960 | + | 1.00396057687336 | 3.41098722619315 | 2.38645642073598 | 1.06710864697714 |
| chr13 | 110720004 | 110720012 | + | 4.01584230749346 | 2.27399148412876 | 1.19322821036799 | 2.13421729395427 |
| chr13 | 110720110 | 110720149 | + | 9.03564519186028 | 6.82197445238629 | 3.57968463110397 | 3.20132594093141 |
| chr13 | 110720241 | 110720246 | + | 3.01188173062009 | 5.68497871032191 | 3.57968463110397 | 1.06710864697714 |
| chr13 | 110720523 | 110720552 | + | 8.03168461498692 | 25.0139063254164 | 23.8645642073598 | 8.53686917581709 |
| chr13 | 110720594 | 110720621 | + | 9.03564519186028 | 15.9179403889013 | 8.35259747257593 | 4.26843458790854 |
| chr13 | 110720644 | 110720686 | + | 10.0396057687336 | 14.780944646837 | 3.57968463110397 | 4.26843458790854 |
| chr13 | 110720969 | 110720971 | + | 1.00396057687336 | 2.27399148412876 | 1.19322821036799 | 2.13421729395427 |
| chr13 | 110879001 | 110879011 | - | 13.0514874993537 | 5.68497871032191 | 2.38645642073598 | 3.20132594093141 |
| chr13 | 110892714 | 110892742 | - | 3.01188173062009 | 2.27399148412876 | 5.96614105183995 | 1.06710864697714 |
| chr13 | 110892762 | 110892953 | - | 24.0950538449607 | 53.438799877026 | 51.3088130458235 | 78.966039876308 |
| chr13 | 110892992 | 110893099 | - | 18.0712903837206 | 12.5069531627082 | 50.1155848354556 | 68.2949534065367 |
| chr13 | 110893123 | 110893139 | - | 1.00396057687336 | 3.41098722619315 | 1.19322821036799 | 1.06710864697714 |
| chr13 | 111305598 | 111305630 | + | 3.01188173062009 | 1.13699574206438 | 3.57968463110397 | 3.20132594093141 |
| chr13 | 112546690 | 112546702 | - | 29.1148567293276 | 28.4248935516096 | 5.96614105183995 | 12.8053037637256 |
| chr13 | 112883877 | 112883963 | + | 20.0792115374673 | 19.3289276150945 | 8.35259747257593 | 5.33554323488568 |
| chr13 | 112886552 | 112886639 | + | 18.0712903837206 | 28.4248935516096 | 2.38645642073598 | 6.40265188186281 |
| chr13 | 112910015 | 112910017 | + | 4.01584230749346 | 1.13699574206438 | 2.38645642073598 | 1.06710864697714 |
| chr13 | 113177637 | 113177649 | - | 112.443584609817 | 173.96034853585 | 15.5119667347839 | 21.3421729395427 |
| chr13 | 113177878 | 113177937 | - | 37.1465413443145 | 50.0278126508328 | 4.77291284147196 | 12.8053037637256 |
| chr13 | 113229965 | 113229993 | + | 2.00792115374673 | 2.27399148412876 | 1.19322821036799 | 8.53686917581709 |
| chr13 | 113263575 | 113263596 | + | 6.02376346124019 | 4.54798296825753 | 2.38645642073598 | 1.06710864697714 |
| chr13 | 113263647 | 113263755 | + | 80.3168461498692 | 54.5757956190903 | 34.6036181006717 | 28.8119334683827 |
| chr13 | 113263779 | 113263794 | + | 2.00792115374673 | 1.13699574206438 | 4.77291284147196 | 2.13421729395427 |
| chr13 | 113263820 | 113263865 | + | 15.0594086531005 | 22.7399148412876 | 8.35259747257593 | 7.46976052883995 |
| chr13 | 113263891 | 113263993 | + | 28.1108961524542 | 15.9179403889013 | 9.54582568294392 | 4.26843458790854 |
| chr13 | 113264007 | 113264077 | + | 15.0594086531005 | 17.0549361309657 | 14.3187385244159 | 27.7448248214055 |
| chr13 | 113264090 | 113264208 | + | 36.1425807674411 | 22.7399148412876 | 9.54582568294392 | 14.9395210576799 |
| chr13 | 113264230 | 113264434 | + | 994.924931681504 | 923.240542556278 | 313.819019326781 | 322.266811387095 |
| chr13 | 113264467 | 113264531 | + | 14.0554480762271 | 7.95897019445067 | 8.35259747257593 | 13.8724124107028 |
| chr13 | 113264816 | 113264842 | + | 36.1425807674411 | 51.1648083928972 | 5.96614105183995 | 6.40265188186281 |
| chr13 | 113322821 | 113322850 | + | 6.02376346124019 | 2.27399148412876 | 1.19322821036799 | 1.06710864697714 |
| chr13 | 113323131 | 113323230 | + | 13.0514874993537 | 12.5069531627082 | 7.15936926220794 | 9.60397782279422 |
| chr13 | 113323276 | 113323404 | + | 59.2336740355285 | 73.9047232341848 | 4.77291284147196 | 17.0737383516342 |
| chr13 | 113323541 | 113323615 | + | 20.0792115374673 | 20.4659233571589 | 2.38645642073598 | 8.53686917581709 |
| chr13 | 113530203 | 113530224 | + | 4.01584230749346 | 2.27399148412876 | 1.19322821036799 | 3.20132594093141 |
| chr13 | 113549764 | 113549809 | + | 5.01980288436682 | 11.3699574206438 | 5.96614105183995 | 8.53686917581709 |
| chr13 | 113549991 | 113550009 | + | 1.00396057687336 | 1.13699574206438 | 2.38645642073598 | 4.26843458790854 |
| chr13 | 113550054 | 113550058 | + | 1.00396057687336 | 4.54798296825753 | 1.19322821036799 | 1.06710864697714 |
| chr13 | 113640214 | 113640310 | + | 51.2019894205416 | 40.9318467143178 | 27.4442488384638 | 48.0198891139711 |
| chr13 | 113640352 | 113640369 | + | 7.02772403811355 | 5.68497871032191 | 4.77291284147196 | 22.4092815865198 |
| chr13 | 113640413 | 113640416 | + | 8.03168461498692 | 5.68497871032191 | 3.57968463110397 | 1.06710864697714 |
| chr13 | 113640470 | 113640551 | + | 21.0831721143407 | 21.6029190992233 | 25.0577924177278 | 21.3421729395427 |
| chr13 | 113640576 | 113640583 | + | 3.01188173062009 | 3.41098722619315 | 2.38645642073598 | 4.26843458790854 |
| chr13 | 113640628 | 113640649 | + | 27.1069355755808 | 21.6029190992233 | 14.3187385244159 | 12.8053037637256 |
| chr13 | 113640689 | 113640691 | + | 26.1029749987075 | 26.1509020674808 | 13.1255103140479 | 21.3421729395427 |
| chr13 | 113640720 | 113640726 | + | 4.01584230749346 | 3.41098722619315 | 1.19322821036799 | 8.53686917581709 |
| chr13 | 113640966 | 113640976 | + | 9.03564519186028 | 5.68497871032191 | 3.57968463110397 | 7.46976052883995 |
| chr13 | 114272563 | 114272594 | + | 2.00792115374673 | 4.54798296825753 | 7.15936926220794 | 10.6710864697714 |
| chr14 | 20289165 | 20289171 | - | 12.0475269224804 | 23.876910583352 | 16.7051949451519 | 17.0737383516342 |
| chr14 | 20289246 | 20289250 | - | 7.02772403811355 | 23.876910583352 | 4.77291284147196 | 3.20132594093141 |
| chr14 | 20311409 | 20311414 | - | 3.01188173062009 | 4.54798296825753 | 7.15936926220794 | 4.26843458790854 |
| chr14 | 20357793 | 20357821 | + | 4.01584230749346 | 1.13699574206438 | 1.19322821036799 | 1.06710864697714 |
| chr14 | 20447088 | 20447128 | - | 9.03564519186028 | 10.2329616785794 | 7.15936926220794 | 6.40265188186281 |
| chr14 | 20457077 | 20457149 | + | 3.01188173062009 | 3.41098722619315 | 2.38645642073598 | 5.33554323488568 |
| chr14 | 20457504 | 20457548 | + | 18.0712903837206 | 13.6439489047726 | 3.57968463110397 | 6.40265188186281 |
| chr14 | 20457581 | 20457584 | + | 1.00396057687336 | 1.13699574206438 | 4.77291284147196 | 2.13421729395427 |
| chr14 | 20457617 | 20457661 | + | 28.1108961524542 | 17.0549361309657 | 7.15936926220794 | 16.006629704657 |
| chr14 | 20457704 | 20457740 | + | 5.01980288436682 | 14.780944646837 | 8.35259747257593 | 2.13421729395427 |
| chr14 | 20457897 | 20457937 | - | 28.1108961524542 | 29.5618892936739 | 2.38645642073598 | 6.40265188186281 |
| chr14 | 20458195 | 20458218 | - | 12.0475269224804 | 11.3699574206438 | 1.19322821036799 | 2.13421729395427 |
| chr14 | 20476695 | 20476710 | + | 6.02376346124019 | 1.13699574206438 | 1.19322821036799 | 4.26843458790854 |
| chr14 | 20476781 | 20476833 | + | 23.0910932680874 | 11.3699574206438 | 4.77291284147196 | 10.6710864697714 |
| chr14 | 20476868 | 20476897 | + | 14.0554480762271 | 7.95897019445067 | 4.77291284147196 | 11.7381951167485 |
| chr14 | 20476972 | 20476980 | + | 147.582204800385 | 133.028501821533 | 79.9462900946553 | 98.1739955218965 |
| chr14 | 20477007 | 20477067 | + | 37.1465413443145 | 52.3018041349616 | 10.7390538933119 | 5.33554323488568 |
| chr14 | 20996933 | 20997010 | + | 78.3089249961224 | 90.9596593651506 | 48.9223566250876 | 48.0198891139711 |
| chr14 | 21089459 | 21089480 | + | 4.01584230749346 | 11.3699574206438 | 1.19322821036799 | 4.26843458790854 |
| chr14 | 21209287 | 21209308 | - | 4.01584230749346 | 4.54798296825753 | 2.38645642073598 | 5.33554323488568 |
| chr14 | 21209423 | 21209438 | - | 2.00792115374673 | 9.09596593651506 | 7.15936926220794 | 1.06710864697714 |
| chr14 | 21209492 | 21209495 | - | 2.00792115374673 | 5.68497871032191 | 2.38645642073598 | 2.13421729395427 |
| chr14 | 21209580 | 21209603 | - | 9.03564519186028 | 18.1919318730301 | 8.35259747257593 | 32.0132594093141 |
| chr14 | 21210203 | 21210204 | - | 5.01980288436682 | 2.27399148412876 | 1.19322821036799 | 1.06710864697714 |
| chr14 | 21210794 | 21210800 | - | 1.00396057687336 | 1.13699574206438 | 2.38645642073598 | 2.13421729395427 |
| chr14 | 21351548 | 21351558 | - | 5.01980288436682 | 9.09596593651506 | 3.57968463110397 | 2.13421729395427 |
| chr14 | 21351589 | 21351611 | - | 111.439624032943 | 79.5897019445067 | 40.5697591525116 | 76.8318225823538 |
| chr14 | 21351842 | 21351846 | - | 7.02772403811355 | 5.68497871032191 | 1.19322821036799 | 5.33554323488568 |
| chr14 | 21351872 | 21351946 | - | 14.0554480762271 | 14.780944646837 | 13.1255103140479 | 12.8053037637256 |
| chr14 | 21352026 | 21352055 | - | 11.043566345607 | 6.82197445238629 | 7.15936926220794 | 5.33554323488568 |
| chr14 | 21352131 | 21352160 | - | 18.0712903837206 | 3.41098722619315 | 3.57968463110397 | 10.6710864697714 |
| chr14 | 21352181 | 21352206 | - | 8.03168461498692 | 9.09596593651506 | 10.7390538933119 | 3.20132594093141 |
| chr14 | 21352310 | 21352322 | - | 2.00792115374673 | 4.54798296825753 | 1.19322821036799 | 5.33554323488568 |
| chr14 | 21352415 | 21352466 | - | 21.0831721143407 | 27.2878978095452 | 14.3187385244159 | 23.476390233497 |
| chr14 | 21352515 | 21352562 | - | 9.03564519186028 | 5.68497871032191 | 2.38645642073598 | 2.13421729395427 |
| chr14 | 21352606 | 21352610 | - | 2.00792115374673 | 2.27399148412876 | 3.57968463110397 | 5.33554323488568 |
| chr14 | 21383014 | 21383015 | - | 1.00396057687336 | 3.41098722619315 | 4.77291284147196 | 3.20132594093141 |
| chr14 | 21385223 | 21385276 | - | 20.0792115374673 | 11.3699574206438 | 7.15936926220794 | 16.006629704657 |
| chr14 | 21459144 | 21459162 | - | 4.01584230749346 | 7.95897019445067 | 2.38645642073598 | 3.20132594093141 |
| chr14 | 21498177 | 21498222 | - | 5.01980288436682 | 13.6439489047726 | 11.9322821036799 | 7.46976052883995 |
| chr14 | 22564925 | 22565140 | - | 26.1029749987075 | 27.2878978095452 | 26.2510206280958 | 26.6777161744284 |
| chr14 | 22575058 | 22575236 | - | 49.1940682667949 | 45.4798296825753 | 28.6374770488317 | 28.8119334683827 |
| chr14 | 22771513 | 22771606 | + | 5.01980288436682 | 5.68497871032191 | 4.77291284147196 | 5.33554323488568 |
| chr14 | 22771633 | 22771767 | + | 65.2574374967687 | 85.2746806548286 | 27.4442488384638 | 25.6106075274513 |
| chr14 | 22834175 | 22834328 | + | 84.3326884573626 | 63.6717615556054 | 95.4582568294392 | 117.381951167485 |
| chr14 | 22834715 | 22834716 | + | 2.00792115374673 | 1.13699574206438 | 3.57968463110397 | 4.26843458790854 |
| chr14 | 22834930 | 22834978 | + | 3.01188173062009 | 3.41098722619315 | 2.38645642073598 | 4.26843458790854 |
| chr14 | 22834999 | 22835011 | + | 10.0396057687336 | 6.82197445238629 | 4.77291284147196 | 16.006629704657 |
| chr14 | 22877931 | 22877956 | + | 2.00792115374673 | 4.54798296825753 | 1.19322821036799 | 4.26843458790854 |
| chr14 | 22877978 | 22878055 | + | 14.0554480762271 | 12.5069531627082 | 4.77291284147196 | 7.46976052883995 |
| chr14 | 22887019 | 22887114 | + | 43.1703048055547 | 25.0139063254164 | 25.0577924177278 | 51.2212150549025 |
| chr14 | 22900939 | 22900943 | - | 5.01980288436682 | 1.13699574206438 | 3.57968463110397 | 13.8724124107028 |
| chr14 | 22901710 | 22901727 | - | 1.00396057687336 | 1.13699574206438 | 3.57968463110397 | 5.33554323488568 |
| chr14 | 22920546 | 22920569 | - | 16.0633692299738 | 14.780944646837 | 16.7051949451519 | 16.006629704657 |
| chr14 | 22920605 | 22920649 | - | 37.1465413443145 | 53.438799877026 | 19.0916513658878 | 13.8724124107028 |
| chr14 | 22920675 | 22920695 | - | 8.03168461498692 | 7.95897019445067 | 11.9322821036799 | 23.476390233497 |
| chr14 | 22929605 | 22929652 | + | 29.1148567293276 | 11.3699574206438 | 7.15936926220794 | 5.33554323488568 |
| chr14 | 22946271 | 22946298 | - | 3.01188173062009 | 2.27399148412876 | 2.38645642073598 | 4.26843458790854 |
| chr14 | 22946417 | 22946462 | - | 11.043566345607 | 14.780944646837 | 8.35259747257593 | 14.9395210576799 |
| chr14 | 22946515 | 22946581 | - | 12.0475269224804 | 9.09596593651506 | 11.9322821036799 | 6.40265188186281 |
| chr14 | 22982649 | 22982792 | + | 58.2297134586551 | 102.329616785794 | 9.54582568294392 | 8.53686917581709 |
| chr14 | 22982838 | 22982852 | + | 9.03564519186028 | 14.780944646837 | 1.19322821036799 | 4.26843458790854 |
| chr14 | 22982883 | 22982988 | + | 52.205949997415 | 100.055625301666 | 38.1833027317757 | 24.5434988804741 |
| chr14 | 23025871 | 23025888 | - | 48.1901076899215 | 55.7127913611547 | 19.0916513658878 | 1.06710864697714 |
| chr14 | 23025917 | 23026377 | - | 3283.95504695278 | 2135.27800359691 | 2089.34259635435 | 2173.70031389243 |
| chr14 | 23047111 | 23047119 | - | 8.03168461498692 | 12.5069531627082 | 9.54582568294392 | 12.8053037637256 |
| chr14 | 23058595 | 23058610 | - | 7.02772403811355 | 21.6029190992233 | 7.15936926220794 | 8.53686917581709 |
| chr14 | 23058648 | 23058782 | - | 47.1861471130481 | 48.8908169087684 | 36.9900745214077 | 25.6106075274513 |
| chr14 | 23059024 | 23059103 | - | 9.03564519186028 | 6.82197445238629 | 7.15936926220794 | 5.33554323488568 |
| chr14 | 23098066 | 23098378 | + | 167.661416337852 | 135.302493305661 | 198.075882921086 | 149.395210576799 |
| chr14 | 23098410 | 23098477 | + | 1.00396057687336 | 3.41098722619315 | 2.38645642073598 | 1.06710864697714 |
| chr14 | 23098599 | 23098607 | + | 4.01584230749346 | 3.41098722619315 | 2.38645642073598 | 3.20132594093141 |
| chr14 | 23098854 | 23098856 | + | 2.00792115374673 | 7.95897019445067 | 4.77291284147196 | 7.46976052883995 |
| chr14 | 23275433 | 23275451 | - | 33.130699036821 | 11.3699574206438 | 9.54582568294392 | 10.6710864697714 |
| chr14 | 23275535 | 23275570 | - | 24.0950538449607 | 29.5618892936739 | 21.4781077866238 | 10.6710864697714 |
| chr14 | 23298602 | 23298604 | - | 1.00396057687336 | 1.13699574206438 | 1.19322821036799 | 1.06710864697714 |
| chr14 | 23308907 | 23309190 | + | 160.633692299738 | 267.19399938513 | 112.163451774591 | 59.7580842307196 |
| chr14 | 23309214 | 23309248 | + | 29.1148567293276 | 46.6168254246397 | 14.3187385244159 | 2.13421729395427 |
| chr14 | 23309393 | 23309414 | + | 4.01584230749346 | 9.09596593651506 | 3.57968463110397 | 2.13421729395427 |
| chr14 | 23309485 | 23309504 | + | 7.02772403811355 | 3.41098722619315 | 3.57968463110397 | 1.06710864697714 |
| chr14 | 23309755 | 23309813 | + | 7.02772403811355 | 9.09596593651506 | 2.38645642073598 | 2.13421729395427 |
| chr14 | 23310046 | 23310085 | + | 2.00792115374673 | 7.95897019445067 | 3.57968463110397 | 2.13421729395427 |
| chr14 | 23310181 | 23310200 | + | 8.03168461498692 | 4.54798296825753 | 4.77291284147196 | 2.13421729395427 |
| chr14 | 23310672 | 23310699 | + | 30.1188173062009 | 57.9867828452835 | 36.9900745214077 | 11.7381951167485 |
| chr14 | 23310732 | 23310810 | + | 8.03168461498692 | 6.82197445238629 | 15.5119667347839 | 11.7381951167485 |
| chr14 | 23310839 | 23310870 | + | 4.01584230749346 | 12.5069531627082 | 7.15936926220794 | 3.20132594093141 |
| chr14 | 23311412 | 23311509 | + | 12.0475269224804 | 21.6029190992233 | 10.7390538933119 | 11.7381951167485 |
| chr14 | 23311536 | 23311541 | + | 2.00792115374673 | 3.41098722619315 | 1.19322821036799 | 1.06710864697714 |
| chr14 | 23324136 | 23324302 | + | 48.1901076899215 | 37.5208594881246 | 38.1833027317757 | 62.959410171651 |
| chr14 | 23324556 | 23324623 | + | 2.00792115374673 | 4.54798296825753 | 3.57968463110397 | 2.13421729395427 |
| chr14 | 23324731 | 23324743 | + | 2.00792115374673 | 2.27399148412876 | 5.96614105183995 | 1.06710864697714 |
| chr14 | 23324897 | 23324984 | + | 22.087132691214 | 23.876910583352 | 23.8645642073598 | 14.9395210576799 |
| chr14 | 23325083 | 23325130 | + | 40.1584230749346 | 54.5757956190903 | 45.3426719939836 | 34.1474767032683 |
| chr14 | 23325166 | 23325216 | + | 205.81191825904 | 241.043097317649 | 464.165773833148 | 117.381951167485 |
| chr14 | 23325244 | 23325424 | + | 870.433820149207 | 822.047921512548 | 1689.61114588107 | 936.921392045925 |
| chr14 | 23325450 | 23325505 | + | 57.2257528817818 | 22.7399148412876 | 73.9801490428153 | 25.6106075274513 |
| chr14 | 23325559 | 23325618 | + | 4.01584230749346 | 3.41098722619315 | 7.15936926220794 | 11.7381951167485 |
| chr14 | 23325884 | 23325978 | + | 64.2534769198953 | 68.2197445238629 | 77.5598336739193 | 81.1002571702623 |
| chr14 | 23326029 | 23326077 | + | 43.1703048055547 | 62.534765813541 | 50.1155848354556 | 55.4896496428111 |
| chr14 | 23326103 | 23326113 | + | 40.1584230749346 | 69.3567402659273 | 68.0140079909754 | 86.435800405148 |
| chr14 | 23478016 | 23478037 | + | 20.0792115374673 | 26.1509020674808 | 13.1255103140479 | 18.1408469986113 |
| chr14 | 23478063 | 23478144 | + | 15.0594086531005 | 21.6029190992233 | 8.35259747257593 | 14.9395210576799 |
| chr14 | 23559575 | 23559629 | - | 5.01980288436682 | 6.82197445238629 | 1.19322821036799 | 3.20132594093141 |
| chr14 | 23645118 | 23645619 | + | 1.00396057687336 | 1.13699574206438 | 10.7390538933119 | 19.2079556455884 |
| chr14 | 24006151 | 24006158 | + | 6.02376346124019 | 4.54798296825753 | 4.77291284147196 | 3.20132594093141 |
| chr14 | 24103969 | 24104043 | + | 2.00792115374673 | 19.3289276150945 | 2.38645642073598 | 3.20132594093141 |
| chr14 | 24104068 | 24104113 | + | 4.01584230749346 | 13.6439489047726 | 2.38645642073598 | 1.06710864697714 |
| chr14 | 24123560 | 24123646 | + | 9.03564519186028 | 10.2329616785794 | 7.15936926220794 | 22.4092815865198 |
| chr14 | 24123706 | 24123779 | + | 5.01980288436682 | 5.68497871032191 | 11.9322821036799 | 22.4092815865198 |
| chr14 | 24123831 | 24124006 | + | 1.00396057687336 | 4.54798296825753 | 3.57968463110397 | 1.06710864697714 |
| chr14 | 24138735 | 24138950 | + | 201.796075951546 | 289.933914226417 | 73.9801490428153 | 73.6304966414224 |
| chr14 | 24143404 | 24143442 | - | 136.538638454778 | 228.536144154941 | 29.8307052591997 | 53.3554323488568 |
| chr14 | 24161232 | 24161319 | + | 390.540664403739 | 388.852543786019 | 93.0718004087032 | 85.3686917581709 |
| chr14 | 24166348 | 24166365 | + | 2.00792115374673 | 1.13699574206438 | 3.57968463110397 | 2.13421729395427 |
| chr14 | 24180240 | 24180322 | - | 25.0990144218341 | 20.4659233571589 | 10.7390538933119 | 36.2816939972226 |
| chr14 | 24182017 | 24182030 | - | 1.00396057687336 | 2.27399148412876 | 2.38645642073598 | 5.33554323488568 |
| chr14 | 24182307 | 24182342 | - | 13.0514874993537 | 6.82197445238629 | 7.15936926220794 | 13.8724124107028 |
| chr14 | 24182807 | 24182841 | - | 4.01584230749346 | 2.27399148412876 | 5.96614105183995 | 23.476390233497 |
| chr14 | 24183069 | 24183117 | - | 5.01980288436682 | 3.41098722619315 | 7.15936926220794 | 16.006629704657 |
| chr14 | 24183491 | 24183516 | - | 2.00792115374673 | 3.41098722619315 | 2.38645642073598 | 19.2079556455884 |
| chr14 | 24183609 | 24183613 | - | 13.0514874993537 | 10.2329616785794 | 2.38645642073598 | 7.46976052883995 |
| chr14 | 24183639 | 24183669 | - | 8.03168461498692 | 2.27399148412876 | 4.77291284147196 | 2.13421729395427 |
| chr14 | 24183802 | 24183891 | - | 39.1544624980612 | 27.2878978095452 | 32.2171616799357 | 44.8185631730397 |
| chr14 | 24184069 | 24184076 | - | 3.01188173062009 | 1.13699574206438 | 2.38645642073598 | 1.06710864697714 |
| chr14 | 24184386 | 24184415 | - | 7.02772403811355 | 3.41098722619315 | 13.1255103140479 | 7.46976052883995 |
| chr14 | 24184973 | 24184976 | - | 3.01188173062009 | 1.13699574206438 | 1.19322821036799 | 2.13421729395427 |
| chr14 | 24185283 | 24185293 | - | 5.01980288436682 | 4.54798296825753 | 4.77291284147196 | 12.8053037637256 |
| chr14 | 24185512 | 24185543 | - | 6.02376346124019 | 2.27399148412876 | 7.15936926220794 | 2.13421729395427 |
| chr14 | 24185940 | 24185963 | - | 1.00396057687336 | 1.13699574206438 | 2.38645642073598 | 5.33554323488568 |
| chr14 | 24186313 | 24186318 | - | 5.01980288436682 | 5.68497871032191 | 7.15936926220794 | 2.13421729395427 |
| chr14 | 24186340 | 24186362 | - | 5.01980288436682 | 7.95897019445067 | 4.77291284147196 | 6.40265188186281 |
| chr14 | 24186923 | 24186980 | - | 14.0554480762271 | 4.54798296825753 | 5.96614105183995 | 5.33554323488568 |
| chr14 | 24187669 | 24187673 | - | 2.00792115374673 | 2.27399148412876 | 1.19322821036799 | 2.13421729395427 |
| chr14 | 24188790 | 24188883 | - | 488.928800937329 | 180.782322988237 | 77.5598336739193 | 267.844270391261 |
| chr14 | 24209645 | 24209744 | - | 3.01188173062009 | 14.780944646837 | 3.57968463110397 | 1.06710864697714 |
| chr14 | 24213986 | 24214064 | - | 6.02376346124019 | 25.0139063254164 | 7.15936926220794 | 7.46976052883995 |
| chr14 | 24216902 | 24216963 | - | 411.623836518079 | 308.125846099448 | 87.1056593568632 | 114.180625226554 |
| chr14 | 24216990 | 24217013 | - | 6.02376346124019 | 3.41098722619315 | 3.57968463110397 | 7.46976052883995 |
| chr14 | 24218046 | 24218059 | - | 4.01584230749346 | 2.27399148412876 | 1.19322821036799 | 2.13421729395427 |
| chr14 | 24238966 | 24238991 | + | 1.00396057687336 | 4.54798296825753 | 1.19322821036799 | 4.26843458790854 |
| chr14 | 24239020 | 24239032 | + | 2.00792115374673 | 3.41098722619315 | 1.19322821036799 | 1.06710864697714 |
| chr14 | 24239063 | 24239225 | + | 7.02772403811355 | 14.780944646837 | 16.7051949451519 | 10.6710864697714 |
| chr14 | 24239672 | 24239733 | - | 13.0514874993537 | 17.0549361309657 | 3.57968463110397 | 3.20132594093141 |
| chr14 | 24240057 | 24240091 | - | 3.01188173062009 | 11.3699574206438 | 2.38645642073598 | 2.13421729395427 |
| chr14 | 24249145 | 24249147 | - | 21.0831721143407 | 28.4248935516096 | 4.77291284147196 | 4.26843458790854 |
| chr14 | 24265555 | 24265612 | - | 3.01188173062009 | 3.41098722619315 | 1.19322821036799 | 1.06710864697714 |
| chr14 | 24290622 | 24290635 | - | 3.01188173062009 | 1.13699574206438 | 4.77291284147196 | 1.06710864697714 |
| chr14 | 24290678 | 24290723 | - | 8.03168461498692 | 12.5069531627082 | 7.15936926220794 | 6.40265188186281 |
| chr14 | 24290771 | 24290816 | - | 72.2851615348822 | 54.5757956190903 | 26.2510206280958 | 21.3421729395427 |
| chr14 | 24290838 | 24290943 | - | 59.2336740355285 | 71.6307317500561 | 40.5697591525116 | 38.4159112911769 |
| chr14 | 24305045 | 24305142 | + | 44.174265382428 | 17.0549361309657 | 5.96614105183995 | 10.6710864697714 |
| chr14 | 24305174 | 24305338 | + | 44.174265382428 | 37.5208594881246 | 7.15936926220794 | 5.33554323488568 |
| chr14 | 24305371 | 24305374 | + | 12.0475269224804 | 9.09596593651506 | 3.57968463110397 | 2.13421729395427 |
| chr14 | 24305401 | 24305430 | + | 6.02376346124019 | 6.82197445238629 | 2.38645642073598 | 4.26843458790854 |
| chr14 | 24305449 | 24305539 | + | 24.0950538449607 | 19.3289276150945 | 4.77291284147196 | 10.6710864697714 |
| chr14 | 24307591 | 24307643 | + | 4.01584230749346 | 1.13699574206438 | 4.77291284147196 | 2.13421729395427 |
| chr14 | 24308021 | 24308167 | + | 32.1267384599477 | 18.1919318730301 | 16.7051949451519 | 29.8790421153598 |
| chr14 | 24438135 | 24438218 | + | 1.00396057687336 | 5.68497871032191 | 2.38645642073598 | 4.26843458790854 |
| chr14 | 24438243 | 24438283 | + | 2.00792115374673 | 1.13699574206438 | 3.57968463110397 | 1.06710864697714 |
| chr14 | 24438889 | 24438935 | + | 6.02376346124019 | 4.54798296825753 | 3.57968463110397 | 2.13421729395427 |
| chr14 | 24439069 | 24439226 | + | 11.043566345607 | 3.41098722619315 | 11.9322821036799 | 4.26843458790854 |
| chr14 | 24440000 | 24440003 | - | 1.00396057687336 | 1.13699574206438 | 1.19322821036799 | 1.06710864697714 |
| chr14 | 24441272 | 24441320 | + | 7.02772403811355 | 10.2329616785794 | 1.19322821036799 | 1.06710864697714 |
| chr14 | 30735640 | 30735780 | + | 7.02772403811355 | 6.82197445238629 | 8.35259747257593 | 5.33554323488568 |
| chr14 | 30894196 | 30894311 | - | 22.087132691214 | 50.0278126508328 | 64.4343233598714 | 53.3554323488568 |
| chr14 | 30894338 | 30894343 | - | 2.00792115374673 | 14.780944646837 | 3.57968463110397 | 5.33554323488568 |
| chr14 | 30894367 | 30894477 | - | 24.0950538449607 | 31.8358807778027 | 33.4103898903037 | 30.9461507623369 |
| chr14 | 30894561 | 30894606 | - | 18.0712903837206 | 39.7948509722534 | 23.8645642073598 | 20.2750642925656 |
| chr14 | 30894718 | 30894740 | - | 1.00396057687336 | 3.41098722619315 | 3.57968463110397 | 4.26843458790854 |
| chr14 | 30894989 | 30895035 | - | 19.0752509605939 | 26.1509020674808 | 64.4343233598714 | 53.3554323488568 |
| chr14 | 31100225 | 31100303 | - | 10.0396057687336 | 11.3699574206438 | 2.38645642073598 | 2.13421729395427 |
| chr14 | 31100830 | 31101065 | - | 3.01188173062009 | 3.41098722619315 | 1.19322821036799 | 7.46976052883995 |
| chr14 | 31400293 | 31400338 | - | 22.087132691214 | 12.5069531627082 | 2.38645642073598 | 1.06710864697714 |
| chr14 | 31400368 | 31400393 | - | 8.03168461498692 | 5.68497871032191 | 1.19322821036799 | 2.13421729395427 |
| chr14 | 31420753 | 31420776 | + | 12.0475269224804 | 14.780944646837 | 5.96614105183995 | 2.13421729395427 |
| chr14 | 31446290 | 31446330 | - | 6.02376346124019 | 5.68497871032191 | 3.57968463110397 | 5.33554323488568 |
| chr14 | 31446351 | 31446409 | - | 15.0594086531005 | 18.1919318730301 | 16.7051949451519 | 7.46976052883995 |
| chr14 | 31446432 | 31446474 | - | 21.0831721143407 | 30.6988850357383 | 17.8984231555198 | 19.2079556455884 |
| chr14 | 31446500 | 31446523 | - | 9.03564519186028 | 21.6029190992233 | 4.77291284147196 | 7.46976052883995 |
| chr14 | 31446554 | 31446557 | - | 2.00792115374673 | 1.13699574206438 | 2.38645642073598 | 1.06710864697714 |
| chr14 | 31446662 | 31446664 | - | 1.00396057687336 | 1.13699574206438 | 2.38645642073598 | 1.06710864697714 |
| chr14 | 31861124 | 31861150 | + | 7.02772403811355 | 5.68497871032191 | 2.38645642073598 | 4.26843458790854 |
| chr14 | 32155009 | 32155025 | + | 11.043566345607 | 7.95897019445067 | 2.38645642073598 | 2.13421729395427 |
| chr14 | 32155985 | 32156072 | + | 12.0475269224804 | 19.3289276150945 | 14.3187385244159 | 6.40265188186281 |
| chr14 | 32156158 | 32156172 | + | 56.2217923049084 | 70.4937360079917 | 7.15936926220794 | 1.06710864697714 |
| chr14 | 32156359 | 32156392 | + | 4.01584230749346 | 5.68497871032191 | 5.96614105183995 | 4.26843458790854 |
| chr14 | 33924871 | 33924897 | - | 2.00792115374673 | 2.27399148412876 | 13.1255103140479 | 17.0737383516342 |
| chr14 | 33925455 | 33925474 | - | 1.00396057687336 | 1.13699574206438 | 4.77291284147196 | 7.46976052883995 |
| chr14 | 33925484 | 33925659 | - | 13.0514874993537 | 18.1919318730301 | 134.834787771583 | 152.59653651773 |
| chr14 | 34434189 | 34434223 | - | 6.02376346124019 | 11.3699574206438 | 1.19322821036799 | 1.06710864697714 |
| chr14 | 34434260 | 34434369 | - | 36.1425807674411 | 47.753821166704 | 9.54582568294392 | 3.20132594093141 |
| chr14 | 34434391 | 34434534 | - | 94.3722942260963 | 101.19262104373 | 9.54582568294392 | 19.2079556455884 |
| chr14 | 34434561 | 34434601 | - | 2.00792115374673 | 1.13699574206438 | 2.38645642073598 | 3.20132594093141 |
| chr14 | 34434661 | 34434665 | - | 9.03564519186028 | 12.5069531627082 | 3.57968463110397 | 4.26843458790854 |
| chr14 | 34434703 | 34434833 | - | 71.2812009580089 | 108.014595496116 | 26.2510206280958 | 16.006629704657 |
| chr14 | 34434906 | 34435066 | - | 8.03168461498692 | 10.2329616785794 | 7.15936926220794 | 5.33554323488568 |
| chr14 | 34516037 | 34516071 | - | 4.01584230749346 | 3.41098722619315 | 1.19322821036799 | 7.46976052883995 |
| chr14 | 34516178 | 34516225 | - | 6.02376346124019 | 3.41098722619315 | 3.57968463110397 | 4.26843458790854 |
| chr14 | 34516466 | 34516493 | - | 6.02376346124019 | 12.5069531627082 | 2.38645642073598 | 3.20132594093141 |
| chr14 | 34562590 | 34562628 | - | 9.03564519186028 | 5.68497871032191 | 2.38645642073598 | 2.13421729395427 |
| chr14 | 34562842 | 34562971 | - | 8.03168461498692 | 6.82197445238629 | 11.9322821036799 | 10.6710864697714 |
| chr14 | 34563108 | 34563176 | - | 2.00792115374673 | 1.13699574206438 | 1.19322821036799 | 1.06710864697714 |
| chr14 | 34753006 | 34753027 | - | 8.03168461498692 | 2.27399148412876 | 1.19322821036799 | 7.46976052883995 |
| chr14 | 34753103 | 34753159 | - | 14.0554480762271 | 19.3289276150945 | 2.38645642073598 | 21.3421729395427 |
| chr14 | 34753225 | 34753239 | - | 5.01980288436682 | 3.41098722619315 | 1.19322821036799 | 6.40265188186281 |
| chr14 | 34753346 | 34753362 | - | 1.00396057687336 | 1.13699574206438 | 2.38645642073598 | 6.40265188186281 |
| chr14 | 34874822 | 34874823 | + | 5.01980288436682 | 9.09596593651506 | 2.38645642073598 | 1.06710864697714 |
| chr14 | 34875572 | 34875575 | + | 1.00396057687336 | 1.13699574206438 | 1.19322821036799 | 3.20132594093141 |
| chr14 | 35029059 | 35029158 | + | 12.0475269224804 | 3.41098722619315 | 7.15936926220794 | 10.6710864697714 |
| chr14 | 35029187 | 35029298 | + | 140.554480762271 | 118.247557174696 | 66.8207797806074 | 97.1068868749194 |
| chr14 | 35029329 | 35029384 | + | 31.1227778830743 | 21.6029190992233 | 15.5119667347839 | 27.7448248214055 |
| chr14 | 35029419 | 35029509 | + | 75.2970432655023 | 70.4937360079917 | 47.7291284147196 | 65.0936274656053 |
| chr14 | 35050219 | 35050224 | + | 15.0594086531005 | 12.5069531627082 | 2.38645642073598 | 10.6710864697714 |
| chr14 | 35050292 | 35050375 | + | 12.0475269224804 | 23.876910583352 | 8.35259747257593 | 10.6710864697714 |
| chr14 | 35081301 | 35081307 | + | 1.00396057687336 | 3.41098722619315 | 1.19322821036799 | 1.06710864697714 |
| chr14 | 35085494 | 35085585 | - | 9.03564519186028 | 12.5069531627082 | 9.54582568294392 | 2.13421729395427 |
| chr14 | 35274020 | 35274022 | + | 4.01584230749346 | 1.13699574206438 | 3.57968463110397 | 2.13421729395427 |
| chr14 | 35308033 | 35308066 | + | 4.01584230749346 | 10.2329616785794 | 8.35259747257593 | 12.8053037637256 |
| chr14 | 35312903 | 35312973 | + | 6.02376346124019 | 2.27399148412876 | 4.77291284147196 | 2.13421729395427 |
| chr14 | 35317258 | 35317271 | + | 1.00396057687336 | 2.27399148412876 | 2.38645642073598 | 6.40265188186281 |
| chr14 | 35317291 | 35317312 | + | 2.00792115374673 | 1.13699574206438 | 2.38645642073598 | 5.33554323488568 |
| chr14 | 35401539 | 35401677 | - | 180.712903837206 | 267.19399938513 | 21.4781077866238 | 42.6843458790854 |
| chr14 | 35401815 | 35401823 | - | 13.0514874993537 | 32.9728765198671 | 1.19322821036799 | 8.53686917581709 |
| chr14 | 35401986 | 35402059 | - | 32.1267384599477 | 39.7948509722534 | 19.0916513658878 | 9.60397782279422 |
| chr14 | 35871355 | 35871383 | + | 6.02376346124019 | 4.54798296825753 | 1.19322821036799 | 1.06710864697714 |
| chr14 | 35871623 | 35871647 | + | 68.2693192273888 | 69.3567402659273 | 25.0577924177278 | 13.8724124107028 |
| chr14 | 35871671 | 35871687 | + | 6.02376346124019 | 3.41098722619315 | 3.57968463110397 | 2.13421729395427 |
| chr14 | 35871731 | 35871755 | + | 8.03168461498692 | 5.68497871032191 | 3.57968463110397 | 2.13421729395427 |
| chr14 | 37172082 | 37172197 | + | 18.0712903837206 | 60.2607743294122 | 96.6514850398071 | 59.7580842307196 |
| chr14 | 39031988 | 39032136 | - | 8.03168461498692 | 13.6439489047726 | 5.96614105183995 | 3.20132594093141 |
| chr14 | 39032243 | 39032329 | - | 10.0396057687336 | 2.27399148412876 | 3.57968463110397 | 1.06710864697714 |
| chr14 | 39032352 | 39032795 | - | 30.1188173062009 | 22.7399148412876 | 15.5119667347839 | 22.4092815865198 |
| chr14 | 39033197 | 39033331 | - | 6.02376346124019 | 9.09596593651506 | 2.38645642073598 | 3.20132594093141 |
| chr14 | 39103290 | 39103292 | + | 1.00396057687336 | 2.27399148412876 | 1.19322821036799 | 1.06710864697714 |
| chr14 | 39136740 | 39136769 | + | 8.03168461498692 | 17.0549361309657 | 8.35259747257593 | 7.46976052883995 |
| chr14 | 39147837 | 39147878 | - | 24.0950538449607 | 63.6717615556054 | 25.0577924177278 | 27.7448248214055 |
| chr14 | 39150025 | 39150083 | - | 1.00396057687336 | 2.27399148412876 | 2.38645642073598 | 9.60397782279422 |
| chr14 | 39150111 | 39150127 | - | 2.00792115374673 | 6.82197445238629 | 1.19322821036799 | 12.8053037637256 |
| chr14 | 39150195 | 39150198 | - | 2.00792115374673 | 4.54798296825753 | 2.38645642073598 | 2.13421729395427 |
| chr14 | 39177610 | 39177701 | + | 69.2732798042621 | 69.3567402659273 | 44.1494437836156 | 56.5567582897882 |
| chr14 | 39177839 | 39177917 | + | 99.3920971104631 | 73.9047232341848 | 56.0817258872955 | 43.7514545260626 |
| chr14 | 39179069 | 39179253 | + | 555.190199010971 | 526.429028575809 | 307.852878274941 | 335.072115150821 |
| chr14 | 39181839 | 39182021 | + | 21.0831721143407 | 20.4659233571589 | 11.9322821036799 | 26.6777161744284 |
| chr14 | 39182042 | 39182088 | + | 9.03564519186028 | 21.6029190992233 | 13.1255103140479 | 4.26843458790854 |
| chr14 | 39182419 | 39182421 | + | 4.01584230749346 | 12.5069531627082 | 1.19322821036799 | 3.20132594093141 |
| chr14 | 39182645 | 39182657 | + | 21.0831721143407 | 38.657855230189 | 7.15936926220794 | 8.53686917581709 |
| chr14 | 39182759 | 39182806 | + | 3.01188173062009 | 5.68497871032191 | 5.96614105183995 | 3.20132594093141 |
| chr14 | 39182834 | 39182858 | + | 2.00792115374673 | 1.13699574206438 | 1.19322821036799 | 2.13421729395427 |
| chr14 | 39182883 | 39182901 | + | 5.01980288436682 | 4.54798296825753 | 2.38645642073598 | 3.20132594093141 |
| chr14 | 39350506 | 39350617 | + | 12.0475269224804 | 26.1509020674808 | 2.38645642073598 | 3.20132594093141 |
| chr14 | 45074211 | 45074222 | + | 38.1505019211879 | 42.0688424563821 | 13.1255103140479 | 8.53686917581709 |
| chr14 | 45114937 | 45114952 | + | 1.00396057687336 | 1.13699574206438 | 5.96614105183995 | 16.006629704657 |
| chr14 | 45115270 | 45115282 | + | 7.02772403811355 | 7.95897019445067 | 10.7390538933119 | 7.46976052883995 |
| chr14 | 45115751 | 45115753 | - | 3.01188173062009 | 4.54798296825753 | 5.96614105183995 | 1.06710864697714 |
| chr14 | 45115973 | 45116044 | - | 5.01980288436682 | 6.82197445238629 | 7.15936926220794 | 4.26843458790854 |
| chr14 | 45116074 | 45116121 | - | 6.02376346124019 | 12.5069531627082 | 4.77291284147196 | 4.26843458790854 |
| chr14 | 45116183 | 45116240 | - | 5.01980288436682 | 5.68497871032191 | 1.19322821036799 | 3.20132594093141 |
| chr14 | 45203903 | 45203906 | - | 3.01188173062009 | 3.41098722619315 | 1.19322821036799 | 1.06710864697714 |
| chr14 | 49577605 | 49577667 | - | 1.00396057687336 | 9.09596593651506 | 35.7968463110397 | 17.0737383516342 |
| chr14 | 49577701 | 49577751 | - | 4.01584230749346 | 3.41098722619315 | 9.54582568294392 | 1.06710864697714 |
| chr14 | 49583601 | 49583678 | - | 712.812009580089 | 1123.35179315961 | 541.725607507067 | 370.286700501066 |
| chr14 | 49614399 | 49614423 | + | 7.02772403811355 | 15.9179403889013 | 3.57968463110397 | 5.33554323488568 |
| chr14 | 49614529 | 49614543 | + | 34.1346596136944 | 39.7948509722534 | 11.9322821036799 | 19.2079556455884 |
| chr14 | 49614577 | 49614641 | + | 109.431702879197 | 184.19331021443 | 77.5598336739193 | 126.985928990279 |
| chr14 | 49618708 | 49618755 | - | 70.2772403811355 | 125.069531627082 | 75.1733772531833 | 60.8251928776967 |
| chr14 | 49622789 | 49622833 | + | 5.01980288436682 | 5.68497871032191 | 2.38645642073598 | 1.06710864697714 |
| chr14 | 49622869 | 49622873 | + | 40.1584230749346 | 35.2468680039958 | 8.35259747257593 | 17.0737383516342 |
| chr14 | 49623055 | 49623065 | + | 6.02376346124019 | 6.82197445238629 | 2.38645642073598 | 4.26843458790854 |
| chr14 | 49625382 | 49625393 | - | 84.3326884573626 | 98.9186295596012 | 19.0916513658878 | 64.0265188186281 |
| chr14 | 49782919 | 49782928 | + | 6.02376346124019 | 15.9179403889013 | 1.19322821036799 | 1.06710864697714 |
| chr14 | 49783012 | 49783109 | + | 4.01584230749346 | 18.1919318730301 | 3.57968463110397 | 1.06710864697714 |
| chr14 | 49894291 | 49894293 | + | 5.01980288436682 | 3.41098722619315 | 2.38645642073598 | 2.13421729395427 |
| chr14 | 49894314 | 49894336 | + | 5.01980288436682 | 5.68497871032191 | 7.15936926220794 | 23.476390233497 |
| chr14 | 49894376 | 49894436 | + | 31.1227778830743 | 54.5757956190903 | 22.6713359969918 | 19.2079556455884 |
| chr14 | 49894475 | 49894495 | + | 4.01584230749346 | 9.09596593651506 | 4.77291284147196 | 3.20132594093141 |
| chr14 | 49894554 | 49894598 | + | 43.1703048055547 | 101.19262104373 | 23.8645642073598 | 39.483019938154 |
| chr14 | 49894624 | 49894629 | + | 1.00396057687336 | 6.82197445238629 | 2.38645642073598 | 1.06710864697714 |
| chr14 | 49895940 | 49895955 | + | 2.00792115374673 | 1.13699574206438 | 3.57968463110397 | 3.20132594093141 |
| chr14 | 49896642 | 49896651 | + | 3.01188173062009 | 6.82197445238629 | 3.57968463110397 | 7.46976052883995 |
| chr14 | 49897005 | 49897011 | + | 2.00792115374673 | 13.6439489047726 | 3.57968463110397 | 4.26843458790854 |
| chr14 | 50108675 | 50108684 | - | 3.01188173062009 | 1.13699574206438 | 3.57968463110397 | 1.06710864697714 |
| chr14 | 50117679 | 50117709 | - | 7.02772403811355 | 14.780944646837 | 1.19322821036799 | 3.20132594093141 |
| chr14 | 50117942 | 50117978 | - | 18.0712903837206 | 19.3289276150945 | 16.7051949451519 | 16.006629704657 |
| chr14 | 50118032 | 50118036 | - | 1.00396057687336 | 4.54798296825753 | 4.77291284147196 | 5.33554323488568 |
| chr14 | 50418637 | 50418658 | - | 1.00396057687336 | 5.68497871032191 | 1.19322821036799 | 1.06710864697714 |
| chr14 | 50418894 | 50418904 | - | 5.01980288436682 | 11.3699574206438 | 2.38645642073598 | 5.33554323488568 |
| chr14 | 50419111 | 50419117 | - | 4.01584230749346 | 7.95897019445067 | 3.57968463110397 | 9.60397782279422 |
| chr14 | 50634129 | 50634150 | - | 6.02376346124019 | 4.54798296825753 | 1.19322821036799 | 4.26843458790854 |
| chr14 | 50634916 | 50634933 | - | 22.087132691214 | 40.9318467143178 | 17.8984231555198 | 13.8724124107028 |
| chr14 | 50635080 | 50635083 | - | 1.00396057687336 | 3.41098722619315 | 1.19322821036799 | 1.06710864697714 |
| chr14 | 50905235 | 50905347 | - | 11.043566345607 | 14.780944646837 | 7.15936926220794 | 3.20132594093141 |
| chr14 | 51254618 | 51254622 | + | 11.043566345607 | 1.13699574206438 | 3.57968463110397 | 1.06710864697714 |
| chr14 | 51255085 | 51255140 | + | 11.043566345607 | 7.95897019445067 | 3.57968463110397 | 6.40265188186281 |
| chr14 | 51255791 | 51255795 | + | 31.1227778830743 | 39.7948509722534 | 9.54582568294392 | 28.8119334683827 |
| chr14 | 52004381 | 52004421 | + | 11.043566345607 | 12.5069531627082 | 1.19322821036799 | 2.13421729395427 |
| chr14 | 52004497 | 52004559 | + | 12.0475269224804 | 27.2878978095452 | 3.57968463110397 | 6.40265188186281 |
| chr14 | 52004605 | 52004657 | + | 163.645574030358 | 280.837948289902 | 81.1395183050233 | 131.254363578188 |
| chr14 | 52639940 | 52639949 | - | 4.01584230749346 | 9.09596593651506 | 11.9322821036799 | 9.60397782279422 |
| chr14 | 52640134 | 52640162 | - | 5.01980288436682 | 6.82197445238629 | 8.35259747257593 | 11.7381951167485 |
| chr14 | 52640267 | 52640274 | - | 7.02772403811355 | 4.54798296825753 | 7.15936926220794 | 6.40265188186281 |
| chr14 | 52641044 | 52641089 | - | 2.00792115374673 | 2.27399148412876 | 4.77291284147196 | 5.33554323488568 |
| chr14 | 52641212 | 52641296 | - | 5.01980288436682 | 7.95897019445067 | 16.7051949451519 | 7.46976052883995 |
| chr14 | 52641321 | 52641328 | - | 20.0792115374673 | 26.1509020674808 | 19.0916513658878 | 37.3488026441998 |
| chr14 | 52641995 | 52642201 | - | 60.2376346124019 | 64.8087572976698 | 38.1833027317757 | 13.8724124107028 |
| chr14 | 52642260 | 52642281 | - | 8.03168461498692 | 6.82197445238629 | 7.15936926220794 | 5.33554323488568 |
| chr14 | 52642394 | 52642494 | - | 12.0475269224804 | 5.68497871032191 | 8.35259747257593 | 17.0737383516342 |
| chr14 | 52642593 | 52642623 | - | 2.00792115374673 | 1.13699574206438 | 3.57968463110397 | 1.06710864697714 |
| chr14 | 52642774 | 52642865 | - | 5.01980288436682 | 11.3699574206438 | 25.0577924177278 | 22.4092815865198 |
| chr14 | 52642970 | 52642990 | - | 14.0554480762271 | 4.54798296825753 | 10.7390538933119 | 1.06710864697714 |
| chr14 | 52643281 | 52643353 | - | 5.01980288436682 | 6.82197445238629 | 7.15936926220794 | 2.13421729395427 |
| chr14 | 52643433 | 52643567 | - | 22.087132691214 | 20.4659233571589 | 22.6713359969918 | 19.2079556455884 |
| chr14 | 52727634 | 52727719 | + | 5.01980288436682 | 4.54798296825753 | 3.57968463110397 | 7.46976052883995 |
| chr14 | 52727813 | 52727859 | + | 13.0514874993537 | 15.9179403889013 | 7.15936926220794 | 14.9395210576799 |
| chr14 | 52771109 | 52771183 | + | 7.02772403811355 | 12.5069531627082 | 16.7051949451519 | 22.4092815865198 |
| chr14 | 52771278 | 52771294 | + | 1.00396057687336 | 1.13699574206438 | 1.19322821036799 | 1.06710864697714 |
| chr14 | 52771473 | 52771515 | + | 18.0712903837206 | 28.4248935516096 | 14.3187385244159 | 6.40265188186281 |
| chr14 | 52773615 | 52773635 | + | 2.00792115374673 | 3.41098722619315 | 9.54582568294392 | 13.8724124107028 |
| chr14 | 52774498 | 52774523 | + | 5.01980288436682 | 1.13699574206438 | 7.15936926220794 | 4.26843458790854 |
| chr14 | 52774556 | 52774581 | + | 5.01980288436682 | 5.68497871032191 | 7.15936926220794 | 9.60397782279422 |
| chr14 | 52774683 | 52774687 | + | 10.0396057687336 | 5.68497871032191 | 14.3187385244159 | 10.6710864697714 |
| chr14 | 52775341 | 52775364 | - | 17.0673298068472 | 26.1509020674808 | 10.7390538933119 | 8.53686917581709 |
| chr14 | 52777357 | 52777461 | - | 21.0831721143407 | 70.4937360079917 | 60.8546387287675 | 5.33554323488568 |
| chr14 | 52777498 | 52777559 | - | 40.1584230749346 | 56.8497871032191 | 10.7390538933119 | 11.7381951167485 |
| chr14 | 52777580 | 52777606 | - | 27.1069355755808 | 38.657855230189 | 7.15936926220794 | 10.6710864697714 |
| chr14 | 52777742 | 52777780 | - | 13.0514874993537 | 15.9179403889013 | 4.77291284147196 | 5.33554323488568 |
| chr14 | 52777871 | 52777874 | - | 25.0990144218341 | 15.9179403889013 | 7.15936926220794 | 4.26843458790854 |
| chr14 | 52777993 | 52778031 | - | 8.03168461498692 | 18.1919318730301 | 1.19322821036799 | 1.06710864697714 |
| chr14 | 52778082 | 52778105 | - | 31.1227778830743 | 37.5208594881246 | 7.15936926220794 | 3.20132594093141 |
| chr14 | 52778204 | 52778291 | - | 106.419821148577 | 68.2197445238629 | 16.7051949451519 | 19.2079556455884 |
| chr14 | 54417925 | 54417934 | + | 4.01584230749346 | 12.5069531627082 | 1.19322821036799 | 7.46976052883995 |
| chr14 | 54420017 | 54420019 | + | 2.00792115374673 | 4.54798296825753 | 7.15936926220794 | 4.26843458790854 |
| chr14 | 54420180 | 54420193 | + | 7.02772403811355 | 15.9179403889013 | 1.19322821036799 | 13.8724124107028 |
| chr14 | 54424061 | 54424075 | - | 6.02376346124019 | 6.82197445238629 | 3.57968463110397 | 1.06710864697714 |
| chr14 | 54424114 | 54424149 | - | 10.0396057687336 | 7.95897019445067 | 4.77291284147196 | 4.26843458790854 |
| chr14 | 54424264 | 54424303 | - | 1.00396057687336 | 5.68497871032191 | 3.57968463110397 | 8.53686917581709 |
| chr14 | 54424477 | 54424500 | - | 3.01188173062009 | 6.82197445238629 | 7.15936926220794 | 2.13421729395427 |
| chr14 | 54425000 | 54425024 | - | 1.00396057687336 | 2.27399148412876 | 4.77291284147196 | 1.06710864697714 |
| chr14 | 54426951 | 54427097 | - | 490.936722091075 | 637.854611298118 | 263.703434491326 | 114.180625226554 |
| chr14 | 54427131 | 54427175 | - | 75.2970432655023 | 119.38455291676 | 99.0379414605431 | 33.0803680562912 |
| chr14 | 54427178 | 54427414 | - | 912.600164377888 | 1071.04998902465 | 439.10798141542 | 430.044784731786 |
| chr14 | 54427435 | 54427573 | - | 93.3683336492229 | 80.7266976865711 | 53.6952694665595 | 41.6172372321083 |
| chr14 | 54427629 | 54427664 | - | 2.00792115374673 | 5.68497871032191 | 4.77291284147196 | 4.26843458790854 |
| chr14 | 54427690 | 54427730 | - | 6.02376346124019 | 5.68497871032191 | 8.35259747257593 | 8.53686917581709 |
| chr14 | 54427754 | 54427839 | - | 14.0554480762271 | 15.9179403889013 | 13.1255103140479 | 4.26843458790854 |
| chr14 | 54476481 | 54476585 | - | 27.1069355755808 | 22.7399148412876 | 11.9322821036799 | 11.7381951167485 |
| chr14 | 54789212 | 54789257 | + | 17.0673298068472 | 81.8636934286355 | 73.9801490428153 | 34.1474767032683 |
| chr14 | 54793278 | 54793285 | + | 1.00396057687336 | 4.54798296825753 | 1.19322821036799 | 7.46976052883995 |
| chr14 | 54842578 | 54842605 | - | 2.00792115374673 | 1.13699574206438 | 2.38645642073598 | 4.26843458790854 |
| chr14 | 54941044 | 54941046 | - | 2.00792115374673 | 3.41098722619315 | 2.38645642073598 | 2.13421729395427 |
| chr14 | 54941469 | 54941498 | - | 3.01188173062009 | 9.09596593651506 | 7.15936926220794 | 6.40265188186281 |
| chr14 | 55044368 | 55044384 | + | 17.0673298068472 | 19.3289276150945 | 9.54582568294392 | 10.6710864697714 |
| chr14 | 55046463 | 55046526 | + | 18.0712903837206 | 44.3428339405109 | 8.35259747257593 | 10.6710864697714 |
| chr14 | 55046701 | 55046724 | + | 47.1861471130481 | 77.315710460378 | 22.6713359969918 | 27.7448248214055 |
| chr14 | 55046802 | 55046822 | + | 14.0554480762271 | 25.0139063254164 | 8.35259747257593 | 5.33554323488568 |
| chr14 | 55065112 | 55065131 | + | 3.01188173062009 | 2.27399148412876 | 3.57968463110397 | 2.13421729395427 |
| chr14 | 55065613 | 55065667 | + | 54.2138711511617 | 80.7266976865711 | 14.3187385244159 | 17.0737383516342 |
| chr14 | 55065788 | 55065820 | + | 2.00792115374673 | 2.27399148412876 | 1.19322821036799 | 4.26843458790854 |
| chr14 | 55065844 | 55065858 | + | 6.02376346124019 | 6.82197445238629 | 1.19322821036799 | 3.20132594093141 |
| chr14 | 55065940 | 55065947 | + | 1.00396057687336 | 2.27399148412876 | 1.19322821036799 | 1.06710864697714 |
| chr14 | 55066131 | 55066166 | + | 6.02376346124019 | 11.3699574206438 | 10.7390538933119 | 8.53686917581709 |
| chr14 | 55066668 | 55066691 | + | 5.01980288436682 | 1.13699574206438 | 2.38645642073598 | 1.06710864697714 |
| chr14 | 55067181 | 55067200 | + | 19.0752509605939 | 19.3289276150945 | 2.38645642073598 | 9.60397782279422 |
| chr14 | 55067368 | 55067383 | + | 9.03564519186028 | 1.13699574206438 | 2.38645642073598 | 8.53686917581709 |
| chr14 | 55067678 | 55067699 | + | 27.1069355755808 | 18.1919318730301 | 19.0916513658878 | 20.2750642925656 |
| chr14 | 55068630 | 55068647 | + | 7.02772403811355 | 3.41098722619315 | 2.38645642073598 | 3.20132594093141 |
| chr14 | 55068851 | 55068887 | + | 8.03168461498692 | 5.68497871032191 | 7.15936926220794 | 3.20132594093141 |
| chr14 | 55068938 | 55069006 | + | 53.2099105742883 | 64.8087572976698 | 38.1833027317757 | 22.4092815865198 |
| chr14 | 55069045 | 55069053 | + | 6.02376346124019 | 7.95897019445067 | 3.57968463110397 | 1.06710864697714 |
| chr14 | 55069074 | 55069093 | + | 48.1901076899215 | 59.1237785873479 | 32.2171616799357 | 24.5434988804741 |
| chr14 | 55069117 | 55069210 | + | 67.2653586505154 | 76.1787147183136 | 36.9900745214077 | 21.3421729395427 |
| chr14 | 55069304 | 55069319 | + | 8.03168461498692 | 15.9179403889013 | 8.35259747257593 | 3.20132594093141 |
| chr14 | 55069440 | 55069453 | + | 6.02376346124019 | 3.41098722619315 | 1.19322821036799 | 2.13421729395427 |
| chr14 | 55148177 | 55148186 | - | 19.0752509605939 | 20.4659233571589 | 11.9322821036799 | 12.8053037637256 |
| chr14 | 55353244 | 55353263 | + | 10.0396057687336 | 10.2329616785794 | 3.57968463110397 | 8.53686917581709 |
| chr14 | 55367065 | 55367080 | - | 9.03564519186028 | 4.54798296825753 | 1.19322821036799 | 1.06710864697714 |
| chr14 | 55369201 | 55369223 | - | 1.00396057687336 | 4.54798296825753 | 5.96614105183995 | 3.20132594093141 |
| chr14 | 55684127 | 55684157 | + | 2.00792115374673 | 4.54798296825753 | 1.19322821036799 | 3.20132594093141 |
| chr14 | 55684257 | 55684262 | + | 8.03168461498692 | 5.68497871032191 | 2.38645642073598 | 1.06710864697714 |
| chr14 | 55684539 | 55684565 | + | 7.02772403811355 | 6.82197445238629 | 3.57968463110397 | 5.33554323488568 |
| chr14 | 56647842 | 56647858 | + | 3.01188173062009 | 2.27399148412876 | 4.77291284147196 | 2.13421729395427 |
| chr14 | 57203896 | 57203898 | - | 6.02376346124019 | 3.41098722619315 | 4.77291284147196 | 1.06710864697714 |
| chr14 | 57203967 | 57203991 | - | 6.02376346124019 | 3.41098722619315 | 15.5119667347839 | 4.26843458790854 |
| chr14 | 57204067 | 57204083 | - | 1.00396057687336 | 2.27399148412876 | 4.77291284147196 | 2.13421729395427 |
| chr14 | 57204111 | 57204121 | - | 8.03168461498692 | 3.41098722619315 | 7.15936926220794 | 6.40265188186281 |
| chr14 | 57204836 | 57204839 | - | 9.03564519186028 | 13.6439489047726 | 1.19322821036799 | 1.06710864697714 |
| chr14 | 57206790 | 57206800 | - | 2.00792115374673 | 5.68497871032191 | 3.57968463110397 | 2.13421729395427 |
| chr14 | 57207015 | 57207036 | - | 6.02376346124019 | 7.95897019445067 | 4.77291284147196 | 5.33554323488568 |
| chr14 | 57409930 | 57409949 | + | 12.0475269224804 | 14.780944646837 | 5.96614105183995 | 4.26843458790854 |
| chr14 | 57410356 | 57410383 | + | 16.0633692299738 | 13.6439489047726 | 2.38645642073598 | 1.06710864697714 |
| chr14 | 57410568 | 57410571 | + | 1.00396057687336 | 1.13699574206438 | 1.19322821036799 | 2.13421729395427 |
| chr14 | 57411024 | 57411056 | + | 19.0752509605939 | 18.1919318730301 | 16.7051949451519 | 48.0198891139711 |
| chr14 | 57411182 | 57411190 | + | 5.01980288436682 | 3.41098722619315 | 3.57968463110397 | 4.26843458790854 |
| chr14 | 57411969 | 57411971 | + | 7.02772403811355 | 7.95897019445067 | 1.19322821036799 | 7.46976052883995 |
| chr14 | 57412114 | 57412135 | + | 16.0633692299738 | 20.4659233571589 | 7.15936926220794 | 8.53686917581709 |
| chr14 | 57412149 | 57412152 | + | 2.00792115374673 | 5.68497871032191 | 2.38645642073598 | 1.06710864697714 |
| chr14 | 57412290 | 57412309 | + | 3.01188173062009 | 5.68497871032191 | 1.19322821036799 | 4.26843458790854 |
| chr14 | 57412346 | 57412383 | + | 15.0594086531005 | 37.5208594881246 | 8.35259747257593 | 10.6710864697714 |
| chr14 | 57412414 | 57412466 | + | 8.03168461498692 | 12.5069531627082 | 5.96614105183995 | 3.20132594093141 |
| chr14 | 57412499 | 57412528 | + | 15.0594086531005 | 20.4659233571589 | 10.7390538933119 | 8.53686917581709 |
| chr14 | 57412554 | 57412589 | + | 8.03168461498692 | 12.5069531627082 | 13.1255103140479 | 5.33554323488568 |
| chr14 | 58234626 | 58234657 | + | 3.01188173062009 | 7.95897019445067 | 2.38645642073598 | 6.40265188186281 |
| chr14 | 58234725 | 58234754 | + | 9.03564519186028 | 18.1919318730301 | 4.77291284147196 | 2.13421729395427 |
| chr14 | 58271891 | 58271981 | + | 147.582204800385 | 142.124467758048 | 68.0140079909754 | 68.2949534065367 |
| chr14 | 58372398 | 58372420 | + | 2.00792115374673 | 5.68497871032191 | 3.57968463110397 | 2.13421729395427 |
| chr14 | 58408693 | 58408743 | - | 12.0475269224804 | 18.1919318730301 | 13.1255103140479 | 3.20132594093141 |
| chr14 | 58408769 | 58408818 | - | 15.0594086531005 | 53.438799877026 | 10.7390538933119 | 10.6710864697714 |
| chr14 | 58408889 | 58408900 | - | 3.01188173062009 | 5.68497871032191 | 2.38645642073598 | 10.6710864697714 |
| chr14 | 58408941 | 58408981 | - | 5.01980288436682 | 6.82197445238629 | 4.77291284147196 | 2.13421729395427 |
| chr14 | 58547899 | 58547937 | + | 4.01584230749346 | 5.68497871032191 | 1.19322821036799 | 3.20132594093141 |
| chr14 | 59369425 | 59369434 | + | 6.02376346124019 | 13.6439489047726 | 7.15936926220794 | 5.33554323488568 |
| chr14 | 59371063 | 59371082 | + | 2.00792115374673 | 4.54798296825753 | 1.19322821036799 | 1.06710864697714 |
| chr14 | 59503870 | 59503880 | + | 1.00396057687336 | 5.68497871032191 | 4.77291284147196 | 1.06710864697714 |
| chr14 | 59504040 | 59504108 | + | 9.03564519186028 | 5.68497871032191 | 3.57968463110397 | 4.26843458790854 |
| chr14 | 59504436 | 59504451 | + | 4.01584230749346 | 14.780944646837 | 1.19322821036799 | 4.26843458790854 |
| chr14 | 59504512 | 59504515 | + | 14.0554480762271 | 7.95897019445067 | 2.38645642073598 | 4.26843458790854 |
| chr14 | 60138658 | 60138659 | + | 5.01980288436682 | 4.54798296825753 | 2.38645642073598 | 4.26843458790854 |
| chr14 | 60144925 | 60144935 | - | 5.01980288436682 | 11.3699574206438 | 4.77291284147196 | 1.06710864697714 |
| chr14 | 60292572 | 60292645 | + | 21.0831721143407 | 36.3838637460602 | 13.1255103140479 | 28.8119334683827 |
| chr14 | 60292887 | 60292921 | + | 16.0633692299738 | 25.0139063254164 | 8.35259747257593 | 12.8053037637256 |
| chr14 | 60292942 | 60292956 | + | 2.00792115374673 | 6.82197445238629 | 2.38645642073598 | 4.26843458790854 |
| chr14 | 60293016 | 60293065 | + | 77.3049644192491 | 134.165497563597 | 50.1155848354556 | 41.6172372321083 |
| chr14 | 60293089 | 60293091 | + | 4.01584230749346 | 1.13699574206438 | 3.57968463110397 | 2.13421729395427 |
| chr14 | 60293146 | 60293183 | + | 107.42378172545 | 130.754510337404 | 75.1733772531833 | 45.8856718200168 |
| chr14 | 60293236 | 60293255 | + | 22.087132691214 | 14.780944646837 | 11.9322821036799 | 6.40265188186281 |
| chr14 | 60294176 | 60294186 | + | 1.00396057687336 | 1.13699574206438 | 2.38645642073598 | 1.06710864697714 |
| chr14 | 60296288 | 60296306 | + | 2.00792115374673 | 5.68497871032191 | 2.38645642073598 | 5.33554323488568 |
| chr14 | 60968514 | 60968517 | + | 2.00792115374673 | 5.68497871032191 | 3.57968463110397 | 1.06710864697714 |
| chr14 | 60968588 | 60968597 | + | 8.03168461498692 | 22.7399148412876 | 7.15936926220794 | 4.26843458790854 |
| chr14 | 60968620 | 60968638 | + | 17.0673298068472 | 48.8908169087684 | 17.8984231555198 | 30.9461507623369 |
| chr14 | 61747032 | 61747085 | + | 75.2970432655023 | 102.329616785794 | 8.35259747257593 | 11.7381951167485 |
| chr14 | 63372639 | 63372651 | - | 41.1623836518079 | 46.6168254246397 | 10.7390538933119 | 7.46976052883995 |
| chr14 | 63373953 | 63373991 | - | 3.01188173062009 | 4.54798296825753 | 5.96614105183995 | 2.13421729395427 |
| chr14 | 63374034 | 63374066 | - | 13.0514874993537 | 9.09596593651506 | 7.15936926220794 | 13.8724124107028 |
| chr14 | 63374144 | 63374175 | - | 5.01980288436682 | 4.54798296825753 | 2.38645642073598 | 2.13421729395427 |
| chr14 | 63374407 | 63374422 | - | 4.01584230749346 | 3.41098722619315 | 5.96614105183995 | 6.40265188186281 |
| chr14 | 63374472 | 63374537 | - | 31.1227778830743 | 20.4659233571589 | 46.5359002043516 | 7.46976052883995 |
| chr14 | 63375326 | 63375356 | - | 20.0792115374673 | 22.7399148412876 | 13.1255103140479 | 4.26843458790854 |
| chr14 | 63375418 | 63375422 | - | 7.02772403811355 | 12.5069531627082 | 9.54582568294392 | 3.20132594093141 |
| chr14 | 63375457 | 63375477 | - | 20.0792115374673 | 20.4659233571589 | 9.54582568294392 | 11.7381951167485 |
| chr14 | 63375808 | 63375823 | - | 2.00792115374673 | 5.68497871032191 | 2.38645642073598 | 1.06710864697714 |
| chr14 | 63597261 | 63597262 | - | 1.00396057687336 | 1.13699574206438 | 1.19322821036799 | 1.06710864697714 |
| chr14 | 63597803 | 63597807 | - | 1.00396057687336 | 1.13699574206438 | 2.38645642073598 | 6.40265188186281 |
| chr14 | 63642123 | 63642167 | + | 25.0990144218341 | 22.7399148412876 | 11.9322821036799 | 12.8053037637256 |
| chr14 | 63684544 | 63684546 | - | 32.1267384599477 | 34.1098722619315 | 3.57968463110397 | 4.26843458790854 |
| chr14 | 64459821 | 64459847 | + | 10.0396057687336 | 11.3699574206438 | 4.77291284147196 | 3.20132594093141 |
| chr14 | 64459869 | 64459903 | + | 13.0514874993537 | 18.1919318730301 | 1.19322821036799 | 1.06710864697714 |
| chr14 | 64481914 | 64481939 | - | 6.02376346124019 | 30.6988850357383 | 3.57968463110397 | 1.06710864697714 |
| chr14 | 64482109 | 64482116 | - | 67.2653586505154 | 236.495114349391 | 22.6713359969918 | 20.2750642925656 |
| chr14 | 64485203 | 64485211 | - | 12.0475269224804 | 21.6029190992233 | 5.96614105183995 | 2.13421729395427 |
| chr14 | 64932146 | 64932201 | + | 7.02772403811355 | 3.41098722619315 | 7.15936926220794 | 4.26843458790854 |
| chr14 | 64932242 | 64932298 | + | 25.0990144218341 | 30.6988850357383 | 13.1255103140479 | 8.53686917581709 |
| chr14 | 64934575 | 64934598 | + | 137.542599031651 | 156.905412404885 | 7.15936926220794 | 7.46976052883995 |
| chr14 | 65075172 | 65075175 | - | 7.02772403811355 | 4.54798296825753 | 7.15936926220794 | 3.20132594093141 |
| chr14 | 65075337 | 65075340 | - | 16.0633692299738 | 10.2329616785794 | 4.77291284147196 | 1.06710864697714 |
| chr14 | 65075664 | 65075736 | - | 31.1227778830743 | 25.0139063254164 | 16.7051949451519 | 19.2079556455884 |
| chr14 | 65075791 | 65075811 | - | 14.0554480762271 | 6.82197445238629 | 10.7390538933119 | 5.33554323488568 |
| chr14 | 65075870 | 65075883 | - | 2.00792115374673 | 2.27399148412876 | 1.19322821036799 | 4.26843458790854 |
| chr14 | 65076006 | 65076009 | - | 2.00792115374673 | 1.13699574206438 | 1.19322821036799 | 1.06710864697714 |
| chr14 | 65076105 | 65076107 | - | 11.043566345607 | 4.54798296825753 | 3.57968463110397 | 1.06710864697714 |
| chr14 | 67181515 | 67181542 | + | 6.02376346124019 | 5.68497871032191 | 3.57968463110397 | 1.06710864697714 |
| chr14 | 67333490 | 67333523 | + | 12.0475269224804 | 7.95897019445067 | 1.19322821036799 | 1.06710864697714 |
| chr14 | 67333857 | 67333885 | + | 3.01188173062009 | 5.68497871032191 | 1.19322821036799 | 2.13421729395427 |
| chr14 | 67335387 | 67335393 | + | 5.01980288436682 | 6.82197445238629 | 1.19322821036799 | 6.40265188186281 |
| chr14 | 67338099 | 67338132 | - | 38.1505019211879 | 39.7948509722534 | 4.77291284147196 | 8.53686917581709 |
| chr14 | 67338239 | 67338242 | - | 2.00792115374673 | 3.41098722619315 | 1.19322821036799 | 3.20132594093141 |
| chr14 | 67338638 | 67338649 | - | 9.03564519186028 | 4.54798296825753 | 1.19322821036799 | 12.8053037637256 |
| chr14 | 67383654 | 67383659 | + | 7.02772403811355 | 13.6439489047726 | 1.19322821036799 | 10.6710864697714 |
| chr14 | 67383777 | 67383792 | + | 19.0752509605939 | 28.4248935516096 | 5.96614105183995 | 35.2145853502455 |
| chr14 | 67384205 | 67384207 | + | 9.03564519186028 | 6.82197445238629 | 1.19322821036799 | 2.13421729395427 |
| chr14 | 67384342 | 67384356 | + | 6.02376346124019 | 11.3699574206438 | 3.57968463110397 | 4.26843458790854 |
| chr14 | 67384944 | 67384966 | + | 1.00396057687336 | 1.13699574206438 | 1.19322821036799 | 12.8053037637256 |
| chr14 | 67385626 | 67385645 | + | 4.01584230749346 | 2.27399148412876 | 3.57968463110397 | 9.60397782279422 |
| chr14 | 67647157 | 67647265 | - | 80.3168461498692 | 88.6856678810218 | 131.255103140479 | 50.1541064079254 |
| chr14 | 67647308 | 67647336 | - | 9.03564519186028 | 13.6439489047726 | 14.3187385244159 | 12.8053037637256 |
| chr14 | 67647429 | 67647447 | - | 15.0594086531005 | 6.82197445238629 | 5.96614105183995 | 2.13421729395427 |
| chr14 | 67647528 | 67647531 | - | 2.00792115374673 | 6.82197445238629 | 7.15936926220794 | 4.26843458790854 |
| chr14 | 67647600 | 67647609 | - | 1.00396057687336 | 2.27399148412876 | 9.54582568294392 | 2.13421729395427 |
| chr14 | 67650978 | 67651009 | + | 5.01980288436682 | 26.1509020674808 | 27.4442488384638 | 22.4092815865198 |
| chr14 | 67651213 | 67651254 | - | 11.043566345607 | 17.0549361309657 | 13.1255103140479 | 19.2079556455884 |
| chr14 | 67651319 | 67651394 | - | 127.502993262917 | 122.795540142953 | 101.424397881279 | 96.0397782279422 |
| chr14 | 67659832 | 67659853 | - | 9.03564519186028 | 3.41098722619315 | 10.7390538933119 | 8.53686917581709 |
| chr14 | 67676882 | 67676896 | - | 9.03564519186028 | 10.2329616785794 | 2.38645642073598 | 3.20132594093141 |
| chr14 | 67676962 | 67676993 | - | 3.01188173062009 | 10.2329616785794 | 5.96614105183995 | 2.13421729395427 |
| chr14 | 67677035 | 67677073 | - | 145.574283646638 | 237.632110091456 | 59.6614105183995 | 172.871600810296 |
| chr14 | 67677288 | 67677309 | - | 3.01188173062009 | 3.41098722619315 | 3.57968463110397 | 1.06710864697714 |
| chr14 | 67677404 | 67677448 | - | 14.0554480762271 | 14.780944646837 | 8.35259747257593 | 17.0737383516342 |
| chr14 | 67677589 | 67677646 | - | 16.0633692299738 | 13.6439489047726 | 4.77291284147196 | 7.46976052883995 |
| chr14 | 67677681 | 67677701 | - | 3.01188173062009 | 1.13699574206438 | 1.19322821036799 | 6.40265188186281 |
| chr14 | 67677736 | 67677747 | - | 2.00792115374673 | 3.41098722619315 | 2.38645642073598 | 3.20132594093141 |
| chr14 | 67747638 | 67747647 | - | 2.00792115374673 | 6.82197445238629 | 1.19322821036799 | 3.20132594093141 |
| chr14 | 68787978 | 68787989 | - | 1.00396057687336 | 1.13699574206438 | 2.38645642073598 | 1.06710864697714 |
| chr14 | 68788677 | 68788716 | - | 5.01980288436682 | 5.68497871032191 | 4.77291284147196 | 1.06710864697714 |
| chr14 | 68788752 | 68788841 | - | 43.1703048055547 | 67.0827487817985 | 25.0577924177278 | 38.4159112911769 |
| chr14 | 68788884 | 68788890 | - | 13.0514874993537 | 15.9179403889013 | 13.1255103140479 | 23.476390233497 |
| chr14 | 68790227 | 68790231 | - | 12.0475269224804 | 14.780944646837 | 13.1255103140479 | 5.33554323488568 |
| chr14 | 68874302 | 68874435 | - | 34.1346596136944 | 28.4248935516096 | 11.9322821036799 | 22.4092815865198 |
| chr14 | 68874500 | 68874607 | - | 45.1782259593014 | 31.8358807778027 | 21.4781077866238 | 24.5434988804741 |
| chr14 | 68874634 | 68874754 | - | 57.2257528817818 | 39.7948509722534 | 17.8984231555198 | 37.3488026441998 |
| chr14 | 68874777 | 68874847 | - | 32.1267384599477 | 20.4659233571589 | 5.96614105183995 | 21.3421729395427 |
| chr14 | 69052039 | 69052049 | - | 3.01188173062009 | 6.82197445238629 | 2.38645642073598 | 1.06710864697714 |
| chr14 | 69052215 | 69052235 | - | 3.01188173062009 | 6.82197445238629 | 7.15936926220794 | 5.33554323488568 |
| chr14 | 69052510 | 69052522 | - | 1.00396057687336 | 2.27399148412876 | 5.96614105183995 | 1.06710864697714 |
| chr14 | 69052609 | 69052627 | - | 1.00396057687336 | 1.13699574206438 | 7.15936926220794 | 2.13421729395427 |
| chr14 | 69052685 | 69052707 | - | 2.00792115374673 | 3.41098722619315 | 8.35259747257593 | 6.40265188186281 |
| chr14 | 69052739 | 69052740 | - | 1.00396057687336 | 1.13699574206438 | 1.19322821036799 | 1.06710864697714 |
| chr14 | 69242738 | 69242739 | + | 3.01188173062009 | 5.68497871032191 | 1.19322821036799 | 1.06710864697714 |
| chr14 | 69380156 | 69380208 | - | 242.958459603354 | 194.426271893009 | 102.617626091647 | 255.038966627535 |
| chr14 | 69380299 | 69380304 | - | 2.00792115374673 | 1.13699574206438 | 4.77291284147196 | 3.20132594093141 |
| chr14 | 69380362 | 69380434 | - | 75.2970432655023 | 46.6168254246397 | 16.7051949451519 | 69.3620620535138 |
| chr14 | 69380472 | 69380517 | - | 241.954499026481 | 102.329616785794 | 119.322821036799 | 321.199702740118 |
| chr14 | 69458722 | 69458734 | + | 38.1505019211879 | 37.5208594881246 | 14.3187385244159 | 10.6710864697714 |
| chr14 | 69459074 | 69459080 | + | 2.00792115374673 | 3.41098722619315 | 1.19322821036799 | 1.06710864697714 |
| chr14 | 69459378 | 69459540 | + | 83.3287278804893 | 73.9047232341848 | 57.2749540976635 | 37.3488026441998 |
| chr14 | 69459667 | 69459672 | + | 1.00396057687336 | 5.68497871032191 | 4.77291284147196 | 3.20132594093141 |
| chr14 | 69459692 | 69459742 | + | 4.01584230749346 | 4.54798296825753 | 7.15936926220794 | 3.20132594093141 |
| chr14 | 69459806 | 69459841 | + | 8.03168461498692 | 10.2329616785794 | 10.7390538933119 | 5.33554323488568 |
| chr14 | 69459934 | 69459959 | + | 1.00396057687336 | 3.41098722619315 | 1.19322821036799 | 2.13421729395427 |
| chr14 | 69460004 | 69460034 | + | 5.01980288436682 | 5.68497871032191 | 8.35259747257593 | 4.26843458790854 |
| chr14 | 69460158 | 69460199 | + | 8.03168461498692 | 9.09596593651506 | 2.38645642073598 | 8.53686917581709 |
| chr14 | 69460428 | 69460463 | + | 9.03564519186028 | 9.09596593651506 | 4.77291284147196 | 7.46976052883995 |
| chr14 | 69460514 | 69460563 | + | 2.00792115374673 | 2.27399148412876 | 5.96614105183995 | 1.06710864697714 |
| chr14 | 69460957 | 69461003 | + | 3.01188173062009 | 5.68497871032191 | 2.38645642073598 | 2.13421729395427 |
| chr14 | 69461447 | 69461484 | + | 6.02376346124019 | 2.27399148412876 | 10.7390538933119 | 6.40265188186281 |
| chr14 | 69461541 | 69461635 | + | 18.0712903837206 | 15.9179403889013 | 22.6713359969918 | 13.8724124107028 |
| chr14 | 69461657 | 69461675 | + | 11.043566345607 | 11.3699574206438 | 5.96614105183995 | 4.26843458790854 |
| chr14 | 69461710 | 69461726 | + | 4.01584230749346 | 3.41098722619315 | 1.19322821036799 | 2.13421729395427 |
| chr14 | 69461752 | 69461775 | + | 5.01980288436682 | 5.68497871032191 | 10.7390538933119 | 2.13421729395427 |
| chr14 | 69461822 | 69461844 | + | 18.0712903837206 | 4.54798296825753 | 2.38645642073598 | 10.6710864697714 |
| chr14 | 69462053 | 69462072 | + | 4.01584230749346 | 5.68497871032191 | 4.77291284147196 | 5.33554323488568 |
| chr14 | 69462289 | 69462299 | + | 3.01188173062009 | 1.13699574206438 | 2.38645642073598 | 1.06710864697714 |
| chr14 | 69711446 | 69711447 | + | 1.00396057687336 | 1.13699574206438 | 2.38645642073598 | 1.06710864697714 |
| chr14 | 69712932 | 69712937 | + | 2.00792115374673 | 2.27399148412876 | 13.1255103140479 | 6.40265188186281 |
| chr14 | 69713020 | 69713028 | + | 1.00396057687336 | 3.41098722619315 | 9.54582568294392 | 3.20132594093141 |
| chr14 | 69769161 | 69769320 | + | 93.3683336492229 | 89.8226636230862 | 126.482190299007 | 109.912190638645 |
| chr14 | 69769351 | 69769364 | + | 3.01188173062009 | 6.82197445238629 | 3.57968463110397 | 1.06710864697714 |
| chr14 | 69771265 | 69771501 | + | 58.2297134586551 | 48.8908169087684 | 91.8785721983352 | 125.918820343302 |
| chr14 | 69771513 | 69771729 | + | 1106.36455571445 | 1006.24123172698 | 693.265590223802 | 829.143418701234 |
| chr14 | 69771761 | 69771810 | + | 11.043566345607 | 10.2329616785794 | 20.2848795762558 | 2.13421729395427 |
| chr14 | 69771851 | 69771927 | + | 673.657547082028 | 440.017352178916 | 509.508445827131 | 459.923826847146 |
| chr14 | 69771949 | 69771985 | + | 6.02376346124019 | 18.1919318730301 | 8.35259747257593 | 11.7381951167485 |
| chr14 | 70326098 | 70326250 | - | 128.506953839791 | 105.740604011988 | 13.1255103140479 | 26.6777161744284 |
| chr14 | 70326325 | 70326430 | - | 287.132724985782 | 206.933225055718 | 105.004082512383 | 185.676904574022 |
| chr14 | 70368247 | 70368253 | - | 15.0594086531005 | 6.82197445238629 | 4.77291284147196 | 1.06710864697714 |
| chr14 | 70368454 | 70368457 | - | 14.0554480762271 | 10.2329616785794 | 1.19322821036799 | 1.06710864697714 |
| chr14 | 70370838 | 70370860 | - | 3.01188173062009 | 3.41098722619315 | 1.19322821036799 | 1.06710864697714 |
| chr14 | 70370888 | 70370909 | - | 19.0752509605939 | 20.4659233571589 | 2.38645642073598 | 5.33554323488568 |
| chr14 | 70581464 | 70581466 | - | 1.00396057687336 | 3.41098722619315 | 2.38645642073598 | 3.20132594093141 |
| chr14 | 70583828 | 70583838 | - | 3.01188173062009 | 9.09596593651506 | 11.9322821036799 | 9.60397782279422 |
| chr14 | 70584306 | 70584330 | - | 4.01584230749346 | 1.13699574206438 | 3.57968463110397 | 2.13421729395427 |
| chr14 | 70584460 | 70584467 | - | 4.01584230749346 | 4.54798296825753 | 3.57968463110397 | 7.46976052883995 |
| chr14 | 71111018 | 71111024 | + | 5.01980288436682 | 3.41098722619315 | 1.19322821036799 | 2.13421729395427 |
| chr14 | 71739340 | 71739341 | + | 4.01584230749346 | 10.2329616785794 | 1.19322821036799 | 1.06710864697714 |
| chr14 | 72959265 | 72959274 | + | 8.03168461498692 | 6.82197445238629 | 16.7051949451519 | 9.60397782279422 |
| chr14 | 73099922 | 73099938 | + | 2.00792115374673 | 2.27399148412876 | 17.8984231555198 | 4.26843458790854 |
| chr14 | 73100092 | 73100096 | + | 1.00396057687336 | 5.68497871032191 | 9.54582568294392 | 2.13421729395427 |
| chr14 | 73100184 | 73100209 | + | 21.0831721143407 | 27.2878978095452 | 11.9322821036799 | 12.8053037637256 |
| chr14 | 73100299 | 73100371 | + | 22.087132691214 | 14.780944646837 | 17.8984231555198 | 12.8053037637256 |
| chr14 | 73100424 | 73100429 | + | 6.02376346124019 | 4.54798296825753 | 9.54582568294392 | 1.06710864697714 |
| chr14 | 73100507 | 73100532 | + | 1.00396057687336 | 1.13699574206438 | 4.77291284147196 | 1.06710864697714 |
| chr14 | 73100589 | 73100606 | + | 2.00792115374673 | 2.27399148412876 | 1.19322821036799 | 2.13421729395427 |
| chr14 | 73119711 | 73120094 | + | 306.207975946376 | 283.111939774031 | 408.084047945852 | 218.757272630313 |
| chr14 | 73121172 | 73121183 | + | 5.01980288436682 | 10.2329616785794 | 3.57968463110397 | 3.20132594093141 |
| chr14 | 73219802 | 73219826 | + | 32.1267384599477 | 28.4248935516096 | 17.8984231555198 | 16.006629704657 |
| chr14 | 73219964 | 73220005 | + | 5.01980288436682 | 10.2329616785794 | 2.38645642073598 | 2.13421729395427 |
| chr14 | 73220197 | 73220284 | + | 45.1782259593014 | 47.753821166704 | 15.5119667347839 | 20.2750642925656 |
| chr14 | 73223490 | 73223509 | + | 10.0396057687336 | 11.3699574206438 | 2.38645642073598 | 5.33554323488568 |
| chr14 | 73275329 | 73275333 | - | 3.01188173062009 | 9.09596593651506 | 2.38645642073598 | 2.13421729395427 |
| chr14 | 73276043 | 73276066 | - | 10.0396057687336 | 12.5069531627082 | 9.54582568294392 | 18.1408469986113 |
| chr14 | 73493315 | 73493324 | + | 12.0475269224804 | 5.68497871032191 | 1.19322821036799 | 9.60397782279422 |
| chr14 | 73711814 | 73711858 | - | 11.043566345607 | 21.6029190992233 | 2.38645642073598 | 1.06710864697714 |
| chr14 | 73711907 | 73711914 | - | 13.0514874993537 | 10.2329616785794 | 2.38645642073598 | 3.20132594093141 |
| chr14 | 73712323 | 73712341 | - | 2.00792115374673 | 6.82197445238629 | 1.19322821036799 | 1.06710864697714 |
| chr14 | 73715497 | 73715499 | - | 2.00792115374673 | 4.54798296825753 | 1.19322821036799 | 6.40265188186281 |
| chr14 | 73716402 | 73716413 | - | 1.00396057687336 | 4.54798296825753 | 1.19322821036799 | 1.06710864697714 |
| chr14 | 73787348 | 73787378 | + | 2.00792115374673 | 11.3699574206438 | 5.96614105183995 | 2.13421729395427 |
| chr14 | 73884715 | 73884724 | + | 2.00792115374673 | 4.54798296825753 | 1.19322821036799 | 1.06710864697714 |
| chr14 | 73886820 | 73886904 | + | 610.408030739006 | 781.11607479823 | 206.428480393662 | 278.515356861032 |
| chr14 | 73896383 | 73896405 | + | 4.01584230749346 | 3.41098722619315 | 2.38645642073598 | 1.06710864697714 |
| chr14 | 73898125 | 73898127 | + | 1.00396057687336 | 1.13699574206438 | 3.57968463110397 | 2.13421729395427 |
| chr14 | 73923503 | 73923523 | + | 7.02772403811355 | 3.41098722619315 | 1.19322821036799 | 2.13421729395427 |
| chr14 | 73932102 | 73932113 | + | 5.01980288436682 | 10.2329616785794 | 1.19322821036799 | 4.26843458790854 |
| chr14 | 73932190 | 73932192 | + | 5.01980288436682 | 13.6439489047726 | 2.38645642073598 | 1.06710864697714 |
| chr14 | 73965220 | 73965223 | - | 9.03564519186028 | 15.9179403889013 | 5.96614105183995 | 6.40265188186281 |
| chr14 | 73966871 | 73966898 | - | 1.00396057687336 | 5.68497871032191 | 5.96614105183995 | 5.33554323488568 |
| chr14 | 74057777 | 74057786 | - | 1.00396057687336 | 1.13699574206438 | 1.19322821036799 | 2.13421729395427 |
| chr14 | 74199624 | 74199632 | + | 1.00396057687336 | 4.54798296825753 | 1.19322821036799 | 1.06710864697714 |
| chr14 | 74286277 | 74286353 | - | 23.0910932680874 | 28.4248935516096 | 8.35259747257593 | 4.26843458790854 |
| chr14 | 74479959 | 74480010 | - | 23.0910932680874 | 38.657855230189 | 5.96614105183995 | 9.60397782279422 |
| chr14 | 74480039 | 74480174 | - | 108.427742302323 | 93.2336508492793 | 21.4781077866238 | 9.60397782279422 |
| chr14 | 74480255 | 74480291 | - | 18.0712903837206 | 21.6029190992233 | 3.57968463110397 | 3.20132594093141 |
| chr14 | 74494974 | 74494977 | + | 1.00396057687336 | 1.13699574206438 | 2.38645642073598 | 2.13421729395427 |
| chr14 | 74661295 | 74661327 | - | 5.01980288436682 | 5.68497871032191 | 9.54582568294392 | 5.33554323488568 |
| chr14 | 74661373 | 74661408 | - | 10.0396057687336 | 6.82197445238629 | 5.96614105183995 | 5.33554323488568 |
| chr14 | 74661438 | 74661465 | - | 9.03564519186028 | 4.54798296825753 | 21.4781077866238 | 7.46976052883995 |
| chr14 | 74661639 | 74661658 | - | 3.01188173062009 | 1.13699574206438 | 4.77291284147196 | 3.20132594093141 |
| chr14 | 74662111 | 74662131 | - | 2.00792115374673 | 3.41098722619315 | 5.96614105183995 | 2.13421729395427 |
| chr14 | 74662561 | 74662581 | - | 5.01980288436682 | 2.27399148412876 | 9.54582568294392 | 6.40265188186281 |
| chr14 | 74662931 | 74662960 | - | 4.01584230749346 | 5.68497871032191 | 1.19322821036799 | 2.13421729395427 |
| chr14 | 74663216 | 74663227 | - | 11.043566345607 | 7.95897019445067 | 1.19322821036799 | 6.40265188186281 |
| chr14 | 74663263 | 74663273 | - | 8.03168461498692 | 7.95897019445067 | 3.57968463110397 | 5.33554323488568 |
| chr14 | 74735149 | 74735182 | + | 20.0792115374673 | 22.7399148412876 | 4.77291284147196 | 14.9395210576799 |
| chr14 | 74735547 | 74735555 | + | 3.01188173062009 | 2.27399148412876 | 1.19322821036799 | 3.20132594093141 |
| chr14 | 74735756 | 74735759 | + | 5.01980288436682 | 7.95897019445067 | 3.57968463110397 | 2.13421729395427 |
| chr14 | 74736440 | 74736442 | + | 5.01980288436682 | 4.54798296825753 | 1.19322821036799 | 1.06710864697714 |
| chr14 | 74835401 | 74835432 | + | 2.00792115374673 | 4.54798296825753 | 7.15936926220794 | 1.06710864697714 |
| chr14 | 74835774 | 74835816 | + | 8.03168461498692 | 13.6439489047726 | 10.7390538933119 | 10.6710864697714 |
| chr14 | 74902924 | 74902955 | + | 3.01188173062009 | 6.82197445238629 | 4.77291284147196 | 4.26843458790854 |
| chr14 | 74903195 | 74903227 | + | 9.03564519186028 | 5.68497871032191 | 7.15936926220794 | 4.26843458790854 |
| chr14 | 74903278 | 74903298 | + | 7.02772403811355 | 3.41098722619315 | 4.77291284147196 | 6.40265188186281 |
| chr14 | 74903393 | 74903441 | + | 21.0831721143407 | 13.6439489047726 | 21.4781077866238 | 14.9395210576799 |
| chr14 | 74903468 | 74903485 | + | 2.00792115374673 | 2.27399148412876 | 1.19322821036799 | 2.13421729395427 |
| chr14 | 75009396 | 75009408 | + | 3.01188173062009 | 7.95897019445067 | 1.19322821036799 | 1.06710864697714 |
| chr14 | 75009515 | 75009562 | + | 10.0396057687336 | 15.9179403889013 | 9.54582568294392 | 2.13421729395427 |
| chr14 | 75053278 | 75053295 | - | 4.01584230749346 | 10.2329616785794 | 2.38645642073598 | 5.33554323488568 |
| chr14 | 75081588 | 75081592 | - | 5.01980288436682 | 13.6439489047726 | 2.38645642073598 | 4.26843458790854 |
| chr14 | 75082152 | 75082243 | - | 14.0554480762271 | 13.6439489047726 | 2.38645642073598 | 4.26843458790854 |
| chr14 | 75082342 | 75082394 | - | 21.0831721143407 | 38.657855230189 | 7.15936926220794 | 9.60397782279422 |
| chr14 | 75082569 | 75082660 | - | 5.01980288436682 | 6.82197445238629 | 4.77291284147196 | 3.20132594093141 |
| chr14 | 75082830 | 75082860 | - | 2.00792115374673 | 2.27399148412876 | 7.15936926220794 | 1.06710864697714 |
| chr14 | 75131701 | 75131727 | - | 1.00396057687336 | 4.54798296825753 | 1.19322821036799 | 3.20132594093141 |
| chr14 | 75132682 | 75132732 | - | 5.01980288436682 | 3.41098722619315 | 13.1255103140479 | 11.7381951167485 |
| chr14 | 75132825 | 75132856 | - | 1.00396057687336 | 1.13699574206438 | 4.77291284147196 | 8.53686917581709 |
| chr14 | 75132889 | 75132898 | - | 3.01188173062009 | 2.27399148412876 | 1.19322821036799 | 3.20132594093141 |
| chr14 | 75132960 | 75132997 | - | 6.02376346124019 | 3.41098722619315 | 14.3187385244159 | 2.13421729395427 |
| chr14 | 75133076 | 75133139 | - | 2.00792115374673 | 3.41098722619315 | 19.0916513658878 | 11.7381951167485 |
| chr14 | 75133341 | 75133431 | - | 8.03168461498692 | 6.82197445238629 | 7.15936926220794 | 5.33554323488568 |
| chr14 | 75133632 | 75133662 | - | 5.01980288436682 | 4.54798296825753 | 3.57968463110397 | 2.13421729395427 |
| chr14 | 75133744 | 75133760 | - | 1.00396057687336 | 3.41098722619315 | 3.57968463110397 | 1.06710864697714 |
| chr14 | 75133829 | 75133844 | - | 49.1940682667949 | 34.1098722619315 | 36.9900745214077 | 22.4092815865198 |
| chr14 | 75133990 | 75134010 | - | 3.01188173062009 | 1.13699574206438 | 9.54582568294392 | 6.40265188186281 |
| chr14 | 75134227 | 75134356 | - | 75.2970432655023 | 87.5486721389574 | 32.2171616799357 | 27.7448248214055 |
| chr14 | 75134369 | 75134397 | - | 5.01980288436682 | 10.2329616785794 | 5.96614105183995 | 2.13421729395427 |
| chr14 | 75134429 | 75134497 | - | 22.087132691214 | 10.2329616785794 | 15.5119667347839 | 8.53686917581709 |
| chr14 | 75134523 | 75134530 | - | 3.01188173062009 | 4.54798296825753 | 2.38645642073598 | 6.40265188186281 |
| chr14 | 75134559 | 75134586 | - | 53.2099105742883 | 64.8087572976698 | 27.4442488384638 | 22.4092815865198 |
| chr14 | 75134622 | 75134685 | - | 6.02376346124019 | 13.6439489047726 | 11.9322821036799 | 3.20132594093141 |
| chr14 | 75134744 | 75134799 | - | 6.02376346124019 | 6.82197445238629 | 20.2848795762558 | 7.46976052883995 |
| chr14 | 75134879 | 75134906 | - | 1.00396057687336 | 1.13699574206438 | 2.38645642073598 | 1.06710864697714 |
| chr14 | 75282010 | 75282013 | + | 14.0554480762271 | 23.876910583352 | 1.19322821036799 | 5.33554323488568 |
| chr14 | 75469465 | 75469575 | + | 37.1465413443145 | 69.3567402659273 | 20.2848795762558 | 2.13421729395427 |
| chr14 | 75650942 | 75650985 | - | 19.0752509605939 | 13.6439489047726 | 20.2848795762558 | 8.53686917581709 |
| chr14 | 75651024 | 75651063 | - | 3.01188173062009 | 1.13699574206438 | 1.19322821036799 | 2.13421729395427 |
| chr14 | 75651109 | 75651144 | - | 17.0673298068472 | 11.3699574206438 | 3.57968463110397 | 16.006629704657 |
| chr14 | 75651193 | 75651220 | - | 9.03564519186028 | 6.82197445238629 | 9.54582568294392 | 9.60397782279422 |
| chr14 | 75651246 | 75651263 | - | 4.01584230749346 | 4.54798296825753 | 3.57968463110397 | 5.33554323488568 |
| chr14 | 75651491 | 75651563 | - | 11.043566345607 | 13.6439489047726 | 28.6374770488317 | 13.8724124107028 |
| chr14 | 75954500 | 75954530 | + | 27.1069355755808 | 21.6029190992233 | 5.96614105183995 | 6.40265188186281 |
| chr14 | 75954675 | 75954689 | + | 9.03564519186028 | 14.780944646837 | 1.19322821036799 | 3.20132594093141 |
| chr14 | 75954946 | 75954955 | + | 4.01584230749346 | 5.68497871032191 | 1.19322821036799 | 1.06710864697714 |
| chr14 | 76210718 | 76210720 | + | 1.00396057687336 | 1.13699574206438 | 2.38645642073598 | 3.20132594093141 |
| chr14 | 76787439 | 76787447 | - | 4.01584230749346 | 3.41098722619315 | 4.77291284147196 | 1.06710864697714 |
| chr14 | 76787690 | 76787712 | - | 6.02376346124019 | 5.68497871032191 | 7.15936926220794 | 5.33554323488568 |
| chr14 | 77024571 | 77024588 | - | 7.02772403811355 | 15.9179403889013 | 5.96614105183995 | 4.26843458790854 |
| chr14 | 77024749 | 77024811 | - | 145.574283646638 | 183.056314472366 | 48.9223566250876 | 52.2883237018796 |
| chr14 | 77025121 | 77025216 | - | 11.043566345607 | 22.7399148412876 | 28.6374770488317 | 14.9395210576799 |
| chr14 | 77025232 | 77025264 | - | 2.00792115374673 | 1.13699574206438 | 3.57968463110397 | 4.26843458790854 |
| chr14 | 77025291 | 77025305 | - | 1.00396057687336 | 3.41098722619315 | 2.38645642073598 | 3.20132594093141 |
| chr14 | 77114907 | 77114916 | + | 3.01188173062009 | 1.13699574206438 | 5.96614105183995 | 3.20132594093141 |
| chr14 | 77115672 | 77115674 | + | 1.00396057687336 | 1.13699574206438 | 2.38645642073598 | 2.13421729395427 |
| chr14 | 77116614 | 77116619 | + | 4.01584230749346 | 2.27399148412876 | 2.38645642073598 | 3.20132594093141 |
| chr14 | 77116678 | 77116712 | + | 5.01980288436682 | 3.41098722619315 | 1.19322821036799 | 6.40265188186281 |
| chr14 | 77116754 | 77116758 | + | 7.02772403811355 | 11.3699574206438 | 8.35259747257593 | 10.6710864697714 |
| chr14 | 77116793 | 77116810 | + | 21.0831721143407 | 20.4659233571589 | 13.1255103140479 | 30.9461507623369 |
| chr14 | 77116832 | 77116864 | + | 13.0514874993537 | 14.780944646837 | 11.9322821036799 | 9.60397782279422 |
| chr14 | 77116920 | 77116921 | + | 3.01188173062009 | 3.41098722619315 | 1.19322821036799 | 1.06710864697714 |
| chr14 | 77275132 | 77275167 | - | 23.0910932680874 | 15.9179403889013 | 10.7390538933119 | 3.20132594093141 |
| chr14 | 77275195 | 77275230 | - | 11.043566345607 | 15.9179403889013 | 7.15936926220794 | 3.20132594093141 |
| chr14 | 77331232 | 77331292 | + | 21.0831721143407 | 25.0139063254164 | 9.54582568294392 | 17.0737383516342 |
| chr14 | 77331371 | 77331408 | + | 11.043566345607 | 17.0549361309657 | 10.7390538933119 | 6.40265188186281 |
| chr14 | 77335306 | 77335311 | - | 5.01980288436682 | 1.13699574206438 | 8.35259747257593 | 9.60397782279422 |
| chr14 | 77335338 | 77335339 | - | 3.01188173062009 | 2.27399148412876 | 2.38645642073598 | 2.13421729395427 |
| chr14 | 77335396 | 77335419 | - | 151.598047107878 | 271.741982353387 | 171.82486229299 | 211.287512101473 |
| chr14 | 77335454 | 77335474 | - | 1.00396057687336 | 1.13699574206438 | 8.35259747257593 | 1.06710864697714 |
| chr14 | 77335553 | 77335568 | - | 20.0792115374673 | 21.6029190992233 | 8.35259747257593 | 6.40265188186281 |
| chr14 | 77335606 | 77335639 | - | 24.0950538449607 | 34.1098722619315 | 14.3187385244159 | 21.3421729395427 |
| chr14 | 77335670 | 77335692 | - | 1.00396057687336 | 1.13699574206438 | 2.38645642073598 | 1.06710864697714 |
| chr14 | 77335716 | 77335728 | - | 6.02376346124019 | 11.3699574206438 | 8.35259747257593 | 5.33554323488568 |
| chr14 | 77336195 | 77336219 | - | 3.01188173062009 | 6.82197445238629 | 3.57968463110397 | 3.20132594093141 |
| chr14 | 77336625 | 77336631 | - | 2.00792115374673 | 3.41098722619315 | 7.15936926220794 | 3.20132594093141 |
| chr14 | 77336734 | 77336737 | - | 9.03564519186028 | 6.82197445238629 | 3.57968463110397 | 1.06710864697714 |
| chr14 | 77336825 | 77336832 | - | 2.00792115374673 | 2.27399148412876 | 3.57968463110397 | 5.33554323488568 |
| chr14 | 77337145 | 77337182 | - | 3.01188173062009 | 2.27399148412876 | 11.9322821036799 | 17.0737383516342 |
| chr14 | 77337258 | 77337264 | - | 4.01584230749346 | 5.68497871032191 | 7.15936926220794 | 2.13421729395427 |
| chr14 | 77339159 | 77339161 | - | 6.02376346124019 | 12.5069531627082 | 8.35259747257593 | 7.46976052883995 |
| chr14 | 77340504 | 77340516 | - | 6.02376346124019 | 2.27399148412876 | 3.57968463110397 | 6.40265188186281 |
| chr14 | 77340953 | 77340967 | - | 9.03564519186028 | 21.6029190992233 | 19.0916513658878 | 16.006629704657 |
| chr14 | 77341000 | 77341027 | - | 14.0554480762271 | 21.6029190992233 | 19.0916513658878 | 30.9461507623369 |
| chr14 | 77341185 | 77341211 | - | 16.0633692299738 | 18.1919318730301 | 4.77291284147196 | 4.26843458790854 |
| chr14 | 77341325 | 77341345 | - | 2.00792115374673 | 7.95897019445067 | 2.38645642073598 | 1.06710864697714 |
| chr14 | 77341477 | 77341482 | - | 3.01188173062009 | 4.54798296825753 | 4.77291284147196 | 2.13421729395427 |
| chr14 | 77341588 | 77341594 | - | 6.02376346124019 | 5.68497871032191 | 2.38645642073598 | 2.13421729395427 |
| chr14 | 77427339 | 77427352 | - | 2.00792115374673 | 1.13699574206438 | 3.57968463110397 | 1.06710864697714 |
| chr14 | 77469273 | 77469447 | + | 727.871418233189 | 235.358118607327 | 195.68942650035 | 374.555135088975 |
| chr14 | 77506038 | 77506074 | - | 6.02376346124019 | 7.95897019445067 | 4.77291284147196 | 4.26843458790854 |
| chr14 | 77506261 | 77506265 | - | 4.01584230749346 | 4.54798296825753 | 3.57968463110397 | 5.33554323488568 |
| chr14 | 77506328 | 77506330 | - | 4.01584230749346 | 1.13699574206438 | 5.96614105183995 | 4.26843458790854 |
| chr14 | 77510395 | 77510410 | - | 6.02376346124019 | 7.95897019445067 | 1.19322821036799 | 6.40265188186281 |
| chr14 | 77511095 | 77511107 | - | 35.1386201905678 | 40.9318467143178 | 8.35259747257593 | 20.2750642925656 |
| chr14 | 77512061 | 77512068 | - | 3.01188173062009 | 4.54798296825753 | 4.77291284147196 | 7.46976052883995 |
| chr14 | 77512153 | 77512163 | - | 4.01584230749346 | 1.13699574206438 | 3.57968463110397 | 2.13421729395427 |
| chr14 | 77717478 | 77717575 | + | 265.045592294568 | 253.550050480357 | 134.834787771583 | 295.589095212667 |
| chr14 | 77717634 | 77717660 | - | 7.02772403811355 | 18.1919318730301 | 13.1255103140479 | 18.1408469986113 |
| chr14 | 77717712 | 77717736 | - | 6.02376346124019 | 4.54798296825753 | 7.15936926220794 | 7.46976052883995 |
| chr14 | 77717938 | 77717942 | - | 2.00792115374673 | 4.54798296825753 | 1.19322821036799 | 7.46976052883995 |
| chr14 | 77933723 | 77933741 | + | 3.01188173062009 | 5.68497871032191 | 2.38645642073598 | 4.26843458790854 |
| chr14 | 77933843 | 77933864 | + | 47.1861471130481 | 32.9728765198671 | 22.6713359969918 | 49.0869977609482 |
| chr14 | 81176268 | 81176291 | - | 3.01188173062009 | 2.27399148412876 | 1.19322821036799 | 2.13421729395427 |
| chr14 | 81176387 | 81176411 | - | 459.813944208001 | 532.114007286131 | 207.62170860403 | 353.212962149432 |
| chr14 | 81176452 | 81176482 | - | 3.01188173062009 | 2.27399148412876 | 1.19322821036799 | 1.06710864697714 |
| chr14 | 81176846 | 81176881 | - | 1.00396057687336 | 1.13699574206438 | 2.38645642073598 | 1.06710864697714 |
| chr14 | 81176902 | 81176917 | - | 12.0475269224804 | 12.5069531627082 | 9.54582568294392 | 1.06710864697714 |
| chr14 | 81177156 | 81177173 | - | 1.00396057687336 | 2.27399148412876 | 2.38645642073598 | 2.13421729395427 |
| chr14 | 81177307 | 81177329 | - | 10.0396057687336 | 17.0549361309657 | 5.96614105183995 | 4.26843458790854 |
| chr14 | 81178331 | 81178371 | - | 7.02772403811355 | 4.54798296825753 | 5.96614105183995 | 1.06710864697714 |
| chr14 | 81178429 | 81178438 | - | 4.01584230749346 | 5.68497871032191 | 2.38645642073598 | 1.06710864697714 |
| chr14 | 81178526 | 81178554 | - | 7.02772403811355 | 3.41098722619315 | 5.96614105183995 | 5.33554323488568 |
| chr14 | 81178654 | 81178680 | - | 2.00792115374673 | 2.27399148412876 | 8.35259747257593 | 13.8724124107028 |
| chr14 | 81178778 | 81178792 | - | 5.01980288436682 | 2.27399148412876 | 1.19322821036799 | 3.20132594093141 |
| chr14 | 81178860 | 81178891 | - | 4.01584230749346 | 10.2329616785794 | 4.77291284147196 | 7.46976052883995 |
| chr14 | 81178925 | 81178944 | - | 3.01188173062009 | 4.54798296825753 | 1.19322821036799 | 2.13421729395427 |
| chr14 | 81179095 | 81179126 | - | 14.0554480762271 | 15.9179403889013 | 17.8984231555198 | 14.9395210576799 |
| chr14 | 81179241 | 81179258 | - | 4.01584230749346 | 6.82197445238629 | 5.96614105183995 | 3.20132594093141 |
| chr14 | 81179993 | 81180006 | - | 37.1465413443145 | 64.8087572976698 | 28.6374770488317 | 10.6710864697714 |
| chr14 | 81180219 | 81180237 | - | 8.03168461498692 | 10.2329616785794 | 4.77291284147196 | 4.26843458790854 |
| chr14 | 81260738 | 81260740 | - | 2.00792115374673 | 2.27399148412876 | 2.38645642073598 | 4.26843458790854 |
| chr14 | 81473025 | 81473101 | - | 3.01188173062009 | 3.41098722619315 | 4.77291284147196 | 6.40265188186281 |
| chr14 | 81476379 | 81476388 | - | 2.00792115374673 | 3.41098722619315 | 1.19322821036799 | 1.06710864697714 |
| chr14 | 85636378 | 85636380 | + | 3.01188173062009 | 3.41098722619315 | 5.96614105183995 | 3.20132594093141 |
| chr14 | 88611767 | 88611807 | + | 8.03168461498692 | 13.6439489047726 | 9.54582568294392 | 7.46976052883995 |
| chr14 | 88611843 | 88611867 | + | 4.01584230749346 | 4.54798296825753 | 2.38645642073598 | 1.06710864697714 |
| chr14 | 88611925 | 88611935 | + | 2.00792115374673 | 1.13699574206438 | 3.57968463110397 | 3.20132594093141 |
| chr14 | 88612311 | 88612319 | + | 2.00792115374673 | 2.27399148412876 | 3.57968463110397 | 2.13421729395427 |
| chr14 | 88612361 | 88612437 | + | 29.1148567293276 | 29.5618892936739 | 9.54582568294392 | 16.006629704657 |
| chr14 | 88612962 | 88612963 | + | 2.00792115374673 | 2.27399148412876 | 1.19322821036799 | 1.06710864697714 |
| chr14 | 88877860 | 88877886 | + | 4.01584230749346 | 9.09596593651506 | 7.15936926220794 | 5.33554323488568 |
| chr14 | 89160212 | 89160226 | - | 12.0475269224804 | 9.09596593651506 | 4.77291284147196 | 3.20132594093141 |
| chr14 | 89797155 | 89797164 | - | 2.00792115374673 | 3.41098722619315 | 5.96614105183995 | 1.06710864697714 |
| chr14 | 89797210 | 89797214 | - | 3.01188173062009 | 1.13699574206438 | 9.54582568294392 | 2.13421729395427 |
| chr14 | 90044620 | 90044630 | + | 10.0396057687336 | 7.95897019445067 | 1.19322821036799 | 11.7381951167485 |
| chr14 | 90404701 | 90404719 | + | 25.0990144218341 | 7.95897019445067 | 7.15936926220794 | 18.1408469986113 |
| chr14 | 90404750 | 90404751 | + | 2.00792115374673 | 2.27399148412876 | 2.38645642073598 | 3.20132594093141 |
| chr14 | 90404893 | 90404910 | + | 11.043566345607 | 10.2329616785794 | 3.57968463110397 | 4.26843458790854 |
| chr14 | 90405012 | 90405017 | + | 4.01584230749346 | 4.54798296825753 | 1.19322821036799 | 4.26843458790854 |
| chr14 | 90405039 | 90405063 | + | 63.249516343022 | 43.2058381984465 | 11.9322821036799 | 13.8724124107028 |
| chr14 | 90405196 | 90405251 | + | 8.03168461498692 | 2.27399148412876 | 4.77291284147196 | 4.26843458790854 |
| chr14 | 90405289 | 90405312 | + | 6.02376346124019 | 4.54798296825753 | 2.38645642073598 | 2.13421729395427 |
| chr14 | 90405397 | 90405431 | + | 10.0396057687336 | 5.68497871032191 | 4.77291284147196 | 13.8724124107028 |
| chr14 | 90405454 | 90405492 | + | 36.1425807674411 | 29.5618892936739 | 19.0916513658878 | 19.2079556455884 |
| chr14 | 90405537 | 90405566 | + | 34.1346596136944 | 36.3838637460602 | 17.8984231555198 | 6.40265188186281 |
| chr14 | 90405826 | 90405840 | + | 29.1148567293276 | 15.9179403889013 | 17.8984231555198 | 9.60397782279422 |
| chr14 | 90406352 | 90406357 | + | 1.00396057687336 | 1.13699574206438 | 2.38645642073598 | 4.26843458790854 |
| chr14 | 90406625 | 90406650 | + | 11.043566345607 | 3.41098722619315 | 1.19322821036799 | 1.06710864697714 |
| chr14 | 90406690 | 90406696 | + | 3.01188173062009 | 2.27399148412876 | 2.38645642073598 | 1.06710864697714 |
| chr14 | 90407176 | 90407200 | + | 8.03168461498692 | 3.41098722619315 | 3.57968463110397 | 2.13421729395427 |
| chr14 | 90407484 | 90407505 | + | 10.0396057687336 | 3.41098722619315 | 2.38645642073598 | 1.06710864697714 |
| chr14 | 90408179 | 90408183 | + | 11.043566345607 | 9.09596593651506 | 4.77291284147196 | 5.33554323488568 |
| chr14 | 91271390 | 91271394 | - | 5.01980288436682 | 11.3699574206438 | 5.96614105183995 | 5.33554323488568 |
| chr14 | 91271509 | 91271513 | - | 16.0633692299738 | 27.2878978095452 | 11.9322821036799 | 16.006629704657 |
| chr14 | 91271733 | 91271736 | - | 4.01584230749346 | 4.54798296825753 | 10.7390538933119 | 1.06710864697714 |
| chr14 | 91457666 | 91457674 | - | 10.0396057687336 | 14.780944646837 | 3.57968463110397 | 4.26843458790854 |
| chr14 | 91457695 | 91457705 | - | 3.01188173062009 | 1.13699574206438 | 1.19322821036799 | 3.20132594093141 |
| chr14 | 91458037 | 91458069 | - | 6.02376346124019 | 4.54798296825753 | 1.19322821036799 | 2.13421729395427 |
| chr14 | 91458122 | 91458152 | - | 24.0950538449607 | 20.4659233571589 | 7.15936926220794 | 7.46976052883995 |
| chr14 | 91458250 | 91458265 | - | 32.1267384599477 | 55.7127913611547 | 5.96614105183995 | 14.9395210576799 |
| chr14 | 91458576 | 91458609 | - | 7.02772403811355 | 9.09596593651506 | 4.77291284147196 | 5.33554323488568 |
| chr14 | 92038723 | 92038733 | - | 6.02376346124019 | 5.68497871032191 | 5.96614105183995 | 5.33554323488568 |
| chr14 | 92039073 | 92039074 | - | 2.00792115374673 | 4.54798296825753 | 1.19322821036799 | 4.26843458790854 |
| chr14 | 92039222 | 92039225 | - | 6.02376346124019 | 18.1919318730301 | 16.7051949451519 | 8.53686917581709 |
| chr14 | 92039356 | 92039365 | - | 6.02376346124019 | 2.27399148412876 | 9.54582568294392 | 5.33554323488568 |
| chr14 | 92040034 | 92040074 | - | 115.455466340437 | 105.740604011988 | 90.6853439879672 | 98.1739955218965 |
| chr14 | 92040857 | 92040875 | - | 1.00396057687336 | 2.27399148412876 | 3.57968463110397 | 1.06710864697714 |
| chr14 | 92042654 | 92042655 | - | 1.00396057687336 | 1.13699574206438 | 1.19322821036799 | 3.20132594093141 |
| chr14 | 92061517 | 92061535 | - | 7.02772403811355 | 2.27399148412876 | 1.19322821036799 | 2.13421729395427 |
| chr14 | 92116148 | 92116156 | - | 36.1425807674411 | 45.4798296825753 | 14.3187385244159 | 12.8053037637256 |
| chr14 | 92116180 | 92116232 | - | 124.491111532297 | 130.754510337404 | 34.6036181006717 | 41.6172372321083 |
| chr14 | 92162295 | 92162312 | + | 3.01188173062009 | 4.54798296825753 | 8.35259747257593 | 5.33554323488568 |
| chr14 | 92164007 | 92164028 | + | 16.0633692299738 | 12.5069531627082 | 4.77291284147196 | 27.7448248214055 |
| chr14 | 92165475 | 92165479 | + | 1.00396057687336 | 3.41098722619315 | 1.19322821036799 | 3.20132594093141 |
| chr14 | 92839695 | 92839770 | + | 7.02772403811355 | 6.82197445238629 | 1.19322821036799 | 1.06710864697714 |
| chr14 | 95086523 | 95086526 | - | 2.00792115374673 | 3.41098722619315 | 1.19322821036799 | 3.20132594093141 |
| chr14 | 95086682 | 95086725 | - | 15.0594086531005 | 31.8358807778027 | 10.7390538933119 | 7.46976052883995 |
| chr14 | 95086755 | 95086758 | - | 1.00396057687336 | 1.13699574206438 | 1.19322821036799 | 1.06710864697714 |
| chr14 | 95086981 | 95087029 | - | 12.0475269224804 | 12.5069531627082 | 1.19322821036799 | 2.13421729395427 |
| chr14 | 95087064 | 95087126 | - | 6.02376346124019 | 2.27399148412876 | 1.19322821036799 | 2.13421729395427 |
| chr14 | 95087657 | 95087669 | - | 1.00396057687336 | 3.41098722619315 | 1.19322821036799 | 1.06710864697714 |
| chr14 | 95087762 | 95087828 | - | 17.0673298068472 | 14.780944646837 | 8.35259747257593 | 11.7381951167485 |
| chr14 | 95088718 | 95088730 | - | 41.1623836518079 | 56.8497871032191 | 13.1255103140479 | 20.2750642925656 |
| chr14 | 95089927 | 95089945 | - | 4.01584230749346 | 4.54798296825753 | 4.77291284147196 | 2.13421729395427 |
| chr14 | 95090359 | 95090450 | - | 19.0752509605939 | 20.4659233571589 | 17.8984231555198 | 9.60397782279422 |
| chr14 | 95090597 | 95090655 | - | 21.0831721143407 | 13.6439489047726 | 14.3187385244159 | 9.60397782279422 |
| chr14 | 95157674 | 95157735 | + | 21.0831721143407 | 29.5618892936739 | 14.3187385244159 | 3.20132594093141 |
| chr14 | 95544194 | 95544336 | + | 7.02772403811355 | 13.6439489047726 | 5.96614105183995 | 11.7381951167485 |
| chr14 | 95544460 | 95544489 | + | 2.00792115374673 | 1.13699574206438 | 1.19322821036799 | 3.20132594093141 |
| chr14 | 95544586 | 95544695 | + | 22.087132691214 | 28.4248935516096 | 27.4442488384638 | 69.3620620535138 |
| chr14 | 95620912 | 95620979 | + | 85.336649034236 | 136.439489047726 | 17.8984231555198 | 16.006629704657 |
| chr14 | 95679204 | 95679207 | + | 2.00792115374673 | 9.09596593651506 | 2.38645642073598 | 1.06710864697714 |
| chr14 | 96281517 | 96281524 | - | 5.01980288436682 | 1.13699574206438 | 2.38645642073598 | 2.13421729395427 |
| chr14 | 96385868 | 96385883 | + | 2.00792115374673 | 1.13699574206438 | 1.19322821036799 | 1.06710864697714 |
| chr14 | 96386671 | 96386687 | + | 10.0396057687336 | 6.82197445238629 | 4.77291284147196 | 2.13421729395427 |
| chr14 | 96386716 | 96386739 | + | 4.01584230749346 | 4.54798296825753 | 8.35259747257593 | 2.13421729395427 |
| chr14 | 96565126 | 96565149 | + | 3.01188173062009 | 1.13699574206438 | 3.57968463110397 | 14.9395210576799 |
| chr14 | 96565564 | 96565567 | + | 6.02376346124019 | 3.41098722619315 | 1.19322821036799 | 1.06710864697714 |
| chr14 | 96565624 | 96565644 | + | 5.01980288436682 | 5.68497871032191 | 1.19322821036799 | 1.06710864697714 |
| chr14 | 96566600 | 96566615 | + | 1.00396057687336 | 1.13699574206438 | 1.19322821036799 | 1.06710864697714 |
| chr14 | 96566802 | 96566806 | + | 4.01584230749346 | 3.41098722619315 | 3.57968463110397 | 7.46976052883995 |
| chr14 | 96566835 | 96566842 | + | 3.01188173062009 | 6.82197445238629 | 3.57968463110397 | 6.40265188186281 |
| chr14 | 96566877 | 96566922 | + | 2.00792115374673 | 3.41098722619315 | 3.57968463110397 | 4.26843458790854 |
| chr14 | 96566955 | 96566991 | + | 6.02376346124019 | 10.2329616785794 | 1.19322821036799 | 4.26843458790854 |
| chr14 | 96881188 | 96881224 | + | 4.01584230749346 | 6.82197445238629 | 35.7968463110397 | 39.483019938154 |
| chr14 | 96881274 | 96881275 | + | 1.00396057687336 | 2.27399148412876 | 1.19322821036799 | 1.06710864697714 |
| chr14 | 96881303 | 96881313 | + | 10.0396057687336 | 18.1919318730301 | 5.96614105183995 | 21.3421729395427 |
| chr14 | 96881385 | 96881387 | + | 1.00396057687336 | 3.41098722619315 | 4.77291284147196 | 4.26843458790854 |
| chr14 | 96881560 | 96881572 | + | 2.00792115374673 | 1.13699574206438 | 4.77291284147196 | 4.26843458790854 |
| chr14 | 99397839 | 99397862 | - | 6.02376346124019 | 9.09596593651506 | 3.57968463110397 | 8.53686917581709 |
| chr14 | 99398133 | 99398148 | - | 6.02376346124019 | 11.3699574206438 | 3.57968463110397 | 5.33554323488568 |
| chr14 | 99409930 | 99409935 | - | 3.01188173062009 | 2.27399148412876 | 1.19322821036799 | 1.06710864697714 |
| chr14 | 99502780 | 99502816 | - | 13.0514874993537 | 10.2329616785794 | 7.15936926220794 | 13.8724124107028 |
| chr14 | 99502861 | 99502882 | - | 43.1703048055547 | 29.5618892936739 | 16.7051949451519 | 24.5434988804741 |
| chr14 | 99502939 | 99502982 | - | 13.0514874993537 | 11.3699574206438 | 2.38645642073598 | 9.60397782279422 |
| chr14 | 99507093 | 99507145 | - | 22.087132691214 | 18.1919318730301 | 9.54582568294392 | 26.6777161744284 |
| chr14 | 99510166 | 99510176 | - | 15.0594086531005 | 37.5208594881246 | 48.9223566250876 | 28.8119334683827 |
| chr14 | 99510243 | 99510254 | - | 2.00792115374673 | 3.41098722619315 | 2.38645642073598 | 1.06710864697714 |
| chr14 | 99510752 | 99510786 | + | 7.02772403811355 | 10.2329616785794 | 16.7051949451519 | 11.7381951167485 |
| chr14 | 99511238 | 99511247 | + | 2.00792115374673 | 1.13699574206438 | 3.57968463110397 | 7.46976052883995 |
| chr14 | 99511302 | 99511314 | + | 26.1029749987075 | 6.82197445238629 | 4.77291284147196 | 21.3421729395427 |
| chr14 | 99511341 | 99511351 | + | 10.0396057687336 | 30.6988850357383 | 13.1255103140479 | 32.0132594093141 |
| chr14 | 99512496 | 99512516 | - | 4.01584230749346 | 1.13699574206438 | 4.77291284147196 | 2.13421729395427 |
| chr14 | 99513873 | 99513888 | - | 6.02376346124019 | 2.27399148412876 | 1.19322821036799 | 1.06710864697714 |
| chr14 | 100144141 | 100144199 | + | 2.00792115374673 | 1.13699574206438 | 1.19322821036799 | 2.13421729395427 |
| chr14 | 100277727 | 100277729 | + | 3.01188173062009 | 4.54798296825753 | 1.19322821036799 | 1.06710864697714 |
| chr14 | 100277857 | 100277863 | + | 7.02772403811355 | 19.3289276150945 | 1.19322821036799 | 9.60397782279422 |
| chr14 | 100278001 | 100278075 | + | 30.1188173062009 | 32.9728765198671 | 4.77291284147196 | 8.53686917581709 |
| chr14 | 100278374 | 100278386 | + | 1.00396057687336 | 2.27399148412876 | 3.57968463110397 | 4.26843458790854 |
| chr14 | 100278435 | 100278443 | + | 3.01188173062009 | 3.41098722619315 | 9.54582568294392 | 6.40265188186281 |
| chr14 | 100279046 | 100279065 | + | 3.01188173062009 | 5.68497871032191 | 4.77291284147196 | 1.06710864697714 |
| chr14 | 100280454 | 100280465 | + | 19.0752509605939 | 13.6439489047726 | 5.96614105183995 | 10.6710864697714 |
| chr14 | 100280543 | 100280554 | + | 2.00792115374673 | 2.27399148412876 | 1.19322821036799 | 1.06710864697714 |
| chr14 | 100280764 | 100280769 | + | 7.02772403811355 | 4.54798296825753 | 3.57968463110397 | 2.13421729395427 |
| chr14 | 100280807 | 100280810 | + | 4.01584230749346 | 1.13699574206438 | 1.19322821036799 | 1.06710864697714 |
| chr14 | 100281191 | 100281195 | + | 4.01584230749346 | 2.27399148412876 | 1.19322821036799 | 1.06710864697714 |
| chr14 | 100281912 | 100281981 | + | 10.0396057687336 | 11.3699574206438 | 25.0577924177278 | 8.53686917581709 |
| chr14 | 100282019 | 100282033 | + | 4.01584230749346 | 15.9179403889013 | 3.57968463110397 | 1.06710864697714 |
| chr14 | 100282100 | 100282111 | + | 7.02772403811355 | 2.27399148412876 | 2.38645642073598 | 8.53686917581709 |
| chr14 | 100291138 | 100291149 | - | 1.00396057687336 | 3.41098722619315 | 2.38645642073598 | 2.13421729395427 |
| chr14 | 100291208 | 100291243 | - | 3.01188173062009 | 2.27399148412876 | 1.19322821036799 | 1.06710864697714 |
| chr14 | 100333798 | 100333809 | + | 3.01188173062009 | 18.1919318730301 | 8.35259747257593 | 3.20132594093141 |
| chr14 | 100333810 | 100333902 | - | 16.0633692299738 | 42.0688424563821 | 33.4103898903037 | 33.0803680562912 |
| chr14 | 100334511 | 100334547 | - | 2.00792115374673 | 4.54798296825753 | 8.35259747257593 | 2.13421729395427 |
| chr14 | 101766210 | 101766330 | + | 17.0673298068472 | 5.68497871032191 | 107.390538933119 | 102.442430109805 |
| chr14 | 101766352 | 101766451 | + | 9.03564519186028 | 3.41098722619315 | 48.9223566250876 | 26.6777161744284 |
| chr14 | 101766481 | 101766508 | + | 10.0396057687336 | 5.68497871032191 | 103.810854302015 | 144.059667341913 |
| chr14 | 101766925 | 101766927 | + | 1.00396057687336 | 1.13699574206438 | 3.57968463110397 | 3.20132594093141 |
| chr14 | 101912554 | 101912562 | + | 30.1188173062009 | 36.3838637460602 | 50.1155848354556 | 85.3686917581709 |
| chr14 | 101926254 | 101926257 | + | 2.00792115374673 | 1.13699574206438 | 1.19322821036799 | 3.20132594093141 |
| chr14 | 101927169 | 101927181 | + | 3.01188173062009 | 3.41098722619315 | 2.38645642073598 | 6.40265188186281 |
| chr14 | 101927221 | 101927272 | + | 1.00396057687336 | 1.13699574206438 | 1.19322821036799 | 2.13421729395427 |
| chr14 | 101948318 | 101948378 | + | 10.0396057687336 | 26.1509020674808 | 17.8984231555198 | 35.2145853502455 |
| chr14 | 101949251 | 101949259 | + | 1.00396057687336 | 1.13699574206438 | 2.38645642073598 | 5.33554323488568 |
| chr14 | 102050532 | 102050774 | + | 106.419821148577 | 95.5076423334081 | 44.1494437836156 | 86.435800405148 |
| chr14 | 102081012 | 102081021 | - | 2.00792115374673 | 3.41098722619315 | 5.96614105183995 | 5.33554323488568 |
| chr14 | 102081097 | 102081108 | - | 11.043566345607 | 5.68497871032191 | 21.4781077866238 | 14.9395210576799 |
| chr14 | 102081154 | 102081195 | - | 393.552546134359 | 226.262152670812 | 420.016330049532 | 577.30577801463 |
| chr14 | 102081347 | 102081426 | - | 74.293082688629 | 32.9728765198671 | 68.0140079909754 | 70.429170700491 |
| chr14 | 102081459 | 102081529 | - | 219.867366335267 | 108.014595496116 | 120.516049247167 | 237.965228275901 |
| chr14 | 102081600 | 102081605 | - | 5.01980288436682 | 1.13699574206438 | 1.19322821036799 | 8.53686917581709 |
| chr14 | 102081654 | 102081787 | - | 238.942617295861 | 152.357429436627 | 230.293044601022 | 357.48139673734 |
| chr14 | 102228949 | 102228993 | - | 6.02376346124019 | 11.3699574206438 | 1.19322821036799 | 3.20132594093141 |
| chr14 | 102229030 | 102229035 | - | 2.00792115374673 | 6.82197445238629 | 2.38645642073598 | 1.06710864697714 |
| chr14 | 102229097 | 102229098 | - | 2.00792115374673 | 2.27399148412876 | 1.19322821036799 | 1.06710864697714 |
| chr14 | 102342217 | 102342246 | + | 3.01188173062009 | 10.2329616785794 | 3.57968463110397 | 2.13421729395427 |
| chr14 | 102348309 | 102348338 | - | 9.03564519186028 | 3.41098722619315 | 3.57968463110397 | 1.06710864697714 |
| chr14 | 102506873 | 102506942 | - | 5.01980288436682 | 6.82197445238629 | 4.77291284147196 | 8.53686917581709 |
| chr14 | 102726933 | 102726944 | + | 2.00792115374673 | 3.41098722619315 | 1.19322821036799 | 2.13421729395427 |
| chr14 | 102727383 | 102727402 | + | 1.00396057687336 | 1.13699574206438 | 5.96614105183995 | 4.26843458790854 |
| chr14 | 102727465 | 102727472 | + | 5.01980288436682 | 1.13699574206438 | 10.7390538933119 | 9.60397782279422 |
| chr14 | 102727513 | 102727535 | + | 5.01980288436682 | 5.68497871032191 | 5.96614105183995 | 5.33554323488568 |
| chr14 | 102727571 | 102727590 | + | 2.00792115374673 | 1.13699574206438 | 3.57968463110397 | 5.33554323488568 |
| chr14 | 102727753 | 102727769 | + | 3.01188173062009 | 3.41098722619315 | 7.15936926220794 | 3.20132594093141 |
| chr14 | 102727847 | 102727851 | + | 9.03564519186028 | 3.41098722619315 | 9.54582568294392 | 2.13421729395427 |
| chr14 | 102727894 | 102727946 | + | 14.0554480762271 | 3.41098722619315 | 9.54582568294392 | 24.5434988804741 |
| chr14 | 102728095 | 102728107 | + | 3.01188173062009 | 2.27399148412876 | 4.77291284147196 | 3.20132594093141 |
| chr14 | 102728135 | 102728146 | + | 19.0752509605939 | 9.09596593651506 | 35.7968463110397 | 11.7381951167485 |
| chr14 | 102729020 | 102729029 | + | 4.01584230749346 | 2.27399148412876 | 2.38645642073598 | 1.06710864697714 |
| chr14 | 102730156 | 102730179 | + | 21.0831721143407 | 29.5618892936739 | 14.3187385244159 | 50.1541064079254 |
| chr14 | 102730250 | 102730278 | + | 1.00396057687336 | 1.13699574206438 | 1.19322821036799 | 3.20132594093141 |
| chr14 | 102730337 | 102730348 | + | 1.00396057687336 | 1.13699574206438 | 3.57968463110397 | 2.13421729395427 |
| chr14 | 102906068 | 102906071 | + | 1.00396057687336 | 1.13699574206438 | 1.19322821036799 | 2.13421729395427 |
| chr14 | 102908650 | 102908651 | + | 4.01584230749346 | 1.13699574206438 | 2.38645642073598 | 1.06710864697714 |
| chr14 | 102932495 | 102932510 | - | 39.1544624980612 | 47.753821166704 | 4.77291284147196 | 7.46976052883995 |
| chr14 | 102933223 | 102933235 | - | 8.03168461498692 | 6.82197445238629 | 3.57968463110397 | 1.06710864697714 |
| chr14 | 103137273 | 103137328 | + | 20.0792115374673 | 37.5208594881246 | 5.96614105183995 | 9.60397782279422 |
| chr14 | 103341051 | 103341063 | + | 7.02772403811355 | 12.5069531627082 | 4.77291284147196 | 8.53686917581709 |
| chr14 | 103341188 | 103341196 | + | 3.01188173062009 | 9.09596593651506 | 4.77291284147196 | 9.60397782279422 |
| chr14 | 103341687 | 103341693 | + | 7.02772403811355 | 9.09596593651506 | 2.38645642073598 | 4.26843458790854 |
| chr14 | 103341853 | 103341855 | + | 1.00396057687336 | 2.27399148412876 | 5.96614105183995 | 4.26843458790854 |
| chr14 | 103342463 | 103342483 | + | 4.01584230749346 | 1.13699574206438 | 3.57968463110397 | 2.13421729395427 |
| chr14 | 103342718 | 103342757 | + | 19.0752509605939 | 14.780944646837 | 1.19322821036799 | 18.1408469986113 |
| chr14 | 103342907 | 103342974 | + | 30.1188173062009 | 61.3977700714766 | 10.7390538933119 | 11.7381951167485 |
| chr14 | 103343083 | 103343086 | + | 5.01980288436682 | 9.09596593651506 | 4.77291284147196 | 1.06710864697714 |
| chr14 | 103343108 | 103343124 | + | 3.01188173062009 | 5.68497871032191 | 3.57968463110397 | 1.06710864697714 |
| chr14 | 103503624 | 103503627 | + | 5.01980288436682 | 29.5618892936739 | 1.19322821036799 | 9.60397782279422 |
| chr14 | 103519679 | 103520045 | - | 1269.00616916793 | 781.11607479823 | 987.992958184695 | 1362.6977421898 |
| chr14 | 103535972 | 103535998 | + | 3.01188173062009 | 3.41098722619315 | 1.19322821036799 | 3.20132594093141 |
| chr14 | 103557273 | 103557275 | - | 9.03564519186028 | 15.9179403889013 | 4.77291284147196 | 9.60397782279422 |
| chr14 | 103557365 | 103557390 | - | 1.00396057687336 | 1.13699574206438 | 1.19322821036799 | 1.06710864697714 |
| chr14 | 103559311 | 103559318 | - | 40.1584230749346 | 56.8497871032191 | 10.7390538933119 | 18.1408469986113 |
| chr14 | 103559438 | 103559445 | - | 13.0514874993537 | 15.9179403889013 | 4.77291284147196 | 5.33554323488568 |
| chr14 | 103559520 | 103559524 | - | 3.01188173062009 | 1.13699574206438 | 1.19322821036799 | 1.06710864697714 |
| chr14 | 103559591 | 103559649 | - | 25.0990144218341 | 23.876910583352 | 16.7051949451519 | 22.4092815865198 |
| chr14 | 103559731 | 103559782 | - | 5.01980288436682 | 10.2329616785794 | 9.54582568294392 | 7.46976052883995 |
| chr14 | 103559813 | 103559836 | - | 4.01584230749346 | 4.54798296825753 | 3.57968463110397 | 10.6710864697714 |
| chr14 | 103590217 | 103590238 | + | 4.01584230749346 | 1.13699574206438 | 5.96614105183995 | 12.8053037637256 |
| chr14 | 103590307 | 103590349 | + | 7.02772403811355 | 4.54798296825753 | 23.8645642073598 | 5.33554323488568 |
| chr14 | 103685721 | 103685748 | + | 5.01980288436682 | 3.41098722619315 | 2.38645642073598 | 3.20132594093141 |
| chr14 | 103686311 | 103686334 | + | 1.00396057687336 | 2.27399148412876 | 2.38645642073598 | 2.13421729395427 |
| chr14 | 103696997 | 103697022 | + | 7.02772403811355 | 4.54798296825753 | 3.57968463110397 | 5.33554323488568 |
| chr14 | 103701437 | 103701456 | + | 3.01188173062009 | 2.27399148412876 | 2.38645642073598 | 5.33554323488568 |
| chr14 | 103701484 | 103701507 | + | 9.03564519186028 | 12.5069531627082 | 3.57968463110397 | 3.20132594093141 |
| chr14 | 103733553 | 103733566 | + | 29.1148567293276 | 59.1237785873479 | 7.15936926220794 | 11.7381951167485 |
| chr14 | 103733602 | 103733621 | + | 1.00396057687336 | 1.13699574206438 | 2.38645642073598 | 3.20132594093141 |
| chr14 | 103912333 | 103912682 | - | 390.540664403739 | 309.262841841512 | 164.665493030783 | 314.797050858255 |
| chr14 | 103912934 | 103912939 | - | 9.03564519186028 | 2.27399148412876 | 2.38645642073598 | 5.33554323488568 |
| chr14 | 103913960 | 103913984 | - | 15.0594086531005 | 5.68497871032191 | 3.57968463110397 | 4.26843458790854 |
| chr14 | 104704610 | 104704614 | + | 1.00396057687336 | 1.13699574206438 | 1.19322821036799 | 5.33554323488568 |
| chr14 | 104704754 | 104704781 | + | 11.043566345607 | 13.6439489047726 | 15.5119667347839 | 9.60397782279422 |
| chr14 | 104719533 | 104719586 | + | 24.0950538449607 | 20.4659233571589 | 2.38645642073598 | 2.13421729395427 |
| chr14 | 104759512 | 104759636 | + | 204.807957682166 | 319.495803520091 | 93.0718004087032 | 119.516168461439 |
| chr14 | 104769374 | 104769397 | - | 9.03564519186028 | 18.1919318730301 | 7.15936926220794 | 10.6710864697714 |
| chr14 | 104769454 | 104769542 | - | 13.0514874993537 | 6.82197445238629 | 9.54582568294392 | 10.6710864697714 |
| chr14 | 104769703 | 104769765 | - | 1.00396057687336 | 1.13699574206438 | 3.57968463110397 | 3.20132594093141 |
| chr14 | 104803075 | 104803082 | + | 5.01980288436682 | 3.41098722619315 | 3.57968463110397 | 4.26843458790854 |
| chr14 | 104896088 | 104896101 | + | 15.0594086531005 | 6.82197445238629 | 7.15936926220794 | 3.20132594093141 |
| chr14 | 104896224 | 104896278 | + | 44.174265382428 | 19.3289276150945 | 5.96614105183995 | 5.33554323488568 |
| chr14 | 104896600 | 104896635 | + | 7.02772403811355 | 7.95897019445067 | 1.19322821036799 | 4.26843458790854 |
| chr14 | 104896658 | 104896711 | + | 5.01980288436682 | 2.27399148412876 | 3.57968463110397 | 1.06710864697714 |
| chr14 | 105010085 | 105010114 | - | 18.0712903837206 | 12.5069531627082 | 5.96614105183995 | 7.46976052883995 |
| chr14 | 105010474 | 105010489 | - | 3.01188173062009 | 4.54798296825753 | 13.1255103140479 | 3.20132594093141 |
| chr14 | 105010604 | 105010609 | - | 2.00792115374673 | 3.41098722619315 | 1.19322821036799 | 2.13421729395427 |
| chr14 | 105010691 | 105010704 | - | 9.03564519186028 | 7.95897019445067 | 15.5119667347839 | 2.13421729395427 |
| chr14 | 105010833 | 105010850 | - | 4.01584230749346 | 12.5069531627082 | 44.1494437836156 | 36.2816939972226 |
| chr14 | 105010874 | 105010881 | - | 3.01188173062009 | 4.54798296825753 | 4.77291284147196 | 1.06710864697714 |
| chr14 | 105010908 | 105010917 | - | 6.02376346124019 | 4.54798296825753 | 2.38645642073598 | 5.33554323488568 |
| chr14 | 105011000 | 105011016 | - | 3.01188173062009 | 2.27399148412876 | 3.57968463110397 | 6.40265188186281 |
| chr14 | 105141128 | 105141137 | - | 9.03564519186028 | 1.13699574206438 | 8.35259747257593 | 7.46976052883995 |
| chr14 | 105142375 | 105142385 | - | 2.00792115374673 | 3.41098722619315 | 5.96614105183995 | 4.26843458790854 |
| chr14 | 105142500 | 105142503 | - | 1.00396057687336 | 2.27399148412876 | 3.57968463110397 | 3.20132594093141 |
| chr14 | 105209308 | 105209355 | - | 12.0475269224804 | 20.4659233571589 | 4.77291284147196 | 8.53686917581709 |
| chr14 | 105470351 | 105470381 | + | 6.02376346124019 | 6.82197445238629 | 4.77291284147196 | 3.20132594093141 |
| chr14 | 105470482 | 105470538 | + | 16.0633692299738 | 4.54798296825753 | 11.9322821036799 | 9.60397782279422 |
| chr14 | 105470582 | 105470655 | + | 17.0673298068472 | 31.8358807778027 | 10.7390538933119 | 13.8724124107028 |
| chr14 | 105480039 | 105480094 | + | 100.396057687336 | 51.1648083928972 | 1.19322821036799 | 3.20132594093141 |
| chr14 | 105499162 | 105499219 | + | 33.130699036821 | 56.8497871032191 | 2.38645642073598 | 12.8053037637256 |
| chr15 | 22826473 | 22826480 | + | 4.01584230749346 | 11.3699574206438 | 2.38645642073598 | 1.06710864697714 |
| chr15 | 22829330 | 22829377 | + | 20.0792115374673 | 27.2878978095452 | 7.15936926220794 | 13.8724124107028 |
| chr15 | 22829445 | 22829496 | + | 7.02772403811355 | 4.54798296825753 | 7.15936926220794 | 1.06710864697714 |
| chr15 | 22829520 | 22829578 | + | 17.0673298068472 | 23.876910583352 | 4.77291284147196 | 8.53686917581709 |
| chr15 | 22829600 | 22829610 | + | 86.3406096111093 | 112.562578464374 | 88.2988875672312 | 84.3015831111937 |
| chr15 | 22829643 | 22829645 | + | 3.01188173062009 | 2.27399148412876 | 3.57968463110397 | 2.13421729395427 |
| chr15 | 22866863 | 22866884 | + | 3.01188173062009 | 2.27399148412876 | 3.57968463110397 | 2.13421729395427 |
| chr15 | 22867109 | 22867124 | + | 4.01584230749346 | 1.13699574206438 | 4.77291284147196 | 2.13421729395427 |
| chr15 | 22867163 | 22867187 | + | 8.03168461498692 | 11.3699574206438 | 13.1255103140479 | 6.40265188186281 |
| chr15 | 22867290 | 22867319 | + | 10.0396057687336 | 9.09596593651506 | 20.2848795762558 | 24.5434988804741 |
| chr15 | 22867342 | 22867371 | + | 19.0752509605939 | 17.0549361309657 | 25.0577924177278 | 16.006629704657 |
| chr15 | 22867398 | 22867473 | + | 22.087132691214 | 19.3289276150945 | 29.8307052591997 | 49.0869977609482 |
| chr15 | 22867524 | 22867531 | + | 4.01584230749346 | 4.54798296825753 | 2.38645642073598 | 6.40265188186281 |
| chr15 | 22867653 | 22867694 | + | 6.02376346124019 | 10.2329616785794 | 10.7390538933119 | 6.40265188186281 |
| chr15 | 22869600 | 22869679 | - | 5.01980288436682 | 12.5069531627082 | 1.19322821036799 | 1.06710864697714 |
| chr15 | 22869736 | 22869744 | - | 5.01980288436682 | 6.82197445238629 | 5.96614105183995 | 4.26843458790854 |
| chr15 | 22869798 | 22869838 | - | 6.02376346124019 | 10.2329616785794 | 13.1255103140479 | 17.0737383516342 |
| chr15 | 22869917 | 22869971 | - | 147.582204800385 | 144.398459242177 | 139.607700613055 | 131.254363578188 |
| chr15 | 22869997 | 22870026 | - | 8.03168461498692 | 6.82197445238629 | 11.9322821036799 | 6.40265188186281 |
| chr15 | 25333776 | 25333778 | - | 2.00792115374673 | 1.13699574206438 | 3.57968463110397 | 2.13421729395427 |
| chr15 | 25333889 | 25333893 | - | 1.00396057687336 | 3.41098722619315 | 7.15936926220794 | 12.8053037637256 |
| chr15 | 25334634 | 25334636 | - | 3.01188173062009 | 1.13699574206438 | 3.57968463110397 | 2.13421729395427 |
| chr15 | 25335901 | 25335904 | - | 2.00792115374673 | 3.41098722619315 | 1.19322821036799 | 1.06710864697714 |
| chr15 | 25337586 | 25337600 | - | 8.03168461498692 | 2.27399148412876 | 3.57968463110397 | 5.33554323488568 |
| chr15 | 25338301 | 25338324 | - | 2.00792115374673 | 3.41098722619315 | 4.77291284147196 | 3.20132594093141 |
| chr15 | 25338344 | 25338353 | - | 1.00396057687336 | 1.13699574206438 | 1.19322821036799 | 4.26843458790854 |
| chr15 | 25338642 | 25338648 | - | 2.00792115374673 | 6.82197445238629 | 17.8984231555198 | 14.9395210576799 |
| chr15 | 25338698 | 25338716 | - | 2.00792115374673 | 1.13699574206438 | 9.54582568294392 | 4.26843458790854 |
| chr15 | 25338830 | 25338856 | - | 9.03564519186028 | 6.82197445238629 | 20.2848795762558 | 27.7448248214055 |
| chr15 | 25338879 | 25338882 | - | 2.00792115374673 | 4.54798296825753 | 2.38645642073598 | 2.13421729395427 |
| chr15 | 25338948 | 25339018 | - | 32.1267384599477 | 37.5208594881246 | 36.9900745214077 | 37.3488026441998 |
| chr15 | 25339055 | 25339077 | - | 13.0514874993537 | 17.0549361309657 | 11.9322821036799 | 16.006629704657 |
| chr15 | 25339145 | 25339208 | - | 6.02376346124019 | 7.95897019445067 | 13.1255103140479 | 7.46976052883995 |
| chr15 | 25339257 | 25339259 | - | 2.00792115374673 | 6.82197445238629 | 15.5119667347839 | 2.13421729395427 |
| chr15 | 25354305 | 25354309 | - | 3.01188173062009 | 5.68497871032191 | 1.19322821036799 | 3.20132594093141 |
| chr15 | 33866379 | 33866384 | - | 4.01584230749346 | 12.5069531627082 | 4.77291284147196 | 4.26843458790854 |
| chr15 | 33866550 | 33866559 | - | 2.00792115374673 | 2.27399148412876 | 1.19322821036799 | 1.06710864697714 |
| chr15 | 33866606 | 33866628 | - | 1.00396057687336 | 9.09596593651506 | 2.38645642073598 | 2.13421729395427 |
| chr15 | 34084052 | 34084176 | - | 215.851524027773 | 230.81013563907 | 88.2988875672312 | 92.8384522870108 |
| chr15 | 34084207 | 34084300 | - | 25.0990144218341 | 38.657855230189 | 14.3187385244159 | 9.60397782279422 |
| chr15 | 34084326 | 34084354 | - | 6.02376346124019 | 6.82197445238629 | 3.57968463110397 | 3.20132594093141 |
| chr15 | 34141499 | 34141507 | - | 1.00396057687336 | 1.13699574206438 | 2.38645642073598 | 2.13421729395427 |
| chr15 | 34142115 | 34142118 | - | 1.00396057687336 | 2.27399148412876 | 3.57968463110397 | 4.26843458790854 |
| chr15 | 34228271 | 34228277 | + | 2.00792115374673 | 1.13699574206438 | 3.57968463110397 | 5.33554323488568 |
| chr15 | 34228435 | 34228587 | + | 89.3524913417294 | 68.2197445238629 | 60.8546387287675 | 67.2278447595595 |
| chr15 | 34229750 | 34229826 | + | 225.891129796507 | 198.974254861267 | 219.55399070771 | 247.569206098695 |
| chr15 | 34229849 | 34230057 | + | 1404.54084704584 | 1696.39764716006 | 1571.48155305464 | 1375.50304595353 |
| chr15 | 34229994 | 34230016 | - | 3.01188173062009 | 2.27399148412876 | 1.19322821036799 | 1.06710864697714 |
| chr15 | 34230076 | 34230110 | + | 6.02376346124019 | 9.09596593651506 | 3.57968463110397 | 2.13421729395427 |
| chr15 | 34230133 | 34230136 | + | 3.01188173062009 | 2.27399148412876 | 4.77291284147196 | 2.13421729395427 |
| chr15 | 34230447 | 34230459 | - | 8.03168461498692 | 11.3699574206438 | 1.19322821036799 | 3.20132594093141 |
| chr15 | 34233567 | 34233575 | - | 2.00792115374673 | 6.82197445238629 | 2.38645642073598 | 1.06710864697714 |
| chr15 | 34341737 | 34341748 | - | 5.01980288436682 | 7.95897019445067 | 1.19322821036799 | 1.06710864697714 |
| chr15 | 34341784 | 34341925 | - | 111.439624032943 | 169.412365567593 | 114.549908195327 | 178.207144045182 |
| chr15 | 34341965 | 34341984 | - | 4.01584230749346 | 6.82197445238629 | 2.38645642073598 | 9.60397782279422 |
| chr15 | 34379099 | 34379259 | - | 419.655521133066 | 428.647394758272 | 1077.48507396229 | 707.493032945841 |
| chr15 | 34379280 | 34379308 | - | 9.03564519186028 | 20.4659233571589 | 26.2510206280958 | 16.006629704657 |
| chr15 | 34379387 | 34379436 | - | 136.538638454778 | 127.343523111211 | 244.611783125438 | 119.516168461439 |
| chr15 | 34379532 | 34379550 | - | 206.815878835913 | 234.221122865263 | 291.147683329789 | 176.072926751227 |
| chr15 | 34379640 | 34379644 | - | 1.00396057687336 | 3.41098722619315 | 3.57968463110397 | 3.20132594093141 |
| chr15 | 34379704 | 34379727 | - | 4.01584230749346 | 3.41098722619315 | 7.15936926220794 | 7.46976052883995 |
| chr15 | 34379888 | 34379895 | - | 3.01188173062009 | 2.27399148412876 | 2.38645642073598 | 8.53686917581709 |
| chr15 | 34380549 | 34380570 | - | 2.00792115374673 | 3.41098722619315 | 8.35259747257593 | 4.26843458790854 |
| chr15 | 34380792 | 34380801 | - | 1.00396057687336 | 6.82197445238629 | 29.8307052591997 | 9.60397782279422 |
| chr15 | 34380887 | 34380891 | - | 2.00792115374673 | 2.27399148412876 | 7.15936926220794 | 1.06710864697714 |
| chr15 | 34525305 | 34525328 | - | 5.01980288436682 | 5.68497871032191 | 8.35259747257593 | 2.13421729395427 |
| chr15 | 34525375 | 34525459 | - | 32.1267384599477 | 30.6988850357383 | 69.2072362013434 | 23.476390233497 |
| chr15 | 34525495 | 34525514 | - | 1.00396057687336 | 3.41098722619315 | 15.5119667347839 | 2.13421729395427 |
| chr15 | 34525556 | 34525565 | - | 4.01584230749346 | 4.54798296825753 | 15.5119667347839 | 2.13421729395427 |
| chr15 | 34525606 | 34525662 | - | 42.1663442286813 | 48.8908169087684 | 65.6275515702394 | 44.8185631730397 |
| chr15 | 34525672 | 34525769 | - | 19.0752509605939 | 21.6029190992233 | 35.7968463110397 | 14.9395210576799 |
| chr15 | 34854635 | 34854648 | - | 6.02376346124019 | 2.27399148412876 | 3.57968463110397 | 3.20132594093141 |
| chr15 | 34856588 | 34856589 | - | 1.00396057687336 | 1.13699574206438 | 3.57968463110397 | 1.06710864697714 |
| chr15 | 34856743 | 34856768 | - | 4.01584230749346 | 6.82197445238629 | 10.7390538933119 | 6.40265188186281 |
| chr15 | 34978384 | 34978416 | - | 7.02772403811355 | 11.3699574206438 | 4.77291284147196 | 2.13421729395427 |
| chr15 | 34978491 | 34978532 | - | 11.043566345607 | 13.6439489047726 | 7.15936926220794 | 17.0737383516342 |
| chr15 | 34978559 | 34978566 | - | 2.00792115374673 | 2.27399148412876 | 4.77291284147196 | 3.20132594093141 |
| chr15 | 34978638 | 34978654 | - | 1.00396057687336 | 1.13699574206438 | 7.15936926220794 | 6.40265188186281 |
| chr15 | 34978786 | 34978793 | - | 6.02376346124019 | 5.68497871032191 | 5.96614105183995 | 10.6710864697714 |
| chr15 | 34979653 | 34979668 | - | 13.0514874993537 | 19.3289276150945 | 5.96614105183995 | 5.33554323488568 |
| chr15 | 34980371 | 34980384 | - | 3.01188173062009 | 5.68497871032191 | 1.19322821036799 | 3.20132594093141 |
| chr15 | 34980443 | 34980448 | - | 2.00792115374673 | 1.13699574206438 | 1.19322821036799 | 13.8724124107028 |
| chr15 | 34980883 | 34980905 | - | 12.0475269224804 | 13.6439489047726 | 15.5119667347839 | 10.6710864697714 |
| chr15 | 34980950 | 34980993 | - | 10.0396057687336 | 34.1098722619315 | 20.2848795762558 | 44.8185631730397 |
| chr15 | 34981071 | 34981076 | - | 3.01188173062009 | 4.54798296825753 | 2.38645642073598 | 1.06710864697714 |
| chr15 | 34981219 | 34981228 | - | 16.0633692299738 | 32.9728765198671 | 28.6374770488317 | 12.8053037637256 |
| chr15 | 36809693 | 36809698 | + | 4.01584230749346 | 9.09596593651506 | 5.96614105183995 | 6.40265188186281 |
| chr15 | 36891300 | 36891311 | - | 3.01188173062009 | 6.82197445238629 | 13.1255103140479 | 4.26843458790854 |
| chr15 | 36891336 | 36891400 | - | 24.0950538449607 | 31.8358807778027 | 38.1833027317757 | 45.8856718200168 |
| chr15 | 36891433 | 36891495 | - | 14.0554480762271 | 19.3289276150945 | 34.6036181006717 | 58.6909755837425 |
| chr15 | 36892279 | 36892284 | - | 3.01188173062009 | 7.95897019445067 | 9.54582568294392 | 13.8724124107028 |
| chr15 | 38352046 | 38352057 | + | 1.00396057687336 | 2.27399148412876 | 2.38645642073598 | 1.06710864697714 |
| chr15 | 38353113 | 38353137 | + | 4.01584230749346 | 2.27399148412876 | 1.19322821036799 | 3.20132594093141 |
| chr15 | 38353504 | 38353536 | + | 1.00396057687336 | 1.13699574206438 | 2.38645642073598 | 3.20132594093141 |
| chr15 | 38353971 | 38353986 | + | 1.00396057687336 | 2.27399148412876 | 2.38645642073598 | 3.20132594093141 |
| chr15 | 38485997 | 38486026 | + | 9.03564519186028 | 11.3699574206438 | 16.7051949451519 | 16.006629704657 |
| chr15 | 38486180 | 38486186 | + | 5.01980288436682 | 3.41098722619315 | 5.96614105183995 | 7.46976052883995 |
| chr15 | 39785485 | 39785497 | + | 18.0712903837206 | 10.2329616785794 | 8.35259747257593 | 7.46976052883995 |
| chr15 | 40035516 | 40035540 | + | 1.00396057687336 | 2.27399148412876 | 1.19322821036799 | 1.06710864697714 |
| chr15 | 40035781 | 40035826 | - | 29.1148567293276 | 46.6168254246397 | 33.4103898903037 | 43.7514545260626 |
| chr15 | 40036095 | 40036112 | - | 16.0633692299738 | 34.1098722619315 | 15.5119667347839 | 3.20132594093141 |
| chr15 | 40036216 | 40036240 | - | 74.293082688629 | 73.9047232341848 | 51.3088130458235 | 39.483019938154 |
| chr15 | 40036270 | 40036276 | - | 3.01188173062009 | 1.13699574206438 | 1.19322821036799 | 8.53686917581709 |
| chr15 | 40036329 | 40036409 | - | 54.2138711511617 | 45.4798296825753 | 25.0577924177278 | 49.0869977609482 |
| chr15 | 40039272 | 40039349 | + | 127.502993262917 | 358.15365875028 | 385.412711948861 | 357.48139673734 |
| chr15 | 40220927 | 40220946 | + | 6.02376346124019 | 4.54798296825753 | 3.57968463110397 | 2.13421729395427 |
| chr15 | 40220986 | 40221003 | + | 11.043566345607 | 15.9179403889013 | 9.54582568294392 | 10.6710864697714 |
| chr15 | 40324527 | 40324542 | + | 1.00396057687336 | 2.27399148412876 | 5.96614105183995 | 1.06710864697714 |
| chr15 | 40324714 | 40324717 | + | 1.00396057687336 | 1.13699574206438 | 4.77291284147196 | 2.13421729395427 |
| chr15 | 40324776 | 40324789 | + | 2.00792115374673 | 7.95897019445067 | 20.2848795762558 | 9.60397782279422 |
| chr15 | 40324871 | 40324894 | + | 5.01980288436682 | 2.27399148412876 | 5.96614105183995 | 6.40265188186281 |
| chr15 | 40325265 | 40325315 | + | 5.01980288436682 | 2.27399148412876 | 4.77291284147196 | 10.6710864697714 |
| chr15 | 40325978 | 40325988 | + | 1.00396057687336 | 1.13699574206438 | 8.35259747257593 | 9.60397782279422 |
| chr15 | 40326184 | 40326188 | + | 2.00792115374673 | 2.27399148412876 | 5.96614105183995 | 11.7381951167485 |
| chr15 | 40326268 | 40326281 | + | 4.01584230749346 | 11.3699574206438 | 14.3187385244159 | 11.7381951167485 |
| chr15 | 40326402 | 40326422 | + | 1.00396057687336 | 3.41098722619315 | 3.57968463110397 | 3.20132594093141 |
| chr15 | 40393960 | 40393982 | + | 4.01584230749346 | 4.54798296825753 | 4.77291284147196 | 7.46976052883995 |
| chr15 | 40394012 | 40394013 | + | 4.01584230749346 | 1.13699574206438 | 1.19322821036799 | 3.20132594093141 |
| chr15 | 40394060 | 40394067 | + | 7.02772403811355 | 6.82197445238629 | 1.19322821036799 | 5.33554323488568 |
| chr15 | 40394127 | 40394147 | + | 19.0752509605939 | 7.95897019445067 | 1.19322821036799 | 8.53686917581709 |
| chr15 | 40420304 | 40420384 | + | 28.1108961524542 | 14.780944646837 | 1.19322821036799 | 5.33554323488568 |
| chr15 | 40421230 | 40421251 | + | 7.02772403811355 | 7.95897019445067 | 3.57968463110397 | 3.20132594093141 |
| chr15 | 40466297 | 40466335 | + | 4.01584230749346 | 6.82197445238629 | 20.2848795762558 | 8.53686917581709 |
| chr15 | 40466455 | 40466467 | + | 4.01584230749346 | 5.68497871032191 | 21.4781077866238 | 4.26843458790854 |
| chr15 | 40466911 | 40466945 | + | 8.03168461498692 | 6.82197445238629 | 21.4781077866238 | 13.8724124107028 |
| chr15 | 40466972 | 40467010 | + | 1.00396057687336 | 1.13699574206438 | 5.96614105183995 | 14.9395210576799 |
| chr15 | 40468140 | 40468146 | + | 2.00792115374673 | 4.54798296825753 | 3.57968463110397 | 8.53686917581709 |
| chr15 | 40553167 | 40553177 | - | 3.01188173062009 | 6.82197445238629 | 5.96614105183995 | 17.0737383516342 |
| chr15 | 40574261 | 40574321 | + | 11.043566345607 | 13.6439489047726 | 1.19322821036799 | 4.26843458790854 |
| chr15 | 40574347 | 40574387 | + | 19.0752509605939 | 12.5069531627082 | 7.15936926220794 | 12.8053037637256 |
| chr15 | 40574413 | 40574435 | + | 12.0475269224804 | 25.0139063254164 | 3.57968463110397 | 3.20132594093141 |
| chr15 | 40574609 | 40574611 | + | 1.00396057687336 | 1.13699574206438 | 1.19322821036799 | 1.06710864697714 |
| chr15 | 40663334 | 40663341 | + | 22.087132691214 | 26.1509020674808 | 32.2171616799357 | 21.3421729395427 |
| chr15 | 40664154 | 40664193 | + | 14.0554480762271 | 10.2329616785794 | 8.35259747257593 | 5.33554323488568 |
| chr15 | 40731522 | 40731528 | + | 1.00396057687336 | 1.13699574206438 | 3.57968463110397 | 8.53686917581709 |
| chr15 | 40735934 | 40735979 | - | 18.0712903837206 | 34.1098722619315 | 15.5119667347839 | 20.2750642925656 |
| chr15 | 40736097 | 40736100 | - | 6.02376346124019 | 10.2329616785794 | 3.57968463110397 | 5.33554323488568 |
| chr15 | 40736220 | 40736230 | - | 6.02376346124019 | 15.9179403889013 | 1.19322821036799 | 9.60397782279422 |
| chr15 | 40736264 | 40736267 | - | 10.0396057687336 | 17.0549361309657 | 2.38645642073598 | 1.06710864697714 |
| chr15 | 40736485 | 40736494 | - | 8.03168461498692 | 28.4248935516096 | 3.57968463110397 | 4.26843458790854 |
| chr15 | 40767226 | 40767348 | + | 41.1623836518079 | 27.2878978095452 | 10.7390538933119 | 8.53686917581709 |
| chr15 | 40767628 | 40767696 | + | 11.043566345607 | 18.1919318730301 | 3.57968463110397 | 4.26843458790854 |
| chr15 | 40767898 | 40767912 | - | 4.01584230749346 | 11.3699574206438 | 7.15936926220794 | 2.13421729395427 |
| chr15 | 40814512 | 40814543 | + | 26.1029749987075 | 28.4248935516096 | 1.19322821036799 | 4.26843458790854 |
| chr15 | 40872577 | 40872698 | - | 89.3524913417294 | 60.2607743294122 | 2.38645642073598 | 2.13421729395427 |
| chr15 | 40903936 | 40903952 | + | 10.0396057687336 | 7.95897019445067 | 1.19322821036799 | 3.20132594093141 |
| chr15 | 41279757 | 41279812 | + | 14.0554480762271 | 3.41098722619315 | 5.96614105183995 | 12.8053037637256 |
| chr15 | 41281630 | 41281700 | + | 8.03168461498692 | 9.09596593651506 | 5.96614105183995 | 11.7381951167485 |
| chr15 | 41283951 | 41284034 | + | 469.853549976735 | 461.620271278139 | 499.962620144188 | 471.662021963894 |
| chr15 | 41284062 | 41284083 | + | 4.01584230749346 | 6.82197445238629 | 9.54582568294392 | 3.20132594093141 |
| chr15 | 41287350 | 41287357 | + | 45.1782259593014 | 22.7399148412876 | 27.4442488384638 | 11.7381951167485 |
| chr15 | 41298925 | 41298948 | + | 7.02772403811355 | 9.09596593651506 | 9.54582568294392 | 10.6710864697714 |
| chr15 | 41299303 | 41299310 | + | 3.01188173062009 | 1.13699574206438 | 3.57968463110397 | 2.13421729395427 |
| chr15 | 41299392 | 41299402 | + | 8.03168461498692 | 19.3289276150945 | 10.7390538933119 | 9.60397782279422 |
| chr15 | 41299546 | 41299569 | + | 12.0475269224804 | 6.82197445238629 | 11.9322821036799 | 20.2750642925656 |
| chr15 | 41301426 | 41301473 | + | 3.01188173062009 | 1.13699574206438 | 1.19322821036799 | 1.06710864697714 |
| chr15 | 41301511 | 41301561 | + | 10.0396057687336 | 13.6439489047726 | 3.57968463110397 | 11.7381951167485 |
| chr15 | 41301681 | 41301740 | + | 9.03564519186028 | 7.95897019445067 | 16.7051949451519 | 7.46976052883995 |
| chr15 | 41302995 | 41303008 | + | 2.00792115374673 | 1.13699574206438 | 1.19322821036799 | 2.13421729395427 |
| chr15 | 41304149 | 41304156 | + | 3.01188173062009 | 1.13699574206438 | 4.77291284147196 | 3.20132594093141 |
| chr15 | 41304760 | 41304765 | + | 1.00396057687336 | 4.54798296825753 | 2.38645642073598 | 1.06710864697714 |
| chr15 | 41309298 | 41309320 | - | 30.1188173062009 | 28.4248935516096 | 2.38645642073598 | 4.26843458790854 |
| chr15 | 41309358 | 41309408 | - | 27.1069355755808 | 39.7948509722534 | 13.1255103140479 | 10.6710864697714 |
| chr15 | 41309577 | 41309589 | - | 3.01188173062009 | 10.2329616785794 | 4.77291284147196 | 5.33554323488568 |
| chr15 | 41309649 | 41309653 | - | 23.0910932680874 | 47.753821166704 | 5.96614105183995 | 11.7381951167485 |
| chr15 | 41380092 | 41380434 | + | 1094.31702879197 | 927.788525524536 | 464.165773833148 | 513.279259196002 |
| chr15 | 41380592 | 41380613 | + | 24.0950538449607 | 22.7399148412876 | 2.38645642073598 | 7.46976052883995 |
| chr15 | 41380651 | 41380660 | + | 25.0990144218341 | 17.0549361309657 | 1.19322821036799 | 10.6710864697714 |
| chr15 | 41380696 | 41380726 | + | 12.0475269224804 | 18.1919318730301 | 7.15936926220794 | 2.13421729395427 |
| chr15 | 41380885 | 41380897 | + | 13.0514874993537 | 14.780944646837 | 2.38645642073598 | 3.20132594093141 |
| chr15 | 41380935 | 41380956 | + | 34.1346596136944 | 10.2329616785794 | 5.96614105183995 | 4.26843458790854 |
| chr15 | 41380987 | 41381019 | + | 43.1703048055547 | 35.2468680039958 | 11.9322821036799 | 7.46976052883995 |
| chr15 | 41387408 | 41387410 | - | 6.02376346124019 | 6.82197445238629 | 2.38645642073598 | 4.26843458790854 |
| chr15 | 41480891 | 41480917 | + | 9.03564519186028 | 4.54798296825753 | 7.15936926220794 | 3.20132594093141 |
| chr15 | 41481249 | 41481301 | + | 67.2653586505154 | 76.1787147183136 | 32.2171616799357 | 40.5501285851312 |
| chr15 | 41481337 | 41481358 | + | 7.02772403811355 | 3.41098722619315 | 3.57968463110397 | 6.40265188186281 |
| chr15 | 41482309 | 41482326 | + | 1.00396057687336 | 1.13699574206438 | 5.96614105183995 | 10.6710864697714 |
| chr15 | 41482420 | 41482447 | + | 26.1029749987075 | 18.1919318730301 | 28.6374770488317 | 58.6909755837425 |
| chr15 | 41482483 | 41482518 | + | 34.1346596136944 | 18.1919318730301 | 14.3187385244159 | 16.006629704657 |
| chr15 | 41482555 | 41482595 | + | 7.02772403811355 | 6.82197445238629 | 4.77291284147196 | 9.60397782279422 |
| chr15 | 41482622 | 41482670 | + | 20.0792115374673 | 23.876910583352 | 13.1255103140479 | 18.1408469986113 |
| chr15 | 41482702 | 41482735 | + | 7.02772403811355 | 6.82197445238629 | 8.35259747257593 | 9.60397782279422 |
| chr15 | 41482784 | 41482808 | + | 5.01980288436682 | 2.27399148412876 | 2.38645642073598 | 2.13421729395427 |
| chr15 | 41482876 | 41482880 | + | 2.00792115374673 | 4.54798296825753 | 2.38645642073598 | 4.26843458790854 |
| chr15 | 41482911 | 41482935 | + | 4.01584230749346 | 3.41098722619315 | 8.35259747257593 | 1.06710864697714 |
| chr15 | 41503494 | 41503527 | + | 4.01584230749346 | 5.68497871032191 | 11.9322821036799 | 9.60397782279422 |
| chr15 | 41517288 | 41517318 | - | 1.00396057687336 | 3.41098722619315 | 1.19322821036799 | 6.40265188186281 |
| chr15 | 41579235 | 41579263 | + | 11.043566345607 | 13.6439489047726 | 1.19322821036799 | 9.60397782279422 |
| chr15 | 41767315 | 41767317 | + | 7.02772403811355 | 2.27399148412876 | 1.19322821036799 | 3.20132594093141 |
| chr15 | 41767419 | 41767427 | + | 14.0554480762271 | 15.9179403889013 | 4.77291284147196 | 6.40265188186281 |
| chr15 | 41767523 | 41767566 | + | 13.0514874993537 | 7.95897019445067 | 2.38645642073598 | 3.20132594093141 |
| chr15 | 41767618 | 41767670 | + | 17.0673298068472 | 7.95897019445067 | 9.54582568294392 | 2.13421729395427 |
| chr15 | 41767703 | 41767733 | + | 7.02772403811355 | 10.2329616785794 | 5.96614105183995 | 13.8724124107028 |
| chr15 | 41767765 | 41767778 | + | 13.0514874993537 | 15.9179403889013 | 2.38645642073598 | 4.26843458790854 |
| chr15 | 41767802 | 41767810 | + | 5.01980288436682 | 3.41098722619315 | 2.38645642073598 | 1.06710864697714 |
| chr15 | 41768152 | 41768159 | + | 13.0514874993537 | 15.9179403889013 | 8.35259747257593 | 13.8724124107028 |
| chr15 | 41768244 | 41768285 | + | 5.01980288436682 | 11.3699574206438 | 7.15936926220794 | 9.60397782279422 |
| chr15 | 41768355 | 41768398 | + | 7.02772403811355 | 12.5069531627082 | 13.1255103140479 | 3.20132594093141 |
| chr15 | 41768426 | 41768458 | + | 3.01188173062009 | 6.82197445238629 | 9.54582568294392 | 2.13421729395427 |
| chr15 | 41768493 | 41768542 | + | 47.1861471130481 | 54.5757956190903 | 33.4103898903037 | 39.483019938154 |
| chr15 | 41769597 | 41769601 | + | 9.03564519186028 | 6.82197445238629 | 1.19322821036799 | 2.13421729395427 |
| chr15 | 41769646 | 41769684 | + | 3.01188173062009 | 11.3699574206438 | 4.77291284147196 | 1.06710864697714 |
| chr15 | 41771126 | 41771141 | + | 8.03168461498692 | 6.82197445238629 | 4.77291284147196 | 9.60397782279422 |
| chr15 | 41771180 | 41771187 | + | 6.02376346124019 | 5.68497871032191 | 3.57968463110397 | 2.13421729395427 |
| chr15 | 41771247 | 41771266 | + | 8.03168461498692 | 7.95897019445067 | 3.57968463110397 | 6.40265188186281 |
| chr15 | 41771300 | 41771301 | + | 4.01584230749346 | 2.27399148412876 | 4.77291284147196 | 2.13421729395427 |
| chr15 | 41771373 | 41771398 | + | 11.043566345607 | 11.3699574206438 | 4.77291284147196 | 2.13421729395427 |
| chr15 | 41837498 | 41837514 | + | 1.00396057687336 | 2.27399148412876 | 1.19322821036799 | 4.26843458790854 |
| chr15 | 41899469 | 41899471 | - | 3.01188173062009 | 1.13699574206438 | 1.19322821036799 | 2.13421729395427 |
| chr15 | 41900611 | 41900615 | - | 1.00396057687336 | 1.13699574206438 | 4.77291284147196 | 2.13421729395427 |
| chr15 | 42158741 | 42158803 | - | 17.0673298068472 | 15.9179403889013 | 7.15936926220794 | 5.33554323488568 |
| chr15 | 42210637 | 42210673 | - | 7.02772403811355 | 14.780944646837 | 4.77291284147196 | 5.33554323488568 |
| chr15 | 42210700 | 42210732 | - | 6.02376346124019 | 6.82197445238629 | 2.38645642073598 | 1.06710864697714 |
| chr15 | 42210741 | 42210784 | - | 36.1425807674411 | 67.0827487817985 | 2.38645642073598 | 3.20132594093141 |
| chr15 | 42210813 | 42210924 | - | 178.704982683459 | 320.632799262156 | 56.0817258872955 | 48.0198891139711 |
| chr15 | 42210954 | 42210975 | - | 33.130699036821 | 47.753821166704 | 8.35259747257593 | 17.0737383516342 |
| chr15 | 42211070 | 42211075 | - | 2.00792115374673 | 3.41098722619315 | 3.57968463110397 | 3.20132594093141 |
| chr15 | 42211113 | 42211139 | - | 5.01980288436682 | 5.68497871032191 | 3.57968463110397 | 2.13421729395427 |
| chr15 | 42211173 | 42211240 | - | 25.0990144218341 | 50.0278126508328 | 13.1255103140479 | 14.9395210576799 |
| chr15 | 42211263 | 42211323 | - | 418.651560556193 | 584.415811421092 | 219.55399070771 | 178.207144045182 |
| chr15 | 42211418 | 42211456 | - | 14.0554480762271 | 10.2329616785794 | 3.57968463110397 | 2.13421729395427 |
| chr15 | 42211499 | 42211751 | - | 201.796075951546 | 342.235718361379 | 112.163451774591 | 151.529427870753 |
| chr15 | 42412845 | 42412850 | - | 6.02376346124019 | 25.0139063254164 | 5.96614105183995 | 4.26843458790854 |
| chr15 | 42412895 | 42412897 | - | 1.00396057687336 | 4.54798296825753 | 1.19322821036799 | 1.06710864697714 |
| chr15 | 42413023 | 42413041 | - | 6.02376346124019 | 2.27399148412876 | 4.77291284147196 | 5.33554323488568 |
| chr15 | 42413066 | 42413095 | - | 67.2653586505154 | 103.466612527859 | 19.0916513658878 | 66.1607361125824 |
| chr15 | 42413241 | 42413254 | - | 3.01188173062009 | 1.13699574206438 | 5.96614105183995 | 11.7381951167485 |
| chr15 | 42413263 | 42413303 | - | 3.01188173062009 | 1.13699574206438 | 5.96614105183995 | 7.46976052883995 |
| chr15 | 42413321 | 42413332 | - | 4.01584230749346 | 5.68497871032191 | 5.96614105183995 | 2.13421729395427 |
| chr15 | 42413413 | 42413416 | - | 1.00396057687336 | 1.13699574206438 | 1.19322821036799 | 2.13421729395427 |
| chr15 | 42413494 | 42413509 | - | 1.00396057687336 | 1.13699574206438 | 2.38645642073598 | 5.33554323488568 |
| chr15 | 42413528 | 42413639 | - | 38.1505019211879 | 30.6988850357383 | 51.3088130458235 | 53.3554323488568 |
| chr15 | 42413696 | 42413736 | - | 6.02376346124019 | 12.5069531627082 | 8.35259747257593 | 24.5434988804741 |
| chr15 | 42413768 | 42413771 | - | 12.0475269224804 | 10.2329616785794 | 7.15936926220794 | 5.33554323488568 |
| chr15 | 42414174 | 42414204 | - | 3.01188173062009 | 2.27399148412876 | 5.96614105183995 | 8.53686917581709 |
| chr15 | 42414648 | 42414657 | - | 1.00396057687336 | 2.27399148412876 | 1.19322821036799 | 5.33554323488568 |
| chr15 | 42415367 | 42415369 | - | 1.00396057687336 | 3.41098722619315 | 2.38645642073598 | 3.20132594093141 |
| chr15 | 42416828 | 42416835 | - | 2.00792115374673 | 2.27399148412876 | 1.19322821036799 | 3.20132594093141 |
| chr15 | 42416861 | 42416863 | - | 8.03168461498692 | 4.54798296825753 | 8.35259747257593 | 1.06710864697714 |
| chr15 | 42416919 | 42416924 | - | 1.00396057687336 | 4.54798296825753 | 1.19322821036799 | 1.06710864697714 |
| chr15 | 42417150 | 42417158 | - | 1.00396057687336 | 1.13699574206438 | 3.57968463110397 | 2.13421729395427 |
| chr15 | 42417182 | 42417219 | - | 7.02772403811355 | 3.41098722619315 | 5.96614105183995 | 9.60397782279422 |
| chr15 | 42531515 | 42531617 | + | 4.01584230749346 | 3.41098722619315 | 20.2848795762558 | 25.6106075274513 |
| chr15 | 42532379 | 42532402 | + | 4.01584230749346 | 3.41098722619315 | 10.7390538933119 | 24.5434988804741 |
| chr15 | 42532626 | 42532631 | + | 2.00792115374673 | 6.82197445238629 | 2.38645642073598 | 3.20132594093141 |
| chr15 | 42533006 | 42533034 | + | 7.02772403811355 | 6.82197445238629 | 26.2510206280958 | 17.0737383516342 |
| chr15 | 42540749 | 42540764 | - | 4.01584230749346 | 3.41098722619315 | 14.3187385244159 | 2.13421729395427 |
| chr15 | 42561325 | 42561403 | + | 24.0950538449607 | 19.3289276150945 | 14.3187385244159 | 17.0737383516342 |
| chr15 | 42561586 | 42561626 | + | 16.0633692299738 | 23.876910583352 | 5.96614105183995 | 7.46976052883995 |
| chr15 | 42743145 | 42743172 | - | 7.02772403811355 | 2.27399148412876 | 1.19322821036799 | 1.06710864697714 |
| chr15 | 42743320 | 42743336 | - | 5.01980288436682 | 3.41098722619315 | 1.19322821036799 | 1.06710864697714 |
| chr15 | 42743533 | 42743541 | - | 7.02772403811355 | 15.9179403889013 | 7.15936926220794 | 5.33554323488568 |
| chr15 | 42945229 | 42945251 | - | 14.0554480762271 | 19.3289276150945 | 14.3187385244159 | 8.53686917581709 |
| chr15 | 43185074 | 43185101 | + | 18.0712903837206 | 18.1919318730301 | 4.77291284147196 | 5.33554323488568 |
| chr15 | 43194883 | 43194890 | + | 2.00792115374673 | 3.41098722619315 | 2.38645642073598 | 4.26843458790854 |
| chr15 | 43194913 | 43194916 | + | 3.01188173062009 | 3.41098722619315 | 5.96614105183995 | 3.20132594093141 |
| chr15 | 43194939 | 43194946 | + | 4.01584230749346 | 6.82197445238629 | 5.96614105183995 | 3.20132594093141 |
| chr15 | 43194988 | 43194996 | + | 2.00792115374673 | 6.82197445238629 | 2.38645642073598 | 7.46976052883995 |
| chr15 | 43195091 | 43195129 | + | 13.0514874993537 | 10.2329616785794 | 11.9322821036799 | 8.53686917581709 |
| chr15 | 43195149 | 43195178 | + | 16.0633692299738 | 12.5069531627082 | 10.7390538933119 | 6.40265188186281 |
| chr15 | 43328067 | 43328069 | - | 4.01584230749346 | 5.68497871032191 | 1.19322821036799 | 1.06710864697714 |
| chr15 | 43328205 | 43328208 | - | 1.00396057687336 | 3.41098722619315 | 1.19322821036799 | 1.06710864697714 |
| chr15 | 43352299 | 43352300 | + | 3.01188173062009 | 2.27399148412876 | 3.57968463110397 | 1.06710864697714 |
| chr15 | 43352604 | 43352615 | + | 2.00792115374673 | 1.13699574206438 | 2.38645642073598 | 2.13421729395427 |
| chr15 | 43352899 | 43352943 | + | 6.02376346124019 | 18.1919318730301 | 14.3187385244159 | 19.2079556455884 |
| chr15 | 43358574 | 43358623 | - | 32.1267384599477 | 32.9728765198671 | 36.9900745214077 | 32.0132594093141 |
| chr15 | 43358648 | 43358778 | - | 267.053513448315 | 349.057692813765 | 181.370687975934 | 209.153294807519 |
| chr15 | 43358856 | 43358998 | - | 51.2019894205416 | 56.8497871032191 | 66.8207797806074 | 66.1607361125824 |
| chr15 | 43359019 | 43359067 | - | 31.1227778830743 | 68.2197445238629 | 56.0817258872955 | 39.483019938154 |
| chr15 | 43359090 | 43359184 | - | 41.1623836518079 | 29.5618892936739 | 45.3426719939836 | 25.6106075274513 |
| chr15 | 43359213 | 43359247 | - | 4.01584230749346 | 3.41098722619315 | 3.57968463110397 | 1.06710864697714 |
| chr15 | 43405313 | 43405327 | + | 7.02772403811355 | 3.41098722619315 | 2.38645642073598 | 2.13421729395427 |
| chr15 | 43405397 | 43405431 | + | 9.03564519186028 | 13.6439489047726 | 10.7390538933119 | 11.7381951167485 |
| chr15 | 43405525 | 43405571 | + | 7.02772403811355 | 7.95897019445067 | 7.15936926220794 | 10.6710864697714 |
| chr15 | 43406163 | 43406176 | + | 5.01980288436682 | 6.82197445238629 | 2.38645642073598 | 3.20132594093141 |
| chr15 | 43407342 | 43407347 | - | 3.01188173062009 | 3.41098722619315 | 3.57968463110397 | 4.26843458790854 |
| chr15 | 43407530 | 43407532 | + | 1.00396057687336 | 1.13699574206438 | 5.96614105183995 | 11.7381951167485 |
| chr15 | 43407584 | 43407598 | + | 5.01980288436682 | 1.13699574206438 | 2.38645642073598 | 1.06710864697714 |
| chr15 | 43409715 | 43409724 | + | 2.00792115374673 | 6.82197445238629 | 2.38645642073598 | 1.06710864697714 |
| chr15 | 43531421 | 43531477 | + | 3.01188173062009 | 5.68497871032191 | 17.8984231555198 | 67.2278447595595 |
| chr15 | 43531511 | 43531519 | + | 3.01188173062009 | 1.13699574206438 | 9.54582568294392 | 11.7381951167485 |
| chr15 | 43533500 | 43533535 | - | 17.0673298068472 | 31.8358807778027 | 10.7390538933119 | 24.5434988804741 |
| chr15 | 43771185 | 43771306 | + | 13.0514874993537 | 7.95897019445067 | 13.1255103140479 | 20.2750642925656 |
| chr15 | 43771414 | 43771524 | + | 85.336649034236 | 109.151591238181 | 51.3088130458235 | 62.959410171651 |
| chr15 | 43773166 | 43773181 | + | 6.02376346124019 | 4.54798296825753 | 10.7390538933119 | 8.53686917581709 |
| chr15 | 43793761 | 43793786 | + | 3.01188173062009 | 1.13699574206438 | 5.96614105183995 | 4.26843458790854 |
| chr15 | 43793819 | 43793881 | + | 8.03168461498692 | 3.41098722619315 | 4.77291284147196 | 19.2079556455884 |
| chr15 | 43793965 | 43793975 | + | 314.239660561363 | 253.550050480357 | 363.934604162237 | 652.00338330303 |
| chr15 | 43794056 | 43794086 | + | 8.03168461498692 | 7.95897019445067 | 7.15936926220794 | 3.20132594093141 |
| chr15 | 43801734 | 43801849 | + | 351.386201905678 | 485.497181861491 | 108.583767143487 | 166.468948928433 |
| chr15 | 43804608 | 43804615 | - | 2.00792115374673 | 9.09596593651506 | 4.77291284147196 | 1.06710864697714 |
| chr15 | 43804642 | 43804659 | - | 1.00396057687336 | 12.5069531627082 | 4.77291284147196 | 4.26843458790854 |
| chr15 | 43804744 | 43804745 | - | 1.00396057687336 | 1.13699574206438 | 1.19322821036799 | 1.06710864697714 |
| chr15 | 43805038 | 43805056 | - | 3.01188173062009 | 1.13699574206438 | 1.19322821036799 | 2.13421729395427 |
| chr15 | 43866599 | 43866611 | + | 18.0712903837206 | 18.1919318730301 | 13.1255103140479 | 16.006629704657 |
| chr15 | 43868176 | 43868180 | + | 7.02772403811355 | 13.6439489047726 | 2.38645642073598 | 1.06710864697714 |
| chr15 | 44413658 | 44413690 | + | 3.01188173062009 | 5.68497871032191 | 3.57968463110397 | 4.26843458790854 |
| chr15 | 44413715 | 44413750 | + | 4.01584230749346 | 2.27399148412876 | 3.57968463110397 | 7.46976052883995 |
| chr15 | 44413783 | 44413801 | + | 4.01584230749346 | 4.54798296825753 | 4.77291284147196 | 19.2079556455884 |
| chr15 | 44414161 | 44414166 | + | 1.00396057687336 | 3.41098722619315 | 1.19322821036799 | 3.20132594093141 |
| chr15 | 44414225 | 44414268 | + | 3.01188173062009 | 4.54798296825753 | 10.7390538933119 | 16.006629704657 |
| chr15 | 44415381 | 44415397 | + | 1.00396057687336 | 2.27399148412876 | 1.19322821036799 | 1.06710864697714 |
| chr15 | 44415433 | 44415465 | + | 8.03168461498692 | 2.27399148412876 | 3.57968463110397 | 3.20132594093141 |
| chr15 | 44524197 | 44524217 | + | 32.1267384599477 | 67.0827487817985 | 29.8307052591997 | 19.2079556455884 |
| chr15 | 44524299 | 44524328 | + | 5.01980288436682 | 4.54798296825753 | 9.54582568294392 | 4.26843458790854 |
| chr15 | 44524380 | 44524399 | + | 1.00396057687336 | 3.41098722619315 | 5.96614105183995 | 3.20132594093141 |
| chr15 | 44524421 | 44524430 | + | 4.01584230749346 | 3.41098722619315 | 5.96614105183995 | 1.06710864697714 |
| chr15 | 44524490 | 44524563 | + | 10.0396057687336 | 13.6439489047726 | 8.35259747257593 | 7.46976052883995 |
| chr15 | 44524602 | 44524619 | + | 4.01584230749346 | 6.82197445238629 | 7.15936926220794 | 4.26843458790854 |
| chr15 | 44524698 | 44524735 | + | 13.0514874993537 | 6.82197445238629 | 29.8307052591997 | 5.33554323488568 |
| chr15 | 44524837 | 44524859 | + | 1.00396057687336 | 7.95897019445067 | 3.57968463110397 | 1.06710864697714 |
| chr15 | 44524916 | 44524943 | + | 2.00792115374673 | 10.2329616785794 | 13.1255103140479 | 4.26843458790854 |
| chr15 | 44524974 | 44524982 | + | 1.00396057687336 | 3.41098722619315 | 8.35259747257593 | 2.13421729395427 |
| chr15 | 44525073 | 44525077 | + | 26.1029749987075 | 42.0688424563821 | 11.9322821036799 | 24.5434988804741 |
| chr15 | 44525384 | 44525446 | + | 22.087132691214 | 35.2468680039958 | 9.54582568294392 | 23.476390233497 |
| chr15 | 44525608 | 44525630 | + | 1.00396057687336 | 9.09596593651506 | 3.57968463110397 | 4.26843458790854 |
| chr15 | 44526842 | 44526849 | + | 5.01980288436682 | 5.68497871032191 | 2.38645642073598 | 9.60397782279422 |
| chr15 | 44527184 | 44527219 | + | 2.00792115374673 | 9.09596593651506 | 10.7390538933119 | 3.20132594093141 |
| chr15 | 44527482 | 44527502 | + | 1.00396057687336 | 4.54798296825753 | 1.19322821036799 | 2.13421729395427 |
| chr15 | 44527770 | 44527786 | + | 4.01584230749346 | 6.82197445238629 | 8.35259747257593 | 3.20132594093141 |
| chr15 | 44528159 | 44528169 | + | 4.01584230749346 | 2.27399148412876 | 8.35259747257593 | 4.26843458790854 |
| chr15 | 44528319 | 44528321 | + | 1.00396057687336 | 1.13699574206438 | 1.19322821036799 | 2.13421729395427 |
| chr15 | 44529002 | 44529014 | + | 5.01980288436682 | 17.0549361309657 | 3.57968463110397 | 2.13421729395427 |
| chr15 | 44561331 | 44561347 | + | 11.043566345607 | 20.4659233571589 | 7.15936926220794 | 5.33554323488568 |
| chr15 | 44561392 | 44561399 | + | 4.01584230749346 | 2.27399148412876 | 1.19322821036799 | 1.06710864697714 |
| chr15 | 44561564 | 44561573 | + | 10.0396057687336 | 7.95897019445067 | 11.9322821036799 | 5.33554323488568 |
| chr15 | 44561829 | 44561852 | + | 54.2138711511617 | 37.5208594881246 | 20.2848795762558 | 41.6172372321083 |
| chr15 | 44561933 | 44561937 | + | 9.03564519186028 | 2.27399148412876 | 3.57968463110397 | 1.06710864697714 |
| chr15 | 44562251 | 44562264 | + | 1.00396057687336 | 1.13699574206438 | 1.19322821036799 | 3.20132594093141 |
| chr15 | 44562659 | 44562661 | + | 1.00396057687336 | 3.41098722619315 | 4.77291284147196 | 6.40265188186281 |
| chr15 | 44562957 | 44562959 | - | 2.00792115374673 | 5.68497871032191 | 5.96614105183995 | 3.20132594093141 |
| chr15 | 44563107 | 44563127 | - | 4.01584230749346 | 6.82197445238629 | 1.19322821036799 | 4.26843458790854 |
| chr15 | 44716326 | 44716354 | + | 20.0792115374673 | 14.780944646837 | 5.96614105183995 | 13.8724124107028 |
| chr15 | 44717788 | 44717814 | + | 37.1465413443145 | 40.9318467143178 | 10.7390538933119 | 16.006629704657 |
| chr15 | 45167511 | 45167549 | - | 5.01980288436682 | 9.09596593651506 | 2.38645642073598 | 4.26843458790854 |
| chr15 | 45606507 | 45606530 | + | 1.00396057687336 | 3.41098722619315 | 5.96614105183995 | 2.13421729395427 |
| chr15 | 45607047 | 45607048 | + | 3.01188173062009 | 3.41098722619315 | 3.57968463110397 | 3.20132594093141 |
| chr15 | 45608119 | 45608121 | + | 2.00792115374673 | 1.13699574206438 | 2.38645642073598 | 1.06710864697714 |
| chr15 | 45608153 | 45608154 | + | 1.00396057687336 | 1.13699574206438 | 2.38645642073598 | 1.06710864697714 |
| chr15 | 45608174 | 45608193 | + | 4.01584230749346 | 3.41098722619315 | 8.35259747257593 | 8.53686917581709 |
| chr15 | 45608684 | 45608713 | + | 2.00792115374673 | 5.68497871032191 | 10.7390538933119 | 3.20132594093141 |
| chr15 | 45608793 | 45608810 | + | 1.00396057687336 | 5.68497871032191 | 5.96614105183995 | 3.20132594093141 |
| chr15 | 45609376 | 45609380 | + | 2.00792115374673 | 3.41098722619315 | 1.19322821036799 | 2.13421729395427 |
| chr15 | 45609424 | 45609451 | + | 1.00396057687336 | 3.41098722619315 | 2.38645642073598 | 2.13421729395427 |
| chr15 | 45609598 | 45609599 | + | 4.01584230749346 | 2.27399148412876 | 2.38645642073598 | 2.13421729395427 |
| chr15 | 48341523 | 48341568 | + | 15.0594086531005 | 13.6439489047726 | 5.96614105183995 | 7.46976052883995 |
| chr15 | 48342027 | 48342059 | + | 128.506953839791 | 170.549361309657 | 25.0577924177278 | 40.5501285851312 |
| chr15 | 48342085 | 48342108 | + | 7.02772403811355 | 10.2329616785794 | 1.19322821036799 | 3.20132594093141 |
| chr15 | 48342150 | 48342162 | + | 10.0396057687336 | 11.3699574206438 | 2.38645642073598 | 3.20132594093141 |
| chr15 | 48342233 | 48342236 | + | 16.0633692299738 | 50.0278126508328 | 20.2848795762558 | 19.2079556455884 |
| chr15 | 48342306 | 48342320 | + | 11.043566345607 | 18.1919318730301 | 3.57968463110397 | 1.06710864697714 |
| chr15 | 48342346 | 48342391 | + | 8.03168461498692 | 7.95897019445067 | 1.19322821036799 | 2.13421729395427 |
| chr15 | 48342508 | 48342528 | + | 9.03564519186028 | 6.82197445238629 | 2.38645642073598 | 2.13421729395427 |
| chr15 | 48343313 | 48343320 | + | 5.01980288436682 | 10.2329616785794 | 1.19322821036799 | 2.13421729395427 |
| chr15 | 48738193 | 48738198 | - | 4.01584230749346 | 9.09596593651506 | 3.57968463110397 | 4.26843458790854 |
| chr15 | 48740899 | 48740905 | - | 5.01980288436682 | 1.13699574206438 | 2.38645642073598 | 2.13421729395427 |
| chr15 | 48741583 | 48741601 | - | 16.0633692299738 | 3.41098722619315 | 2.38645642073598 | 1.06710864697714 |
| chr15 | 48878758 | 48878781 | + | 7.02772403811355 | 5.68497871032191 | 19.0916513658878 | 9.60397782279422 |
| chr15 | 48879027 | 48879046 | + | 10.0396057687336 | 11.3699574206438 | 4.77291284147196 | 5.33554323488568 |
| chr15 | 48879159 | 48879207 | + | 8.03168461498692 | 11.3699574206438 | 2.38645642073598 | 3.20132594093141 |
| chr15 | 48879229 | 48879235 | + | 4.01584230749346 | 3.41098722619315 | 1.19322821036799 | 1.06710864697714 |
| chr15 | 48879302 | 48879336 | + | 9.03564519186028 | 20.4659233571589 | 4.77291284147196 | 8.53686917581709 |
| chr15 | 48879573 | 48879578 | + | 1.00396057687336 | 9.09596593651506 | 1.19322821036799 | 2.13421729395427 |
| chr15 | 48879600 | 48879638 | + | 13.0514874993537 | 32.9728765198671 | 15.5119667347839 | 8.53686917581709 |
| chr15 | 48879751 | 48879780 | + | 25.0990144218341 | 29.5618892936739 | 14.3187385244159 | 16.006629704657 |
| chr15 | 48988666 | 48988678 | - | 4.01584230749346 | 4.54798296825753 | 1.19322821036799 | 1.06710864697714 |
| chr15 | 48989249 | 48989299 | - | 500.976327859809 | 502.552117992457 | 365.127832372605 | 335.072115150821 |
| chr15 | 48989414 | 48989461 | - | 4.01584230749346 | 9.09596593651506 | 3.57968463110397 | 9.60397782279422 |
| chr15 | 48989693 | 48989707 | - | 8.03168461498692 | 6.82197445238629 | 2.38645642073598 | 1.06710864697714 |
| chr15 | 48989751 | 48989757 | - | 10.0396057687336 | 13.6439489047726 | 4.77291284147196 | 2.13421729395427 |
| chr15 | 48989848 | 48989862 | - | 2.00792115374673 | 1.13699574206438 | 1.19322821036799 | 5.33554323488568 |
| chr15 | 48990718 | 48990740 | - | 8.03168461498692 | 5.68497871032191 | 4.77291284147196 | 2.13421729395427 |
| chr15 | 49125814 | 49125816 | - | 18.0712903837206 | 18.1919318730301 | 1.19322821036799 | 10.6710864697714 |
| chr15 | 49126237 | 49126239 | - | 1.00396057687336 | 3.41098722619315 | 2.38645642073598 | 2.13421729395427 |
| chr15 | 49126474 | 49126504 | - | 2.00792115374673 | 1.13699574206438 | 3.57968463110397 | 12.8053037637256 |
| chr15 | 49126570 | 49126576 | - | 4.01584230749346 | 2.27399148412876 | 10.7390538933119 | 4.26843458790854 |
| chr15 | 49126695 | 49126712 | - | 1.00396057687336 | 1.13699574206438 | 1.19322821036799 | 6.40265188186281 |
| chr15 | 49126770 | 49126791 | - | 1.00396057687336 | 1.13699574206438 | 4.77291284147196 | 1.06710864697714 |
| chr15 | 49127001 | 49127005 | - | 1.00396057687336 | 1.13699574206438 | 3.57968463110397 | 1.06710864697714 |
| chr15 | 49127133 | 49127152 | - | 1.00396057687336 | 4.54798296825753 | 13.1255103140479 | 8.53686917581709 |
| chr15 | 49127174 | 49127196 | - | 5.01980288436682 | 10.2329616785794 | 27.4442488384638 | 2.13421729395427 |
| chr15 | 49127225 | 49127239 | - | 4.01584230749346 | 2.27399148412876 | 8.35259747257593 | 5.33554323488568 |
| chr15 | 49127333 | 49127378 | - | 8.03168461498692 | 6.82197445238629 | 14.3187385244159 | 11.7381951167485 |
| chr15 | 49127394 | 49127403 | - | 8.03168461498692 | 6.82197445238629 | 1.19322821036799 | 1.06710864697714 |
| chr15 | 49127443 | 49127466 | - | 5.01980288436682 | 2.27399148412876 | 5.96614105183995 | 1.06710864697714 |
| chr15 | 49127657 | 49127678 | - | 3.01188173062009 | 1.13699574206438 | 13.1255103140479 | 16.006629704657 |
| chr15 | 49127822 | 49127980 | - | 226.89509037338 | 241.043097317649 | 174.211318713726 | 129.120146284233 |
| chr15 | 49328476 | 49328482 | + | 1.00396057687336 | 2.27399148412876 | 1.19322821036799 | 1.06710864697714 |
| chr15 | 49644049 | 49644052 | + | 7.02772403811355 | 3.41098722619315 | 3.57968463110397 | 1.06710864697714 |
| chr15 | 49644094 | 49644097 | + | 2.00792115374673 | 1.13699574206438 | 3.57968463110397 | 2.13421729395427 |
| chr15 | 49645016 | 49645033 | + | 4.01584230749346 | 12.5069531627082 | 2.38645642073598 | 5.33554323488568 |
| chr15 | 50236106 | 50236142 | + | 29.1148567293276 | 60.2607743294122 | 2.38645642073598 | 4.26843458790854 |
| chr15 | 50236175 | 50236263 | + | 43.1703048055547 | 50.0278126508328 | 1.19322821036799 | 4.26843458790854 |
| chr15 | 50236330 | 50236337 | + | 34.1346596136944 | 61.3977700714766 | 7.15936926220794 | 5.33554323488568 |
| chr15 | 50277459 | 50277466 | - | 2.00792115374673 | 5.68497871032191 | 10.7390538933119 | 6.40265188186281 |
| chr15 | 50355450 | 50355551 | + | 153.605968261625 | 164.864382599335 | 150.346754506367 | 129.120146284233 |
| chr15 | 50356099 | 50356115 | + | 9.03564519186028 | 10.2329616785794 | 3.57968463110397 | 2.13421729395427 |
| chr15 | 50356335 | 50356380 | + | 5.01980288436682 | 6.82197445238629 | 13.1255103140479 | 17.0737383516342 |
| chr15 | 50356610 | 50356633 | + | 5.01980288436682 | 6.82197445238629 | 10.7390538933119 | 7.46976052883995 |
| chr15 | 50356635 | 50356686 | + | 3.01188173062009 | 3.41098722619315 | 1.19322821036799 | 2.13421729395427 |
| chr15 | 50356757 | 50356836 | + | 21.0831721143407 | 13.6439489047726 | 15.5119667347839 | 10.6710864697714 |
| chr15 | 50356928 | 50357023 | + | 20.0792115374673 | 20.4659233571589 | 29.8307052591997 | 17.0737383516342 |
| chr15 | 50357045 | 50357157 | + | 31.1227778830743 | 32.9728765198671 | 58.4681823080315 | 44.8185631730397 |
| chr15 | 50358114 | 50358123 | + | 1.00396057687336 | 1.13699574206438 | 1.19322821036799 | 1.06710864697714 |
| chr15 | 50359595 | 50359605 | + | 7.02772403811355 | 4.54798296825753 | 8.35259747257593 | 8.53686917581709 |
| chr15 | 50359902 | 50359917 | + | 18.0712903837206 | 7.95897019445067 | 10.7390538933119 | 5.33554323488568 |
| chr15 | 50362621 | 50362654 | + | 1.00396057687336 | 5.68497871032191 | 7.15936926220794 | 1.06710864697714 |
| chr15 | 50362773 | 50362791 | + | 6.02376346124019 | 4.54798296825753 | 5.96614105183995 | 7.46976052883995 |
| chr15 | 50363085 | 50363106 | + | 1.00396057687336 | 2.27399148412876 | 4.77291284147196 | 3.20132594093141 |
| chr15 | 50365344 | 50365347 | + | 1.00396057687336 | 4.54798296825753 | 2.38645642073598 | 1.06710864697714 |
| chr15 | 50365951 | 50365989 | + | 5.01980288436682 | 3.41098722619315 | 5.96614105183995 | 2.13421729395427 |
| chr15 | 50367558 | 50367578 | + | 4.01584230749346 | 3.41098722619315 | 1.19322821036799 | 1.06710864697714 |
| chr15 | 50371036 | 50371093 | + | 8.03168461498692 | 21.6029190992233 | 8.35259747257593 | 4.26843458790854 |
| chr15 | 50371119 | 50371148 | + | 2.00792115374673 | 4.54798296825753 | 3.57968463110397 | 1.06710864697714 |
| chr15 | 50459009 | 50459021 | + | 5.01980288436682 | 5.68497871032191 | 7.15936926220794 | 2.13421729395427 |
| chr15 | 50459047 | 50459054 | + | 2.00792115374673 | 4.54798296825753 | 2.38645642073598 | 2.13421729395427 |
| chr15 | 50459095 | 50459125 | + | 6.02376346124019 | 5.68497871032191 | 2.38645642073598 | 1.06710864697714 |
| chr15 | 50471686 | 50471770 | + | 28.1108961524542 | 30.6988850357383 | 31.0239334695677 | 24.5434988804741 |
| chr15 | 50476849 | 50476871 | + | 1.00396057687336 | 1.13699574206438 | 2.38645642073598 | 6.40265188186281 |
| chr15 | 50476875 | 50476913 | + | 72.2851615348822 | 44.3428339405109 | 59.6614105183995 | 75.7647139353766 |
| chr15 | 50476941 | 50476981 | + | 99.3920971104631 | 81.8636934286355 | 34.6036181006717 | 56.5567582897882 |
| chr15 | 50477379 | 50477401 | + | 8.03168461498692 | 14.780944646837 | 13.1255103140479 | 7.46976052883995 |
| chr15 | 50477472 | 50477475 | + | 4.01584230749346 | 5.68497871032191 | 5.96614105183995 | 1.06710864697714 |
| chr15 | 50481479 | 50481524 | + | 11.043566345607 | 9.09596593651506 | 17.8984231555198 | 4.26843458790854 |
| chr15 | 50481552 | 50481614 | + | 159.629731722865 | 122.795540142953 | 193.302970079614 | 124.851711696325 |
| chr15 | 50481635 | 50481760 | + | 175.693100952839 | 191.015284666816 | 219.55399070771 | 182.47557863309 |
| chr15 | 50505138 | 50505141 | + | 2.00792115374673 | 1.13699574206438 | 1.19322821036799 | 1.06710864697714 |
| chr15 | 50560372 | 50560393 | - | 16.0633692299738 | 35.2468680039958 | 11.9322821036799 | 17.0737383516342 |
| chr15 | 50560490 | 50560492 | - | 8.03168461498692 | 17.0549361309657 | 11.9322821036799 | 12.8053037637256 |
| chr15 | 50560563 | 50560625 | - | 4.01584230749346 | 15.9179403889013 | 8.35259747257593 | 20.2750642925656 |
| chr15 | 50561051 | 50561056 | - | 6.02376346124019 | 1.13699574206438 | 2.38645642073598 | 4.26843458790854 |
| chr15 | 50561077 | 50561103 | - | 17.0673298068472 | 14.780944646837 | 15.5119667347839 | 18.1408469986113 |
| chr15 | 50561381 | 50561386 | - | 3.01188173062009 | 13.6439489047726 | 5.96614105183995 | 1.06710864697714 |
| chr15 | 50706117 | 50706128 | - | 72.2851615348822 | 70.4937360079917 | 31.0239334695677 | 14.9395210576799 |
| chr15 | 50707028 | 50707050 | - | 4.01584230749346 | 3.41098722619315 | 5.96614105183995 | 1.06710864697714 |
| chr15 | 51447833 | 51447837 | - | 2.00792115374673 | 4.54798296825753 | 1.19322821036799 | 3.20132594093141 |
| chr15 | 51723090 | 51723093 | - | 5.01980288436682 | 6.82197445238629 | 2.38645642073598 | 2.13421729395427 |
| chr15 | 51723361 | 51723363 | - | 2.00792115374673 | 12.5069531627082 | 2.38645642073598 | 5.33554323488568 |
| chr15 | 51723594 | 51723612 | - | 21.0831721143407 | 39.7948509722534 | 5.96614105183995 | 14.9395210576799 |
| chr15 | 51908819 | 51908929 | + | 212.839642297153 | 252.413054738293 | 103.810854302015 | 148.328101929822 |
| chr15 | 51909047 | 51909060 | + | 5.01980288436682 | 4.54798296825753 | 3.57968463110397 | 4.26843458790854 |
| chr15 | 51909116 | 51909125 | + | 6.02376346124019 | 4.54798296825753 | 3.57968463110397 | 2.13421729395427 |
| chr15 | 51909259 | 51909267 | + | 14.0554480762271 | 14.780944646837 | 2.38645642073598 | 11.7381951167485 |
| chr15 | 51909341 | 51909432 | + | 69.2732798042621 | 39.7948509722534 | 13.1255103140479 | 32.0132594093141 |
| chr15 | 51909476 | 51909549 | + | 47.1861471130481 | 38.657855230189 | 8.35259747257593 | 25.6106075274513 |
| chr15 | 51909612 | 51909641 | + | 20.0792115374673 | 20.4659233571589 | 10.7390538933119 | 5.33554323488568 |
| chr15 | 51909763 | 51909805 | + | 17.0673298068472 | 15.9179403889013 | 4.77291284147196 | 6.40265188186281 |
| chr15 | 51910252 | 51910259 | + | 2.00792115374673 | 1.13699574206438 | 1.19322821036799 | 3.20132594093141 |
| chr15 | 51911076 | 51911100 | + | 4.01584230749346 | 3.41098722619315 | 1.19322821036799 | 9.60397782279422 |
| chr15 | 51911445 | 51911448 | + | 3.01188173062009 | 2.27399148412876 | 1.19322821036799 | 1.06710864697714 |
| chr15 | 51912025 | 51912028 | + | 1.00396057687336 | 4.54798296825753 | 2.38645642073598 | 1.06710864697714 |
| chr15 | 51913780 | 51913783 | + | 6.02376346124019 | 4.54798296825753 | 4.77291284147196 | 8.53686917581709 |
| chr15 | 51938146 | 51938260 | - | 23.0910932680874 | 28.4248935516096 | 19.0916513658878 | 18.1408469986113 |
| chr15 | 52065511 | 52065513 | + | 1.00396057687336 | 4.54798296825753 | 5.96614105183995 | 3.20132594093141 |
| chr15 | 52066115 | 52066153 | + | 4.01584230749346 | 2.27399148412876 | 2.38645642073598 | 4.26843458790854 |
| chr15 | 52547068 | 52547087 | - | 4.01584230749346 | 3.41098722619315 | 1.19322821036799 | 2.13421729395427 |
| chr15 | 52547126 | 52547139 | - | 1.00396057687336 | 2.27399148412876 | 1.19322821036799 | 1.06710864697714 |
| chr15 | 52547381 | 52547406 | - | 43.1703048055547 | 59.1237785873479 | 33.4103898903037 | 46.952780466994 |
| chr15 | 52547586 | 52547670 | - | 21.0831721143407 | 23.876910583352 | 27.4442488384638 | 8.53686917581709 |
| chr15 | 52547705 | 52547787 | - | 26.1029749987075 | 20.4659233571589 | 14.3187385244159 | 23.476390233497 |
| chr15 | 52547846 | 52547888 | - | 39.1544624980612 | 48.8908169087684 | 21.4781077866238 | 17.0737383516342 |
| chr15 | 52547949 | 52548118 | - | 164.649534607232 | 144.398459242177 | 136.028015981951 | 77.8989312293309 |
| chr15 | 52548130 | 52548153 | - | 8.03168461498692 | 5.68497871032191 | 3.57968463110397 | 7.46976052883995 |
| chr15 | 52548198 | 52548221 | - | 11.043566345607 | 27.2878978095452 | 7.15936926220794 | 4.26843458790854 |
| chr15 | 52548515 | 52548529 | - | 6.02376346124019 | 3.41098722619315 | 27.4442488384638 | 28.8119334683827 |
| chr15 | 52548550 | 52548558 | - | 1.00396057687336 | 1.13699574206438 | 5.96614105183995 | 9.60397782279422 |
| chr15 | 52548580 | 52548620 | - | 17.0673298068472 | 10.2329616785794 | 19.0916513658878 | 14.9395210576799 |
| chr15 | 52548999 | 52549019 | - | 7.02772403811355 | 2.27399148412876 | 1.19322821036799 | 1.06710864697714 |
| chr15 | 52549381 | 52549395 | - | 24.0950538449607 | 18.1919318730301 | 14.3187385244159 | 8.53686917581709 |
| chr15 | 52549448 | 52549466 | - | 1.00396057687336 | 1.13699574206438 | 1.19322821036799 | 6.40265188186281 |
| chr15 | 52549640 | 52549661 | - | 4.01584230749346 | 4.54798296825753 | 2.38645642073598 | 2.13421729395427 |
| chr15 | 52549777 | 52549856 | - | 18.0712903837206 | 13.6439489047726 | 29.8307052591997 | 6.40265188186281 |
| chr15 | 52549882 | 52549906 | - | 11.043566345607 | 12.5069531627082 | 10.7390538933119 | 7.46976052883995 |
| chr15 | 52549946 | 52549972 | - | 21.0831721143407 | 14.780944646837 | 9.54582568294392 | 4.26843458790854 |
| chr15 | 52550018 | 52550022 | - | 4.01584230749346 | 4.54798296825753 | 3.57968463110397 | 2.13421729395427 |
| chr15 | 52550135 | 52550178 | - | 9.03564519186028 | 12.5069531627082 | 8.35259747257593 | 4.26843458790854 |
| chr15 | 52550222 | 52550231 | - | 10.0396057687336 | 6.82197445238629 | 4.77291284147196 | 7.46976052883995 |
| chr15 | 52550372 | 52550378 | - | 2.00792115374673 | 1.13699574206438 | 1.19322821036799 | 1.06710864697714 |
| chr15 | 52550478 | 52550480 | - | 1.00396057687336 | 2.27399148412876 | 2.38645642073598 | 2.13421729395427 |
| chr15 | 52550639 | 52550672 | - | 10.0396057687336 | 13.6439489047726 | 4.77291284147196 | 3.20132594093141 |
| chr15 | 52550702 | 52550808 | - | 9.03564519186028 | 12.5069531627082 | 11.9322821036799 | 13.8724124107028 |
| chr15 | 52551249 | 52551286 | - | 6.02376346124019 | 1.13699574206438 | 2.38645642073598 | 3.20132594093141 |
| chr15 | 52551458 | 52551468 | - | 16.0633692299738 | 21.6029190992233 | 1.19322821036799 | 9.60397782279422 |
| chr15 | 52551739 | 52551749 | - | 3.01188173062009 | 5.68497871032191 | 4.77291284147196 | 3.20132594093141 |
| chr15 | 55181358 | 55181384 | - | 10.0396057687336 | 18.1919318730301 | 15.5119667347839 | 5.33554323488568 |
| chr15 | 55181439 | 55181444 | - | 10.0396057687336 | 7.95897019445067 | 13.1255103140479 | 13.8724124107028 |
| chr15 | 55181519 | 55181523 | - | 2.00792115374673 | 1.13699574206438 | 8.35259747257593 | 4.26843458790854 |
| chr15 | 55181721 | 55181777 | - | 6.02376346124019 | 20.4659233571589 | 25.0577924177278 | 40.5501285851312 |
| chr15 | 55181856 | 55181863 | - | 1.00396057687336 | 2.27399148412876 | 7.15936926220794 | 4.26843458790854 |
| chr15 | 55181886 | 55181911 | - | 9.03564519186028 | 10.2329616785794 | 7.15936926220794 | 5.33554323488568 |
| chr15 | 55204133 | 55204134 | - | 1.00396057687336 | 1.13699574206438 | 4.77291284147196 | 3.20132594093141 |
| chr15 | 55205418 | 55205439 | - | 5.01980288436682 | 2.27399148412876 | 5.96614105183995 | 13.8724124107028 |
| chr15 | 55205464 | 55205493 | - | 2.00792115374673 | 3.41098722619315 | 8.35259747257593 | 5.33554323488568 |
| chr15 | 55317292 | 55317293 | - | 1.00396057687336 | 1.13699574206438 | 1.19322821036799 | 1.06710864697714 |
| chr15 | 55317394 | 55317397 | - | 7.02772403811355 | 5.68497871032191 | 1.19322821036799 | 3.20132594093141 |
| chr15 | 55356729 | 55356733 | - | 4.01584230749346 | 1.13699574206438 | 4.77291284147196 | 1.06710864697714 |
| chr15 | 55356831 | 55356839 | - | 2.00792115374673 | 1.13699574206438 | 3.57968463110397 | 6.40265188186281 |
| chr15 | 55356902 | 55356910 | - | 2.00792115374673 | 6.82197445238629 | 7.15936926220794 | 5.33554323488568 |
| chr15 | 55357016 | 55357020 | - | 9.03564519186028 | 4.54798296825753 | 11.9322821036799 | 13.8724124107028 |
| chr15 | 55357400 | 55357401 | - | 3.01188173062009 | 2.27399148412876 | 1.19322821036799 | 2.13421729395427 |
| chr15 | 55358980 | 55359026 | - | 21.0831721143407 | 35.2468680039958 | 14.3187385244159 | 9.60397782279422 |
| chr15 | 55359430 | 55359491 | - | 22.087132691214 | 38.657855230189 | 5.96614105183995 | 2.13421729395427 |
| chr15 | 56090677 | 56090716 | - | 5.01980288436682 | 3.41098722619315 | 5.96614105183995 | 5.33554323488568 |
| chr15 | 56090768 | 56090821 | - | 13.0514874993537 | 17.0549361309657 | 59.6614105183995 | 12.8053037637256 |
| chr15 | 56090933 | 56090961 | - | 3.01188173062009 | 4.54798296825753 | 3.57968463110397 | 3.20132594093141 |
| chr15 | 56090989 | 56091025 | - | 6.02376346124019 | 7.95897019445067 | 7.15936926220794 | 11.7381951167485 |
| chr15 | 56091059 | 56091105 | - | 25.0990144218341 | 12.5069531627082 | 19.0916513658878 | 12.8053037637256 |
| chr15 | 56091155 | 56091160 | - | 4.01584230749346 | 7.95897019445067 | 8.35259747257593 | 13.8724124107028 |
| chr15 | 56091195 | 56091214 | - | 4.01584230749346 | 7.95897019445067 | 1.19322821036799 | 4.26843458790854 |
| chr15 | 56091766 | 56091771 | - | 4.01584230749346 | 3.41098722619315 | 3.57968463110397 | 1.06710864697714 |
| chr15 | 56630433 | 56630437 | - | 3.01188173062009 | 3.41098722619315 | 13.1255103140479 | 7.46976052883995 |
| chr15 | 56630756 | 56630769 | - | 2.00792115374673 | 7.95897019445067 | 10.7390538933119 | 6.40265188186281 |
| chr15 | 56630844 | 56630846 | - | 2.00792115374673 | 1.13699574206438 | 3.57968463110397 | 3.20132594093141 |
| chr15 | 56630989 | 56631016 | - | 1.00396057687336 | 1.13699574206438 | 5.96614105183995 | 2.13421729395427 |
| chr15 | 56631065 | 56631066 | - | 1.00396057687336 | 2.27399148412876 | 10.7390538933119 | 3.20132594093141 |
| chr15 | 56699995 | 56700031 | - | 5.01980288436682 | 4.54798296825753 | 1.19322821036799 | 1.06710864697714 |
| chr15 | 56700067 | 56700112 | - | 52.205949997415 | 65.9457530397342 | 13.1255103140479 | 16.006629704657 |
| chr15 | 56700749 | 56700754 | - | 2.00792115374673 | 7.95897019445067 | 4.77291284147196 | 6.40265188186281 |
| chr15 | 57288254 | 57288313 | + | 6.02376346124019 | 28.4248935516096 | 32.2171616799357 | 18.1408469986113 |
| chr15 | 57288414 | 57288423 | + | 2.00792115374673 | 5.68497871032191 | 3.57968463110397 | 1.06710864697714 |
| chr15 | 57288465 | 57288493 | + | 3.01188173062009 | 11.3699574206438 | 1.19322821036799 | 2.13421729395427 |
| chr15 | 57714797 | 57714800 | + | 1.00396057687336 | 1.13699574206438 | 1.19322821036799 | 1.06710864697714 |
| chr15 | 58590005 | 58590020 | - | 1.00396057687336 | 2.27399148412876 | 4.77291284147196 | 1.06710864697714 |
| chr15 | 58595785 | 58595830 | - | 27.1069355755808 | 31.8358807778027 | 3.57968463110397 | 19.2079556455884 |
| chr15 | 58596120 | 58596189 | - | 11.043566345607 | 2.27399148412876 | 16.7051949451519 | 18.1408469986113 |
| chr15 | 58596223 | 58596238 | - | 1.00396057687336 | 1.13699574206438 | 5.96614105183995 | 1.06710864697714 |
| chr15 | 58596254 | 58596261 | - | 5.01980288436682 | 5.68497871032191 | 2.38645642073598 | 1.06710864697714 |
| chr15 | 58596269 | 58596285 | - | 1.00396057687336 | 1.13699574206438 | 1.19322821036799 | 6.40265188186281 |
| chr15 | 58596424 | 58596479 | - | 3.01188173062009 | 4.54798296825753 | 3.57968463110397 | 4.26843458790854 |
| chr15 | 58596717 | 58596747 | - | 4.01584230749346 | 2.27399148412876 | 8.35259747257593 | 6.40265188186281 |
| chr15 | 58596857 | 58596880 | - | 5.01980288436682 | 6.82197445238629 | 1.19322821036799 | 5.33554323488568 |
| chr15 | 58597315 | 58597358 | - | 8.03168461498692 | 6.82197445238629 | 2.38645642073598 | 3.20132594093141 |
| chr15 | 58854736 | 58854742 | + | 1.00396057687336 | 1.13699574206438 | 1.19322821036799 | 4.26843458790854 |
| chr15 | 58858154 | 58858165 | + | 1.00396057687336 | 6.82197445238629 | 1.19322821036799 | 1.06710864697714 |
| chr15 | 58879092 | 58879103 | - | 7.02772403811355 | 30.6988850357383 | 11.9322821036799 | 7.46976052883995 |
| chr15 | 58879491 | 58879511 | - | 10.0396057687336 | 15.9179403889013 | 10.7390538933119 | 7.46976052883995 |
| chr15 | 58879648 | 58879663 | - | 8.03168461498692 | 4.54798296825753 | 1.19322821036799 | 3.20132594093141 |
| chr15 | 58879842 | 58879846 | - | 8.03168461498692 | 9.09596593651506 | 3.57968463110397 | 5.33554323488568 |
| chr15 | 58879902 | 58879939 | - | 9.03564519186028 | 11.3699574206438 | 7.15936926220794 | 7.46976052883995 |
| chr15 | 58898802 | 58898844 | - | 16.0633692299738 | 5.68497871032191 | 16.7051949451519 | 23.476390233497 |
| chr15 | 58899471 | 58899684 | - | 86.3406096111093 | 103.466612527859 | 102.617626091647 | 100.308212815851 |
| chr15 | 58899722 | 58899785 | - | 15.0594086531005 | 27.2878978095452 | 40.5697591525116 | 22.4092815865198 |
| chr15 | 58899809 | 58899819 | - | 3.01188173062009 | 6.82197445238629 | 11.9322821036799 | 5.33554323488568 |
| chr15 | 58899840 | 58899885 | - | 12.0475269224804 | 11.3699574206438 | 31.0239334695677 | 11.7381951167485 |
| chr15 | 58901258 | 58901288 | - | 19.0752509605939 | 9.09596593651506 | 17.8984231555198 | 10.6710864697714 |
| chr15 | 58912583 | 58912613 | - | 15.0594086531005 | 10.2329616785794 | 21.4781077866238 | 13.8724124107028 |
| chr15 | 58913498 | 58913699 | - | 52.205949997415 | 61.3977700714766 | 121.709277457535 | 82.1673658172394 |
| chr15 | 58916934 | 58917002 | - | 34.1346596136944 | 55.7127913611547 | 71.5936926220794 | 38.4159112911769 |
| chr15 | 59094969 | 59094970 | + | 2.00792115374673 | 1.13699574206438 | 1.19322821036799 | 2.13421729395427 |
| chr15 | 59095184 | 59095237 | + | 9.03564519186028 | 7.95897019445067 | 20.2848795762558 | 16.006629704657 |
| chr15 | 59095485 | 59095506 | + | 22.087132691214 | 18.1919318730301 | 9.54582568294392 | 8.53686917581709 |
| chr15 | 59096852 | 59096873 | + | 7.02772403811355 | 6.82197445238629 | 2.38645642073598 | 5.33554323488568 |
| chr15 | 59096910 | 59096913 | + | 6.02376346124019 | 1.13699574206438 | 1.19322821036799 | 1.06710864697714 |
| chr15 | 59124869 | 59124913 | + | 187.740627875319 | 147.80944646837 | 116.936364616063 | 171.804492163319 |
| chr15 | 59124961 | 59125026 | + | 150.594086531005 | 173.96034853585 | 101.424397881279 | 105.643756050736 |
| chr15 | 59638895 | 59638948 | - | 23.0910932680874 | 45.4798296825753 | 27.4442488384638 | 30.9461507623369 |
| chr15 | 59638980 | 59638996 | - | 2.00792115374673 | 3.41098722619315 | 2.38645642073598 | 3.20132594093141 |
| chr15 | 59639073 | 59639079 | - | 3.01188173062009 | 2.27399148412876 | 1.19322821036799 | 2.13421729395427 |
| chr15 | 59660933 | 59660935 | - | 10.0396057687336 | 4.54798296825753 | 1.19322821036799 | 1.06710864697714 |
| chr15 | 59661058 | 59661059 | - | 1.00396057687336 | 1.13699574206438 | 3.57968463110397 | 2.13421729395427 |
| chr15 | 59661545 | 59661550 | - | 10.0396057687336 | 7.95897019445067 | 11.9322821036799 | 7.46976052883995 |
| chr15 | 59661716 | 59661726 | - | 4.01584230749346 | 5.68497871032191 | 7.15936926220794 | 1.06710864697714 |
| chr15 | 59661813 | 59661828 | - | 16.0633692299738 | 12.5069531627082 | 17.8984231555198 | 3.20132594093141 |
| chr15 | 59661914 | 59661939 | - | 10.0396057687336 | 5.68497871032191 | 3.57968463110397 | 3.20132594093141 |
| chr15 | 59662168 | 59662190 | - | 20.0792115374673 | 13.6439489047726 | 3.57968463110397 | 3.20132594093141 |
| chr15 | 59663104 | 59663107 | - | 2.00792115374673 | 1.13699574206438 | 1.19322821036799 | 1.06710864697714 |
| chr15 | 59663128 | 59663132 | - | 4.01584230749346 | 11.3699574206438 | 4.77291284147196 | 4.26843458790854 |
| chr15 | 59663246 | 59663247 | - | 1.00396057687336 | 1.13699574206438 | 1.19322821036799 | 2.13421729395427 |
| chr15 | 59663280 | 59663353 | - | 4.01584230749346 | 2.27399148412876 | 2.38645642073598 | 4.26843458790854 |
| chr15 | 59664000 | 59664040 | - | 30.1188173062009 | 38.657855230189 | 32.2171616799357 | 27.7448248214055 |
| chr15 | 60347420 | 60347446 | - | 38.1505019211879 | 57.9867828452835 | 17.8984231555198 | 22.4092815865198 |
| chr15 | 60347469 | 60347560 | - | 497.964446129189 | 557.127913611547 | 114.549908195327 | 197.41509969077 |
| chr15 | 60423264 | 60423299 | - | 17.0673298068472 | 7.95897019445067 | 9.54582568294392 | 9.60397782279422 |
| chr15 | 60423340 | 60423351 | - | 2.00792115374673 | 3.41098722619315 | 10.7390538933119 | 1.06710864697714 |
| chr15 | 60423431 | 60423591 | - | 140.554480762271 | 194.426271893009 | 169.438405872254 | 160.06629704657 |
| chr15 | 60423621 | 60423634 | - | 4.01584230749346 | 4.54798296825753 | 8.35259747257593 | 7.46976052883995 |
| chr15 | 60451296 | 60451298 | - | 2.00792115374673 | 7.95897019445067 | 2.38645642073598 | 3.20132594093141 |
| chr15 | 60452414 | 60452422 | - | 9.03564519186028 | 15.9179403889013 | 2.38645642073598 | 4.26843458790854 |
| chr15 | 60479192 | 60479323 | + | 47.1861471130481 | 73.9047232341848 | 99.0379414605431 | 88.5700176991023 |
| chr15 | 61852922 | 61852925 | - | 2.00792115374673 | 1.13699574206438 | 1.19322821036799 | 2.13421729395427 |
| chr15 | 61852954 | 61852973 | - | 2.00792115374673 | 4.54798296825753 | 2.38645642073598 | 8.53686917581709 |
| chr15 | 63071677 | 63071693 | + | 3.01188173062009 | 1.13699574206438 | 1.19322821036799 | 1.06710864697714 |
| chr15 | 63071732 | 63071771 | + | 8.03168461498692 | 12.5069531627082 | 3.57968463110397 | 2.13421729395427 |
| chr15 | 63071798 | 63071848 | + | 15.0594086531005 | 21.6029190992233 | 3.57968463110397 | 5.33554323488568 |
| chr15 | 63141972 | 63141993 | + | 3.01188173062009 | 2.27399148412876 | 1.19322821036799 | 1.06710864697714 |
| chr15 | 63153547 | 63153582 | - | 7.02772403811355 | 10.2329616785794 | 4.77291284147196 | 3.20132594093141 |
| chr15 | 63153882 | 63153931 | - | 47.1861471130481 | 36.3838637460602 | 33.4103898903037 | 54.4225409958339 |
| chr15 | 63153963 | 63154063 | - | 22.087132691214 | 28.4248935516096 | 32.2171616799357 | 14.9395210576799 |
| chr15 | 63265693 | 63265698 | + | 5.01980288436682 | 3.41098722619315 | 3.57968463110397 | 4.26843458790854 |
| chr15 | 63265729 | 63265734 | + | 1.00396057687336 | 5.68497871032191 | 8.35259747257593 | 6.40265188186281 |
| chr15 | 63266633 | 63266644 | + | 4.01584230749346 | 6.82197445238629 | 11.9322821036799 | 13.8724124107028 |
| chr15 | 63267623 | 63267647 | + | 1.00396057687336 | 1.13699574206438 | 1.19322821036799 | 1.06710864697714 |
| chr15 | 63590748 | 63590828 | + | 57.2257528817818 | 104.603608269923 | 72.7869208324474 | 27.7448248214055 |
| chr15 | 63590950 | 63590997 | + | 24.0950538449607 | 26.1509020674808 | 36.9900745214077 | 16.006629704657 |
| chr15 | 63591018 | 63591088 | + | 14.0554480762271 | 38.657855230189 | 26.2510206280958 | 16.006629704657 |
| chr15 | 63591125 | 63591137 | + | 23.0910932680874 | 42.0688424563821 | 25.0577924177278 | 27.7448248214055 |
| chr15 | 63591231 | 63591291 | + | 25.0990144218341 | 27.2878978095452 | 21.4781077866238 | 28.8119334683827 |
| chr15 | 63591313 | 63591375 | + | 55.217831728035 | 102.329616785794 | 63.2410951495034 | 36.2816939972226 |
| chr15 | 63591396 | 63591400 | + | 5.01980288436682 | 1.13699574206438 | 2.38645642073598 | 3.20132594093141 |
| chr15 | 63593333 | 63593369 | + | 1.00396057687336 | 6.82197445238629 | 4.77291284147196 | 2.13421729395427 |
| chr15 | 63593422 | 63593461 | + | 6.02376346124019 | 7.95897019445067 | 4.77291284147196 | 3.20132594093141 |
| chr15 | 63593506 | 63593576 | + | 16.0633692299738 | 22.7399148412876 | 36.9900745214077 | 6.40265188186281 |
| chr15 | 63593603 | 63593637 | + | 3.01188173062009 | 3.41098722619315 | 2.38645642073598 | 2.13421729395427 |
| chr15 | 63608784 | 63608791 | - | 4.01584230749346 | 1.13699574206438 | 2.38645642073598 | 2.13421729395427 |
| chr15 | 64072611 | 64072642 | - | 21.0831721143407 | 21.6029190992233 | 15.5119667347839 | 37.3488026441998 |
| chr15 | 64072664 | 64072689 | - | 13.0514874993537 | 18.1919318730301 | 10.7390538933119 | 10.6710864697714 |
| chr15 | 64072785 | 64072864 | - | 30.1188173062009 | 39.7948509722534 | 23.8645642073598 | 37.3488026441998 |
| chr15 | 64072904 | 64073024 | - | 53.2099105742883 | 63.6717615556054 | 13.1255103140479 | 58.6909755837425 |
| chr15 | 64075077 | 64075122 | - | 17.0673298068472 | 5.68497871032191 | 17.8984231555198 | 9.60397782279422 |
| chr15 | 64075491 | 64075517 | - | 8.03168461498692 | 10.2329616785794 | 5.96614105183995 | 14.9395210576799 |
| chr15 | 64137851 | 64137875 | + | 24.0950538449607 | 26.1509020674808 | 15.5119667347839 | 19.2079556455884 |
| chr15 | 64137916 | 64137939 | + | 6.02376346124019 | 3.41098722619315 | 3.57968463110397 | 7.46976052883995 |
| chr15 | 64137960 | 64137984 | + | 16.0633692299738 | 36.3838637460602 | 7.15936926220794 | 9.60397782279422 |
| chr15 | 64138304 | 64138311 | + | 9.03564519186028 | 5.68497871032191 | 8.35259747257593 | 8.53686917581709 |
| chr15 | 64155852 | 64155898 | - | 8.03168461498692 | 7.95897019445067 | 4.77291284147196 | 9.60397782279422 |
| chr15 | 64155920 | 64155964 | - | 16.0633692299738 | 4.54798296825753 | 10.7390538933119 | 14.9395210576799 |
| chr15 | 64155997 | 64156148 | - | 19.0752509605939 | 10.2329616785794 | 23.8645642073598 | 23.476390233497 |
| chr15 | 64156076 | 64156110 | + | 7.02772403811355 | 3.41098722619315 | 5.96614105183995 | 4.26843458790854 |
| chr15 | 64156863 | 64156889 | + | 2.00792115374673 | 1.13699574206438 | 1.19322821036799 | 1.06710864697714 |
| chr15 | 64165788 | 64165798 | - | 6.02376346124019 | 11.3699574206438 | 3.57968463110397 | 12.8053037637256 |
| chr15 | 64166539 | 64166566 | - | 6.02376346124019 | 4.54798296825753 | 5.96614105183995 | 8.53686917581709 |
| chr15 | 64166598 | 64166615 | - | 3.01188173062009 | 18.1919318730301 | 11.9322821036799 | 8.53686917581709 |
| chr15 | 64167302 | 64167305 | - | 9.03564519186028 | 15.9179403889013 | 7.15936926220794 | 6.40265188186281 |
| chr15 | 64167363 | 64167398 | - | 10.0396057687336 | 13.6439489047726 | 3.57968463110397 | 10.6710864697714 |
| chr15 | 64167423 | 64167433 | - | 2.00792115374673 | 4.54798296825753 | 1.19322821036799 | 1.06710864697714 |
| chr15 | 64364424 | 64364427 | - | 7.02772403811355 | 10.2329616785794 | 2.38645642073598 | 2.13421729395427 |
| chr15 | 64364517 | 64364536 | - | 7.02772403811355 | 1.13699574206438 | 2.38645642073598 | 3.20132594093141 |
| chr15 | 64365616 | 64365635 | - | 27.1069355755808 | 34.1098722619315 | 3.57968463110397 | 10.6710864697714 |
| chr15 | 64365657 | 64366081 | - | 2030.00828643794 | 2127.31903340246 | 1310.16457498405 | 665.875795713733 |
| chr15 | 64455252 | 64455257 | + | 13.0514874993537 | 17.0549361309657 | 11.9322821036799 | 9.60397782279422 |
| chr15 | 64682699 | 64682734 | + | 3.01188173062009 | 1.13699574206438 | 1.19322821036799 | 3.20132594093141 |
| chr15 | 64682897 | 64682918 | + | 2.00792115374673 | 2.27399148412876 | 2.38645642073598 | 2.13421729395427 |
| chr15 | 64683123 | 64683147 | + | 7.02772403811355 | 11.3699574206438 | 8.35259747257593 | 2.13421729395427 |
| chr15 | 64684455 | 64684456 | + | 2.00792115374673 | 1.13699574206438 | 2.38645642073598 | 1.06710864697714 |
| chr15 | 64685013 | 64685063 | + | 7.02772403811355 | 25.0139063254164 | 26.2510206280958 | 27.7448248214055 |
| chr15 | 64685833 | 64685840 | + | 2.00792115374673 | 3.41098722619315 | 2.38645642073598 | 9.60397782279422 |
| chr15 | 64687600 | 64687665 | - | 79.3128855729958 | 87.5486721389574 | 60.8546387287675 | 89.6371263460794 |
| chr15 | 64687724 | 64687747 | - | 8.03168461498692 | 3.41098722619315 | 3.57968463110397 | 2.13421729395427 |
| chr15 | 64688350 | 64688423 | - | 32.1267384599477 | 11.3699574206438 | 19.0916513658878 | 30.9461507623369 |
| chr15 | 64688559 | 64688600 | - | 23.0910932680874 | 12.5069531627082 | 23.8645642073598 | 16.006629704657 |
| chr15 | 64688626 | 64688636 | - | 3.01188173062009 | 3.41098722619315 | 3.57968463110397 | 3.20132594093141 |
| chr15 | 64688658 | 64688672 | - | 13.0514874993537 | 4.54798296825753 | 4.77291284147196 | 7.46976052883995 |
| chr15 | 64815692 | 64815717 | - | 8.03168461498692 | 5.68497871032191 | 1.19322821036799 | 2.13421729395427 |
| chr15 | 64963050 | 64963084 | - | 9.03564519186028 | 12.5069531627082 | 4.77291284147196 | 5.33554323488568 |
| chr15 | 64963156 | 64963159 | - | 7.02772403811355 | 4.54798296825753 | 10.7390538933119 | 7.46976052883995 |
| chr15 | 64963292 | 64963310 | - | 29.1148567293276 | 23.876910583352 | 14.3187385244159 | 18.1408469986113 |
| chr15 | 64963335 | 64963337 | - | 5.01980288436682 | 1.13699574206438 | 1.19322821036799 | 3.20132594093141 |
| chr15 | 64963362 | 64963370 | - | 10.0396057687336 | 6.82197445238629 | 1.19322821036799 | 8.53686917581709 |
| chr15 | 64963438 | 64963466 | - | 11.043566345607 | 10.2329616785794 | 3.57968463110397 | 14.9395210576799 |
| chr15 | 65117485 | 65117498 | - | 2.00792115374673 | 10.2329616785794 | 4.77291284147196 | 2.13421729395427 |
| chr15 | 65150475 | 65150477 | - | 4.01584230749346 | 12.5069531627082 | 2.38645642073598 | 2.13421729395427 |
| chr15 | 65150691 | 65150741 | - | 2.00792115374673 | 3.41098722619315 | 5.96614105183995 | 8.53686917581709 |
| chr15 | 65258302 | 65258311 | - | 10.0396057687336 | 10.2329616785794 | 8.35259747257593 | 10.6710864697714 |
| chr15 | 65258984 | 65259016 | - | 3.01188173062009 | 7.95897019445067 | 9.54582568294392 | 5.33554323488568 |
| chr15 | 65259185 | 65259189 | - | 15.0594086531005 | 27.2878978095452 | 8.35259747257593 | 5.33554323488568 |
| chr15 | 65442532 | 65442603 | - | 30.1188173062009 | 36.3838637460602 | 39.3765309421437 | 9.60397782279422 |
| chr15 | 65442630 | 65442632 | - | 4.01584230749346 | 2.27399148412876 | 5.96614105183995 | 1.06710864697714 |
| chr15 | 65442659 | 65442671 | - | 3.01188173062009 | 4.54798296825753 | 4.77291284147196 | 1.06710864697714 |
| chr15 | 65442729 | 65442740 | - | 7.02772403811355 | 4.54798296825753 | 13.1255103140479 | 6.40265188186281 |
| chr15 | 65442767 | 65442800 | - | 107.42378172545 | 62.534765813541 | 64.4343233598714 | 44.8185631730397 |
| chr15 | 65442877 | 65442899 | - | 43.1703048055547 | 21.6029190992233 | 11.9322821036799 | 29.8790421153598 |
| chr15 | 65442960 | 65443030 | - | 24.0950538449607 | 29.5618892936739 | 19.0916513658878 | 13.8724124107028 |
| chr15 | 65443242 | 65443264 | - | 2.00792115374673 | 3.41098722619315 | 7.15936926220794 | 1.06710864697714 |
| chr15 | 65443566 | 65443585 | - | 12.0475269224804 | 1.13699574206438 | 3.57968463110397 | 2.13421729395427 |
| chr15 | 65446094 | 65446113 | - | 1.00396057687336 | 4.54798296825753 | 4.77291284147196 | 3.20132594093141 |
| chr15 | 65576492 | 65576499 | + | 8.03168461498692 | 12.5069531627082 | 4.77291284147196 | 14.9395210576799 |
| chr15 | 65576788 | 65576927 | + | 63.249516343022 | 100.055625301666 | 72.7869208324474 | 44.8185631730397 |
| chr15 | 65576974 | 65577023 | + | 59.2336740355285 | 73.9047232341848 | 35.7968463110397 | 35.2145853502455 |
| chr15 | 65577177 | 65577198 | + | 13.0514874993537 | 18.1919318730301 | 13.1255103140479 | 21.3421729395427 |
| chr15 | 65577320 | 65577337 | + | 23.0910932680874 | 18.1919318730301 | 23.8645642073598 | 17.0737383516342 |
| chr15 | 65577372 | 65577407 | + | 9.03564519186028 | 9.09596593651506 | 5.96614105183995 | 16.006629704657 |
| chr15 | 65577474 | 65577519 | + | 12.0475269224804 | 7.95897019445067 | 28.6374770488317 | 30.9461507623369 |
| chr15 | 65577574 | 65577599 | + | 3.01188173062009 | 2.27399148412876 | 3.57968463110397 | 4.26843458790854 |
| chr15 | 65577639 | 65577707 | + | 24.0950538449607 | 20.4659233571589 | 15.5119667347839 | 12.8053037637256 |
| chr15 | 65577793 | 65577805 | + | 10.0396057687336 | 3.41098722619315 | 1.19322821036799 | 2.13421729395427 |
| chr15 | 65578010 | 65578013 | + | 17.0673298068472 | 28.4248935516096 | 15.5119667347839 | 25.6106075274513 |
| chr15 | 65578091 | 65578101 | + | 4.01584230749346 | 2.27399148412876 | 3.57968463110397 | 3.20132594093141 |
| chr15 | 65578196 | 65578321 | + | 25.0990144218341 | 40.9318467143178 | 14.3187385244159 | 11.7381951167485 |
| chr15 | 65579221 | 65579257 | - | 19.0752509605939 | 12.5069531627082 | 21.4781077866238 | 10.6710864697714 |
| chr15 | 65579296 | 65579299 | - | 2.00792115374673 | 1.13699574206438 | 3.57968463110397 | 4.26843458790854 |
| chr15 | 65579393 | 65579448 | - | 10.0396057687336 | 10.2329616785794 | 16.7051949451519 | 18.1408469986113 |
| chr15 | 65661593 | 65661595 | - | 2.00792115374673 | 2.27399148412876 | 1.19322821036799 | 3.20132594093141 |
| chr15 | 65661673 | 65661678 | - | 7.02772403811355 | 6.82197445238629 | 2.38645642073598 | 5.33554323488568 |
| chr15 | 65887731 | 65887776 | + | 9.03564519186028 | 12.5069531627082 | 7.15936926220794 | 8.53686917581709 |
| chr15 | 65887807 | 65887929 | + | 42.1663442286813 | 48.8908169087684 | 50.1155848354556 | 91.7713436400337 |
| chr15 | 65888010 | 65888066 | + | 14.0554480762271 | 10.2329616785794 | 8.35259747257593 | 14.9395210576799 |
| chr15 | 65888425 | 65888433 | + | 7.02772403811355 | 2.27399148412876 | 3.57968463110397 | 3.20132594093141 |
| chr15 | 65889155 | 65889160 | + | 3.01188173062009 | 3.41098722619315 | 1.19322821036799 | 3.20132594093141 |
| chr15 | 65889234 | 65889269 | + | 18.0712903837206 | 12.5069531627082 | 8.35259747257593 | 5.33554323488568 |
| chr15 | 65891545 | 65891546 | + | 2.00792115374673 | 1.13699574206438 | 1.19322821036799 | 1.06710864697714 |
| chr15 | 65891735 | 65891736 | + | 1.00396057687336 | 1.13699574206438 | 1.19322821036799 | 2.13421729395427 |
| chr15 | 65891848 | 65891892 | + | 32.1267384599477 | 39.7948509722534 | 35.7968463110397 | 54.4225409958339 |
| chr15 | 66490607 | 66490618 | + | 4.01584230749346 | 3.41098722619315 | 3.57968463110397 | 2.13421729395427 |
| chr15 | 66490829 | 66490904 | + | 12.0475269224804 | 17.0549361309657 | 19.0916513658878 | 17.0737383516342 |
| chr15 | 66491009 | 66491035 | + | 10.0396057687336 | 19.3289276150945 | 10.7390538933119 | 18.1408469986113 |
| chr15 | 66491064 | 66491112 | + | 3.01188173062009 | 1.13699574206438 | 5.96614105183995 | 6.40265188186281 |
| chr15 | 66491179 | 66491243 | + | 44.174265382428 | 29.5618892936739 | 14.3187385244159 | 27.7448248214055 |
| chr15 | 66491277 | 66491300 | + | 6.02376346124019 | 17.0549361309657 | 17.8984231555198 | 6.40265188186281 |
| chr15 | 66493901 | 66493928 | - | 1.00396057687336 | 3.41098722619315 | 2.38645642073598 | 2.13421729395427 |
| chr15 | 66494185 | 66494204 | - | 2.00792115374673 | 5.68497871032191 | 1.19322821036799 | 5.33554323488568 |
| chr15 | 66494302 | 66494308 | - | 5.01980288436682 | 3.41098722619315 | 1.19322821036799 | 2.13421729395427 |
| chr15 | 66494415 | 66494435 | - | 9.03564519186028 | 12.5069531627082 | 5.96614105183995 | 9.60397782279422 |
| chr15 | 66499173 | 66499176 | - | 1.00396057687336 | 5.68497871032191 | 2.38645642073598 | 4.26843458790854 |
| chr15 | 66548547 | 66548548 | + | 8.03168461498692 | 1.13699574206438 | 3.57968463110397 | 2.13421729395427 |
| chr15 | 66548944 | 66548947 | + | 2.00792115374673 | 11.3699574206438 | 2.38645642073598 | 2.13421729395427 |
| chr15 | 66549318 | 66549322 | + | 17.0673298068472 | 26.1509020674808 | 5.96614105183995 | 14.9395210576799 |
| chr15 | 66549410 | 66549426 | + | 5.01980288436682 | 6.82197445238629 | 1.19322821036799 | 3.20132594093141 |
| chr15 | 66549453 | 66549465 | + | 8.03168461498692 | 31.8358807778027 | 4.77291284147196 | 6.40265188186281 |
| chr15 | 66781599 | 66781630 | + | 22.087132691214 | 20.4659233571589 | 1.19322821036799 | 3.20132594093141 |
| chr15 | 66781748 | 66781784 | + | 14.0554480762271 | 11.3699574206438 | 4.77291284147196 | 2.13421729395427 |
| chr15 | 67191155 | 67191210 | + | 17.0673298068472 | 17.0549361309657 | 5.96614105183995 | 26.6777161744284 |
| chr15 | 67191340 | 67191435 | + | 6.02376346124019 | 6.82197445238629 | 2.38645642073598 | 5.33554323488568 |
| chr15 | 67191624 | 67191649 | + | 2.00792115374673 | 3.41098722619315 | 1.19322821036799 | 1.06710864697714 |
| chr15 | 67191838 | 67191863 | + | 1.00396057687336 | 2.27399148412876 | 1.19322821036799 | 1.06710864697714 |
| chr15 | 67193978 | 67193985 | + | 2.00792115374673 | 4.54798296825753 | 2.38645642073598 | 2.13421729395427 |
| chr15 | 67194327 | 67194411 | + | 5.01980288436682 | 10.2329616785794 | 8.35259747257593 | 8.53686917581709 |
| chr15 | 67194807 | 67194854 | + | 22.087132691214 | 28.4248935516096 | 29.8307052591997 | 44.8185631730397 |
| chr15 | 67194932 | 67194941 | + | 2.00792115374673 | 4.54798296825753 | 1.19322821036799 | 5.33554323488568 |
| chr15 | 67201695 | 67201764 | - | 33.130699036821 | 55.7127913611547 | 35.7968463110397 | 38.4159112911769 |
| chr15 | 67201800 | 67201820 | - | 7.02772403811355 | 13.6439489047726 | 4.77291284147196 | 9.60397782279422 |
| chr15 | 67201845 | 67201906 | - | 64.2534769198953 | 65.9457530397342 | 44.1494437836156 | 43.7514545260626 |
| chr15 | 67202071 | 67202270 | - | 66.2613980736421 | 93.2336508492793 | 84.7192029361272 | 67.2278447595595 |
| chr15 | 67202297 | 67202442 | - | 40.1584230749346 | 52.3018041349616 | 46.5359002043516 | 71.4962793474681 |
| chr15 | 67202491 | 67202514 | - | 6.02376346124019 | 3.41098722619315 | 1.19322821036799 | 3.20132594093141 |
| chr15 | 67202537 | 67202562 | - | 11.043566345607 | 5.68497871032191 | 9.54582568294392 | 12.8053037637256 |
| chr15 | 67202594 | 67202632 | - | 22.087132691214 | 34.1098722619315 | 26.2510206280958 | 67.2278447595595 |
| chr15 | 67202660 | 67202674 | - | 23.0910932680874 | 15.9179403889013 | 20.2848795762558 | 27.7448248214055 |
| chr15 | 67202697 | 67202720 | - | 7.02772403811355 | 7.95897019445067 | 13.1255103140479 | 17.0737383516342 |
| chr15 | 67526524 | 67526555 | + | 6.02376346124019 | 3.41098722619315 | 1.19322821036799 | 1.06710864697714 |
| chr15 | 67807085 | 67807088 | + | 4.01584230749346 | 18.1919318730301 | 1.19322821036799 | 4.26843458790854 |
| chr15 | 68187875 | 68187905 | + | 2.00792115374673 | 3.41098722619315 | 15.5119667347839 | 3.20132594093141 |
| chr15 | 68188275 | 68188284 | + | 8.03168461498692 | 13.6439489047726 | 8.35259747257593 | 5.33554323488568 |
| chr15 | 68188857 | 68188878 | + | 5.01980288436682 | 2.27399148412876 | 1.19322821036799 | 4.26843458790854 |
| chr15 | 68188912 | 68188915 | + | 4.01584230749346 | 1.13699574206438 | 2.38645642073598 | 3.20132594093141 |
| chr15 | 68189200 | 68189230 | + | 7.02772403811355 | 5.68497871032191 | 22.6713359969918 | 7.46976052883995 |
| chr15 | 68189269 | 68189290 | + | 9.03564519186028 | 6.82197445238629 | 10.7390538933119 | 14.9395210576799 |
| chr15 | 68190298 | 68190303 | + | 8.03168461498692 | 4.54798296825753 | 3.57968463110397 | 5.33554323488568 |
| chr15 | 68190866 | 68190870 | + | 1.00396057687336 | 2.27399148412876 | 3.57968463110397 | 2.13421729395427 |
| chr15 | 68190931 | 68190944 | + | 6.02376346124019 | 22.7399148412876 | 13.1255103140479 | 9.60397782279422 |
| chr15 | 68190971 | 68190977 | + | 3.01188173062009 | 6.82197445238629 | 4.77291284147196 | 2.13421729395427 |
| chr15 | 68191119 | 68191142 | + | 19.0752509605939 | 9.09596593651506 | 3.57968463110397 | 14.9395210576799 |
| chr15 | 68191191 | 68191194 | + | 8.03168461498692 | 6.82197445238629 | 3.57968463110397 | 5.33554323488568 |
| chr15 | 68193881 | 68193883 | - | 1.00396057687336 | 1.13699574206438 | 1.19322821036799 | 3.20132594093141 |
| chr15 | 68193999 | 68194011 | - | 13.0514874993537 | 6.82197445238629 | 5.96614105183995 | 11.7381951167485 |
| chr15 | 68207029 | 68207059 | - | 7.02772403811355 | 12.5069531627082 | 3.57968463110397 | 5.33554323488568 |
| chr15 | 68291197 | 68291302 | + | 16.0633692299738 | 23.876910583352 | 9.54582568294392 | 6.40265188186281 |
| chr15 | 68291401 | 68291419 | + | 5.01980288436682 | 6.82197445238629 | 2.38645642073598 | 5.33554323488568 |
| chr15 | 68291440 | 68291475 | + | 9.03564519186028 | 12.5069531627082 | 11.9322821036799 | 3.20132594093141 |
| chr15 | 68292378 | 68292401 | + | 4.01584230749346 | 4.54798296825753 | 1.19322821036799 | 11.7381951167485 |
| chr15 | 68292456 | 68292459 | + | 2.00792115374673 | 4.54798296825753 | 1.19322821036799 | 1.06710864697714 |
| chr15 | 68292611 | 68292634 | + | 6.02376346124019 | 6.82197445238629 | 13.1255103140479 | 10.6710864697714 |
| chr15 | 68293017 | 68293050 | + | 4.01584230749346 | 4.54798296825753 | 1.19322821036799 | 10.6710864697714 |
| chr15 | 68293171 | 68293186 | + | 13.0514874993537 | 12.5069531627082 | 10.7390538933119 | 11.7381951167485 |
| chr15 | 68293235 | 68293292 | + | 14.0554480762271 | 11.3699574206438 | 10.7390538933119 | 5.33554323488568 |
| chr15 | 68293531 | 68293562 | + | 6.02376346124019 | 7.95897019445067 | 4.77291284147196 | 6.40265188186281 |
| chr15 | 68293621 | 68293622 | + | 3.01188173062009 | 4.54798296825753 | 2.38645642073598 | 1.06710864697714 |
| chr15 | 68293687 | 68293697 | + | 3.01188173062009 | 2.27399148412876 | 4.77291284147196 | 2.13421729395427 |
| chr15 | 68294142 | 68294143 | + | 3.01188173062009 | 3.41098722619315 | 2.38645642073598 | 2.13421729395427 |
| chr15 | 68295482 | 68295484 | + | 8.03168461498692 | 4.54798296825753 | 1.19322821036799 | 1.06710864697714 |
| chr15 | 68295606 | 68295610 | + | 6.02376346124019 | 13.6439489047726 | 3.57968463110397 | 4.26843458790854 |
| chr15 | 68778825 | 68778828 | - | 3.01188173062009 | 2.27399148412876 | 1.19322821036799 | 2.13421729395427 |
| chr15 | 68778935 | 68778999 | - | 6.02376346124019 | 7.95897019445067 | 2.38645642073598 | 2.13421729395427 |
| chr15 | 68779042 | 68779061 | - | 10.0396057687336 | 12.5069531627082 | 8.35259747257593 | 11.7381951167485 |
| chr15 | 68779082 | 68779096 | - | 3.01188173062009 | 4.54798296825753 | 1.19322821036799 | 2.13421729395427 |
| chr15 | 68779148 | 68779161 | - | 1.00396057687336 | 1.13699574206438 | 4.77291284147196 | 5.33554323488568 |
| chr15 | 68779191 | 68779229 | - | 5.01980288436682 | 9.09596593651506 | 3.57968463110397 | 10.6710864697714 |
| chr15 | 68779469 | 68779492 | - | 8.03168461498692 | 10.2329616785794 | 1.19322821036799 | 7.46976052883995 |
| chr15 | 68779663 | 68779666 | - | 12.0475269224804 | 4.54798296825753 | 1.19322821036799 | 5.33554323488568 |
| chr15 | 68779913 | 68779966 | - | 40.1584230749346 | 34.1098722619315 | 16.7051949451519 | 42.6843458790854 |
| chr15 | 68780436 | 68780462 | - | 40.1584230749346 | 29.5618892936739 | 38.1833027317757 | 54.4225409958339 |
| chr15 | 69448041 | 69448067 | + | 22.087132691214 | 9.09596593651506 | 2.38645642073598 | 8.53686917581709 |
| chr15 | 69448191 | 69448208 | + | 7.02772403811355 | 3.41098722619315 | 7.15936926220794 | 2.13421729395427 |
| chr15 | 69455509 | 69455521 | + | 3.01188173062009 | 11.3699574206438 | 14.3187385244159 | 4.26843458790854 |
| chr15 | 70047879 | 70047886 | - | 2.00792115374673 | 11.3699574206438 | 1.19322821036799 | 3.20132594093141 |
| chr15 | 70047937 | 70047940 | - | 13.0514874993537 | 25.0139063254164 | 9.54582568294392 | 7.46976052883995 |
| chr15 | 70048292 | 70048319 | - | 30.1188173062009 | 50.0278126508328 | 7.15936926220794 | 10.6710864697714 |
| chr15 | 70048559 | 70048603 | - | 18.0712903837206 | 19.3289276150945 | 9.54582568294392 | 8.53686917581709 |
| chr15 | 70048678 | 70048791 | - | 92.3643730723495 | 90.9596593651506 | 21.4781077866238 | 33.0803680562912 |
| chr15 | 70048812 | 70048840 | - | 17.0673298068472 | 26.1509020674808 | 8.35259747257593 | 5.33554323488568 |
| chr15 | 70048870 | 70048913 | - | 72.2851615348822 | 83.0006891706999 | 27.4442488384638 | 11.7381951167485 |
| chr15 | 70049021 | 70049057 | - | 20.0792115374673 | 32.9728765198671 | 5.96614105183995 | 24.5434988804741 |
| chr15 | 70049252 | 70049257 | - | 8.03168461498692 | 3.41098722619315 | 1.19322821036799 | 1.06710864697714 |
| chr15 | 70049299 | 70049334 | - | 12.0475269224804 | 15.9179403889013 | 2.38645642073598 | 7.46976052883995 |
| chr15 | 70049439 | 70049493 | - | 19.0752509605939 | 21.6029190992233 | 8.35259747257593 | 4.26843458790854 |
| chr15 | 70049594 | 70049745 | - | 126.499032686044 | 136.439489047726 | 41.7629873628796 | 19.2079556455884 |
| chr15 | 70049841 | 70049861 | - | 10.0396057687336 | 9.09596593651506 | 1.19322821036799 | 7.46976052883995 |
| chr15 | 70049886 | 70049915 | - | 34.1346596136944 | 42.0688424563821 | 2.38645642073598 | 1.06710864697714 |
| chr15 | 70050003 | 70050027 | - | 11.043566345607 | 6.82197445238629 | 3.57968463110397 | 2.13421729395427 |
| chr15 | 71824128 | 71824134 | - | 4.01584230749346 | 9.09596593651506 | 1.19322821036799 | 3.20132594093141 |
| chr15 | 71825832 | 71825851 | - | 2.00792115374673 | 3.41098722619315 | 1.19322821036799 | 1.06710864697714 |
| chr15 | 71826131 | 71826140 | - | 9.03564519186028 | 4.54798296825753 | 4.77291284147196 | 1.06710864697714 |
| chr15 | 71826229 | 71826268 | - | 64.2534769198953 | 69.3567402659273 | 13.1255103140479 | 10.6710864697714 |
| chr15 | 72199076 | 72199109 | - | 14.0554480762271 | 13.6439489047726 | 3.57968463110397 | 6.40265188186281 |
| chr15 | 72199135 | 72199156 | - | 5.01980288436682 | 5.68497871032191 | 3.57968463110397 | 7.46976052883995 |
| chr15 | 72199160 | 72199311 | - | 58.2297134586551 | 25.0139063254164 | 19.0916513658878 | 19.2079556455884 |
| chr15 | 72199328 | 72199380 | - | 20.0792115374673 | 15.9179403889013 | 2.38645642073598 | 4.26843458790854 |
| chr15 | 72199496 | 72199594 | - | 27.1069355755808 | 15.9179403889013 | 17.8984231555198 | 51.2212150549025 |
| chr15 | 72241260 | 72241270 | - | 17.0673298068472 | 21.6029190992233 | 1.19322821036799 | 8.53686917581709 |
| chr15 | 72343555 | 72343591 | - | 24.0950538449607 | 19.3289276150945 | 26.2510206280958 | 22.4092815865198 |
| chr15 | 72343880 | 72343967 | - | 16.0633692299738 | 17.0549361309657 | 11.9322821036799 | 11.7381951167485 |
| chr15 | 72343996 | 72344003 | - | 2.00792115374673 | 1.13699574206438 | 1.19322821036799 | 1.06710864697714 |
| chr15 | 72583566 | 72583575 | + | 1.00396057687336 | 2.27399148412876 | 3.57968463110397 | 5.33554323488568 |
| chr15 | 72583812 | 72583830 | + | 2.00792115374673 | 7.95897019445067 | 2.38645642073598 | 8.53686917581709 |
| chr15 | 72583943 | 72583953 | + | 1.00396057687336 | 11.3699574206438 | 1.19322821036799 | 1.06710864697714 |
| chr15 | 72585846 | 72585866 | + | 6.02376346124019 | 6.82197445238629 | 11.9322821036799 | 4.26843458790854 |
| chr15 | 72585912 | 72585932 | + | 2.00792115374673 | 4.54798296825753 | 5.96614105183995 | 2.13421729395427 |
| chr15 | 72585938 | 72585980 | + | 11.043566345607 | 12.5069531627082 | 20.2848795762558 | 9.60397782279422 |
| chr15 | 72586011 | 72586026 | + | 2.00792115374673 | 2.27399148412876 | 4.77291284147196 | 6.40265188186281 |
| chr15 | 72586073 | 72586092 | + | 2.00792115374673 | 9.09596593651506 | 1.19322821036799 | 1.06710864697714 |
| chr15 | 72586212 | 72586218 | + | 4.01584230749346 | 10.2329616785794 | 4.77291284147196 | 11.7381951167485 |
| chr15 | 72586361 | 72586397 | + | 4.01584230749346 | 7.95897019445067 | 1.19322821036799 | 3.20132594093141 |
| chr15 | 72586509 | 72586519 | + | 5.01980288436682 | 35.2468680039958 | 20.2848795762558 | 14.9395210576799 |
| chr15 | 72737536 | 72737597 | + | 2.00792115374673 | 5.68497871032191 | 14.3187385244159 | 19.2079556455884 |
| chr15 | 72737621 | 72737644 | + | 4.01584230749346 | 3.41098722619315 | 19.0916513658878 | 8.53686917581709 |
| chr15 | 72737666 | 72737696 | + | 2.00792115374673 | 1.13699574206438 | 5.96614105183995 | 7.46976052883995 |
| chr15 | 72737764 | 72737769 | + | 3.01188173062009 | 1.13699574206438 | 29.8307052591997 | 23.476390233497 |
| chr15 | 72738355 | 72738356 | + | 1.00396057687336 | 1.13699574206438 | 1.19322821036799 | 3.20132594093141 |
| chr15 | 72738414 | 72738446 | + | 6.02376346124019 | 9.09596593651506 | 7.15936926220794 | 11.7381951167485 |
| chr15 | 72751445 | 72751487 | - | 3.01188173062009 | 7.95897019445067 | 2.38645642073598 | 3.20132594093141 |
| chr15 | 72751516 | 72751536 | - | 2.00792115374673 | 4.54798296825753 | 3.57968463110397 | 3.20132594093141 |
| chr15 | 72751579 | 72751667 | - | 4.01584230749346 | 3.41098722619315 | 2.38645642073598 | 4.26843458790854 |
| chr15 | 73560395 | 73560539 | - | 10.0396057687336 | 15.9179403889013 | 2.38645642073598 | 3.20132594093141 |
| chr15 | 73560602 | 73560616 | - | 1060.18236917827 | 859.568781000673 | 201.65556755219 | 317.998376799186 |
| chr15 | 73560684 | 73560706 | - | 8.03168461498692 | 4.54798296825753 | 2.38645642073598 | 2.13421729395427 |
| chr15 | 73560734 | 73560926 | - | 199.7881547978 | 171.686357051722 | 69.2072362013434 | 41.6172372321083 |
| chr15 | 73560950 | 73561043 | - | 13.0514874993537 | 11.3699574206438 | 8.35259747257593 | 4.26843458790854 |
| chr15 | 73713016 | 73713173 | + | 87.3445701879827 | 68.2197445238629 | 17.8984231555198 | 5.33554323488568 |
| chr15 | 73714421 | 73714492 | + | 40.1584230749346 | 47.753821166704 | 2.38645642073598 | 4.26843458790854 |
| chr15 | 74036333 | 74036340 | + | 1.00396057687336 | 1.13699574206438 | 3.57968463110397 | 2.13421729395427 |
| chr15 | 74446018 | 74446060 | - | 27.1069355755808 | 10.2329616785794 | 20.2848795762558 | 22.4092815865198 |
| chr15 | 74461260 | 74461415 | + | 259.021828833328 | 285.38593125816 | 360.354919531133 | 392.695982087586 |
| chr15 | 74461789 | 74461791 | + | 1.00396057687336 | 1.13699574206438 | 5.96614105183995 | 3.20132594093141 |
| chr15 | 74630585 | 74630601 | - | 3.01188173062009 | 5.68497871032191 | 3.57968463110397 | 3.20132594093141 |
| chr15 | 74630625 | 74630632 | - | 14.0554480762271 | 4.54798296825753 | 2.38645642073598 | 8.53686917581709 |
| chr15 | 74632055 | 74632061 | - | 14.0554480762271 | 18.1919318730301 | 4.77291284147196 | 3.20132594093141 |
| chr15 | 74836145 | 74836172 | - | 20.0792115374673 | 26.1509020674808 | 16.7051949451519 | 20.2750642925656 |
| chr15 | 74836194 | 74836201 | - | 1.00396057687336 | 14.780944646837 | 2.38645642073598 | 5.33554323488568 |
| chr15 | 74836234 | 74836244 | - | 19.0752509605939 | 23.876910583352 | 2.38645642073598 | 7.46976052883995 |
| chr15 | 74836283 | 74836319 | - | 2.00792115374673 | 6.82197445238629 | 1.19322821036799 | 5.33554323488568 |
| chr15 | 74843760 | 74843841 | - | 21.0831721143407 | 26.1509020674808 | 9.54582568294392 | 8.53686917581709 |
| chr15 | 74844914 | 74844926 | - | 7.02772403811355 | 12.5069531627082 | 7.15936926220794 | 18.1408469986113 |
| chr15 | 74897580 | 74897582 | + | 3.01188173062009 | 3.41098722619315 | 22.6713359969918 | 2.13421729395427 |
| chr15 | 74898140 | 74898179 | + | 4.01584230749346 | 3.41098722619315 | 7.15936926220794 | 16.006629704657 |
| chr15 | 74899161 | 74899165 | + | 1.00396057687336 | 2.27399148412876 | 2.38645642073598 | 3.20132594093141 |
| chr15 | 74900071 | 74900122 | - | 22.087132691214 | 34.1098722619315 | 5.96614105183995 | 10.6710864697714 |
| chr15 | 74900241 | 74900269 | - | 2.00792115374673 | 3.41098722619315 | 4.77291284147196 | 2.13421729395427 |
| chr15 | 74900295 | 74900315 | - | 1.00396057687336 | 2.27399148412876 | 3.57968463110397 | 2.13421729395427 |
| chr15 | 74900995 | 74901012 | - | 1.00396057687336 | 1.13699574206438 | 2.38645642073598 | 1.06710864697714 |
| chr15 | 74901752 | 74901764 | - | 1.00396057687336 | 1.13699574206438 | 8.35259747257593 | 1.06710864697714 |
| chr15 | 74901949 | 74901956 | - | 5.01980288436682 | 11.3699574206438 | 1.19322821036799 | 1.06710864697714 |
| chr15 | 74902253 | 74902301 | - | 1.00396057687336 | 2.27399148412876 | 3.57968463110397 | 2.13421729395427 |
| chr15 | 74919920 | 74919939 | - | 2.00792115374673 | 2.27399148412876 | 1.19322821036799 | 1.06710864697714 |
| chr15 | 74920321 | 74920329 | - | 15.0594086531005 | 23.876910583352 | 13.1255103140479 | 11.7381951167485 |
| chr15 | 74920395 | 74920445 | - | 30.1188173062009 | 48.8908169087684 | 50.1155848354556 | 43.7514545260626 |
| chr15 | 74923650 | 74923676 | - | 8.03168461498692 | 10.2329616785794 | 10.7390538933119 | 12.8053037637256 |
| chr15 | 74954477 | 74954494 | - | 234.926774988367 | 214.892195250168 | 20.2848795762558 | 11.7381951167485 |
| chr15 | 74954530 | 74954565 | - | 98.3881365335897 | 59.1237785873479 | 7.15936926220794 | 7.46976052883995 |
| chr15 | 74955334 | 74955341 | - | 9.03564519186028 | 12.5069531627082 | 3.57968463110397 | 2.13421729395427 |
| chr15 | 74955433 | 74955451 | - | 52.205949997415 | 31.8358807778027 | 2.38645642073598 | 3.20132594093141 |
| chr15 | 74955489 | 74955523 | - | 58.2297134586551 | 52.3018041349616 | 13.1255103140479 | 7.46976052883995 |
| chr15 | 74955557 | 74955576 | - | 43.1703048055547 | 26.1509020674808 | 8.35259747257593 | 4.26843458790854 |
| chr15 | 74955655 | 74955735 | - | 23.0910932680874 | 27.2878978095452 | 13.1255103140479 | 3.20132594093141 |
| chr15 | 74955756 | 74955777 | - | 10.0396057687336 | 2.27399148412876 | 2.38645642073598 | 2.13421729395427 |
| chr15 | 75049710 | 75049719 | + | 20.0792115374673 | 208.070220797782 | 32.2171616799357 | 39.483019938154 |
| chr15 | 75211193 | 75211236 | + | 26.1029749987075 | 10.2329616785794 | 23.8645642073598 | 10.6710864697714 |
| chr15 | 75211258 | 75211311 | + | 21.0831721143407 | 10.2329616785794 | 42.9562155732476 | 54.4225409958339 |
| chr15 | 75340174 | 75340185 | + | 5.01980288436682 | 5.68497871032191 | 4.77291284147196 | 3.20132594093141 |
| chr15 | 75340218 | 75340248 | + | 95.3762548029696 | 170.549361309657 | 20.2848795762558 | 44.8185631730397 |
| chr15 | 75369059 | 75369065 | + | 6.02376346124019 | 7.95897019445067 | 2.38645642073598 | 4.26843458790854 |
| chr15 | 75369441 | 75369442 | + | 1.00396057687336 | 1.13699574206438 | 1.19322821036799 | 1.06710864697714 |
| chr15 | 75369604 | 75369608 | - | 7.02772403811355 | 4.54798296825753 | 2.38645642073598 | 2.13421729395427 |
| chr15 | 75369639 | 75369640 | - | 1.00396057687336 | 2.27399148412876 | 1.19322821036799 | 3.20132594093141 |
| chr15 | 75369735 | 75369755 | - | 4.01584230749346 | 2.27399148412876 | 1.19322821036799 | 2.13421729395427 |
| chr15 | 75370647 | 75370649 | - | 1.00396057687336 | 2.27399148412876 | 1.19322821036799 | 3.20132594093141 |
| chr15 | 75371284 | 75371319 | - | 4.01584230749346 | 5.68497871032191 | 1.19322821036799 | 8.53686917581709 |
| chr15 | 75371382 | 75371402 | - | 5.01980288436682 | 7.95897019445067 | 2.38645642073598 | 4.26843458790854 |
| chr15 | 75371433 | 75371434 | - | 1.00396057687336 | 1.13699574206438 | 2.38645642073598 | 2.13421729395427 |
| chr15 | 75371573 | 75371580 | - | 6.02376346124019 | 4.54798296825753 | 2.38645642073598 | 7.46976052883995 |
| chr15 | 75371628 | 75371665 | - | 6.02376346124019 | 3.41098722619315 | 1.19322821036799 | 2.13421729395427 |
| chr15 | 75371723 | 75371772 | - | 1.00396057687336 | 1.13699574206438 | 1.19322821036799 | 1.06710864697714 |
| chr15 | 75421987 | 75421989 | - | 2.00792115374673 | 14.780944646837 | 8.35259747257593 | 5.33554323488568 |
| chr15 | 75467310 | 75467326 | - | 3.01188173062009 | 4.54798296825753 | 1.19322821036799 | 1.06710864697714 |
| chr15 | 75467471 | 75467494 | - | 5.01980288436682 | 3.41098722619315 | 2.38645642073598 | 4.26843458790854 |
| chr15 | 75598168 | 75598203 | - | 13.0514874993537 | 19.3289276150945 | 7.15936926220794 | 9.60397782279422 |
| chr15 | 75598278 | 75598299 | - | 1.00396057687336 | 4.54798296825753 | 1.19322821036799 | 7.46976052883995 |
| chr15 | 75639113 | 75639224 | - | 37.1465413443145 | 40.9318467143178 | 20.2848795762558 | 26.6777161744284 |
| chr15 | 75639256 | 75639267 | - | 9.03564519186028 | 7.95897019445067 | 2.38645642073598 | 11.7381951167485 |
| chr15 | 75639332 | 75639362 | - | 9.03564519186028 | 5.68497871032191 | 2.38645642073598 | 1.06710864697714 |
| chr15 | 75658559 | 75658587 | + | 12.0475269224804 | 18.1919318730301 | 3.57968463110397 | 9.60397782279422 |
| chr15 | 75758525 | 75758556 | + | 5.01980288436682 | 12.5069531627082 | 9.54582568294392 | 6.40265188186281 |
| chr15 | 75933071 | 75933175 | + | 19.0752509605939 | 40.9318467143178 | 46.5359002043516 | 21.3421729395427 |
| chr15 | 75933228 | 75933249 | + | 4.01584230749346 | 6.82197445238629 | 9.54582568294392 | 8.53686917581709 |
| chr15 | 75933271 | 75933322 | + | 4.01584230749346 | 5.68497871032191 | 11.9322821036799 | 11.7381951167485 |
| chr15 | 75933526 | 75933548 | + | 3.01188173062009 | 4.54798296825753 | 1.19322821036799 | 5.33554323488568 |
| chr15 | 75934152 | 75934155 | + | 1.00396057687336 | 1.13699574206438 | 2.38645642073598 | 1.06710864697714 |
| chr15 | 75934343 | 75934350 | + | 1.00396057687336 | 1.13699574206438 | 4.77291284147196 | 5.33554323488568 |
| chr15 | 75936203 | 75936204 | + | 6.02376346124019 | 1.13699574206438 | 3.57968463110397 | 3.20132594093141 |
| chr15 | 75938315 | 75938321 | + | 1.00396057687336 | 9.09596593651506 | 7.15936926220794 | 7.46976052883995 |
| chr15 | 75939046 | 75939050 | + | 90.3564519186028 | 81.8636934286355 | 94.2650286190712 | 46.952780466994 |
| chr15 | 75939245 | 75939254 | + | 2.00792115374673 | 5.68497871032191 | 8.35259747257593 | 3.20132594093141 |
| chr15 | 75940398 | 75940443 | + | 7.02772403811355 | 1.13699574206438 | 5.96614105183995 | 10.6710864697714 |
| chr15 | 75941249 | 75941252 | + | 11.043566345607 | 5.68497871032191 | 10.7390538933119 | 6.40265188186281 |
| chr15 | 75941292 | 75941310 | + | 5.01980288436682 | 6.82197445238629 | 9.54582568294392 | 11.7381951167485 |
| chr15 | 75941749 | 75941784 | + | 20.0792115374673 | 17.0549361309657 | 23.8645642073598 | 9.60397782279422 |
| chr15 | 75942001 | 75942008 | + | 6.02376346124019 | 4.54798296825753 | 4.77291284147196 | 4.26843458790854 |
| chr15 | 75942163 | 75942201 | + | 6.02376346124019 | 3.41098722619315 | 1.19322821036799 | 3.20132594093141 |
| chr15 | 75942431 | 75942447 | + | 4.01584230749346 | 5.68497871032191 | 2.38645642073598 | 4.26843458790854 |
| chr15 | 76216394 | 76216502 | - | 115.455466340437 | 151.220433694563 | 76.3666054635513 | 135.522798166096 |
| chr15 | 76216553 | 76216599 | - | 6.02376346124019 | 2.27399148412876 | 2.38645642073598 | 2.13421729395427 |
| chr15 | 76341977 | 76341978 | + | 2.00792115374673 | 2.27399148412876 | 2.38645642073598 | 3.20132594093141 |
| chr15 | 76949066 | 76949374 | + | 24.0950538449607 | 22.7399148412876 | 7.15936926220794 | 19.2079556455884 |
| chr15 | 76949437 | 76949654 | + | 62.2455557661486 | 60.2607743294122 | 19.0916513658878 | 30.9461507623369 |
| chr15 | 76949711 | 76949725 | + | 5.01980288436682 | 5.68497871032191 | 1.19322821036799 | 2.13421729395427 |
| chr15 | 76949829 | 76949849 | + | 4.01584230749346 | 4.54798296825753 | 3.57968463110397 | 1.06710864697714 |
| chr15 | 77046060 | 77046105 | - | 2.00792115374673 | 4.54798296825753 | 2.38645642073598 | 3.20132594093141 |
| chr15 | 77046126 | 77046313 | - | 48.1901076899215 | 57.9867828452835 | 29.8307052591997 | 22.4092815865198 |
| chr15 | 77046384 | 77046442 | - | 24.0950538449607 | 21.6029190992233 | 9.54582568294392 | 18.1408469986113 |
| chr15 | 77046792 | 77046923 | - | 44.174265382428 | 30.6988850357383 | 29.8307052591997 | 43.7514545260626 |
| chr15 | 77485413 | 77485425 | + | 3.01188173062009 | 10.2329616785794 | 7.15936926220794 | 5.33554323488568 |
| chr15 | 77995717 | 77995718 | - | 4.01584230749346 | 3.41098722619315 | 2.38645642073598 | 1.06710864697714 |
| chr15 | 78166296 | 78166297 | + | 8.03168461498692 | 1.13699574206438 | 2.38645642073598 | 1.06710864697714 |
| chr15 | 78169156 | 78169176 | + | 12.0475269224804 | 6.82197445238629 | 3.57968463110397 | 6.40265188186281 |
| chr15 | 78169321 | 78169334 | + | 4.01584230749346 | 1.13699574206438 | 1.19322821036799 | 1.06710864697714 |
| chr15 | 78169471 | 78169492 | + | 5.01980288436682 | 13.6439489047726 | 9.54582568294392 | 8.53686917581709 |
| chr15 | 78169740 | 78169765 | + | 3.01188173062009 | 5.68497871032191 | 7.15936926220794 | 6.40265188186281 |
| chr15 | 78169855 | 78169921 | + | 37.1465413443145 | 48.8908169087684 | 60.8546387287675 | 50.1541064079254 |
| chr15 | 78169949 | 78169961 | + | 7.02772403811355 | 4.54798296825753 | 9.54582568294392 | 4.26843458790854 |
| chr15 | 78170113 | 78170115 | + | 4.01584230749346 | 2.27399148412876 | 2.38645642073598 | 2.13421729395427 |
| chr15 | 78170250 | 78170251 | + | 3.01188173062009 | 2.27399148412876 | 1.19322821036799 | 3.20132594093141 |
| chr15 | 78170347 | 78170428 | + | 71.2812009580089 | 70.4937360079917 | 34.6036181006717 | 39.483019938154 |
| chr15 | 78283266 | 78283279 | - | 2.00792115374673 | 15.9179403889013 | 9.54582568294392 | 11.7381951167485 |
| chr15 | 78283388 | 78283392 | - | 3.01188173062009 | 4.54798296825753 | 7.15936926220794 | 6.40265188186281 |
| chr15 | 78498398 | 78498436 | + | 24.0950538449607 | 51.1648083928972 | 15.5119667347839 | 23.476390233497 |
| chr15 | 78499086 | 78499090 | + | 3.01188173062009 | 5.68497871032191 | 2.38645642073598 | 1.06710864697714 |
| chr15 | 78499378 | 78499405 | + | 12.0475269224804 | 3.41098722619315 | 2.38645642073598 | 5.33554323488568 |
| chr15 | 78499535 | 78499537 | + | 4.01584230749346 | 2.27399148412876 | 1.19322821036799 | 1.06710864697714 |
| chr15 | 78499693 | 78499712 | + | 3.01188173062009 | 5.68497871032191 | 3.57968463110397 | 2.13421729395427 |
| chr15 | 78500129 | 78500147 | + | 2.00792115374673 | 9.09596593651506 | 4.77291284147196 | 3.20132594093141 |
| chr15 | 78500185 | 78500194 | + | 6.02376346124019 | 14.780944646837 | 9.54582568294392 | 5.33554323488568 |
| chr15 | 78500376 | 78500407 | + | 11.043566345607 | 5.68497871032191 | 5.96614105183995 | 1.06710864697714 |
| chr15 | 78500939 | 78501002 | + | 9.03564519186028 | 7.95897019445067 | 17.8984231555198 | 16.006629704657 |
| chr15 | 78501066 | 78501091 | + | 4.01584230749346 | 7.95897019445067 | 2.38645642073598 | 4.26843458790854 |
| chr15 | 78501122 | 78501141 | + | 27.1069355755808 | 32.9728765198671 | 19.0916513658878 | 42.6843458790854 |
| chr15 | 78501372 | 78501415 | + | 14.0554480762271 | 18.1919318730301 | 13.1255103140479 | 22.4092815865198 |
| chr15 | 78548787 | 78548908 | + | 61.2415951892752 | 53.438799877026 | 69.2072362013434 | 76.8318225823538 |
| chr15 | 78549013 | 78549075 | + | 37.1465413443145 | 34.1098722619315 | 21.4781077866238 | 33.0803680562912 |
| chr15 | 78549113 | 78549137 | + | 43.1703048055547 | 42.0688424563821 | 27.4442488384638 | 21.3421729395427 |
| chr15 | 78594024 | 78594025 | + | 7.02772403811355 | 3.41098722619315 | 2.38645642073598 | 1.06710864697714 |
| chr15 | 78897199 | 78897231 | + | 227.899050950254 | 245.591080285907 | 77.5598336739193 | 167.53605757541 |
| chr15 | 78897467 | 78897480 | + | 7.02772403811355 | 3.41098722619315 | 4.77291284147196 | 4.26843458790854 |
| chr15 | 78897624 | 78897643 | + | 22.087132691214 | 17.0549361309657 | 8.35259747257593 | 16.006629704657 |
| chr15 | 79322197 | 79322219 | + | 2.00792115374673 | 1.13699574206438 | 2.38645642073598 | 4.26843458790854 |
| chr15 | 79322249 | 79322252 | + | 1.00396057687336 | 2.27399148412876 | 2.38645642073598 | 1.06710864697714 |
| chr15 | 79322282 | 79322302 | + | 2.00792115374673 | 3.41098722619315 | 3.57968463110397 | 4.26843458790854 |
| chr15 | 79322458 | 79322510 | + | 4.01584230749346 | 4.54798296825753 | 1.19322821036799 | 2.13421729395427 |
| chr15 | 79322537 | 79322689 | + | 36.1425807674411 | 21.6029190992233 | 8.35259747257593 | 9.60397782279422 |
| chr15 | 79322712 | 79322817 | + | 94.3722942260963 | 183.056314472366 | 64.4343233598714 | 54.4225409958339 |
| chr15 | 79845069 | 79845075 | - | 3.01188173062009 | 4.54798296825753 | 2.38645642073598 | 1.06710864697714 |
| chr15 | 79845124 | 79845170 | - | 12.0475269224804 | 17.0549361309657 | 2.38645642073598 | 1.06710864697714 |
| chr15 | 79923142 | 79923172 | + | 3.01188173062009 | 5.68497871032191 | 1.19322821036799 | 5.33554323488568 |
| chr15 | 79923341 | 79923342 | + | 4.01584230749346 | 5.68497871032191 | 1.19322821036799 | 1.06710864697714 |
| chr15 | 80137634 | 80137656 | + | 3.01188173062009 | 3.41098722619315 | 2.38645642073598 | 1.06710864697714 |
| chr15 | 80137700 | 80137742 | + | 8.03168461498692 | 9.09596593651506 | 23.8645642073598 | 23.476390233497 |
| chr15 | 80137809 | 80137836 | + | 3.01188173062009 | 3.41098722619315 | 1.19322821036799 | 8.53686917581709 |
| chr15 | 80137894 | 80137914 | + | 33.130699036821 | 45.4798296825753 | 23.8645642073598 | 42.6843458790854 |
| chr15 | 80138029 | 80138032 | + | 6.02376346124019 | 2.27399148412876 | 3.57968463110397 | 3.20132594093141 |
| chr15 | 80138155 | 80138189 | + | 3.01188173062009 | 7.95897019445067 | 13.1255103140479 | 22.4092815865198 |
| chr15 | 80138283 | 80138292 | + | 7.02772403811355 | 9.09596593651506 | 8.35259747257593 | 3.20132594093141 |
| chr15 | 80755445 | 80755461 | + | 23.0910932680874 | 7.95897019445067 | 1.19322821036799 | 1.06710864697714 |
| chr15 | 80977057 | 80977102 | - | 8.03168461498692 | 6.82197445238629 | 10.7390538933119 | 9.60397782279422 |
| chr15 | 80977378 | 80977380 | - | 2.00792115374673 | 1.13699574206438 | 3.57968463110397 | 1.06710864697714 |
| chr15 | 80978242 | 80978252 | - | 1.00396057687336 | 1.13699574206438 | 1.19322821036799 | 2.13421729395427 |
| chr15 | 80978434 | 80978491 | - | 4.01584230749346 | 9.09596593651506 | 5.96614105183995 | 7.46976052883995 |
| chr15 | 80978513 | 80978523 | - | 8.03168461498692 | 5.68497871032191 | 2.38645642073598 | 4.26843458790854 |
| chr15 | 80978558 | 80978612 | - | 235.930735565241 | 200.111250603331 | 53.6952694665595 | 57.6238669367653 |
| chr15 | 80978677 | 80978796 | - | 60.2376346124019 | 67.0827487817985 | 50.1155848354556 | 72.5633879944452 |
| chr15 | 80978829 | 80978843 | - | 2.00792115374673 | 3.41098722619315 | 3.57968463110397 | 3.20132594093141 |
| chr15 | 80978871 | 80978886 | - | 7.02772403811355 | 4.54798296825753 | 2.38645642073598 | 4.26843458790854 |
| chr15 | 80978925 | 80978971 | - | 6.02376346124019 | 9.09596593651506 | 3.57968463110397 | 10.6710864697714 |
| chr15 | 80978992 | 80979258 | - | 137.542599031651 | 104.603608269923 | 69.2072362013434 | 99.2411041688736 |
| chr15 | 81003485 | 81003507 | + | 2.00792115374673 | 2.27399148412876 | 2.38645642073598 | 9.60397782279422 |
| chr15 | 81003676 | 81003693 | + | 5.01980288436682 | 3.41098722619315 | 4.77291284147196 | 14.9395210576799 |
| chr15 | 81003871 | 81003891 | + | 25.0990144218341 | 39.7948509722534 | 10.7390538933119 | 73.6304966414224 |
| chr15 | 81324389 | 81324412 | + | 2.00792115374673 | 2.27399148412876 | 5.96614105183995 | 16.006629704657 |
| chr15 | 81324446 | 81324496 | + | 1.00396057687336 | 3.41098722619315 | 25.0577924177278 | 27.7448248214055 |
| chr15 | 82749700 | 82749704 | + | 2.00792115374673 | 3.41098722619315 | 3.57968463110397 | 5.33554323488568 |
| chr15 | 82750528 | 82750608 | + | 586.312976894045 | 674.238475044178 | 578.715682028475 | 348.944527561523 |
| chr15 | 82750677 | 82750722 | + | 29.1148567293276 | 44.3428339405109 | 36.9900745214077 | 17.0737383516342 |
| chr15 | 82750751 | 82750760 | + | 2.00792115374673 | 3.41098722619315 | 3.57968463110397 | 2.13421729395427 |
| chr15 | 82750840 | 82750849 | + | 4.01584230749346 | 3.41098722619315 | 3.57968463110397 | 1.06710864697714 |
| chr15 | 82756728 | 82756739 | + | 17.0673298068472 | 12.5069531627082 | 17.8984231555198 | 11.7381951167485 |
| chr15 | 82756916 | 82756940 | + | 22.087132691214 | 6.82197445238629 | 2.38645642073598 | 9.60397782279422 |
| chr15 | 82757011 | 82757031 | - | 3.01188173062009 | 6.82197445238629 | 2.38645642073598 | 1.06710864697714 |
| chr15 | 82834417 | 82834484 | + | 21.0831721143407 | 25.0139063254164 | 3.57968463110397 | 5.33554323488568 |
| chr15 | 82990699 | 82990719 | + | 35.1386201905678 | 47.753821166704 | 20.2848795762558 | 27.7448248214055 |
| chr15 | 82990834 | 82990848 | + | 3.01188173062009 | 3.41098722619315 | 3.57968463110397 | 3.20132594093141 |
| chr15 | 82990874 | 82990894 | + | 26.1029749987075 | 25.0139063254164 | 26.2510206280958 | 33.0803680562912 |
| chr15 | 83004836 | 83004839 | - | 16.0633692299738 | 13.6439489047726 | 1.19322821036799 | 5.33554323488568 |
| chr15 | 83005289 | 83005326 | - | 13.0514874993537 | 18.1919318730301 | 9.54582568294392 | 7.46976052883995 |
| chr15 | 83016472 | 83016477 | - | 3.01188173062009 | 2.27399148412876 | 2.38645642073598 | 4.26843458790854 |
| chr15 | 83016506 | 83016508 | - | 1.00396057687336 | 2.27399148412876 | 2.38645642073598 | 4.26843458790854 |
| chr15 | 83016570 | 83016595 | - | 8.03168461498692 | 12.5069531627082 | 9.54582568294392 | 9.60397782279422 |
| chr15 | 83016690 | 83016692 | - | 2.00792115374673 | 2.27399148412876 | 1.19322821036799 | 2.13421729395427 |
| chr15 | 83016742 | 83016797 | - | 7.02772403811355 | 7.95897019445067 | 4.77291284147196 | 3.20132594093141 |
| chr15 | 83017219 | 83017296 | - | 21.0831721143407 | 21.6029190992233 | 25.0577924177278 | 21.3421729395427 |
| chr15 | 83017364 | 83017389 | - | 3.01188173062009 | 6.82197445238629 | 5.96614105183995 | 6.40265188186281 |
| chr15 | 83017474 | 83017489 | - | 1.00396057687336 | 1.13699574206438 | 7.15936926220794 | 9.60397782279422 |
| chr15 | 83017557 | 83017585 | - | 35.1386201905678 | 44.3428339405109 | 53.6952694665595 | 20.2750642925656 |
| chr15 | 83017614 | 83017711 | - | 87.3445701879827 | 86.411676396893 | 110.970223564223 | 99.2411041688736 |
| chr15 | 83017726 | 83017750 | - | 10.0396057687336 | 11.3699574206438 | 11.9322821036799 | 13.8724124107028 |
| chr15 | 83017766 | 83017866 | - | 14.0554480762271 | 15.9179403889013 | 52.5020412561915 | 21.3421729395427 |
| chr15 | 83017981 | 83018008 | - | 7.02772403811355 | 5.68497871032191 | 17.8984231555198 | 4.26843458790854 |
| chr15 | 83018098 | 83018131 | - | 9.03564519186028 | 9.09596593651506 | 7.15936926220794 | 4.26843458790854 |
| chr15 | 83018156 | 83018169 | - | 3.01188173062009 | 6.82197445238629 | 13.1255103140479 | 12.8053037637256 |
| chr15 | 83018196 | 83018226 | - | 13.0514874993537 | 14.780944646837 | 11.9322821036799 | 16.006629704657 |
| chr15 | 83018706 | 83018840 | - | 36.1425807674411 | 31.8358807778027 | 81.1395183050233 | 61.8923015246739 |
| chr15 | 83135997 | 83135998 | - | 2.00792115374673 | 2.27399148412876 | 3.57968463110397 | 1.06710864697714 |
| chr15 | 83139102 | 83139104 | - | 6.02376346124019 | 7.95897019445067 | 7.15936926220794 | 1.06710864697714 |
| chr15 | 84639398 | 84639401 | - | 9.03564519186028 | 10.2329616785794 | 1.19322821036799 | 3.20132594093141 |
| chr15 | 84639429 | 84639440 | - | 15.0594086531005 | 18.1919318730301 | 1.19322821036799 | 3.20132594093141 |
| chr15 | 84669564 | 84669647 | - | 42.1663442286813 | 47.753821166704 | 34.6036181006717 | 22.4092815865198 |
| chr15 | 84669722 | 84669770 | - | 3.01188173062009 | 4.54798296825753 | 3.57968463110397 | 7.46976052883995 |
| chr15 | 84669787 | 84669872 | - | 59.2336740355285 | 86.411676396893 | 26.2510206280958 | 45.8856718200168 |
| chr15 | 84669884 | 84670007 | - | 8.03168461498692 | 15.9179403889013 | 11.9322821036799 | 12.8053037637256 |
| chr15 | 84670047 | 84670072 | - | 9.03564519186028 | 2.27399148412876 | 1.19322821036799 | 3.20132594093141 |
| chr15 | 84670724 | 84670728 | - | 21.0831721143407 | 10.2329616785794 | 13.1255103140479 | 24.5434988804741 |
| chr15 | 84805832 | 84805834 | + | 7.02772403811355 | 6.82197445238629 | 1.19322821036799 | 4.26843458790854 |
| chr15 | 85137904 | 85137920 | + | 2.00792115374673 | 6.82197445238629 | 1.19322821036799 | 2.13421729395427 |
| chr15 | 85380568 | 85380696 | + | 863.406096111093 | 1395.093775513 | 2.38645642073598 | 2.13421729395427 |
| chr15 | 88459513 | 88459611 | - | 54.2138711511617 | 103.466612527859 | 16.7051949451519 | 27.7448248214055 |
| chr15 | 88478202 | 88478238 | + | 56.2217923049084 | 47.753821166704 | 10.7390538933119 | 23.476390233497 |
| chr15 | 88478681 | 88478683 | + | 3.01188173062009 | 2.27399148412876 | 1.19322821036799 | 2.13421729395427 |
| chr15 | 88631513 | 88631574 | + | 12.0475269224804 | 12.5069531627082 | 9.54582568294392 | 11.7381951167485 |
| chr15 | 88631621 | 88631651 | + | 8.03168461498692 | 7.95897019445067 | 8.35259747257593 | 7.46976052883995 |
| chr15 | 88632177 | 88632234 | + | 27.1069355755808 | 59.1237785873479 | 20.2848795762558 | 44.8185631730397 |
| chr15 | 89196232 | 89196236 | + | 1.00396057687336 | 1.13699574206438 | 2.38645642073598 | 2.13421729395427 |
| chr15 | 89196514 | 89196545 | + | 7.02772403811355 | 1.13699574206438 | 2.38645642073598 | 6.40265188186281 |
| chr15 | 89198275 | 89198283 | + | 1.00396057687336 | 2.27399148412876 | 4.77291284147196 | 1.06710864697714 |
| chr15 | 89201420 | 89201422 | + | 1.00396057687336 | 1.13699574206438 | 4.77291284147196 | 1.06710864697714 |
| chr15 | 89201630 | 89201641 | + | 1.00396057687336 | 1.13699574206438 | 9.54582568294392 | 2.13421729395427 |
| chr15 | 89201845 | 89201900 | + | 48.1901076899215 | 31.8358807778027 | 133.641559561215 | 130.187254931211 |
| chr15 | 89201931 | 89201933 | + | 1.00396057687336 | 1.13699574206438 | 1.19322821036799 | 3.20132594093141 |
| chr15 | 89202097 | 89202100 | + | 12.0475269224804 | 6.82197445238629 | 5.96614105183995 | 8.53686917581709 |
| chr15 | 89202261 | 89202267 | + | 3.01188173062009 | 3.41098722619315 | 4.77291284147196 | 5.33554323488568 |
| chr15 | 89316342 | 89316366 | - | 11.043566345607 | 12.5069531627082 | 5.96614105183995 | 4.26843458790854 |
| chr15 | 89316387 | 89316408 | - | 18.0712903837206 | 13.6439489047726 | 5.96614105183995 | 21.3421729395427 |
| chr15 | 89316395 | 89316462 | + | 38.1505019211879 | 38.657855230189 | 17.8984231555198 | 23.476390233497 |
| chr15 | 89316487 | 89316491 | + | 6.02376346124019 | 1.13699574206438 | 1.19322821036799 | 1.06710864697714 |
| chr15 | 89316511 | 89316549 | - | 6.02376346124019 | 3.41098722619315 | 3.57968463110397 | 8.53686917581709 |
| chr15 | 89316723 | 89316765 | + | 33.130699036821 | 23.876910583352 | 3.57968463110397 | 13.8724124107028 |
| chr15 | 89316931 | 89316977 | + | 19.0752509605939 | 19.3289276150945 | 22.6713359969918 | 16.006629704657 |
| chr15 | 89317091 | 89317111 | + | 10.0396057687336 | 20.4659233571589 | 14.3187385244159 | 3.20132594093141 |
| chr15 | 89317150 | 89317152 | + | 3.01188173062009 | 2.27399148412876 | 2.38645642073598 | 1.06710864697714 |
| chr15 | 89627398 | 89627409 | + | 5.01980288436682 | 1.13699574206438 | 9.54582568294392 | 3.20132594093141 |
| chr15 | 89627483 | 89627505 | + | 6.02376346124019 | 2.27399148412876 | 3.57968463110397 | 5.33554323488568 |
| chr15 | 89627518 | 89627539 | + | 1.00396057687336 | 1.13699574206438 | 1.19322821036799 | 5.33554323488568 |
| chr15 | 89627747 | 89627750 | + | 4.01584230749346 | 6.82197445238629 | 3.57968463110397 | 7.46976052883995 |
| chr15 | 89627843 | 89627863 | + | 21.0831721143407 | 21.6029190992233 | 7.15936926220794 | 32.0132594093141 |
| chr15 | 89830685 | 89830710 | - | 36.1425807674411 | 21.6029190992233 | 8.35259747257593 | 11.7381951167485 |
| chr15 | 89834373 | 89834406 | - | 3.01188173062009 | 3.41098722619315 | 3.57968463110397 | 3.20132594093141 |
| chr15 | 90083996 | 90084100 | - | 4.01584230749346 | 6.82197445238629 | 31.0239334695677 | 23.476390233497 |
| chr15 | 90084144 | 90084168 | - | 2.00792115374673 | 4.54798296825753 | 13.1255103140479 | 16.006629704657 |
| chr15 | 90084190 | 90084195 | - | 1.00396057687336 | 1.13699574206438 | 4.77291284147196 | 6.40265188186281 |
| chr15 | 90228655 | 90228701 | + | 5.01980288436682 | 2.27399148412876 | 14.3187385244159 | 16.006629704657 |
| chr15 | 90229472 | 90229494 | + | 6.02376346124019 | 5.68497871032191 | 1.19322821036799 | 8.53686917581709 |
| chr15 | 90229571 | 90229576 | + | 5.01980288436682 | 3.41098722619315 | 10.7390538933119 | 11.7381951167485 |
| chr15 | 90230278 | 90230400 | - | 181.716864414079 | 94.3706465913437 | 127.675418509375 | 277.448248214055 |
| chr15 | 90230453 | 90230473 | - | 9.03564519186028 | 7.95897019445067 | 5.96614105183995 | 2.13421729395427 |
| chr15 | 90242945 | 90242948 | + | 6.02376346124019 | 7.95897019445067 | 2.38645642073598 | 1.06710864697714 |
| chr15 | 90266359 | 90266399 | + | 7.02772403811355 | 3.41098722619315 | 3.57968463110397 | 3.20132594093141 |
| chr15 | 90271301 | 90271345 | + | 4.01584230749346 | 1.13699574206438 | 4.77291284147196 | 6.40265188186281 |
| chr15 | 90271372 | 90271378 | + | 4.01584230749346 | 2.27399148412876 | 5.96614105183995 | 1.06710864697714 |
| chr15 | 90271526 | 90271569 | + | 8.03168461498692 | 12.5069531627082 | 5.96614105183995 | 11.7381951167485 |
| chr15 | 90271739 | 90271747 | + | 5.01980288436682 | 11.3699574206438 | 1.19322821036799 | 4.26843458790854 |
| chr15 | 90271784 | 90271795 | + | 2.00792115374673 | 3.41098722619315 | 1.19322821036799 | 1.06710864697714 |
| chr15 | 90271879 | 90271903 | + | 5.01980288436682 | 4.54798296825753 | 1.19322821036799 | 4.26843458790854 |
| chr15 | 90271936 | 90271956 | + | 1.00396057687336 | 7.95897019445067 | 1.19322821036799 | 1.06710864697714 |
| chr15 | 90271984 | 90272043 | + | 11.043566345607 | 15.9179403889013 | 4.77291284147196 | 3.20132594093141 |
| chr15 | 90272138 | 90272175 | + | 160.633692299738 | 308.125846099448 | 91.8785721983352 | 138.724124107028 |
| chr15 | 90500129 | 90500131 | + | 1.00396057687336 | 2.27399148412876 | 1.19322821036799 | 1.06710864697714 |
| chr15 | 90500264 | 90500273 | + | 10.0396057687336 | 7.95897019445067 | 5.96614105183995 | 1.06710864697714 |
| chr15 | 90500309 | 90500329 | + | 31.1227778830743 | 17.0549361309657 | 13.1255103140479 | 7.46976052883995 |
| chr15 | 90500354 | 90500420 | + | 17.0673298068472 | 14.780944646837 | 25.0577924177278 | 11.7381951167485 |
| chr15 | 90500432 | 90500461 | + | 19.0752509605939 | 9.09596593651506 | 5.96614105183995 | 4.26843458790854 |
| chr15 | 90500621 | 90500639 | + | 27.1069355755808 | 23.876910583352 | 10.7390538933119 | 2.13421729395427 |
| chr15 | 90500665 | 90500684 | + | 10.0396057687336 | 6.82197445238629 | 8.35259747257593 | 2.13421729395427 |
| chr15 | 90500771 | 90500948 | + | 245.970341333974 | 183.056314472366 | 176.597775134462 | 193.146665102862 |
| chr15 | 90500972 | 90501036 | + | 19.0752509605939 | 21.6029190992233 | 16.7051949451519 | 14.9395210576799 |
| chr15 | 90501058 | 90501072 | + | 9.03564519186028 | 11.3699574206438 | 8.35259747257593 | 6.40265188186281 |
| chr15 | 90501099 | 90501126 | + | 13.0514874993537 | 10.2329616785794 | 5.96614105183995 | 1.06710864697714 |
| chr15 | 90501149 | 90501241 | + | 120.475269224804 | 101.19262104373 | 215.974306076606 | 266.777161744284 |
| chr15 | 90501267 | 90501334 | + | 426.68324517118 | 422.96241604795 | 190.916513658878 | 99.2411041688736 |
| chr15 | 90501368 | 90501378 | + | 12.0475269224804 | 9.09596593651506 | 1.19322821036799 | 1.06710864697714 |
| chr15 | 90501662 | 90501678 | + | 4.01584230749346 | 3.41098722619315 | 1.19322821036799 | 6.40265188186281 |
| chr15 | 90502030 | 90502046 | + | 6.02376346124019 | 11.3699574206438 | 1.19322821036799 | 2.13421729395427 |
| chr15 | 90642220 | 90642223 | + | 2.00792115374673 | 2.27399148412876 | 2.38645642073598 | 1.06710864697714 |
| chr15 | 90642613 | 90642652 | + | 32.1267384599477 | 23.876910583352 | 1.19322821036799 | 2.13421729395427 |
| chr15 | 90644571 | 90644588 | + | 5.01980288436682 | 19.3289276150945 | 8.35259747257593 | 3.20132594093141 |
| chr15 | 90644692 | 90644818 | + | 47.1861471130481 | 64.8087572976698 | 16.7051949451519 | 10.6710864697714 |
| chr15 | 90644850 | 90644864 | + | 1.00396057687336 | 4.54798296825753 | 1.19322821036799 | 1.06710864697714 |
| chr15 | 90644940 | 90644994 | + | 293.156488447022 | 337.687735393121 | 14.3187385244159 | 25.6106075274513 |
| chr15 | 90815415 | 90815418 | + | 9.03564519186028 | 22.7399148412876 | 20.2848795762558 | 9.60397782279422 |
| chr15 | 90882664 | 90882705 | + | 4.01584230749346 | 3.41098722619315 | 7.15936926220794 | 7.46976052883995 |
| chr15 | 90883312 | 90883345 | + | 8.03168461498692 | 7.95897019445067 | 11.9322821036799 | 19.2079556455884 |
| chr15 | 90883371 | 90883438 | + | 4.01584230749346 | 4.54798296825753 | 14.3187385244159 | 8.53686917581709 |
| chr15 | 90919973 | 90920029 | + | 9.03564519186028 | 13.6439489047726 | 44.1494437836156 | 24.5434988804741 |
| chr15 | 90920051 | 90920199 | + | 16.0633692299738 | 14.780944646837 | 90.6853439879672 | 74.6976052883995 |
| chr15 | 90920225 | 90920313 | + | 9.03564519186028 | 3.41098722619315 | 34.6036181006717 | 51.2212150549025 |
| chr15 | 90920414 | 90920512 | + | 7.02772403811355 | 3.41098722619315 | 46.5359002043516 | 44.8185631730397 |
| chr15 | 90920529 | 90920551 | + | 7.02772403811355 | 7.95897019445067 | 35.7968463110397 | 16.006629704657 |
| chr15 | 90920792 | 90920840 | + | 6.02376346124019 | 6.82197445238629 | 11.9322821036799 | 7.46976052883995 |
| chr15 | 90920867 | 90920884 | + | 2.00792115374673 | 11.3699574206438 | 10.7390538933119 | 5.33554323488568 |
| chr15 | 90921069 | 90921072 | + | 7.02772403811355 | 4.54798296825753 | 7.15936926220794 | 9.60397782279422 |
| chr15 | 90921113 | 90921140 | + | 3.01188173062009 | 6.82197445238629 | 11.9322821036799 | 8.53686917581709 |
| chr15 | 90922290 | 90922291 | + | 1.00396057687336 | 2.27399148412876 | 1.19322821036799 | 2.13421729395427 |
| chr15 | 90922525 | 90922546 | + | 2.00792115374673 | 5.68497871032191 | 2.38645642073598 | 3.20132594093141 |
| chr15 | 90930100 | 90930131 | - | 16.0633692299738 | 29.5618892936739 | 10.7390538933119 | 10.6710864697714 |
| chr15 | 90930989 | 90931022 | - | 5.01980288436682 | 9.09596593651506 | 1.19322821036799 | 6.40265188186281 |
| chr15 | 90953716 | 90953739 | + | 3.01188173062009 | 1.13699574206438 | 1.19322821036799 | 2.13421729395427 |
| chr15 | 90953959 | 90953974 | + | 5.01980288436682 | 1.13699574206438 | 3.57968463110397 | 1.06710864697714 |
| chr15 | 90954045 | 90954062 | + | 2.00792115374673 | 1.13699574206438 | 1.19322821036799 | 3.20132594093141 |
| chr15 | 90961748 | 90961775 | + | 4.01584230749346 | 9.09596593651506 | 9.54582568294392 | 8.53686917581709 |
| chr15 | 90961847 | 90961919 | + | 35.1386201905678 | 30.6988850357383 | 15.5119667347839 | 27.7448248214055 |
| chr15 | 90962066 | 90962124 | + | 18.0712903837206 | 4.54798296825753 | 3.57968463110397 | 5.33554323488568 |
| chr15 | 90962198 | 90962212 | + | 4.01584230749346 | 2.27399148412876 | 3.57968463110397 | 3.20132594093141 |
| chr15 | 90962277 | 90962311 | + | 4.01584230749346 | 9.09596593651506 | 3.57968463110397 | 3.20132594093141 |
| chr15 | 90962661 | 90962685 | + | 20.0792115374673 | 27.2878978095452 | 2.38645642073598 | 7.46976052883995 |
| chr15 | 90966059 | 90966062 | - | 3.01188173062009 | 5.68497871032191 | 2.38645642073598 | 1.06710864697714 |
| chr15 | 90966106 | 90966151 | - | 56.2217923049084 | 37.5208594881246 | 25.0577924177278 | 19.2079556455884 |
| chr15 | 90966271 | 90966337 | - | 209.827760566533 | 264.920007901001 | 150.346754506367 | 172.871600810296 |
| chr15 | 90966358 | 90966399 | - | 8.03168461498692 | 7.95897019445067 | 1.19322821036799 | 2.13421729395427 |
| chr15 | 90966508 | 90966510 | - | 1.00396057687336 | 1.13699574206438 | 1.19322821036799 | 1.06710864697714 |
| chr15 | 90967068 | 90967086 | - | 6.02376346124019 | 4.54798296825753 | 2.38645642073598 | 4.26843458790854 |
| chr15 | 90998732 | 90998741 | - | 6.02376346124019 | 7.95897019445067 | 8.35259747257593 | 12.8053037637256 |
| chr15 | 91022603 | 91022671 | + | 6.02376346124019 | 7.95897019445067 | 13.1255103140479 | 13.8724124107028 |
| chr15 | 92882662 | 92882876 | + | 531.09514516601 | 2633.28213862111 | 400.924678683644 | 550.628061840202 |
| chr15 | 92883598 | 92883629 | + | 10.0396057687336 | 18.1919318730301 | 1.19322821036799 | 5.33554323488568 |
| chr15 | 92885542 | 92885586 | + | 13.0514874993537 | 6.82197445238629 | 13.1255103140479 | 10.6710864697714 |
| chr15 | 92886189 | 92886220 | + | 7.02772403811355 | 21.6029190992233 | 2.38645642073598 | 6.40265188186281 |
| chr15 | 92892058 | 92892073 | + | 19.0752509605939 | 72.7677274921204 | 13.1255103140479 | 11.7381951167485 |
| chr15 | 92897774 | 92897823 | + | 26.1029749987075 | 88.6856678810218 | 11.9322821036799 | 12.8053037637256 |
| chr15 | 92897829 | 92897853 | + | 3.01188173062009 | 28.4248935516096 | 3.57968463110397 | 4.26843458790854 |
| chr15 | 92897875 | 92897887 | + | 2.00792115374673 | 4.54798296825753 | 1.19322821036799 | 2.13421729395427 |
| chr15 | 92897912 | 92898047 | + | 45.1782259593014 | 185.330305956494 | 27.4442488384638 | 26.6777161744284 |
| chr15 | 92901721 | 92901722 | + | 1.00396057687336 | 3.41098722619315 | 1.19322821036799 | 1.06710864697714 |
| chr15 | 92902266 | 92902267 | + | 1.00396057687336 | 2.27399148412876 | 1.19322821036799 | 1.06710864697714 |
| chr15 | 93024953 | 93024982 | + | 1.00396057687336 | 3.41098722619315 | 2.38645642073598 | 1.06710864697714 |
| chr15 | 93025019 | 93025035 | + | 1.00396057687336 | 1.13699574206438 | 7.15936926220794 | 3.20132594093141 |
| chr15 | 93025059 | 93025061 | + | 4.01584230749346 | 5.68497871032191 | 2.38645642073598 | 2.13421729395427 |
| chr15 | 93025122 | 93025147 | + | 2.00792115374673 | 6.82197445238629 | 2.38645642073598 | 3.20132594093141 |
| chr15 | 93025216 | 93025242 | + | 16.0633692299738 | 23.876910583352 | 20.2848795762558 | 7.46976052883995 |
| chr15 | 93025269 | 93025341 | + | 8.03168461498692 | 12.5069531627082 | 14.3187385244159 | 5.33554323488568 |
| chr15 | 93025502 | 93025599 | + | 95.3762548029696 | 139.850476273919 | 110.970223564223 | 58.6909755837425 |
| chr15 | 93025623 | 93025624 | + | 1.00396057687336 | 1.13699574206438 | 1.19322821036799 | 1.06710864697714 |
| chr15 | 93025700 | 93025764 | + | 11.043566345607 | 12.5069531627082 | 8.35259747257593 | 8.53686917581709 |
| chr15 | 93027454 | 93027473 | + | 3.01188173062009 | 5.68497871032191 | 8.35259747257593 | 1.06710864697714 |
| chr15 | 93027687 | 93027918 | + | 196.776273067179 | 228.536144154941 | 279.21540122611 | 167.53605757541 |
| chr15 | 96337639 | 96337659 | + | 13.0514874993537 | 18.1919318730301 | 14.3187385244159 | 9.60397782279422 |
| chr15 | 96337744 | 96337765 | + | 68.2693192273888 | 59.1237785873479 | 56.0817258872955 | 56.5567582897882 |
| chr15 | 96337846 | 96337852 | + | 8.03168461498692 | 5.68497871032191 | 1.19322821036799 | 4.26843458790854 |
| chr15 | 96337950 | 96337977 | + | 15.0594086531005 | 12.5069531627082 | 8.35259747257593 | 13.8724124107028 |
| chr15 | 96338472 | 96338480 | + | 1.00396057687336 | 1.13699574206438 | 1.19322821036799 | 1.06710864697714 |
| chr15 | 96338730 | 96338776 | + | 15.0594086531005 | 25.0139063254164 | 7.15936926220794 | 4.26843458790854 |
| chr15 | 96339863 | 96339867 | + | 4.01584230749346 | 3.41098722619315 | 1.19322821036799 | 1.06710864697714 |
| chr15 | 97973536 | 97973538 | + | 2.00792115374673 | 2.27399148412876 | 10.7390538933119 | 6.40265188186281 |
| chr15 | 98660430 | 98660437 | + | 65.2574374967687 | 73.9047232341848 | 2.38645642073598 | 5.33554323488568 |
| chr15 | 98660468 | 98660500 | + | 12.0475269224804 | 19.3289276150945 | 3.57968463110397 | 1.06710864697714 |
| chr15 | 98957492 | 98957508 | + | 37.1465413443145 | 60.2607743294122 | 2.38645642073598 | 2.13421729395427 |
| chr15 | 98958437 | 98958455 | + | 28.1108961524542 | 46.6168254246397 | 2.38645642073598 | 1.06710864697714 |
| chr15 | 98959620 | 98959633 | + | 33.130699036821 | 34.1098722619315 | 2.38645642073598 | 1.06710864697714 |
| chr15 | 99713274 | 99713313 | + | 14.0554480762271 | 12.5069531627082 | 5.96614105183995 | 3.20132594093141 |
| chr15 | 99713369 | 99713390 | + | 8.03168461498692 | 5.68497871032191 | 1.19322821036799 | 3.20132594093141 |
| chr15 | 100650442 | 100650468 | + | 9.03564519186028 | 1.13699574206438 | 5.96614105183995 | 4.26843458790854 |
| chr15 | 100651543 | 100651565 | + | 5.01980288436682 | 2.27399148412876 | 2.38645642073598 | 3.20132594093141 |
| chr15 | 101176293 | 101176317 | - | 10.0396057687336 | 20.4659233571589 | 4.77291284147196 | 11.7381951167485 |
| chr15 | 101176347 | 101176353 | - | 8.03168461498692 | 14.780944646837 | 1.19322821036799 | 3.20132594093141 |
| chr15 | 101272222 | 101272270 | - | 5.01980288436682 | 9.09596593651506 | 4.77291284147196 | 1.06710864697714 |
| chr15 | 101272340 | 101272435 | - | 28.1108961524542 | 28.4248935516096 | 8.35259747257593 | 13.8724124107028 |
| chr15 | 101272475 | 101272477 | - | 6.02376346124019 | 4.54798296825753 | 2.38645642073598 | 3.20132594093141 |
| chr15 | 101272515 | 101272533 | - | 4.01584230749346 | 5.68497871032191 | 1.19322821036799 | 1.06710864697714 |
| chr15 | 101281706 | 101281720 | - | 234.926774988367 | 302.440867389126 | 686.106220961594 | 376.689352382929 |
| chr15 | 101641877 | 101641890 | - | 10.0396057687336 | 23.876910583352 | 11.9322821036799 | 18.1408469986113 |
| chr15 | 101641969 | 101642065 | - | 16.0633692299738 | 29.5618892936739 | 19.0916513658878 | 19.2079556455884 |
| chr15 | 101642099 | 101642109 | - | 6.02376346124019 | 7.95897019445067 | 4.77291284147196 | 6.40265188186281 |
| chr15 | 101642313 | 101642364 | - | 1.00396057687336 | 3.41098722619315 | 1.19322821036799 | 4.26843458790854 |
| chr15 | 101642398 | 101642423 | - | 3.01188173062009 | 2.27399148412876 | 3.57968463110397 | 4.26843458790854 |
| chr16 | 47213 | 47263 | - | 9.03564519186028 | 15.9179403889013 | 17.8984231555198 | 35.2145853502455 |
| chr16 | 47289 | 47291 | - | 2.00792115374673 | 3.41098722619315 | 1.19322821036799 | 8.53686917581709 |
| chr16 | 57296 | 57320 | + | 2.00792115374673 | 10.2329616785794 | 4.77291284147196 | 10.6710864697714 |
| chr16 | 57353 | 57363 | + | 1.00396057687336 | 2.27399148412876 | 2.38645642073598 | 3.20132594093141 |
| chr16 | 57394 | 57436 | + | 28.1108961524542 | 19.3289276150945 | 41.7629873628796 | 71.4962793474681 |
| chr16 | 58080 | 58085 | - | 3.01188173062009 | 4.54798296825753 | 1.19322821036799 | 1.06710864697714 |
| chr16 | 85731 | 85825 | + | 65.2574374967687 | 36.3838637460602 | 50.1155848354556 | 80.0331485232852 |
| chr16 | 85852 | 85905 | - | 3.01188173062009 | 10.2329616785794 | 14.3187385244159 | 25.6106075274513 |
| chr16 | 189018 | 189093 | - | 3.01188173062009 | 23.876910583352 | 20.2848795762558 | 9.60397782279422 |
| chr16 | 189182 | 189293 | - | 19.0752509605939 | 19.3289276150945 | 19.0916513658878 | 25.6106075274513 |
| chr16 | 189980 | 189981 | - | 1.00396057687336 | 2.27399148412876 | 1.19322821036799 | 2.13421729395427 |
| chr16 | 190106 | 190107 | - | 6.02376346124019 | 5.68497871032191 | 9.54582568294392 | 4.26843458790854 |
| chr16 | 192926 | 192990 | - | 10.0396057687336 | 7.95897019445067 | 9.54582568294392 | 17.0737383516342 |
| chr16 | 206075 | 206085 | - | 3.01188173062009 | 3.41098722619315 | 3.57968463110397 | 1.06710864697714 |
| chr16 | 208093 | 208149 | - | 5.01980288436682 | 2.27399148412876 | 15.5119667347839 | 13.8724124107028 |
| chr16 | 220648 | 220702 | - | 6.02376346124019 | 2.27399148412876 | 9.54582568294392 | 9.60397782279422 |
| chr16 | 227297 | 227339 | - | 82.3247673036159 | 50.0278126508328 | 41.7629873628796 | 64.0265188186281 |
| chr16 | 265995 | 266069 | + | 3.01188173062009 | 5.68497871032191 | 5.96614105183995 | 3.20132594093141 |
| chr16 | 287514 | 287555 | - | 6.02376346124019 | 7.95897019445067 | 8.35259747257593 | 13.8724124107028 |
| chr16 | 288019 | 288039 | - | 3.01188173062009 | 2.27399148412876 | 3.57968463110397 | 3.20132594093141 |
| chr16 | 367403 | 367479 | - | 39.1544624980612 | 23.876910583352 | 13.1255103140479 | 51.2212150549025 |
| chr16 | 370880 | 370917 | - | 4.01584230749346 | 4.54798296825753 | 8.35259747257593 | 4.26843458790854 |
| chr16 | 371092 | 371094 | - | 1.00396057687336 | 1.13699574206438 | 2.38645642073598 | 1.06710864697714 |
| chr16 | 371889 | 371927 | - | 3.01188173062009 | 2.27399148412876 | 4.77291284147196 | 13.8724124107028 |
| chr16 | 400562 | 400588 | + | 12.0475269224804 | 12.5069531627082 | 38.1833027317757 | 68.2949534065367 |
| chr16 | 400616 | 400737 | + | 33.130699036821 | 32.9728765198671 | 116.936364616063 | 160.06629704657 |
| chr16 | 553580 | 553594 | + | 6.02376346124019 | 2.27399148412876 | 1.19322821036799 | 6.40265188186281 |
| chr16 | 553986 | 554002 | + | 4.01584230749346 | 2.27399148412876 | 10.7390538933119 | 1.06710864697714 |
| chr16 | 554412 | 554447 | + | 9.03564519186028 | 4.54798296825753 | 10.7390538933119 | 7.46976052883995 |
| chr16 | 568937 | 568967 | + | 4.01584230749346 | 7.95897019445067 | 5.96614105183995 | 8.53686917581709 |
| chr16 | 569719 | 569728 | + | 2.00792115374673 | 3.41098722619315 | 1.19322821036799 | 2.13421729395427 |
| chr16 | 577151 | 577154 | + | 3.01188173062009 | 4.54798296825753 | 11.9322821036799 | 6.40265188186281 |
| chr16 | 583445 | 583465 | + | 7.02772403811355 | 9.09596593651506 | 3.57968463110397 | 4.26843458790854 |
| chr16 | 583590 | 583604 | + | 2.00792115374673 | 3.41098722619315 | 2.38645642073598 | 3.20132594093141 |
| chr16 | 583847 | 583883 | + | 13.0514874993537 | 13.6439489047726 | 3.57968463110397 | 1.06710864697714 |
| chr16 | 583952 | 584083 | + | 45.1782259593014 | 44.3428339405109 | 8.35259747257593 | 13.8724124107028 |
| chr16 | 627895 | 627906 | + | 3.01188173062009 | 4.54798296825753 | 1.19322821036799 | 3.20132594093141 |
| chr16 | 628879 | 628897 | + | 4.01584230749346 | 2.27399148412876 | 1.19322821036799 | 1.06710864697714 |
| chr16 | 628918 | 628942 | + | 5.01980288436682 | 3.41098722619315 | 10.7390538933119 | 1.06710864697714 |
| chr16 | 629104 | 629128 | + | 15.0594086531005 | 6.82197445238629 | 19.0916513658878 | 13.8724124107028 |
| chr16 | 629150 | 629161 | + | 13.0514874993537 | 3.41098722619315 | 10.7390538933119 | 11.7381951167485 |
| chr16 | 634458 | 634662 | - | 438.73077209366 | 319.495803520091 | 251.771152387646 | 334.005006503844 |
| chr16 | 648428 | 648453 | + | 16.0633692299738 | 13.6439489047726 | 16.7051949451519 | 17.0737383516342 |
| chr16 | 674028 | 674151 | + | 16.0633692299738 | 23.876910583352 | 25.0577924177278 | 34.1474767032683 |
| chr16 | 681722 | 681726 | - | 7.02772403811355 | 5.68497871032191 | 2.38645642073598 | 5.33554323488568 |
| chr16 | 681748 | 681788 | - | 27.1069355755808 | 20.4659233571589 | 5.96614105183995 | 16.006629704657 |
| chr16 | 681890 | 681893 | - | 26.1029749987075 | 6.82197445238629 | 2.38645642073598 | 9.60397782279422 |
| chr16 | 681990 | 681993 | - | 5.01980288436682 | 4.54798296825753 | 3.57968463110397 | 4.26843458790854 |
| chr16 | 682145 | 682155 | - | 1.00396057687336 | 1.13699574206438 | 2.38645642073598 | 2.13421729395427 |
| chr16 | 682335 | 682498 | + | 33.130699036821 | 42.0688424563821 | 79.9462900946553 | 94.9726695809651 |
| chr16 | 682591 | 682592 | - | 86.3406096111093 | 43.2058381984465 | 22.6713359969918 | 32.0132594093141 |
| chr16 | 682682 | 682718 | + | 12.0475269224804 | 13.6439489047726 | 16.7051949451519 | 26.6777161744284 |
| chr16 | 692667 | 692686 | - | 10.0396057687336 | 18.1919318730301 | 2.38645642073598 | 6.40265188186281 |
| chr16 | 692963 | 692988 | - | 8.03168461498692 | 3.41098722619315 | 2.38645642073598 | 3.20132594093141 |
| chr16 | 717371 | 717453 | + | 12.0475269224804 | 10.2329616785794 | 44.1494437836156 | 44.8185631730397 |
| chr16 | 722532 | 722559 | + | 9.03564519186028 | 12.5069531627082 | 8.35259747257593 | 17.0737383516342 |
| chr16 | 729645 | 729689 | + | 7.02772403811355 | 4.54798296825753 | 4.77291284147196 | 14.9395210576799 |
| chr16 | 785008 | 785020 | - | 4.01584230749346 | 2.27399148412876 | 5.96614105183995 | 5.33554323488568 |
| chr16 | 785604 | 785630 | - | 3.01188173062009 | 6.82197445238629 | 3.57968463110397 | 5.33554323488568 |
| chr16 | 785926 | 785929 | - | 7.02772403811355 | 3.41098722619315 | 4.77291284147196 | 1.06710864697714 |
| chr16 | 786051 | 786052 | - | 1.00396057687336 | 2.27399148412876 | 2.38645642073598 | 1.06710864697714 |
| chr16 | 797988 | 797989 | + | 1.00396057687336 | 1.13699574206438 | 1.19322821036799 | 5.33554323488568 |
| chr16 | 798017 | 798053 | + | 1.00396057687336 | 4.54798296825753 | 7.15936926220794 | 8.53686917581709 |
| chr16 | 1324737 | 1324840 | + | 38.1505019211879 | 30.6988850357383 | 50.1155848354556 | 37.3488026441998 |
| chr16 | 1324900 | 1325025 | + | 87.3445701879827 | 63.6717615556054 | 169.438405872254 | 181.408469986113 |
| chr16 | 1325084 | 1325172 | + | 5.01980288436682 | 3.41098722619315 | 2.38645642073598 | 4.26843458790854 |
| chr16 | 1325229 | 1325233 | + | 21.0831721143407 | 22.7399148412876 | 10.7390538933119 | 39.483019938154 |
| chr16 | 1325259 | 1325323 | + | 31.1227778830743 | 37.5208594881246 | 14.3187385244159 | 11.7381951167485 |
| chr16 | 1349377 | 1349519 | - | 175.693100952839 | 94.3706465913437 | 72.7869208324474 | 124.851711696325 |
| chr16 | 1363148 | 1363197 | + | 12.0475269224804 | 3.41098722619315 | 2.38645642073598 | 4.26843458790854 |
| chr16 | 1363210 | 1363231 | - | 1.00396057687336 | 2.27399148412876 | 2.38645642073598 | 1.06710864697714 |
| chr16 | 1363439 | 1363465 | - | 2.00792115374673 | 1.13699574206438 | 3.57968463110397 | 4.26843458790854 |
| chr16 | 1363559 | 1363583 | - | 3.01188173062009 | 1.13699574206438 | 4.77291284147196 | 5.33554323488568 |
| chr16 | 1419802 | 1419831 | - | 12.0475269224804 | 9.09596593651506 | 7.15936926220794 | 11.7381951167485 |
| chr16 | 1419994 | 1419999 | - | 5.01980288436682 | 1.13699574206438 | 2.38645642073598 | 2.13421729395427 |
| chr16 | 1445364 | 1445434 | - | 2.00792115374673 | 18.1919318730301 | 3.57968463110397 | 7.46976052883995 |
| chr16 | 1446440 | 1446477 | - | 1.00396057687336 | 1.13699574206438 | 2.38645642073598 | 1.06710864697714 |
| chr16 | 1446481 | 1446484 | - | 5.01980288436682 | 3.41098722619315 | 3.57968463110397 | 1.06710864697714 |
| chr16 | 1510110 | 1510122 | + | 15.0594086531005 | 4.54798296825753 | 2.38645642073598 | 2.13421729395427 |
| chr16 | 1510144 | 1510164 | + | 5.01980288436682 | 5.68497871032191 | 7.15936926220794 | 6.40265188186281 |
| chr16 | 1510190 | 1510215 | + | 28.1108961524542 | 14.780944646837 | 28.6374770488317 | 12.8053037637256 |
| chr16 | 1510402 | 1510432 | + | 13.0514874993537 | 7.95897019445067 | 13.1255103140479 | 26.6777161744284 |
| chr16 | 1685437 | 1685472 | + | 11.043566345607 | 10.2329616785794 | 50.1155848354556 | 12.8053037637256 |
| chr16 | 1685509 | 1685526 | + | 3.01188173062009 | 2.27399148412876 | 4.77291284147196 | 3.20132594093141 |
| chr16 | 1685568 | 1685588 | + | 2.00792115374673 | 1.13699574206438 | 3.57968463110397 | 4.26843458790854 |
| chr16 | 1697829 | 1697850 | + | 1.00396057687336 | 1.13699574206438 | 7.15936926220794 | 1.06710864697714 |
| chr16 | 1698926 | 1699099 | + | 39.1544624980612 | 26.1509020674808 | 50.1155848354556 | 60.8251928776967 |
| chr16 | 1699178 | 1699332 | + | 1446.70719127452 | 1049.44706992542 | 652.69583107129 | 735.237857767247 |
| chr16 | 1700183 | 1700221 | + | 3.01188173062009 | 4.54798296825753 | 10.7390538933119 | 8.53686917581709 |
| chr16 | 1700264 | 1700290 | + | 4.01584230749346 | 3.41098722619315 | 4.77291284147196 | 5.33554323488568 |
| chr16 | 1700363 | 1700400 | + | 4.01584230749346 | 3.41098722619315 | 7.15936926220794 | 4.26843458790854 |
| chr16 | 1700523 | 1700878 | + | 88.3485307648561 | 72.7677274921204 | 173.018090503358 | 201.683534278679 |
| chr16 | 1700901 | 1700986 | + | 37.1465413443145 | 36.3838637460602 | 66.8207797806074 | 84.3015831111937 |
| chr16 | 1701034 | 1701085 | + | 23.0910932680874 | 17.0549361309657 | 28.6374770488317 | 34.1474767032683 |
| chr16 | 1701336 | 1701397 | + | 17.0673298068472 | 25.0139063254164 | 19.0916513658878 | 20.2750642925656 |
| chr16 | 1770224 | 1770294 | + | 14.0554480762271 | 15.9179403889013 | 10.7390538933119 | 20.2750642925656 |
| chr16 | 1770587 | 1770595 | - | 1.00396057687336 | 1.13699574206438 | 10.7390538933119 | 1.06710864697714 |
| chr16 | 1771913 | 1771953 | - | 32.1267384599477 | 48.8908169087684 | 19.0916513658878 | 25.6106075274513 |
| chr16 | 1771975 | 1772003 | - | 30.1188173062009 | 21.6029190992233 | 10.7390538933119 | 28.8119334683827 |
| chr16 | 1772054 | 1772206 | - | 318.255502868857 | 180.782322988237 | 188.530057238142 | 208.086186160541 |
| chr16 | 1776729 | 1776766 | - | 3.01188173062009 | 5.68497871032191 | 1.19322821036799 | 4.26843458790854 |
| chr16 | 1777081 | 1777119 | - | 4.01584230749346 | 4.54798296825753 | 10.7390538933119 | 6.40265188186281 |
| chr16 | 1788741 | 1788773 | + | 6.02376346124019 | 2.27399148412876 | 8.35259747257593 | 40.5501285851312 |
| chr16 | 1788811 | 1788836 | + | 7.02772403811355 | 2.27399148412876 | 10.7390538933119 | 7.46976052883995 |
| chr16 | 1788943 | 1788965 | + | 17.0673298068472 | 7.95897019445067 | 7.15936926220794 | 1.06710864697714 |
| chr16 | 1789049 | 1789164 | + | 250.990144218341 | 246.728076027971 | 220.747218918078 | 248.636314745673 |
| chr16 | 1809124 | 1809223 | - | 6.02376346124019 | 17.0549361309657 | 10.7390538933119 | 10.6710864697714 |
| chr16 | 1809246 | 1809350 | - | 358.413925943791 | 350.19468855583 | 46.5359002043516 | 28.8119334683827 |
| chr16 | 1809372 | 1809385 | - | 2.00792115374673 | 3.41098722619315 | 2.38645642073598 | 2.13421729395427 |
| chr16 | 1828842 | 1828864 | + | 2.00792115374673 | 9.09596593651506 | 1.19322821036799 | 2.13421729395427 |
| chr16 | 1938260 | 1938275 | - | 5.01980288436682 | 3.41098722619315 | 5.96614105183995 | 9.60397782279422 |
| chr16 | 1938306 | 1938340 | - | 1.00396057687336 | 2.27399148412876 | 1.19322821036799 | 6.40265188186281 |
| chr16 | 1938800 | 1938829 | - | 3.01188173062009 | 1.13699574206438 | 10.7390538933119 | 5.33554323488568 |
| chr16 | 1961799 | 1961953 | + | 135.534677877904 | 78.4527062024424 | 127.675418509375 | 205.951968866587 |
| chr16 | 1962063 | 1962087 | - | 4.01584230749346 | 4.54798296825753 | 7.15936926220794 | 5.33554323488568 |
| chr16 | 1962176 | 1962186 | - | 36.1425807674411 | 25.0139063254164 | 31.0239334695677 | 51.2212150549025 |
| chr16 | 1964994 | 1965084 | + | 119.47130864793 | 93.2336508492793 | 77.5598336739193 | 151.529427870753 |
| chr16 | 1965449 | 1965477 | + | 23.0910932680874 | 31.8358807778027 | 13.1255103140479 | 18.1408469986113 |
| chr16 | 1978335 | 1978356 | + | 3.01188173062009 | 1.13699574206438 | 1.19322821036799 | 5.33554323488568 |
| chr16 | 1986030 | 1986086 | + | 11.043566345607 | 10.2329616785794 | 9.54582568294392 | 9.60397782279422 |
| chr16 | 1986464 | 1986484 | + | 4.01584230749346 | 3.41098722619315 | 7.15936926220794 | 1.06710864697714 |
| chr16 | 1986767 | 1986798 | + | 2.00792115374673 | 4.54798296825753 | 1.19322821036799 | 7.46976052883995 |
| chr16 | 1986848 | 1986855 | + | 2.00792115374673 | 3.41098722619315 | 10.7390538933119 | 5.33554323488568 |
| chr16 | 1987035 | 1987062 | + | 4.01584230749346 | 3.41098722619315 | 13.1255103140479 | 9.60397782279422 |
| chr16 | 1987098 | 1987160 | + | 4.01584230749346 | 5.68497871032191 | 10.7390538933119 | 24.5434988804741 |
| chr16 | 1987242 | 1987246 | + | 4.01584230749346 | 5.68497871032191 | 2.38645642073598 | 3.20132594093141 |
| chr16 | 1987403 | 1987405 | + | 3.01188173062009 | 3.41098722619315 | 5.96614105183995 | 1.06710864697714 |
| chr16 | 1987495 | 1987524 | + | 5.01980288436682 | 5.68497871032191 | 5.96614105183995 | 4.26843458790854 |
| chr16 | 1997712 | 1997730 | - | 6.02376346124019 | 7.95897019445067 | 7.15936926220794 | 5.33554323488568 |
| chr16 | 2038003 | 2038029 | + | 6.02376346124019 | 1.13699574206438 | 4.77291284147196 | 11.7381951167485 |
| chr16 | 2038051 | 2038118 | + | 31.1227778830743 | 19.3289276150945 | 46.5359002043516 | 35.2145853502455 |
| chr16 | 2038147 | 2038203 | + | 52.205949997415 | 53.438799877026 | 106.197310722751 | 66.1607361125824 |
| chr16 | 2038437 | 2038444 | + | 11.043566345607 | 6.82197445238629 | 8.35259747257593 | 13.8724124107028 |
| chr16 | 2039834 | 2039939 | - | 16.0633692299738 | 18.1919318730301 | 2.38645642073598 | 14.9395210576799 |
| chr16 | 2089087 | 2089106 | - | 1.00396057687336 | 5.68497871032191 | 7.15936926220794 | 5.33554323488568 |
| chr16 | 2089270 | 2089288 | - | 3.01188173062009 | 6.82197445238629 | 17.8984231555198 | 20.2750642925656 |
| chr16 | 2089625 | 2089645 | - | 2.00792115374673 | 3.41098722619315 | 8.35259747257593 | 9.60397782279422 |
| chr16 | 2089681 | 2089707 | - | 4.01584230749346 | 5.68497871032191 | 15.5119667347839 | 1.06710864697714 |
| chr16 | 2176840 | 2176854 | + | 4.01584230749346 | 6.82197445238629 | 17.8984231555198 | 19.2079556455884 |
| chr16 | 2176875 | 2176912 | + | 7.02772403811355 | 11.3699574206438 | 8.35259747257593 | 9.60397782279422 |
| chr16 | 2177272 | 2177287 | + | 1.00396057687336 | 1.13699574206438 | 3.57968463110397 | 1.06710864697714 |
| chr16 | 2177396 | 2177411 | + | 1.00396057687336 | 1.13699574206438 | 5.96614105183995 | 6.40265188186281 |
| chr16 | 2177943 | 2178019 | + | 12.0475269224804 | 19.3289276150945 | 19.0916513658878 | 9.60397782279422 |
| chr16 | 2208779 | 2208863 | + | 13.0514874993537 | 6.82197445238629 | 39.3765309421437 | 55.4896496428111 |
| chr16 | 2209011 | 2209046 | + | 4.01584230749346 | 3.41098722619315 | 15.5119667347839 | 6.40265188186281 |
| chr16 | 2209103 | 2209106 | + | 4.01584230749346 | 3.41098722619315 | 4.77291284147196 | 6.40265188186281 |
| chr16 | 2209335 | 2209393 | + | 23.0910932680874 | 38.657855230189 | 15.5119667347839 | 21.3421729395427 |
| chr16 | 2212026 | 2212044 | - | 8.03168461498692 | 7.95897019445067 | 7.15936926220794 | 9.60397782279422 |
| chr16 | 2213704 | 2213763 | - | 79.3128855729958 | 129.61751459534 | 64.4343233598714 | 38.4159112911769 |
| chr16 | 2235653 | 2235692 | + | 10.0396057687336 | 10.2329616785794 | 8.35259747257593 | 20.2750642925656 |
| chr16 | 2239922 | 2239962 | - | 47.1861471130481 | 77.315710460378 | 46.5359002043516 | 61.8923015246739 |
| chr16 | 2239971 | 2240013 | - | 4.01584230749346 | 10.2329616785794 | 2.38645642073598 | 8.53686917581709 |
| chr16 | 2253239 | 2253246 | - | 4.01584230749346 | 2.27399148412876 | 1.19322821036799 | 5.33554323488568 |
| chr16 | 2253322 | 2253381 | - | 48.1901076899215 | 55.7127913611547 | 26.2510206280958 | 38.4159112911769 |
| chr16 | 2276355 | 2276397 | - | 3.01188173062009 | 1.13699574206438 | 13.1255103140479 | 9.60397782279422 |
| chr16 | 2276635 | 2276636 | - | 1.00396057687336 | 1.13699574206438 | 3.57968463110397 | 1.06710864697714 |
| chr16 | 2457059 | 2457093 | + | 8.03168461498692 | 2.27399148412876 | 22.6713359969918 | 6.40265188186281 |
| chr16 | 2457540 | 2457605 | + | 25.0990144218341 | 5.68497871032191 | 9.54582568294392 | 19.2079556455884 |
| chr16 | 2457667 | 2457682 | + | 8.03168461498692 | 7.95897019445067 | 23.8645642073598 | 2.13421729395427 |
| chr16 | 2457750 | 2457800 | + | 3.01188173062009 | 4.54798296825753 | 7.15936926220794 | 9.60397782279422 |
| chr16 | 2457904 | 2457925 | + | 1.00396057687336 | 2.27399148412876 | 10.7390538933119 | 30.9461507623369 |
| chr16 | 2458234 | 2458248 | + | 7.02772403811355 | 4.54798296825753 | 9.54582568294392 | 30.9461507623369 |
| chr16 | 2458794 | 2458796 | + | 4.01584230749346 | 2.27399148412876 | 2.38645642073598 | 3.20132594093141 |
| chr16 | 2458817 | 2458828 | + | 10.0396057687336 | 3.41098722619315 | 10.7390538933119 | 10.6710864697714 |
| chr16 | 2464697 | 2464712 | + | 6.02376346124019 | 4.54798296825753 | 9.54582568294392 | 7.46976052883995 |
| chr16 | 2502307 | 2502328 | + | 2.00792115374673 | 3.41098722619315 | 7.15936926220794 | 1.06710864697714 |
| chr16 | 2502701 | 2502713 | + | 4.01584230749346 | 2.27399148412876 | 10.7390538933119 | 4.26843458790854 |
| chr16 | 2502864 | 2502865 | + | 1.00396057687336 | 1.13699574206438 | 1.19322821036799 | 2.13421729395427 |
| chr16 | 2502897 | 2502901 | + | 1.00396057687336 | 1.13699574206438 | 1.19322821036799 | 3.20132594093141 |
| chr16 | 2503841 | 2503857 | + | 30.1188173062009 | 43.2058381984465 | 10.7390538933119 | 7.46976052883995 |
| chr16 | 2519617 | 2519636 | + | 2.00792115374673 | 1.13699574206438 | 4.77291284147196 | 6.40265188186281 |
| chr16 | 2519657 | 2519695 | + | 2.00792115374673 | 1.13699574206438 | 4.77291284147196 | 4.26843458790854 |
| chr16 | 2519848 | 2519884 | + | 2.00792115374673 | 1.13699574206438 | 2.38645642073598 | 5.33554323488568 |
| chr16 | 2519912 | 2520013 | + | 14.0554480762271 | 13.6439489047726 | 7.15936926220794 | 25.6106075274513 |
| chr16 | 2520039 | 2520197 | + | 203.803997105293 | 176.234340019979 | 128.868646719743 | 220.891489924267 |
| chr16 | 2529619 | 2529627 | + | 1.00396057687336 | 1.13699574206438 | 2.38645642073598 | 1.06710864697714 |
| chr16 | 2530511 | 2530521 | + | 3.01188173062009 | 1.13699574206438 | 8.35259747257593 | 38.4159112911769 |
| chr16 | 2531214 | 2531244 | + | 12.0475269224804 | 7.95897019445067 | 2.38645642073598 | 4.26843458790854 |
| chr16 | 2531251 | 2531293 | + | 28.1108961524542 | 11.3699574206438 | 7.15936926220794 | 20.2750642925656 |
| chr16 | 2531335 | 2531360 | + | 10.0396057687336 | 14.780944646837 | 5.96614105183995 | 7.46976052883995 |
| chr16 | 2597778 | 2597838 | + | 1.00396057687336 | 1.13699574206438 | 2.38645642073598 | 1.06710864697714 |
| chr16 | 2597870 | 2597898 | + | 30.1188173062009 | 27.2878978095452 | 15.5119667347839 | 13.8724124107028 |
| chr16 | 2602576 | 2602590 | + | 1.00396057687336 | 1.13699574206438 | 1.19322821036799 | 1.06710864697714 |
| chr16 | 2602939 | 2603076 | + | 5.01980288436682 | 12.5069531627082 | 8.35259747257593 | 6.40265188186281 |
| chr16 | 2708513 | 2708525 | + | 3.01188173062009 | 2.27399148412876 | 7.15936926220794 | 9.60397782279422 |
| chr16 | 2708561 | 2708596 | + | 2.00792115374673 | 3.41098722619315 | 11.9322821036799 | 8.53686917581709 |
| chr16 | 2708631 | 2708638 | + | 1.00396057687336 | 1.13699574206438 | 3.57968463110397 | 10.6710864697714 |
| chr16 | 2708684 | 2708691 | + | 1.00396057687336 | 1.13699574206438 | 8.35259747257593 | 2.13421729395427 |
| chr16 | 2708780 | 2708830 | + | 16.0633692299738 | 5.68497871032191 | 14.3187385244159 | 17.0737383516342 |
| chr16 | 2770935 | 2770947 | + | 2.00792115374673 | 1.13699574206438 | 1.19322821036799 | 3.20132594093141 |
| chr16 | 2770969 | 2770996 | + | 7.02772403811355 | 2.27399148412876 | 9.54582568294392 | 4.26843458790854 |
| chr16 | 2771085 | 2771123 | + | 23.0910932680874 | 25.0139063254164 | 15.5119667347839 | 26.6777161744284 |
| chr16 | 2771160 | 2771230 | + | 6.02376346124019 | 5.68497871032191 | 7.15936926220794 | 24.5434988804741 |
| chr16 | 2771249 | 2771328 | + | 18.0712903837206 | 21.6029190992233 | 5.96614105183995 | 21.3421729395427 |
| chr16 | 2771349 | 2771388 | + | 14.0554480762271 | 21.6029190992233 | 22.6713359969918 | 17.0737383516342 |
| chr16 | 2771455 | 2771481 | - | 8.03168461498692 | 4.54798296825753 | 3.57968463110397 | 10.6710864697714 |
| chr16 | 2771927 | 2771947 | - | 4.01584230749346 | 3.41098722619315 | 5.96614105183995 | 2.13421729395427 |
| chr16 | 2771974 | 2771999 | - | 22.087132691214 | 27.2878978095452 | 14.3187385244159 | 45.8856718200168 |
| chr16 | 2777375 | 2777389 | + | 11.043566345607 | 7.95897019445067 | 7.15936926220794 | 5.33554323488568 |
| chr16 | 2899221 | 2899254 | + | 10.0396057687336 | 21.6029190992233 | 4.77291284147196 | 8.53686917581709 |
| chr16 | 2949752 | 2949754 | + | 9.03564519186028 | 2.27399148412876 | 1.19322821036799 | 3.20132594093141 |
| chr16 | 2972452 | 2972475 | + | 5.01980288436682 | 12.5069531627082 | 2.38645642073598 | 2.13421729395427 |
| chr16 | 2972504 | 2972517 | + | 3.01188173062009 | 4.54798296825753 | 4.77291284147196 | 1.06710864697714 |
| chr16 | 2972635 | 2972640 | + | 7.02772403811355 | 5.68497871032191 | 3.57968463110397 | 1.06710864697714 |
| chr16 | 2972845 | 2972885 | - | 7.02772403811355 | 14.780944646837 | 7.15936926220794 | 8.53686917581709 |
| chr16 | 2973434 | 2973459 | + | 46.1821865361748 | 102.329616785794 | 11.9322821036799 | 9.60397782279422 |
| chr16 | 3021938 | 3021966 | + | 20.0792115374673 | 43.2058381984465 | 3.57968463110397 | 1.06710864697714 |
| chr16 | 3022307 | 3022361 | + | 244.966380757101 | 859.568781000673 | 84.7192029361272 | 212.35462074845 |
| chr16 | 3022649 | 3022676 | - | 20.0792115374673 | 14.780944646837 | 2.38645642073598 | 9.60397782279422 |
| chr16 | 3120464 | 3120482 | + | 4.01584230749346 | 9.09596593651506 | 2.38645642073598 | 2.13421729395427 |
| chr16 | 3263795 | 3263828 | + | 11.043566345607 | 15.9179403889013 | 4.77291284147196 | 1.06710864697714 |
| chr16 | 3291388 | 3291406 | + | 9.03564519186028 | 5.68497871032191 | 3.57968463110397 | 2.13421729395427 |
| chr16 | 3291432 | 3291435 | + | 5.01980288436682 | 15.9179403889013 | 3.57968463110397 | 1.06710864697714 |
| chr16 | 3382509 | 3382510 | - | 1.00396057687336 | 1.13699574206438 | 1.19322821036799 | 3.20132594093141 |
| chr16 | 3382531 | 3382532 | - | 2.00792115374673 | 1.13699574206438 | 1.19322821036799 | 4.26843458790854 |
| chr16 | 3457979 | 3458164 | + | 183.724785567826 | 261.509020674808 | 105.004082512383 | 191.012447808907 |
| chr16 | 3485785 | 3485786 | + | 3.01188173062009 | 2.27399148412876 | 3.57968463110397 | 1.06710864697714 |
| chr16 | 3485840 | 3485850 | + | 3.01188173062009 | 4.54798296825753 | 5.96614105183995 | 1.06710864697714 |
| chr16 | 3486406 | 3486438 | + | 5.01980288436682 | 1.13699574206438 | 8.35259747257593 | 5.33554323488568 |
| chr16 | 3486888 | 3486931 | + | 3.01188173062009 | 10.2329616785794 | 1.19322821036799 | 3.20132594093141 |
| chr16 | 3658073 | 3658139 | - | 52.205949997415 | 47.753821166704 | 50.1155848354556 | 116.314842520508 |
| chr16 | 3660957 | 3660964 | + | 3.01188173062009 | 1.13699574206438 | 8.35259747257593 | 13.8724124107028 |
| chr16 | 4257230 | 4257255 | - | 10.0396057687336 | 9.09596593651506 | 4.77291284147196 | 4.26843458790854 |
| chr16 | 4257364 | 4257371 | - | 2.00792115374673 | 3.41098722619315 | 1.19322821036799 | 1.06710864697714 |
| chr16 | 4257774 | 4257828 | - | 28.1108961524542 | 20.4659233571589 | 17.8984231555198 | 32.0132594093141 |
| chr16 | 4257844 | 4257869 | - | 7.02772403811355 | 3.41098722619315 | 4.77291284147196 | 7.46976052883995 |
| chr16 | 4257898 | 4257933 | - | 5.01980288436682 | 2.27399148412876 | 9.54582568294392 | 1.06710864697714 |
| chr16 | 4257978 | 4258028 | - | 45.1782259593014 | 19.3289276150945 | 29.8307052591997 | 34.1474767032683 |
| chr16 | 4608903 | 4608904 | - | 3.01188173062009 | 1.13699574206438 | 2.38645642073598 | 1.06710864697714 |
| chr16 | 4608935 | 4608946 | - | 45.1782259593014 | 29.5618892936739 | 15.5119667347839 | 23.476390233497 |
| chr16 | 4609010 | 4609051 | - | 4.01584230749346 | 4.54798296825753 | 3.57968463110397 | 5.33554323488568 |
| chr16 | 4609383 | 4609403 | - | 7.02772403811355 | 2.27399148412876 | 5.96614105183995 | 2.13421729395427 |
| chr16 | 4689495 | 4689509 | + | 5.01980288436682 | 6.82197445238629 | 2.38645642073598 | 1.06710864697714 |
| chr16 | 4689692 | 4689756 | + | 6.02376346124019 | 4.54798296825753 | 2.38645642073598 | 3.20132594093141 |
| chr16 | 4689977 | 4689992 | + | 1.00396057687336 | 5.68497871032191 | 7.15936926220794 | 2.13421729395427 |
| chr16 | 4690058 | 4690103 | + | 10.0396057687336 | 6.82197445238629 | 11.9322821036799 | 5.33554323488568 |
| chr16 | 4690127 | 4690195 | + | 26.1029749987075 | 10.2329616785794 | 25.0577924177278 | 21.3421729395427 |
| chr16 | 4690861 | 4690929 | + | 2.00792115374673 | 9.09596593651506 | 5.96614105183995 | 2.13421729395427 |
| chr16 | 4694738 | 4694740 | + | 1.00396057687336 | 2.27399148412876 | 2.38645642073598 | 2.13421729395427 |
| chr16 | 4695782 | 4695785 | + | 18.0712903837206 | 18.1919318730301 | 8.35259747257593 | 17.0737383516342 |
| chr16 | 4696552 | 4696595 | - | 2.00792115374673 | 3.41098722619315 | 2.38645642073598 | 2.13421729395427 |
| chr16 | 4696623 | 4696625 | - | 7.02772403811355 | 9.09596593651506 | 3.57968463110397 | 4.26843458790854 |
| chr16 | 4796995 | 4797045 | - | 5.01980288436682 | 12.5069531627082 | 3.57968463110397 | 7.46976052883995 |
| chr16 | 4797067 | 4797104 | - | 11.043566345607 | 7.95897019445067 | 8.35259747257593 | 14.9395210576799 |
| chr16 | 4803230 | 4803231 | - | 2.00792115374673 | 1.13699574206438 | 2.38645642073598 | 1.06710864697714 |
| chr16 | 4803546 | 4803562 | - | 9.03564519186028 | 1.13699574206438 | 4.77291284147196 | 6.40265188186281 |
| chr16 | 4805049 | 4805058 | - | 2.00792115374673 | 4.54798296825753 | 4.77291284147196 | 4.26843458790854 |
| chr16 | 4805148 | 4805179 | - | 27.1069355755808 | 11.3699574206438 | 44.1494437836156 | 19.2079556455884 |
| chr16 | 4881621 | 4881634 | + | 1.00396057687336 | 6.82197445238629 | 1.19322821036799 | 12.8053037637256 |
| chr16 | 4882257 | 4882312 | + | 15.0594086531005 | 23.876910583352 | 23.8645642073598 | 42.6843458790854 |
| chr16 | 5024946 | 5025019 | - | 96.380215379843 | 61.3977700714766 | 4.77291284147196 | 5.33554323488568 |
| chr16 | 5025045 | 5025108 | - | 49.1940682667949 | 21.6029190992233 | 17.8984231555198 | 32.0132594093141 |
| chr16 | 5025159 | 5025208 | - | 4.01584230749346 | 2.27399148412876 | 4.77291284147196 | 4.26843458790854 |
| chr16 | 5033939 | 5033986 | + | 23.0910932680874 | 18.1919318730301 | 14.3187385244159 | 9.60397782279422 |
| chr16 | 5084949 | 5084966 | + | 8.03168461498692 | 6.82197445238629 | 2.38645642073598 | 5.33554323488568 |
| chr16 | 5085028 | 5085087 | + | 33.130699036821 | 29.5618892936739 | 5.96614105183995 | 10.6710864697714 |
| chr16 | 8646162 | 8646188 | + | 12.0475269224804 | 4.54798296825753 | 3.57968463110397 | 1.06710864697714 |
| chr16 | 8795483 | 8795496 | - | 11.043566345607 | 23.876910583352 | 4.77291284147196 | 17.0737383516342 |
| chr16 | 8795559 | 8795569 | - | 3.01188173062009 | 2.27399148412876 | 4.77291284147196 | 5.33554323488568 |
| chr16 | 8795599 | 8795619 | - | 2.00792115374673 | 7.95897019445067 | 5.96614105183995 | 11.7381951167485 |
| chr16 | 8848165 | 8848191 | + | 4.01584230749346 | 1.13699574206438 | 1.19322821036799 | 1.06710864697714 |
| chr16 | 8848351 | 8848385 | + | 4.01584230749346 | 1.13699574206438 | 4.77291284147196 | 4.26843458790854 |
| chr16 | 8849188 | 8849202 | + | 7.02772403811355 | 7.95897019445067 | 5.96614105183995 | 4.26843458790854 |
| chr16 | 8849232 | 8849266 | + | 1.00396057687336 | 2.27399148412876 | 1.19322821036799 | 3.20132594093141 |
| chr16 | 8849292 | 8849297 | + | 2.00792115374673 | 7.95897019445067 | 2.38645642073598 | 3.20132594093141 |
| chr16 | 8852978 | 8853103 | - | 15.0594086531005 | 17.0549361309657 | 5.96614105183995 | 10.6710864697714 |
| chr16 | 8853509 | 8853529 | - | 9.03564519186028 | 2.27399148412876 | 3.57968463110397 | 3.20132594093141 |
| chr16 | 8853564 | 8853618 | - | 5.01980288436682 | 1.13699574206438 | 4.77291284147196 | 5.33554323488568 |
| chr16 | 8853706 | 8853717 | - | 1.00396057687336 | 2.27399148412876 | 2.38645642073598 | 1.06710864697714 |
| chr16 | 8853802 | 8853814 | - | 15.0594086531005 | 18.1919318730301 | 3.57968463110397 | 4.26843458790854 |
| chr16 | 8854183 | 8854240 | - | 28.1108961524542 | 30.6988850357383 | 1.19322821036799 | 9.60397782279422 |
| chr16 | 8854580 | 8854635 | - | 18.0712903837206 | 26.1509020674808 | 8.35259747257593 | 9.60397782279422 |
| chr16 | 8854915 | 8855076 | - | 42.1663442286813 | 59.1237785873479 | 19.0916513658878 | 23.476390233497 |
| chr16 | 8855156 | 8855254 | - | 2.00792115374673 | 2.27399148412876 | 3.57968463110397 | 1.06710864697714 |
| chr16 | 8892801 | 8892859 | - | 11.043566345607 | 12.5069531627082 | 17.8984231555198 | 33.0803680562912 |
| chr16 | 8893046 | 8893092 | - | 7.02772403811355 | 2.27399148412876 | 3.57968463110397 | 7.46976052883995 |
| chr16 | 8893340 | 8893379 | - | 58.2297134586551 | 15.9179403889013 | 62.0478669391355 | 32.0132594093141 |
| chr16 | 8893438 | 8893444 | - | 8.03168461498692 | 9.09596593651506 | 4.77291284147196 | 10.6710864697714 |
| chr16 | 8893530 | 8893576 | - | 3.01188173062009 | 1.13699574206438 | 1.19322821036799 | 2.13421729395427 |
| chr16 | 8893598 | 8893756 | - | 14.0554480762271 | 4.54798296825753 | 16.7051949451519 | 17.0737383516342 |
| chr16 | 8893846 | 8893888 | - | 2.00792115374673 | 1.13699574206438 | 2.38645642073598 | 2.13421729395427 |
| chr16 | 8893912 | 8894047 | - | 109.431702879197 | 77.315710460378 | 114.549908195327 | 245.434988804741 |
| chr16 | 9104918 | 9104936 | + | 5.01980288436682 | 3.41098722619315 | 2.38645642073598 | 1.06710864697714 |
| chr16 | 9105191 | 9105196 | + | 5.01980288436682 | 7.95897019445067 | 1.19322821036799 | 1.06710864697714 |
| chr16 | 9105372 | 9105389 | + | 2.00792115374673 | 1.13699574206438 | 1.19322821036799 | 3.20132594093141 |
| chr16 | 9105523 | 9105582 | + | 19.0752509605939 | 22.7399148412876 | 15.5119667347839 | 12.8053037637256 |
| chr16 | 9105741 | 9105779 | + | 13.0514874993537 | 7.95897019445067 | 8.35259747257593 | 4.26843458790854 |
| chr16 | 9105829 | 9105845 | + | 2.00792115374673 | 1.13699574206438 | 1.19322821036799 | 4.26843458790854 |
| chr16 | 9105870 | 9105890 | + | 25.0990144218341 | 17.0549361309657 | 26.2510206280958 | 22.4092815865198 |
| chr16 | 9107444 | 9107450 | + | 2.00792115374673 | 1.13699574206438 | 3.57968463110397 | 1.06710864697714 |
| chr16 | 9107849 | 9107853 | + | 1.00396057687336 | 2.27399148412876 | 2.38645642073598 | 3.20132594093141 |
| chr16 | 9109717 | 9109747 | + | 5.01980288436682 | 4.54798296825753 | 2.38645642073598 | 8.53686917581709 |
| chr16 | 9110057 | 9110153 | + | 11.043566345607 | 9.09596593651506 | 7.15936926220794 | 17.0737383516342 |
| chr16 | 9111058 | 9111114 | + | 7.02772403811355 | 20.4659233571589 | 17.8984231555198 | 17.0737383516342 |
| chr16 | 9111123 | 9111127 | + | 2.00792115374673 | 5.68497871032191 | 4.77291284147196 | 5.33554323488568 |
| chr16 | 9111212 | 9111273 | + | 4.01584230749346 | 10.2329616785794 | 23.8645642073598 | 12.8053037637256 |
| chr16 | 9111298 | 9111299 | + | 2.00792115374673 | 1.13699574206438 | 2.38645642073598 | 1.06710864697714 |
| chr16 | 9111659 | 9111664 | + | 1.00396057687336 | 1.13699574206438 | 3.57968463110397 | 2.13421729395427 |
| chr16 | 9111941 | 9111942 | + | 1.00396057687336 | 1.13699574206438 | 1.19322821036799 | 1.06710864697714 |
| chr16 | 9111975 | 9111999 | + | 1.00396057687336 | 7.95897019445067 | 7.15936926220794 | 2.13421729395427 |
| chr16 | 9112026 | 9112040 | + | 3.01188173062009 | 3.41098722619315 | 2.38645642073598 | 1.06710864697714 |
| chr16 | 9112107 | 9112118 | + | 14.0554480762271 | 11.3699574206438 | 8.35259747257593 | 4.26843458790854 |
| chr16 | 9112327 | 9112329 | + | 1.00396057687336 | 1.13699574206438 | 3.57968463110397 | 3.20132594093141 |
| chr16 | 9112547 | 9112580 | + | 5.01980288436682 | 9.09596593651506 | 9.54582568294392 | 6.40265188186281 |
| chr16 | 9112600 | 9112621 | + | 2.00792115374673 | 3.41098722619315 | 4.77291284147196 | 2.13421729395427 |
| chr16 | 9112765 | 9112784 | + | 5.01980288436682 | 10.2329616785794 | 9.54582568294392 | 4.26843458790854 |
| chr16 | 9113086 | 9113114 | + | 3.01188173062009 | 5.68497871032191 | 1.19322821036799 | 2.13421729395427 |
| chr16 | 9117025 | 9117066 | + | 2.00792115374673 | 1.13699574206438 | 9.54582568294392 | 11.7381951167485 |
| chr16 | 9117183 | 9117193 | + | 4.01584230749346 | 1.13699574206438 | 4.77291284147196 | 5.33554323488568 |
| chr16 | 9117216 | 9117252 | + | 22.087132691214 | 31.8358807778027 | 40.5697591525116 | 73.6304966414224 |
| chr16 | 9117795 | 9117807 | + | 3.01188173062009 | 2.27399148412876 | 3.57968463110397 | 3.20132594093141 |
| chr16 | 9117869 | 9117874 | + | 1.00396057687336 | 2.27399148412876 | 1.19322821036799 | 3.20132594093141 |
| chr16 | 9118826 | 9118849 | + | 1.00396057687336 | 4.54798296825753 | 2.38645642073598 | 2.13421729395427 |
| chr16 | 9119405 | 9119414 | + | 5.01980288436682 | 1.13699574206438 | 1.19322821036799 | 1.06710864697714 |
| chr16 | 10529484 | 10529555 | - | 147.582204800385 | 87.5486721389574 | 2.38645642073598 | 2.13421729395427 |
| chr16 | 10529973 | 10530023 | - | 51.2019894205416 | 37.5208594881246 | 3.57968463110397 | 1.06710864697714 |
| chr16 | 10530026 | 10530054 | - | 7.02772403811355 | 4.54798296825753 | 1.19322821036799 | 8.53686917581709 |
| chr16 | 10532797 | 10533087 | - | 402.588191326219 | 286.522927000224 | 11.9322821036799 | 26.6777161744284 |
| chr16 | 10769188 | 10769265 | + | 16.0633692299738 | 9.09596593651506 | 5.96614105183995 | 9.60397782279422 |
| chr16 | 10928929 | 10928952 | - | 5.01980288436682 | 6.82197445238629 | 9.54582568294392 | 5.33554323488568 |
| chr16 | 10929049 | 10929050 | - | 3.01188173062009 | 1.13699574206438 | 5.96614105183995 | 4.26843458790854 |
| chr16 | 10929089 | 10929110 | - | 9.03564519186028 | 1.13699574206438 | 7.15936926220794 | 9.60397782279422 |
| chr16 | 11180491 | 11180560 | + | 11.043566345607 | 9.09596593651506 | 14.3187385244159 | 7.46976052883995 |
| chr16 | 11180594 | 11180596 | + | 3.01188173062009 | 2.27399148412876 | 1.19322821036799 | 1.06710864697714 |
| chr16 | 11254510 | 11254561 | - | 3.01188173062009 | 1.13699574206438 | 50.1155848354556 | 96.0397782279422 |
| chr16 | 11350640 | 11350663 | + | 2.00792115374673 | 3.41098722619315 | 3.57968463110397 | 5.33554323488568 |
| chr16 | 11350700 | 11350716 | + | 1.00396057687336 | 5.68497871032191 | 10.7390538933119 | 12.8053037637256 |
| chr16 | 11350750 | 11350806 | + | 3.01188173062009 | 6.82197445238629 | 15.5119667347839 | 4.26843458790854 |
| chr16 | 11350839 | 11350864 | + | 3.01188173062009 | 1.13699574206438 | 3.57968463110397 | 7.46976052883995 |
| chr16 | 11350904 | 11350958 | + | 7.02772403811355 | 7.95897019445067 | 19.0916513658878 | 10.6710864697714 |
| chr16 | 11351044 | 11351062 | + | 2.00792115374673 | 5.68497871032191 | 7.15936926220794 | 4.26843458790854 |
| chr16 | 11351294 | 11351297 | + | 2.00792115374673 | 4.54798296825753 | 1.19322821036799 | 1.06710864697714 |
| chr16 | 11351328 | 11351330 | + | 1.00396057687336 | 4.54798296825753 | 1.19322821036799 | 3.20132594093141 |
| chr16 | 11351410 | 11351469 | + | 15.0594086531005 | 14.780944646837 | 35.7968463110397 | 38.4159112911769 |
| chr16 | 11351476 | 11351514 | + | 2.00792115374673 | 6.82197445238629 | 15.5119667347839 | 5.33554323488568 |
| chr16 | 11351538 | 11351605 | + | 32.1267384599477 | 117.110561432631 | 70.4004644117114 | 71.4962793474681 |
| chr16 | 11351680 | 11351723 | + | 1.00396057687336 | 11.3699574206438 | 3.57968463110397 | 1.06710864697714 |
| chr16 | 11548308 | 11548642 | - | 360.421847097538 | 281.974944031967 | 28.6374770488317 | 16.006629704657 |
| chr16 | 11548671 | 11548734 | - | 104.41189999483 | 64.8087572976698 | 13.1255103140479 | 6.40265188186281 |
| chr16 | 11548840 | 11549008 | - | 51.2019894205416 | 46.6168254246397 | 5.96614105183995 | 13.8724124107028 |
| chr16 | 11549039 | 11549049 | - | 14.0554480762271 | 19.3289276150945 | 1.19322821036799 | 3.20132594093141 |
| chr16 | 11549079 | 11549292 | - | 23.0910932680874 | 9.09596593651506 | 4.77291284147196 | 2.13421729395427 |
| chr16 | 11549600 | 11549623 | - | 1.00396057687336 | 1.13699574206438 | 1.19322821036799 | 1.06710864697714 |
| chr16 | 11679136 | 11679240 | - | 8.03168461498692 | 6.82197445238629 | 7.15936926220794 | 2.13421729395427 |
| chr16 | 11743053 | 11743055 | + | 2.00792115374673 | 1.13699574206438 | 1.19322821036799 | 1.06710864697714 |
| chr16 | 11750619 | 11750638 | - | 1.00396057687336 | 3.41098722619315 | 3.57968463110397 | 1.06710864697714 |
| chr16 | 11750794 | 11750806 | - | 1.00396057687336 | 3.41098722619315 | 1.19322821036799 | 2.13421729395427 |
| chr16 | 11750946 | 11750966 | - | 5.01980288436682 | 4.54798296825753 | 1.19322821036799 | 4.26843458790854 |
| chr16 | 11750989 | 11750990 | - | 2.00792115374673 | 3.41098722619315 | 1.19322821036799 | 3.20132594093141 |
| chr16 | 11797522 | 11797552 | + | 7.02772403811355 | 1.13699574206438 | 1.19322821036799 | 4.26843458790854 |
| chr16 | 11837368 | 11837428 | - | 8.03168461498692 | 10.2329616785794 | 2.38645642073598 | 11.7381951167485 |
| chr16 | 11868604 | 11868633 | - | 6.02376346124019 | 6.82197445238629 | 11.9322821036799 | 8.53686917581709 |
| chr16 | 11868701 | 11868718 | - | 14.0554480762271 | 9.09596593651506 | 5.96614105183995 | 12.8053037637256 |
| chr16 | 11868910 | 11868964 | - | 1.00396057687336 | 1.13699574206438 | 2.38645642073598 | 3.20132594093141 |
| chr16 | 11870137 | 11870148 | - | 1.00396057687336 | 1.13699574206438 | 1.19322821036799 | 1.06710864697714 |
| chr16 | 11872409 | 11872536 | - | 17.0673298068472 | 25.0139063254164 | 11.9322821036799 | 12.8053037637256 |
| chr16 | 11872701 | 11872792 | - | 44.174265382428 | 63.6717615556054 | 47.7291284147196 | 54.4225409958339 |
| chr16 | 11872817 | 11872870 | - | 16.0633692299738 | 9.09596593651506 | 9.54582568294392 | 14.9395210576799 |
| chr16 | 11872935 | 11873008 | - | 24.0950538449607 | 19.3289276150945 | 9.54582568294392 | 29.8790421153598 |
| chr16 | 11873034 | 11873164 | - | 23.0910932680874 | 22.7399148412876 | 13.1255103140479 | 29.8790421153598 |
| chr16 | 12662474 | 12662570 | - | 2.00792115374673 | 1.13699574206438 | 4.77291284147196 | 3.20132594093141 |
| chr16 | 12663876 | 12664031 | - | 1.00396057687336 | 1.13699574206438 | 7.15936926220794 | 11.7381951167485 |
| chr16 | 12664065 | 12664181 | - | 1.00396057687336 | 2.27399148412876 | 3.57968463110397 | 2.13421729395427 |
| chr16 | 13948687 | 13948713 | + | 15.0594086531005 | 12.5069531627082 | 5.96614105183995 | 7.46976052883995 |
| chr16 | 14263343 | 14263359 | + | 1.00396057687336 | 2.27399148412876 | 1.19322821036799 | 1.06710864697714 |
| chr16 | 14265143 | 14265166 | + | 7.02772403811355 | 4.54798296825753 | 1.19322821036799 | 3.20132594093141 |
| chr16 | 14302228 | 14302331 | + | 43.1703048055547 | 43.2058381984465 | 13.1255103140479 | 6.40265188186281 |
| chr16 | 14895966 | 14895987 | + | 11.043566345607 | 21.6029190992233 | 4.77291284147196 | 2.13421729395427 |
| chr16 | 18784737 | 18784738 | - | 4.01584230749346 | 2.27399148412876 | 3.57968463110397 | 6.40265188186281 |
| chr16 | 18791693 | 18791703 | - | 366.445610558778 | 552.57993064329 | 190.916513658878 | 171.804492163319 |
| chr16 | 18791758 | 18791776 | - | 1.00396057687336 | 2.27399148412876 | 1.19322821036799 | 2.13421729395427 |
| chr16 | 18791799 | 18791834 | - | 2.00792115374673 | 3.41098722619315 | 4.77291284147196 | 2.13421729395427 |
| chr16 | 18791871 | 18791987 | - | 26.1029749987075 | 18.1919318730301 | 15.5119667347839 | 20.2750642925656 |
| chr16 | 18792031 | 18792078 | - | 8.03168461498692 | 4.54798296825753 | 8.35259747257593 | 3.20132594093141 |
| chr16 | 18792445 | 18792525 | - | 7.02772403811355 | 1.13699574206438 | 3.57968463110397 | 4.26843458790854 |
| chr16 | 18792548 | 18792645 | - | 5.01980288436682 | 5.68497871032191 | 9.54582568294392 | 7.46976052883995 |
| chr16 | 18792676 | 18792780 | - | 4.01584230749346 | 2.27399148412876 | 9.54582568294392 | 4.26843458790854 |
| chr16 | 18792855 | 18792940 | - | 14.0554480762271 | 2.27399148412876 | 3.57968463110397 | 7.46976052883995 |
| chr16 | 18792990 | 18793109 | - | 21.0831721143407 | 12.5069531627082 | 13.1255103140479 | 12.8053037637256 |
| chr16 | 18804893 | 18804943 | - | 7.02772403811355 | 4.54798296825753 | 7.15936926220794 | 6.40265188186281 |
| chr16 | 18805406 | 18805474 | - | 2.00792115374673 | 4.54798296825753 | 1.19322821036799 | 2.13421729395427 |
| chr16 | 19079777 | 19079799 | + | 13.0514874993537 | 5.68497871032191 | 1.19322821036799 | 2.13421729395427 |
| chr16 | 19119130 | 19119152 | + | 14.0554480762271 | 12.5069531627082 | 3.57968463110397 | 2.13421729395427 |
| chr16 | 19501762 | 19501797 | - | 21.0831721143407 | 13.6439489047726 | 4.77291284147196 | 8.53686917581709 |
| chr16 | 19502601 | 19502609 | - | 2.00792115374673 | 6.82197445238629 | 1.19322821036799 | 12.8053037637256 |
| chr16 | 19502681 | 19502696 | - | 14.0554480762271 | 11.3699574206438 | 1.19322821036799 | 4.26843458790854 |
| chr16 | 19551276 | 19551282 | + | 1.00396057687336 | 5.68497871032191 | 4.77291284147196 | 2.13421729395427 |
| chr16 | 19700481 | 19700504 | + | 10.0396057687336 | 5.68497871032191 | 7.15936926220794 | 10.6710864697714 |
| chr16 | 19703279 | 19703289 | - | 3.01188173062009 | 3.41098722619315 | 7.15936926220794 | 2.13421729395427 |
| chr16 | 19703394 | 19703412 | - | 6.02376346124019 | 6.82197445238629 | 2.38645642073598 | 2.13421729395427 |
| chr16 | 19703890 | 19703896 | - | 23.0910932680874 | 11.3699574206438 | 5.96614105183995 | 6.40265188186281 |
| chr16 | 19704057 | 19704073 | - | 3.01188173062009 | 3.41098722619315 | 1.19322821036799 | 5.33554323488568 |
| chr16 | 19857265 | 19857268 | + | 4.01584230749346 | 3.41098722619315 | 2.38645642073598 | 1.06710864697714 |
| chr16 | 19857543 | 19857545 | + | 2.00792115374673 | 2.27399148412876 | 1.19322821036799 | 2.13421729395427 |
| chr16 | 20733727 | 20733739 | - | 2.00792115374673 | 1.13699574206438 | 5.96614105183995 | 5.33554323488568 |
| chr16 | 20733832 | 20733835 | - | 3.01188173062009 | 9.09596593651506 | 11.9322821036799 | 2.13421729395427 |
| chr16 | 20733929 | 20733950 | - | 10.0396057687336 | 18.1919318730301 | 22.6713359969918 | 37.3488026441998 |
| chr16 | 20734155 | 20734250 | - | 4.01584230749346 | 3.41098722619315 | 3.57968463110397 | 9.60397782279422 |
| chr16 | 20734315 | 20734380 | - | 1.00396057687336 | 2.27399148412876 | 14.3187385244159 | 4.26843458790854 |
| chr16 | 20735866 | 20735887 | - | 8.03168461498692 | 17.0549361309657 | 10.7390538933119 | 17.0737383516342 |
| chr16 | 20735914 | 20735938 | - | 2.00792115374673 | 6.82197445238629 | 8.35259747257593 | 6.40265188186281 |
| chr16 | 20736398 | 20736478 | - | 2.00792115374673 | 2.27399148412876 | 36.9900745214077 | 57.6238669367653 |
| chr16 | 20736509 | 20736558 | - | 4.01584230749346 | 1.13699574206438 | 7.15936926220794 | 24.5434988804741 |
| chr16 | 20736632 | 20736655 | - | 11.043566345607 | 18.1919318730301 | 10.7390538933119 | 11.7381951167485 |
| chr16 | 20736820 | 20736835 | - | 4.01584230749346 | 1.13699574206438 | 1.19322821036799 | 5.33554323488568 |
| chr16 | 20736857 | 20736882 | - | 6.02376346124019 | 2.27399148412876 | 17.8984231555198 | 6.40265188186281 |
| chr16 | 20796791 | 20796800 | - | 2.00792115374673 | 1.13699574206438 | 1.19322821036799 | 4.26843458790854 |
| chr16 | 20855032 | 20855052 | - | 2.00792115374673 | 4.54798296825753 | 3.57968463110397 | 2.13421729395427 |
| chr16 | 20855173 | 20855202 | - | 14.0554480762271 | 11.3699574206438 | 21.4781077866238 | 11.7381951167485 |
| chr16 | 20855230 | 20855271 | - | 12.0475269224804 | 21.6029190992233 | 10.7390538933119 | 5.33554323488568 |
| chr16 | 20855936 | 20855937 | - | 1.00396057687336 | 1.13699574206438 | 1.19322821036799 | 1.06710864697714 |
| chr16 | 20859505 | 20859516 | - | 14.0554480762271 | 14.780944646837 | 2.38645642073598 | 2.13421729395427 |
| chr16 | 20924876 | 20924930 | + | 6.02376346124019 | 4.54798296825753 | 5.96614105183995 | 7.46976052883995 |
| chr16 | 23388517 | 23388537 | - | 1.00396057687336 | 3.41098722619315 | 2.38645642073598 | 1.06710864697714 |
| chr16 | 23388754 | 23388756 | - | 2.00792115374673 | 1.13699574206438 | 11.9322821036799 | 1.06710864697714 |
| chr16 | 23452724 | 23452816 | + | 20.0792115374673 | 21.6029190992233 | 1.19322821036799 | 4.26843458790854 |
| chr16 | 23465662 | 23465677 | - | 3.01188173062009 | 2.27399148412876 | 4.77291284147196 | 4.26843458790854 |
| chr16 | 23466023 | 23466037 | - | 3.01188173062009 | 5.68497871032191 | 2.38645642073598 | 1.06710864697714 |
| chr16 | 23466189 | 23466231 | - | 10.0396057687336 | 7.95897019445067 | 7.15936926220794 | 14.9395210576799 |
| chr16 | 23466258 | 23466300 | - | 2.00792115374673 | 4.54798296825753 | 2.38645642073598 | 2.13421729395427 |
| chr16 | 23466399 | 23466430 | - | 3.01188173062009 | 1.13699574206438 | 5.96614105183995 | 6.40265188186281 |
| chr16 | 23466884 | 23466893 | - | 2.00792115374673 | 1.13699574206438 | 1.19322821036799 | 1.06710864697714 |
| chr16 | 23467222 | 23467281 | - | 9.03564519186028 | 3.41098722619315 | 14.3187385244159 | 11.7381951167485 |
| chr16 | 23467347 | 23467402 | - | 8.03168461498692 | 9.09596593651506 | 15.5119667347839 | 18.1408469986113 |
| chr16 | 23467505 | 23467552 | - | 4.01584230749346 | 4.54798296825753 | 5.96614105183995 | 12.8053037637256 |
| chr16 | 23522221 | 23522243 | - | 1.00396057687336 | 1.13699574206438 | 2.38645642073598 | 5.33554323488568 |
| chr16 | 23523053 | 23523056 | - | 1.00396057687336 | 2.27399148412876 | 1.19322821036799 | 1.06710864697714 |
| chr16 | 23524047 | 23524070 | - | 3.01188173062009 | 7.95897019445067 | 9.54582568294392 | 5.33554323488568 |
| chr16 | 23524085 | 23524109 | - | 2.00792115374673 | 2.27399148412876 | 9.54582568294392 | 4.26843458790854 |
| chr16 | 23570702 | 23570743 | + | 7.02772403811355 | 2.27399148412876 | 9.54582568294392 | 5.33554323488568 |
| chr16 | 23570918 | 23570922 | + | 3.01188173062009 | 4.54798296825753 | 2.38645642073598 | 6.40265188186281 |
| chr16 | 23570990 | 23571035 | + | 4.01584230749346 | 7.95897019445067 | 5.96614105183995 | 8.53686917581709 |
| chr16 | 23571062 | 23571081 | + | 19.0752509605939 | 11.3699574206438 | 8.35259747257593 | 5.33554323488568 |
| chr16 | 23571837 | 23571898 | + | 11.043566345607 | 4.54798296825753 | 9.54582568294392 | 10.6710864697714 |
| chr16 | 23572176 | 23572211 | + | 4.01584230749346 | 2.27399148412876 | 1.19322821036799 | 4.26843458790854 |
| chr16 | 23572678 | 23572730 | + | 8.03168461498692 | 3.41098722619315 | 3.57968463110397 | 5.33554323488568 |
| chr16 | 23572944 | 23572962 | + | 1.00396057687336 | 1.13699574206438 | 1.19322821036799 | 3.20132594093141 |
| chr16 | 23573106 | 23573109 | + | 4.01584230749346 | 2.27399148412876 | 2.38645642073598 | 7.46976052883995 |
| chr16 | 23573205 | 23573238 | + | 2.00792115374673 | 1.13699574206438 | 1.19322821036799 | 2.13421729395427 |
| chr16 | 23573494 | 23573521 | + | 2.00792115374673 | 1.13699574206438 | 5.96614105183995 | 5.33554323488568 |
| chr16 | 23573766 | 23573788 | + | 1.00396057687336 | 2.27399148412876 | 3.57968463110397 | 1.06710864697714 |
| chr16 | 23573821 | 23573874 | + | 5.01980288436682 | 2.27399148412876 | 7.15936926220794 | 2.13421729395427 |
| chr16 | 23573937 | 23573976 | + | 6.02376346124019 | 4.54798296825753 | 17.8984231555198 | 17.0737383516342 |
| chr16 | 23573998 | 23574005 | + | 6.02376346124019 | 1.13699574206438 | 4.77291284147196 | 1.06710864697714 |
| chr16 | 23574037 | 23574069 | + | 68.2693192273888 | 15.9179403889013 | 23.8645642073598 | 34.1474767032683 |
| chr16 | 23574091 | 23574143 | + | 157.621810569118 | 104.603608269923 | 110.970223564223 | 178.207144045182 |
| chr16 | 23574170 | 23574285 | + | 581.293174009678 | 383.167565075697 | 573.942769187003 | 579.439995308585 |
| chr16 | 23581035 | 23581173 | - | 107.42378172545 | 153.494425178692 | 151.539982716735 | 208.086186160541 |
| chr16 | 23667694 | 23667701 | + | 10.0396057687336 | 4.54798296825753 | 8.35259747257593 | 6.40265188186281 |
| chr16 | 23667747 | 23667757 | + | 3.01188173062009 | 2.27399148412876 | 1.19322821036799 | 1.06710864697714 |
| chr16 | 23667808 | 23667845 | + | 3.01188173062009 | 1.13699574206438 | 4.77291284147196 | 2.13421729395427 |
| chr16 | 23668046 | 23668074 | + | 4.01584230749346 | 1.13699574206438 | 2.38645642073598 | 2.13421729395427 |
| chr16 | 23668181 | 23668206 | + | 8.03168461498692 | 5.68497871032191 | 8.35259747257593 | 4.26843458790854 |
| chr16 | 23668311 | 23668332 | + | 1.00396057687336 | 2.27399148412876 | 1.19322821036799 | 5.33554323488568 |
| chr16 | 23668424 | 23668461 | + | 1.00396057687336 | 2.27399148412876 | 2.38645642073598 | 5.33554323488568 |
| chr16 | 23668567 | 23668576 | + | 2.00792115374673 | 1.13699574206438 | 4.77291284147196 | 9.60397782279422 |
| chr16 | 23668642 | 23668649 | + | 7.02772403811355 | 1.13699574206438 | 7.15936926220794 | 5.33554323488568 |
| chr16 | 23668789 | 23668793 | + | 4.01584230749346 | 1.13699574206438 | 4.77291284147196 | 2.13421729395427 |
| chr16 | 23669013 | 23669044 | + | 2.00792115374673 | 1.13699574206438 | 5.96614105183995 | 8.53686917581709 |
| chr16 | 23669049 | 23669095 | + | 7.02772403811355 | 6.82197445238629 | 3.57968463110397 | 5.33554323488568 |
| chr16 | 23669154 | 23669172 | + | 1.00396057687336 | 5.68497871032191 | 1.19322821036799 | 3.20132594093141 |
| chr16 | 23669200 | 23669219 | + | 11.043566345607 | 5.68497871032191 | 10.7390538933119 | 10.6710864697714 |
| chr16 | 23669242 | 23669261 | + | 4.01584230749346 | 5.68497871032191 | 13.1255103140479 | 6.40265188186281 |
| chr16 | 23669313 | 23669336 | + | 4.01584230749346 | 2.27399148412876 | 3.57968463110397 | 6.40265188186281 |
| chr16 | 23669543 | 23669659 | + | 31.1227778830743 | 14.780944646837 | 34.6036181006717 | 29.8790421153598 |
| chr16 | 23669802 | 23669812 | + | 11.043566345607 | 13.6439489047726 | 7.15936926220794 | 8.53686917581709 |
| chr16 | 23673910 | 23673920 | + | 2.00792115374673 | 3.41098722619315 | 4.77291284147196 | 7.46976052883995 |
| chr16 | 23690174 | 23690180 | + | 1.00396057687336 | 4.54798296825753 | 2.38645642073598 | 3.20132594093141 |
| chr16 | 23690240 | 23690267 | + | 44.174265382428 | 18.1919318730301 | 15.5119667347839 | 19.2079556455884 |
| chr16 | 23690317 | 23690347 | + | 13.0514874993537 | 11.3699574206438 | 1.19322821036799 | 9.60397782279422 |
| chr16 | 23690327 | 23690354 | - | 38.1505019211879 | 31.8358807778027 | 27.4442488384638 | 70.429170700491 |
| chr16 | 24215253 | 24215257 | + | 3.01188173062009 | 6.82197445238629 | 7.15936926220794 | 23.476390233497 |
| chr16 | 24215420 | 24215454 | + | 11.043566345607 | 15.9179403889013 | 34.6036181006717 | 74.6976052883995 |
| chr16 | 24215569 | 24215572 | + | 1.00396057687336 | 6.82197445238629 | 31.0239334695677 | 34.1474767032683 |
| chr16 | 24215667 | 24215668 | + | 1.00396057687336 | 1.13699574206438 | 3.57968463110397 | 1.06710864697714 |
| chr16 | 24215753 | 24215845 | + | 4.01584230749346 | 12.5069531627082 | 76.3666054635513 | 84.3015831111937 |
| chr16 | 24215908 | 24215977 | + | 2.00792115374673 | 15.9179403889013 | 33.4103898903037 | 43.7514545260626 |
| chr16 | 24216196 | 24216218 | + | 1.00396057687336 | 1.13699574206438 | 17.8984231555198 | 7.46976052883995 |
| chr16 | 24216915 | 24216964 | + | 1.00396057687336 | 1.13699574206438 | 28.6374770488317 | 67.2278447595595 |
| chr16 | 24219209 | 24219231 | + | 2.00792115374673 | 3.41098722619315 | 14.3187385244159 | 38.4159112911769 |
| chr16 | 24572699 | 24572703 | + | 19.0752509605939 | 12.5069531627082 | 16.7051949451519 | 18.1408469986113 |
| chr16 | 24824239 | 24824243 | + | 1.00396057687336 | 2.27399148412876 | 1.19322821036799 | 2.13421729395427 |
| chr16 | 24824286 | 24824294 | + | 6.02376346124019 | 7.95897019445067 | 3.57968463110397 | 4.26843458790854 |
| chr16 | 24824488 | 24824553 | + | 13.0514874993537 | 6.82197445238629 | 5.96614105183995 | 5.33554323488568 |
| chr16 | 24919506 | 24919574 | - | 1.00396057687336 | 5.68497871032191 | 3.57968463110397 | 7.46976052883995 |
| chr16 | 24919620 | 24919632 | - | 7.02772403811355 | 9.09596593651506 | 1.19322821036799 | 5.33554323488568 |
| chr16 | 24919656 | 24919682 | - | 3.01188173062009 | 7.95897019445067 | 7.15936926220794 | 8.53686917581709 |
| chr16 | 24919894 | 24919932 | - | 4.01584230749346 | 2.27399148412876 | 1.19322821036799 | 10.6710864697714 |
| chr16 | 25178143 | 25178170 | + | 5.01980288436682 | 3.41098722619315 | 9.54582568294392 | 7.46976052883995 |
| chr16 | 25239320 | 25239328 | - | 1.00396057687336 | 1.13699574206438 | 3.57968463110397 | 3.20132594093141 |
| chr16 | 27225021 | 27225086 | - | 5.01980288436682 | 14.780944646837 | 4.77291284147196 | 8.53686917581709 |
| chr16 | 27225119 | 27225139 | - | 4.01584230749346 | 3.41098722619315 | 1.19322821036799 | 2.13421729395427 |
| chr16 | 27268839 | 27268892 | + | 6.02376346124019 | 10.2329616785794 | 5.96614105183995 | 13.8724124107028 |
| chr16 | 27364638 | 27364665 | + | 11.043566345607 | 21.6029190992233 | 1.19322821036799 | 1.06710864697714 |
| chr16 | 27460719 | 27460809 | - | 9.03564519186028 | 2.27399148412876 | 7.15936926220794 | 9.60397782279422 |
| chr16 | 28097995 | 28098046 | - | 1.00396057687336 | 13.6439489047726 | 2.38645642073598 | 3.20132594093141 |
| chr16 | 28098144 | 28098169 | - | 10.0396057687336 | 11.3699574206438 | 9.54582568294392 | 7.46976052883995 |
| chr16 | 28098208 | 28098214 | - | 1.00396057687336 | 1.13699574206438 | 1.19322821036799 | 1.06710864697714 |
| chr16 | 28477301 | 28477358 | - | 4.01584230749346 | 6.82197445238629 | 2.38645642073598 | 4.26843458790854 |
| chr16 | 28591757 | 28591770 | + | 2.00792115374673 | 10.2329616785794 | 7.15936926220794 | 1.06710864697714 |
| chr16 | 28605635 | 28605655 | - | 2.00792115374673 | 5.68497871032191 | 1.19322821036799 | 3.20132594093141 |
| chr16 | 28836311 | 28836338 | + | 1.00396057687336 | 1.13699574206438 | 5.96614105183995 | 5.33554323488568 |
| chr16 | 28836682 | 28836695 | + | 1.00396057687336 | 3.41098722619315 | 4.77291284147196 | 3.20132594093141 |
| chr16 | 28836957 | 28836999 | + | 29.1148567293276 | 21.6029190992233 | 8.35259747257593 | 18.1408469986113 |
| chr16 | 28837035 | 28837045 | + | 16.0633692299738 | 13.6439489047726 | 20.2848795762558 | 14.9395210576799 |
| chr16 | 28837126 | 28837168 | + | 16.0633692299738 | 37.5208594881246 | 57.2749540976635 | 41.6172372321083 |
| chr16 | 28842806 | 28842892 | - | 755.982314385643 | 803.855989639518 | 622.86512581209 | 728.835205885384 |
| chr16 | 28842960 | 28843015 | - | 30.1188173062009 | 20.4659233571589 | 16.7051949451519 | 21.3421729395427 |
| chr16 | 28874112 | 28874143 | + | 32.1267384599477 | 39.7948509722534 | 26.2510206280958 | 21.3421729395427 |
| chr16 | 28964428 | 28964477 | + | 10.0396057687336 | 7.95897019445067 | 3.57968463110397 | 7.46976052883995 |
| chr16 | 28966220 | 28966242 | + | 6.02376346124019 | 2.27399148412876 | 2.38645642073598 | 2.13421729395427 |
| chr16 | 28966419 | 28966422 | + | 1.00396057687336 | 2.27399148412876 | 1.19322821036799 | 3.20132594093141 |
| chr16 | 28966521 | 28966522 | + | 1.00396057687336 | 1.13699574206438 | 2.38645642073598 | 1.06710864697714 |
| chr16 | 28966544 | 28966545 | + | 3.01188173062009 | 3.41098722619315 | 7.15936926220794 | 3.20132594093141 |
| chr16 | 28966983 | 28966995 | + | 1.00396057687336 | 1.13699574206438 | 3.57968463110397 | 3.20132594093141 |
| chr16 | 28975325 | 28975341 | + | 6.02376346124019 | 4.54798296825753 | 2.38645642073598 | 3.20132594093141 |
| chr16 | 28975508 | 28975529 | + | 5.01980288436682 | 6.82197445238629 | 2.38645642073598 | 2.13421729395427 |
| chr16 | 28977961 | 28978022 | + | 39.1544624980612 | 20.4659233571589 | 10.7390538933119 | 21.3421729395427 |
| chr16 | 28979192 | 28979203 | + | 1.00396057687336 | 3.41098722619315 | 5.96614105183995 | 1.06710864697714 |
| chr16 | 28981586 | 28981608 | + | 8.03168461498692 | 7.95897019445067 | 3.57968463110397 | 2.13421729395427 |
| chr16 | 28981998 | 28982010 | + | 6.02376346124019 | 4.54798296825753 | 2.38645642073598 | 2.13421729395427 |
| chr16 | 28982353 | 28982366 | + | 1.00396057687336 | 1.13699574206438 | 2.38645642073598 | 4.26843458790854 |
| chr16 | 28982408 | 28982483 | + | 23.0910932680874 | 10.2329616785794 | 9.54582568294392 | 3.20132594093141 |
| chr16 | 28982522 | 28982538 | + | 5.01980288436682 | 7.95897019445067 | 3.57968463110397 | 4.26843458790854 |
| chr16 | 28983866 | 28983889 | + | 4.01584230749346 | 1.13699574206438 | 4.77291284147196 | 4.26843458790854 |
| chr16 | 28984450 | 28984524 | + | 28.1108961524542 | 40.9318467143178 | 16.7051949451519 | 22.4092815865198 |
| chr16 | 29697302 | 29697537 | + | 1.00396057687336 | 1.13699574206438 | 99.0379414605431 | 85.3686917581709 |
| chr16 | 29698026 | 29698091 | + | 6.02376346124019 | 5.68497871032191 | 3580.87785931434 | 2437.27614969578 |
| chr16 | 29805293 | 29805316 | + | 3.01188173062009 | 1.13699574206438 | 4.77291284147196 | 8.53686917581709 |
| chr16 | 29805352 | 29805360 | + | 4.01584230749346 | 2.27399148412876 | 3.57968463110397 | 4.26843458790854 |
| chr16 | 29808573 | 29808702 | + | 38.1505019211879 | 37.5208594881246 | 29.8307052591997 | 26.6777161744284 |
| chr16 | 29809740 | 29809747 | + | 6.02376346124019 | 13.6439489047726 | 5.96614105183995 | 4.26843458790854 |
| chr16 | 29810075 | 29810401 | + | 1642.47950376482 | 1312.0930863423 | 1471.25038338373 | 600.782168248127 |
| chr16 | 29810424 | 29810587 | + | 165.653495184105 | 143.261463500112 | 288.761226909053 | 370.286700501066 |
| chr16 | 29810619 | 29810651 | + | 193.764391336559 | 156.905412404885 | 188.530057238142 | 290.253551977781 |
| chr16 | 29810677 | 29810902 | + | 60.2376346124019 | 68.2197445238629 | 100.231169670911 | 165.401840281456 |
| chr16 | 29810930 | 29811142 | + | 71.2812009580089 | 71.6307317500561 | 58.4681823080315 | 83.2344744642166 |
| chr16 | 29816357 | 29816633 | + | 686.709034581381 | 487.77117334562 | 503.542304775292 | 745.908944237018 |
| chr16 | 29816654 | 29816769 | + | 57.2257528817818 | 48.8908169087684 | 38.1833027317757 | 53.3554323488568 |
| chr16 | 29816798 | 29817004 | + | 405.600073056839 | 210.344212281911 | 60.8546387287675 | 39.483019938154 |
| chr16 | 29817213 | 29817274 | + | 16.0633692299738 | 7.95897019445067 | 14.3187385244159 | 21.3421729395427 |
| chr16 | 29819583 | 29819613 | + | 9.03564519186028 | 4.54798296825753 | 4.77291284147196 | 10.6710864697714 |
| chr16 | 29819634 | 29819658 | + | 17.0673298068472 | 10.2329616785794 | 13.1255103140479 | 5.33554323488568 |
| chr16 | 29819710 | 29819745 | + | 29.1148567293276 | 21.6029190992233 | 21.4781077866238 | 11.7381951167485 |
| chr16 | 29819842 | 29819843 | + | 2.00792115374673 | 2.27399148412876 | 1.19322821036799 | 1.06710864697714 |
| chr16 | 29819882 | 29819909 | + | 10.0396057687336 | 6.82197445238629 | 8.35259747257593 | 8.53686917581709 |
| chr16 | 29819980 | 29819986 | + | 3.01188173062009 | 1.13699574206438 | 1.19322821036799 | 4.26843458790854 |
| chr16 | 29820018 | 29820030 | + | 5.01980288436682 | 3.41098722619315 | 1.19322821036799 | 1.06710864697714 |
| chr16 | 29820055 | 29820066 | + | 4.01584230749346 | 3.41098722619315 | 2.38645642073598 | 5.33554323488568 |
| chr16 | 29858380 | 29858419 | - | 8.03168461498692 | 17.0549361309657 | 1.19322821036799 | 2.13421729395427 |
| chr16 | 29858449 | 29858471 | - | 10.0396057687336 | 11.3699574206438 | 2.38645642073598 | 8.53686917581709 |
| chr16 | 29859227 | 29859231 | - | 1.00396057687336 | 2.27399148412876 | 1.19322821036799 | 3.20132594093141 |
| chr16 | 29971537 | 29971550 | + | 7.02772403811355 | 11.3699574206438 | 17.8984231555198 | 27.7448248214055 |
| chr16 | 29971624 | 29971644 | + | 10.0396057687336 | 11.3699574206438 | 10.7390538933119 | 7.46976052883995 |
| chr16 | 29972948 | 29973026 | + | 27.1069355755808 | 45.4798296825753 | 31.0239334695677 | 36.2816939972226 |
| chr16 | 29988264 | 29988281 | + | 7.02772403811355 | 7.95897019445067 | 7.15936926220794 | 13.8724124107028 |
| chr16 | 29992175 | 29992228 | + | 2.00792115374673 | 7.95897019445067 | 4.77291284147196 | 7.46976052883995 |
| chr16 | 30005490 | 30005604 | + | 14.0554480762271 | 6.82197445238629 | 25.0577924177278 | 64.0265188186281 |
| chr16 | 30005726 | 30005765 | + | 9.03564519186028 | 15.9179403889013 | 8.35259747257593 | 20.2750642925656 |
| chr16 | 30070114 | 30070395 | + | 43.1703048055547 | 29.5618892936739 | 159.892580189311 | 229.428359100084 |
| chr16 | 30082749 | 30082811 | + | 27.1069355755808 | 12.5069531627082 | 27.4442488384638 | 27.7448248214055 |
| chr16 | 30083392 | 30083523 | + | 21.0831721143407 | 19.3289276150945 | 27.4442488384638 | 22.4092815865198 |
| chr16 | 30083663 | 30083688 | + | 3.01188173062009 | 2.27399148412876 | 11.9322821036799 | 6.40265188186281 |
| chr16 | 30083709 | 30083757 | + | 6.02376346124019 | 4.54798296825753 | 8.35259747257593 | 29.8790421153598 |
| chr16 | 30083766 | 30083800 | + | 10.0396057687336 | 12.5069531627082 | 8.35259747257593 | 4.26843458790854 |
| chr16 | 30084673 | 30084853 | + | 36.1425807674411 | 17.0549361309657 | 81.1395183050233 | 91.7713436400337 |
| chr16 | 30084930 | 30084935 | + | 2.00792115374673 | 1.13699574206438 | 5.96614105183995 | 7.46976052883995 |
| chr16 | 30084996 | 30085045 | + | 8.03168461498692 | 5.68497871032191 | 10.7390538933119 | 23.476390233497 |
| chr16 | 30085070 | 30085153 | + | 15.0594086531005 | 6.82197445238629 | 26.2510206280958 | 44.8185631730397 |
| chr16 | 30085191 | 30085275 | + | 29.1148567293276 | 26.1509020674808 | 42.9562155732476 | 64.0265188186281 |
| chr16 | 30085327 | 30085346 | + | 4.01584230749346 | 18.1919318730301 | 8.35259747257593 | 4.26843458790854 |
| chr16 | 30114140 | 30114181 | - | 3.01188173062009 | 4.54798296825753 | 1.19322821036799 | 4.26843458790854 |
| chr16 | 30114613 | 30114626 | - | 8.03168461498692 | 1.13699574206438 | 10.7390538933119 | 5.33554323488568 |
| chr16 | 30350910 | 30350932 | - | 4.01584230749346 | 3.41098722619315 | 2.38645642073598 | 1.06710864697714 |
| chr16 | 30351502 | 30351506 | - | 4.01584230749346 | 4.54798296825753 | 3.57968463110397 | 3.20132594093141 |
| chr16 | 30351585 | 30351588 | - | 1.00396057687336 | 1.13699574206438 | 4.77291284147196 | 2.13421729395427 |
| chr16 | 30351669 | 30351707 | - | 9.03564519186028 | 4.54798296825753 | 4.77291284147196 | 6.40265188186281 |
| chr16 | 30352503 | 30352507 | - | 3.01188173062009 | 1.13699574206438 | 3.57968463110397 | 1.06710864697714 |
| chr16 | 30352928 | 30352974 | - | 3.01188173062009 | 3.41098722619315 | 2.38645642073598 | 1.06710864697714 |
| chr16 | 30355420 | 30355449 | + | 20.0792115374673 | 84.1376849127643 | 14.3187385244159 | 14.9395210576799 |
| chr16 | 30357132 | 30357235 | - | 372.469374020018 | 341.098722619315 | 473.711599516092 | 646.667840068144 |
| chr16 | 30357269 | 30357285 | - | 5.01980288436682 | 6.82197445238629 | 5.96614105183995 | 8.53686917581709 |
| chr16 | 30357298 | 30357336 | - | 7.02772403811355 | 3.41098722619315 | 10.7390538933119 | 19.2079556455884 |
| chr16 | 30357428 | 30357482 | - | 10.0396057687336 | 11.3699574206438 | 32.2171616799357 | 22.4092815865198 |
| chr16 | 30357503 | 30357523 | - | 6.02376346124019 | 1.13699574206438 | 13.1255103140479 | 8.53686917581709 |
| chr16 | 30357569 | 30357583 | - | 6.02376346124019 | 2.27399148412876 | 20.2848795762558 | 14.9395210576799 |
| chr16 | 30357612 | 30357738 | - | 20.0792115374673 | 13.6439489047726 | 45.3426719939836 | 34.1474767032683 |
| chr16 | 30357930 | 30357932 | - | 7.02772403811355 | 1.13699574206438 | 1.19322821036799 | 3.20132594093141 |
| chr16 | 30399185 | 30399220 | + | 20.0792115374673 | 36.3838637460602 | 19.0916513658878 | 52.2883237018796 |
| chr16 | 30423727 | 30423752 | - | 5.01980288436682 | 7.95897019445067 | 4.77291284147196 | 1.06710864697714 |
| chr16 | 30423775 | 30423843 | - | 14.0554480762271 | 15.9179403889013 | 8.35259747257593 | 23.476390233497 |
| chr16 | 30423967 | 30423972 | - | 2.00792115374673 | 6.82197445238629 | 4.77291284147196 | 11.7381951167485 |
| chr16 | 30424009 | 30424012 | - | 2.00792115374673 | 1.13699574206438 | 1.19322821036799 | 1.06710864697714 |
| chr16 | 30424184 | 30424220 | - | 1.00396057687336 | 1.13699574206438 | 8.35259747257593 | 9.60397782279422 |
| chr16 | 30443649 | 30443772 | - | 15.0594086531005 | 11.3699574206438 | 20.2848795762558 | 37.3488026441998 |
| chr16 | 30444092 | 30444115 | - | 3.01188173062009 | 2.27399148412876 | 1.19322821036799 | 1.06710864697714 |
| chr16 | 30444149 | 30444231 | - | 6.02376346124019 | 4.54798296825753 | 7.15936926220794 | 10.6710864697714 |
| chr16 | 30444253 | 30444327 | - | 3.01188173062009 | 1.13699574206438 | 1.19322821036799 | 5.33554323488568 |
| chr16 | 30524029 | 30524052 | - | 2.00792115374673 | 4.54798296825753 | 2.38645642073598 | 5.33554323488568 |
| chr16 | 30524074 | 30524104 | - | 4.01584230749346 | 2.27399148412876 | 3.57968463110397 | 5.33554323488568 |
| chr16 | 30524193 | 30524252 | - | 123.487150955424 | 130.754510337404 | 89.4921157775992 | 99.2411041688736 |
| chr16 | 30524276 | 30524345 | - | 15.0594086531005 | 12.5069531627082 | 20.2848795762558 | 22.4092815865198 |
| chr16 | 30524353 | 30524503 | - | 33.130699036821 | 40.9318467143178 | 119.322821036799 | 92.8384522870108 |
| chr16 | 30525985 | 30526005 | - | 2.00792115374673 | 3.41098722619315 | 3.57968463110397 | 4.26843458790854 |
| chr16 | 30535025 | 30535119 | + | 290.144606716402 | 453.661301083688 | 508.315217616764 | 390.561764793632 |
| chr16 | 30535440 | 30535462 | + | 2.00792115374673 | 2.27399148412876 | 1.19322821036799 | 4.26843458790854 |
| chr16 | 30537035 | 30537036 | + | 2.00792115374673 | 1.13699574206438 | 2.38645642073598 | 3.20132594093141 |
| chr16 | 30553949 | 30553952 | - | 4.01584230749346 | 3.41098722619315 | 2.38645642073598 | 3.20132594093141 |
| chr16 | 30554876 | 30554898 | - | 14.0554480762271 | 13.6439489047726 | 16.7051949451519 | 5.33554323488568 |
| chr16 | 30585883 | 30585947 | + | 22.087132691214 | 21.6029190992233 | 56.0817258872955 | 48.0198891139711 |
| chr16 | 30602583 | 30602619 | - | 1.00396057687336 | 5.68497871032191 | 5.96614105183995 | 4.26843458790854 |
| chr16 | 30603407 | 30603481 | - | 31.1227778830743 | 34.1098722619315 | 29.8307052591997 | 40.5501285851312 |
| chr16 | 30604258 | 30604280 | - | 5.01980288436682 | 3.41098722619315 | 7.15936926220794 | 3.20132594093141 |
| chr16 | 30656325 | 30656361 | + | 14.0554480762271 | 14.780944646837 | 10.7390538933119 | 19.2079556455884 |
| chr16 | 30669655 | 30669667 | + | 6.02376346124019 | 7.95897019445067 | 3.57968463110397 | 1.06710864697714 |
| chr16 | 30670707 | 30670792 | + | 10.0396057687336 | 18.1919318730301 | 5.96614105183995 | 8.53686917581709 |
| chr16 | 30740044 | 30740062 | + | 45.1782259593014 | 43.2058381984465 | 85.9124311464952 | 51.2212150549025 |
| chr16 | 30740157 | 30740207 | + | 13.0514874993537 | 10.2329616785794 | 13.1255103140479 | 5.33554323488568 |
| chr16 | 30744209 | 30744229 | + | 6.02376346124019 | 9.09596593651506 | 1.19322821036799 | 5.33554323488568 |
| chr16 | 30748372 | 30748451 | + | 535.110987473503 | 494.593147798006 | 257.737293439486 | 417.23948096806 |
| chr16 | 30748816 | 30748820 | + | 6.02376346124019 | 3.41098722619315 | 1.19322821036799 | 3.20132594093141 |
| chr16 | 30757253 | 30757340 | + | 39.1544624980612 | 14.780944646837 | 9.54582568294392 | 25.6106075274513 |
| chr16 | 30775063 | 30775104 | + | 8.03168461498692 | 5.68497871032191 | 4.77291284147196 | 7.46976052883995 |
| chr16 | 30775444 | 30775624 | + | 129.510914416664 | 222.851165444619 | 190.916513658878 | 167.53605757541 |
| chr16 | 30778497 | 30778500 | - | 3.01188173062009 | 2.27399148412876 | 1.19322821036799 | 1.06710864697714 |
| chr16 | 30778531 | 30778625 | - | 2.00792115374673 | 4.54798296825753 | 4.77291284147196 | 2.13421729395427 |
| chr16 | 30779750 | 30779812 | - | 13.0514874993537 | 11.3699574206438 | 20.2848795762558 | 7.46976052883995 |
| chr16 | 30875418 | 30875437 | + | 9.03564519186028 | 7.95897019445067 | 5.96614105183995 | 1.06710864697714 |
| chr16 | 30875455 | 30875492 | + | 30.1188173062009 | 34.1098722619315 | 5.96614105183995 | 4.26843458790854 |
| chr16 | 30887828 | 30887835 | - | 2.00792115374673 | 4.54798296825753 | 3.57968463110397 | 10.6710864697714 |
| chr16 | 30895083 | 30895114 | + | 5.01980288436682 | 12.5069531627082 | 7.15936926220794 | 10.6710864697714 |
| chr16 | 30895172 | 30895173 | + | 1.00396057687336 | 3.41098722619315 | 2.38645642073598 | 1.06710864697714 |
| chr16 | 30947615 | 30947642 | + | 4.01584230749346 | 1.13699574206438 | 13.1255103140479 | 5.33554323488568 |
| chr16 | 30947678 | 30947712 | + | 113.44754518669 | 71.6307317500561 | 66.8207797806074 | 71.4962793474681 |
| chr16 | 30947779 | 30947799 | + | 1.00396057687336 | 1.13699574206438 | 4.77291284147196 | 3.20132594093141 |
| chr16 | 30948225 | 30948303 | + | 9.03564519186028 | 9.09596593651506 | 28.6374770488317 | 25.6106075274513 |
| chr16 | 30948338 | 30948359 | + | 2.00792115374673 | 1.13699574206438 | 3.57968463110397 | 14.9395210576799 |
| chr16 | 30948623 | 30948631 | + | 2.00792115374673 | 1.13699574206438 | 4.77291284147196 | 8.53686917581709 |
| chr16 | 30948714 | 30948755 | + | 2.00792115374673 | 3.41098722619315 | 7.15936926220794 | 13.8724124107028 |
| chr16 | 30954826 | 30954857 | + | 2.00792115374673 | 5.68497871032191 | 5.96614105183995 | 6.40265188186281 |
| chr16 | 30984567 | 30984588 | + | 14.0554480762271 | 7.95897019445067 | 8.35259747257593 | 13.8724124107028 |
| chr16 | 30984612 | 30984645 | + | 1.00396057687336 | 4.54798296825753 | 2.38645642073598 | 1.06710864697714 |
| chr16 | 30989091 | 30989129 | + | 2.00792115374673 | 2.27399148412876 | 4.77291284147196 | 2.13421729395427 |
| chr16 | 31040090 | 31040097 | + | 6.02376346124019 | 9.09596593651506 | 3.57968463110397 | 8.53686917581709 |
| chr16 | 31040118 | 31040124 | + | 7.02772403811355 | 6.82197445238629 | 9.54582568294392 | 22.4092815865198 |
| chr16 | 31083126 | 31083168 | + | 11.043566345607 | 19.3289276150945 | 27.4442488384638 | 12.8053037637256 |
| chr16 | 31083220 | 31083254 | + | 6.02376346124019 | 5.68497871032191 | 19.0916513658878 | 19.2079556455884 |
| chr16 | 31083278 | 31083280 | + | 1.00396057687336 | 3.41098722619315 | 5.96614105183995 | 2.13421729395427 |
| chr16 | 31083331 | 31083418 | + | 266.049552871442 | 342.235718361379 | 352.002322058557 | 377.756461029906 |
| chr16 | 31090872 | 31090989 | - | 109.431702879197 | 112.562578464374 | 45.3426719939836 | 75.7647139353766 |
| chr16 | 31091031 | 31091192 | - | 16.0633692299738 | 9.09596593651506 | 5.96614105183995 | 9.60397782279422 |
| chr16 | 31091223 | 31091306 | - | 11.043566345607 | 13.6439489047726 | 22.6713359969918 | 26.6777161744284 |
| chr16 | 31112518 | 31112547 | + | 36.1425807674411 | 37.5208594881246 | 27.4442488384638 | 39.483019938154 |
| chr16 | 31123781 | 31123796 | + | 4.01584230749346 | 22.7399148412876 | 31.0239334695677 | 21.3421729395427 |
| chr16 | 31123934 | 31123956 | + | 4.01584230749346 | 7.95897019445067 | 9.54582568294392 | 3.20132594093141 |
| chr16 | 31123972 | 31124020 | + | 1.00396057687336 | 7.95897019445067 | 1.19322821036799 | 3.20132594093141 |
| chr16 | 31131193 | 31131239 | + | 10.0396057687336 | 4.54798296825753 | 19.0916513658878 | 14.9395210576799 |
| chr16 | 31131279 | 31131324 | + | 1.00396057687336 | 1.13699574206438 | 2.38645642073598 | 2.13421729395427 |
| chr16 | 31191398 | 31191581 | + | 119.47130864793 | 122.795540142953 | 93.0718004087032 | 126.985928990279 |
| chr16 | 31489512 | 31489522 | - | 7.02772403811355 | 28.4248935516096 | 8.35259747257593 | 7.46976052883995 |
| chr16 | 31489558 | 31489561 | - | 2.00792115374673 | 7.95897019445067 | 10.7390538933119 | 3.20132594093141 |
| chr16 | 31489858 | 31489868 | - | 3.01188173062009 | 2.27399148412876 | 1.19322821036799 | 3.20132594093141 |
| chr16 | 31489948 | 31489973 | - | 3.01188173062009 | 1.13699574206438 | 2.38645642073598 | 2.13421729395427 |
| chr16 | 31490072 | 31490113 | - | 20.0792115374673 | 15.9179403889013 | 19.0916513658878 | 8.53686917581709 |
| chr16 | 31490749 | 31490768 | - | 8.03168461498692 | 2.27399148412876 | 9.54582568294392 | 9.60397782279422 |
| chr16 | 31490806 | 31490955 | - | 17.0673298068472 | 14.780944646837 | 56.0817258872955 | 18.1408469986113 |
| chr16 | 31759465 | 31759478 | + | 10.0396057687336 | 11.3699574206438 | 14.3187385244159 | 13.8724124107028 |
| chr16 | 46580811 | 46580828 | - | 75.2970432655023 | 30.6988850357383 | 32.2171616799357 | 32.0132594093141 |
| chr16 | 46580853 | 46580863 | - | 38.1505019211879 | 21.6029190992233 | 14.3187385244159 | 30.9461507623369 |
| chr16 | 46581059 | 46581075 | - | 3.01188173062009 | 2.27399148412876 | 2.38645642073598 | 6.40265188186281 |
| chr16 | 46581146 | 46581156 | - | 5.01980288436682 | 1.13699574206438 | 1.19322821036799 | 2.13421729395427 |
| chr16 | 46581395 | 46581405 | - | 3.01188173062009 | 5.68497871032191 | 4.77291284147196 | 2.13421729395427 |
| chr16 | 46581580 | 46581608 | - | 10.0396057687336 | 2.27399148412876 | 2.38645642073598 | 5.33554323488568 |
| chr16 | 46658543 | 46658561 | - | 1.00396057687336 | 1.13699574206438 | 2.38645642073598 | 3.20132594093141 |
| chr16 | 46659701 | 46659705 | - | 2.00792115374673 | 7.95897019445067 | 4.77291284147196 | 3.20132594093141 |
| chr16 | 46659734 | 46659744 | - | 6.02376346124019 | 11.3699574206438 | 9.54582568294392 | 3.20132594093141 |
| chr16 | 46660071 | 46660096 | - | 75.2970432655023 | 76.1787147183136 | 34.6036181006717 | 45.8856718200168 |
| chr16 | 46660152 | 46660180 | - | 3.01188173062009 | 5.68497871032191 | 2.38645642073598 | 11.7381951167485 |
| chr16 | 46660278 | 46660433 | - | 18.0712903837206 | 23.876910583352 | 26.2510206280958 | 19.2079556455884 |
| chr16 | 46660466 | 46660542 | - | 84.3326884573626 | 75.0417189762492 | 42.9562155732476 | 94.9726695809651 |
| chr16 | 46698088 | 46698102 | + | 7.02772403811355 | 9.09596593651506 | 3.57968463110397 | 5.33554323488568 |
| chr16 | 46703727 | 46703734 | - | 1.00396057687336 | 1.13699574206438 | 5.96614105183995 | 3.20132594093141 |
| chr16 | 46802422 | 46802446 | - | 2.00792115374673 | 10.2329616785794 | 2.38645642073598 | 2.13421729395427 |
| chr16 | 46802633 | 46802648 | - | 9.03564519186028 | 14.780944646837 | 2.38645642073598 | 7.46976052883995 |
| chr16 | 46928981 | 46929026 | + | 3.01188173062009 | 5.68497871032191 | 2.38645642073598 | 1.06710864697714 |
| chr16 | 46929050 | 46929428 | + | 11.043566345607 | 45.4798296825753 | 13.1255103140479 | 7.46976052883995 |
| chr16 | 46930704 | 46930763 | + | 7.02772403811355 | 15.9179403889013 | 2.38645642073598 | 4.26843458790854 |
| chr16 | 46930838 | 46931258 | + | 88.3485307648561 | 301.303871647061 | 47.7291284147196 | 27.7448248214055 |
| chr16 | 46956496 | 46956565 | - | 8.03168461498692 | 5.68497871032191 | 11.9322821036799 | 9.60397782279422 |
| chr16 | 46956613 | 46956686 | - | 9.03564519186028 | 7.95897019445067 | 5.96614105183995 | 7.46976052883995 |
| chr16 | 46956710 | 46956852 | - | 25.0990144218341 | 17.0549361309657 | 22.6713359969918 | 22.4092815865198 |
| chr16 | 46956873 | 46957080 | - | 79.3128855729958 | 68.2197445238629 | 17.8984231555198 | 58.6909755837425 |
| chr16 | 47082191 | 47082197 | - | 8.03168461498692 | 2.27399148412876 | 1.19322821036799 | 1.06710864697714 |
| chr16 | 47083035 | 47083036 | - | 1.00396057687336 | 1.13699574206438 | 1.19322821036799 | 1.06710864697714 |
| chr16 | 47155007 | 47155021 | - | 4.01584230749346 | 5.68497871032191 | 1.19322821036799 | 5.33554323488568 |
| chr16 | 47155100 | 47155102 | - | 4.01584230749346 | 7.95897019445067 | 1.19322821036799 | 3.20132594093141 |
| chr16 | 47155406 | 47155421 | - | 12.0475269224804 | 15.9179403889013 | 7.15936926220794 | 4.26843458790854 |
| chr16 | 47155447 | 47155478 | - | 11.043566345607 | 6.82197445238629 | 7.15936926220794 | 2.13421729395427 |
| chr16 | 47155555 | 47155776 | - | 30.1188173062009 | 37.5208594881246 | 17.8984231555198 | 30.9461507623369 |
| chr16 | 47699474 | 47699536 | + | 2.00792115374673 | 6.82197445238629 | 1.19322821036799 | 6.40265188186281 |
| chr16 | 47699753 | 47699798 | + | 6.02376346124019 | 4.54798296825753 | 9.54582568294392 | 4.26843458790854 |
| chr16 | 47699961 | 47699971 | + | 5.01980288436682 | 5.68497871032191 | 3.57968463110397 | 7.46976052883995 |
| chr16 | 48351778 | 48351823 | + | 16.0633692299738 | 10.2329616785794 | 8.35259747257593 | 3.20132594093141 |
| chr16 | 48352822 | 48352859 | + | 7.02772403811355 | 15.9179403889013 | 11.9322821036799 | 12.8053037637256 |
| chr16 | 48354104 | 48354115 | + | 14.0554480762271 | 13.6439489047726 | 20.2848795762558 | 13.8724124107028 |
| chr16 | 48354486 | 48354505 | + | 9.03564519186028 | 7.95897019445067 | 23.8645642073598 | 12.8053037637256 |
| chr16 | 48355318 | 48355327 | + | 3.01188173062009 | 4.54798296825753 | 1.19322821036799 | 4.26843458790854 |
| chr16 | 48356059 | 48356065 | + | 4.01584230749346 | 10.2329616785794 | 7.15936926220794 | 17.0737383516342 |
| chr16 | 48356271 | 48356273 | + | 5.01980288436682 | 1.13699574206438 | 1.19322821036799 | 1.06710864697714 |
| chr16 | 48361162 | 48361181 | - | 71.2812009580089 | 87.5486721389574 | 26.2510206280958 | 33.0803680562912 |
| chr16 | 48361216 | 48361346 | - | 171.677258645345 | 208.070220797782 | 75.1733772531833 | 75.7647139353766 |
| chr16 | 48361445 | 48361512 | - | 16.0633692299738 | 42.0688424563821 | 13.1255103140479 | 5.33554323488568 |
| chr16 | 48361557 | 48361559 | - | 1.00396057687336 | 1.13699574206438 | 1.19322821036799 | 1.06710864697714 |
| chr16 | 48542560 | 48542583 | - | 1.00396057687336 | 5.68497871032191 | 1.19322821036799 | 1.06710864697714 |
| chr16 | 50105852 | 50105863 | + | 23.0910932680874 | 38.657855230189 | 65.6275515702394 | 41.6172372321083 |
| chr16 | 50106348 | 50106358 | + | 4.01584230749346 | 12.5069531627082 | 5.96614105183995 | 6.40265188186281 |
| chr16 | 50229338 | 50229348 | + | 8.03168461498692 | 5.68497871032191 | 7.15936926220794 | 3.20132594093141 |
| chr16 | 50229392 | 50229433 | + | 3.01188173062009 | 3.41098722619315 | 1.19322821036799 | 1.06710864697714 |
| chr16 | 50229583 | 50229588 | + | 2.00792115374673 | 5.68497871032191 | 5.96614105183995 | 3.20132594093141 |
| chr16 | 50229667 | 50229681 | + | 3.01188173062009 | 7.95897019445067 | 4.77291284147196 | 5.33554323488568 |
| chr16 | 50229703 | 50229775 | + | 222.879248065887 | 226.262152670812 | 137.221244192319 | 160.06629704657 |
| chr16 | 50229970 | 50229989 | + | 7.02772403811355 | 7.95897019445067 | 5.96614105183995 | 3.20132594093141 |
| chr16 | 50231310 | 50231330 | + | 5.01980288436682 | 2.27399148412876 | 5.96614105183995 | 4.26843458790854 |
| chr16 | 50231334 | 50231354 | + | 22.087132691214 | 46.6168254246397 | 7.15936926220794 | 16.006629704657 |
| chr16 | 50231420 | 50231425 | + | 3.01188173062009 | 4.54798296825753 | 1.19322821036799 | 5.33554323488568 |
| chr16 | 50231507 | 50231615 | + | 165.653495184105 | 219.440178218426 | 106.197310722751 | 129.120146284233 |
| chr16 | 50317808 | 50317816 | + | 3.01188173062009 | 2.27399148412876 | 4.77291284147196 | 14.9395210576799 |
| chr16 | 50317990 | 50318009 | + | 2.00792115374673 | 1.13699574206438 | 2.38645642073598 | 2.13421729395427 |
| chr16 | 50800284 | 50800308 | + | 2.00792115374673 | 18.1919318730301 | 4.77291284147196 | 5.33554323488568 |
| chr16 | 50801648 | 50801689 | + | 55.217831728035 | 128.480518853275 | 9.54582568294392 | 14.9395210576799 |
| chr16 | 52280517 | 52280518 | + | 46.1821865361748 | 38.657855230189 | 9.54582568294392 | 1.06710864697714 |
| chr16 | 53326352 | 53326355 | + | 4.01584230749346 | 6.82197445238629 | 1.19322821036799 | 3.20132594093141 |
| chr16 | 53327098 | 53327100 | + | 6.02376346124019 | 11.3699574206438 | 5.96614105183995 | 14.9395210576799 |
| chr16 | 53490863 | 53491011 | + | 19.0752509605939 | 7.95897019445067 | 19.0916513658878 | 18.1408469986113 |
| chr16 | 53491096 | 53491115 | + | 1.00396057687336 | 2.27399148412876 | 3.57968463110397 | 4.26843458790854 |
| chr16 | 53491181 | 53491378 | + | 449.774338439267 | 384.304560817761 | 233.872729232126 | 169.670274869365 |
| chr16 | 53491512 | 53491579 | + | 24.0950538449607 | 57.9867828452835 | 91.8785721983352 | 30.9461507623369 |
| chr16 | 54114197 | 54114230 | + | 3.01188173062009 | 3.41098722619315 | 1.19322821036799 | 9.60397782279422 |
| chr16 | 54933878 | 54933893 | + | 3.01188173062009 | 7.95897019445067 | 1.19322821036799 | 2.13421729395427 |
| chr16 | 54934009 | 54934017 | + | 3.01188173062009 | 4.54798296825753 | 2.38645642073598 | 3.20132594093141 |
| chr16 | 56361552 | 56361570 | - | 1.00396057687336 | 1.13699574206438 | 3.57968463110397 | 1.06710864697714 |
| chr16 | 56361603 | 56361624 | - | 2.00792115374673 | 3.41098722619315 | 1.19322821036799 | 1.06710864697714 |
| chr16 | 56361692 | 56361717 | - | 1.00396057687336 | 4.54798296825753 | 7.15936926220794 | 10.6710864697714 |
| chr16 | 56362018 | 56362037 | - | 3.01188173062009 | 9.09596593651506 | 4.77291284147196 | 2.13421729395427 |
| chr16 | 56362122 | 56362152 | - | 4.01584230749346 | 4.54798296825753 | 8.35259747257593 | 19.2079556455884 |
| chr16 | 56362530 | 56362556 | - | 4.01584230749346 | 4.54798296825753 | 9.54582568294392 | 56.5567582897882 |
| chr16 | 56362598 | 56362647 | - | 4.01584230749346 | 2.27399148412876 | 11.9322821036799 | 22.4092815865198 |
| chr16 | 56362680 | 56362701 | - | 3.01188173062009 | 5.68497871032191 | 10.7390538933119 | 5.33554323488568 |
| chr16 | 56362762 | 56362828 | - | 14.0554480762271 | 15.9179403889013 | 59.6614105183995 | 36.2816939972226 |
| chr16 | 56362852 | 56362869 | - | 2.00792115374673 | 3.41098722619315 | 7.15936926220794 | 10.6710864697714 |
| chr16 | 56429388 | 56429398 | - | 2.00792115374673 | 3.41098722619315 | 2.38645642073598 | 2.13421729395427 |
| chr16 | 56429431 | 56429476 | - | 5.01980288436682 | 5.68497871032191 | 5.96614105183995 | 7.46976052883995 |
| chr16 | 56429498 | 56429500 | - | 8.03168461498692 | 3.41098722619315 | 1.19322821036799 | 2.13421729395427 |
| chr16 | 56430511 | 56430514 | - | 3.01188173062009 | 1.13699574206438 | 1.19322821036799 | 2.13421729395427 |
| chr16 | 56430700 | 56430713 | - | 2.00792115374673 | 1.13699574206438 | 1.19322821036799 | 2.13421729395427 |
| chr16 | 56431777 | 56431791 | - | 1.00396057687336 | 4.54798296825753 | 3.57968463110397 | 7.46976052883995 |
| chr16 | 56431931 | 56431963 | - | 3.01188173062009 | 5.68497871032191 | 5.96614105183995 | 5.33554323488568 |
| chr16 | 56432010 | 56432029 | - | 12.0475269224804 | 15.9179403889013 | 10.7390538933119 | 2.13421729395427 |
| chr16 | 56432598 | 56432736 | - | 32.1267384599477 | 29.5618892936739 | 40.5697591525116 | 52.2883237018796 |
| chr16 | 56477343 | 56477356 | + | 4.01584230749346 | 5.68497871032191 | 3.57968463110397 | 5.33554323488568 |
| chr16 | 56477463 | 56477472 | + | 2.00792115374673 | 2.27399148412876 | 1.19322821036799 | 2.13421729395427 |
| chr16 | 56484600 | 56484679 | - | 2.00792115374673 | 19.3289276150945 | 3.57968463110397 | 1.06710864697714 |
| chr16 | 56609322 | 56609445 | + | 152.602007684751 | 122.795540142953 | 65.6275515702394 | 45.8856718200168 |
| chr16 | 56626974 | 56627017 | + | 36.1425807674411 | 65.9457530397342 | 2.38645642073598 | 2.13421729395427 |
| chr16 | 56682407 | 56682533 | + | 437.726811516787 | 483.223190377362 | 2811.24566362698 | 1143.94046955949 |
| chr16 | 56844625 | 56844716 | + | 40.1584230749346 | 20.4659233571589 | 45.3426719939836 | 44.8185631730397 |
| chr16 | 56844741 | 56844744 | + | 3.01188173062009 | 3.41098722619315 | 1.19322821036799 | 1.06710864697714 |
| chr16 | 56913706 | 56913707 | + | 21.0831721143407 | 17.0549361309657 | 29.8307052591997 | 34.1474767032683 |
| chr16 | 56943288 | 56943523 | + | 75.2970432655023 | 137.57648478979 | 66.8207797806074 | 41.6172372321083 |
| chr16 | 56943545 | 56943767 | + | 75.2970432655023 | 117.110561432631 | 35.7968463110397 | 26.6777161744284 |
| chr16 | 56943793 | 56943862 | + | 12.0475269224804 | 25.0139063254164 | 4.77291284147196 | 6.40265188186281 |
| chr16 | 57153590 | 57153592 | - | 1.00396057687336 | 1.13699574206438 | 1.19322821036799 | 1.06710864697714 |
| chr16 | 57153619 | 57153639 | - | 6.02376346124019 | 6.82197445238629 | 10.7390538933119 | 10.6710864697714 |
| chr16 | 57153676 | 57153733 | - | 40.1584230749346 | 39.7948509722534 | 13.1255103140479 | 30.9461507623369 |
| chr16 | 57153827 | 57153830 | - | 39.1544624980612 | 35.2468680039958 | 16.7051949451519 | 28.8119334683827 |
| chr16 | 57153895 | 57153899 | - | 3.01188173062009 | 1.13699574206438 | 3.57968463110397 | 3.20132594093141 |
| chr16 | 57153961 | 57154005 | - | 29.1148567293276 | 18.1919318730301 | 35.7968463110397 | 21.3421729395427 |
| chr16 | 57154246 | 57154272 | - | 5.01980288436682 | 7.95897019445067 | 4.77291284147196 | 7.46976052883995 |
| chr16 | 57240283 | 57240320 | + | 4.01584230749346 | 4.54798296825753 | 11.9322821036799 | 5.33554323488568 |
| chr16 | 57240413 | 57240419 | + | 2.00792115374673 | 1.13699574206438 | 1.19322821036799 | 3.20132594093141 |
| chr16 | 57252304 | 57252393 | + | 6.02376346124019 | 2.27399148412876 | 13.1255103140479 | 3.20132594093141 |
| chr16 | 57253321 | 57253382 | + | 12.0475269224804 | 27.2878978095452 | 10.7390538933119 | 21.3421729395427 |
| chr16 | 57253407 | 57253458 | + | 3.01188173062009 | 3.41098722619315 | 8.35259747257593 | 9.60397782279422 |
| chr16 | 57253531 | 57253572 | + | 4.01584230749346 | 7.95897019445067 | 10.7390538933119 | 4.26843458790854 |
| chr16 | 57429067 | 57429109 | - | 4.01584230749346 | 11.3699574206438 | 4.77291284147196 | 18.1408469986113 |
| chr16 | 57429189 | 57429212 | - | 11.043566345607 | 9.09596593651506 | 19.0916513658878 | 16.006629704657 |
| chr16 | 57460599 | 57460725 | + | 51.2019894205416 | 84.1376849127643 | 72.7869208324474 | 41.6172372321083 |
| chr16 | 57460785 | 57460824 | + | 4.01584230749346 | 5.68497871032191 | 13.1255103140479 | 10.6710864697714 |
| chr16 | 57460871 | 57460909 | + | 9.03564519186028 | 9.09596593651506 | 11.9322821036799 | 5.33554323488568 |
| chr16 | 57460985 | 57461028 | + | 21.0831721143407 | 18.1919318730301 | 8.35259747257593 | 17.0737383516342 |
| chr16 | 57461051 | 57461124 | + | 503.988209590429 | 397.948509722534 | 156.312895558207 | 369.219591854089 |
| chr16 | 57461206 | 57461238 | + | 2.00792115374673 | 4.54798296825753 | 1.19322821036799 | 1.06710864697714 |
| chr16 | 57462634 | 57462724 | + | 765.017959577504 | 594.648773099672 | 492.80325088198 | 581.574212602539 |
| chr16 | 57471146 | 57471212 | + | 12.0475269224804 | 15.9179403889013 | 26.2510206280958 | 22.4092815865198 |
| chr16 | 57471257 | 57471306 | + | 15.0594086531005 | 6.82197445238629 | 19.0916513658878 | 27.7448248214055 |
| chr16 | 57471531 | 57471538 | + | 5.01980288436682 | 12.5069531627082 | 7.15936926220794 | 7.46976052883995 |
| chr16 | 57471565 | 57471574 | + | 1.00396057687336 | 2.27399148412876 | 2.38645642073598 | 2.13421729395427 |
| chr16 | 57471752 | 57471761 | + | 14.0554480762271 | 5.68497871032191 | 5.96614105183995 | 11.7381951167485 |
| chr16 | 57471896 | 57471958 | + | 34.1346596136944 | 61.3977700714766 | 48.9223566250876 | 34.1474767032683 |
| chr16 | 57471985 | 57472131 | - | 6.02376346124019 | 11.3699574206438 | 3.57968463110397 | 1.06710864697714 |
| chr16 | 57472492 | 57472537 | - | 10.0396057687336 | 5.68497871032191 | 1.19322821036799 | 3.20132594093141 |
| chr16 | 57473165 | 57473185 | - | 13.0514874993537 | 4.54798296825753 | 4.77291284147196 | 1.06710864697714 |
| chr16 | 57757067 | 57757123 | + | 48.1901076899215 | 34.1098722619315 | 31.0239334695677 | 51.2212150549025 |
| chr16 | 57757145 | 57757218 | + | 11.043566345607 | 7.95897019445067 | 4.77291284147196 | 14.9395210576799 |
| chr16 | 57758311 | 57758341 | - | 5.01980288436682 | 4.54798296825753 | 2.38645642073598 | 2.13421729395427 |
| chr16 | 57758369 | 57758447 | - | 43.1703048055547 | 32.9728765198671 | 19.0916513658878 | 44.8185631730397 |
| chr16 | 57758481 | 57758505 | - | 11.043566345607 | 10.2329616785794 | 9.54582568294392 | 44.8185631730397 |
| chr16 | 57758528 | 57758543 | - | 6.02376346124019 | 5.68497871032191 | 9.54582568294392 | 16.006629704657 |
| chr16 | 57758604 | 57758635 | - | 20.0792115374673 | 10.2329616785794 | 20.2848795762558 | 17.0737383516342 |
| chr16 | 57758686 | 57758702 | - | 3.01188173062009 | 3.41098722619315 | 3.57968463110397 | 20.2750642925656 |
| chr16 | 57758717 | 57758763 | - | 12.0475269224804 | 9.09596593651506 | 14.3187385244159 | 37.3488026441998 |
| chr16 | 57758795 | 57758823 | - | 13.0514874993537 | 6.82197445238629 | 8.35259747257593 | 10.6710864697714 |
| chr16 | 57758861 | 57758912 | - | 8.03168461498692 | 9.09596593651506 | 4.77291284147196 | 8.53686917581709 |
| chr16 | 57759135 | 57759148 | - | 7.02772403811355 | 1.13699574206438 | 2.38645642073598 | 1.06710864697714 |
| chr16 | 58020289 | 58020314 | + | 1.00396057687336 | 2.27399148412876 | 1.19322821036799 | 6.40265188186281 |
| chr16 | 58020353 | 58020406 | + | 7.02772403811355 | 4.54798296825753 | 14.3187385244159 | 14.9395210576799 |
| chr16 | 58021062 | 58021101 | + | 1.00396057687336 | 6.82197445238629 | 17.8984231555198 | 14.9395210576799 |
| chr16 | 58021160 | 58021173 | + | 3.01188173062009 | 2.27399148412876 | 5.96614105183995 | 3.20132594093141 |
| chr16 | 58021195 | 58021208 | + | 7.02772403811355 | 1.13699574206438 | 10.7390538933119 | 2.13421729395427 |
| chr16 | 58021255 | 58021279 | + | 4.01584230749346 | 3.41098722619315 | 4.77291284147196 | 11.7381951167485 |
| chr16 | 58021351 | 58021390 | + | 17.0673298068472 | 3.41098722619315 | 3.57968463110397 | 13.8724124107028 |
| chr16 | 58021436 | 58021461 | + | 4.01584230749346 | 4.54798296825753 | 4.77291284147196 | 5.33554323488568 |
| chr16 | 58021501 | 58021574 | + | 8.03168461498692 | 6.82197445238629 | 7.15936926220794 | 4.26843458790854 |
| chr16 | 58046339 | 58046440 | + | 17.0673298068472 | 4.54798296825753 | 16.7051949451519 | 7.46976052883995 |
| chr16 | 58046477 | 58046533 | + | 6.02376346124019 | 3.41098722619315 | 5.96614105183995 | 7.46976052883995 |
| chr16 | 58046561 | 58046570 | + | 3.01188173062009 | 1.13699574206438 | 7.15936926220794 | 1.06710864697714 |
| chr16 | 58046671 | 58046703 | + | 5.01980288436682 | 1.13699574206438 | 2.38645642073598 | 8.53686917581709 |
| chr16 | 58113643 | 58113649 | - | 1.00396057687336 | 2.27399148412876 | 1.19322821036799 | 2.13421729395427 |
| chr16 | 58113761 | 58113837 | - | 28.1108961524542 | 44.3428339405109 | 21.4781077866238 | 37.3488026441998 |
| chr16 | 58113950 | 58114023 | - | 9.03564519186028 | 9.09596593651506 | 35.7968463110397 | 17.0737383516342 |
| chr16 | 58129517 | 58129580 | + | 2.00792115374673 | 3.41098722619315 | 16.7051949451519 | 19.2079556455884 |
| chr16 | 58157961 | 58157993 | - | 1.00396057687336 | 13.6439489047726 | 13.1255103140479 | 8.53686917581709 |
| chr16 | 58405882 | 58405884 | + | 1.00396057687336 | 4.54798296825753 | 2.38645642073598 | 1.06710864697714 |
| chr16 | 58519103 | 58519114 | + | 15.0594086531005 | 23.876910583352 | 5.96614105183995 | 6.40265188186281 |
| chr16 | 58519144 | 58519194 | + | 37.1465413443145 | 40.9318467143178 | 10.7390538933119 | 7.46976052883995 |
| chr16 | 58519223 | 58519232 | + | 3.01188173062009 | 2.27399148412876 | 3.57968463110397 | 2.13421729395427 |
| chr16 | 58519352 | 58519354 | + | 9.03564519186028 | 2.27399148412876 | 1.19322821036799 | 1.06710864697714 |
| chr16 | 58519668 | 58519672 | + | 15.0594086531005 | 22.7399148412876 | 8.35259747257593 | 6.40265188186281 |
| chr16 | 58520684 | 58520758 | - | 7.02772403811355 | 14.780944646837 | 8.35259747257593 | 18.1408469986113 |
| chr16 | 58520791 | 58520842 | - | 5.01980288436682 | 7.95897019445067 | 4.77291284147196 | 4.26843458790854 |
| chr16 | 58665139 | 58665145 | - | 73.2891221117556 | 55.7127913611547 | 7.15936926220794 | 39.483019938154 |
| chr16 | 58665184 | 58665193 | - | 36.1425807674411 | 36.3838637460602 | 4.77291284147196 | 8.53686917581709 |
| chr16 | 58665300 | 58665310 | - | 1.00396057687336 | 1.13699574206438 | 2.38645642073598 | 1.06710864697714 |
| chr16 | 58666481 | 58666515 | - | 5.01980288436682 | 9.09596593651506 | 1.19322821036799 | 1.06710864697714 |
| chr16 | 58707151 | 58707188 | - | 27.1069355755808 | 18.1919318730301 | 16.7051949451519 | 11.7381951167485 |
| chr16 | 58707200 | 58707213 | - | 2.00792115374673 | 4.54798296825753 | 4.77291284147196 | 4.26843458790854 |
| chr16 | 58707240 | 58707270 | - | 9.03564519186028 | 2.27399148412876 | 3.57968463110397 | 2.13421729395427 |
| chr16 | 58707498 | 58707529 | - | 10.0396057687336 | 5.68497871032191 | 7.15936926220794 | 6.40265188186281 |
| chr16 | 58707553 | 58707626 | - | 39.1544624980612 | 23.876910583352 | 40.5697591525116 | 36.2816939972226 |
| chr16 | 58707660 | 58707841 | - | 55.217831728035 | 34.1098722619315 | 68.0140079909754 | 33.0803680562912 |
| chr16 | 58707892 | 58707909 | - | 2.00792115374673 | 1.13699574206438 | 4.77291284147196 | 3.20132594093141 |
| chr16 | 58707945 | 58707998 | - | 61.2415951892752 | 31.8358807778027 | 40.5697591525116 | 44.8185631730397 |
| chr16 | 58708051 | 58708100 | - | 24.0950538449607 | 9.09596593651506 | 23.8645642073598 | 16.006629704657 |
| chr16 | 58708103 | 58708120 | - | 3.01188173062009 | 2.27399148412876 | 1.19322821036799 | 2.13421729395427 |
| chr16 | 58708156 | 58708175 | - | 8.03168461498692 | 10.2329616785794 | 9.54582568294392 | 11.7381951167485 |
| chr16 | 66510772 | 66510807 | - | 8.03168461498692 | 10.2329616785794 | 2.38645642073598 | 2.13421729395427 |
| chr16 | 66566022 | 66566050 | + | 2.00792115374673 | 7.95897019445067 | 3.57968463110397 | 1.06710864697714 |
| chr16 | 66614778 | 66614781 | - | 2.00792115374673 | 3.41098722619315 | 1.19322821036799 | 3.20132594093141 |
| chr16 | 66616527 | 66616549 | - | 10.0396057687336 | 5.68497871032191 | 7.15936926220794 | 3.20132594093141 |
| chr16 | 66617230 | 66617236 | - | 3.01188173062009 | 2.27399148412876 | 7.15936926220794 | 2.13421729395427 |
| chr16 | 66720913 | 66720937 | + | 15.0594086531005 | 5.68497871032191 | 1.19322821036799 | 5.33554323488568 |
| chr16 | 66720924 | 66721041 | - | 130.514874993537 | 127.343523111211 | 23.8645642073598 | 34.1474767032683 |
| chr16 | 66721243 | 66721323 | - | 8.03168461498692 | 6.82197445238629 | 1.19322821036799 | 3.20132594093141 |
| chr16 | 66721595 | 66721623 | - | 2.00792115374673 | 1.13699574206438 | 1.19322821036799 | 4.26843458790854 |
| chr16 | 66721642 | 66721713 | - | 3.01188173062009 | 10.2329616785794 | 4.77291284147196 | 6.40265188186281 |
| chr16 | 66721792 | 66721840 | - | 2.00792115374673 | 10.2329616785794 | 2.38645642073598 | 2.13421729395427 |
| chr16 | 66721875 | 66721916 | - | 11.043566345607 | 4.54798296825753 | 1.19322821036799 | 1.06710864697714 |
| chr16 | 66722205 | 66722232 | - | 8.03168461498692 | 10.2329616785794 | 1.19322821036799 | 1.06710864697714 |
| chr16 | 66722872 | 66723004 | - | 52.205949997415 | 83.0006891706999 | 42.9562155732476 | 43.7514545260626 |
| chr16 | 66723030 | 66723076 | - | 14.0554480762271 | 21.6029190992233 | 9.54582568294392 | 10.6710864697714 |
| chr16 | 66723210 | 66723326 | - | 20.0792115374673 | 22.7399148412876 | 4.77291284147196 | 8.53686917581709 |
| chr16 | 66723402 | 66723569 | - | 59.2336740355285 | 92.0966551072149 | 41.7629873628796 | 20.2750642925656 |
| chr16 | 66723610 | 66723619 | - | 16.0633692299738 | 47.753821166704 | 20.2848795762558 | 5.33554323488568 |
| chr16 | 66723652 | 66723678 | - | 2.00792115374673 | 3.41098722619315 | 3.57968463110397 | 3.20132594093141 |
| chr16 | 66802908 | 66802926 | - | 19.0752509605939 | 21.6029190992233 | 45.3426719939836 | 27.7448248214055 |
| chr16 | 66802951 | 66803016 | - | 10.0396057687336 | 9.09596593651506 | 14.3187385244159 | 12.8053037637256 |
| chr16 | 66932078 | 66932274 | - | 55.217831728035 | 75.0417189762492 | 41.7629873628796 | 67.2278447595595 |
| chr16 | 66942760 | 66942786 | + | 3.01188173062009 | 5.68497871032191 | 2.38645642073598 | 3.20132594093141 |
| chr16 | 66944221 | 66944284 | + | 5.01980288436682 | 3.41098722619315 | 2.38645642073598 | 2.13421729395427 |
| chr16 | 66944981 | 66945069 | + | 12.0475269224804 | 18.1919318730301 | 9.54582568294392 | 7.46976052883995 |
| chr16 | 66975069 | 66975073 | + | 6.02376346124019 | 4.54798296825753 | 1.19322821036799 | 1.06710864697714 |
| chr16 | 67098708 | 67098756 | + | 33.130699036821 | 34.1098722619315 | 51.3088130458235 | 19.2079556455884 |
| chr16 | 67098792 | 67098824 | + | 4.01584230749346 | 1.13699574206438 | 4.77291284147196 | 7.46976052883995 |
| chr16 | 67098875 | 67098914 | + | 4.01584230749346 | 13.6439489047726 | 11.9322821036799 | 9.60397782279422 |
| chr16 | 67098976 | 67098988 | + | 3.01188173062009 | 7.95897019445067 | 4.77291284147196 | 7.46976052883995 |
| chr16 | 67099026 | 67099044 | + | 23.0910932680874 | 17.0549361309657 | 11.9322821036799 | 14.9395210576799 |
| chr16 | 67099094 | 67099149 | + | 78.3089249961224 | 102.329616785794 | 93.0718004087032 | 74.6976052883995 |
| chr16 | 67099176 | 67099219 | + | 10.0396057687336 | 5.68497871032191 | 16.7051949451519 | 16.006629704657 |
| chr16 | 67099398 | 67099410 | + | 1.00396057687336 | 1.13699574206438 | 3.57968463110397 | 2.13421729395427 |
| chr16 | 67100306 | 67100349 | + | 5.01980288436682 | 7.95897019445067 | 1.19322821036799 | 7.46976052883995 |
| chr16 | 67100581 | 67100606 | + | 3.01188173062009 | 2.27399148412876 | 3.57968463110397 | 3.20132594093141 |
| chr16 | 67100808 | 67100812 | + | 2.00792115374673 | 2.27399148412876 | 1.19322821036799 | 2.13421729395427 |
| chr16 | 67147189 | 67147243 | + | 2.00792115374673 | 2.27399148412876 | 5.96614105183995 | 3.20132594093141 |
| chr16 | 67147989 | 67148045 | + | 3.01188173062009 | 3.41098722619315 | 1.19322821036799 | 6.40265188186281 |
| chr16 | 67148438 | 67148444 | + | 2.00792115374673 | 1.13699574206438 | 1.19322821036799 | 5.33554323488568 |
| chr16 | 67176168 | 67176187 | - | 5.01980288436682 | 10.2329616785794 | 20.2848795762558 | 6.40265188186281 |
| chr16 | 67176309 | 67176325 | - | 6.02376346124019 | 2.27399148412876 | 2.38645642073598 | 3.20132594093141 |
| chr16 | 67176345 | 67176369 | - | 7.02772403811355 | 7.95897019445067 | 8.35259747257593 | 5.33554323488568 |
| chr16 | 67198208 | 67198232 | + | 9.03564519186028 | 1.13699574206438 | 7.15936926220794 | 2.13421729395427 |
| chr16 | 67198396 | 67198472 | + | 21.0831721143407 | 9.09596593651506 | 42.9562155732476 | 50.1541064079254 |
| chr16 | 67198505 | 67198537 | + | 7.02772403811355 | 5.68497871032191 | 17.8984231555198 | 12.8053037637256 |
| chr16 | 67198615 | 67198622 | + | 3.01188173062009 | 2.27399148412876 | 1.19322821036799 | 2.13421729395427 |
| chr16 | 67198641 | 67198707 | + | 88.3485307648561 | 48.8908169087684 | 52.5020412561915 | 103.509538756782 |
| chr16 | 67229185 | 67229255 | + | 49.1940682667949 | 63.6717615556054 | 26.2510206280958 | 35.2145853502455 |
| chr16 | 67229417 | 67229425 | - | 7.02772403811355 | 2.27399148412876 | 11.9322821036799 | 12.8053037637256 |
| chr16 | 67481321 | 67481408 | + | 38.1505019211879 | 29.5618892936739 | 48.9223566250876 | 59.7580842307196 |
| chr16 | 67546705 | 67546726 | + | 4.01584230749346 | 4.54798296825753 | 1.19322821036799 | 1.06710864697714 |
| chr16 | 67638345 | 67638366 | + | 4.01584230749346 | 2.27399148412876 | 2.38645642073598 | 2.13421729395427 |
| chr16 | 67638466 | 67638487 | + | 3.01188173062009 | 2.27399148412876 | 2.38645642073598 | 5.33554323488568 |
| chr16 | 67638610 | 67638658 | + | 28.1108961524542 | 32.9728765198671 | 10.7390538933119 | 19.2079556455884 |
| chr16 | 67638762 | 67638783 | + | 21.0831721143407 | 35.2468680039958 | 16.7051949451519 | 16.006629704657 |
| chr16 | 67638791 | 67638846 | + | 52.205949997415 | 76.1787147183136 | 46.5359002043516 | 43.7514545260626 |
| chr16 | 67638902 | 67638958 | + | 5.01980288436682 | 14.780944646837 | 14.3187385244159 | 6.40265188186281 |
| chr16 | 67638974 | 67639053 | + | 32.1267384599477 | 34.1098722619315 | 32.2171616799357 | 26.6777161744284 |
| chr16 | 67639074 | 67639132 | + | 44.174265382428 | 64.8087572976698 | 17.8984231555198 | 23.476390233497 |
| chr16 | 67657580 | 67657634 | - | 8.03168461498692 | 14.780944646837 | 2.38645642073598 | 8.53686917581709 |
| chr16 | 67662992 | 67663009 | - | 1.00396057687336 | 11.3699574206438 | 4.77291284147196 | 11.7381951167485 |
| chr16 | 67726137 | 67726158 | - | 8.03168461498692 | 3.41098722619315 | 7.15936926220794 | 5.33554323488568 |
| chr16 | 67828181 | 67828204 | - | 5.01980288436682 | 2.27399148412876 | 2.38645642073598 | 8.53686917581709 |
| chr16 | 67828229 | 67828231 | - | 3.01188173062009 | 3.41098722619315 | 1.19322821036799 | 1.06710864697714 |
| chr16 | 67828670 | 67828706 | - | 8.03168461498692 | 9.09596593651506 | 22.6713359969918 | 26.6777161744284 |
| chr16 | 67828722 | 67828738 | - | 3.01188173062009 | 3.41098722619315 | 1.19322821036799 | 1.06710864697714 |
| chr16 | 67829880 | 67829902 | - | 4.01584230749346 | 2.27399148412876 | 5.96614105183995 | 18.1408469986113 |
| chr16 | 67829925 | 67830065 | - | 50.1980288436682 | 67.0827487817985 | 242.225326704702 | 105.643756050736 |
| chr16 | 67830396 | 67830461 | - | 6.02376346124019 | 7.95897019445067 | 15.5119667347839 | 16.006629704657 |
| chr16 | 67830540 | 67830544 | - | 5.01980288436682 | 1.13699574206438 | 1.19322821036799 | 3.20132594093141 |
| chr16 | 67831239 | 67831276 | - | 1.00396057687336 | 3.41098722619315 | 5.96614105183995 | 3.20132594093141 |
| chr16 | 67843541 | 67843556 | + | 3.01188173062009 | 3.41098722619315 | 8.35259747257593 | 5.33554323488568 |
| chr16 | 67843585 | 67843642 | + | 15.0594086531005 | 7.95897019445067 | 14.3187385244159 | 4.26843458790854 |
| chr16 | 67843650 | 67843765 | + | 24.0950538449607 | 11.3699574206438 | 14.3187385244159 | 3.20132594093141 |
| chr16 | 67843779 | 67843833 | + | 23.0910932680874 | 14.780944646837 | 21.4781077866238 | 28.8119334683827 |
| chr16 | 67843856 | 67843916 | + | 12.0475269224804 | 6.82197445238629 | 4.77291284147196 | 3.20132594093141 |
| chr16 | 67843930 | 67844020 | + | 95.3762548029696 | 53.438799877026 | 17.8984231555198 | 37.3488026441998 |
| chr16 | 67870892 | 67870929 | + | 13.0514874993537 | 4.54798296825753 | 11.9322821036799 | 20.2750642925656 |
| chr16 | 67871142 | 67871154 | + | 7.02772403811355 | 1.13699574206438 | 7.15936926220794 | 20.2750642925656 |
| chr16 | 67871296 | 67871298 | + | 1.00396057687336 | 1.13699574206438 | 4.77291284147196 | 1.06710864697714 |
| chr16 | 67872310 | 67872321 | + | 15.0594086531005 | 6.82197445238629 | 14.3187385244159 | 9.60397782279422 |
| chr16 | 67872411 | 67872415 | + | 1.00396057687336 | 1.13699574206438 | 2.38645642073598 | 6.40265188186281 |
| chr16 | 67882726 | 67882737 | + | 1.00396057687336 | 1.13699574206438 | 4.77291284147196 | 7.46976052883995 |
| chr16 | 67882759 | 67882762 | + | 2.00792115374673 | 1.13699574206438 | 2.38645642073598 | 2.13421729395427 |
| chr16 | 67884268 | 67884297 | + | 1.00396057687336 | 1.13699574206438 | 1.19322821036799 | 2.13421729395427 |
| chr16 | 67884320 | 67884433 | + | 12.0475269224804 | 11.3699574206438 | 16.7051949451519 | 14.9395210576799 |
| chr16 | 67928506 | 67928568 | + | 5.01980288436682 | 6.82197445238629 | 3.57968463110397 | 4.26843458790854 |
| chr16 | 67929461 | 67929463 | + | 5.01980288436682 | 2.27399148412876 | 4.77291284147196 | 7.46976052883995 |
| chr16 | 67929655 | 67929658 | + | 1.00396057687336 | 4.54798296825753 | 2.38645642073598 | 3.20132594093141 |
| chr16 | 67934530 | 67934602 | - | 34.1346596136944 | 43.2058381984465 | 17.8984231555198 | 27.7448248214055 |
| chr16 | 67934876 | 67934907 | - | 14.0554480762271 | 5.68497871032191 | 11.9322821036799 | 7.46976052883995 |
| chr16 | 67939901 | 67939968 | - | 2.00792115374673 | 2.27399148412876 | 2.38645642073598 | 2.13421729395427 |
| chr16 | 67944360 | 67944413 | - | 1.00396057687336 | 1.13699574206438 | 2.38645642073598 | 4.26843458790854 |
| chr16 | 68021457 | 68021519 | - | 166.657455760979 | 194.426271893009 | 62.0478669391355 | 125.918820343302 |
| chr16 | 68226558 | 68226888 | + | 269.061434602062 | 355.879667266152 | 863.897224306424 | 886.767285638 |
| chr16 | 68227087 | 68227090 | + | 1.00396057687336 | 2.27399148412876 | 2.38645642073598 | 1.06710864697714 |
| chr16 | 68227263 | 68227278 | + | 5.01980288436682 | 9.09596593651506 | 4.77291284147196 | 4.26843458790854 |
| chr16 | 68227452 | 68227558 | + | 5.01980288436682 | 3.41098722619315 | 27.4442488384638 | 30.9461507623369 |
| chr16 | 68228044 | 68228121 | + | 16.0633692299738 | 13.6439489047726 | 82.3327465153913 | 51.2212150549025 |
| chr16 | 68228170 | 68228227 | + | 3.01188173062009 | 3.41098722619315 | 11.9322821036799 | 16.006629704657 |
| chr16 | 68228265 | 68228282 | + | 2.00792115374673 | 1.13699574206438 | 2.38645642073598 | 7.46976052883995 |
| chr16 | 68228480 | 68228488 | + | 1.00396057687336 | 1.13699574206438 | 2.38645642073598 | 3.20132594093141 |
| chr16 | 68228630 | 68228665 | + | 6.02376346124019 | 4.54798296825753 | 7.15936926220794 | 6.40265188186281 |
| chr16 | 68228799 | 68228868 | + | 3.01188173062009 | 3.41098722619315 | 21.4781077866238 | 20.2750642925656 |
| chr16 | 68228976 | 68229051 | + | 5.01980288436682 | 5.68497871032191 | 23.8645642073598 | 36.2816939972226 |
| chr16 | 68229156 | 68229227 | + | 5.01980288436682 | 4.54798296825753 | 26.2510206280958 | 17.0737383516342 |
| chr16 | 68260986 | 68261026 | + | 1.00396057687336 | 3.41098722619315 | 3.57968463110397 | 5.33554323488568 |
| chr16 | 68297302 | 68297315 | + | 1.00396057687336 | 1.13699574206438 | 1.19322821036799 | 3.20132594093141 |
| chr16 | 68297535 | 68297563 | + | 37.1465413443145 | 28.4248935516096 | 22.6713359969918 | 8.53686917581709 |
| chr16 | 68297631 | 68297658 | + | 3.01188173062009 | 2.27399148412876 | 1.19322821036799 | 1.06710864697714 |
| chr16 | 68297786 | 68297808 | + | 41.1623836518079 | 29.5618892936739 | 9.54582568294392 | 14.9395210576799 |
| chr16 | 68299503 | 68299509 | + | 1.00396057687336 | 1.13699574206438 | 1.19322821036799 | 3.20132594093141 |
| chr16 | 68300678 | 68300680 | - | 3.01188173062009 | 4.54798296825753 | 5.96614105183995 | 4.26843458790854 |
| chr16 | 68300907 | 68300963 | + | 4.01584230749346 | 5.68497871032191 | 11.9322821036799 | 12.8053037637256 |
| chr16 | 68300931 | 68300946 | - | 34.1346596136944 | 34.1098722619315 | 9.54582568294392 | 12.8053037637256 |
| chr16 | 68300985 | 68301007 | + | 13.0514874993537 | 6.82197445238629 | 2.38645642073598 | 4.26843458790854 |
| chr16 | 68301046 | 68301051 | + | 1.00396057687336 | 1.13699574206438 | 4.77291284147196 | 3.20132594093141 |
| chr16 | 68301094 | 68301148 | + | 9.03564519186028 | 5.68497871032191 | 5.96614105183995 | 2.13421729395427 |
| chr16 | 68301104 | 68301146 | - | 6.02376346124019 | 7.95897019445067 | 10.7390538933119 | 7.46976052883995 |
| chr16 | 68301237 | 68301259 | + | 1.00396057687336 | 2.27399148412876 | 3.57968463110397 | 1.06710864697714 |
| chr16 | 68301292 | 68301299 | + | 4.01584230749346 | 4.54798296825753 | 2.38645642073598 | 9.60397782279422 |
| chr16 | 69118055 | 69118092 | - | 6.02376346124019 | 10.2329616785794 | 7.15936926220794 | 6.40265188186281 |
| chr16 | 69118114 | 69118153 | - | 49.1940682667949 | 114.836569948503 | 21.4781077866238 | 36.2816939972226 |
| chr16 | 69118651 | 69118732 | - | 13.0514874993537 | 11.3699574206438 | 31.0239334695677 | 26.6777161744284 |
| chr16 | 69118754 | 69118820 | - | 48.1901076899215 | 43.2058381984465 | 54.8884976769275 | 42.6843458790854 |
| chr16 | 69118881 | 69118909 | - | 8.03168461498692 | 3.41098722619315 | 10.7390538933119 | 13.8724124107028 |
| chr16 | 69118970 | 69118977 | - | 5.01980288436682 | 3.41098722619315 | 5.96614105183995 | 16.006629704657 |
| chr16 | 69119168 | 69119188 | - | 1.00396057687336 | 2.27399148412876 | 2.38645642073598 | 12.8053037637256 |
| chr16 | 69119335 | 69119339 | - | 6.02376346124019 | 6.82197445238629 | 16.7051949451519 | 9.60397782279422 |
| chr16 | 69119581 | 69119591 | - | 4.01584230749346 | 1.13699574206438 | 1.19322821036799 | 2.13421729395427 |
| chr16 | 69119645 | 69119673 | - | 4.01584230749346 | 6.82197445238629 | 5.96614105183995 | 9.60397782279422 |
| chr16 | 69119847 | 69119868 | - | 2.00792115374673 | 5.68497871032191 | 7.15936926220794 | 1.06710864697714 |
| chr16 | 69119915 | 69119946 | - | 3.01188173062009 | 2.27399148412876 | 7.15936926220794 | 3.20132594093141 |
| chr16 | 69305053 | 69305085 | + | 27.1069355755808 | 22.7399148412876 | 8.35259747257593 | 2.13421729395427 |
| chr16 | 69305684 | 69305755 | + | 30.1188173062009 | 34.1098722619315 | 11.9322821036799 | 17.0737383516342 |
| chr16 | 69307269 | 69307309 | + | 50.1980288436682 | 63.6717615556054 | 5.96614105183995 | 8.53686917581709 |
| chr16 | 69307335 | 69307361 | + | 46.1821865361748 | 45.4798296825753 | 14.3187385244159 | 13.8724124107028 |
| chr16 | 69324323 | 69324326 | + | 4.01584230749346 | 2.27399148412876 | 2.38645642073598 | 1.06710864697714 |
| chr16 | 69324962 | 69325014 | + | 8.03168461498692 | 26.1509020674808 | 16.7051949451519 | 14.9395210576799 |
| chr16 | 69327881 | 69327894 | - | 4.01584230749346 | 3.41098722619315 | 3.57968463110397 | 5.33554323488568 |
| chr16 | 69328573 | 69328592 | - | 14.0554480762271 | 3.41098722619315 | 5.96614105183995 | 5.33554323488568 |
| chr16 | 69328637 | 69328732 | - | 5.01980288436682 | 18.1919318730301 | 5.96614105183995 | 11.7381951167485 |
| chr16 | 69328753 | 69328769 | - | 16.0633692299738 | 15.9179403889013 | 2.38645642073598 | 10.6710864697714 |
| chr16 | 69328789 | 69328832 | - | 6.02376346124019 | 9.09596593651506 | 4.77291284147196 | 3.20132594093141 |
| chr16 | 69329134 | 69329143 | - | 4.01584230749346 | 6.82197445238629 | 2.38645642073598 | 2.13421729395427 |
| chr16 | 69342548 | 69342654 | + | 17.0673298068472 | 23.876910583352 | 15.5119667347839 | 38.4159112911769 |
| chr16 | 69342684 | 69342793 | + | 27.1069355755808 | 13.6439489047726 | 11.9322821036799 | 25.6106075274513 |
| chr16 | 69342881 | 69342887 | + | 1.00396057687336 | 5.68497871032191 | 3.57968463110397 | 5.33554323488568 |
| chr16 | 69356208 | 69356209 | - | 1.00396057687336 | 1.13699574206438 | 2.38645642073598 | 1.06710864697714 |
| chr16 | 69462730 | 69462778 | + | 2.00792115374673 | 4.54798296825753 | 2.38645642073598 | 4.26843458790854 |
| chr16 | 69462885 | 69462902 | + | 7.02772403811355 | 2.27399148412876 | 4.77291284147196 | 4.26843458790854 |
| chr16 | 69465916 | 69465918 | + | 9.03564519186028 | 10.2329616785794 | 3.57968463110397 | 3.20132594093141 |
| chr16 | 69466005 | 69466022 | + | 4.01584230749346 | 3.41098722619315 | 3.57968463110397 | 2.13421729395427 |
| chr16 | 69466045 | 69466049 | + | 10.0396057687336 | 14.780944646837 | 13.1255103140479 | 12.8053037637256 |
| chr16 | 69697912 | 69697927 | + | 3.01188173062009 | 6.82197445238629 | 1.19322821036799 | 3.20132594093141 |
| chr16 | 69698475 | 69698512 | + | 2.00792115374673 | 1.13699574206438 | 3.57968463110397 | 3.20132594093141 |
| chr16 | 69699424 | 69699441 | + | 11.043566345607 | 3.41098722619315 | 7.15936926220794 | 20.2750642925656 |
| chr16 | 69700907 | 69700939 | + | 8.03168461498692 | 4.54798296825753 | 1.19322821036799 | 11.7381951167485 |
| chr16 | 69701228 | 69701251 | + | 1.00396057687336 | 2.27399148412876 | 1.19322821036799 | 4.26843458790854 |
| chr16 | 69702064 | 69702071 | + | 2.00792115374673 | 2.27399148412876 | 2.38645642073598 | 2.13421729395427 |
| chr16 | 69702705 | 69702757 | + | 2.00792115374673 | 2.27399148412876 | 5.96614105183995 | 6.40265188186281 |
| chr16 | 69703175 | 69703244 | + | 10.0396057687336 | 6.82197445238629 | 9.54582568294392 | 20.2750642925656 |
| chr16 | 69703434 | 69703507 | + | 5.01980288436682 | 3.41098722619315 | 3.57968463110397 | 4.26843458790854 |
| chr16 | 69703686 | 69703755 | + | 2.00792115374673 | 3.41098722619315 | 15.5119667347839 | 6.40265188186281 |
| chr16 | 69703942 | 69703965 | + | 1.00396057687336 | 6.82197445238629 | 2.38645642073598 | 2.13421729395427 |
| chr16 | 69704408 | 69704471 | + | 1.00396057687336 | 4.54798296825753 | 2.38645642073598 | 1.06710864697714 |
| chr16 | 69704496 | 69704508 | + | 12.0475269224804 | 20.4659233571589 | 2.38645642073598 | 3.20132594093141 |
| chr16 | 69709456 | 69709514 | - | 19.0752509605939 | 7.95897019445067 | 1.19322821036799 | 1.06710864697714 |
| chr16 | 69710833 | 69710938 | - | 115.455466340437 | 73.9047232341848 | 9.54582568294392 | 18.1408469986113 |
| chr16 | 69710960 | 69711228 | - | 79.3128855729958 | 51.1648083928972 | 10.7390538933119 | 27.7448248214055 |
| chr16 | 69727005 | 69727039 | + | 92.3643730723495 | 46.6168254246397 | 20.2848795762558 | 27.7448248214055 |
| chr16 | 69741928 | 69741981 | - | 11.043566345607 | 25.0139063254164 | 13.1255103140479 | 6.40265188186281 |
| chr16 | 69742011 | 69742026 | - | 1.00396057687336 | 1.13699574206438 | 2.38645642073598 | 2.13421729395427 |
| chr16 | 69742061 | 69742222 | - | 8.03168461498692 | 10.2329616785794 | 7.15936926220794 | 18.1408469986113 |
| chr16 | 69941054 | 69941075 | + | 3.01188173062009 | 1.13699574206438 | 3.57968463110397 | 3.20132594093141 |
| chr16 | 69941552 | 69941579 | + | 2.00792115374673 | 3.41098722619315 | 2.38645642073598 | 2.13421729395427 |
| chr16 | 70158671 | 70158690 | + | 11.043566345607 | 6.82197445238629 | 2.38645642073598 | 1.06710864697714 |
| chr16 | 70161003 | 70161011 | + | 24.0950538449607 | 23.876910583352 | 22.6713359969918 | 22.4092815865198 |
| chr16 | 70161120 | 70161136 | + | 4.01584230749346 | 2.27399148412876 | 7.15936926220794 | 6.40265188186281 |
| chr16 | 70248951 | 70248966 | - | 3.01188173062009 | 4.54798296825753 | 3.57968463110397 | 2.13421729395427 |
| chr16 | 70249703 | 70249717 | - | 4.01584230749346 | 2.27399148412876 | 7.15936926220794 | 25.6106075274513 |
| chr16 | 70250240 | 70250276 | - | 10.0396057687336 | 11.3699574206438 | 10.7390538933119 | 7.46976052883995 |
| chr16 | 70252321 | 70252503 | - | 168.665376914725 | 270.604986611323 | 178.984231555198 | 198.482208337747 |
| chr16 | 70252526 | 70252528 | - | 23.0910932680874 | 21.6029190992233 | 4.77291284147196 | 9.60397782279422 |
| chr16 | 70252550 | 70252894 | - | 144.570323069764 | 242.180093059713 | 130.061874930111 | 168.603166222387 |
| chr16 | 70333753 | 70333764 | + | 4.01584230749346 | 5.68497871032191 | 3.57968463110397 | 7.46976052883995 |
| chr16 | 70333784 | 70333798 | + | 5.01980288436682 | 2.27399148412876 | 1.19322821036799 | 3.20132594093141 |
| chr16 | 70372159 | 70372166 | + | 4.01584230749346 | 2.27399148412876 | 7.15936926220794 | 6.40265188186281 |
| chr16 | 70372187 | 70372197 | + | 1.00396057687336 | 5.68497871032191 | 1.19322821036799 | 1.06710864697714 |
| chr16 | 70372254 | 70372306 | + | 9.03564519186028 | 1.13699574206438 | 7.15936926220794 | 10.6710864697714 |
| chr16 | 70372485 | 70372497 | + | 11.043566345607 | 10.2329616785794 | 8.35259747257593 | 9.60397782279422 |
| chr16 | 70372684 | 70372691 | + | 6.02376346124019 | 2.27399148412876 | 1.19322821036799 | 4.26843458790854 |
| chr16 | 70372816 | 70372894 | + | 7.02772403811355 | 7.95897019445067 | 17.8984231555198 | 35.2145853502455 |
| chr16 | 70373003 | 70373008 | + | 8.03168461498692 | 3.41098722619315 | 7.15936926220794 | 6.40265188186281 |
| chr16 | 70373140 | 70373147 | + | 18.0712903837206 | 15.9179403889013 | 5.96614105183995 | 14.9395210576799 |
| chr16 | 70373207 | 70373315 | + | 26.1029749987075 | 25.0139063254164 | 10.7390538933119 | 7.46976052883995 |
| chr16 | 70379983 | 70379990 | - | 1.00396057687336 | 1.13699574206438 | 8.35259747257593 | 8.53686917581709 |
| chr16 | 70380025 | 70380044 | - | 2.00792115374673 | 1.13699574206438 | 2.38645642073598 | 6.40265188186281 |
| chr16 | 70380502 | 70380528 | - | 1.00396057687336 | 2.27399148412876 | 4.77291284147196 | 4.26843458790854 |
| chr16 | 70380726 | 70380730 | - | 5.01980288436682 | 1.13699574206438 | 34.6036181006717 | 4.26843458790854 |
| chr16 | 70381060 | 70381061 | - | 1.00396057687336 | 1.13699574206438 | 3.57968463110397 | 1.06710864697714 |
| chr16 | 70480592 | 70480657 | - | 18.0712903837206 | 31.8358807778027 | 10.7390538933119 | 33.0803680562912 |
| chr16 | 70572054 | 70572069 | + | 8.03168461498692 | 3.41098722619315 | 1.19322821036799 | 8.53686917581709 |
| chr16 | 70572095 | 70572136 | + | 5.01980288436682 | 7.95897019445067 | 4.77291284147196 | 4.26843458790854 |
| chr16 | 70572193 | 70572278 | + | 62.2455557661486 | 63.6717615556054 | 22.6713359969918 | 53.3554323488568 |
| chr16 | 70574129 | 70574144 | + | 15.0594086531005 | 9.09596593651506 | 10.7390538933119 | 8.53686917581709 |
| chr16 | 70576069 | 70576070 | + | 12.0475269224804 | 5.68497871032191 | 7.15936926220794 | 4.26843458790854 |
| chr16 | 70661239 | 70661261 | - | 1.00396057687336 | 1.13699574206438 | 2.38645642073598 | 3.20132594093141 |
| chr16 | 70661292 | 70661521 | - | 58.2297134586551 | 36.3838637460602 | 14.3187385244159 | 26.6777161744284 |
| chr16 | 70661685 | 70661704 | - | 13.0514874993537 | 11.3699574206438 | 3.57968463110397 | 2.13421729395427 |
| chr16 | 70661804 | 70661901 | - | 8.03168461498692 | 12.5069531627082 | 2.38645642073598 | 10.6710864697714 |
| chr16 | 70662057 | 70662288 | - | 11.043566345607 | 6.82197445238629 | 2.38645642073598 | 5.33554323488568 |
| chr16 | 70662562 | 70662611 | - | 2.00792115374673 | 1.13699574206438 | 1.19322821036799 | 2.13421729395427 |
| chr16 | 70662633 | 70662723 | - | 16.0633692299738 | 22.7399148412876 | 11.9322821036799 | 18.1408469986113 |
| chr16 | 70662745 | 70662979 | - | 30.1188173062009 | 28.4248935516096 | 21.4781077866238 | 10.6710864697714 |
| chr16 | 71447806 | 71447808 | - | 1.00396057687336 | 1.13699574206438 | 1.19322821036799 | 1.06710864697714 |
| chr16 | 71462185 | 71462255 | - | 225.891129796507 | 230.81013563907 | 429.562155732476 | 337.206332444775 |
| chr16 | 71462270 | 71462294 | + | 15.0594086531005 | 21.6029190992233 | 8.35259747257593 | 6.40265188186281 |
| chr16 | 71646532 | 71646538 | - | 1.00396057687336 | 2.27399148412876 | 2.38645642073598 | 1.06710864697714 |
| chr16 | 71646675 | 71646677 | - | 1.00396057687336 | 1.13699574206438 | 1.19322821036799 | 1.06710864697714 |
| chr16 | 71646786 | 71646804 | - | 6.02376346124019 | 3.41098722619315 | 1.19322821036799 | 3.20132594093141 |
| chr16 | 71648627 | 71648647 | - | 101.40001826421 | 343.372714103443 | 3.57968463110397 | 4.26843458790854 |
| chr16 | 71729110 | 71729147 | - | 68.2693192273888 | 46.6168254246397 | 89.4921157775992 | 66.1607361125824 |
| chr16 | 71729177 | 71729221 | - | 44.174265382428 | 27.2878978095452 | 70.4004644117114 | 36.2816939972226 |
| chr16 | 71729243 | 71729251 | - | 5.01980288436682 | 4.54798296825753 | 1.19322821036799 | 1.06710864697714 |
| chr16 | 71729289 | 71729328 | - | 12.0475269224804 | 15.9179403889013 | 23.8645642073598 | 16.006629704657 |
| chr16 | 71729372 | 71729375 | - | 4.01584230749346 | 2.27399148412876 | 2.38645642073598 | 1.06710864697714 |
| chr16 | 71729402 | 71729416 | - | 15.0594086531005 | 9.09596593651506 | 4.77291284147196 | 12.8053037637256 |
| chr16 | 71729464 | 71729492 | - | 17.0673298068472 | 3.41098722619315 | 17.8984231555198 | 23.476390233497 |
| chr16 | 71729534 | 71729570 | - | 5.01980288436682 | 2.27399148412876 | 5.96614105183995 | 21.3421729395427 |
| chr16 | 71729753 | 71729795 | - | 6.02376346124019 | 5.68497871032191 | 15.5119667347839 | 8.53686917581709 |
| chr16 | 71729856 | 71729885 | - | 9.03564519186028 | 5.68497871032191 | 23.8645642073598 | 10.6710864697714 |
| chr16 | 71729924 | 71729953 | - | 2.00792115374673 | 1.13699574206438 | 8.35259747257593 | 4.26843458790854 |
| chr16 | 71730061 | 71730067 | - | 3.01188173062009 | 4.54798296825753 | 5.96614105183995 | 10.6710864697714 |
| chr16 | 71730088 | 71730106 | - | 43.1703048055547 | 31.8358807778027 | 17.8984231555198 | 22.4092815865198 |
| chr16 | 71730451 | 71730463 | - | 3.01188173062009 | 1.13699574206438 | 3.57968463110397 | 3.20132594093141 |
| chr16 | 71730492 | 71730493 | - | 3.01188173062009 | 1.13699574206438 | 2.38645642073598 | 1.06710864697714 |
| chr16 | 71731002 | 71731029 | - | 1.00396057687336 | 3.41098722619315 | 4.77291284147196 | 2.13421729395427 |
| chr16 | 71731304 | 71731328 | - | 4.01584230749346 | 1.13699574206438 | 2.38645642073598 | 2.13421729395427 |
| chr16 | 71731787 | 71731817 | - | 4.01584230749346 | 2.27399148412876 | 4.77291284147196 | 2.13421729395427 |
| chr16 | 71855705 | 71855717 | + | 1.00396057687336 | 3.41098722619315 | 3.57968463110397 | 2.13421729395427 |
| chr16 | 71856846 | 71856871 | + | 3.01188173062009 | 4.54798296825753 | 1.19322821036799 | 1.06710864697714 |
| chr16 | 71857057 | 71857077 | + | 2.00792115374673 | 2.27399148412876 | 2.38645642073598 | 1.06710864697714 |
| chr16 | 71928514 | 71928603 | + | 2.00792115374673 | 2.27399148412876 | 1.19322821036799 | 4.26843458790854 |
| chr16 | 71928659 | 71928771 | + | 1.00396057687336 | 1.13699574206438 | 4.77291284147196 | 2.13421729395427 |
| chr16 | 71928862 | 71928931 | + | 21.0831721143407 | 37.5208594881246 | 21.4781077866238 | 36.2816939972226 |
| chr16 | 72024341 | 72024344 | + | 10.0396057687336 | 2.27399148412876 | 2.38645642073598 | 6.40265188186281 |
| chr16 | 72084930 | 72084931 | + | 41.1623836518079 | 81.8636934286355 | 23.8645642073598 | 19.2079556455884 |
| chr16 | 72085206 | 72085214 | - | 17.0673298068472 | 22.7399148412876 | 8.35259747257593 | 11.7381951167485 |
| chr16 | 72085741 | 72085745 | - | 4.01584230749346 | 4.54798296825753 | 2.38645642073598 | 3.20132594093141 |
| chr16 | 72086028 | 72086039 | - | 8.03168461498692 | 23.876910583352 | 14.3187385244159 | 12.8053037637256 |
| chr16 | 72112555 | 72112572 | + | 7.02772403811355 | 2.27399148412876 | 2.38645642073598 | 5.33554323488568 |
| chr16 | 72112695 | 72112745 | + | 11.043566345607 | 12.5069531627082 | 3.57968463110397 | 12.8053037637256 |
| chr16 | 72112769 | 72112827 | + | 8.03168461498692 | 4.54798296825753 | 2.38645642073598 | 8.53686917581709 |
| chr16 | 72665125 | 72665190 | + | 62.2455557661486 | 120.521548658824 | 143.187385244159 | 166.468948928433 |
| chr16 | 72790219 | 72790226 | + | 6.02376346124019 | 7.95897019445067 | 13.1255103140479 | 13.8724124107028 |
| chr16 | 74305465 | 74305561 | + | 75.2970432655023 | 60.2607743294122 | 124.095733878271 | 98.1739955218965 |
| chr16 | 74305589 | 74305743 | + | 209.827760566533 | 208.070220797782 | 292.340911540157 | 383.092004264792 |
| chr16 | 74305787 | 74305893 | + | 88.3485307648561 | 68.2197445238629 | 66.8207797806074 | 82.1673658172394 |
| chr16 | 74306044 | 74306047 | + | 1.00396057687336 | 6.82197445238629 | 1.19322821036799 | 3.20132594093141 |
| chr16 | 74306070 | 74306081 | + | 4.01584230749346 | 1.13699574206438 | 2.38645642073598 | 3.20132594093141 |
| chr16 | 74306199 | 74306231 | + | 33.130699036821 | 47.753821166704 | 21.4781077866238 | 27.7448248214055 |
| chr16 | 74447464 | 74447474 | - | 2.00792115374673 | 1.13699574206438 | 2.38645642073598 | 2.13421729395427 |
| chr16 | 74447736 | 74447798 | - | 3.01188173062009 | 3.41098722619315 | 2.38645642073598 | 3.20132594093141 |
| chr16 | 74451962 | 74452040 | - | 20.0792115374673 | 35.2468680039958 | 9.54582568294392 | 9.60397782279422 |
| chr16 | 74452157 | 74452246 | - | 2.00792115374673 | 3.41098722619315 | 2.38645642073598 | 3.20132594093141 |
| chr16 | 74452372 | 74452449 | - | 6.02376346124019 | 2.27399148412876 | 14.3187385244159 | 7.46976052883995 |
| chr16 | 74452478 | 74452511 | - | 2.00792115374673 | 3.41098722619315 | 2.38645642073598 | 3.20132594093141 |
| chr16 | 74452668 | 74452703 | - | 2.00792115374673 | 1.13699574206438 | 3.57968463110397 | 4.26843458790854 |
| chr16 | 74452728 | 74452798 | - | 3.01188173062009 | 6.82197445238629 | 4.77291284147196 | 6.40265188186281 |
| chr16 | 74452828 | 74452851 | - | 1.00396057687336 | 1.13699574206438 | 2.38645642073598 | 2.13421729395427 |
| chr16 | 74452909 | 74453008 | - | 7.02772403811355 | 9.09596593651506 | 5.96614105183995 | 6.40265188186281 |
| chr16 | 74453030 | 74453087 | - | 2.00792115374673 | 4.54798296825753 | 5.96614105183995 | 2.13421729395427 |
| chr16 | 74453114 | 74453258 | - | 21.0831721143407 | 22.7399148412876 | 29.8307052591997 | 16.006629704657 |
| chr16 | 74621459 | 74621484 | - | 11.043566345607 | 12.5069531627082 | 10.7390538933119 | 12.8053037637256 |
| chr16 | 74621618 | 74621634 | - | 5.01980288436682 | 4.54798296825753 | 2.38645642073598 | 3.20132594093141 |
| chr16 | 74622562 | 74622566 | - | 13.0514874993537 | 15.9179403889013 | 1.19322821036799 | 1.06710864697714 |
| chr16 | 74623167 | 74623178 | - | 2.00792115374673 | 5.68497871032191 | 3.57968463110397 | 6.40265188186281 |
| chr16 | 74873612 | 74873638 | - | 2.00792115374673 | 2.27399148412876 | 2.38645642073598 | 4.26843458790854 |
| chr16 | 74873674 | 74873830 | - | 3.01188173062009 | 1.13699574206438 | 5.96614105183995 | 6.40265188186281 |
| chr16 | 75107777 | 75107826 | + | 5.01980288436682 | 2.27399148412876 | 3.57968463110397 | 6.40265188186281 |
| chr16 | 75107940 | 75107948 | + | 2.00792115374673 | 10.2329616785794 | 3.57968463110397 | 11.7381951167485 |
| chr16 | 75107973 | 75107986 | + | 4.01584230749346 | 6.82197445238629 | 1.19322821036799 | 10.6710864697714 |
| chr16 | 75108648 | 75108653 | + | 2.00792115374673 | 1.13699574206438 | 1.19322821036799 | 1.06710864697714 |
| chr16 | 75172058 | 75172061 | + | 3.01188173062009 | 2.27399148412876 | 2.38645642073598 | 1.06710864697714 |
| chr16 | 75229065 | 75229200 | - | 15.0594086531005 | 14.780944646837 | 4.77291284147196 | 2.13421729395427 |
| chr16 | 75293818 | 75293945 | - | 3.01188173062009 | 6.82197445238629 | 5.96614105183995 | 1.06710864697714 |
| chr16 | 75443395 | 75443452 | - | 56.2217923049084 | 85.2746806548286 | 42.9562155732476 | 37.3488026441998 |
| chr16 | 75443526 | 75443530 | - | 1.00396057687336 | 1.13699574206438 | 1.19322821036799 | 3.20132594093141 |
| chr16 | 75443624 | 75443659 | - | 12.0475269224804 | 17.0549361309657 | 7.15936926220794 | 12.8053037637256 |
| chr16 | 75443775 | 75443860 | - | 3.01188173062009 | 4.54798296825753 | 3.57968463110397 | 3.20132594093141 |
| chr16 | 75446996 | 75447005 | - | 3.01188173062009 | 10.2329616785794 | 2.38645642073598 | 4.26843458790854 |
| chr16 | 75447161 | 75447168 | - | 6.02376346124019 | 5.68497871032191 | 4.77291284147196 | 3.20132594093141 |
| chr16 | 75447258 | 75447262 | - | 4.01584230749346 | 4.54798296825753 | 1.19322821036799 | 4.26843458790854 |
| chr16 | 75447432 | 75447462 | - | 2.00792115374673 | 5.68497871032191 | 7.15936926220794 | 4.26843458790854 |
| chr16 | 75577345 | 75577387 | + | 6.02376346124019 | 3.41098722619315 | 8.35259747257593 | 13.8724124107028 |
| chr16 | 75577446 | 75577552 | + | 6.02376346124019 | 4.54798296825753 | 9.54582568294392 | 8.53686917581709 |
| chr16 | 75577651 | 75577685 | + | 7.02772403811355 | 20.4659233571589 | 5.96614105183995 | 26.6777161744284 |
| chr16 | 75577758 | 75577855 | + | 10.0396057687336 | 21.6029190992233 | 16.7051949451519 | 22.4092815865198 |
| chr16 | 75627756 | 75627881 | - | 22.087132691214 | 23.876910583352 | 42.9562155732476 | 24.5434988804741 |
| chr16 | 75656621 | 75656670 | + | 11.043566345607 | 15.9179403889013 | 32.2171616799357 | 20.2750642925656 |
| chr16 | 75656765 | 75656784 | + | 1.00396057687336 | 1.13699574206438 | 2.38645642073598 | 2.13421729395427 |
| chr16 | 75657163 | 75657188 | + | 8.03168461498692 | 4.54798296825753 | 7.15936926220794 | 10.6710864697714 |
| chr16 | 75657204 | 75657232 | + | 6.02376346124019 | 5.68497871032191 | 8.35259747257593 | 7.46976052883995 |
| chr16 | 75657266 | 75657409 | + | 8.03168461498692 | 10.2329616785794 | 16.7051949451519 | 12.8053037637256 |
| chr16 | 77199255 | 77199289 | + | 2.00792115374673 | 1.13699574206438 | 1.19322821036799 | 12.8053037637256 |
| chr16 | 77199352 | 77199393 | + | 3.01188173062009 | 3.41098722619315 | 1.19322821036799 | 7.46976052883995 |
| chr16 | 77199565 | 77199571 | + | 2.00792115374673 | 7.95897019445067 | 3.57968463110397 | 6.40265188186281 |
| chr16 | 77201922 | 77201926 | + | 10.0396057687336 | 11.3699574206438 | 2.38645642073598 | 3.20132594093141 |
| chr16 | 77202238 | 77202258 | + | 5.01980288436682 | 4.54798296825753 | 2.38645642073598 | 2.13421729395427 |
| chr16 | 79212481 | 79212499 | + | 13.0514874993537 | 14.780944646837 | 13.1255103140479 | 7.46976052883995 |
| chr16 | 79212532 | 79212551 | + | 25.0990144218341 | 18.1919318730301 | 54.8884976769275 | 37.3488026441998 |
| chr16 | 80975823 | 80975858 | - | 23.0910932680874 | 57.9867828452835 | 7.15936926220794 | 4.26843458790854 |
| chr16 | 80975883 | 80975925 | - | 72.2851615348822 | 143.261463500112 | 26.2510206280958 | 24.5434988804741 |
| chr16 | 80975961 | 80975963 | - | 2.00792115374673 | 2.27399148412876 | 1.19322821036799 | 2.13421729395427 |
| chr16 | 81022849 | 81022870 | + | 15.0594086531005 | 17.0549361309657 | 9.54582568294392 | 3.20132594093141 |
| chr16 | 81378341 | 81378365 | + | 12.0475269224804 | 1.13699574206438 | 3.57968463110397 | 5.33554323488568 |
| chr16 | 81383774 | 81383776 | + | 3.01188173062009 | 2.27399148412876 | 1.19322821036799 | 1.06710864697714 |
| chr16 | 81383865 | 81383910 | + | 4.01584230749346 | 4.54798296825753 | 10.7390538933119 | 2.13421729395427 |
| chr16 | 81385062 | 81385065 | + | 22.087132691214 | 29.5618892936739 | 20.2848795762558 | 13.8724124107028 |
| chr16 | 81386919 | 81386931 | + | 2.00792115374673 | 1.13699574206438 | 1.19322821036799 | 1.06710864697714 |
| chr16 | 81390533 | 81390534 | + | 1.00396057687336 | 4.54798296825753 | 1.19322821036799 | 6.40265188186281 |
| chr16 | 81709947 | 81709960 | + | 6.02376346124019 | 2.27399148412876 | 1.19322821036799 | 1.06710864697714 |
| chr16 | 81710081 | 81710119 | + | 31.1227778830743 | 22.7399148412876 | 66.8207797806074 | 51.2212150549025 |
| chr16 | 81710448 | 81710490 | + | 13.0514874993537 | 15.9179403889013 | 32.2171616799357 | 25.6106075274513 |
| chr16 | 81711150 | 81711363 | + | 26.1029749987075 | 23.876910583352 | 82.3327465153913 | 124.851711696325 |
| chr16 | 81711381 | 81711457 | + | 20.0792115374673 | 13.6439489047726 | 50.1155848354556 | 57.6238669367653 |
| chr16 | 81711524 | 81711527 | + | 1.00396057687336 | 1.13699574206438 | 7.15936926220794 | 4.26843458790854 |
| chr16 | 81711552 | 81711565 | + | 5.01980288436682 | 6.82197445238629 | 11.9322821036799 | 20.2750642925656 |
| chr16 | 81711616 | 81711741 | + | 3.01188173062009 | 4.54798296825753 | 13.1255103140479 | 10.6710864697714 |
| chr16 | 82148199 | 82148297 | - | 11.043566345607 | 14.780944646837 | 33.4103898903037 | 8.53686917581709 |
| chr16 | 82148486 | 82148491 | - | 8.03168461498692 | 7.95897019445067 | 10.7390538933119 | 9.60397782279422 |
| chr16 | 82148529 | 82148576 | - | 6.02376346124019 | 11.3699574206438 | 2.38645642073598 | 5.33554323488568 |
| chr16 | 82148614 | 82148727 | - | 4.01584230749346 | 4.54798296825753 | 2.38645642073598 | 10.6710864697714 |
| chr16 | 83809410 | 83809437 | + | 11.043566345607 | 10.2329616785794 | 14.3187385244159 | 5.33554323488568 |
| chr16 | 83811418 | 83811699 | + | 75.2970432655023 | 120.521548658824 | 97.8447132501751 | 116.314842520508 |
| chr16 | 83811721 | 83811763 | + | 5.01980288436682 | 21.6029190992233 | 29.8307052591997 | 14.9395210576799 |
| chr16 | 83812819 | 83812952 | + | 29.1148567293276 | 27.2878978095452 | 17.8984231555198 | 24.5434988804741 |
| chr16 | 84053807 | 84053849 | - | 8.03168461498692 | 14.780944646837 | 4.77291284147196 | 2.13421729395427 |
| chr16 | 84053962 | 84053967 | - | 5.01980288436682 | 4.54798296825753 | 4.77291284147196 | 2.13421729395427 |
| chr16 | 84053999 | 84054079 | - | 17.0673298068472 | 20.4659233571589 | 11.9322821036799 | 20.2750642925656 |
| chr16 | 84054136 | 84054189 | - | 10.0396057687336 | 25.0139063254164 | 2.38645642073598 | 11.7381951167485 |
| chr16 | 84117022 | 84117100 | + | 72.2851615348822 | 61.3977700714766 | 97.8447132501751 | 96.0397782279422 |
| chr16 | 84177886 | 84177889 | - | 2.00792115374673 | 2.27399148412876 | 2.38645642073598 | 2.13421729395427 |
| chr16 | 84178544 | 84178580 | - | 15.0594086531005 | 4.54798296825753 | 8.35259747257593 | 6.40265188186281 |
| chr16 | 84178778 | 84178786 | - | 2.00792115374673 | 3.41098722619315 | 7.15936926220794 | 14.9395210576799 |
| chr16 | 84565624 | 84565689 | - | 85.336649034236 | 110.288586980245 | 35.7968463110397 | 48.0198891139711 |
| chr16 | 84565725 | 84565765 | - | 32.1267384599477 | 22.7399148412876 | 19.0916513658878 | 26.6777161744284 |
| chr16 | 84565790 | 84565796 | - | 4.01584230749346 | 3.41098722619315 | 4.77291284147196 | 4.26843458790854 |
| chr16 | 84565833 | 84565868 | - | 8.03168461498692 | 10.2329616785794 | 5.96614105183995 | 13.8724124107028 |
| chr16 | 84565895 | 84565898 | - | 8.03168461498692 | 3.41098722619315 | 3.57968463110397 | 2.13421729395427 |
| chr16 | 84779078 | 84779116 | + | 9.03564519186028 | 10.2329616785794 | 13.1255103140479 | 18.1408469986113 |
| chr16 | 84779218 | 84779260 | + | 68.2693192273888 | 53.438799877026 | 64.4343233598714 | 42.6843458790854 |
| chr16 | 84779282 | 84779311 | + | 4.01584230749346 | 9.09596593651506 | 8.35259747257593 | 5.33554323488568 |
| chr16 | 84909224 | 84909225 | + | 25.0990144218341 | 21.6029190992233 | 5.96614105183995 | 8.53686917581709 |
| chr16 | 84974638 | 84974752 | - | 17.0673298068472 | 10.2329616785794 | 4.77291284147196 | 13.8724124107028 |
| chr16 | 84974923 | 84975010 | - | 10.0396057687336 | 7.95897019445067 | 22.6713359969918 | 7.46976052883995 |
| chr16 | 84975065 | 84975087 | - | 9.03564519186028 | 3.41098722619315 | 11.9322821036799 | 9.60397782279422 |
| chr16 | 84975127 | 84975148 | - | 5.01980288436682 | 2.27399148412876 | 9.54582568294392 | 6.40265188186281 |
| chr16 | 84975187 | 84975262 | - | 4.01584230749346 | 9.09596593651506 | 4.77291284147196 | 9.60397782279422 |
| chr16 | 84975282 | 84975364 | - | 2.00792115374673 | 6.82197445238629 | 16.7051949451519 | 7.46976052883995 |
| chr16 | 84975432 | 84975445 | - | 2.00792115374673 | 1.13699574206438 | 3.57968463110397 | 4.26843458790854 |
| chr16 | 84975468 | 84975487 | - | 19.0752509605939 | 15.9179403889013 | 20.2848795762558 | 19.2079556455884 |
| chr16 | 84975524 | 84975629 | - | 2.00792115374673 | 7.95897019445067 | 11.9322821036799 | 12.8053037637256 |
| chr16 | 84975698 | 84975791 | - | 4.01584230749346 | 4.54798296825753 | 4.77291284147196 | 3.20132594093141 |
| chr16 | 84975906 | 84975913 | - | 2.00792115374673 | 3.41098722619315 | 2.38645642073598 | 4.26843458790854 |
| chr16 | 84976176 | 84976216 | - | 1.00396057687336 | 1.13699574206438 | 7.15936926220794 | 5.33554323488568 |
| chr16 | 84976332 | 84976392 | - | 5.01980288436682 | 2.27399148412876 | 7.15936926220794 | 2.13421729395427 |
| chr16 | 85672506 | 85672561 | + | 3.01188173062009 | 3.41098722619315 | 11.9322821036799 | 13.8724124107028 |
| chr16 | 85672617 | 85672668 | + | 2.00792115374673 | 4.54798296825753 | 10.7390538933119 | 9.60397782279422 |
| chr16 | 85672881 | 85672890 | + | 3.01188173062009 | 1.13699574206438 | 5.96614105183995 | 5.33554323488568 |
| chr16 | 85672962 | 85672969 | + | 7.02772403811355 | 10.2329616785794 | 27.4442488384638 | 50.1541064079254 |
| chr16 | 85672991 | 85673040 | + | 6.02376346124019 | 7.95897019445067 | 22.6713359969918 | 16.006629704657 |
| chr16 | 85673096 | 85673109 | + | 3.01188173062009 | 1.13699574206438 | 2.38645642073598 | 6.40265188186281 |
| chr16 | 85673173 | 85673347 | + | 44.174265382428 | 51.1648083928972 | 189.72328544851 | 354.280070796409 |
| chr16 | 85673375 | 85673477 | + | 28.1108961524542 | 35.2468680039958 | 136.028015981951 | 110.979299285622 |
| chr16 | 85673522 | 85673543 | + | 2.00792115374673 | 1.13699574206438 | 5.96614105183995 | 7.46976052883995 |
| chr16 | 85676005 | 85676085 | + | 11.043566345607 | 17.0549361309657 | 32.2171616799357 | 60.8251928776967 |
| chr16 | 85677716 | 85677748 | - | 34.1346596136944 | 32.9728765198671 | 17.8984231555198 | 35.2145853502455 |
| chr16 | 85677794 | 85677796 | - | 6.02376346124019 | 1.13699574206438 | 8.35259747257593 | 9.60397782279422 |
| chr16 | 85678025 | 85678028 | - | 4.01584230749346 | 5.68497871032191 | 1.19322821036799 | 5.33554323488568 |
| chr16 | 85678094 | 85678099 | - | 1.00396057687336 | 1.13699574206438 | 1.19322821036799 | 3.20132594093141 |
| chr16 | 85678154 | 85678201 | - | 11.043566345607 | 11.3699574206438 | 5.96614105183995 | 7.46976052883995 |
| chr16 | 85779429 | 85779452 | - | 3.01188173062009 | 1.13699574206438 | 2.38645642073598 | 4.26843458790854 |
| chr16 | 85779474 | 85779528 | - | 14.0554480762271 | 11.3699574206438 | 7.15936926220794 | 6.40265188186281 |
| chr16 | 85779554 | 85779624 | - | 47.1861471130481 | 37.5208594881246 | 10.7390538933119 | 29.8790421153598 |
| chr16 | 85806736 | 85806967 | + | 144.570323069764 | 114.836569948503 | 70.4004644117114 | 179.274252692159 |
| chr16 | 87317491 | 87317526 | + | 18.0712903837206 | 19.3289276150945 | 2.38645642073598 | 5.33554323488568 |
| chr16 | 87330990 | 87331039 | - | 2.00792115374673 | 3.41098722619315 | 2.38645642073598 | 1.06710864697714 |
| chr16 | 87403508 | 87403583 | + | 8.03168461498692 | 6.82197445238629 | 15.5119667347839 | 19.2079556455884 |
| chr16 | 87403713 | 87403720 | + | 4.01584230749346 | 9.09596593651506 | 1.19322821036799 | 9.60397782279422 |
| chr16 | 87404674 | 87404720 | + | 21.0831721143407 | 31.8358807778027 | 47.7291284147196 | 36.2816939972226 |
| chr16 | 87407650 | 87407735 | - | 18.0712903837206 | 7.95897019445067 | 2.38645642073598 | 4.26843458790854 |
| chr16 | 87409405 | 87409408 | - | 89.3524913417294 | 35.2468680039958 | 13.1255103140479 | 45.8856718200168 |
| chr16 | 87698175 | 87698207 | - | 3.01188173062009 | 6.82197445238629 | 3.57968463110397 | 4.26843458790854 |
| chr16 | 87707838 | 87707895 | - | 24.0950538449607 | 30.6988850357383 | 20.2848795762558 | 17.0737383516342 |
| chr16 | 87708349 | 87708350 | - | 1.00396057687336 | 1.13699574206438 | 1.19322821036799 | 1.06710864697714 |
| chr16 | 87830041 | 87830203 | - | 241.954499026481 | 504.826109476586 | 45.3426719939836 | 57.6238669367653 |
| chr16 | 87830241 | 87830371 | - | 29.1148567293276 | 23.876910583352 | 2.38645642073598 | 7.46976052883995 |
| chr16 | 87830401 | 87830438 | - | 8.03168461498692 | 11.3699574206438 | 1.19322821036799 | 1.06710864697714 |
| chr16 | 87830464 | 87830529 | - | 6.02376346124019 | 20.4659233571589 | 1.19322821036799 | 5.33554323488568 |
| chr16 | 87830552 | 87830798 | - | 292.152527870149 | 247.865071770035 | 69.2072362013434 | 70.429170700491 |
| chr16 | 87830836 | 87830919 | - | 84.3326884573626 | 105.740604011988 | 34.6036181006717 | 28.8119334683827 |
| chr16 | 87830941 | 87831486 | - | 330.303029791337 | 470.716237214654 | 161.085808399679 | 71.4962793474681 |
| chr16 | 87831534 | 87831555 | - | 6.02376346124019 | 17.0549361309657 | 1.19322821036799 | 1.06710864697714 |
| chr16 | 87831585 | 87831693 | - | 591.332779778412 | 303.57786313119 | 83.5259747257593 | 27.7448248214055 |
| chr16 | 87831733 | 87831818 | - | 47.1861471130481 | 76.1787147183136 | 1.19322821036799 | 6.40265188186281 |
| chr16 | 87832722 | 87832784 | - | 13.0514874993537 | 7.95897019445067 | 3.57968463110397 | 4.26843458790854 |
| chr16 | 87832917 | 87833001 | - | 45.1782259593014 | 95.5076423334081 | 3.57968463110397 | 3.20132594093141 |
| chr16 | 88535110 | 88535136 | + | 8.03168461498692 | 9.09596593651506 | 5.96614105183995 | 21.3421729395427 |
| chr16 | 88631275 | 88631339 | + | 7.02772403811355 | 10.2329616785794 | 15.5119667347839 | 14.9395210576799 |
| chr16 | 88631881 | 88631916 | + | 31.1227778830743 | 38.657855230189 | 19.0916513658878 | 68.2949534065367 |
| chr16 | 88643322 | 88643420 | - | 6.02376346124019 | 4.54798296825753 | 10.7390538933119 | 9.60397782279422 |
| chr16 | 88651960 | 88652037 | - | 8.03168461498692 | 5.68497871032191 | 16.7051949451519 | 4.26843458790854 |
| chr16 | 88652094 | 88652256 | - | 8.03168461498692 | 6.82197445238629 | 175.404546924094 | 199.549316984724 |
| chr16 | 88652278 | 88652513 | - | 16.0633692299738 | 21.6029190992233 | 318.591932168253 | 242.23366286381 |
| chr16 | 88663310 | 88663376 | + | 32.1267384599477 | 36.3838637460602 | 68.0140079909754 | 83.2344744642166 |
| chr16 | 88696582 | 88696598 | - | 3.01188173062009 | 5.68497871032191 | 11.9322821036799 | 29.8790421153598 |
| chr16 | 88696930 | 88696943 | - | 3.01188173062009 | 3.41098722619315 | 17.8984231555198 | 26.6777161744284 |
| chr16 | 88696984 | 88697011 | - | 2.00792115374673 | 4.54798296825753 | 10.7390538933119 | 6.40265188186281 |
| chr16 | 88697042 | 88697045 | - | 1.00396057687336 | 1.13699574206438 | 7.15936926220794 | 2.13421729395427 |
| chr16 | 88697077 | 88697083 | - | 2.00792115374673 | 1.13699574206438 | 7.15936926220794 | 6.40265188186281 |
| chr16 | 88697131 | 88697146 | - | 2.00792115374673 | 1.13699574206438 | 5.96614105183995 | 2.13421729395427 |
| chr16 | 88697298 | 88697322 | - | 1.00396057687336 | 2.27399148412876 | 9.54582568294392 | 2.13421729395427 |
| chr16 | 88715307 | 88715349 | + | 8.03168461498692 | 14.780944646837 | 26.2510206280958 | 29.8790421153598 |
| chr16 | 88715398 | 88715420 | - | 5.01980288436682 | 1.13699574206438 | 10.7390538933119 | 13.8724124107028 |
| chr16 | 88715487 | 88715564 | - | 18.0712903837206 | 4.54798296825753 | 28.6374770488317 | 52.2883237018796 |
| chr16 | 88808349 | 88808358 | + | 2.00792115374673 | 3.41098722619315 | 1.19322821036799 | 2.13421729395427 |
| chr16 | 88808464 | 88808481 | + | 5.01980288436682 | 19.3289276150945 | 10.7390538933119 | 26.6777161744284 |
| chr16 | 88809179 | 88809194 | + | 3.01188173062009 | 7.95897019445067 | 5.96614105183995 | 3.20132594093141 |
| chr16 | 88809487 | 88809621 | - | 27.1069355755808 | 37.5208594881246 | 23.8645642073598 | 70.429170700491 |
| chr16 | 88813754 | 88813760 | - | 10.0396057687336 | 12.5069531627082 | 1.19322821036799 | 1.06710864697714 |
| chr16 | 88861056 | 88861080 | + | 6.02376346124019 | 6.82197445238629 | 3.57968463110397 | 11.7381951167485 |
| chr16 | 89154535 | 89154549 | + | 1.00396057687336 | 3.41098722619315 | 1.19322821036799 | 1.06710864697714 |
| chr16 | 89160604 | 89160605 | + | 9.03564519186028 | 14.780944646837 | 9.54582568294392 | 19.2079556455884 |
| chr16 | 89285270 | 89285309 | - | 9.03564519186028 | 13.6439489047726 | 14.3187385244159 | 5.33554323488568 |
| chr16 | 89517900 | 89517916 | + | 2.00792115374673 | 3.41098722619315 | 4.77291284147196 | 2.13421729395427 |
| chr16 | 89557489 | 89557553 | + | 12.0475269224804 | 6.82197445238629 | 4.77291284147196 | 8.53686917581709 |
| chr16 | 89557576 | 89557612 | + | 9.03564519186028 | 5.68497871032191 | 2.38645642073598 | 6.40265188186281 |
| chr16 | 89557632 | 89557733 | + | 18.0712903837206 | 13.6439489047726 | 13.1255103140479 | 18.1408469986113 |
| chr16 | 89561426 | 89561434 | + | 56.2217923049084 | 45.4798296825753 | 83.5259747257593 | 128.053037637256 |
| chr16 | 89563042 | 89563301 | + | 181.716864414079 | 184.19331021443 | 93.0718004087032 | 105.643756050736 |
| chr16 | 89563388 | 89563415 | + | 9.03564519186028 | 4.54798296825753 | 3.57968463110397 | 10.6710864697714 |
| chr16 | 89564185 | 89564225 | + | 7.02772403811355 | 6.82197445238629 | 2.38645642073598 | 1.06710864697714 |
| chr16 | 89596638 | 89596659 | + | 5.01980288436682 | 5.68497871032191 | 11.9322821036799 | 2.13421729395427 |
| chr16 | 89596854 | 89596889 | + | 35.1386201905678 | 12.5069531627082 | 32.2171616799357 | 34.1474767032683 |
| chr16 | 89597017 | 89597020 | + | 10.0396057687336 | 7.95897019445067 | 3.57968463110397 | 3.20132594093141 |
| chr16 | 89597041 | 89597055 | + | 15.0594086531005 | 7.95897019445067 | 3.57968463110397 | 4.26843458790854 |
| chr16 | 89597169 | 89597179 | + | 8.03168461498692 | 3.41098722619315 | 3.57968463110397 | 2.13421729395427 |
| chr16 | 89597204 | 89597220 | + | 11.043566345607 | 28.4248935516096 | 9.54582568294392 | 6.40265188186281 |
| chr16 | 89644464 | 89644499 | - | 6.02376346124019 | 15.9179403889013 | 8.35259747257593 | 11.7381951167485 |
| chr16 | 89644981 | 89644999 | - | 3.01188173062009 | 3.41098722619315 | 1.19322821036799 | 1.06710864697714 |
| chr16 | 89645018 | 89645023 | - | 5.01980288436682 | 1.13699574206438 | 1.19322821036799 | 1.06710864697714 |
| chr16 | 89645049 | 89645096 | - | 2.00792115374673 | 5.68497871032191 | 9.54582568294392 | 5.33554323488568 |
| chr16 | 89645225 | 89645276 | - | 7.02772403811355 | 3.41098722619315 | 8.35259747257593 | 6.40265188186281 |
| chr16 | 89645467 | 89645526 | - | 14.0554480762271 | 5.68497871032191 | 5.96614105183995 | 9.60397782279422 |
| chr16 | 89645717 | 89645728 | - | 2.00792115374673 | 1.13699574206438 | 3.57968463110397 | 7.46976052883995 |
| chr16 | 89645791 | 89645818 | - | 3.01188173062009 | 1.13699574206438 | 2.38645642073598 | 5.33554323488568 |
| chr16 | 89645861 | 89645889 | - | 8.03168461498692 | 2.27399148412876 | 4.77291284147196 | 2.13421729395427 |
| chr16 | 89661109 | 89661112 | + | 5.01980288436682 | 13.6439489047726 | 7.15936926220794 | 1.06710864697714 |
| chr16 | 89661363 | 89661366 | + | 4.01584230749346 | 14.780944646837 | 2.38645642073598 | 1.06710864697714 |
| chr16 | 89670389 | 89670414 | + | 4.01584230749346 | 7.95897019445067 | 3.57968463110397 | 3.20132594093141 |
| chr16 | 89696306 | 89696334 | + | 7.02772403811355 | 10.2329616785794 | 4.77291284147196 | 8.53686917581709 |
| chr16 | 89696386 | 89696398 | - | 3.01188173062009 | 5.68497871032191 | 3.57968463110397 | 4.26843458790854 |
| chr16 | 89707174 | 89707252 | - | 25.0990144218341 | 18.1919318730301 | 13.1255103140479 | 12.8053037637256 |
| chr16 | 89707344 | 89707382 | - | 3.01188173062009 | 4.54798296825753 | 3.57968463110397 | 10.6710864697714 |
| chr16 | 89711841 | 89711895 | + | 217.85944518152 | 213.755199508104 | 204.042023972926 | 304.125964388484 |
| chr16 | 89716802 | 89716805 | + | 3.01188173062009 | 1.13699574206438 | 9.54582568294392 | 10.6710864697714 |
| chr16 | 89716831 | 89716833 | + | 1.00396057687336 | 1.13699574206438 | 2.38645642073598 | 1.06710864697714 |
| chr16 | 89717811 | 89717813 | + | 2.00792115374673 | 6.82197445238629 | 3.57968463110397 | 2.13421729395427 |
| chr16 | 89718020 | 89718022 | + | 5.01980288436682 | 3.41098722619315 | 2.38645642073598 | 1.06710864697714 |
| chr16 | 89738369 | 89738370 | + | 1.00396057687336 | 1.13699574206438 | 1.19322821036799 | 1.06710864697714 |
| chr16 | 89738449 | 89738455 | + | 1.00396057687336 | 1.13699574206438 | 1.19322821036799 | 5.33554323488568 |
| chr16 | 89738762 | 89738790 | + | 4.01584230749346 | 3.41098722619315 | 8.35259747257593 | 6.40265188186281 |
| chr16 | 89739166 | 89739168 | + | 1.00396057687336 | 3.41098722619315 | 5.96614105183995 | 1.06710864697714 |
| chr16 | 89739428 | 89739429 | + | 3.01188173062009 | 3.41098722619315 | 4.77291284147196 | 2.13421729395427 |
| chr16 | 89739731 | 89739774 | + | 4.01584230749346 | 9.09596593651506 | 8.35259747257593 | 5.33554323488568 |
| chr16 | 89798544 | 89798552 | - | 3.01188173062009 | 3.41098722619315 | 3.57968463110397 | 2.13421729395427 |
| chr16 | 89871238 | 89871262 | + | 10.0396057687336 | 21.6029190992233 | 2.38645642073598 | 5.33554323488568 |
| chr16 | 89910657 | 89910661 | + | 1.00396057687336 | 2.27399148412876 | 2.38645642073598 | 4.26843458790854 |
| chr16 | 89911203 | 89911269 | + | 9.03564519186028 | 6.82197445238629 | 2.38645642073598 | 10.6710864697714 |
| chr16 | 89911292 | 89911352 | + | 6.02376346124019 | 7.95897019445067 | 13.1255103140479 | 16.006629704657 |
| chr16 | 89917875 | 89917907 | + | 7.02772403811355 | 5.68497871032191 | 9.54582568294392 | 7.46976052883995 |
| chr16 | 89936003 | 89936060 | + | 2.00792115374673 | 5.68497871032191 | 1.19322821036799 | 1.06710864697714 |
| chr16 | 89959306 | 89959343 | + | 62.2455557661486 | 38.657855230189 | 102.617626091647 | 104.576647403759 |
| chr16 | 89967908 | 89967993 | + | 15.0594086531005 | 21.6029190992233 | 59.6614105183995 | 55.4896496428111 |
| chr16 | 89970501 | 89970530 | - | 2.00792115374673 | 1.13699574206438 | 1.19322821036799 | 5.33554323488568 |
| chr16 | 89971101 | 89971128 | - | 4.01584230749346 | 10.2329616785794 | 4.77291284147196 | 9.60397782279422 |
| chr16 | 89972543 | 89972626 | + | 245.970341333974 | 185.330305956494 | 68.0140079909754 | 160.06629704657 |
| chr16 | 90004959 | 90005011 | - | 10.0396057687336 | 3.41098722619315 | 9.54582568294392 | 22.4092815865198 |
| chr16 | 90005297 | 90005328 | - | 3.01188173062009 | 1.13699574206438 | 2.38645642073598 | 5.33554323488568 |
| chr16 | 90005363 | 90005418 | - | 5.01980288436682 | 3.41098722619315 | 20.2848795762558 | 7.46976052883995 |
| chr16 | 90005577 | 90005581 | - | 2.00792115374673 | 2.27399148412876 | 2.38645642073598 | 1.06710864697714 |
| chr17 | 744655 | 744657 | - | 3.01188173062009 | 6.82197445238629 | 1.19322821036799 | 1.06710864697714 |
| chr17 | 744748 | 744759 | - | 10.0396057687336 | 10.2329616785794 | 4.77291284147196 | 17.0737383516342 |
| chr17 | 759366 | 759380 | - | 2.00792115374673 | 2.27399148412876 | 2.38645642073598 | 1.06710864697714 |
| chr17 | 759552 | 759623 | - | 20.0792115374673 | 11.3699574206438 | 5.96614105183995 | 14.9395210576799 |
| chr17 | 760046 | 760070 | - | 12.0475269224804 | 9.09596593651506 | 11.9322821036799 | 10.6710864697714 |
| chr17 | 760107 | 760131 | - | 2.00792115374673 | 2.27399148412876 | 2.38645642073598 | 6.40265188186281 |
| chr17 | 792385 | 792422 | + | 1.00396057687336 | 5.68497871032191 | 8.35259747257593 | 4.26843458790854 |
| chr17 | 792472 | 792479 | + | 21.0831721143407 | 20.4659233571589 | 14.3187385244159 | 8.53686917581709 |
| chr17 | 1464274 | 1464309 | - | 19.0752509605939 | 5.68497871032191 | 3.57968463110397 | 5.33554323488568 |
| chr17 | 1464413 | 1464422 | - | 7.02772403811355 | 2.27399148412876 | 1.19322821036799 | 18.1408469986113 |
| chr17 | 1464466 | 1464499 | - | 15.0594086531005 | 7.95897019445067 | 7.15936926220794 | 30.9461507623369 |
| chr17 | 1464542 | 1464576 | - | 8.03168461498692 | 10.2329616785794 | 1.19322821036799 | 12.8053037637256 |
| chr17 | 1464724 | 1464765 | - | 17.0673298068472 | 10.2329616785794 | 1.19322821036799 | 1.06710864697714 |
| chr17 | 1465064 | 1465068 | - | 22.087132691214 | 15.9179403889013 | 3.57968463110397 | 4.26843458790854 |
| chr17 | 1465145 | 1465167 | - | 12.0475269224804 | 4.54798296825753 | 1.19322821036799 | 1.06710864697714 |
| chr17 | 1465362 | 1465575 | - | 25.0990144218341 | 27.2878978095452 | 11.9322821036799 | 11.7381951167485 |
| chr17 | 1516885 | 1516957 | + | 55.217831728035 | 84.1376849127643 | 15.5119667347839 | 35.2145853502455 |
| chr17 | 1518039 | 1518062 | + | 9.03564519186028 | 10.2329616785794 | 2.38645642073598 | 2.13421729395427 |
| chr17 | 1518196 | 1518268 | - | 164.649534607232 | 119.38455291676 | 71.5936926220794 | 93.9055609339879 |
| chr17 | 1518346 | 1518361 | - | 4.01584230749346 | 1.13699574206438 | 5.96614105183995 | 2.13421729395427 |
| chr17 | 1518450 | 1518467 | - | 5.01980288436682 | 3.41098722619315 | 3.57968463110397 | 2.13421729395427 |
| chr17 | 1518833 | 1518847 | - | 2.00792115374673 | 2.27399148412876 | 1.19322821036799 | 1.06710864697714 |
| chr17 | 1519030 | 1519041 | - | 12.0475269224804 | 5.68497871032191 | 2.38645642073598 | 4.26843458790854 |
| chr17 | 1519157 | 1519186 | - | 3.01188173062009 | 6.82197445238629 | 3.57968463110397 | 10.6710864697714 |
| chr17 | 1519684 | 1519685 | - | 2.00792115374673 | 1.13699574206438 | 1.19322821036799 | 1.06710864697714 |
| chr17 | 1520323 | 1520354 | - | 1.00396057687336 | 1.13699574206438 | 3.57968463110397 | 3.20132594093141 |
| chr17 | 1897593 | 1897609 | + | 9.03564519186028 | 6.82197445238629 | 1.19322821036799 | 3.20132594093141 |
| chr17 | 1897782 | 1897795 | + | 11.043566345607 | 11.3699574206438 | 3.57968463110397 | 16.006629704657 |
| chr17 | 1897867 | 1897872 | + | 4.01584230749346 | 2.27399148412876 | 1.19322821036799 | 3.20132594093141 |
| chr17 | 1897923 | 1897952 | + | 7.02772403811355 | 13.6439489047726 | 4.77291284147196 | 12.8053037637256 |
| chr17 | 1897990 | 1898053 | + | 5.01980288436682 | 9.09596593651506 | 4.77291284147196 | 4.26843458790854 |
| chr17 | 2042633 | 2042718 | + | 23.0910932680874 | 6.82197445238629 | 10.7390538933119 | 13.8724124107028 |
| chr17 | 2042742 | 2042760 | + | 3.01188173062009 | 5.68497871032191 | 7.15936926220794 | 2.13421729395427 |
| chr17 | 2042795 | 2042846 | + | 5.01980288436682 | 9.09596593651506 | 8.35259747257593 | 14.9395210576799 |
| chr17 | 2042880 | 2042898 | + | 5.01980288436682 | 9.09596593651506 | 7.15936926220794 | 7.46976052883995 |
| chr17 | 2042973 | 2043007 | + | 9.03564519186028 | 6.82197445238629 | 23.8645642073598 | 13.8724124107028 |
| chr17 | 2043063 | 2043078 | + | 1.00396057687336 | 1.13699574206438 | 4.77291284147196 | 4.26843458790854 |
| chr17 | 2043092 | 2043114 | + | 7.02772403811355 | 5.68497871032191 | 2.38645642073598 | 1.06710864697714 |
| chr17 | 2043340 | 2043396 | + | 6.02376346124019 | 15.9179403889013 | 4.77291284147196 | 5.33554323488568 |
| chr17 | 2323590 | 2323639 | - | 16.0633692299738 | 19.3289276150945 | 9.54582568294392 | 7.46976052883995 |
| chr17 | 2324123 | 2324160 | - | 14.0554480762271 | 5.68497871032191 | 3.57968463110397 | 4.26843458790854 |
| chr17 | 2380935 | 2381011 | + | 9.03564519186028 | 14.780944646837 | 9.54582568294392 | 5.33554323488568 |
| chr17 | 2384340 | 2384414 | - | 130.514874993537 | 46.6168254246397 | 42.9562155732476 | 56.5567582897882 |
| chr17 | 2384829 | 2384847 | - | 12.0475269224804 | 9.09596593651506 | 10.7390538933119 | 2.13421729395427 |
| chr17 | 2416341 | 2416402 | - | 4.01584230749346 | 15.9179403889013 | 2.38645642073598 | 20.2750642925656 |
| chr17 | 2417104 | 2417105 | - | 10.0396057687336 | 7.95897019445067 | 11.9322821036799 | 1.06710864697714 |
| chr17 | 2418746 | 2418765 | - | 5.01980288436682 | 1.13699574206438 | 4.77291284147196 | 2.13421729395427 |
| chr17 | 2681840 | 2681871 | + | 6.02376346124019 | 7.95897019445067 | 2.38645642073598 | 1.06710864697714 |
| chr17 | 2682750 | 2682774 | + | 9.03564519186028 | 9.09596593651506 | 2.38645642073598 | 2.13421729395427 |
| chr17 | 2683261 | 2683288 | + | 2.00792115374673 | 1.13699574206438 | 2.38645642073598 | 2.13421729395427 |
| chr17 | 2683456 | 2683491 | + | 11.043566345607 | 26.1509020674808 | 7.15936926220794 | 4.26843458790854 |
| chr17 | 2683506 | 2683550 | + | 10.0396057687336 | 10.2329616785794 | 1.19322821036799 | 13.8724124107028 |
| chr17 | 2683848 | 2683850 | + | 2.00792115374673 | 1.13699574206438 | 1.19322821036799 | 3.20132594093141 |
| chr17 | 2684166 | 2684185 | + | 3.01188173062009 | 3.41098722619315 | 1.19322821036799 | 3.20132594093141 |
| chr17 | 2684218 | 2684230 | + | 26.1029749987075 | 31.8358807778027 | 9.54582568294392 | 26.6777161744284 |
| chr17 | 2684362 | 2684408 | + | 35.1386201905678 | 62.534765813541 | 17.8984231555198 | 11.7381951167485 |
| chr17 | 2689461 | 2689499 | - | 13.0514874993537 | 13.6439489047726 | 10.7390538933119 | 5.33554323488568 |
| chr17 | 2689741 | 2689812 | - | 43.1703048055547 | 15.9179403889013 | 28.6374770488317 | 99.2411041688736 |
| chr17 | 2689856 | 2689881 | - | 22.087132691214 | 7.95897019445067 | 11.9322821036799 | 13.8724124107028 |
| chr17 | 2689940 | 2689968 | - | 19.0752509605939 | 9.09596593651506 | 11.9322821036799 | 5.33554323488568 |
| chr17 | 2690002 | 2690080 | - | 77.3049644192491 | 20.4659233571589 | 39.3765309421437 | 24.5434988804741 |
| chr17 | 2690176 | 2690224 | - | 15.0594086531005 | 10.2329616785794 | 9.54582568294392 | 9.60397782279422 |
| chr17 | 2690274 | 2690299 | - | 13.0514874993537 | 4.54798296825753 | 8.35259747257593 | 19.2079556455884 |
| chr17 | 2690387 | 2690450 | - | 13.0514874993537 | 5.68497871032191 | 11.9322821036799 | 14.9395210576799 |
| chr17 | 3662917 | 3662954 | - | 120.475269224804 | 129.61751459534 | 10.7390538933119 | 16.006629704657 |
| chr17 | 3663543 | 3663570 | - | 80.3168461498692 | 143.261463500112 | 36.9900745214077 | 16.006629704657 |
| chr17 | 3663874 | 3663887 | - | 1.00396057687336 | 4.54798296825753 | 1.19322821036799 | 2.13421729395427 |
| chr17 | 3664763 | 3664795 | - | 9.03564519186028 | 2.27399148412876 | 4.77291284147196 | 1.06710864697714 |
| chr17 | 3669463 | 3669588 | + | 24.0950538449607 | 48.8908169087684 | 4.77291284147196 | 17.0737383516342 |
| chr17 | 3669621 | 3669631 | + | 32.1267384599477 | 64.8087572976698 | 15.5119667347839 | 25.6106075274513 |
| chr17 | 3714663 | 3714685 | - | 8.03168461498692 | 10.2329616785794 | 1.19322821036799 | 1.06710864697714 |
| chr17 | 3714798 | 3714831 | - | 10.0396057687336 | 7.95897019445067 | 9.54582568294392 | 2.13421729395427 |
| chr17 | 3807348 | 3807358 | - | 4.01584230749346 | 5.68497871032191 | 7.15936926220794 | 6.40265188186281 |
| chr17 | 3807447 | 3807449 | - | 1.00396057687336 | 3.41098722619315 | 1.19322821036799 | 3.20132594093141 |
| chr17 | 3807703 | 3807740 | - | 7.02772403811355 | 12.5069531627082 | 8.35259747257593 | 10.6710864697714 |
| chr17 | 3808025 | 3808040 | - | 5.01980288436682 | 3.41098722619315 | 3.57968463110397 | 4.26843458790854 |
| chr17 | 3812310 | 3812342 | - | 13.0514874993537 | 9.09596593651506 | 1.19322821036799 | 3.20132594093141 |
| chr17 | 3924459 | 3924541 | - | 1.00396057687336 | 1.13699574206438 | 44.1494437836156 | 38.4159112911769 |
| chr17 | 4004478 | 4004500 | - | 1.00396057687336 | 6.82197445238629 | 1.19322821036799 | 3.20132594093141 |
| chr17 | 4157262 | 4157272 | + | 13.0514874993537 | 11.3699574206438 | 2.38645642073598 | 3.20132594093141 |
| chr17 | 4157357 | 4157369 | + | 28.1108961524542 | 67.0827487817985 | 3.57968463110397 | 10.6710864697714 |
| chr17 | 4164007 | 4164060 | - | 9.03564519186028 | 11.3699574206438 | 1.19322821036799 | 7.46976052883995 |
| chr17 | 4270678 | 4270716 | - | 2.00792115374673 | 2.27399148412876 | 3.57968463110397 | 5.33554323488568 |
| chr17 | 4539036 | 4539043 | - | 9.03564519186028 | 4.54798296825753 | 8.35259747257593 | 7.46976052883995 |
| chr17 | 4671420 | 4671478 | - | 10.0396057687336 | 10.2329616785794 | 10.7390538933119 | 9.60397782279422 |
| chr17 | 4721342 | 4721355 | + | 15.0594086531005 | 13.6439489047726 | 2.38645642073598 | 4.26843458790854 |
| chr17 | 4721426 | 4721475 | + | 9.03564519186028 | 18.1919318730301 | 1.19322821036799 | 3.20132594093141 |
| chr17 | 4733119 | 4733125 | + | 2.00792115374673 | 5.68497871032191 | 1.19322821036799 | 1.06710864697714 |
| chr17 | 4733195 | 4733217 | + | 9.03564519186028 | 5.68497871032191 | 2.38645642073598 | 2.13421729395427 |
| chr17 | 4733315 | 4733351 | + | 21.0831721143407 | 11.3699574206438 | 2.38645642073598 | 7.46976052883995 |
| chr17 | 4733384 | 4733408 | + | 76.3010038423757 | 47.753821166704 | 19.0916513658878 | 19.2079556455884 |
| chr17 | 4733424 | 4733451 | + | 174.689140375965 | 140.987472015983 | 57.2749540976635 | 51.2212150549025 |
| chr17 | 4798353 | 4798472 | + | 220.87132691214 | 218.303182476361 | 119.322821036799 | 170.737383516342 |
| chr17 | 4897303 | 4897306 | + | 4.01584230749346 | 1.13699574206438 | 7.15936926220794 | 4.26843458790854 |
| chr17 | 4897359 | 4897379 | + | 1.00396057687336 | 1.13699574206438 | 5.96614105183995 | 1.06710864697714 |
| chr17 | 4898005 | 4898027 | + | 3.01188173062009 | 7.95897019445067 | 3.57968463110397 | 5.33554323488568 |
| chr17 | 4937267 | 4937273 | - | 7.02772403811355 | 12.5069531627082 | 1.19322821036799 | 2.13421729395427 |
| chr17 | 4937332 | 4937371 | - | 42.1663442286813 | 67.0827487817985 | 17.8984231555198 | 26.6777161744284 |
| chr17 | 4937408 | 4937418 | - | 2.00792115374673 | 3.41098722619315 | 3.57968463110397 | 2.13421729395427 |
| chr17 | 4945149 | 4945194 | + | 55.217831728035 | 75.0417189762492 | 17.8984231555198 | 18.1408469986113 |
| chr17 | 4945688 | 4945767 | - | 200.792115374673 | 204.659233571589 | 118.129592826431 | 234.76390233497 |
| chr17 | 4945802 | 4945908 | - | 23.0910932680874 | 11.3699574206438 | 25.0577924177278 | 49.0869977609482 |
| chr17 | 4959256 | 4959268 | - | 24.0950538449607 | 23.876910583352 | 23.8645642073598 | 17.0737383516342 |
| chr17 | 4959703 | 4959704 | - | 4.01584230749346 | 7.95897019445067 | 3.57968463110397 | 5.33554323488568 |
| chr17 | 4968015 | 4968082 | - | 6.02376346124019 | 25.0139063254164 | 4.77291284147196 | 2.13421729395427 |
| chr17 | 4987687 | 4987753 | + | 9.03564519186028 | 3.41098722619315 | 54.8884976769275 | 65.0936274656053 |
| chr17 | 5024163 | 5024217 | + | 10.0396057687336 | 5.68497871032191 | 4.77291284147196 | 8.53686917581709 |
| chr17 | 5024227 | 5024359 | + | 18.0712903837206 | 14.780944646837 | 16.7051949451519 | 27.7448248214055 |
| chr17 | 5024384 | 5024406 | + | 4.01584230749346 | 4.54798296825753 | 1.19322821036799 | 1.06710864697714 |
| chr17 | 5024437 | 5024454 | + | 39.1544624980612 | 27.2878978095452 | 10.7390538933119 | 17.0737383516342 |
| chr17 | 5105812 | 5105818 | - | 12.0475269224804 | 21.6029190992233 | 2.38645642073598 | 3.20132594093141 |
| chr17 | 5111912 | 5112020 | + | 156.617849992245 | 163.727386857271 | 83.5259747257593 | 260.374509862421 |
| chr17 | 5112094 | 5112111 | + | 4.01584230749346 | 7.95897019445067 | 7.15936926220794 | 12.8053037637256 |
| chr17 | 5114298 | 5114327 | + | 3.01188173062009 | 6.82197445238629 | 1.19322821036799 | 2.13421729395427 |
| chr17 | 5383398 | 5383402 | + | 6.02376346124019 | 3.41098722619315 | 2.38645642073598 | 3.20132594093141 |
| chr17 | 5383881 | 5383884 | + | 4.01584230749346 | 5.68497871032191 | 5.96614105183995 | 1.06710864697714 |
| chr17 | 5384143 | 5384160 | + | 5.01980288436682 | 4.54798296825753 | 4.77291284147196 | 5.33554323488568 |
| chr17 | 5385508 | 5385520 | + | 1.00396057687336 | 1.13699574206438 | 2.38645642073598 | 4.26843458790854 |
| chr17 | 5385651 | 5385675 | + | 15.0594086531005 | 22.7399148412876 | 31.0239334695677 | 21.3421729395427 |
| chr17 | 5385719 | 5385776 | + | 74.293082688629 | 52.3018041349616 | 93.0718004087032 | 67.2278447595595 |
| chr17 | 5386089 | 5386094 | - | 3.01188173062009 | 3.41098722619315 | 2.38645642073598 | 3.20132594093141 |
| chr17 | 5386136 | 5386140 | - | 6.02376346124019 | 6.82197445238629 | 4.77291284147196 | 5.33554323488568 |
| chr17 | 5386161 | 5386197 | - | 6.02376346124019 | 10.2329616785794 | 5.96614105183995 | 9.60397782279422 |
| chr17 | 5432748 | 5432786 | + | 28.1108961524542 | 77.315710460378 | 19.0916513658878 | 19.2079556455884 |
| chr17 | 5432865 | 5432922 | - | 89.3524913417294 | 137.57648478979 | 45.3426719939836 | 68.2949534065367 |
| chr17 | 5432953 | 5432979 | - | 2.00792115374673 | 4.54798296825753 | 4.77291284147196 | 5.33554323488568 |
| chr17 | 5433000 | 5433025 | - | 7.02772403811355 | 2.27399148412876 | 8.35259747257593 | 7.46976052883995 |
| chr17 | 5441735 | 5441744 | - | 1.00396057687336 | 5.68497871032191 | 3.57968463110397 | 1.06710864697714 |
| chr17 | 5441793 | 5441810 | - | 14.0554480762271 | 3.41098722619315 | 5.96614105183995 | 6.40265188186281 |
| chr17 | 5441873 | 5441875 | - | 10.0396057687336 | 6.82197445238629 | 2.38645642073598 | 4.26843458790854 |
| chr17 | 5442132 | 5442137 | - | 5.01980288436682 | 2.27399148412876 | 11.9322821036799 | 3.20132594093141 |
| chr17 | 5442186 | 5442189 | - | 35.1386201905678 | 25.0139063254164 | 4.77291284147196 | 21.3421729395427 |
| chr17 | 5442332 | 5442343 | - | 17.0673298068472 | 36.3838637460602 | 7.15936926220794 | 4.26843458790854 |
| chr17 | 5442455 | 5442704 | - | 151.598047107878 | 120.521548658824 | 100.231169670911 | 149.395210576799 |
| chr17 | 5442747 | 5442767 | - | 3.01188173062009 | 3.41098722619315 | 2.38645642073598 | 2.13421729395427 |
| chr17 | 5443114 | 5443131 | - | 294.160449023896 | 180.782322988237 | 165.858721241151 | 183.542687280067 |
| chr17 | 5443213 | 5443235 | - | 39.1544624980612 | 43.2058381984465 | 11.9322821036799 | 5.33554323488568 |
| chr17 | 5471346 | 5471351 | - | 4.01584230749346 | 7.95897019445067 | 3.57968463110397 | 1.06710864697714 |
| chr17 | 5474303 | 5474363 | - | 64.2534769198953 | 125.069531627082 | 13.1255103140479 | 14.9395210576799 |
| chr17 | 5474384 | 5474391 | - | 2.00792115374673 | 6.82197445238629 | 1.19322821036799 | 3.20132594093141 |
| chr17 | 5474628 | 5474653 | - | 10.0396057687336 | 12.5069531627082 | 1.19322821036799 | 9.60397782279422 |
| chr17 | 5490588 | 5490596 | + | 2.00792115374673 | 5.68497871032191 | 1.19322821036799 | 2.13421729395427 |
| chr17 | 5490680 | 5490707 | + | 5.01980288436682 | 13.6439489047726 | 4.77291284147196 | 2.13421729395427 |
| chr17 | 6424019 | 6424020 | - | 28.1108961524542 | 18.1919318730301 | 8.35259747257593 | 19.2079556455884 |
| chr17 | 6451010 | 6451015 | + | 10.0396057687336 | 1.13699574206438 | 1.19322821036799 | 1.06710864697714 |
| chr17 | 6642961 | 6643147 | + | 311.227778830743 | 256.96103770655 | 182.563916186302 | 177.140035398205 |
| chr17 | 6644341 | 6644360 | - | 3.01188173062009 | 6.82197445238629 | 3.57968463110397 | 10.6710864697714 |
| chr17 | 6644356 | 6644382 | + | 20.0792115374673 | 14.780944646837 | 5.96614105183995 | 8.53686917581709 |
| chr17 | 6644442 | 6644446 | + | 4.01584230749346 | 5.68497871032191 | 1.19322821036799 | 2.13421729395427 |
| chr17 | 6644508 | 6644537 | + | 31.1227778830743 | 19.3289276150945 | 2.38645642073598 | 3.20132594093141 |
| chr17 | 6644524 | 6644534 | - | 6.02376346124019 | 4.54798296825753 | 1.19322821036799 | 6.40265188186281 |
| chr17 | 6644581 | 6644600 | - | 2.00792115374673 | 4.54798296825753 | 1.19322821036799 | 1.06710864697714 |
| chr17 | 7014447 | 7014504 | + | 233.922814411494 | 236.495114349391 | 27.4442488384638 | 133.388580872142 |
| chr17 | 7017319 | 7017322 | + | 1.00396057687336 | 1.13699574206438 | 1.19322821036799 | 1.06710864697714 |
| chr17 | 7017418 | 7017431 | + | 13.0514874993537 | 2.27399148412876 | 4.77291284147196 | 11.7381951167485 |
| chr17 | 7017461 | 7017482 | + | 2.00792115374673 | 2.27399148412876 | 2.38645642073598 | 2.13421729395427 |
| chr17 | 7189953 | 7190009 | - | 11.043566345607 | 7.95897019445067 | 14.3187385244159 | 11.7381951167485 |
| chr17 | 7225147 | 7225247 | + | 36.1425807674411 | 28.4248935516096 | 11.9322821036799 | 22.4092815865198 |
| chr17 | 7225408 | 7225437 | - | 5.01980288436682 | 9.09596593651506 | 5.96614105183995 | 2.13421729395427 |
| chr17 | 7225459 | 7225485 | - | 8.03168461498692 | 5.68497871032191 | 3.57968463110397 | 4.26843458790854 |
| chr17 | 7225750 | 7225753 | - | 2.00792115374673 | 1.13699574206438 | 1.19322821036799 | 2.13421729395427 |
| chr17 | 7235314 | 7235348 | - | 8.03168461498692 | 5.68497871032191 | 1.19322821036799 | 6.40265188186281 |
| chr17 | 7235405 | 7235414 | - | 4.01584230749346 | 2.27399148412876 | 1.19322821036799 | 2.13421729395427 |
| chr17 | 7240435 | 7240498 | - | 61.2415951892752 | 69.3567402659273 | 31.0239334695677 | 64.0265188186281 |
| chr17 | 7240669 | 7240698 | - | 198.784194220926 | 234.221122865263 | 102.617626091647 | 90.7042349930565 |
| chr17 | 7240719 | 7240749 | - | 52.205949997415 | 71.6307317500561 | 81.1395183050233 | 62.959410171651 |
| chr17 | 7240789 | 7240850 | - | 98.3881365335897 | 92.0966551072149 | 192.109741869246 | 73.6304966414224 |
| chr17 | 7240877 | 7240922 | - | 17.0673298068472 | 14.780944646837 | 34.6036181006717 | 21.3421729395427 |
| chr17 | 7241341 | 7241403 | - | 16.0633692299738 | 19.3289276150945 | 4.77291284147196 | 11.7381951167485 |
| chr17 | 7241436 | 7241493 | - | 6.02376346124019 | 7.95897019445067 | 27.4442488384638 | 11.7381951167485 |
| chr17 | 7241604 | 7241647 | - | 18.0712903837206 | 19.3289276150945 | 26.2510206280958 | 20.2750642925656 |
| chr17 | 7243701 | 7243735 | - | 14.0554480762271 | 5.68497871032191 | 7.15936926220794 | 7.46976052883995 |
| chr17 | 7243828 | 7243840 | - | 5.01980288436682 | 9.09596593651506 | 3.57968463110397 | 2.13421729395427 |
| chr17 | 7243883 | 7243902 | - | 3.01188173062009 | 3.41098722619315 | 3.57968463110397 | 5.33554323488568 |
| chr17 | 7243927 | 7243963 | - | 12.0475269224804 | 4.54798296825753 | 2.38645642073598 | 4.26843458790854 |
| chr17 | 7243993 | 7244032 | - | 1.00396057687336 | 2.27399148412876 | 1.19322821036799 | 1.06710864697714 |
| chr17 | 7246832 | 7246862 | - | 17.0673298068472 | 13.6439489047726 | 9.54582568294392 | 6.40265188186281 |
| chr17 | 7247095 | 7247122 | - | 24.0950538449607 | 13.6439489047726 | 9.54582568294392 | 11.7381951167485 |
| chr17 | 7247175 | 7247180 | - | 7.02772403811355 | 4.54798296825753 | 4.77291284147196 | 10.6710864697714 |
| chr17 | 7259718 | 7259737 | + | 5.01980288436682 | 9.09596593651506 | 2.38645642073598 | 9.60397782279422 |
| chr17 | 7259854 | 7259907 | + | 55.217831728035 | 106.877599754052 | 63.2410951495034 | 78.966039876308 |
| chr17 | 7287712 | 7287720 | + | 1.00396057687336 | 1.13699574206438 | 3.57968463110397 | 12.8053037637256 |
| chr17 | 7311825 | 7311842 | + | 8.03168461498692 | 4.54798296825753 | 8.35259747257593 | 9.60397782279422 |
| chr17 | 7311873 | 7311903 | + | 19.0752509605939 | 5.68497871032191 | 9.54582568294392 | 7.46976052883995 |
| chr17 | 7312001 | 7312018 | + | 8.03168461498692 | 4.54798296825753 | 21.4781077866238 | 23.476390233497 |
| chr17 | 7312097 | 7312099 | + | 6.02376346124019 | 5.68497871032191 | 23.8645642073598 | 20.2750642925656 |
| chr17 | 7312354 | 7312359 | + | 9.03564519186028 | 5.68497871032191 | 5.96614105183995 | 20.2750642925656 |
| chr17 | 7312681 | 7312712 | - | 73.2891221117556 | 60.2607743294122 | 10.7390538933119 | 10.6710864697714 |
| chr17 | 7314520 | 7314548 | - | 3.01188173062009 | 10.2329616785794 | 2.38645642073598 | 1.06710864697714 |
| chr17 | 7314580 | 7314599 | - | 36.1425807674411 | 34.1098722619315 | 16.7051949451519 | 12.8053037637256 |
| chr17 | 7314961 | 7315017 | - | 68.2693192273888 | 45.4798296825753 | 35.7968463110397 | 33.0803680562912 |
| chr17 | 7315067 | 7315178 | - | 95.3762548029696 | 73.9047232341848 | 13.1255103140479 | 13.8724124107028 |
| chr17 | 7315274 | 7315410 | - | 1449.71907300514 | 803.855989639518 | 341.263268165245 | 614.65458065883 |
| chr17 | 7571768 | 7571811 | + | 15.0594086531005 | 25.0139063254164 | 4.77291284147196 | 2.13421729395427 |
| chr17 | 7571927 | 7571949 | + | 6.02376346124019 | 13.6439489047726 | 2.38645642073598 | 2.13421729395427 |
| chr17 | 7572726 | 7572865 | + | 27150.1058803864 | 54839.5786312493 | 23852.6319252561 | 16167.7631103506 |
| chr17 | 7574584 | 7574593 | + | 3.01188173062009 | 5.68497871032191 | 3.57968463110397 | 4.26843458790854 |
| chr17 | 7574668 | 7574676 | + | 11.043566345607 | 19.3289276150945 | 25.0577924177278 | 46.952780466994 |
| chr17 | 7575097 | 7575159 | + | 86.3406096111093 | 97.7816338175368 | 120.516049247167 | 57.6238669367653 |
| chr17 | 7576536 | 7576592 | + | 96.380215379843 | 53.438799877026 | 54.8884976769275 | 89.6371263460794 |
| chr17 | 7576637 | 7576693 | + | 79.3128855729958 | 61.3977700714766 | 70.4004644117114 | 59.7580842307196 |
| chr17 | 7576809 | 7576811 | + | 1.00396057687336 | 2.27399148412876 | 1.19322821036799 | 1.06710864697714 |
| chr17 | 7576874 | 7576929 | + | 13.0514874993537 | 4.54798296825753 | 2.38645642073598 | 2.13421729395427 |
| chr17 | 7577321 | 7577352 | + | 30.1188173062009 | 31.8358807778027 | 36.9900745214077 | 21.3421729395427 |
| chr17 | 7577437 | 7577462 | + | 59.2336740355285 | 77.315710460378 | 83.5259747257593 | 45.8856718200168 |
| chr17 | 7577486 | 7577488 | + | 9.03564519186028 | 4.54798296825753 | 9.54582568294392 | 1.06710864697714 |
| chr17 | 7577675 | 7577696 | + | 8.03168461498692 | 6.82197445238629 | 14.3187385244159 | 11.7381951167485 |
| chr17 | 7577829 | 7577864 | + | 102.403978841083 | 77.315710460378 | 120.516049247167 | 93.9055609339879 |
| chr17 | 7578512 | 7578551 | + | 105.415860571703 | 80.7266976865711 | 35.7968463110397 | 86.435800405148 |
| chr17 | 7578664 | 7578691 | + | 16.0633692299738 | 23.876910583352 | 4.77291284147196 | 8.53686917581709 |
| chr17 | 7578801 | 7578974 | + | 252.998065372088 | 345.646705587572 | 76.3666054635513 | 106.710864697714 |
| chr17 | 7579565 | 7579673 | + | 649.562493237067 | 476.401215924976 | 2046.3863807811 | 1611.33405693548 |
| chr17 | 7579703 | 7579716 | + | 3.01188173062009 | 2.27399148412876 | 7.15936926220794 | 2.13421729395427 |
| chr17 | 7579812 | 7579823 | + | 4.01584230749346 | 2.27399148412876 | 23.8645642073598 | 4.26843458790854 |
| chr17 | 7579891 | 7579909 | + | 4.01584230749346 | 1.13699574206438 | 7.15936926220794 | 1.06710864697714 |
| chr17 | 7582034 | 7582094 | + | 2.00792115374673 | 3.41098722619315 | 7.15936926220794 | 3.20132594093141 |
| chr17 | 7587431 | 7587452 | + | 13.0514874993537 | 1.13699574206438 | 10.7390538933119 | 7.46976052883995 |
| chr17 | 7587499 | 7587564 | + | 9.03564519186028 | 15.9179403889013 | 9.54582568294392 | 5.33554323488568 |
| chr17 | 7587626 | 7587630 | + | 12.0475269224804 | 12.5069531627082 | 4.77291284147196 | 6.40265188186281 |
| chr17 | 7587659 | 7587744 | + | 80.3168461498692 | 56.8497871032191 | 46.5359002043516 | 33.0803680562912 |
| chr17 | 7587787 | 7587808 | + | 3.01188173062009 | 7.95897019445067 | 5.96614105183995 | 4.26843458790854 |
| chr17 | 7587852 | 7587874 | + | 3.01188173062009 | 5.68497871032191 | 3.57968463110397 | 9.60397782279422 |
| chr17 | 7587930 | 7587967 | + | 13.0514874993537 | 1.13699574206438 | 5.96614105183995 | 18.1408469986113 |
| chr17 | 7587989 | 7588177 | + | 730.883299963809 | 667.416500591792 | 225.52013175955 | 425.776350143877 |
| chr17 | 7591268 | 7591355 | - | 23.0910932680874 | 38.657855230189 | 14.3187385244159 | 19.2079556455884 |
| chr17 | 7626262 | 7626343 | - | 32.1267384599477 | 71.6307317500561 | 8.35259747257593 | 6.40265188186281 |
| chr17 | 7669100 | 7669129 | - | 8.03168461498692 | 9.09596593651506 | 7.15936926220794 | 9.60397782279422 |
| chr17 | 7669326 | 7669345 | - | 10.0396057687336 | 7.95897019445067 | 2.38645642073598 | 1.06710864697714 |
| chr17 | 7669394 | 7669430 | - | 44.174265382428 | 69.3567402659273 | 15.5119667347839 | 1.06710864697714 |
| chr17 | 7670626 | 7670628 | - | 3.01188173062009 | 6.82197445238629 | 1.19322821036799 | 1.06710864697714 |
| chr17 | 7853650 | 7853661 | + | 1.00396057687336 | 1.13699574206438 | 2.38645642073598 | 1.06710864697714 |
| chr17 | 7853922 | 7853971 | + | 10.0396057687336 | 7.95897019445067 | 4.77291284147196 | 2.13421729395427 |
| chr17 | 7856705 | 7856768 | - | 301.188173062009 | 317.221812035963 | 69.2072362013434 | 108.845081991668 |
| chr17 | 7859926 | 7859935 | + | 9.03564519186028 | 4.54798296825753 | 1.19322821036799 | 1.06710864697714 |
| chr17 | 7862055 | 7862080 | + | 11.043566345607 | 6.82197445238629 | 1.19322821036799 | 1.06710864697714 |
| chr17 | 7911698 | 7911705 | + | 3.01188173062009 | 4.54798296825753 | 1.19322821036799 | 2.13421729395427 |
| chr17 | 7911744 | 7911747 | + | 2.00792115374673 | 2.27399148412876 | 1.19322821036799 | 1.06710864697714 |
| chr17 | 7911889 | 7911918 | + | 27.1069355755808 | 14.780944646837 | 7.15936926220794 | 5.33554323488568 |
| chr17 | 7911942 | 7912097 | + | 165.653495184105 | 114.836569948503 | 73.9801490428153 | 86.435800405148 |
| chr17 | 7912121 | 7912133 | + | 349.378280751931 | 181.919318730301 | 44.1494437836156 | 115.247733873531 |
| chr17 | 7912612 | 7912613 | + | 2.00792115374673 | 1.13699574206438 | 1.19322821036799 | 3.20132594093141 |
| chr17 | 7915154 | 7915161 | - | 3.01188173062009 | 1.13699574206438 | 1.19322821036799 | 4.26843458790854 |
| chr17 | 7915183 | 7915211 | - | 3.01188173062009 | 7.95897019445067 | 3.57968463110397 | 18.1408469986113 |
| chr17 | 7930369 | 7930393 | - | 18.0712903837206 | 23.876910583352 | 13.1255103140479 | 20.2750642925656 |
| chr17 | 7949494 | 7949516 | + | 10.0396057687336 | 5.68497871032191 | 2.38645642073598 | 11.7381951167485 |
| chr17 | 8080115 | 8080116 | + | 1.00396057687336 | 1.13699574206438 | 1.19322821036799 | 4.26843458790854 |
| chr17 | 8159180 | 8159197 | - | 5.01980288436682 | 4.54798296825753 | 2.38645642073598 | 3.20132594093141 |
| chr17 | 8159331 | 8159333 | - | 2.00792115374673 | 2.27399148412876 | 1.19322821036799 | 1.06710864697714 |
| chr17 | 8159943 | 8159957 | - | 3.01188173062009 | 4.54798296825753 | 2.38645642073598 | 1.06710864697714 |
| chr17 | 8160174 | 8160176 | - | 1.00396057687336 | 1.13699574206438 | 1.19322821036799 | 2.13421729395427 |
| chr17 | 8160275 | 8160306 | - | 2.00792115374673 | 9.09596593651506 | 4.77291284147196 | 3.20132594093141 |
| chr17 | 8160319 | 8160330 | - | 3.01188173062009 | 2.27399148412876 | 1.19322821036799 | 3.20132594093141 |
| chr17 | 8160425 | 8160432 | - | 9.03564519186028 | 6.82197445238629 | 2.38645642073598 | 8.53686917581709 |
| chr17 | 8160461 | 8160485 | - | 15.0594086531005 | 19.3289276150945 | 4.77291284147196 | 2.13421729395427 |
| chr17 | 8160610 | 8160757 | - | 48.1901076899215 | 56.8497871032191 | 20.2848795762558 | 11.7381951167485 |
| chr17 | 8160783 | 8160849 | - | 19.0752509605939 | 13.6439489047726 | 4.77291284147196 | 10.6710864697714 |
| chr17 | 8173577 | 8173590 | - | 12.0475269224804 | 27.2878978095452 | 1.19322821036799 | 3.20132594093141 |
| chr17 | 8174115 | 8174165 | - | 156.617849992245 | 164.864382599335 | 50.1155848354556 | 115.247733873531 |
| chr17 | 8174189 | 8174190 | - | 6.02376346124019 | 9.09596593651506 | 1.19322821036799 | 5.33554323488568 |
| chr17 | 8174224 | 8174260 | - | 11.043566345607 | 12.5069531627082 | 2.38645642073598 | 4.26843458790854 |
| chr17 | 8174546 | 8174556 | - | 5.01980288436682 | 3.41098722619315 | 1.19322821036799 | 1.06710864697714 |
| chr17 | 8188376 | 8188516 | - | 239.946577872734 | 205.796229313653 | 101.424397881279 | 106.710864697714 |
| chr17 | 8189459 | 8189497 | + | 30.1188173062009 | 20.4659233571589 | 4.77291284147196 | 1.06710864697714 |
| chr17 | 8204783 | 8204818 | - | 16.0633692299738 | 14.780944646837 | 11.9322821036799 | 11.7381951167485 |
| chr17 | 8204845 | 8204870 | - | 8.03168461498692 | 12.5069531627082 | 5.96614105183995 | 12.8053037637256 |
| chr17 | 8269272 | 8269287 | + | 9.03564519186028 | 4.54798296825753 | 14.3187385244159 | 7.46976052883995 |
| chr17 | 8289975 | 8290065 | + | 83.3287278804893 | 127.343523111211 | 9.54582568294392 | 23.476390233497 |
| chr17 | 8467851 | 8467892 | + | 7.02772403811355 | 7.95897019445067 | 1.19322821036799 | 4.26843458790854 |
| chr17 | 8468034 | 8468136 | + | 4.01584230749346 | 4.54798296825753 | 1.19322821036799 | 3.20132594093141 |
| chr17 | 9250499 | 9250602 | - | 7.02772403811355 | 5.68497871032191 | 3.57968463110397 | 8.53686917581709 |
| chr17 | 10672518 | 10672539 | - | 2.00792115374673 | 1.13699574206438 | 1.19322821036799 | 1.06710864697714 |
| chr17 | 10675790 | 10675814 | - | 3.01188173062009 | 2.27399148412876 | 5.96614105183995 | 9.60397782279422 |
| chr17 | 10676670 | 10676671 | - | 3.01188173062009 | 2.27399148412876 | 1.19322821036799 | 1.06710864697714 |
| chr17 | 10680396 | 10680433 | - | 2.00792115374673 | 3.41098722619315 | 1.19322821036799 | 1.06710864697714 |
| chr17 | 10680495 | 10680529 | - | 23.0910932680874 | 13.6439489047726 | 7.15936926220794 | 4.26843458790854 |
| chr17 | 11977502 | 11977504 | - | 2.00792115374673 | 4.54798296825753 | 2.38645642073598 | 3.20132594093141 |
| chr17 | 12141916 | 12141938 | + | 1.00396057687336 | 2.27399148412876 | 1.19322821036799 | 1.06710864697714 |
| chr17 | 12142019 | 12142023 | + | 2.00792115374673 | 2.27399148412876 | 2.38645642073598 | 2.13421729395427 |
| chr17 | 12992430 | 12992471 | - | 26.1029749987075 | 31.8358807778027 | 9.54582568294392 | 14.9395210576799 |
| chr17 | 12992522 | 12992526 | - | 4.01584230749346 | 4.54798296825753 | 5.96614105183995 | 12.8053037637256 |
| chr17 | 13011768 | 13011781 | - | 6.02376346124019 | 4.54798296825753 | 9.54582568294392 | 3.20132594093141 |
| chr17 | 14207310 | 14207347 | + | 8.03168461498692 | 9.09596593651506 | 7.15936926220794 | 5.33554323488568 |
| chr17 | 14208218 | 14208235 | + | 2.00792115374673 | 1.13699574206438 | 1.19322821036799 | 1.06710864697714 |
| chr17 | 14208267 | 14208292 | + | 9.03564519186028 | 5.68497871032191 | 4.77291284147196 | 3.20132594093141 |
| chr17 | 15230779 | 15230784 | - | 2.00792115374673 | 1.13699574206438 | 4.77291284147196 | 1.06710864697714 |
| chr17 | 15502781 | 15502802 | - | 2.00792115374673 | 3.41098722619315 | 1.19322821036799 | 1.06710864697714 |
| chr17 | 15976626 | 15976671 | - | 9.03564519186028 | 9.09596593651506 | 4.77291284147196 | 10.6710864697714 |
| chr17 | 15977778 | 15977806 | - | 7.02772403811355 | 18.1919318730301 | 4.77291284147196 | 3.20132594093141 |
| chr17 | 16027534 | 16027564 | + | 11.043566345607 | 3.41098722619315 | 5.96614105183995 | 2.13421729395427 |
| chr17 | 16027657 | 16027664 | + | 17.0673298068472 | 7.95897019445067 | 16.7051949451519 | 18.1408469986113 |
| chr17 | 16027790 | 16027863 | + | 16.0633692299738 | 34.1098722619315 | 10.7390538933119 | 20.2750642925656 |
| chr17 | 16027958 | 16027972 | + | 12.0475269224804 | 5.68497871032191 | 11.9322821036799 | 11.7381951167485 |
| chr17 | 16028011 | 16028038 | + | 4.01584230749346 | 4.54798296825753 | 2.38645642073598 | 5.33554323488568 |
| chr17 | 16028072 | 16028118 | + | 16.0633692299738 | 20.4659233571589 | 19.0916513658878 | 19.2079556455884 |
| chr17 | 16028170 | 16028201 | + | 101.40001826421 | 145.535454984241 | 54.8884976769275 | 71.4962793474681 |
| chr17 | 16028315 | 16028347 | + | 11.043566345607 | 15.9179403889013 | 9.54582568294392 | 11.7381951167485 |
| chr17 | 16028377 | 16028408 | + | 8.03168461498692 | 14.780944646837 | 11.9322821036799 | 17.0737383516342 |
| chr17 | 16028456 | 16028495 | + | 123.487150955424 | 209.207216539846 | 122.902505667903 | 216.623055336359 |
| chr17 | 16028578 | 16028586 | + | 2.00792115374673 | 5.68497871032191 | 3.57968463110397 | 4.26843458790854 |
| chr17 | 16028692 | 16028714 | + | 30.1188173062009 | 43.2058381984465 | 34.6036181006717 | 64.0265188186281 |
| chr17 | 16031064 | 16031085 | - | 4.01584230749346 | 1.13699574206438 | 2.38645642073598 | 2.13421729395427 |
| chr17 | 16032024 | 16032064 | - | 7.02772403811355 | 5.68497871032191 | 1.19322821036799 | 6.40265188186281 |
| chr17 | 16342560 | 16342665 | - | 13.0514874993537 | 11.3699574206438 | 4.77291284147196 | 2.13421729395427 |
| chr17 | 16342694 | 16342715 | - | 18.0712903837206 | 9.09596593651506 | 8.35259747257593 | 8.53686917581709 |
| chr17 | 16342737 | 16342943 | - | 93.3683336492229 | 53.438799877026 | 57.2749540976635 | 89.6371263460794 |
| chr17 | 16382582 | 16382610 | + | 260.025789410201 | 268.330995127194 | 177.79100334483 | 304.125964388484 |
| chr17 | 16382707 | 16382727 | + | 32.1267384599477 | 35.2468680039958 | 21.4781077866238 | 5.33554323488568 |
| chr17 | 16438961 | 16439085 | + | 1256.95864224545 | 1824.87816601333 | 1202.77403605093 | 416.172372321083 |
| chr17 | 16439134 | 16439137 | + | 19.0752509605939 | 11.3699574206438 | 3.57968463110397 | 3.20132594093141 |
| chr17 | 16439305 | 16439412 | + | 44.174265382428 | 36.3838637460602 | 145.573841664895 | 75.7647139353766 |
| chr17 | 16439539 | 16439724 | + | 30.1188173062009 | 43.2058381984465 | 14.3187385244159 | 16.006629704657 |
| chr17 | 16440167 | 16440254 | + | 47.1861471130481 | 38.657855230189 | 103.810854302015 | 60.8251928776967 |
| chr17 | 16441071 | 16441140 | + | 21.0831721143407 | 34.1098722619315 | 101.424397881279 | 71.4962793474681 |
| chr17 | 16441365 | 16441457 | + | 28.1108961524542 | 56.8497871032191 | 126.482190299007 | 112.046407932599 |
| chr17 | 16441480 | 16441564 | + | 26.1029749987075 | 46.6168254246397 | 78.7530618842873 | 45.8856718200168 |
| chr17 | 16441628 | 16441670 | + | 1.00396057687336 | 1.13699574206438 | 11.9322821036799 | 4.26843458790854 |
| chr17 | 16441692 | 16441825 | + | 25.0990144218341 | 46.6168254246397 | 51.3088130458235 | 52.2883237018796 |
| chr17 | 16441847 | 16441872 | + | 10.0396057687336 | 10.2329616785794 | 7.15936926220794 | 5.33554323488568 |
| chr17 | 16441897 | 16442013 | + | 76.3010038423757 | 154.631420920756 | 85.9124311464952 | 108.845081991668 |
| chr17 | 17184962 | 17184993 | + | 4.01584230749346 | 2.27399148412876 | 2.38645642073598 | 4.26843458790854 |
| chr17 | 17192437 | 17192450 | + | 468.849589399861 | 293.344901452611 | 44.1494437836156 | 56.5567582897882 |
| chr17 | 17246858 | 17246869 | - | 4.01584230749346 | 12.5069531627082 | 16.7051949451519 | 1.06710864697714 |
| chr17 | 17246890 | 17246899 | - | 7.02772403811355 | 7.95897019445067 | 3.57968463110397 | 9.60397782279422 |
| chr17 | 17246922 | 17246924 | - | 2.00792115374673 | 10.2329616785794 | 2.38645642073598 | 9.60397782279422 |
| chr17 | 17246997 | 17247034 | - | 23.0910932680874 | 39.7948509722534 | 23.8645642073598 | 45.8856718200168 |
| chr17 | 17491571 | 17491573 | + | 12.0475269224804 | 15.9179403889013 | 10.7390538933119 | 3.20132594093141 |
| chr17 | 17492761 | 17492784 | + | 2.00792115374673 | 1.13699574206438 | 1.19322821036799 | 2.13421729395427 |
| chr17 | 17493124 | 17493170 | + | 137.542599031651 | 102.329616785794 | 76.3666054635513 | 100.308212815851 |
| chr17 | 17505619 | 17505643 | - | 5.01980288436682 | 3.41098722619315 | 2.38645642073598 | 4.26843458790854 |
| chr17 | 17505676 | 17505695 | - | 12.0475269224804 | 5.68497871032191 | 2.38645642073598 | 8.53686917581709 |
| chr17 | 17505766 | 17505803 | - | 7.02772403811355 | 4.54798296825753 | 3.57968463110397 | 6.40265188186281 |
| chr17 | 17812104 | 17812120 | - | 2.00792115374673 | 3.41098722619315 | 2.38645642073598 | 2.13421729395427 |
| chr17 | 17812153 | 17812271 | - | 16.0633692299738 | 40.9318467143178 | 16.7051949451519 | 23.476390233497 |
| chr17 | 17812572 | 17812579 | - | 3.01188173062009 | 3.41098722619315 | 8.35259747257593 | 2.13421729395427 |
| chr17 | 18016863 | 18016865 | + | 3.01188173062009 | 6.82197445238629 | 9.54582568294392 | 3.20132594093141 |
| chr17 | 18018298 | 18018381 | - | 15.0594086531005 | 22.7399148412876 | 9.54582568294392 | 17.0737383516342 |
| chr17 | 18018448 | 18018470 | - | 5.01980288436682 | 2.27399148412876 | 1.19322821036799 | 3.20132594093141 |
| chr17 | 18065298 | 18065302 | + | 2.00792115374673 | 4.54798296825753 | 3.57968463110397 | 3.20132594093141 |
| chr17 | 18065424 | 18065432 | + | 1.00396057687336 | 3.41098722619315 | 1.19322821036799 | 1.06710864697714 |
| chr17 | 18066401 | 18066409 | + | 9.03564519186028 | 3.41098722619315 | 5.96614105183995 | 6.40265188186281 |
| chr17 | 18067805 | 18067824 | + | 6.02376346124019 | 2.27399148412876 | 1.19322821036799 | 2.13421729395427 |
| chr17 | 18068302 | 18068336 | + | 2.00792115374673 | 7.95897019445067 | 4.77291284147196 | 3.20132594093141 |
| chr17 | 18107284 | 18107301 | + | 3.01188173062009 | 2.27399148412876 | 2.38645642073598 | 5.33554323488568 |
| chr17 | 18107882 | 18107905 | + | 2.00792115374673 | 4.54798296825753 | 1.19322821036799 | 6.40265188186281 |
| chr17 | 18208351 | 18208483 | + | 34.1346596136944 | 30.6988850357383 | 65.6275515702394 | 56.5567582897882 |
| chr17 | 18208515 | 18208532 | + | 5.01980288436682 | 2.27399148412876 | 1.19322821036799 | 1.06710864697714 |
| chr17 | 18208555 | 18208593 | + | 6.02376346124019 | 1.13699574206438 | 5.96614105183995 | 3.20132594093141 |
| chr17 | 18208635 | 18208647 | + | 1.00396057687336 | 1.13699574206438 | 2.38645642073598 | 5.33554323488568 |
| chr17 | 18208702 | 18208715 | + | 5.01980288436682 | 5.68497871032191 | 8.35259747257593 | 4.26843458790854 |
| chr17 | 18208770 | 18208787 | + | 4.01584230749346 | 2.27399148412876 | 4.77291284147196 | 16.006629704657 |
| chr17 | 18208946 | 18208985 | + | 13.0514874993537 | 7.95897019445067 | 5.96614105183995 | 11.7381951167485 |
| chr17 | 18209026 | 18209028 | + | 3.01188173062009 | 3.41098722619315 | 3.57968463110397 | 1.06710864697714 |
| chr17 | 18209158 | 18209195 | + | 4.01584230749346 | 9.09596593651506 | 1.19322821036799 | 1.06710864697714 |
| chr17 | 18209222 | 18209228 | + | 4.01584230749346 | 3.41098722619315 | 2.38645642073598 | 1.06710864697714 |
| chr17 | 18209776 | 18209781 | + | 5.01980288436682 | 7.95897019445067 | 4.77291284147196 | 8.53686917581709 |
| chr17 | 18209884 | 18209916 | + | 5.01980288436682 | 13.6439489047726 | 11.9322821036799 | 11.7381951167485 |
| chr17 | 18805742 | 18805782 | + | 5.01980288436682 | 7.95897019445067 | 4.77291284147196 | 10.6710864697714 |
| chr17 | 18930832 | 18930882 | + | 19.0752509605939 | 47.753821166704 | 9.54582568294392 | 26.6777161744284 |
| chr17 | 18931181 | 18931199 | + | 3.01188173062009 | 5.68497871032191 | 1.19322821036799 | 2.13421729395427 |
| chr17 | 18970834 | 18970867 | - | 10.0396057687336 | 11.3699574206438 | 2.38645642073598 | 8.53686917581709 |
| chr17 | 19334796 | 19334797 | + | 6.02376346124019 | 5.68497871032191 | 1.19322821036799 | 2.13421729395427 |
| chr17 | 19335876 | 19335881 | + | 1.00396057687336 | 2.27399148412876 | 2.38645642073598 | 1.06710864697714 |
| chr17 | 20238661 | 20238736 | + | 14.0554480762271 | 25.0139063254164 | 2.38645642073598 | 1.06710864697714 |
| chr17 | 20999624 | 20999671 | - | 9.03564519186028 | 7.95897019445067 | 1.19322821036799 | 7.46976052883995 |
| chr17 | 20999838 | 20999871 | - | 12.0475269224804 | 13.6439489047726 | 8.35259747257593 | 11.7381951167485 |
| chr17 | 20999916 | 20999930 | - | 5.01980288436682 | 6.82197445238629 | 1.19322821036799 | 8.53686917581709 |
| chr17 | 20999955 | 20999995 | - | 3.01188173062009 | 2.27399148412876 | 1.19322821036799 | 2.13421729395427 |
| chr17 | 21000310 | 21000367 | - | 5.01980288436682 | 5.68497871032191 | 7.15936926220794 | 6.40265188186281 |
| chr17 | 21000421 | 21000517 | - | 30.1188173062009 | 22.7399148412876 | 25.0577924177278 | 26.6777161744284 |
| chr17 | 21000556 | 21000589 | - | 17.0673298068472 | 2.27399148412876 | 3.57968463110397 | 4.26843458790854 |
| chr17 | 21000606 | 21000791 | - | 54.2138711511617 | 46.6168254246397 | 29.8307052591997 | 41.6172372321083 |
| chr17 | 21000880 | 21000884 | - | 6.02376346124019 | 6.82197445238629 | 4.77291284147196 | 4.26843458790854 |
| chr17 | 21001067 | 21001109 | - | 37.1465413443145 | 21.6029190992233 | 66.8207797806074 | 68.2949534065367 |
| chr17 | 21001133 | 21001165 | - | 8.03168461498692 | 6.82197445238629 | 9.54582568294392 | 27.7448248214055 |
| chr17 | 21001223 | 21001275 | - | 38.1505019211879 | 38.657855230189 | 52.5020412561915 | 14.9395210576799 |
| chr17 | 21001323 | 21001367 | - | 24.0950538449607 | 7.95897019445067 | 45.3426719939836 | 65.0936274656053 |
| chr17 | 21001397 | 21001502 | - | 18.0712903837206 | 10.2329616785794 | 29.8307052591997 | 22.4092815865198 |
| chr17 | 21001531 | 21001546 | - | 15.0594086531005 | 7.95897019445067 | 11.9322821036799 | 17.0737383516342 |
| chr17 | 21001566 | 21001646 | - | 19.0752509605939 | 13.6439489047726 | 11.9322821036799 | 22.4092815865198 |
| chr17 | 21001692 | 21001946 | - | 62.2455557661486 | 83.0006891706999 | 87.1056593568632 | 59.7580842307196 |
| chr17 | 21001967 | 21002038 | - | 51.2019894205416 | 38.657855230189 | 48.9223566250876 | 11.7381951167485 |
| chr17 | 21002113 | 21002177 | - | 20.0792115374673 | 18.1919318730301 | 23.8645642073598 | 16.006629704657 |
| chr17 | 21002246 | 21002270 | - | 12.0475269224804 | 4.54798296825753 | 4.77291284147196 | 1.06710864697714 |
| chr17 | 21002372 | 21002441 | - | 3.01188173062009 | 5.68497871032191 | 7.15936926220794 | 8.53686917581709 |
| chr17 | 21002543 | 21002584 | - | 20.0792115374673 | 4.54798296825753 | 4.77291284147196 | 7.46976052883995 |
| chr17 | 21002636 | 21002662 | - | 1.00396057687336 | 4.54798296825753 | 2.38645642073598 | 1.06710864697714 |
| chr17 | 21002728 | 21002756 | - | 2.00792115374673 | 1.13699574206438 | 2.38645642073598 | 6.40265188186281 |
| chr17 | 21002787 | 21002923 | - | 60.2376346124019 | 39.7948509722534 | 28.6374770488317 | 13.8724124107028 |
| chr17 | 21191159 | 21191217 | + | 27.1069355755808 | 42.0688424563821 | 15.5119667347839 | 38.4159112911769 |
| chr17 | 21191244 | 21191265 | + | 18.0712903837206 | 21.6029190992233 | 9.54582568294392 | 12.8053037637256 |
| chr17 | 21191320 | 21191348 | + | 7.02772403811355 | 7.95897019445067 | 3.57968463110397 | 7.46976052883995 |
| chr17 | 21197983 | 21197999 | - | 24.0950538449607 | 35.2468680039958 | 14.3187385244159 | 10.6710864697714 |
| chr17 | 21198020 | 21198072 | - | 166.657455760979 | 234.221122865263 | 58.4681823080315 | 72.5633879944452 |
| chr17 | 21198110 | 21198120 | - | 8.03168461498692 | 5.68497871032191 | 2.38645642073598 | 5.33554323488568 |
| chr17 | 21198223 | 21198232 | - | 4.01584230749346 | 2.27399148412876 | 1.19322821036799 | 3.20132594093141 |
| chr17 | 21214319 | 21214371 | + | 22.087132691214 | 20.4659233571589 | 9.54582568294392 | 17.0737383516342 |
| chr17 | 21315163 | 21315202 | + | 7.02772403811355 | 7.95897019445067 | 1.19322821036799 | 1.06710864697714 |
| chr17 | 27307009 | 27307029 | + | 3.01188173062009 | 1.13699574206438 | 1.19322821036799 | 9.60397782279422 |
| chr17 | 27307906 | 27307941 | + | 4.01584230749346 | 7.95897019445067 | 35.7968463110397 | 34.1474767032683 |
| chr17 | 27308041 | 27308043 | + | 1.00396057687336 | 1.13699574206438 | 2.38645642073598 | 2.13421729395427 |
| chr17 | 27308226 | 27308247 | + | 1.00396057687336 | 1.13699574206438 | 4.77291284147196 | 4.26843458790854 |
| chr17 | 27308719 | 27308751 | + | 4.01584230749346 | 4.54798296825753 | 16.7051949451519 | 11.7381951167485 |
| chr17 | 27312207 | 27312559 | + | 58.2297134586551 | 38.657855230189 | 163.472264820415 | 257.17318392149 |
| chr17 | 27312590 | 27312600 | + | 2.00792115374673 | 1.13699574206438 | 14.3187385244159 | 5.33554323488568 |
| chr17 | 27312621 | 27312818 | + | 59.2336740355285 | 79.5897019445067 | 381.833027317757 | 255.038966627535 |
| chr17 | 27313054 | 27313250 | + | 619.443675930866 | 847.061827837965 | 2053.54575004331 | 3030.58855741507 |
| chr17 | 27313309 | 27313376 | + | 4.01584230749346 | 3.41098722619315 | 8.35259747257593 | 2.13421729395427 |
| chr17 | 27313407 | 27313414 | + | 12.0475269224804 | 20.4659233571589 | 13.1255103140479 | 22.4092815865198 |
| chr17 | 27313469 | 27313600 | + | 11.043566345607 | 15.9179403889013 | 25.0577924177278 | 26.6777161744284 |
| chr17 | 27333299 | 27333318 | + | 18.0712903837206 | 28.4248935516096 | 14.3187385244159 | 24.5434988804741 |
| chr17 | 27626105 | 27626139 | + | 3.01188173062009 | 11.3699574206438 | 7.15936926220794 | 18.1408469986113 |
| chr17 | 28195151 | 28195169 | + | 3.01188173062009 | 2.27399148412876 | 2.38645642073598 | 8.53686917581709 |
| chr17 | 28196149 | 28196151 | + | 2.00792115374673 | 4.54798296825753 | 1.19322821036799 | 4.26843458790854 |
| chr17 | 28196350 | 28196354 | + | 2.00792115374673 | 4.54798296825753 | 4.77291284147196 | 8.53686917581709 |
| chr17 | 28326744 | 28326886 | + | 36.1425807674411 | 20.4659233571589 | 32.2171616799357 | 22.4092815865198 |
| chr17 | 28326912 | 28326959 | + | 12.0475269224804 | 3.41098722619315 | 8.35259747257593 | 11.7381951167485 |
| chr17 | 28327051 | 28327065 | + | 1.00396057687336 | 7.95897019445067 | 2.38645642073598 | 4.26843458790854 |
| chr17 | 28327093 | 28327124 | + | 15.0594086531005 | 11.3699574206438 | 13.1255103140479 | 38.4159112911769 |
| chr17 | 28327179 | 28327185 | + | 5.01980288436682 | 10.2329616785794 | 2.38645642073598 | 3.20132594093141 |
| chr17 | 28327321 | 28327378 | + | 17.0673298068472 | 17.0549361309657 | 15.5119667347839 | 16.006629704657 |
| chr17 | 28327389 | 28327432 | + | 2.00792115374673 | 9.09596593651506 | 1.19322821036799 | 5.33554323488568 |
| chr17 | 28327540 | 28327589 | + | 9.03564519186028 | 9.09596593651506 | 11.9322821036799 | 13.8724124107028 |
| chr17 | 28327640 | 28327643 | + | 4.01584230749346 | 1.13699574206438 | 3.57968463110397 | 3.20132594093141 |
| chr17 | 28327651 | 28327802 | + | 82.3247673036159 | 90.9596593651506 | 53.6952694665595 | 103.509538756782 |
| chr17 | 28327830 | 28327857 | + | 3.01188173062009 | 11.3699574206438 | 14.3187385244159 | 12.8053037637256 |
| chr17 | 28327875 | 28327947 | + | 38.1505019211879 | 40.9318467143178 | 22.6713359969918 | 36.2816939972226 |
| chr17 | 28327949 | 28327956 | + | 7.02772403811355 | 4.54798296825753 | 3.57968463110397 | 2.13421729395427 |
| chr17 | 28327978 | 28328197 | + | 857.382332649853 | 803.855989639518 | 696.845274854906 | 675.479773536527 |
| chr17 | 28328349 | 28328364 | - | 23.0910932680874 | 31.8358807778027 | 7.15936926220794 | 7.46976052883995 |
| chr17 | 28328442 | 28328454 | - | 4.01584230749346 | 1.13699574206438 | 1.19322821036799 | 2.13421729395427 |
| chr17 | 28328487 | 28328542 | - | 147.582204800385 | 173.96034853585 | 63.2410951495034 | 118.449059814462 |
| chr17 | 28328622 | 28328632 | - | 14.0554480762271 | 22.7399148412876 | 5.96614105183995 | 7.46976052883995 |
| chr17 | 28328694 | 28328736 | - | 22.087132691214 | 32.9728765198671 | 13.1255103140479 | 21.3421729395427 |
| chr17 | 28344620 | 28344627 | + | 6.02376346124019 | 3.41098722619315 | 1.19322821036799 | 3.20132594093141 |
| chr17 | 28344711 | 28344712 | + | 2.00792115374673 | 1.13699574206438 | 1.19322821036799 | 1.06710864697714 |
| chr17 | 28345858 | 28345879 | + | 18.0712903837206 | 17.0549361309657 | 7.15936926220794 | 1.06710864697714 |
| chr17 | 28345959 | 28345966 | + | 29.1148567293276 | 6.82197445238629 | 2.38645642073598 | 3.20132594093141 |
| chr17 | 28346542 | 28346620 | + | 34.1346596136944 | 37.5208594881246 | 3.57968463110397 | 8.53686917581709 |
| chr17 | 28346657 | 28346730 | + | 43.1703048055547 | 27.2878978095452 | 4.77291284147196 | 13.8724124107028 |
| chr17 | 28346753 | 28346784 | + | 313.23569998449 | 230.81013563907 | 17.8984231555198 | 46.952780466994 |
| chr17 | 28346893 | 28346911 | + | 3.01188173062009 | 7.95897019445067 | 1.19322821036799 | 5.33554323488568 |
| chr17 | 28347216 | 28347245 | - | 7.02772403811355 | 13.6439489047726 | 4.77291284147196 | 7.46976052883995 |
| chr17 | 28347270 | 28347299 | - | 3.01188173062009 | 2.27399148412876 | 2.38645642073598 | 8.53686917581709 |
| chr17 | 28347339 | 28347374 | - | 4.01584230749346 | 9.09596593651506 | 2.38645642073598 | 5.33554323488568 |
| chr17 | 28347430 | 28347469 | - | 11.043566345607 | 4.54798296825753 | 1.19322821036799 | 2.13421729395427 |
| chr17 | 28347974 | 28348017 | - | 4.01584230749346 | 6.82197445238629 | 3.57968463110397 | 2.13421729395427 |
| chr17 | 28357628 | 28357738 | + | 996.932852835251 | 1275.70922259624 | 337.683583534141 | 959.330673632445 |
| chr17 | 28357783 | 28357803 | + | 2.00792115374673 | 2.27399148412876 | 1.19322821036799 | 2.13421729395427 |
| chr17 | 28360543 | 28360563 | + | 4.01584230749346 | 4.54798296825753 | 2.38645642073598 | 2.13421729395427 |
| chr17 | 28361288 | 28361307 | + | 18.0712903837206 | 18.1919318730301 | 13.1255103140479 | 20.2750642925656 |
| chr17 | 28361395 | 28361399 | + | 5.01980288436682 | 11.3699574206438 | 1.19322821036799 | 6.40265188186281 |
| chr17 | 28361422 | 28361450 | + | 8.03168461498692 | 13.6439489047726 | 2.38645642073598 | 12.8053037637256 |
| chr17 | 28399159 | 28399161 | - | 13.0514874993537 | 11.3699574206438 | 1.19322821036799 | 1.06710864697714 |
| chr17 | 28546734 | 28546790 | - | 36.1425807674411 | 60.2607743294122 | 28.6374770488317 | 27.7448248214055 |
| chr17 | 28553407 | 28553444 | - | 25.0990144218341 | 23.876910583352 | 19.0916513658878 | 38.4159112911769 |
| chr17 | 28553466 | 28553505 | - | 8.03168461498692 | 10.2329616785794 | 13.1255103140479 | 20.2750642925656 |
| chr17 | 28553530 | 28553572 | - | 7.02772403811355 | 10.2329616785794 | 2.38645642073598 | 2.13421729395427 |
| chr17 | 28553686 | 28553701 | - | 3.01188173062009 | 1.13699574206438 | 2.38645642073598 | 5.33554323488568 |
| chr17 | 28553863 | 28553890 | - | 2.00792115374673 | 3.41098722619315 | 11.9322821036799 | 16.006629704657 |
| chr17 | 28553937 | 28553950 | - | 1.00396057687336 | 1.13699574206438 | 4.77291284147196 | 7.46976052883995 |
| chr17 | 28553983 | 28554010 | - | 5.01980288436682 | 2.27399148412876 | 11.9322821036799 | 39.483019938154 |
| chr17 | 28554121 | 28554222 | - | 10.0396057687336 | 10.2329616785794 | 23.8645642073598 | 39.483019938154 |
| chr17 | 28577587 | 28577634 | - | 14.0554480762271 | 13.6439489047726 | 4.77291284147196 | 9.60397782279422 |
| chr17 | 28599131 | 28599152 | + | 6.02376346124019 | 2.27399148412876 | 3.57968463110397 | 8.53686917581709 |
| chr17 | 28614485 | 28614541 | - | 22.087132691214 | 26.1509020674808 | 4.77291284147196 | 22.4092815865198 |
| chr17 | 28614755 | 28614795 | - | 16.0633692299738 | 26.1509020674808 | 16.7051949451519 | 18.1408469986113 |
| chr17 | 28614988 | 28614993 | - | 2.00792115374673 | 1.13699574206438 | 2.38645642073598 | 1.06710864697714 |
| chr17 | 28648622 | 28648714 | - | 18.0712903837206 | 31.8358807778027 | 8.35259747257593 | 16.006629704657 |
| chr17 | 28648751 | 28648792 | - | 20.0792115374673 | 15.9179403889013 | 2.38645642073598 | 20.2750642925656 |
| chr17 | 28701839 | 28701912 | + | 12.0475269224804 | 7.95897019445067 | 13.1255103140479 | 27.7448248214055 |
| chr17 | 28702001 | 28702072 | + | 111.439624032943 | 76.1787147183136 | 63.2410951495034 | 100.308212815851 |
| chr17 | 28702102 | 28702156 | + | 38.1505019211879 | 20.4659233571589 | 7.15936926220794 | 26.6777161744284 |
| chr17 | 28703953 | 28703975 | - | 3.01188173062009 | 2.27399148412876 | 5.96614105183995 | 13.8724124107028 |
| chr17 | 28723429 | 28723430 | + | 1.00396057687336 | 2.27399148412876 | 2.38645642073598 | 2.13421729395427 |
| chr17 | 28724200 | 28724254 | + | 12.0475269224804 | 11.3699574206438 | 4.77291284147196 | 9.60397782279422 |
| chr17 | 28749560 | 28749652 | + | 35.1386201905678 | 28.4248935516096 | 29.8307052591997 | 41.6172372321083 |
| chr17 | 28749831 | 28749834 | + | 4.01584230749346 | 5.68497871032191 | 2.38645642073598 | 4.26843458790854 |
| chr17 | 28749865 | 28749888 | + | 3.01188173062009 | 6.82197445238629 | 2.38645642073598 | 5.33554323488568 |
| chr17 | 28749913 | 28749940 | + | 5.01980288436682 | 15.9179403889013 | 2.38645642073598 | 4.26843458790854 |
| chr17 | 28750037 | 28750045 | + | 39.1544624980612 | 145.535454984241 | 23.8645642073598 | 45.8856718200168 |
| chr17 | 28757616 | 28757619 | - | 3.01188173062009 | 1.13699574206438 | 2.38645642073598 | 5.33554323488568 |
| chr17 | 28860823 | 28860931 | + | 22.087132691214 | 12.5069531627082 | 9.54582568294392 | 14.9395210576799 |
| chr17 | 28860958 | 28860990 | + | 12.0475269224804 | 7.95897019445067 | 4.77291284147196 | 11.7381951167485 |
| chr17 | 28879362 | 28879425 | - | 23.0910932680874 | 17.0549361309657 | 15.5119667347839 | 19.2079556455884 |
| chr17 | 28879861 | 28879895 | - | 2.00792115374673 | 2.27399148412876 | 2.38645642073598 | 14.9395210576799 |
| chr17 | 28879997 | 28880043 | - | 3.01188173062009 | 3.41098722619315 | 34.6036181006717 | 28.8119334683827 |
| chr17 | 28880107 | 28880136 | - | 7.02772403811355 | 2.27399148412876 | 8.35259747257593 | 8.53686917581709 |
| chr17 | 28880190 | 28880218 | - | 3.01188173062009 | 3.41098722619315 | 8.35259747257593 | 17.0737383516342 |
| chr17 | 28880258 | 28880273 | - | 1.00396057687336 | 2.27399148412876 | 4.77291284147196 | 7.46976052883995 |
| chr17 | 28880302 | 28880371 | - | 21.0831721143407 | 14.780944646837 | 35.7968463110397 | 20.2750642925656 |
| chr17 | 28880404 | 28880486 | - | 18.0712903837206 | 14.780944646837 | 28.6374770488317 | 32.0132594093141 |
| chr17 | 28880513 | 28880526 | - | 1.00396057687336 | 1.13699574206438 | 2.38645642073598 | 9.60397782279422 |
| chr17 | 28880548 | 28880588 | - | 10.0396057687336 | 6.82197445238629 | 11.9322821036799 | 10.6710864697714 |
| chr17 | 28897822 | 28897840 | - | 14.0554480762271 | 11.3699574206438 | 7.15936926220794 | 4.26843458790854 |
| chr17 | 28906370 | 28906372 | - | 1.00396057687336 | 1.13699574206438 | 1.19322821036799 | 1.06710864697714 |
| chr17 | 29073982 | 29074038 | - | 1.00396057687336 | 1.13699574206438 | 1.19322821036799 | 2.13421729395427 |
| chr17 | 29256015 | 29256032 | - | 1.00396057687336 | 2.27399148412876 | 1.19322821036799 | 4.26843458790854 |
| chr17 | 29256185 | 29256223 | - | 8.03168461498692 | 18.1919318730301 | 2.38645642073598 | 14.9395210576799 |
| chr17 | 29256245 | 29256256 | - | 5.01980288436682 | 2.27399148412876 | 2.38645642073598 | 6.40265188186281 |
| chr17 | 29257894 | 29257935 | - | 8.03168461498692 | 18.1919318730301 | 9.54582568294392 | 16.006629704657 |
| chr17 | 29258937 | 29258951 | - | 8.03168461498692 | 3.41098722619315 | 1.19322821036799 | 8.53686917581709 |
| chr17 | 29258992 | 29259012 | - | 6.02376346124019 | 1.13699574206438 | 3.57968463110397 | 2.13421729395427 |
| chr17 | 29259097 | 29259130 | - | 9.03564519186028 | 4.54798296825753 | 3.57968463110397 | 5.33554323488568 |
| chr17 | 29259180 | 29259190 | - | 4.01584230749346 | 2.27399148412876 | 3.57968463110397 | 4.26843458790854 |
| chr17 | 29259302 | 29259318 | - | 17.0673298068472 | 39.7948509722534 | 9.54582568294392 | 12.8053037637256 |
| chr17 | 29260893 | 29260925 | - | 4.01584230749346 | 7.95897019445067 | 10.7390538933119 | 13.8724124107028 |
| chr17 | 29261473 | 29261475 | - | 13.0514874993537 | 3.41098722619315 | 1.19322821036799 | 5.33554323488568 |
| chr17 | 29262431 | 29262436 | - | 4.01584230749346 | 3.41098722619315 | 2.38645642073598 | 2.13421729395427 |
| chr17 | 29262490 | 29262499 | - | 9.03564519186028 | 5.68497871032191 | 4.77291284147196 | 2.13421729395427 |
| chr17 | 29262685 | 29262702 | - | 2.00792115374673 | 4.54798296825753 | 1.19322821036799 | 9.60397782279422 |
| chr17 | 29262920 | 29262955 | - | 9.03564519186028 | 10.2329616785794 | 8.35259747257593 | 3.20132594093141 |
| chr17 | 29262966 | 29263012 | - | 3.01188173062009 | 4.54798296825753 | 3.57968463110397 | 5.33554323488568 |
| chr17 | 29263502 | 29263504 | - | 3.01188173062009 | 1.13699574206438 | 1.19322821036799 | 1.06710864697714 |
| chr17 | 29263660 | 29263663 | - | 7.02772403811355 | 5.68497871032191 | 1.19322821036799 | 1.06710864697714 |
| chr17 | 29263853 | 29263875 | - | 48.1901076899215 | 72.7677274921204 | 20.2848795762558 | 11.7381951167485 |
| chr17 | 29263910 | 29263929 | - | 10.0396057687336 | 14.780944646837 | 2.38645642073598 | 6.40265188186281 |
| chr17 | 29264282 | 29264335 | - | 15.0594086531005 | 13.6439489047726 | 8.35259747257593 | 34.1474767032683 |
| chr17 | 29264362 | 29264363 | - | 2.00792115374673 | 1.13699574206438 | 1.19322821036799 | 1.06710864697714 |
| chr17 | 29264418 | 29264452 | - | 6.02376346124019 | 9.09596593651506 | 4.77291284147196 | 14.9395210576799 |
| chr17 | 29264473 | 29264488 | - | 10.0396057687336 | 19.3289276150945 | 11.9322821036799 | 18.1408469986113 |
| chr17 | 29543786 | 29543803 | + | 9.03564519186028 | 6.82197445238629 | 2.38645642073598 | 2.13421729395427 |
| chr17 | 29543861 | 29543908 | + | 20.0792115374673 | 20.4659233571589 | 4.77291284147196 | 2.13421729395427 |
| chr17 | 29543930 | 29543945 | + | 1.00396057687336 | 2.27399148412876 | 3.57968463110397 | 2.13421729395427 |
| chr17 | 29547355 | 29547365 | + | 18.0712903837206 | 15.9179403889013 | 7.15936926220794 | 8.53686917581709 |
| chr17 | 29548166 | 29548172 | + | 1.00396057687336 | 4.54798296825753 | 2.38645642073598 | 5.33554323488568 |
| chr17 | 29548314 | 29548336 | + | 8.03168461498692 | 11.3699574206438 | 7.15936926220794 | 5.33554323488568 |
| chr17 | 29548394 | 29548402 | + | 1.00396057687336 | 3.41098722619315 | 7.15936926220794 | 59.7580842307196 |
| chr17 | 29549644 | 29549651 | + | 4.01584230749346 | 4.54798296825753 | 1.19322821036799 | 2.13421729395427 |
| chr17 | 29549686 | 29549690 | + | 10.0396057687336 | 2.27399148412876 | 5.96614105183995 | 1.06710864697714 |
| chr17 | 29550915 | 29550935 | + | 4.01584230749346 | 4.54798296825753 | 1.19322821036799 | 8.53686917581709 |
| chr17 | 29551563 | 29551575 | + | 6.02376346124019 | 6.82197445238629 | 2.38645642073598 | 3.20132594093141 |
| chr17 | 29573514 | 29573560 | - | 19.0752509605939 | 19.3289276150945 | 4.77291284147196 | 5.33554323488568 |
| chr17 | 29573815 | 29573817 | - | 4.01584230749346 | 2.27399148412876 | 2.38645642073598 | 2.13421729395427 |
| chr17 | 29574149 | 29574172 | - | 24.0950538449607 | 7.95897019445067 | 9.54582568294392 | 35.2145853502455 |
| chr17 | 29574266 | 29574307 | - | 13.0514874993537 | 7.95897019445067 | 9.54582568294392 | 5.33554323488568 |
| chr17 | 29614206 | 29614228 | + | 4.01584230749346 | 2.27399148412876 | 4.77291284147196 | 12.8053037637256 |
| chr17 | 30118148 | 30118174 | + | 6.02376346124019 | 10.2329616785794 | 5.96614105183995 | 12.8053037637256 |
| chr17 | 30248224 | 30248250 | - | 9.03564519186028 | 7.95897019445067 | 7.15936926220794 | 21.3421729395427 |
| chr17 | 30248271 | 30248344 | - | 103.407939417957 | 78.4527062024424 | 65.6275515702394 | 227.29414180613 |
| chr17 | 30248380 | 30248453 | - | 23.0910932680874 | 9.09596593651506 | 8.35259747257593 | 36.2816939972226 |
| chr17 | 30248606 | 30248625 | - | 31.1227778830743 | 32.9728765198671 | 13.1255103140479 | 88.5700176991023 |
| chr17 | 30248651 | 30248694 | - | 12.0475269224804 | 12.5069531627082 | 21.4781077866238 | 65.0936274656053 |
| chr17 | 30248742 | 30248778 | - | 3.01188173062009 | 3.41098722619315 | 19.0916513658878 | 12.8053037637256 |
| chr17 | 30248807 | 30248825 | - | 3.01188173062009 | 7.95897019445067 | 27.4442488384638 | 37.3488026441998 |
| chr17 | 30248855 | 30248883 | - | 2.00792115374673 | 1.13699574206438 | 7.15936926220794 | 16.006629704657 |
| chr17 | 30248929 | 30248967 | - | 3.01188173062009 | 2.27399148412876 | 3.57968463110397 | 13.8724124107028 |
| chr17 | 30464819 | 30464976 | + | 10.0396057687336 | 9.09596593651506 | 8.35259747257593 | 7.46976052883995 |
| chr17 | 30465025 | 30465152 | + | 28.1108961524542 | 29.5618892936739 | 17.8984231555198 | 23.476390233497 |
| chr17 | 30465245 | 30465373 | + | 32.1267384599477 | 28.4248935516096 | 8.35259747257593 | 19.2079556455884 |
| chr17 | 30465404 | 30465522 | + | 5.01980288436682 | 10.2329616785794 | 10.7390538933119 | 5.33554323488568 |
| chr17 | 30465870 | 30465950 | + | 32.1267384599477 | 20.4659233571589 | 11.9322821036799 | 12.8053037637256 |
| chr17 | 30466089 | 30466126 | + | 2.00792115374673 | 1.13699574206438 | 1.19322821036799 | 4.26843458790854 |
| chr17 | 30466323 | 30466386 | + | 4.01584230749346 | 4.54798296825753 | 1.19322821036799 | 1.06710864697714 |
| chr17 | 30466411 | 30466491 | + | 1.00396057687336 | 2.27399148412876 | 1.19322821036799 | 3.20132594093141 |
| chr17 | 30466606 | 30466622 | + | 4.01584230749346 | 2.27399148412876 | 2.38645642073598 | 5.33554323488568 |
| chr17 | 30466673 | 30466685 | + | 2.00792115374673 | 1.13699574206438 | 1.19322821036799 | 1.06710864697714 |
| chr17 | 30467118 | 30467154 | + | 2.00792115374673 | 3.41098722619315 | 1.19322821036799 | 5.33554323488568 |
| chr17 | 30467262 | 30467289 | + | 6.02376346124019 | 7.95897019445067 | 1.19322821036799 | 1.06710864697714 |
| chr17 | 30467329 | 30467406 | + | 6.02376346124019 | 1.13699574206438 | 3.57968463110397 | 6.40265188186281 |
| chr17 | 30467442 | 30467593 | + | 13.0514874993537 | 11.3699574206438 | 9.54582568294392 | 6.40265188186281 |
| chr17 | 30467618 | 30467697 | + | 7.02772403811355 | 3.41098722619315 | 1.19322821036799 | 2.13421729395427 |
| chr17 | 30469035 | 30469086 | + | 7.02772403811355 | 9.09596593651506 | 4.77291284147196 | 5.33554323488568 |
| chr17 | 30522446 | 30522465 | + | 5.01980288436682 | 6.82197445238629 | 10.7390538933119 | 13.8724124107028 |
| chr17 | 30522511 | 30522569 | + | 8.03168461498692 | 6.82197445238629 | 7.15936926220794 | 4.26843458790854 |
| chr17 | 30524911 | 30524913 | + | 1.00396057687336 | 3.41098722619315 | 1.19322821036799 | 2.13421729395427 |
| chr17 | 30526562 | 30526569 | + | 4.01584230749346 | 5.68497871032191 | 2.38645642073598 | 4.26843458790854 |
| chr17 | 30782712 | 30782718 | - | 1.00396057687336 | 4.54798296825753 | 1.19322821036799 | 4.26843458790854 |
| chr17 | 30783340 | 30783359 | - | 1.00396057687336 | 9.09596593651506 | 7.15936926220794 | 12.8053037637256 |
| chr17 | 30895050 | 30895051 | + | 1.00396057687336 | 2.27399148412876 | 2.38645642073598 | 4.26843458790854 |
| chr17 | 31374194 | 31374208 | + | 3.01188173062009 | 2.27399148412876 | 1.19322821036799 | 1.06710864697714 |
| chr17 | 31374420 | 31374422 | + | 2.00792115374673 | 1.13699574206438 | 1.19322821036799 | 2.13421729395427 |
| chr17 | 31374740 | 31374770 | + | 13.0514874993537 | 21.6029190992233 | 8.35259747257593 | 1.06710864697714 |
| chr17 | 31375611 | 31375658 | + | 17.0673298068472 | 12.5069531627082 | 2.38645642073598 | 28.8119334683827 |
| chr17 | 31377557 | 31377564 | + | 6.02376346124019 | 4.54798296825753 | 1.19322821036799 | 2.13421729395427 |
| chr17 | 31851903 | 31851980 | - | 35.1386201905678 | 45.4798296825753 | 14.3187385244159 | 16.006629704657 |
| chr17 | 31852070 | 31852082 | - | 3.01188173062009 | 5.68497871032191 | 1.19322821036799 | 1.06710864697714 |
| chr17 | 31861015 | 31861024 | - | 3.01188173062009 | 5.68497871032191 | 5.96614105183995 | 4.26843458790854 |
| chr17 | 31861081 | 31861100 | - | 2.00792115374673 | 3.41098722619315 | 9.54582568294392 | 4.26843458790854 |
| chr17 | 31861190 | 31861194 | - | 10.0396057687336 | 4.54798296825753 | 3.57968463110397 | 9.60397782279422 |
| chr17 | 31863236 | 31863262 | - | 5.01980288436682 | 13.6439489047726 | 8.35259747257593 | 21.3421729395427 |
| chr17 | 31999338 | 31999469 | + | 58.2297134586551 | 70.4937360079917 | 45.3426719939836 | 72.5633879944452 |
| chr17 | 31999632 | 31999635 | + | 2.00792115374673 | 2.27399148412876 | 5.96614105183995 | 1.06710864697714 |
| chr17 | 31999650 | 31999771 | + | 71.2812009580089 | 70.4937360079917 | 70.4004644117114 | 73.6304966414224 |
| chr17 | 31999854 | 31999863 | + | 3.01188173062009 | 2.27399148412876 | 2.38645642073598 | 1.06710864697714 |
| chr17 | 31999878 | 31999968 | + | 147.582204800385 | 193.289276150945 | 122.902505667903 | 170.737383516342 |
| chr17 | 31999995 | 32000118 | + | 25.0990144218341 | 31.8358807778027 | 19.0916513658878 | 46.952780466994 |
| chr17 | 32000139 | 32000154 | + | 163.645574030358 | 183.056314472366 | 63.2410951495034 | 147.260993282845 |
| chr17 | 32000308 | 32000311 | + | 2.00792115374673 | 1.13699574206438 | 2.38645642073598 | 1.06710864697714 |
| chr17 | 32000488 | 32000492 | + | 2.00792115374673 | 4.54798296825753 | 2.38645642073598 | 10.6710864697714 |
| chr17 | 32000544 | 32000620 | + | 9.03564519186028 | 21.6029190992233 | 5.96614105183995 | 13.8724124107028 |
| chr17 | 32000656 | 32000674 | + | 16.0633692299738 | 35.2468680039958 | 7.15936926220794 | 17.0737383516342 |
| chr17 | 32225089 | 32225122 | + | 2.00792115374673 | 3.41098722619315 | 5.96614105183995 | 3.20132594093141 |
| chr17 | 32225320 | 32225424 | + | 99.3920971104631 | 110.288586980245 | 72.7869208324474 | 118.449059814462 |
| chr17 | 32225533 | 32225544 | + | 4.01584230749346 | 3.41098722619315 | 10.7390538933119 | 7.46976052883995 |
| chr17 | 32225565 | 32225628 | + | 31.1227778830743 | 36.3838637460602 | 52.5020412561915 | 34.1474767032683 |
| chr17 | 32225648 | 32225683 | + | 8.03168461498692 | 14.780944646837 | 11.9322821036799 | 3.20132594093141 |
| chr17 | 32322504 | 32322506 | + | 22.087132691214 | 15.9179403889013 | 7.15936926220794 | 7.46976052883995 |
| chr17 | 32350118 | 32350228 | + | 2747.8400989024 | 2293.32041174386 | 1557.16281453023 | 1942.13773749839 |
| chr17 | 32350271 | 32350297 | + | 11.043566345607 | 2.27399148412876 | 9.54582568294392 | 3.20132594093141 |
| chr17 | 32350321 | 32350327 | + | 6.02376346124019 | 4.54798296825753 | 8.35259747257593 | 1.06710864697714 |
| chr17 | 32351787 | 32351813 | + | 7.02772403811355 | 6.82197445238629 | 8.35259747257593 | 5.33554323488568 |
| chr17 | 32358544 | 32358562 | + | 8.03168461498692 | 4.54798296825753 | 7.15936926220794 | 10.6710864697714 |
| chr17 | 32369887 | 32369957 | + | 6.02376346124019 | 7.95897019445067 | 8.35259747257593 | 26.6777161744284 |
| chr17 | 32370040 | 32370049 | + | 2.00792115374673 | 2.27399148412876 | 3.57968463110397 | 2.13421729395427 |
| chr17 | 32370160 | 32370180 | + | 7.02772403811355 | 6.82197445238629 | 4.77291284147196 | 10.6710864697714 |
| chr17 | 32370253 | 32370314 | + | 11.043566345607 | 23.876910583352 | 2.38645642073598 | 4.26843458790854 |
| chr17 | 32372343 | 32372346 | + | 2.00792115374673 | 3.41098722619315 | 2.38645642073598 | 4.26843458790854 |
| chr17 | 32378281 | 32378302 | + | 10.0396057687336 | 7.95897019445067 | 1.19322821036799 | 5.33554323488568 |
| chr17 | 32378323 | 32378329 | + | 1.00396057687336 | 3.41098722619315 | 7.15936926220794 | 17.0737383516342 |
| chr17 | 32381833 | 32381863 | + | 1.00396057687336 | 2.27399148412876 | 1.19322821036799 | 2.13421729395427 |
| chr17 | 32480656 | 32480685 | + | 14.0554480762271 | 9.09596593651506 | 5.96614105183995 | 32.0132594093141 |
| chr17 | 32480751 | 32480908 | + | 1297.11706532039 | 1027.8441508262 | 603.773474446203 | 1052.16912591946 |
| chr17 | 32480934 | 32480957 | + | 233.922814411494 | 250.139063254164 | 210.008165024766 | 178.207144045182 |
| chr17 | 32481006 | 32481034 | + | 14.0554480762271 | 5.68497871032191 | 1.19322821036799 | 4.26843458790854 |
| chr17 | 32481055 | 32481182 | + | 81.3208067267425 | 46.6168254246397 | 54.8884976769275 | 60.8251928776967 |
| chr17 | 32481215 | 32481216 | + | 2.00792115374673 | 1.13699574206438 | 2.38645642073598 | 1.06710864697714 |
| chr17 | 32481241 | 32481272 | + | 6.02376346124019 | 6.82197445238629 | 1.19322821036799 | 2.13421729395427 |
| chr17 | 32481428 | 32481431 | + | 2.00792115374673 | 3.41098722619315 | 1.19322821036799 | 11.7381951167485 |
| chr17 | 32481455 | 32481459 | + | 1.00396057687336 | 1.13699574206438 | 2.38645642073598 | 11.7381951167485 |
| chr17 | 32481482 | 32481500 | + | 19.0752509605939 | 3.41098722619315 | 13.1255103140479 | 50.1541064079254 |
| chr17 | 32481527 | 32481548 | + | 10.0396057687336 | 4.54798296825753 | 2.38645642073598 | 11.7381951167485 |
| chr17 | 32481739 | 32481743 | + | 2.00792115374673 | 1.13699574206438 | 5.96614105183995 | 35.2145853502455 |
| chr17 | 32481776 | 32481799 | + | 328.29510863759 | 192.152280408881 | 195.68942650035 | 392.695982087586 |
| chr17 | 32481841 | 32481931 | + | 167.661416337852 | 133.028501821533 | 171.82486229299 | 232.629685041016 |
| chr17 | 32481953 | 32482034 | + | 409.615915364333 | 278.563956805774 | 215.974306076606 | 465.259370082031 |
| chr17 | 32482303 | 32482349 | + | 32.1267384599477 | 46.6168254246397 | 11.9322821036799 | 16.006629704657 |
| chr17 | 34962601 | 34962686 | + | 66.2613980736421 | 75.0417189762492 | 46.5359002043516 | 54.4225409958339 |
| chr17 | 35004596 | 35004605 | + | 8.03168461498692 | 7.95897019445067 | 1.19322821036799 | 2.13421729395427 |
| chr17 | 35004726 | 35004734 | + | 4.01584230749346 | 6.82197445238629 | 2.38645642073598 | 7.46976052883995 |
| chr17 | 35005877 | 35005901 | + | 8.03168461498692 | 6.82197445238629 | 2.38645642073598 | 6.40265188186281 |
| chr17 | 35009154 | 35009194 | - | 5.01980288436682 | 2.27399148412876 | 1.19322821036799 | 1.06710864697714 |
| chr17 | 35010468 | 35010494 | - | 5.01980288436682 | 5.68497871032191 | 7.15936926220794 | 6.40265188186281 |
| chr17 | 35011123 | 35011136 | - | 5.01980288436682 | 3.41098722619315 | 3.57968463110397 | 1.06710864697714 |
| chr17 | 35100571 | 35100573 | - | 5.01980288436682 | 7.95897019445067 | 1.19322821036799 | 6.40265188186281 |
| chr17 | 35131411 | 35131414 | - | 5.01980288436682 | 6.82197445238629 | 1.19322821036799 | 2.13421729395427 |
| chr17 | 35132158 | 35132161 | - | 1.00396057687336 | 1.13699574206438 | 1.19322821036799 | 4.26843458790854 |
| chr17 | 35568072 | 35568178 | + | 189.748549029066 | 206.933225055718 | 204.042023972926 | 439.64876255458 |
| chr17 | 35568442 | 35568470 | + | 8.03168461498692 | 7.95897019445067 | 9.54582568294392 | 13.8724124107028 |
| chr17 | 35568500 | 35568502 | + | 7.02772403811355 | 2.27399148412876 | 4.77291284147196 | 10.6710864697714 |
| chr17 | 35574240 | 35574281 | + | 2.00792115374673 | 1.13699574206438 | 2.38645642073598 | 5.33554323488568 |
| chr17 | 35575094 | 35575096 | - | 9.03564519186028 | 22.7399148412876 | 9.54582568294392 | 8.53686917581709 |
| chr17 | 35671859 | 35671878 | + | 1.00396057687336 | 1.13699574206438 | 2.38645642073598 | 1.06710864697714 |
| chr17 | 35674202 | 35674234 | + | 6.02376346124019 | 4.54798296825753 | 4.77291284147196 | 1.06710864697714 |
| chr17 | 35709221 | 35709264 | + | 4.01584230749346 | 4.54798296825753 | 5.96614105183995 | 5.33554323488568 |
| chr17 | 35710231 | 35710262 | + | 6.02376346124019 | 3.41098722619315 | 20.2848795762558 | 7.46976052883995 |
| chr17 | 35710301 | 35710321 | + | 5.01980288436682 | 2.27399148412876 | 9.54582568294392 | 9.60397782279422 |
| chr17 | 35717225 | 35717262 | + | 5.01980288436682 | 2.27399148412876 | 14.3187385244159 | 16.006629704657 |
| chr17 | 35717310 | 35717347 | + | 5.01980288436682 | 2.27399148412876 | 13.1255103140479 | 12.8053037637256 |
| chr17 | 35723697 | 35723802 | + | 26.1029749987075 | 28.4248935516096 | 53.6952694665595 | 62.959410171651 |
| chr17 | 35723835 | 35723865 | + | 16.0633692299738 | 12.5069531627082 | 13.1255103140479 | 20.2750642925656 |
| chr17 | 35725484 | 35725486 | + | 4.01584230749346 | 3.41098722619315 | 1.19322821036799 | 3.20132594093141 |
| chr17 | 35726282 | 35726296 | + | 23.0910932680874 | 13.6439489047726 | 7.15936926220794 | 19.2079556455884 |
| chr17 | 35726330 | 35726389 | + | 140.554480762271 | 175.097344277915 | 81.1395183050233 | 223.025707218221 |
| chr17 | 37611769 | 37611781 | - | 9.03564519186028 | 4.54798296825753 | 17.8984231555198 | 8.53686917581709 |
| chr17 | 38848031 | 38848034 | - | 1.00396057687336 | 2.27399148412876 | 2.38645642073598 | 2.13421729395427 |
| chr17 | 38848258 | 38848281 | - | 6.02376346124019 | 1.13699574206438 | 1.19322821036799 | 5.33554323488568 |
| chr17 | 38850088 | 38850123 | - | 7.02772403811355 | 12.5069531627082 | 5.96614105183995 | 2.13421729395427 |
| chr17 | 38850151 | 38850183 | - | 138.546559608524 | 97.7816338175368 | 66.8207797806074 | 172.871600810296 |
| chr17 | 38918815 | 38918855 | + | 13.0514874993537 | 5.68497871032191 | 2.38645642073598 | 2.13421729395427 |
| chr17 | 38918948 | 38918993 | + | 4.01584230749346 | 1.13699574206438 | 1.19322821036799 | 1.06710864697714 |
| chr17 | 38919174 | 38919178 | + | 2.00792115374673 | 1.13699574206438 | 1.19322821036799 | 5.33554323488568 |
| chr17 | 38919199 | 38919266 | + | 15.0594086531005 | 6.82197445238629 | 2.38645642073598 | 3.20132594093141 |
| chr17 | 38919291 | 38919311 | + | 15.0594086531005 | 4.54798296825753 | 5.96614105183995 | 4.26843458790854 |
| chr17 | 38919369 | 38919412 | + | 8.03168461498692 | 6.82197445238629 | 4.77291284147196 | 2.13421729395427 |
| chr17 | 38919444 | 38919535 | + | 18.0712903837206 | 11.3699574206438 | 8.35259747257593 | 9.60397782279422 |
| chr17 | 38919578 | 38919653 | + | 21.0831721143407 | 18.1919318730301 | 11.9322821036799 | 10.6710864697714 |
| chr17 | 38919676 | 38919730 | + | 14.0554480762271 | 9.09596593651506 | 14.3187385244159 | 12.8053037637256 |
| chr17 | 38919755 | 38919959 | + | 95.3762548029696 | 46.6168254246397 | 73.9801490428153 | 62.959410171651 |
| chr17 | 38919998 | 38920037 | + | 38.1505019211879 | 22.7399148412876 | 32.2171616799357 | 7.46976052883995 |
| chr17 | 38920105 | 38920147 | + | 19.0752509605939 | 5.68497871032191 | 10.7390538933119 | 7.46976052883995 |
| chr17 | 38920170 | 38920207 | + | 12.0475269224804 | 11.3699574206438 | 11.9322821036799 | 9.60397782279422 |
| chr17 | 38920232 | 38920286 | + | 23.0910932680874 | 5.68497871032191 | 9.54582568294392 | 10.6710864697714 |
| chr17 | 38920339 | 38920340 | + | 1.00396057687336 | 2.27399148412876 | 1.19322821036799 | 1.06710864697714 |
| chr17 | 38920351 | 38920400 | + | 23.0910932680874 | 9.09596593651506 | 17.8984231555198 | 6.40265188186281 |
| chr17 | 38920447 | 38920488 | + | 7.02772403811355 | 1.13699574206438 | 4.77291284147196 | 1.06710864697714 |
| chr17 | 38920551 | 38920608 | + | 33.130699036821 | 4.54798296825753 | 9.54582568294392 | 9.60397782279422 |
| chr17 | 38920637 | 38920710 | + | 115.455466340437 | 55.7127913611547 | 40.5697591525116 | 44.8185631730397 |
| chr17 | 38920752 | 38920828 | + | 63.249516343022 | 35.2468680039958 | 101.424397881279 | 49.0869977609482 |
| chr17 | 38920916 | 38920981 | + | 16.0633692299738 | 3.41098722619315 | 4.77291284147196 | 17.0737383516342 |
| chr17 | 38921042 | 38921093 | + | 20.0792115374673 | 7.95897019445067 | 16.7051949451519 | 34.1474767032683 |
| chr17 | 38921185 | 38921237 | + | 4.01584230749346 | 2.27399148412876 | 3.57968463110397 | 5.33554323488568 |
| chr17 | 38921319 | 38921346 | + | 1.00396057687336 | 1.13699574206438 | 1.19322821036799 | 7.46976052883995 |
| chr17 | 38921381 | 38921406 | + | 6.02376346124019 | 4.54798296825753 | 4.77291284147196 | 8.53686917581709 |
| chr17 | 38921552 | 38921688 | + | 28.1108961524542 | 28.4248935516096 | 13.1255103140479 | 27.7448248214055 |
| chr17 | 38921732 | 38921747 | + | 4.01584230749346 | 12.5069531627082 | 3.57968463110397 | 6.40265188186281 |
| chr17 | 39204630 | 39204657 | + | 286.128764408909 | 444.565335147173 | 262.510206280958 | 436.447436613649 |
| chr17 | 39204696 | 39204710 | + | 7.02772403811355 | 42.0688424563821 | 11.9322821036799 | 6.40265188186281 |
| chr17 | 39252859 | 39252862 | - | 5.01980288436682 | 3.41098722619315 | 1.19322821036799 | 4.26843458790854 |
| chr17 | 39253073 | 39253106 | - | 19.0752509605939 | 19.3289276150945 | 11.9322821036799 | 23.476390233497 |
| chr17 | 39253955 | 39253967 | - | 7.02772403811355 | 9.09596593651506 | 19.0916513658878 | 4.26843458790854 |
| chr17 | 39261143 | 39261164 | - | 11.043566345607 | 3.41098722619315 | 8.35259747257593 | 7.46976052883995 |
| chr17 | 39404344 | 39404345 | - | 1.00396057687336 | 4.54798296825753 | 2.38645642073598 | 6.40265188186281 |
| chr17 | 39404464 | 39404479 | - | 4.01584230749346 | 10.2329616785794 | 5.96614105183995 | 13.8724124107028 |
| chr17 | 39404568 | 39404575 | - | 3.01188173062009 | 6.82197445238629 | 1.19322821036799 | 6.40265188186281 |
| chr17 | 39404846 | 39404868 | - | 69.2732798042621 | 69.3567402659273 | 35.7968463110397 | 54.4225409958339 |
| chr17 | 39405200 | 39405212 | - | 8.03168461498692 | 15.9179403889013 | 4.77291284147196 | 4.26843458790854 |
| chr17 | 39406771 | 39406807 | - | 9.03564519186028 | 10.2329616785794 | 13.1255103140479 | 8.53686917581709 |
| chr17 | 39406902 | 39406924 | - | 30.1188173062009 | 56.8497871032191 | 23.8645642073598 | 48.0198891139711 |
| chr17 | 39407161 | 39407214 | - | 242.958459603354 | 285.38593125816 | 107.390538933119 | 178.207144045182 |
| chr17 | 39531655 | 39531680 | + | 2.00792115374673 | 1.13699574206438 | 2.38645642073598 | 6.40265188186281 |
| chr17 | 39531706 | 39531722 | + | 20.0792115374673 | 15.9179403889013 | 11.9322821036799 | 12.8053037637256 |
| chr17 | 39532027 | 39532080 | + | 5.01980288436682 | 1.13699574206438 | 11.9322821036799 | 23.476390233497 |
| chr17 | 39532182 | 39532225 | + | 4.01584230749346 | 5.68497871032191 | 2.38645642073598 | 7.46976052883995 |
| chr17 | 39532983 | 39532985 | + | 2.00792115374673 | 1.13699574206438 | 1.19322821036799 | 2.13421729395427 |
| chr17 | 39663189 | 39663204 | + | 7.02772403811355 | 3.41098722619315 | 4.77291284147196 | 2.13421729395427 |
| chr17 | 39663223 | 39663251 | + | 7.02772403811355 | 4.54798296825753 | 5.96614105183995 | 8.53686917581709 |
| chr17 | 39663272 | 39663316 | + | 7.02772403811355 | 6.82197445238629 | 7.15936926220794 | 14.9395210576799 |
| chr17 | 39663374 | 39663461 | + | 30.1188173062009 | 30.6988850357383 | 19.0916513658878 | 28.8119334683827 |
| chr17 | 39728605 | 39728625 | + | 12.0475269224804 | 13.6439489047726 | 1.19322821036799 | 1.06710864697714 |
| chr17 | 39729180 | 39729201 | - | 6.02376346124019 | 5.68497871032191 | 5.96614105183995 | 19.2079556455884 |
| chr17 | 39729248 | 39729289 | - | 9.03564519186028 | 6.82197445238629 | 2.38645642073598 | 14.9395210576799 |
| chr17 | 39729337 | 39729348 | - | 1.00396057687336 | 1.13699574206438 | 3.57968463110397 | 10.6710864697714 |
| chr17 | 39921104 | 39921331 | - | 5.01980288436682 | 4.54798296825753 | 3.57968463110397 | 11.7381951167485 |
| chr17 | 39921610 | 39921895 | - | 64.2534769198953 | 79.5897019445067 | 90.6853439879672 | 157.932079752616 |
| chr17 | 39921912 | 39922387 | - | 43.1703048055547 | 44.3428339405109 | 39.3765309421437 | 96.0397782279422 |
| chr17 | 39922480 | 39922524 | - | 6.02376346124019 | 6.82197445238629 | 5.96614105183995 | 2.13421729395427 |
| chr17 | 39927730 | 39927793 | + | 2.00792115374673 | 2.27399148412876 | 7.15936926220794 | 8.53686917581709 |
| chr17 | 39997648 | 39997654 | + | 1.00396057687336 | 1.13699574206438 | 1.19322821036799 | 5.33554323488568 |
| chr17 | 39997685 | 39997712 | + | 6.02376346124019 | 2.27399148412876 | 9.54582568294392 | 23.476390233497 |
| chr17 | 39997757 | 39997846 | + | 57.2257528817818 | 84.1376849127643 | 53.6952694665595 | 187.811121867976 |
| chr17 | 39997868 | 39997941 | + | 86.3406096111093 | 94.3706465913437 | 78.7530618842873 | 182.47557863309 |
| chr17 | 40019135 | 40019154 | - | 14.0554480762271 | 26.1509020674808 | 23.8645642073598 | 45.8856718200168 |
| chr17 | 40019226 | 40019250 | - | 10.0396057687336 | 4.54798296825753 | 4.77291284147196 | 21.3421729395427 |
| chr17 | 40019355 | 40019420 | - | 21.0831721143407 | 10.2329616785794 | 7.15936926220794 | 27.7448248214055 |
| chr17 | 40019442 | 40019511 | - | 7.02772403811355 | 10.2329616785794 | 9.54582568294392 | 49.0869977609482 |
| chr17 | 40091744 | 40091755 | + | 4.01584230749346 | 1.13699574206438 | 3.57968463110397 | 1.06710864697714 |
| chr17 | 40092192 | 40092200 | + | 19.0752509605939 | 28.4248935516096 | 2.38645642073598 | 12.8053037637256 |
| chr17 | 40092321 | 40092331 | + | 4.01584230749346 | 1.13699574206438 | 2.38645642073598 | 1.06710864697714 |
| chr17 | 40092438 | 40092467 | + | 10.0396057687336 | 6.82197445238629 | 1.19322821036799 | 6.40265188186281 |
| chr17 | 40092527 | 40092542 | + | 1.00396057687336 | 5.68497871032191 | 1.19322821036799 | 2.13421729395427 |
| chr17 | 40130154 | 40130170 | + | 5.01980288436682 | 10.2329616785794 | 3.57968463110397 | 4.26843458790854 |
| chr17 | 40134782 | 40134787 | + | 2.00792115374673 | 1.13699574206438 | 2.38645642073598 | 2.13421729395427 |
| chr17 | 40134906 | 40134930 | + | 2.00792115374673 | 3.41098722619315 | 7.15936926220794 | 1.06710864697714 |
| chr17 | 40134971 | 40134991 | + | 7.02772403811355 | 3.41098722619315 | 8.35259747257593 | 9.60397782279422 |
| chr17 | 40135239 | 40135262 | + | 3.01188173062009 | 4.54798296825753 | 3.57968463110397 | 8.53686917581709 |
| chr17 | 40135623 | 40135722 | + | 11.043566345607 | 10.2329616785794 | 25.0577924177278 | 43.7514545260626 |
| chr17 | 40135739 | 40135762 | + | 1.00396057687336 | 3.41098722619315 | 2.38645642073598 | 2.13421729395427 |
| chr17 | 40135848 | 40135869 | + | 2.00792115374673 | 6.82197445238629 | 5.96614105183995 | 9.60397782279422 |
| chr17 | 40136482 | 40136490 | + | 18.0712903837206 | 12.5069531627082 | 7.15936926220794 | 16.006629704657 |
| chr17 | 40136673 | 40136733 | + | 1.00396057687336 | 2.27399148412876 | 2.38645642073598 | 1.06710864697714 |
| chr17 | 40169606 | 40169625 | + | 5.01980288436682 | 5.68497871032191 | 5.96614105183995 | 3.20132594093141 |
| chr17 | 40170479 | 40170519 | + | 9.03564519186028 | 2.27399148412876 | 3.57968463110397 | 8.53686917581709 |
| chr17 | 40170638 | 40170688 | + | 13.0514874993537 | 5.68497871032191 | 1.19322821036799 | 3.20132594093141 |
| chr17 | 40170741 | 40170788 | + | 6.02376346124019 | 2.27399148412876 | 3.57968463110397 | 6.40265188186281 |
| chr17 | 40170809 | 40170819 | + | 3.01188173062009 | 5.68497871032191 | 5.96614105183995 | 3.20132594093141 |
| chr17 | 40171105 | 40171147 | + | 2.00792115374673 | 3.41098722619315 | 3.57968463110397 | 4.26843458790854 |
| chr17 | 40171216 | 40171251 | + | 8.03168461498692 | 5.68497871032191 | 4.77291284147196 | 8.53686917581709 |
| chr17 | 40171268 | 40171305 | + | 14.0554480762271 | 7.95897019445067 | 7.15936926220794 | 13.8724124107028 |
| chr17 | 40171325 | 40171359 | + | 17.0673298068472 | 10.2329616785794 | 4.77291284147196 | 5.33554323488568 |
| chr17 | 40171447 | 40171528 | + | 9.03564519186028 | 6.82197445238629 | 11.9322821036799 | 10.6710864697714 |
| chr17 | 40171850 | 40171970 | + | 19.0752509605939 | 17.0549361309657 | 4.77291284147196 | 18.1408469986113 |
| chr17 | 40172026 | 40172089 | + | 11.043566345607 | 10.2329616785794 | 4.77291284147196 | 5.33554323488568 |
| chr17 | 40172111 | 40172145 | + | 7.02772403811355 | 29.5618892936739 | 10.7390538933119 | 11.7381951167485 |
| chr17 | 40195106 | 40195133 | + | 27.1069355755808 | 20.4659233571589 | 1.19322821036799 | 1.06710864697714 |
| chr17 | 40278602 | 40278779 | + | 78.3089249961224 | 97.7816338175368 | 56.0817258872955 | 64.0265188186281 |
| chr17 | 40279425 | 40279445 | + | 8.03168461498692 | 36.3838637460602 | 5.96614105183995 | 10.6710864697714 |
| chr17 | 40279588 | 40279626 | + | 16.0633692299738 | 11.3699574206438 | 5.96614105183995 | 7.46976052883995 |
| chr17 | 40279648 | 40279657 | + | 14.0554480762271 | 11.3699574206438 | 5.96614105183995 | 6.40265188186281 |
| chr17 | 40279705 | 40279773 | + | 35.1386201905678 | 42.0688424563821 | 29.8307052591997 | 10.6710864697714 |
| chr17 | 40279937 | 40279941 | + | 2.00792115374673 | 1.13699574206438 | 1.19322821036799 | 1.06710864697714 |
| chr17 | 40280511 | 40280516 | + | 1.00396057687336 | 1.13699574206438 | 1.19322821036799 | 5.33554323488568 |
| chr17 | 40280558 | 40280573 | + | 2.00792115374673 | 4.54798296825753 | 2.38645642073598 | 2.13421729395427 |
| chr17 | 40280844 | 40280859 | + | 13.0514874993537 | 18.1919318730301 | 4.77291284147196 | 6.40265188186281 |
| chr17 | 40281048 | 40281067 | + | 4.01584230749346 | 2.27399148412876 | 2.38645642073598 | 1.06710864697714 |
| chr17 | 40281227 | 40281250 | + | 1.00396057687336 | 1.13699574206438 | 7.15936926220794 | 4.26843458790854 |
| chr17 | 40281695 | 40281714 | + | 2.00792115374673 | 5.68497871032191 | 4.77291284147196 | 3.20132594093141 |
| chr17 | 40281768 | 40281795 | + | 71.2812009580089 | 69.3567402659273 | 8.35259747257593 | 65.0936274656053 |
| chr17 | 40281819 | 40281843 | + | 104.41189999483 | 137.57648478979 | 19.0916513658878 | 87.5029090521251 |
| chr17 | 40281963 | 40281982 | + | 15.0594086531005 | 11.3699574206438 | 3.57968463110397 | 11.7381951167485 |
| chr17 | 40282013 | 40282033 | + | 103.407939417957 | 103.466612527859 | 145.573841664895 | 88.5700176991023 |
| chr17 | 40282152 | 40282162 | + | 27.1069355755808 | 43.2058381984465 | 3.57968463110397 | 5.33554323488568 |
| chr17 | 40302083 | 40302087 | + | 3.01188173062009 | 5.68497871032191 | 3.57968463110397 | 1.06710864697714 |
| chr17 | 40302273 | 40302297 | + | 3.01188173062009 | 4.54798296825753 | 1.19322821036799 | 4.26843458790854 |
| chr17 | 40302856 | 40302886 | + | 21.0831721143407 | 37.5208594881246 | 4.77291284147196 | 16.006629704657 |
| chr17 | 40303252 | 40303255 | + | 2.00792115374673 | 7.95897019445067 | 2.38645642073598 | 4.26843458790854 |
| chr17 | 40304074 | 40304076 | + | 3.01188173062009 | 5.68497871032191 | 1.19322821036799 | 1.06710864697714 |
| chr17 | 40356678 | 40356682 | + | 5.01980288436682 | 2.27399148412876 | 4.77291284147196 | 3.20132594093141 |
| chr17 | 40357545 | 40357578 | + | 5.01980288436682 | 2.27399148412876 | 3.57968463110397 | 4.26843458790854 |
| chr17 | 40388605 | 40388657 | - | 15.0594086531005 | 17.0549361309657 | 10.7390538933119 | 10.6710864697714 |
| chr17 | 40388691 | 40388723 | - | 15.0594086531005 | 36.3838637460602 | 17.8984231555198 | 13.8724124107028 |
| chr17 | 40388748 | 40388839 | - | 25.0990144218341 | 22.7399148412876 | 14.3187385244159 | 39.483019938154 |
| chr17 | 40389302 | 40389341 | - | 8.03168461498692 | 2.27399148412876 | 1.19322821036799 | 8.53686917581709 |
| chr17 | 40389417 | 40389421 | - | 7.02772403811355 | 2.27399148412876 | 1.19322821036799 | 2.13421729395427 |
| chr17 | 40627739 | 40627761 | - | 5.01980288436682 | 5.68497871032191 | 2.38645642073598 | 1.06710864697714 |
| chr17 | 40628705 | 40628803 | - | 19.0752509605939 | 11.3699574206438 | 19.0916513658878 | 16.006629704657 |
| chr17 | 40628851 | 40628860 | - | 6.02376346124019 | 6.82197445238629 | 11.9322821036799 | 12.8053037637256 |
| chr17 | 40628904 | 40628926 | - | 10.0396057687336 | 4.54798296825753 | 8.35259747257593 | 23.476390233497 |
| chr17 | 40630718 | 40630778 | - | 6.02376346124019 | 2.27399148412876 | 16.7051949451519 | 14.9395210576799 |
| chr17 | 40630800 | 40630862 | - | 29.1148567293276 | 20.4659233571589 | 78.7530618842873 | 77.8989312293309 |
| chr17 | 40630917 | 40630919 | - | 13.0514874993537 | 4.54798296825753 | 15.5119667347839 | 7.46976052883995 |
| chr17 | 40631591 | 40631604 | - | 4.01584230749346 | 2.27399148412876 | 5.96614105183995 | 2.13421729395427 |
| chr17 | 40631638 | 40631639 | - | 4.01584230749346 | 4.54798296825753 | 11.9322821036799 | 7.46976052883995 |
| chr17 | 40632222 | 40632239 | - | 3.01188173062009 | 5.68497871032191 | 3.57968463110397 | 3.20132594093141 |
| chr17 | 40632318 | 40632330 | - | 1.00396057687336 | 1.13699574206438 | 2.38645642073598 | 3.20132594093141 |
| chr17 | 40635982 | 40636014 | - | 35.1386201905678 | 19.3289276150945 | 29.8307052591997 | 56.5567582897882 |
| chr17 | 40636059 | 40636083 | - | 53.2099105742883 | 31.8358807778027 | 33.4103898903037 | 55.4896496428111 |
| chr17 | 40636406 | 40636425 | - | 3.01188173062009 | 3.41098722619315 | 7.15936926220794 | 3.20132594093141 |
| chr17 | 40818135 | 40818276 | - | 107.42378172545 | 169.412365567593 | 56.0817258872955 | 78.966039876308 |
| chr17 | 40818341 | 40818366 | - | 28.1108961524542 | 39.7948509722534 | 16.7051949451519 | 51.2212150549025 |
| chr17 | 40818391 | 40818478 | - | 98.3881365335897 | 196.700263377138 | 53.6952694665595 | 171.804492163319 |
| chr17 | 41523656 | 41523886 | - | 26.1029749987075 | 25.0139063254164 | 96.6514850398071 | 138.724124107028 |
| chr17 | 41690867 | 41690902 | + | 11.043566345607 | 31.8358807778027 | 27.4442488384638 | 16.006629704657 |
| chr17 | 41691026 | 41691108 | + | 6.02376346124019 | 12.5069531627082 | 8.35259747257593 | 23.476390233497 |
| chr17 | 41691157 | 41691469 | + | 244.966380757101 | 889.130670294347 | 328.137757851197 | 462.0580441411 |
| chr17 | 41691497 | 41691512 | + | 1.00396057687336 | 5.68497871032191 | 1.19322821036799 | 8.53686917581709 |
| chr17 | 41691541 | 41691626 | + | 37.1465413443145 | 137.57648478979 | 39.3765309421437 | 25.6106075274513 |
| chr17 | 41754629 | 41754643 | - | 1.00396057687336 | 9.09596593651506 | 2.38645642073598 | 8.53686917581709 |
| chr17 | 41754675 | 41754716 | - | 62.2455557661486 | 62.534765813541 | 27.4442488384638 | 141.925450047959 |
| chr17 | 41754770 | 41754779 | - | 7.02772403811355 | 4.54798296825753 | 1.19322821036799 | 3.20132594093141 |
| chr17 | 41755377 | 41755400 | - | 2.00792115374673 | 3.41098722619315 | 5.96614105183995 | 7.46976052883995 |
| chr17 | 41755476 | 41755519 | - | 9.03564519186028 | 3.41098722619315 | 4.77291284147196 | 25.6106075274513 |
| chr17 | 41823125 | 41823193 | + | 61.2415951892752 | 69.3567402659273 | 2.38645642073598 | 1.06710864697714 |
| chr17 | 41825205 | 41825240 | - | 9.03564519186028 | 21.6029190992233 | 8.35259747257593 | 11.7381951167485 |
| chr17 | 41825263 | 41825336 | - | 42.1663442286813 | 64.8087572976698 | 35.7968463110397 | 53.3554323488568 |
| chr17 | 41866938 | 41867000 | - | 4.01584230749346 | 13.6439489047726 | 5.96614105183995 | 25.6106075274513 |
| chr17 | 41867033 | 41867048 | - | 5.01980288436682 | 1.13699574206438 | 3.57968463110397 | 8.53686917581709 |
| chr17 | 41867209 | 41867227 | - | 14.0554480762271 | 5.68497871032191 | 7.15936926220794 | 16.006629704657 |
| chr17 | 41867488 | 41867519 | - | 11.043566345607 | 20.4659233571589 | 8.35259747257593 | 23.476390233497 |
| chr17 | 41867592 | 41867779 | - | 118.467348071057 | 72.7677274921204 | 428.368927522108 | 366.018265913158 |
| chr17 | 41867800 | 41867830 | - | 7.02772403811355 | 1.13699574206438 | 15.5119667347839 | 22.4092815865198 |
| chr17 | 41974326 | 41974424 | + | 24.0950538449607 | 5.68497871032191 | 3.57968463110397 | 1.06710864697714 |
| chr17 | 41975015 | 41975144 | + | 8.03168461498692 | 9.09596593651506 | 3.57968463110397 | 6.40265188186281 |
| chr17 | 41976491 | 41976538 | - | 60.2376346124019 | 76.1787147183136 | 32.2171616799357 | 74.6976052883995 |
| chr17 | 41976569 | 41976588 | - | 7.02772403811355 | 18.1919318730301 | 8.35259747257593 | 12.8053037637256 |
| chr17 | 42024320 | 42024324 | + | 1.00396057687336 | 1.13699574206438 | 4.77291284147196 | 2.13421729395427 |
| chr17 | 42024436 | 42024459 | + | 8.03168461498692 | 6.82197445238629 | 4.77291284147196 | 11.7381951167485 |
| chr17 | 42024749 | 42024750 | + | 1.00396057687336 | 1.13699574206438 | 1.19322821036799 | 1.06710864697714 |
| chr17 | 42024833 | 42024846 | + | 8.03168461498692 | 7.95897019445067 | 38.1833027317757 | 8.53686917581709 |
| chr17 | 42025520 | 42025613 | + | 21.0831721143407 | 23.876910583352 | 13.1255103140479 | 21.3421729395427 |
| chr17 | 42113136 | 42113189 | - | 69.2732798042621 | 64.8087572976698 | 22.6713359969918 | 29.8790421153598 |
| chr17 | 42125071 | 42125107 | - | 478.889195168595 | 359.290654492345 | 150.346754506367 | 331.870789209889 |
| chr17 | 42125270 | 42125364 | - | 22.087132691214 | 15.9179403889013 | 16.7051949451519 | 32.0132594093141 |
| chr17 | 42125388 | 42125465 | - | 16.0633692299738 | 4.54798296825753 | 17.8984231555198 | 13.8724124107028 |
| chr17 | 42125549 | 42125647 | - | 24.0950538449607 | 19.3289276150945 | 17.8984231555198 | 20.2750642925656 |
| chr17 | 42125660 | 42125682 | - | 39.1544624980612 | 31.8358807778027 | 52.5020412561915 | 39.483019938154 |
| chr17 | 42189109 | 42189120 | - | 2.00792115374673 | 4.54798296825753 | 1.19322821036799 | 4.26843458790854 |
| chr17 | 42199432 | 42199441 | - | 3.01188173062009 | 6.82197445238629 | 5.96614105183995 | 18.1408469986113 |
| chr17 | 42199813 | 42199852 | - | 4.01584230749346 | 5.68497871032191 | 10.7390538933119 | 36.2816939972226 |
| chr17 | 42199884 | 42199900 | - | 1.00396057687336 | 2.27399148412876 | 2.38645642073598 | 6.40265188186281 |
| chr17 | 42199936 | 42200011 | - | 2.00792115374673 | 3.41098722619315 | 14.3187385244159 | 26.6777161744284 |
| chr17 | 42200104 | 42200163 | - | 2.00792115374673 | 3.41098722619315 | 25.0577924177278 | 16.006629704657 |
| chr17 | 42200239 | 42200297 | - | 10.0396057687336 | 9.09596593651506 | 34.6036181006717 | 50.1541064079254 |
| chr17 | 42200326 | 42200367 | - | 6.02376346124019 | 1.13699574206438 | 7.15936926220794 | 18.1408469986113 |
| chr17 | 42200493 | 42200499 | - | 4.01584230749346 | 4.54798296825753 | 9.54582568294392 | 4.26843458790854 |
| chr17 | 42200533 | 42200548 | - | 3.01188173062009 | 7.95897019445067 | 8.35259747257593 | 12.8053037637256 |
| chr17 | 42200588 | 42200613 | - | 12.0475269224804 | 5.68497871032191 | 11.9322821036799 | 16.006629704657 |
| chr17 | 42200739 | 42200819 | - | 8.03168461498692 | 17.0549361309657 | 56.0817258872955 | 132.321472225165 |
| chr17 | 42200842 | 42200855 | - | 1.00396057687336 | 2.27399148412876 | 4.77291284147196 | 6.40265188186281 |
| chr17 | 42200949 | 42200952 | - | 2.00792115374673 | 1.13699574206438 | 1.19322821036799 | 3.20132594093141 |
| chr17 | 42200976 | 42201007 | - | 1.00396057687336 | 3.41098722619315 | 17.8984231555198 | 11.7381951167485 |
| chr17 | 42201102 | 42201139 | - | 2.00792115374673 | 7.95897019445067 | 13.1255103140479 | 27.7448248214055 |
| chr17 | 42201173 | 42201183 | - | 1.00396057687336 | 2.27399148412876 | 4.77291284147196 | 3.20132594093141 |
| chr17 | 42201311 | 42201372 | - | 5.01980288436682 | 3.41098722619315 | 16.7051949451519 | 13.8724124107028 |
| chr17 | 42201397 | 42201418 | - | 3.01188173062009 | 1.13699574206438 | 2.38645642073598 | 10.6710864697714 |
| chr17 | 42201508 | 42201538 | - | 3.01188173062009 | 5.68497871032191 | 14.3187385244159 | 12.8053037637256 |
| chr17 | 42201567 | 42201605 | - | 4.01584230749346 | 7.95897019445067 | 17.8984231555198 | 33.0803680562912 |
| chr17 | 42201677 | 42201699 | - | 2.00792115374673 | 2.27399148412876 | 7.15936926220794 | 8.53686917581709 |
| chr17 | 42201731 | 42201740 | - | 2.00792115374673 | 1.13699574206438 | 4.77291284147196 | 4.26843458790854 |
| chr17 | 42310705 | 42310722 | + | 1.00396057687336 | 1.13699574206438 | 1.19322821036799 | 9.60397782279422 |
| chr17 | 42311731 | 42311824 | + | 1.00396057687336 | 2.27399148412876 | 44.1494437836156 | 77.8989312293309 |
| chr17 | 42311848 | 42311857 | + | 2.00792115374673 | 7.95897019445067 | 28.6374770488317 | 145.12677598889 |
| chr17 | 42311911 | 42311921 | + | 3.01188173062009 | 1.13699574206438 | 3.57968463110397 | 6.40265188186281 |
| chr17 | 42313498 | 42313533 | - | 7.02772403811355 | 6.82197445238629 | 10.7390538933119 | 14.9395210576799 |
| chr17 | 42313557 | 42313624 | - | 17.0673298068472 | 5.68497871032191 | 21.4781077866238 | 37.3488026441998 |
| chr17 | 42313647 | 42313799 | - | 47.1861471130481 | 44.3428339405109 | 126.482190299007 | 131.254363578188 |
| chr17 | 42313823 | 42313861 | - | 11.043566345607 | 9.09596593651506 | 21.4781077866238 | 23.476390233497 |
| chr17 | 42313881 | 42313932 | - | 14.0554480762271 | 11.3699574206438 | 35.7968463110397 | 20.2750642925656 |
| chr17 | 42313955 | 42313975 | - | 5.01980288436682 | 3.41098722619315 | 15.5119667347839 | 21.3421729395427 |
| chr17 | 42313997 | 42314008 | - | 2.00792115374673 | 3.41098722619315 | 8.35259747257593 | 3.20132594093141 |
| chr17 | 42314030 | 42314083 | - | 19.0752509605939 | 12.5069531627082 | 17.8984231555198 | 22.4092815865198 |
| chr17 | 42314109 | 42314141 | - | 5.01980288436682 | 3.41098722619315 | 5.96614105183995 | 5.33554323488568 |
| chr17 | 42314181 | 42314406 | - | 53.2099105742883 | 57.9867828452835 | 161.085808399679 | 126.985928990279 |
| chr17 | 42314428 | 42314456 | - | 8.03168461498692 | 3.41098722619315 | 19.0916513658878 | 2.13421729395427 |
| chr17 | 42314519 | 42314568 | - | 25.0990144218341 | 3.41098722619315 | 25.0577924177278 | 39.483019938154 |
| chr17 | 42314608 | 42314649 | - | 23.0910932680874 | 23.876910583352 | 42.9562155732476 | 55.4896496428111 |
| chr17 | 42314687 | 42314741 | - | 17.0673298068472 | 19.3289276150945 | 53.6952694665595 | 35.2145853502455 |
| chr17 | 42314764 | 42314843 | - | 15.0594086531005 | 19.3289276150945 | 29.8307052591997 | 43.7514545260626 |
| chr17 | 42314893 | 42314905 | - | 13.0514874993537 | 11.3699574206438 | 21.4781077866238 | 5.33554323488568 |
| chr17 | 42315110 | 42315112 | - | 6.02376346124019 | 4.54798296825753 | 3.57968463110397 | 16.006629704657 |
| chr17 | 42315205 | 42315207 | - | 2.00792115374673 | 6.82197445238629 | 1.19322821036799 | 5.33554323488568 |
| chr17 | 42315249 | 42315265 | - | 9.03564519186028 | 4.54798296825753 | 3.57968463110397 | 1.06710864697714 |
| chr17 | 42315297 | 42315303 | - | 3.01188173062009 | 2.27399148412876 | 2.38645642073598 | 1.06710864697714 |
| chr17 | 42315451 | 42315566 | - | 51.2019894205416 | 55.7127913611547 | 157.506123768575 | 51.2212150549025 |
| chr17 | 42315587 | 42315612 | - | 9.03564519186028 | 7.95897019445067 | 25.0577924177278 | 6.40265188186281 |
| chr17 | 42315636 | 42315643 | - | 5.01980288436682 | 3.41098722619315 | 4.77291284147196 | 4.26843458790854 |
| chr17 | 42315664 | 42315742 | - | 18.0712903837206 | 18.1919318730301 | 38.1833027317757 | 20.2750642925656 |
| chr17 | 42315750 | 42315808 | - | 7.02772403811355 | 2.27399148412876 | 17.8984231555198 | 12.8053037637256 |
| chr17 | 42316794 | 42316854 | - | 15.0594086531005 | 5.68497871032191 | 27.4442488384638 | 18.1408469986113 |
| chr17 | 42402473 | 42402555 | - | 100.396057687336 | 45.4798296825753 | 32.2171616799357 | 69.3620620535138 |
| chr17 | 42402585 | 42402603 | - | 1.00396057687336 | 9.09596593651506 | 1.19322821036799 | 6.40265188186281 |
| chr17 | 42402699 | 42402723 | - | 9.03564519186028 | 9.09596593651506 | 4.77291284147196 | 10.6710864697714 |
| chr17 | 42402952 | 42402981 | - | 5.01980288436682 | 2.27399148412876 | 4.77291284147196 | 4.26843458790854 |
| chr17 | 42403297 | 42403361 | - | 7.02772403811355 | 7.95897019445067 | 2.38645642073598 | 20.2750642925656 |
| chr17 | 42403469 | 42403518 | - | 14.0554480762271 | 3.41098722619315 | 11.9322821036799 | 39.483019938154 |
| chr17 | 42403599 | 42403662 | - | 21.0831721143407 | 14.780944646837 | 14.3187385244159 | 21.3421729395427 |
| chr17 | 42404035 | 42404054 | - | 7.02772403811355 | 1.13699574206438 | 4.77291284147196 | 4.26843458790854 |
| chr17 | 42404083 | 42404099 | - | 2.00792115374673 | 1.13699574206438 | 1.19322821036799 | 4.26843458790854 |
| chr17 | 42404178 | 42404236 | - | 3.01188173062009 | 9.09596593651506 | 4.77291284147196 | 4.26843458790854 |
| chr17 | 42404283 | 42404284 | - | 1.00396057687336 | 1.13699574206438 | 5.96614105183995 | 3.20132594093141 |
| chr17 | 42404373 | 42404481 | - | 82.3247673036159 | 43.2058381984465 | 50.1155848354556 | 188.878230514953 |
| chr17 | 42404590 | 42404611 | - | 4.01584230749346 | 7.95897019445067 | 3.57968463110397 | 1.06710864697714 |
| chr17 | 42522491 | 42522561 | + | 6.02376346124019 | 1.13699574206438 | 4.77291284147196 | 11.7381951167485 |
| chr17 | 42566093 | 42566259 | + | 20.0792115374673 | 19.3289276150945 | 7.15936926220794 | 13.8724124107028 |
| chr17 | 42571621 | 42571640 | + | 5.01980288436682 | 3.41098722619315 | 3.57968463110397 | 5.33554323488568 |
| chr17 | 42572385 | 42572429 | - | 21.0831721143407 | 43.2058381984465 | 9.54582568294392 | 10.6710864697714 |
| chr17 | 42572536 | 42572560 | + | 8.03168461498692 | 4.54798296825753 | 1.19322821036799 | 5.33554323488568 |
| chr17 | 42572550 | 42572553 | - | 2.00792115374673 | 1.13699574206438 | 1.19322821036799 | 1.06710864697714 |
| chr17 | 42572577 | 42572619 | - | 17.0673298068472 | 36.3838637460602 | 11.9322821036799 | 22.4092815865198 |
| chr17 | 42572631 | 42572653 | + | 12.0475269224804 | 10.2329616785794 | 1.19322821036799 | 3.20132594093141 |
| chr17 | 42572643 | 42572663 | - | 2.00792115374673 | 6.82197445238629 | 1.19322821036799 | 1.06710864697714 |
| chr17 | 42572781 | 42572804 | - | 12.0475269224804 | 45.4798296825753 | 26.2510206280958 | 32.0132594093141 |
| chr17 | 42572811 | 42572840 | - | 6.02376346124019 | 9.09596593651506 | 2.38645642073598 | 3.20132594093141 |
| chr17 | 42572918 | 42572944 | - | 6.02376346124019 | 6.82197445238629 | 4.77291284147196 | 7.46976052883995 |
| chr17 | 42579535 | 42579556 | - | 10.0396057687336 | 20.4659233571589 | 5.96614105183995 | 13.8724124107028 |
| chr17 | 42579685 | 42579724 | - | 7.02772403811355 | 2.27399148412876 | 1.19322821036799 | 1.06710864697714 |
| chr17 | 42579804 | 42579850 | - | 2.00792115374673 | 4.54798296825753 | 2.38645642073598 | 1.06710864697714 |
| chr17 | 42580474 | 42580577 | - | 12.0475269224804 | 11.3699574206438 | 10.7390538933119 | 5.33554323488568 |
| chr17 | 42580600 | 42580644 | - | 7.02772403811355 | 4.54798296825753 | 1.19322821036799 | 5.33554323488568 |
| chr17 | 42580756 | 42580779 | - | 22.087132691214 | 25.0139063254164 | 3.57968463110397 | 17.0737383516342 |
| chr17 | 42581082 | 42581131 | - | 12.0475269224804 | 6.82197445238629 | 2.38645642073598 | 12.8053037637256 |
| chr17 | 42581533 | 42581536 | - | 2.00792115374673 | 1.13699574206438 | 2.38645642073598 | 7.46976052883995 |
| chr17 | 42581559 | 42581586 | - | 9.03564519186028 | 3.41098722619315 | 1.19322821036799 | 3.20132594093141 |
| chr17 | 42581737 | 42581756 | - | 3.01188173062009 | 4.54798296825753 | 9.54582568294392 | 8.53686917581709 |
| chr17 | 42581781 | 42581825 | - | 9.03564519186028 | 1.13699574206438 | 10.7390538933119 | 7.46976052883995 |
| chr17 | 42615056 | 42615175 | + | 38.1505019211879 | 39.7948509722534 | 21.4781077866238 | 43.7514545260626 |
| chr17 | 42667991 | 42667999 | - | 16.0633692299738 | 25.0139063254164 | 7.15936926220794 | 5.33554323488568 |
| chr17 | 42679936 | 42680000 | + | 3.01188173062009 | 10.2329616785794 | 79.9462900946553 | 45.8856718200168 |
| chr17 | 42699678 | 42699704 | + | 1.00396057687336 | 6.82197445238629 | 7.15936926220794 | 1.06710864697714 |
| chr17 | 42700347 | 42700389 | - | 3.01188173062009 | 6.82197445238629 | 1.19322821036799 | 4.26843458790854 |
| chr17 | 42700515 | 42700533 | - | 2.00792115374673 | 3.41098722619315 | 3.57968463110397 | 3.20132594093141 |
| chr17 | 42700631 | 42700640 | - | 2.00792115374673 | 4.54798296825753 | 5.96614105183995 | 1.06710864697714 |
| chr17 | 42779009 | 42779074 | + | 4.01584230749346 | 6.82197445238629 | 14.3187385244159 | 25.6106075274513 |
| chr17 | 42779169 | 42779280 | + | 17.0673298068472 | 27.2878978095452 | 35.7968463110397 | 51.2212150549025 |
| chr17 | 42779314 | 42779345 | + | 8.03168461498692 | 2.27399148412876 | 3.57968463110397 | 13.8724124107028 |
| chr17 | 42779435 | 42779572 | + | 6.02376346124019 | 4.54798296825753 | 7.15936926220794 | 19.2079556455884 |
| chr17 | 42796783 | 42796867 | + | 1.00396057687336 | 4.54798296825753 | 20.2848795762558 | 25.6106075274513 |
| chr17 | 42798053 | 42798078 | - | 8.03168461498692 | 13.6439489047726 | 3.57968463110397 | 9.60397782279422 |
| chr17 | 42810154 | 42810234 | - | 13.0514874993537 | 18.1919318730301 | 7.15936926220794 | 10.6710864697714 |
| chr17 | 42810281 | 42810371 | - | 22.087132691214 | 23.876910583352 | 9.54582568294392 | 13.8724124107028 |
| chr17 | 42810505 | 42810572 | - | 3.01188173062009 | 9.09596593651506 | 3.57968463110397 | 3.20132594093141 |
| chr17 | 42810741 | 42810748 | - | 10.0396057687336 | 11.3699574206438 | 2.38645642073598 | 12.8053037637256 |
| chr17 | 42841584 | 42841630 | + | 25.0990144218341 | 11.3699574206438 | 5.96614105183995 | 18.1408469986113 |
| chr17 | 42841644 | 42841677 | + | 39.1544624980612 | 52.3018041349616 | 9.54582568294392 | 42.6843458790854 |
| chr17 | 42841707 | 42841729 | + | 5.01980288436682 | 5.68497871032191 | 2.38645642073598 | 2.13421729395427 |
| chr17 | 42841842 | 42841851 | + | 10.0396057687336 | 9.09596593651506 | 2.38645642073598 | 9.60397782279422 |
| chr17 | 42841893 | 42841905 | + | 5.01980288436682 | 11.3699574206438 | 3.57968463110397 | 2.13421729395427 |
| chr17 | 42841950 | 42841968 | + | 16.0633692299738 | 15.9179403889013 | 4.77291284147196 | 12.8053037637256 |
| chr17 | 42841995 | 42842104 | + | 122.48319037855 | 134.165497563597 | 57.2749540976635 | 86.435800405148 |
| chr17 | 42842124 | 42842167 | + | 10.0396057687336 | 7.95897019445067 | 10.7390538933119 | 17.0737383516342 |
| chr17 | 42842311 | 42842325 | + | 7.02772403811355 | 3.41098722619315 | 2.38645642073598 | 2.13421729395427 |
| chr17 | 42842381 | 42842539 | + | 29.1148567293276 | 39.7948509722534 | 21.4781077866238 | 43.7514545260626 |
| chr17 | 42843111 | 42843135 | + | 8.03168461498692 | 6.82197445238629 | 1.19322821036799 | 8.53686917581709 |
| chr17 | 42843196 | 42843223 | + | 2.00792115374673 | 10.2329616785794 | 5.96614105183995 | 1.06710864697714 |
| chr17 | 42950581 | 42950592 | - | 13.0514874993537 | 18.1919318730301 | 9.54582568294392 | 16.006629704657 |
| chr17 | 42993158 | 42993177 | + | 2.00792115374673 | 2.27399148412876 | 2.38645642073598 | 3.20132594093141 |
| chr17 | 43002869 | 43002938 | + | 244.966380757101 | 329.728765198671 | 175.404546924094 | 139.791232754005 |
| chr17 | 43014626 | 43014751 | - | 25.0990144218341 | 15.9179403889013 | 23.8645642073598 | 41.6172372321083 |
| chr17 | 43015816 | 43015889 | - | 7.02772403811355 | 2.27399148412876 | 9.54582568294392 | 16.006629704657 |
| chr17 | 43015922 | 43016170 | - | 36.1425807674411 | 22.7399148412876 | 22.6713359969918 | 19.2079556455884 |
| chr17 | 43209986 | 43210081 | + | 17.0673298068472 | 17.0549361309657 | 15.5119667347839 | 23.476390233497 |
| chr17 | 43210105 | 43210192 | + | 27.1069355755808 | 18.1919318730301 | 14.3187385244159 | 22.4092815865198 |
| chr17 | 43210392 | 43210475 | + | 4.01584230749346 | 1.13699574206438 | 2.38645642073598 | 3.20132594093141 |
| chr17 | 43210717 | 43210772 | + | 3.01188173062009 | 3.41098722619315 | 4.77291284147196 | 3.20132594093141 |
| chr17 | 43210795 | 43210870 | + | 5.01980288436682 | 10.2329616785794 | 4.77291284147196 | 9.60397782279422 |
| chr17 | 43210891 | 43211049 | + | 7.02772403811355 | 5.68497871032191 | 11.9322821036799 | 13.8724124107028 |
| chr17 | 43211127 | 43211142 | + | 7.02772403811355 | 5.68497871032191 | 7.15936926220794 | 25.6106075274513 |
| chr17 | 43211304 | 43211385 | + | 5.01980288436682 | 22.7399148412876 | 5.96614105183995 | 24.5434988804741 |
| chr17 | 43211603 | 43211634 | + | 4.01584230749346 | 9.09596593651506 | 3.57968463110397 | 7.46976052883995 |
| chr17 | 43523889 | 43523911 | + | 4.01584230749346 | 3.41098722619315 | 1.19322821036799 | 2.13421729395427 |
| chr17 | 43766231 | 43766297 | - | 31.1227778830743 | 32.9728765198671 | 21.4781077866238 | 10.6710864697714 |
| chr17 | 43766319 | 43766346 | - | 7.02772403811355 | 7.95897019445067 | 5.96614105183995 | 6.40265188186281 |
| chr17 | 43766402 | 43766414 | - | 2.00792115374673 | 6.82197445238629 | 2.38645642073598 | 2.13421729395427 |
| chr17 | 43766532 | 43766637 | - | 3.01188173062009 | 4.54798296825753 | 4.77291284147196 | 2.13421729395427 |
| chr17 | 43766681 | 43766686 | - | 26.1029749987075 | 7.95897019445067 | 9.54582568294392 | 6.40265188186281 |
| chr17 | 43766725 | 43766730 | - | 11.043566345607 | 6.82197445238629 | 4.77291284147196 | 4.26843458790854 |
| chr17 | 43766760 | 43766804 | - | 1.00396057687336 | 2.27399148412876 | 1.19322821036799 | 1.06710864697714 |
| chr17 | 43766900 | 43766964 | - | 1.00396057687336 | 1.13699574206438 | 1.19322821036799 | 2.13421729395427 |
| chr17 | 43769040 | 43769088 | - | 14.0554480762271 | 15.9179403889013 | 14.3187385244159 | 4.26843458790854 |
| chr17 | 43769265 | 43769299 | - | 7.02772403811355 | 6.82197445238629 | 2.38645642073598 | 2.13421729395427 |
| chr17 | 43769342 | 43769362 | - | 2.00792115374673 | 1.13699574206438 | 5.96614105183995 | 4.26843458790854 |
| chr17 | 43769414 | 43769423 | - | 9.03564519186028 | 3.41098722619315 | 3.57968463110397 | 2.13421729395427 |
| chr17 | 44011219 | 44011254 | - | 7.02772403811355 | 22.7399148412876 | 20.2848795762558 | 16.006629704657 |
| chr17 | 44011313 | 44011334 | - | 3.01188173062009 | 1.13699574206438 | 3.57968463110397 | 2.13421729395427 |
| chr17 | 44011383 | 44011454 | - | 3.01188173062009 | 6.82197445238629 | 21.4781077866238 | 25.6106075274513 |
| chr17 | 44011485 | 44011538 | - | 2.00792115374673 | 5.68497871032191 | 14.3187385244159 | 8.53686917581709 |
| chr17 | 44011560 | 44011690 | - | 26.1029749987075 | 12.5069531627082 | 51.3088130458235 | 27.7448248214055 |
| chr17 | 44011712 | 44011755 | - | 3.01188173062009 | 4.54798296825753 | 20.2848795762558 | 11.7381951167485 |
| chr17 | 44011819 | 44011855 | - | 5.01980288436682 | 1.13699574206438 | 2.38645642073598 | 4.26843458790854 |
| chr17 | 44011875 | 44011952 | - | 37.1465413443145 | 29.5618892936739 | 25.0577924177278 | 55.4896496428111 |
| chr17 | 44035773 | 44035788 | - | 13.0514874993537 | 4.54798296825753 | 15.5119667347839 | 28.8119334683827 |
| chr17 | 44076264 | 44076320 | + | 14.0554480762271 | 13.6439489047726 | 4.77291284147196 | 6.40265188186281 |
| chr17 | 44078350 | 44078413 | - | 7.02772403811355 | 2.27399148412876 | 1.19322821036799 | 7.46976052883995 |
| chr17 | 44162318 | 44162349 | + | 6.02376346124019 | 2.27399148412876 | 1.19322821036799 | 3.20132594093141 |
| chr17 | 44162430 | 44162437 | + | 3.01188173062009 | 2.27399148412876 | 5.96614105183995 | 6.40265188186281 |
| chr17 | 44191226 | 44191240 | + | 3.01188173062009 | 2.27399148412876 | 3.57968463110397 | 1.06710864697714 |
| chr17 | 44191274 | 44191276 | + | 5.01980288436682 | 2.27399148412876 | 1.19322821036799 | 3.20132594093141 |
| chr17 | 44191320 | 44191341 | + | 4.01584230749346 | 1.13699574206438 | 2.38645642073598 | 3.20132594093141 |
| chr17 | 44191409 | 44191443 | + | 7.02772403811355 | 3.41098722619315 | 14.3187385244159 | 5.33554323488568 |
| chr17 | 44191484 | 44191502 | + | 2.00792115374673 | 1.13699574206438 | 3.57968463110397 | 9.60397782279422 |
| chr17 | 44191639 | 44191700 | + | 2.00792115374673 | 6.82197445238629 | 3.57968463110397 | 1.06710864697714 |
| chr17 | 44191830 | 44191838 | - | 1.00396057687336 | 4.54798296825753 | 2.38645642073598 | 2.13421729395427 |
| chr17 | 44191860 | 44191890 | - | 56.2217923049084 | 31.8358807778027 | 54.8884976769275 | 30.9461507623369 |
| chr17 | 44191923 | 44191964 | - | 409.615915364333 | 392.263531012212 | 421.2095582599 | 400.165742616426 |
| chr17 | 44192045 | 44192085 | - | 1.00396057687336 | 4.54798296825753 | 1.19322821036799 | 1.06710864697714 |
| chr17 | 44192145 | 44192155 | - | 3.01188173062009 | 5.68497871032191 | 3.57968463110397 | 8.53686917581709 |
| chr17 | 44192185 | 44192240 | - | 3.01188173062009 | 9.09596593651506 | 11.9322821036799 | 35.2145853502455 |
| chr17 | 44192268 | 44192285 | - | 2.00792115374673 | 7.95897019445067 | 8.35259747257593 | 7.46976052883995 |
| chr17 | 44192360 | 44192387 | - | 9.03564519186028 | 5.68497871032191 | 14.3187385244159 | 10.6710864697714 |
| chr17 | 44192413 | 44192445 | - | 14.0554480762271 | 13.6439489047726 | 39.3765309421437 | 13.8724124107028 |
| chr17 | 44192478 | 44192508 | - | 10.0396057687336 | 7.95897019445067 | 11.9322821036799 | 7.46976052883995 |
| chr17 | 44192534 | 44192555 | - | 3.01188173062009 | 10.2329616785794 | 11.9322821036799 | 10.6710864697714 |
| chr17 | 44192722 | 44192726 | - | 1.00396057687336 | 4.54798296825753 | 5.96614105183995 | 2.13421729395427 |
| chr17 | 44194012 | 44194036 | - | 4.01584230749346 | 1.13699574206438 | 2.38645642073598 | 1.06710864697714 |
| chr17 | 44205098 | 44205132 | - | 4.01584230749346 | 4.54798296825753 | 1.19322821036799 | 3.20132594093141 |
| chr17 | 44206407 | 44206428 | - | 20.0792115374673 | 12.5069531627082 | 3.57968463110397 | 2.13421729395427 |
| chr17 | 44206539 | 44206562 | - | 107.42378172545 | 118.247557174696 | 58.4681823080315 | 17.0737383516342 |
| chr17 | 44206647 | 44206704 | - | 136.538638454778 | 109.151591238181 | 56.0817258872955 | 62.959410171651 |
| chr17 | 44207201 | 44207210 | - | 6.02376346124019 | 3.41098722619315 | 4.77291284147196 | 10.6710864697714 |
| chr17 | 44319658 | 44319768 | - | 70.2772403811355 | 76.1787147183136 | 69.2072362013434 | 87.5029090521251 |
| chr17 | 44319796 | 44319799 | - | 47.1861471130481 | 21.6029190992233 | 1.19322821036799 | 2.13421729395427 |
| chr17 | 44319846 | 44320119 | - | 195.772312490306 | 134.165497563597 | 221.940447128446 | 331.870789209889 |
| chr17 | 44352926 | 44353081 | + | 164.649534607232 | 136.439489047726 | 28.6374770488317 | 33.0803680562912 |
| chr17 | 44353269 | 44353271 | - | 5.01980288436682 | 2.27399148412876 | 2.38645642073598 | 1.06710864697714 |
| chr17 | 44353345 | 44353353 | - | 4.01584230749346 | 5.68497871032191 | 5.96614105183995 | 3.20132594093141 |
| chr17 | 44353378 | 44353381 | - | 9.03564519186028 | 4.54798296825753 | 7.15936926220794 | 4.26843458790854 |
| chr17 | 44353529 | 44353534 | - | 2.00792115374673 | 1.13699574206438 | 1.19322821036799 | 1.06710864697714 |
| chr17 | 44677526 | 44677540 | - | 4.01584230749346 | 11.3699574206438 | 9.54582568294392 | 3.20132594093141 |
| chr17 | 44677635 | 44677640 | - | 4.01584230749346 | 11.3699574206438 | 9.54582568294392 | 11.7381951167485 |
| chr17 | 44678317 | 44678364 | - | 98.3881365335897 | 100.055625301666 | 22.6713359969918 | 58.6909755837425 |
| chr17 | 44678392 | 44678403 | - | 4.01584230749346 | 2.27399148412876 | 2.38645642073598 | 1.06710864697714 |
| chr17 | 44678610 | 44678628 | - | 11.043566345607 | 6.82197445238629 | 1.19322821036799 | 1.06710864697714 |
| chr17 | 44678848 | 44678879 | - | 4.01584230749346 | 6.82197445238629 | 1.19322821036799 | 5.33554323488568 |
| chr17 | 44678933 | 44678935 | - | 2.00792115374673 | 1.13699574206438 | 4.77291284147196 | 1.06710864697714 |
| chr17 | 44678975 | 44678995 | - | 6.02376346124019 | 6.82197445238629 | 2.38645642073598 | 1.06710864697714 |
| chr17 | 44850829 | 44850830 | - | 3.01188173062009 | 3.41098722619315 | 3.57968463110397 | 1.06710864697714 |
| chr17 | 44850930 | 44851045 | - | 30.1188173062009 | 35.2468680039958 | 28.6374770488317 | 50.1541064079254 |
| chr17 | 44851187 | 44851320 | - | 38.1505019211879 | 20.4659233571589 | 36.9900745214077 | 78.966039876308 |
| chr17 | 44925965 | 44925990 | - | 8.03168461498692 | 2.27399148412876 | 10.7390538933119 | 9.60397782279422 |
| chr17 | 45023382 | 45023406 | - | 1.00396057687336 | 2.27399148412876 | 2.38645642073598 | 2.13421729395427 |
| chr17 | 45023718 | 45023740 | - | 3.01188173062009 | 1.13699574206438 | 3.57968463110397 | 5.33554323488568 |
| chr17 | 45023987 | 45024002 | - | 1.00396057687336 | 3.41098722619315 | 3.57968463110397 | 5.33554323488568 |
| chr17 | 45024089 | 45024109 | - | 6.02376346124019 | 7.95897019445067 | 8.35259747257593 | 5.33554323488568 |
| chr17 | 45024376 | 45024384 | - | 2.00792115374673 | 1.13699574206438 | 7.15936926220794 | 4.26843458790854 |
| chr17 | 45024414 | 45024419 | - | 2.00792115374673 | 1.13699574206438 | 4.77291284147196 | 4.26843458790854 |
| chr17 | 45105649 | 45105660 | + | 5.01980288436682 | 2.27399148412876 | 3.57968463110397 | 5.33554323488568 |
| chr17 | 45105706 | 45105866 | + | 491.940682667949 | 302.440867389126 | 661.048428543866 | 712.828576180727 |
| chr17 | 45105903 | 45105948 | + | 36.1425807674411 | 47.753821166704 | 33.4103898903037 | 19.2079556455884 |
| chr17 | 45106952 | 45107007 | + | 3.01188173062009 | 4.54798296825753 | 4.77291284147196 | 3.20132594093141 |
| chr17 | 45107036 | 45107075 | + | 2.00792115374673 | 5.68497871032191 | 9.54582568294392 | 6.40265188186281 |
| chr17 | 45107181 | 45107186 | + | 1.00396057687336 | 2.27399148412876 | 2.38645642073598 | 10.6710864697714 |
| chr17 | 45108492 | 45108551 | + | 4.01584230749346 | 2.27399148412876 | 17.8984231555198 | 5.33554323488568 |
| chr17 | 45108627 | 45108675 | + | 1.00396057687336 | 1.13699574206438 | 5.96614105183995 | 11.7381951167485 |
| chr17 | 45108910 | 45108944 | + | 34.1346596136944 | 31.8358807778027 | 20.2848795762558 | 65.0936274656053 |
| chr17 | 45150287 | 45150355 | + | 39.1544624980612 | 45.4798296825753 | 14.3187385244159 | 54.4225409958339 |
| chr17 | 45150378 | 45150473 | + | 22.087132691214 | 17.0549361309657 | 11.9322821036799 | 24.5434988804741 |
| chr17 | 45150576 | 45150601 | + | 9.03564519186028 | 10.2329616785794 | 1.19322821036799 | 2.13421729395427 |
| chr17 | 45151824 | 45151832 | + | 1.00396057687336 | 1.13699574206438 | 2.38645642073598 | 1.06710864697714 |
| chr17 | 45151877 | 45151899 | + | 2.00792115374673 | 2.27399148412876 | 2.38645642073598 | 3.20132594093141 |
| chr17 | 45152036 | 45152040 | + | 1.00396057687336 | 1.13699574206438 | 3.57968463110397 | 1.06710864697714 |
| chr17 | 45152067 | 45152069 | + | 2.00792115374673 | 2.27399148412876 | 1.19322821036799 | 2.13421729395427 |
| chr17 | 45247875 | 45247974 | + | 125.495072109171 | 128.480518853275 | 943.84351440108 | 714.962793474681 |
| chr17 | 45268387 | 45268394 | + | 2.00792115374673 | 1.13699574206438 | 5.96614105183995 | 2.13421729395427 |
| chr17 | 46936779 | 46936782 | + | 3.01188173062009 | 7.95897019445067 | 1.19322821036799 | 1.06710864697714 |
| chr17 | 46938659 | 46938675 | + | 1.00396057687336 | 2.27399148412876 | 2.38645642073598 | 8.53686917581709 |
| chr17 | 47117999 | 47118002 | - | 6.02376346124019 | 14.780944646837 | 14.3187385244159 | 17.0737383516342 |
| chr17 | 47118326 | 47118482 | - | 207.819839412786 | 196.700263377138 | 285.181542277949 | 166.468948928433 |
| chr17 | 47118520 | 47118529 | - | 9.03564519186028 | 11.3699574206438 | 76.3666054635513 | 3.20132594093141 |
| chr17 | 47118623 | 47118654 | - | 5.01980288436682 | 5.68497871032191 | 7.15936926220794 | 10.6710864697714 |
| chr17 | 47118753 | 47118814 | - | 9.03564519186028 | 6.82197445238629 | 29.8307052591997 | 19.2079556455884 |
| chr17 | 47118843 | 47118885 | - | 63.249516343022 | 63.6717615556054 | 26.2510206280958 | 58.6909755837425 |
| chr17 | 47118935 | 47118975 | - | 16.0633692299738 | 15.9179403889013 | 11.9322821036799 | 12.8053037637256 |
| chr17 | 47118993 | 47118997 | - | 4.01584230749346 | 1.13699574206438 | 1.19322821036799 | 1.06710864697714 |
| chr17 | 47119069 | 47119096 | - | 1.00396057687336 | 10.2329616785794 | 10.7390538933119 | 19.2079556455884 |
| chr17 | 47119144 | 47119197 | - | 6.02376346124019 | 15.9179403889013 | 20.2848795762558 | 9.60397782279422 |
| chr17 | 47119224 | 47119236 | - | 9.03564519186028 | 11.3699574206438 | 20.2848795762558 | 32.0132594093141 |
| chr17 | 47119304 | 47119333 | - | 9.03564519186028 | 7.95897019445067 | 16.7051949451519 | 20.2750642925656 |
| chr17 | 47119437 | 47119467 | - | 9.03564519186028 | 4.54798296825753 | 5.96614105183995 | 7.46976052883995 |
| chr17 | 47119492 | 47119545 | - | 12.0475269224804 | 11.3699574206438 | 17.8984231555198 | 4.26843458790854 |
| chr17 | 47119583 | 47119594 | - | 3.01188173062009 | 4.54798296825753 | 2.38645642073598 | 2.13421729395427 |
| chr17 | 47119738 | 47119835 | - | 84.3326884573626 | 94.3706465913437 | 56.0817258872955 | 27.7448248214055 |
| chr17 | 47119863 | 47119883 | - | 3.01188173062009 | 11.3699574206438 | 7.15936926220794 | 7.46976052883995 |
| chr17 | 47119891 | 47119930 | - | 4.01584230749346 | 9.09596593651506 | 2.38645642073598 | 1.06710864697714 |
| chr17 | 47120059 | 47120082 | - | 3.01188173062009 | 2.27399148412876 | 3.57968463110397 | 10.6710864697714 |
| chr17 | 47120348 | 47120372 | - | 2.00792115374673 | 1.13699574206438 | 1.19322821036799 | 1.06710864697714 |
| chr17 | 47120438 | 47120473 | - | 4.01584230749346 | 2.27399148412876 | 1.19322821036799 | 2.13421729395427 |
| chr17 | 47120614 | 47120646 | - | 8.03168461498692 | 10.2329616785794 | 8.35259747257593 | 12.8053037637256 |
| chr17 | 47120693 | 47120712 | - | 46.1821865361748 | 71.6307317500561 | 36.9900745214077 | 66.1607361125824 |
| chr17 | 47621892 | 47622061 | + | 90.3564519186028 | 64.8087572976698 | 94.2650286190712 | 83.2344744642166 |
| chr17 | 47622091 | 47622117 | + | 3.01188173062009 | 7.95897019445067 | 9.54582568294392 | 2.13421729395427 |
| chr17 | 47622149 | 47622192 | + | 7.02772403811355 | 7.95897019445067 | 7.15936926220794 | 1.06710864697714 |
| chr17 | 47622232 | 47622241 | + | 1.00396057687336 | 1.13699574206438 | 2.38645642073598 | 3.20132594093141 |
| chr17 | 47622278 | 47622457 | + | 40.1584230749346 | 29.5618892936739 | 93.0718004087032 | 107.777973344691 |
| chr17 | 47622530 | 47622546 | + | 4.01584230749346 | 2.27399148412876 | 8.35259747257593 | 11.7381951167485 |
| chr17 | 47622583 | 47622608 | + | 4.01584230749346 | 2.27399148412876 | 8.35259747257593 | 1.06710864697714 |
| chr17 | 47622629 | 47622668 | + | 14.0554480762271 | 13.6439489047726 | 26.2510206280958 | 19.2079556455884 |
| chr17 | 47622690 | 47622704 | + | 15.0594086531005 | 21.6029190992233 | 46.5359002043516 | 26.6777161744284 |
| chr17 | 47622761 | 47622765 | + | 5.01980288436682 | 3.41098722619315 | 5.96614105183995 | 1.06710864697714 |
| chr17 | 47622794 | 47622895 | + | 67.2653586505154 | 57.9867828452835 | 36.9900745214077 | 23.476390233497 |
| chr17 | 47622929 | 47622955 | + | 4.01584230749346 | 3.41098722619315 | 2.38645642073598 | 4.26843458790854 |
| chr17 | 47623140 | 47623190 | + | 4.01584230749346 | 4.54798296825753 | 7.15936926220794 | 1.06710864697714 |
| chr17 | 47623241 | 47623251 | + | 12.0475269224804 | 12.5069531627082 | 4.77291284147196 | 5.33554323488568 |
| chr17 | 47682413 | 47682443 | + | 6.02376346124019 | 9.09596593651506 | 11.9322821036799 | 11.7381951167485 |
| chr17 | 47682472 | 47682487 | + | 5.01980288436682 | 6.82197445238629 | 11.9322821036799 | 16.006629704657 |
| chr17 | 47682568 | 47682594 | + | 6.02376346124019 | 5.68497871032191 | 7.15936926220794 | 11.7381951167485 |
| chr17 | 47682664 | 47682690 | + | 1.00396057687336 | 11.3699574206438 | 3.57968463110397 | 4.26843458790854 |
| chr17 | 47682693 | 47682719 | + | 2.00792115374673 | 2.27399148412876 | 5.96614105183995 | 4.26843458790854 |
| chr17 | 47682743 | 47682758 | + | 3.01188173062009 | 3.41098722619315 | 2.38645642073598 | 4.26843458790854 |
| chr17 | 47682850 | 47682862 | + | 7.02772403811355 | 3.41098722619315 | 3.57968463110397 | 6.40265188186281 |
| chr17 | 47682892 | 47682927 | + | 187.740627875319 | 236.495114349391 | 171.82486229299 | 401.232851263403 |
| chr17 | 47682962 | 47682989 | + | 133.526756724157 | 138.713480531855 | 206.428480393662 | 101.375321462828 |
| chr17 | 47683030 | 47683037 | + | 3.01188173062009 | 2.27399148412876 | 2.38645642073598 | 4.26843458790854 |
| chr17 | 47683118 | 47683172 | + | 35.1386201905678 | 39.7948509722534 | 28.6374770488317 | 48.0198891139711 |
| chr17 | 47683292 | 47683377 | + | 53.2099105742883 | 95.5076423334081 | 46.5359002043516 | 38.4159112911769 |
| chr17 | 47683442 | 47683449 | + | 1.00396057687336 | 1.13699574206438 | 2.38645642073598 | 3.20132594093141 |
| chr17 | 47683936 | 47683945 | + | 12.0475269224804 | 2.27399148412876 | 3.57968463110397 | 16.006629704657 |
| chr17 | 47684227 | 47684233 | + | 6.02376346124019 | 3.41098722619315 | 3.57968463110397 | 3.20132594093141 |
| chr17 | 47684319 | 47684327 | + | 4.01584230749346 | 3.41098722619315 | 3.57968463110397 | 2.13421729395427 |
| chr17 | 47684428 | 47684472 | + | 7.02772403811355 | 14.780944646837 | 16.7051949451519 | 6.40265188186281 |
| chr17 | 47684515 | 47684563 | + | 31.1227778830743 | 27.2878978095452 | 21.4781077866238 | 28.8119334683827 |
| chr17 | 47684618 | 47684651 | + | 8.03168461498692 | 13.6439489047726 | 2.38645642073598 | 10.6710864697714 |
| chr17 | 47823485 | 47823539 | - | 28.1108961524542 | 32.9728765198671 | 21.4781077866238 | 39.483019938154 |
| chr17 | 47823677 | 47823713 | - | 1.00396057687336 | 1.13699574206438 | 1.19322821036799 | 2.13421729395427 |
| chr17 | 47823763 | 47823824 | - | 10.0396057687336 | 12.5069531627082 | 10.7390538933119 | 18.1408469986113 |
| chr17 | 47823898 | 47823958 | - | 9.03564519186028 | 2.27399148412876 | 8.35259747257593 | 9.60397782279422 |
| chr17 | 47824045 | 47824139 | - | 22.087132691214 | 20.4659233571589 | 27.4442488384638 | 20.2750642925656 |
| chr17 | 47837865 | 47837874 | - | 6.02376346124019 | 3.41098722619315 | 2.38645642073598 | 2.13421729395427 |
| chr17 | 47838105 | 47838107 | - | 1.00396057687336 | 1.13699574206438 | 5.96614105183995 | 4.26843458790854 |
| chr17 | 47845100 | 47845118 | - | 1.00396057687336 | 5.68497871032191 | 39.3765309421437 | 93.9055609339879 |
| chr17 | 47928543 | 47928550 | + | 4.01584230749346 | 5.68497871032191 | 2.38645642073598 | 5.33554323488568 |
| chr17 | 47946814 | 47946834 | + | 1.00396057687336 | 2.27399148412876 | 4.77291284147196 | 1.06710864697714 |
| chr17 | 47947393 | 47947409 | + | 3.01188173062009 | 2.27399148412876 | 2.38645642073598 | 10.6710864697714 |
| chr17 | 47947472 | 47947500 | + | 5.01980288436682 | 2.27399148412876 | 4.77291284147196 | 4.26843458790854 |
| chr17 | 47947803 | 47947809 | + | 3.01188173062009 | 1.13699574206438 | 1.19322821036799 | 2.13421729395427 |
| chr17 | 47947923 | 47947943 | + | 2.00792115374673 | 4.54798296825753 | 1.19322821036799 | 1.06710864697714 |
| chr17 | 47948051 | 47948057 | + | 1.00396057687336 | 1.13699574206438 | 3.57968463110397 | 1.06710864697714 |
| chr17 | 47948144 | 47948179 | + | 7.02772403811355 | 7.95897019445067 | 8.35259747257593 | 13.8724124107028 |
| chr17 | 47949074 | 47949086 | + | 10.0396057687336 | 3.41098722619315 | 2.38645642073598 | 5.33554323488568 |
| chr17 | 47971343 | 47971368 | + | 4.01584230749346 | 11.3699574206438 | 7.15936926220794 | 4.26843458790854 |
| chr17 | 47974021 | 47974032 | + | 2.00792115374673 | 1.13699574206438 | 1.19322821036799 | 5.33554323488568 |
| chr17 | 47974403 | 47974409 | + | 2.00792115374673 | 2.27399148412876 | 4.77291284147196 | 2.13421729395427 |
| chr17 | 47975102 | 47975179 | + | 8.03168461498692 | 9.09596593651506 | 46.5359002043516 | 9.60397782279422 |
| chr17 | 47975267 | 47975284 | + | 9.03564519186028 | 3.41098722619315 | 2.38645642073598 | 12.8053037637256 |
| chr17 | 47975510 | 47975515 | + | 2.00792115374673 | 1.13699574206438 | 2.38645642073598 | 4.26843458790854 |
| chr17 | 47975573 | 47975654 | + | 32.1267384599477 | 10.2329616785794 | 36.9900745214077 | 14.9395210576799 |
| chr17 | 47975874 | 47975882 | + | 2.00792115374673 | 2.27399148412876 | 3.57968463110397 | 3.20132594093141 |
| chr17 | 47975936 | 47975984 | + | 4.01584230749346 | 10.2329616785794 | 29.8307052591997 | 9.60397782279422 |
| chr17 | 47976729 | 47976757 | + | 5.01980288436682 | 7.95897019445067 | 9.54582568294392 | 5.33554323488568 |
| chr17 | 47976793 | 47976823 | + | 5.01980288436682 | 4.54798296825753 | 7.15936926220794 | 2.13421729395427 |
| chr17 | 47977829 | 47977857 | + | 5.01980288436682 | 4.54798296825753 | 8.35259747257593 | 9.60397782279422 |
| chr17 | 47978847 | 47978855 | + | 4.01584230749346 | 1.13699574206438 | 5.96614105183995 | 2.13421729395427 |
| chr17 | 47978878 | 47978884 | + | 1.00396057687336 | 1.13699574206438 | 2.38645642073598 | 1.06710864697714 |
| chr17 | 47980611 | 47980635 | + | 2.00792115374673 | 5.68497871032191 | 19.0916513658878 | 7.46976052883995 |
| chr17 | 47980700 | 47980743 | + | 7.02772403811355 | 4.54798296825753 | 23.8645642073598 | 12.8053037637256 |
| chr17 | 47981538 | 47981561 | + | 2.00792115374673 | 2.27399148412876 | 1.19322821036799 | 1.06710864697714 |
| chr17 | 47981683 | 47981712 | + | 12.0475269224804 | 14.780944646837 | 9.54582568294392 | 16.006629704657 |
| chr17 | 48059604 | 48060150 | + | 169.669337491599 | 244.454084543842 | 137.221244192319 | 120.583277108416 |
| chr17 | 48060182 | 48060229 | + | 101.40001826421 | 155.76841666282 | 75.1733772531833 | 36.2816939972226 |
| chr17 | 48060256 | 48060404 | + | 19.0752509605939 | 20.4659233571589 | 29.8307052591997 | 6.40265188186281 |
| chr17 | 48060461 | 48060610 | + | 21.0831721143407 | 26.1509020674808 | 19.0916513658878 | 24.5434988804741 |
| chr17 | 48060641 | 48060753 | + | 18.0712903837206 | 21.6029190992233 | 17.8984231555198 | 24.5434988804741 |
| chr17 | 48060793 | 48061041 | + | 80.3168461498692 | 96.6446380754725 | 40.5697591525116 | 32.0132594093141 |
| chr17 | 48061136 | 48061152 | + | 4.01584230749346 | 10.2329616785794 | 3.57968463110397 | 7.46976052883995 |
| chr17 | 48061338 | 48061355 | + | 5.01980288436682 | 13.6439489047726 | 5.96614105183995 | 8.53686917581709 |
| chr17 | 48070167 | 48070173 | - | 2.00792115374673 | 2.27399148412876 | 2.38645642073598 | 1.06710864697714 |
| chr17 | 48070964 | 48070988 | - | 206.815878835913 | 221.714169702554 | 249.38469596691 | 335.072115150821 |
| chr17 | 48121956 | 48121964 | + | 3.01188173062009 | 2.27399148412876 | 2.38645642073598 | 2.13421729395427 |
| chr17 | 48121988 | 48122007 | + | 10.0396057687336 | 13.6439489047726 | 22.6713359969918 | 23.476390233497 |
| chr17 | 48122232 | 48122233 | + | 3.01188173062009 | 5.68497871032191 | 19.0916513658878 | 20.2750642925656 |
| chr17 | 48122389 | 48122400 | + | 1.00396057687336 | 2.27399148412876 | 1.19322821036799 | 2.13421729395427 |
| chr17 | 48122712 | 48122722 | + | 4.01584230749346 | 7.95897019445067 | 5.96614105183995 | 1.06710864697714 |
| chr17 | 48549976 | 48550041 | - | 1.00396057687336 | 1.13699574206438 | 17.8984231555198 | 21.3421729395427 |
| chr17 | 48576559 | 48576575 | - | 3.01188173062009 | 1.13699574206438 | 115.743136405695 | 77.8989312293309 |
| chr17 | 48576620 | 48576665 | - | 1.00396057687336 | 1.13699574206438 | 34.6036181006717 | 36.2816939972226 |
| chr17 | 48607246 | 48607263 | - | 18.0712903837206 | 36.3838637460602 | 3.57968463110397 | 7.46976052883995 |
| chr17 | 48607346 | 48607384 | - | 160.633692299738 | 139.850476273919 | 21.4781077866238 | 33.0803680562912 |
| chr17 | 48607425 | 48607445 | - | 10.0396057687336 | 9.09596593651506 | 1.19322821036799 | 1.06710864697714 |
| chr17 | 48607602 | 48607631 | - | 17.0673298068472 | 19.3289276150945 | 4.77291284147196 | 4.26843458790854 |
| chr17 | 48607653 | 48607666 | - | 22.087132691214 | 47.753821166704 | 5.96614105183995 | 7.46976052883995 |
| chr17 | 48607747 | 48607933 | - | 193.764391336559 | 249.0020675121 | 94.2650286190712 | 70.429170700491 |
| chr17 | 48863626 | 48863639 | + | 3.01188173062009 | 3.41098722619315 | 1.19322821036799 | 1.06710864697714 |
| chr17 | 48863705 | 48863770 | + | 4.01584230749346 | 3.41098722619315 | 3.57968463110397 | 1.06710864697714 |
| chr17 | 48895780 | 48895798 | + | 20.0792115374673 | 22.7399148412876 | 23.8645642073598 | 43.7514545260626 |
| chr17 | 48895823 | 48895837 | + | 128.506953839791 | 98.9186295596012 | 82.3327465153913 | 125.918820343302 |
| chr17 | 48927287 | 48927308 | + | 5.01980288436682 | 2.27399148412876 | 7.15936926220794 | 1.06710864697714 |
| chr17 | 48927365 | 48927405 | + | 13.0514874993537 | 18.1919318730301 | 7.15936926220794 | 18.1408469986113 |
| chr17 | 48927443 | 48927623 | + | 142.562401916018 | 168.275369825529 | 107.390538933119 | 68.2949534065367 |
| chr17 | 48927664 | 48927670 | + | 2.00792115374673 | 3.41098722619315 | 1.19322821036799 | 2.13421729395427 |
| chr17 | 48928176 | 48928200 | + | 11.043566345607 | 2.27399148412876 | 1.19322821036799 | 2.13421729395427 |
| chr17 | 48928942 | 48928988 | + | 9.03564519186028 | 10.2329616785794 | 5.96614105183995 | 6.40265188186281 |
| chr17 | 48930410 | 48930467 | - | 288.136685562656 | 278.563956805774 | 151.539982716735 | 317.998376799186 |
| chr17 | 49049663 | 49049757 | + | 8.03168461498692 | 6.82197445238629 | 9.54582568294392 | 18.1408469986113 |
| chr17 | 49049791 | 49049828 | + | 2.00792115374673 | 3.41098722619315 | 7.15936926220794 | 20.2750642925656 |
| chr17 | 49049873 | 49049877 | + | 3.01188173062009 | 2.27399148412876 | 4.77291284147196 | 1.06710864697714 |
| chr17 | 49050285 | 49050332 | + | 4.01584230749346 | 5.68497871032191 | 10.7390538933119 | 4.26843458790854 |
| chr17 | 49050453 | 49050623 | + | 9.03564519186028 | 10.2329616785794 | 39.3765309421437 | 13.8724124107028 |
| chr17 | 49050673 | 49050754 | + | 6.02376346124019 | 7.95897019445067 | 20.2848795762558 | 24.5434988804741 |
| chr17 | 49050781 | 49050826 | + | 4.01584230749346 | 2.27399148412876 | 16.7051949451519 | 28.8119334683827 |
| chr17 | 49050877 | 49051138 | + | 145.574283646638 | 202.38524208746 | 620.478669391355 | 1443.79799936006 |
| chr17 | 49051214 | 49051224 | + | 1.00396057687336 | 1.13699574206438 | 2.38645642073598 | 1.06710864697714 |
| chr17 | 49052480 | 49052504 | + | 1.00396057687336 | 2.27399148412876 | 1.19322821036799 | 3.20132594093141 |
| chr17 | 49052573 | 49052629 | + | 1.00396057687336 | 2.27399148412876 | 1.19322821036799 | 4.26843458790854 |
| chr17 | 49055099 | 49055119 | + | 1.00396057687336 | 1.13699574206438 | 8.35259747257593 | 6.40265188186281 |
| chr17 | 49055483 | 49055532 | + | 10.0396057687336 | 17.0549361309657 | 90.6853439879672 | 94.9726695809651 |
| chr17 | 49222859 | 49222860 | + | 3.01188173062009 | 5.68497871032191 | 7.15936926220794 | 11.7381951167485 |
| chr17 | 49294660 | 49294744 | - | 97.3841759567164 | 105.740604011988 | 47.7291284147196 | 23.476390233497 |
| chr17 | 49294987 | 49295008 | - | 2.00792115374673 | 5.68497871032191 | 1.19322821036799 | 1.06710864697714 |
| chr17 | 49297423 | 49297443 | - | 15.0594086531005 | 5.68497871032191 | 2.38645642073598 | 2.13421729395427 |
| chr17 | 49361159 | 49361186 | + | 6.02376346124019 | 5.68497871032191 | 1.19322821036799 | 2.13421729395427 |
| chr17 | 49404115 | 49404149 | - | 226.89509037338 | 89.8226636230862 | 78.7530618842873 | 69.3620620535138 |
| chr17 | 49404174 | 49404205 | - | 39.1544624980612 | 10.2329616785794 | 14.3187385244159 | 38.4159112911769 |
| chr17 | 49404280 | 49404382 | - | 39.1544624980612 | 23.876910583352 | 17.8984231555198 | 20.2750642925656 |
| chr17 | 49404474 | 49404500 | - | 5.01980288436682 | 3.41098722619315 | 7.15936926220794 | 5.33554323488568 |
| chr17 | 49404588 | 49404676 | - | 9.03564519186028 | 5.68497871032191 | 23.8645642073598 | 20.2750642925656 |
| chr17 | 49404939 | 49404990 | - | 208.82379998966 | 54.5757956190903 | 58.4681823080315 | 184.609795927044 |
| chr17 | 49599336 | 49599366 | - | 95.3762548029696 | 259.235029190679 | 76.3666054635513 | 100.308212815851 |
| chr17 | 49599574 | 49599590 | - | 1.00396057687336 | 6.82197445238629 | 1.19322821036799 | 8.53686917581709 |
| chr17 | 49599657 | 49599659 | - | 5.01980288436682 | 6.82197445238629 | 3.57968463110397 | 4.26843458790854 |
| chr17 | 49599712 | 49599735 | - | 6.02376346124019 | 11.3699574206438 | 15.5119667347839 | 12.8053037637256 |
| chr17 | 49599968 | 49600002 | - | 11.043566345607 | 12.5069531627082 | 7.15936926220794 | 10.6710864697714 |
| chr17 | 49600086 | 49600092 | - | 11.043566345607 | 6.82197445238629 | 5.96614105183995 | 6.40265188186281 |
| chr17 | 49600261 | 49600271 | - | 2.00792115374673 | 3.41098722619315 | 3.57968463110397 | 2.13421729395427 |
| chr17 | 49701358 | 49701407 | - | 9.03564519186028 | 13.6439489047726 | 3.57968463110397 | 5.33554323488568 |
| chr17 | 49701448 | 49701455 | - | 2.00792115374673 | 2.27399148412876 | 4.77291284147196 | 4.26843458790854 |
| chr17 | 49828894 | 49828901 | + | 4.01584230749346 | 1.13699574206438 | 3.57968463110397 | 1.06710864697714 |
| chr17 | 49828985 | 49829002 | + | 3.01188173062009 | 11.3699574206438 | 3.57968463110397 | 1.06710864697714 |
| chr17 | 49829042 | 49829053 | + | 13.0514874993537 | 32.9728765198671 | 15.5119667347839 | 7.46976052883995 |
| chr17 | 50134300 | 50134302 | - | 2.00792115374673 | 5.68497871032191 | 1.19322821036799 | 2.13421729395427 |
| chr17 | 50134332 | 50134353 | - | 11.043566345607 | 3.41098722619315 | 8.35259747257593 | 7.46976052883995 |
| chr17 | 50134443 | 50134487 | - | 6.02376346124019 | 1.13699574206438 | 10.7390538933119 | 12.8053037637256 |
| chr17 | 50134536 | 50134549 | - | 4.01584230749346 | 2.27399148412876 | 2.38645642073598 | 3.20132594093141 |
| chr17 | 50134725 | 50134760 | - | 5.01980288436682 | 2.27399148412876 | 1.19322821036799 | 1.06710864697714 |
| chr17 | 50185373 | 50185397 | - | 19.0752509605939 | 42.0688424563821 | 2.38645642073598 | 1.06710864697714 |
| chr17 | 50185445 | 50185497 | - | 68.2693192273888 | 43.2058381984465 | 1.19322821036799 | 1.06710864697714 |
| chr17 | 50361103 | 50361111 | + | 2.00792115374673 | 1.13699574206438 | 1.19322821036799 | 1.06710864697714 |
| chr17 | 50367884 | 50367948 | - | 14.0554480762271 | 13.6439489047726 | 2.38645642073598 | 16.006629704657 |
| chr17 | 50367974 | 50368145 | - | 272.073316332682 | 205.796229313653 | 158.699351978943 | 268.911379038238 |
| chr17 | 50368164 | 50368301 | - | 81.3208067267425 | 71.6307317500561 | 71.5936926220794 | 96.0397782279422 |
| chr17 | 50381301 | 50381358 | - | 11.043566345607 | 6.82197445238629 | 9.54582568294392 | 23.476390233497 |
| chr17 | 50381551 | 50381661 | - | 67.2653586505154 | 39.7948509722534 | 139.607700613055 | 137.657015460051 |
| chr17 | 50381737 | 50381747 | - | 3.01188173062009 | 2.27399148412876 | 8.35259747257593 | 8.53686917581709 |
| chr17 | 50381784 | 50381925 | - | 40.1584230749346 | 14.780944646837 | 53.6952694665595 | 35.2145853502455 |
| chr17 | 50381973 | 50382025 | - | 7.02772403811355 | 2.27399148412876 | 4.77291284147196 | 23.476390233497 |
| chr17 | 50382091 | 50382123 | - | 3.01188173062009 | 2.27399148412876 | 5.96614105183995 | 9.60397782279422 |
| chr17 | 50382210 | 50382249 | - | 3.01188173062009 | 4.54798296825753 | 11.9322821036799 | 34.1474767032683 |
| chr17 | 50382314 | 50382357 | - | 12.0475269224804 | 4.54798296825753 | 11.9322821036799 | 7.46976052883995 |
| chr17 | 50382436 | 50382449 | - | 4.01584230749346 | 2.27399148412876 | 2.38645642073598 | 2.13421729395427 |
| chr17 | 50382479 | 50382537 | - | 24.0950538449607 | 14.780944646837 | 76.3666054635513 | 30.9461507623369 |
| chr17 | 50382564 | 50382589 | - | 36.1425807674411 | 22.7399148412876 | 51.3088130458235 | 56.5567582897882 |
| chr17 | 50382649 | 50382688 | - | 14.0554480762271 | 4.54798296825753 | 3.57968463110397 | 9.60397782279422 |
| chr17 | 50382719 | 50382797 | - | 115.455466340437 | 73.9047232341848 | 156.312895558207 | 136.589906813073 |
| chr17 | 50382818 | 50382823 | - | 2.00792115374673 | 2.27399148412876 | 9.54582568294392 | 4.26843458790854 |
| chr17 | 50382859 | 50382880 | - | 5.01980288436682 | 2.27399148412876 | 8.35259747257593 | 26.6777161744284 |
| chr17 | 50382950 | 50382962 | - | 3.01188173062009 | 6.82197445238629 | 8.35259747257593 | 12.8053037637256 |
| chr17 | 50485075 | 50485099 | + | 8.03168461498692 | 4.54798296825753 | 9.54582568294392 | 10.6710864697714 |
| chr17 | 50485273 | 50485301 | + | 1.00396057687336 | 4.54798296825753 | 10.7390538933119 | 18.1408469986113 |
| chr17 | 50485335 | 50485372 | + | 3.01188173062009 | 1.13699574206438 | 7.15936926220794 | 9.60397782279422 |
| chr17 | 50485621 | 50485648 | + | 4.01584230749346 | 1.13699574206438 | 7.15936926220794 | 6.40265188186281 |
| chr17 | 50485838 | 50485889 | + | 5.01980288436682 | 6.82197445238629 | 7.15936926220794 | 17.0737383516342 |
| chr17 | 50693379 | 50693400 | - | 3.01188173062009 | 5.68497871032191 | 1.19322821036799 | 1.06710864697714 |
| chr17 | 50693537 | 50693568 | - | 1.00396057687336 | 9.09596593651506 | 1.19322821036799 | 3.20132594093141 |
| chr17 | 50695617 | 50695624 | - | 12.0475269224804 | 12.5069531627082 | 4.77291284147196 | 4.26843458790854 |
| chr17 | 50695662 | 50695670 | - | 9.03564519186028 | 9.09596593651506 | 2.38645642073598 | 3.20132594093141 |
| chr17 | 50695734 | 50695758 | - | 6.02376346124019 | 6.82197445238629 | 1.19322821036799 | 3.20132594093141 |
| chr17 | 50695795 | 50695803 | - | 2.00792115374673 | 1.13699574206438 | 3.57968463110397 | 1.06710864697714 |
| chr17 | 50695825 | 50695827 | - | 3.01188173062009 | 2.27399148412876 | 1.19322821036799 | 1.06710864697714 |
| chr17 | 50750690 | 50750736 | + | 6.02376346124019 | 10.2329616785794 | 10.7390538933119 | 9.60397782279422 |
| chr17 | 50750817 | 50750855 | + | 5.01980288436682 | 4.54798296825753 | 11.9322821036799 | 2.13421729395427 |
| chr17 | 50750885 | 50750987 | + | 25.0990144218341 | 44.3428339405109 | 69.2072362013434 | 29.8790421153598 |
| chr17 | 50751009 | 50751018 | + | 4.01584230749346 | 11.3699574206438 | 14.3187385244159 | 1.06710864697714 |
| chr17 | 50751156 | 50751173 | + | 1.00396057687336 | 4.54798296825753 | 1.19322821036799 | 2.13421729395427 |
| chr17 | 50751324 | 50751376 | + | 2.00792115374673 | 1.13699574206438 | 14.3187385244159 | 18.1408469986113 |
| chr17 | 50751472 | 50751497 | + | 5.01980288436682 | 3.41098722619315 | 7.15936926220794 | 8.53686917581709 |
| chr17 | 50751902 | 50751979 | + | 3.01188173062009 | 2.27399148412876 | 1.19322821036799 | 4.26843458790854 |
| chr17 | 50752190 | 50752313 | + | 40.1584230749346 | 52.3018041349616 | 29.8307052591997 | 30.9461507623369 |
| chr17 | 50752353 | 50752355 | + | 2.00792115374673 | 1.13699574206438 | 2.38645642073598 | 1.06710864697714 |
| chr17 | 50752385 | 50752425 | + | 14.0554480762271 | 5.68497871032191 | 10.7390538933119 | 2.13421729395427 |
| chr17 | 50752460 | 50752670 | + | 20.0792115374673 | 30.6988850357383 | 23.8645642073598 | 7.46976052883995 |
| chr17 | 50862262 | 50862294 | - | 60.2376346124019 | 117.110561432631 | 5.96614105183995 | 5.33554323488568 |
| chr17 | 50862444 | 50862486 | - | 16.0633692299738 | 25.0139063254164 | 1.19322821036799 | 3.20132594093141 |
| chr17 | 50862629 | 50862648 | - | 22.087132691214 | 55.7127913611547 | 5.96614105183995 | 2.13421729395427 |
| chr17 | 50862683 | 50862693 | - | 33.130699036821 | 34.1098722619315 | 2.38645642073598 | 3.20132594093141 |
| chr17 | 50862728 | 50862741 | - | 8.03168461498692 | 27.2878978095452 | 4.77291284147196 | 3.20132594093141 |
| chr17 | 50862933 | 50862951 | - | 5.01980288436682 | 2.27399148412876 | 1.19322821036799 | 7.46976052883995 |
| chr17 | 50866742 | 50866773 | + | 8.03168461498692 | 9.09596593651506 | 3.57968463110397 | 6.40265188186281 |
| chr17 | 50962219 | 50962230 | - | 4.01584230749346 | 6.82197445238629 | 1.19322821036799 | 2.13421729395427 |
| chr17 | 50962310 | 50962312 | - | 2.00792115374673 | 2.27399148412876 | 1.19322821036799 | 3.20132594093141 |
| chr17 | 50964614 | 50964621 | - | 2.00792115374673 | 3.41098722619315 | 3.57968463110397 | 11.7381951167485 |
| chr17 | 50965143 | 50965160 | - | 1.00396057687336 | 1.13699574206438 | 3.57968463110397 | 3.20132594093141 |
| chr17 | 50965288 | 50965326 | - | 9.03564519186028 | 9.09596593651506 | 13.1255103140479 | 9.60397782279422 |
| chr17 | 50965419 | 50965432 | - | 1.00396057687336 | 1.13699574206438 | 3.57968463110397 | 3.20132594093141 |
| chr17 | 50965463 | 50965492 | - | 3.01188173062009 | 10.2329616785794 | 8.35259747257593 | 5.33554323488568 |
| chr17 | 50965668 | 50965692 | - | 9.03564519186028 | 17.0549361309657 | 21.4781077866238 | 6.40265188186281 |
| chr17 | 50966221 | 50966278 | - | 7.02772403811355 | 4.54798296825753 | 14.3187385244159 | 7.46976052883995 |
| chr17 | 51161170 | 51161210 | + | 12.0475269224804 | 2.27399148412876 | 10.7390538933119 | 22.4092815865198 |
| chr17 | 51161233 | 51161269 | + | 37.1465413443145 | 20.4659233571589 | 13.1255103140479 | 23.476390233497 |
| chr17 | 51161725 | 51161827 | + | 19.0752509605939 | 13.6439489047726 | 15.5119667347839 | 43.7514545260626 |
| chr17 | 51161846 | 51161860 | + | 13.0514874993537 | 23.876910583352 | 13.1255103140479 | 52.2883237018796 |
| chr17 | 51161889 | 51162013 | + | 105.415860571703 | 136.439489047726 | 89.4921157775992 | 187.811121867976 |
| chr17 | 51162032 | 51162052 | + | 1.00396057687336 | 1.13699574206438 | 4.77291284147196 | 1.06710864697714 |
| chr17 | 51171518 | 51171542 | + | 5.01980288436682 | 3.41098722619315 | 1.19322821036799 | 3.20132594093141 |
| chr17 | 51171591 | 51171672 | + | 80.3168461498692 | 73.9047232341848 | 41.7629873628796 | 66.1607361125824 |
| chr17 | 51171714 | 51171726 | + | 13.0514874993537 | 13.6439489047726 | 7.15936926220794 | 1.06710864697714 |
| chr17 | 51177738 | 51177760 | - | 4.01584230749346 | 2.27399148412876 | 2.38645642073598 | 3.20132594093141 |
| chr17 | 51178304 | 51178324 | - | 1.00396057687336 | 2.27399148412876 | 9.54582568294392 | 2.13421729395427 |
| chr17 | 51178463 | 51178495 | - | 2.00792115374673 | 1.13699574206438 | 9.54582568294392 | 8.53686917581709 |
| chr17 | 51180401 | 51180414 | - | 11.043566345607 | 20.4659233571589 | 56.0817258872955 | 67.2278447595595 |
| chr17 | 54937419 | 54937426 | + | 12.0475269224804 | 10.2329616785794 | 1.19322821036799 | 5.33554323488568 |
| chr17 | 54937577 | 54937616 | + | 7.02772403811355 | 7.95897019445067 | 2.38645642073598 | 5.33554323488568 |
| chr17 | 54937906 | 54937908 | + | 1.00396057687336 | 4.54798296825753 | 1.19322821036799 | 2.13421729395427 |
| chr17 | 55160647 | 55160652 | + | 1.00396057687336 | 1.13699574206438 | 1.19322821036799 | 1.06710864697714 |
| chr17 | 55392815 | 55393237 | - | 3.01188173062009 | 2.27399148412876 | 7.15936926220794 | 7.46976052883995 |
| chr17 | 55393848 | 55394111 | - | 1.00396057687336 | 4.54798296825753 | 11.9322821036799 | 9.60397782279422 |
| chr17 | 55394252 | 55394359 | - | 6.02376346124019 | 2.27399148412876 | 14.3187385244159 | 13.8724124107028 |
| chr17 | 55776372 | 55776433 | + | 3.01188173062009 | 2.27399148412876 | 23.8645642073598 | 18.1408469986113 |
| chr17 | 55777204 | 55777206 | + | 1.00396057687336 | 1.13699574206438 | 3.57968463110397 | 1.06710864697714 |
| chr17 | 55777297 | 55777300 | + | 1.00396057687336 | 2.27399148412876 | 2.38645642073598 | 3.20132594093141 |
| chr17 | 56863178 | 56863181 | + | 5.01980288436682 | 2.27399148412876 | 3.57968463110397 | 1.06710864697714 |
| chr17 | 56863657 | 56863684 | + | 3.01188173062009 | 4.54798296825753 | 4.77291284147196 | 1.06710864697714 |
| chr17 | 56863735 | 56863760 | + | 15.0594086531005 | 28.4248935516096 | 20.2848795762558 | 3.20132594093141 |
| chr17 | 56864496 | 56864534 | + | 4.01584230749346 | 3.41098722619315 | 5.96614105183995 | 6.40265188186281 |
| chr17 | 56864566 | 56864582 | + | 11.043566345607 | 14.780944646837 | 7.15936926220794 | 7.46976052883995 |
| chr17 | 56887941 | 56888015 | - | 23.0910932680874 | 13.6439489047726 | 7.15936926220794 | 11.7381951167485 |
| chr17 | 56888791 | 56888804 | - | 28.1108961524542 | 19.3289276150945 | 1.19322821036799 | 4.26843458790854 |
| chr17 | 56891284 | 56891307 | - | 12.0475269224804 | 5.68497871032191 | 11.9322821036799 | 5.33554323488568 |
| chr17 | 56891472 | 56891479 | - | 4.01584230749346 | 2.27399148412876 | 3.57968463110397 | 5.33554323488568 |
| chr17 | 56891504 | 56891522 | - | 18.0712903837206 | 9.09596593651506 | 5.96614105183995 | 3.20132594093141 |
| chr17 | 56891540 | 56891580 | - | 18.0712903837206 | 7.95897019445067 | 4.77291284147196 | 3.20132594093141 |
| chr17 | 57120752 | 57120791 | + | 2.00792115374673 | 2.27399148412876 | 3.57968463110397 | 2.13421729395427 |
| chr17 | 57120921 | 57120933 | + | 16.0633692299738 | 6.82197445238629 | 2.38645642073598 | 5.33554323488568 |
| chr17 | 57121126 | 57121214 | + | 22.087132691214 | 6.82197445238629 | 4.77291284147196 | 14.9395210576799 |
| chr17 | 57679567 | 57679590 | + | 2.00792115374673 | 6.82197445238629 | 1.19322821036799 | 7.46976052883995 |
| chr17 | 57679654 | 57679685 | + | 15.0594086531005 | 14.780944646837 | 3.57968463110397 | 8.53686917581709 |
| chr17 | 57680094 | 57680117 | + | 76.3010038423757 | 78.4527062024424 | 47.7291284147196 | 49.0869977609482 |
| chr17 | 57680138 | 57680302 | + | 14.0554480762271 | 11.3699574206438 | 25.0577924177278 | 26.6777161744284 |
| chr17 | 57680454 | 57680457 | + | 1.00396057687336 | 2.27399148412876 | 1.19322821036799 | 1.06710864697714 |
| chr17 | 57680765 | 57680778 | + | 610.408030739006 | 662.868517623535 | 326.944529640829 | 194.213773749839 |
| chr17 | 57680904 | 57680939 | + | 23.0910932680874 | 15.9179403889013 | 31.0239334695677 | 28.8119334683827 |
| chr17 | 57680972 | 57681001 | + | 9.03564519186028 | 7.95897019445067 | 9.54582568294392 | 3.20132594093141 |
| chr17 | 57681023 | 57681089 | + | 62.2455557661486 | 43.2058381984465 | 69.2072362013434 | 134.455689519119 |
| chr17 | 57681123 | 57681228 | + | 538.122869204123 | 624.210662393346 | 442.687666046524 | 679.748208124435 |
| chr17 | 57681257 | 57681332 | + | 100.396057687336 | 89.8226636230862 | 121.709277457535 | 104.576647403759 |
| chr17 | 57681405 | 57681462 | + | 3.01188173062009 | 7.95897019445067 | 9.54582568294392 | 7.46976052883995 |
| chr17 | 57681519 | 57681558 | + | 28.1108961524542 | 38.657855230189 | 21.4781077866238 | 43.7514545260626 |
| chr17 | 57681674 | 57681691 | + | 9.03564519186028 | 4.54798296825753 | 5.96614105183995 | 4.26843458790854 |
| chr17 | 57682332 | 57682384 | + | 14.0554480762271 | 7.95897019445067 | 15.5119667347839 | 50.1541064079254 |
| chr17 | 57683216 | 57683281 | + | 1.00396057687336 | 4.54798296825753 | 3.57968463110397 | 10.6710864697714 |
| chr17 | 57683428 | 57683439 | + | 5.01980288436682 | 5.68497871032191 | 2.38645642073598 | 5.33554323488568 |
| chr17 | 57683654 | 57683672 | + | 25.0990144218341 | 26.1509020674808 | 17.8984231555198 | 39.483019938154 |
| chr17 | 57684348 | 57684355 | + | 5.01980288436682 | 2.27399148412876 | 2.38645642073598 | 7.46976052883995 |
| chr17 | 57839509 | 57839519 | - | 1.00396057687336 | 2.27399148412876 | 5.96614105183995 | 3.20132594093141 |
| chr17 | 57839556 | 57839837 | - | 20.0792115374673 | 26.1509020674808 | 20.2848795762558 | 22.4092815865198 |
| chr17 | 57974077 | 57974083 | - | 3.01188173062009 | 3.41098722619315 | 3.57968463110397 | 1.06710864697714 |
| chr17 | 57989058 | 57989147 | - | 3.01188173062009 | 7.95897019445067 | 35.7968463110397 | 9.60397782279422 |
| chr17 | 58003515 | 58003565 | - | 21.0831721143407 | 14.780944646837 | 14.3187385244159 | 1.06710864697714 |
| chr17 | 58003589 | 58003726 | - | 116.45942691731 | 92.0966551072149 | 50.1155848354556 | 117.381951167485 |
| chr17 | 58003747 | 58003774 | - | 15.0594086531005 | 11.3699574206438 | 4.77291284147196 | 19.2079556455884 |
| chr17 | 58003801 | 58003996 | - | 96.380215379843 | 69.3567402659273 | 99.0379414605431 | 136.589906813073 |
| chr17 | 58004019 | 58004067 | - | 2.00792115374673 | 4.54798296825753 | 9.54582568294392 | 6.40265188186281 |
| chr17 | 58004315 | 58004334 | - | 3.01188173062009 | 1.13699574206438 | 3.57968463110397 | 3.20132594093141 |
| chr17 | 58004346 | 58004348 | - | 1.00396057687336 | 3.41098722619315 | 2.38645642073598 | 1.06710864697714 |
| chr17 | 58004441 | 58004465 | - | 9.03564519186028 | 3.41098722619315 | 5.96614105183995 | 2.13421729395427 |
| chr17 | 58004519 | 58004545 | - | 5.01980288436682 | 5.68497871032191 | 3.57968463110397 | 2.13421729395427 |
| chr17 | 58004576 | 58004623 | - | 17.0673298068472 | 10.2329616785794 | 3.57968463110397 | 19.2079556455884 |
| chr17 | 58004651 | 58004699 | - | 26.1029749987075 | 17.0549361309657 | 31.0239334695677 | 21.3421729395427 |
| chr17 | 58004726 | 58004766 | - | 7.02772403811355 | 7.95897019445067 | 10.7390538933119 | 5.33554323488568 |
| chr17 | 58004797 | 58004899 | - | 19.0752509605939 | 17.0549361309657 | 50.1155848354556 | 44.8185631730397 |
| chr17 | 58004920 | 58005169 | - | 431.703048055547 | 326.317777972478 | 457.00640457094 | 299.857529800575 |
| chr17 | 58005191 | 58005243 | - | 34.1346596136944 | 28.4248935516096 | 32.2171616799357 | 19.2079556455884 |
| chr17 | 58005285 | 58005531 | - | 119.47130864793 | 93.2336508492793 | 141.994157033791 | 88.5700176991023 |
| chr17 | 58005557 | 58005602 | - | 31.1227778830743 | 26.1509020674808 | 34.6036181006717 | 12.8053037637256 |
| chr17 | 58006663 | 58006704 | + | 13.0514874993537 | 11.3699574206438 | 2.38645642073598 | 12.8053037637256 |
| chr17 | 58007413 | 58007421 | + | 3.01188173062009 | 4.54798296825753 | 1.19322821036799 | 1.06710864697714 |
| chr17 | 58007661 | 58007662 | + | 18.0712903837206 | 14.780944646837 | 10.7390538933119 | 12.8053037637256 |
| chr17 | 58089326 | 58089340 | + | 3.01188173062009 | 1.13699574206438 | 2.38645642073598 | 3.20132594093141 |
| chr17 | 58089469 | 58089500 | + | 4.01584230749346 | 5.68497871032191 | 7.15936926220794 | 20.2750642925656 |
| chr17 | 58089609 | 58089628 | + | 1.00396057687336 | 1.13699574206438 | 8.35259747257593 | 4.26843458790854 |
| chr17 | 58089690 | 58089737 | + | 12.0475269224804 | 7.95897019445067 | 10.7390538933119 | 18.1408469986113 |
| chr17 | 58089760 | 58089798 | + | 10.0396057687336 | 2.27399148412876 | 5.96614105183995 | 8.53686917581709 |
| chr17 | 58089840 | 58089868 | + | 64.2534769198953 | 36.3838637460602 | 89.4921157775992 | 76.8318225823538 |
| chr17 | 58089891 | 58089912 | + | 14.0554480762271 | 19.3289276150945 | 48.9223566250876 | 37.3488026441998 |
| chr17 | 58090001 | 58090025 | + | 30.1188173062009 | 40.9318467143178 | 66.8207797806074 | 66.1607361125824 |
| chr17 | 58090205 | 58090347 | + | 27.1069355755808 | 39.7948509722534 | 79.9462900946553 | 107.777973344691 |
| chr17 | 58090465 | 58090723 | + | 76.3010038423757 | 68.2197445238629 | 174.211318713726 | 240.099445569856 |
| chr17 | 58090738 | 58090766 | + | 180.712903837206 | 117.110561432631 | 72.7869208324474 | 167.53605757541 |
| chr17 | 58090962 | 58090986 | + | 2.00792115374673 | 2.27399148412876 | 1.19322821036799 | 6.40265188186281 |
| chr17 | 58205507 | 58205549 | - | 5.01980288436682 | 11.3699574206438 | 28.6374770488317 | 8.53686917581709 |
| chr17 | 58205578 | 58205665 | - | 19.0752509605939 | 14.780944646837 | 8.35259747257593 | 21.3421729395427 |
| chr17 | 58206003 | 58206037 | - | 2.00792115374673 | 3.41098722619315 | 5.96614105183995 | 3.20132594093141 |
| chr17 | 58269865 | 58269866 | - | 3.01188173062009 | 4.54798296825753 | 4.77291284147196 | 10.6710864697714 |
| chr17 | 58345220 | 58345231 | - | 3.01188173062009 | 2.27399148412876 | 3.57968463110397 | 2.13421729395427 |
| chr17 | 58345395 | 58345423 | - | 1.00396057687336 | 3.41098722619315 | 1.19322821036799 | 2.13421729395427 |
| chr17 | 58345932 | 58346024 | - | 11.043566345607 | 13.6439489047726 | 14.3187385244159 | 9.60397782279422 |
| chr17 | 58346100 | 58346129 | - | 2.00792115374673 | 4.54798296825753 | 5.96614105183995 | 4.26843458790854 |
| chr17 | 58489561 | 58489659 | - | 16.0633692299738 | 18.1919318730301 | 10.7390538933119 | 16.006629704657 |
| chr17 | 58489680 | 58489712 | - | 3.01188173062009 | 1.13699574206438 | 2.38645642073598 | 6.40265188186281 |
| chr17 | 58489776 | 58489813 | - | 9.03564519186028 | 4.54798296825753 | 3.57968463110397 | 3.20132594093141 |
| chr17 | 58490453 | 58490559 | - | 7.02772403811355 | 3.41098722619315 | 3.57968463110397 | 4.26843458790854 |
| chr17 | 58490614 | 58490688 | - | 37.1465413443145 | 19.3289276150945 | 44.1494437836156 | 33.0803680562912 |
| chr17 | 58490762 | 58490796 | - | 3.01188173062009 | 1.13699574206438 | 5.96614105183995 | 6.40265188186281 |
| chr17 | 58491011 | 58491051 | - | 7.02772403811355 | 3.41098722619315 | 17.8984231555198 | 11.7381951167485 |
| chr17 | 58695215 | 58695242 | + | 2.00792115374673 | 4.54798296825753 | 2.38645642073598 | 2.13421729395427 |
| chr17 | 58695263 | 58695311 | + | 5.01980288436682 | 7.95897019445067 | 2.38645642073598 | 2.13421729395427 |
| chr17 | 58734187 | 58734228 | + | 1.00396057687336 | 2.27399148412876 | 2.38645642073598 | 8.53686917581709 |
| chr17 | 58734283 | 58734295 | + | 9.03564519186028 | 2.27399148412876 | 3.57968463110397 | 6.40265188186281 |
| chr17 | 58998381 | 58998460 | - | 34.1346596136944 | 52.3018041349616 | 28.6374770488317 | 14.9395210576799 |
| chr17 | 58998498 | 58998535 | - | 110.43566345607 | 170.549361309657 | 77.5598336739193 | 58.6909755837425 |
| chr17 | 58998565 | 58998568 | - | 1.00396057687336 | 3.41098722619315 | 2.38645642073598 | 2.13421729395427 |
| chr17 | 58998626 | 58998638 | - | 4.01584230749346 | 12.5069531627082 | 7.15936926220794 | 2.13421729395427 |
| chr17 | 58998683 | 58998734 | - | 26.1029749987075 | 45.4798296825753 | 7.15936926220794 | 11.7381951167485 |
| chr17 | 58998758 | 58998796 | - | 7.02772403811355 | 19.3289276150945 | 8.35259747257593 | 11.7381951167485 |
| chr17 | 59107095 | 59107300 | + | 62.2455557661486 | 88.6856678810218 | 17.8984231555198 | 6.40265188186281 |
| chr17 | 59110146 | 59110163 | - | 1.00396057687336 | 2.27399148412876 | 1.19322821036799 | 1.06710864697714 |
| chr17 | 59110301 | 59110323 | - | 3.01188173062009 | 4.54798296825753 | 1.19322821036799 | 2.13421729395427 |
| chr17 | 59110351 | 59110367 | - | 2.00792115374673 | 4.54798296825753 | 3.57968463110397 | 3.20132594093141 |
| chr17 | 59111214 | 59111230 | - | 5.01980288436682 | 3.41098722619315 | 3.57968463110397 | 3.20132594093141 |
| chr17 | 59111884 | 59111893 | - | 10.0396057687336 | 3.41098722619315 | 5.96614105183995 | 2.13421729395427 |
| chr17 | 59112156 | 59112188 | - | 90.3564519186028 | 89.8226636230862 | 29.8307052591997 | 40.5501285851312 |
| chr17 | 59112229 | 59112245 | - | 3.01188173062009 | 4.54798296825753 | 5.96614105183995 | 2.13421729395427 |
| chr17 | 59202134 | 59202159 | + | 5.01980288436682 | 3.41098722619315 | 4.77291284147196 | 5.33554323488568 |
| chr17 | 59202369 | 59202386 | + | 2.00792115374673 | 2.27399148412876 | 2.38645642073598 | 3.20132594093141 |
| chr17 | 59202469 | 59202491 | + | 13.0514874993537 | 2.27399148412876 | 15.5119667347839 | 4.26843458790854 |
| chr17 | 59607499 | 59607518 | + | 2.00792115374673 | 5.68497871032191 | 1.19322821036799 | 3.20132594093141 |
| chr17 | 59608217 | 59608235 | + | 3.01188173062009 | 13.6439489047726 | 3.57968463110397 | 4.26843458790854 |
| chr17 | 59693726 | 59693891 | + | 5.01980288436682 | 2.27399148412876 | 15.5119667347839 | 22.4092815865198 |
| chr17 | 59693921 | 59694094 | + | 35.1386201905678 | 44.3428339405109 | 39.3765309421437 | 69.3620620535138 |
| chr17 | 59694427 | 59694697 | + | 60.2376346124019 | 59.1237785873479 | 62.0478669391355 | 86.435800405148 |
| chr17 | 59694755 | 59694774 | + | 4.01584230749346 | 2.27399148412876 | 3.57968463110397 | 1.06710864697714 |
| chr17 | 59694847 | 59694900 | + | 2.00792115374673 | 2.27399148412876 | 2.38645642073598 | 6.40265188186281 |
| chr17 | 59695023 | 59695033 | + | 2.00792115374673 | 1.13699574206438 | 1.19322821036799 | 1.06710864697714 |
| chr17 | 59697327 | 59697395 | - | 13.0514874993537 | 20.4659233571589 | 21.4781077866238 | 22.4092815865198 |
| chr17 | 59697424 | 59697440 | - | 2.00792115374673 | 7.95897019445067 | 7.15936926220794 | 2.13421729395427 |
| chr17 | 59839882 | 59839912 | + | 4.01584230749346 | 1.13699574206438 | 1.19322821036799 | 2.13421729395427 |
| chr17 | 59839944 | 59839988 | + | 4.01584230749346 | 7.95897019445067 | 1.19322821036799 | 5.33554323488568 |
| chr17 | 59840133 | 59840175 | + | 5.01980288436682 | 7.95897019445067 | 2.38645642073598 | 7.46976052883995 |
| chr17 | 59840417 | 59840596 | + | 115.455466340437 | 147.80944646837 | 54.8884976769275 | 70.429170700491 |
| chr17 | 59840643 | 59840654 | + | 2.00792115374673 | 3.41098722619315 | 1.19322821036799 | 1.06710864697714 |
| chr17 | 59840842 | 59840863 | + | 1.00396057687336 | 2.27399148412876 | 2.38645642073598 | 3.20132594093141 |
| chr17 | 59859616 | 59859619 | - | 2.00792115374673 | 5.68497871032191 | 3.57968463110397 | 9.60397782279422 |
| chr17 | 59860308 | 59860310 | - | 1.00396057687336 | 4.54798296825753 | 2.38645642073598 | 5.33554323488568 |
| chr17 | 59946809 | 59946856 | + | 4.01584230749346 | 5.68497871032191 | 9.54582568294392 | 6.40265188186281 |
| chr17 | 59947483 | 59947486 | + | 1.00396057687336 | 3.41098722619315 | 2.38645642073598 | 1.06710864697714 |
| chr17 | 59950309 | 59950359 | + | 5.01980288436682 | 13.6439489047726 | 7.15936926220794 | 9.60397782279422 |
| chr17 | 60179216 | 60179224 | - | 4.01584230749346 | 6.82197445238629 | 9.54582568294392 | 9.60397782279422 |
| chr17 | 60445502 | 60445572 | - | 4.01584230749346 | 4.54798296825753 | 4.77291284147196 | 1.06710864697714 |
| chr17 | 60526241 | 60526329 | + | 67.2653586505154 | 98.9186295596012 | 102.617626091647 | 106.710864697714 |
| chr17 | 60663596 | 60663640 | + | 3.01188173062009 | 3.41098722619315 | 2.38645642073598 | 7.46976052883995 |
| chr17 | 61681406 | 61681409 | - | 5.01980288436682 | 17.0549361309657 | 2.38645642073598 | 5.33554323488568 |
| chr17 | 61685353 | 61685361 | - | 1.00396057687336 | 1.13699574206438 | 5.96614105183995 | 2.13421729395427 |
| chr17 | 61865608 | 61865625 | - | 3.01188173062009 | 3.41098722619315 | 8.35259747257593 | 5.33554323488568 |
| chr17 | 61865650 | 61865755 | - | 27.1069355755808 | 37.5208594881246 | 105.004082512383 | 32.0132594093141 |
| chr17 | 61865821 | 61865843 | - | 3.01188173062009 | 3.41098722619315 | 2.38645642073598 | 2.13421729395427 |
| chr17 | 61865880 | 61865961 | - | 10.0396057687336 | 9.09596593651506 | 23.8645642073598 | 19.2079556455884 |
| chr17 | 61866042 | 61866056 | - | 2.00792115374673 | 1.13699574206438 | 5.96614105183995 | 5.33554323488568 |
| chr17 | 61866228 | 61866241 | - | 87.3445701879827 | 37.5208594881246 | 1.19322821036799 | 2.13421729395427 |
| chr17 | 61866386 | 61866395 | - | 2.00792115374673 | 1.13699574206438 | 10.7390538933119 | 24.5434988804741 |
| chr17 | 61866725 | 61866730 | - | 8.03168461498692 | 5.68497871032191 | 14.3187385244159 | 19.2079556455884 |
| chr17 | 61866790 | 61866814 | - | 1.00396057687336 | 1.13699574206438 | 5.96614105183995 | 5.33554323488568 |
| chr17 | 61867110 | 61867147 | - | 3.01188173062009 | 5.68497871032191 | 5.96614105183995 | 7.46976052883995 |
| chr17 | 61867199 | 61867210 | - | 3.01188173062009 | 2.27399148412876 | 3.57968463110397 | 1.06710864697714 |
| chr17 | 61867324 | 61867429 | - | 1.00396057687336 | 3.41098722619315 | 25.0577924177278 | 24.5434988804741 |
| chr17 | 61942682 | 61942694 | - | 3.01188173062009 | 7.95897019445067 | 2.38645642073598 | 2.13421729395427 |
| chr17 | 61943070 | 61943128 | - | 2.00792115374673 | 3.41098722619315 | 1.19322821036799 | 1.06710864697714 |
| chr17 | 61943815 | 61943820 | - | 23.0910932680874 | 21.6029190992233 | 2.38645642073598 | 11.7381951167485 |
| chr17 | 61944787 | 61944818 | - | 6.02376346124019 | 2.27399148412876 | 3.57968463110397 | 9.60397782279422 |
| chr17 | 61945520 | 61945522 | - | 2.00792115374673 | 1.13699574206438 | 2.38645642073598 | 1.06710864697714 |
| chr17 | 61946314 | 61946324 | - | 20.0792115374673 | 22.7399148412876 | 2.38645642073598 | 6.40265188186281 |
| chr17 | 62448839 | 62448854 | + | 26.1029749987075 | 65.9457530397342 | 15.5119667347839 | 39.483019938154 |
| chr17 | 62448872 | 62448913 | + | 47.1861471130481 | 63.6717615556054 | 38.1833027317757 | 53.3554323488568 |
| chr17 | 62613359 | 62613372 | + | 2.00792115374673 | 4.54798296825753 | 1.19322821036799 | 1.06710864697714 |
| chr17 | 62692722 | 62692742 | + | 6.02376346124019 | 4.54798296825753 | 4.77291284147196 | 5.33554323488568 |
| chr17 | 62692990 | 62693000 | + | 7.02772403811355 | 2.27399148412876 | 13.1255103140479 | 8.53686917581709 |
| chr17 | 63432328 | 63432337 | - | 4.01584230749346 | 5.68497871032191 | 2.38645642073598 | 3.20132594093141 |
| chr17 | 63433543 | 63433567 | - | 11.043566345607 | 5.68497871032191 | 7.15936926220794 | 11.7381951167485 |
| chr17 | 63433898 | 63433925 | - | 4.01584230749346 | 7.95897019445067 | 2.38645642073598 | 2.13421729395427 |
| chr17 | 63434026 | 63434034 | - | 2.00792115374673 | 1.13699574206438 | 1.19322821036799 | 4.26843458790854 |
| chr17 | 63434290 | 63434300 | - | 3.01188173062009 | 3.41098722619315 | 10.7390538933119 | 2.13421729395427 |
| chr17 | 63434513 | 63434517 | - | 15.0594086531005 | 12.5069531627082 | 1.19322821036799 | 3.20132594093141 |
| chr17 | 63589361 | 63589393 | + | 7.02772403811355 | 5.68497871032191 | 5.96614105183995 | 5.33554323488568 |
| chr17 | 63589426 | 63589519 | + | 6.02376346124019 | 6.82197445238629 | 7.15936926220794 | 5.33554323488568 |
| chr17 | 63589572 | 63589625 | + | 1.00396057687336 | 1.13699574206438 | 1.19322821036799 | 2.13421729395427 |
| chr17 | 63590009 | 63590046 | + | 22.087132691214 | 25.0139063254164 | 52.5020412561915 | 29.8790421153598 |
| chr17 | 63590175 | 63590226 | + | 4.01584230749346 | 7.95897019445067 | 5.96614105183995 | 5.33554323488568 |
| chr17 | 63590301 | 63590340 | + | 3.01188173062009 | 1.13699574206438 | 5.96614105183995 | 7.46976052883995 |
| chr17 | 63590387 | 63590406 | + | 5.01980288436682 | 1.13699574206438 | 3.57968463110397 | 5.33554323488568 |
| chr17 | 63590556 | 63590610 | + | 1.00396057687336 | 7.95897019445067 | 9.54582568294392 | 14.9395210576799 |
| chr17 | 63590656 | 63590667 | + | 6.02376346124019 | 3.41098722619315 | 2.38645642073598 | 5.33554323488568 |
| chr17 | 63590733 | 63590767 | + | 4.01584230749346 | 1.13699574206438 | 11.9322821036799 | 16.006629704657 |
| chr17 | 63590908 | 63590947 | + | 1.00396057687336 | 4.54798296825753 | 7.15936926220794 | 2.13421729395427 |
| chr17 | 63591072 | 63591079 | + | 1.00396057687336 | 1.13699574206438 | 1.19322821036799 | 2.13421729395427 |
| chr17 | 63591107 | 63591148 | + | 4.01584230749346 | 3.41098722619315 | 8.35259747257593 | 9.60397782279422 |
| chr17 | 63591178 | 63591226 | + | 4.01584230749346 | 3.41098722619315 | 9.54582568294392 | 12.8053037637256 |
| chr17 | 63591934 | 63591949 | + | 11.043566345607 | 1.13699574206438 | 4.77291284147196 | 8.53686917581709 |
| chr17 | 63591984 | 63592117 | + | 37.1465413443145 | 19.3289276150945 | 56.0817258872955 | 73.6304966414224 |
| chr17 | 63592163 | 63592174 | + | 24.0950538449607 | 18.1919318730301 | 32.2171616799357 | 57.6238669367653 |
| chr17 | 63592263 | 63592265 | + | 6.02376346124019 | 2.27399148412876 | 4.77291284147196 | 5.33554323488568 |
| chr17 | 63592430 | 63592484 | + | 7.02772403811355 | 5.68497871032191 | 16.7051949451519 | 17.0737383516342 |
| chr17 | 63592498 | 63592634 | + | 15.0594086531005 | 20.4659233571589 | 50.1155848354556 | 66.1607361125824 |
| chr17 | 63592663 | 63592768 | + | 13.0514874993537 | 7.95897019445067 | 33.4103898903037 | 28.8119334683827 |
| chr17 | 63592797 | 63592893 | + | 13.0514874993537 | 18.1919318730301 | 54.8884976769275 | 41.6172372321083 |
| chr17 | 63593002 | 63593024 | + | 2.00792115374673 | 1.13699574206438 | 15.5119667347839 | 9.60397782279422 |
| chr17 | 63593087 | 63593177 | + | 15.0594086531005 | 9.09596593651506 | 25.0577924177278 | 27.7448248214055 |
| chr17 | 63593211 | 63593234 | + | 5.01980288436682 | 2.27399148412876 | 5.96614105183995 | 9.60397782279422 |
| chr17 | 63593259 | 63593284 | + | 3.01188173062009 | 3.41098722619315 | 3.57968463110397 | 3.20132594093141 |
| chr17 | 63593950 | 63593988 | + | 15.0594086531005 | 12.5069531627082 | 11.9322821036799 | 19.2079556455884 |
| chr17 | 63594064 | 63594099 | + | 10.0396057687336 | 11.3699574206438 | 13.1255103140479 | 21.3421729395427 |
| chr17 | 63594110 | 63594134 | + | 6.02376346124019 | 9.09596593651506 | 22.6713359969918 | 18.1408469986113 |
| chr17 | 63594203 | 63594258 | + | 5.01980288436682 | 9.09596593651506 | 3.57968463110397 | 1.06710864697714 |
| chr17 | 63608005 | 63608044 | + | 2.00792115374673 | 4.54798296825753 | 3.57968463110397 | 4.26843458790854 |
| chr17 | 63608084 | 63608138 | + | 4.01584230749346 | 10.2329616785794 | 4.77291284147196 | 8.53686917581709 |
| chr17 | 63608238 | 63608332 | + | 18.0712903837206 | 22.7399148412876 | 9.54582568294392 | 14.9395210576799 |
| chr17 | 63694916 | 63694940 | + | 32.1267384599477 | 36.3838637460602 | 14.3187385244159 | 14.9395210576799 |
| chr17 | 63694969 | 63694997 | + | 3.01188173062009 | 7.95897019445067 | 10.7390538933119 | 1.06710864697714 |
| chr17 | 63695189 | 63695190 | + | 2.00792115374673 | 1.13699574206438 | 4.77291284147196 | 1.06710864697714 |
| chr17 | 63702867 | 63702921 | - | 16.0633692299738 | 15.9179403889013 | 14.3187385244159 | 9.60397782279422 |
| chr17 | 63704004 | 63704005 | - | 1.00396057687336 | 1.13699574206438 | 1.19322821036799 | 2.13421729395427 |
| chr17 | 63745278 | 63745332 | - | 2.00792115374673 | 6.82197445238629 | 4.77291284147196 | 6.40265188186281 |
| chr17 | 63745431 | 63745434 | - | 9.03564519186028 | 4.54798296825753 | 4.77291284147196 | 13.8724124107028 |
| chr17 | 63745635 | 63745763 | - | 618.439715353992 | 512.785079671036 | 590.647964132155 | 994.54525898269 |
| chr17 | 63745804 | 63745957 | - | 31.1227778830743 | 25.0139063254164 | 36.9900745214077 | 55.4896496428111 |
| chr17 | 63745987 | 63746053 | - | 11.043566345607 | 7.95897019445067 | 16.7051949451519 | 12.8053037637256 |
| chr17 | 63746079 | 63746100 | - | 7.02772403811355 | 3.41098722619315 | 4.77291284147196 | 5.33554323488568 |
| chr17 | 63746134 | 63746181 | - | 6.02376346124019 | 2.27399148412876 | 26.2510206280958 | 10.6710864697714 |
| chr17 | 63746217 | 63746274 | - | 10.0396057687336 | 7.95897019445067 | 17.8984231555198 | 10.6710864697714 |
| chr17 | 63746332 | 63746338 | - | 1.00396057687336 | 1.13699574206438 | 8.35259747257593 | 18.1408469986113 |
| chr17 | 63746467 | 63746488 | - | 5.01980288436682 | 4.54798296825753 | 5.96614105183995 | 8.53686917581709 |
| chr17 | 63746511 | 63746540 | - | 2.00792115374673 | 4.54798296825753 | 5.96614105183995 | 4.26843458790854 |
| chr17 | 63746610 | 63746636 | - | 2.00792115374673 | 2.27399148412876 | 9.54582568294392 | 3.20132594093141 |
| chr17 | 63746684 | 63746739 | - | 80.3168461498692 | 67.0827487817985 | 101.424397881279 | 86.435800405148 |
| chr17 | 63818135 | 63818238 | + | 15.0594086531005 | 21.6029190992233 | 41.7629873628796 | 38.4159112911769 |
| chr17 | 63819016 | 63819068 | + | 10.0396057687336 | 29.5618892936739 | 9.54582568294392 | 20.2750642925656 |
| chr17 | 63819214 | 63819297 | + | 7.02772403811355 | 15.9179403889013 | 19.0916513658878 | 16.006629704657 |
| chr17 | 63819780 | 63819800 | - | 5.01980288436682 | 1.13699574206438 | 2.38645642073598 | 7.46976052883995 |
| chr17 | 63831915 | 63831987 | + | 11.043566345607 | 17.0549361309657 | 10.7390538933119 | 20.2750642925656 |
| chr17 | 63832110 | 63832158 | - | 10.0396057687336 | 5.68497871032191 | 8.35259747257593 | 11.7381951167485 |
| chr17 | 63832182 | 63832243 | - | 4.01584230749346 | 2.27399148412876 | 3.57968463110397 | 10.6710864697714 |
| chr17 | 63832269 | 63832276 | - | 2.00792115374673 | 1.13699574206438 | 1.19322821036799 | 4.26843458790854 |
| chr17 | 63832522 | 63832632 | - | 14.0554480762271 | 6.82197445238629 | 46.5359002043516 | 41.6172372321083 |
| chr17 | 63832737 | 63832812 | - | 4.01584230749346 | 4.54798296825753 | 10.7390538933119 | 16.006629704657 |
| chr17 | 63832854 | 63832866 | - | 1.00396057687336 | 1.13699574206438 | 2.38645642073598 | 9.60397782279422 |
| chr17 | 63832929 | 63832974 | - | 1.00396057687336 | 1.13699574206438 | 13.1255103140479 | 13.8724124107028 |
| chr17 | 64039250 | 64039261 | - | 3.01188173062009 | 51.1648083928972 | 16.7051949451519 | 8.53686917581709 |
| chr17 | 64039340 | 64039350 | - | 2.00792115374673 | 7.95897019445067 | 11.9322821036799 | 4.26843458790854 |
| chr17 | 64039377 | 64039416 | - | 3.01188173062009 | 34.1098722619315 | 16.7051949451519 | 1.06710864697714 |
| chr17 | 64039459 | 64039515 | - | 31.1227778830743 | 156.905412404885 | 20.2848795762558 | 23.476390233497 |
| chr17 | 64039748 | 64039752 | - | 3.01188173062009 | 5.68497871032191 | 1.19322821036799 | 1.06710864697714 |
| chr17 | 64041690 | 64041691 | - | 1.00396057687336 | 3.41098722619315 | 1.19322821036799 | 1.06710864697714 |
| chr17 | 64145963 | 64146015 | + | 713.815970156962 | 343.372714103443 | 215.974306076606 | 431.111893378763 |
| chr17 | 64146080 | 64146088 | + | 1.00396057687336 | 2.27399148412876 | 1.19322821036799 | 1.06710864697714 |
| chr17 | 64146182 | 64146287 | + | 52.205949997415 | 54.5757956190903 | 16.7051949451519 | 36.2816939972226 |
| chr17 | 64147897 | 64147916 | - | 8.03168461498692 | 5.68497871032191 | 3.57968463110397 | 3.20132594093141 |
| chr17 | 64148965 | 64148967 | - | 1.00396057687336 | 1.13699574206438 | 1.19322821036799 | 2.13421729395427 |
| chr17 | 64499612 | 64499674 | - | 42.1663442286813 | 46.6168254246397 | 46.5359002043516 | 61.8923015246739 |
| chr17 | 64499751 | 64499913 | - | 40.1584230749346 | 46.6168254246397 | 77.5598336739193 | 58.6909755837425 |
| chr17 | 64537800 | 64537846 | + | 1.00396057687336 | 14.780944646837 | 5.96614105183995 | 3.20132594093141 |
| chr17 | 64545689 | 64545697 | - | 8.03168461498692 | 12.5069531627082 | 4.77291284147196 | 12.8053037637256 |
| chr17 | 65011216 | 65011306 | - | 37.1465413443145 | 23.876910583352 | 17.8984231555198 | 28.8119334683827 |
| chr17 | 65012233 | 65012261 | - | 1.00396057687336 | 1.13699574206438 | 1.19322821036799 | 1.06710864697714 |
| chr17 | 65012715 | 65012765 | - | 1.00396057687336 | 2.27399148412876 | 1.19322821036799 | 2.13421729395427 |
| chr17 | 65013100 | 65013137 | - | 1.00396057687336 | 1.13699574206438 | 5.96614105183995 | 1.06710864697714 |
| chr17 | 65013216 | 65013257 | - | 2.00792115374673 | 4.54798296825753 | 5.96614105183995 | 2.13421729395427 |
| chr17 | 65014242 | 65014262 | - | 1.00396057687336 | 1.13699574206438 | 2.38645642073598 | 4.26843458790854 |
| chr17 | 66808143 | 66808153 | + | 2.00792115374673 | 2.27399148412876 | 2.38645642073598 | 1.06710864697714 |
| chr17 | 66808990 | 66809014 | + | 8.03168461498692 | 1.13699574206438 | 2.38645642073598 | 2.13421729395427 |
| chr17 | 66810504 | 66810519 | + | 9.03564519186028 | 2.27399148412876 | 3.57968463110397 | 10.6710864697714 |
| chr17 | 66810603 | 66810615 | + | 6.02376346124019 | 1.13699574206438 | 8.35259747257593 | 3.20132594093141 |
| chr17 | 67070513 | 67070573 | - | 85.336649034236 | 73.9047232341848 | 73.9801490428153 | 89.6371263460794 |
| chr17 | 67070600 | 67070622 | - | 3.01188173062009 | 1.13699574206438 | 1.19322821036799 | 1.06710864697714 |
| chr17 | 67070647 | 67070701 | - | 3.01188173062009 | 5.68497871032191 | 4.77291284147196 | 10.6710864697714 |
| chr17 | 67070728 | 67070763 | - | 34.1346596136944 | 26.1509020674808 | 16.7051949451519 | 30.9461507623369 |
| chr17 | 67070797 | 67070817 | - | 23.0910932680874 | 17.0549361309657 | 8.35259747257593 | 19.2079556455884 |
| chr17 | 67070844 | 67070882 | - | 16.0633692299738 | 25.0139063254164 | 5.96614105183995 | 23.476390233497 |
| chr17 | 67071645 | 67071653 | - | 3.01188173062009 | 6.82197445238629 | 1.19322821036799 | 3.20132594093141 |
| chr17 | 67071815 | 67071821 | - | 2.00792115374673 | 1.13699574206438 | 2.38645642073598 | 4.26843458790854 |
| chr17 | 67071994 | 67072016 | - | 8.03168461498692 | 9.09596593651506 | 22.6713359969918 | 10.6710864697714 |
| chr17 | 67072039 | 67072068 | - | 7.02772403811355 | 12.5069531627082 | 11.9322821036799 | 9.60397782279422 |
| chr17 | 67073220 | 67073239 | - | 9.03564519186028 | 4.54798296825753 | 20.2848795762558 | 12.8053037637256 |
| chr17 | 67073329 | 67073344 | - | 10.0396057687336 | 12.5069531627082 | 20.2848795762558 | 6.40265188186281 |
| chr17 | 67073434 | 67073492 | - | 2.00792115374673 | 4.54798296825753 | 15.5119667347839 | 14.9395210576799 |
| chr17 | 67073612 | 67073617 | - | 3.01188173062009 | 6.82197445238629 | 8.35259747257593 | 3.20132594093141 |
| chr17 | 67074273 | 67074282 | - | 3.01188173062009 | 1.13699574206438 | 2.38645642073598 | 1.06710864697714 |
| chr17 | 67074495 | 67074499 | - | 1.00396057687336 | 2.27399148412876 | 2.38645642073598 | 1.06710864697714 |
| chr17 | 67074884 | 67074935 | - | 14.0554480762271 | 15.9179403889013 | 33.4103898903037 | 27.7448248214055 |
| chr17 | 67075015 | 67075080 | - | 4.01584230749346 | 7.95897019445067 | 7.15936926220794 | 1.06710864697714 |
| chr17 | 67075336 | 67075361 | - | 2.00792115374673 | 2.27399148412876 | 3.57968463110397 | 1.06710864697714 |
| chr17 | 67078226 | 67078235 | - | 27.1069355755808 | 22.7399148412876 | 27.4442488384638 | 20.2750642925656 |
| chr17 | 67268220 | 67268222 | + | 24.0950538449607 | 39.7948509722534 | 34.6036181006717 | 10.6710864697714 |
| chr17 | 67337944 | 67337985 | - | 1.00396057687336 | 6.82197445238629 | 1.19322821036799 | 1.06710864697714 |
| chr17 | 67338193 | 67338205 | - | 5.01980288436682 | 2.27399148412876 | 3.57968463110397 | 2.13421729395427 |
| chr17 | 67338253 | 67338283 | - | 6.02376346124019 | 9.09596593651506 | 4.77291284147196 | 11.7381951167485 |
| chr17 | 67340123 | 67340144 | - | 7.02772403811355 | 5.68497871032191 | 4.77291284147196 | 17.0737383516342 |
| chr17 | 67340282 | 67340312 | - | 13.0514874993537 | 7.95897019445067 | 3.57968463110397 | 3.20132594093141 |
| chr17 | 67340460 | 67340463 | - | 3.01188173062009 | 4.54798296825753 | 8.35259747257593 | 9.60397782279422 |
| chr17 | 67340654 | 67340658 | - | 10.0396057687336 | 6.82197445238629 | 1.19322821036799 | 12.8053037637256 |
| chr17 | 67340767 | 67340844 | - | 41.1623836518079 | 30.6988850357383 | 31.0239334695677 | 8.53686917581709 |
| chr17 | 67744010 | 67744020 | + | 18.0712903837206 | 17.0549361309657 | 10.7390538933119 | 24.5434988804741 |
| chr17 | 67744045 | 67744066 | + | 10.0396057687336 | 19.3289276150945 | 10.7390538933119 | 22.4092815865198 |
| chr17 | 67894143 | 67894156 | + | 3.01188173062009 | 3.41098722619315 | 4.77291284147196 | 5.33554323488568 |
| chr17 | 67983018 | 67983047 | + | 1.00396057687336 | 2.27399148412876 | 1.19322821036799 | 1.06710864697714 |
| chr17 | 67991150 | 67991170 | - | 2.00792115374673 | 2.27399148412876 | 4.77291284147196 | 2.13421729395427 |
| chr17 | 67991275 | 67991298 | - | 3.01188173062009 | 1.13699574206438 | 7.15936926220794 | 6.40265188186281 |
| chr17 | 68046595 | 68046623 | + | 8.03168461498692 | 1.13699574206438 | 14.3187385244159 | 6.40265188186281 |
| chr17 | 68129388 | 68129406 | + | 3.01188173062009 | 4.54798296825753 | 1.19322821036799 | 1.06710864697714 |
| chr17 | 68256823 | 68256835 | + | 5.01980288436682 | 10.2329616785794 | 4.77291284147196 | 4.26843458790854 |
| chr17 | 68256881 | 68256952 | + | 12.0475269224804 | 5.68497871032191 | 5.96614105183995 | 6.40265188186281 |
| chr17 | 68257028 | 68257047 | + | 8.03168461498692 | 6.82197445238629 | 1.19322821036799 | 4.26843458790854 |
| chr17 | 68530533 | 68530592 | + | 8.03168461498692 | 3.41098722619315 | 2.38645642073598 | 1.06710864697714 |
| chr17 | 68530659 | 68530681 | + | 5.01980288436682 | 4.54798296825753 | 2.38645642073598 | 2.13421729395427 |
| chr17 | 68530692 | 68530752 | + | 16.0633692299738 | 11.3699574206438 | 7.15936926220794 | 13.8724124107028 |
| chr17 | 68530770 | 68530881 | + | 9.03564519186028 | 14.780944646837 | 4.77291284147196 | 14.9395210576799 |
| chr17 | 68530904 | 68531020 | + | 67.2653586505154 | 64.8087572976698 | 27.4442488384638 | 44.8185631730397 |
| chr17 | 68531040 | 68531089 | + | 4.01584230749346 | 5.68497871032191 | 4.77291284147196 | 2.13421729395427 |
| chr17 | 68531238 | 68531342 | + | 24.0950538449607 | 17.0549361309657 | 7.15936926220794 | 8.53686917581709 |
| chr17 | 68531369 | 68531557 | + | 157.621810569118 | 163.727386857271 | 56.0817258872955 | 78.966039876308 |
| chr17 | 68531580 | 68531701 | + | 52.205949997415 | 85.2746806548286 | 32.2171616799357 | 45.8856718200168 |
| chr17 | 68531730 | 68531803 | + | 3.01188173062009 | 5.68497871032191 | 2.38645642073598 | 6.40265188186281 |
| chr17 | 68531831 | 68531959 | + | 5.01980288436682 | 4.54798296825753 | 1.19322821036799 | 5.33554323488568 |
| chr17 | 68532039 | 68532132 | + | 13.0514874993537 | 18.1919318730301 | 8.35259747257593 | 8.53686917581709 |
| chr17 | 68532152 | 68532182 | + | 9.03564519186028 | 25.0139063254164 | 8.35259747257593 | 4.26843458790854 |
| chr17 | 68532224 | 68532426 | + | 10.0396057687336 | 11.3699574206438 | 2.38645642073598 | 3.20132594093141 |
| chr17 | 68532496 | 68532606 | + | 11.043566345607 | 20.4659233571589 | 8.35259747257593 | 10.6710864697714 |
| chr17 | 68532627 | 68532691 | + | 6.02376346124019 | 5.68497871032191 | 4.77291284147196 | 7.46976052883995 |
| chr17 | 68532717 | 68532748 | + | 17.0673298068472 | 45.4798296825753 | 11.9322821036799 | 17.0737383516342 |
| chr17 | 73207416 | 73207451 | - | 2.00792115374673 | 5.68497871032191 | 2.38645642073598 | 7.46976052883995 |
| chr17 | 73207953 | 73207980 | - | 4.01584230749346 | 4.54798296825753 | 11.9322821036799 | 19.2079556455884 |
| chr17 | 73207994 | 73208040 | - | 1.00396057687336 | 2.27399148412876 | 1.19322821036799 | 1.06710864697714 |
| chr17 | 73208424 | 73208476 | + | 2.00792115374673 | 1.13699574206438 | 3.57968463110397 | 5.33554323488568 |
| chr17 | 73209159 | 73209178 | - | 8.03168461498692 | 7.95897019445067 | 8.35259747257593 | 8.53686917581709 |
| chr17 | 73209272 | 73209304 | - | 3.01188173062009 | 6.82197445238629 | 2.38645642073598 | 2.13421729395427 |
| chr17 | 73247433 | 73247457 | + | 3.01188173062009 | 1.13699574206438 | 1.19322821036799 | 1.06710864697714 |
| chr17 | 73248756 | 73248762 | + | 7.02772403811355 | 5.68497871032191 | 9.54582568294392 | 6.40265188186281 |
| chr17 | 73248791 | 73248812 | + | 1.00396057687336 | 3.41098722619315 | 4.77291284147196 | 2.13421729395427 |
| chr17 | 73283650 | 73283658 | - | 10.0396057687336 | 14.780944646837 | 8.35259747257593 | 8.53686917581709 |
| chr17 | 73285252 | 73285301 | - | 7.02772403811355 | 13.6439489047726 | 3.57968463110397 | 2.13421729395427 |
| chr17 | 74209801 | 74209857 | + | 1587.26167203679 | 1947.67370615629 | 1831.60530291486 | 1970.94967096677 |
| chr17 | 74447262 | 74447402 | + | 12.0475269224804 | 10.2329616785794 | 17.8984231555198 | 6.40265188186281 |
| chr17 | 74748578 | 74748663 | + | 3161.47185657422 | 1388.27180106061 | 637.183864336506 | 1273.06061584372 |
| chr17 | 74748678 | 74748703 | + | 12.0475269224804 | 2.27399148412876 | 5.96614105183995 | 7.46976052883995 |
| chr17 | 74768685 | 74768694 | + | 6.02376346124019 | 3.41098722619315 | 5.96614105183995 | 12.8053037637256 |
| chr17 | 74769247 | 74769334 | + | 17.0673298068472 | 9.09596593651506 | 8.35259747257593 | 7.46976052883995 |
| chr17 | 74770577 | 74770601 | - | 6.02376346124019 | 10.2329616785794 | 5.96614105183995 | 5.33554323488568 |
| chr17 | 74770645 | 74770712 | - | 3.01188173062009 | 12.5069531627082 | 15.5119667347839 | 13.8724124107028 |
| chr17 | 74771390 | 74771422 | - | 3.01188173062009 | 3.41098722619315 | 15.5119667347839 | 18.1408469986113 |
| chr17 | 74838619 | 74838622 | + | 1.00396057687336 | 1.13699574206438 | 1.19322821036799 | 2.13421729395427 |
| chr17 | 74862520 | 74862552 | - | 3.01188173062009 | 2.27399148412876 | 9.54582568294392 | 17.0737383516342 |
| chr17 | 74862585 | 74862719 | - | 132.522796147284 | 61.3977700714766 | 115.743136405695 | 448.185631730397 |
| chr17 | 74862751 | 74862793 | - | 4.01584230749346 | 3.41098722619315 | 20.2848795762558 | 54.4225409958339 |
| chr17 | 75004414 | 75004417 | + | 3.01188173062009 | 5.68497871032191 | 2.38645642073598 | 2.13421729395427 |
| chr17 | 75004546 | 75004547 | + | 2.00792115374673 | 1.13699574206438 | 1.19322821036799 | 1.06710864697714 |
| chr17 | 75005635 | 75005654 | + | 3.01188173062009 | 6.82197445238629 | 1.19322821036799 | 2.13421729395427 |
| chr17 | 75021023 | 75021066 | + | 12.0475269224804 | 11.3699574206438 | 13.1255103140479 | 6.40265188186281 |
| chr17 | 75021161 | 75021241 | + | 13.0514874993537 | 13.6439489047726 | 16.7051949451519 | 12.8053037637256 |
| chr17 | 75038913 | 75038923 | - | 75.2970432655023 | 102.329616785794 | 51.3088130458235 | 91.7713436400337 |
| chr17 | 75064503 | 75064524 | + | 3.01188173062009 | 3.41098722619315 | 2.38645642073598 | 1.06710864697714 |
| chr17 | 75065336 | 75065351 | + | 3.01188173062009 | 3.41098722619315 | 3.57968463110397 | 3.20132594093141 |
| chr17 | 75065776 | 75065833 | + | 7.02772403811355 | 9.09596593651506 | 1.19322821036799 | 6.40265188186281 |
| chr17 | 75130254 | 75130323 | - | 61.2415951892752 | 121.658544400889 | 38.1833027317757 | 55.4896496428111 |
| chr17 | 75130634 | 75130639 | - | 6.02376346124019 | 4.54798296825753 | 5.96614105183995 | 6.40265188186281 |
| chr17 | 75130666 | 75130667 | - | 19.0752509605939 | 7.95897019445067 | 10.7390538933119 | 8.53686917581709 |
| chr17 | 75130752 | 75130861 | - | 26.1029749987075 | 28.4248935516096 | 39.3765309421437 | 40.5501285851312 |
| chr17 | 75130870 | 75130898 | + | 11.043566345607 | 29.5618892936739 | 2.38645642073598 | 2.13421729395427 |
| chr17 | 75131085 | 75131092 | + | 29.1148567293276 | 9.09596593651506 | 1.19322821036799 | 1.06710864697714 |
| chr17 | 75131743 | 75131744 | + | 27.1069355755808 | 4.54798296825753 | 2.38645642073598 | 7.46976052883995 |
| chr17 | 75135270 | 75135305 | - | 3.01188173062009 | 9.09596593651506 | 3.57968463110397 | 1.06710864697714 |
| chr17 | 75135326 | 75135362 | - | 7.02772403811355 | 3.41098722619315 | 4.77291284147196 | 3.20132594093141 |
| chr17 | 75135383 | 75135416 | - | 14.0554480762271 | 10.2329616785794 | 4.77291284147196 | 10.6710864697714 |
| chr17 | 75136012 | 75136055 | - | 32.1267384599477 | 27.2878978095452 | 29.8307052591997 | 59.7580842307196 |
| chr17 | 75136078 | 75136131 | - | 70.2772403811355 | 65.9457530397342 | 46.5359002043516 | 73.6304966414224 |
| chr17 | 75136216 | 75136221 | - | 21.0831721143407 | 7.95897019445067 | 10.7390538933119 | 9.60397782279422 |
| chr17 | 75146665 | 75146689 | - | 21.0831721143407 | 21.6029190992233 | 42.9562155732476 | 30.9461507623369 |
| chr17 | 75168151 | 75168159 | - | 5.01980288436682 | 4.54798296825753 | 7.15936926220794 | 9.60397782279422 |
| chr17 | 75235576 | 75235711 | + | 79.3128855729958 | 72.7677274921204 | 31.0239334695677 | 78.966039876308 |
| chr17 | 75236628 | 75236797 | - | 79.3128855729958 | 85.2746806548286 | 194.496198289982 | 133.388580872142 |
| chr17 | 75236845 | 75236862 | - | 6.02376346124019 | 4.54798296825753 | 25.0577924177278 | 17.0737383516342 |
| chr17 | 75236921 | 75236930 | - | 13.0514874993537 | 14.780944646837 | 8.35259747257593 | 11.7381951167485 |
| chr17 | 75236951 | 75236956 | - | 1.00396057687336 | 1.13699574206438 | 5.96614105183995 | 9.60397782279422 |
| chr17 | 75237037 | 75237057 | - | 5.01980288436682 | 9.09596593651506 | 9.54582568294392 | 3.20132594093141 |
| chr17 | 75237187 | 75237216 | - | 2.00792115374673 | 1.13699574206438 | 3.57968463110397 | 5.33554323488568 |
| chr17 | 75237222 | 75237244 | - | 3.01188173062009 | 3.41098722619315 | 2.38645642073598 | 3.20132594093141 |
| chr17 | 75237482 | 75237490 | - | 5.01980288436682 | 1.13699574206438 | 7.15936926220794 | 5.33554323488568 |
| chr17 | 75237894 | 75237932 | - | 92.3643730723495 | 76.1787147183136 | 35.7968463110397 | 17.0737383516342 |
| chr17 | 75238270 | 75238292 | - | 7.02772403811355 | 5.68497871032191 | 40.5697591525116 | 17.0737383516342 |
| chr17 | 75265981 | 75266040 | + | 11.043566345607 | 14.780944646837 | 5.96614105183995 | 13.8724124107028 |
| chr17 | 75266131 | 75266134 | + | 4.01584230749346 | 6.82197445238629 | 5.96614105183995 | 3.20132594093141 |
| chr17 | 75266191 | 75266226 | + | 4.01584230749346 | 9.09596593651506 | 2.38645642073598 | 2.13421729395427 |
| chr17 | 75266259 | 75266283 | - | 7.02772403811355 | 15.9179403889013 | 1.19322821036799 | 8.53686917581709 |
| chr17 | 75266267 | 75266335 | + | 16.0633692299738 | 34.1098722619315 | 13.1255103140479 | 21.3421729395427 |
| chr17 | 75266383 | 75266395 | - | 1.00396057687336 | 3.41098722619315 | 7.15936926220794 | 3.20132594093141 |
| chr17 | 75271333 | 75271392 | + | 2.00792115374673 | 4.54798296825753 | 3.57968463110397 | 8.53686917581709 |
| chr17 | 75273038 | 75273057 | - | 1.00396057687336 | 4.54798296825753 | 8.35259747257593 | 7.46976052883995 |
| chr17 | 75273128 | 75273132 | - | 22.087132691214 | 18.1919318730301 | 11.9322821036799 | 14.9395210576799 |
| chr17 | 75273266 | 75273295 | - | 1.00396057687336 | 1.13699574206438 | 3.57968463110397 | 14.9395210576799 |
| chr17 | 75318099 | 75318136 | - | 12.0475269224804 | 12.5069531627082 | 8.35259747257593 | 4.26843458790854 |
| chr17 | 75318201 | 75318238 | - | 3.01188173062009 | 3.41098722619315 | 2.38645642073598 | 2.13421729395427 |
| chr17 | 75318264 | 75318302 | - | 6.02376346124019 | 2.27399148412876 | 5.96614105183995 | 6.40265188186281 |
| chr17 | 75318350 | 75318445 | - | 12.0475269224804 | 5.68497871032191 | 5.96614105183995 | 21.3421729395427 |
| chr17 | 75318757 | 75318912 | - | 11.043566345607 | 10.2329616785794 | 34.6036181006717 | 42.6843458790854 |
| chr17 | 75318991 | 75319015 | - | 1.00396057687336 | 5.68497871032191 | 7.15936926220794 | 6.40265188186281 |
| chr17 | 75319047 | 75319050 | - | 3.01188173062009 | 1.13699574206438 | 3.57968463110397 | 2.13421729395427 |
| chr17 | 75319270 | 75319286 | - | 3.01188173062009 | 3.41098722619315 | 3.57968463110397 | 14.9395210576799 |
| chr17 | 75319345 | 75319440 | - | 69.2732798042621 | 57.9867828452835 | 170.631634082622 | 783.257746881218 |
| chr17 | 75319788 | 75319803 | - | 5.01980288436682 | 9.09596593651506 | 5.96614105183995 | 12.8053037637256 |
| chr17 | 75319854 | 75319869 | - | 10.0396057687336 | 13.6439489047726 | 17.8984231555198 | 19.2079556455884 |
| chr17 | 75319891 | 75319909 | - | 7.02772403811355 | 1.13699574206438 | 9.54582568294392 | 3.20132594093141 |
| chr17 | 75319939 | 75319992 | - | 1.00396057687336 | 3.41098722619315 | 9.54582568294392 | 5.33554323488568 |
| chr17 | 75320035 | 75320075 | - | 16.0633692299738 | 4.54798296825753 | 13.1255103140479 | 6.40265188186281 |
| chr17 | 75320114 | 75320135 | - | 4.01584230749346 | 6.82197445238629 | 4.77291284147196 | 3.20132594093141 |
| chr17 | 75320158 | 75320245 | - | 668.637744197661 | 516.196066897229 | 310.239334695677 | 599.71505960115 |
| chr17 | 75320266 | 75320275 | - | 12.0475269224804 | 2.27399148412876 | 10.7390538933119 | 6.40265188186281 |
| chr17 | 75320319 | 75320555 | - | 96.380215379843 | 113.699574206438 | 134.834787771583 | 43.7514545260626 |
| chr17 | 75499973 | 75500066 | + | 7.02772403811355 | 17.0549361309657 | 25.0577924177278 | 20.2750642925656 |
| chr17 | 75524376 | 75524483 | + | 32.1267384599477 | 36.3838637460602 | 39.3765309421437 | 58.6909755837425 |
| chr17 | 75524510 | 75524566 | + | 61.2415951892752 | 32.9728765198671 | 40.5697591525116 | 36.2816939972226 |
| chr17 | 75524621 | 75524714 | + | 55.217831728035 | 29.5618892936739 | 33.4103898903037 | 49.0869977609482 |
| chr17 | 75575028 | 75575090 | + | 7.02772403811355 | 6.82197445238629 | 2.38645642073598 | 3.20132594093141 |
| chr17 | 75575131 | 75575183 | + | 8.03168461498692 | 18.1919318730301 | 3.57968463110397 | 4.26843458790854 |
| chr17 | 75706702 | 75706747 | + | 10.0396057687336 | 7.95897019445067 | 5.96614105183995 | 9.60397782279422 |
| chr17 | 75707529 | 75707558 | + | 8.03168461498692 | 4.54798296825753 | 1.19322821036799 | 3.20132594093141 |
| chr17 | 75708004 | 75708018 | + | 2.00792115374673 | 4.54798296825753 | 1.19322821036799 | 1.06710864697714 |
| chr17 | 75776459 | 75776485 | - | 3.01188173062009 | 11.3699574206438 | 9.54582568294392 | 4.26843458790854 |
| chr17 | 75777640 | 75777656 | - | 84.3326884573626 | 101.19262104373 | 27.4442488384638 | 93.9055609339879 |
| chr17 | 75777706 | 75777713 | - | 2.00792115374673 | 2.27399148412876 | 1.19322821036799 | 6.40265188186281 |
| chr17 | 75777775 | 75777788 | - | 4.01584230749346 | 4.54798296825753 | 3.57968463110397 | 2.13421729395427 |
| chr17 | 75777814 | 75777844 | - | 18.0712903837206 | 19.3289276150945 | 16.7051949451519 | 8.53686917581709 |
| chr17 | 75777866 | 75777947 | - | 8.03168461498692 | 12.5069531627082 | 19.0916513658878 | 22.4092815865198 |
| chr17 | 75778056 | 75778120 | - | 6.02376346124019 | 7.95897019445067 | 22.6713359969918 | 5.33554323488568 |
| chr17 | 75778149 | 75778193 | - | 28.1108961524542 | 65.9457530397342 | 78.7530618842873 | 61.8923015246739 |
| chr17 | 75778288 | 75778326 | - | 33.130699036821 | 40.9318467143178 | 39.3765309421437 | 44.8185631730397 |
| chr17 | 75778368 | 75778467 | - | 421.663442286813 | 691.293411175144 | 1258.85576193823 | 673.345556242573 |
| chr17 | 75778509 | 75778552 | - | 8.03168461498692 | 21.6029190992233 | 38.1833027317757 | 2.13421729395427 |
| chr17 | 75778574 | 75778600 | - | 62.2455557661486 | 73.9047232341848 | 81.1395183050233 | 123.784603049348 |
| chr17 | 75778689 | 75778717 | - | 18.0712903837206 | 13.6439489047726 | 13.1255103140479 | 19.2079556455884 |
| chr17 | 75827265 | 75827314 | - | 10.0396057687336 | 20.4659233571589 | 22.6713359969918 | 46.952780466994 |
| chr17 | 75827345 | 75827593 | - | 73.2891221117556 | 44.3428339405109 | 143.187385244159 | 157.932079752616 |
| chr17 | 75827656 | 75827661 | - | 23.0910932680874 | 14.780944646837 | 75.1733772531833 | 24.5434988804741 |
| chr17 | 75827711 | 75827723 | - | 2.00792115374673 | 1.13699574206438 | 5.96614105183995 | 32.0132594093141 |
| chr17 | 75827783 | 75827858 | - | 6.02376346124019 | 6.82197445238629 | 23.8645642073598 | 38.4159112911769 |
| chr17 | 75827879 | 75827975 | - | 40.1584230749346 | 29.5618892936739 | 75.1733772531833 | 30.9461507623369 |
| chr17 | 75845834 | 75845851 | - | 1.00396057687336 | 1.13699574206438 | 1.19322821036799 | 2.13421729395427 |
| chr17 | 75845876 | 75845938 | - | 8.03168461498692 | 5.68497871032191 | 7.15936926220794 | 13.8724124107028 |
| chr17 | 75846011 | 75846049 | - | 6.02376346124019 | 5.68497871032191 | 11.9322821036799 | 13.8724124107028 |
| chr17 | 75846079 | 75846196 | - | 32.1267384599477 | 11.3699574206438 | 64.4343233598714 | 53.3554323488568 |
| chr17 | 75846259 | 75846294 | - | 2.00792115374673 | 1.13699574206438 | 4.77291284147196 | 9.60397782279422 |
| chr17 | 75846451 | 75846583 | - | 24.0950538449607 | 11.3699574206438 | 51.3088130458235 | 18.1408469986113 |
| chr17 | 75846596 | 75846681 | - | 66.2613980736421 | 36.3838637460602 | 113.356679984959 | 62.959410171651 |
| chr17 | 75846727 | 75846756 | - | 4.01584230749346 | 1.13699574206438 | 4.77291284147196 | 4.26843458790854 |
| chr17 | 75874238 | 75874276 | - | 2.00792115374673 | 3.41098722619315 | 1.19322821036799 | 1.06710864697714 |
| chr17 | 75889049 | 75889056 | - | 3.01188173062009 | 1.13699574206438 | 4.77291284147196 | 3.20132594093141 |
| chr17 | 75898672 | 75898726 | - | 33.130699036821 | 54.5757956190903 | 10.7390538933119 | 26.6777161744284 |
| chr17 | 75904675 | 75904677 | - | 1.00396057687336 | 2.27399148412876 | 2.38645642073598 | 1.06710864697714 |
| chr17 | 75904688 | 75904713 | - | 6.02376346124019 | 3.41098722619315 | 5.96614105183995 | 4.26843458790854 |
| chr17 | 75942611 | 75942660 | - | 7.02772403811355 | 5.68497871032191 | 3.57968463110397 | 1.06710864697714 |
| chr17 | 75943816 | 75943855 | - | 2.00792115374673 | 1.13699574206438 | 1.19322821036799 | 1.06710864697714 |
| chr17 | 75945602 | 75945644 | - | 3.01188173062009 | 7.95897019445067 | 1.19322821036799 | 1.06710864697714 |
| chr17 | 76039180 | 76039208 | - | 9.03564519186028 | 1.13699574206438 | 3.57968463110397 | 2.13421729395427 |
| chr17 | 76039492 | 76039539 | - | 7.02772403811355 | 1.13699574206438 | 8.35259747257593 | 11.7381951167485 |
| chr17 | 76039578 | 76039619 | - | 8.03168461498692 | 1.13699574206438 | 15.5119667347839 | 7.46976052883995 |
| chr17 | 76039674 | 76039690 | - | 21.0831721143407 | 30.6988850357383 | 46.5359002043516 | 29.8790421153598 |
| chr17 | 76081159 | 76081165 | - | 1.00396057687336 | 1.13699574206438 | 1.19322821036799 | 2.13421729395427 |
| chr17 | 76081246 | 76081271 | - | 1.00396057687336 | 2.27399148412876 | 1.19322821036799 | 1.06710864697714 |
| chr17 | 76081465 | 76081526 | - | 25.0990144218341 | 6.82197445238629 | 3.57968463110397 | 8.53686917581709 |
| chr17 | 76081638 | 76081645 | - | 8.03168461498692 | 3.41098722619315 | 9.54582568294392 | 3.20132594093141 |
| chr17 | 76081675 | 76081710 | - | 6.02376346124019 | 11.3699574206438 | 7.15936926220794 | 1.06710864697714 |
| chr17 | 76081736 | 76081818 | - | 30.1188173062009 | 14.780944646837 | 23.8645642073598 | 28.8119334683827 |
| chr17 | 76081851 | 76081879 | - | 3.01188173062009 | 3.41098722619315 | 3.57968463110397 | 12.8053037637256 |
| chr17 | 76083370 | 76083382 | - | 1.00396057687336 | 2.27399148412876 | 1.19322821036799 | 4.26843458790854 |
| chr17 | 76083601 | 76083618 | - | 2.00792115374673 | 1.13699574206438 | 2.38645642073598 | 1.06710864697714 |
| chr17 | 76270528 | 76270625 | + | 50.1980288436682 | 26.1509020674808 | 58.4681823080315 | 39.483019938154 |
| chr17 | 76270652 | 76270708 | + | 34.1346596136944 | 30.6988850357383 | 76.3666054635513 | 99.2411041688736 |
| chr17 | 76270805 | 76270826 | + | 7.02772403811355 | 1.13699574206438 | 20.2848795762558 | 13.8724124107028 |
| chr17 | 76270847 | 76270867 | + | 3.01188173062009 | 1.13699574206438 | 8.35259747257593 | 9.60397782279422 |
| chr17 | 76270940 | 76271015 | + | 7.02772403811355 | 6.82197445238629 | 19.0916513658878 | 13.8724124107028 |
| chr17 | 76271192 | 76271218 | + | 5.01980288436682 | 3.41098722619315 | 2.38645642073598 | 5.33554323488568 |
| chr17 | 76271252 | 76271256 | + | 34.1346596136944 | 62.534765813541 | 29.8307052591997 | 21.3421729395427 |
| chr17 | 76311102 | 76311216 | - | 6.02376346124019 | 10.2329616785794 | 4.77291284147196 | 6.40265188186281 |
| chr17 | 76311359 | 76311555 | - | 118.467348071057 | 134.165497563597 | 63.2410951495034 | 78.966039876308 |
| chr17 | 76387727 | 76387824 | + | 8.03168461498692 | 4.54798296825753 | 2.38645642073598 | 13.8724124107028 |
| chr17 | 76389590 | 76389715 | - | 24.0950538449607 | 15.9179403889013 | 17.8984231555198 | 40.5501285851312 |
| chr17 | 76389995 | 76390007 | - | 8.03168461498692 | 3.41098722619315 | 7.15936926220794 | 8.53686917581709 |
| chr17 | 76390275 | 76390304 | - | 2.00792115374673 | 1.13699574206438 | 10.7390538933119 | 4.26843458790854 |
| chr17 | 76557713 | 76557854 | + | 2097.27364508846 | 1569.05412404885 | 1907.97190837842 | 1896.25206567837 |
| chr17 | 76558792 | 76558800 | + | 8.03168461498692 | 7.95897019445067 | 13.1255103140479 | 17.0737383516342 |
| chr17 | 76558951 | 76558955 | + | 1.00396057687336 | 1.13699574206438 | 2.38645642073598 | 4.26843458790854 |
| chr17 | 76561111 | 76561113 | + | 3.01188173062009 | 1.13699574206438 | 3.57968463110397 | 3.20132594093141 |
| chr17 | 76561387 | 76561399 | + | 20.0792115374673 | 9.09596593651506 | 21.4781077866238 | 10.6710864697714 |
| chr17 | 76563194 | 76563214 | + | 2.00792115374673 | 7.95897019445067 | 4.77291284147196 | 1.06710864697714 |
| chr17 | 76563437 | 76563461 | + | 8.03168461498692 | 13.6439489047726 | 10.7390538933119 | 6.40265188186281 |
| chr17 | 76563493 | 76563517 | + | 1.00396057687336 | 4.54798296825753 | 5.96614105183995 | 10.6710864697714 |
| chr17 | 76563570 | 76563604 | + | 13.0514874993537 | 3.41098722619315 | 15.5119667347839 | 11.7381951167485 |
| chr17 | 76563698 | 76563714 | + | 15.0594086531005 | 12.5069531627082 | 59.6614105183995 | 71.4962793474681 |
| chr17 | 76564027 | 76564104 | + | 25.0990144218341 | 27.2878978095452 | 84.7192029361272 | 120.583277108416 |
| chr17 | 76564442 | 76564456 | + | 1.00396057687336 | 1.13699574206438 | 5.96614105183995 | 6.40265188186281 |
| chr17 | 76564482 | 76564508 | + | 28.1108961524542 | 43.2058381984465 | 54.8884976769275 | 49.0869977609482 |
| chr17 | 76564541 | 76564562 | + | 7.02772403811355 | 7.95897019445067 | 4.77291284147196 | 3.20132594093141 |
| chr17 | 76564591 | 76564595 | + | 4.01584230749346 | 1.13699574206438 | 7.15936926220794 | 2.13421729395427 |
| chr17 | 76564736 | 76564808 | + | 9.03564519186028 | 9.09596593651506 | 39.3765309421437 | 25.6106075274513 |
| chr17 | 76564831 | 76564878 | + | 44.174265382428 | 38.657855230189 | 54.8884976769275 | 48.0198891139711 |
| chr17 | 76564936 | 76564957 | + | 5.01980288436682 | 2.27399148412876 | 15.5119667347839 | 8.53686917581709 |
| chr17 | 76564983 | 76565009 | + | 15.0594086531005 | 13.6439489047726 | 29.8307052591997 | 11.7381951167485 |
| chr17 | 76565120 | 76565144 | + | 2.00792115374673 | 5.68497871032191 | 10.7390538933119 | 8.53686917581709 |
| chr17 | 76671922 | 76671991 | + | 22.087132691214 | 52.3018041349616 | 23.8645642073598 | 28.8119334683827 |
| chr17 | 76715703 | 76715704 | - | 1.00396057687336 | 2.27399148412876 | 1.19322821036799 | 1.06710864697714 |
| chr17 | 76718485 | 76718536 | - | 14.0554480762271 | 17.0549361309657 | 34.6036181006717 | 24.5434988804741 |
| chr17 | 76718596 | 76718690 | - | 83.3287278804893 | 30.6988850357383 | 31.0239334695677 | 82.1673658172394 |
| chr17 | 76733655 | 76733679 | + | 3.01188173062009 | 1.13699574206438 | 1.19322821036799 | 3.20132594093141 |
| chr17 | 76733773 | 76733795 | + | 20.0792115374673 | 40.9318467143178 | 15.5119667347839 | 12.8053037637256 |
| chr17 | 76734177 | 76734245 | - | 13.0514874993537 | 12.5069531627082 | 19.0916513658878 | 5.33554323488568 |
| chr17 | 76734271 | 76734305 | - | 3.01188173062009 | 9.09596593651506 | 9.54582568294392 | 1.06710864697714 |
| chr17 | 76734333 | 76734410 | - | 65.2574374967687 | 46.6168254246397 | 23.8645642073598 | 62.959410171651 |
| chr17 | 76734489 | 76734565 | - | 60.2376346124019 | 45.4798296825753 | 29.8307052591997 | 33.0803680562912 |
| chr17 | 76734640 | 76734643 | - | 9.03564519186028 | 2.27399148412876 | 2.38645642073598 | 5.33554323488568 |
| chr17 | 76734667 | 76734720 | - | 11.043566345607 | 2.27399148412876 | 1.19322821036799 | 4.26843458790854 |
| chr17 | 76734878 | 76734954 | - | 72.2851615348822 | 72.7677274921204 | 41.7629873628796 | 54.4225409958339 |
| chr17 | 76734977 | 76735017 | - | 22.087132691214 | 12.5069531627082 | 5.96614105183995 | 9.60397782279422 |
| chr17 | 76735043 | 76735163 | - | 126.499032686044 | 76.1787147183136 | 38.1833027317757 | 59.7580842307196 |
| chr17 | 76735273 | 76735277 | - | 1.00396057687336 | 1.13699574206438 | 1.19322821036799 | 3.20132594093141 |
| chr17 | 76735420 | 76735460 | - | 1.00396057687336 | 3.41098722619315 | 11.9322821036799 | 11.7381951167485 |
| chr17 | 76735588 | 76735620 | - | 3.01188173062009 | 4.54798296825753 | 7.15936926220794 | 13.8724124107028 |
| chr17 | 76735771 | 76735841 | - | 6.02376346124019 | 1.13699574206438 | 10.7390538933119 | 9.60397782279422 |
| chr17 | 76735963 | 76736051 | - | 13.0514874993537 | 12.5069531627082 | 21.4781077866238 | 11.7381951167485 |
| chr17 | 76736075 | 76736112 | - | 22.087132691214 | 23.876910583352 | 17.8984231555198 | 13.8724124107028 |
| chr17 | 76736143 | 76736288 | - | 47.1861471130481 | 30.6988850357383 | 15.5119667347839 | 37.3488026441998 |
| chr17 | 76778547 | 76778567 | + | 2.00792115374673 | 2.27399148412876 | 1.19322821036799 | 1.06710864697714 |
| chr17 | 76778666 | 76778697 | + | 3.01188173062009 | 3.41098722619315 | 3.57968463110397 | 2.13421729395427 |
| chr17 | 77088639 | 77088880 | + | 464.833747092368 | 628.758645361603 | 687.299449171962 | 584.77553854347 |
| chr17 | 77088914 | 77088933 | + | 7.02772403811355 | 7.95897019445067 | 7.15936926220794 | 5.33554323488568 |
| chr17 | 77093222 | 77093244 | + | 2.00792115374673 | 1.13699574206438 | 5.96614105183995 | 5.33554323488568 |
| chr17 | 77214631 | 77214648 | + | 1.00396057687336 | 2.27399148412876 | 2.38645642073598 | 1.06710864697714 |
| chr17 | 77214762 | 77214800 | + | 5.01980288436682 | 10.2329616785794 | 1.19322821036799 | 5.33554323488568 |
| chr17 | 77215112 | 77215136 | + | 16.0633692299738 | 23.876910583352 | 22.6713359969918 | 7.46976052883995 |
| chr17 | 77215524 | 77215575 | + | 15.0594086531005 | 19.3289276150945 | 8.35259747257593 | 11.7381951167485 |
| chr17 | 77215602 | 77215627 | + | 2.00792115374673 | 5.68497871032191 | 2.38645642073598 | 2.13421729395427 |
| chr17 | 77215934 | 77215955 | + | 12.0475269224804 | 11.3699574206438 | 7.15936926220794 | 11.7381951167485 |
| chr17 | 77216505 | 77216514 | + | 10.0396057687336 | 5.68497871032191 | 8.35259747257593 | 9.60397782279422 |
| chr17 | 77216600 | 77216869 | + | 271.069355755808 | 259.235029190679 | 291.147683329789 | 385.226221558746 |
| chr17 | 77487421 | 77487546 | + | 54.2138711511617 | 57.9867828452835 | 46.5359002043516 | 9.60397782279422 |
| chr17 | 77498595 | 77498763 | + | 25.0990144218341 | 15.9179403889013 | 8.35259747257593 | 6.40265188186281 |
| chr17 | 77498785 | 77498837 | + | 2.00792115374673 | 1.13699574206438 | 3.57968463110397 | 9.60397782279422 |
| chr17 | 77498854 | 77498914 | + | 12.0475269224804 | 7.95897019445067 | 4.77291284147196 | 7.46976052883995 |
| chr17 | 77498941 | 77498991 | + | 12.0475269224804 | 5.68497871032191 | 4.77291284147196 | 9.60397782279422 |
| chr17 | 77499014 | 77499061 | + | 12.0475269224804 | 7.95897019445067 | 14.3187385244159 | 11.7381951167485 |
| chr17 | 77499092 | 77499176 | + | 34.1346596136944 | 10.2329616785794 | 21.4781077866238 | 18.1408469986113 |
| chr17 | 77499194 | 77499249 | + | 4.01584230749346 | 2.27399148412876 | 2.38645642073598 | 20.2750642925656 |
| chr17 | 77499260 | 77499351 | + | 25.0990144218341 | 14.780944646837 | 40.5697591525116 | 26.6777161744284 |
| chr17 | 77499481 | 77499687 | + | 3.01188173062009 | 4.54798296825753 | 4.77291284147196 | 5.33554323488568 |
| chr17 | 77499861 | 77499950 | + | 1.00396057687336 | 1.13699574206438 | 1.19322821036799 | 3.20132594093141 |
| chr17 | 77499972 | 77500008 | + | 2.00792115374673 | 1.13699574206438 | 1.19322821036799 | 1.06710864697714 |
| chr17 | 77500101 | 77500205 | + | 5.01980288436682 | 4.54798296825753 | 3.57968463110397 | 3.20132594093141 |
| chr17 | 77500383 | 77500576 | + | 42.1663442286813 | 44.3428339405109 | 15.5119667347839 | 41.6172372321083 |
| chr17 | 78112979 | 78113037 | - | 3.01188173062009 | 2.27399148412876 | 5.96614105183995 | 12.8053037637256 |
| chr17 | 78113106 | 78113232 | - | 12.0475269224804 | 5.68497871032191 | 38.1833027317757 | 91.7713436400337 |
| chr17 | 78120652 | 78120805 | - | 17.0673298068472 | 6.82197445238629 | 60.8546387287675 | 56.5567582897882 |
| chr17 | 78171777 | 78171783 | + | 1.00396057687336 | 1.13699574206438 | 3.57968463110397 | 4.26843458790854 |
| chr17 | 78171941 | 78171943 | + | 4.01584230749346 | 2.27399148412876 | 7.15936926220794 | 4.26843458790854 |
| chr17 | 78172049 | 78172097 | + | 14.0554480762271 | 14.780944646837 | 84.7192029361272 | 56.5567582897882 |
| chr17 | 78172166 | 78172182 | + | 2.00792115374673 | 4.54798296825753 | 8.35259747257593 | 12.8053037637256 |
| chr17 | 78172277 | 78172350 | + | 10.0396057687336 | 7.95897019445067 | 33.4103898903037 | 33.0803680562912 |
| chr17 | 78172634 | 78172701 | + | 105.415860571703 | 86.411676396893 | 114.549908195327 | 202.750642925656 |
| chr17 | 78174113 | 78174335 | - | 526.075342281643 | 532.114007286131 | 233.872729232126 | 385.226221558746 |
| chr17 | 78174366 | 78174394 | - | 7.02772403811355 | 4.54798296825753 | 10.7390538933119 | 19.2079556455884 |
| chr17 | 78174437 | 78174547 | - | 24.0950538449607 | 13.6439489047726 | 25.0577924177278 | 33.0803680562912 |
| chr17 | 78174571 | 78174578 | - | 2.00792115374673 | 3.41098722619315 | 8.35259747257593 | 35.2145853502455 |
| chr17 | 78174614 | 78174680 | - | 44.174265382428 | 19.3289276150945 | 38.1833027317757 | 37.3488026441998 |
| chr17 | 78174736 | 78174761 | - | 9.03564519186028 | 6.82197445238629 | 11.9322821036799 | 12.8053037637256 |
| chr17 | 78174806 | 78174953 | - | 47.1861471130481 | 45.4798296825753 | 99.0379414605431 | 75.7647139353766 |
| chr17 | 78175052 | 78175168 | - | 48.1901076899215 | 54.5757956190903 | 101.424397881279 | 86.435800405148 |
| chr17 | 78175532 | 78175550 | - | 5.01980288436682 | 4.54798296825753 | 7.15936926220794 | 4.26843458790854 |
| chr17 | 78207561 | 78207619 | + | 1.00396057687336 | 2.27399148412876 | 7.15936926220794 | 7.46976052883995 |
| chr17 | 78223539 | 78223606 | + | 31.1227778830743 | 5.68497871032191 | 22.6713359969918 | 22.4092815865198 |
| chr17 | 78223633 | 78223729 | + | 39.1544624980612 | 28.4248935516096 | 63.2410951495034 | 39.483019938154 |
| chr17 | 78223750 | 78223766 | + | 6.02376346124019 | 5.68497871032191 | 5.96614105183995 | 9.60397782279422 |
| chr17 | 78223794 | 78223801 | + | 13.0514874993537 | 11.3699574206438 | 13.1255103140479 | 8.53686917581709 |
| chr17 | 78223824 | 78224036 | + | 202.80003652842 | 167.138374083464 | 190.916513658878 | 198.482208337747 |
| chr17 | 78224076 | 78224135 | + | 20.0792115374673 | 13.6439489047726 | 36.9900745214077 | 20.2750642925656 |
| chr17 | 78224188 | 78224296 | + | 57.2257528817818 | 32.9728765198671 | 97.8447132501751 | 121.650385755393 |
| chr17 | 78224331 | 78224351 | + | 18.0712903837206 | 5.68497871032191 | 20.2848795762558 | 26.6777161744284 |
| chr17 | 78224373 | 78224395 | + | 11.043566345607 | 6.82197445238629 | 26.2510206280958 | 24.5434988804741 |
| chr17 | 78224495 | 78224502 | + | 25.0990144218341 | 12.5069531627082 | 10.7390538933119 | 23.476390233497 |
| chr17 | 78224537 | 78224581 | + | 23.0910932680874 | 18.1919318730301 | 11.9322821036799 | 27.7448248214055 |
| chr17 | 78224616 | 78224678 | + | 110.43566345607 | 72.7677274921204 | 81.1395183050233 | 68.2949534065367 |
| chr17 | 78357031 | 78357040 | - | 6.02376346124019 | 10.2329616785794 | 10.7390538933119 | 14.9395210576799 |
| chr17 | 78357105 | 78357108 | - | 2.00792115374673 | 14.780944646837 | 2.38645642073598 | 4.26843458790854 |
| chr17 | 78357161 | 78357171 | - | 1.00396057687336 | 4.54798296825753 | 9.54582568294392 | 77.8989312293309 |
| chr17 | 78357395 | 78357425 | - | 15.0594086531005 | 23.876910583352 | 33.4103898903037 | 21.3421729395427 |
| chr17 | 78357584 | 78357589 | - | 1.00396057687336 | 2.27399148412876 | 5.96614105183995 | 2.13421729395427 |
| chr17 | 78358219 | 78358227 | - | 5.01980288436682 | 6.82197445238629 | 7.15936926220794 | 1.06710864697714 |
| chr17 | 78360416 | 78360472 | + | 51.2019894205416 | 89.8226636230862 | 4.77291284147196 | 7.46976052883995 |
| chr17 | 78424444 | 78424478 | + | 2.00792115374673 | 5.68497871032191 | 2.38645642073598 | 5.33554323488568 |
| chr17 | 78674262 | 78674266 | - | 5.01980288436682 | 4.54798296825753 | 5.96614105183995 | 5.33554323488568 |
| chr17 | 78675907 | 78675941 | - | 1.00396057687336 | 2.27399148412876 | 1.19322821036799 | 1.06710864697714 |
| chr17 | 78796275 | 78796318 | - | 7.02772403811355 | 6.82197445238629 | 5.96614105183995 | 5.33554323488568 |
| chr17 | 78796538 | 78796621 | - | 21.0831721143407 | 40.9318467143178 | 26.2510206280958 | 27.7448248214055 |
| chr17 | 78796745 | 78796768 | - | 3.01188173062009 | 10.2329616785794 | 3.57968463110397 | 5.33554323488568 |
| chr17 | 78797014 | 78797019 | - | 1.00396057687336 | 2.27399148412876 | 2.38645642073598 | 2.13421729395427 |
| chr17 | 78797120 | 78797151 | - | 12.0475269224804 | 19.3289276150945 | 8.35259747257593 | 8.53686917581709 |
| chr17 | 78797228 | 78797269 | - | 11.043566345607 | 12.5069531627082 | 4.77291284147196 | 9.60397782279422 |
| chr17 | 78797391 | 78797413 | - | 1.00396057687336 | 1.13699574206438 | 3.57968463110397 | 1.06710864697714 |
| chr17 | 78798317 | 78798336 | - | 8.03168461498692 | 3.41098722619315 | 2.38645642073598 | 1.06710864697714 |
| chr17 | 78855699 | 78855700 | + | 7.02772403811355 | 2.27399148412876 | 1.19322821036799 | 1.06710864697714 |
| chr17 | 78971894 | 78972414 | - | 226.89509037338 | 146.672450726305 | 1.19322821036799 | 6.40265188186281 |
| chr17 | 78991823 | 78991891 | - | 62.2455557661486 | 40.9318467143178 | 50.1155848354556 | 64.0265188186281 |
| chr17 | 78991912 | 78991987 | - | 41.1623836518079 | 28.4248935516096 | 29.8307052591997 | 43.7514545260626 |
| chr17 | 78991992 | 78992033 | - | 6.02376346124019 | 2.27399148412876 | 3.57968463110397 | 3.20132594093141 |
| chr17 | 78992072 | 78992145 | - | 11.043566345607 | 10.2329616785794 | 13.1255103140479 | 11.7381951167485 |
| chr17 | 78992171 | 78992191 | - | 9.03564519186028 | 2.27399148412876 | 4.77291284147196 | 12.8053037637256 |
| chr17 | 78992267 | 78992284 | - | 1.00396057687336 | 2.27399148412876 | 1.19322821036799 | 2.13421729395427 |
| chr17 | 79086466 | 79086490 | + | 6.02376346124019 | 4.54798296825753 | 5.96614105183995 | 4.26843458790854 |
| chr17 | 79086515 | 79086518 | + | 2.00792115374673 | 3.41098722619315 | 2.38645642073598 | 1.06710864697714 |
| chr17 | 79086571 | 79086579 | + | 2.00792115374673 | 4.54798296825753 | 5.96614105183995 | 5.33554323488568 |
| chr17 | 79086625 | 79086650 | + | 11.043566345607 | 13.6439489047726 | 11.9322821036799 | 4.26843458790854 |
| chr17 | 79786421 | 79786428 | + | 1.00396057687336 | 1.13699574206438 | 5.96614105183995 | 2.13421729395427 |
| chr17 | 79786613 | 79786695 | + | 18.0712903837206 | 17.0549361309657 | 11.9322821036799 | 11.7381951167485 |
| chr17 | 79786784 | 79786796 | + | 15.0594086531005 | 12.5069531627082 | 17.8984231555198 | 9.60397782279422 |
| chr17 | 79786818 | 79786855 | + | 7.02772403811355 | 5.68497871032191 | 2.38645642073598 | 4.26843458790854 |
| chr17 | 79786890 | 79786905 | + | 5.01980288436682 | 4.54798296825753 | 2.38645642073598 | 1.06710864697714 |
| chr17 | 79787478 | 79787527 | + | 8.03168461498692 | 14.780944646837 | 3.57968463110397 | 5.33554323488568 |
| chr17 | 79794462 | 79794465 | - | 4.01584230749346 | 5.68497871032191 | 2.38645642073598 | 2.13421729395427 |
| chr17 | 79833305 | 79833418 | - | 182.720824990952 | 219.440178218426 | 141.994157033791 | 74.6976052883995 |
| chr17 | 79833446 | 79833716 | - | 113.44754518669 | 230.81013563907 | 126.482190299007 | 117.381951167485 |
| chr17 | 79833811 | 79833835 | - | 14.0554480762271 | 12.5069531627082 | 5.96614105183995 | 4.26843458790854 |
| chr17 | 79835012 | 79835076 | - | 12.0475269224804 | 7.95897019445067 | 4.77291284147196 | 7.46976052883995 |
| chr17 | 79835192 | 79835233 | - | 9.03564519186028 | 7.95897019445067 | 4.77291284147196 | 7.46976052883995 |
| chr17 | 79835286 | 79835310 | - | 4.01584230749346 | 9.09596593651506 | 1.19322821036799 | 1.06710864697714 |
| chr17 | 80135290 | 80135508 | - | 112.443584609817 | 173.96034853585 | 44.1494437836156 | 51.2212150549025 |
| chr17 | 80147343 | 80147382 | + | 4.01584230749346 | 12.5069531627082 | 2.38645642073598 | 5.33554323488568 |
| chr17 | 80321340 | 80321353 | + | 27.1069355755808 | 42.0688424563821 | 4.77291284147196 | 4.26843458790854 |
| chr17 | 80966293 | 80966311 | + | 2.00792115374673 | 1.13699574206438 | 2.38645642073598 | 2.13421729395427 |
| chr17 | 81117496 | 81117598 | - | 400.580270172472 | 177.371335762044 | 2.38645642073598 | 12.8053037637256 |
| chr17 | 81117720 | 81117742 | - | 12.0475269224804 | 4.54798296825753 | 1.19322821036799 | 3.20132594093141 |
| chr17 | 81241117 | 81241149 | + | 8.03168461498692 | 7.95897019445067 | 5.96614105183995 | 17.0737383516342 |
| chr17 | 81241178 | 81241250 | + | 13.0514874993537 | 23.876910583352 | 8.35259747257593 | 16.006629704657 |
| chr17 | 81464413 | 81464432 | + | 1.00396057687336 | 1.13699574206438 | 1.19322821036799 | 2.13421729395427 |
| chr17 | 81464451 | 81464474 | + | 9.03564519186028 | 2.27399148412876 | 5.96614105183995 | 3.20132594093141 |
| chr17 | 81464640 | 81464661 | + | 3.01188173062009 | 2.27399148412876 | 3.57968463110397 | 1.06710864697714 |
| chr17 | 81466173 | 81466178 | + | 1.00396057687336 | 1.13699574206438 | 1.19322821036799 | 2.13421729395427 |
| chr17 | 81509989 | 81510003 | - | 141.558441339144 | 210.344212281911 | 138.414472402687 | 56.5567582897882 |
| chr17 | 81510091 | 81510371 | - | 839.311042266133 | 620.799675167153 | 570.363084555899 | 586.909755837425 |
| chr17 | 81510406 | 81510525 | - | 35.1386201905678 | 28.4248935516096 | 101.424397881279 | 59.7580842307196 |
| chr17 | 81510579 | 81510756 | - | 179.708943260332 | 170.549361309657 | 156.312895558207 | 237.965228275901 |
| chr17 | 81539931 | 81540041 | - | 74.293082688629 | 48.8908169087684 | 65.6275515702394 | 157.932079752616 |
| chr17 | 81556912 | 81556927 | - | 4.01584230749346 | 6.82197445238629 | 5.96614105183995 | 6.40265188186281 |
| chr17 | 81557336 | 81557369 | - | 10.0396057687336 | 6.82197445238629 | 20.2848795762558 | 21.3421729395427 |
| chr17 | 81557440 | 81557474 | - | 3.01188173062009 | 2.27399148412876 | 2.38645642073598 | 10.6710864697714 |
| chr17 | 81557566 | 81557609 | - | 1.00396057687336 | 3.41098722619315 | 9.54582568294392 | 6.40265188186281 |
| chr17 | 81557775 | 81557802 | - | 7.02772403811355 | 2.27399148412876 | 9.54582568294392 | 10.6710864697714 |
| chr17 | 81557914 | 81557922 | - | 3.01188173062009 | 1.13699574206438 | 2.38645642073598 | 9.60397782279422 |
| chr17 | 81557955 | 81557958 | - | 4.01584230749346 | 6.82197445238629 | 4.77291284147196 | 14.9395210576799 |
| chr17 | 81558607 | 81558674 | - | 13.0514874993537 | 11.3699574206438 | 35.7968463110397 | 25.6106075274513 |
| chr17 | 81558734 | 81558768 | - | 2.00792115374673 | 1.13699574206438 | 1.19322821036799 | 5.33554323488568 |
| chr17 | 81665093 | 81665106 | - | 6.02376346124019 | 10.2329616785794 | 7.15936926220794 | 8.53686917581709 |
| chr17 | 81665251 | 81665272 | - | 1.00396057687336 | 1.13699574206438 | 4.77291284147196 | 10.6710864697714 |
| chr17 | 81665293 | 81665311 | - | 3.01188173062009 | 9.09596593651506 | 7.15936926220794 | 12.8053037637256 |
| chr17 | 81666268 | 81666317 | - | 11.043566345607 | 40.9318467143178 | 5.96614105183995 | 12.8053037637256 |
| chr17 | 81672842 | 81672862 | + | 1.00396057687336 | 1.13699574206438 | 4.77291284147196 | 3.20132594093141 |
| chr17 | 81673152 | 81673154 | + | 3.01188173062009 | 5.68497871032191 | 1.19322821036799 | 2.13421729395427 |
| chr17 | 81673207 | 81673258 | + | 9.03564519186028 | 1.13699574206438 | 20.2848795762558 | 9.60397782279422 |
| chr17 | 81673470 | 81673483 | + | 1.00396057687336 | 2.27399148412876 | 8.35259747257593 | 2.13421729395427 |
| chr17 | 81673821 | 81673839 | + | 5.01980288436682 | 3.41098722619315 | 1.19322821036799 | 1.06710864697714 |
| chr17 | 81681232 | 81681235 | - | 3.01188173062009 | 5.68497871032191 | 4.77291284147196 | 2.13421729395427 |
| chr17 | 81701967 | 81702004 | + | 3.01188173062009 | 3.41098722619315 | 1.19322821036799 | 8.53686917581709 |
| chr17 | 81702025 | 81702096 | + | 44.174265382428 | 22.7399148412876 | 20.2848795762558 | 22.4092815865198 |
| chr17 | 81707121 | 81707304 | + | 77.3049644192491 | 46.6168254246397 | 73.9801490428153 | 139.791232754005 |
| chr17 | 81707331 | 81707390 | + | 39.1544624980612 | 29.5618892936739 | 36.9900745214077 | 43.7514545260626 |
| chr17 | 81707418 | 81707496 | + | 199.7881547978 | 131.891506079468 | 63.2410951495034 | 146.193884635868 |
| chr17 | 81720108 | 81720128 | + | 2.00792115374673 | 1.13699574206438 | 1.19322821036799 | 6.40265188186281 |
| chr17 | 81720345 | 81720365 | + | 4.01584230749346 | 2.27399148412876 | 7.15936926220794 | 19.2079556455884 |
| chr17 | 81720392 | 81720402 | + | 11.043566345607 | 6.82197445238629 | 34.6036181006717 | 6.40265188186281 |
| chr17 | 81720428 | 81720448 | + | 2.00792115374673 | 1.13699574206438 | 1.19322821036799 | 4.26843458790854 |
| chr17 | 81720963 | 81720984 | + | 5.01980288436682 | 6.82197445238629 | 3.57968463110397 | 8.53686917581709 |
| chr17 | 81843160 | 81843207 | + | 6.02376346124019 | 5.68497871032191 | 3.57968463110397 | 1.06710864697714 |
| chr17 | 81843176 | 81843932 | - | 197.780233644053 | 222.851165444619 | 233.872729232126 | 219.82438127729 |
| chr17 | 81843955 | 81843993 | - | 1.00396057687336 | 2.27399148412876 | 1.19322821036799 | 1.06710864697714 |
| chr17 | 81867740 | 81867783 | - | 18.0712903837206 | 29.5618892936739 | 4.77291284147196 | 13.8724124107028 |
| chr17 | 81867805 | 81867880 | - | 3.01188173062009 | 3.41098722619315 | 5.96614105183995 | 7.46976052883995 |
| chr17 | 81868279 | 81868491 | - | 87.3445701879827 | 47.753821166704 | 66.8207797806074 | 68.2949534065367 |
| chr17 | 81868583 | 81868608 | - | 5.01980288436682 | 9.09596593651506 | 3.57968463110397 | 7.46976052883995 |
| chr17 | 81868629 | 81868633 | - | 47.1861471130481 | 25.0139063254164 | 32.2171616799357 | 12.8053037637256 |
| chr17 | 81868663 | 81868704 | - | 7.02772403811355 | 2.27399148412876 | 4.77291284147196 | 13.8724124107028 |
| chr17 | 81868809 | 81868862 | - | 7.02772403811355 | 6.82197445238629 | 1.19322821036799 | 4.26843458790854 |
| chr17 | 81887873 | 81887959 | - | 28.1108961524542 | 39.7948509722534 | 31.0239334695677 | 59.7580842307196 |
| chr17 | 81888040 | 81888070 | - | 21.0831721143407 | 3.41098722619315 | 17.8984231555198 | 36.2816939972226 |
| chr17 | 81899919 | 81899962 | + | 28.1108961524542 | 21.6029190992233 | 33.4103898903037 | 43.7514545260626 |
| chr17 | 81900009 | 81900101 | + | 118.467348071057 | 128.480518853275 | 178.984231555198 | 274.246922273124 |
| chr17 | 81900133 | 81900236 | + | 55.217831728035 | 112.562578464374 | 141.994157033791 | 132.321472225165 |
| chr17 | 81904273 | 81904277 | - | 3.01188173062009 | 2.27399148412876 | 5.96614105183995 | 4.26843458790854 |
| chr17 | 81904308 | 81904398 | - | 3.01188173062009 | 2.27399148412876 | 9.54582568294392 | 7.46976052883995 |
| chr17 | 81904420 | 81904485 | - | 3.01188173062009 | 2.27399148412876 | 4.77291284147196 | 5.33554323488568 |
| chr17 | 81911983 | 81912007 | - | 3.01188173062009 | 1.13699574206438 | 1.19322821036799 | 5.33554323488568 |
| chr17 | 81912316 | 81912360 | - | 10.0396057687336 | 10.2329616785794 | 14.3187385244159 | 5.33554323488568 |
| chr17 | 81918333 | 81918378 | - | 5.01980288436682 | 4.54798296825753 | 2.38645642073598 | 3.20132594093141 |
| chr17 | 81918831 | 81918842 | - | 4.01584230749346 | 3.41098722619315 | 1.19322821036799 | 2.13421729395427 |
| chr17 | 81919059 | 81919069 | - | 10.0396057687336 | 2.27399148412876 | 7.15936926220794 | 5.33554323488568 |
| chr17 | 81919666 | 81919684 | - | 9.03564519186028 | 5.68497871032191 | 11.9322821036799 | 7.46976052883995 |
| chr17 | 81919731 | 81919780 | - | 2.00792115374673 | 3.41098722619315 | 4.77291284147196 | 4.26843458790854 |
| chr17 | 81921444 | 81921459 | - | 5.01980288436682 | 3.41098722619315 | 1.19322821036799 | 1.06710864697714 |
| chr17 | 81921634 | 81921645 | - | 6.02376346124019 | 1.13699574206438 | 9.54582568294392 | 3.20132594093141 |
| chr17 | 81921873 | 81921877 | - | 10.0396057687336 | 7.95897019445067 | 13.1255103140479 | 1.06710864697714 |
| chr17 | 81921901 | 81922052 | - | 82.3247673036159 | 79.5897019445067 | 41.7629873628796 | 21.3421729395427 |
| chr17 | 81932404 | 81932522 | - | 28.1108961524542 | 60.2607743294122 | 106.197310722751 | 83.2344744642166 |
| chr17 | 81932570 | 81932628 | - | 3.01188173062009 | 6.82197445238629 | 3.57968463110397 | 5.33554323488568 |
| chr17 | 81932676 | 81932990 | - | 25.0990144218341 | 27.2878978095452 | 189.72328544851 | 182.47557863309 |
| chr17 | 81932998 | 81933171 | - | 23.0910932680874 | 28.4248935516096 | 121.709277457535 | 106.710864697714 |
| chr17 | 81933210 | 81933376 | - | 6.02376346124019 | 9.09596593651506 | 33.4103898903037 | 22.4092815865198 |
| chr17 | 82017309 | 82017388 | + | 12.0475269224804 | 14.780944646837 | 8.35259747257593 | 5.33554323488568 |
| chr17 | 82018730 | 82018810 | - | 91.3604124954762 | 117.110561432631 | 81.1395183050233 | 117.381951167485 |
| chr17 | 82018840 | 82018857 | - | 7.02772403811355 | 2.27399148412876 | 3.57968463110397 | 6.40265188186281 |
| chr17 | 82018884 | 82018925 | - | 1.00396057687336 | 3.41098722619315 | 3.57968463110397 | 13.8724124107028 |
| chr17 | 82018947 | 82018996 | - | 8.03168461498692 | 10.2329616785794 | 14.3187385244159 | 9.60397782279422 |
| chr17 | 82019017 | 82019050 | - | 1.00396057687336 | 2.27399148412876 | 7.15936926220794 | 13.8724124107028 |
| chr17 | 82019083 | 82019123 | - | 5.01980288436682 | 1.13699574206438 | 14.3187385244159 | 9.60397782279422 |
| chr17 | 82031111 | 82031133 | + | 11.043566345607 | 17.0549361309657 | 7.15936926220794 | 4.26843458790854 |
| chr17 | 82034124 | 82034134 | + | 1.00396057687336 | 2.27399148412876 | 4.77291284147196 | 10.6710864697714 |
| chr17 | 82034155 | 82034170 | + | 2.00792115374673 | 2.27399148412876 | 47.7291284147196 | 56.5567582897882 |
| chr17 | 82035903 | 82036030 | - | 447.766417285521 | 272.878978095452 | 835.259747257593 | 893.169937519863 |
| chr17 | 82047923 | 82047942 | - | 6.02376346124019 | 4.54798296825753 | 5.96614105183995 | 4.26843458790854 |
| chr17 | 82057355 | 82057449 | + | 61.2415951892752 | 84.1376849127643 | 44.1494437836156 | 76.8318225823538 |
| chr17 | 82057919 | 82057948 | - | 5.01980288436682 | 5.68497871032191 | 3.57968463110397 | 6.40265188186281 |
| chr17 | 82058068 | 82058085 | - | 2.00792115374673 | 1.13699574206438 | 1.19322821036799 | 3.20132594093141 |
| chr17 | 82058112 | 82058182 | - | 9.03564519186028 | 4.54798296825753 | 9.54582568294392 | 7.46976052883995 |
| chr17 | 82078494 | 82078547 | - | 40.1584230749346 | 25.0139063254164 | 97.8447132501751 | 118.449059814462 |
| chr17 | 82078735 | 82078801 | - | 20.0792115374673 | 6.82197445238629 | 81.1395183050233 | 81.1002571702623 |
| chr17 | 82078862 | 82078973 | - | 32.1267384599477 | 9.09596593651506 | 159.892580189311 | 207.019077513564 |
| chr17 | 82078997 | 82079013 | - | 8.03168461498692 | 7.95897019445067 | 17.8984231555198 | 13.8724124107028 |
| chr17 | 82079035 | 82079079 | - | 15.0594086531005 | 6.82197445238629 | 32.2171616799357 | 57.6238669367653 |
| chr17 | 82079113 | 82079215 | - | 10.0396057687336 | 4.54798296825753 | 72.7869208324474 | 35.2145853502455 |
| chr17 | 82101598 | 82101610 | - | 6.02376346124019 | 3.41098722619315 | 3.57968463110397 | 1.06710864697714 |
| chr17 | 82238655 | 82239200 | + | 325.28322690697 | 227.399148412876 | 57.2749540976635 | 23.476390233497 |
| chr17 | 82239376 | 82239478 | + | 44.174265382428 | 51.1648083928972 | 2.38645642073598 | 2.13421729395427 |
| chr17 | 82242701 | 82242759 | - | 2.00792115374673 | 3.41098722619315 | 7.15936926220794 | 5.33554323488568 |
| chr17 | 82243052 | 82243074 | - | 5.01980288436682 | 3.41098722619315 | 5.96614105183995 | 22.4092815865198 |
| chr17 | 82243627 | 82243644 | - | 2.00792115374673 | 2.27399148412876 | 4.77291284147196 | 6.40265188186281 |
| chr17 | 82244180 | 82244322 | - | 11.043566345607 | 14.780944646837 | 46.5359002043516 | 71.4962793474681 |
| chr17 | 82244366 | 82244550 | - | 12.0475269224804 | 7.95897019445067 | 27.4442488384638 | 13.8724124107028 |
| chr17 | 82244668 | 82244696 | - | 1.00396057687336 | 2.27399148412876 | 5.96614105183995 | 3.20132594093141 |
| chr17 | 82245356 | 82245357 | + | 5.01980288436682 | 4.54798296825753 | 5.96614105183995 | 1.06710864697714 |
| chr17 | 82294047 | 82294054 | + | 1.00396057687336 | 3.41098722619315 | 1.19322821036799 | 1.06710864697714 |
| chr17 | 82294159 | 82294163 | + | 1.00396057687336 | 1.13699574206438 | 4.77291284147196 | 4.26843458790854 |
| chr17 | 82294392 | 82294393 | + | 4.01584230749346 | 3.41098722619315 | 2.38645642073598 | 3.20132594093141 |
| chr17 | 82294689 | 82294777 | + | 11.043566345607 | 20.4659233571589 | 35.7968463110397 | 11.7381951167485 |
| chr17 | 82321377 | 82321394 | - | 5.01980288436682 | 5.68497871032191 | 1.19322821036799 | 1.06710864697714 |
| chr17 | 82392244 | 82392268 | - | 2.00792115374673 | 3.41098722619315 | 5.96614105183995 | 2.13421729395427 |
| chr17 | 82442521 | 82442609 | + | 2.00792115374673 | 7.95897019445067 | 1.19322821036799 | 4.26843458790854 |
| chr17 | 82442666 | 82442684 | - | 9.03564519186028 | 11.3699574206438 | 19.0916513658878 | 11.7381951167485 |
| chr17 | 82488152 | 82488190 | + | 3.01188173062009 | 2.27399148412876 | 5.96614105183995 | 6.40265188186281 |
| chr17 | 82488215 | 82488238 | + | 15.0594086531005 | 14.780944646837 | 60.8546387287675 | 85.3686917581709 |
| chr17 | 82489984 | 82489985 | + | 2.00792115374673 | 3.41098722619315 | 1.19322821036799 | 1.06710864697714 |
| chr17 | 82601468 | 82601527 | + | 12.0475269224804 | 10.2329616785794 | 33.4103898903037 | 13.8724124107028 |
| chr17 | 82601555 | 82601596 | + | 2.00792115374673 | 10.2329616785794 | 16.7051949451519 | 5.33554323488568 |
| chr17 | 82601854 | 82601859 | + | 2.00792115374673 | 2.27399148412876 | 1.19322821036799 | 1.06710864697714 |
| chr17 | 82601950 | 82601958 | + | 1.00396057687336 | 2.27399148412876 | 3.57968463110397 | 2.13421729395427 |
| chr17 | 82602205 | 82602223 | + | 6.02376346124019 | 5.68497871032191 | 2.38645642073598 | 5.33554323488568 |
| chr17 | 82602244 | 82602310 | + | 11.043566345607 | 17.0549361309657 | 8.35259747257593 | 13.8724124107028 |
| chr17 | 82602334 | 82602355 | + | 2.00792115374673 | 2.27399148412876 | 3.57968463110397 | 2.13421729395427 |
| chr17 | 82602812 | 82602872 | + | 13.0514874993537 | 12.5069531627082 | 20.2848795762558 | 28.8119334683827 |
| chr17 | 82602957 | 82602978 | + | 2.00792115374673 | 3.41098722619315 | 5.96614105183995 | 1.06710864697714 |
| chr17 | 82603477 | 82603499 | + | 2.00792115374673 | 3.41098722619315 | 5.96614105183995 | 7.46976052883995 |
| chr17 | 82603567 | 82603573 | + | 5.01980288436682 | 3.41098722619315 | 5.96614105183995 | 5.33554323488568 |
| chr17 | 82603628 | 82603645 | + | 1.00396057687336 | 3.41098722619315 | 3.57968463110397 | 7.46976052883995 |
| chr17 | 82603667 | 82603710 | + | 10.0396057687336 | 15.9179403889013 | 19.0916513658878 | 33.0803680562912 |
| chr17 | 82604419 | 82604447 | + | 17.0673298068472 | 20.4659233571589 | 10.7390538933119 | 26.6777161744284 |
| chr17 | 82604516 | 82604541 | + | 3.01188173062009 | 3.41098722619315 | 1.19322821036799 | 3.20132594093141 |
| chr17 | 82614660 | 82614760 | - | 34.1346596136944 | 39.7948509722534 | 16.7051949451519 | 20.2750642925656 |
| chr17 | 82614852 | 82614885 | - | 6.02376346124019 | 9.09596593651506 | 1.19322821036799 | 10.6710864697714 |
| chr17 | 82614910 | 82614937 | - | 6.02376346124019 | 14.780944646837 | 1.19322821036799 | 7.46976052883995 |
| chr17 | 82614970 | 82615041 | - | 1.00396057687336 | 6.82197445238629 | 1.19322821036799 | 3.20132594093141 |
| chr17 | 82615159 | 82615186 | - | 3.01188173062009 | 1.13699574206438 | 5.96614105183995 | 1.06710864697714 |
| chr17 | 82615563 | 82615616 | - | 10.0396057687336 | 7.95897019445067 | 10.7390538933119 | 11.7381951167485 |
| chr17 | 82727245 | 82727285 | + | 2.00792115374673 | 1.13699574206438 | 7.15936926220794 | 11.7381951167485 |
| chr17 | 82727301 | 82727335 | + | 10.0396057687336 | 3.41098722619315 | 13.1255103140479 | 7.46976052883995 |
| chr17 | 82727381 | 82727412 | + | 5.01980288436682 | 4.54798296825753 | 2.38645642073598 | 20.2750642925656 |
| chr17 | 82727483 | 82727564 | + | 18.0712903837206 | 7.95897019445067 | 4.77291284147196 | 4.26843458790854 |
| chr17 | 82727573 | 82727690 | + | 64.2534769198953 | 77.315710460378 | 48.9223566250876 | 50.1541064079254 |
| chr17 | 82727715 | 82727734 | + | 12.0475269224804 | 7.95897019445067 | 2.38645642073598 | 21.3421729395427 |
| chr18 | 211237 | 211312 | + | 14.0554480762271 | 10.2329616785794 | 33.4103898903037 | 27.7448248214055 |
| chr18 | 211350 | 211527 | + | 56.2217923049084 | 54.5757956190903 | 112.163451774591 | 116.314842520508 |
| chr18 | 211552 | 211590 | + | 67.2653586505154 | 39.7948509722534 | 81.1395183050233 | 59.7580842307196 |
| chr18 | 211622 | 211699 | + | 37.1465413443145 | 20.4659233571589 | 56.0817258872955 | 57.6238669367653 |
| chr18 | 211734 | 211755 | + | 118.467348071057 | 105.740604011988 | 220.747218918078 | 193.146665102862 |
| chr18 | 211947 | 211949 | + | 2.00792115374673 | 5.68497871032191 | 4.77291284147196 | 4.26843458790854 |
| chr18 | 212393 | 212394 | + | 3.01188173062009 | 1.13699574206438 | 1.19322821036799 | 1.06710864697714 |
| chr18 | 212968 | 212999 | + | 2.00792115374673 | 5.68497871032191 | 4.77291284147196 | 4.26843458790854 |
| chr18 | 213068 | 213092 | + | 1.00396057687336 | 1.13699574206438 | 4.77291284147196 | 9.60397782279422 |
| chr18 | 213726 | 213729 | + | 2.00792115374673 | 3.41098722619315 | 4.77291284147196 | 3.20132594093141 |
| chr18 | 214544 | 214594 | - | 15.0594086531005 | 17.0549361309657 | 26.2510206280958 | 20.2750642925656 |
| chr18 | 214571 | 214577 | + | 1.00396057687336 | 1.13699574206438 | 3.57968463110397 | 3.20132594093141 |
| chr18 | 214628 | 214641 | - | 4.01584230749346 | 4.54798296825753 | 11.9322821036799 | 14.9395210576799 |
| chr18 | 214726 | 214731 | - | 14.0554480762271 | 9.09596593651506 | 8.35259747257593 | 11.7381951167485 |
| chr18 | 214763 | 214782 | - | 2.00792115374673 | 3.41098722619315 | 2.38645642073598 | 2.13421729395427 |
| chr18 | 216580 | 216610 | - | 4.01584230749346 | 2.27399148412876 | 11.9322821036799 | 7.46976052883995 |
| chr18 | 218948 | 218971 | - | 1.00396057687336 | 4.54798296825753 | 14.3187385244159 | 7.46976052883995 |
| chr18 | 223439 | 223446 | - | 1.00396057687336 | 1.13699574206438 | 17.8984231555198 | 2.13421729395427 |
| chr18 | 252549 | 252580 | - | 4.01584230749346 | 1.13699574206438 | 13.1255103140479 | 3.20132594093141 |
| chr18 | 673036 | 673060 | + | 35.1386201905678 | 30.6988850357383 | 20.2848795762558 | 17.0737383516342 |
| chr18 | 673089 | 673114 | + | 8.03168461498692 | 4.54798296825753 | 4.77291284147196 | 2.13421729395427 |
| chr18 | 673116 | 673152 | + | 4.01584230749346 | 3.41098722619315 | 4.77291284147196 | 5.33554323488568 |
| chr18 | 673188 | 673262 | + | 27.1069355755808 | 23.876910583352 | 38.1833027317757 | 38.4159112911769 |
| chr18 | 673300 | 673331 | + | 15.0594086531005 | 19.3289276150945 | 21.4781077866238 | 28.8119334683827 |
| chr18 | 673390 | 673485 | + | 209.827760566533 | 246.728076027971 | 288.761226909053 | 476.99756519878 |
| chr18 | 674066 | 674071 | - | 7.02772403811355 | 17.0549361309657 | 2.38645642073598 | 2.13421729395427 |
| chr18 | 674165 | 674166 | - | 2.00792115374673 | 2.27399148412876 | 3.57968463110397 | 5.33554323488568 |
| chr18 | 722321 | 722371 | - | 1.00396057687336 | 3.41098722619315 | 1.19322821036799 | 8.53686917581709 |
| chr18 | 722676 | 722737 | - | 2.00792115374673 | 1.13699574206438 | 9.54582568294392 | 11.7381951167485 |
| chr18 | 722780 | 722791 | - | 1.00396057687336 | 1.13699574206438 | 1.19322821036799 | 2.13421729395427 |
| chr18 | 722814 | 722829 | - | 2.00792115374673 | 4.54798296825753 | 4.77291284147196 | 8.53686917581709 |
| chr18 | 723342 | 723395 | - | 1.00396057687336 | 1.13699574206438 | 4.77291284147196 | 5.33554323488568 |
| chr18 | 723429 | 723454 | - | 1.00396057687336 | 1.13699574206438 | 15.5119667347839 | 8.53686917581709 |
| chr18 | 723510 | 723545 | - | 3.01188173062009 | 2.27399148412876 | 20.2848795762558 | 7.46976052883995 |
| chr18 | 723617 | 723627 | - | 25.0990144218341 | 35.2468680039958 | 73.9801490428153 | 54.4225409958339 |
| chr18 | 723669 | 723720 | - | 10.0396057687336 | 14.780944646837 | 33.4103898903037 | 19.2079556455884 |
| chr18 | 723757 | 723764 | - | 1.00396057687336 | 4.54798296825753 | 1.19322821036799 | 1.06710864697714 |
| chr18 | 724123 | 724215 | - | 6.02376346124019 | 9.09596593651506 | 29.8307052591997 | 17.0737383516342 |
| chr18 | 724273 | 724306 | - | 1.00396057687336 | 3.41098722619315 | 7.15936926220794 | 7.46976052883995 |
| chr18 | 813277 | 813305 | + | 1.00396057687336 | 1.13699574206438 | 3.57968463110397 | 2.13421729395427 |
| chr18 | 2802795 | 2802836 | + | 1.00396057687336 | 3.41098722619315 | 5.96614105183995 | 3.20132594093141 |
| chr18 | 2802991 | 2803012 | + | 3.01188173062009 | 3.41098722619315 | 5.96614105183995 | 4.26843458790854 |
| chr18 | 2803128 | 2803151 | + | 2.00792115374673 | 2.27399148412876 | 1.19322821036799 | 3.20132594093141 |
| chr18 | 2917026 | 2917110 | - | 1.00396057687336 | 7.95897019445067 | 3.57968463110397 | 1.06710864697714 |
| chr18 | 2917224 | 2917234 | - | 13.0514874993537 | 55.7127913611547 | 9.54582568294392 | 20.2750642925656 |
| chr18 | 2917345 | 2917387 | - | 2.00792115374673 | 7.95897019445067 | 1.19322821036799 | 3.20132594093141 |
| chr18 | 2920174 | 2920192 | - | 1.00396057687336 | 1.13699574206438 | 1.19322821036799 | 1.06710864697714 |
| chr18 | 3247249 | 3247302 | + | 10.0396057687336 | 7.95897019445067 | 45.3426719939836 | 50.1541064079254 |
| chr18 | 3255935 | 3255972 | + | 4.01584230749346 | 5.68497871032191 | 14.3187385244159 | 8.53686917581709 |
| chr18 | 3256016 | 3256070 | + | 2.00792115374673 | 5.68497871032191 | 13.1255103140479 | 5.33554323488568 |
| chr18 | 3256111 | 3256214 | + | 31.1227778830743 | 67.0827487817985 | 89.4921157775992 | 56.5567582897882 |
| chr18 | 3278106 | 3278178 | + | 82.3247673036159 | 137.57648478979 | 65.6275515702394 | 36.2816939972226 |
| chr18 | 3278182 | 3278214 | + | 8.03168461498692 | 13.6439489047726 | 8.35259747257593 | 2.13421729395427 |
| chr18 | 3457964 | 3457989 | + | 10.0396057687336 | 26.1509020674808 | 2.38645642073598 | 4.26843458790854 |
| chr18 | 3458162 | 3458191 | + | 10.0396057687336 | 29.5618892936739 | 5.96614105183995 | 4.26843458790854 |
| chr18 | 3458244 | 3458317 | + | 12.0475269224804 | 34.1098722619315 | 8.35259747257593 | 10.6710864697714 |
| chr18 | 3594029 | 3594235 | + | 1928.60826817373 | 1921.52280408881 | 127.675418509375 | 129.120146284233 |
| chr18 | 3594359 | 3594511 | + | 783.089249961224 | 710.622338790239 | 328.137757851197 | 205.951968866587 |
| chr18 | 3597018 | 3597056 | + | 27.1069355755808 | 15.9179403889013 | 5.96614105183995 | 13.8724124107028 |
| chr18 | 3603320 | 3603351 | + | 4.01584230749346 | 1.13699574206438 | 13.1255103140479 | 20.2750642925656 |
| chr18 | 3603609 | 3603752 | + | 50.1980288436682 | 96.6446380754725 | 171.82486229299 | 165.401840281456 |
| chr18 | 3603884 | 3603908 | + | 2.00792115374673 | 2.27399148412876 | 3.57968463110397 | 6.40265188186281 |
| chr18 | 3604102 | 3604134 | + | 14.0554480762271 | 26.1509020674808 | 1.19322821036799 | 4.26843458790854 |
| chr18 | 3604181 | 3604236 | + | 17.0673298068472 | 22.7399148412876 | 19.0916513658878 | 27.7448248214055 |
| chr18 | 3607114 | 3607125 | + | 2.00792115374673 | 3.41098722619315 | 4.77291284147196 | 5.33554323488568 |
| chr18 | 3607435 | 3607455 | + | 2.00792115374673 | 6.82197445238629 | 2.38645642073598 | 2.13421729395427 |
| chr18 | 3655887 | 3655913 | + | 8.03168461498692 | 6.82197445238629 | 1.19322821036799 | 2.13421729395427 |
| chr18 | 3656132 | 3656135 | + | 3.01188173062009 | 4.54798296825753 | 1.19322821036799 | 3.20132594093141 |
| chr18 | 5238057 | 5238189 | + | 349.378280751931 | 577.593836968706 | 723.096295483002 | 758.714248000744 |
| chr18 | 8638839 | 8638844 | + | 2.00792115374673 | 1.13699574206438 | 2.38645642073598 | 2.13421729395427 |
| chr18 | 9102595 | 9102789 | + | 3042.00054792629 | 2543.45947499802 | 4630.91868443817 | 5431.58301311362 |
| chr18 | 9117835 | 9117848 | + | 3.01188173062009 | 3.41098722619315 | 7.15936926220794 | 6.40265188186281 |
| chr18 | 9126878 | 9126908 | + | 6.02376346124019 | 6.82197445238629 | 27.4442488384638 | 24.5434988804741 |
| chr18 | 9134204 | 9134262 | + | 85.336649034236 | 102.329616785794 | 143.187385244159 | 136.589906813073 |
| chr18 | 9134304 | 9134324 | + | 55.217831728035 | 56.8497871032191 | 62.0478669391355 | 56.5567582897882 |
| chr18 | 9399764 | 9399800 | + | 1.00396057687336 | 1.13699574206438 | 13.1255103140479 | 4.26843458790854 |
| chr18 | 9400390 | 9400453 | + | 10.0396057687336 | 10.2329616785794 | 44.1494437836156 | 23.476390233497 |
| chr18 | 9400549 | 9400557 | + | 1.00396057687336 | 1.13699574206438 | 11.9322821036799 | 3.20132594093141 |
| chr18 | 9401145 | 9401191 | + | 7.02772403811355 | 4.54798296825753 | 39.3765309421437 | 17.0737383516342 |
| chr18 | 9402076 | 9402090 | + | 1.00396057687336 | 3.41098722619315 | 7.15936926220794 | 3.20132594093141 |
| chr18 | 9536002 | 9536010 | + | 1.00396057687336 | 3.41098722619315 | 20.2848795762558 | 6.40265188186281 |
| chr18 | 9536073 | 9536116 | + | 4.01584230749346 | 5.68497871032191 | 25.0577924177278 | 7.46976052883995 |
| chr18 | 9536147 | 9536179 | + | 3.01188173062009 | 6.82197445238629 | 9.54582568294392 | 9.60397782279422 |
| chr18 | 9536618 | 9536626 | + | 8.03168461498692 | 3.41098722619315 | 14.3187385244159 | 16.006629704657 |
| chr18 | 9536709 | 9536740 | + | 2.00792115374673 | 3.41098722619315 | 8.35259747257593 | 13.8724124107028 |
| chr18 | 9536797 | 9536799 | + | 2.00792115374673 | 2.27399148412876 | 5.96614105183995 | 7.46976052883995 |
| chr18 | 9546824 | 9546829 | - | 14.0554480762271 | 12.5069531627082 | 13.1255103140479 | 9.60397782279422 |
| chr18 | 9546947 | 9546996 | - | 6.02376346124019 | 7.95897019445067 | 9.54582568294392 | 12.8053037637256 |
| chr18 | 9547037 | 9547046 | - | 133.526756724157 | 128.480518853275 | 177.79100334483 | 184.609795927044 |
| chr18 | 9547139 | 9547182 | - | 10.0396057687336 | 6.82197445238629 | 4.77291284147196 | 6.40265188186281 |
| chr18 | 9547204 | 9547264 | - | 7.02772403811355 | 4.54798296825753 | 1.19322821036799 | 4.26843458790854 |
| chr18 | 9859314 | 9859338 | + | 126.499032686044 | 81.8636934286355 | 15.5119667347839 | 89.6371263460794 |
| chr18 | 9859466 | 9859474 | + | 21.0831721143407 | 21.6029190992233 | 17.8984231555198 | 11.7381951167485 |
| chr18 | 9859566 | 9859570 | + | 10.0396057687336 | 22.7399148412876 | 4.77291284147196 | 10.6710864697714 |
| chr18 | 9859764 | 9859783 | + | 23.0910932680874 | 12.5069531627082 | 2.38645642073598 | 23.476390233497 |
| chr18 | 9860211 | 9860213 | + | 4.01584230749346 | 11.3699574206438 | 1.19322821036799 | 4.26843458790854 |
| chr18 | 9860651 | 9860654 | + | 7.02772403811355 | 7.95897019445067 | 4.77291284147196 | 9.60397782279422 |
| chr18 | 9860793 | 9860810 | + | 4.01584230749346 | 9.09596593651506 | 2.38645642073598 | 5.33554323488568 |
| chr18 | 9860899 | 9860909 | + | 5.01980288436682 | 4.54798296825753 | 1.19322821036799 | 2.13421729395427 |
| chr18 | 9861001 | 9861003 | + | 6.02376346124019 | 18.1919318730301 | 1.19322821036799 | 1.06710864697714 |
| chr18 | 9954112 | 9954230 | + | 35.1386201905678 | 51.1648083928972 | 127.675418509375 | 451.386957671328 |
| chr18 | 9954297 | 9954341 | + | 3.01188173062009 | 5.68497871032191 | 3.57968463110397 | 6.40265188186281 |
| chr18 | 9954396 | 9954408 | + | 4.01584230749346 | 10.2329616785794 | 9.54582568294392 | 12.8053037637256 |
| chr18 | 9954430 | 9954453 | + | 1.00396057687336 | 7.95897019445067 | 5.96614105183995 | 9.60397782279422 |
| chr18 | 9954479 | 9954498 | + | 1.00396057687336 | 1.13699574206438 | 1.19322821036799 | 1.06710864697714 |
| chr18 | 9954506 | 9954527 | + | 1.00396057687336 | 4.54798296825753 | 3.57968463110397 | 11.7381951167485 |
| chr18 | 9954607 | 9954627 | + | 4.01584230749346 | 3.41098722619315 | 3.57968463110397 | 1.06710864697714 |
| chr18 | 9954631 | 9954670 | + | 76.3010038423757 | 142.124467758048 | 68.0140079909754 | 178.207144045182 |
| chr18 | 9954691 | 9954781 | + | 19.0752509605939 | 39.7948509722534 | 25.0577924177278 | 51.2212150549025 |
| chr18 | 9956205 | 9956220 | + | 1.00396057687336 | 1.13699574206438 | 2.38645642073598 | 7.46976052883995 |
| chr18 | 9956371 | 9956372 | + | 5.01980288436682 | 2.27399148412876 | 1.19322821036799 | 1.06710864697714 |
| chr18 | 10550277 | 10550300 | + | 2.00792115374673 | 1.13699574206438 | 4.77291284147196 | 5.33554323488568 |
| chr18 | 10552302 | 10552367 | + | 8.03168461498692 | 14.780944646837 | 4.77291284147196 | 10.6710864697714 |
| chr18 | 11852294 | 11852326 | + | 11.043566345607 | 9.09596593651506 | 1.19322821036799 | 1.06710864697714 |
| chr18 | 11852369 | 11852535 | + | 109.431702879197 | 202.38524208746 | 32.2171616799357 | 35.2145853502455 |
| chr18 | 11852559 | 11852599 | + | 37.1465413443145 | 57.9867828452835 | 8.35259747257593 | 10.6710864697714 |
| chr18 | 11852681 | 11852686 | + | 14.0554480762271 | 20.4659233571589 | 1.19322821036799 | 2.13421729395427 |
| chr18 | 11853054 | 11853066 | + | 5.01980288436682 | 9.09596593651506 | 2.38645642073598 | 1.06710864697714 |
| chr18 | 12030502 | 12030759 | + | 835.295199958639 | 811.814959833969 | 22.6713359969918 | 17.0737383516342 |
| chr18 | 12030816 | 12030857 | + | 51.2019894205416 | 55.7127913611547 | 1.19322821036799 | 1.06710864697714 |
| chr18 | 12326082 | 12326116 | + | 10.0396057687336 | 3.41098722619315 | 16.7051949451519 | 10.6710864697714 |
| chr18 | 12326188 | 12326192 | + | 3.01188173062009 | 3.41098722619315 | 4.77291284147196 | 4.26843458790854 |
| chr18 | 12326216 | 12326235 | + | 4.01584230749346 | 1.13699574206438 | 4.77291284147196 | 1.06710864697714 |
| chr18 | 12326307 | 12326318 | + | 2.00792115374673 | 3.41098722619315 | 3.57968463110397 | 3.20132594093141 |
| chr18 | 12326343 | 12326347 | + | 1.00396057687336 | 1.13699574206438 | 2.38645642073598 | 11.7381951167485 |
| chr18 | 12326374 | 12326385 | + | 7.02772403811355 | 5.68497871032191 | 2.38645642073598 | 5.33554323488568 |
| chr18 | 12326424 | 12326449 | + | 18.0712903837206 | 26.1509020674808 | 15.5119667347839 | 21.3421729395427 |
| chr18 | 12326493 | 12326494 | + | 5.01980288436682 | 3.41098722619315 | 10.7390538933119 | 9.60397782279422 |
| chr18 | 12326516 | 12326536 | + | 21.0831721143407 | 17.0549361309657 | 20.2848795762558 | 6.40265188186281 |
| chr18 | 12329034 | 12329069 | - | 5.01980288436682 | 10.2329616785794 | 7.15936926220794 | 8.53686917581709 |
| chr18 | 12329143 | 12329187 | - | 14.0554480762271 | 17.0549361309657 | 2.38645642073598 | 17.0737383516342 |
| chr18 | 12329206 | 12329221 | - | 14.0554480762271 | 31.8358807778027 | 10.7390538933119 | 26.6777161744284 |
| chr18 | 12329243 | 12329286 | - | 8.03168461498692 | 20.4659233571589 | 7.15936926220794 | 16.006629704657 |
| chr18 | 12329720 | 12329721 | + | 2.00792115374673 | 1.13699574206438 | 1.19322821036799 | 2.13421729395427 |
| chr18 | 12444240 | 12444241 | + | 9.03564519186028 | 14.780944646837 | 1.19322821036799 | 6.40265188186281 |
| chr18 | 12447747 | 12447748 | + | 2.00792115374673 | 1.13699574206438 | 1.19322821036799 | 1.06710864697714 |
| chr18 | 12448735 | 12448766 | - | 4.01584230749346 | 2.27399148412876 | 1.19322821036799 | 5.33554323488568 |
| chr18 | 12448995 | 12449018 | - | 6.02376346124019 | 5.68497871032191 | 4.77291284147196 | 2.13421729395427 |
| chr18 | 12725660 | 12725717 | + | 14.0554480762271 | 25.0139063254164 | 16.7051949451519 | 10.6710864697714 |
| chr18 | 12785499 | 12785506 | - | 1.00396057687336 | 5.68497871032191 | 7.15936926220794 | 5.33554323488568 |
| chr18 | 12785616 | 12785670 | - | 18.0712903837206 | 32.9728765198671 | 11.9322821036799 | 18.1408469986113 |
| chr18 | 12985310 | 12985318 | + | 11.043566345607 | 10.2329616785794 | 13.1255103140479 | 16.006629704657 |
| chr18 | 12985402 | 12985418 | + | 1.00396057687336 | 1.13699574206438 | 4.77291284147196 | 11.7381951167485 |
| chr18 | 12985457 | 12985494 | + | 8.03168461498692 | 4.54798296825753 | 4.77291284147196 | 4.26843458790854 |
| chr18 | 12985537 | 12985563 | + | 7.02772403811355 | 5.68497871032191 | 4.77291284147196 | 3.20132594093141 |
| chr18 | 12985587 | 12985617 | + | 138.546559608524 | 209.207216539846 | 141.994157033791 | 209.153294807519 |
| chr18 | 12985666 | 12985700 | + | 12.0475269224804 | 18.1919318730301 | 21.4781077866238 | 11.7381951167485 |
| chr18 | 12985720 | 12985731 | + | 1.00396057687336 | 1.13699574206438 | 10.7390538933119 | 2.13421729395427 |
| chr18 | 12985822 | 12985826 | + | 2.00792115374673 | 4.54798296825753 | 7.15936926220794 | 1.06710864697714 |
| chr18 | 12985854 | 12985932 | + | 14.0554480762271 | 10.2329616785794 | 22.6713359969918 | 24.5434988804741 |
| chr18 | 12985963 | 12986027 | + | 2.00792115374673 | 4.54798296825753 | 19.0916513658878 | 8.53686917581709 |
| chr18 | 12986230 | 12986295 | + | 8.03168461498692 | 3.41098722619315 | 21.4781077866238 | 25.6106075274513 |
| chr18 | 12987470 | 12987491 | + | 1.00396057687336 | 1.13699574206438 | 5.96614105183995 | 3.20132594093141 |
| chr18 | 13124973 | 13124984 | + | 1.00396057687336 | 3.41098722619315 | 3.57968463110397 | 2.13421729395427 |
| chr18 | 13664910 | 13664929 | - | 1.00396057687336 | 2.27399148412876 | 4.77291284147196 | 1.06710864697714 |
| chr18 | 13665805 | 13665808 | - | 2.00792115374673 | 1.13699574206438 | 1.19322821036799 | 1.06710864697714 |
| chr18 | 13666295 | 13666301 | - | 1.00396057687336 | 2.27399148412876 | 1.19322821036799 | 11.7381951167485 |
| chr18 | 13666421 | 13666425 | - | 1.00396057687336 | 1.13699574206438 | 2.38645642073598 | 3.20132594093141 |
| chr18 | 13762328 | 13762345 | + | 1.00396057687336 | 1.13699574206438 | 7.15936926220794 | 3.20132594093141 |
| chr18 | 20949962 | 20949984 | - | 2.00792115374673 | 2.27399148412876 | 2.38645642073598 | 1.06710864697714 |
| chr18 | 20950723 | 20950736 | - | 2.00792115374673 | 9.09596593651506 | 3.57968463110397 | 3.20132594093141 |
| chr18 | 20951013 | 20951038 | - | 8.03168461498692 | 7.95897019445067 | 7.15936926220794 | 11.7381951167485 |
| chr18 | 21529409 | 21529472 | - | 15.0594086531005 | 20.4659233571589 | 28.6374770488317 | 9.60397782279422 |
| chr18 | 21529494 | 21529589 | - | 38.1505019211879 | 46.6168254246397 | 96.6514850398071 | 30.9461507623369 |
| chr18 | 21529641 | 21529725 | - | 52.205949997415 | 86.411676396893 | 71.5936926220794 | 75.7647139353766 |
| chr18 | 21529761 | 21529815 | - | 14.0554480762271 | 20.4659233571589 | 11.9322821036799 | 14.9395210576799 |
| chr18 | 21529853 | 21529883 | - | 5.01980288436682 | 18.1919318730301 | 14.3187385244159 | 22.4092815865198 |
| chr18 | 21530012 | 21530086 | - | 3.01188173062009 | 5.68497871032191 | 17.8984231555198 | 38.4159112911769 |
| chr18 | 21530221 | 21530243 | - | 1.00396057687336 | 2.27399148412876 | 11.9322821036799 | 8.53686917581709 |
| chr18 | 21623933 | 21623940 | + | 4.01584230749346 | 2.27399148412876 | 7.15936926220794 | 10.6710864697714 |
| chr18 | 21629065 | 21629080 | + | 9.03564519186028 | 3.41098722619315 | 13.1255103140479 | 17.0737383516342 |
| chr18 | 21629122 | 21629238 | + | 73.2891221117556 | 112.562578464374 | 82.3327465153913 | 106.710864697714 |
| chr18 | 21629296 | 21629345 | + | 22.087132691214 | 14.780944646837 | 40.5697591525116 | 44.8185631730397 |
| chr18 | 21629917 | 21629924 | + | 6.02376346124019 | 7.95897019445067 | 2.38645642073598 | 4.26843458790854 |
| chr18 | 21630005 | 21630033 | + | 3.01188173062009 | 2.27399148412876 | 1.19322821036799 | 1.06710864697714 |
| chr18 | 21630180 | 21630219 | + | 7.02772403811355 | 4.54798296825753 | 1.19322821036799 | 4.26843458790854 |
| chr18 | 21650943 | 21650947 | - | 17.0673298068472 | 14.780944646837 | 4.77291284147196 | 2.13421729395427 |
| chr18 | 21651104 | 21651105 | - | 1.00396057687336 | 1.13699574206438 | 1.19322821036799 | 2.13421729395427 |
| chr18 | 21651142 | 21651173 | - | 20.0792115374673 | 28.4248935516096 | 7.15936926220794 | 1.06710864697714 |
| chr18 | 21651475 | 21651485 | - | 24.0950538449607 | 28.4248935516096 | 2.38645642073598 | 2.13421729395427 |
| chr18 | 21865683 | 21865726 | + | 1.00396057687336 | 2.27399148412876 | 8.35259747257593 | 14.9395210576799 |
| chr18 | 21865786 | 21865794 | + | 2.00792115374673 | 2.27399148412876 | 4.77291284147196 | 3.20132594093141 |
| chr18 | 21865921 | 21865929 | + | 2.00792115374673 | 1.13699574206438 | 3.57968463110397 | 3.20132594093141 |
| chr18 | 21866007 | 21866059 | + | 12.0475269224804 | 4.54798296825753 | 7.15936926220794 | 14.9395210576799 |
| chr18 | 21866255 | 21866277 | + | 2.00792115374673 | 2.27399148412876 | 5.96614105183995 | 5.33554323488568 |
| chr18 | 21866572 | 21866622 | + | 4.01584230749346 | 4.54798296825753 | 7.15936926220794 | 13.8724124107028 |
| chr18 | 21867140 | 21867155 | + | 2.00792115374673 | 3.41098722619315 | 4.77291284147196 | 3.20132594093141 |
| chr18 | 21867884 | 21867950 | + | 8.03168461498692 | 7.95897019445067 | 15.5119667347839 | 6.40265188186281 |
| chr18 | 21868472 | 21868511 | + | 6.02376346124019 | 3.41098722619315 | 5.96614105183995 | 8.53686917581709 |
| chr18 | 21869705 | 21869712 | + | 1.00396057687336 | 2.27399148412876 | 1.19322821036799 | 1.06710864697714 |
| chr18 | 21869789 | 21869796 | + | 5.01980288436682 | 4.54798296825753 | 2.38645642073598 | 3.20132594093141 |
| chr18 | 21870652 | 21870654 | + | 3.01188173062009 | 1.13699574206438 | 1.19322821036799 | 1.06710864697714 |
| chr18 | 22933313 | 22933448 | + | 310.22381825387 | 478.675207409105 | 101.424397881279 | 162.200514340525 |
| chr18 | 23026233 | 23026269 | + | 8.03168461498692 | 20.4659233571589 | 10.7390538933119 | 4.26843458790854 |
| chr18 | 23026411 | 23026414 | + | 9.03564519186028 | 17.0549361309657 | 2.38645642073598 | 1.06710864697714 |
| chr18 | 23297581 | 23297601 | - | 1.00396057687336 | 1.13699574206438 | 3.57968463110397 | 14.9395210576799 |
| chr18 | 23370260 | 23370264 | - | 1.00396057687336 | 3.41098722619315 | 5.96614105183995 | 3.20132594093141 |
| chr18 | 23481370 | 23481389 | + | 3.01188173062009 | 4.54798296825753 | 5.96614105183995 | 9.60397782279422 |
| chr18 | 23481525 | 23481555 | + | 3.01188173062009 | 2.27399148412876 | 4.77291284147196 | 2.13421729395427 |
| chr18 | 23481607 | 23481642 | + | 4.01584230749346 | 7.95897019445067 | 11.9322821036799 | 24.5434988804741 |
| chr18 | 23481910 | 23481945 | + | 1.00396057687336 | 3.41098722619315 | 7.15936926220794 | 7.46976052883995 |
| chr18 | 23481996 | 23482030 | + | 1.00396057687336 | 9.09596593651506 | 23.8645642073598 | 19.2079556455884 |
| chr18 | 23482366 | 23482371 | + | 3.01188173062009 | 9.09596593651506 | 11.9322821036799 | 14.9395210576799 |
| chr18 | 23482479 | 23482515 | + | 9.03564519186028 | 18.1919318730301 | 70.4004644117114 | 26.6777161744284 |
| chr18 | 23482561 | 23482605 | + | 12.0475269224804 | 26.1509020674808 | 31.0239334695677 | 27.7448248214055 |
| chr18 | 23482626 | 23482666 | + | 8.03168461498692 | 40.9318467143178 | 31.0239334695677 | 25.6106075274513 |
| chr18 | 23482687 | 23482715 | + | 2.00792115374673 | 18.1919318730301 | 15.5119667347839 | 16.006629704657 |
| chr18 | 23482736 | 23482835 | + | 25.0990144218341 | 51.1648083928972 | 102.617626091647 | 50.1541064079254 |
| chr18 | 23482854 | 23482925 | + | 92.3643730723495 | 212.618203766039 | 369.900745214077 | 197.41509969077 |
| chr18 | 23482949 | 23483010 | + | 52.205949997415 | 131.891506079468 | 227.906588180286 | 84.3015831111937 |
| chr18 | 23531541 | 23531579 | - | 14.0554480762271 | 21.6029190992233 | 4.77291284147196 | 7.46976052883995 |
| chr18 | 23531671 | 23531689 | + | 1.00396057687336 | 4.54798296825753 | 3.57968463110397 | 5.33554323488568 |
| chr18 | 23532005 | 23532030 | - | 1.00396057687336 | 5.68497871032191 | 1.19322821036799 | 1.06710864697714 |
| chr18 | 26016275 | 26016279 | - | 3.01188173062009 | 6.82197445238629 | 1.19322821036799 | 1.06710864697714 |
| chr18 | 26016442 | 26016459 | - | 10.0396057687336 | 6.82197445238629 | 15.5119667347839 | 22.4092815865198 |
| chr18 | 26016768 | 26016778 | - | 2.00792115374673 | 1.13699574206438 | 4.77291284147196 | 22.4092815865198 |
| chr18 | 26017281 | 26017284 | - | 1.00396057687336 | 2.27399148412876 | 3.57968463110397 | 3.20132594093141 |
| chr18 | 26017371 | 26017373 | - | 8.03168461498692 | 5.68497871032191 | 1.19322821036799 | 3.20132594093141 |
| chr18 | 26017704 | 26017706 | - | 2.00792115374673 | 2.27399148412876 | 1.19322821036799 | 2.13421729395427 |
| chr18 | 26017780 | 26017788 | - | 12.0475269224804 | 11.3699574206438 | 4.77291284147196 | 10.6710864697714 |
| chr18 | 26017898 | 26017920 | - | 21.0831721143407 | 25.0139063254164 | 7.15936926220794 | 7.46976052883995 |
| chr18 | 26017948 | 26017980 | - | 32.1267384599477 | 30.6988850357383 | 31.0239334695677 | 30.9461507623369 |
| chr18 | 26018007 | 26018100 | - | 75.2970432655023 | 60.2607743294122 | 105.004082512383 | 72.5633879944452 |
| chr18 | 26018114 | 26018139 | - | 6.02376346124019 | 3.41098722619315 | 22.6713359969918 | 7.46976052883995 |
| chr18 | 26018173 | 26018336 | - | 56.2217923049084 | 52.3018041349616 | 39.3765309421437 | 13.8724124107028 |
| chr18 | 26390806 | 26390817 | + | 4.01584230749346 | 9.09596593651506 | 3.57968463110397 | 9.60397782279422 |
| chr18 | 31546770 | 31546843 | + | 3.01188173062009 | 1.13699574206438 | 9.54582568294392 | 3.20132594093141 |
| chr18 | 31546912 | 31546916 | + | 3.01188173062009 | 2.27399148412876 | 2.38645642073598 | 8.53686917581709 |
| chr18 | 31546959 | 31546987 | + | 5.01980288436682 | 2.27399148412876 | 4.77291284147196 | 4.26843458790854 |
| chr18 | 31547111 | 31547115 | + | 3.01188173062009 | 5.68497871032191 | 5.96614105183995 | 1.06710864697714 |
| chr18 | 31547835 | 31547851 | + | 7.02772403811355 | 5.68497871032191 | 17.8984231555198 | 10.6710864697714 |
| chr18 | 31548100 | 31548125 | + | 9.03564519186028 | 3.41098722619315 | 2.38645642073598 | 7.46976052883995 |
| chr18 | 31548347 | 31548372 | + | 3.01188173062009 | 1.13699574206438 | 1.19322821036799 | 5.33554323488568 |
| chr18 | 31548456 | 31548468 | + | 1.00396057687336 | 1.13699574206438 | 2.38645642073598 | 3.20132594093141 |
| chr18 | 31548491 | 31548530 | + | 6.02376346124019 | 2.27399148412876 | 3.57968463110397 | 1.06710864697714 |
| chr18 | 31548662 | 31548708 | + | 7.02772403811355 | 3.41098722619315 | 3.57968463110397 | 6.40265188186281 |
| chr18 | 31548751 | 31548761 | + | 2.00792115374673 | 1.13699574206438 | 1.19322821036799 | 1.06710864697714 |
| chr18 | 31830238 | 31830240 | - | 1.00396057687336 | 3.41098722619315 | 1.19322821036799 | 3.20132594093141 |
| chr18 | 31830507 | 31830551 | - | 4.01584230749346 | 4.54798296825753 | 7.15936926220794 | 5.33554323488568 |
| chr18 | 31830585 | 31830618 | - | 25.0990144218341 | 10.2329616785794 | 40.5697591525116 | 21.3421729395427 |
| chr18 | 31830661 | 31830670 | - | 46.1821865361748 | 48.8908169087684 | 53.6952694665595 | 28.8119334683827 |
| chr18 | 31943330 | 31943411 | + | 11.043566345607 | 7.95897019445067 | 25.0577924177278 | 33.0803680562912 |
| chr18 | 32068787 | 32068788 | + | 2.00792115374673 | 3.41098722619315 | 3.57968463110397 | 8.53686917581709 |
| chr18 | 32129237 | 32129239 | + | 1.00396057687336 | 3.41098722619315 | 5.96614105183995 | 9.60397782279422 |
| chr18 | 32130184 | 32130189 | + | 3.01188173062009 | 15.9179403889013 | 5.96614105183995 | 4.26843458790854 |
| chr18 | 32130492 | 32130503 | + | 1.00396057687336 | 1.13699574206438 | 4.77291284147196 | 1.06710864697714 |
| chr18 | 32772106 | 32772107 | + | 2.00792115374673 | 1.13699574206438 | 8.35259747257593 | 10.6710864697714 |
| chr18 | 35140891 | 35140910 | + | 1.00396057687336 | 1.13699574206438 | 2.38645642073598 | 1.06710864697714 |
| chr18 | 35141414 | 35141444 | + | 7.02772403811355 | 6.82197445238629 | 9.54582568294392 | 5.33554323488568 |
| chr18 | 35141513 | 35141652 | + | 5.01980288436682 | 4.54798296825753 | 15.5119667347839 | 11.7381951167485 |
| chr18 | 35141771 | 35141829 | + | 5.01980288436682 | 4.54798296825753 | 13.1255103140479 | 13.8724124107028 |
| chr18 | 35142891 | 35142909 | + | 6.02376346124019 | 1.13699574206438 | 2.38645642073598 | 3.20132594093141 |
| chr18 | 35243768 | 35243779 | + | 4.01584230749346 | 3.41098722619315 | 4.77291284147196 | 8.53686917581709 |
| chr18 | 35243811 | 35243817 | + | 5.01980288436682 | 3.41098722619315 | 7.15936926220794 | 4.26843458790854 |
| chr18 | 35243877 | 35243902 | + | 2.00792115374673 | 3.41098722619315 | 21.4781077866238 | 10.6710864697714 |
| chr18 | 35243969 | 35243976 | + | 2.00792115374673 | 2.27399148412876 | 3.57968463110397 | 4.26843458790854 |
| chr18 | 35244000 | 35244005 | + | 3.01188173062009 | 2.27399148412876 | 2.38645642073598 | 4.26843458790854 |
| chr18 | 35244044 | 35244064 | + | 2.00792115374673 | 1.13699574206438 | 3.57968463110397 | 7.46976052883995 |
| chr18 | 35246756 | 35246766 | + | 1.00396057687336 | 1.13699574206438 | 1.19322821036799 | 1.06710864697714 |
| chr18 | 35332627 | 35332653 | - | 4.01584230749346 | 4.54798296825753 | 1.19322821036799 | 3.20132594093141 |
| chr18 | 35332687 | 35332703 | - | 2.00792115374673 | 1.13699574206438 | 4.77291284147196 | 7.46976052883995 |
| chr18 | 35332739 | 35332768 | - | 6.02376346124019 | 2.27399148412876 | 11.9322821036799 | 5.33554323488568 |
| chr18 | 35332816 | 35332834 | - | 6.02376346124019 | 3.41098722619315 | 10.7390538933119 | 7.46976052883995 |
| chr18 | 35332854 | 35332908 | - | 4.01584230749346 | 4.54798296825753 | 23.8645642073598 | 6.40265188186281 |
| chr18 | 35333079 | 35333100 | - | 6.02376346124019 | 1.13699574206438 | 8.35259747257593 | 6.40265188186281 |
| chr18 | 35333566 | 35333583 | - | 2.00792115374673 | 1.13699574206438 | 14.3187385244159 | 11.7381951167485 |
| chr18 | 35333638 | 35333645 | - | 2.00792115374673 | 1.13699574206438 | 1.19322821036799 | 1.06710864697714 |
| chr18 | 35334269 | 35334276 | - | 7.02772403811355 | 4.54798296825753 | 13.1255103140479 | 7.46976052883995 |
| chr18 | 35335428 | 35335455 | - | 11.043566345607 | 11.3699574206438 | 8.35259747257593 | 10.6710864697714 |
| chr18 | 35335503 | 35335527 | - | 13.0514874993537 | 11.3699574206438 | 8.35259747257593 | 18.1408469986113 |
| chr18 | 35335596 | 35335625 | - | 5.01980288436682 | 2.27399148412876 | 5.96614105183995 | 1.06710864697714 |
| chr18 | 35335710 | 35335715 | - | 1.00396057687336 | 4.54798296825753 | 2.38645642073598 | 1.06710864697714 |
| chr18 | 35335841 | 35335875 | - | 4.01584230749346 | 6.82197445238629 | 16.7051949451519 | 4.26843458790854 |
| chr18 | 35335938 | 35335967 | - | 16.0633692299738 | 11.3699574206438 | 13.1255103140479 | 22.4092815865198 |
| chr18 | 35335985 | 35336018 | - | 7.02772403811355 | 3.41098722619315 | 8.35259747257593 | 8.53686917581709 |
| chr18 | 35336286 | 35336292 | - | 3.01188173062009 | 1.13699574206438 | 2.38645642073598 | 4.26843458790854 |
| chr18 | 35337322 | 35337347 | - | 18.0712903837206 | 11.3699574206438 | 21.4781077866238 | 9.60397782279422 |
| chr18 | 35337375 | 35337378 | - | 5.01980288436682 | 3.41098722619315 | 1.19322821036799 | 2.13421729395427 |
| chr18 | 35337484 | 35337520 | - | 2.00792115374673 | 2.27399148412876 | 16.7051949451519 | 11.7381951167485 |
| chr18 | 35468360 | 35468393 | - | 3.01188173062009 | 5.68497871032191 | 2.38645642073598 | 3.20132594093141 |
| chr18 | 35468442 | 35468446 | - | 4.01584230749346 | 2.27399148412876 | 1.19322821036799 | 2.13421729395427 |
| chr18 | 35468468 | 35468525 | - | 16.0633692299738 | 29.5618892936739 | 64.4343233598714 | 32.0132594093141 |
| chr18 | 35468562 | 35468627 | - | 25.0990144218341 | 18.1919318730301 | 29.8307052591997 | 30.9461507623369 |
| chr18 | 35709798 | 35709813 | + | 3.01188173062009 | 1.13699574206438 | 15.5119667347839 | 1.06710864697714 |
| chr18 | 35709857 | 35709891 | + | 8.03168461498692 | 3.41098722619315 | 22.6713359969918 | 6.40265188186281 |
| chr18 | 35709956 | 35709983 | + | 6.02376346124019 | 6.82197445238629 | 8.35259747257593 | 7.46976052883995 |
| chr18 | 35710010 | 35710051 | + | 4.01584230749346 | 7.95897019445067 | 5.96614105183995 | 4.26843458790854 |
| chr18 | 35710084 | 35710104 | + | 5.01980288436682 | 5.68497871032191 | 15.5119667347839 | 9.60397782279422 |
| chr18 | 35710131 | 35710150 | + | 3.01188173062009 | 4.54798296825753 | 2.38645642073598 | 4.26843458790854 |
| chr18 | 35710375 | 35710382 | + | 5.01980288436682 | 6.82197445238629 | 7.15936926220794 | 5.33554323488568 |
| chr18 | 35710423 | 35710502 | + | 27.1069355755808 | 32.9728765198671 | 56.0817258872955 | 30.9461507623369 |
| chr18 | 35710525 | 35710552 | + | 19.0752509605939 | 26.1509020674808 | 23.8645642073598 | 13.8724124107028 |
| chr18 | 35710576 | 35710660 | + | 29.1148567293276 | 30.6988850357383 | 26.2510206280958 | 20.2750642925656 |
| chr18 | 35710718 | 35710774 | + | 3.01188173062009 | 4.54798296825753 | 17.8984231555198 | 8.53686917581709 |
| chr18 | 35710813 | 35710880 | + | 33.130699036821 | 32.9728765198671 | 105.004082512383 | 38.4159112911769 |
| chr18 | 35710901 | 35710935 | + | 5.01980288436682 | 9.09596593651506 | 9.54582568294392 | 2.13421729395427 |
| chr18 | 35711165 | 35711188 | + | 2.00792115374673 | 3.41098722619315 | 2.38645642073598 | 2.13421729395427 |
| chr18 | 35711441 | 35711456 | + | 4.01584230749346 | 4.54798296825753 | 3.57968463110397 | 4.26843458790854 |
| chr18 | 35972628 | 35972735 | + | 64.2534769198953 | 85.2746806548286 | 93.0718004087032 | 87.5029090521251 |
| chr18 | 35978988 | 35978991 | + | 1.00396057687336 | 2.27399148412876 | 2.38645642073598 | 3.20132594093141 |
| chr18 | 35991329 | 35991348 | - | 7.02772403811355 | 3.41098722619315 | 2.38645642073598 | 1.06710864697714 |
| chr18 | 35992103 | 35992115 | - | 21.0831721143407 | 37.5208594881246 | 14.3187385244159 | 13.8724124107028 |
| chr18 | 35992186 | 35992197 | - | 6.02376346124019 | 6.82197445238629 | 9.54582568294392 | 5.33554323488568 |
| chr18 | 35992367 | 35992399 | - | 15.0594086531005 | 23.876910583352 | 14.3187385244159 | 7.46976052883995 |
| chr18 | 35992468 | 35992498 | - | 6.02376346124019 | 6.82197445238629 | 1.19322821036799 | 3.20132594093141 |
| chr18 | 35992554 | 35992616 | - | 9.03564519186028 | 7.95897019445067 | 11.9322821036799 | 2.13421729395427 |
| chr18 | 35992749 | 35992772 | - | 14.0554480762271 | 15.9179403889013 | 8.35259747257593 | 7.46976052883995 |
| chr18 | 35992849 | 35992871 | - | 3.01188173062009 | 6.82197445238629 | 3.57968463110397 | 2.13421729395427 |
| chr18 | 35993031 | 35993064 | - | 1.00396057687336 | 1.13699574206438 | 3.57968463110397 | 2.13421729395427 |
| chr18 | 35993094 | 35993115 | - | 7.02772403811355 | 9.09596593651506 | 5.96614105183995 | 4.26843458790854 |
| chr18 | 36109085 | 36109087 | - | 31.1227778830743 | 53.438799877026 | 2.38645642073598 | 4.26843458790854 |
| chr18 | 36109309 | 36109325 | - | 92.3643730723495 | 79.5897019445067 | 5.96614105183995 | 5.33554323488568 |
| chr18 | 36109348 | 36109404 | - | 14.0554480762271 | 28.4248935516096 | 2.38645642073598 | 2.13421729395427 |
| chr18 | 36109433 | 36109617 | - | 241.954499026481 | 303.57786313119 | 22.6713359969918 | 20.2750642925656 |
| chr18 | 36174699 | 36174702 | + | 2.00792115374673 | 5.68497871032191 | 2.38645642073598 | 3.20132594093141 |
| chr18 | 36174974 | 36174985 | + | 4.01584230749346 | 1.13699574206438 | 1.19322821036799 | 2.13421729395427 |
| chr18 | 36271771 | 36271804 | + | 40.1584230749346 | 51.1648083928972 | 5.96614105183995 | 5.33554323488568 |
| chr18 | 36786821 | 36786861 | - | 26.1029749987075 | 23.876910583352 | 21.4781077866238 | 13.8724124107028 |
| chr18 | 36786895 | 36786916 | - | 7.02772403811355 | 4.54798296825753 | 1.19322821036799 | 3.20132594093141 |
| chr18 | 36796089 | 36796121 | - | 3.01188173062009 | 5.68497871032191 | 10.7390538933119 | 2.13421729395427 |
| chr18 | 36796143 | 36796144 | - | 1.00396057687336 | 1.13699574206438 | 1.19322821036799 | 1.06710864697714 |
| chr18 | 36796149 | 36796207 | - | 8.03168461498692 | 13.6439489047726 | 15.5119667347839 | 8.53686917581709 |
| chr18 | 36796493 | 36796497 | - | 4.01584230749346 | 3.41098722619315 | 3.57968463110397 | 5.33554323488568 |
| chr18 | 36796550 | 36796569 | - | 6.02376346124019 | 5.68497871032191 | 1.19322821036799 | 3.20132594093141 |
| chr18 | 36796591 | 36796594 | - | 1.00396057687336 | 4.54798296825753 | 1.19322821036799 | 2.13421729395427 |
| chr18 | 36796717 | 36796741 | - | 3.01188173062009 | 1.13699574206438 | 1.19322821036799 | 3.20132594093141 |
| chr18 | 36885998 | 36886009 | + | 3.01188173062009 | 2.27399148412876 | 1.19322821036799 | 1.06710864697714 |
| chr18 | 36886080 | 36886084 | + | 5.01980288436682 | 4.54798296825753 | 2.38645642073598 | 8.53686917581709 |
| chr18 | 36886228 | 36886262 | + | 4.01584230749346 | 3.41098722619315 | 15.5119667347839 | 11.7381951167485 |
| chr18 | 37244791 | 37244793 | - | 4.01584230749346 | 6.82197445238629 | 1.19322821036799 | 2.13421729395427 |
| chr18 | 42081127 | 42081165 | + | 2.00792115374673 | 5.68497871032191 | 1.19322821036799 | 1.06710864697714 |
| chr18 | 42081230 | 42081236 | + | 5.01980288436682 | 5.68497871032191 | 3.57968463110397 | 4.26843458790854 |
| chr18 | 42081330 | 42081332 | + | 6.02376346124019 | 6.82197445238629 | 4.77291284147196 | 5.33554323488568 |
| chr18 | 45847697 | 45847699 | - | 3.01188173062009 | 2.27399148412876 | 1.19322821036799 | 1.06710864697714 |
| chr18 | 46084167 | 46084193 | - | 11.043566345607 | 14.780944646837 | 8.35259747257593 | 5.33554323488568 |
| chr18 | 46084234 | 46084257 | - | 2.00792115374673 | 2.27399148412876 | 1.19322821036799 | 1.06710864697714 |
| chr18 | 46264837 | 46264889 | + | 6.02376346124019 | 5.68497871032191 | 3.57968463110397 | 6.40265188186281 |
| chr18 | 46265206 | 46265222 | + | 5.01980288436682 | 2.27399148412876 | 8.35259747257593 | 1.06710864697714 |
| chr18 | 46265243 | 46265278 | + | 5.01980288436682 | 3.41098722619315 | 5.96614105183995 | 3.20132594093141 |
| chr18 | 46266282 | 46266286 | + | 15.0594086531005 | 11.3699574206438 | 3.57968463110397 | 4.26843458790854 |
| chr18 | 46266455 | 46266462 | + | 10.0396057687336 | 15.9179403889013 | 4.77291284147196 | 3.20132594093141 |
| chr18 | 46266550 | 46266564 | + | 15.0594086531005 | 12.5069531627082 | 11.9322821036799 | 9.60397782279422 |
| chr18 | 46266597 | 46266634 | + | 21.0831721143407 | 28.4248935516096 | 17.8984231555198 | 8.53686917581709 |
| chr18 | 46266660 | 46266667 | + | 6.02376346124019 | 9.09596593651506 | 1.19322821036799 | 8.53686917581709 |
| chr18 | 46679020 | 46679021 | - | 6.02376346124019 | 7.95897019445067 | 20.2848795762558 | 12.8053037637256 |
| chr18 | 46810559 | 46810588 | - | 1.00396057687336 | 3.41098722619315 | 1.19322821036799 | 6.40265188186281 |
| chr18 | 46811485 | 46811509 | - | 1.00396057687336 | 1.13699574206438 | 11.9322821036799 | 1.06710864697714 |
| chr18 | 46811574 | 46811576 | - | 1.00396057687336 | 2.27399148412876 | 1.19322821036799 | 1.06710864697714 |
| chr18 | 46812409 | 46812412 | - | 5.01980288436682 | 2.27399148412876 | 1.19322821036799 | 1.06710864697714 |
| chr18 | 47107433 | 47107449 | - | 2.00792115374673 | 5.68497871032191 | 2.38645642073598 | 5.33554323488568 |
| chr18 | 47107547 | 47107594 | - | 5.01980288436682 | 9.09596593651506 | 19.0916513658878 | 4.26843458790854 |
| chr18 | 47107644 | 47107647 | - | 1.00396057687336 | 3.41098722619315 | 2.38645642073598 | 1.06710864697714 |
| chr18 | 47108395 | 47108417 | - | 8.03168461498692 | 5.68497871032191 | 10.7390538933119 | 14.9395210576799 |
| chr18 | 47108442 | 47108460 | - | 1.00396057687336 | 1.13699574206438 | 1.19322821036799 | 2.13421729395427 |
| chr18 | 47155070 | 47155165 | - | 18.0712903837206 | 32.9728765198671 | 8.35259747257593 | 14.9395210576799 |
| chr18 | 47155200 | 47155259 | - | 14.0554480762271 | 6.82197445238629 | 8.35259747257593 | 4.26843458790854 |
| chr18 | 47155287 | 47155318 | - | 12.0475269224804 | 13.6439489047726 | 16.7051949451519 | 9.60397782279422 |
| chr18 | 47155408 | 47155430 | - | 1.00396057687336 | 1.13699574206438 | 2.38645642073598 | 1.06710864697714 |
| chr18 | 47155583 | 47155603 | - | 5.01980288436682 | 18.1919318730301 | 4.77291284147196 | 3.20132594093141 |
| chr18 | 47155832 | 47155839 | - | 3.01188173062009 | 4.54798296825753 | 1.19322821036799 | 2.13421729395427 |
| chr18 | 47155880 | 47155956 | - | 2.00792115374673 | 10.2329616785794 | 5.96614105183995 | 2.13421729395427 |
| chr18 | 47156041 | 47156089 | - | 8.03168461498692 | 6.82197445238629 | 3.57968463110397 | 11.7381951167485 |
| chr18 | 47156116 | 47156187 | - | 5.01980288436682 | 2.27399148412876 | 1.19322821036799 | 6.40265188186281 |
| chr18 | 47833146 | 47833149 | - | 1.00396057687336 | 4.54798296825753 | 1.19322821036799 | 3.20132594093141 |
| chr18 | 47836943 | 47836951 | - | 2.00792115374673 | 1.13699574206438 | 2.38645642073598 | 16.006629704657 |
| chr18 | 47840506 | 47840531 | - | 21.0831721143407 | 61.3977700714766 | 3.57968463110397 | 5.33554323488568 |
| chr18 | 47841186 | 47841209 | - | 23.0910932680874 | 39.7948509722534 | 15.5119667347839 | 32.0132594093141 |
| chr18 | 47841242 | 47841257 | - | 12.0475269224804 | 18.1919318730301 | 3.57968463110397 | 3.20132594093141 |
| chr18 | 47841420 | 47841453 | - | 17.0673298068472 | 25.0139063254164 | 7.15936926220794 | 7.46976052883995 |
| chr18 | 47841517 | 47841537 | - | 4.01584230749346 | 5.68497871032191 | 1.19322821036799 | 1.06710864697714 |
| chr18 | 47841608 | 47841613 | - | 9.03564519186028 | 6.82197445238629 | 4.77291284147196 | 2.13421729395427 |
| chr18 | 47841638 | 47841654 | - | 3.01188173062009 | 17.0549361309657 | 3.57968463110397 | 1.06710864697714 |
| chr18 | 49043512 | 49043513 | - | 1.00396057687336 | 2.27399148412876 | 1.19322821036799 | 1.06710864697714 |
| chr18 | 49043841 | 49043901 | - | 30.1188173062009 | 31.8358807778027 | 35.7968463110397 | 20.2750642925656 |
| chr18 | 49044014 | 49044055 | - | 9.03564519186028 | 7.95897019445067 | 1.19322821036799 | 3.20132594093141 |
| chr18 | 49481750 | 49481782 | - | 24.0950538449607 | 35.2468680039958 | 15.5119667347839 | 11.7381951167485 |
| chr18 | 49783672 | 49783687 | - | 10.0396057687336 | 9.09596593651506 | 2.38645642073598 | 2.13421729395427 |
| chr18 | 49795836 | 49795873 | - | 3.01188173062009 | 1.13699574206438 | 5.96614105183995 | 5.33554323488568 |
| chr18 | 49802690 | 49802744 | - | 7.02772403811355 | 4.54798296825753 | 1.19322821036799 | 4.26843458790854 |
| chr18 | 49802774 | 49802856 | - | 42.1663442286813 | 25.0139063254164 | 4.77291284147196 | 6.40265188186281 |
| chr18 | 49813961 | 49814132 | + | 814.212027844299 | 820.910925770484 | 78.7530618842873 | 151.529427870753 |
| chr18 | 49814201 | 49814260 | + | 28.1108961524542 | 20.4659233571589 | 1.19322821036799 | 2.13421729395427 |
| chr18 | 50266922 | 50266926 | - | 2.00792115374673 | 2.27399148412876 | 1.19322821036799 | 1.06710864697714 |
| chr18 | 50282405 | 50282416 | - | 4.01584230749346 | 4.54798296825753 | 5.96614105183995 | 6.40265188186281 |
| chr18 | 50947291 | 50947302 | + | 4.01584230749346 | 4.54798296825753 | 3.57968463110397 | 1.06710864697714 |
| chr18 | 50947337 | 50947347 | + | 6.02376346124019 | 13.6439489047726 | 16.7051949451519 | 27.7448248214055 |
| chr18 | 50947417 | 50947449 | + | 11.043566345607 | 9.09596593651506 | 7.15936926220794 | 8.53686917581709 |
| chr18 | 50947495 | 50947535 | + | 75.2970432655023 | 90.9596593651506 | 45.3426719939836 | 52.2883237018796 |
| chr18 | 50947584 | 50947602 | + | 7.02772403811355 | 15.9179403889013 | 7.15936926220794 | 2.13421729395427 |
| chr18 | 50947627 | 50947629 | + | 2.00792115374673 | 4.54798296825753 | 9.54582568294392 | 4.26843458790854 |
| chr18 | 50947666 | 50947678 | + | 3.01188173062009 | 3.41098722619315 | 4.77291284147196 | 5.33554323488568 |
| chr18 | 50947702 | 50947720 | + | 35.1386201905678 | 39.7948509722534 | 64.4343233598714 | 26.6777161744284 |
| chr18 | 50987242 | 50987245 | + | 2.00792115374673 | 21.6029190992233 | 3.57968463110397 | 4.26843458790854 |
| chr18 | 51079376 | 51079380 | + | 4.01584230749346 | 4.54798296825753 | 3.57968463110397 | 1.06710864697714 |
| chr18 | 51079585 | 51079588 | + | 4.01584230749346 | 3.41098722619315 | 4.77291284147196 | 1.06710864697714 |
| chr18 | 51079744 | 51079751 | + | 2.00792115374673 | 7.95897019445067 | 5.96614105183995 | 1.06710864697714 |
| chr18 | 51082659 | 51082718 | + | 6.02376346124019 | 11.3699574206438 | 4.77291284147196 | 4.26843458790854 |
| chr18 | 51083318 | 51083374 | + | 17.0673298068472 | 13.6439489047726 | 1.19322821036799 | 2.13421729395427 |
| chr18 | 51083402 | 51083436 | + | 13.0514874993537 | 13.6439489047726 | 7.15936926220794 | 10.6710864697714 |
| chr18 | 51084145 | 51084146 | + | 3.01188173062009 | 2.27399148412876 | 2.38645642073598 | 6.40265188186281 |
| chr18 | 51084172 | 51084207 | + | 20.0792115374673 | 28.4248935516096 | 7.15936926220794 | 3.20132594093141 |
| chr18 | 51084501 | 51084522 | + | 45.1782259593014 | 39.7948509722534 | 9.54582568294392 | 13.8724124107028 |
| chr18 | 51174625 | 51174681 | - | 3.01188173062009 | 1.13699574206438 | 1.19322821036799 | 2.13421729395427 |
| chr18 | 51174831 | 51174857 | - | 2.00792115374673 | 12.5069531627082 | 3.57968463110397 | 3.20132594093141 |
| chr18 | 51175417 | 51175432 | - | 2.00792115374673 | 2.27399148412876 | 2.38645642073598 | 1.06710864697714 |
| chr18 | 54154895 | 54154899 | - | 20.0792115374673 | 31.8358807778027 | 5.96614105183995 | 9.60397782279422 |
| chr18 | 54155031 | 54155080 | - | 5.01980288436682 | 5.68497871032191 | 1.19322821036799 | 1.06710864697714 |
| chr18 | 56602843 | 56602863 | - | 2.00792115374673 | 5.68497871032191 | 1.19322821036799 | 3.20132594093141 |
| chr18 | 56602976 | 56603012 | - | 1.00396057687336 | 4.54798296825753 | 1.19322821036799 | 6.40265188186281 |
| chr18 | 57478296 | 57478363 | + | 18.0712903837206 | 15.9179403889013 | 1.19322821036799 | 4.26843458790854 |
| chr18 | 57491063 | 57491146 | + | 52.205949997415 | 51.1648083928972 | 1.19322821036799 | 4.26843458790854 |
| chr18 | 57548397 | 57548432 | - | 1.00396057687336 | 5.68497871032191 | 3.57968463110397 | 2.13421729395427 |
| chr18 | 57548611 | 57548652 | - | 9.03564519186028 | 14.780944646837 | 10.7390538933119 | 16.006629704657 |
| chr18 | 57549058 | 57549059 | - | 2.00792115374673 | 4.54798296825753 | 5.96614105183995 | 6.40265188186281 |
| chr18 | 57549892 | 57549912 | - | 12.0475269224804 | 6.82197445238629 | 5.96614105183995 | 11.7381951167485 |
| chr18 | 57549978 | 57549995 | - | 4.01584230749346 | 3.41098722619315 | 3.57968463110397 | 9.60397782279422 |
| chr18 | 57550079 | 57550081 | - | 2.00792115374673 | 1.13699574206438 | 1.19322821036799 | 2.13421729395427 |
| chr18 | 57550150 | 57550154 | - | 6.02376346124019 | 1.13699574206438 | 8.35259747257593 | 2.13421729395427 |
| chr18 | 57550312 | 57550366 | - | 10.0396057687336 | 3.41098722619315 | 20.2848795762558 | 35.2145853502455 |
| chr18 | 57550574 | 57550611 | - | 2.00792115374673 | 3.41098722619315 | 10.7390538933119 | 7.46976052883995 |
| chr18 | 57550645 | 57550651 | - | 4.01584230749346 | 3.41098722619315 | 7.15936926220794 | 7.46976052883995 |
| chr18 | 57550688 | 57550704 | - | 10.0396057687336 | 2.27399148412876 | 5.96614105183995 | 9.60397782279422 |
| chr18 | 57600673 | 57600717 | - | 9.03564519186028 | 22.7399148412876 | 11.9322821036799 | 16.006629704657 |
| chr18 | 57600736 | 57600837 | - | 14.0554480762271 | 40.9318467143178 | 35.7968463110397 | 43.7514545260626 |
| chr18 | 57600858 | 57600862 | - | 3.01188173062009 | 1.13699574206438 | 5.96614105183995 | 3.20132594093141 |
| chr18 | 57600909 | 57600937 | - | 11.043566345607 | 27.2878978095452 | 22.6713359969918 | 22.4092815865198 |
| chr18 | 57600964 | 57601082 | - | 20.0792115374673 | 35.2468680039958 | 38.1833027317757 | 30.9461507623369 |
| chr18 | 57601182 | 57601207 | - | 309.219857676996 | 569.634866774255 | 143.187385244159 | 233.696793687993 |
| chr18 | 57601286 | 57601306 | - | 20.0792115374673 | 19.3289276150945 | 5.96614105183995 | 3.20132594093141 |
| chr18 | 57601330 | 57601495 | - | 29.1148567293276 | 43.2058381984465 | 21.4781077866238 | 16.006629704657 |
| chr18 | 57601551 | 57601585 | - | 13.0514874993537 | 9.09596593651506 | 7.15936926220794 | 5.33554323488568 |
| chr18 | 57601607 | 57601654 | - | 21.0831721143407 | 14.780944646837 | 13.1255103140479 | 14.9395210576799 |
| chr18 | 57630329 | 57630394 | + | 20.0792115374673 | 9.09596593651506 | 2.38645642073598 | 12.8053037637256 |
| chr18 | 58045407 | 58045440 | + | 2.00792115374673 | 11.3699574206438 | 4.77291284147196 | 19.2079556455884 |
| chr18 | 58396194 | 58396393 | + | 3.01188173062009 | 4.54798296825753 | 1.19322821036799 | 1.06710864697714 |
| chr18 | 58396582 | 58396621 | + | 1.00396057687336 | 7.95897019445067 | 2.38645642073598 | 1.06710864697714 |
| chr18 | 58396643 | 58396749 | + | 2.00792115374673 | 10.2329616785794 | 3.57968463110397 | 1.06710864697714 |
| chr18 | 58397083 | 58397107 | + | 7.02772403811355 | 4.54798296825753 | 1.19322821036799 | 2.13421729395427 |
| chr18 | 58397129 | 58397143 | + | 2.00792115374673 | 2.27399148412876 | 1.19322821036799 | 2.13421729395427 |
| chr18 | 58397311 | 58397341 | + | 20.0792115374673 | 32.9728765198671 | 1.19322821036799 | 4.26843458790854 |
| chr18 | 58397965 | 58398147 | + | 7.02772403811355 | 7.95897019445067 | 5.96614105183995 | 1.06710864697714 |
| chr18 | 58750410 | 58750411 | + | 16.0633692299738 | 12.5069531627082 | 7.15936926220794 | 6.40265188186281 |
| chr18 | 59158639 | 59158705 | + | 13.0514874993537 | 25.0139063254164 | 3.57968463110397 | 9.60397782279422 |
| chr18 | 59158772 | 59158810 | + | 26.1029749987075 | 45.4798296825753 | 11.9322821036799 | 18.1408469986113 |
| chr18 | 59327844 | 59327890 | - | 5.01980288436682 | 11.3699574206438 | 19.0916513658878 | 11.7381951167485 |
| chr18 | 59327920 | 59328026 | - | 17.0673298068472 | 23.876910583352 | 19.0916513658878 | 6.40265188186281 |
| chr18 | 59328054 | 59328149 | - | 15.0594086531005 | 15.9179403889013 | 14.3187385244159 | 19.2079556455884 |
| chr18 | 59328192 | 59328314 | - | 28.1108961524542 | 18.1919318730301 | 25.0577924177278 | 29.8790421153598 |
| chr18 | 59328378 | 59328446 | - | 3.01188173062009 | 1.13699574206438 | 5.96614105183995 | 3.20132594093141 |
| chr18 | 59328535 | 59328630 | - | 12.0475269224804 | 13.6439489047726 | 20.2848795762558 | 12.8053037637256 |
| chr18 | 59328655 | 59328673 | - | 7.02772403811355 | 2.27399148412876 | 2.38645642073598 | 3.20132594093141 |
| chr18 | 59328701 | 59328771 | - | 1.00396057687336 | 2.27399148412876 | 5.96614105183995 | 3.20132594093141 |
| chr18 | 59329121 | 59329144 | - | 1.00396057687336 | 2.27399148412876 | 9.54582568294392 | 1.06710864697714 |
| chr18 | 59329832 | 59329992 | - | 4.01584230749346 | 2.27399148412876 | 4.77291284147196 | 3.20132594093141 |
| chr18 | 59697437 | 59697438 | + | 12.0475269224804 | 2.27399148412876 | 1.19322821036799 | 16.006629704657 |
| chr18 | 59903125 | 59903148 | + | 7.02772403811355 | 19.3289276150945 | 4.77291284147196 | 4.26843458790854 |
| chr18 | 59903961 | 59903963 | + | 1.00396057687336 | 6.82197445238629 | 1.19322821036799 | 3.20132594093141 |
| chr18 | 59904097 | 59904117 | + | 8.03168461498692 | 22.7399148412876 | 4.77291284147196 | 6.40265188186281 |
| chr18 | 59904198 | 59904200 | + | 4.01584230749346 | 13.6439489047726 | 4.77291284147196 | 3.20132594093141 |
| chr18 | 62044646 | 62044649 | - | 4.01584230749346 | 3.41098722619315 | 1.19322821036799 | 3.20132594093141 |
| chr18 | 62045383 | 62045405 | - | 1.00396057687336 | 6.82197445238629 | 3.57968463110397 | 4.26843458790854 |
| chr18 | 62305571 | 62305603 | + | 25.0990144218341 | 25.0139063254164 | 16.7051949451519 | 12.8053037637256 |
| chr18 | 62577241 | 62577259 | + | 1.00396057687336 | 2.27399148412876 | 1.19322821036799 | 2.13421729395427 |
| chr18 | 62577916 | 62577962 | + | 18.0712903837206 | 18.1919318730301 | 14.3187385244159 | 10.6710864697714 |
| chr18 | 62578157 | 62578213 | + | 34.1346596136944 | 32.9728765198671 | 16.7051949451519 | 25.6106075274513 |
| chr18 | 62979783 | 62979801 | + | 1.00396057687336 | 3.41098722619315 | 3.57968463110397 | 1.06710864697714 |
| chr18 | 62979836 | 62979840 | + | 21.0831721143407 | 32.9728765198671 | 22.6713359969918 | 23.476390233497 |
| chr18 | 62980160 | 62980243 | + | 18.0712903837206 | 10.2329616785794 | 9.54582568294392 | 16.006629704657 |
| chr18 | 63330870 | 63330926 | - | 66.2613980736421 | 86.411676396893 | 47.7291284147196 | 38.4159112911769 |
| chr18 | 63330947 | 63331129 | - | 998.940773988998 | 1383.72381809235 | 381.833027317757 | 362.816939972226 |
| chr18 | 63331150 | 63331239 | - | 92.3643730723495 | 86.411676396893 | 14.3187385244159 | 27.7448248214055 |
| chr18 | 63331391 | 63331412 | - | 16.0633692299738 | 22.7399148412876 | 3.57968463110397 | 2.13421729395427 |
| chr18 | 63331483 | 63331551 | - | 25.0990144218341 | 37.5208594881246 | 3.57968463110397 | 7.46976052883995 |
| chr18 | 63331628 | 63331666 | - | 26.1029749987075 | 29.5618892936739 | 5.96614105183995 | 8.53686917581709 |
| chr18 | 63389338 | 63389353 | - | 4.01584230749346 | 3.41098722619315 | 1.19322821036799 | 3.20132594093141 |
| chr18 | 63389540 | 63389556 | - | 1.00396057687336 | 4.54798296825753 | 2.38645642073598 | 1.06710864697714 |
| chr18 | 63390652 | 63390672 | - | 25.0990144218341 | 12.5069531627082 | 4.77291284147196 | 2.13421729395427 |
| chr18 | 63390740 | 63390795 | - | 8.03168461498692 | 14.780944646837 | 2.38645642073598 | 1.06710864697714 |
| chr18 | 68676184 | 68676210 | - | 10.0396057687336 | 7.95897019445067 | 1.19322821036799 | 3.20132594093141 |
| chr18 | 68676596 | 68676598 | - | 1.00396057687336 | 2.27399148412876 | 1.19322821036799 | 1.06710864697714 |
| chr18 | 68676685 | 68676705 | - | 6.02376346124019 | 3.41098722619315 | 1.19322821036799 | 1.06710864697714 |
| chr18 | 68676752 | 68676780 | - | 8.03168461498692 | 22.7399148412876 | 16.7051949451519 | 8.53686917581709 |
| chr18 | 74158487 | 74158516 | + | 11.043566345607 | 6.82197445238629 | 8.35259747257593 | 16.006629704657 |
| chr18 | 74158567 | 74158583 | + | 5.01980288436682 | 1.13699574206438 | 3.57968463110397 | 1.06710864697714 |
| chr18 | 74520937 | 74520980 | + | 3.01188173062009 | 1.13699574206438 | 2.38645642073598 | 4.26843458790854 |
| chr18 | 74521054 | 74521105 | + | 5.01980288436682 | 9.09596593651506 | 2.38645642073598 | 5.33554323488568 |
| chr18 | 75198028 | 75198035 | - | 6.02376346124019 | 3.41098722619315 | 1.19322821036799 | 1.06710864697714 |
| chr18 | 75198700 | 75198761 | - | 10.0396057687336 | 9.09596593651506 | 15.5119667347839 | 5.33554323488568 |
| chr18 | 75198793 | 75198801 | - | 27.1069355755808 | 19.3289276150945 | 15.5119667347839 | 13.8724124107028 |
| chr18 | 75198895 | 75198905 | - | 7.02772403811355 | 15.9179403889013 | 3.57968463110397 | 4.26843458790854 |
| chr18 | 75199346 | 75199349 | - | 1.00396057687336 | 1.13699574206438 | 1.19322821036799 | 3.20132594093141 |
| chr18 | 75199541 | 75199544 | - | 20.0792115374673 | 32.9728765198671 | 13.1255103140479 | 22.4092815865198 |
| chr18 | 75200144 | 75200179 | - | 6.02376346124019 | 2.27399148412876 | 9.54582568294392 | 2.13421729395427 |
| chr18 | 75200237 | 75200238 | - | 1.00396057687336 | 1.13699574206438 | 1.19322821036799 | 1.06710864697714 |
| chr18 | 75200655 | 75200677 | - | 2.00792115374673 | 4.54798296825753 | 1.19322821036799 | 4.26843458790854 |
| chr18 | 75201196 | 75201199 | - | 4.01584230749346 | 15.9179403889013 | 3.57968463110397 | 1.06710864697714 |
| chr18 | 79753906 | 79753923 | + | 1.00396057687336 | 2.27399148412876 | 3.57968463110397 | 5.33554323488568 |
| chr18 | 79754076 | 79754090 | + | 18.0712903837206 | 7.95897019445067 | 4.77291284147196 | 8.53686917581709 |
| chr18 | 79754381 | 79754403 | + | 3.01188173062009 | 7.95897019445067 | 2.38645642073598 | 2.13421729395427 |
| chr18 | 79902514 | 79902542 | - | 18.0712903837206 | 44.3428339405109 | 11.9322821036799 | 5.33554323488568 |
| chr18 | 79902563 | 79902685 | - | 172.681219222219 | 236.495114349391 | 78.7530618842873 | 118.449059814462 |
| chr18 | 79902711 | 79902722 | - | 4.01584230749346 | 2.27399148412876 | 4.77291284147196 | 2.13421729395427 |
| chr18 | 79903102 | 79903145 | - | 9.03564519186028 | 7.95897019445067 | 1.19322821036799 | 4.26843458790854 |
| chr18 | 79903718 | 79903820 | - | 19.0752509605939 | 19.3289276150945 | 13.1255103140479 | 8.53686917581709 |
| chr18 | 79973488 | 79973589 | - | 43.1703048055547 | 43.2058381984465 | 33.4103898903037 | 46.952780466994 |
| chr18 | 79973636 | 79973693 | - | 4.01584230749346 | 6.82197445238629 | 2.38645642073598 | 2.13421729395427 |
| chr18 | 80045959 | 80046057 | + | 7.02772403811355 | 4.54798296825753 | 2.38645642073598 | 2.13421729395427 |
| chr18 | 80046132 | 80046256 | + | 20.0792115374673 | 12.5069531627082 | 5.96614105183995 | 18.1408469986113 |
| chr18 | 80046363 | 80046374 | + | 4.01584230749346 | 5.68497871032191 | 5.96614105183995 | 2.13421729395427 |
| chr18 | 80139916 | 80139928 | + | 2.00792115374673 | 1.13699574206438 | 1.19322821036799 | 1.06710864697714 |
| chr19 | 541586 | 541730 | + | 29.1148567293276 | 22.7399148412876 | 29.8307052591997 | 81.1002571702623 |
| chr19 | 541756 | 541779 | + | 13.0514874993537 | 4.54798296825753 | 3.57968463110397 | 6.40265188186281 |
| chr19 | 541808 | 541854 | + | 11.043566345607 | 9.09596593651506 | 20.2848795762558 | 27.7448248214055 |
| chr19 | 541938 | 542067 | + | 17.0673298068472 | 29.5618892936739 | 16.7051949451519 | 26.6777161744284 |
| chr19 | 582512 | 582576 | + | 36.1425807674411 | 31.8358807778027 | 19.0916513658878 | 18.1408469986113 |
| chr19 | 582747 | 582944 | + | 202.80003652842 | 186.467301698559 | 326.944529640829 | 348.944527561523 |
| chr19 | 582980 | 583279 | + | 38.1505019211879 | 26.1509020674808 | 19.0916513658878 | 22.4092815865198 |
| chr19 | 583306 | 583473 | + | 29.1148567293276 | 21.6029190992233 | 22.6713359969918 | 21.3421729395427 |
| chr19 | 647553 | 647581 | - | 10.0396057687336 | 13.6439489047726 | 5.96614105183995 | 3.20132594093141 |
| chr19 | 647601 | 647656 | - | 26.1029749987075 | 27.2878978095452 | 13.1255103140479 | 29.8790421153598 |
| chr19 | 647868 | 647873 | - | 16.0633692299738 | 22.7399148412876 | 15.5119667347839 | 22.4092815865198 |
| chr19 | 647922 | 647947 | - | 13.0514874993537 | 2.27399148412876 | 3.57968463110397 | 11.7381951167485 |
| chr19 | 682837 | 682850 | + | 12.0475269224804 | 9.09596593651506 | 1.19322821036799 | 3.20132594093141 |
| chr19 | 810856 | 810869 | + | 2.00792115374673 | 1.13699574206438 | 3.57968463110397 | 3.20132594093141 |
| chr19 | 811384 | 811424 | + | 5.01980288436682 | 4.54798296825753 | 3.57968463110397 | 9.60397782279422 |
| chr19 | 811473 | 811484 | + | 1.00396057687336 | 2.27399148412876 | 7.15936926220794 | 26.6777161744284 |
| chr19 | 811544 | 811600 | + | 4.01584230749346 | 1.13699574206438 | 4.77291284147196 | 13.8724124107028 |
| chr19 | 811739 | 811745 | + | 5.01980288436682 | 3.41098722619315 | 1.19322821036799 | 6.40265188186281 |
| chr19 | 811966 | 812012 | + | 33.130699036821 | 32.9728765198671 | 17.8984231555198 | 42.6843458790854 |
| chr19 | 812055 | 812087 | + | 9.03564519186028 | 6.82197445238629 | 1.19322821036799 | 9.60397782279422 |
| chr19 | 812104 | 812200 | + | 95.3762548029696 | 105.740604011988 | 57.2749540976635 | 120.583277108416 |
| chr19 | 812244 | 812294 | + | 29.1148567293276 | 38.657855230189 | 20.2848795762558 | 28.8119334683827 |
| chr19 | 896530 | 896563 | - | 8.03168461498692 | 15.9179403889013 | 38.1833027317757 | 58.6909755837425 |
| chr19 | 896789 | 896799 | - | 3.01188173062009 | 2.27399148412876 | 4.77291284147196 | 6.40265188186281 |
| chr19 | 897094 | 897131 | - | 7.02772403811355 | 5.68497871032191 | 26.2510206280958 | 14.9395210576799 |
| chr19 | 897165 | 897199 | - | 1.00396057687336 | 3.41098722619315 | 2.38645642073598 | 7.46976052883995 |
| chr19 | 972112 | 972202 | + | 1.00396057687336 | 3.41098722619315 | 44.1494437836156 | 40.5501285851312 |
| chr19 | 972455 | 972517 | + | 1.00396057687336 | 1.13699574206438 | 10.7390538933119 | 3.20132594093141 |
| chr19 | 974694 | 974809 | + | 1.00396057687336 | 2.27399148412876 | 23.8645642073598 | 24.5434988804741 |
| chr19 | 975351 | 975446 | + | 1.00396057687336 | 2.27399148412876 | 32.2171616799357 | 40.5501285851312 |
| chr19 | 975654 | 975693 | + | 2.00792115374673 | 1.13699574206438 | 15.5119667347839 | 20.2750642925656 |
| chr19 | 975888 | 975914 | + | 1.00396057687336 | 1.13699574206438 | 8.35259747257593 | 12.8053037637256 |
| chr19 | 994410 | 994450 | + | 1.00396057687336 | 2.27399148412876 | 1.19322821036799 | 1.06710864697714 |
| chr19 | 994479 | 994550 | + | 10.0396057687336 | 23.876910583352 | 22.6713359969918 | 26.6777161744284 |
| chr19 | 1009683 | 1009802 | - | 54.2138711511617 | 61.3977700714766 | 33.4103898903037 | 32.0132594093141 |
| chr19 | 1010079 | 1010095 | - | 3.01188173062009 | 2.27399148412876 | 1.19322821036799 | 2.13421729395427 |
| chr19 | 1010106 | 1010155 | - | 37.1465413443145 | 36.3838637460602 | 14.3187385244159 | 11.7381951167485 |
| chr19 | 1010199 | 1010243 | - | 10.0396057687336 | 11.3699574206438 | 5.96614105183995 | 9.60397782279422 |
| chr19 | 1010356 | 1010396 | - | 14.0554480762271 | 12.5069531627082 | 5.96614105183995 | 20.2750642925656 |
| chr19 | 1010413 | 1010419 | - | 3.01188173062009 | 2.27399148412876 | 5.96614105183995 | 2.13421729395427 |
| chr19 | 1010564 | 1010623 | - | 11.043566345607 | 10.2329616785794 | 5.96614105183995 | 1.06710864697714 |
| chr19 | 1037879 | 1037937 | + | 12.0475269224804 | 1.13699574206438 | 53.6952694665595 | 67.2278447595595 |
| chr19 | 1037995 | 1038049 | + | 5.01980288436682 | 3.41098722619315 | 32.2171616799357 | 45.8856718200168 |
| chr19 | 1038095 | 1038122 | + | 4.01584230749346 | 4.54798296825753 | 7.15936926220794 | 9.60397782279422 |
| chr19 | 1038324 | 1038346 | + | 1.00396057687336 | 1.13699574206438 | 2.38645642073598 | 10.6710864697714 |
| chr19 | 1038915 | 1038949 | + | 2.00792115374673 | 2.27399148412876 | 1.19322821036799 | 3.20132594093141 |
| chr19 | 1038977 | 1039039 | + | 12.0475269224804 | 10.2329616785794 | 14.3187385244159 | 36.2816939972226 |
| chr19 | 1065507 | 1065553 | + | 8.03168461498692 | 9.09596593651506 | 4.77291284147196 | 7.46976052883995 |
| chr19 | 1086715 | 1086740 | - | 6.02376346124019 | 6.82197445238629 | 2.38645642073598 | 3.20132594093141 |
| chr19 | 1087509 | 1087545 | - | 14.0554480762271 | 15.9179403889013 | 3.57968463110397 | 1.06710864697714 |
| chr19 | 1088186 | 1088287 | - | 200.792115374673 | 198.974254861267 | 165.858721241151 | 326.535245975004 |
| chr19 | 1088309 | 1088343 | - | 3.01188173062009 | 4.54798296825753 | 1.19322821036799 | 5.33554323488568 |
| chr19 | 1088678 | 1088710 | - | 18.0712903837206 | 4.54798296825753 | 21.4781077866238 | 53.3554323488568 |
| chr19 | 1106537 | 1106637 | + | 33.130699036821 | 18.1919318730301 | 40.5697591525116 | 49.0869977609482 |
| chr19 | 1106658 | 1106769 | + | 340.342635560071 | 407.044475659049 | 221.940447128446 | 306.260181682438 |
| chr19 | 1227959 | 1227968 | + | 5.01980288436682 | 1.13699574206438 | 2.38645642073598 | 1.06710864697714 |
| chr19 | 1244744 | 1244804 | + | 20.0792115374673 | 7.95897019445067 | 1.19322821036799 | 13.8724124107028 |
| chr19 | 1257365 | 1257402 | + | 13.0514874993537 | 13.6439489047726 | 3.57968463110397 | 55.4896496428111 |
| chr19 | 1257831 | 1257886 | + | 7.02772403811355 | 4.54798296825753 | 14.3187385244159 | 21.3421729395427 |
| chr19 | 1257932 | 1257998 | + | 6.02376346124019 | 6.82197445238629 | 20.2848795762558 | 32.0132594093141 |
| chr19 | 1258070 | 1258221 | + | 28.1108961524542 | 18.1919318730301 | 54.8884976769275 | 81.1002571702623 |
| chr19 | 1258297 | 1258358 | + | 9.03564519186028 | 10.2329616785794 | 26.2510206280958 | 32.0132594093141 |
| chr19 | 1258449 | 1258493 | + | 1.00396057687336 | 3.41098722619315 | 2.38645642073598 | 18.1408469986113 |
| chr19 | 1258726 | 1258765 | + | 182.720824990952 | 139.850476273919 | 50.1155848354556 | 121.650385755393 |
| chr19 | 1258819 | 1258865 | + | 5.01980288436682 | 2.27399148412876 | 3.57968463110397 | 9.60397782279422 |
| chr19 | 1259019 | 1259066 | + | 40.1584230749346 | 20.4659233571589 | 13.1255103140479 | 44.8185631730397 |
| chr19 | 1272438 | 1272519 | + | 4.01584230749346 | 6.82197445238629 | 3.57968463110397 | 2.13421729395427 |
| chr19 | 1272586 | 1272616 | + | 5.01980288436682 | 5.68497871032191 | 3.57968463110397 | 6.40265188186281 |
| chr19 | 1272728 | 1272737 | + | 1.00396057687336 | 7.95897019445067 | 4.77291284147196 | 5.33554323488568 |
| chr19 | 1272919 | 1272960 | + | 4.01584230749346 | 1.13699574206438 | 2.38645642073598 | 1.06710864697714 |
| chr19 | 1272985 | 1273038 | + | 14.0554480762271 | 25.0139063254164 | 15.5119667347839 | 23.476390233497 |
| chr19 | 1273064 | 1273149 | + | 32.1267384599477 | 53.438799877026 | 41.7629873628796 | 78.966039876308 |
| chr19 | 1278793 | 1278825 | + | 13.0514874993537 | 2.27399148412876 | 9.54582568294392 | 13.8724124107028 |
| chr19 | 1279150 | 1279205 | + | 39.1544624980612 | 47.753821166704 | 16.7051949451519 | 20.2750642925656 |
| chr19 | 1376664 | 1376681 | + | 3.01188173062009 | 2.27399148412876 | 2.38645642073598 | 2.13421729395427 |
| chr19 | 1378203 | 1378206 | + | 1.00396057687336 | 1.13699574206438 | 3.57968463110397 | 3.20132594093141 |
| chr19 | 1378229 | 1378327 | + | 39.1544624980612 | 38.657855230189 | 42.9562155732476 | 43.7514545260626 |
| chr19 | 1395507 | 1395557 | + | 5.01980288436682 | 9.09596593651506 | 2.38645642073598 | 9.60397782279422 |
| chr19 | 1397108 | 1397162 | - | 5.01980288436682 | 7.95897019445067 | 3.57968463110397 | 4.26843458790854 |
| chr19 | 1435016 | 1435076 | + | 231.914893257747 | 134.165497563597 | 76.3666054635513 | 231.562576394038 |
| chr19 | 1435122 | 1435159 | + | 3.01188173062009 | 3.41098722619315 | 8.35259747257593 | 13.8724124107028 |
| chr19 | 1435368 | 1435376 | + | 3.01188173062009 | 2.27399148412876 | 9.54582568294392 | 7.46976052883995 |
| chr19 | 1435535 | 1435664 | + | 22.087132691214 | 28.4248935516096 | 27.4442488384638 | 39.483019938154 |
| chr19 | 1441873 | 1441894 | + | 4.01584230749346 | 1.13699574206438 | 1.19322821036799 | 1.06710864697714 |
| chr19 | 1473231 | 1473257 | - | 4.01584230749346 | 6.82197445238629 | 2.38645642073598 | 2.13421729395427 |
| chr19 | 1474200 | 1474223 | - | 7.02772403811355 | 9.09596593651506 | 4.77291284147196 | 12.8053037637256 |
| chr19 | 1474276 | 1474290 | - | 4.01584230749346 | 2.27399148412876 | 2.38645642073598 | 2.13421729395427 |
| chr19 | 1474890 | 1474891 | - | 1.00396057687336 | 2.27399148412876 | 2.38645642073598 | 1.06710864697714 |
| chr19 | 1497873 | 1497903 | + | 5.01980288436682 | 1.13699574206438 | 29.8307052591997 | 28.8119334683827 |
| chr19 | 1554958 | 1554962 | - | 1.00396057687336 | 2.27399148412876 | 1.19322821036799 | 1.06710864697714 |
| chr19 | 1555014 | 1555024 | - | 24.0950538449607 | 18.1919318730301 | 7.15936926220794 | 7.46976052883995 |
| chr19 | 1555108 | 1555113 | - | 1.00396057687336 | 1.13699574206438 | 1.19322821036799 | 1.06710864697714 |
| chr19 | 1555328 | 1555329 | - | 6.02376346124019 | 5.68497871032191 | 3.57968463110397 | 1.06710864697714 |
| chr19 | 1555426 | 1555460 | - | 9.03564519186028 | 13.6439489047726 | 5.96614105183995 | 3.20132594093141 |
| chr19 | 1576690 | 1576735 | - | 16.0633692299738 | 7.95897019445067 | 7.15936926220794 | 20.2750642925656 |
| chr19 | 1576985 | 1577038 | - | 16.0633692299738 | 6.82197445238629 | 8.35259747257593 | 28.8119334683827 |
| chr19 | 1577071 | 1577105 | - | 8.03168461498692 | 4.54798296825753 | 15.5119667347839 | 17.0737383516342 |
| chr19 | 1577132 | 1577168 | - | 8.03168461498692 | 5.68497871032191 | 15.5119667347839 | 19.2079556455884 |
| chr19 | 1577240 | 1577320 | - | 23.0910932680874 | 7.95897019445067 | 38.1833027317757 | 57.6238669367653 |
| chr19 | 1577366 | 1577395 | - | 8.03168461498692 | 6.82197445238629 | 17.8984231555198 | 40.5501285851312 |
| chr19 | 1577429 | 1577444 | - | 5.01980288436682 | 1.13699574206438 | 1.19322821036799 | 1.06710864697714 |
| chr19 | 1577467 | 1577492 | - | 4.01584230749346 | 7.95897019445067 | 4.77291284147196 | 7.46976052883995 |
| chr19 | 1577521 | 1577532 | - | 1.00396057687336 | 2.27399148412876 | 3.57968463110397 | 4.26843458790854 |
| chr19 | 1577568 | 1577603 | - | 5.01980288436682 | 1.13699574206438 | 4.77291284147196 | 11.7381951167485 |
| chr19 | 1577634 | 1577646 | - | 3.01188173062009 | 1.13699574206438 | 1.19322821036799 | 4.26843458790854 |
| chr19 | 1577723 | 1577812 | - | 32.1267384599477 | 27.2878978095452 | 53.6952694665595 | 21.3421729395427 |
| chr19 | 1577870 | 1577941 | - | 21.0831721143407 | 14.780944646837 | 27.4442488384638 | 21.3421729395427 |
| chr19 | 1577966 | 1578001 | - | 10.0396057687336 | 9.09596593651506 | 11.9322821036799 | 8.53686917581709 |
| chr19 | 1578041 | 1578054 | - | 2.00792115374673 | 5.68497871032191 | 1.19322821036799 | 11.7381951167485 |
| chr19 | 1578355 | 1578463 | - | 42.1663442286813 | 36.3838637460602 | 20.2848795762558 | 28.8119334683827 |
| chr19 | 1578490 | 1578496 | - | 4.01584230749346 | 7.95897019445067 | 2.38645642073598 | 1.06710864697714 |
| chr19 | 1578513 | 1578534 | - | 3.01188173062009 | 2.27399148412876 | 3.57968463110397 | 6.40265188186281 |
| chr19 | 1581104 | 1581125 | - | 7.02772403811355 | 5.68497871032191 | 2.38645642073598 | 3.20132594093141 |
| chr19 | 1581148 | 1581191 | - | 37.1465413443145 | 29.5618892936739 | 22.6713359969918 | 12.8053037637256 |
| chr19 | 1581263 | 1581271 | - | 29.1148567293276 | 29.5618892936739 | 10.7390538933119 | 11.7381951167485 |
| chr19 | 1582621 | 1582625 | - | 1.00396057687336 | 1.13699574206438 | 4.77291284147196 | 2.13421729395427 |
| chr19 | 1582667 | 1582704 | - | 10.0396057687336 | 4.54798296825753 | 9.54582568294392 | 3.20132594093141 |
| chr19 | 1584555 | 1584559 | - | 10.0396057687336 | 4.54798296825753 | 1.19322821036799 | 2.13421729395427 |
| chr19 | 1585112 | 1585128 | - | 7.02772403811355 | 3.41098722619315 | 4.77291284147196 | 1.06710864697714 |
| chr19 | 1585204 | 1585208 | - | 7.02772403811355 | 2.27399148412876 | 4.77291284147196 | 1.06710864697714 |
| chr19 | 1598068 | 1598174 | - | 44.174265382428 | 50.0278126508328 | 79.9462900946553 | 60.8251928776967 |
| chr19 | 1598204 | 1598216 | - | 38.1505019211879 | 12.5069531627082 | 25.0577924177278 | 46.952780466994 |
| chr19 | 1609355 | 1609358 | - | 2.00792115374673 | 7.95897019445067 | 1.19322821036799 | 6.40265188186281 |
| chr19 | 1609380 | 1609397 | - | 1.00396057687336 | 3.41098722619315 | 4.77291284147196 | 2.13421729395427 |
| chr19 | 1609457 | 1609482 | - | 41.1623836518079 | 29.5618892936739 | 16.7051949451519 | 24.5434988804741 |
| chr19 | 1609587 | 1609592 | - | 2.00792115374673 | 1.13699574206438 | 5.96614105183995 | 3.20132594093141 |
| chr19 | 1609708 | 1609720 | - | 3.01188173062009 | 6.82197445238629 | 8.35259747257593 | 2.13421729395427 |
| chr19 | 1610522 | 1610541 | - | 4.01584230749346 | 3.41098722619315 | 3.57968463110397 | 2.13421729395427 |
| chr19 | 1610859 | 1610891 | - | 16.0633692299738 | 5.68497871032191 | 1.19322821036799 | 4.26843458790854 |
| chr19 | 1610989 | 1611030 | - | 2.00792115374673 | 4.54798296825753 | 7.15936926220794 | 3.20132594093141 |
| chr19 | 1611076 | 1611083 | - | 8.03168461498692 | 5.68497871032191 | 11.9322821036799 | 14.9395210576799 |
| chr19 | 1611116 | 1611335 | - | 522.05949997415 | 392.263531012212 | 551.271433190011 | 685.083751359321 |
| chr19 | 1611390 | 1611534 | - | 86.3406096111093 | 54.5757956190903 | 101.424397881279 | 220.891489924267 |
| chr19 | 1611639 | 1611741 | - | 14.0554480762271 | 6.82197445238629 | 19.0916513658878 | 11.7381951167485 |
| chr19 | 1748263 | 1748301 | + | 3.01188173062009 | 5.68497871032191 | 8.35259747257593 | 4.26843458790854 |
| chr19 | 1799487 | 1799489 | - | 2.00792115374673 | 2.27399148412876 | 1.19322821036799 | 1.06710864697714 |
| chr19 | 1815274 | 1815282 | - | 3.01188173062009 | 3.41098722619315 | 4.77291284147196 | 11.7381951167485 |
| chr19 | 1815676 | 1815760 | - | 71.2812009580089 | 73.9047232341848 | 189.72328544851 | 80.0331485232852 |
| chr19 | 1852445 | 1852493 | - | 4.01584230749346 | 2.27399148412876 | 4.77291284147196 | 12.8053037637256 |
| chr19 | 1853171 | 1853245 | - | 26.1029749987075 | 11.3699574206438 | 26.2510206280958 | 42.6843458790854 |
| chr19 | 1853270 | 1853326 | - | 26.1029749987075 | 14.780944646837 | 33.4103898903037 | 122.717494402371 |
| chr19 | 1853350 | 1853400 | - | 10.0396057687336 | 5.68497871032191 | 14.3187385244159 | 6.40265188186281 |
| chr19 | 1853648 | 1853659 | - | 12.0475269224804 | 3.41098722619315 | 4.77291284147196 | 6.40265188186281 |
| chr19 | 1853716 | 1853736 | - | 7.02772403811355 | 6.82197445238629 | 5.96614105183995 | 33.0803680562912 |
| chr19 | 1853761 | 1853784 | - | 16.0633692299738 | 12.5069531627082 | 14.3187385244159 | 3.20132594093141 |
| chr19 | 1853811 | 1853852 | - | 10.0396057687336 | 6.82197445238629 | 3.57968463110397 | 1.06710864697714 |
| chr19 | 1854119 | 1854120 | - | 12.0475269224804 | 5.68497871032191 | 7.15936926220794 | 2.13421729395427 |
| chr19 | 1854233 | 1854257 | - | 4.01584230749346 | 4.54798296825753 | 2.38645642073598 | 2.13421729395427 |
| chr19 | 1876998 | 1877000 | - | 7.02772403811355 | 10.2329616785794 | 7.15936926220794 | 4.26843458790854 |
| chr19 | 1925040 | 1925077 | + | 12.0475269224804 | 22.7399148412876 | 11.9322821036799 | 6.40265188186281 |
| chr19 | 1925399 | 1925468 | + | 55.217831728035 | 18.1919318730301 | 36.9900745214077 | 22.4092815865198 |
| chr19 | 1925531 | 1925577 | + | 10.0396057687336 | 1.13699574206438 | 10.7390538933119 | 6.40265188186281 |
| chr19 | 1925930 | 1925980 | + | 3.01188173062009 | 3.41098722619315 | 5.96614105183995 | 4.26843458790854 |
| chr19 | 1980367 | 1980393 | + | 19.0752509605939 | 22.7399148412876 | 53.6952694665595 | 21.3421729395427 |
| chr19 | 1980443 | 1980462 | + | 1.00396057687336 | 2.27399148412876 | 8.35259747257593 | 14.9395210576799 |
| chr19 | 1980484 | 1980486 | + | 1.00396057687336 | 1.13699574206438 | 1.19322821036799 | 8.53686917581709 |
| chr19 | 1980639 | 1980670 | + | 11.043566345607 | 6.82197445238629 | 13.1255103140479 | 10.6710864697714 |
| chr19 | 1980703 | 1980727 | + | 2.00792115374673 | 4.54798296825753 | 7.15936926220794 | 40.5501285851312 |
| chr19 | 1981157 | 1981216 | + | 22.087132691214 | 18.1919318730301 | 21.4781077866238 | 32.0132594093141 |
| chr19 | 1981286 | 1981304 | + | 1.00396057687336 | 7.95897019445067 | 2.38645642073598 | 2.13421729395427 |
| chr19 | 1985470 | 1985514 | - | 12.0475269224804 | 12.5069531627082 | 4.77291284147196 | 6.40265188186281 |
| chr19 | 2037495 | 2037610 | - | 11.043566345607 | 9.09596593651506 | 29.8307052591997 | 20.2750642925656 |
| chr19 | 2037668 | 2037700 | - | 10.0396057687336 | 10.2329616785794 | 8.35259747257593 | 14.9395210576799 |
| chr19 | 2038182 | 2038185 | - | 13.0514874993537 | 5.68497871032191 | 13.1255103140479 | 2.13421729395427 |
| chr19 | 2038228 | 2038242 | - | 2.00792115374673 | 4.54798296825753 | 17.8984231555198 | 33.0803680562912 |
| chr19 | 2038284 | 2038362 | - | 18.0712903837206 | 9.09596593651506 | 39.3765309421437 | 22.4092815865198 |
| chr19 | 2038402 | 2038428 | - | 10.0396057687336 | 10.2329616785794 | 42.9562155732476 | 13.8724124107028 |
| chr19 | 2038497 | 2038517 | - | 6.02376346124019 | 1.13699574206438 | 13.1255103140479 | 7.46976052883995 |
| chr19 | 2038562 | 2038642 | - | 6.02376346124019 | 4.54798296825753 | 17.8984231555198 | 37.3488026441998 |
| chr19 | 2038699 | 2038722 | - | 4.01584230749346 | 5.68497871032191 | 19.0916513658878 | 43.7514545260626 |
| chr19 | 2038997 | 2039047 | - | 4.01584230749346 | 4.54798296825753 | 11.9322821036799 | 6.40265188186281 |
| chr19 | 2039464 | 2039481 | - | 4.01584230749346 | 10.2329616785794 | 13.1255103140479 | 26.6777161744284 |
| chr19 | 2071065 | 2071066 | - | 2.00792115374673 | 1.13699574206438 | 1.19322821036799 | 3.20132594093141 |
| chr19 | 2071900 | 2071909 | - | 2.00792115374673 | 7.95897019445067 | 7.15936926220794 | 9.60397782279422 |
| chr19 | 2073332 | 2073336 | - | 3.01188173062009 | 1.13699574206438 | 2.38645642073598 | 1.06710864697714 |
| chr19 | 2101027 | 2101139 | - | 52.205949997415 | 73.9047232341848 | 54.8884976769275 | 61.8923015246739 |
| chr19 | 2101162 | 2101182 | - | 12.0475269224804 | 13.6439489047726 | 20.2848795762558 | 5.33554323488568 |
| chr19 | 2101200 | 2101328 | - | 237.938656718987 | 222.851165444619 | 326.944529640829 | 446.051414436443 |
| chr19 | 2101355 | 2101393 | - | 5.01980288436682 | 2.27399148412876 | 9.54582568294392 | 21.3421729395427 |
| chr19 | 2101522 | 2101576 | - | 41.1623836518079 | 20.4659233571589 | 34.6036181006717 | 20.2750642925656 |
| chr19 | 2101634 | 2101681 | - | 7.02772403811355 | 1.13699574206438 | 11.9322821036799 | 28.8119334683827 |
| chr19 | 2101721 | 2101741 | - | 2.00792115374673 | 5.68497871032191 | 2.38645642073598 | 5.33554323488568 |
| chr19 | 2101774 | 2101808 | - | 5.01980288436682 | 4.54798296825753 | 9.54582568294392 | 12.8053037637256 |
| chr19 | 2229932 | 2229996 | + | 5.01980288436682 | 10.2329616785794 | 9.54582568294392 | 4.26843458790854 |
| chr19 | 2230503 | 2230522 | + | 5.01980288436682 | 3.41098722619315 | 3.57968463110397 | 6.40265188186281 |
| chr19 | 2230580 | 2230645 | + | 17.0673298068472 | 19.3289276150945 | 23.8645642073598 | 41.6172372321083 |
| chr19 | 2230746 | 2230748 | + | 2.00792115374673 | 4.54798296825753 | 1.19322821036799 | 1.06710864697714 |
| chr19 | 2232171 | 2232185 | + | 7.02772403811355 | 2.27399148412876 | 3.57968463110397 | 3.20132594093141 |
| chr19 | 2233173 | 2233237 | - | 39.1544624980612 | 70.4937360079917 | 45.3426719939836 | 76.8318225823538 |
| chr19 | 2233502 | 2233531 | - | 17.0673298068472 | 11.3699574206438 | 31.0239334695677 | 29.8790421153598 |
| chr19 | 2233569 | 2233625 | - | 9.03564519186028 | 19.3289276150945 | 20.2848795762558 | 28.8119334683827 |
| chr19 | 2233763 | 2233771 | - | 1.00396057687336 | 5.68497871032191 | 7.15936926220794 | 6.40265188186281 |
| chr19 | 2233805 | 2233807 | - | 2.00792115374673 | 1.13699574206438 | 2.38645642073598 | 1.06710864697714 |
| chr19 | 2233873 | 2233898 | - | 3.01188173062009 | 1.13699574206438 | 9.54582568294392 | 8.53686917581709 |
| chr19 | 2234000 | 2234052 | - | 4.01584230749346 | 7.95897019445067 | 14.3187385244159 | 12.8053037637256 |
| chr19 | 2248574 | 2248585 | + | 3.01188173062009 | 7.95897019445067 | 5.96614105183995 | 10.6710864697714 |
| chr19 | 2248609 | 2248636 | + | 12.0475269224804 | 10.2329616785794 | 2.38645642073598 | 13.8724124107028 |
| chr19 | 2272999 | 2273309 | + | 371.465413443145 | 454.798296825753 | 486.83710983014 | 659.47314383187 |
| chr19 | 2273371 | 2273469 | + | 19.0752509605939 | 12.5069531627082 | 32.2171616799357 | 33.0803680562912 |
| chr19 | 2321559 | 2321692 | - | 107.42378172545 | 68.2197445238629 | 131.255103140479 | 325.468137328026 |
| chr19 | 2321718 | 2321729 | - | 3.01188173062009 | 3.41098722619315 | 5.96614105183995 | 8.53686917581709 |
| chr19 | 2324127 | 2324198 | - | 7.02772403811355 | 4.54798296825753 | 33.4103898903037 | 21.3421729395427 |
| chr19 | 2426588 | 2426635 | - | 36.1425807674411 | 50.0278126508328 | 16.7051949451519 | 41.6172372321083 |
| chr19 | 2426657 | 2426668 | - | 8.03168461498692 | 5.68497871032191 | 4.77291284147196 | 11.7381951167485 |
| chr19 | 2426934 | 2426946 | - | 2.00792115374673 | 2.27399148412876 | 1.19322821036799 | 2.13421729395427 |
| chr19 | 2428187 | 2428212 | - | 2.00792115374673 | 3.41098722619315 | 1.19322821036799 | 5.33554323488568 |
| chr19 | 2428260 | 2428357 | - | 31.1227778830743 | 19.3289276150945 | 31.0239334695677 | 29.8790421153598 |
| chr19 | 2428705 | 2428927 | - | 104.41189999483 | 81.8636934286355 | 145.573841664895 | 91.7713436400337 |
| chr19 | 2429074 | 2429173 | - | 20.0792115374673 | 17.0549361309657 | 44.1494437836156 | 30.9461507623369 |
| chr19 | 2429269 | 2429316 | - | 12.0475269224804 | 10.2329616785794 | 17.8984231555198 | 13.8724124107028 |
| chr19 | 2429467 | 2429486 | - | 5.01980288436682 | 3.41098722619315 | 4.77291284147196 | 2.13421729395427 |
| chr19 | 2429691 | 2429705 | - | 3.01188173062009 | 3.41098722619315 | 5.96614105183995 | 1.06710864697714 |
| chr19 | 2429747 | 2429769 | - | 1.00396057687336 | 3.41098722619315 | 2.38645642073598 | 5.33554323488568 |
| chr19 | 2429844 | 2429845 | - | 1.00396057687336 | 1.13699574206438 | 1.19322821036799 | 1.06710864697714 |
| chr19 | 2430020 | 2430058 | - | 13.0514874993537 | 1.13699574206438 | 3.57968463110397 | 3.20132594093141 |
| chr19 | 2430130 | 2430147 | - | 10.0396057687336 | 11.3699574206438 | 8.35259747257593 | 5.33554323488568 |
| chr19 | 2430197 | 2430224 | - | 11.043566345607 | 18.1919318730301 | 9.54582568294392 | 4.26843458790854 |
| chr19 | 2430257 | 2430303 | - | 19.0752509605939 | 10.2329616785794 | 5.96614105183995 | 5.33554323488568 |
| chr19 | 2430522 | 2430544 | - | 7.02772403811355 | 4.54798296825753 | 2.38645642073598 | 6.40265188186281 |
| chr19 | 2732566 | 2732675 | - | 13.0514874993537 | 10.2329616785794 | 15.5119667347839 | 38.4159112911769 |
| chr19 | 2754737 | 2754787 | - | 54.2138711511617 | 63.6717615556054 | 25.0577924177278 | 49.0869977609482 |
| chr19 | 2755207 | 2755220 | - | 4.01584230749346 | 1.13699574206438 | 5.96614105183995 | 50.1541064079254 |
| chr19 | 2755272 | 2755337 | - | 18.0712903837206 | 6.82197445238629 | 26.2510206280958 | 65.0936274656053 |
| chr19 | 2755395 | 2755420 | - | 6.02376346124019 | 9.09596593651506 | 26.2510206280958 | 17.0737383516342 |
| chr19 | 2755453 | 2755463 | - | 1.00396057687336 | 1.13699574206438 | 4.77291284147196 | 1.06710864697714 |
| chr19 | 2755508 | 2755525 | - | 2.00792115374673 | 4.54798296825753 | 1.19322821036799 | 9.60397782279422 |
| chr19 | 2755587 | 2755677 | - | 9.03564519186028 | 12.5069531627082 | 7.15936926220794 | 6.40265188186281 |
| chr19 | 2755809 | 2755858 | - | 8.03168461498692 | 7.95897019445067 | 15.5119667347839 | 9.60397782279422 |
| chr19 | 2813374 | 2813395 | + | 2.00792115374673 | 1.13699574206438 | 1.19322821036799 | 3.20132594093141 |
| chr19 | 2813455 | 2813511 | + | 6.02376346124019 | 3.41098722619315 | 1.19322821036799 | 3.20132594093141 |
| chr19 | 2813551 | 2813577 | + | 30.1188173062009 | 35.2468680039958 | 21.4781077866238 | 16.006629704657 |
| chr19 | 2933464 | 2933467 | - | 1.00396057687336 | 1.13699574206438 | 1.19322821036799 | 1.06710864697714 |
| chr19 | 2997665 | 2997691 | - | 6.02376346124019 | 6.82197445238629 | 5.96614105183995 | 16.006629704657 |
| chr19 | 2997734 | 2997757 | - | 7.02772403811355 | 3.41098722619315 | 3.57968463110397 | 1.06710864697714 |
| chr19 | 2997839 | 2997858 | - | 4.01584230749346 | 4.54798296825753 | 3.57968463110397 | 5.33554323488568 |
| chr19 | 2997879 | 2997885 | - | 2.00792115374673 | 9.09596593651506 | 4.77291284147196 | 2.13421729395427 |
| chr19 | 3052937 | 3052977 | - | 97.3841759567164 | 75.0417189762492 | 35.7968463110397 | 64.0265188186281 |
| chr19 | 3053088 | 3053091 | - | 5.01980288436682 | 4.54798296825753 | 3.57968463110397 | 3.20132594093141 |
| chr19 | 3053155 | 3053260 | - | 184.728746144699 | 109.151591238181 | 64.4343233598714 | 134.455689519119 |
| chr19 | 3053460 | 3053500 | - | 5.01980288436682 | 3.41098722619315 | 3.57968463110397 | 6.40265188186281 |
| chr19 | 3053542 | 3053596 | - | 8.03168461498692 | 2.27399148412876 | 10.7390538933119 | 6.40265188186281 |
| chr19 | 3053650 | 3053683 | - | 2.00792115374673 | 4.54798296825753 | 7.15936926220794 | 1.06710864697714 |
| chr19 | 3053764 | 3053822 | - | 19.0752509605939 | 9.09596593651506 | 14.3187385244159 | 14.9395210576799 |
| chr19 | 3121253 | 3121301 | + | 354.398083636298 | 401.359496948727 | 60.8546387287675 | 129.120146284233 |
| chr19 | 3121350 | 3121356 | + | 99.3920971104631 | 84.1376849127643 | 7.15936926220794 | 7.46976052883995 |
| chr19 | 3121406 | 3121428 | + | 7.02772403811355 | 5.68497871032191 | 1.19322821036799 | 1.06710864697714 |
| chr19 | 3121821 | 3121832 | + | 33.130699036821 | 29.5618892936739 | 9.54582568294392 | 16.006629704657 |
| chr19 | 3123855 | 3123875 | + | 1.00396057687336 | 2.27399148412876 | 1.19322821036799 | 3.20132594093141 |
| chr19 | 3207810 | 3207821 | + | 4.01584230749346 | 4.54798296825753 | 5.96614105183995 | 5.33554323488568 |
| chr19 | 3209202 | 3209204 | + | 3.01188173062009 | 4.54798296825753 | 15.5119667347839 | 4.26843458790854 |
| chr19 | 3209518 | 3209550 | + | 31.1227778830743 | 22.7399148412876 | 17.8984231555198 | 26.6777161744284 |
| chr19 | 3456592 | 3456630 | + | 13.0514874993537 | 38.657855230189 | 3.57968463110397 | 3.20132594093141 |
| chr19 | 3462762 | 3462774 | + | 23.0910932680874 | 10.2329616785794 | 4.77291284147196 | 3.20132594093141 |
| chr19 | 3462801 | 3462973 | + | 418.651560556193 | 458.209284051946 | 119.322821036799 | 74.6976052883995 |
| chr19 | 3464879 | 3464910 | + | 21.0831721143407 | 9.09596593651506 | 1.19322821036799 | 3.20132594093141 |
| chr19 | 3464950 | 3464994 | + | 45.1782259593014 | 47.753821166704 | 8.35259747257593 | 23.476390233497 |
| chr19 | 3465068 | 3465140 | + | 139.550520185398 | 79.5897019445067 | 17.8984231555198 | 28.8119334683827 |
| chr19 | 3465172 | 3465182 | + | 9.03564519186028 | 14.780944646837 | 1.19322821036799 | 1.06710864697714 |
| chr19 | 3465203 | 3465258 | + | 45.1782259593014 | 61.3977700714766 | 4.77291284147196 | 1.06710864697714 |
| chr19 | 3465285 | 3465422 | + | 147.582204800385 | 193.289276150945 | 31.0239334695677 | 10.6710864697714 |
| chr19 | 3465458 | 3465467 | + | 2.00792115374673 | 6.82197445238629 | 1.19322821036799 | 1.06710864697714 |
| chr19 | 3466008 | 3466024 | + | 3.01188173062009 | 6.82197445238629 | 2.38645642073598 | 2.13421729395427 |
| chr19 | 3466102 | 3466198 | + | 85.336649034236 | 86.411676396893 | 9.54582568294392 | 20.2750642925656 |
| chr19 | 3466618 | 3466646 | + | 8.03168461498692 | 9.09596593651506 | 8.35259747257593 | 25.6106075274513 |
| chr19 | 3466961 | 3466995 | + | 4.01584230749346 | 3.41098722619315 | 1.19322821036799 | 1.06710864697714 |
| chr19 | 3467058 | 3467091 | + | 71.2812009580089 | 65.9457530397342 | 17.8984231555198 | 42.6843458790854 |
| chr19 | 3467891 | 3467900 | + | 16.0633692299738 | 11.3699574206438 | 9.54582568294392 | 1.06710864697714 |
| chr19 | 3468032 | 3468035 | + | 9.03564519186028 | 17.0549361309657 | 1.19322821036799 | 1.06710864697714 |
| chr19 | 3468060 | 3468077 | + | 5.01980288436682 | 6.82197445238629 | 2.38645642073598 | 2.13421729395427 |
| chr19 | 3468570 | 3468608 | + | 43.1703048055547 | 38.657855230189 | 21.4781077866238 | 51.2212150549025 |
| chr19 | 3468670 | 3468676 | + | 18.0712903837206 | 22.7399148412876 | 8.35259747257593 | 3.20132594093141 |
| chr19 | 3469123 | 3469127 | + | 20.0792115374673 | 67.0827487817985 | 19.0916513658878 | 24.5434988804741 |
| chr19 | 3490846 | 3490907 | - | 18.0712903837206 | 32.9728765198671 | 9.54582568294392 | 18.1408469986113 |
| chr19 | 3491371 | 3491374 | - | 1.00396057687336 | 1.13699574206438 | 1.19322821036799 | 2.13421729395427 |
| chr19 | 3534950 | 3534983 | + | 6.02376346124019 | 7.95897019445067 | 9.54582568294392 | 2.13421729395427 |
| chr19 | 3535632 | 3535645 | + | 2.00792115374673 | 1.13699574206438 | 2.38645642073598 | 2.13421729395427 |
| chr19 | 3535673 | 3535692 | + | 1.00396057687336 | 1.13699574206438 | 2.38645642073598 | 5.33554323488568 |
| chr19 | 3535766 | 3535772 | + | 2.00792115374673 | 2.27399148412876 | 16.7051949451519 | 2.13421729395427 |
| chr19 | 3536148 | 3536174 | + | 4.01584230749346 | 4.54798296825753 | 9.54582568294392 | 17.0737383516342 |
| chr19 | 3544218 | 3544576 | - | 347.370359598184 | 447.976322373366 | 138.414472402687 | 136.589906813073 |
| chr19 | 3544636 | 3544688 | - | 31.1227778830743 | 56.8497871032191 | 4.77291284147196 | 16.006629704657 |
| chr19 | 3578509 | 3578674 | + | 125.495072109171 | 110.288586980245 | 34.6036181006717 | 32.0132594093141 |
| chr19 | 3578699 | 3578810 | + | 52.205949997415 | 36.3838637460602 | 20.2848795762558 | 23.476390233497 |
| chr19 | 3578856 | 3578942 | + | 42.1663442286813 | 17.0549361309657 | 10.7390538933119 | 10.6710864697714 |
| chr19 | 3578963 | 3579045 | + | 186.736667298446 | 173.96034853585 | 32.2171616799357 | 43.7514545260626 |
| chr19 | 3630905 | 3630940 | - | 1.00396057687336 | 1.13699574206438 | 9.54582568294392 | 6.40265188186281 |
| chr19 | 3767515 | 3767545 | + | 14.0554480762271 | 23.876910583352 | 9.54582568294392 | 13.8724124107028 |
| chr19 | 3777994 | 3778027 | - | 15.0594086531005 | 13.6439489047726 | 8.35259747257593 | 10.6710864697714 |
| chr19 | 3958479 | 3958535 | - | 5.01980288436682 | 4.54798296825753 | 2.38645642073598 | 4.26843458790854 |
| chr19 | 3958608 | 3958629 | - | 17.0673298068472 | 6.82197445238629 | 2.38645642073598 | 7.46976052883995 |
| chr19 | 3958652 | 3958697 | - | 33.130699036821 | 29.5618892936739 | 13.1255103140479 | 36.2816939972226 |
| chr19 | 3958734 | 3958758 | - | 6.02376346124019 | 3.41098722619315 | 4.77291284147196 | 10.6710864697714 |
| chr19 | 3958913 | 3958975 | - | 46.1821865361748 | 43.2058381984465 | 28.6374770488317 | 30.9461507623369 |
| chr19 | 3976076 | 3976769 | - | 1146.52297878938 | 1287.07918001688 | 1150.27199479474 | 1737.25287727878 |
| chr19 | 4037920 | 4037952 | + | 5.01980288436682 | 9.09596593651506 | 14.3187385244159 | 13.8724124107028 |
| chr19 | 4038290 | 4038301 | + | 9.03564519186028 | 1.13699574206438 | 1.19322821036799 | 3.20132594093141 |
| chr19 | 4044784 | 4044812 | - | 9.03564519186028 | 9.09596593651506 | 9.54582568294392 | 7.46976052883995 |
| chr19 | 4044925 | 4044974 | - | 18.0712903837206 | 12.5069531627082 | 19.0916513658878 | 20.2750642925656 |
| chr19 | 4045176 | 4045180 | - | 21.0831721143407 | 13.6439489047726 | 19.0916513658878 | 10.6710864697714 |
| chr19 | 4045280 | 4045284 | - | 15.0594086531005 | 2.27399148412876 | 3.57968463110397 | 11.7381951167485 |
| chr19 | 4045848 | 4045878 | - | 11.043566345607 | 7.95897019445067 | 2.38645642073598 | 4.26843458790854 |
| chr19 | 4045916 | 4045949 | - | 11.043566345607 | 5.68497871032191 | 4.77291284147196 | 1.06710864697714 |
| chr19 | 4046392 | 4046414 | - | 4.01584230749346 | 6.82197445238629 | 3.57968463110397 | 3.20132594093141 |
| chr19 | 4046528 | 4046536 | - | 33.130699036821 | 23.876910583352 | 27.4442488384638 | 5.33554323488568 |
| chr19 | 4046663 | 4046748 | - | 256.009947102708 | 222.851165444619 | 147.960298085631 | 130.187254931211 |
| chr19 | 4046854 | 4046866 | - | 21.0831721143407 | 9.09596593651506 | 5.96614105183995 | 9.60397782279422 |
| chr19 | 4046897 | 4046928 | - | 32.1267384599477 | 9.09596593651506 | 7.15936926220794 | 11.7381951167485 |
| chr19 | 4047325 | 4047342 | - | 4.01584230749346 | 3.41098722619315 | 2.38645642073598 | 2.13421729395427 |
| chr19 | 4090368 | 4090482 | - | 117.463387494184 | 102.329616785794 | 60.8546387287675 | 74.6976052883995 |
| chr19 | 4090555 | 4090597 | - | 10.0396057687336 | 10.2329616785794 | 19.0916513658878 | 5.33554323488568 |
| chr19 | 4174134 | 4174175 | - | 21.0831721143407 | 27.2878978095452 | 8.35259747257593 | 12.8053037637256 |
| chr19 | 4174210 | 4174235 | - | 5.01980288436682 | 4.54798296825753 | 1.19322821036799 | 2.13421729395427 |
| chr19 | 4359995 | 4360051 | + | 5.01980288436682 | 3.41098722619315 | 2.38645642073598 | 2.13421729395427 |
| chr19 | 4360421 | 4360468 | - | 17.0673298068472 | 21.6029190992233 | 13.1255103140479 | 33.0803680562912 |
| chr19 | 4361141 | 4361178 | - | 2.00792115374673 | 5.68497871032191 | 13.1255103140479 | 3.20132594093141 |
| chr19 | 4361370 | 4361430 | - | 20.0792115374673 | 9.09596593651506 | 29.8307052591997 | 9.60397782279422 |
| chr19 | 4443014 | 4443071 | + | 5.01980288436682 | 17.0549361309657 | 5.96614105183995 | 8.53686917581709 |
| chr19 | 4443347 | 4443372 | + | 2.00792115374673 | 13.6439489047726 | 7.15936926220794 | 5.33554323488568 |
| chr19 | 4445035 | 4445050 | - | 2.00792115374673 | 3.41098722619315 | 2.38645642073598 | 2.13421729395427 |
| chr19 | 4445072 | 4445086 | - | 1.00396057687336 | 2.27399148412876 | 8.35259747257593 | 11.7381951167485 |
| chr19 | 4502078 | 4502114 | + | 5.01980288436682 | 7.95897019445067 | 16.7051949451519 | 5.33554323488568 |
| chr19 | 4654569 | 4654632 | + | 4.01584230749346 | 1.13699574206438 | 19.0916513658878 | 8.53686917581709 |
| chr19 | 4657574 | 4657728 | - | 97.3841759567164 | 135.302493305661 | 54.8884976769275 | 64.0265188186281 |
| chr19 | 4657802 | 4658085 | - | 72.2851615348822 | 56.8497871032191 | 99.0379414605431 | 87.5029090521251 |
| chr19 | 4675252 | 4675270 | - | 4.01584230749346 | 3.41098722619315 | 1.19322821036799 | 4.26843458790854 |
| chr19 | 4675377 | 4675384 | - | 13.0514874993537 | 9.09596593651506 | 2.38645642073598 | 4.26843458790854 |
| chr19 | 4675451 | 4675469 | - | 7.02772403811355 | 3.41098722619315 | 3.57968463110397 | 4.26843458790854 |
| chr19 | 4675606 | 4675624 | - | 10.0396057687336 | 3.41098722619315 | 1.19322821036799 | 16.006629704657 |
| chr19 | 4675713 | 4675724 | - | 5.01980288436682 | 1.13699574206438 | 1.19322821036799 | 1.06710864697714 |
| chr19 | 4676101 | 4676108 | - | 7.02772403811355 | 2.27399148412876 | 4.77291284147196 | 1.06710864697714 |
| chr19 | 4694386 | 4694426 | - | 11.043566345607 | 6.82197445238629 | 7.15936926220794 | 7.46976052883995 |
| chr19 | 4815964 | 4815999 | - | 22.087132691214 | 29.5618892936739 | 4.77291284147196 | 5.33554323488568 |
| chr19 | 4838350 | 4838385 | + | 9.03564519186028 | 4.54798296825753 | 5.96614105183995 | 7.46976052883995 |
| chr19 | 4838366 | 4838512 | - | 71.2812009580089 | 32.9728765198671 | 38.1833027317757 | 52.2883237018796 |
| chr19 | 5151725 | 5151727 | + | 6.02376346124019 | 7.95897019445067 | 7.15936926220794 | 4.26843458790854 |
| chr19 | 5152007 | 5152013 | + | 1.00396057687336 | 1.13699574206438 | 3.57968463110397 | 7.46976052883995 |
| chr19 | 5152780 | 5152795 | + | 3.01188173062009 | 3.41098722619315 | 16.7051949451519 | 8.53686917581709 |
| chr19 | 5153169 | 5153177 | + | 1.00396057687336 | 2.27399148412876 | 9.54582568294392 | 7.46976052883995 |
| chr19 | 5153198 | 5153232 | + | 5.01980288436682 | 2.27399148412876 | 23.8645642073598 | 20.2750642925656 |
| chr19 | 5153264 | 5153313 | + | 2.00792115374673 | 1.13699574206438 | 25.0577924177278 | 3.20132594093141 |
| chr19 | 5153415 | 5153446 | + | 1.00396057687336 | 2.27399148412876 | 2.38645642073598 | 8.53686917581709 |
| chr19 | 5206279 | 5206406 | - | 57.2257528817818 | 28.4248935516096 | 33.4103898903037 | 70.429170700491 |
| chr19 | 5206441 | 5206465 | - | 5.01980288436682 | 2.27399148412876 | 4.77291284147196 | 23.476390233497 |
| chr19 | 5206488 | 5206506 | - | 20.0792115374673 | 19.3289276150945 | 8.35259747257593 | 11.7381951167485 |
| chr19 | 5587035 | 5587050 | - | 5.01980288436682 | 5.68497871032191 | 8.35259747257593 | 4.26843458790854 |
| chr19 | 5587166 | 5587187 | - | 17.0673298068472 | 12.5069531627082 | 4.77291284147196 | 4.26843458790854 |
| chr19 | 5668193 | 5668359 | + | 155.613889415371 | 140.987472015983 | 274.442488384638 | 232.629685041016 |
| chr19 | 5678443 | 5678469 | - | 36.1425807674411 | 39.7948509722534 | 41.7629873628796 | 139.791232754005 |
| chr19 | 5691530 | 5691563 | + | 267.053513448315 | 219.440178218426 | 398.538222262909 | 541.024084017408 |
| chr19 | 5691600 | 5691640 | + | 35.1386201905678 | 30.6988850357383 | 47.7291284147196 | 78.966039876308 |
| chr19 | 5691790 | 5691846 | + | 44.174265382428 | 35.2468680039958 | 33.4103898903037 | 30.9461507623369 |
| chr19 | 5691852 | 5691966 | - | 12.0475269224804 | 15.9179403889013 | 25.0577924177278 | 32.0132594093141 |
| chr19 | 5785164 | 5785186 | - | 65.2574374967687 | 73.9047232341848 | 56.0817258872955 | 66.1607361125824 |
| chr19 | 5894697 | 5894814 | - | 20.0792115374673 | 18.1919318730301 | 10.7390538933119 | 37.3488026441998 |
| chr19 | 5916163 | 5916246 | - | 5.01980288436682 | 9.09596593651506 | 3.57968463110397 | 4.26843458790854 |
| chr19 | 5916277 | 5916319 | - | 4.01584230749346 | 2.27399148412876 | 3.57968463110397 | 5.33554323488568 |
| chr19 | 5917040 | 5917044 | - | 33.130699036821 | 36.3838637460602 | 14.3187385244159 | 17.0737383516342 |
| chr19 | 5917336 | 5917445 | - | 5.01980288436682 | 6.82197445238629 | 2.38645642073598 | 7.46976052883995 |
| chr19 | 5917512 | 5917517 | - | 2.00792115374673 | 2.27399148412876 | 19.0916513658878 | 1.06710864697714 |
| chr19 | 5978343 | 5978463 | + | 58.2297134586551 | 84.1376849127643 | 468.93868667462 | 310.528616270347 |
| chr19 | 6210406 | 6210428 | - | 2.00792115374673 | 14.780944646837 | 3.57968463110397 | 7.46976052883995 |
| chr19 | 6211206 | 6211249 | - | 12.0475269224804 | 23.876910583352 | 23.8645642073598 | 8.53686917581709 |
| chr19 | 6211345 | 6211362 | - | 11.043566345607 | 10.2329616785794 | 7.15936926220794 | 4.26843458790854 |
| chr19 | 6211503 | 6211597 | - | 27.1069355755808 | 34.1098722619315 | 33.4103898903037 | 10.6710864697714 |
| chr19 | 6211749 | 6211789 | - | 9.03564519186028 | 7.95897019445067 | 7.15936926220794 | 1.06710864697714 |
| chr19 | 6212129 | 6212157 | - | 6.02376346124019 | 6.82197445238629 | 1.19322821036799 | 4.26843458790854 |
| chr19 | 6368794 | 6368873 | + | 74.293082688629 | 121.658544400889 | 69.2072362013434 | 75.7647139353766 |
| chr19 | 6379600 | 6379616 | - | 5.01980288436682 | 13.6439489047726 | 5.96614105183995 | 10.6710864697714 |
| chr19 | 6413448 | 6413459 | - | 3.01188173062009 | 1.13699574206438 | 2.38645642073598 | 7.46976052883995 |
| chr19 | 6413480 | 6413524 | - | 6.02376346124019 | 1.13699574206438 | 3.57968463110397 | 14.9395210576799 |
| chr19 | 6413680 | 6413709 | - | 6.02376346124019 | 6.82197445238629 | 3.57968463110397 | 6.40265188186281 |
| chr19 | 6413852 | 6413895 | - | 14.0554480762271 | 6.82197445238629 | 10.7390538933119 | 13.8724124107028 |
| chr19 | 6413920 | 6414019 | - | 121.479229801677 | 50.0278126508328 | 113.356679984959 | 155.797862458662 |
| chr19 | 6414042 | 6414058 | - | 5.01980288436682 | 2.27399148412876 | 3.57968463110397 | 1.06710864697714 |
| chr19 | 6414082 | 6414089 | - | 7.02772403811355 | 3.41098722619315 | 5.96614105183995 | 14.9395210576799 |
| chr19 | 6440085 | 6440101 | - | 1.00396057687336 | 3.41098722619315 | 1.19322821036799 | 2.13421729395427 |
| chr19 | 6441229 | 6441283 | - | 20.0792115374673 | 12.5069531627082 | 38.1833027317757 | 37.3488026441998 |
| chr19 | 6441748 | 6441771 | - | 1.00396057687336 | 1.13699574206438 | 4.77291284147196 | 6.40265188186281 |
| chr19 | 6663516 | 6663544 | - | 2.00792115374673 | 1.13699574206438 | 1.19322821036799 | 2.13421729395427 |
| chr19 | 6663570 | 6663611 | - | 6.02376346124019 | 1.13699574206438 | 4.77291284147196 | 3.20132594093141 |
| chr19 | 6677854 | 6678021 | - | 40.1584230749346 | 31.8358807778027 | 2.38645642073598 | 2.13421729395427 |
| chr19 | 6751265 | 6751276 | + | 18.0712903837206 | 11.3699574206438 | 8.35259747257593 | 16.006629704657 |
| chr19 | 6751370 | 6751381 | + | 3.01188173062009 | 5.68497871032191 | 1.19322821036799 | 4.26843458790854 |
| chr19 | 6751482 | 6751484 | + | 25.0990144218341 | 36.3838637460602 | 13.1255103140479 | 5.33554323488568 |
| chr19 | 6752208 | 6752215 | - | 2.00792115374673 | 7.95897019445067 | 7.15936926220794 | 2.13421729395427 |
| chr19 | 7520959 | 7521003 | + | 11.043566345607 | 13.6439489047726 | 2.38645642073598 | 4.26843458790854 |
| chr19 | 7533961 | 7533967 | + | 6.02376346124019 | 10.2329616785794 | 3.57968463110397 | 6.40265188186281 |
| chr19 | 7534161 | 7534198 | + | 15.0594086531005 | 11.3699574206438 | 3.57968463110397 | 4.26843458790854 |
| chr19 | 7534711 | 7534771 | + | 87.3445701879827 | 48.8908169087684 | 1.19322821036799 | 2.13421729395427 |
| chr19 | 7638732 | 7638767 | + | 4.01584230749346 | 3.41098722619315 | 16.7051949451519 | 20.2750642925656 |
| chr19 | 7639800 | 7639808 | + | 1.00396057687336 | 1.13699574206438 | 3.57968463110397 | 2.13421729395427 |
| chr19 | 7640741 | 7640769 | + | 1.00396057687336 | 3.41098722619315 | 9.54582568294392 | 5.33554323488568 |
| chr19 | 7647821 | 7647846 | + | 3.01188173062009 | 6.82197445238629 | 7.15936926220794 | 10.6710864697714 |
| chr19 | 7682810 | 7682832 | + | 9.03564519186028 | 12.5069531627082 | 1.19322821036799 | 4.26843458790854 |
| chr19 | 7913076 | 7913083 | + | 7.02772403811355 | 10.2329616785794 | 9.54582568294392 | 26.6777161744284 |
| chr19 | 7913123 | 7913180 | + | 37.1465413443145 | 42.0688424563821 | 16.7051949451519 | 14.9395210576799 |
| chr19 | 7913402 | 7913488 | + | 27.1069355755808 | 19.3289276150945 | 17.8984231555198 | 19.2079556455884 |
| chr19 | 7913517 | 7913547 | + | 19.0752509605939 | 11.3699574206438 | 9.54582568294392 | 3.20132594093141 |
| chr19 | 7913679 | 7913718 | + | 13.0514874993537 | 17.0549361309657 | 7.15936926220794 | 7.46976052883995 |
| chr19 | 7914433 | 7914457 | + | 6.02376346124019 | 14.780944646837 | 7.15936926220794 | 4.26843458790854 |
| chr19 | 7922948 | 7922984 | + | 5.01980288436682 | 1.13699574206438 | 13.1255103140479 | 25.6106075274513 |
| chr19 | 7923065 | 7923106 | + | 15.0594086531005 | 5.68497871032191 | 39.3765309421437 | 35.2145853502455 |
| chr19 | 7923129 | 7923144 | + | 21.0831721143407 | 31.8358807778027 | 34.6036181006717 | 74.6976052883995 |
| chr19 | 7923165 | 7923229 | + | 11.043566345607 | 17.0549361309657 | 9.54582568294392 | 8.53686917581709 |
| chr19 | 7924529 | 7924530 | - | 2.00792115374673 | 1.13699574206438 | 2.38645642073598 | 4.26843458790854 |
| chr19 | 7926749 | 7926768 | - | 3.01188173062009 | 7.95897019445067 | 3.57968463110397 | 9.60397782279422 |
| chr19 | 7926805 | 7926836 | - | 1.00396057687336 | 4.54798296825753 | 4.77291284147196 | 4.26843458790854 |
| chr19 | 7959378 | 7959387 | - | 5.01980288436682 | 6.82197445238629 | 3.57968463110397 | 6.40265188186281 |
| chr19 | 7962360 | 7962408 | - | 46.1821865361748 | 38.657855230189 | 31.0239334695677 | 27.7448248214055 |
| chr19 | 7962453 | 7962477 | - | 3.01188173062009 | 1.13699574206438 | 14.3187385244159 | 7.46976052883995 |
| chr19 | 7962518 | 7962578 | - | 15.0594086531005 | 21.6029190992233 | 68.0140079909754 | 35.2145853502455 |
| chr19 | 7962607 | 7962629 | - | 7.02772403811355 | 4.54798296825753 | 17.8984231555198 | 24.5434988804741 |
| chr19 | 7963423 | 7963502 | - | 5.01980288436682 | 9.09596593651506 | 13.1255103140479 | 21.3421729395427 |
| chr19 | 7963691 | 7963709 | - | 2.00792115374673 | 1.13699574206438 | 1.19322821036799 | 1.06710864697714 |
| chr19 | 8302148 | 8302169 | - | 8.03168461498692 | 12.5069531627082 | 2.38645642073598 | 8.53686917581709 |
| chr19 | 8302192 | 8302229 | - | 603.380306700892 | 342.235718361379 | 63.2410951495034 | 148.328101929822 |
| chr19 | 8302251 | 8302319 | - | 37.1465413443145 | 34.1098722619315 | 13.1255103140479 | 22.4092815865198 |
| chr19 | 8302340 | 8302344 | - | 1.00396057687336 | 1.13699574206438 | 4.77291284147196 | 6.40265188186281 |
| chr19 | 8302364 | 8302436 | - | 38.1505019211879 | 13.6439489047726 | 11.9322821036799 | 32.0132594093141 |
| chr19 | 8308280 | 8308395 | - | 3730.71750366142 | 2247.84058206128 | 658.66197212313 | 1206.89987973114 |
| chr19 | 8311404 | 8311454 | - | 89.3524913417294 | 96.6446380754725 | 159.892580189311 | 140.858341400982 |
| chr19 | 8311482 | 8311501 | - | 1.00396057687336 | 2.27399148412876 | 1.19322821036799 | 3.20132594093141 |
| chr19 | 8403996 | 8404010 | + | 3.01188173062009 | 1.13699574206438 | 2.38645642073598 | 1.06710864697714 |
| chr19 | 8404144 | 8404162 | + | 7.02772403811355 | 1.13699574206438 | 2.38645642073598 | 3.20132594093141 |
| chr19 | 8404187 | 8404255 | + | 4.01584230749346 | 3.41098722619315 | 2.38645642073598 | 7.46976052883995 |
| chr19 | 8404366 | 8404408 | + | 8.03168461498692 | 7.95897019445067 | 5.96614105183995 | 2.13421729395427 |
| chr19 | 8488713 | 8488882 | + | 81.3208067267425 | 69.3567402659273 | 39.3765309421437 | 150.462319223776 |
| chr19 | 8488924 | 8489090 | + | 74.293082688629 | 98.9186295596012 | 63.2410951495034 | 81.1002571702623 |
| chr19 | 8510612 | 8510636 | - | 3.01188173062009 | 2.27399148412876 | 1.19322821036799 | 4.26843458790854 |
| chr19 | 8809837 | 8809850 | - | 18.0712903837206 | 32.9728765198671 | 78.7530618842873 | 53.3554323488568 |
| chr19 | 8809871 | 8809961 | - | 14.0554480762271 | 23.876910583352 | 42.9562155732476 | 59.7580842307196 |
| chr19 | 8810016 | 8810054 | - | 1.00396057687336 | 4.54798296825753 | 9.54582568294392 | 9.60397782279422 |
| chr19 | 9162187 | 9162192 | + | 6.02376346124019 | 2.27399148412876 | 26.2510206280958 | 38.4159112911769 |
| chr19 | 9162576 | 9162608 | + | 3.01188173062009 | 3.41098722619315 | 2.38645642073598 | 5.33554323488568 |
| chr19 | 9412856 | 9412875 | - | 1.00396057687336 | 1.13699574206438 | 3.57968463110397 | 1.06710864697714 |
| chr19 | 9523333 | 9523335 | - | 1.00396057687336 | 4.54798296825753 | 1.19322821036799 | 2.13421729395427 |
| chr19 | 9538691 | 9538739 | + | 7.02772403811355 | 9.09596593651506 | 7.15936926220794 | 13.8724124107028 |
| chr19 | 9560374 | 9560377 | - | 5.01980288436682 | 18.1919318730301 | 8.35259747257593 | 10.6710864697714 |
| chr19 | 9560503 | 9560528 | - | 3.01188173062009 | 6.82197445238629 | 8.35259747257593 | 5.33554323488568 |
| chr19 | 9560890 | 9560891 | - | 7.02772403811355 | 5.68497871032191 | 9.54582568294392 | 12.8053037637256 |
| chr19 | 9561182 | 9561298 | - | 20.0792115374673 | 19.3289276150945 | 54.8884976769275 | 29.8790421153598 |
| chr19 | 9561471 | 9561477 | - | 12.0475269224804 | 9.09596593651506 | 19.0916513658878 | 14.9395210576799 |
| chr19 | 9561501 | 9561507 | - | 5.01980288436682 | 6.82197445238629 | 9.54582568294392 | 10.6710864697714 |
| chr19 | 9561541 | 9561545 | - | 1.00396057687336 | 4.54798296825753 | 4.77291284147196 | 2.13421729395427 |
| chr19 | 9561622 | 9561627 | - | 2.00792115374673 | 2.27399148412876 | 44.1494437836156 | 24.5434988804741 |
| chr19 | 9561925 | 9561949 | - | 9.03564519186028 | 3.41098722619315 | 27.4442488384638 | 26.6777161744284 |
| chr19 | 9561977 | 9561994 | - | 4.01584230749346 | 2.27399148412876 | 10.7390538933119 | 5.33554323488568 |
| chr19 | 9562565 | 9562584 | - | 5.01980288436682 | 1.13699574206438 | 8.35259747257593 | 1.06710864697714 |
| chr19 | 9564415 | 9564430 | - | 3.01188173062009 | 3.41098722619315 | 2.38645642073598 | 2.13421729395427 |
| chr19 | 9621485 | 9621568 | + | 53.2099105742883 | 65.9457530397342 | 54.8884976769275 | 51.2212150549025 |
| chr19 | 9632319 | 9632321 | + | 4.01584230749346 | 1.13699574206438 | 3.57968463110397 | 1.06710864697714 |
| chr19 | 9633763 | 9633766 | + | 3.01188173062009 | 1.13699574206438 | 2.38645642073598 | 3.20132594093141 |
| chr19 | 9648326 | 9648337 | - | 1.00396057687336 | 5.68497871032191 | 4.77291284147196 | 6.40265188186281 |
| chr19 | 9651453 | 9651458 | - | 10.0396057687336 | 10.2329616785794 | 16.7051949451519 | 7.46976052883995 |
| chr19 | 9810399 | 9810417 | - | 1.00396057687336 | 6.82197445238629 | 5.96614105183995 | 1.06710864697714 |
| chr19 | 9829979 | 9830058 | + | 70.2772403811355 | 43.2058381984465 | 42.9562155732476 | 80.0331485232852 |
| chr19 | 9830084 | 9830095 | + | 1.00396057687336 | 2.27399148412876 | 5.96614105183995 | 1.06710864697714 |
| chr19 | 9849475 | 9849511 | + | 3.01188173062009 | 3.41098722619315 | 1.19322821036799 | 6.40265188186281 |
| chr19 | 9849534 | 9849577 | + | 17.0673298068472 | 17.0549361309657 | 8.35259747257593 | 16.006629704657 |
| chr19 | 9849599 | 9849664 | + | 85.336649034236 | 69.3567402659273 | 51.3088130458235 | 66.1607361125824 |
| chr19 | 10092257 | 10092334 | + | 8.03168461498692 | 14.780944646837 | 7.15936926220794 | 10.6710864697714 |
| chr19 | 10111209 | 10111260 | + | 2.00792115374673 | 7.95897019445067 | 5.96614105183995 | 8.53686917581709 |
| chr19 | 10115035 | 10115093 | - | 35.1386201905678 | 32.9728765198671 | 10.7390538933119 | 36.2816939972226 |
| chr19 | 10133411 | 10133422 | - | 9.03564519186028 | 21.6029190992233 | 4.77291284147196 | 7.46976052883995 |
| chr19 | 10133472 | 10133498 | - | 3.01188173062009 | 3.41098722619315 | 3.57968463110397 | 2.13421729395427 |
| chr19 | 10133521 | 10133575 | - | 48.1901076899215 | 69.3567402659273 | 33.4103898903037 | 33.0803680562912 |
| chr19 | 10133632 | 10133691 | - | 19.0752509605939 | 19.3289276150945 | 10.7390538933119 | 21.3421729395427 |
| chr19 | 10222229 | 10222230 | - | 4.01584230749346 | 3.41098722619315 | 2.38645642073598 | 3.20132594093141 |
| chr19 | 10258897 | 10258899 | + | 2.00792115374673 | 3.41098722619315 | 1.19322821036799 | 2.13421729395427 |
| chr19 | 10258995 | 10259029 | + | 26.1029749987075 | 12.5069531627082 | 38.1833027317757 | 23.476390233497 |
| chr19 | 10259939 | 10259999 | + | 45.1782259593014 | 53.438799877026 | 48.9223566250876 | 58.6909755837425 |
| chr19 | 10285285 | 10285360 | + | 1.00396057687336 | 1.13699574206438 | 7.15936926220794 | 6.40265188186281 |
| chr19 | 10285506 | 10285733 | + | 22.087132691214 | 25.0139063254164 | 54.8884976769275 | 50.1541064079254 |
| chr19 | 10286491 | 10286525 | + | 1.00396057687336 | 4.54798296825753 | 2.38645642073598 | 5.33554323488568 |
| chr19 | 10286554 | 10286589 | + | 3.01188173062009 | 9.09596593651506 | 1.19322821036799 | 7.46976052883995 |
| chr19 | 10296727 | 10296735 | + | 1.00396057687336 | 1.13699574206438 | 8.35259747257593 | 6.40265188186281 |
| chr19 | 10310239 | 10310273 | - | 6.02376346124019 | 11.3699574206438 | 7.15936926220794 | 13.8724124107028 |
| chr19 | 10310284 | 10310318 | - | 11.043566345607 | 12.5069531627082 | 2.38645642073598 | 3.20132594093141 |
| chr19 | 10315732 | 10315777 | - | 35.1386201905678 | 18.1919318730301 | 4.77291284147196 | 13.8724124107028 |
| chr19 | 10315961 | 10316030 | - | 51.2019894205416 | 67.0827487817985 | 34.6036181006717 | 45.8856718200168 |
| chr19 | 10316246 | 10316315 | - | 6.02376346124019 | 6.82197445238629 | 9.54582568294392 | 19.2079556455884 |
| chr19 | 10316606 | 10316715 | - | 44.174265382428 | 12.5069531627082 | 28.6374770488317 | 45.8856718200168 |
| chr19 | 10316735 | 10316765 | - | 4.01584230749346 | 2.27399148412876 | 14.3187385244159 | 14.9395210576799 |
| chr19 | 10316824 | 10316868 | - | 5.01980288436682 | 1.13699574206438 | 10.7390538933119 | 20.2750642925656 |
| chr19 | 10316903 | 10316956 | - | 16.0633692299738 | 6.82197445238629 | 76.3666054635513 | 14.9395210576799 |
| chr19 | 10317039 | 10317173 | - | 79.3128855729958 | 54.5757956190903 | 156.312895558207 | 113.113516579576 |
| chr19 | 10317196 | 10317215 | - | 11.043566345607 | 6.82197445238629 | 2.38645642073598 | 8.53686917581709 |
| chr19 | 10317251 | 10317291 | - | 7.02772403811355 | 3.41098722619315 | 5.96614105183995 | 3.20132594093141 |
| chr19 | 10317397 | 10317442 | - | 20.0792115374673 | 20.4659233571589 | 15.5119667347839 | 23.476390233497 |
| chr19 | 10333697 | 10333742 | + | 10.0396057687336 | 4.54798296825753 | 15.5119667347839 | 17.0737383516342 |
| chr19 | 10333814 | 10333831 | - | 1.00396057687336 | 4.54798296825753 | 8.35259747257593 | 1.06710864697714 |
| chr19 | 10350554 | 10350580 | - | 11.043566345607 | 15.9179403889013 | 10.7390538933119 | 11.7381951167485 |
| chr19 | 10350602 | 10350657 | - | 19.0752509605939 | 14.780944646837 | 19.0916513658878 | 14.9395210576799 |
| chr19 | 10350739 | 10350760 | - | 9.03564519186028 | 5.68497871032191 | 5.96614105183995 | 6.40265188186281 |
| chr19 | 10391159 | 10391185 | - | 2.00792115374673 | 3.41098722619315 | 7.15936926220794 | 4.26843458790854 |
| chr19 | 10391215 | 10391374 | - | 2820.12526043728 | 1166.55763135806 | 1156.23813584658 | 2361.5114357604 |
| chr19 | 10391399 | 10391491 | - | 22.087132691214 | 10.2329616785794 | 15.5119667347839 | 23.476390233497 |
| chr19 | 10391511 | 10391533 | - | 12.0475269224804 | 5.68497871032191 | 10.7390538933119 | 8.53686917581709 |
| chr19 | 10467802 | 10467850 | + | 3.01188173062009 | 2.27399148412876 | 28.6374770488317 | 36.2816939972226 |
| chr19 | 10486148 | 10486159 | - | 7.02772403811355 | 2.27399148412876 | 14.3187385244159 | 2.13421729395427 |
| chr19 | 10486197 | 10486297 | - | 11.043566345607 | 4.54798296825753 | 8.35259747257593 | 17.0737383516342 |
| chr19 | 10553068 | 10553088 | + | 20.0792115374673 | 25.0139063254164 | 48.9223566250876 | 19.2079556455884 |
| chr19 | 10553212 | 10553233 | + | 2.00792115374673 | 3.41098722619315 | 1.19322821036799 | 11.7381951167485 |
| chr19 | 10553272 | 10553349 | + | 7.02772403811355 | 11.3699574206438 | 9.54582568294392 | 19.2079556455884 |
| chr19 | 10553376 | 10553397 | + | 5.01980288436682 | 19.3289276150945 | 14.3187385244159 | 8.53686917581709 |
| chr19 | 10553929 | 10553946 | - | 35.1386201905678 | 15.9179403889013 | 11.9322821036799 | 18.1408469986113 |
| chr19 | 10566646 | 10566656 | - | 11.043566345607 | 9.09596593651506 | 3.57968463110397 | 9.60397782279422 |
| chr19 | 10566894 | 10566951 | - | 15.0594086531005 | 31.8358807778027 | 8.35259747257593 | 8.53686917581709 |
| chr19 | 10566999 | 10567023 | - | 5.01980288436682 | 9.09596593651506 | 5.96614105183995 | 3.20132594093141 |
| chr19 | 10643408 | 10643443 | + | 8.03168461498692 | 7.95897019445067 | 2.38645642073598 | 9.60397782279422 |
| chr19 | 10643577 | 10643607 | + | 5.01980288436682 | 3.41098722619315 | 1.19322821036799 | 1.06710864697714 |
| chr19 | 10643697 | 10643718 | + | 4.01584230749346 | 4.54798296825753 | 1.19322821036799 | 1.06710864697714 |
| chr19 | 10643797 | 10643850 | + | 12.0475269224804 | 14.780944646837 | 2.38645642073598 | 21.3421729395427 |
| chr19 | 10643883 | 10643898 | + | 3.01188173062009 | 2.27399148412876 | 3.57968463110397 | 2.13421729395427 |
| chr19 | 10643929 | 10643953 | + | 10.0396057687336 | 7.95897019445067 | 2.38645642073598 | 1.06710864697714 |
| chr19 | 10643976 | 10644065 | + | 65.2574374967687 | 46.6168254246397 | 15.5119667347839 | 12.8053037637256 |
| chr19 | 10644511 | 10644514 | + | 11.043566345607 | 17.0549361309657 | 1.19322821036799 | 1.06710864697714 |
| chr19 | 10684414 | 10684416 | + | 1.00396057687336 | 1.13699574206438 | 2.38645642073598 | 1.06710864697714 |
| chr19 | 10684438 | 10684469 | + | 18.0712903837206 | 6.82197445238629 | 14.3187385244159 | 9.60397782279422 |
| chr19 | 10684532 | 10684865 | + | 96.380215379843 | 121.658544400889 | 262.510206280958 | 235.831010981947 |
| chr19 | 10684903 | 10684926 | + | 2.00792115374673 | 4.54798296825753 | 9.54582568294392 | 3.20132594093141 |
| chr19 | 10684946 | 10685010 | + | 23.0910932680874 | 20.4659233571589 | 32.2171616799357 | 35.2145853502455 |
| chr19 | 10685057 | 10685090 | + | 15.0594086531005 | 7.95897019445067 | 14.3187385244159 | 10.6710864697714 |
| chr19 | 10685099 | 10685112 | + | 1.00396057687336 | 1.13699574206438 | 4.77291284147196 | 1.06710864697714 |
| chr19 | 10685118 | 10685180 | + | 22.087132691214 | 14.780944646837 | 20.2848795762558 | 32.0132594093141 |
| chr19 | 10685209 | 10685295 | + | 39.1544624980612 | 31.8358807778027 | 56.0817258872955 | 110.979299285622 |
| chr19 | 10685354 | 10685387 | + | 24.0950538449607 | 14.780944646837 | 16.7051949451519 | 17.0737383516342 |
| chr19 | 10685448 | 10685506 | + | 2.00792115374673 | 2.27399148412876 | 4.77291284147196 | 4.26843458790854 |
| chr19 | 10685544 | 10685632 | + | 29.1148567293276 | 47.753821166704 | 17.8984231555198 | 19.2079556455884 |
| chr19 | 10685662 | 10685726 | + | 24.0950538449607 | 22.7399148412876 | 20.2848795762558 | 17.0737383516342 |
| chr19 | 10689320 | 10689326 | + | 2.00792115374673 | 9.09596593651506 | 2.38645642073598 | 1.06710864697714 |
| chr19 | 10689530 | 10689554 | + | 7.02772403811355 | 7.95897019445067 | 8.35259747257593 | 3.20132594093141 |
| chr19 | 10689641 | 10689687 | + | 12.0475269224804 | 11.3699574206438 | 16.7051949451519 | 14.9395210576799 |
| chr19 | 10689750 | 10689824 | + | 13.0514874993537 | 9.09596593651506 | 14.3187385244159 | 17.0737383516342 |
| chr19 | 10690184 | 10690202 | + | 5.01980288436682 | 1.13699574206438 | 7.15936926220794 | 1.06710864697714 |
| chr19 | 10690330 | 10690349 | + | 14.0554480762271 | 13.6439489047726 | 4.77291284147196 | 5.33554323488568 |
| chr19 | 10690375 | 10690415 | + | 5.01980288436682 | 4.54798296825753 | 5.96614105183995 | 13.8724124107028 |
| chr19 | 10690437 | 10690482 | + | 8.03168461498692 | 4.54798296825753 | 2.38645642073598 | 5.33554323488568 |
| chr19 | 10690538 | 10690572 | + | 10.0396057687336 | 6.82197445238629 | 2.38645642073598 | 6.40265188186281 |
| chr19 | 10690617 | 10690650 | + | 9.03564519186028 | 5.68497871032191 | 4.77291284147196 | 12.8053037637256 |
| chr19 | 10690810 | 10690868 | + | 18.0712903837206 | 10.2329616785794 | 13.1255103140479 | 1.06710864697714 |
| chr19 | 10690900 | 10690926 | + | 6.02376346124019 | 5.68497871032191 | 3.57968463110397 | 3.20132594093141 |
| chr19 | 10690937 | 10690958 | + | 3.01188173062009 | 6.82197445238629 | 3.57968463110397 | 3.20132594093141 |
| chr19 | 10690990 | 10691014 | + | 11.043566345607 | 11.3699574206438 | 19.0916513658878 | 21.3421729395427 |
| chr19 | 10691074 | 10691083 | + | 7.02772403811355 | 2.27399148412876 | 4.77291284147196 | 5.33554323488568 |
| chr19 | 10691154 | 10691163 | + | 2.00792115374673 | 2.27399148412876 | 4.77291284147196 | 2.13421729395427 |
| chr19 | 10691698 | 10691785 | + | 42.1663442286813 | 42.0688424563821 | 45.3426719939836 | 46.952780466994 |
| chr19 | 10691805 | 10692001 | + | 90.3564519186028 | 63.6717615556054 | 75.1733772531833 | 86.435800405148 |
| chr19 | 10692022 | 10692139 | + | 75.2970432655023 | 52.3018041349616 | 51.3088130458235 | 55.4896496428111 |
| chr19 | 10692146 | 10692241 | + | 96.380215379843 | 52.3018041349616 | 40.5697591525116 | 74.6976052883995 |
| chr19 | 10692343 | 10692359 | + | 24.0950538449607 | 20.4659233571589 | 13.1255103140479 | 22.4092815865198 |
| chr19 | 10713288 | 10713331 | + | 27.1069355755808 | 35.2468680039958 | 28.6374770488317 | 24.5434988804741 |
| chr19 | 10831250 | 10831270 | + | 40.1584230749346 | 13.6439489047726 | 66.8207797806074 | 10.6710864697714 |
| chr19 | 10831300 | 10831316 | + | 4.01584230749346 | 4.54798296825753 | 38.1833027317757 | 7.46976052883995 |
| chr19 | 10831334 | 10831568 | + | 83.3287278804893 | 56.8497871032191 | 180.177459765566 | 103.509538756782 |
| chr19 | 10831613 | 10831617 | + | 1.00396057687336 | 2.27399148412876 | 1.19322821036799 | 22.4092815865198 |
| chr19 | 10831648 | 10831668 | + | 5.01980288436682 | 1.13699574206438 | 11.9322821036799 | 4.26843458790854 |
| chr19 | 10831687 | 10831748 | + | 22.087132691214 | 13.6439489047726 | 25.0577924177278 | 34.1474767032683 |
| chr19 | 10831778 | 10831864 | + | 368.453531712525 | 271.741982353387 | 403.31113510438 | 416.172372321083 |
| chr19 | 10832462 | 10832538 | - | 44.174265382428 | 88.6856678810218 | 32.2171616799357 | 25.6106075274513 |
| chr19 | 10921821 | 10921916 | + | 23.0910932680874 | 15.9179403889013 | 42.9562155732476 | 25.6106075274513 |
| chr19 | 10921943 | 10921980 | + | 4.01584230749346 | 6.82197445238629 | 10.7390538933119 | 9.60397782279422 |
| chr19 | 10922119 | 10922192 | + | 33.130699036821 | 25.0139063254164 | 48.9223566250876 | 80.0331485232852 |
| chr19 | 10922231 | 10922268 | - | 10.0396057687336 | 22.7399148412876 | 23.8645642073598 | 27.7448248214055 |
| chr19 | 10922267 | 10922402 | + | 24.0950538449607 | 14.780944646837 | 48.9223566250876 | 60.8251928776967 |
| chr19 | 10922355 | 10922436 | - | 9.03564519186028 | 10.2329616785794 | 38.1833027317757 | 55.4896496428111 |
| chr19 | 10922448 | 10922471 | - | 32.1267384599477 | 21.6029190992233 | 38.1833027317757 | 49.0869977609482 |
| chr19 | 10922593 | 10922637 | - | 5.01980288436682 | 3.41098722619315 | 15.5119667347839 | 20.2750642925656 |
| chr19 | 10922690 | 10922708 | + | 13.0514874993537 | 9.09596593651506 | 16.7051949451519 | 16.006629704657 |
| chr19 | 10922759 | 10922821 | - | 20.0792115374673 | 28.4248935516096 | 36.9900745214077 | 29.8790421153598 |
| chr19 | 10922953 | 10922967 | + | 8.03168461498692 | 5.68497871032191 | 2.38645642073598 | 2.13421729395427 |
| chr19 | 10922995 | 10923011 | + | 2.00792115374673 | 1.13699574206438 | 1.19322821036799 | 1.06710864697714 |
| chr19 | 10923028 | 10923041 | - | 1.00396057687336 | 1.13699574206438 | 2.38645642073598 | 4.26843458790854 |
| chr19 | 10923044 | 10923066 | - | 2.00792115374673 | 6.82197445238629 | 4.77291284147196 | 6.40265188186281 |
| chr19 | 10923160 | 10923166 | - | 1.00396057687336 | 1.13699574206438 | 1.19322821036799 | 1.06710864697714 |
| chr19 | 10929830 | 10929834 | + | 2.00792115374673 | 1.13699574206438 | 1.19322821036799 | 1.06710864697714 |
| chr19 | 10929898 | 10929907 | + | 3.01188173062009 | 5.68497871032191 | 1.19322821036799 | 1.06710864697714 |
| chr19 | 10930157 | 10930167 | + | 1.00396057687336 | 3.41098722619315 | 1.19322821036799 | 1.06710864697714 |
| chr19 | 11061841 | 11061917 | + | 23.0910932680874 | 13.6439489047726 | 32.2171616799357 | 43.7514545260626 |
| chr19 | 11062044 | 11062065 | + | 5.01980288436682 | 3.41098722619315 | 3.57968463110397 | 5.33554323488568 |
| chr19 | 11062207 | 11062242 | + | 41.1623836518079 | 42.0688424563821 | 42.9562155732476 | 62.959410171651 |
| chr19 | 11131541 | 11131544 | + | 2.00792115374673 | 6.82197445238629 | 8.35259747257593 | 3.20132594093141 |
| chr19 | 11145586 | 11145607 | - | 113.44754518669 | 110.288586980245 | 51.3088130458235 | 86.435800405148 |
| chr19 | 11145630 | 11145687 | - | 14.0554480762271 | 13.6439489047726 | 19.0916513658878 | 19.2079556455884 |
| chr19 | 11145728 | 11145733 | - | 3.01188173062009 | 4.54798296825753 | 1.19322821036799 | 5.33554323488568 |
| chr19 | 11145786 | 11145834 | - | 9.03564519186028 | 15.9179403889013 | 9.54582568294392 | 3.20132594093141 |
| chr19 | 11145874 | 11145897 | - | 20.0792115374673 | 15.9179403889013 | 23.8645642073598 | 11.7381951167485 |
| chr19 | 11146254 | 11146371 | - | 56.2217923049084 | 61.3977700714766 | 89.4921157775992 | 96.0397782279422 |
| chr19 | 11146392 | 11146414 | - | 13.0514874993537 | 9.09596593651506 | 25.0577924177278 | 16.006629704657 |
| chr19 | 11146794 | 11146813 | - | 16.0633692299738 | 14.780944646837 | 2.38645642073598 | 20.2750642925656 |
| chr19 | 11147139 | 11147170 | - | 6.02376346124019 | 3.41098722619315 | 4.77291284147196 | 8.53686917581709 |
| chr19 | 11164372 | 11164373 | - | 5.01980288436682 | 6.82197445238629 | 2.38645642073598 | 2.13421729395427 |
| chr19 | 11322372 | 11322403 | - | 21.0831721143407 | 7.95897019445067 | 1.19322821036799 | 4.26843458790854 |
| chr19 | 11324283 | 11324310 | - | 7.02772403811355 | 4.54798296825753 | 2.38645642073598 | 3.20132594093141 |
| chr19 | 11342798 | 11342877 | - | 6.02376346124019 | 7.95897019445067 | 10.7390538933119 | 19.2079556455884 |
| chr19 | 11364597 | 11364616 | + | 4.01584230749346 | 2.27399148412876 | 4.77291284147196 | 3.20132594093141 |
| chr19 | 11364641 | 11364643 | + | 6.02376346124019 | 3.41098722619315 | 3.57968463110397 | 2.13421729395427 |
| chr19 | 11376024 | 11376029 | + | 2.00792115374673 | 3.41098722619315 | 2.38645642073598 | 2.13421729395427 |
| chr19 | 11377880 | 11377900 | - | 4.01584230749346 | 2.27399148412876 | 1.19322821036799 | 8.53686917581709 |
| chr19 | 11450858 | 11450934 | + | 86.3406096111093 | 81.8636934286355 | 34.6036181006717 | 40.5501285851312 |
| chr19 | 11453881 | 11453883 | - | 29.1148567293276 | 20.4659233571589 | 41.7629873628796 | 16.006629704657 |
| chr19 | 11505945 | 11505955 | - | 21.0831721143407 | 13.6439489047726 | 7.15936926220794 | 20.2750642925656 |
| chr19 | 11553083 | 11553111 | - | 25.0990144218341 | 28.4248935516096 | 11.9322821036799 | 27.7448248214055 |
| chr19 | 11553647 | 11553669 | - | 3.01188173062009 | 4.54798296825753 | 4.77291284147196 | 5.33554323488568 |
| chr19 | 11617101 | 11617106 | + | 2.00792115374673 | 1.13699574206438 | 1.19322821036799 | 1.06710864697714 |
| chr19 | 11639727 | 11639824 | + | 23.0910932680874 | 35.2468680039958 | 64.4343233598714 | 66.1607361125824 |
| chr19 | 12430042 | 12430049 | - | 5.01980288436682 | 1.13699574206438 | 3.57968463110397 | 3.20132594093141 |
| chr19 | 12646569 | 12646578 | - | 20.0792115374673 | 19.3289276150945 | 19.0916513658878 | 30.9461507623369 |
| chr19 | 12646605 | 12646613 | - | 3.01188173062009 | 2.27399148412876 | 2.38645642073598 | 4.26843458790854 |
| chr19 | 12668093 | 12668158 | - | 120.475269224804 | 178.508331504108 | 56.0817258872955 | 88.5700176991023 |
| chr19 | 12668183 | 12668290 | - | 151.598047107878 | 119.38455291676 | 89.4921157775992 | 104.576647403759 |
| chr19 | 12668333 | 12668372 | - | 5.01980288436682 | 10.2329616785794 | 14.3187385244159 | 6.40265188186281 |
| chr19 | 12675735 | 12675736 | - | 1.00396057687336 | 2.27399148412876 | 2.38645642073598 | 1.06710864697714 |
| chr19 | 12688946 | 12688953 | - | 2.00792115374673 | 3.41098722619315 | 1.19322821036799 | 2.13421729395427 |
| chr19 | 12699227 | 12699280 | - | 22.087132691214 | 30.6988850357383 | 10.7390538933119 | 19.2079556455884 |
| chr19 | 12699301 | 12699313 | - | 9.03564519186028 | 4.54798296825753 | 5.96614105183995 | 7.46976052883995 |
| chr19 | 12699496 | 12699511 | - | 8.03168461498692 | 5.68497871032191 | 5.96614105183995 | 8.53686917581709 |
| chr19 | 12699680 | 12699731 | - | 29.1148567293276 | 17.0549361309657 | 20.2848795762558 | 26.6777161744284 |
| chr19 | 12699759 | 12699776 | - | 5.01980288436682 | 5.68497871032191 | 14.3187385244159 | 9.60397782279422 |
| chr19 | 12699811 | 12699833 | - | 2.00792115374673 | 3.41098722619315 | 5.96614105183995 | 6.40265188186281 |
| chr19 | 12699854 | 12699871 | - | 17.0673298068472 | 12.5069531627082 | 5.96614105183995 | 10.6710864697714 |
| chr19 | 12699944 | 12700050 | - | 62.2455557661486 | 44.3428339405109 | 50.1155848354556 | 65.0936274656053 |
| chr19 | 12700159 | 12700176 | - | 11.043566345607 | 4.54798296825753 | 4.77291284147196 | 6.40265188186281 |
| chr19 | 12700217 | 12700239 | - | 9.03564519186028 | 7.95897019445067 | 7.15936926220794 | 5.33554323488568 |
| chr19 | 12700345 | 12700366 | - | 4.01584230749346 | 2.27399148412876 | 4.77291284147196 | 2.13421729395427 |
| chr19 | 12700399 | 12700415 | - | 4.01584230749346 | 4.54798296825753 | 7.15936926220794 | 2.13421729395427 |
| chr19 | 12700461 | 12700466 | - | 5.01980288436682 | 2.27399148412876 | 4.77291284147196 | 2.13421729395427 |
| chr19 | 12700521 | 12700543 | - | 3.01188173062009 | 5.68497871032191 | 3.57968463110397 | 5.33554323488568 |
| chr19 | 12700598 | 12700664 | - | 196.776273067179 | 115.973565690567 | 48.9223566250876 | 34.1474767032683 |
| chr19 | 12700814 | 12700838 | - | 10.0396057687336 | 3.41098722619315 | 1.19322821036799 | 1.06710864697714 |
| chr19 | 12700873 | 12700910 | - | 23.0910932680874 | 30.6988850357383 | 23.8645642073598 | 20.2750642925656 |
| chr19 | 12700941 | 12700958 | - | 10.0396057687336 | 23.876910583352 | 10.7390538933119 | 9.60397782279422 |
| chr19 | 12701004 | 12701042 | - | 27.1069355755808 | 15.9179403889013 | 13.1255103140479 | 4.26843458790854 |
| chr19 | 12701068 | 12701075 | - | 12.0475269224804 | 6.82197445238629 | 10.7390538933119 | 2.13421729395427 |
| chr19 | 12701144 | 12701189 | - | 13.0514874993537 | 21.6029190992233 | 5.96614105183995 | 4.26843458790854 |
| chr19 | 12701334 | 12701365 | - | 11.043566345607 | 11.3699574206438 | 9.54582568294392 | 4.26843458790854 |
| chr19 | 12730659 | 12730960 | - | 21.0831721143407 | 21.6029190992233 | 21.4781077866238 | 24.5434988804741 |
| chr19 | 12730993 | 12731043 | - | 5.01980288436682 | 2.27399148412876 | 3.57968463110397 | 3.20132594093141 |
| chr19 | 12748258 | 12748296 | + | 53.2099105742883 | 32.9728765198671 | 9.54582568294392 | 25.6106075274513 |
| chr19 | 12763125 | 12763134 | - | 5.01980288436682 | 12.5069531627082 | 1.19322821036799 | 5.33554323488568 |
| chr19 | 12792806 | 12792989 | + | 57.2257528817818 | 53.438799877026 | 14.3187385244159 | 39.483019938154 |
| chr19 | 12793155 | 12793259 | + | 34.1346596136944 | 77.315710460378 | 9.54582568294392 | 19.2079556455884 |
| chr19 | 12796844 | 12796875 | - | 33.130699036821 | 52.3018041349616 | 14.3187385244159 | 50.1541064079254 |
| chr19 | 12796888 | 12796934 | - | 8.03168461498692 | 9.09596593651506 | 2.38645642073598 | 8.53686917581709 |
| chr19 | 12797107 | 12797151 | - | 4.01584230749346 | 3.41098722619315 | 2.38645642073598 | 11.7381951167485 |
| chr19 | 12813479 | 12813499 | + | 14.0554480762271 | 18.1919318730301 | 7.15936926220794 | 7.46976052883995 |
| chr19 | 12813565 | 12813594 | + | 19.0752509605939 | 45.4798296825753 | 13.1255103140479 | 17.0737383516342 |
| chr19 | 12875479 | 12875480 | - | 4.01584230749346 | 3.41098722619315 | 3.57968463110397 | 2.13421729395427 |
| chr19 | 12875585 | 12875602 | - | 6.02376346124019 | 13.6439489047726 | 3.57968463110397 | 3.20132594093141 |
| chr19 | 12899554 | 12899557 | + | 3.01188173062009 | 1.13699574206438 | 2.38645642073598 | 1.06710864697714 |
| chr19 | 12899902 | 12899905 | + | 9.03564519186028 | 3.41098722619315 | 3.57968463110397 | 3.20132594093141 |
| chr19 | 12899939 | 12899946 | + | 15.0594086531005 | 18.1919318730301 | 11.9322821036799 | 14.9395210576799 |
| chr19 | 12922493 | 12922533 | - | 5.01980288436682 | 1.13699574206438 | 2.38645642073598 | 6.40265188186281 |
| chr19 | 12922743 | 12922760 | - | 6.02376346124019 | 1.13699574206438 | 7.15936926220794 | 18.1408469986113 |
| chr19 | 12940538 | 12940652 | + | 378.493137481258 | 296.755888678804 | 257.737293439486 | 243.300771510787 |
| chr19 | 12943692 | 12944000 | + | 1049.13880283267 | 750.417189762492 | 914.01280914188 | 797.13015929192 |
| chr19 | 12944026 | 12944046 | + | 12.0475269224804 | 2.27399148412876 | 5.96614105183995 | 4.26843458790854 |
| chr19 | 12944106 | 12944188 | + | 10.0396057687336 | 6.82197445238629 | 16.7051949451519 | 9.60397782279422 |
| chr19 | 12944228 | 12944253 | + | 7.02772403811355 | 3.41098722619315 | 7.15936926220794 | 10.6710864697714 |
| chr19 | 12944302 | 12944341 | + | 5.01980288436682 | 2.27399148412876 | 3.57968463110397 | 5.33554323488568 |
| chr19 | 12944394 | 12944427 | + | 65.2574374967687 | 51.1648083928972 | 29.8307052591997 | 59.7580842307196 |
| chr19 | 12953055 | 12953089 | + | 3.01188173062009 | 4.54798296825753 | 3.57968463110397 | 9.60397782279422 |
| chr19 | 12953147 | 12953150 | + | 5.01980288436682 | 4.54798296825753 | 1.19322821036799 | 1.06710864697714 |
| chr19 | 12953188 | 12953203 | - | 2.00792115374673 | 2.27399148412876 | 1.19322821036799 | 3.20132594093141 |
| chr19 | 12953420 | 12953438 | + | 1.00396057687336 | 3.41098722619315 | 2.38645642073598 | 1.06710864697714 |
| chr19 | 12953465 | 12953502 | + | 1.00396057687336 | 4.54798296825753 | 1.19322821036799 | 3.20132594093141 |
| chr19 | 12953541 | 12953621 | + | 76.3010038423757 | 110.288586980245 | 62.0478669391355 | 66.1607361125824 |
| chr19 | 12953631 | 12953642 | - | 6.02376346124019 | 39.7948509722534 | 14.3187385244159 | 2.13421729395427 |
| chr19 | 12954179 | 12954529 | - | 344.358477867564 | 221.714169702554 | 424.789242891004 | 425.776350143877 |
| chr19 | 13096863 | 13096908 | + | 5.01980288436682 | 3.41098722619315 | 9.54582568294392 | 23.476390233497 |
| chr19 | 13097103 | 13097159 | + | 3.01188173062009 | 4.54798296825753 | 4.77291284147196 | 9.60397782279422 |
| chr19 | 13104920 | 13104981 | - | 9.03564519186028 | 17.0549361309657 | 14.3187385244159 | 26.6777161744284 |
| chr19 | 13139096 | 13139121 | + | 15.0594086531005 | 10.2329616785794 | 1.19322821036799 | 2.13421729395427 |
| chr19 | 13139472 | 13139539 | + | 45.1782259593014 | 26.1509020674808 | 4.77291284147196 | 2.13421729395427 |
| chr19 | 13139718 | 13139734 | + | 19.0752509605939 | 5.68497871032191 | 11.9322821036799 | 25.6106075274513 |
| chr19 | 13139801 | 13139819 | + | 5.01980288436682 | 10.2329616785794 | 1.19322821036799 | 2.13421729395427 |
| chr19 | 13140206 | 13140209 | + | 7.02772403811355 | 3.41098722619315 | 1.19322821036799 | 1.06710864697714 |
| chr19 | 13140428 | 13140448 | + | 32.1267384599477 | 6.82197445238629 | 21.4781077866238 | 9.60397782279422 |
| chr19 | 13140631 | 13140639 | + | 7.02772403811355 | 1.13699574206438 | 3.57968463110397 | 1.06710864697714 |
| chr19 | 13140661 | 13140734 | + | 76.3010038423757 | 30.6988850357383 | 64.4343233598714 | 60.8251928776967 |
| chr19 | 13141080 | 13141121 | + | 26.1029749987075 | 30.6988850357383 | 14.3187385244159 | 18.1408469986113 |
| chr19 | 13144078 | 13144135 | - | 23.0910932680874 | 12.5069531627082 | 7.15936926220794 | 6.40265188186281 |
| chr19 | 13144162 | 13144187 | - | 1.00396057687336 | 13.6439489047726 | 2.38645642073598 | 3.20132594093141 |
| chr19 | 13153892 | 13153930 | + | 7.02772403811355 | 5.68497871032191 | 20.2848795762558 | 7.46976052883995 |
| chr19 | 13153952 | 13153955 | + | 1.00396057687336 | 1.13699574206438 | 1.19322821036799 | 1.06710864697714 |
| chr19 | 13153994 | 13154205 | + | 117.463387494184 | 89.8226636230862 | 180.177459765566 | 160.06629704657 |
| chr19 | 13154233 | 13154291 | + | 12.0475269224804 | 17.0549361309657 | 44.1494437836156 | 115.247733873531 |
| chr19 | 13154313 | 13154317 | + | 4.01584230749346 | 3.41098722619315 | 4.77291284147196 | 4.26843458790854 |
| chr19 | 13154349 | 13154486 | + | 112.443584609817 | 101.19262104373 | 95.4582568294392 | 199.549316984724 |
| chr19 | 13154546 | 13154560 | + | 1.00396057687336 | 3.41098722619315 | 2.38645642073598 | 3.20132594093141 |
| chr19 | 13154683 | 13154707 | + | 5.01980288436682 | 9.09596593651506 | 3.57968463110397 | 6.40265188186281 |
| chr19 | 13154785 | 13154805 | + | 42.1663442286813 | 72.7677274921204 | 8.35259747257593 | 34.1474767032683 |
| chr19 | 13206981 | 13206986 | - | 2.00792115374673 | 4.54798296825753 | 3.57968463110397 | 3.20132594093141 |
| chr19 | 13763242 | 13763270 | + | 3.01188173062009 | 12.5069531627082 | 4.77291284147196 | 2.13421729395427 |
| chr19 | 13774053 | 13774064 | + | 8.03168461498692 | 3.41098722619315 | 1.19322821036799 | 3.20132594093141 |
| chr19 | 13774171 | 13774179 | + | 3.01188173062009 | 5.68497871032191 | 2.38645642073598 | 3.20132594093141 |
| chr19 | 13778333 | 13778438 | + | 29.1148567293276 | 23.876910583352 | 17.8984231555198 | 29.8790421153598 |
| chr19 | 13832104 | 13832106 | + | 1.00396057687336 | 1.13699574206438 | 3.57968463110397 | 3.20132594093141 |
| chr19 | 13832185 | 13832220 | + | 2.00792115374673 | 3.41098722619315 | 2.38645642073598 | 6.40265188186281 |
| chr19 | 13961384 | 13961395 | + | 7.02772403811355 | 5.68497871032191 | 8.35259747257593 | 19.2079556455884 |
| chr19 | 13961591 | 13961599 | - | 3.01188173062009 | 1.13699574206438 | 5.96614105183995 | 5.33554323488568 |
| chr19 | 13961629 | 13961657 | - | 4.01584230749346 | 1.13699574206438 | 4.77291284147196 | 4.26843458790854 |
| chr19 | 14052634 | 14052642 | + | 2.00792115374673 | 3.41098722619315 | 3.57968463110397 | 11.7381951167485 |
| chr19 | 14052674 | 14052675 | + | 1.00396057687336 | 3.41098722619315 | 1.19322821036799 | 5.33554323488568 |
| chr19 | 14053121 | 14053131 | + | 2.00792115374673 | 7.95897019445067 | 1.19322821036799 | 10.6710864697714 |
| chr19 | 14087960 | 14087988 | - | 66.2613980736421 | 70.4937360079917 | 10.7390538933119 | 40.5501285851312 |
| chr19 | 14088222 | 14088223 | - | 3.01188173062009 | 3.41098722619315 | 3.57968463110397 | 3.20132594093141 |
| chr19 | 14091718 | 14091802 | - | 43.1703048055547 | 47.753821166704 | 17.8984231555198 | 38.4159112911769 |
| chr19 | 14092122 | 14092127 | - | 3.01188173062009 | 2.27399148412876 | 11.9322821036799 | 1.06710864697714 |
| chr19 | 14092146 | 14092185 | - | 1.00396057687336 | 1.13699574206438 | 5.96614105183995 | 9.60397782279422 |
| chr19 | 14092227 | 14092264 | - | 36.1425807674411 | 31.8358807778027 | 34.6036181006717 | 25.6106075274513 |
| chr19 | 14092319 | 14092365 | - | 5.01980288436682 | 11.3699574206438 | 20.2848795762558 | 8.53686917581709 |
| chr19 | 14092417 | 14092464 | - | 18.0712903837206 | 15.9179403889013 | 42.9562155732476 | 13.8724124107028 |
| chr19 | 14092592 | 14092648 | - | 24.0950538449607 | 21.6029190992233 | 25.0577924177278 | 14.9395210576799 |
| chr19 | 14093093 | 14093095 | - | 1.00396057687336 | 1.13699574206438 | 1.19322821036799 | 10.6710864697714 |
| chr19 | 14119537 | 14119543 | - | 18.0712903837206 | 34.1098722619315 | 3.57968463110397 | 6.40265188186281 |
| chr19 | 14119578 | 14119669 | - | 77.3049644192491 | 84.1376849127643 | 25.0577924177278 | 30.9461507623369 |
| chr19 | 14119982 | 14120005 | - | 13.0514874993537 | 18.1919318730301 | 3.57968463110397 | 10.6710864697714 |
| chr19 | 14120174 | 14120194 | - | 6.02376346124019 | 9.09596593651506 | 4.77291284147196 | 3.20132594093141 |
| chr19 | 14120223 | 14120320 | - | 50.1980288436682 | 44.3428339405109 | 35.7968463110397 | 55.4896496428111 |
| chr19 | 14120350 | 14120394 | - | 49.1940682667949 | 27.2878978095452 | 22.6713359969918 | 21.3421729395427 |
| chr19 | 14120421 | 14120447 | - | 9.03564519186028 | 20.4659233571589 | 11.9322821036799 | 8.53686917581709 |
| chr19 | 14137134 | 14137208 | + | 95.3762548029696 | 93.2336508492793 | 26.2510206280958 | 49.0869977609482 |
| chr19 | 14148742 | 14148763 | - | 1.00396057687336 | 1.13699574206438 | 4.77291284147196 | 7.46976052883995 |
| chr19 | 14149465 | 14149466 | - | 5.01980288436682 | 1.13699574206438 | 1.19322821036799 | 1.06710864697714 |
| chr19 | 14150017 | 14150099 | - | 47.1861471130481 | 32.9728765198671 | 100.231169670911 | 82.1673658172394 |
| chr19 | 14170170 | 14170174 | + | 3.01188173062009 | 5.68497871032191 | 3.57968463110397 | 2.13421729395427 |
| chr19 | 14170849 | 14170863 | + | 12.0475269224804 | 15.9179403889013 | 8.35259747257593 | 6.40265188186281 |
| chr19 | 14171019 | 14171038 | + | 13.0514874993537 | 7.95897019445067 | 2.38645642073598 | 5.33554323488568 |
| chr19 | 14408162 | 14408181 | + | 9.03564519186028 | 2.27399148412876 | 5.96614105183995 | 7.46976052883995 |
| chr19 | 14408615 | 14408652 | + | 68.2693192273888 | 85.2746806548286 | 15.5119667347839 | 14.9395210576799 |
| chr19 | 14408839 | 14408873 | - | 53.2099105742883 | 60.2607743294122 | 39.3765309421437 | 58.6909755837425 |
| chr19 | 14408897 | 14408910 | - | 11.043566345607 | 2.27399148412876 | 2.38645642073598 | 14.9395210576799 |
| chr19 | 14477782 | 14477859 | - | 1619.38841049674 | 1343.9289671201 | 486.83710983014 | 478.064673845757 |
| chr19 | 14477895 | 14477908 | - | 7.02772403811355 | 3.41098722619315 | 1.19322821036799 | 2.13421729395427 |
| chr19 | 14477933 | 14477961 | - | 29.1148567293276 | 27.2878978095452 | 7.15936926220794 | 11.7381951167485 |
| chr19 | 14478002 | 14478105 | - | 106.419821148577 | 101.19262104373 | 36.9900745214077 | 49.0869977609482 |
| chr19 | 14478127 | 14478159 | - | 28.1108961524542 | 14.780944646837 | 2.38645642073598 | 11.7381951167485 |
| chr19 | 14478262 | 14478310 | - | 14.0554480762271 | 2.27399148412876 | 4.77291284147196 | 3.20132594093141 |
| chr19 | 14478323 | 14478359 | - | 8.03168461498692 | 7.95897019445067 | 4.77291284147196 | 4.26843458790854 |
| chr19 | 14514802 | 14514830 | - | 11.043566345607 | 7.95897019445067 | 13.1255103140479 | 10.6710864697714 |
| chr19 | 14514871 | 14514887 | - | 6.02376346124019 | 11.3699574206438 | 5.96614105183995 | 10.6710864697714 |
| chr19 | 14514917 | 14514927 | - | 7.02772403811355 | 4.54798296825753 | 1.19322821036799 | 7.46976052883995 |
| chr19 | 14515277 | 14515279 | - | 1.00396057687336 | 1.13699574206438 | 2.38645642073598 | 1.06710864697714 |
| chr19 | 14515833 | 14515855 | - | 20.0792115374673 | 11.3699574206438 | 5.96614105183995 | 6.40265188186281 |
| chr19 | 14515883 | 14515924 | - | 20.0792115374673 | 5.68497871032191 | 14.3187385244159 | 14.9395210576799 |
| chr19 | 14529531 | 14529701 | + | 25645.1689756532 | 18763.8407312885 | 5608.17258872955 | 8065.20715385319 |
| chr19 | 14566101 | 14566163 | - | 164.649534607232 | 153.494425178692 | 81.1395183050233 | 172.871600810296 |
| chr19 | 14733616 | 14733634 | - | 1.00396057687336 | 1.13699574206438 | 4.77291284147196 | 4.26843458790854 |
| chr19 | 15114995 | 15115123 | - | 14.0554480762271 | 23.876910583352 | 1.19322821036799 | 5.33554323488568 |
| chr19 | 15236860 | 15236922 | - | 7.02772403811355 | 19.3289276150945 | 4.77291284147196 | 3.20132594093141 |
| chr19 | 15237051 | 15237052 | - | 1.00396057687336 | 1.13699574206438 | 1.19322821036799 | 1.06710864697714 |
| chr19 | 15237203 | 15237238 | - | 5.01980288436682 | 10.2329616785794 | 13.1255103140479 | 73.6304966414224 |
| chr19 | 15237408 | 15237427 | - | 2.00792115374673 | 3.41098722619315 | 1.19322821036799 | 2.13421729395427 |
| chr19 | 15237464 | 15237480 | - | 1.00396057687336 | 2.27399148412876 | 1.19322821036799 | 1.06710864697714 |
| chr19 | 15237647 | 15237735 | - | 65.2574374967687 | 38.657855230189 | 40.5697591525116 | 152.59653651773 |
| chr19 | 15237757 | 15237781 | - | 2.00792115374673 | 1.13699574206438 | 1.19322821036799 | 1.06710864697714 |
| chr19 | 15247176 | 15247192 | - | 15.0594086531005 | 29.5618892936739 | 5.96614105183995 | 10.6710864697714 |
| chr19 | 15247246 | 15247248 | - | 4.01584230749346 | 6.82197445238629 | 1.19322821036799 | 2.13421729395427 |
| chr19 | 15247907 | 15247908 | - | 3.01188173062009 | 2.27399148412876 | 1.19322821036799 | 1.06710864697714 |
| chr19 | 15248108 | 15248121 | - | 57.2257528817818 | 64.8087572976698 | 40.5697591525116 | 9.60397782279422 |
| chr19 | 15248260 | 15248279 | - | 9.03564519186028 | 10.2329616785794 | 3.57968463110397 | 3.20132594093141 |
| chr19 | 15253236 | 15253237 | - | 13.0514874993537 | 5.68497871032191 | 20.2848795762558 | 86.435800405148 |
| chr19 | 15353582 | 15353627 | - | 8.03168461498692 | 2.27399148412876 | 8.35259747257593 | 9.60397782279422 |
| chr19 | 15353808 | 15353809 | - | 3.01188173062009 | 2.27399148412876 | 2.38645642073598 | 1.06710864697714 |
| chr19 | 15353843 | 15353865 | - | 7.02772403811355 | 3.41098722619315 | 4.77291284147196 | 3.20132594093141 |
| chr19 | 15353989 | 15354000 | - | 6.02376346124019 | 4.54798296825753 | 2.38645642073598 | 1.06710864697714 |
| chr19 | 15354135 | 15354143 | - | 5.01980288436682 | 4.54798296825753 | 4.77291284147196 | 5.33554323488568 |
| chr19 | 15354624 | 15354647 | - | 11.043566345607 | 9.09596593651506 | 4.77291284147196 | 5.33554323488568 |
| chr19 | 15354734 | 15354757 | - | 6.02376346124019 | 4.54798296825753 | 1.19322821036799 | 4.26843458790854 |
| chr19 | 15354870 | 15354890 | - | 6.02376346124019 | 3.41098722619315 | 1.19322821036799 | 1.06710864697714 |
| chr19 | 15380081 | 15380108 | - | 15.0594086531005 | 11.3699574206438 | 14.3187385244159 | 20.2750642925656 |
| chr19 | 15422186 | 15422191 | - | 1.00396057687336 | 7.95897019445067 | 4.77291284147196 | 8.53686917581709 |
| chr19 | 15422345 | 15422351 | - | 17.0673298068472 | 22.7399148412876 | 13.1255103140479 | 17.0737383516342 |
| chr19 | 15422748 | 15422751 | - | 2.00792115374673 | 3.41098722619315 | 13.1255103140479 | 2.13421729395427 |
| chr19 | 15422781 | 15422799 | - | 21.0831721143407 | 10.2329616785794 | 25.0577924177278 | 7.46976052883995 |
| chr19 | 15422864 | 15422885 | - | 5.01980288436682 | 6.82197445238629 | 8.35259747257593 | 6.40265188186281 |
| chr19 | 15422911 | 15422943 | - | 12.0475269224804 | 5.68497871032191 | 20.2848795762558 | 6.40265188186281 |
| chr19 | 15828975 | 15829049 | + | 235.930735565241 | 297.892884420868 | 29.8307052591997 | 23.476390233497 |
| chr19 | 15829224 | 15829227 | + | 7.02772403811355 | 9.09596593651506 | 2.38645642073598 | 2.13421729395427 |
| chr19 | 15834583 | 15834599 | + | 5.01980288436682 | 1.13699574206438 | 5.96614105183995 | 1.06710864697714 |
| chr19 | 15835043 | 15835066 | + | 25.0990144218341 | 17.0549361309657 | 21.4781077866238 | 14.9395210576799 |
| chr19 | 15835135 | 15835143 | + | 13.0514874993537 | 12.5069531627082 | 19.0916513658878 | 9.60397782279422 |
| chr19 | 15835175 | 15835182 | + | 15.0594086531005 | 5.68497871032191 | 14.3187385244159 | 13.8724124107028 |
| chr19 | 15835271 | 15835307 | + | 31.1227778830743 | 26.1509020674808 | 21.4781077866238 | 20.2750642925656 |
| chr19 | 15835331 | 15835343 | + | 21.0831721143407 | 23.876910583352 | 31.0239334695677 | 18.1408469986113 |
| chr19 | 15835450 | 15835462 | + | 3.01188173062009 | 5.68497871032191 | 3.57968463110397 | 1.06710864697714 |
| chr19 | 15835504 | 15835524 | + | 2.00792115374673 | 6.82197445238629 | 4.77291284147196 | 5.33554323488568 |
| chr19 | 15835552 | 15835630 | + | 110.43566345607 | 137.57648478979 | 144.380613454527 | 76.8318225823538 |
| chr19 | 15835704 | 15835711 | + | 5.01980288436682 | 5.68497871032191 | 13.1255103140479 | 1.06710864697714 |
| chr19 | 16101474 | 16101526 | + | 32.1267384599477 | 28.4248935516096 | 20.2848795762558 | 23.476390233497 |
| chr19 | 16101547 | 16101587 | + | 17.0673298068472 | 10.2329616785794 | 5.96614105183995 | 3.20132594093141 |
| chr19 | 16102249 | 16102294 | + | 15.0594086531005 | 35.2468680039958 | 9.54582568294392 | 7.46976052883995 |
| chr19 | 16102562 | 16102578 | + | 7.02772403811355 | 10.2329616785794 | 2.38645642073598 | 1.06710864697714 |
| chr19 | 16102765 | 16102783 | + | 9.03564519186028 | 22.7399148412876 | 7.15936926220794 | 3.20132594093141 |
| chr19 | 16132380 | 16132450 | + | 32.1267384599477 | 17.0549361309657 | 23.8645642073598 | 23.476390233497 |
| chr19 | 16132492 | 16132517 | + | 5.01980288436682 | 4.54798296825753 | 4.77291284147196 | 2.13421729395427 |
| chr19 | 16132737 | 16132757 | + | 3.01188173062009 | 2.27399148412876 | 3.57968463110397 | 4.26843458790854 |
| chr19 | 16132759 | 16132812 | + | 31.1227778830743 | 11.3699574206438 | 10.7390538933119 | 9.60397782279422 |
| chr19 | 16132837 | 16132845 | + | 5.01980288436682 | 1.13699574206438 | 1.19322821036799 | 3.20132594093141 |
| chr19 | 16132909 | 16132931 | + | 9.03564519186028 | 10.2329616785794 | 4.77291284147196 | 1.06710864697714 |
| chr19 | 16132953 | 16132976 | + | 23.0910932680874 | 18.1919318730301 | 15.5119667347839 | 20.2750642925656 |
| chr19 | 16133015 | 16133262 | + | 130.514874993537 | 115.973565690567 | 170.631634082622 | 87.5029090521251 |
| chr19 | 16133322 | 16133355 | + | 4.01584230749346 | 10.2329616785794 | 20.2848795762558 | 10.6710864697714 |
| chr19 | 16133383 | 16133432 | + | 4832.0622564915 | 2887.96918484353 | 1486.76235011851 | 2300.6862428827 |
| chr19 | 16133535 | 16133557 | + | 13.0514874993537 | 6.82197445238629 | 1.19322821036799 | 9.60397782279422 |
| chr19 | 16133599 | 16133600 | + | 6.02376346124019 | 2.27399148412876 | 1.19322821036799 | 2.13421729395427 |
| chr19 | 16190970 | 16190988 | + | 3.01188173062009 | 1.13699574206438 | 1.19322821036799 | 6.40265188186281 |
| chr19 | 16191697 | 16191699 | + | 1.00396057687336 | 1.13699574206438 | 3.57968463110397 | 5.33554323488568 |
| chr19 | 16191823 | 16191901 | + | 147.582204800385 | 77.315710460378 | 32.2171616799357 | 87.5029090521251 |
| chr19 | 16191918 | 16191947 | + | 13.0514874993537 | 14.780944646837 | 19.0916513658878 | 11.7381951167485 |
| chr19 | 16191967 | 16192029 | + | 32.1267384599477 | 64.8087572976698 | 8.35259747257593 | 18.1408469986113 |
| chr19 | 16235235 | 16235311 | + | 9.03564519186028 | 18.1919318730301 | 8.35259747257593 | 10.6710864697714 |
| chr19 | 16355299 | 16355314 | - | 1.00396057687336 | 3.41098722619315 | 3.57968463110397 | 5.33554323488568 |
| chr19 | 16517935 | 16517958 | - | 1.00396057687336 | 4.54798296825753 | 1.19322821036799 | 1.06710864697714 |
| chr19 | 16518120 | 16518142 | - | 10.0396057687336 | 7.95897019445067 | 3.57968463110397 | 10.6710864697714 |
| chr19 | 16543413 | 16543428 | + | 1.00396057687336 | 3.41098722619315 | 4.77291284147196 | 3.20132594093141 |
| chr19 | 16550908 | 16550922 | - | 3.01188173062009 | 10.2329616785794 | 4.77291284147196 | 5.33554323488568 |
| chr19 | 16550966 | 16551005 | - | 10.0396057687336 | 17.0549361309657 | 20.2848795762558 | 4.26843458790854 |
| chr19 | 16551022 | 16551035 | - | 1.00396057687336 | 1.13699574206438 | 5.96614105183995 | 2.13421729395427 |
| chr19 | 16551060 | 16551142 | - | 76.3010038423757 | 103.466612527859 | 76.3666054635513 | 68.2949534065367 |
| chr19 | 16551193 | 16551236 | - | 9.03564519186028 | 15.9179403889013 | 9.54582568294392 | 19.2079556455884 |
| chr19 | 16551272 | 16551327 | - | 23.0910932680874 | 7.95897019445067 | 3.57968463110397 | 10.6710864697714 |
| chr19 | 16551442 | 16551473 | - | 5.01980288436682 | 3.41098722619315 | 4.77291284147196 | 4.26843458790854 |
| chr19 | 16551518 | 16551541 | - | 4.01584230749346 | 6.82197445238629 | 9.54582568294392 | 5.33554323488568 |
| chr19 | 16551586 | 16551602 | - | 3.01188173062009 | 2.27399148412876 | 3.57968463110397 | 3.20132594093141 |
| chr19 | 16551842 | 16551844 | - | 1.00396057687336 | 3.41098722619315 | 3.57968463110397 | 1.06710864697714 |
| chr19 | 16551990 | 16552000 | - | 5.01980288436682 | 2.27399148412876 | 1.19322821036799 | 4.26843458790854 |
| chr19 | 16552786 | 16552807 | - | 3.01188173062009 | 6.82197445238629 | 2.38645642073598 | 1.06710864697714 |
| chr19 | 16552902 | 16552929 | - | 12.0475269224804 | 7.95897019445067 | 4.77291284147196 | 8.53686917581709 |
| chr19 | 16552968 | 16552979 | - | 17.0673298068472 | 7.95897019445067 | 8.35259747257593 | 13.8724124107028 |
| chr19 | 16566531 | 16566538 | - | 18.0712903837206 | 6.82197445238629 | 14.3187385244159 | 7.46976052883995 |
| chr19 | 16572596 | 16572684 | + | 17.0673298068472 | 17.0549361309657 | 13.1255103140479 | 25.6106075274513 |
| chr19 | 16574992 | 16574995 | - | 6.02376346124019 | 3.41098722619315 | 3.57968463110397 | 4.26843458790854 |
| chr19 | 16575143 | 16575146 | - | 5.01980288436682 | 2.27399148412876 | 3.57968463110397 | 4.26843458790854 |
| chr19 | 16645328 | 16645329 | - | 1.00396057687336 | 1.13699574206438 | 1.19322821036799 | 1.06710864697714 |
| chr19 | 16646145 | 16646190 | - | 6.02376346124019 | 13.6439489047726 | 33.4103898903037 | 10.6710864697714 |
| chr19 | 16646216 | 16646234 | - | 3.01188173062009 | 3.41098722619315 | 7.15936926220794 | 3.20132594093141 |
| chr19 | 16646337 | 16646340 | - | 1.00396057687336 | 3.41098722619315 | 1.19322821036799 | 3.20132594093141 |
| chr19 | 16646361 | 16646364 | - | 8.03168461498692 | 2.27399148412876 | 2.38645642073598 | 5.33554323488568 |
| chr19 | 16646620 | 16646649 | - | 10.0396057687336 | 13.6439489047726 | 5.96614105183995 | 18.1408469986113 |
| chr19 | 16646671 | 16646688 | - | 3.01188173062009 | 3.41098722619315 | 2.38645642073598 | 1.06710864697714 |
| chr19 | 16646746 | 16646773 | - | 1.00396057687336 | 1.13699574206438 | 2.38645642073598 | 1.06710864697714 |
| chr19 | 16854282 | 16854296 | + | 7.02772403811355 | 4.54798296825753 | 3.57968463110397 | 1.06710864697714 |
| chr19 | 16880226 | 16880295 | + | 22.087132691214 | 14.780944646837 | 11.9322821036799 | 22.4092815865198 |
| chr19 | 17049804 | 17049889 | - | 35.1386201905678 | 62.534765813541 | 23.8645642073598 | 28.8119334683827 |
| chr19 | 17212519 | 17212558 | + | 10.0396057687336 | 3.41098722619315 | 8.35259747257593 | 10.6710864697714 |
| chr19 | 17219737 | 17219740 | + | 2.00792115374673 | 4.54798296825753 | 1.19322821036799 | 1.06710864697714 |
| chr19 | 17231911 | 17232012 | - | 34.1346596136944 | 23.876910583352 | 16.7051949451519 | 21.3421729395427 |
| chr19 | 17232239 | 17232268 | - | 4.01584230749346 | 3.41098722619315 | 3.57968463110397 | 9.60397782279422 |
| chr19 | 17273965 | 17274004 | + | 5.01980288436682 | 3.41098722619315 | 9.54582568294392 | 5.33554323488568 |
| chr19 | 17274104 | 17274160 | + | 6.02376346124019 | 9.09596593651506 | 17.8984231555198 | 9.60397782279422 |
| chr19 | 17276592 | 17276625 | + | 6.02376346124019 | 3.41098722619315 | 8.35259747257593 | 9.60397782279422 |
| chr19 | 17276877 | 17276910 | + | 20.0792115374673 | 14.780944646837 | 23.8645642073598 | 14.9395210576799 |
| chr19 | 17279141 | 17279231 | + | 15.0594086531005 | 14.780944646837 | 5.96614105183995 | 16.006629704657 |
| chr19 | 17279260 | 17279300 | + | 4.01584230749346 | 1.13699574206438 | 3.57968463110397 | 2.13421729395427 |
| chr19 | 17292172 | 17292202 | - | 3.01188173062009 | 1.13699574206438 | 2.38645642073598 | 2.13421729395427 |
| chr19 | 17306456 | 17306532 | + | 64.2534769198953 | 63.6717615556054 | 15.5119667347839 | 27.7448248214055 |
| chr19 | 17306559 | 17306696 | + | 33.130699036821 | 37.5208594881246 | 8.35259747257593 | 14.9395210576799 |
| chr19 | 17306735 | 17306785 | + | 45.1782259593014 | 62.534765813541 | 23.8645642073598 | 32.0132594093141 |
| chr19 | 17306814 | 17306818 | + | 42.1663442286813 | 55.7127913611547 | 5.96614105183995 | 12.8053037637256 |
| chr19 | 17319799 | 17319804 | + | 8.03168461498692 | 2.27399148412876 | 2.38645642073598 | 2.13421729395427 |
| chr19 | 17323367 | 17323373 | - | 1.00396057687336 | 3.41098722619315 | 1.19322821036799 | 2.13421729395427 |
| chr19 | 17342609 | 17342627 | + | 12.0475269224804 | 4.54798296825753 | 3.57968463110397 | 5.33554323488568 |
| chr19 | 17342663 | 17342713 | + | 11.043566345607 | 4.54798296825753 | 3.57968463110397 | 1.06710864697714 |
| chr19 | 17402970 | 17403011 | - | 95.3762548029696 | 79.5897019445067 | 9.54582568294392 | 16.006629704657 |
| chr19 | 17403039 | 17403053 | - | 18.0712903837206 | 12.5069531627082 | 4.77291284147196 | 2.13421729395427 |
| chr19 | 17405669 | 17405774 | + | 602.376346124019 | 553.716926385354 | 122.902505667903 | 84.3015831111937 |
| chr19 | 17405819 | 17405851 | + | 85.336649034236 | 48.8908169087684 | 38.1833027317757 | 33.0803680562912 |
| chr19 | 17406028 | 17406151 | + | 84.3326884573626 | 64.8087572976698 | 7.15936926220794 | 5.33554323488568 |
| chr19 | 17425277 | 17425301 | + | 6.02376346124019 | 5.68497871032191 | 3.57968463110397 | 3.20132594093141 |
| chr19 | 17521139 | 17521159 | + | 9.03564519186028 | 2.27399148412876 | 4.77291284147196 | 1.06710864697714 |
| chr19 | 17521227 | 17521266 | + | 32.1267384599477 | 36.3838637460602 | 13.1255103140479 | 22.4092815865198 |
| chr19 | 17581383 | 17581512 | + | 39.1544624980612 | 27.2878978095452 | 31.0239334695677 | 18.1408469986113 |
| chr19 | 17581545 | 17581575 | + | 59.2336740355285 | 45.4798296825753 | 33.4103898903037 | 10.6710864697714 |
| chr19 | 17581584 | 17581611 | + | 6.02376346124019 | 2.27399148412876 | 4.77291284147196 | 4.26843458790854 |
| chr19 | 17581647 | 17581667 | + | 22.087132691214 | 17.0549361309657 | 9.54582568294392 | 6.40265188186281 |
| chr19 | 17581715 | 17581723 | + | 7.02772403811355 | 3.41098722619315 | 4.77291284147196 | 1.06710864697714 |
| chr19 | 17581744 | 17581775 | + | 14.0554480762271 | 15.9179403889013 | 7.15936926220794 | 49.0869977609482 |
| chr19 | 17581856 | 17581900 | + | 49.1940682667949 | 37.5208594881246 | 38.1833027317757 | 122.717494402371 |
| chr19 | 17581927 | 17581933 | + | 5.01980288436682 | 3.41098722619315 | 3.57968463110397 | 2.13421729395427 |
| chr19 | 17581959 | 17581973 | + | 4.01584230749346 | 3.41098722619315 | 1.19322821036799 | 3.20132594093141 |
| chr19 | 17582040 | 17582056 | + | 17.0673298068472 | 9.09596593651506 | 8.35259747257593 | 7.46976052883995 |
| chr19 | 17582190 | 17582297 | + | 44.174265382428 | 28.4248935516096 | 33.4103898903037 | 41.6172372321083 |
| chr19 | 17582352 | 17582390 | + | 12.0475269224804 | 5.68497871032191 | 2.38645642073598 | 6.40265188186281 |
| chr19 | 17582966 | 17583072 | + | 31.1227778830743 | 15.9179403889013 | 5.96614105183995 | 18.1408469986113 |
| chr19 | 17583091 | 17583128 | + | 84.3326884573626 | 103.466612527859 | 40.5697591525116 | 35.2145853502455 |
| chr19 | 17734462 | 17734467 | + | 3.01188173062009 | 3.41098722619315 | 1.19322821036799 | 2.13421729395427 |
| chr19 | 17943921 | 17943943 | + | 10.0396057687336 | 5.68497871032191 | 8.35259747257593 | 8.53686917581709 |
| chr19 | 17943965 | 17943966 | + | 7.02772403811355 | 4.54798296825753 | 1.19322821036799 | 10.6710864697714 |
| chr19 | 17999029 | 17999054 | + | 1.00396057687336 | 1.13699574206438 | 27.4442488384638 | 19.2079556455884 |
| chr19 | 18014009 | 18014041 | + | 8.03168461498692 | 11.3699574206438 | 1.19322821036799 | 3.20132594093141 |
| chr19 | 18149950 | 18149960 | + | 2.00792115374673 | 1.13699574206438 | 3.57968463110397 | 5.33554323488568 |
| chr19 | 18169358 | 18169426 | + | 17.0673298068472 | 9.09596593651506 | 7.15936926220794 | 11.7381951167485 |
| chr19 | 18170474 | 18170494 | + | 6.02376346124019 | 9.09596593651506 | 11.9322821036799 | 4.26843458790854 |
| chr19 | 18173749 | 18173841 | + | 3871.27198442369 | 1781.67232781489 | 760.086370004409 | 1638.0117731099 |
| chr19 | 18175058 | 18175075 | + | 9.03564519186028 | 5.68497871032191 | 1.19322821036799 | 1.06710864697714 |
| chr19 | 18175310 | 18175383 | + | 25.0990144218341 | 6.82197445238629 | 3.57968463110397 | 8.53686917581709 |
| chr19 | 18175610 | 18175617 | + | 7.02772403811355 | 1.13699574206438 | 1.19322821036799 | 1.06710864697714 |
| chr19 | 18177236 | 18177253 | + | 14.0554480762271 | 3.41098722619315 | 1.19322821036799 | 1.06710864697714 |
| chr19 | 18177877 | 18177963 | + | 1626.41613453485 | 472.990228698783 | 72.7869208324474 | 274.246922273124 |
| chr19 | 18177989 | 18178040 | + | 20.0792115374673 | 32.9728765198671 | 3.57968463110397 | 16.006629704657 |
| chr19 | 18178088 | 18178092 | + | 26.1029749987075 | 65.9457530397342 | 4.77291284147196 | 5.33554323488568 |
| chr19 | 18196064 | 18196151 | + | 15.0594086531005 | 17.0549361309657 | 95.4582568294392 | 32.0132594093141 |
| chr19 | 18196179 | 18196221 | + | 3.01188173062009 | 4.54798296825753 | 34.6036181006717 | 8.53686917581709 |
| chr19 | 18196422 | 18196481 | + | 6.02376346124019 | 7.95897019445067 | 17.8984231555198 | 17.0737383516342 |
| chr19 | 18196535 | 18196550 | + | 5.01980288436682 | 5.68497871032191 | 4.77291284147196 | 3.20132594093141 |
| chr19 | 18279778 | 18279830 | - | 11.043566345607 | 30.6988850357383 | 23.8645642073598 | 19.2079556455884 |
| chr19 | 18279864 | 18280050 | - | 51.2019894205416 | 55.7127913611547 | 44.1494437836156 | 105.643756050736 |
| chr19 | 18280080 | 18280115 | - | 9.03564519186028 | 21.6029190992233 | 42.9562155732476 | 88.5700176991023 |
| chr19 | 18280249 | 18280264 | - | 1.00396057687336 | 2.27399148412876 | 1.19322821036799 | 5.33554323488568 |
| chr19 | 18280427 | 18280428 | - | 1.00396057687336 | 1.13699574206438 | 4.77291284147196 | 2.13421729395427 |
| chr19 | 18282089 | 18282095 | + | 34.1346596136944 | 83.0006891706999 | 10.7390538933119 | 21.3421729395427 |
| chr19 | 18306938 | 18307010 | - | 77.3049644192491 | 84.1376849127643 | 94.2650286190712 | 124.851711696325 |
| chr19 | 18307105 | 18307118 | - | 2.00792115374673 | 5.68497871032191 | 1.19322821036799 | 6.40265188186281 |
| chr19 | 18307201 | 18307234 | - | 5.01980288436682 | 1.13699574206438 | 10.7390538933119 | 7.46976052883995 |
| chr19 | 18307269 | 18307287 | - | 3.01188173062009 | 7.95897019445067 | 9.54582568294392 | 2.13421729395427 |
| chr19 | 18307339 | 18307368 | - | 6.02376346124019 | 1.13699574206438 | 5.96614105183995 | 5.33554323488568 |
| chr19 | 18367052 | 18367086 | + | 4.01584230749346 | 1.13699574206438 | 10.7390538933119 | 10.6710864697714 |
| chr19 | 18367346 | 18367393 | + | 4.01584230749346 | 3.41098722619315 | 10.7390538933119 | 7.46976052883995 |
| chr19 | 18367568 | 18367594 | + | 1.00396057687336 | 1.13699574206438 | 7.15936926220794 | 6.40265188186281 |
| chr19 | 18368199 | 18368210 | + | 3.01188173062009 | 2.27399148412876 | 4.77291284147196 | 4.26843458790854 |
| chr19 | 18368556 | 18368559 | + | 1.00396057687336 | 1.13699574206438 | 3.57968463110397 | 7.46976052883995 |
| chr19 | 18368928 | 18368953 | + | 1.00396057687336 | 4.54798296825753 | 20.2848795762558 | 9.60397782279422 |
| chr19 | 18369857 | 18369928 | + | 1.00396057687336 | 3.41098722619315 | 8.35259747257593 | 20.2750642925656 |
| chr19 | 18388964 | 18389156 | + | 19.0752509605939 | 71.6307317500561 | 40.5697591525116 | 76.8318225823538 |
| chr19 | 18434517 | 18434543 | + | 5.01980288436682 | 11.3699574206438 | 14.3187385244159 | 32.0132594093141 |
| chr19 | 18442826 | 18442834 | - | 5.01980288436682 | 4.54798296825753 | 3.57968463110397 | 10.6710864697714 |
| chr19 | 18442936 | 18442939 | - | 6.02376346124019 | 2.27399148412876 | 2.38645642073598 | 2.13421729395427 |
| chr19 | 18443033 | 18443045 | - | 2.00792115374673 | 2.27399148412876 | 8.35259747257593 | 14.9395210576799 |
| chr19 | 18443688 | 18443736 | - | 4.01584230749346 | 3.41098722619315 | 25.0577924177278 | 16.006629704657 |
| chr19 | 18531785 | 18531827 | - | 66.2613980736421 | 57.9867828452835 | 33.4103898903037 | 39.483019938154 |
| chr19 | 18531958 | 18531975 | - | 13.0514874993537 | 4.54798296825753 | 26.2510206280958 | 24.5434988804741 |
| chr19 | 18532036 | 18532079 | - | 12.0475269224804 | 18.1919318730301 | 8.35259747257593 | 8.53686917581709 |
| chr19 | 18532090 | 18532104 | - | 1.00396057687336 | 1.13699574206438 | 2.38645642073598 | 8.53686917581709 |
| chr19 | 18569221 | 18569355 | + | 38.1505019211879 | 29.5618892936739 | 41.7629873628796 | 81.1002571702623 |
| chr19 | 18575056 | 18575192 | + | 74.293082688629 | 68.2197445238629 | 84.7192029361272 | 172.871600810296 |
| chr19 | 18575213 | 18575215 | + | 20.0792115374673 | 32.9728765198671 | 38.1833027317757 | 21.3421729395427 |
| chr19 | 18575333 | 18575340 | + | 11.043566345607 | 11.3699574206438 | 10.7390538933119 | 18.1408469986113 |
| chr19 | 18575398 | 18575421 | + | 8.03168461498692 | 12.5069531627082 | 3.57968463110397 | 4.26843458790854 |
| chr19 | 18576383 | 18576389 | + | 2.00792115374673 | 1.13699574206438 | 2.38645642073598 | 4.26843458790854 |
| chr19 | 18576453 | 18576471 | + | 5.01980288436682 | 4.54798296825753 | 2.38645642073598 | 4.26843458790854 |
| chr19 | 18576526 | 18576529 | + | 42.1663442286813 | 84.1376849127643 | 2.38645642073598 | 1.06710864697714 |
| chr19 | 18576878 | 18576888 | + | 5.01980288436682 | 7.95897019445067 | 9.54582568294392 | 2.13421729395427 |
| chr19 | 18576954 | 18576967 | + | 16.0633692299738 | 37.5208594881246 | 22.6713359969918 | 59.7580842307196 |
| chr19 | 18577073 | 18577092 | + | 5.01980288436682 | 4.54798296825753 | 4.77291284147196 | 5.33554323488568 |
| chr19 | 18577156 | 18577182 | + | 22.087132691214 | 10.2329616785794 | 10.7390538933119 | 8.53686917581709 |
| chr19 | 18577289 | 18577422 | + | 121.479229801677 | 98.9186295596012 | 62.0478669391355 | 87.5029090521251 |
| chr19 | 18592213 | 18592232 | + | 1.00396057687336 | 2.27399148412876 | 2.38645642073598 | 9.60397782279422 |
| chr19 | 18592279 | 18592304 | + | 5.01980288436682 | 2.27399148412876 | 14.3187385244159 | 8.53686917581709 |
| chr19 | 18670206 | 18670231 | + | 14.0554480762271 | 11.3699574206438 | 20.2848795762558 | 11.7381951167485 |
| chr19 | 18866523 | 18866570 | + | 9.03564519186028 | 10.2329616785794 | 17.8984231555198 | 12.8053037637256 |
| chr19 | 18866732 | 18866809 | + | 26.1029749987075 | 18.1919318730301 | 23.8645642073598 | 28.8119334683827 |
| chr19 | 18866869 | 18866891 | + | 7.02772403811355 | 1.13699574206438 | 5.96614105183995 | 2.13421729395427 |
| chr19 | 18866953 | 18866972 | + | 2.00792115374673 | 1.13699574206438 | 2.38645642073598 | 2.13421729395427 |
| chr19 | 18866994 | 18867055 | + | 34.1346596136944 | 22.7399148412876 | 28.6374770488317 | 25.6106075274513 |
| chr19 | 18867140 | 18867161 | + | 5.01980288436682 | 3.41098722619315 | 14.3187385244159 | 3.20132594093141 |
| chr19 | 18867225 | 18867247 | + | 14.0554480762271 | 9.09596593651506 | 14.3187385244159 | 14.9395210576799 |
| chr19 | 18867292 | 18867294 | + | 14.0554480762271 | 10.2329616785794 | 5.96614105183995 | 19.2079556455884 |
| chr19 | 18867646 | 18867667 | + | 4.01584230749346 | 2.27399148412876 | 5.96614105183995 | 4.26843458790854 |
| chr19 | 18867891 | 18867928 | + | 1.00396057687336 | 3.41098722619315 | 8.35259747257593 | 6.40265188186281 |
| chr19 | 18868066 | 18868075 | + | 5.01980288436682 | 7.95897019445067 | 2.38645642073598 | 5.33554323488568 |
| chr19 | 18868097 | 18868100 | + | 3.01188173062009 | 1.13699574206438 | 2.38645642073598 | 5.33554323488568 |
| chr19 | 18868142 | 18868210 | + | 11.043566345607 | 15.9179403889013 | 13.1255103140479 | 21.3421729395427 |
| chr19 | 18899548 | 18899582 | - | 15.0594086531005 | 13.6439489047726 | 10.7390538933119 | 18.1408469986113 |
| chr19 | 18906997 | 18907016 | - | 7.02772403811355 | 2.27399148412876 | 10.7390538933119 | 14.9395210576799 |
| chr19 | 18907061 | 18907101 | - | 7.02772403811355 | 6.82197445238629 | 21.4781077866238 | 18.1408469986113 |
| chr19 | 18907109 | 18907130 | - | 5.01980288436682 | 4.54798296825753 | 1.19322821036799 | 3.20132594093141 |
| chr19 | 18910970 | 18910982 | - | 5.01980288436682 | 2.27399148412876 | 5.96614105183995 | 7.46976052883995 |
| chr19 | 18911001 | 18911031 | - | 1.00396057687336 | 3.41098722619315 | 4.77291284147196 | 1.06710864697714 |
| chr19 | 18911045 | 18911075 | - | 14.0554480762271 | 10.2329616785794 | 14.3187385244159 | 6.40265188186281 |
| chr19 | 18912989 | 18913045 | - | 20.0792115374673 | 26.1509020674808 | 28.6374770488317 | 33.0803680562912 |
| chr19 | 18922467 | 18922471 | + | 1.00396057687336 | 1.13699574206438 | 1.19322821036799 | 3.20132594093141 |
| chr19 | 18928525 | 18928550 | + | 4.01584230749346 | 5.68497871032191 | 1.19322821036799 | 4.26843458790854 |
| chr19 | 18928581 | 18928612 | + | 59.2336740355285 | 78.4527062024424 | 27.4442488384638 | 27.7448248214055 |
| chr19 | 18929263 | 18929304 | - | 6.02376346124019 | 4.54798296825753 | 2.38645642073598 | 1.06710864697714 |
| chr19 | 18990933 | 18990955 | - | 1.00396057687336 | 4.54798296825753 | 2.38645642073598 | 3.20132594093141 |
| chr19 | 18990984 | 18991061 | - | 3.01188173062009 | 5.68497871032191 | 17.8984231555198 | 16.006629704657 |
| chr19 | 18991074 | 18991144 | - | 10.0396057687336 | 10.2329616785794 | 8.35259747257593 | 17.0737383516342 |
| chr19 | 18993112 | 18993119 | - | 2.00792115374673 | 1.13699574206438 | 4.77291284147196 | 2.13421729395427 |
| chr19 | 18993637 | 18993648 | - | 4.01584230749346 | 5.68497871032191 | 11.9322821036799 | 6.40265188186281 |
| chr19 | 18993828 | 18993894 | - | 7.02772403811355 | 12.5069531627082 | 25.0577924177278 | 30.9461507623369 |
| chr19 | 19057642 | 19057655 | + | 2.00792115374673 | 2.27399148412876 | 1.19322821036799 | 9.60397782279422 |
| chr19 | 19057682 | 19057693 | + | 7.02772403811355 | 4.54798296825753 | 17.8984231555198 | 7.46976052883995 |
| chr19 | 19057720 | 19057813 | + | 6.02376346124019 | 6.82197445238629 | 27.4442488384638 | 22.4092815865198 |
| chr19 | 19057906 | 19057933 | + | 4.01584230749346 | 1.13699574206438 | 5.96614105183995 | 7.46976052883995 |
| chr19 | 19058129 | 19058142 | + | 4.01584230749346 | 3.41098722619315 | 2.38645642073598 | 3.20132594093141 |
| chr19 | 19112696 | 19112706 | + | 3.01188173062009 | 1.13699574206438 | 5.96614105183995 | 19.2079556455884 |
| chr19 | 19119635 | 19119638 | - | 1.00396057687336 | 1.13699574206438 | 4.77291284147196 | 4.26843458790854 |
| chr19 | 19119659 | 19119677 | - | 34.1346596136944 | 36.3838637460602 | 28.6374770488317 | 28.8119334683827 |
| chr19 | 19119713 | 19119786 | - | 5.01980288436682 | 4.54798296825753 | 9.54582568294392 | 11.7381951167485 |
| chr19 | 19176987 | 19177012 | - | 2.00792115374673 | 6.82197445238629 | 1.19322821036799 | 4.26843458790854 |
| chr19 | 19181993 | 19181995 | - | 2.00792115374673 | 4.54798296825753 | 1.19322821036799 | 2.13421729395427 |
| chr19 | 19201505 | 19201515 | - | 1.00396057687336 | 4.54798296825753 | 3.57968463110397 | 5.33554323488568 |
| chr19 | 19201821 | 19201843 | + | 11.043566345607 | 7.95897019445067 | 16.7051949451519 | 36.2816939972226 |
| chr19 | 19276542 | 19276551 | - | 7.02772403811355 | 6.82197445238629 | 3.57968463110397 | 4.26843458790854 |
| chr19 | 19356862 | 19356865 | + | 1.00396057687336 | 1.13699574206438 | 2.38645642073598 | 3.20132594093141 |
| chr19 | 19505698 | 19505711 | + | 1.00396057687336 | 3.41098722619315 | 2.38645642073598 | 4.26843458790854 |
| chr19 | 19505885 | 19505894 | + | 7.02772403811355 | 5.68497871032191 | 3.57968463110397 | 11.7381951167485 |
| chr19 | 19505950 | 19505968 | + | 6.02376346124019 | 7.95897019445067 | 8.35259747257593 | 20.2750642925656 |
| chr19 | 19506259 | 19506309 | + | 8.03168461498692 | 2.27399148412876 | 1.19322821036799 | 17.0737383516342 |
| chr19 | 19506353 | 19506384 | + | 11.043566345607 | 6.82197445238629 | 4.77291284147196 | 7.46976052883995 |
| chr19 | 19506520 | 19506544 | + | 3.01188173062009 | 2.27399148412876 | 3.57968463110397 | 24.5434988804741 |
| chr19 | 19506584 | 19506601 | + | 10.0396057687336 | 6.82197445238629 | 8.35259747257593 | 41.6172372321083 |
| chr19 | 19506665 | 19506676 | + | 13.0514874993537 | 4.54798296825753 | 3.57968463110397 | 22.4092815865198 |
| chr19 | 19506771 | 19506797 | + | 5.01980288436682 | 1.13699574206438 | 3.57968463110397 | 5.33554323488568 |
| chr19 | 19506815 | 19506896 | + | 55.217831728035 | 20.4659233571589 | 36.9900745214077 | 69.3620620535138 |
| chr19 | 19506920 | 19506922 | + | 4.01584230749346 | 3.41098722619315 | 1.19322821036799 | 5.33554323488568 |
| chr19 | 19506963 | 19507012 | + | 7.02772403811355 | 4.54798296825753 | 5.96614105183995 | 7.46976052883995 |
| chr19 | 19507140 | 19507164 | + | 22.087132691214 | 13.6439489047726 | 11.9322821036799 | 39.483019938154 |
| chr19 | 19507185 | 19507194 | + | 4.01584230749346 | 2.27399148412876 | 3.57968463110397 | 6.40265188186281 |
| chr19 | 19507229 | 19507237 | + | 69.2732798042621 | 62.534765813541 | 48.9223566250876 | 38.4159112911769 |
| chr19 | 19507709 | 19507728 | + | 2.00792115374673 | 1.13699574206438 | 1.19322821036799 | 16.006629704657 |
| chr19 | 19508862 | 19508886 | + | 3.01188173062009 | 2.27399148412876 | 2.38645642073598 | 3.20132594093141 |
| chr19 | 19528116 | 19528180 | + | 67.2653586505154 | 101.19262104373 | 38.1833027317757 | 108.845081991668 |
| chr19 | 19561750 | 19561751 | - | 1.00396057687336 | 1.13699574206438 | 2.38645642073598 | 1.06710864697714 |
| chr19 | 19645229 | 19645260 | - | 3.01188173062009 | 2.27399148412876 | 2.38645642073598 | 6.40265188186281 |
| chr19 | 19645350 | 19645377 | - | 15.0594086531005 | 2.27399148412876 | 13.1255103140479 | 12.8053037637256 |
| chr19 | 21120702 | 21120721 | + | 2.00792115374673 | 1.13699574206438 | 8.35259747257593 | 1.06710864697714 |
| chr19 | 21122027 | 21122034 | + | 13.0514874993537 | 5.68497871032191 | 7.15936926220794 | 14.9395210576799 |
| chr19 | 21122139 | 21122140 | + | 7.02772403811355 | 5.68497871032191 | 7.15936926220794 | 4.26843458790854 |
| chr19 | 21186863 | 21186870 | + | 3.01188173062009 | 1.13699574206438 | 5.96614105183995 | 2.13421729395427 |
| chr19 | 21190947 | 21190952 | + | 3.01188173062009 | 2.27399148412876 | 7.15936926220794 | 3.20132594093141 |
| chr19 | 21191577 | 21191596 | + | 10.0396057687336 | 13.6439489047726 | 14.3187385244159 | 3.20132594093141 |
| chr19 | 24033399 | 24033549 | + | 196.776273067179 | 293.344901452611 | 138.414472402687 | 108.845081991668 |
| chr19 | 24128066 | 24128071 | + | 10.0396057687336 | 7.95897019445067 | 28.6374770488317 | 50.1541064079254 |
| chr19 | 27793419 | 27793491 | + | 375.481255750638 | 336.550739651057 | 137.221244192319 | 334.005006503844 |
| chr19 | 27793997 | 27794071 | + | 68.2693192273888 | 71.6307317500561 | 78.7530618842873 | 131.254363578188 |
| chr19 | 27798156 | 27798177 | + | 3.01188173062009 | 3.41098722619315 | 13.1255103140479 | 10.6710864697714 |
| chr19 | 27806184 | 27806186 | + | 9.03564519186028 | 1.13699574206438 | 9.54582568294392 | 20.2750642925656 |
| chr19 | 29615532 | 29615536 | + | 1.00396057687336 | 1.13699574206438 | 5.96614105183995 | 1.06710864697714 |
| chr19 | 29615647 | 29615675 | + | 22.087132691214 | 29.5618892936739 | 17.8984231555198 | 16.006629704657 |
| chr19 | 29615711 | 29615717 | + | 4.01584230749346 | 5.68497871032191 | 1.19322821036799 | 4.26843458790854 |
| chr19 | 29617085 | 29617088 | + | 2.00792115374673 | 2.27399148412876 | 1.19322821036799 | 1.06710864697714 |
| chr19 | 29824193 | 29824201 | + | 9.03564519186028 | 10.2329616785794 | 3.57968463110397 | 17.0737383516342 |
| chr19 | 30015394 | 30015420 | + | 2.00792115374673 | 1.13699574206438 | 2.38645642073598 | 1.06710864697714 |
| chr19 | 30015600 | 30015652 | + | 4.01584230749346 | 3.41098722619315 | 4.77291284147196 | 3.20132594093141 |
| chr19 | 30016277 | 30016281 | + | 4.01584230749346 | 1.13699574206438 | 3.57968463110397 | 1.06710864697714 |
| chr19 | 32384449 | 32384467 | + | 4.01584230749346 | 2.27399148412876 | 3.57968463110397 | 1.06710864697714 |
| chr19 | 32384531 | 32384538 | + | 12.0475269224804 | 3.41098722619315 | 1.19322821036799 | 1.06710864697714 |
| chr19 | 32384675 | 32384693 | + | 18.0712903837206 | 11.3699574206438 | 5.96614105183995 | 12.8053037637256 |
| chr19 | 32385997 | 32386005 | + | 4.01584230749346 | 2.27399148412876 | 4.77291284147196 | 6.40265188186281 |
| chr19 | 32386044 | 32386048 | + | 4.01584230749346 | 1.13699574206438 | 3.57968463110397 | 2.13421729395427 |
| chr19 | 32386090 | 32386095 | + | 1.00396057687336 | 2.27399148412876 | 4.77291284147196 | 1.06710864697714 |
| chr19 | 32386228 | 32386233 | + | 1.00396057687336 | 1.13699574206438 | 1.19322821036799 | 1.06710864697714 |
| chr19 | 32386687 | 32386689 | + | 3.01188173062009 | 6.82197445238629 | 1.19322821036799 | 6.40265188186281 |
| chr19 | 32482952 | 32482998 | + | 7.02772403811355 | 18.1919318730301 | 10.7390538933119 | 22.4092815865198 |
| chr19 | 32587262 | 32587287 | + | 14.0554480762271 | 6.82197445238629 | 9.54582568294392 | 19.2079556455884 |
| chr19 | 32587308 | 32587391 | + | 117.463387494184 | 160.316399631078 | 59.6614105183995 | 62.959410171651 |
| chr19 | 32597106 | 32597112 | - | 2.00792115374673 | 10.2329616785794 | 3.57968463110397 | 5.33554323488568 |
| chr19 | 32597309 | 32597312 | - | 3.01188173062009 | 5.68497871032191 | 4.77291284147196 | 7.46976052883995 |
| chr19 | 32597381 | 32597397 | - | 1.00396057687336 | 1.13699574206438 | 2.38645642073598 | 4.26843458790854 |
| chr19 | 32712666 | 32712692 | + | 10.0396057687336 | 11.3699574206438 | 5.96614105183995 | 7.46976052883995 |
| chr19 | 32712705 | 32712720 | + | 10.0396057687336 | 6.82197445238629 | 11.9322821036799 | 12.8053037637256 |
| chr19 | 32712820 | 32712826 | + | 2.00792115374673 | 5.68497871032191 | 5.96614105183995 | 40.5501285851312 |
| chr19 | 32713533 | 32713541 | + | 3.01188173062009 | 3.41098722619315 | 3.57968463110397 | 1.06710864697714 |
| chr19 | 32968963 | 32968988 | - | 1.00396057687336 | 11.3699574206438 | 3.57968463110397 | 3.20132594093141 |
| chr19 | 32970459 | 32970487 | - | 13.0514874993537 | 14.780944646837 | 11.9322821036799 | 1.06710864697714 |
| chr19 | 32970881 | 32970895 | - | 8.03168461498692 | 6.82197445238629 | 2.38645642073598 | 2.13421729395427 |
| chr19 | 32976678 | 32976685 | + | 3.01188173062009 | 1.13699574206438 | 3.57968463110397 | 5.33554323488568 |
| chr19 | 32976770 | 32976774 | + | 4.01584230749346 | 3.41098722619315 | 1.19322821036799 | 3.20132594093141 |
| chr19 | 32977017 | 32977033 | + | 5.01980288436682 | 9.09596593651506 | 4.77291284147196 | 2.13421729395427 |
| chr19 | 32977122 | 32977133 | + | 3.01188173062009 | 3.41098722619315 | 1.19322821036799 | 1.06710864697714 |
| chr19 | 32977184 | 32977187 | + | 4.01584230749346 | 5.68497871032191 | 7.15936926220794 | 7.46976052883995 |
| chr19 | 32977210 | 32977224 | + | 9.03564519186028 | 9.09596593651506 | 10.7390538933119 | 11.7381951167485 |
| chr19 | 32977271 | 32977290 | + | 3.01188173062009 | 2.27399148412876 | 9.54582568294392 | 7.46976052883995 |
| chr19 | 32977338 | 32977342 | + | 5.01980288436682 | 1.13699574206438 | 3.57968463110397 | 5.33554323488568 |
| chr19 | 32978629 | 32978797 | - | 29.1148567293276 | 15.9179403889013 | 11.9322821036799 | 8.53686917581709 |
| chr19 | 32978856 | 32978889 | - | 16.0633692299738 | 12.5069531627082 | 11.9322821036799 | 4.26843458790854 |
| chr19 | 32978950 | 32978979 | - | 39.1544624980612 | 29.5618892936739 | 7.15936926220794 | 5.33554323488568 |
| chr19 | 33130204 | 33130213 | + | 9.03564519186028 | 6.82197445238629 | 2.38645642073598 | 4.26843458790854 |
| chr19 | 33379768 | 33379837 | + | 6.02376346124019 | 14.780944646837 | 7.15936926220794 | 4.26843458790854 |
| chr19 | 33379873 | 33379969 | + | 9.03564519186028 | 54.5757956190903 | 10.7390538933119 | 5.33554323488568 |
| chr19 | 33380007 | 33380078 | + | 4.01584230749346 | 15.9179403889013 | 3.57968463110397 | 9.60397782279422 |
| chr19 | 33380306 | 33380307 | + | 2.00792115374673 | 1.13699574206438 | 1.19322821036799 | 1.06710864697714 |
| chr19 | 33380582 | 33380594 | + | 2.00792115374673 | 18.1919318730301 | 4.77291284147196 | 1.06710864697714 |
| chr19 | 33380642 | 33380709 | + | 12.0475269224804 | 36.3838637460602 | 4.77291284147196 | 7.46976052883995 |
| chr19 | 33380988 | 33381051 | + | 8.03168461498692 | 26.1509020674808 | 13.1255103140479 | 17.0737383516342 |
| chr19 | 33381142 | 33381156 | + | 3.01188173062009 | 5.68497871032191 | 3.57968463110397 | 2.13421729395427 |
| chr19 | 33381362 | 33381381 | + | 3.01188173062009 | 3.41098722619315 | 1.19322821036799 | 1.06710864697714 |
| chr19 | 33381412 | 33381432 | + | 5.01980288436682 | 12.5069531627082 | 5.96614105183995 | 1.06710864697714 |
| chr19 | 33381455 | 33381470 | + | 11.043566345607 | 19.3289276150945 | 15.5119667347839 | 5.33554323488568 |
| chr19 | 33381768 | 33381780 | + | 7.02772403811355 | 19.3289276150945 | 11.9322821036799 | 6.40265188186281 |
| chr19 | 33381909 | 33381943 | + | 11.043566345607 | 19.3289276150945 | 10.7390538933119 | 4.26843458790854 |
| chr19 | 33381980 | 33382051 | + | 12.0475269224804 | 48.8908169087684 | 17.8984231555198 | 19.2079556455884 |
| chr19 | 33382096 | 33382131 | + | 5.01980288436682 | 10.2329616785794 | 5.96614105183995 | 3.20132594093141 |
| chr19 | 33382160 | 33382198 | + | 9.03564519186028 | 11.3699574206438 | 11.9322821036799 | 4.26843458790854 |
| chr19 | 33382517 | 33382568 | + | 5.01980288436682 | 17.0549361309657 | 5.96614105183995 | 4.26843458790854 |
| chr19 | 33382591 | 33382611 | + | 6.02376346124019 | 13.6439489047726 | 4.77291284147196 | 2.13421729395427 |
| chr19 | 33387182 | 33387190 | - | 21.0831721143407 | 19.3289276150945 | 1.19322821036799 | 4.26843458790854 |
| chr19 | 33387313 | 33387317 | - | 5.01980288436682 | 5.68497871032191 | 1.19322821036799 | 1.06710864697714 |
| chr19 | 33813414 | 33813423 | + | 2.00792115374673 | 2.27399148412876 | 1.19322821036799 | 1.06710864697714 |
| chr19 | 33813564 | 33813581 | + | 25.0990144218341 | 36.3838637460602 | 15.5119667347839 | 6.40265188186281 |
| chr19 | 33814085 | 33814093 | + | 4.01584230749346 | 13.6439489047726 | 2.38645642073598 | 3.20132594093141 |
| chr19 | 33814127 | 33814171 | + | 5.01980288436682 | 12.5069531627082 | 1.19322821036799 | 1.06710864697714 |
| chr19 | 33814302 | 33814319 | + | 10.0396057687336 | 2.27399148412876 | 1.19322821036799 | 1.06710864697714 |
| chr19 | 33814635 | 33814652 | + | 1.00396057687336 | 5.68497871032191 | 8.35259747257593 | 2.13421729395427 |
| chr19 | 33815007 | 33815033 | + | 20.0792115374673 | 28.4248935516096 | 4.77291284147196 | 11.7381951167485 |
| chr19 | 33815640 | 33815654 | + | 6.02376346124019 | 9.09596593651506 | 2.38645642073598 | 5.33554323488568 |
| chr19 | 34399898 | 34399991 | + | 8.03168461498692 | 1.13699574206438 | 13.1255103140479 | 6.40265188186281 |
| chr19 | 34400145 | 34400343 | + | 26.1029749987075 | 23.876910583352 | 36.9900745214077 | 57.6238669367653 |
| chr19 | 34400491 | 34400528 | + | 5.01980288436682 | 2.27399148412876 | 2.38645642073598 | 5.33554323488568 |
| chr19 | 34400584 | 34400723 | + | 8.03168461498692 | 4.54798296825753 | 10.7390538933119 | 23.476390233497 |
| chr19 | 34401020 | 34401042 | + | 18.0712903837206 | 54.5757956190903 | 16.7051949451519 | 10.6710864697714 |
| chr19 | 34401153 | 34401175 | + | 3.01188173062009 | 1.13699574206438 | 3.57968463110397 | 7.46976052883995 |
| chr19 | 34401378 | 34401380 | + | 15.0594086531005 | 9.09596593651506 | 11.9322821036799 | 9.60397782279422 |
| chr19 | 34401517 | 34401527 | + | 8.03168461498692 | 1.13699574206438 | 2.38645642073598 | 1.06710864697714 |
| chr19 | 34402090 | 34402106 | + | 16.0633692299738 | 15.9179403889013 | 10.7390538933119 | 11.7381951167485 |
| chr19 | 34469224 | 34469281 | + | 7.02772403811355 | 9.09596593651506 | 20.2848795762558 | 29.8790421153598 |
| chr19 | 34469302 | 34469322 | + | 14.0554480762271 | 6.82197445238629 | 11.9322821036799 | 17.0737383516342 |
| chr19 | 34469383 | 34469405 | + | 1.00396057687336 | 4.54798296825753 | 2.38645642073598 | 3.20132594093141 |
| chr19 | 34469569 | 34469590 | + | 2.00792115374673 | 6.82197445238629 | 39.3765309421437 | 13.8724124107028 |
| chr19 | 34469654 | 34469667 | + | 1.00396057687336 | 1.13699574206438 | 10.7390538933119 | 9.60397782279422 |
| chr19 | 34469744 | 34469769 | + | 3.01188173062009 | 12.5069531627082 | 15.5119667347839 | 19.2079556455884 |
| chr19 | 34742438 | 34742441 | + | 1.00396057687336 | 2.27399148412876 | 2.38645642073598 | 2.13421729395427 |
| chr19 | 35026119 | 35026178 | + | 4.01584230749346 | 1.13699574206438 | 11.9322821036799 | 13.8724124107028 |
| chr19 | 35026367 | 35026450 | + | 2.00792115374673 | 3.41098722619315 | 7.15936926220794 | 7.46976052883995 |
| chr19 | 35153925 | 35153956 | + | 6.02376346124019 | 4.54798296825753 | 5.96614105183995 | 9.60397782279422 |
| chr19 | 35169564 | 35169614 | + | 12.0475269224804 | 6.82197445238629 | 13.1255103140479 | 37.3488026441998 |
| chr19 | 35169755 | 35169763 | + | 3.01188173062009 | 2.27399148412876 | 2.38645642073598 | 3.20132594093141 |
| chr19 | 35169803 | 35169831 | + | 69.2732798042621 | 70.4937360079917 | 58.4681823080315 | 149.395210576799 |
| chr19 | 35267822 | 35267878 | + | 22.087132691214 | 35.2468680039958 | 17.8984231555198 | 33.0803680562912 |
| chr19 | 35267905 | 35267943 | + | 19.0752509605939 | 30.6988850357383 | 14.3187385244159 | 18.1408469986113 |
| chr19 | 35279539 | 35279546 | + | 24.0950538449607 | 9.09596593651506 | 1.19322821036799 | 4.26843458790854 |
| chr19 | 35279658 | 35279666 | + | 8.03168461498692 | 4.54798296825753 | 1.19322821036799 | 14.9395210576799 |
| chr19 | 35279692 | 35279727 | + | 50.1980288436682 | 38.657855230189 | 27.4442488384638 | 28.8119334683827 |
| chr19 | 35279766 | 35279779 | + | 3.01188173062009 | 3.41098722619315 | 2.38645642073598 | 1.06710864697714 |
| chr19 | 35547347 | 35547473 | + | 85.336649034236 | 39.7948509722534 | 29.8307052591997 | 94.9726695809651 |
| chr19 | 35622967 | 35623016 | + | 6.02376346124019 | 6.82197445238629 | 11.9322821036799 | 9.60397782279422 |
| chr19 | 35623245 | 35623254 | + | 8.03168461498692 | 5.68497871032191 | 8.35259747257593 | 2.13421729395427 |
| chr19 | 35623298 | 35623338 | + | 6.02376346124019 | 13.6439489047726 | 9.54582568294392 | 3.20132594093141 |
| chr19 | 35623557 | 35623581 | + | 36.1425807674411 | 57.9867828452835 | 31.0239334695677 | 33.0803680562912 |
| chr19 | 35624386 | 35624396 | + | 3.01188173062009 | 22.7399148412876 | 4.77291284147196 | 4.26843458790854 |
| chr19 | 35624423 | 35624439 | + | 5.01980288436682 | 5.68497871032191 | 4.77291284147196 | 3.20132594093141 |
| chr19 | 35624728 | 35624731 | + | 15.0594086531005 | 14.780944646837 | 4.77291284147196 | 1.06710864697714 |
| chr19 | 35624926 | 35624935 | + | 11.043566345607 | 1.13699574206438 | 8.35259747257593 | 11.7381951167485 |
| chr19 | 35637616 | 35637658 | + | 9.03564519186028 | 5.68497871032191 | 5.96614105183995 | 10.6710864697714 |
| chr19 | 35658633 | 35658758 | + | 322.27134517635 | 203.522237829524 | 151.539982716735 | 509.010824608094 |
| chr19 | 35738838 | 35738844 | + | 2.00792115374673 | 4.54798296825753 | 2.38645642073598 | 5.33554323488568 |
| chr19 | 35739522 | 35739523 | - | 1.00396057687336 | 2.27399148412876 | 2.38645642073598 | 2.13421729395427 |
| chr19 | 35740473 | 35740625 | - | 367.449571135651 | 478.675207409105 | 286.374770488317 | 420.440806908991 |
| chr19 | 35742568 | 35742586 | - | 1.00396057687336 | 3.41098722619315 | 1.19322821036799 | 2.13421729395427 |
| chr19 | 35746713 | 35746762 | + | 5.01980288436682 | 3.41098722619315 | 7.15936926220794 | 4.26843458790854 |
| chr19 | 35746797 | 35746868 | + | 14.0554480762271 | 23.876910583352 | 23.8645642073598 | 34.1474767032683 |
| chr19 | 35746891 | 35746946 | + | 1701.71317780035 | 1247.28432904463 | 1072.71216112082 | 1420.32160912657 |
| chr19 | 35788736 | 35788745 | + | 3.01188173062009 | 3.41098722619315 | 5.96614105183995 | 3.20132594093141 |
| chr19 | 36009192 | 36009211 | - | 3.01188173062009 | 5.68497871032191 | 4.77291284147196 | 7.46976052883995 |
| chr19 | 36014486 | 36014562 | + | 49.1940682667949 | 53.438799877026 | 46.5359002043516 | 58.6909755837425 |
| chr19 | 36105013 | 36105033 | + | 22.087132691214 | 3.41098722619315 | 3.57968463110397 | 10.6710864697714 |
| chr19 | 36113734 | 36113815 | - | 63.249516343022 | 65.9457530397342 | 57.2749540976635 | 117.381951167485 |
| chr19 | 36114965 | 36114972 | + | 6.02376346124019 | 4.54798296825753 | 1.19322821036799 | 1.06710864697714 |
| chr19 | 36125665 | 36125869 | + | 110.43566345607 | 81.8636934286355 | 44.1494437836156 | 89.6371263460794 |
| chr19 | 36125895 | 36125922 | + | 10.0396057687336 | 5.68497871032191 | 10.7390538933119 | 1.06710864697714 |
| chr19 | 36149813 | 36150037 | + | 122.48319037855 | 54.5757956190903 | 72.7869208324474 | 64.0265188186281 |
| chr19 | 36150182 | 36150282 | + | 123.487150955424 | 56.8497871032191 | 9.54582568294392 | 51.2212150549025 |
| chr19 | 36150304 | 36150335 | + | 53.2099105742883 | 92.0966551072149 | 23.8645642073598 | 51.2212150549025 |
| chr19 | 36237343 | 36237346 | + | 7.02772403811355 | 13.6439489047726 | 17.8984231555198 | 9.60397782279422 |
| chr19 | 36237411 | 36237421 | + | 4.01584230749346 | 3.41098722619315 | 10.7390538933119 | 2.13421729395427 |
| chr19 | 36237512 | 36237539 | + | 1.00396057687336 | 2.27399148412876 | 5.96614105183995 | 13.8724124107028 |
| chr19 | 36237567 | 36237586 | + | 1.00396057687336 | 2.27399148412876 | 16.7051949451519 | 6.40265188186281 |
| chr19 | 36237842 | 36237864 | + | 7.02772403811355 | 5.68497871032191 | 16.7051949451519 | 8.53686917581709 |
| chr19 | 36237954 | 36237977 | + | 8.03168461498692 | 1.13699574206438 | 2.38645642073598 | 7.46976052883995 |
| chr19 | 36238053 | 36238066 | + | 2.00792115374673 | 7.95897019445067 | 2.38645642073598 | 2.13421729395427 |
| chr19 | 36238114 | 36238115 | + | 1.00396057687336 | 3.41098722619315 | 1.19322821036799 | 2.13421729395427 |
| chr19 | 36238189 | 36238235 | + | 10.0396057687336 | 12.5069531627082 | 22.6713359969918 | 16.006629704657 |
| chr19 | 36238290 | 36238343 | + | 9.03564519186028 | 21.6029190992233 | 7.15936926220794 | 12.8053037637256 |
| chr19 | 36238372 | 36238437 | + | 38.1505019211879 | 72.7677274921204 | 52.5020412561915 | 69.3620620535138 |
| chr19 | 36238500 | 36238532 | + | 14.0554480762271 | 22.7399148412876 | 19.0916513658878 | 21.3421729395427 |
| chr19 | 36238588 | 36238623 | + | 23.0910932680874 | 30.6988850357383 | 23.8645642073598 | 55.4896496428111 |
| chr19 | 36238694 | 36238715 | + | 9.03564519186028 | 11.3699574206438 | 15.5119667347839 | 12.8053037637256 |
| chr19 | 36511246 | 36511253 | - | 8.03168461498692 | 19.3289276150945 | 1.19322821036799 | 3.20132594093141 |
| chr19 | 36544497 | 36544502 | - | 1.00396057687336 | 1.13699574206438 | 4.77291284147196 | 2.13421729395427 |
| chr19 | 36545961 | 36545995 | - | 10.0396057687336 | 12.5069531627082 | 66.8207797806074 | 9.60397782279422 |
| chr19 | 36573340 | 36573386 | + | 30.1188173062009 | 54.5757956190903 | 36.9900745214077 | 52.2883237018796 |
| chr19 | 36573409 | 36573460 | + | 15.0594086531005 | 12.5069531627082 | 11.9322821036799 | 13.8724124107028 |
| chr19 | 36577399 | 36577444 | + | 1.00396057687336 | 1.13699574206438 | 3.57968463110397 | 8.53686917581709 |
| chr19 | 36720670 | 36720676 | + | 5.01980288436682 | 7.95897019445067 | 2.38645642073598 | 1.06710864697714 |
| chr19 | 36720795 | 36720805 | + | 4.01584230749346 | 3.41098722619315 | 1.19322821036799 | 2.13421729395427 |
| chr19 | 36720849 | 36720869 | + | 2.00792115374673 | 4.54798296825753 | 4.77291284147196 | 3.20132594093141 |
| chr19 | 36721036 | 36721092 | + | 19.0752509605939 | 22.7399148412876 | 8.35259747257593 | 11.7381951167485 |
| chr19 | 36721155 | 36721156 | + | 4.01584230749346 | 3.41098722619315 | 3.57968463110397 | 3.20132594093141 |
| chr19 | 36721205 | 36721214 | + | 24.0950538449607 | 23.876910583352 | 23.8645642073598 | 30.9461507623369 |
| chr19 | 36721254 | 36721276 | + | 7.02772403811355 | 11.3699574206438 | 2.38645642073598 | 2.13421729395427 |
| chr19 | 37696490 | 37696496 | - | 15.0594086531005 | 20.4659233571589 | 20.2848795762558 | 10.6710864697714 |
| chr19 | 38211084 | 38211144 | - | 16.0633692299738 | 10.2329616785794 | 15.5119667347839 | 23.476390233497 |
| chr19 | 38211197 | 38211220 | - | 4.01584230749346 | 3.41098722619315 | 16.7051949451519 | 6.40265188186281 |
| chr19 | 38211775 | 38211781 | - | 8.03168461498692 | 5.68497871032191 | 20.2848795762558 | 10.6710864697714 |
| chr19 | 38292430 | 38292432 | + | 7.02772403811355 | 3.41098722619315 | 1.19322821036799 | 1.06710864697714 |
| chr19 | 38305140 | 38305159 | - | 5.01980288436682 | 4.54798296825753 | 9.54582568294392 | 22.4092815865198 |
| chr19 | 38305384 | 38305510 | - | 10.0396057687336 | 4.54798296825753 | 71.5936926220794 | 62.959410171651 |
| chr19 | 38383366 | 38383470 | + | 73.2891221117556 | 64.8087572976698 | 137.221244192319 | 204.88486021961 |
| chr19 | 38383674 | 38383715 | + | 17.0673298068472 | 17.0549361309657 | 17.8984231555198 | 59.7580842307196 |
| chr19 | 38383761 | 38383772 | + | 63.249516343022 | 78.4527062024424 | 120.516049247167 | 216.623055336359 |
| chr19 | 38383794 | 38383805 | + | 22.087132691214 | 37.5208594881246 | 62.0478669391355 | 89.6371263460794 |
| chr19 | 39383613 | 39383635 | + | 6.02376346124019 | 11.3699574206438 | 13.1255103140479 | 4.26843458790854 |
| chr19 | 39384709 | 39384734 | + | 12.0475269224804 | 10.2329616785794 | 5.96614105183995 | 5.33554323488568 |
| chr19 | 39385613 | 39385662 | + | 6.02376346124019 | 4.54798296825753 | 2.38645642073598 | 3.20132594093141 |
| chr19 | 39385874 | 39385928 | - | 3.01188173062009 | 4.54798296825753 | 3.57968463110397 | 1.06710864697714 |
| chr19 | 39409246 | 39409256 | + | 7.02772403811355 | 7.95897019445067 | 1.19322821036799 | 3.20132594093141 |
| chr19 | 39425460 | 39425483 | + | 1.00396057687336 | 9.09596593651506 | 4.77291284147196 | 6.40265188186281 |
| chr19 | 39433249 | 39433366 | - | 214.8475634509 | 229.673139897005 | 151.539982716735 | 148.328101929822 |
| chr19 | 39476575 | 39476622 | + | 3.01188173062009 | 5.68497871032191 | 1.19322821036799 | 8.53686917581709 |
| chr19 | 39489826 | 39489901 | + | 31.1227778830743 | 13.6439489047726 | 31.0239334695677 | 30.9461507623369 |
| chr19 | 39508359 | 39508370 | + | 3.01188173062009 | 1.13699574206438 | 1.19322821036799 | 1.06710864697714 |
| chr19 | 39531565 | 39531571 | - | 1.00396057687336 | 2.27399148412876 | 2.38645642073598 | 1.06710864697714 |
| chr19 | 39538944 | 39538992 | - | 13.0514874993537 | 9.09596593651506 | 5.96614105183995 | 6.40265188186281 |
| chr19 | 39539069 | 39539091 | - | 4.01584230749346 | 5.68497871032191 | 1.19322821036799 | 3.20132594093141 |
| chr19 | 39539236 | 39539266 | - | 109.431702879197 | 112.562578464374 | 8.35259747257593 | 46.952780466994 |
| chr19 | 40215428 | 40215429 | + | 5.01980288436682 | 5.68497871032191 | 2.38645642073598 | 1.06710864697714 |
| chr19 | 40231262 | 40231269 | - | 2.00792115374673 | 1.13699574206438 | 2.38645642073598 | 6.40265188186281 |
| chr19 | 40232353 | 40232476 | - | 110.43566345607 | 83.0006891706999 | 46.5359002043516 | 58.6909755837425 |
| chr19 | 40233395 | 40233399 | - | 11.043566345607 | 7.95897019445067 | 1.19322821036799 | 3.20132594093141 |
| chr19 | 40233792 | 40233842 | - | 9.03564519186028 | 7.95897019445067 | 8.35259747257593 | 1.06710864697714 |
| chr19 | 40320360 | 40320381 | - | 7.02772403811355 | 11.3699574206438 | 2.38645642073598 | 1.06710864697714 |
| chr19 | 40321118 | 40321153 | - | 41.1623836518079 | 44.3428339405109 | 19.0916513658878 | 38.4159112911769 |
| chr19 | 40321195 | 40321210 | - | 5.01980288436682 | 6.82197445238629 | 2.38645642073598 | 5.33554323488568 |
| chr19 | 40321586 | 40321589 | - | 4.01584230749346 | 3.41098722619315 | 2.38645642073598 | 2.13421729395427 |
| chr19 | 40356865 | 40356868 | + | 1.00396057687336 | 4.54798296825753 | 1.19322821036799 | 3.20132594093141 |
| chr19 | 40357011 | 40357014 | + | 2.00792115374673 | 2.27399148412876 | 2.38645642073598 | 4.26843458790854 |
| chr19 | 40369906 | 40369927 | + | 1.00396057687336 | 4.54798296825753 | 10.7390538933119 | 8.53686917581709 |
| chr19 | 40378395 | 40378457 | + | 20.0792115374673 | 29.5618892936739 | 29.8307052591997 | 50.1541064079254 |
| chr19 | 40422551 | 40422574 | - | 2.00792115374673 | 4.54798296825753 | 2.38645642073598 | 6.40265188186281 |
| chr19 | 40440916 | 40440923 | - | 4.01584230749346 | 2.27399148412876 | 1.19322821036799 | 4.26843458790854 |
| chr19 | 40447809 | 40447874 | - | 85.336649034236 | 118.247557174696 | 213.58784965587 | 426.843458790854 |
| chr19 | 40591278 | 40591321 | + | 9.03564519186028 | 4.54798296825753 | 2.38645642073598 | 4.26843458790854 |
| chr19 | 40591358 | 40591361 | + | 2.00792115374673 | 6.82197445238629 | 4.77291284147196 | 10.6710864697714 |
| chr19 | 40629744 | 40629769 | + | 11.043566345607 | 7.95897019445067 | 21.4781077866238 | 22.4092815865198 |
| chr19 | 40666515 | 40666533 | - | 5.01980288436682 | 2.27399148412876 | 5.96614105183995 | 11.7381951167485 |
| chr19 | 40666645 | 40666673 | - | 5.01980288436682 | 1.13699574206438 | 7.15936926220794 | 9.60397782279422 |
| chr19 | 40666781 | 40666820 | - | 31.1227778830743 | 25.0139063254164 | 19.0916513658878 | 42.6843458790854 |
| chr19 | 40691568 | 40691580 | - | 2.00792115374673 | 5.68497871032191 | 7.15936926220794 | 9.60397782279422 |
| chr19 | 40740701 | 40740712 | + | 4.01584230749346 | 1.13699574206438 | 5.96614105183995 | 7.46976052883995 |
| chr19 | 40740994 | 40741009 | - | 3.01188173062009 | 2.27399148412876 | 2.38645642073598 | 2.13421729395427 |
| chr19 | 40765236 | 40765291 | + | 11.043566345607 | 17.0549361309657 | 14.3187385244159 | 16.006629704657 |
| chr19 | 40765315 | 40765373 | + | 19.0752509605939 | 34.1098722619315 | 19.0916513658878 | 21.3421729395427 |
| chr19 | 40780448 | 40780450 | + | 2.00792115374673 | 1.13699574206438 | 4.77291284147196 | 2.13421729395427 |
| chr19 | 40796649 | 40796650 | + | 1.00396057687336 | 1.13699574206438 | 5.96614105183995 | 1.06710864697714 |
| chr19 | 40800494 | 40800531 | + | 8.03168461498692 | 3.41098722619315 | 5.96614105183995 | 5.33554323488568 |
| chr19 | 40800750 | 40800752 | + | 4.01584230749346 | 1.13699574206438 | 3.57968463110397 | 2.13421729395427 |
| chr19 | 40800875 | 40800881 | + | 3.01188173062009 | 3.41098722619315 | 3.57968463110397 | 5.33554323488568 |
| chr19 | 40800903 | 40800910 | + | 2.00792115374673 | 1.13699574206438 | 4.77291284147196 | 3.20132594093141 |
| chr19 | 40800983 | 40801020 | + | 3.01188173062009 | 2.27399148412876 | 4.77291284147196 | 12.8053037637256 |
| chr19 | 40801181 | 40801182 | + | 1.00396057687336 | 1.13699574206438 | 5.96614105183995 | 1.06710864697714 |
| chr19 | 40801282 | 40801301 | + | 5.01980288436682 | 6.82197445238629 | 3.57968463110397 | 1.06710864697714 |
| chr19 | 40801380 | 40801406 | + | 1.00396057687336 | 2.27399148412876 | 10.7390538933119 | 10.6710864697714 |
| chr19 | 40806559 | 40806590 | + | 4.01584230749346 | 11.3699574206438 | 8.35259747257593 | 8.53686917581709 |
| chr19 | 40806620 | 40806640 | + | 4.01584230749346 | 4.54798296825753 | 4.77291284147196 | 4.26843458790854 |
| chr19 | 40807170 | 40807207 | + | 8.03168461498692 | 10.2329616785794 | 19.0916513658878 | 8.53686917581709 |
| chr19 | 40807210 | 40807252 | + | 2.00792115374673 | 14.780944646837 | 5.96614105183995 | 4.26843458790854 |
| chr19 | 40807481 | 40807552 | + | 10.0396057687336 | 5.68497871032191 | 17.8984231555198 | 21.3421729395427 |
| chr19 | 40807875 | 40807921 | + | 5.01980288436682 | 3.41098722619315 | 5.96614105183995 | 17.0737383516342 |
| chr19 | 40808367 | 40808410 | + | 13.0514874993537 | 26.1509020674808 | 15.5119667347839 | 36.2816939972226 |
| chr19 | 41261732 | 41261735 | + | 18.0712903837206 | 18.1919318730301 | 4.77291284147196 | 16.006629704657 |
| chr19 | 41306646 | 41306743 | + | 14.0554480762271 | 3.41098722619315 | 13.1255103140479 | 17.0737383516342 |
| chr19 | 41306766 | 41306804 | + | 2.00792115374673 | 3.41098722619315 | 1.19322821036799 | 11.7381951167485 |
| chr19 | 41306882 | 41306903 | + | 13.0514874993537 | 5.68497871032191 | 9.54582568294392 | 19.2079556455884 |
| chr19 | 41306955 | 41307016 | + | 17.0673298068472 | 12.5069531627082 | 17.8984231555198 | 26.6777161744284 |
| chr19 | 41307064 | 41307109 | + | 95.3762548029696 | 72.7677274921204 | 131.255103140479 | 217.690163983336 |
| chr19 | 41307146 | 41307171 | + | 6.02376346124019 | 11.3699574206438 | 17.8984231555198 | 21.3421729395427 |
| chr19 | 41307202 | 41307209 | + | 2.00792115374673 | 4.54798296825753 | 2.38645642073598 | 8.53686917581709 |
| chr19 | 41307324 | 41307358 | + | 7.02772403811355 | 5.68497871032191 | 11.9322821036799 | 8.53686917581709 |
| chr19 | 41307410 | 41307445 | + | 8.03168461498692 | 7.95897019445067 | 15.5119667347839 | 39.483019938154 |
| chr19 | 41307492 | 41307537 | + | 11.043566345607 | 10.2329616785794 | 13.1255103140479 | 7.46976052883995 |
| chr19 | 41330975 | 41330995 | - | 3.01188173062009 | 2.27399148412876 | 9.54582568294392 | 6.40265188186281 |
| chr19 | 41386395 | 41386458 | - | 95.3762548029696 | 112.562578464374 | 112.163451774591 | 126.985928990279 |
| chr19 | 41424711 | 41424713 | + | 1.00396057687336 | 1.13699574206438 | 2.38645642073598 | 9.60397782279422 |
| chr19 | 41424863 | 41424924 | + | 2.00792115374673 | 9.09596593651506 | 7.15936926220794 | 19.2079556455884 |
| chr19 | 41424946 | 41424980 | + | 11.043566345607 | 19.3289276150945 | 23.8645642073598 | 13.8724124107028 |
| chr19 | 41431352 | 41431391 | - | 12.0475269224804 | 19.3289276150945 | 7.15936926220794 | 3.20132594093141 |
| chr19 | 41431427 | 41431483 | - | 2.00792115374673 | 3.41098722619315 | 1.19322821036799 | 1.06710864697714 |
| chr19 | 41433180 | 41433199 | - | 42.1663442286813 | 28.4248935516096 | 64.4343233598714 | 40.5501285851312 |
| chr19 | 41869696 | 41869741 | + | 181.716864414079 | 219.440178218426 | 156.312895558207 | 346.810310267569 |
| chr19 | 41871388 | 41871421 | + | 58.2297134586551 | 384.304560817761 | 497.576163723452 | 78.966039876308 |
| chr19 | 41872384 | 41872389 | + | 3.01188173062009 | 3.41098722619315 | 2.38645642073598 | 2.13421729395427 |
| chr19 | 41872453 | 41872456 | + | 2.00792115374673 | 1.13699574206438 | 3.57968463110397 | 1.06710864697714 |
| chr19 | 41872487 | 41872528 | + | 10.0396057687336 | 1.13699574206438 | 3.57968463110397 | 2.13421729395427 |
| chr19 | 41872824 | 41872850 | + | 9.03564519186028 | 6.82197445238629 | 2.38645642073598 | 10.6710864697714 |
| chr19 | 41872879 | 41872882 | + | 3.01188173062009 | 2.27399148412876 | 2.38645642073598 | 5.33554323488568 |
| chr19 | 41956705 | 41956736 | - | 14.0554480762271 | 18.1919318730301 | 7.15936926220794 | 8.53686917581709 |
| chr19 | 41956797 | 41956876 | - | 22.087132691214 | 9.09596593651506 | 17.8984231555198 | 13.8724124107028 |
| chr19 | 42199027 | 42199056 | - | 4.01584230749346 | 3.41098722619315 | 9.54582568294392 | 2.13421729395427 |
| chr19 | 42199161 | 42199193 | - | 9.03564519186028 | 1.13699574206438 | 29.8307052591997 | 23.476390233497 |
| chr19 | 42230214 | 42230285 | - | 26.1029749987075 | 26.1509020674808 | 35.7968463110397 | 40.5501285851312 |
| chr19 | 42230423 | 42230428 | - | 3.01188173062009 | 3.41098722619315 | 13.1255103140479 | 11.7381951167485 |
| chr19 | 42243287 | 42243338 | + | 23.0910932680874 | 37.5208594881246 | 35.7968463110397 | 49.0869977609482 |
| chr19 | 42247588 | 42247633 | - | 33.130699036821 | 43.2058381984465 | 29.8307052591997 | 39.483019938154 |
| chr19 | 42247655 | 42247682 | - | 10.0396057687336 | 6.82197445238629 | 10.7390538933119 | 17.0737383516342 |
| chr19 | 42247859 | 42247900 | - | 5.01980288436682 | 3.41098722619315 | 3.57968463110397 | 4.26843458790854 |
| chr19 | 42247976 | 42247996 | - | 5.01980288436682 | 5.68497871032191 | 1.19322821036799 | 3.20132594093141 |
| chr19 | 42248149 | 42248234 | - | 22.087132691214 | 31.8358807778027 | 21.4781077866238 | 16.006629704657 |
| chr19 | 42295566 | 42295567 | + | 4.01584230749346 | 4.54798296825753 | 1.19322821036799 | 2.13421729395427 |
| chr19 | 42295737 | 42295766 | + | 3.01188173062009 | 2.27399148412876 | 1.19322821036799 | 5.33554323488568 |
| chr19 | 42297054 | 42297071 | - | 2.00792115374673 | 1.13699574206438 | 4.77291284147196 | 8.53686917581709 |
| chr19 | 42377844 | 42377969 | + | 1.00396057687336 | 1.13699574206438 | 11.9322821036799 | 8.53686917581709 |
| chr19 | 42377994 | 42378060 | + | 1.00396057687336 | 1.13699574206438 | 2.38645642073598 | 2.13421729395427 |
| chr19 | 42397146 | 42397193 | + | 20.0792115374673 | 20.4659233571589 | 7.15936926220794 | 10.6710864697714 |
| chr19 | 43475325 | 43475326 | - | 1.00396057687336 | 1.13699574206438 | 2.38645642073598 | 2.13421729395427 |
| chr19 | 43506743 | 43506796 | - | 51.2019894205416 | 63.6717615556054 | 21.4781077866238 | 24.5434988804741 |
| chr19 | 43543338 | 43543342 | - | 1.00396057687336 | 2.27399148412876 | 2.38645642073598 | 2.13421729395427 |
| chr19 | 43585963 | 43585994 | - | 5.01980288436682 | 5.68497871032191 | 3.57968463110397 | 5.33554323488568 |
| chr19 | 43592012 | 43592014 | - | 2.00792115374673 | 2.27399148412876 | 3.57968463110397 | 2.13421729395427 |
| chr19 | 43601120 | 43601123 | + | 2.00792115374673 | 4.54798296825753 | 1.19322821036799 | 6.40265188186281 |
| chr19 | 43607509 | 43607599 | - | 118.467348071057 | 162.590391115207 | 190.916513658878 | 442.850088495511 |
| chr19 | 43622954 | 43622970 | - | 5.01980288436682 | 3.41098722619315 | 8.35259747257593 | 4.26843458790854 |
| chr19 | 43646300 | 43646311 | - | 9.03564519186028 | 13.6439489047726 | 1.19322821036799 | 1.06710864697714 |
| chr19 | 43648679 | 43648704 | - | 4.01584230749346 | 14.780944646837 | 2.38645642073598 | 5.33554323488568 |
| chr19 | 43731177 | 43731179 | - | 6.02376346124019 | 27.2878978095452 | 3.57968463110397 | 1.06710864697714 |
| chr19 | 43731204 | 43731213 | - | 2.00792115374673 | 5.68497871032191 | 7.15936926220794 | 2.13421729395427 |
| chr19 | 43766588 | 43766596 | - | 14.0554480762271 | 13.6439489047726 | 9.54582568294392 | 11.7381951167485 |
| chr19 | 43851010 | 43851011 | + | 1.00396057687336 | 4.54798296825753 | 2.38645642073598 | 2.13421729395427 |
| chr19 | 44663085 | 44663143 | + | 10.0396057687336 | 18.1919318730301 | 1.19322821036799 | 1.06710864697714 |
| chr19 | 44888504 | 44888561 | + | 2.00792115374673 | 1.13699574206438 | 8.35259747257593 | 13.8724124107028 |
| chr19 | 44888604 | 44888612 | + | 3.01188173062009 | 2.27399148412876 | 25.0577924177278 | 12.8053037637256 |
| chr19 | 44889137 | 44889168 | + | 3.01188173062009 | 3.41098722619315 | 9.54582568294392 | 11.7381951167485 |
| chr19 | 44903371 | 44903395 | + | 1.00396057687336 | 1.13699574206438 | 2.38645642073598 | 3.20132594093141 |
| chr19 | 44903539 | 44903576 | + | 33.130699036821 | 19.3289276150945 | 20.2848795762558 | 36.2816939972226 |
| chr19 | 44903631 | 44903643 | + | 7.02772403811355 | 15.9179403889013 | 13.1255103140479 | 5.33554323488568 |
| chr19 | 44993197 | 44993199 | + | 3.01188173062009 | 1.13699574206438 | 1.19322821036799 | 5.33554323488568 |
| chr19 | 45070830 | 45070906 | + | 159.629731722865 | 129.61751459534 | 102.617626091647 | 158.999188399593 |
| chr19 | 45146795 | 45146820 | + | 7.02772403811355 | 7.95897019445067 | 9.54582568294392 | 5.33554323488568 |
| chr19 | 45146940 | 45146981 | + | 11.043566345607 | 6.82197445238629 | 8.35259747257593 | 4.26843458790854 |
| chr19 | 45147016 | 45147099 | + | 57.2257528817818 | 47.753821166704 | 56.0817258872955 | 48.0198891139711 |
| chr19 | 45147215 | 45147237 | + | 2.00792115374673 | 1.13699574206438 | 1.19322821036799 | 1.06710864697714 |
| chr19 | 45150512 | 45150513 | - | 9.03564519186028 | 6.82197445238629 | 11.9322821036799 | 4.26843458790854 |
| chr19 | 45181755 | 45181774 | + | 6.02376346124019 | 7.95897019445067 | 13.1255103140479 | 13.8724124107028 |
| chr19 | 45280383 | 45280401 | + | 1.00396057687336 | 2.27399148412876 | 1.19322821036799 | 2.13421729395427 |
| chr19 | 45303002 | 45303038 | + | 3.01188173062009 | 1.13699574206438 | 25.0577924177278 | 6.40265188186281 |
| chr19 | 45349870 | 45349925 | - | 44.174265382428 | 40.9318467143178 | 68.0140079909754 | 122.717494402371 |
| chr19 | 45350406 | 45350498 | - | 7.02772403811355 | 6.82197445238629 | 27.4442488384638 | 24.5434988804741 |
| chr19 | 45350838 | 45350881 | - | 2.00792115374673 | 1.13699574206438 | 8.35259747257593 | 39.483019938154 |
| chr19 | 45351240 | 45351247 | - | 9.03564519186028 | 3.41098722619315 | 34.6036181006717 | 13.8724124107028 |
| chr19 | 45351612 | 45351622 | - | 3.01188173062009 | 4.54798296825753 | 2.38645642073598 | 3.20132594093141 |
| chr19 | 45407366 | 45407388 | - | 5.01980288436682 | 17.0549361309657 | 11.9322821036799 | 35.2145853502455 |
| chr19 | 45408707 | 45408709 | - | 1.00396057687336 | 1.13699574206438 | 1.19322821036799 | 2.13421729395427 |
| chr19 | 45408796 | 45408824 | - | 1.00396057687336 | 2.27399148412876 | 8.35259747257593 | 9.60397782279422 |
| chr19 | 45409647 | 45409719 | - | 12.0475269224804 | 12.5069531627082 | 14.3187385244159 | 20.2750642925656 |
| chr19 | 45485503 | 45485542 | - | 1.00396057687336 | 1.13699574206438 | 2.38645642073598 | 13.8724124107028 |
| chr19 | 45502261 | 45502264 | + | 7.02772403811355 | 5.68497871032191 | 19.0916513658878 | 8.53686917581709 |
| chr19 | 45502298 | 45502303 | + | 1.00396057687336 | 13.6439489047726 | 3.57968463110397 | 5.33554323488568 |
| chr19 | 45526179 | 45526194 | + | 1.00396057687336 | 1.13699574206438 | 5.96614105183995 | 6.40265188186281 |
| chr19 | 45526255 | 45526280 | + | 2.00792115374673 | 2.27399148412876 | 13.1255103140479 | 51.2212150549025 |
| chr19 | 45526310 | 45526329 | + | 3.01188173062009 | 1.13699574206438 | 8.35259747257593 | 11.7381951167485 |
| chr19 | 45526351 | 45526377 | + | 12.0475269224804 | 7.95897019445067 | 14.3187385244159 | 28.8119334683827 |
| chr19 | 45526463 | 45526484 | + | 4.01584230749346 | 1.13699574206438 | 16.7051949451519 | 24.5434988804741 |
| chr19 | 45526571 | 45526593 | + | 3.01188173062009 | 4.54798296825753 | 17.8984231555198 | 29.8790421153598 |
| chr19 | 45526715 | 45526759 | + | 6.02376346124019 | 2.27399148412876 | 14.3187385244159 | 11.7381951167485 |
| chr19 | 45526886 | 45526971 | + | 13.0514874993537 | 9.09596593651506 | 17.8984231555198 | 35.2145853502455 |
| chr19 | 45549180 | 45549188 | - | 4.01584230749346 | 9.09596593651506 | 17.8984231555198 | 10.6710864697714 |
| chr19 | 45549281 | 45549310 | - | 3.01188173062009 | 3.41098722619315 | 2.38645642073598 | 7.46976052883995 |
| chr19 | 45550300 | 45550307 | - | 1.00396057687336 | 2.27399148412876 | 1.19322821036799 | 1.06710864697714 |
| chr19 | 45609455 | 45609509 | - | 30.1188173062009 | 31.8358807778027 | 14.3187385244159 | 18.1408469986113 |
| chr19 | 45609539 | 45609565 | - | 27.1069355755808 | 38.657855230189 | 13.1255103140479 | 33.0803680562912 |
| chr19 | 45609616 | 45609674 | - | 6.02376346124019 | 9.09596593651506 | 8.35259747257593 | 8.53686917581709 |
| chr19 | 45641438 | 45641673 | + | 510.011973051669 | 603.744739036187 | 76.3666054635513 | 97.1068868749194 |
| chr19 | 45687475 | 45687604 | - | 354.398083636298 | 418.414433079693 | 541.725607507067 | 733.103640473292 |
| chr19 | 45687655 | 45687719 | - | 8.03168461498692 | 5.68497871032191 | 14.3187385244159 | 13.8724124107028 |
| chr19 | 45710680 | 45710702 | - | 1.00396057687336 | 6.82197445238629 | 2.38645642073598 | 2.13421729395427 |
| chr19 | 45710835 | 45710857 | - | 12.0475269224804 | 9.09596593651506 | 4.77291284147196 | 6.40265188186281 |
| chr19 | 45711461 | 45711482 | - | 4.01584230749346 | 6.82197445238629 | 2.38645642073598 | 9.60397782279422 |
| chr19 | 45765758 | 45765766 | - | 10.0396057687336 | 3.41098722619315 | 2.38645642073598 | 2.13421729395427 |
| chr19 | 45770613 | 45770636 | - | 2.00792115374673 | 1.13699574206438 | 2.38645642073598 | 2.13421729395427 |
| chr19 | 45779496 | 45779512 | - | 3.01188173062009 | 2.27399148412876 | 1.19322821036799 | 1.06710864697714 |
| chr19 | 45815470 | 45815514 | - | 18.0712903837206 | 22.7399148412876 | 17.8984231555198 | 23.476390233497 |
| chr19 | 45818948 | 45818955 | - | 3.01188173062009 | 7.95897019445067 | 2.38645642073598 | 1.06710864697714 |
| chr19 | 45890048 | 45890060 | - | 6.02376346124019 | 4.54798296825753 | 5.96614105183995 | 5.33554323488568 |
| chr19 | 46390780 | 46390815 | + | 23.0910932680874 | 11.3699574206438 | 4.77291284147196 | 13.8724124107028 |
| chr19 | 46609123 | 46609422 | + | 45.1782259593014 | 30.6988850357383 | 122.902505667903 | 109.912190638645 |
| chr19 | 46609445 | 46609503 | + | 9.03564519186028 | 6.82197445238629 | 42.9562155732476 | 17.0737383516342 |
| chr19 | 46609529 | 46609533 | + | 6.02376346124019 | 1.13699574206438 | 7.15936926220794 | 3.20132594093141 |
| chr19 | 46609565 | 46609785 | + | 47.1861471130481 | 27.2878978095452 | 187.336829027774 | 107.777973344691 |
| chr19 | 46609849 | 46609857 | + | 2.00792115374673 | 1.13699574206438 | 3.57968463110397 | 6.40265188186281 |
| chr19 | 46609894 | 46609934 | + | 1.00396057687336 | 1.13699574206438 | 4.77291284147196 | 9.60397782279422 |
| chr19 | 46609965 | 46610032 | + | 12.0475269224804 | 4.54798296825753 | 22.6713359969918 | 14.9395210576799 |
| chr19 | 46610386 | 46610435 | + | 1.00396057687336 | 2.27399148412876 | 11.9322821036799 | 20.2750642925656 |
| chr19 | 46610575 | 46610677 | + | 63.249516343022 | 43.2058381984465 | 96.6514850398071 | 220.891489924267 |
| chr19 | 46610704 | 46610753 | + | 19.0752509605939 | 26.1509020674808 | 41.7629873628796 | 56.5567582897882 |
| chr19 | 46719536 | 46719637 | - | 5.01980288436682 | 2.27399148412876 | 2.38645642073598 | 3.20132594093141 |
| chr19 | 46719659 | 46719700 | - | 2.00792115374673 | 2.27399148412876 | 1.19322821036799 | 1.06710864697714 |
| chr19 | 46774949 | 46775072 | - | 126.499032686044 | 160.316399631078 | 161.085808399679 | 196.347991043793 |
| chr19 | 46775107 | 46775240 | - | 995.928892258378 | 1783.94631929902 | 946.229970821816 | 1171.68529438089 |
| chr19 | 46775261 | 46775484 | - | 192.760430759686 | 267.19399938513 | 350.809093848189 | 498.339738138322 |
| chr19 | 46838195 | 46838274 | - | 86.3406096111093 | 77.315710460378 | 63.2410951495034 | 130.187254931211 |
| chr19 | 46838303 | 46838308 | - | 1.00396057687336 | 2.27399148412876 | 1.19322821036799 | 3.20132594093141 |
| chr19 | 47001355 | 47001386 | + | 4.01584230749346 | 4.54798296825753 | 1.19322821036799 | 1.06710864697714 |
| chr19 | 47001783 | 47001849 | + | 11.043566345607 | 13.6439489047726 | 5.96614105183995 | 1.06710864697714 |
| chr19 | 47003414 | 47003434 | + | 3.01188173062009 | 1.13699574206438 | 3.57968463110397 | 1.06710864697714 |
| chr19 | 47003619 | 47003644 | + | 4.01584230749346 | 4.54798296825753 | 2.38645642073598 | 5.33554323488568 |
| chr19 | 47003749 | 47003766 | + | 2.00792115374673 | 3.41098722619315 | 2.38645642073598 | 3.20132594093141 |
| chr19 | 47003799 | 47003808 | + | 4.01584230749346 | 5.68497871032191 | 2.38645642073598 | 2.13421729395427 |
| chr19 | 47004098 | 47004114 | + | 39.1544624980612 | 31.8358807778027 | 22.6713359969918 | 8.53686917581709 |
| chr19 | 47004153 | 47004211 | + | 20.0792115374673 | 11.3699574206438 | 14.3187385244159 | 5.33554323488568 |
| chr19 | 47004349 | 47004366 | + | 3.01188173062009 | 2.27399148412876 | 4.77291284147196 | 13.8724124107028 |
| chr19 | 47004429 | 47004454 | + | 4.01584230749346 | 1.13699574206438 | 4.77291284147196 | 2.13421729395427 |
| chr19 | 47004492 | 47004532 | + | 4.01584230749346 | 3.41098722619315 | 1.19322821036799 | 2.13421729395427 |
| chr19 | 47004897 | 47004931 | + | 8.03168461498692 | 10.2329616785794 | 5.96614105183995 | 4.26843458790854 |
| chr19 | 47045933 | 47045943 | - | 3.01188173062009 | 11.3699574206438 | 2.38645642073598 | 6.40265188186281 |
| chr19 | 47064435 | 47064439 | - | 2.00792115374673 | 1.13699574206438 | 1.19322821036799 | 1.06710864697714 |
| chr19 | 47065241 | 47065257 | - | 2.00792115374673 | 1.13699574206438 | 1.19322821036799 | 7.46976052883995 |
| chr19 | 47065307 | 47065332 | - | 2.00792115374673 | 2.27399148412876 | 2.38645642073598 | 6.40265188186281 |
| chr19 | 47065363 | 47065393 | - | 3.01188173062009 | 2.27399148412876 | 1.19322821036799 | 14.9395210576799 |
| chr19 | 47066256 | 47066278 | - | 43.1703048055547 | 18.1919318730301 | 28.6374770488317 | 14.9395210576799 |
| chr19 | 47209203 | 47209225 | + | 4.01584230749346 | 1.13699574206438 | 7.15936926220794 | 4.26843458790854 |
| chr19 | 47209828 | 47209869 | + | 4.01584230749346 | 2.27399148412876 | 2.38645642073598 | 2.13421729395427 |
| chr19 | 47209996 | 47210011 | + | 3.01188173062009 | 4.54798296825753 | 3.57968463110397 | 4.26843458790854 |
| chr19 | 47210041 | 47210136 | + | 34.1346596136944 | 64.8087572976698 | 63.2410951495034 | 78.966039876308 |
| chr19 | 47220887 | 47221034 | - | 321.267384599477 | 578.73083271077 | 767.245739266617 | 904.908132636611 |
| chr19 | 47221075 | 47221128 | - | 3.01188173062009 | 11.3699574206438 | 15.5119667347839 | 45.8856718200168 |
| chr19 | 47221227 | 47221239 | - | 2.00792115374673 | 3.41098722619315 | 11.9322821036799 | 9.60397782279422 |
| chr19 | 47221294 | 47221300 | - | 1.00396057687336 | 1.13699574206438 | 1.19322821036799 | 3.20132594093141 |
| chr19 | 47221427 | 47221437 | - | 10.0396057687336 | 10.2329616785794 | 21.4781077866238 | 8.53686917581709 |
| chr19 | 47221569 | 47221585 | - | 1.00396057687336 | 1.13699574206438 | 4.77291284147196 | 2.13421729395427 |
| chr19 | 47221762 | 47221765 | - | 2.00792115374673 | 1.13699574206438 | 3.57968463110397 | 2.13421729395427 |
| chr19 | 47275302 | 47275366 | + | 4.01584230749346 | 6.82197445238629 | 35.7968463110397 | 20.2750642925656 |
| chr19 | 47275490 | 47275514 | + | 3.01188173062009 | 2.27399148412876 | 20.2848795762558 | 32.0132594093141 |
| chr19 | 47275544 | 47275565 | + | 8.03168461498692 | 20.4659233571589 | 60.8546387287675 | 78.966039876308 |
| chr19 | 47382204 | 47382206 | + | 1.00396057687336 | 1.13699574206438 | 1.19322821036799 | 1.06710864697714 |
| chr19 | 47382237 | 47382250 | + | 1.00396057687336 | 1.13699574206438 | 3.57968463110397 | 4.26843458790854 |
| chr19 | 47484277 | 47484299 | + | 10.0396057687336 | 12.5069531627082 | 9.54582568294392 | 18.1408469986113 |
| chr19 | 47484364 | 47484410 | + | 18.0712903837206 | 26.1509020674808 | 19.0916513658878 | 18.1408469986113 |
| chr19 | 47487664 | 47487692 | - | 9.03564519186028 | 9.09596593651506 | 4.77291284147196 | 6.40265188186281 |
| chr19 | 47487748 | 47487788 | - | 3.01188173062009 | 1.13699574206438 | 2.38645642073598 | 1.06710864697714 |
| chr19 | 47487923 | 47487994 | - | 16.0633692299738 | 9.09596593651506 | 32.2171616799357 | 73.6304966414224 |
| chr19 | 47488008 | 47488028 | - | 2.00792115374673 | 1.13699574206438 | 4.77291284147196 | 4.26843458790854 |
| chr19 | 47488090 | 47488117 | - | 18.0712903837206 | 4.54798296825753 | 40.5697591525116 | 8.53686917581709 |
| chr19 | 47488152 | 47488154 | - | 1.00396057687336 | 1.13699574206438 | 2.38645642073598 | 1.06710864697714 |
| chr19 | 47488213 | 47488236 | - | 7.02772403811355 | 4.54798296825753 | 10.7390538933119 | 6.40265188186281 |
| chr19 | 47741550 | 47741553 | + | 13.0514874993537 | 4.54798296825753 | 8.35259747257593 | 4.26843458790854 |
| chr19 | 47741595 | 47741631 | + | 64.2534769198953 | 27.2878978095452 | 59.6614105183995 | 84.3015831111937 |
| chr19 | 47741971 | 47741999 | + | 2.00792115374673 | 2.27399148412876 | 3.57968463110397 | 6.40265188186281 |
| chr19 | 47742022 | 47742115 | + | 110.43566345607 | 63.6717615556054 | 181.370687975934 | 252.904749333581 |
| chr19 | 47755851 | 47755855 | + | 10.0396057687336 | 6.82197445238629 | 89.4921157775992 | 85.3686917581709 |
| chr19 | 47756998 | 47757039 | + | 34.1346596136944 | 35.2468680039958 | 97.8447132501751 | 54.4225409958339 |
| chr19 | 47781288 | 47781383 | + | 14.0554480762271 | 9.09596593651506 | 29.8307052591997 | 24.5434988804741 |
| chr19 | 47784292 | 47784295 | + | 6.02376346124019 | 1.13699574206438 | 14.3187385244159 | 33.0803680562912 |
| chr19 | 47784317 | 47784335 | + | 4.01584230749346 | 1.13699574206438 | 5.96614105183995 | 7.46976052883995 |
| chr19 | 47784558 | 47784560 | + | 8.03168461498692 | 6.82197445238629 | 5.96614105183995 | 5.33554323488568 |
| chr19 | 47784611 | 47784649 | + | 25.0990144218341 | 11.3699574206438 | 9.54582568294392 | 14.9395210576799 |
| chr19 | 48115479 | 48115535 | - | 29.1148567293276 | 34.1098722619315 | 11.9322821036799 | 32.0132594093141 |
| chr19 | 48115595 | 48115634 | - | 31.1227778830743 | 38.657855230189 | 23.8645642073598 | 27.7448248214055 |
| chr19 | 48115665 | 48115691 | - | 8.03168461498692 | 5.68497871032191 | 13.1255103140479 | 24.5434988804741 |
| chr19 | 48115719 | 48115735 | - | 11.043566345607 | 7.95897019445067 | 14.3187385244159 | 14.9395210576799 |
| chr19 | 48115937 | 48115968 | - | 8.03168461498692 | 4.54798296825753 | 20.2848795762558 | 12.8053037637256 |
| chr19 | 48117686 | 48117696 | - | 5.01980288436682 | 3.41098722619315 | 1.19322821036799 | 3.20132594093141 |
| chr19 | 48117733 | 48117754 | - | 6.02376346124019 | 3.41098722619315 | 9.54582568294392 | 9.60397782279422 |
| chr19 | 48197005 | 48197395 | + | 827.263515343652 | 1481.50545190989 | 157.506123768575 | 217.690163983336 |
| chr19 | 48204058 | 48204135 | + | 55.217831728035 | 65.9457530397342 | 68.0140079909754 | 85.3686917581709 |
| chr19 | 48204160 | 48204163 | + | 2.00792115374673 | 2.27399148412876 | 5.96614105183995 | 3.20132594093141 |
| chr19 | 48255668 | 48255732 | + | 43.1703048055547 | 70.4937360079917 | 87.1056593568632 | 60.8251928776967 |
| chr19 | 48258086 | 48258105 | + | 3.01188173062009 | 4.54798296825753 | 1.19322821036799 | 2.13421729395427 |
| chr19 | 48330518 | 48330524 | + | 5.01980288436682 | 7.95897019445067 | 7.15936926220794 | 9.60397782279422 |
| chr19 | 48332493 | 48332503 | - | 1.00396057687336 | 3.41098722619315 | 1.19322821036799 | 1.06710864697714 |
| chr19 | 48382596 | 48382741 | - | 130.514874993537 | 172.823352793786 | 93.0718004087032 | 85.3686917581709 |
| chr19 | 48382764 | 48382849 | - | 8.03168461498692 | 2.27399148412876 | 5.96614105183995 | 7.46976052883995 |
| chr19 | 48382942 | 48382971 | - | 6.02376346124019 | 11.3699574206438 | 3.57968463110397 | 1.06710864697714 |
| chr19 | 48382994 | 48383165 | - | 90.3564519186028 | 84.1376849127643 | 180.177459765566 | 141.925450047959 |
| chr19 | 48383198 | 48383261 | - | 41.1623836518079 | 29.5618892936739 | 41.7629873628796 | 43.7514545260626 |
| chr19 | 48383288 | 48383319 | - | 7.02772403811355 | 4.54798296825753 | 14.3187385244159 | 6.40265188186281 |
| chr19 | 48479251 | 48479307 | + | 13.0514874993537 | 10.2329616785794 | 14.3187385244159 | 9.60397782279422 |
| chr19 | 48479350 | 48479360 | + | 5.01980288436682 | 5.68497871032191 | 3.57968463110397 | 1.06710864697714 |
| chr19 | 48615352 | 48615421 | - | 182.720824990952 | 212.618203766039 | 208.814936814398 | 255.038966627535 |
| chr19 | 48638063 | 48638065 | - | 1.00396057687336 | 11.3699574206438 | 4.77291284147196 | 4.26843458790854 |
| chr19 | 48720602 | 48720657 | - | 115.455466340437 | 194.426271893009 | 2.38645642073598 | 1.06710864697714 |
| chr19 | 48749647 | 48749653 | - | 1.00396057687336 | 4.54798296825753 | 2.38645642073598 | 17.0737383516342 |
| chr19 | 48795121 | 48795141 | - | 12.0475269224804 | 19.3289276150945 | 4.77291284147196 | 8.53686917581709 |
| chr19 | 48875962 | 48875986 | + | 15.0594086531005 | 39.7948509722534 | 32.2171616799357 | 76.8318225823538 |
| chr19 | 48922454 | 48922544 | + | 20.0792115374673 | 20.4659233571589 | 29.8307052591997 | 18.1408469986113 |
| chr19 | 48922570 | 48922589 | + | 3.01188173062009 | 1.13699574206438 | 1.19322821036799 | 2.13421729395427 |
| chr19 | 48922602 | 48922683 | + | 12.0475269224804 | 6.82197445238629 | 28.6374770488317 | 21.3421729395427 |
| chr19 | 48922706 | 48922708 | + | 1.00396057687336 | 3.41098722619315 | 1.19322821036799 | 1.06710864697714 |
| chr19 | 48922730 | 48922763 | + | 8.03168461498692 | 3.41098722619315 | 2.38645642073598 | 4.26843458790854 |
| chr19 | 48922785 | 48922798 | + | 1.00396057687336 | 4.54798296825753 | 3.57968463110397 | 3.20132594093141 |
| chr19 | 48922836 | 48922889 | + | 21.0831721143407 | 9.09596593651506 | 22.6713359969918 | 16.006629704657 |
| chr19 | 48922917 | 48922932 | + | 3.01188173062009 | 2.27399148412876 | 3.57968463110397 | 6.40265188186281 |
| chr19 | 48922946 | 48923047 | + | 23.0910932680874 | 20.4659233571589 | 34.6036181006717 | 32.0132594093141 |
| chr19 | 48923072 | 48923193 | + | 100.396057687336 | 59.1237785873479 | 65.6275515702394 | 107.777973344691 |
| chr19 | 48923217 | 48923247 | + | 10.0396057687336 | 5.68497871032191 | 3.57968463110397 | 4.26843458790854 |
| chr19 | 48961629 | 48961660 | + | 6.02376346124019 | 4.54798296825753 | 1.19322821036799 | 2.13421729395427 |
| chr19 | 48961699 | 48961780 | + | 147.582204800385 | 177.371335762044 | 36.9900745214077 | 52.2883237018796 |
| chr19 | 48966809 | 48966859 | + | 171.677258645345 | 156.905412404885 | 329.330986061565 | 383.092004264792 |
| chr19 | 48968155 | 48968197 | - | 39.1544624980612 | 35.2468680039958 | 22.6713359969918 | 18.1408469986113 |
| chr19 | 49009970 | 49010067 | + | 46.1821865361748 | 36.3838637460602 | 45.3426719939836 | 90.7042349930565 |
| chr19 | 49015881 | 49015921 | + | 9.03564519186028 | 1.13699574206438 | 10.7390538933119 | 7.46976052883995 |
| chr19 | 49107826 | 49108500 | + | 579.285252855931 | 319.495803520091 | 457.00640457094 | 816.338114937509 |
| chr19 | 49108521 | 49108577 | + | 42.1663442286813 | 31.8358807778027 | 31.0239334695677 | 46.952780466994 |
| chr19 | 49340804 | 49340808 | - | 3.01188173062009 | 3.41098722619315 | 2.38645642073598 | 3.20132594093141 |
| chr19 | 49446322 | 49446425 | - | 267.053513448315 | 260.372024932743 | 451.0402635191 | 512.212150549025 |
| chr19 | 49470511 | 49470552 | + | 9.03564519186028 | 6.82197445238629 | 9.54582568294392 | 12.8053037637256 |
| chr19 | 49478997 | 49479041 | + | 1.00396057687336 | 2.27399148412876 | 31.0239334695677 | 16.006629704657 |
| chr19 | 49487543 | 49487642 | + | 4958.56128917755 | 3971.52612703089 | 4329.03194721507 | 3172.51400746302 |
| chr19 | 49489965 | 49489968 | + | 18.0712903837206 | 9.09596593651506 | 33.4103898903037 | 16.006629704657 |
| chr19 | 49490615 | 49490629 | + | 11.043566345607 | 4.54798296825753 | 11.9322821036799 | 16.006629704657 |
| chr19 | 49491821 | 49491866 | + | 3.01188173062009 | 2.27399148412876 | 4.77291284147196 | 8.53686917581709 |
| chr19 | 49492029 | 49492053 | + | 5.01980288436682 | 4.54798296825753 | 17.8984231555198 | 11.7381951167485 |
| chr19 | 49492074 | 49492105 | + | 51.2019894205416 | 52.3018041349616 | 59.6614105183995 | 49.0869977609482 |
| chr19 | 49492138 | 49492153 | + | 5.01980288436682 | 9.09596593651506 | 4.77291284147196 | 2.13421729395427 |
| chr19 | 49492235 | 49492283 | + | 159.629731722865 | 179.645327246172 | 208.814936814398 | 178.207144045182 |
| chr19 | 49497718 | 49497742 | + | 63.249516343022 | 29.5618892936739 | 139.607700613055 | 87.5029090521251 |
| chr19 | 49499540 | 49499672 | + | 183.724785567826 | 231.947131381134 | 207.62170860403 | 325.468137328026 |
| chr19 | 49555738 | 49555820 | - | 10.0396057687336 | 9.09596593651506 | 9.54582568294392 | 19.2079556455884 |
| chr19 | 49626146 | 49626198 | + | 9.03564519186028 | 6.82197445238629 | 1.19322821036799 | 2.13421729395427 |
| chr19 | 49635415 | 49635424 | - | 7.02772403811355 | 1.13699574206438 | 1.19322821036799 | 1.06710864697714 |
| chr19 | 49635452 | 49635473 | - | 6.02376346124019 | 3.41098722619315 | 9.54582568294392 | 11.7381951167485 |
| chr19 | 49635539 | 49635569 | - | 2.00792115374673 | 3.41098722619315 | 4.77291284147196 | 7.46976052883995 |
| chr19 | 49658539 | 49658612 | + | 8.03168461498692 | 3.41098722619315 | 3.57968463110397 | 4.26843458790854 |
| chr19 | 49659606 | 49659681 | - | 26.1029749987075 | 13.6439489047726 | 5.96614105183995 | 21.3421729395427 |
| chr19 | 49669084 | 49669116 | + | 7.02772403811355 | 4.54798296825753 | 10.7390538933119 | 11.7381951167485 |
| chr19 | 49673699 | 49673730 | + | 32.1267384599477 | 5.68497871032191 | 16.7051949451519 | 46.952780466994 |
| chr19 | 49673836 | 49673868 | + | 32.1267384599477 | 28.4248935516096 | 59.6614105183995 | 147.260993282845 |
| chr19 | 49682193 | 49682226 | + | 4.01584230749346 | 4.54798296825753 | 1.19322821036799 | 2.13421729395427 |
| chr19 | 49688355 | 49688429 | + | 47.1861471130481 | 44.3428339405109 | 33.4103898903037 | 36.2816939972226 |
| chr19 | 49806743 | 49806893 | + | 83.3287278804893 | 40.9318467143178 | 108.583767143487 | 138.724124107028 |
| chr19 | 49807069 | 49807092 | + | 22.087132691214 | 31.8358807778027 | 34.6036181006717 | 58.6909755837425 |
| chr19 | 49836908 | 49836933 | + | 7.02772403811355 | 7.95897019445067 | 3.57968463110397 | 8.53686917581709 |
| chr19 | 49860272 | 49860281 | + | 3.01188173062009 | 1.13699574206438 | 1.19322821036799 | 2.13421729395427 |
| chr19 | 49860632 | 49860708 | + | 60.2376346124019 | 34.1098722619315 | 27.4442488384638 | 49.0869977609482 |
| chr19 | 49861237 | 49861253 | - | 4.01584230749346 | 1.13699574206438 | 2.38645642073598 | 4.26843458790854 |
| chr19 | 49869065 | 49869116 | - | 69.2732798042621 | 39.7948509722534 | 15.5119667347839 | 44.8185631730397 |
| chr19 | 49869143 | 49869158 | - | 118.467348071057 | 46.6168254246397 | 7.15936926220794 | 25.6106075274513 |
| chr19 | 49869365 | 49869405 | - | 21.0831721143407 | 20.4659233571589 | 4.77291284147196 | 2.13421729395427 |
| chr19 | 49869436 | 49869444 | - | 7.02772403811355 | 5.68497871032191 | 1.19322821036799 | 2.13421729395427 |
| chr19 | 49888718 | 49888723 | + | 1.00396057687336 | 6.82197445238629 | 1.19322821036799 | 1.06710864697714 |
| chr19 | 49906855 | 49906954 | - | 18.0712903837206 | 23.876910583352 | 11.9322821036799 | 25.6106075274513 |
| chr19 | 49906988 | 49907026 | - | 14.0554480762271 | 7.95897019445067 | 3.57968463110397 | 8.53686917581709 |
| chr19 | 49908155 | 49908223 | - | 8.03168461498692 | 7.95897019445067 | 7.15936926220794 | 11.7381951167485 |
| chr19 | 49909554 | 49909557 | - | 12.0475269224804 | 5.68497871032191 | 5.96614105183995 | 5.33554323488568 |
| chr19 | 49909635 | 49909637 | - | 2.00792115374673 | 2.27399148412876 | 1.19322821036799 | 2.13421729395427 |
| chr19 | 49909764 | 49909860 | - | 17.0673298068472 | 5.68497871032191 | 11.9322821036799 | 6.40265188186281 |
| chr19 | 49933853 | 49933907 | + | 34.1346596136944 | 34.1098722619315 | 5.96614105183995 | 8.53686917581709 |
| chr19 | 49976516 | 49976544 | - | 1.00396057687336 | 5.68497871032191 | 3.57968463110397 | 6.40265188186281 |
| chr19 | 49981742 | 49981775 | - | 2.00792115374673 | 1.13699574206438 | 5.96614105183995 | 5.33554323488568 |
| chr19 | 50048387 | 50048392 | + | 1.00396057687336 | 3.41098722619315 | 4.77291284147196 | 3.20132594093141 |
| chr19 | 50048672 | 50048675 | + | 3.01188173062009 | 2.27399148412876 | 4.77291284147196 | 3.20132594093141 |
| chr19 | 50382933 | 50382943 | + | 47.1861471130481 | 44.3428339405109 | 14.3187385244159 | 18.1408469986113 |
| chr19 | 50417965 | 50417985 | + | 39.1544624980612 | 55.7127913611547 | 39.3765309421437 | 66.1607361125824 |
| chr19 | 50482226 | 50482283 | + | 151.598047107878 | 77.315710460378 | 13.1255103140479 | 19.2079556455884 |
| chr19 | 50482317 | 50482348 | + | 11.043566345607 | 1.13699574206438 | 2.38645642073598 | 2.13421729395427 |
| chr19 | 50483043 | 50483077 | + | 49.1940682667949 | 36.3838637460602 | 2.38645642073598 | 8.53686917581709 |
| chr19 | 50488380 | 50488381 | + | 15.0594086531005 | 3.41098722619315 | 5.96614105183995 | 2.13421729395427 |
| chr19 | 50488552 | 50488569 | + | 10.0396057687336 | 3.41098722619315 | 4.77291284147196 | 2.13421729395427 |
| chr19 | 50488615 | 50488623 | + | 8.03168461498692 | 4.54798296825753 | 11.9322821036799 | 3.20132594093141 |
| chr19 | 50488755 | 50488780 | + | 13.0514874993537 | 1.13699574206438 | 9.54582568294392 | 5.33554323488568 |
| chr19 | 50488941 | 50488966 | + | 7.02772403811355 | 2.27399148412876 | 2.38645642073598 | 1.06710864697714 |
| chr19 | 50489032 | 50489091 | + | 27.1069355755808 | 14.780944646837 | 25.0577924177278 | 16.006629704657 |
| chr19 | 50489271 | 50489304 | + | 14.0554480762271 | 6.82197445238629 | 8.35259747257593 | 3.20132594093141 |
| chr19 | 50506031 | 50506065 | - | 17.0673298068472 | 18.1919318730301 | 4.77291284147196 | 4.26843458790854 |
| chr19 | 50506165 | 50506228 | - | 19.0752509605939 | 21.6029190992233 | 7.15936926220794 | 4.26843458790854 |
| chr19 | 50797731 | 50797797 | - | 49.1940682667949 | 134.165497563597 | 23.8645642073598 | 43.7514545260626 |
| chr19 | 50797960 | 50797986 | - | 3.01188173062009 | 11.3699574206438 | 3.57968463110397 | 4.26843458790854 |
| chr19 | 51098557 | 51098580 | - | 11.043566345607 | 14.780944646837 | 16.7051949451519 | 12.8053037637256 |
| chr19 | 51345187 | 51345383 | - | 46.1821865361748 | 65.9457530397342 | 38.1833027317757 | 53.3554323488568 |
| chr19 | 51353165 | 51353318 | - | 65.2574374967687 | 46.6168254246397 | 60.8546387287675 | 114.180625226554 |
| chr19 | 51354149 | 51354908 | - | 68.2693192273888 | 51.1648083928972 | 121.709277457535 | 137.657015460051 |
| chr19 | 51394516 | 51394531 | + | 10.0396057687336 | 4.54798296825753 | 15.5119667347839 | 16.006629704657 |
| chr19 | 51693326 | 51693328 | + | 5.01980288436682 | 2.27399148412876 | 2.38645642073598 | 2.13421729395427 |
| chr19 | 52049445 | 52049450 | + | 2.00792115374673 | 3.41098722619315 | 1.19322821036799 | 2.13421729395427 |
| chr19 | 52225986 | 52226248 | + | 171.677258645345 | 87.5486721389574 | 218.360762497342 | 268.911379038238 |
| chr19 | 52226273 | 52226318 | + | 29.1148567293276 | 14.780944646837 | 7.15936926220794 | 21.3421729395427 |
| chr19 | 52292189 | 52292212 | + | 3.01188173062009 | 4.54798296825753 | 14.3187385244159 | 7.46976052883995 |
| chr19 | 52292439 | 52292457 | + | 4.01584230749346 | 1.13699574206438 | 2.38645642073598 | 5.33554323488568 |
| chr19 | 52292484 | 52292496 | + | 12.0475269224804 | 10.2329616785794 | 8.35259747257593 | 19.2079556455884 |
| chr19 | 52292570 | 52292597 | + | 1.00396057687336 | 2.27399148412876 | 5.96614105183995 | 10.6710864697714 |
| chr19 | 52323998 | 52324000 | + | 8.03168461498692 | 2.27399148412876 | 4.77291284147196 | 5.33554323488568 |
| chr19 | 52612470 | 52612474 | - | 1.00396057687336 | 2.27399148412876 | 3.57968463110397 | 4.26843458790854 |
| chr19 | 52612677 | 52612681 | - | 2.00792115374673 | 1.13699574206438 | 1.19322821036799 | 2.13421729395427 |
| chr19 | 52838577 | 52838579 | - | 1.00396057687336 | 1.13699574206438 | 3.57968463110397 | 1.06710864697714 |
| chr19 | 53066757 | 53066781 | - | 4.01584230749346 | 5.68497871032191 | 5.96614105183995 | 5.33554323488568 |
| chr19 | 53411689 | 53411694 | + | 1.00396057687336 | 2.27399148412876 | 2.38645642073598 | 1.06710864697714 |
| chr19 | 53874321 | 53874583 | + | 27.1069355755808 | 42.0688424563821 | 46.5359002043516 | 39.483019938154 |
| chr19 | 53874650 | 53874672 | + | 2.00792115374673 | 2.27399148412876 | 1.19322821036799 | 1.06710864697714 |
| chr19 | 53874698 | 53874751 | + | 7.02772403811355 | 5.68497871032191 | 19.0916513658878 | 43.7514545260626 |
| chr19 | 53875236 | 53875314 | + | 22.087132691214 | 11.3699574206438 | 51.3088130458235 | 49.0869977609482 |
| chr19 | 53875344 | 53875389 | + | 3.01188173062009 | 6.82197445238629 | 3.57968463110397 | 10.6710864697714 |
| chr19 | 53875493 | 53875503 | + | 9.03564519186028 | 10.2329616785794 | 34.6036181006717 | 26.6777161744284 |
| chr19 | 53875719 | 53875747 | + | 10.0396057687336 | 22.7399148412876 | 25.0577924177278 | 11.7381951167485 |
| chr19 | 53875777 | 53875793 | + | 4.01584230749346 | 6.82197445238629 | 2.38645642073598 | 5.33554323488568 |
| chr19 | 53875860 | 53876009 | + | 2.00792115374673 | 6.82197445238629 | 47.7291284147196 | 57.6238669367653 |
| chr19 | 53876046 | 53876197 | + | 73.2891221117556 | 86.411676396893 | 134.834787771583 | 228.361250453107 |
| chr19 | 53876321 | 53876391 | + | 4.01584230749346 | 1.13699574206438 | 2.38645642073598 | 5.33554323488568 |
| chr19 | 54108031 | 54108033 | - | 2.00792115374673 | 4.54798296825753 | 1.19322821036799 | 1.06710864697714 |
| chr19 | 55090947 | 55090989 | - | 24.0950538449607 | 38.657855230189 | 26.2510206280958 | 27.7448248214055 |
| chr19 | 55091379 | 55091400 | - | 1.00396057687336 | 2.27399148412876 | 1.19322821036799 | 6.40265188186281 |
| chr19 | 55132821 | 55132903 | - | 10.0396057687336 | 9.09596593651506 | 53.6952694665595 | 82.1673658172394 |
| chr19 | 55158691 | 55158729 | - | 4.01584230749346 | 5.68497871032191 | 11.9322821036799 | 19.2079556455884 |
| chr19 | 55159264 | 55159324 | + | 7.02772403811355 | 2.27399148412876 | 51.3088130458235 | 37.3488026441998 |
| chr19 | 55229812 | 55229859 | - | 16.0633692299738 | 21.6029190992233 | 10.7390538933119 | 13.8724124107028 |
| chr19 | 55229906 | 55229947 | - | 7.02772403811355 | 3.41098722619315 | 3.57968463110397 | 5.33554323488568 |
| chr19 | 55230003 | 55230075 | - | 6.02376346124019 | 4.54798296825753 | 2.38645642073598 | 5.33554323488568 |
| chr19 | 55230271 | 55230307 | - | 2.00792115374673 | 4.54798296825753 | 2.38645642073598 | 5.33554323488568 |
| chr19 | 55230326 | 55230388 | - | 18.0712903837206 | 14.780944646837 | 34.6036181006717 | 16.006629704657 |
| chr19 | 55230433 | 55230469 | - | 7.02772403811355 | 3.41098722619315 | 9.54582568294392 | 3.20132594093141 |
| chr19 | 55262238 | 55262311 | - | 24.0950538449607 | 25.0139063254164 | 17.8984231555198 | 25.6106075274513 |
| chr19 | 55262364 | 55262369 | - | 11.043566345607 | 13.6439489047726 | 2.38645642073598 | 1.06710864697714 |
| chr19 | 55262399 | 55262460 | - | 18.0712903837206 | 27.2878978095452 | 5.96614105183995 | 13.8724124107028 |
| chr19 | 55262515 | 55262517 | - | 2.00792115374673 | 1.13699574206438 | 1.19322821036799 | 1.06710864697714 |
| chr19 | 55347455 | 55347484 | + | 1.00396057687336 | 7.95897019445067 | 15.5119667347839 | 5.33554323488568 |
| chr19 | 55348056 | 55348079 | + | 14.0554480762271 | 23.876910583352 | 36.9900745214077 | 28.8119334683827 |
| chr19 | 55387985 | 55388015 | + | 2.00792115374673 | 2.27399148412876 | 7.15936926220794 | 25.6106075274513 |
| chr19 | 55388041 | 55388049 | + | 19.0752509605939 | 10.2329616785794 | 14.3187385244159 | 12.8053037637256 |
| chr19 | 55388242 | 55388362 | + | 248.982223064594 | 210.344212281911 | 210.008165024766 | 317.998376799186 |
| chr19 | 55391490 | 55391496 | + | 3.01188173062009 | 3.41098722619315 | 3.57968463110397 | 1.06710864697714 |
| chr19 | 55392001 | 55392027 | + | 6.02376346124019 | 3.41098722619315 | 1.19322821036799 | 2.13421729395427 |
| chr19 | 55404287 | 55404288 | - | 12.0475269224804 | 1.13699574206438 | 2.38645642073598 | 8.53686917581709 |
| chr19 | 55453004 | 55453060 | - | 24.0950538449607 | 25.0139063254164 | 76.3666054635513 | 94.9726695809651 |
| chr19 | 55484436 | 55484457 | + | 6.02376346124019 | 12.5069531627082 | 4.77291284147196 | 5.33554323488568 |
| chr19 | 55487488 | 55487509 | + | 2.00792115374673 | 3.41098722619315 | 1.19322821036799 | 6.40265188186281 |
| chr19 | 55577575 | 55577599 | - | 11.043566345607 | 6.82197445238629 | 3.57968463110397 | 9.60397782279422 |
| chr19 | 55591446 | 55591460 | - | 2.00792115374673 | 2.27399148412876 | 1.19322821036799 | 5.33554323488568 |
| chr19 | 55603102 | 55603111 | + | 4.01584230749346 | 5.68497871032191 | 1.19322821036799 | 6.40265188186281 |
| chr19 | 55617208 | 55617241 | + | 7.02772403811355 | 12.5069531627082 | 5.96614105183995 | 10.6710864697714 |
| chr19 | 55643367 | 55643450 | + | 6.02376346124019 | 7.95897019445067 | 4.77291284147196 | 2.13421729395427 |
| chr19 | 55653058 | 55653080 | + | 2.00792115374673 | 6.82197445238629 | 4.77291284147196 | 13.8724124107028 |
| chr19 | 55653104 | 55653106 | + | 1.00396057687336 | 2.27399148412876 | 1.19322821036799 | 4.26843458790854 |
| chr19 | 55671574 | 55671575 | + | 6.02376346124019 | 1.13699574206438 | 3.57968463110397 | 13.8724124107028 |
| chr19 | 55674224 | 55674292 | + | 101.40001826421 | 106.877599754052 | 41.7629873628796 | 45.8856718200168 |
| chr19 | 55674321 | 55674378 | + | 18.0712903837206 | 12.5069531627082 | 14.3187385244159 | 28.8119334683827 |
| chr19 | 55674440 | 55674461 | + | 1.00396057687336 | 3.41098722619315 | 4.77291284147196 | 14.9395210576799 |
| chr19 | 55674546 | 55674581 | + | 13.0514874993537 | 9.09596593651506 | 5.96614105183995 | 12.8053037637256 |
| chr19 | 55674608 | 55674689 | + | 31.1227778830743 | 52.3018041349616 | 38.1833027317757 | 32.0132594093141 |
| chr19 | 55695683 | 55695746 | + | 7.02772403811355 | 9.09596593651506 | 4.77291284147196 | 4.26843458790854 |
| chr19 | 56087393 | 56087467 | - | 27.1069355755808 | 46.6168254246397 | 29.8307052591997 | 41.6172372321083 |
| chr19 | 56087648 | 56087679 | - | 6.02376346124019 | 3.41098722619315 | 7.15936926220794 | 1.06710864697714 |
| chr19 | 56160249 | 56160270 | + | 4.01584230749346 | 1.13699574206438 | 2.38645642073598 | 1.06710864697714 |
| chr19 | 56160794 | 56160857 | + | 3.01188173062009 | 4.54798296825753 | 4.77291284147196 | 4.26843458790854 |
| chr19 | 56221462 | 56221487 | - | 8.03168461498692 | 2.27399148412876 | 2.38645642073598 | 3.20132594093141 |
| chr19 | 56315273 | 56315295 | + | 9.03564519186028 | 14.780944646837 | 11.9322821036799 | 6.40265188186281 |
| chr19 | 57293098 | 57293119 | + | 6.02376346124019 | 10.2329616785794 | 2.38645642073598 | 7.46976052883995 |
| chr19 | 57293396 | 57293428 | + | 6.02376346124019 | 5.68497871032191 | 16.7051949451519 | 5.33554323488568 |
| chr19 | 57307018 | 57307036 | + | 1.00396057687336 | 1.13699574206438 | 14.3187385244159 | 4.26843458790854 |
| chr19 | 57308421 | 57308427 | + | 1.00396057687336 | 2.27399148412876 | 1.19322821036799 | 1.06710864697714 |
| chr19 | 57329803 | 57329818 | + | 15.0594086531005 | 14.780944646837 | 7.15936926220794 | 11.7381951167485 |
| chr19 | 57359678 | 57359685 | + | 1.00396057687336 | 5.68497871032191 | 2.38645642073598 | 2.13421729395427 |
| chr19 | 57400922 | 57400928 | + | 2.00792115374673 | 4.54798296825753 | 1.19322821036799 | 1.06710864697714 |
| chr19 | 57546456 | 57546457 | - | 1.00396057687336 | 1.13699574206438 | 3.57968463110397 | 1.06710864697714 |
| chr19 | 57622635 | 57622669 | + | 3.01188173062009 | 2.27399148412876 | 2.38645642073598 | 1.06710864697714 |
| chr19 | 57622804 | 57622806 | + | 4.01584230749346 | 15.9179403889013 | 7.15936926220794 | 3.20132594093141 |
| chr19 | 57623114 | 57623131 | + | 4.01584230749346 | 10.2329616785794 | 7.15936926220794 | 2.13421729395427 |
| chr19 | 57807191 | 57807200 | - | 7.02772403811355 | 1.13699574206438 | 5.96614105183995 | 5.33554323488568 |
| chr19 | 57807279 | 57807300 | - | 36.1425807674411 | 15.9179403889013 | 36.9900745214077 | 39.483019938154 |
| chr19 | 57819694 | 57819821 | + | 453.790180746761 | 462.757267020203 | 947.423199032184 | 663.741578419778 |
| chr19 | 57846178 | 57846181 | + | 4.01584230749346 | 3.41098722619315 | 1.19322821036799 | 2.13421729395427 |
| chr19 | 58213148 | 58213161 | + | 2.00792115374673 | 2.27399148412876 | 2.38645642073598 | 4.26843458790854 |
| chr19 | 58213321 | 58213326 | + | 5.01980288436682 | 6.82197445238629 | 8.35259747257593 | 2.13421729395427 |
| chr19 | 58277419 | 58277423 | + | 10.0396057687336 | 7.95897019445067 | 8.35259747257593 | 3.20132594093141 |
| chr19 | 58301767 | 58301781 | + | 8.03168461498692 | 10.2329616785794 | 17.8984231555198 | 21.3421729395427 |
| chr19 | 58301805 | 58301819 | + | 1.00396057687336 | 5.68497871032191 | 4.77291284147196 | 2.13421729395427 |
| chr19 | 58306188 | 58306205 | + | 3.01188173062009 | 3.41098722619315 | 1.19322821036799 | 1.06710864697714 |
| chr19 | 58312351 | 58312357 | + | 1.00396057687336 | 3.41098722619315 | 2.38645642073598 | 1.06710864697714 |
| chr19 | 58314978 | 58315001 | + | 17.0673298068472 | 5.68497871032191 | 7.15936926220794 | 2.13421729395427 |
| chr19 | 58315301 | 58315320 | + | 27.1069355755808 | 38.657855230189 | 8.35259747257593 | 7.46976052883995 |
| chr19 | 58315389 | 58315467 | + | 19.0752509605939 | 63.6717615556054 | 7.15936926220794 | 1.06710864697714 |
| chr19 | 58315538 | 58315540 | + | 3.01188173062009 | 6.82197445238629 | 1.19322821036799 | 1.06710864697714 |
| chr19 | 58315583 | 58315607 | + | 112.443584609817 | 181.919318730301 | 14.3187385244159 | 11.7381951167485 |
| chr19 | 58367670 | 58367671 | - | 1.00396057687336 | 1.13699574206438 | 2.38645642073598 | 2.13421729395427 |
| chr19 | 58394660 | 58394786 | + | 593.340700932158 | 482.086194635298 | 721.903067272634 | 857.955352169617 |
| chr19 | 58417746 | 58417800 | + | 1.00396057687336 | 3.41098722619315 | 14.3187385244159 | 14.9395210576799 |
| chr19 | 58473324 | 58473327 | + | 3.01188173062009 | 13.6439489047726 | 9.54582568294392 | 5.33554323488568 |
| chr19 | 58498358 | 58498395 | - | 12.0475269224804 | 20.4659233571589 | 7.15936926220794 | 27.7448248214055 |
| chr19 | 58550183 | 58550285 | + | 40.1584230749346 | 27.2878978095452 | 70.4004644117114 | 167.53605757541 |
| chr19 | 58550375 | 58550440 | + | 13.0514874993537 | 10.2329616785794 | 21.4781077866238 | 39.483019938154 |
| chr19 | 58550557 | 58550601 | + | 8.03168461498692 | 7.95897019445067 | 4.77291284147196 | 11.7381951167485 |
| chr19 | 58550608 | 58550694 | + | 28.1108961524542 | 14.780944646837 | 20.2848795762558 | 49.0869977609482 |
| chr19 | 58555871 | 58555880 | - | 4.01584230749346 | 3.41098722619315 | 2.38645642073598 | 6.40265188186281 |
| chr19 | 58555903 | 58555921 | - | 5.01980288436682 | 4.54798296825753 | 5.96614105183995 | 18.1408469986113 |
| chr19 | 58555942 | 58555956 | - | 47.1861471130481 | 40.9318467143178 | 35.7968463110397 | 43.7514545260626 |
| chr19 | 58556033 | 58556044 | - | 11.043566345607 | 6.82197445238629 | 4.77291284147196 | 8.53686917581709 |
| chr19 | 58561955 | 58561983 | - | 7.02772403811355 | 9.09596593651506 | 1.19322821036799 | 3.20132594093141 |
| chr19 | 58573486 | 58573504 | + | 7.02772403811355 | 3.41098722619315 | 2.38645642073598 | 6.40265188186281 |
| chr19 | 58573633 | 58573694 | + | 6.02376346124019 | 14.780944646837 | 20.2848795762558 | 27.7448248214055 |
| chr19 | 58573782 | 58573806 | + | 3.01188173062009 | 1.13699574206438 | 5.96614105183995 | 5.33554323488568 |
| chr19 | 58573935 | 58573969 | + | 4.01584230749346 | 10.2329616785794 | 7.15936926220794 | 2.13421729395427 |
| chr2 | 272300 | 272306 | + | 2.00792115374673 | 1.13699574206438 | 1.19322821036799 | 1.06710864697714 |
| chr2 | 272552 | 272565 | + | 7.02772403811355 | 4.54798296825753 | 5.96614105183995 | 3.20132594093141 |
| chr2 | 277320 | 277394 | + | 11.043566345607 | 12.5069531627082 | 7.15936926220794 | 14.9395210576799 |
| chr2 | 277444 | 277482 | + | 5.01980288436682 | 17.0549361309657 | 14.3187385244159 | 23.476390233497 |
| chr2 | 277828 | 277946 | + | 11.043566345607 | 14.780944646837 | 11.9322821036799 | 18.1408469986113 |
| chr2 | 277992 | 278083 | + | 30.1188173062009 | 27.2878978095452 | 13.1255103140479 | 20.2750642925656 |
| chr2 | 278107 | 278136 | + | 35.1386201905678 | 53.438799877026 | 26.2510206280958 | 23.476390233497 |
| chr2 | 668181 | 668187 | - | 31.1227778830743 | 71.6307317500561 | 41.7629873628796 | 28.8119334683827 |
| chr2 | 668327 | 668350 | - | 4.01584230749346 | 2.27399148412876 | 1.19322821036799 | 4.26843458790854 |
| chr2 | 669284 | 669353 | - | 13.0514874993537 | 34.1098722619315 | 4.77291284147196 | 7.46976052883995 |
| chr2 | 3519655 | 3519673 | + | 1.00396057687336 | 1.13699574206438 | 2.38645642073598 | 1.06710864697714 |
| chr2 | 3545613 | 3545653 | - | 4.01584230749346 | 12.5069531627082 | 7.15936926220794 | 5.33554323488568 |
| chr2 | 3558621 | 3558757 | + | 169.669337491599 | 135.302493305661 | 27.4442488384638 | 58.6909755837425 |
| chr2 | 3559455 | 3559465 | + | 3.01188173062009 | 3.41098722619315 | 1.19322821036799 | 5.33554323488568 |
| chr2 | 8729030 | 8729043 | - | 2.00792115374673 | 2.27399148412876 | 8.35259747257593 | 9.60397782279422 |
| chr2 | 8729320 | 8729323 | - | 1.00396057687336 | 2.27399148412876 | 1.19322821036799 | 3.20132594093141 |
| chr2 | 8729362 | 8729393 | - | 1.00396057687336 | 3.41098722619315 | 4.77291284147196 | 4.26843458790854 |
| chr2 | 8852850 | 8852853 | - | 2.00792115374673 | 1.13699574206438 | 3.57968463110397 | 1.06710864697714 |
| chr2 | 8855946 | 8855948 | - | 3.01188173062009 | 3.41098722619315 | 7.15936926220794 | 4.26843458790854 |
| chr2 | 8857136 | 8857140 | - | 5.01980288436682 | 1.13699574206438 | 13.1255103140479 | 10.6710864697714 |
| chr2 | 9405649 | 9405651 | + | 2.00792115374673 | 2.27399148412876 | 1.19322821036799 | 1.06710864697714 |
| chr2 | 9406696 | 9406798 | - | 22.087132691214 | 29.5618892936739 | 4.77291284147196 | 10.6710864697714 |
| chr2 | 9472969 | 9473041 | + | 10.0396057687336 | 13.6439489047726 | 14.3187385244159 | 20.2750642925656 |
| chr2 | 9488643 | 9488676 | - | 5.01980288436682 | 13.6439489047726 | 11.9322821036799 | 6.40265188186281 |
| chr2 | 9488851 | 9488854 | - | 1.00396057687336 | 3.41098722619315 | 1.19322821036799 | 1.06710864697714 |
| chr2 | 9489087 | 9489090 | - | 6.02376346124019 | 2.27399148412876 | 4.77291284147196 | 3.20132594093141 |
| chr2 | 9489760 | 9489769 | - | 1.00396057687336 | 2.27399148412876 | 2.38645642073598 | 14.9395210576799 |
| chr2 | 9489834 | 9489848 | - | 1.00396057687336 | 1.13699574206438 | 3.57968463110397 | 12.8053037637256 |
| chr2 | 9489985 | 9489992 | - | 3.01188173062009 | 1.13699574206438 | 5.96614105183995 | 5.33554323488568 |
| chr2 | 9555963 | 9556054 | + | 29.1148567293276 | 28.4248935516096 | 20.2848795762558 | 25.6106075274513 |
| chr2 | 9556491 | 9556494 | + | 6.02376346124019 | 11.3699574206438 | 2.38645642073598 | 1.06710864697714 |
| chr2 | 9583992 | 9584000 | - | 5.01980288436682 | 9.09596593651506 | 4.77291284147196 | 7.46976052883995 |
| chr2 | 9584104 | 9584113 | - | 5.01980288436682 | 2.27399148412876 | 2.38645642073598 | 3.20132594093141 |
| chr2 | 9584160 | 9584163 | - | 1.00396057687336 | 1.13699574206438 | 3.57968463110397 | 1.06710864697714 |
| chr2 | 9584380 | 9584399 | - | 30.1188173062009 | 47.753821166704 | 45.3426719939836 | 24.5434988804741 |
| chr2 | 9584494 | 9584523 | - | 18.0712903837206 | 38.657855230189 | 35.7968463110397 | 27.7448248214055 |
| chr2 | 9584643 | 9584666 | - | 6.02376346124019 | 7.95897019445067 | 4.77291284147196 | 2.13421729395427 |
| chr2 | 9584851 | 9584869 | - | 93.3683336492229 | 96.6446380754725 | 35.7968463110397 | 46.952780466994 |
| chr2 | 9584912 | 9584951 | - | 8.03168461498692 | 19.3289276150945 | 13.1255103140479 | 9.60397782279422 |
| chr2 | 9585034 | 9585035 | - | 1.00396057687336 | 1.13699574206438 | 1.19322821036799 | 1.06710864697714 |
| chr2 | 9585059 | 9585083 | - | 10.0396057687336 | 21.6029190992233 | 17.8984231555198 | 4.26843458790854 |
| chr2 | 9585112 | 9585206 | - | 29.1148567293276 | 31.8358807778027 | 41.7629873628796 | 29.8790421153598 |
| chr2 | 9585269 | 9585330 | - | 82.3247673036159 | 86.411676396893 | 126.482190299007 | 110.979299285622 |
| chr2 | 9692789 | 9692791 | + | 64.2534769198953 | 63.6717615556054 | 143.187385244159 | 69.3620620535138 |
| chr2 | 10001958 | 10001961 | + | 1.00396057687336 | 5.68497871032191 | 1.19322821036799 | 3.20132594093141 |
| chr2 | 10052604 | 10052644 | + | 27.1069355755808 | 62.534765813541 | 11.9322821036799 | 9.60397782279422 |
| chr2 | 10052675 | 10052706 | + | 5.01980288436682 | 5.68497871032191 | 3.57968463110397 | 13.8724124107028 |
| chr2 | 10054619 | 10054672 | + | 12.0475269224804 | 9.09596593651506 | 3.57968463110397 | 6.40265188186281 |
| chr2 | 10054780 | 10054787 | + | 6.02376346124019 | 15.9179403889013 | 1.19322821036799 | 1.06710864697714 |
| chr2 | 10129388 | 10129413 | + | 10.0396057687336 | 9.09596593651506 | 4.77291284147196 | 1.06710864697714 |
| chr2 | 10129621 | 10129672 | + | 152.602007684751 | 171.686357051722 | 22.6713359969918 | 74.6976052883995 |
| chr2 | 10129710 | 10129734 | + | 4.01584230749346 | 2.27399148412876 | 2.38645642073598 | 5.33554323488568 |
| chr2 | 10130245 | 10130272 | + | 3.01188173062009 | 13.6439489047726 | 1.19322821036799 | 1.06710864697714 |
| chr2 | 10130354 | 10130371 | + | 4.01584230749346 | 6.82197445238629 | 3.57968463110397 | 5.33554323488568 |
| chr2 | 10130388 | 10130403 | + | 7.02772403811355 | 10.2329616785794 | 1.19322821036799 | 1.06710864697714 |
| chr2 | 10130551 | 10130582 | + | 11.043566345607 | 9.09596593651506 | 5.96614105183995 | 11.7381951167485 |
| chr2 | 10131056 | 10131058 | + | 6.02376346124019 | 9.09596593651506 | 1.19322821036799 | 9.60397782279422 |
| chr2 | 10131240 | 10131260 | + | 3.01188173062009 | 4.54798296825753 | 1.19322821036799 | 1.06710864697714 |
| chr2 | 10131337 | 10131388 | + | 8.03168461498692 | 27.2878978095452 | 5.96614105183995 | 8.53686917581709 |
| chr2 | 10198784 | 10198846 | + | 4.01584230749346 | 9.09596593651506 | 36.9900745214077 | 33.0803680562912 |
| chr2 | 10198867 | 10198886 | + | 8.03168461498692 | 6.82197445238629 | 40.5697591525116 | 19.2079556455884 |
| chr2 | 10427225 | 10427254 | + | 7.02772403811355 | 3.41098722619315 | 1.19322821036799 | 3.20132594093141 |
| chr2 | 10427295 | 10427309 | + | 4.01584230749346 | 2.27399148412876 | 2.38645642073598 | 1.06710864697714 |
| chr2 | 10427339 | 10427366 | + | 5.01980288436682 | 4.54798296825753 | 2.38645642073598 | 3.20132594093141 |
| chr2 | 10427398 | 10427445 | + | 13.0514874993537 | 4.54798296825753 | 3.57968463110397 | 4.26843458790854 |
| chr2 | 10427497 | 10427583 | + | 65.2574374967687 | 71.6307317500561 | 41.7629873628796 | 45.8856718200168 |
| chr2 | 10440402 | 10440479 | - | 29.1148567293276 | 39.7948509722534 | 42.9562155732476 | 48.0198891139711 |
| chr2 | 10440502 | 10440862 | - | 3106.25402484619 | 3773.68886791168 | 4045.04363314748 | 5089.04113743396 |
| chr2 | 10448703 | 10448785 | + | 23.0910932680874 | 84.1376849127643 | 44.1494437836156 | 76.8318225823538 |
| chr2 | 10783418 | 10783553 | - | 36.1425807674411 | 88.6856678810218 | 58.4681823080315 | 57.6238669367653 |
| chr2 | 10783581 | 10783684 | - | 15.0594086531005 | 27.2878978095452 | 38.1833027317757 | 21.3421729395427 |
| chr2 | 10783714 | 10783741 | - | 5.01980288436682 | 4.54798296825753 | 11.9322821036799 | 12.8053037637256 |
| chr2 | 10783770 | 10783825 | - | 1.00396057687336 | 1.13699574206438 | 5.96614105183995 | 7.46976052883995 |
| chr2 | 10783849 | 10784361 | - | 1755.92704895151 | 1281.39420130656 | 1584.60706336869 | 1618.80381746431 |
| chr2 | 11179923 | 11179966 | - | 1.00396057687336 | 4.54798296825753 | 2.38645642073598 | 1.06710864697714 |
| chr2 | 11179991 | 11179999 | - | 29.1148567293276 | 34.1098722619315 | 5.96614105183995 | 10.6710864697714 |
| chr2 | 11180375 | 11180425 | - | 7.02772403811355 | 4.54798296825753 | 1.19322821036799 | 1.06710864697714 |
| chr2 | 11180615 | 11180673 | - | 4.01584230749346 | 7.95897019445067 | 8.35259747257593 | 2.13421729395427 |
| chr2 | 11180742 | 11180902 | - | 4.01584230749346 | 10.2329616785794 | 3.57968463110397 | 1.06710864697714 |
| chr2 | 11180924 | 11181016 | - | 10.0396057687336 | 15.9179403889013 | 1.19322821036799 | 3.20132594093141 |
| chr2 | 11181221 | 11181300 | - | 6.02376346124019 | 4.54798296825753 | 3.57968463110397 | 1.06710864697714 |
| chr2 | 11181350 | 11181445 | - | 33.130699036821 | 35.2468680039958 | 1.19322821036799 | 10.6710864697714 |
| chr2 | 11181474 | 11181541 | - | 9.03564519186028 | 9.09596593651506 | 3.57968463110397 | 4.26843458790854 |
| chr2 | 11181572 | 11181632 | - | 3.01188173062009 | 6.82197445238629 | 1.19322821036799 | 1.06710864697714 |
| chr2 | 11182040 | 11182137 | - | 89.3524913417294 | 61.3977700714766 | 31.0239334695677 | 66.1607361125824 |
| chr2 | 11182165 | 11182253 | - | 46.1821865361748 | 61.3977700714766 | 20.2848795762558 | 24.5434988804741 |
| chr2 | 11182343 | 11182387 | - | 2.00792115374673 | 4.54798296825753 | 1.19322821036799 | 3.20132594093141 |
| chr2 | 11182419 | 11182569 | - | 8.03168461498692 | 10.2329616785794 | 11.9322821036799 | 2.13421729395427 |
| chr2 | 11182760 | 11182860 | - | 5.01980288436682 | 6.82197445238629 | 2.38645642073598 | 1.06710864697714 |
| chr2 | 11444565 | 11444587 | - | 10.0396057687336 | 18.1919318730301 | 17.8984231555198 | 23.476390233497 |
| chr2 | 11825714 | 11825718 | + | 3.01188173062009 | 4.54798296825753 | 2.38645642073598 | 10.6710864697714 |
| chr2 | 11825879 | 11825899 | + | 2.00792115374673 | 5.68497871032191 | 2.38645642073598 | 3.20132594093141 |
| chr2 | 15630931 | 15631072 | + | 10.0396057687336 | 18.1919318730301 | 23.8645642073598 | 25.6106075274513 |
| chr2 | 15631074 | 15631080 | + | 3.01188173062009 | 6.82197445238629 | 3.57968463110397 | 5.33554323488568 |
| chr2 | 17664241 | 17664251 | - | 2.00792115374673 | 5.68497871032191 | 3.57968463110397 | 6.40265188186281 |
| chr2 | 17665451 | 17665454 | - | 1.00396057687336 | 2.27399148412876 | 1.19322821036799 | 4.26843458790854 |
| chr2 | 17782741 | 17782761 | + | 3.01188173062009 | 9.09596593651506 | 8.35259747257593 | 3.20132594093141 |
| chr2 | 17784578 | 17784589 | + | 6.02376346124019 | 10.2329616785794 | 9.54582568294392 | 6.40265188186281 |
| chr2 | 18554750 | 18554793 | - | 4.01584230749346 | 9.09596593651506 | 11.9322821036799 | 6.40265188186281 |
| chr2 | 18555055 | 18555086 | - | 3.01188173062009 | 3.41098722619315 | 2.38645642073598 | 4.26843458790854 |
| chr2 | 18555122 | 18555134 | - | 5.01980288436682 | 9.09596593651506 | 2.38645642073598 | 4.26843458790854 |
| chr2 | 18555172 | 18555229 | - | 4.01584230749346 | 12.5069531627082 | 1.19322821036799 | 4.26843458790854 |
| chr2 | 19902035 | 19902050 | + | 12.0475269224804 | 6.82197445238629 | 4.77291284147196 | 4.26843458790854 |
| chr2 | 20032672 | 20033211 | - | 311.227778830743 | 391.126535270147 | 115.743136405695 | 137.657015460051 |
| chr2 | 20202126 | 20202228 | - | 11.043566345607 | 5.68497871032191 | 1.19322821036799 | 3.20132594093141 |
| chr2 | 20202251 | 20202387 | - | 6.02376346124019 | 3.41098722619315 | 2.38645642073598 | 2.13421729395427 |
| chr2 | 20248836 | 20248920 | - | 3.01188173062009 | 7.95897019445067 | 2.38645642073598 | 2.13421729395427 |
| chr2 | 20249112 | 20249119 | - | 10.0396057687336 | 10.2329616785794 | 4.77291284147196 | 8.53686917581709 |
| chr2 | 20249941 | 20249946 | - | 1.00396057687336 | 3.41098722619315 | 2.38645642073598 | 1.06710864697714 |
| chr2 | 20250077 | 20250123 | - | 7.02772403811355 | 17.0549361309657 | 2.38645642073598 | 12.8053037637256 |
| chr2 | 20250346 | 20250359 | - | 7.02772403811355 | 3.41098722619315 | 2.38645642073598 | 6.40265188186281 |
| chr2 | 20250395 | 20250501 | - | 1.00396057687336 | 5.68497871032191 | 1.19322821036799 | 19.2079556455884 |
| chr2 | 20250522 | 20250536 | - | 3.01188173062009 | 1.13699574206438 | 3.57968463110397 | 1.06710864697714 |
| chr2 | 20250563 | 20250577 | - | 1.00396057687336 | 4.54798296825753 | 4.77291284147196 | 4.26843458790854 |
| chr2 | 20250715 | 20250780 | - | 8.03168461498692 | 9.09596593651506 | 21.4781077866238 | 11.7381951167485 |
| chr2 | 20250812 | 20250854 | - | 1.00396057687336 | 1.13699574206438 | 3.57968463110397 | 10.6710864697714 |
| chr2 | 20251403 | 20251412 | - | 4.01584230749346 | 5.68497871032191 | 2.38645642073598 | 6.40265188186281 |
| chr2 | 20448213 | 20448358 | + | 74.293082688629 | 104.603608269923 | 2.38645642073598 | 2.13421729395427 |
| chr2 | 20448776 | 20448935 | + | 72.2851615348822 | 196.700263377138 | 1.19322821036799 | 2.13421729395427 |
| chr2 | 20448992 | 20449169 | + | 87.3445701879827 | 271.741982353387 | 1.19322821036799 | 4.26843458790854 |
| chr2 | 20617845 | 20617903 | - | 5.01980288436682 | 4.54798296825753 | 1.19322821036799 | 2.13421729395427 |
| chr2 | 20701584 | 20701633 | - | 1.00396057687336 | 1.13699574206438 | 2.38645642073598 | 5.33554323488568 |
| chr2 | 23748729 | 23748768 | - | 16.0633692299738 | 19.3289276150945 | 14.3187385244159 | 8.53686917581709 |
| chr2 | 23748798 | 23748861 | - | 13.0514874993537 | 28.4248935516096 | 40.5697591525116 | 10.6710864697714 |
| chr2 | 23748885 | 23748924 | - | 7.02772403811355 | 31.8358807778027 | 38.1833027317757 | 11.7381951167485 |
| chr2 | 23748957 | 23748970 | - | 2.00792115374673 | 5.68497871032191 | 7.15936926220794 | 3.20132594093141 |
| chr2 | 23749009 | 23749061 | - | 3.01188173062009 | 10.2329616785794 | 13.1255103140479 | 12.8053037637256 |
| chr2 | 23749090 | 23749104 | - | 54.2138711511617 | 95.5076423334081 | 54.8884976769275 | 57.6238669367653 |
| chr2 | 23749200 | 23749241 | - | 8.03168461498692 | 7.95897019445067 | 4.77291284147196 | 5.33554323488568 |
| chr2 | 23749271 | 23749283 | - | 1.00396057687336 | 3.41098722619315 | 4.77291284147196 | 1.06710864697714 |
| chr2 | 23749784 | 23749801 | - | 2.00792115374673 | 2.27399148412876 | 7.15936926220794 | 8.53686917581709 |
| chr2 | 24000682 | 24000693 | + | 1.00396057687336 | 5.68497871032191 | 1.19322821036799 | 2.13421729395427 |
| chr2 | 24003549 | 24003552 | + | 1.00396057687336 | 2.27399148412876 | 2.38645642073598 | 3.20132594093141 |
| chr2 | 24029495 | 24029496 | - | 3.01188173062009 | 1.13699574206438 | 1.19322821036799 | 2.13421729395427 |
| chr2 | 24029634 | 24029652 | - | 4.01584230749346 | 6.82197445238629 | 1.19322821036799 | 5.33554323488568 |
| chr2 | 24029687 | 24029691 | - | 1.00396057687336 | 3.41098722619315 | 2.38645642073598 | 4.26843458790854 |
| chr2 | 24030156 | 24030184 | - | 5.01980288436682 | 5.68497871032191 | 3.57968463110397 | 4.26843458790854 |
| chr2 | 24067603 | 24067663 | - | 12.0475269224804 | 34.1098722619315 | 7.15936926220794 | 10.6710864697714 |
| chr2 | 24067718 | 24067741 | - | 15.0594086531005 | 29.5618892936739 | 21.4781077866238 | 36.2816939972226 |
| chr2 | 24790292 | 24790306 | - | 25.0990144218341 | 30.6988850357383 | 4.77291284147196 | 2.13421729395427 |
| chr2 | 24819364 | 24819372 | - | 28.1108961524542 | 27.2878978095452 | 3.57968463110397 | 9.60397782279422 |
| chr2 | 24821270 | 24821323 | + | 1.00396057687336 | 2.27399148412876 | 3.57968463110397 | 13.8724124107028 |
| chr2 | 24822258 | 24822261 | + | 4.01584230749346 | 3.41098722619315 | 2.38645642073598 | 6.40265188186281 |
| chr2 | 24822341 | 24822343 | + | 17.0673298068472 | 34.1098722619315 | 10.7390538933119 | 17.0737383516342 |
| chr2 | 24972121 | 24972188 | + | 8.03168461498692 | 14.780944646837 | 25.0577924177278 | 16.006629704657 |
| chr2 | 25233001 | 25233031 | - | 1.00396057687336 | 1.13699574206438 | 3.57968463110397 | 5.33554323488568 |
| chr2 | 25233841 | 25233871 | - | 1.00396057687336 | 1.13699574206438 | 1.19322821036799 | 17.0737383516342 |
| chr2 | 25234089 | 25234121 | - | 1.00396057687336 | 1.13699574206438 | 4.77291284147196 | 2.13421729395427 |
| chr2 | 25737848 | 25737875 | - | 5.01980288436682 | 9.09596593651506 | 2.38645642073598 | 1.06710864697714 |
| chr2 | 25739698 | 25739740 | - | 7.02772403811355 | 7.95897019445067 | 3.57968463110397 | 3.20132594093141 |
| chr2 | 25739864 | 25739895 | - | 4.01584230749346 | 9.09596593651506 | 1.19322821036799 | 6.40265188186281 |
| chr2 | 25740695 | 25740736 | - | 9.03564519186028 | 13.6439489047726 | 8.35259747257593 | 9.60397782279422 |
| chr2 | 25740931 | 25740951 | - | 5.01980288436682 | 2.27399148412876 | 5.96614105183995 | 5.33554323488568 |
| chr2 | 25741085 | 25741094 | - | 8.03168461498692 | 6.82197445238629 | 15.5119667347839 | 19.2079556455884 |
| chr2 | 25741254 | 25741270 | - | 9.03564519186028 | 7.95897019445067 | 1.19322821036799 | 9.60397782279422 |
| chr2 | 25741984 | 25741998 | - | 58.2297134586551 | 56.8497871032191 | 42.9562155732476 | 36.2816939972226 |
| chr2 | 26134935 | 26135087 | + | 71.2812009580089 | 78.4527062024424 | 41.7629873628796 | 96.0397782279422 |
| chr2 | 26135272 | 26135301 | + | 1.00396057687336 | 2.27399148412876 | 4.77291284147196 | 2.13421729395427 |
| chr2 | 26135478 | 26135546 | + | 6.02376346124019 | 5.68497871032191 | 7.15936926220794 | 9.60397782279422 |
| chr2 | 26136113 | 26136156 | + | 6.02376346124019 | 2.27399148412876 | 1.19322821036799 | 1.06710864697714 |
| chr2 | 26136274 | 26136348 | + | 3.01188173062009 | 5.68497871032191 | 9.54582568294392 | 4.26843458790854 |
| chr2 | 26136818 | 26136821 | + | 5.01980288436682 | 1.13699574206438 | 2.38645642073598 | 2.13421729395427 |
| chr2 | 26136989 | 26137012 | + | 6.02376346124019 | 3.41098722619315 | 1.19322821036799 | 2.13421729395427 |
| chr2 | 26137065 | 26137107 | + | 35.1386201905678 | 38.657855230189 | 29.8307052591997 | 43.7514545260626 |
| chr2 | 26137194 | 26137199 | + | 7.02772403811355 | 18.1919318730301 | 5.96614105183995 | 5.33554323488568 |
| chr2 | 26137307 | 26137308 | + | 1.00396057687336 | 2.27399148412876 | 7.15936926220794 | 2.13421729395427 |
| chr2 | 26190662 | 26190672 | - | 4.01584230749346 | 5.68497871032191 | 1.19322821036799 | 2.13421729395427 |
| chr2 | 26190894 | 26190943 | - | 58.2297134586551 | 51.1648083928972 | 69.2072362013434 | 93.9055609339879 |
| chr2 | 26290178 | 26290194 | + | 5.01980288436682 | 7.95897019445067 | 5.96614105183995 | 12.8053037637256 |
| chr2 | 26290231 | 26290259 | + | 3.01188173062009 | 4.54798296825753 | 8.35259747257593 | 5.33554323488568 |
| chr2 | 26290428 | 26290441 | + | 11.043566345607 | 30.6988850357383 | 21.4781077866238 | 18.1408469986113 |
| chr2 | 26389196 | 26389232 | + | 4.01584230749346 | 3.41098722619315 | 3.57968463110397 | 3.20132594093141 |
| chr2 | 26389380 | 26389419 | + | 11.043566345607 | 7.95897019445067 | 9.54582568294392 | 10.6710864697714 |
| chr2 | 26389731 | 26389931 | + | 30.1188173062009 | 39.7948509722534 | 28.6374770488317 | 22.4092815865198 |
| chr2 | 26389941 | 26389985 | + | 9.03564519186028 | 17.0549361309657 | 14.3187385244159 | 4.26843458790854 |
| chr2 | 26390013 | 26390099 | + | 56.2217923049084 | 76.1787147183136 | 59.6614105183995 | 42.6843458790854 |
| chr2 | 26390129 | 26390218 | + | 18.0712903837206 | 23.876910583352 | 11.9322821036799 | 19.2079556455884 |
| chr2 | 26390229 | 26390361 | + | 251.994104795214 | 410.455462885242 | 178.984231555198 | 165.401840281456 |
| chr2 | 26390364 | 26390453 | + | 56.2217923049084 | 79.5897019445067 | 33.4103898903037 | 18.1408469986113 |
| chr2 | 26390485 | 26390510 | + | 2.00792115374673 | 3.41098722619315 | 1.19322821036799 | 3.20132594093141 |
| chr2 | 26390632 | 26390756 | + | 33.130699036821 | 25.0139063254164 | 31.0239334695677 | 45.8856718200168 |
| chr2 | 26390811 | 26390848 | + | 1.00396057687336 | 2.27399148412876 | 7.15936926220794 | 1.06710864697714 |
| chr2 | 26390866 | 26390902 | + | 12.0475269224804 | 22.7399148412876 | 13.1255103140479 | 3.20132594093141 |
| chr2 | 26391081 | 26391129 | + | 26.1029749987075 | 38.657855230189 | 22.6713359969918 | 32.0132594093141 |
| chr2 | 26391569 | 26391597 | + | 3.01188173062009 | 9.09596593651506 | 1.19322821036799 | 3.20132594093141 |
| chr2 | 26391757 | 26391767 | + | 2.00792115374673 | 2.27399148412876 | 2.38645642073598 | 1.06710864697714 |
| chr2 | 26393217 | 26393258 | + | 6.02376346124019 | 15.9179403889013 | 4.77291284147196 | 3.20132594093141 |
| chr2 | 26393427 | 26393620 | + | 23.0910932680874 | 38.657855230189 | 28.6374770488317 | 19.2079556455884 |
| chr2 | 26393687 | 26393753 | + | 5.01980288436682 | 5.68497871032191 | 8.35259747257593 | 7.46976052883995 |
| chr2 | 26393805 | 26393860 | + | 8.03168461498692 | 11.3699574206438 | 8.35259747257593 | 5.33554323488568 |
| chr2 | 26394524 | 26394569 | + | 6.02376346124019 | 3.41098722619315 | 9.54582568294392 | 21.3421729395427 |
| chr2 | 26394585 | 26394627 | + | 1.00396057687336 | 3.41098722619315 | 3.57968463110397 | 5.33554323488568 |
| chr2 | 26394772 | 26394781 | + | 2.00792115374673 | 1.13699574206438 | 1.19322821036799 | 2.13421729395427 |
| chr2 | 26394825 | 26394851 | + | 1.00396057687336 | 1.13699574206438 | 8.35259747257593 | 6.40265188186281 |
| chr2 | 26394981 | 26395018 | + | 7.02772403811355 | 6.82197445238629 | 9.54582568294392 | 6.40265188186281 |
| chr2 | 26395229 | 26395257 | + | 3.01188173062009 | 2.27399148412876 | 5.96614105183995 | 8.53686917581709 |
| chr2 | 26395428 | 26395642 | + | 478.889195168595 | 754.96517273075 | 483.257425199036 | 668.010013007687 |
| chr2 | 26395673 | 26395743 | + | 68.2693192273888 | 77.315710460378 | 97.8447132501751 | 72.5633879944452 |
| chr2 | 26395773 | 26395799 | + | 1.00396057687336 | 4.54798296825753 | 2.38645642073598 | 1.06710864697714 |
| chr2 | 26778581 | 26778585 | + | 11.043566345607 | 7.95897019445067 | 3.57968463110397 | 2.13421729395427 |
| chr2 | 26779929 | 26779930 | + | 1.00396057687336 | 13.6439489047726 | 1.19322821036799 | 1.06710864697714 |
| chr2 | 26780950 | 26780962 | + | 2.00792115374673 | 5.68497871032191 | 1.19322821036799 | 1.06710864697714 |
| chr2 | 26781049 | 26781076 | + | 7.02772403811355 | 7.95897019445067 | 1.19322821036799 | 1.06710864697714 |
| chr2 | 26793291 | 26793324 | + | 1.00396057687336 | 2.27399148412876 | 3.57968463110397 | 11.7381951167485 |
| chr2 | 26793791 | 26793905 | + | 77.3049644192491 | 44.3428339405109 | 65.6275515702394 | 133.388580872142 |
| chr2 | 26793936 | 26793939 | + | 3.01188173062009 | 1.13699574206438 | 2.38645642073598 | 1.06710864697714 |
| chr2 | 26794203 | 26794245 | + | 9.03564519186028 | 7.95897019445067 | 1.19322821036799 | 6.40265188186281 |
| chr2 | 26794296 | 26794397 | + | 40.1584230749346 | 23.876910583352 | 23.8645642073598 | 27.7448248214055 |
| chr2 | 26794445 | 26794449 | + | 2.00792115374673 | 5.68497871032191 | 3.57968463110397 | 1.06710864697714 |
| chr2 | 26794481 | 26794501 | + | 7.02772403811355 | 11.3699574206438 | 4.77291284147196 | 2.13421729395427 |
| chr2 | 26794523 | 26794608 | + | 24.0950538449607 | 43.2058381984465 | 21.4781077866238 | 19.2079556455884 |
| chr2 | 27026808 | 27026814 | + | 2.00792115374673 | 3.41098722619315 | 31.0239334695677 | 11.7381951167485 |
| chr2 | 27040998 | 27041018 | + | 1.00396057687336 | 3.41098722619315 | 2.38645642073598 | 3.20132594093141 |
| chr2 | 27041511 | 27041528 | + | 1.00396057687336 | 2.27399148412876 | 4.77291284147196 | 4.26843458790854 |
| chr2 | 27041579 | 27041625 | + | 7.02772403811355 | 2.27399148412876 | 11.9322821036799 | 7.46976052883995 |
| chr2 | 27041627 | 27041677 | + | 27.1069355755808 | 22.7399148412876 | 19.0916513658878 | 5.33554323488568 |
| chr2 | 27059398 | 27059508 | + | 38.1505019211879 | 47.753821166704 | 27.4442488384638 | 21.3421729395427 |
| chr2 | 27070508 | 27070567 | - | 133.526756724157 | 89.8226636230862 | 83.5259747257593 | 141.925450047959 |
| chr2 | 27070677 | 27070721 | - | 45.1782259593014 | 40.9318467143178 | 22.6713359969918 | 46.952780466994 |
| chr2 | 27099903 | 27099948 | + | 4.01584230749346 | 3.41098722619315 | 5.96614105183995 | 4.26843458790854 |
| chr2 | 27130774 | 27130955 | - | 18.0712903837206 | 45.4798296825753 | 15.5119667347839 | 33.0803680562912 |
| chr2 | 27131094 | 27131131 | - | 1.00396057687336 | 2.27399148412876 | 8.35259747257593 | 8.53686917581709 |
| chr2 | 27131161 | 27131362 | - | 17.0673298068472 | 11.3699574206438 | 26.2510206280958 | 33.0803680562912 |
| chr2 | 27199608 | 27199615 | - | 22.087132691214 | 35.2468680039958 | 7.15936926220794 | 6.40265188186281 |
| chr2 | 27217005 | 27217011 | + | 3.01188173062009 | 10.2329616785794 | 3.57968463110397 | 7.46976052883995 |
| chr2 | 27217055 | 27217134 | + | 123.487150955424 | 151.220433694563 | 33.4103898903037 | 19.2079556455884 |
| chr2 | 27243487 | 27243520 | + | 10.0396057687336 | 4.54798296825753 | 1.19322821036799 | 16.006629704657 |
| chr2 | 27243677 | 27243711 | + | 18.0712903837206 | 15.9179403889013 | 5.96614105183995 | 22.4092815865198 |
| chr2 | 27309514 | 27309546 | - | 9.03564519186028 | 14.780944646837 | 16.7051949451519 | 18.1408469986113 |
| chr2 | 27309787 | 27309858 | - | 9.03564519186028 | 5.68497871032191 | 2.38645642073598 | 3.20132594093141 |
| chr2 | 27325877 | 27325909 | - | 3.01188173062009 | 6.82197445238629 | 3.57968463110397 | 4.26843458790854 |
| chr2 | 27326108 | 27326109 | - | 5.01980288436682 | 1.13699574206438 | 5.96614105183995 | 3.20132594093141 |
| chr2 | 27326174 | 27326402 | - | 30.1188173062009 | 15.9179403889013 | 5.96614105183995 | 8.53686917581709 |
| chr2 | 27356272 | 27356300 | + | 14.0554480762271 | 15.9179403889013 | 5.96614105183995 | 14.9395210576799 |
| chr2 | 27356801 | 27356802 | + | 4.01584230749346 | 2.27399148412876 | 1.19322821036799 | 1.06710864697714 |
| chr2 | 27356828 | 27356880 | + | 20.0792115374673 | 14.780944646837 | 15.5119667347839 | 19.2079556455884 |
| chr2 | 27357023 | 27357198 | + | 131.518835570411 | 146.672450726305 | 127.675418509375 | 138.724124107028 |
| chr2 | 27358372 | 27358387 | + | 4.01584230749346 | 1.13699574206438 | 3.57968463110397 | 1.06710864697714 |
| chr2 | 27364383 | 27364410 | - | 5.01980288436682 | 5.68497871032191 | 2.38645642073598 | 3.20132594093141 |
| chr2 | 27366921 | 27366938 | - | 1.00396057687336 | 2.27399148412876 | 2.38645642073598 | 4.26843458790854 |
| chr2 | 27367185 | 27367220 | - | 4.01584230749346 | 2.27399148412876 | 4.77291284147196 | 5.33554323488568 |
| chr2 | 27367454 | 27367487 | - | 2.00792115374673 | 7.95897019445067 | 4.77291284147196 | 9.60397782279422 |
| chr2 | 27376662 | 27376764 | + | 11.043566345607 | 11.3699574206438 | 11.9322821036799 | 21.3421729395427 |
| chr2 | 27376782 | 27376821 | + | 31.1227778830743 | 28.4248935516096 | 20.2848795762558 | 27.7448248214055 |
| chr2 | 27376848 | 27376856 | + | 6.02376346124019 | 5.68497871032191 | 5.96614105183995 | 8.53686917581709 |
| chr2 | 27376884 | 27376922 | + | 5.01980288436682 | 7.95897019445067 | 9.54582568294392 | 6.40265188186281 |
| chr2 | 27376936 | 27377099 | + | 429.6951269018 | 302.440867389126 | 269.669575543166 | 271.045596332192 |
| chr2 | 27381244 | 27381284 | - | 11.043566345607 | 14.780944646837 | 11.9322821036799 | 9.60397782279422 |
| chr2 | 27381313 | 27381380 | - | 19.0752509605939 | 21.6029190992233 | 7.15936926220794 | 9.60397782279422 |
| chr2 | 27381387 | 27381392 | - | 1.00396057687336 | 1.13699574206438 | 1.19322821036799 | 3.20132594093141 |
| chr2 | 27381452 | 27381522 | - | 39.1544624980612 | 28.4248935516096 | 38.1833027317757 | 97.1068868749194 |
| chr2 | 27381544 | 27381711 | - | 107.42378172545 | 63.6717615556054 | 151.539982716735 | 247.569206098695 |
| chr2 | 27441823 | 27441960 | + | 39.1544624980612 | 17.0549361309657 | 51.3088130458235 | 33.0803680562912 |
| chr2 | 27442058 | 27442234 | + | 336.326793252577 | 200.111250603331 | 110.970223564223 | 212.35462074845 |
| chr2 | 27444404 | 27444430 | - | 2.00792115374673 | 1.13699574206438 | 2.38645642073598 | 3.20132594093141 |
| chr2 | 27582970 | 27583131 | + | 1302.13686820475 | 1217.72243975095 | 2454.47042872695 | 2102.20403454496 |
| chr2 | 27599582 | 27599630 | + | 16.0633692299738 | 27.2878978095452 | 72.7869208324474 | 46.952780466994 |
| chr2 | 27599969 | 27599997 | + | 3.01188173062009 | 2.27399148412876 | 8.35259747257593 | 7.46976052883995 |
| chr2 | 27622995 | 27623068 | + | 9.03564519186028 | 26.1509020674808 | 22.6713359969918 | 36.2816939972226 |
| chr2 | 27631852 | 27631857 | + | 10.0396057687336 | 3.41098722619315 | 3.57968463110397 | 4.26843458790854 |
| chr2 | 27634867 | 27634916 | + | 12.0475269224804 | 3.41098722619315 | 3.57968463110397 | 5.33554323488568 |
| chr2 | 27650245 | 27650257 | + | 1.00396057687336 | 4.54798296825753 | 4.77291284147196 | 5.33554323488568 |
| chr2 | 27650284 | 27650299 | + | 14.0554480762271 | 14.780944646837 | 3.57968463110397 | 8.53686917581709 |
| chr2 | 27650363 | 27650397 | + | 42.1663442286813 | 61.3977700714766 | 22.6713359969918 | 24.5434988804741 |
| chr2 | 27650433 | 27650451 | + | 12.0475269224804 | 6.82197445238629 | 15.5119667347839 | 19.2079556455884 |
| chr2 | 27650651 | 27650658 | + | 5.01980288436682 | 1.13699574206438 | 2.38645642073598 | 5.33554323488568 |
| chr2 | 27650682 | 27650690 | + | 3.01188173062009 | 1.13699574206438 | 2.38645642073598 | 3.20132594093141 |
| chr2 | 27650755 | 27650779 | + | 5.01980288436682 | 2.27399148412876 | 2.38645642073598 | 6.40265188186281 |
| chr2 | 27650815 | 27650826 | + | 8.03168461498692 | 11.3699574206438 | 3.57968463110397 | 1.06710864697714 |
| chr2 | 27652901 | 27652906 | - | 4.01584230749346 | 4.54798296825753 | 3.57968463110397 | 3.20132594093141 |
| chr2 | 27653085 | 27653134 | - | 2.00792115374673 | 10.2329616785794 | 9.54582568294392 | 2.13421729395427 |
| chr2 | 27653329 | 27653353 | - | 8.03168461498692 | 11.3699574206438 | 5.96614105183995 | 5.33554323488568 |
| chr2 | 27653415 | 27653501 | - | 15.0594086531005 | 20.4659233571589 | 17.8984231555198 | 19.2079556455884 |
| chr2 | 27779484 | 27779711 | + | 73.2891221117556 | 76.1787147183136 | 26.2510206280958 | 51.2212150549025 |
| chr2 | 28338575 | 28338617 | + | 5.01980288436682 | 10.2329616785794 | 7.15936926220794 | 4.26843458790854 |
| chr2 | 28338796 | 28338824 | + | 5.01980288436682 | 2.27399148412876 | 1.19322821036799 | 3.20132594093141 |
| chr2 | 28799279 | 28799343 | + | 9.03564519186028 | 14.780944646837 | 7.15936926220794 | 8.53686917581709 |
| chr2 | 28799365 | 28799484 | + | 99.3920971104631 | 126.206527369146 | 27.4442488384638 | 22.4092815865198 |
| chr2 | 28799578 | 28799735 | + | 55.217831728035 | 57.9867828452835 | 29.8307052591997 | 21.3421729395427 |
| chr2 | 28799769 | 28799786 | + | 9.03564519186028 | 2.27399148412876 | 5.96614105183995 | 7.46976052883995 |
| chr2 | 28799953 | 28800013 | + | 4.01584230749346 | 14.780944646837 | 4.77291284147196 | 6.40265188186281 |
| chr2 | 28800079 | 28800131 | + | 10.0396057687336 | 7.95897019445067 | 4.77291284147196 | 5.33554323488568 |
| chr2 | 28800167 | 28800200 | + | 3.01188173062009 | 4.54798296825753 | 4.77291284147196 | 23.476390233497 |
| chr2 | 28800342 | 28800587 | + | 120.475269224804 | 108.014595496116 | 34.6036181006717 | 57.6238669367653 |
| chr2 | 28800600 | 28800673 | + | 12.0475269224804 | 22.7399148412876 | 7.15936926220794 | 16.006629704657 |
| chr2 | 28800707 | 28800769 | + | 12.0475269224804 | 12.5069531627082 | 3.57968463110397 | 4.26843458790854 |
| chr2 | 28800885 | 28800954 | + | 11.043566345607 | 36.3838637460602 | 13.1255103140479 | 33.0803680562912 |
| chr2 | 28801324 | 28801326 | + | 10.0396057687336 | 12.5069531627082 | 2.38645642073598 | 6.40265188186281 |
| chr2 | 28801386 | 28801420 | + | 15.0594086531005 | 13.6439489047726 | 9.54582568294392 | 24.5434988804741 |
| chr2 | 28801506 | 28801514 | + | 1.00396057687336 | 1.13699574206438 | 1.19322821036799 | 2.13421729395427 |
| chr2 | 28801608 | 28801622 | + | 10.0396057687336 | 5.68497871032191 | 2.38645642073598 | 1.06710864697714 |
| chr2 | 28802902 | 28802913 | + | 4.01584230749346 | 1.13699574206438 | 1.19322821036799 | 3.20132594093141 |
| chr2 | 28849947 | 28849949 | - | 2.00792115374673 | 5.68497871032191 | 4.77291284147196 | 5.33554323488568 |
| chr2 | 28946833 | 28946842 | + | 3.01188173062009 | 1.13699574206438 | 2.38645642073598 | 3.20132594093141 |
| chr2 | 28946916 | 28946956 | + | 2.00792115374673 | 2.27399148412876 | 3.57968463110397 | 3.20132594093141 |
| chr2 | 28947114 | 28947134 | + | 5.01980288436682 | 4.54798296825753 | 1.19322821036799 | 1.06710864697714 |
| chr2 | 28947440 | 28947475 | + | 9.03564519186028 | 5.68497871032191 | 2.38645642073598 | 1.06710864697714 |
| chr2 | 28947560 | 28947584 | + | 7.02772403811355 | 2.27399148412876 | 2.38645642073598 | 4.26843458790854 |
| chr2 | 28947612 | 28947622 | + | 10.0396057687336 | 3.41098722619315 | 1.19322821036799 | 2.13421729395427 |
| chr2 | 28947749 | 28947762 | + | 11.043566345607 | 7.95897019445067 | 3.57968463110397 | 12.8053037637256 |
| chr2 | 28947898 | 28948098 | + | 239.946577872734 | 249.0020675121 | 175.404546924094 | 177.140035398205 |
| chr2 | 28948119 | 28948170 | + | 9.03564519186028 | 7.95897019445067 | 5.96614105183995 | 6.40265188186281 |
| chr2 | 30158748 | 30158874 | + | 6.02376346124019 | 19.3289276150945 | 21.4781077866238 | 11.7381951167485 |
| chr2 | 30159176 | 30159225 | + | 6.02376346124019 | 20.4659233571589 | 19.0916513658878 | 6.40265188186281 |
| chr2 | 30159336 | 30159372 | + | 6.02376346124019 | 19.3289276150945 | 19.0916513658878 | 22.4092815865198 |
| chr2 | 30159401 | 30159416 | + | 2.00792115374673 | 2.27399148412876 | 5.96614105183995 | 5.33554323488568 |
| chr2 | 30159444 | 30159519 | + | 7.02772403811355 | 13.6439489047726 | 13.1255103140479 | 16.006629704657 |
| chr2 | 30640763 | 30640769 | + | 2.00792115374673 | 13.6439489047726 | 2.38645642073598 | 2.13421729395427 |
| chr2 | 30640842 | 30640844 | + | 5.01980288436682 | 9.09596593651506 | 1.19322821036799 | 4.26843458790854 |
| chr2 | 30641266 | 30641268 | + | 8.03168461498692 | 6.82197445238629 | 2.38645642073598 | 2.13421729395427 |
| chr2 | 30641858 | 30641880 | + | 9.03564519186028 | 11.3699574206438 | 2.38645642073598 | 7.46976052883995 |
| chr2 | 30642179 | 30642230 | + | 10.0396057687336 | 11.3699574206438 | 3.57968463110397 | 4.26843458790854 |
| chr2 | 30642624 | 30642654 | + | 3.01188173062009 | 3.41098722619315 | 1.19322821036799 | 1.06710864697714 |
| chr2 | 30643852 | 30643854 | + | 1.00396057687336 | 2.27399148412876 | 1.19322821036799 | 2.13421729395427 |
| chr2 | 30643905 | 30643934 | + | 7.02772403811355 | 15.9179403889013 | 1.19322821036799 | 13.8724124107028 |
| chr2 | 30644135 | 30644172 | + | 3.01188173062009 | 3.41098722619315 | 2.38645642073598 | 2.13421729395427 |
| chr2 | 31867528 | 31867535 | - | 2.00792115374673 | 4.54798296825753 | 2.38645642073598 | 3.20132594093141 |
| chr2 | 31867784 | 31867813 | - | 3.01188173062009 | 3.41098722619315 | 4.77291284147196 | 1.06710864697714 |
| chr2 | 31867838 | 31867840 | - | 6.02376346124019 | 2.27399148412876 | 5.96614105183995 | 4.26843458790854 |
| chr2 | 31868106 | 31868113 | - | 4.01584230749346 | 6.82197445238629 | 4.77291284147196 | 5.33554323488568 |
| chr2 | 32023959 | 32024003 | - | 14.0554480762271 | 11.3699574206438 | 4.77291284147196 | 6.40265188186281 |
| chr2 | 32024042 | 32024075 | - | 6.02376346124019 | 14.780944646837 | 10.7390538933119 | 17.0737383516342 |
| chr2 | 32024110 | 32024130 | - | 6.02376346124019 | 3.41098722619315 | 3.57968463110397 | 4.26843458790854 |
| chr2 | 32024172 | 32024196 | - | 10.0396057687336 | 3.41098722619315 | 5.96614105183995 | 12.8053037637256 |
| chr2 | 32154562 | 32154597 | + | 5.01980288436682 | 1.13699574206438 | 3.57968463110397 | 4.26843458790854 |
| chr2 | 32154835 | 32154848 | + | 3.01188173062009 | 3.41098722619315 | 3.57968463110397 | 7.46976052883995 |
| chr2 | 32154944 | 32154959 | + | 6.02376346124019 | 1.13699574206438 | 2.38645642073598 | 4.26843458790854 |
| chr2 | 32155043 | 32155065 | + | 4.01584230749346 | 6.82197445238629 | 4.77291284147196 | 2.13421729395427 |
| chr2 | 32155181 | 32155254 | + | 14.0554480762271 | 15.9179403889013 | 31.0239334695677 | 22.4092815865198 |
| chr2 | 32222063 | 32222072 | + | 3.01188173062009 | 4.54798296825753 | 8.35259747257593 | 2.13421729395427 |
| chr2 | 32305672 | 32305717 | + | 1.00396057687336 | 3.41098722619315 | 1.19322821036799 | 1.06710864697714 |
| chr2 | 32305739 | 32305754 | + | 6.02376346124019 | 13.6439489047726 | 4.77291284147196 | 4.26843458790854 |
| chr2 | 32305917 | 32305961 | + | 5.01980288436682 | 13.6439489047726 | 3.57968463110397 | 2.13421729395427 |
| chr2 | 32306371 | 32306403 | + | 4.01584230749346 | 7.95897019445067 | 1.19322821036799 | 4.26843458790854 |
| chr2 | 32306493 | 32306551 | + | 12.0475269224804 | 22.7399148412876 | 19.0916513658878 | 33.0803680562912 |
| chr2 | 32307306 | 32307322 | + | 4.01584230749346 | 7.95897019445067 | 1.19322821036799 | 6.40265188186281 |
| chr2 | 32618007 | 32618076 | + | 3.01188173062009 | 4.54798296825753 | 3.57968463110397 | 25.6106075274513 |
| chr2 | 32618167 | 32618207 | + | 1.00396057687336 | 3.41098722619315 | 4.77291284147196 | 29.8790421153598 |
| chr2 | 32618250 | 32618254 | + | 3.01188173062009 | 3.41098722619315 | 1.19322821036799 | 2.13421729395427 |
| chr2 | 32820945 | 32820981 | + | 4.01584230749346 | 1.13699574206438 | 7.15936926220794 | 13.8724124107028 |
| chr2 | 33583686 | 33583713 | - | 20.0792115374673 | 10.2329616785794 | 29.8307052591997 | 35.2145853502455 |
| chr2 | 33584850 | 33584892 | - | 2.00792115374673 | 2.27399148412876 | 9.54582568294392 | 10.6710864697714 |
| chr2 | 33585060 | 33585120 | - | 7.02772403811355 | 3.41098722619315 | 20.2848795762558 | 33.0803680562912 |
| chr2 | 33585131 | 33585288 | - | 52.205949997415 | 17.0549361309657 | 82.3327465153913 | 74.6976052883995 |
| chr2 | 33585303 | 33585327 | - | 40.1584230749346 | 22.7399148412876 | 45.3426719939836 | 34.1474767032683 |
| chr2 | 33585359 | 33585360 | - | 1.00396057687336 | 2.27399148412876 | 13.1255103140479 | 1.06710864697714 |
| chr2 | 33585414 | 33585436 | - | 5.01980288436682 | 2.27399148412876 | 10.7390538933119 | 5.33554323488568 |
| chr2 | 36552285 | 36552334 | - | 2.00792115374673 | 5.68497871032191 | 3.57968463110397 | 5.33554323488568 |
| chr2 | 36552565 | 36552626 | - | 13.0514874993537 | 15.9179403889013 | 15.5119667347839 | 22.4092815865198 |
| chr2 | 36552765 | 36552817 | - | 2.00792115374673 | 4.54798296825753 | 1.19322821036799 | 2.13421729395427 |
| chr2 | 36552979 | 36553003 | - | 2.00792115374673 | 1.13699574206438 | 2.38645642073598 | 1.06710864697714 |
| chr2 | 36844137 | 36844141 | - | 5.01980288436682 | 2.27399148412876 | 4.77291284147196 | 9.60397782279422 |
| chr2 | 36844187 | 36844200 | - | 12.0475269224804 | 2.27399148412876 | 2.38645642073598 | 10.6710864697714 |
| chr2 | 36844217 | 36844243 | - | 6.02376346124019 | 2.27399148412876 | 2.38645642073598 | 1.06710864697714 |
| chr2 | 36844406 | 36844412 | - | 3.01188173062009 | 5.68497871032191 | 15.5119667347839 | 4.26843458790854 |
| chr2 | 36844598 | 36844600 | - | 7.02772403811355 | 5.68497871032191 | 20.2848795762558 | 18.1408469986113 |
| chr2 | 36845283 | 36845294 | - | 1.00396057687336 | 3.41098722619315 | 3.57968463110397 | 4.26843458790854 |
| chr2 | 36845676 | 36845678 | - | 1.00396057687336 | 2.27399148412876 | 2.38645642073598 | 2.13421729395427 |
| chr2 | 36845966 | 36845980 | - | 18.0712903837206 | 22.7399148412876 | 42.9562155732476 | 54.4225409958339 |
| chr2 | 36846622 | 36846627 | - | 6.02376346124019 | 6.82197445238629 | 1.19322821036799 | 4.26843458790854 |
| chr2 | 36981146 | 36981158 | - | 1.00396057687336 | 2.27399148412876 | 2.38645642073598 | 3.20132594093141 |
| chr2 | 37096941 | 37096944 | + | 3.01188173062009 | 3.41098722619315 | 1.19322821036799 | 3.20132594093141 |
| chr2 | 37097456 | 37097472 | + | 1.00396057687336 | 2.27399148412876 | 5.96614105183995 | 1.06710864697714 |
| chr2 | 37098808 | 37098821 | + | 4.01584230749346 | 2.27399148412876 | 2.38645642073598 | 8.53686917581709 |
| chr2 | 37098855 | 37098886 | + | 4.01584230749346 | 4.54798296825753 | 1.19322821036799 | 8.53686917581709 |
| chr2 | 37099391 | 37099428 | - | 252.998065372088 | 151.220433694563 | 131.255103140479 | 113.113516579576 |
| chr2 | 37099635 | 37099663 | - | 47.1861471130481 | 38.657855230189 | 16.7051949451519 | 23.476390233497 |
| chr2 | 37099686 | 37099690 | - | 18.0712903837206 | 15.9179403889013 | 1.19322821036799 | 6.40265188186281 |
| chr2 | 37100164 | 37100198 | - | 5.01980288436682 | 2.27399148412876 | 4.77291284147196 | 6.40265188186281 |
| chr2 | 37100274 | 37100299 | - | 15.0594086531005 | 7.95897019445067 | 1.19322821036799 | 9.60397782279422 |
| chr2 | 37100437 | 37100463 | - | 58.2297134586551 | 47.753821166704 | 29.8307052591997 | 5.33554323488568 |
| chr2 | 37104948 | 37104950 | - | 19.0752509605939 | 4.54798296825753 | 2.38645642073598 | 2.13421729395427 |
| chr2 | 37105673 | 37105675 | - | 31.1227778830743 | 10.2329616785794 | 2.38645642073598 | 6.40265188186281 |
| chr2 | 37106084 | 37106124 | - | 21.0831721143407 | 11.3699574206438 | 16.7051949451519 | 8.53686917581709 |
| chr2 | 37106422 | 37106425 | - | 7.02772403811355 | 7.95897019445067 | 2.38645642073598 | 1.06710864697714 |
| chr2 | 37106769 | 37106784 | - | 19.0752509605939 | 14.780944646837 | 3.57968463110397 | 6.40265188186281 |
| chr2 | 37107069 | 37107073 | - | 9.03564519186028 | 3.41098722619315 | 3.57968463110397 | 3.20132594093141 |
| chr2 | 37201658 | 37201727 | + | 18.0712903837206 | 6.82197445238629 | 23.8645642073598 | 17.0737383516342 |
| chr2 | 37202334 | 37202335 | + | 1.00396057687336 | 1.13699574206438 | 1.19322821036799 | 1.06710864697714 |
| chr2 | 37202541 | 37202553 | + | 5.01980288436682 | 4.54798296825753 | 21.4781077866238 | 3.20132594093141 |
| chr2 | 37202841 | 37202861 | + | 1.00396057687336 | 1.13699574206438 | 2.38645642073598 | 3.20132594093141 |
| chr2 | 37202956 | 37202975 | + | 10.0396057687336 | 11.3699574206438 | 20.2848795762558 | 7.46976052883995 |
| chr2 | 37203371 | 37203376 | + | 10.0396057687336 | 6.82197445238629 | 7.15936926220794 | 12.8053037637256 |
| chr2 | 37203403 | 37203405 | + | 2.00792115374673 | 2.27399148412876 | 2.38645642073598 | 1.06710864697714 |
| chr2 | 37203550 | 37203562 | + | 1.00396057687336 | 1.13699574206438 | 1.19322821036799 | 13.8724124107028 |
| chr2 | 37203708 | 37203723 | + | 5.01980288436682 | 4.54798296825753 | 3.57968463110397 | 2.13421729395427 |
| chr2 | 37203770 | 37203809 | + | 25.0990144218341 | 14.780944646837 | 58.4681823080315 | 23.476390233497 |
| chr2 | 37203885 | 37203886 | + | 2.00792115374673 | 1.13699574206438 | 3.57968463110397 | 1.06710864697714 |
| chr2 | 37203909 | 37203945 | + | 5.01980288436682 | 11.3699574206438 | 3.57968463110397 | 1.06710864697714 |
| chr2 | 37204557 | 37204568 | + | 7.02772403811355 | 11.3699574206438 | 7.15936926220794 | 9.60397782279422 |
| chr2 | 37204599 | 37204658 | + | 22.087132691214 | 9.09596593651506 | 22.6713359969918 | 10.6710864697714 |
| chr2 | 37249118 | 37249140 | + | 6.02376346124019 | 11.3699574206438 | 5.96614105183995 | 5.33554323488568 |
| chr2 | 37252972 | 37252975 | - | 3.01188173062009 | 2.27399148412876 | 1.19322821036799 | 5.33554323488568 |
| chr2 | 37253010 | 37253012 | - | 5.01980288436682 | 17.0549361309657 | 4.77291284147196 | 7.46976052883995 |
| chr2 | 38294916 | 38295014 | - | 5.01980288436682 | 7.95897019445067 | 3.57968463110397 | 8.53686917581709 |
| chr2 | 38295068 | 38295154 | - | 3.01188173062009 | 7.95897019445067 | 8.35259747257593 | 7.46976052883995 |
| chr2 | 38295305 | 38295306 | - | 4.01584230749346 | 5.68497871032191 | 3.57968463110397 | 3.20132594093141 |
| chr2 | 38562289 | 38562290 | - | 4.01584230749346 | 3.41098722619315 | 1.19322821036799 | 4.26843458790854 |
| chr2 | 38562396 | 38562411 | - | 3.01188173062009 | 2.27399148412876 | 5.96614105183995 | 10.6710864697714 |
| chr2 | 38564086 | 38564102 | - | 1.00396057687336 | 9.09596593651506 | 1.19322821036799 | 2.13421729395427 |
| chr2 | 38564128 | 38564132 | - | 1.00396057687336 | 2.27399148412876 | 4.77291284147196 | 3.20132594093141 |
| chr2 | 38564167 | 38564194 | - | 4.01584230749346 | 11.3699574206438 | 11.9322821036799 | 21.3421729395427 |
| chr2 | 38733519 | 38733595 | + | 3.01188173062009 | 2.27399148412876 | 3.57968463110397 | 4.26843458790854 |
| chr2 | 38733630 | 38733744 | + | 1.00396057687336 | 3.41098722619315 | 9.54582568294392 | 16.006629704657 |
| chr2 | 38733781 | 38733810 | + | 4.01584230749346 | 3.41098722619315 | 5.96614105183995 | 2.13421729395427 |
| chr2 | 38734014 | 38734024 | + | 2.00792115374673 | 1.13699574206438 | 1.19322821036799 | 5.33554323488568 |
| chr2 | 38734434 | 38734454 | + | 2.00792115374673 | 1.13699574206438 | 4.77291284147196 | 7.46976052883995 |
| chr2 | 38734588 | 38734635 | + | 6.02376346124019 | 2.27399148412876 | 7.15936926220794 | 12.8053037637256 |
| chr2 | 38734663 | 38734681 | + | 1.00396057687336 | 2.27399148412876 | 1.19322821036799 | 2.13421729395427 |
| chr2 | 38734718 | 38734748 | + | 2.00792115374673 | 3.41098722619315 | 1.19322821036799 | 3.20132594093141 |
| chr2 | 38743931 | 38743987 | - | 53.2099105742883 | 64.8087572976698 | 52.5020412561915 | 78.966039876308 |
| chr2 | 38744016 | 38744022 | - | 5.01980288436682 | 7.95897019445067 | 8.35259747257593 | 12.8053037637256 |
| chr2 | 38744043 | 38744048 | - | 5.01980288436682 | 9.09596593651506 | 8.35259747257593 | 7.46976052883995 |
| chr2 | 38744070 | 38744167 | - | 12.0475269224804 | 29.5618892936739 | 46.5359002043516 | 33.0803680562912 |
| chr2 | 38744200 | 38744228 | - | 4.01584230749346 | 5.68497871032191 | 5.96614105183995 | 1.06710864697714 |
| chr2 | 38744338 | 38744345 | - | 4.01584230749346 | 3.41098722619315 | 5.96614105183995 | 1.06710864697714 |
| chr2 | 38744400 | 38744424 | - | 65.2574374967687 | 68.2197445238629 | 81.1395183050233 | 67.2278447595595 |
| chr2 | 38744449 | 38744465 | - | 2.00792115374673 | 4.54798296825753 | 9.54582568294392 | 5.33554323488568 |
| chr2 | 38744576 | 38744606 | - | 1.00396057687336 | 7.95897019445067 | 3.57968463110397 | 4.26843458790854 |
| chr2 | 38744918 | 38744993 | - | 36.1425807674411 | 95.5076423334081 | 81.1395183050233 | 119.516168461439 |
| chr2 | 38745012 | 38745021 | - | 6.02376346124019 | 17.0549361309657 | 14.3187385244159 | 5.33554323488568 |
| chr2 | 38745047 | 38745190 | - | 169.669337491599 | 380.893573591568 | 431.948612153212 | 330.803680562912 |
| chr2 | 38781775 | 38781783 | + | 8.03168461498692 | 14.780944646837 | 5.96614105183995 | 6.40265188186281 |
| chr2 | 38781894 | 38781928 | + | 26.1029749987075 | 19.3289276150945 | 23.8645642073598 | 30.9461507623369 |
| chr2 | 38797853 | 38797855 | - | 1.00396057687336 | 2.27399148412876 | 2.38645642073598 | 5.33554323488568 |
| chr2 | 38797924 | 38797948 | - | 3.01188173062009 | 2.27399148412876 | 7.15936926220794 | 4.26843458790854 |
| chr2 | 38882567 | 38882597 | + | 7.02772403811355 | 7.95897019445067 | 2.38645642073598 | 1.06710864697714 |
| chr2 | 38981747 | 38981752 | - | 1.00396057687336 | 3.41098722619315 | 1.19322821036799 | 2.13421729395427 |
| chr2 | 38981831 | 38981843 | - | 3.01188173062009 | 13.6439489047726 | 3.57968463110397 | 4.26843458790854 |
| chr2 | 38981874 | 38981939 | - | 12.0475269224804 | 14.780944646837 | 10.7390538933119 | 9.60397782279422 |
| chr2 | 38982867 | 38982879 | - | 2.00792115374673 | 9.09596593651506 | 4.77291284147196 | 1.06710864697714 |
| chr2 | 38983149 | 38983170 | - | 7.02772403811355 | 17.0549361309657 | 3.57968463110397 | 6.40265188186281 |
| chr2 | 38984525 | 38984531 | - | 5.01980288436682 | 2.27399148412876 | 7.15936926220794 | 2.13421729395427 |
| chr2 | 38984767 | 38984776 | - | 3.01188173062009 | 3.41098722619315 | 4.77291284147196 | 3.20132594093141 |
| chr2 | 38984850 | 38984905 | - | 1.00396057687336 | 3.41098722619315 | 4.77291284147196 | 4.26843458790854 |
| chr2 | 38985120 | 38985127 | - | 1.00396057687336 | 4.54798296825753 | 1.19322821036799 | 4.26843458790854 |
| chr2 | 39250456 | 39250512 | - | 4.01584230749346 | 3.41098722619315 | 2.38645642073598 | 1.06710864697714 |
| chr2 | 39437355 | 39437475 | + | 29.1148567293276 | 44.3428339405109 | 175.404546924094 | 181.408469986113 |
| chr2 | 39736392 | 39736393 | - | 2.00792115374673 | 7.95897019445067 | 3.57968463110397 | 5.33554323488568 |
| chr2 | 39736595 | 39736612 | - | 8.03168461498692 | 1.13699574206438 | 3.57968463110397 | 8.53686917581709 |
| chr2 | 42330029 | 42330314 | + | 6.02376346124019 | 12.5069531627082 | 13.1255103140479 | 6.40265188186281 |
| chr2 | 42330368 | 42330614 | + | 14.0554480762271 | 14.780944646837 | 15.5119667347839 | 30.9461507623369 |
| chr2 | 42331114 | 42331273 | + | 4.01584230749346 | 3.41098722619315 | 3.57968463110397 | 9.60397782279422 |
| chr2 | 42332020 | 42332279 | + | 3.01188173062009 | 10.2329616785794 | 5.96614105183995 | 11.7381951167485 |
| chr2 | 42332304 | 42332528 | + | 7.02772403811355 | 9.09596593651506 | 8.35259747257593 | 10.6710864697714 |
| chr2 | 42350523 | 42350583 | - | 5.01980288436682 | 21.6029190992233 | 9.54582568294392 | 8.53686917581709 |
| chr2 | 42350610 | 42350656 | - | 16.0633692299738 | 17.0549361309657 | 21.4781077866238 | 10.6710864697714 |
| chr2 | 42350684 | 42350737 | - | 20.0792115374673 | 27.2878978095452 | 19.0916513658878 | 18.1408469986113 |
| chr2 | 42350784 | 42350946 | - | 10.0396057687336 | 20.4659233571589 | 3.57968463110397 | 17.0737383516342 |
| chr2 | 42351160 | 42351222 | - | 15.0594086531005 | 14.780944646837 | 28.6374770488317 | 30.9461507623369 |
| chr2 | 42709054 | 42709157 | + | 5.01980288436682 | 2.27399148412876 | 7.15936926220794 | 2.13421729395427 |
| chr2 | 43222428 | 43222510 | - | 1.00396057687336 | 1.13699574206438 | 2.38645642073598 | 1.06710864697714 |
| chr2 | 43222610 | 43222629 | - | 2.00792115374673 | 4.54798296825753 | 2.38645642073598 | 3.20132594093141 |
| chr2 | 43222794 | 43222858 | - | 1.00396057687336 | 9.09596593651506 | 3.57968463110397 | 25.6106075274513 |
| chr2 | 43222887 | 43222942 | - | 12.0475269224804 | 7.95897019445067 | 13.1255103140479 | 14.9395210576799 |
| chr2 | 43223564 | 43223622 | - | 16.0633692299738 | 7.95897019445067 | 10.7390538933119 | 32.0132594093141 |
| chr2 | 43223643 | 43223658 | - | 7.02772403811355 | 3.41098722619315 | 2.38645642073598 | 6.40265188186281 |
| chr2 | 43223679 | 43223695 | - | 18.0712903837206 | 9.09596593651506 | 25.0577924177278 | 37.3488026441998 |
| chr2 | 43223788 | 43223832 | - | 4.01584230749346 | 3.41098722619315 | 3.57968463110397 | 10.6710864697714 |
| chr2 | 43223851 | 43223897 | - | 6.02376346124019 | 4.54798296825753 | 4.77291284147196 | 14.9395210576799 |
| chr2 | 43223951 | 43223980 | - | 4.01584230749346 | 3.41098722619315 | 4.77291284147196 | 10.6710864697714 |
| chr2 | 43224011 | 43224097 | - | 13.0514874993537 | 12.5069531627082 | 25.0577924177278 | 16.006629704657 |
| chr2 | 43224123 | 43224241 | - | 83.3287278804893 | 103.466612527859 | 108.583767143487 | 126.985928990279 |
| chr2 | 43224270 | 43224295 | - | 5.01980288436682 | 5.68497871032191 | 3.57968463110397 | 20.2750642925656 |
| chr2 | 43809883 | 43809910 | + | 1.00396057687336 | 1.13699574206438 | 2.38645642073598 | 5.33554323488568 |
| chr2 | 43809942 | 43809964 | + | 3.01188173062009 | 6.82197445238629 | 10.7390538933119 | 7.46976052883995 |
| chr2 | 43886552 | 43886557 | - | 4.01584230749346 | 6.82197445238629 | 3.57968463110397 | 5.33554323488568 |
| chr2 | 43887778 | 43887879 | - | 6.02376346124019 | 9.09596593651506 | 2.38645642073598 | 8.53686917581709 |
| chr2 | 43888018 | 43888043 | - | 13.0514874993537 | 4.54798296825753 | 1.19322821036799 | 2.13421729395427 |
| chr2 | 43888288 | 43888657 | - | 335.322832675704 | 476.401215924976 | 390.185624790333 | 313.729942211278 |
| chr2 | 44233499 | 44233519 | + | 1.00396057687336 | 9.09596593651506 | 4.77291284147196 | 3.20132594093141 |
| chr2 | 44233560 | 44233564 | + | 2.00792115374673 | 2.27399148412876 | 2.38645642073598 | 1.06710864697714 |
| chr2 | 44233891 | 44233912 | + | 6.02376346124019 | 11.3699574206438 | 7.15936926220794 | 1.06710864697714 |
| chr2 | 44234011 | 44234032 | + | 1.00396057687336 | 2.27399148412876 | 1.19322821036799 | 2.13421729395427 |
| chr2 | 44318992 | 44319016 | - | 9.03564519186028 | 10.2329616785794 | 8.35259747257593 | 7.46976052883995 |
| chr2 | 44319173 | 44319231 | - | 25.0990144218341 | 59.1237785873479 | 23.8645642073598 | 30.9461507623369 |
| chr2 | 44320202 | 44320216 | - | 3.01188173062009 | 3.41098722619315 | 1.19322821036799 | 2.13421729395427 |
| chr2 | 45388893 | 45388903 | - | 2.00792115374673 | 1.13699574206438 | 1.19322821036799 | 6.40265188186281 |
| chr2 | 46385242 | 46385333 | + | 16.0633692299738 | 22.7399148412876 | 2.38645642073598 | 3.20132594093141 |
| chr2 | 46385748 | 46385764 | + | 23.0910932680874 | 62.534765813541 | 3.57968463110397 | 2.13421729395427 |
| chr2 | 46581910 | 46581920 | + | 3.01188173062009 | 1.13699574206438 | 1.19322821036799 | 2.13421729395427 |
| chr2 | 46582269 | 46582270 | + | 48.1901076899215 | 250.139063254164 | 114.549908195327 | 108.845081991668 |
| chr2 | 46582370 | 46582377 | + | 3.01188173062009 | 6.82197445238629 | 5.96614105183995 | 18.1408469986113 |
| chr2 | 46624537 | 46624560 | + | 14.0554480762271 | 12.5069531627082 | 3.57968463110397 | 5.33554323488568 |
| chr2 | 46624781 | 46624841 | + | 17.0673298068472 | 18.1919318730301 | 17.8984231555198 | 10.6710864697714 |
| chr2 | 46624948 | 46624969 | + | 9.03564519186028 | 10.2329616785794 | 3.57968463110397 | 6.40265188186281 |
| chr2 | 46760654 | 46760676 | + | 6.02376346124019 | 4.54798296825753 | 4.77291284147196 | 7.46976052883995 |
| chr2 | 46760783 | 46760829 | + | 10.0396057687336 | 9.09596593651506 | 10.7390538933119 | 5.33554323488568 |
| chr2 | 46760864 | 46760877 | + | 4.01584230749346 | 4.54798296825753 | 1.19322821036799 | 1.06710864697714 |
| chr2 | 46761074 | 46761077 | + | 4.01584230749346 | 2.27399148412876 | 1.19322821036799 | 2.13421729395427 |
| chr2 | 46901895 | 46901963 | - | 4.01584230749346 | 10.2329616785794 | 5.96614105183995 | 1.06710864697714 |
| chr2 | 46901987 | 46902005 | - | 2.00792115374673 | 3.41098722619315 | 4.77291284147196 | 4.26843458790854 |
| chr2 | 46902057 | 46902243 | - | 42.1663442286813 | 62.534765813541 | 38.1833027317757 | 44.8185631730397 |
| chr2 | 46902270 | 46902327 | - | 25.0990144218341 | 22.7399148412876 | 23.8645642073598 | 18.1408469986113 |
| chr2 | 46902370 | 46902449 | - | 24.0950538449607 | 26.1509020674808 | 7.15936926220794 | 5.33554323488568 |
| chr2 | 46902475 | 46902519 | - | 9.03564519186028 | 9.09596593651506 | 13.1255103140479 | 1.06710864697714 |
| chr2 | 46902576 | 46902627 | - | 4.01584230749346 | 6.82197445238629 | 4.77291284147196 | 3.20132594093141 |
| chr2 | 46902745 | 46902828 | - | 3.01188173062009 | 1.13699574206438 | 1.19322821036799 | 2.13421729395427 |
| chr2 | 46903546 | 46903571 | - | 3.01188173062009 | 2.27399148412876 | 4.77291284147196 | 2.13421729395427 |
| chr2 | 46903960 | 46903982 | - | 2.00792115374673 | 5.68497871032191 | 2.38645642073598 | 7.46976052883995 |
| chr2 | 46905236 | 46905358 | - | 26.1029749987075 | 39.7948509722534 | 44.1494437836156 | 50.1541064079254 |
| chr2 | 46905394 | 46905470 | - | 19.0752509605939 | 14.780944646837 | 20.2848795762558 | 25.6106075274513 |
| chr2 | 47074888 | 47074896 | + | 28.1108961524542 | 21.6029190992233 | 3.57968463110397 | 3.20132594093141 |
| chr2 | 47075706 | 47075727 | + | 3.01188173062009 | 2.27399148412876 | 2.38645642073598 | 1.06710864697714 |
| chr2 | 47075935 | 47076006 | + | 48.1901076899215 | 30.6988850357383 | 8.35259747257593 | 16.006629704657 |
| chr2 | 47160187 | 47160262 | - | 33.130699036821 | 51.1648083928972 | 17.8984231555198 | 17.0737383516342 |
| chr2 | 47160296 | 47160309 | - | 47.1861471130481 | 46.6168254246397 | 34.6036181006717 | 14.9395210576799 |
| chr2 | 47160336 | 47160697 | - | 16234.0425280423 | 14672.9300513408 | 8788.12576936024 | 8613.70099839944 |
| chr2 | 47239117 | 47239118 | + | 1.00396057687336 | 2.27399148412876 | 2.38645642073598 | 1.06710864697714 |
| chr2 | 47483051 | 47483060 | + | 2.00792115374673 | 2.27399148412876 | 5.96614105183995 | 3.20132594093141 |
| chr2 | 47483134 | 47483187 | + | 15.0594086531005 | 25.0139063254164 | 38.1833027317757 | 33.0803680562912 |
| chr2 | 47806900 | 47806926 | + | 21.0831721143407 | 65.9457530397342 | 19.0916513658878 | 13.8724124107028 |
| chr2 | 47806986 | 47807007 | - | 3.01188173062009 | 4.54798296825753 | 2.38645642073598 | 1.06710864697714 |
| chr2 | 47807247 | 47807275 | - | 4.01584230749346 | 9.09596593651506 | 4.77291284147196 | 6.40265188186281 |
| chr2 | 47807306 | 47807314 | - | 9.03564519186028 | 19.3289276150945 | 2.38645642073598 | 3.20132594093141 |
| chr2 | 47807350 | 47807369 | - | 23.0910932680874 | 35.2468680039958 | 14.3187385244159 | 21.3421729395427 |
| chr2 | 47807498 | 47807501 | - | 2.00792115374673 | 3.41098722619315 | 2.38645642073598 | 2.13421729395427 |
| chr2 | 47807618 | 47807665 | - | 7.02772403811355 | 3.41098722619315 | 4.77291284147196 | 1.06710864697714 |
| chr2 | 47807689 | 47807755 | - | 8.03168461498692 | 7.95897019445067 | 14.3187385244159 | 12.8053037637256 |
| chr2 | 47807888 | 47807972 | - | 6.02376346124019 | 15.9179403889013 | 16.7051949451519 | 4.26843458790854 |
| chr2 | 48375525 | 48375544 | + | 11.043566345607 | 31.8358807778027 | 21.4781077866238 | 12.8053037637256 |
| chr2 | 48376750 | 48376812 | + | 3.01188173062009 | 4.54798296825753 | 10.7390538933119 | 5.33554323488568 |
| chr2 | 48376883 | 48376938 | + | 10.0396057687336 | 18.1919318730301 | 8.35259747257593 | 16.006629704657 |
| chr2 | 48377015 | 48377018 | + | 5.01980288436682 | 3.41098722619315 | 4.77291284147196 | 2.13421729395427 |
| chr2 | 48377064 | 48377071 | + | 1.00396057687336 | 3.41098722619315 | 8.35259747257593 | 1.06710864697714 |
| chr2 | 48377198 | 48377211 | + | 4.01584230749346 | 2.27399148412876 | 29.8307052591997 | 18.1408469986113 |
| chr2 | 48378314 | 48378369 | + | 1.00396057687336 | 1.13699574206438 | 9.54582568294392 | 4.26843458790854 |
| chr2 | 48378650 | 48378676 | + | 1.00396057687336 | 2.27399148412876 | 4.77291284147196 | 10.6710864697714 |
| chr2 | 48379241 | 48379268 | + | 1.00396057687336 | 1.13699574206438 | 1.19322821036799 | 1.06710864697714 |
| chr2 | 48515261 | 48515271 | + | 1.00396057687336 | 3.41098722619315 | 2.38645642073598 | 2.13421729395427 |
| chr2 | 50619877 | 50619878 | - | 8.03168461498692 | 12.5069531627082 | 5.96614105183995 | 4.26843458790854 |
| chr2 | 53670059 | 53670080 | - | 1.00396057687336 | 1.13699574206438 | 3.57968463110397 | 4.26843458790854 |
| chr2 | 53670347 | 53670451 | - | 7.02772403811355 | 31.8358807778027 | 25.0577924177278 | 40.5501285851312 |
| chr2 | 53818112 | 53818158 | + | 4.01584230749346 | 5.68497871032191 | 2.38645642073598 | 1.06710864697714 |
| chr2 | 53818488 | 53818516 | + | 6.02376346124019 | 7.95897019445067 | 3.57968463110397 | 4.26843458790854 |
| chr2 | 53818627 | 53818674 | + | 3.01188173062009 | 2.27399148412876 | 2.38645642073598 | 2.13421729395427 |
| chr2 | 53864895 | 53864979 | - | 26.1029749987075 | 34.1098722619315 | 35.7968463110397 | 29.8790421153598 |
| chr2 | 53865000 | 53865016 | - | 1.00396057687336 | 1.13699574206438 | 2.38645642073598 | 3.20132594093141 |
| chr2 | 53865060 | 53865151 | - | 13.0514874993537 | 13.6439489047726 | 17.8984231555198 | 17.0737383516342 |
| chr2 | 53865237 | 53865282 | - | 4.01584230749346 | 4.54798296825753 | 7.15936926220794 | 10.6710864697714 |
| chr2 | 53865368 | 53865414 | - | 4.01584230749346 | 2.27399148412876 | 15.5119667347839 | 17.0737383516342 |
| chr2 | 53865459 | 53865475 | - | 1.00396057687336 | 2.27399148412876 | 1.19322821036799 | 5.33554323488568 |
| chr2 | 53866084 | 53866114 | - | 4.01584230749346 | 3.41098722619315 | 8.35259747257593 | 4.26843458790854 |
| chr2 | 54660206 | 54660275 | + | 1.00396057687336 | 1.13699574206438 | 1.19322821036799 | 3.20132594093141 |
| chr2 | 54668502 | 54668676 | + | 10.0396057687336 | 13.6439489047726 | 8.35259747257593 | 13.8724124107028 |
| chr2 | 54668704 | 54668784 | + | 63.249516343022 | 79.5897019445067 | 35.7968463110397 | 25.6106075274513 |
| chr2 | 54669021 | 54669028 | + | 7.02772403811355 | 1.13699574206438 | 1.19322821036799 | 1.06710864697714 |
| chr2 | 54669137 | 54669229 | + | 13.0514874993537 | 7.95897019445067 | 1.19322821036799 | 2.13421729395427 |
| chr2 | 54669555 | 54669655 | + | 17.0673298068472 | 25.0139063254164 | 3.57968463110397 | 9.60397782279422 |
| chr2 | 54972214 | 54972318 | - | 15.0594086531005 | 22.7399148412876 | 20.2848795762558 | 13.8724124107028 |
| chr2 | 54972343 | 54972494 | - | 11.043566345607 | 21.6029190992233 | 20.2848795762558 | 20.2750642925656 |
| chr2 | 54972546 | 54972556 | - | 2.00792115374673 | 1.13699574206438 | 2.38645642073598 | 2.13421729395427 |
| chr2 | 54972632 | 54972782 | - | 35.1386201905678 | 50.0278126508328 | 19.0916513658878 | 40.5501285851312 |
| chr2 | 54972802 | 54972859 | - | 14.0554480762271 | 18.1919318730301 | 10.7390538933119 | 12.8053037637256 |
| chr2 | 54972891 | 54972903 | - | 8.03168461498692 | 4.54798296825753 | 11.9322821036799 | 8.53686917581709 |
| chr2 | 54972925 | 54973015 | - | 10.0396057687336 | 9.09596593651506 | 8.35259747257593 | 13.8724124107028 |
| chr2 | 54973037 | 54973183 | - | 280.105000947669 | 285.38593125816 | 143.187385244159 | 252.904749333581 |
| chr2 | 55175766 | 55175770 | - | 1.00396057687336 | 7.95897019445067 | 2.38645642073598 | 5.33554323488568 |
| chr2 | 55288381 | 55288385 | - | 3.01188173062009 | 2.27399148412876 | 3.57968463110397 | 4.26843458790854 |
| chr2 | 55288433 | 55288437 | - | 9.03564519186028 | 6.82197445238629 | 4.77291284147196 | 17.0737383516342 |
| chr2 | 55548618 | 55548636 | - | 6.02376346124019 | 2.27399148412876 | 2.38645642073598 | 5.33554323488568 |
| chr2 | 55549566 | 55549663 | - | 17.0673298068472 | 6.82197445238629 | 11.9322821036799 | 19.2079556455884 |
| chr2 | 55617870 | 55617953 | + | 318.255502868857 | 250.139063254164 | 164.665493030783 | 257.17318392149 |
| chr2 | 55618101 | 55618103 | + | 1.00396057687336 | 1.13699574206438 | 3.57968463110397 | 9.60397782279422 |
| chr2 | 55636157 | 55636197 | - | 6.02376346124019 | 13.6439489047726 | 10.7390538933119 | 13.8724124107028 |
| chr2 | 58159267 | 58159315 | - | 12.0475269224804 | 13.6439489047726 | 13.1255103140479 | 8.53686917581709 |
| chr2 | 58159373 | 58159417 | - | 10.0396057687336 | 23.876910583352 | 19.0916513658878 | 22.4092815865198 |
| chr2 | 58159440 | 58159479 | - | 11.043566345607 | 21.6029190992233 | 8.35259747257593 | 5.33554323488568 |
| chr2 | 58159542 | 58159546 | + | 1.00396057687336 | 1.13699574206438 | 1.19322821036799 | 2.13421729395427 |
| chr2 | 58159580 | 58159593 | - | 5.01980288436682 | 9.09596593651506 | 1.19322821036799 | 3.20132594093141 |
| chr2 | 60798223 | 60798232 | + | 1.00396057687336 | 1.13699574206438 | 1.19322821036799 | 3.20132594093141 |
| chr2 | 60924808 | 60924811 | + | 6.02376346124019 | 3.41098722619315 | 1.19322821036799 | 2.13421729395427 |
| chr2 | 60925676 | 60925691 | + | 9.03564519186028 | 3.41098722619315 | 1.19322821036799 | 3.20132594093141 |
| chr2 | 60927220 | 60927241 | + | 8.03168461498692 | 2.27399148412876 | 3.57968463110397 | 1.06710864697714 |
| chr2 | 60931212 | 60931216 | + | 3.01188173062009 | 7.95897019445067 | 1.19322821036799 | 3.20132594093141 |
| chr2 | 60931272 | 60931276 | + | 2.00792115374673 | 7.95897019445067 | 2.38645642073598 | 1.06710864697714 |
| chr2 | 61018294 | 61018295 | + | 1.00396057687336 | 2.27399148412876 | 1.19322821036799 | 1.06710864697714 |
| chr2 | 61018347 | 61018372 | + | 5.01980288436682 | 4.54798296825753 | 1.19322821036799 | 1.06710864697714 |
| chr2 | 61018504 | 61018620 | + | 20.0792115374673 | 39.7948509722534 | 32.2171616799357 | 18.1408469986113 |
| chr2 | 61018792 | 61018802 | + | 4.01584230749346 | 2.27399148412876 | 3.57968463110397 | 5.33554323488568 |
| chr2 | 61019056 | 61019059 | + | 1.00396057687336 | 1.13699574206438 | 5.96614105183995 | 1.06710864697714 |
| chr2 | 61048786 | 61048808 | + | 3.01188173062009 | 2.27399148412876 | 3.57968463110397 | 3.20132594093141 |
| chr2 | 61048864 | 61048899 | + | 5.01980288436682 | 14.780944646837 | 10.7390538933119 | 3.20132594093141 |
| chr2 | 61048921 | 61048942 | + | 7.02772403811355 | 9.09596593651506 | 5.96614105183995 | 3.20132594093141 |
| chr2 | 61049163 | 61049194 | + | 4.01584230749346 | 11.3699574206438 | 1.19322821036799 | 9.60397782279422 |
| chr2 | 61051521 | 61051530 | + | 2.00792115374673 | 2.27399148412876 | 1.19322821036799 | 1.06710864697714 |
| chr2 | 61183545 | 61183576 | + | 5.01980288436682 | 5.68497871032191 | 3.57968463110397 | 1.06710864697714 |
| chr2 | 61184718 | 61184725 | + | 3.01188173062009 | 13.6439489047726 | 1.19322821036799 | 3.20132594093141 |
| chr2 | 61185522 | 61185593 | + | 24.0950538449607 | 19.3289276150945 | 9.54582568294392 | 5.33554323488568 |
| chr2 | 61185609 | 61185619 | + | 14.0554480762271 | 20.4659233571589 | 9.54582568294392 | 2.13421729395427 |
| chr2 | 61186080 | 61186118 | + | 15.0594086531005 | 5.68497871032191 | 8.35259747257593 | 2.13421729395427 |
| chr2 | 61186470 | 61186497 | + | 8.03168461498692 | 1.13699574206438 | 2.38645642073598 | 4.26843458790854 |
| chr2 | 61186587 | 61186631 | + | 4.01584230749346 | 3.41098722619315 | 1.19322821036799 | 3.20132594093141 |
| chr2 | 61186804 | 61186901 | + | 10.0396057687336 | 20.4659233571589 | 13.1255103140479 | 5.33554323488568 |
| chr2 | 61187231 | 61187247 | + | 4.01584230749346 | 6.82197445238629 | 3.57968463110397 | 1.06710864697714 |
| chr2 | 61187552 | 61187575 | - | 4.01584230749346 | 7.95897019445067 | 1.19322821036799 | 4.26843458790854 |
| chr2 | 61471354 | 61471384 | + | 7.02772403811355 | 7.95897019445067 | 10.7390538933119 | 6.40265188186281 |
| chr2 | 61477958 | 61477975 | - | 1.00396057687336 | 3.41098722619315 | 8.35259747257593 | 1.06710864697714 |
| chr2 | 61478106 | 61478112 | - | 1.00396057687336 | 1.13699574206438 | 2.38645642073598 | 1.06710864697714 |
| chr2 | 61478142 | 61478150 | - | 7.02772403811355 | 4.54798296825753 | 4.77291284147196 | 4.26843458790854 |
| chr2 | 61478418 | 61478422 | - | 2.00792115374673 | 2.27399148412876 | 1.19322821036799 | 2.13421729395427 |
| chr2 | 61478452 | 61478456 | - | 6.02376346124019 | 3.41098722619315 | 8.35259747257593 | 6.40265188186281 |
| chr2 | 61478689 | 61478711 | - | 5.01980288436682 | 3.41098722619315 | 4.77291284147196 | 4.26843458790854 |
| chr2 | 61478737 | 61478777 | - | 1.00396057687336 | 5.68497871032191 | 9.54582568294392 | 4.26843458790854 |
| chr2 | 61868196 | 61868248 | - | 33.130699036821 | 19.3289276150945 | 27.4442488384638 | 23.476390233497 |
| chr2 | 61868269 | 61868320 | - | 59.2336740355285 | 37.5208594881246 | 84.7192029361272 | 65.0936274656053 |
| chr2 | 61868361 | 61868408 | - | 19.0752509605939 | 28.4248935516096 | 10.7390538933119 | 14.9395210576799 |
| chr2 | 61868454 | 61868522 | - | 23.0910932680874 | 25.0139063254164 | 31.0239334695677 | 26.6777161744284 |
| chr2 | 61868546 | 61868586 | - | 35.1386201905678 | 31.8358807778027 | 40.5697591525116 | 86.435800405148 |
| chr2 | 61868637 | 61868659 | - | 11.043566345607 | 12.5069531627082 | 25.0577924177278 | 20.2750642925656 |
| chr2 | 61868680 | 61868697 | - | 4.01584230749346 | 1.13699574206438 | 2.38645642073598 | 1.06710864697714 |
| chr2 | 62224174 | 62224278 | + | 5.01980288436682 | 6.82197445238629 | 2.38645642073598 | 8.53686917581709 |
| chr2 | 62224569 | 62224572 | + | 2.00792115374673 | 9.09596593651506 | 5.96614105183995 | 27.7448248214055 |
| chr2 | 63045904 | 63045926 | + | 2.00792115374673 | 3.41098722619315 | 1.19322821036799 | 1.06710864697714 |
| chr2 | 63046109 | 63046182 | + | 7.02772403811355 | 6.82197445238629 | 3.57968463110397 | 2.13421729395427 |
| chr2 | 63056527 | 63056530 | + | 3.01188173062009 | 4.54798296825753 | 20.2848795762558 | 8.53686917581709 |
| chr2 | 63607020 | 63607042 | + | 3.01188173062009 | 2.27399148412876 | 2.38645642073598 | 4.26843458790854 |
| chr2 | 63607065 | 63607173 | + | 33.130699036821 | 47.753821166704 | 41.7629873628796 | 33.0803680562912 |
| chr2 | 63891302 | 63891335 | + | 4.01584230749346 | 4.54798296825753 | 1.19322821036799 | 2.13421729395427 |
| chr2 | 63891369 | 63891398 | + | 12.0475269224804 | 19.3289276150945 | 7.15936926220794 | 12.8053037637256 |
| chr2 | 63891447 | 63891538 | + | 47.1861471130481 | 79.5897019445067 | 10.7390538933119 | 20.2750642925656 |
| chr2 | 63892470 | 63892478 | - | 6.02376346124019 | 1.13699574206438 | 7.15936926220794 | 3.20132594093141 |
| chr2 | 63892519 | 63892560 | - | 5.01980288436682 | 2.27399148412876 | 3.57968463110397 | 1.06710864697714 |
| chr2 | 63892590 | 63892646 | - | 14.0554480762271 | 32.9728765198671 | 21.4781077866238 | 24.5434988804741 |
| chr2 | 63893209 | 63893327 | - | 6.02376346124019 | 6.82197445238629 | 15.5119667347839 | 6.40265188186281 |
| chr2 | 63893367 | 63893451 | - | 2.00792115374673 | 6.82197445238629 | 3.57968463110397 | 3.20132594093141 |
| chr2 | 64093082 | 64093128 | - | 3.01188173062009 | 3.41098722619315 | 25.0577924177278 | 4.26843458790854 |
| chr2 | 64093352 | 64093382 | - | 2.00792115374673 | 1.13699574206438 | 19.0916513658878 | 10.6710864697714 |
| chr2 | 64093416 | 64093436 | - | 1.00396057687336 | 2.27399148412876 | 8.35259747257593 | 5.33554323488568 |
| chr2 | 64093457 | 64093482 | - | 2.00792115374673 | 1.13699574206438 | 7.15936926220794 | 13.8724124107028 |
